# Core cysteine residues in the PAN domain are critical for HGF/c-MET signaling

**DOI:** 10.1101/2021.09.20.460979

**Authors:** Debjani Pal, Kuntal De, Carly M. Shanks, Kai Feng, Timothy B. Yates, Jennifer Morrell-Falvey, Russell B. Davidson, Jerry M. Parks, Wellington Muchero

## Abstract

The Plasminogen-Apple-Nematode (PAN) domain, with a core of four to six cysteine residues, is found in > 28,000 proteins across 959 genera but its role in protein function is not fully understood. The PAN domain was initially characterized to be present in numerous proteins including hepatocyte growth factor (HGF). Dysregulation of HGF-mediated signaling results in numerous deadly cancers. All biological impacts of HGF in cell proliferation are triggered by binding of HGF to its cell surface receptor, cellular mesenchymal-epidermal transition (c-MET). Here, we show that four PAN domain cysteine residues are essential for HGF/c-MET signaling. Mutating these residues resulted in retardation of perinuclear localization, cellular internalization of HGF and its receptor, c-MET, and c-MET ubiquitination. Our observations indicate that the PAN domain of HGF is required for the c-MET binding and subsequent c-MET autophosphorylation and phosphorylation of its downstream targets, protein kinase B (AKT), extracellular signal-regulated kinase (ERK), and signal transducer and activator of transcription 3 (STAT3). Furthermore, transcriptional activation of HGF/c-MET signaling-related genes including matrix metalloproteinase-9 (MMP9), ETS translocation variant 1, 4, and 5 (ETV1, ETV4, ETV5), and early growth response 1 (EGR1) was impaired and cell proliferation was attenuated. These results suggest that core cysteine residues in the PAN domain are critical for HGF/c-MET interaction, c-MET mediated signal transduction, and cell survival. Thus, targeting the PAN domain of HGF may represent a mechanism for selectively regulating the binding and activation of the c-MET pathway.

**Significance:** HGF/c-MET signaling induces multifunctional cellular responses. Dysregulation of HGF/c-MET signaling cascade can lead to tumorigenesis by transforming normal cells to tumor cells. This work defines the importance of core cysteine residues in the PAN domain of HGF in downstream activation of HGF/c-MET signaling. To understand the role of cysteines in the PAN domain, PAN mutants of HGF were used to stimulate c-MET signaling in cells and the impact was delineated by determining phosphorylation and transcription of downstream targets. Mutations in core cysteines in the HGF-PAN domain completely blocked downstream phosphorylation and perinuclear accumulation of c-MET. These results suggest an indispensable role for the cysteine-rich PAN domain in HGF/c-MET interaction and could set the stage for future therapies that selectively disrupt the MET signaling cascade with limited off-target effects in tumors overexpressing HGF/c-MET.

## Introduction

Despite its prevalence in a wide variety of organisms, the role of the Plasminogen-Apple-Nematode (PAN) domain in protein function has largely remained elusive. A key contributing factor is that this domain is found in many phylogenetically unrelated proteins across divergent organisms falling into categories including Alveolata, Archea, Amoebazoa, Bacteria, Cryptophyta, Euglenozoa, Haptophyceae, Opisthokonta, Rhizaria, Rhodophyta, Stramenopilles, Viridplantae, and Viruses. The PAN domain was first characterized by Tordai et al. (*1*) in which they noted that the domain was shared by the plasminogen/hepatocyte growth factor protein family, the prekallikrein/coagulation factor XI protein family, and nematode proteins. The domain possesses the characteristic 4-6-cysteine residues in its core that are strictly conserved. These cysteine residues have been proposed to engage in two or three disulfide bridges to form a hairpin loop structure (*1, 2*). The PAN domain has been suggested to mediate protein-protein or carbohydrate-protein interactions and facilitate receptor dimerization (*2–6*). HGF is secreted by mesenchymal cells as a single-chain, biologically inert precursor and is converted into its bioactive form when extracellular proteases cleave the bond between Arg494 and Val495 (*7*). The mature form of HGF consists of an α- and β-chain, which are held together by a disulfide bond (*8–10*). The α-subunit of HGF includes N-terminal hairpin loop structure and four kringle domains (K1-K4) whereas the β-subunit consists of serine protease homology (SPH) domain (*11*).

To provide evidence for its role in signal transduction, we characterized the functional role of the PAN domain in the heparin binding glycoprotein HGF, which functions as a ligand for the high-affinity receptor, c-MET. HGF/c-MET signaling has been shown to mediate cellular processes including angiogenesis, anti-apoptosis, mitogenesis, morphogenesis, mitogenesis and neurite extension(*12*). Dysregulation of the HGF/c-MET signaling cascade can lead to tumorigenesis by transforming normal cells into tumor cells and c-MET hyperactivation has been reported in cancers including lung cancer, colorectal cancer, glioblastoma, and acute myeloid lymphoma among others(*13, 14*).

An essential aspect of the HGF/c-MET signaling cascade is proteolytic processing of both HGF and c-MET into their active forms, which can then form ligand-receptor complexes in the ECM. Internalization and ubiquitin-dependent degradation of activated HGF and MET triggers signaling cascades via a lysosomal degradative pathway that result in the biological cellular processes described above. (*15, 16*). Previously, Okigaki et al. reported that deleting the N-terminal domain, the first, second, or both kringle domains made HGF functionally inactive as the deletion mutants were unable to trigger c-MET signaling (*17*).

## Results

### PAN domain-carrying proteins are enriched in cell recognition and cellular signaling processes

PAN domain-carrying proteins do not share a clear evolutionary or phylogenetic trajectory suggesting that this domain may have been co-opted independently by organisms to serve protein functions that are yet to be revealed. To provide evidence of its putative function, we scanned the InterPro (https://www.ebi.ac.uk/interpro/) and UniProt (https://G-LecRK.uniprot.org) databases and found 28,300 proteins across 2,496 organisms falling into 959 genera (Table S1). Based on predicted cellular localization, PAN domain proteins predominantly are present in the extracellular matrix (ECM) either as cell surface receptors or ligands for cell surface receptors. Gene Ontology (GO) enrichment analyses using the 28,300 proteins revealed 39 unique GO-terms that were enriched at p < 0.05 (Table S2). These included terms such as cell recognition (*p-value =* 1E-30), cell communication (*p-value =* 1E-30), proteolysis (*p-value =* 1E-30), pollen-pistil interaction (*p-value =* 1E-30), reproduction (*p-value =* 1E-30), response to stimulus (*p-value =* 1E-30) and response to stress (*p-value =* 1E-30). The predominant occurrence of these proteins in the ECM and their enrichment in processes typically associated with cellular signal transduction suggested that the PAN domain may play essential roles in immune response, albeit in different pathways for divergent organisms. Based on these results, we hypothesized that the PAN domain would be essential for HGF/c-MET signaling. To test this hypothesis, we designed experiments to mutate core cysteine residues and assess implications on downstream signaling cascades.

### Core PAN domain cysteine residues modulate HGF overall-abundance

Alignment of the PAN domain of proteins from 14 model organisms revealed four strictly conserved cysteine residues occurring at amino acid positions 70, 74, 84, and 96 of the HGF protein (Fig. S1A&B; Fig. S2). To evaluate the functional significance of these residues, we sequentially mutated single cysteines and also simultaneously mutated all four cysteine residues and examined HGF stability in 293T cells. Mutant HGFs, in which cysteines 70 (C70A), 74 (C74A), 84 (C84A) and 96 (C96A) were substituted with alanine, led to a marked increase in the protein expression of exogenously expressed HGFs in cells compared to their wild-type (WT) counterpart (Fig. 1A-B). Furthermore, mutating all four core cysteines in the PAN domain dramatically increased the abundance of exogenously expressed HGF-4Cys-4Ala in the cells (Fig. 1C). Moreover, the c-MET receptor exhibited a similar increase in abundance in response to stimulation by the HGF 4Cys-4Ala mutant compared to WT HGF. These results suggest that mutant HGF and its c-MET receptor did not undergo the expected lysosomal degradation when the core cysteines were mutated.

**Fig. 1:**
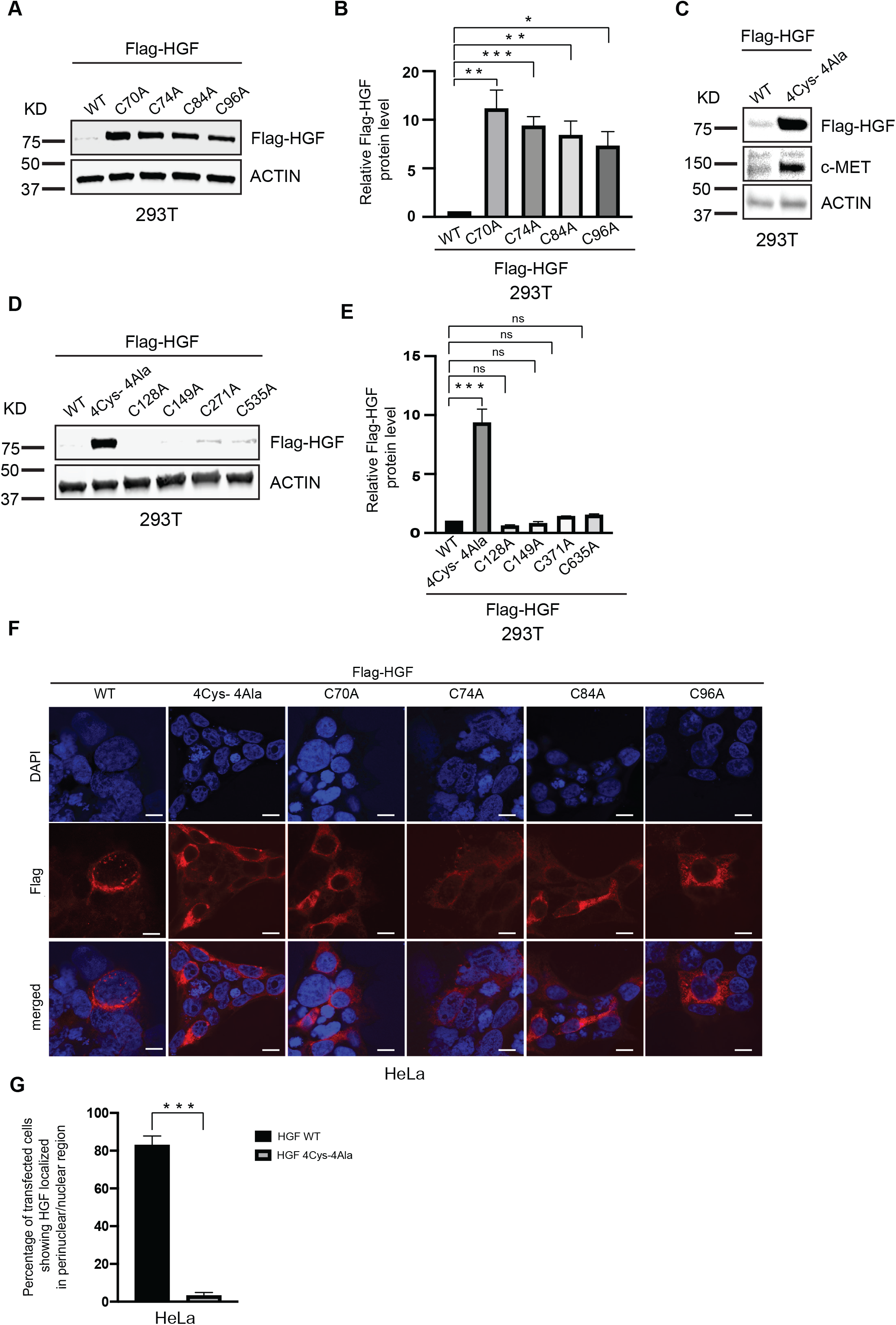
PAN domain modulates HGF stability. (A and B) Core cysteines in PAN domain are crucial for HGF stability. Immunoblot analysis of whole cell lysates derived from 293T cells, transfected with Flag-HGF WT and different single cysteine mutants of Flag-HGF constructs as indicated. 30 h post-transfection, whole-cell lysates were prepared for immunoblot analysis. Representative image of n=3 biological replicates. (B) Quantification of the band intensities in (A). The intensities of Flag-HGF (WT and mutants) bands were normalized to actin and then normalized to Flag-HGF WT. Data are represented as mean ± SD, n = 3, and *p < 0.05, **p < 0.005, *** p < 0.0005 (student’s t-test). (C) Mutation in all four core cysteine residues in PAN domain significantly alters HGF stability. Immunoblot analysis of whole cell lysates derived from 293T cells, transfected with Flag-HGF WT and Flag-HGF 4Cys-4Ala constructs as indicated. 30 h post-transfection, whole-cell lysates were prepared for immunoblot analysis. (D) Cysteines in kringle domains and SPH domain of HGF have no impact on the protein abundance in cells. Immunoblot analysis of whole cell lysates derived from 293T cells, transfected with Flag-HGF WT, Flag-HGF 4Cys-4Ala and different single cysteine mutants of Flag-HGF constructs as indicated. 30 hours post-transfection, whole-cell lysates were prepared for immunoblot analysis. Representative image of n=3 biological replicates. (E) Quantification of the band intensities in (D). The intensities of Flag-HGF (WT and mutants) bands were normalized to actin and then normalized to Flag-HGF WT. Data are represented as mean ± SD, n = 3, and *p < 0.05, **p < 0.005, ***p < 0.0005 (student’s t-test). (F) Localization of Flag-HGF WT and different Flag-HGF mutants’ expression by confocal immunofluorescence microscopy in HeLa cells. The cells were transiently transfected with Flag-HGF WT and different mutants of Flag-HGFs as indicated. 30 h post-transfection cells were fixed, mounted and protein expression patterns were visualized using a Zeiss LSM 710 confocal microscope outfitted with a 63x objective. Scale bars represent 20 μm. The images shown are representative from three independent biological experiments (average 100 cells were observed per experimental condition per replicate). (G) Percentage of transfected HeLa cells showing perinuclear/nuclear staining for Flag-HGF WT and Flag-HGF 4Cys-4Ala were quantified. Data are represented as mean ± SD, n = 3 (average 100 cells were observed for each condition per experiment), and *p<0.05, **p < 0.005, *** p < 0.0005 (Student’s t test).

The kringle domains of HGF have been reported to be crucial for protein-protein interactions. In particular, the first and second kringle domains are especially important for the proper biological function of the protein (*11*). Kringle domains are usually disulfide crosslinked domain and previous studies have established that kringle domains in the α-subunit and the SPH domain in the β-subunit provide c-MET binding sites on HGF (*18, 19*). Recently, Uchikawa et al. determined structures of c-MET/HGF complex mimicking their active state at 4.8 Å resolution using cryo-electron microscopy (*19*). They identified multiple distinctive c-MET binding sites on the HGF protein including the N-terminal and kringle domains but did not implicate the PAN domain in the HGF/c-MET interaction (*19*). Here, we exclusively investigated the role of the PAN domain of HGF on the c-MET signaling cascade.

As we established a connection between the core cysteines in the PAN domain of HGF with HGF abundance, we introduced four more mutations of additional cysteines located in the K1, K2, and SPH domains of HGF (C128A, C149A, C271A, and C535A). None of these mutant HGFs showed increased protein expression when expressed exogenously in cells as did the HGF 4Cys-4Ala (Fig. 1D and 1E). Thus, cysteine residues in the kringle and SPH domains are apparently not involved in HGF activity. Rather, activity is specifically defined by the four strictly conserved cysteine residues in PAN domain.

We hypothesized that the enhanced HGF expression of the 4Cys-4Ala mutant may results from an intra-PAN domain structural change that prevents HGF degradation. To test this hypothesis, we performed molecular dynamics (MD) simulations of the WT and 4Cys-4Ala PAN domains using models generated with AlphaFold2 (*20*) and compared the resulting structural ensembles. Five models were generated for each system and used as starting structures for the simulations. A 200-ns simulation was then performed for each model. Thus, the cumulative simulation time was 1 μs for each system. Analysis of the time evolution of the root-mean-square deviation (RMSD) of each system revealed only minor structural deviations in both sets of simulations. Using the top WT model as a reference structure, the average RMSDs and standard errors of the means were 1.61 +/− 0.02 Å for the WT and 1.71 +/− 0.01 Å for the 4Cys-4Ala system, indicating highly similar overall structures despite the four Cys substitutions in the mutant (Figure S3). The structural analysis indicates that no large-scale structural changes occurred in the 4Cys-4Ala mutant PAN domain compared to the WT on this time scale (Figure S4).

Previous findings have suggested that, upon binding with its receptor, MET, HGF internalization and degradation precedes activation of MET signaling. Translocation of MET-bound HGF toward the perinuclear region is a key step for the degradation of HGF and recycling of the receptor (*7, 16, 21, 22*). The existence of a parallel pathway has also been reported in which HGF-activated MET translocates to the nucleus to initiate calcium signaling (*23*). In agreement with a role for the PAN domain in HGF abundance, we found that mutating the core cysteine residues reduced the perinuclear signal for all mutated HGFs in HeLa cells (Fig. 1F). HeLa cells transfected with HGF 4Cys-4Ala showed less than 5% of the total perinuclear staining of recombinant HGF. However, under the same transfection efficiency 80% of cells transfected with WT HGF showed perinuclear staining for HGF under confocal microscopy (Fig. 1G). As such, both biochemical and immunofluorescence results suggest a critical role for core PAN domain cysteine residues in HGF stability and cellular uptake. Because all single cysteine mutants yielded the same results as the 4Cys-4Ala mutant, all subsequent studies were performed using only the recombinant HGF 4Cys-4Ala variant.

### PAN domain cysteine residues are essential for c-MET, AKT and ERK phosphorylation

Following binding of HGF at the MET semaphoring homology (SEMA) domain, MET homodimerizes and autophosphorylates at two tyrosine residues (Y1234 and Y1235), followed by subsequent phosphorylation of two additional tyrosines in the carboxy-terminal tail (Y1349 and Y1356). This series of events creates a multifunctional docking site for downstream adaptors and effectors that lead to the activation of the Rat sarcoma (RAS)/ERK and phosphatidylinositol 3-kinase (PI3K)/AKT axis via phosphorylation (*24, 25*). To determine the influence of core cysteines on MET phosphorylation, both 293T and glioblastoma U-87 MG cells were stimulated with purified wild-type HGF and HGF 4Cys-4Ala proteins. Western blot analyses revealed the absence of phosphorylated MET, AKT, and ERK in unstimulated controls and in cells stimulated with the HGF 4Cys-4Ala protein in both cell types (Fig. 2A).

**Fig. 2:**
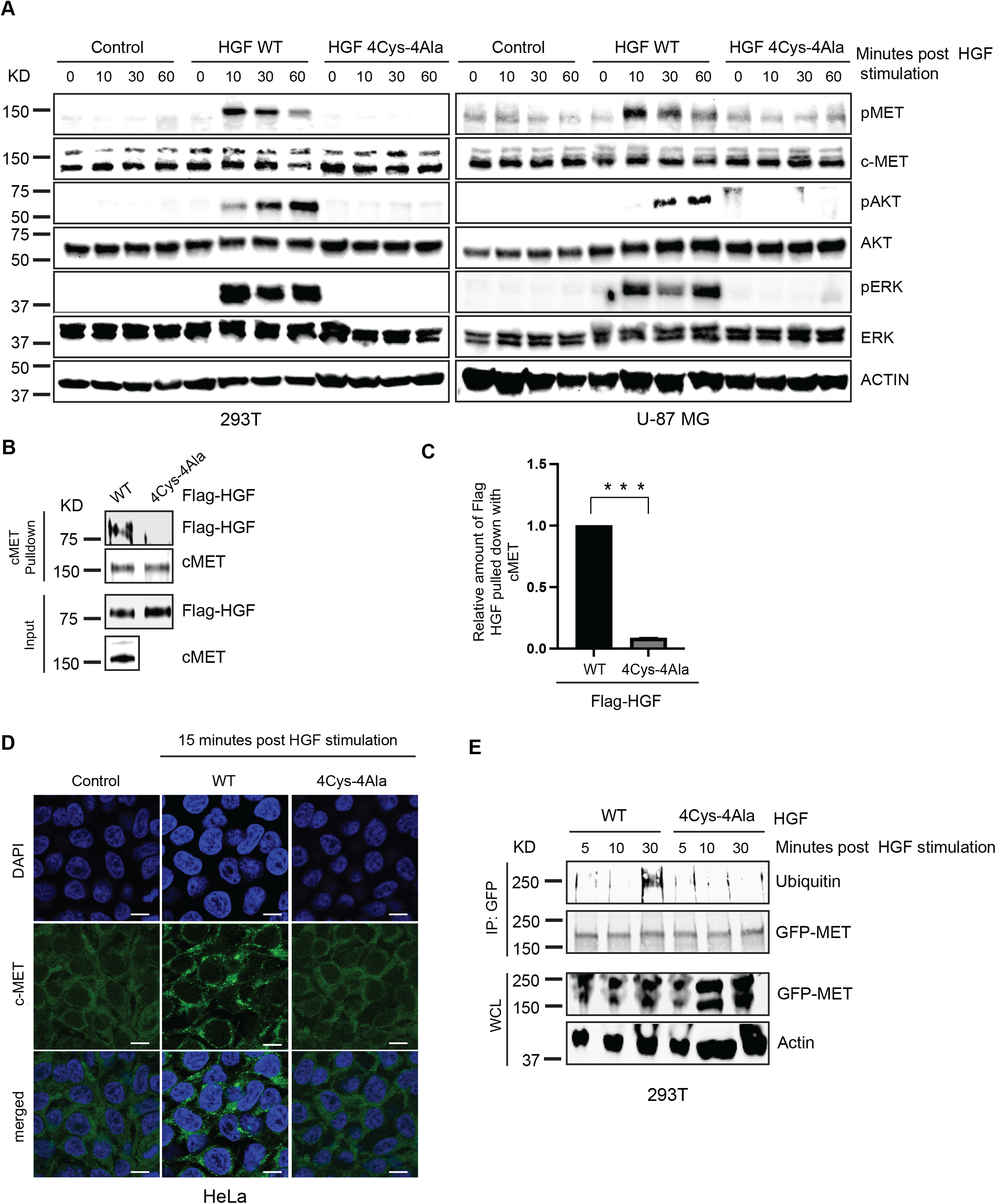
HGF PAN domain regulates MET signaling cascade via four core cysteines. (A) Mutation of the core cysteines in HGF PAN domain blocks HGF induced c-MET signaling. Both 293T and U-87 MG cells were stimulated with HGF WT and HGF 4Cys-4Ala proteins for indicated amount of time. Cells were harvested and immunoblot analysis shows the absence of phosphorylation for MET, AKT and ERK in presence of HGF 4Cys-4Ala. Representative blot images from n=2 experiments for individual cell line. (B) Immunoblot showing that the core cysteines on the HGF PAN domain regulate its binding with c-MET. c-MET was immunoprecipitated from 293T cells on anti-MET bound beads. *In-vitro* translated Flag-tagged HGF WT and HGF 4Cys-4Ala were added to the beads as indicated to detect the interaction between endogenous c-MET and Flag-HGF WT and Flag-HGF 4Cys-4Ala. (C)Right panel, quantification of the band intensities (*n* = 2; *** p < 0.0005 (Student’s t test)). Immunoprecipiated Flag-HGFs band intensities were normalized to the respective c-MET IP bands and then further normalized to HGF-WT. (D) HGF PAN domain defines perinuclear translocation of MET in cells. HeLa cells were stimulated with HGF WT and HGF 4Cys-4Ala proteins for indicated amount of time following serum starvation. Post stimulation cells were fixed, mounted and endogenous c-MET localization pattern was visualized using Zeiss LSM 710 at 63x objective. Scale bars represent 20 μm. The images shown are representative from three independent biological replicates (average 100 cells were observed for each condition per replicate). (E) PAN domain regulates c-MET ubiquitination. In vivo ubiquitination assay shows that HGF WT promotes c-MET ubiquitination in a PAN-dependent manner. 293T cells were transfected with the construct c-MET-C-GFPSpark. After serum starvation, cells were stimulated with HGF WT and HGF 4Cys-4Ala as indicated. The lysates were collected at specific time points and incubated with anti-GFP protein G beads. Ubiquitinated-MET proteins were eluted, resolved by SDS-PAGE and immunoblotted with the indicated antibodies.

To characterize the incompetence of PAN mutant HGF in turning on the phosphorylation cascade of c-MET, we performed a direct interaction assay between endogenous c-MET and *in-vitro* translated HGF proteins and showed that HGF 4Cys-4Ala was unable to interact with c-MET (Fig. 2B and 2C). Our results thus far indicate a central role for the PAN domain in its initial recognition by c-MET. In addition, HGF 4Cys-4Ala, like the unstimulated controls, failed to promote MET perinuclear translocation compared to wild-type HGF based on stimulation assays using HeLa cells (Fig. 2D).

To gain additional insight into the impact of core cysteines in the HGF PAN domain on MET degradation, we performed an *in-vivo* ubiquitination assay with c-MET. Ubiquitination was inhibited with the alanine mutants of HGF compared to wild type (Fig. 2E). Based on these observations, we propose that MET receptors are unable to become internalized by endocytosis, suggesting that the mutated cysteine residues impose an overall retardation of MET activity and its endocytic trafficking.

### Mutating core cysteine residues suppresses STAT3 phosphorylation and nuclear translocation

STAT3 is a transcription factor that is present in the cytoplasm, forms dimers upon activation, and functions as a downstream effector molecule of the HGF/c-MET signaling pathway (*26–28*). STAT3 is reported to be constitutively active in several cancers, which leads to malignant transformation by playing a critical role in stimulating cell proliferation and arresting apoptosis (*28*). We tested the impact of the core cysteines in the HGF PAN domain on STAT3 activation by following its phosphorylation in U-87 MG cells. STAT3 phosphorylation was significantly reduced in cells post-stimulation with the HGF 4Cys-4Ala mutant compared to the wild type (Fig. 3A). Delayed time points were chosen for this assay because HGF induces delayed STAT3 phosphorylation (*29*). Confocal imaging confirmed that HGF 4Cys-4Ala, like the unstimulated controls, was unable to promote STAT3 nuclear translocation, while wild-type HGF promoted normal STAT3 nuclear localization (Fig, 3B). These results are consistent with the above observations, suggesting that mutating the cysteine residues in the PAN domain leads to disrupted HGF/c-MET signaling.

**Fig. 3:**
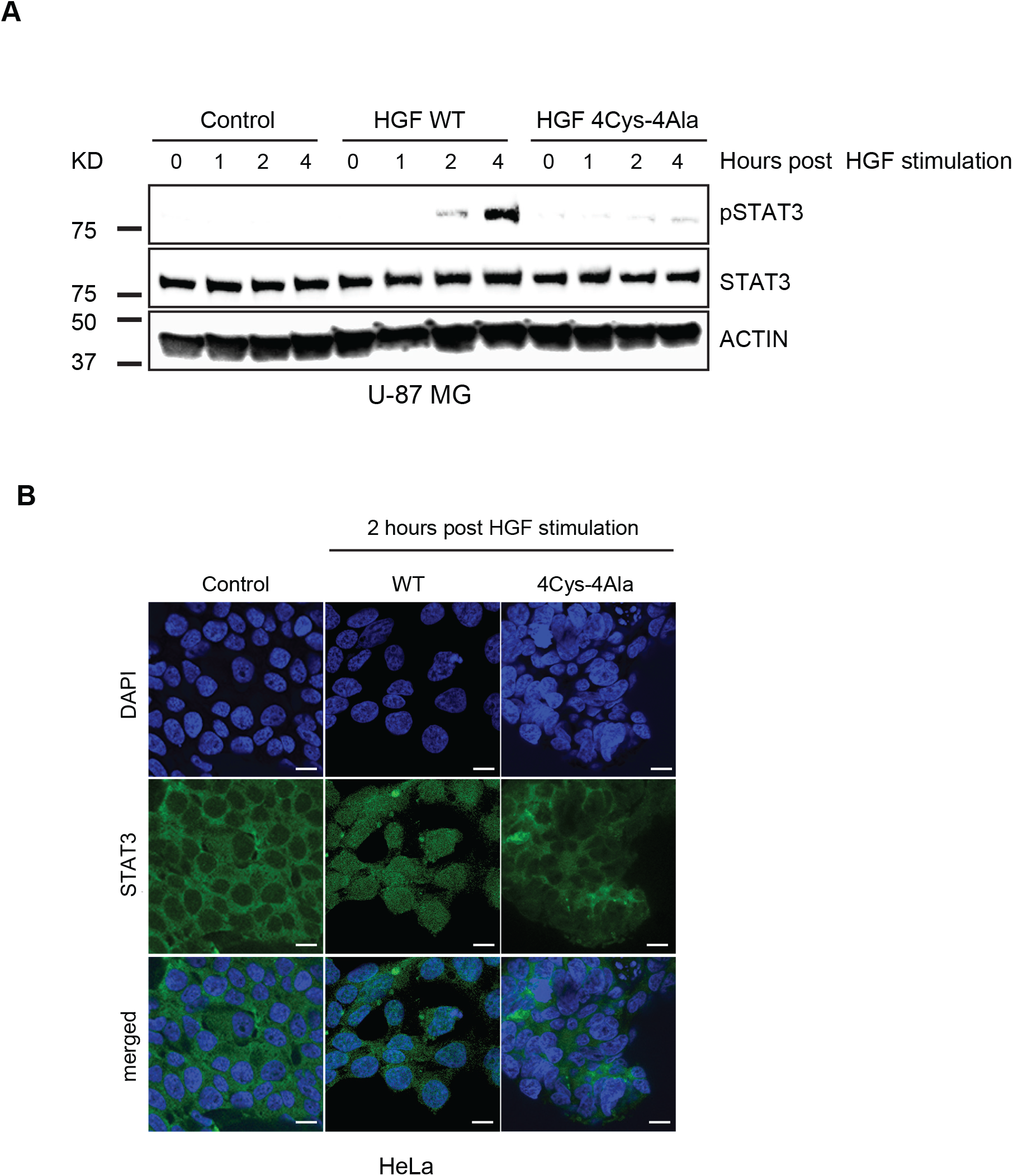
Core cysteines in HGF PAN domain is essential for STAT3 phosphorylation and nuclear translocation. (A) Impaired HGF PAN domain is unable to initiate STAT3 phosphorylation. U-87 MG cells were treated with HGF WT and HGF 4Cys-4Ala where indicated for 1, 2 and 4 h. Cell extracts were prepared and probed for phosphor-STAT3 and total STAT3. (B) STAT3 nuclear localization is suppressed by PAN mutant HGF. HeLa cells were stimulated with HGF WT and HGF 4Cys-4Ala as indicated. Cells were fixed and STAT3 was immunostained with a STAT3-specific antibody. The localization of STAT3 (green) and 4,6-diamidino-2-phenylindole (DAPI) (blue) in U-87 MG cells. Images were visualized using Zeiss LSM 710 at 63x objective. Scale bars represent 20 μm. The images shown are representative of three independent biological replicates (an average of 100 cells were observed for each condition per replicate).

### PAN mutations downregulate HGF/c-MET-dependent cell proliferation and alters expression of genes essential for diverse cellular responses

Dysregulated expression of HGF/c-MET acts as a catalyst in many cancers, with overexpression of HGF often leading to aberrant cell proliferation and extracellular matrix invasion (*14*). Direct evidence has been established connecting a primary role for HGF with increased expression of MMP9, which is crucial for angiogenesis (*30*). HGF facilitates MMP9 expression via the PI3K/AKT and p38 mitogen-activated protein kinases (MAPK) axis. Given the reported role of HGF in modulating expression of MMP9, we evaluated the impact of mutated cysteine residues on its transcriptional response. To evaluate the role of PAN domain mutation on the pattern of the signature gene expression as well as cell proliferation, both 293T and U-87 MG cells were stimulated with wild-type HGF or 4Cys-4Ala HGF for 24 hours following serum starvation. Cell viabilities, as assessed by the MTT assay, were notably reduced by treatment with HGF 4Cys-4Ala, whereas wild-type HGF could overcome serum starvation-mediated growth inhibition in both cell types (Fig. S5A). 293T cells transiently transfected with Flag-WT HGF showed a significant increase in cell proliferation over time compared to cells transfected with Flag-HGF 4Cys-4Ala (Fig. S6A). Quantitative real-time PCR (qRT-PCR) was performed to determine whether MMP9 expression was similarly impacted. mRNA levels of MMP9 were markedly decreased in HGF 4Cys-4Ala-stiumlated 293T and U-87 MG cells compared to the wild type in both cell types (Fig. S5B). Similarly, we observed a significant decline in MET mRNA expression in both cell types stimulated with HGF 4Cys-4Ala (Fig. S5B). Increased c-MET expression might be a crucial determinant for the overall balance of the HGF/c-MET cascade in cells and could be a trigger for the malignant transformation of normal cells. These data suggests that induction of the MMP9 receptor is correlated with overall MET expression in an HGF-dependent manner and could be regulated by minimally mutating the four core cysteines in the PAN domain of HGF.

Based on the apparent PAN domain-dependent transcriptional modulation of MMP9 expression by HGF, we characterized the transcriptional responses of additional downstream genes in the HGF/c-MET signaling pathway. Total RNA was extracted from 293T cells following a 24-hour post treatment with wild-type HGF or 4Cys-4Ala HGF. Based on this analysis, we identified significant differences in the expression of genes previously implicated in MET signaling, cell cycle regulation, and cancer-related processes (Fig. 4; Fig. S6B). Specifically, we observed a significant reduction in the expression levels of the downstream targets of the MET signaling cascade, ETV1, ETV4 and ETV5, in cells stimulated with HGF 4Cys-4Ala compared to wild type HGF. These transcription factors are members of the polyoma enhancer activator 3 (PEA3) subgroup of the E–twenty six (ETS) family and confer resistance to early growth factor response (EGFR) targeted therapy in lung cancer (*31*). Some known target genes of PEA3 are matrix metalloproteinase-2 (MMP2), matrix metalloproteinase-7 (MMP7) and MMP9, which are well recognized for their role in the invasiveness of cancer (*32*). Mutating the core cysteine residues in the PAN domain led to reduced expression of EGR1, as well as the metalloproteinases, A disintegrin and metalloprotease domain-containing protein 9 and 10 (ADAM9 and ADAM10), all of which were previously linked to increased adhesion and cancer progression including, hepatocellular carcinoma, and triple negative breast cancer via the AKT/NF-κB axis (*33–35*). Hyperactivation of focal adhesion kinase (FAK) and PI3K/receptor for activated C kinase 1 (RAC1) are responsible for metastasis progression and mutations of the core cysteines in the HGF PAN domain also resulted in their reduced expression. Apart from these examples, relative expression of proteins involved in DNA damage repair (double-strand-break repair protein or RAD21), chromosomal integrity and cohesion (structural maintenance of chromosomes protein 1A or SMC1A), cell cycle progression (kinesin family member 2A or KIF2A), posttranslational modification, and RNA splicing (protein arginine N-methyltransferase 5 or PRTM5) were also reduced. (Fig. 4) (*36–39*). Furthermore, this analysis revealed well-known cancer biomarkers whose expression levels were affected by mutating the core cysteine residues. These biomarkers included calponin-3 or CNN3 (marker for colorectal cancer), dolichyl-diphosphooligosaccharide-protein glycosyltransferase subunit 2 or RPN2 (which inhibits autophagy and upregulates MMP9 expression), branched chain amino acid transaminase 1 or BCAT1 (which promotes hepatocellular carcinoma and chemoresistance), ski interacting protein or SKIP, epithelial cell transforming sequence 2 oncogene or ECT2 (overexpressed in different cancers including non-small cell lung cancer), and elastin microfibril interfacer 2 or EMILIN2 (which promotes angiogenesis and inflammation) (*40–45*).

**Fig. 4:**
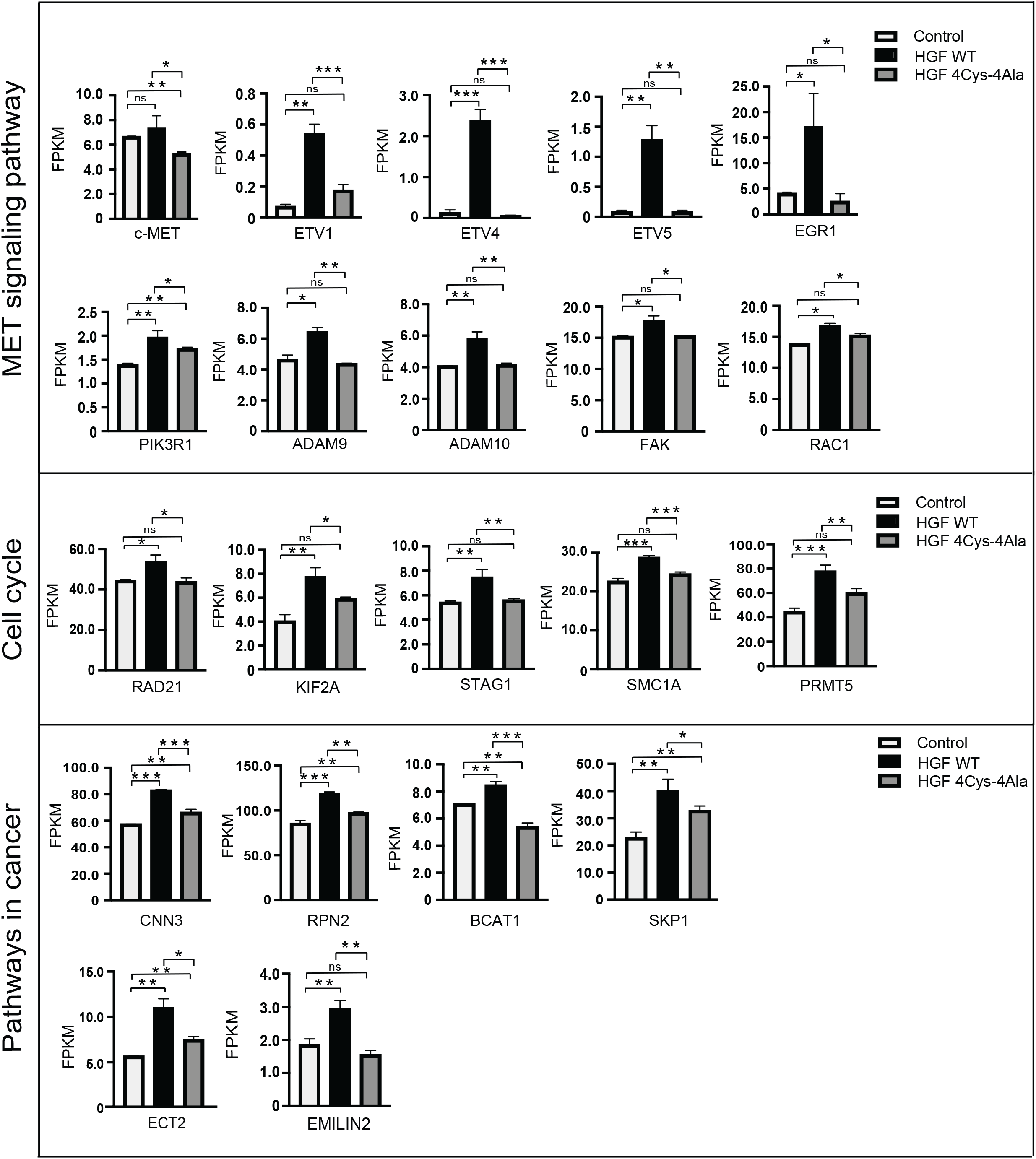
Transcriptome analysis post HGF stimulation in 293T cells. Differential expression analysis by RNA seq in 293T cells following HGF WT and HGF 4Cys-4Ala treatment as indicated confirms that core cysteines in HGF PAN domain are necessary for the expression of a wide range of genes. Responsive genes were normalized to FPKM value for non-treated cells and then normalized to HGF WT treated cells. Data represents the average of three independent biological replicates and *p < 0.05, **p < 0.005 and ***p < 0.0005 were calculated with a student’s t-test.

## Discussion

In summary, this study provides evidence that the PAN domain of HGF provides the functional catalytic core and plays a crucial role in c-MET binding. Further, GO enrichment analysis predicted the strong association with cellular recognition and signaling processes of all the PAN domain containing proteins, suggesting a role in immune signaling. To support these observations, we demonstrated that mutating core cysteine residues in the HGF PAN domain results in a cascade of negative regulation of HGF/c-MET signaling, which starts with retarded HGF degradation, impaired c-MET interaction, disrupted perinuclear localization for both HGF and its receptor c-MET, and is followed by a subsequent lack of phosphorylation for c-MET and its downstream targets AKT, ERK and STAT3 and c-MET ubiquitination.

This disruption of upstream events was confirmed by the lack of transcriptional activation of markers genes associated with cell migration and invasion including ADAM9/10, EGR1, ETV1, ETV4, ETVE5 and MMP9 (Fig. S6B). Targeting the PAN domain in HGF could act as a multi-pathway target as it entirely shuts down the c-MET signaling cascade and downstream expression of relevant proteins known for their role in cancer prognosis and other diseases (Fig. S7). Taken together, the PAN domain and its core cysteine residues are essential for HGF/c-MET signaling in human cells. The observed enhanced stability of HGF with mutated cysteine residues is due to the lack of its ability to bind with c-MET. These results are consistent with predictions based on GO enrichment analysis in which the GO term *proteolysis* was highly enriched in the organismal categories Alveolata, Amoebozoa, bacteria, opisthokonta, stramenopilles and viruses. This finding suggests that the PAN domain may have been co-opted into different proteins along these lineages to facilitate similar functions in the proteolytic processing of proteins involved in immune signaling.

## Materials and Methods

### PAN domain sequence alignment

Proteins with PAN domains were identified from Uniprot and were selected from 14 model organisms according to Bateman et al (*46*). The PAN domain coordinates from UniProt were used to extract the corresponding sequences from the full-length proteins, which were then aligned with MAFFT linsi(*47*). The alignment was visualized with Geneious^41^ and a phylogenetic tree accompanying the alignment was constructed via the neighbor-joining method with Geneious (Table S3).

### GO enrichment of 14 PAN domain categories

GO terms of each category were extracted from InterPro2GO database from InterPro (*48*). GO enrichment analysis for each category against all PAN domain genes was performed by Fisher’s exact test via the TopGO package (*49*). Only the biological process GO category was used for GO enrichment.

### Mammalian cell culture, transfection and drug treatment

HeLa, HEK 293T and Glioblastoma U87 cells were obtained from ATCC and maintained in a humidified atmosphere at 5% CO_2_in Dulbecco’s Modified Eagle’s (DMEM) complete medium (Corning) supplemented with 10% fetal bovine serum (FBS; Seradigm) in 37°C. Plasmid transfections were done with TransIT-LT1 (Mirus Bio) per the manufacturer’s instructions.

### Plasmids and Recombinant Proteins

Flag-HGF (GeneBankTM accession number NM_000601.5) and various mutants of Flag-HGF, cloned into pcDNA 3.1 were obtained from GenScript. Purified HGF protein and HGF 4Cys-4Ala protein were synthesized in GenScript. c-MET-C-GFPSpark® Clone was obtained from SinoBiological.

### Immunofluorescence and confocal microscopy

HeLa cells were seeded on coverslips in 24 well plates. Where indicated, cells were transfected with Flag-HGF WT and different PAN domain mutants for 36 hours, followed by fixation in 4% paraformaldehyde. Next, the cells were permeabilized with 0.5% Triton X-100 in PBS, washed and then blocked for 30 minutes at room temperature with 5% BSA in PBS. Cells were incubated with primary antibodies in 5% BSA in 1X-PBS with 0.5% Triton X-100 for 1 hour at room temperature. After washing the cells were incubated with appropriate secondary antibodies in 5% BSA in PBST for 30 minutes at room temperature. DNA was counterstained with 1 μg/mL Hoechst 33342 and mounted with Fluorimount G (Southern Biotech). Cells were imaged using a Zeiss LSM 710 confocal microscope. For MET and STAT3 localization assays, cells were serum starved for 24 hours before the stimulations. Cells were then stimulated with 100 ng mL^−1^ of WT HGF and or 4Cys-4Ala HGF purified proteins for the indicated amount of time at 37°C and treated as described above.

### Antibodies

The following commercial antibodies and the indicated concentrations were used in this study. HA antibody (HA.C5 #18181; 1:1000) was purchased from Abcam. Flag (#2368S; 1:1000), (Met (clone 25H2 #3127; 1:1000), Phospho-Met (Tyr 1234/1235) (clone D26 # 3077; 1:1000), Stat3 (clone 124H6 #9139; 1:1000), Phospho-Stat3 (Tyr 705) (clone D3H7 #9145; 1:1000), Akt (pan) (clone 40D4 #2920; 1:1000), Phospho Akt (Thr 308) (clone 244F9 #4056; 1:1000), p44/42 MAPK (Erk1/2) (#9102; 1:1000), Ubiquitin (clone E4I2J) (#43124; 1:1000), Met (clone D1C2) (#8198; 1:1000) and Phospho-p44/42 MAPK (Erk1/2) (Thr202/Tyr204) (clone D13.14.4E) (#4370; 1:1000) were purchased from Cell signaling. M2 anti Flag Mouse antibody (#SLBT7654; 1:5000) and Actin (#087M4850; 1:10,000) were purchased from Sigma. HA (#902302; 1:1000) antibody was purchased from Biolegend. Invitrogen GFP (clone A-11122) (1:1000) was purchased from Thermo Fisher Scientific. Met Antibody (clone D-4) (#sc-514148; 1:500) was purchased from Santa Cruz Biotechnology. Chemiluminescence detection was performed according to the manufacturer’s instructions (Amersham ECL Western Blotting Detection Reagent kit) followed by exposure using Chemidoc gel-documentation system (BioRad). For imaging using Li-Cor, secondary antibodies for western blotting were purchased from LI-COR Biosciences.

### Western Blotting and immunoprecipitation

For in vivo stimulation experiments, cells were grown for 36 hours and then stimulated with HGF WT and HGF 4Cys-4Ala where indicated (100 ng mL^−1^), washed with PBS, and lysed. Briefly, cell extracts were generated on ice in EBC buffer, 50mM Tris (pH 8.0), 120mM NaCl, 0.5% NP40, 1mM DTT, and protease and phosphatase inhibitors tablets (Thermo Fisher Scientific). Extracted proteins were quantified using the PierceTM BCA Protein assay kit (Thermo Fisher). Proteins were separated by SDS acrylamide gel electrophoresis and transferred to IMMOBILON-FL 26 PVDF membrane (Millipore) probed with the indicated antibodies and visualized either by chemiluminescence (according to the manufacturer’s instructions) or using a LiCor Odyssey infrared imaging system.

For immunoprecipitation, endogenous c-MET was immunoprecipitated on MET antibody-bound beads (Protein G from Thermo Fisher) and Flag-tagged HGF WT and HGF 4Cys-4Ala were *in-vitro* translated (T_N_T quick coupled Transcription/Translation system, Promega) and were incubated with the bead bound c-MET for 4h at 4 °C. Beads were then washed and proteins resolved by SDS-PAGE and analyzed by western blotting as above.

### In vivo ubiquitination

293T cells were transfected with the construct encoding c-MET-C-GFPSpark. Cells were stimulated with either HGF WT or HGF 4Cys-4Ala as indicated following serum starvation. Cells were collected at indicated time points and washed with ice-cold PBS, lysed in ice-cold buffer containing 10 mM Tris-HCl (pH 8), 150 mM NaCl, 0.1% SDS, 20mM NEM (*N*-ethylmaleimide) supplemented with protease and phosphatase inhibitors tablets (Thermo Fisher Scientific). The lysates were cleared by centrifugation at 10,000 × *g* at 4°C for 20 min, followed by preclearance using protein G-beads (protein G from Thermo Fisher). 500 μg of each precleared lysate was incubated with anti-MET antibody and protein-G beads for overnight at 4°C with rotation. The samples were then washed three times with the lysis buffer and eluted with SDS-gel loading buffer (with reducing agent added). Proteins were resolved by SDS-PAGE and immunoblotted with the indicated antibodies.

### Cell Proliferation Assay

3-(4, 5-dimethylthiazol-2-yl)-2, 5-diphenyl tetrazolium bromide (MTT) assay was used to determine cell viability. HEK293T and U-87-MG cells were plated in 96-well plates with 500 cells per well in triplicate in serum-free medium for 24 hours prior to HGF stimulations. 100 ng mL^−1^ HGF WT, HGF 4Cys-4Ala, or both were added to the cells and incubated for 24 hours prior to the addition of MTT solution (Abcam, Inc # ab211091) and cell viability was measured according to the manufacturer’s instruction. The assays were performed in triplicate and the experiment was repeated three times. Data were expressed as the mean ± SD. Statistical analyses were performed. *P < 0.05 was considered to indicate a statistically significant difference.

### Quantitative Real-Time PCR and RNA-Seq Analysis

Total RNA was extracted from glioblastoma U87 and HEK293T cell line 24 hours post HGF stimulation using TRIzol reagent (Invitrogen), and reverse transcription was performed using the SuperScript II RT kit (Integrative DNA Technologies) with total RNA (1 μg) according to the manufacturer’s instructions. The MET and MMP9 mRNA expression levels were detected by conventional RT-PCR with Taq DNA Polymerase, Recombinant (Invitrogen, no. 10342-020). Glyceraldehyde-3-phosphate dehydrogenase (GAPDH) was used as the internal control. The specific primers for MET, MMP9 and GAPDH were designed with Primer Premier software. The primers used were MET, forward: 5’-TTAAAGGAGACCTCACCATGTAATC-3’ and reverse: 5’-CCTGATCGAGAAACCACAACCT −3’; MMP9, forward: 5’-GATCCAAAACTACTCGGAAGACTTG-3’ and reverse: 5’-GAAGGCGCGGGCAAA-3’ and GAPDH, forward: 5’-TTGCCATCAATGACCCCTTCA-3’ and reverse: 5’-CGCCCCACTTGATTTTGGA-3’. The PCR reaction was performed according to the manufacturer’s instructions. The PCR conditions were as follows: amplification reaction protocol was performed for 35 cycles consisting of 30 s at 94°C (denaturation), annealing 30 s at 45°C and extension 30 s at 72°C.

For RNA-seq analysis, Total RNA was extracted from the HEK293T cell line 24 h post HGF stimulation using TRIzol reagent (Invitrogen) according to the manufacturer’s instructions. Quantification and quality control of isolated RNA was performed by measuring absorbance at 260 nm and 280 nm on a NANODROP ONEC spectrophotometer (Thermo Scientific, USA). The RNA-seq run was performed with four biological replicates. Library prep and sequencing was performed by BGI using the DNBSEQ-G400 platform which generated 100 bp paired-end reads. The raw RNA-seq reads have been deposited at NCBI under BioProject ID PRJNA718097. Clean reads were aligned to the human reference genome GRCh38. Reads were mapped with bowtie2 v2.2.5. (*50*) Expression levels for RNAs were calculated using fragments per kilobase per million reads (FPKM) values with RSEM v1.2.8. (*51*) Differential expression analysis was performed with DESeq2 and genes with an adjusted p-value less than 0.05 were considered differentially expressed (*52*).

### Molecular Dynamics Simulations

MD simulations were initiated from the top five models generated with ColabFold (*20, 53*) of the WT and 4Cys-4Ala mutant PAN domain. The program *tleap* from AmberTools20 (*54*) was used to prepare the parameter and coordinate files for each structure. The ff14SB force field (*55*) and TIP3P water model (*56*) were used to describe the protein and solvent, respectively. Energy minimization was performed using *sander* from AmberTools20. At least a 12 Å solvent buffer between the protein and the periodic images. Sodium and chloride ions were added to neutralize charge and maintain a 0.10 M ion concentration. The simulations were performed with OpenMM version 7.5.1 (*57*)) on the Cuda platform (version 11.0.3) using Python 3.8.0. ParmEd was used to incorporate the force field parameters into the OpenMM platform (*58*). The Langevin integrator and Monte Carlo barostat were used to maintain the systems at 300 K and 1 bar, respectively. Direct non-bonded interactions were calculated up to a 12-Å distance cutoff. All bonds involving hydrogen atom were constrained to their equilibrium values. The particle mesh Ewald method was used to compute long-range Coulombic interactions (*59*). A 2-fs integration time step was used with energies and positions written every 2 ps.

### Simulation Analysis

Analyses of MD trajectories were performed using Python 3.8.0 and the MDAnalysis version 1.0.1 (*60, 61*). Matplotlib was used to plot the data.

### Statistical Analysis

Statistical analyses were performed on individual experiments, as indicated, with GraphPad Prism 8 Software using an unpaired t-test, equal variance, for comparisons between two groups. A P-value of *P < 0.05 was considered statistically significant.

## Acknowledgments

Bioinformatics analyses of PAN domain distribution and functional inference was supported by the United States Department of Energy, Office of Science, Early Career Research Program under the Biological and Environmental Research office. Biochemical, immunofluorescence and transcriptome analyses in human cell lines, and protein modeling and simulation were supported by the Lab Directed Research Development program at Oak Ridge National Laboratory (ORNL). Part of this research used resources at the Oak Ridge Leadership Computing Facility (OLCF) and the Compute and Data Environment for Science (CADES) at ORNL, which is managed by UT-Battelle, LLC for the U.S. Department of Energy under Contract Number DE-AC05-00OR22725.

## Competing interests

The authors declare that they have no competing interests.

## Notes

### Competing Interest Statement

The authors have declared no competing interest.

### Summary of Updates

Supplemental materials

