## Supplemental figures 1 to 7 for "Core cysteine residues in the PAN domain are critical for HGF/c-MET signaling": supplemental.pdf

**This PDF file includes:**

Figs. S1 to S7

Tables S1 to S3

Legends for Figs.S1 to S7

Legends for Tables S1 to S3

**Fig. S1. The HGF PAN domain is comprised of a core of four conserved cysteine residues.** (A) Multiple alignment of the sequences of PAN domain of representative proteins from different organisms highlights the position of four conserved cysteines. (B) Schematic diagram of amino acid sequence represents HGF PAN domain along with the four marked conserved cysteines (Cys70, Cys74, Cys84 and Cys96) and the subsequent mutant version where conserved cysteines were mutated to alanine (Ala70, Ala74, Ala84 and Ala96). Computational secondary structure prediction (Chimera) for HGF PAN domain identifies the residues forming β-strands and α-helix in PAN domain.

**Fig. S2.** The 3D conformation of four reported mutation sites located in the PAN domain of HGF.

**Fig. S3. Root-mean-square deviation (RMSD) time evolution plots from WT and 4Cys-4Ala MD simulations.** RMSD profiles and histograms for (A) Wild-type and (B) 4Cys-4Ala mutant. For both datasets, the reference structure was the initial coordinates of the top WT model. RMSDs were calculated for all heavy atoms after alignment of all Ca atoms.

**Fig. S4. Representative structures of the WT and 4Cys-4Ala PAN domain from MD simulations.** (Left) Ca alignment of both structures to the initial coordinates of the top WT model. (Right) Close-up of the mutation positions for the two systems. In the absence of the disulfide pairs, the 4Cys-4Ala mutant maintained near equivalent positioning of the mutated residues.

**Fig. S5. HGF stimulates cell proliferation and MMP9 expression in PAN domain dependent manner.** (A) Mutations on the conserved cysteines in HGF PAN domain downregulate cell proliferation. 293T and U-87 MG cells were plated in 96-well plates in serum-free medium for 24 hours. HGF WT and HGF 4Cys-4Ala were added to the cells where indicated and plates were incubated for 24 h. MTT reagents were added, and the absorbance was read at 492 nm. A significant increase in the proliferation for both cells were observed when treated with HGF WT (*$p<0.05$). (B) Quantitative PCR analysis for MMP9 and MET mRNA expression. 293T and U-87 MG cells were serum starved before HGF treatment. Treatment was given for 24 hours and MMP9 and MET mRNA expressions were detected by conventional RT PCR. The graph represents the relative mRNA expression normalized to GAPDH control. Data (A and B) are represented as mean +/- SD, n=3 independent biological replicates and *$p<0.05$ (Student's t test).

**Fig. S6.** (A) MTT cell proliferation assay. 293T cells were transfected with Flag-HGF WT and Flag-HGF 4Cys-4Ala. 24 hr post transfection cells were splitted and re-plated in 96-well plates in serum-free medium for 24 hours. Plates were incubated for indicated time. MTT reagents were added, and the absorbance was read at 492 nm. A significant increase in the proliferation for cells was observed when transfected with HGF WT (*$p<0.05$). (B) Heatmap of selected genes. Heat maps displaying pattern of expression for the candidates involved in Met signaling, cell cycle and invasion. Genes are depicted based on their expression ratios across three RNA seq comparison. Colors range from bright green (upregulation; $\log_2$ ratio over control) to bright red (downregulation; $\log_2$ ratio over control).

**Fig. S7.** A proposed model showing how conserved cysteines in HGF PAN domain determines the fate of MET signaling cascade. PAN domain provides the catalytic core to HGF for c-MET interaction leading towards initiation of entire downstream HGF/c-MET axis. Alteration in conserved cysteine residues blocks the activation of several transcription factors and effector molecules which otherwise stimulate cell migration, cell invasion, proliferation, and cell motility. Figure created with BioRender.com.

**Table S1.** Species of origin, protein identification number (ID) and functional annotation of PAN domain-containing proteins.

**Table S2.** GO terms exhibiting significant enrichment among 28,300 PAN domain-containing proteins.

**Table S3.** Proteins with PAN domain in alignment.

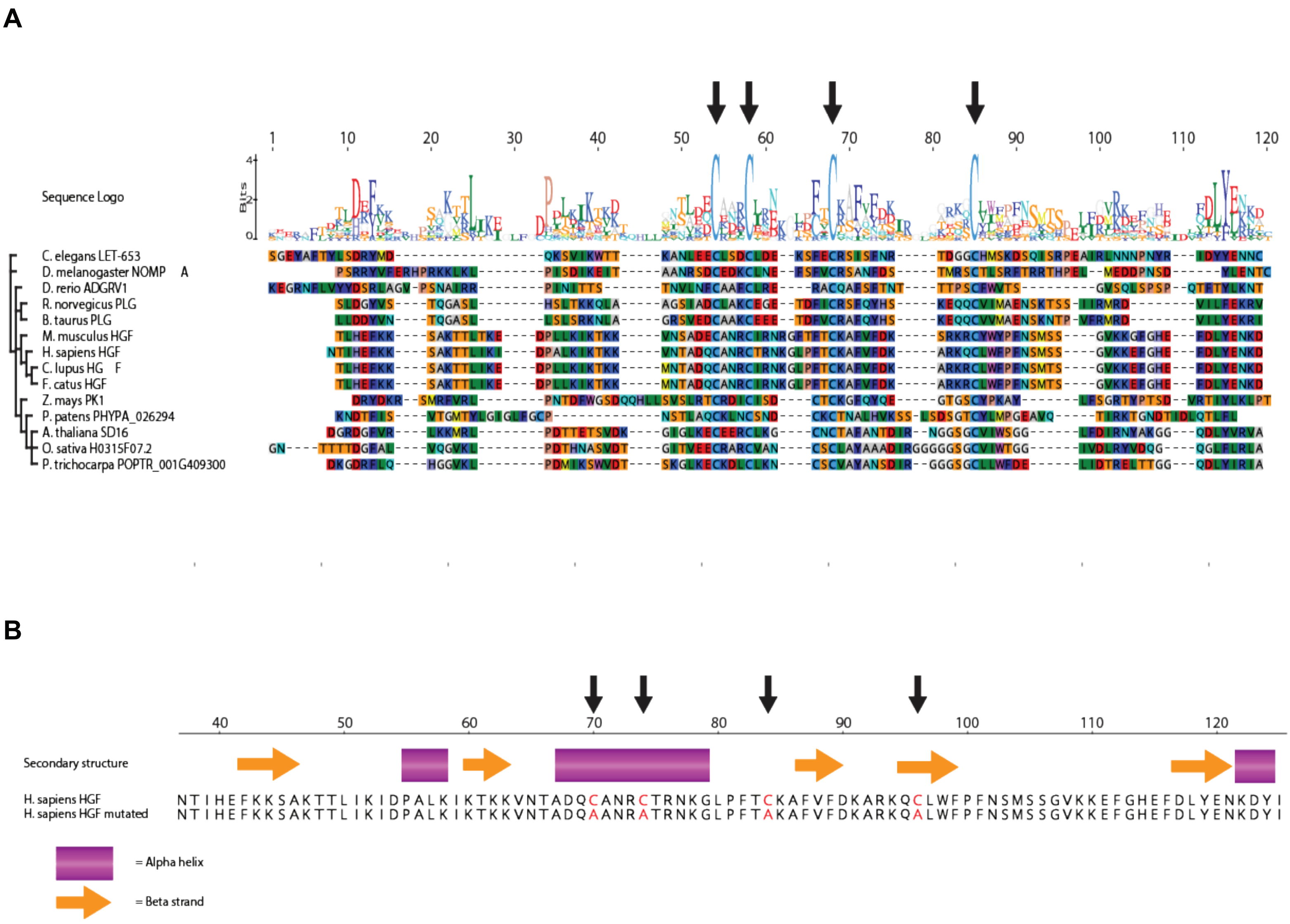
A
Sequence Logo
Bits
C. elegans LET-653
D. melanogaster NOMPA
D. rerio ADGRV1
R. norvegicus PLG
B. taurus PLG
M. musculus HGF
H. sapiens HGF
C. lupus HGF
F. catus HGF
Z. mays PK1
P. patens PHYPA_026294
A. thaliana SD16
O. sativa H0315F07.2
P. trichocarpa POPTR_001G409300
B
Secondary structure
H. sapiens HGF NTIHEFKKSAKTTLIKIDPALKIKTKKVNTADQCANRCTRNKGLPFTCKAFVFDKARKQCLWFPFNSMSSGVKKEFGHEFDLYENKDYI
H. sapiens HGF mutated NTIHEFKKSAKTTLIKIDPALKIKTKKVNTADQAANRATRNKGLPFTAKAFVFDKARKQALWFPFNSMSSGVKKEFGHEFDLYENKDYI
= Alpha helix
= Beta strand

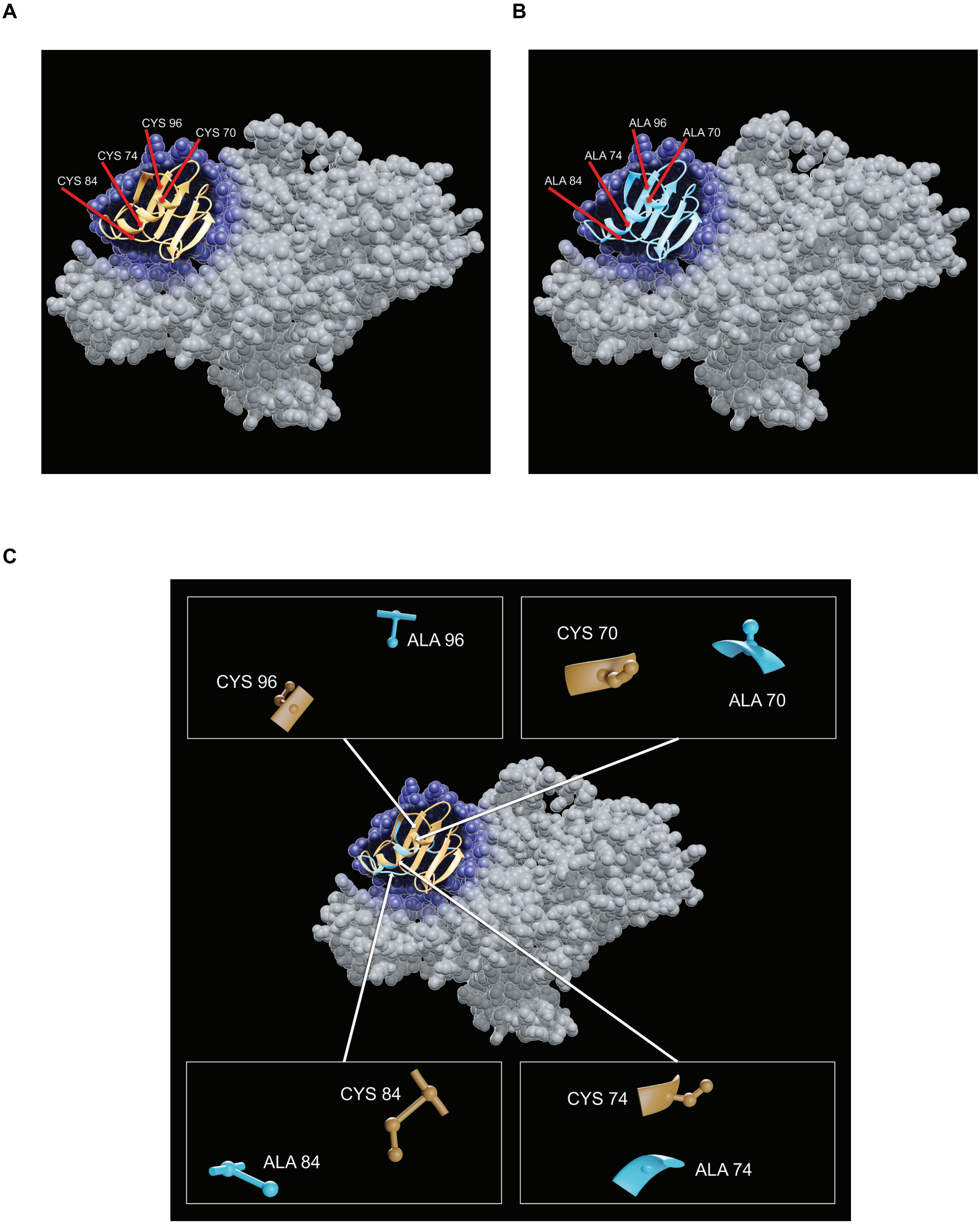
A
CYS 96
CYS 70
CYS 74
CYS 84
B
ALA 96
ALA 70
ALA 74
ALA 84
C
ALA 96
CYS 96
CYS 70
ALA 70
CYS 84
ALA 84
CYS 74
ALA 74

Pal et al., Supplemental Figure 3

A

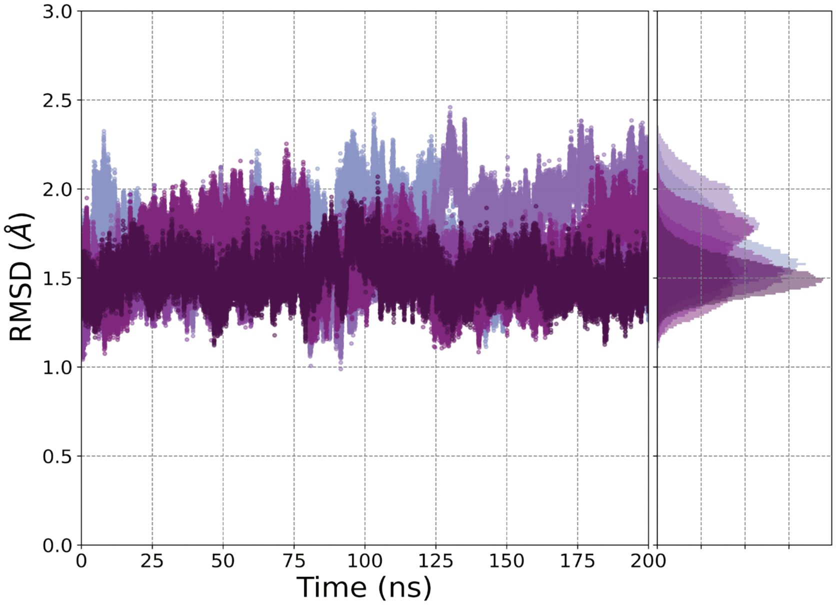


B

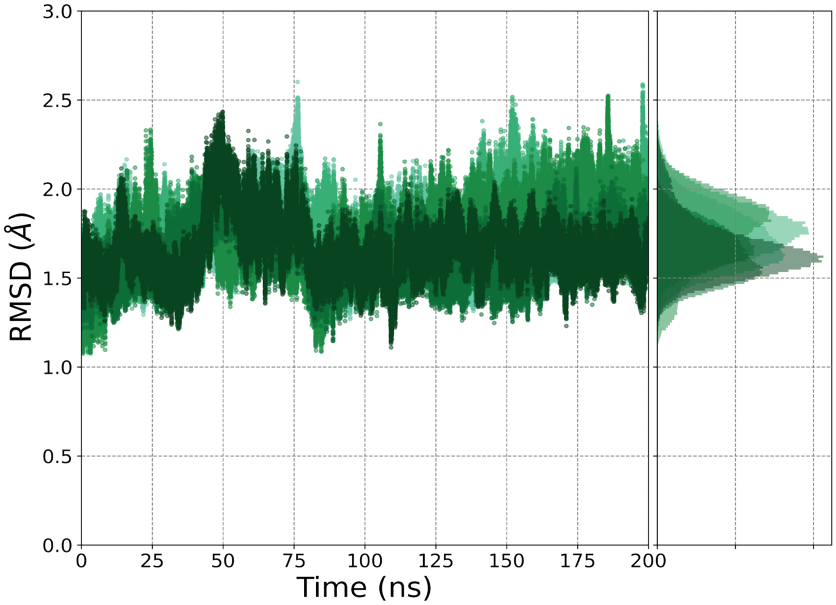

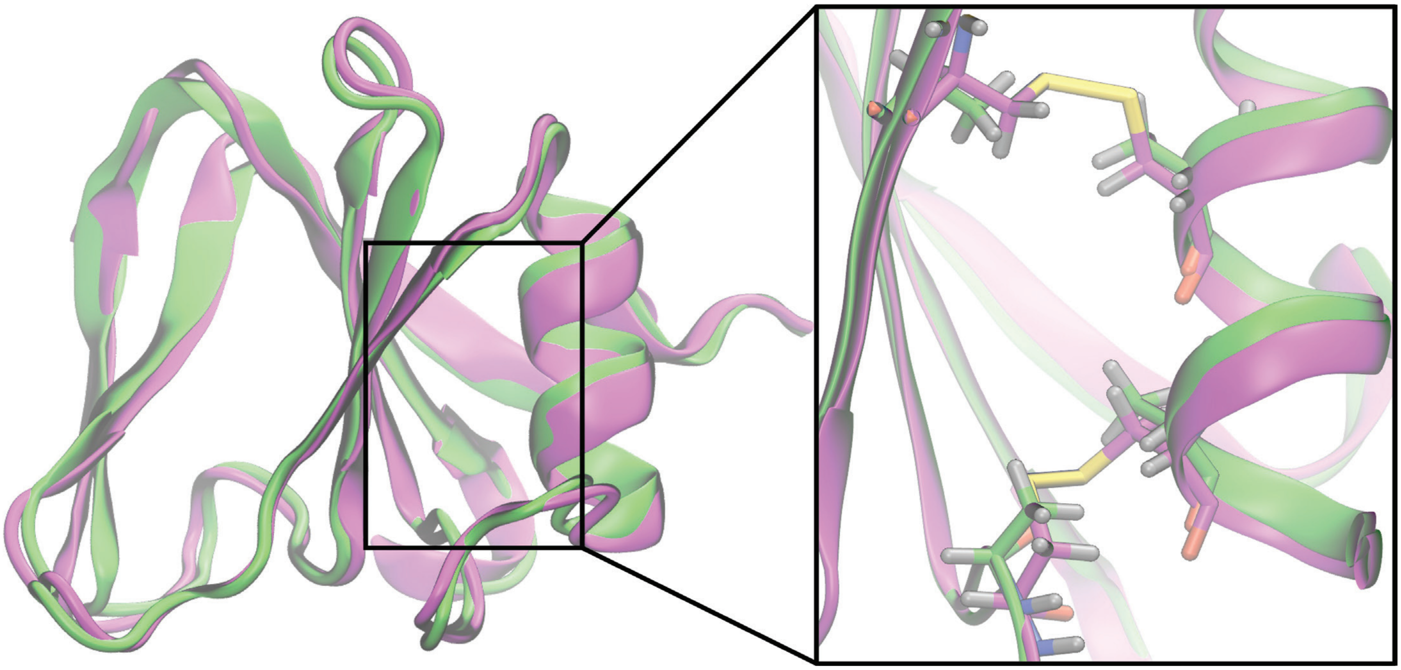

**A**

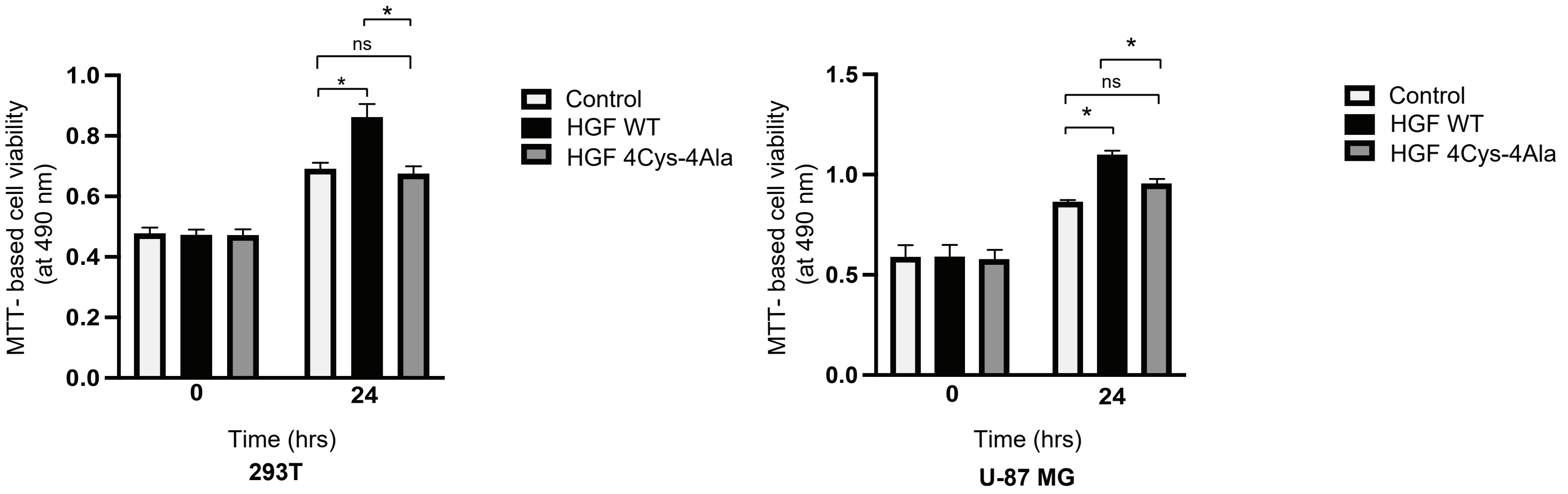

MTT- based cell viability (at 490 nm)
Time (hrs)
293T
U-87 MG
Control
HGF WT
HGF 4Cys-4Ala
ns
*


**B**

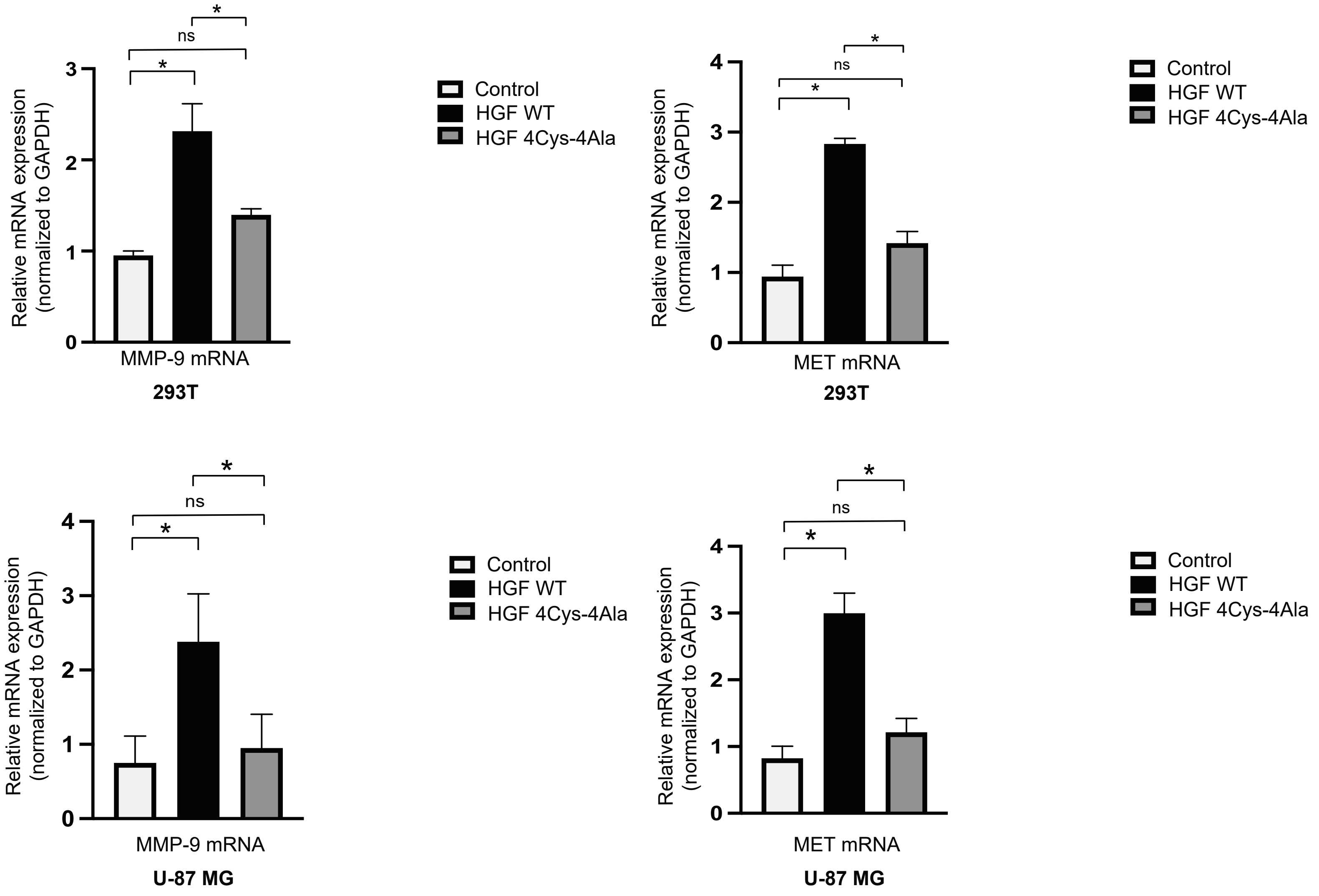

Relative mRNA expression (normalized to GAPDH)
MMP-9 mRNA
MET mRNA
293T
U-87 MG
Control
HGF WT
HGF 4Cys-4Ala
ns
*

A

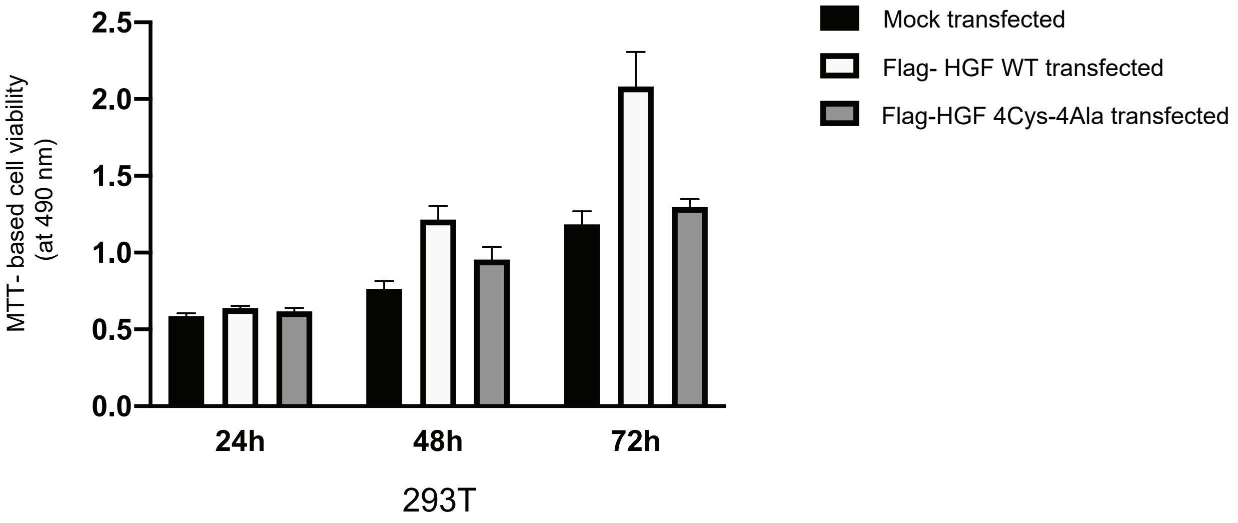


B

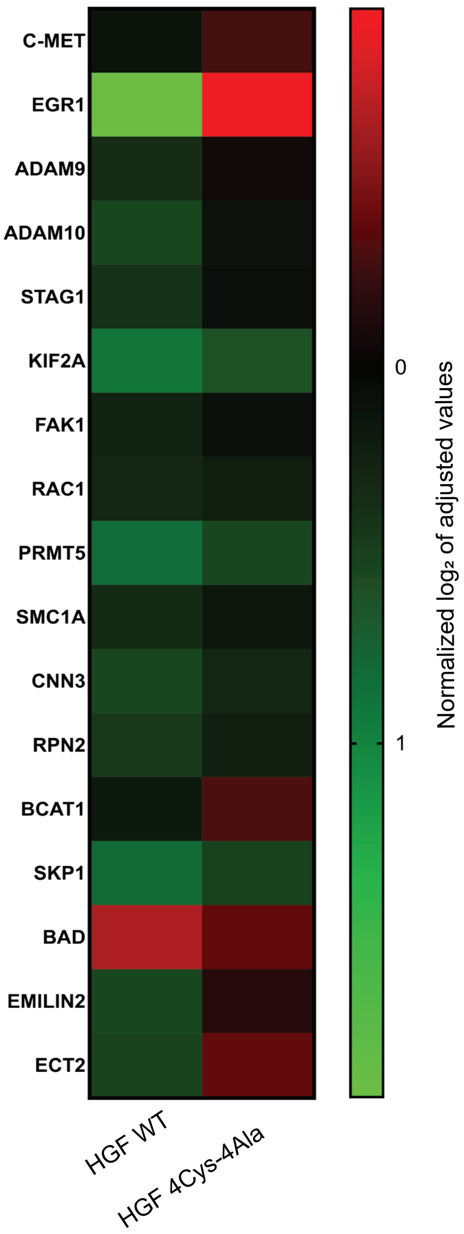


Selected genes involve in MET signaling and cell cycle

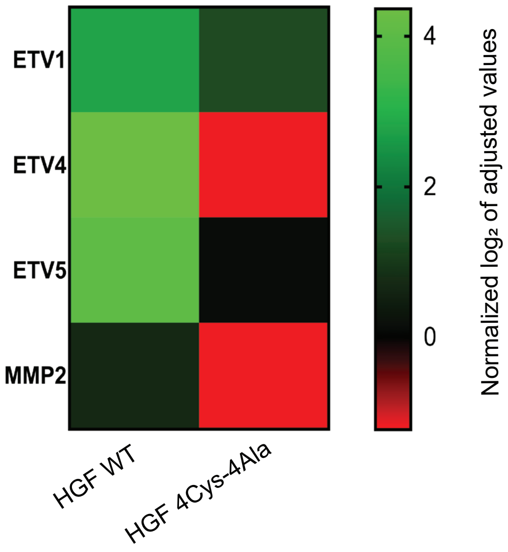


Selected invasive genes

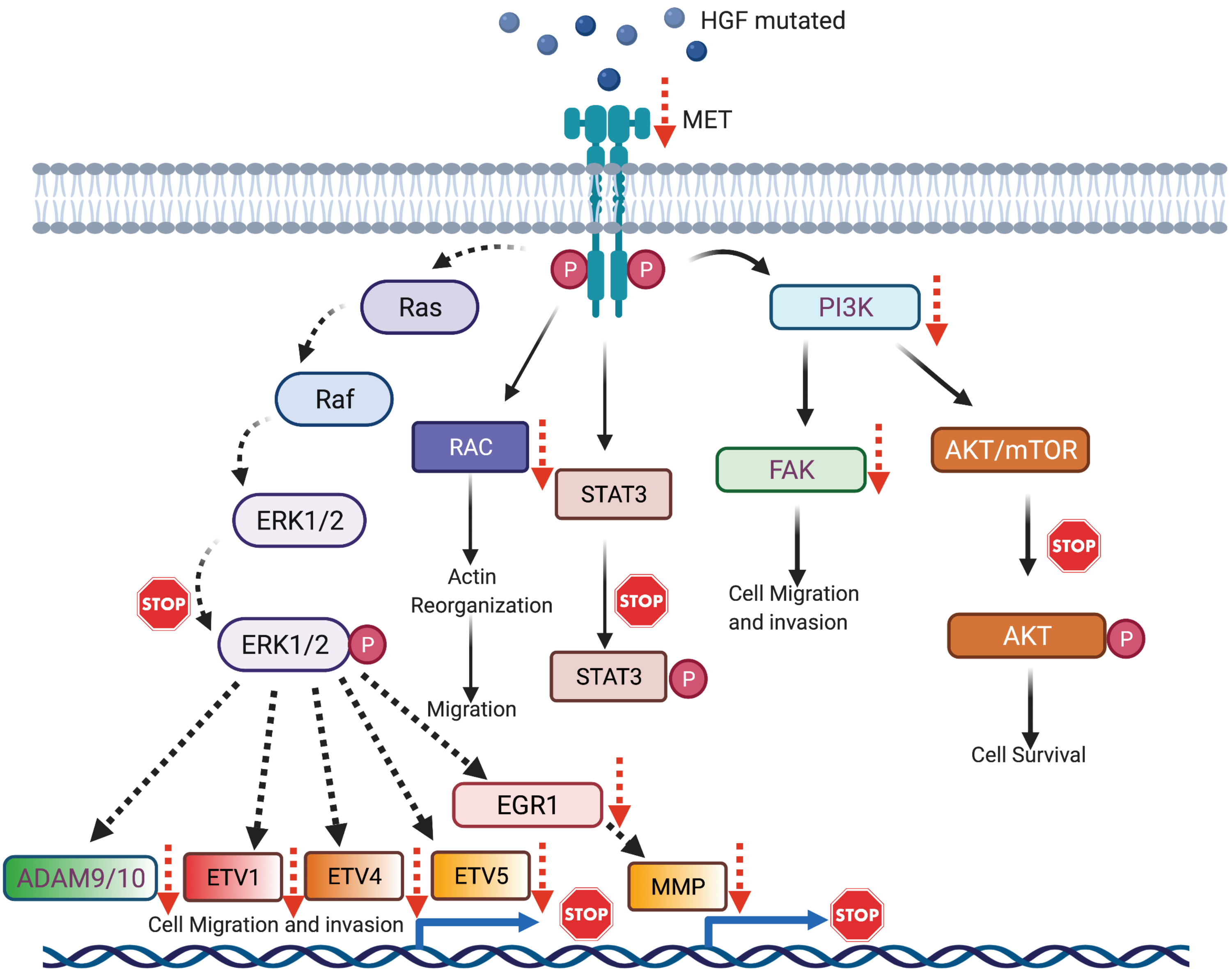
HGF mutated
MET
P
P
Ras
Raf
ERK1/2
STOP
ERK1/2
P
RAC
Actin
Reorganization
Migration
STAT3
STOP
STAT3
P
PI3K
FAK
Cell Migration
and invasion
AKT/mTOR
STOP
AKT
P
Cell Survival
EGR1
ADAM9/10
ETV1
ETV4
ETV5
MMP
STOP
STOP
Cell Migration and invasion

**Table S1: Species of origin, prc**

| Species | ID |
|---|---|
| Abdopus acul | R4G2C1 |
| Abdopus acul | R4G7D0 |
| Acanthamoel | G8EBH2 |
| Acanthamoel | G8EBN0 |
| Acanthamoel | G8EDK5 |
| Acanthamoel | A0A3T1CXD4 |
| Acanthamoel | A0A3T1CXP1 |
| Acanthamoel | A0A1E1EXB1 |
| Acanthamoel | A0A1E1ETK2 |
| Acanthamoel | A0A1E1EUC2 |
| Acanthamoel | L8GVM1 |
| Acanthamoel | L8GRF2 |
| Acanthamoel | L8GX08 |
| Acanthamoel | L8GY48 |
| Acanthamoel | L8H5C9 |
| Acanthamoel | L8GYA1 |
| Acanthamoel | J3E810 |
| Acanthamoel | J2YWZ6 |
| Acanthamoel | J3IYU5 |
| Acanthamoel | A0A2L2DLV0 |
| Acanthamoel | A0A2L2DIZ7 |
| Acanthamoel | A0A0G2Y717 |
| Acanthamoel | E3VZX8 |
| Acanthamoel | E3VYF9 |
| Acanthamoel | F8V6R0 |
| Acanthamoel | A0A0G2Y378 |
| Acanthamoel | A0A0G2Y8U4 |
| Acanthamoel | A0A2L2DJ09 |
| Acanthamoel | Q5UPR5 |
| Acanthamoel | Q5UNW8 |
| Acanthamoel | Q5UQG0 |
| Acanthamoel | A0A2L2DM38 |
| Acanthamoel | A0A0G2Y3S5 |
| Acanthamoel | A0A0G2YB83 |
| Acanthamoel | A0A0G2Y7I9 |
| Acanthamoel | L7RC65 |
| Acanthamoel | L7RC22 |
| Acanthamoel | L7RCU4 |
| Acanthisitta c | A0A091MSS1 |

| | |
|---|---|
| Acanthisitta c | A0A091MQE1 |
| Acanthochroı | A0A3Q1FYN5 |
| Acanthochroı | A0A3Q1FW77 |
| Acanthochroı | A0A3Q1EDF5 |
| Acanthochroı | A0A3Q1G8B6 |
| Acanthochroı | A0A3Q1F1Z0 |
| Acanthochroı | A0A3Q1I407 |
| Acaromyces i | A0A316YDH8 |
| Acaryochloris | A8ZR36 |
| Achlya hypog | A0A0A7CPG5 |
| Achlya hypog | A0A0A7CP60 |
| Achlya hypog | A0A0A7CN07 |
| Achlya hypog | A0A0A7CNR8 |
| Achlya hypog | A0A0A7CN57 |
| Achlya hypog | A0A0A7CP35 |
| Achlya hypog | A0A0A7CMN8 |
| Achlya hypog | A0A0A7CP65 |
| Achlya hypog | A0A0A7CP71 |
| Achlya hypog | A0A0A7CN98 |
| Achlya hypog | A0A0A7CP31 |
| Achlya hypog | A0A1V9ZRA5 |
| Achlya hypog | A0A1V9ZDE5 |
| Achlya hypog | A0A1V9YLY7 |
| Achlya hypog | A0A1V9Z1A6 |
| Achlya hypog | A0A1V9ZKZ6 |
| Achlya hypog | A0A1V9Z451 |
| Achlya hypog | A0A0A7CNG8 |
| Achlya hypog | A0A1V9YJ62 |
| Achlya hypog | A0A0A7CMR5 |
| Achlya hypog | A0A0A7CPD7 |
| Achlya hypog | A0A1V9Z629 |
| Achlya hypog | A0A0A7CPH5 |
| Achlya hypog | A0A1V9YJK3 |
| Achlya hypog | A0A1V9YSE1 |
| Achlya hypog | A0A1V9YDS4 |
| Achlya hypog | A0A1V9YHW9 |
| Achlya hypog | A0A0A7CMJ7 |
| Achlya hypog | A0A0A7CN31 |
| Achlya hypog | A0A1V9ZMC6 |
| Achlya hypog | A0A0A7CNG9 |
| Achlya hypog | A0A1V9YH26 |

| | |
|---|---|
| Achlya hypog | A0A1V9YUS3 |
| Achlya hypog | A0A1V9ZRD0 |
| Achlya hypog | A0A0A7CNY9 |
| Achlya hypog | A0A0A7CM59 |
| Achlya hypog | A0A1V9ZKR0 |
| Achlya hypog | A0A1V9ZUM1 |
| Achlya hypog | A0A1V9YUE0 |
| Achlya hypog | A0A1V9YRF9 |
| Achlya hypog | A0A1V9ZB07 |
| Achlya hypog | A0A0A7CN88 |
| Achlya hypog | A0A1V9YXW4 |
| Achlya hypog | A0A1V9Z9X3 |
| Achlya hypog | A0A0A7CLM9 |
| Achlya hypog | A0A1V9Z933 |
| Achlya hypog | A0A1V9YWJ8 |
| Acidimicrobia | A0A2E4CB18 |
| Acidimicrobia | A0A2E4CBG1 |
| Acidimicrobia | A0A2E4CB13 |
| Acidimicrobia | A0A2E4CBJ3 |
| Acidobacteria | A0A2V9ZGI7 |
| Acidomyces r | A0A150V788 |
| Acidomyces r | A0A150VFP3 |
| Acidovorax ca | A0A3A1WDE8 |
| Acipenser rut | A0A444V1E0 |
| Acipenser rut | A0A444V1E2 |
| Acromyrmex | F4X7R3 |
| Acromyrmex | F4X293 |
| Acromyrmex | F4X6U6 |
| Acromyrmex | F4X815 |
| Acromyrmex | F4WMQ5 |
| Acromyrmex | F4X291 |
| Actibacterium | A0A058ZM17 |
| Actibacterium | A0A037ZPD6 |
| Actinidia chin | A0A2R6Q4L0 |
| Actinidia chin | A0A2R6RAZ9 |
| Actinidia chin | A0A2R6Q6W9 |
| Actinidia chin | A0A2R6QV86 |
| Actinidia chin | A0A2R6QWW6 |
| Actinidia chin | A0A2R6PXQ8 |
| Actinidia chin | A0A2R6R4I5 |
| Actinidia chin | A0A2R6PR71 |

| | |
|---|---|
| Actinidia chin | A0A2R6RB30 |
| Actinidia chin | A0A2R6P2X1 |
| Actinidia chin | A0A2R6R7U2 |
| Actinidia chin | A0A2R6QA16 |
| Actinidia chin | A0A2R6PTF8 |
| Actinidia chin | A0A2R6Q077 |
| Actinidia chin | A0A2R6QTI6 |
| Actinidia chin | A0A2R6R4H6 |
| Actinidia chin | A0A2R6QBV2 |
| Actinidia chin | A0A2R6PQK3 |
| Actinidia chin | A0A2R6QCM9 |
| Actinidia chin | A0A2R6RF49 |
| Actinidia chin | A0A2R6QDU5 |
| Actinidia chin | A0A2R6QLP2 |
| Actinidia chin | A0A2R6PTF3 |
| Actinidia chin | A0A2R6PGN0 |
| Actinidia chin | A0A2R6RRI6 |
| Actinidia chin | A0A2R6RGA0 |
| Actinidia chin | A0A2R6PWC1 |
| Actinidia chin | A0A2R6PXW3 |
| Actinidia chin | A0A2R6Q3G8 |
| Actinidia chin | A0A2R6Q7Z3 |
| Actinidia chin | A0A2R6QBX6 |
| Actinidia chin | A0A2R6QLN4 |
| Actinidia chin | A0A2R6PTE6 |
| Actinidia chin | A0A2R6QWB9 |
| Actinidia chin | A0A2R6PJ76 |
| Actinidia chin | A0A2R6RP34 |
| Actinidia chin | A0A2R6RB22 |
| Actinidia chin | A0A2R6QXS9 |
| Actinidia chin | A0A2R6PEE6 |
| Actinidia chin | A0A2R6NZR5 |
| Actinidia chin | A0A2R6RB09 |
| Actinidia chin | A0A2R6PQU7 |
| Actinidia chin | A0A2R6QLP8 |
| Actinidia chin | A0A2R6QC53 |
| Actinidia chin | A0A2R6PUZ3 |
| Actinidia chin | A0A2R6PTE5 |
| Actinidia chin | A0A2R6QBV5 |
| Actinidia chin | A0A2R6RB02 |
| Actinidia chin | A0A2R6QCD9 |

| | |
|---|---|
| Actinidia chin | A0A2R6QBW7 |
| Actinidia chin | A0A2R6R4H9 |
| Actinidia chin | A0A2R6QWW7 |
| Actinidia chin | A0A2R6PK86 |
| Actinidia chin | A0A2R6NKZ3 |
| Actinidia chin | A0A2R6PRE4 |
| Actinidia chin | A0A2R6PY69 |
| Actinidia chin | A0A2R6RB03 |
| Actinidia chin | A0A2R6QDU9 |
| Actinidia chin | A0A2R6P887 |
| Actinidia chin | A0A2R6QXB5 |
| Actinidia chin | A0A2R6QA21 |
| Actinidia chin | A0A2R6PRM8 |
| Actinidia chin | A0A2R6QMJ8 |
| Acyrthosipho | X1WJB0 |
| Acyrthosipho | J9K2R9 |
| Acyrthosipho | J9LS79 |
| Acyrthosipho | J9KAI5 |
| Acyrthosipho | J9KPT6 |
| Acyrthosipho | J9JPB7 |
| Acyrthosipho | J9K0U0 |
| Acyrthosipho | J9JMX3 |
| Adineta vaga | B3G4G6 |
| Adineta vaga | B3G448 |
| Adineta vaga | B3G4F0 |
| Aedes aegypt | A0A1S4G3R5 |
| Aedes aegypt | Q1HRG8 |
| Aedes aegypt | A0A1S4FK22 |
| Aedes aegypt | Q16Y69 |
| Aedes aegypt | A0A1S4FG50 |
| Aedes aegypt | Q16VE9 |
| Aedes aegypt | Q17JC8 |
| Aedes aegypt | Q171P7 |
| Aedes aegypt | Q17FY1 |
| Aedes aegypt | Q16VE8 |
| Aedes albopi | A0A182H4K1 |
| Aedes albopi | A0A182H5Q1 |
| Aedes albopi | A0A182G6M4 |
| Aedes albopi | A0A182G9D3 |
| Aedes albopi | A0A182G9D4 |
| Aegilops taus | A0A452XH92 |

Aegilops taus A0A452ZGN2
Aegilops taus A0A453BYP1
Aegilops taus A0A453LJQ2
Aegilops taus A0A452ZGI7
Aegilops taus A0A453HQH7
Aegilops taus A0A453AAT9
Aegilops taus A0A453DBA5
Aegilops taus A0A453BBE0
Aegilops taus A0A453BYQ0
Aegilops taus A0A453LK80
Aegilops taus A0A453ADR6
Aegilops taus A0A453Q2G0
Aegilops taus A0A452Z039
Aegilops taus A0A452ZI33
Aegilops taus A0A453D3G9
Aegilops taus A0A453DYZ0
Aegilops taus A0A453JHI0
Aegilops taus A0A453Q2D1
Aegilops taus A0A453LKE2
Aegilops taus A0A453F7F0
Aegilops taus A0A453GM45
Aegilops taus A0A452Y396
Aegilops taus A0A453BYT2
Aegilops taus A0A453MEH1
Aegilops taus A0A453EGG7
Aegilops taus A0A453D372
Aegilops taus A0A453SRN1
Aegilops taus A0A453FPM7
Aegilops taus A0A453BYQ9
Aegilops taus A0A453ADP5
Aegilops taus A0A453JI06
Aegilops taus A0A453DD86
Aegilops taus A0A453LKD6
Aegilops taus A0A453DYQ0
Aegilops taus A0A453ADU5
Aegilops taus A0A452XJR0
Aegilops taus A0A453JGU6
Aegilops taus A0A453NKE5
Aegilops taus A0A453LKC6
Aegilops taus A0A453D3I3
Aegilops taus A0A453JIH7

Aegilops taus A0A453LKE3
Aegilops taus A0A453DBY7
Aegilops taus A0A453SVX0
Aegilops taus A0A453FPN7
Aegilops taus A0A453DDB2
Aegilops taus A0A453DB66
Aegilops taus A0A453FPT3
Aegilops taus A0A453DAW9
Aegilops taus A0A452XJW5
Aegilops taus A0A453DYR4
Aegilops taus A0A453LKH1
Aegilops taus A0A453DBP7
Aegilops taus A0A453NKT3
Aegilops taus A0A453ADR2
Aegilops taus A0A453D3G3
Aegilops taus A0A453AAM1
Aegilops taus A0A453D3A4
Aegilops taus A0A453PQK2
Aegilops taus A0A453Q1Y2
Aegilops taus A0A453LKA1
Aegilops taus A0A453MGE8
Aegilops taus A0A453F7I6
Aegilops taus A0A453FPR4
Aegilops taus A0A453D383
Aegilops taus A0A452Z010
Aegilops taus A0A453QTP8
Aegilops taus A0A453FPN3
Aegilops taus A0A453JHT4
Aegilops taus A0A453FPT0
Aegilops taus A0A453FPK6
Aegilops taus A0A453EGB1
Aegilops taus A0A453QZE9
Aegilops taus A0A453BYN4
Aegilops taus A0A453KBQ0
Aegilops taus A0A452Z4D8
Aegilops taus A0A452ZEX2
Aegilops taus A0A453SHI0
Aegilops taus A0A453D306
Aegilops taus A0A453FPP1
Aegilops taus A0A453F7D3
Aegilops taus A0A453LN14

Aegilops taus A0A453LK85
Aegilops taus A0A453LK71
Aegilops taus A0A453H3Z1
Aegilops taus A0A453DYX5
Aegilops taus A0A453FPQ5
Aegilops taus A0A453DBC1
Aegilops taus A0A452ZGJ2
Aegilops taus A0A452ZGK8
Aegilops taus A0A452ZEZ6
Aegilops taus A0A453LK88
Aegilops taus A0A452YZZ8
Aegilops taus A0A453D3A8
Aegilops taus A0A453LKC7
Aegilops taus A0A453GLW6
Aegilops taus A0A453LK95
Aegilops taus A0A453LK89
Aegilops taus A0A453D358
Aegilops taus A0A453QZ76
Aegilops taus A0A453GMC5
Aegilops taus A0A453BYQ7
Aegilops taus A0A453T0W6
Aegilops taus A0A453DD79
Aegilops taus A0A453BBQ0
Aegilops taus A0A453Q244
Aegilops taus A0A453DBN0
Aegilops taus A0A453GM60
Aegilops taus A0A453D325
Aegilops taus A0A453Q771
Aegilops taus A0A453BCU3
Aegilops taus A0A452XK72
Aegilops taus A0A453DBI2
Aegilops taus A0A453QZN5
Aegilops taus A0A453HAH1
Aegilops taus A0A453FPI6
Aegilops taus A0A452Z002
Aegilops taus A0A453LKB1
Aegilops taus A0A452XK48
Aegilops taus A0A453BBD4
Aegilops taus A0A453Q2E7
Aegilops taus A0A453QZF2
Aegilops taus A0A453DBB2

Aegilops taus A0A453JEW1
Aegilops taus A0A453SU32
Aegilops taus A0A453FPN4
Aegilops taus A0A453DBG7
Aegilops taus A0A453D337
Aegilops taus A0A453LKF9
Aegilops taus A0A453F7G5
Aegilops taus A0A453SVW0
Aegilops taus A0A453BC35
Aegilops taus A0A453D3B0
Aegilops taus A0A453DA54
Aegilops taus A0A453TAB1
Aegilops taus A0A453D3G4
Aegilops taus A0A453JHJ1
Aegilops taus A0A453SRL9
Aegilops taus A0A453ADQ2
Aegilops taus A0A453D349
Aegilops taus A0A453PFF2
Aegilops taus A0A453DA76
Aegilops taus A0A452XH56
Aegilops taus A0A453DC09
Aegilops taus A0A453QTY7
Aegilops taus A0A453DB70
Aegilops taus A0A453DBM7
Aegilops taus A0A453FPK2
Aegilops taus A0A453DBH5
Aegilops taus A0A453Q2F9
Aegilops taus A0A453ADR3
Aegilops taus A0A453DDD0
Aegilops taus A0A452Z027
Aegilops taus A0A453FPS3
Aegilops taus A0A453D6J6
Aegilops taus A0A453J9S0
Aegilops taus A0A453Q2M7
Aegilops taus A0A453LJR3
Aegilops taus A0A453D329
Aegilops taus A0A453GM00
Aegilops taus A0A453BYP5
Aegilops taus A0A453BBD9
Aegilops taus A0A453BYM4
Aegilops taus A0A453Q291

Aegilops taus A0A453QU71
Aegilops taus A0A453H3Z4
Aegilops taus A0A453SGZ7
Aegilops taus A0A453ADQ4
Aegilops taus A0A453LKC4
Aegilops taus A0A453LKG3
Aegilops taus A0A453DBI3
Aegilops taus A0A453DC06
Aegilops taus A0A453SRP6
Aegilops taus A0A453SRK1
Aegilops taus A0A453ADR9
Aegilops taus A0A453QZE3
Aegilops taus A0A453D3H5
Aegilops taus A0A452Z019
Aegilops taus A0A452Y3L8
Aegilops taus A0A453DB32
Aegilops taus A0A453D353
Aegilops taus A0A453H9U8
Aegilops taus A0A453DBK3
Aegilops taus A0A453DBC7
Aegilops taus A0A453NLT6
Aegilops taus A0A453ILU0
Aegilops taus A0A452ZQ21
Aegilops taus A0A453DBV6
Aegilops taus A0A452Z009
Aegilops taus A0A453DDA0
Aegilops taus A0A453G5Z3
Aegilops taus A0A452ZGN5
Aegilops taus A0A453D6L7
Aegilops taus A0A453BYQ2
Aegilops taus A0A453D2Z9
Aegilops taus A0A453DBL2
Aegilops taus A0A453DBD7
Aegilops taus A0A453FPV1
Aegilops taus A0A453D352
Aegilops taus A0A453Q591
Aegilops taus A0A452ZI59
Aegilops taus A0A453Q2L2
Aegilops taus A0A453D333
Aegilops taus A0A453TA51
Aegilops taus A0A453DDL7

Aegilops taus A0A453G617
Aegilops taus A0A453D2X4
Aegilops taus A0A452XD19
Aegilops taus A0A453Q1X4
Aegilops taus A0A453MGQ3
Aegilops taus A0A453HRF8
Aegilops taus A0A453EGH4
Aegilops taus A0A453HQJ5
Aegilops taus A0A453LN67
Aegilops taus A0A453DB22
Aegilops taus A0A453BDB4
Aegilops taus A0A453LK93
Aegilops taus A0A453DDM1
Aegilops taus A0A453D369
Aegilops taus A0A453DBK6
Aegilops taus A0A453SRN7
Aegilops taus A0A452ZGM5
Aegilops taus A0A453D6Q2
Aegilops taus A0A453LKB4
Aegilops taus A0A453BYR3
Aegilops taus A0A453D3C0
Aegilops taus A0A452XD37
Aegilops taus A0A453DDM6
Aegilops taus A0A453ADV9
Aegilops taus A0A453FPM3
Aegilops taus A0A453TAE6
Aegilops taus A0A453BC47
Aegilops taus A0A452XK40
Aegilops taus A0A453F7D6
Aegilops taus A0A453NKD5
Aegilops taus A0A453DC51
Aegilops taus A0A453DBE4
Aegilops taus A0A453T114
Aegilops taus A0A453BYS4
Aegilops taus A0A453H3Z7
Aegilops taus A0A453ATP0
Aegilops taus A0A453G6T0
Aegilops taus A0A452XZ51
Aegilops taus A0A453DBW3
Aegilops taus A0A452XD28
Aegilops taus A0A453LJR8

Aegilops taus A0A453BB79
Aegilops taus A0A453D382
Aegilops taus A0A453LK83
Aegilops taus A0A452ZI63
Aegilops taus A0A452XZD5
Aegilops taus A0A452Z4E2
Aegilops taus A0A453T139
Aegilops taus A0A453DB02
Aegilops taus A0A453FPR7
Aegilops taus A0A453PFD9
Aegilops taus A0A453BBJ6
Aegilops taus A0A453LK79
Aegilops taus A0A453BBV1
Aegilops taus A0A453DBL5
Aegilops taus A0A453FPR9
Aegilops taus A0A453D361
Aegilops taus A0A452XD46
Aegilops taus A0A453QZU7
Aegilops taus A0A453ATS1
Aegilops taus A0A453FPJ1
Aegilops taus A0A452XB26
Aegilops taus A0A453QTW5
Aegilops taus A0A453BYP4
Aegilops taus A0A453D323
Aegilops taus A0A453ADR1
Aegilops taus A0A452XJS2
Aegilops taus A0A453FPS0
Aegilops taus A0A453FPQ0
Aegilops taus A0A453JB99
Aegilops taus A0A453DBF2
Aegilops taus A0A453KC08
Aegilops taus A0A452XJG0
Aegilops taus A0A453DBT2
Aegilops taus A0A453IFX4
Aegilops taus A0A452XK35
Aegilops taus A0A453D3B3
Aegilops taus A0A453LJU7
Aegilops taus A0A453JH21
Aegilops taus A0A453S9X2
Aegilops taus A0A452XJT3
Aegilops taus A0A453DBV3

Aegilops taus A0A452Z033
Aegilops taus A0A452XH89
Aegilops taus A0A453SAP2
Aegilops taus A0A453BBW1
Aegilops taus A0A453DC15
Aegilops taus A0A453DBY3
Aegilops taus A0A453DDL1
Aegilops taus A0A453D3A1
Aegilops taus A0A453D396
Aegilops taus A0A453H3Z3
Aegilops taus A0A453EGJ2
Aegilops taus A0A453DBW8
Aegilops taus A0A453LKB3
Aegilops taus A0A453EGJ7
Aegilops taus A0A453G694
Aegilops taus A0A453Q4I2
Aegilops taus A0A453CF81
Aegilops taus A0A453FPL3
Aegilops taus A0A453GM65
Aegilops taus A0A453GMJ9
Aegilops taus A0A452Y3H0
Aegilops taus A0A453D3A3
Aegilops taus A0A453DDC2
Aegilops taus A0A452ZNP1
Aegilops taus A0A452ZGK3
Aegilops taus A0A453DBU3
Aegilops taus A0A453GQY1
Aegilops taus A0A453IM88
Aegilops taus A0A453D363
Aegilops taus A0A453SRL5
Aegilops taus A0A453Q2A6
Aegilops taus A0A453JHH1
Aegilops taus A0A453BBU6
Aegilops taus A0A453JFZ5
Aegilops taus A0A453H3Z2
Aegilops taus A0A453DBR3
Aegilops taus A0A453DBP2
Aegilops taus A0A453LK56
Aegilops taus A0A453TAD7
Aegilops taus A0A453JH11
Aegilops taus A0A453FPS4

Aegilops taus A0A452ZGP0
Aegilops taus A0A453QZV2
Aegilops taus A0A453Q1W1
Aegilops taus A0A452XH97
Aegilops taus A0A453D2W3
Aegilops taus A0A453SVQ7
Aegilops taus A0A453DBR6
Aegilops taus A0A453EGJ0
Aegilops taus A0A453MGF3
Aegilops taus A0A453D3B5
Aegilops taus A0A453BC42
Aegilops taus A0A453Q2K8
Aegilops taus A0A453K9H8
Aegilops taus A0A453LJN7
Aegilops taus A0A453DBK2
Aegilops taus A0A453BBR3
Aegilops taus A0A453AAL0
Aegilops taus A0A453SRP1
Aegilops taus A0A453FPQ4
Aegilops taus A0A453DBE5
Aegilops taus A0A453QU60
Aegilops taus A0A453SRQ7
Aegilops taus A0A453QUA8
Aegilops taus A0A453SUC2
Aegilops taus A0A453A3H4
Aegilops taus A0A453BYQ1
Aegilops taus A0A452XD39
Aegilops taus A0A453TAB6
Aegilops taus A0A453FPV0
Aegilops taus A0A452ZGM7
Aegilops taus A0A452ZI86
Aegilops taus A0A453SAJ3
Aegilops taus A0A453DBS2
Aegilops taus A0A453D394
Aegilops taus A0A453DB45
Aegilops taus A0A452ZGN4
Aegilops taus A0A452Z015
Aegilops taus A0A452ZNQ3
Aegilops taus A0A453BBE7
Aegilops taus A0A453LK78
Aegilops taus A0A453F7K0

Aegilops taus A0A453LKB7
Aegilops taus A0A453D375
Aegilops taus A0A453DB42
Aegilops taus A0A453FPS1
Aegilops taus A0A453D385
Aegilops taus A0A453LKJ9
Aegilops taus A0A453SRH1
Aegilops taus A0A453SVF0
Aegilops taus A0A453BBF4
Aegilops taus A0A453QU28
Aegilops taus A0A453SGI2
Aegilops taus A0A453DBL7
Aegilops taus A0A453AAP4
Aegilops taus A0A453KCJ5
Aegilops taus A0A453D310
Aegilops taus A0A453DC24
Aegilops taus A0A453D3E6
Aegilops taus A0A453H3Z9
Aegilops taus A0A453TA82
Aegilops taus A0A453FPU6
Aegilops taus A0A453SGY8
Aegilops taus A0A453LK90
Aegilops taus A0A452ZQ32
Aegilops taus A0A453EGC1
Aegilops taus A0A453D390
Aegilops taus A0A452XB18
Aegilops taus A0A453LKE6
Aegilops taus A0A452ZRM6
Aegilops taus A0A453LKA2
Aegilops taus A0A453EGK8
Aegilops taus A0A452Z021
Aegilops taus A0A453DDC6
Aegilops taus A0A453HR14
Aegilops taus A0A453DB15
Aegilops taus A0A453GLY9
Aegilops taus A0A453LJT7
Aegilops taus A0A453FPS5
Aegilops taus A0A453BYR6
Aegilops taus A0A453JIE9
Aegilops taus A0A453FPU5
Aegilops taus A0A453T0S7

Aegilops taus A0A452Z4F2
Aegilops taus A0A453Q2M4
Aegilops taus A0A453ILU7
Aegilops taus A0A452ZGI2
Aegilops taus A0A453SFC7
Aegilops taus A0A453LJW8
Aegilops taus A0A453Q1Y8
Aegilops taus A0A452ZGM9
Aegilops taus A0A452XB15
Aegilops taus A0A453MGP2
Aegilops taus A0A452ZQ12
Aegilops taus A0A453DC10
Aegilops taus A0A453LKG5
Aegilops taus A0A453BBC4
Aegilops taus A0A453D2Z4
Aegilops taus A0A453Q471
Aegilops taus A0A453H401
Aegilops taus A0A453SRN8
Aegilops taus A0A453DYZ4
Aegilops taus A0A453DBH0
Aegilops taus A0A453SVS6
Aegilops taus A0A453LKB9
Aegilops taus A0A453ADT9
Aegilops taus A0A453SRL4
Aegilops taus A0A452XJY3
Aegilops taus A0A453EGB6
Aegilops taus A0A453DBX4
Aegilops taus A0A453D3D9
Aegilops taus A0A453QZF5
Aegilops taus A0A452ZEW0
Aegilops taus A0A453DBG6
Aegilops taus A0A453QZE7
Aegilops taus A0A453LJY8
Aegilops taus A0A453ADP8
Aegilops taus A0A453SRL2
Aegilops taus A0A453DDC8
Aegilops taus A0A453DC41
Aegilops taus A0A453D315
Aegilops taus A0A453DBP0
Aegilops taus A0A453LKE7
Aegilops taus A0A453D3A5

Aegilops taus A0A453JGZ8
Aegilops taus A0A452XHA9
Aegilops taus A0A453DB67
Aegilops taus A0A452ZGL3
Aegilops taus A0A453Q276
Aegilops taus A0A453QZ80
Aegilops taus A0A453MGQ6
Aegilops taus A0A453FPJ7
Aegilops taus A0A453FPR0
Aegilops taus A0A453DBN2
Aegilops taus A0A453FPS6
Aegilops taus A0A452XH94
Aegilops taus A0A453BBV6
Aegilops taus A0A453LN06
Aegilops taus A0A453DBE9
Aegilops taus A0A453ADV4
Aegilops taus A0A453DDC4
Aegilops taus A0A453EGH0
Aegilops taus A0A453D3F8
Aegilops taus A0A453IG18
Aegilops taus A0A453ADQ8
Aegilops taus A0A453DBM5
Aegilops taus A0A453F7C5
Aegilops taus A0A453LKD9
Aegilops taus A0A453DBX3
Aegilops taus A0A453D3H9
Aegilops taus A0A453D302
Aegilops taus A0A453BBC9
Aegilops taus A0A452XD12
Aegilops taus A0A453DBK1
Aegilops taus A0A453SKA3
Aegilops taus A0A453HRC9
Aegilops taus A0A453DBG1
Aegilops taus A0A453DBE1
Aegilops taus A0A453D359
Aegilops taus A0A453HAB3
Aegilops taus A0A453DDN3
Aegilops taus A0A453D6N7
Aegilops taus A0A452XK26
Aegilops taus A0A453T0X8
Aegilops taus A0A452ZGM1

Aegilops taus A0A453EGH5
Aegilops taus A0A453BYM8
Aegilops taus A0A453D395
Aegilops taus A0A453LKC3
Aegilops taus A0A453MGP0
Aegilops taus A0A453LK53
Aegilops taus A0A452XJQ5
Aegilops taus A0A453D3A0
Aegilops taus A0A453D3C4
Aegilops taus A0A452XK20
Aegilops taus A0A452XHA4
Aegilops taus A0A453JG11
Aegilops taus A0A453DBP9
Aegilops taus A0A453DBW4
Aegilops taus A0A453JHF1
Aegilops taus A0A452XHB4
Aegilops taus A0A453SRP8
Aegilops taus A0A453DBL1
Aegilops taus A0A453D309
Aegilops taus A0A452YZZ3
Aegilops taus A0A453DBN5
Aegilops taus A0A453HAG7
Aegilops taus A0A453DBJ1
Aegilops taus A0A453DDM2
Aegilops taus A0A452XB20
Aegilops taus A0A453D2X9
Aegilops taus A0A453SRM5
Aegilops taus A0A453Q4I8
Aegilops taus A0A453DIV2
Aegilops taus A0A453BDH9
Aegilops taus A0A452XB13
Aegilops taus A0A453DDB4
Aegilops taus A0A453GMC0
Aegilops taus A0A453DBP5
Aegilops taus A0A453SRE5
Aegilops taus A0A452ZGQ0
Aegilops taus A0A453LKG2
Aegilops taus A0A453BDI1
Aegilops taus A0A452ZNP3
Aegilops taus A0A453FPR3
Aegilops taus A0A453JIE4

Aegilops taus A0A453BD61
Aegilops taus A0A453D398
Aegilops taus A0A453FPL9
Aegilops taus A0A452ZEW7
Aegilops taus M8C3G1
Aegilops taus N1QZA4
Aegilops taus M8BLT1
Aegilops taus M8CDU3
Aegilops taus M8BQF5
Aegilops taus R7W3S1
Aegilops taus M8CA09
Aegilops taus M8C8E1
Aegilops taus R7W6C2
Aegilops taus M8BRL3
Aegilops taus M8BNQ4
Aegilops taus M8BSU9
Aegilops taus M8B659
Aegilops taus M8B657
Aegilops taus M8B6T4
Aegilops taus R7W524
Aegilops taus R7W6Q0
Aegilops taus M8ARV2
Aegilops taus N1R0P4
Aegilops taus N1QWT2
Aegilops taus M8C3Q0
Aegilops taus M8BS71
Aegilops taus M8C8A2
Aegilops taus M8CGA2
Aegilops taus M8D7P7
Aegilops taus M8BGE8
Aegilops taus M8AXL9
Aegilops taus N1QQ18
Aegilops taus R7W4G4
Aegilops taus N1R0J5
Aegilops taus M8CMY7
Aegilops taus M8BFG4
Aegilops taus M8CQW7
Aegilops taus R7W4J2
Aegilops taus M8BZH7
Aegilops taus R7W0V2
Aegilops taus M8BFQ4

Aegilops taus N1R4A5
Aegilops taus M8BVW7
Aegilops taus R7W466
Aegilops taus N1QUE8
Aegilops taus N1QPR6
Aegilops taus R7W2M1
Aegilops taus M8AQG2
Aegilops taus M8D5R8
Aegilops taus M8B4Z6
Aegilops taus R7WBZ0
Aegilops taus N1R267
Aegilops taus M8BEL9
Aegilops taus M8AXV6
Aegilops taus M8BHI2
Aegilops taus R7W702
Aegilops taus N1QZP8
Aegilops taus M8BWJ0
Aegilops taus M8AR11
Aegilops taus R7W5U0
Aegilops taus M8BKQ6
Aegilops taus M8C7I9
Aegilops taus R7W886
Aegilops taus N1QTT0
Aegilops taus R7W6D7
Aegilops taus M8AR15
Aegilops taus M8BFJ7
Aegilops taus M8B3Y2
Aegilops taus M8BW41
Aegilops taus M8BV04
Aegilops taus M8BSB0
Aegilops taus M8C2A6
Aegilops taus N1QZS0
Aegilops taus M8B4G8
Aegilops taus M8BIP4
Aegilops taus N1QVU3
Aegilops taus M8CU22
Aegilops taus R7WB01
Aegilops taus M8C5G3
Aegilops taus M8BSH4
Aegilops taus M8BH12
Aegilops taus M8CC39

Aegilops taus N1QZA9
Aegilops taus R7W2H2
Aegilops taus M8AWU4
Aegilops taus R7W5J6
Aegilops taus M8BVP9
Aegilops taus M8B6H7
Aegilops taus M8CQN8
Aegilops taus N1QPF3
Aegilops taus N1QVJ3
Aegilops taus M8CV03
Aegilops taus M8C2T7
Aegilops taus M8BJJ3
Aegilops taus M8BBM6
Aegilops taus M8C398
Aegilops taus M8C374
Aegilops taus M8BRL2
Aegilops taus R7W3B4
Aegilops taus M8CMA1
Aegilops taus M8BMX3
Aegilops taus R7WBP2
Aegilops taus M8BAQ4
Aegilops taus M8BCF9
Aegilops taus M8AI36
Aegilops taus M8BZV5
Aegilops taus N1QRV4
Aegilops taus M8AI44
Aegilops taus M8CDI5
Aegilops taus M8BRQ9
Aegilops taus M8BS62
Aegilops taus N1QTT3
Aegilops taus M8CXN2
Aegilops taus R7WAL8
Aegilops taus R7W389
Aegilops taus M8CS97
Aegilops taus R7W2A0
Aegilops taus M8BMF1
Aegilops taus M8CLL1
Aegilops taus M8BBG0
Aegilops taus N1R416
Aegilops taus M8C0P3
Aegilops taus N1QSH2

| | |
|---|---|
| Aegilops taus | M8C8P5 |
| Aegilops taus | M8BC75 |
| Aegilops taus | R7W9K6 |
| Aegilops taus | M8BZ85 |
| Aegilops taus | M8AQX6 |
| Aegilops taus | M8BRA9 |
| Aegilops taus | M8BV83 |
| Aegilops taus | N1QUU5 |
| Aegilops taus | R7W017 |
| Aegilops taus | M8AWM2 |
| Aegilops taus | M8C1A2 |
| Aegilops taus | M8CIT9 |
| Aegilops taus | M8BR67 |
| Aegilops taus | N1R3B9 |
| Aegilops taus | M8BVQ9 |
| Aegilops taus | M8APF0 |
| Aegilops taus | R7W5F9 |
| Aegilops taus | N1QR69 |
| Aegilops taus | N1QWL3 |
| Aegilops taus | R7W6I7 |
| Afifella marin | A0A1G5P0D6 |
| Afipia broom | K8P0P5 |
| Afipia broom | A0A2M8ZRI2 |
| Afipia clevela | K8P4L2 |
| Afipia felis.xls | A0A090MQ82 |
| Afipia felis.xls | A0A380WCA0 |
| Afipia massili | A0A0M3DZ05 |
| Afipia sp. 1NI | D6V6Y8 |
| Afipia sp. 62-7 | A0A1M2ZI81 |
| Afipia sp. 64- | A0A1Q3W3V1 |
| Afipia sp. GAS | A0A1H0JMN5 |
| Afipia sp. P52 | W3RJC5 |
| Afipia sp. Roc | A0A0Q6ZHE5 |
| Afipia sp..xlsx | A0A2D6Z1S1 |
| African swine | P0C9W5 |
| African swine | A0A0C5AWG9 |
| Agrilus planip | A0A1W4XI26 |
| Agrilus planip | A0A1W4WER7 |
| Agrilus planip | A0A1W4WCW5 |
| Agrilus planip | A0A1W4WMX9 |
| Agrobacteriu | K2PXK1 |

| | |
|---|---|
| Agrobacteriu | A0A0D8KTY2 |
| Agrobacteriu | A0A135P8H9 |
| Agrobacteriu | A0A1S7TYJ4 |
| Agrobacteriu | A0A1S7R665 |
| Agrobacteriu | A0A1S7QSC7 |
| Agrobacteriu | Q7CT39 |
| Agrobacteriu | A0A1S7R1K3 |
| Agrobacteriu | A0A2W5HEG2 |
| Agrobacteriu | A0A3G2D9W1 |
| Agrobacteriu | A0A3G2CW11 |
| Agrobacteriu | A0A1S7N2M1 |
| Agrobacteriu | A0A1S7N2Q8 |
| Agrobacteriu | A0A1S7S8C0 |
| Agrobacteriu | A0A1S7MTQ7 |
| Agrobacteriu | A0A1S7N332 |
| Agrobacteriu | A0A1S7R6H0 |
| Agrobacteriu | A0A1S7S0T3 |
| Agrobacteriu | B9J7G8 |
| Agrobacteriu | W1KEF2 |
| Agrobacteriu | W1LDB8 |
| Agrobacteriu | A0A061MPX5 |
| Agrobacteriu | A0A2T7V3G2 |
| Agrobacteriu | A0A2T7VDF4 |
| Agrobacteriu | A0A178H301 |
| Agrobacteriu | A0A071HXB3 |
| Agrobacteriu | A0A071HYF2 |
| Agrobacteriu | A0A2S4E9Q9 |
| Agrobacteriu | A0A081CTJ4 |
| Agrobacteriu | A0A1B9TQL4 |
| Agrobacteriu | A0A1S7UAP0 |
| Agrobacteriu | A0A1S9E9P5 |
| Agrobacteriu | A0A1S9E6I0 |
| Agrobacteriu | F0LCJ2 |
| Agrobacteriu | F5J606 |
| Agrobacteriu | A0A3S2U7W8 |
| Agrobacteriu | A0A1R3TJ16 |
| Agrobacteriu | A0A1R3TLG8 |
| Agrobacteriu | A0A427QHU1 |
| Agrobacteriu | A0A0X6ZW52 |
| Agrobacteriu | A0A0X6ZP94 |
| Agrobacteriu | A0A328F968 |

| | |
|---|---|
| Agrobacteriu | A0A328ESI4 |
| Agrobacteriu | A0A1S7RVT6 |
| Agrobacteriu | A0A1B3P384 |
| Agrobacteriu | A0A1E3YAW1 |
| Agrobacteriu | A0A0L6K5Z5 |
| Agrobacteriu | A0A349XP92 |
| Agrobacteriu | A0A357X7H8 |
| Agrobacteriu | H0H495 |
| Agrobacteriu | H0HI51 |
| Agrobacteriu | F7UCM7 |
| Agrobacteriu | A0A1S7QYX0 |
| Agrobacteriu | A0A1S7RCW1 |
| Agrobacteriu | A0A1S7T9V4 |
| Agrobacteriu | M8B9N1 |
| Agrobacteriu | A0A1S7RU05 |
| Agrobacteriu | B9JSY8 |
| Agrobacteriu | A0A202F108 |
| Agrobacteriu | A0A1S2DVC4 |
| Agrobacteriu | A7XED0 |
| Agrobacteriu | A0A1S2E625 |
| Agrobacteriu | A0A368NW97 |
| Ahrensia sp..: | A0A2E0IUI6 |
| Ahrensia sp..: | A0A2E2RQX8 |
| Ahrensia sp..: | A0A2E0IF36 |
| Ahrensia sp..: | A0A2E2RKN7 |
| Ailuropoda m | G1M174 |
| Ailuropoda m | G1LEP0 |
| Ailuropoda m | G1MBX3 |
| Ailuropoda m | G1LET2 |
| Ailuropoda m | G1M194 |
| Ailuropoda m | D2HM20 |
| Ailuropoda m | D2HEU5 |
| Ailuropoda m | G1MDT1 |
| Ailuropoda m | D2H7K2 |
| Albuca bracte | A0A0A7LYW8 |
| Albugo candi | A0A024GUK6 |
| Albugo candi | A0A024GTY2 |
| Albugo candi | A0A024FUV0 |
| Albugo candi | A0A024FUS4 |
| Albugo laibac | F0WGZ2 |
| Albugo laibac | F0W170 |

| | |
|---|---|
| Albugo laibac | F0WDJ8 |
| Albugo laibac | F0WDJ9 |
| Albugo laibac | F0WGX7 |
| Aliivibrio sp. 1 | A0A1B9PIN4 |
| Alligator miss | A0A151M2C0 |
| Alligator miss | A0A151NPI2 |
| Alligator miss | A0A151PBF9 |
| Alligator miss | A0A151NPX2 |
| Alligator miss | A0A151M2C2 |
| Alligator miss | A0A151MTI6 |
| Alligator miss | A0A151PAG7 |
| Alligator sine | A0A1U7S3Q2 |
| Alligator sine | A0A1U7R1W7 |
| Alligator sine | A0A3Q0GI02 |
| Alligator sine | A0A1U8DHU0 |
| Allochromati | D3RPG7 |
| Allochromati | A0A1H3HNT0 |
| Allomyces ma | A0A0L0S2D3 |
| Allomyces ma | A0A0L0RXQ9 |
| allotetraploid | A0A0U2S302 |
| allotetraploid | A0A0U2JFZ1 |
| allotetraploid | A0A0U2RH63 |
| alpha proteol | A0A1V1PR56 |
| Alphaproteol | A0A1Q7BNX2 |
| Alphaproteol | A0A1Q3K2D4 |
| Alphaproteol | A0A1L9Q6D6 |
| Alphaproteol | A0A2N3CAF3 |
| Alphaproteol | A0A2N3CN55 |
| Alphaproteol | A0A2N3BHU6 |
| Alphaproteol | A0A2N3BWC8 |
| Alphaproteol | A0A3M1A8N1 |
| Alphaproteol | A0A3M1CCK9 |
| Alphaproteol | A0A3B9YVH5 |
| Alphaproteol | A0A3B9YUR4 |
| Alphaproteol | A0A3C0MDP2 |
| Alternaria alt | A0A177E5Y0 |
| Alternaria alt | A0A177DMN5 |
| Alternaria alt | A0A177DB16 |
| Alternaria alt | A0A177DW37 |
| Alternaria alt | A0A177D586 |
| Alternaria alt | A0A177DA37 |

| | |
|---|---|
| Alternaria sp. | A0A399GD40 |
| Alternaria sp. | A0A399HQ58 |
| Alternaria sp. | A0A399G9V7 |
| Amaricoccus | A0A2M9PBP4 |
| Amazona aes | A0A0Q3U1E2 |
| Amazona aes | A0A0Q3UR44 |
| Amazona aes | A0A0Q3MLN8 |
| Amazona aes | A0A0Q3MGK6 |
| Amblyomma | A0A1E1X339 |
| Amblyomma | A0A1E1XND8 |
| Amborella tri | W1PSU2 |
| Amborella tri | W1NHI6 |
| Amborella tri | W1NS07 |
| Amborella tri | W1PUY6 |
| Amborella tri | U5D4B7 |
| Amborella tri | W1PK20 |
| Amborella tri | W1NP94 |
| Amborella tri | W1PWB1 |
| Amborella tri | W1P1F1 |
| Amborella tri | W1PJ97 |
| Amborella tri | U5CZI6 |
| Amborella tri | W1PNM0 |
| Amborella tri | W1PYM5 |
| Amborella tri | U5D3T0 |
| Amborella tri | W1NII4 |
| Amborella tri | W1NTK9 |
| Amborella tri | W1PI96 |
| Amborella tri | W1NKX3 |
| Amborella tri | W1PU85 |
| Amborella tri | W1PHD9 |
| Amborella tri | W1P824 |
| Amborella tri | W1P1U3 |
| Amborella tri | W1PS15 |
| Amborella tri | W1P261 |
| Aminobacter | A0A380WPF7 |
| Aminobacter | A0A380WLF7 |
| Aminobacter | A0A142M5P6 |
| Aminobacter | A0A316H1V4 |
| Aminobacter | A0A2S0XJL2 |
| Aminobacter | A0A0Q6M906 |
| Amphilophus | A0A3Q0QWF0 |

| | |
|---|---|
| Amphilophus | A0A3Q0RP91 |
| Amphilophus | A0A3Q0QW88 |
| Amphilophus | A0A3Q0QZA4 |
| Amphilophus | A0A3Q0QXI1 |
| Amphilophus | A0A3Q0QYJ9 |
| Amphilophus | A0A3Q0QX28 |
| Amphilophus | A0A3Q0RIQ3 |
| Amphilophus | A0A3Q0QX05 |
| Amphilophus | A0A3Q0RNB8 |
| Amphilophus | A0A3Q0QX18 |
| Amphilophus | A0A3Q0RNE4 |
| Amphilophus | A0A3Q0T1F7 |
| Amphimedor | A0A1X7VF86 |
| Amphimedor | A0A1X7VJI8 |
| Amphiprion c | A0A3Q1CUE1 |
| Amphiprion c | A0A3Q1AZ85 |
| Amphiprion c | A0A3Q1BNV0 |
| Amphiprion c | A0A3Q1ASI9 |
| Amphiprion c | A0A3Q1CEN8 |
| Amphiprion p | A0A3P8SBR5 |
| Amphiprion p | A0A3P8U3P7 |
| Amphiprion p | A0A3P8RV74 |
| Amphiprion p | A0A3P8SZJ0 |
| Amphiprion p | A0A3P8T1W7 |
| Ampulex com | A0A1W6EWC0 |
| Ampulex com | A0A1W6EWB2 |
| Amylibacter c | A0A2N5IM96 |
| Anabaena sp. | A0A1B7WGH8 |
| Anabas testu | A0A3Q1JM01 |
| Anabas testu | A0A3Q1JGW1 |
| Anabas testu | A0A3Q1HBQ8 |
| Anabas testu | A0A3Q1JNC7 |
| Anabas testu | A0A3Q1HBU2 |
| Anabas testu | A0A3Q1K4Z1 |
| Anabas testu | A0A3Q1HBM8 |
| Anabas testu | A0A3Q1JWC6 |
| Anabas testu | A0A3Q1H376 |
| Anabas testu | A0A3Q1JWD0 |
| Anabas testu | A0A3Q1H8P6 |
| Anaeromyxol | A0A1F3C361 |
| Ananas comc | A0A199VKR8 |

| | |
|---|---|
| Ananas comc | A0A199W1P5 |
| Ananas comc | A0A199VMM6 |
| Ananas comc | A0A199V5G5 |
| Ananas comc | A0A199V0P4 |
| Ananas comc | A0A199VXD4 |
| Ananas comc | A0A199UUR3 |
| Ananas comc | A0A199UNR1 |
| Ananas comc | A0A199UFB6 |
| Ananas comc | A0A199V3G9 |
| Ananas comc | A0A199UF66 |
| Ananas comc | A0A199VMS2 |
| Ananas comc | A0A199W3M8 |
| Ananas comc | A0A199VXV5 |
| Ananas comc | A0A199UW40 |
| Ananas comc | A0A199VG00 |
| Ananas comc | A0A199VR05 |
| Ananas comc | A0A199UW09 |
| Ananas comc | A0A199VZH8 |
| Ananas comc | A0A199VCY3 |
| Ananas comc | A0A199UMA3 |
| Ananas comc | A0A199VZ99 |
| Ananas comc | A0A199VI10 |
| Ananas comc | A0A199UUQ7 |
| Ananas comc | A0A199UV66 |
| Ananas comc | A0A199UQ24 |
| Anas platyrh | U3IRP5 |
| Anas platyrh | U3I6X0 |
| Anas platyrh | U3II31 |
| Anas platyrh | R0K1Z7 |
| Anas platyrh | R0JPB7 |
| Anas platyrh | R0M4Q0 |
| Anas platyrh | R0LQI0 |
| Ancylostoma | A0A368GRP9 |
| Ancylostoma | A0A368GP95 |
| Ancylostoma | A0A368GMN0 |
| Ancylostoma | A0A368H349 |
| Ancylostoma | A0A368GJB9 |
| Ancylostoma | A0A368GWF9 |
| Ancylostoma | A0A368HB09 |
| Ancylostoma | A0A368G5Y2 |
| Ancylostoma | A0A368GRQ0 |

Ancylostoma A0A368GVT4
Ancylostoma A0A368GWX4
Ancylostoma A0A368H7D6
Ancylostoma A0A368H3B6
Ancylostoma A0A368FPC8
Ancylostoma A0A368GP49
Ancylostoma A0A368GXY9
Ancylostoma A0A368EXZ3
Ancylostoma A0A368GJW8
Ancylostoma A0A368FA73
Ancylostoma A0A368GWI0
Ancylostoma A0A368GX96
Ancylostoma A0A368GTK6
Ancylostoma A0A368H705
Ancylostoma A0A368GFH7
Ancylostoma A0A368H7B8
Ancylostoma A0A368HDM2
Ancylostoma A0A368GFI5
Ancylostoma A0A368HEE4
Ancylostoma A0A368GI28
Ancylostoma A0A368GRE3
Ancylostoma A0A368FCY8
Ancylostoma A0A368FCZ3
Ancylostoma A0A368G2Q2
Ancylostoma A0A016VKR6
Ancylostoma A0A0D6M3P1
Ancylostoma A0A016WI38
Ancylostoma A0A016V0U2
Ancylostoma A0A016S814
Ancylostoma A0A016RWJ5
Ancylostoma A0A0D6LC48
Ancylostoma A0A016WHK2
Ancylostoma A0A016SGK6
Ancylostoma A0A016T8L1
Ancylostoma A0A016SHF1
Ancylostoma A0A016WSY7
Ancylostoma A0A016S7S9
Ancylostoma A0A016S6Y5
Ancylostoma A0A0D6LU43
Ancylostoma A0A016TUW3
Ancylostoma A0A016T0N8

Ancylostoma A0A016W3A0
Ancylostoma A0A0D6LRJ3
Ancylostoma A0A016WIX9
Ancylostoma A0A016WRA4
Ancylostoma A0A016RV21
Ancylostoma A0A016VJZ2
Ancylostoma A0A016STN6
Ancylostoma A0A0D6LUD3
Ancylostoma A0A016SU80
Ancylostoma A0A0D6M9L2
Ancylostoma A0A016SMB0
Ancylostoma A0A016VCU4
Ancylostoma A0A016SS42
Ancylostoma A0A016W8B6
Ancylostoma A0A016V243
Ancylostoma A0A016SU41
Ancylostoma A0A016V216
Ancylostoma A0A016SRZ7
Ancylostoma A0A016V3K2
Ancylostoma A0A016W7N7
Ancylostoma A0A016UI89
Ancylostoma A0A016WS77
Ancylostoma A0A016UHN2
Ancylostoma A0A016TCL6
Ancylostoma A0A016T8A6
Ancylostoma A0A016WA14
Ancylostoma A0A0D6MAD7
Ancylostoma A0A016VDI1
Ancylostoma A0A016T206
Ancylostoma A0A016V2S3
Ancylostoma A0A016WI79
Ancylostoma A0A016W8D8
Ancylostoma A0A0D6M851
Ancylostoma A0A016V2R6
Ancylostoma A0A016W998
Ancylostoma A0A016SR62
Ancylostoma A0A016V233
Ancylostoma A0A016WTI9
Ancylostoma A0A016UJN5
Ancylostoma A0A016V3J7
Ancylostoma A0A016TTS6

Ancylostoma A0A0D6LG45
Ancylostoma A0A016VDY5
Ancylostoma A0A016WNP3
Ancylostoma A0A016STB4
Ancylostoma A0A016VIC1
Ancylostoma A0A016V2R1
Ancylostoma A0A016SS83
Ancylostoma A0A016WSZ0
Ancylostoma A0A016V1H8
Ancylostoma A0A0D6M4L0
Ancylostoma A0A016TCJ1
Ancylostoma A0A016VFG3
Ancylostoma A0A016VEL5
Ancylostoma A0A016WGP1
Ancylostoma A0A016UG75
Ancylostoma A0A0D6LIH3
Ancylostoma A0A016TCI8
Ancylostoma A0A016VL99
Ancylostoma A0A016SU43
Ancylostoma A0A016SHA2
Ancylostoma A0A016TVN9
Ancylostoma A0A016WI62
Ancylostoma A0A016T0X6
Ancylostoma A0A016WQP5
Ancylostoma A0A016SU38
Ancylostoma A0A016VJ86
Ancylostoma A0A016SL82
Ancylostoma A0A016UGP0
Ancylostoma A0A016VA51
Ancylostoma A0A016WJ45
Ancylostoma A0A016WEZ6
Ancylostoma A0A016T7T4
Ancylostoma A0A016TXR8
Ancylostoma A0A0D6LZI8
Ancylostoma A0A0D6LVB3
Ancylostoma A0A016WN18
Ancylostoma A0A016WV69
Ancylostoma A0A016RVX3
Ancylostoma A0A0D6LXD0
Ancylostoma A0A016WHK9
Ancylostoma A0A016UHG4

| | |
|---|---|
| Ancylostoma | A0A016V2P8 |
| Ancylostoma | A0A0D6M3G6 |
| Ancylostoma | A0A016WUK9 |
| Ancylostoma | A0A0D6M3L0 |
| Ancylostoma | A0A0D6M584 |
| Ancylostoma | A0A016RVW0 |
| Ancylostoma | A0A0C2DVX2 |
| Ancylostoma | A0A0C2FMV1 |
| Ancylostoma | A0A0C2G7Y5 |
| Ancylostoma | A0A0C2CRJ1 |
| Ancylostoma | A0A0C2GDQ6 |
| Ancylostoma | A0A0C2H4Q3 |
| Ancylostoma | A0A0C2GLS5 |
| Ancylostoma | A0A0C2G2S8 |
| Ancylostoma | A0A0C2DFM8 |
| Ancylostoma | A0A0C2DQD8 |
| Ancylostoma | A0A0C2H0D7 |
| Ancylostoma | A0A0C2CKU6 |
| Ancylostoma | A0A0C2C4H3 |
| Ancylostoma | A0A0C2HCQ8 |
| Ancylostoma | A0A0C2BR75 |
| Ancylostoma | A0A0C2FCW8 |
| Ancylostoma | A0A0C2FEG0 |
| Ancylostoma | A0A0C2D4J6 |
| Ancylostoma | A0A0C2F8I9 |
| Ancylostoma | A0A0C2FGE8 |
| Ancylostoma | A0A0C2FP16 |
| Ancylostoma | A0A0C2GFH0 |
| Ancylostoma | A0A0C2G7Z8 |
| Ancylostoma | A0A0C2H638 |
| Ancylostoma | A0A0C2FGY3 |
| Ancylostoma | A0A0C2CYI0 |
| Ancylostoma | A0A0C2FI47 |
| Ancylostoma | A0A0C2F9X1 |
| Ancylostoma | A0A0C2GVY3 |
| Ancylostoma | A0A0C2DNE1 |
| Angiostrongy | A0A0K0DQK0 |
| Angiostrongy | A0A0K0CXV8 |
| Angiostrongy | A0A0K0DEV9 |
| Angiostrongy | A0A158PCD7 |
| Angiostrongy | A0A0K0D0D1 |

| | |
|---|---|
| Angiostrongy | A0A0K0D3I0 |
| Angiostrongy | A0A0K0CUG2 |
| Angiostrongy | A0A0K0DH19 |
| Angiostrongy | A0A0K0D5J4 |
| Angiostrongy | A0A0K0D1B4 |
| Angiostrongy | A0A0K0DJR6 |
| Angiostrongy | A0A0K0D008 |
| Angiostrongy | A0A158PB05 |
| Angiostrongy | A0A158P5X6 |
| Angiostrongy | A0A158P8Q5 |
| Angiostrongy | A0A158PAM2 |
| Angiostrongy | A0A0K0DL22 |
| Angiostrongy | A0A3P7JT81 |
| Angiostrongy | A0A158PIZ8 |
| Angiostrongy | A0A3P7HR75 |
| Angiostrongy | A0A158PEZ2 |
| Angiostrongy | A0A3P7J3A0 |
| Angiostrongy | A0A158PKV1 |
| Angiostrongy | A0A0R3PDF2 |
| Angiostrongy | A0A158PGM2 |
| Angiostrongy | A0A158PE13 |
| Angiostrongy | A0A158PFJ6 |
| Angiostrongy | A0A0R3PME5 |
| Angiostrongy | A0A0R3PZF2 |
| Angiostrongy | A0A0R3PL23 |
| Angiostrongy | A0A0R3PMU4 |
| Angiostrongy | A0A0R3PYM1 |
| Angiostrongy | A0A158PJ68 |
| Angiostrongy | A0A158PLM5 |
| Angiostrongy | A0A0R3PB90 |
| Angiostrongy | A0A0R3Q188 |
| Angiostrongy | A0A158PGG5 |
| Angiostrongy | A0A0R3Q0R2 |
| Anguilla angu | A0A0E9X3V9 |
| Anguilla angu | A0A0E9RJH8 |
| Anisakis simp | A0A3P6NGQ5 |
| Anisakis simp | A0A0M3JX10 |
| Anisakis simp | A0A0M3K2F0 |
| Anisakis simp | A0A3P6SFK2 |
| Anisakis simp | A0A3P6RLJ8 |
| Anisakis simp | A0A0M3JD59 |

| | |
|---|---|
| Anisakis simp | A0A0M3KAT5 |
| Anisakis simp | A0A0M3KD76 |
| Anisakis simp | A0A3P6T6E9 |
| Anisakis simp | A0A3P6RTW8 |
| Anisakis simp | A0A0M3K8I9 |
| Anisakis simp | A0A0M3K2Z5 |
| Anisakis simp | A0A0M3J4P1 |
| Anisakis simp | A0A0M3J2X5 |
| Anisakis simp | A0A0M3JVW0 |
| Anisakis simp | A0A0M3J5I5 |
| Anisakis simp | A0A0M3KA02 |
| Anisakis simp | A0A0M3KCU0 |
| Anisakis simp | A0A0M3JKG2 |
| Anisakis simp | A0A0M3K200 |
| Anisakis simp | A0A0M3JKG6 |
| Anisakis simp | A0A0M3K1Z4 |
| Anisakis simp | A0A0M3KIH9 |
| Anisakis simp | A0A0M3K2C6 |
| Anisakis simp | A0A0M3JY76 |
| Anisakis simp | A0A158PN22 |
| Anisakis simp | A0A0M3K2T5 |
| Anisakis simp | A0A0M3J493 |
| Anolis carolin | G1KNL2 |
| Anolis carolin | G1K9G8 |
| Anolis carolin | G1KBV2 |
| Anolis carolin | G1KIB0 |
| Anolis carolin | R4GD12 |
| Anopheles all | A0A182F6L9 |
| Anopheles all | A0A182F5Z9 |
| Anopheles all | A0A182F6L8 |
| Anopheles all | A0A182FRE6 |
| Anopheles all | A0A182F6M0 |
| Anopheles all | A0A182F5Z8 |
| Anopheles all | A0A182FDG2 |
| Anopheles ar | A0A182HU77 |
| Anopheles ar | B2G0D9 |
| Anopheles ar | B2G0D5 |
| Anopheles ar | A0A182HTF9 |
| Anopheles ar | B2G0D4 |
| Anopheles ar | A0A182HTG0 |
| Anopheles ar | B2G0D3 |

| | |
|---|---|
| Anopheles ar | A0A182I7L3 |
| Anopheles ar | A0A2C9GRZ8 |
| Anopheles ar | A0A2C9GRP2 |
| Anopheles ar | A0A182I7L4 |
| Anopheles ar | A0A182HTF8 |
| Anopheles at | A0A182IT52 |
| Anopheles at | A0A182ITQ2 |
| Anopheles at | A0A182JLI3 |
| Anopheles at | A0A182ISL2 |
| Anopheles at | A0A182J7H9 |
| Anopheles br | A0A2M3Z8D8 |
| Anopheles ch | A0A182JNP2 |
| Anopheles ch | A0A182KB58 |
| Anopheles ch | A0A182JQG9 |
| Anopheles ch | A0A182KEB4 |
| Anopheles ch | A0A182KAY0 |
| Anopheles ch | A0A182JNP1 |
| Anopheles co | A0A182LFU3 |
| Anopheles co | A0A182LFM7 |
| Anopheles co | A0A182KQB0 |
| Anopheles co | A0A182LMG3 |
| Anopheles co | A0A182LFM6 |
| Anopheles cu | A0A182M3D4 |
| Anopheles cu | A0A182MI18 |
| Anopheles cu | A0A182MAX7 |
| Anopheles cu | A0A182M333 |
| Anopheles cu | A0A182M9X2 |
| Anopheles cu | A0A182LRC3 |
| Anopheles da | W5J6L5 |
| Anopheles da | W5J5H0 |
| Anopheles da | W5JK04 |
| Anopheles da | W5JQZ0 |
| Anopheles di | A0A182NGA9 |
| Anopheles di | A0A2Y9D1G5 |
| Anopheles di | A0A182NBM5 |
| Anopheles di | A0A182NBM6 |
| Anopheles di | A0A2Y9D1N7 |
| Anopheles di | A0A182N642 |
| Anopheles di | A0A182NGA8 |
| Anopheles di | A0A182NGA7 |
| Anopheles ep | A0A182P472 |

| | |
|---|---|
| Anopheles ep | A0A182PEK4 |
| Anopheles ep | A0A182NZH6 |
| Anopheles ep | A0A182NZH5 |
| Anopheles ep | A0A182PEK5 |
| Anopheles ep | A0A182PEK6 |
| Anopheles ep | A0A182P5T7 |
| Anopheles fa | A0A182QEI7 |
| Anopheles fa | A0A182QKG0 |
| Anopheles fa | A0A182QSK0 |
| Anopheles fa | A0A182QWD4 |
| Anopheles fa | A0A182QWK0 |
| Anopheles fa | A0A182Q5P2 |
| Anopheles fu | A0A182RNR5 |
| Anopheles fu | A0A182RHD6 |
| Anopheles fu | A0A182RF54 |
| Anopheles fu | A0A182R769 |
| Anopheles fu | A0A182RHD7 |
| Anopheles fu | A0A182R770 |
| Anopheles fu | A0A182R771 |
| Anopheles ga | Q7PTF3 |
| Anopheles ga | B2G0E1 |
| Anopheles ga | Q7QBU4 |
| Anopheles ga | Q7QD96 |
| Anopheles ga | Q7PWD7 |
| Anopheles ga | A0A1S4H0L9 |
| Anopheles ga | A0A1S4GZU1 |
| Anopheles ga | B2G0F5 |
| Anopheles ga | B2G0E5 |
| Anopheles m | A0A182T9U0 |
| Anopheles m | A0A182SV91 |
| Anopheles m | A0A182SSP0 |
| Anopheles m | A0A182SAF1 |
| Anopheles m | A0A182T826 |
| Anopheles m | A0A182T015 |
| Anopheles m | A0A182TPY7 |
| Anopheles m | A0A182TF95 |
| Anopheles m | A0A182UCD3 |
| Anopheles m | A0A182TQP9 |
| Anopheles m | A0A182TPR8 |
| Anopheles m | A0A182UC50 |
| Anopheles m | A0A182VE60 |

| | |
|---|---|
| Anopheles m | A0A182VK07 |
| Anopheles m | A0A2C9H5R2 |
| Anopheles m | A0A182V7G1 |
| Anopheles m | A0A182WGE9 |
| Anopheles m | A0A182WLF8 |
| Anopheles m | A0A182W062 |
| Anopheles m | A0A182W2N8 |
| Anopheles m | A0A182WGF0 |
| Anopheles m | A0A182W2N6 |
| Anopheles m | A0A182W2N7 |
| Anopheles qu | A0A182WWW6 |
| Anopheles qu | A0A182WXP9 |
| Anopheles qu | A0A182XM10 |
| Anopheles qu | A0A182XM08 |
| Anopheles qu | A0A182WXQ0 |
| Anopheles qu | A0A182XM09 |
| Anopheles qu | A0A182WRJ9 |
| Anopheles sir | A0A084VR87 |
| Anopheles sir | A0A084VZ46 |
| Anopheles sir | A0A084W3V7 |
| Anopheles sir | A0A084W3V8 |
| Anopheles st | A0A182XVF9 |
| Anopheles st | A0A182Y0J9 |
| Anopheles st | A0A182Y0K0 |
| Anopheles st | A0A182Y4A5 |
| Anopheles st | A0A182YEI6 |
| Anopheles st | A0A182XVG0 |
| Antarctobact | A0A222E9G5 |
| Antarctobact | A0A239JZP1 |
| Antarctobact | A0A239G1U7 |
| Antarctobact | A0A222E1N3 |
| Antarctobact | A0A222E9D9 |
| Anthurium ar | A0A1D1YV81 |
| Anthurium ar | A0A1D1YNY0 |
| Anthurium ar | A0A1D1Z1M7 |
| Anthurium ar | A0A1D1ZCW2 |
| Anthurium ar | A0A1D1XWH9 |
| Anthurium ar | A0A1D1XFS1 |
| Anthurium ar | A0A1D1ZCH5 |
| Anthurium ar | A0A1D1YQR8 |
| Anthurium ar | A0A1D1Y9Q8 |

| | |
|---|---|
| Anthurium ar | A0A1D1Y7P0 |
| Anthurium ar | A0A1D1YD96 |
| Anthurium ar | A0A1D1YYX4 |
| Anthurium ar | A0A1D1YCC3 |
| Anthurium ar | A0A1D1Z0E3 |
| Anthurium ar | A0A1D1Z3R3 |
| Anthurium ar | A0A1D1YIR8 |
| Anthurium ar | A0A1D1YXD0 |
| Anthurium ar | A0A1D1ZLQ1 |
| Anthurium ar | A0A1D1Y6E9 |
| Anthurium ar | A0A1D1XRS7 |
| Anthurium ar | A0A1D1XD27 |
| Anthurium ar | A0A1D1ZCJ4 |
| Anthurium ar | A0A1D1XNX6 |
| Anthurium ar | A0A1D1ZGK0 |
| Anthurium ar | A0A1D1YPM1 |
| Anthurium ar | A0A1D1XWC5 |
| Anthurium ar | A0A1D1YK26 |
| Anthurium ar | A0A1D1XXZ3 |
| Anthurium ar | A0A1D1YPA4 |
| Anthurium ar | A0A1D1Y8U2 |
| Anthurium ar | A0A1D1XS70 |
| Anthurium ar | A0A1D1ZHQ4 |
| Anthurium ar | A0A1D1ZBF3 |
| Anthurium ar | A0A1D1YFK6 |
| Anthurium ar | A0A1D1Y7K5 |
| Anthurium ar | A0A1D1YYY6 |
| Anthurium ar | A0A1D1YYC2 |
| Anthurium ar | A0A1D1Z4A8 |
| Anthurium ar | A0A1D1XX47 |
| Anthurium ar | A0A1D1YX08 |
| Anthurium ar | A0A1D1YFZ2 |
| Anthurium ar | A0A1D1ZJN6 |
| Anthurium ar | A0A1D1XYK2 |
| Anthurium ar | A0A1D1ZM53 |
| Anthurium ar | A0A1D1ZFV4 |
| Anthurium ar | A0A1D1YAP6 |
| Anthurium ar | A0A1D1Y229 |
| Anthurium ar | A0A1D1XT90 |
| Antrostomus | A0A094NGJ6 |
| Antrostomus | A0A094L822 |

| | |
|---|---|
| Antrostomus | A0A094K4U5 |
| Aotus nancyr | A0A2K5DLN0 |
| Aotus nancyr | A0A2K5CCL7 |
| Aotus nancyr | A0A2K5DLM5 |
| Aotus nancyr | A0A2K5DWJ7 |
| Aotus nancyr | A0A2K5DLU9 |
| Aotus nancyr | A0A2K5D8Z0 |
| Aotus nancyr | A0A2K5D8Y5 |
| Aotus nancyr | A0A2K5D923 |
| Apaloderma ' | A0A091N7G4 |
| Aphanomyce | A0A397C4L9 |
| Aphanomyce | A0A3R8DBH7 |
| Aphanomyce | A0A3R6Z9I5 |
| Aphanomyce | A0A3R6ZA35 |
| Aphanomyce | A0A397EGH5 |
| Aphanomyce | A0A418CK86 |
| Aphanomyce | W4H175 |
| Aphanomyce | W4FC62 |
| Aphanomyce | A0A3R7YB83 |
| Aphanomyce | A0A397C6I0 |
| Aphanomyce | W4H5C6 |
| Aphanomyce | W4G226 |
| Aphanomyce | A0A3R6Y2Z4 |
| Aphanomyce | A0A397D222 |
| Aphanomyce | A0A3L6VV49 |
| Aphanomyce | A0A3L6UWB2 |
| Aphanomyce | A0A397DZ46 |
| Aphanomyce | A0A3R7Y206 |
| Aphanomyce | W4G2B7 |
| Aphanomyce | A0A397EC98 |
| Aphanomyce | A0A396ZRW0 |
| Aphanomyce | A0A397CJH5 |
| Aphanomyce | A0A397D947 |
| Aphanomyce | W4F959 |
| Aphanomyce | A0A3R7B8C5 |
| Aphanomyce | W4G4H3 |
| Aphanomyce | A0A3L6V4T6 |
| Aphanomyce | A0A3R6XZD2 |
| Aphanomyce | A0A397EMC9 |
| Aphanomyce | A0A3L6VXF2 |
| Aphanomyce | A0A3L6VVV3 |

Aphanomyce A0A397FE69
Aphanomyce A0A397ER22
Aphanomyce A0A3R6ZU09
Aphanomyce W4GPE9
Aphanomyce A0A3R6Y1Q0
Aphanomyce A0A397DAZ4
Aphanomyce A0A397FMV1
Aphanomyce A0A418C2V2
Aphanomyce W4G2D3
Aphanomyce A0A425D7X0
Aphanomyce A0A3R7BUC6
Aphanomyce W4G048
Aphanomyce A0A418DV78
Aphanomyce A0A418FR37
Aphanomyce W4H736
Aphanomyce A0A397DY85
Aphanomyce A0A3R6Z315
Aphanomyce A0A418EI16
Aphanomyce A0A397BNB4
Aphanomyce A0A396ZQC4
Aphanomyce A0A3R7BMT9
Aphanomyce A0A397EYS8
Aphanomyce A0A396ZVZ8
Aphanomyce A0A397ELC4
Aphanomyce A0A418FSX8
Aphanomyce A0A418BVT6
Aphanomyce A0A3R7ABK1
Aphanomyce A0A3R7DKS6
Aphanomyce W4H234
Aphanomyce A0A397B129
Aphanomyce A0A425CKN8
Aphanomyce A0A418DLV8
Aphanomyce A0A418E0M0
Aphanomyce W4G2I9
Aphanomyce W4FM40
Aphanomyce A0A3L6VNC7
Aphanomyce A0A418EMF9
Aphanomyce W4FJ95
Aphanomyce A0A3L6UZY8
Aphanomyce A0A418BS39
Aphanomyce A0A3R6Y6L2

Aphanomyce W4FBD4
Aphanomyce A0A397BC55
Aphanomyce A0A3R6Z9Q1
Aphanomyce W4FAU3
Aphanomyce A0A396ZUR5
Aphanomyce A0A3R7B511
Aphanomyce A0A397CHM2
Aphanomyce W4FIU7
Aphanomyce A0A397AJZ3
Aphanomyce W4FDQ7
Aphanomyce A0A3L6V775
Aphanomyce A0A397BDL6
Aphanomyce A0A3R7AC59
Aphanomyce A0A397A4C7
Aphanomyce A0A3R8CZN9
Aphanomyce W4FZU6
Aphanomyce W4G6A2
Aphanomyce W4FEN0
Aphanomyce A0A3L6VU69
Aphanomyce A0A3L6VUC0
Aphanomyce A0A3R8D3V5
Aphanomyce A0A397DNW1
Aphanomyce A0A397E6J1
Aphanomyce A0A3R7AVF3
Aphanomyce A0A397CYV4
Aphanomyce A0A3R6WWC2
Aphanomyce A0A397EBU1
Aphanomyce A0A397C7D3
Aphanomyce W4GYA7
Aphanomyce A0A3L6VPA3
Aphanomyce A0A3R7CXJ7
Aphanomyce A0A397DAQ9
Aphanomyce A0A397B2Z6
Aphanomyce A0A397C063
Aphanomyce A0A397EVE7
Aphanomyce A0A397E7V6
Aphanomyce A0A397E2F2
Aphanomyce W4GZF9
Aphanomyce A0A397EAD8
Aphanomyce A0A397AMN7
Aphanomyce A0A397FFG6

Aphanomyce A0A3R7WJU8
Aphanomyce A0A397BRS4
Aphanomyce A0A397CFN8
Aphanomyce A0A397ATR5
Aphanomyce A0A397BCR8
Aphanomyce A0A3L6VBL8
Aphanomyce A0A397DIQ3
Aphanomyce A0A397CN17
Aphanomyce A0A3R6WP37
Aphanomyce A0A397D709
Aphanomyce A0A397FYU7
Aphanomyce A0A3R7C4F4
Aphanomyce W4GZ73
Aphanomyce W4H0H9
Aphanomyce A0A397G0U1
Aphanomyce A0A3L6VNX6
Aphanomyce A0A396ZPP3
Aphanomyce W4FES1
Aphanomyce A0A418D021
Aphanomyce A0A418FQK5
Aphanomyce A0A397CWE3
Aphanomyce A0A3R6XQ00
Aphanomyce A0A418F500
Aphanomyce A0A3R7E8Z5
Aphanomyce A0A3R6XAD8
Aphanomyce A0A3R7EK41
Aphanomyce A0A3R7AZA1
Aphanomyce A0A397EH51
Aphanomyce W4GXF5
Aphanomyce A0A3R6Y8H5
Aphanomyce A0A3R6ZZZ6
Aphanomyce W4GZE8
Aphanomyce A0A397DGM4
Aphanomyce A0A3R7BA06
Aphanomyce A0A397FI29
Aphanomyce A0A3R7F475
Aphanomyce W4FYV5
Aphanomyce A0A397FE42
Aphanomyce W4GBI6
Aphanomyce A0A397BA52
Aphanomyce A0A3L6VUC9

Aphanomyce A0A3R6WPT4
Aphanomyce A0A397BC03
Aphanomyce A0A397D6D7
Aphanomyce A0A397BR53
Aphanomyce A0A397FF82
Aphanomyce W4FKP1
Aphanomyce A0A396ZTJ8
Aphanomyce A0A397ASG3
Aphanomyce A0A418FQ73
Aphanomyce W4FWW2
Aphanomyce A0A418CSY7
Aphanomyce A0A418FU02
Aphanomyce A0A397CX05
Aphanomyce A0A418CAJ3
Aphanomyce A0A3R6XZB8
Aphanomyce A0A397EM03
Aphanomyce W4FWJ0
Aphanomyce A0A425DPY9
Aphanomyce A0A418DX93
Aphanomyce A0A397DI86
Aphanomyce A0A3R7AM78
Aphanomyce A0A418CFB1
Aphanomyce A0A3R6ZM03
Aphanomyce A0A397D336
Aphanomyce A0A397D2R9
Aphanomyce A0A397C5U1
Aphanomyce A0A3R6ZVR7
Aphanomyce A0A3R7WGA7
Aphanomyce A0A397C1U3
Aphanomyce A0A418BP49
Aphanomyce A0A397EL04
Aphanomyce A0A418DXN6
Aphanomyce A0A397B3A6
Aphanomyce A0A3R6Y8I4
Aphanomyce A0A3R7WIS6
Aphanomyce A0A418E5K5
Aphanomyce W4FMA4
Aphanomyce A0A425DM30
Aphanomyce A0A3R6XLJ9
Aphanomyce A0A3R7ZIF7
Aphanomyce A0A3L6UPJ1

Aphanomyce W4FYU2
Aphanomyce A0A418FGP0
Aphanomyce A0A3R7YVR4
Aphanomyce A0A3R6XCY1
Aphanomyce A0A397A0Q9
Aphanomyce A0A397DYY2
Aphanomyce A0A397A3C6
Aphanomyce A0A425CZ29
Aphanomyce A0A418EHM7
Aphanomyce A0A418CJF8
Aphanomyce A0A425DJP4
Aphanomyce A0A397DYB5
Aphanomyce A0A397B9P2
Aphanomyce A0A425D1E4
Aphanomyce W4G0V0
Aphanomyce A0A418CJF6
Aphanomyce A0A397B5E3
Aphanomyce A0A3L6VJM5
Aphanomyce A0A3R7Y395
Aphanomyce A0A418CUV0
Aphanomyce A0A418BSV9
Aphanomyce W4GEU8
Aphanomyce W4GHR1
Aphanomyce A0A425CPR1
Aphanomyce A0A3R7APT9
Aphanomyce A0A418FS09
Aphanomyce A0A397ES50
Aphanomyce A0A425CS30
Aphanomyce A0A418E063
Aphanomyce A0A397EAV1
Aphanomyce A0A3R7WQH8
Aphanomyce A0A3L6VZA1
Aphanomyce A0A418BJ04
Aphanomyce A0A3R6WYY8
Aphanomyce A0A397C1F4
Aphanomyce A0A397FHZ3
Aphanomyce A0A3R6WVI8
Aphanomyce A0A397CVV7
Aphanomyce A0A3L6VF87
Aphanomyce A0A397C940
Aphanomyce A0A397F613

Aphanomyce A0A3R7EHY7
Aphanomyce A0A397DXJ3
Aphanomyce A0A418DCN0
Aphanomyce A0A397ER77
Aphanomyce A0A418CWS7
Aphanomyce A0A3R7AHK3
Aphanomyce A0A3R6WXR4
Aphanomyce W4H7L0
Aphanomyce A0A418BZY3
Aphanomyce A0A397D456
Aphanomyce W4FRR2
Aphanomyce A0A397EV56
Aphanomyce W4FGT5
Aphanomyce A0A3L6VXT1
Aphanomyce A0A397BP25
Aphanomyce W4FGX0
Aphanomyce W4H101
Aphanomyce A0A397FDH7
Aphanomyce A0A425DDM5
Aphanomyce A0A3R7A4K4
Aphanomyce A0A397CJR0
Aphanomyce A0A3L6VMY5
Aphanomyce A0A418E4P5
Aphanomyce A0A418G3N5
Aphanomyce A0A397FAR0
Aphanomyce A0A3R6XZM3
Aphanomyce A0A3R7A7P4
Aphanomyce A0A425CZ94
Aphanomyce A0A418D8J3
Aphanomyce W4GL89
Aphanomyce A0A418FA61
Aphanomyce A0A397BAP7
Aphanomyce A0A396ZQ03
Aphanomyce A0A396ZZ96
Aphanomyce W4FL12
Aphanomyce A0A397ASM2
Aphanomyce A0A418C327
Aphanomyce A0A397EKX9
Aphanomyce A0A3R7AKW5
Aphanomyce A0A3R6XRX3
Aphanomyce A0A418FLE6

Aphanomyce W4FEW7
Aphanomyce A0A418E1G2
Aphanomyce A0A418DBB7
Aphanomyce A0A418CCH5
Aphanomyce A0A3R7Z8J7
Aphanomyce A0A418FGH1
Aphanomyce A0A418FYZ6
Aphanomyce A0A397EMD9
Aphanomyce A0A3R6XJW2
Aphanomyce A0A3R7Z6P2
Aphanomyce W4H6A6
Aphanomyce A0A418E9V7
Aphanomyce W4GDP9
Aphanomyce A0A3R7B8T5
Aphanomyce A0A397E2H2
Aphanomyce A0A418BIJ8
Aphanomyce A0A418E419
Aphanomyce A0A397EUF6
Aphanomyce A0A397D0N4
Aphanomyce A0A396ZWI2
Aphanomyce A0A425CZR5
Aphanomyce A0A397BFX1
Aphanomyce A0A3R6WLH3
Aphanomyce A0A397F2W6
Aphanomyce A0A418FPV6
Aphanomyce A0A397CC55
Aphanomyce A0A397EVK5
Aphanomyce A0A397CFM8
Aphanomyce A0A425DEZ5
Aphanomyce A0A397CGA6
Aphanomyce A0A3L6VYT2
Aphanomyce W4FBP1
Aphanomyce A0A3R6XF63
Aphanomyce A0A3L6VSI2
Aphanomyce A0A3L6V4Z0
Aphanomyce W4FLQ6
Aphanomyce A0A418BUX3
Aphanomyce A0A397D6T6
Aphanomyce A0A396ZRZ2
Aphanomyce A0A418E7S8
Aphanomyce A0A396ZT37

Aphanomyce A0A397ANE1
Aphanomyce A0A3L6VA85
Aphanomyce A0A397A932
Aphanomyce A0A3L6VJ13
Aphanomyce A0A3R6W2X3
Aphanomyce A0A3L6V2L6
Aphanomyce A0A3R6ZFR3
Aphanomyce A0A397A1S1
Aphanomyce W4FPQ1
Aphanomyce A0A3L6VS68
Aphanomyce A0A397BWJ0
Aphanomyce A0A397A9C7
Aphanomyce A0A3R6YID9
Aphanomyce A0A3R7BR36
Aphanomyce A0A3L6UUI8
Aphanomyce A0A3L6V0T7
Aphanomyce A0A3R7AMU4
Aphanomyce A0A397C9E7
Aphanomyce W4G1X8
Aphanomyce A0A3R6X795
Aphanomyce A0A397E6K8
Aphanomyce A0A3L6UW97
Aphanomyce A0A397AFG4
Aphanomyce W4FCE8
Aphanomyce A0A3R6VM88
Aphanomyce A0A3L6VTJ7
Aphanomyce A0A3L6V0S4
Aphanomyce A0A3L6VQP2
Aphanomyce W4F8J2
Aphanomyce A0A397BNT6
Aphanomyce A0A397BQ30
Aphanomyce A0A397AP79
Aphanomyce A0A3L6V0R4
Aphanomyce A0A397BQC2
Aphanomyce W4FI44
Aphanomyce A0A397B2U7
Aphanomyce A0A397CA05
Aphanomyce W4FB83
Aphanomyce A0A397ESK0
Aphanomyce A0A3R6W590
Aphanomyce A0A397BBH9

Aphanomyce W4GB84
Aphanomyce W4FRB0
Aphanomyce A0A397ESI1
Aphanomyce A0A397AHE1
Aphanomyce A0A397CD13
Aphanomyce A0A397ALA4
Aphanomyce A0A3R7XTQ9
Aphanomyce A0A397E2L9
Aphanomyce A0A397AY23
Aphanomyce A0A397BNE7
Aphanomyce A0A397A1D7
Aphanomyce W4G074
Aphanomyce W4GF49
Aphanomyce A0A3R7WNJ7
Aphanomyce A0A3L6V0A9
Aphanomyce W4FTM9
Aphanomyce A0A397C9P7
Aphanomyce W4FC56
Aphanomyce A0A397D5B2
Aphanomyce A0A3L6UVS9
Aphanomyce A0A3R8DIH2
Aphanomyce W4G054
Aphanomyce A0A397AF63
Aphanomyce A0A3R7AA07
Aphanomyce A0A3L6VW95
Aphanomyce W4G9H4
Aphanomyce A0A397DQE0
Aphanomyce A0A397CE78
Aphanomyce A0A397BWP1
Aphanomyce A0A3R7AS62
Aphanomyce A0A397BZC2
Aphanomyce A0A397F5K6
Aphanomyce A0A397CQE7
Aphanomyce W4FUT9
Aphanomyce A0A397BX76
Aphanomyce A0A397EWY4
Aphanomyce A0A3L6UTC6
Aphanomyce W4FCI5
Aphanomyce A0A3R7XUT3
Aphanomyce A0A397AF69
Aphanomyce A0A397FDL6

Aphanomyce A0A3R6W9N8
Aphanomyce A0A3R7FC57
Aphanomyce A0A3L6V861
Aphanomyce A0A397F9B1
Aphanomyce A0A397DBN1
Aphanomyce W4GGA2
Aphanomyce A0A397AHP6
Aphanomyce A0A397DW29
Aphanomyce A0A397A7S0
Aphanomyce A0A3L6VJZ3
Aphanomyce A0A397ED42
Aphanomyce A0A3R6WAM0
Aphanomyce W4GH53
Aphanomyce A0A397CR86
Aphanomyce A0A397BPQ1
Aphanomyce W4GZ49
Aphanomyce A0A397EYN3
Aphanomyce A0A397DRN3
Aphanomyce A0A397DXZ3
Aphanomyce A0A397A3K7
Aphanomyce W4FCU5
Aphanomyce A0A397B480
Aphanomyce A0A397C1F3
Aphanomyce A0A397EXX4
Aphanomyce W4H088
Aphanomyce A0A418D470
Aphanomyce A0A3R7A612
Aphanomyce A0A397ES75
Aphanomyce W4GXA1
Aphanomyce A0A397AYL9
Aphanomyce A0A3R6WDP1
Aphanomyce A0A397EYB4
Aphanomyce A0A418CY42
Aphanomyce A0A397G009
Aphanomyce A0A418FZC5
Aphanomyce A0A3L6V1Y4
Aphanomyce A0A3L6UQC2
Aphanomyce W4GFA8
Aphanomyce A0A3R6XZQ9
Aphanomyce A0A3R6Y2R4
Aphanomyce A0A024UHK3

| | |
|---|---|
| Aphanomyce | A0A024T9L5 |
| Aphanomyce | A0A3R7CTQ5 |
| Aphanomyce | A0A024TVG3 |
| Aphanomyce | A0A024UH93 |
| Aphanomyce | A0A3R7A1Q5 |
| Aphanomyce | A0A418AHC0 |
| Aphanomyce | A0A024TRP2 |
| Aphanomyce | A0A024TEL4 |
| Aphanomyce | A0A418AME6 |
| Aphanomyce | A0A3R6Y3Y5 |
| Aphanomyce | A0A3R6WGF2 |
| Aphanomyce | A0A3R6V2T3 |
| Aphanomyce | A0A024TPQ1 |
| Aphanomyce | A0A024UDS5 |
| Aphanomyce | A0A418AM46 |
| Aphanomyce | A0A024U7C9 |
| Aphanomyce | A0A3R6V2B0 |
| Aphanomyce | A0A024UQH4 |
| Aphanomyce | A0A024TK95 |
| Aphanomyce | A0A024UD11 |
| Aphanomyce | A0A024UDS2 |
| Aphanomyce | A0A418AND4 |
| Aphanomyce | A0A024UAK4 |
| Aphanomyce | A0A418ARH0 |
| Aphanomyce | A0A024UD26 |
| Aphanomyce | A0A418AHK8 |
| Aphanomyce | A0A418AMH7 |
| Aphanomyce | A0A418AJY7 |
| Aphanomyce | A0A3R6VR99 |
| Aphanomyce | A0A024TS56 |
| Aphanomyce | A0A024TUE0 |
| Aphanomyce | A0A024TWF3 |
| Aphanomyce | A0A3R6VF93 |
| Aphanomyce | A0A3R7A275 |
| Aphanomyce | A0A024UR54 |
| Aphanomyce | A0A3R6WDL0 |
| Aphanomyce | A0A024T8G1 |
| Aphanomyce | A0A024TWY4 |
| Aphanomyce | A0A024TCZ9 |
| Aphanomyce | A0A024UMA7 |
| Aphanomyce | A0A024UH78 |

| | |
|---|---|
| Aphanomyce | A0A024TCZ7 |
| Aphanomyce | A0A024TSD7 |
| Aphanomyce | A0A024TBE3 |
| Aphanomyce | A0A3R6Z6F7 |
| Aphanomyce | A0A024TFF2 |
| Aphanomyce | A0A024UC37 |
| Aphanomyce | A0A024U0X8 |
| Aphanomyce | A0A418B1G9 |
| Aphanomyce | A0A3R6VQU0 |
| Aphanomyce | A0A418ASS1 |
| Aphanomyce | A0A024UP64 |
| Aphanomyce | A0A3R6WNC5 |
| Aphanomyce | A0A3R7A440 |
| Aphanomyce | A0A418AQE9 |
| Aphanomyce | A0A024UQE9 |
| Aphanomyce | A0A024UBX1 |
| Aphanomyce | A0A024UNS3 |
| Aphanomyce | A0A024TDF0 |
| Aphanomyce | A0A024U4R8 |
| Aphanomyce | A0A3R6ZIP0 |
| Aphanomyce | A0A024U8U9 |
| Aphanomyce | A0A418AI42 |
| Aphanomyce | A0A024URA0 |
| Aphanomyce | A0A024T9U4 |
| Aphanomyce | A0A418B226 |
| Aphanomyce | A0A3R6VRZ5 |
| Aphanomyce | A0A3R7CVB0 |
| Aphanomyce | A0A3R6VPV8 |
| Apis cerana c | A0A2A3EFQ3 |
| Apis cerana c | A0A2A3EQR0 |
| Apis cerana c | A0A2A3EB22 |
| Apis cerana c | A0A2A3E6U1 |
| Apis cerana c | A0A2A3EPT1 |
| Apis cerana c | A0A2A3EC69 |
| Apis cerana.x | V9IEE8 |
| Apis mellifera | A0A087ZXY2 |
| Apis mellifera | A0A088AFH9 |
| Apis mellifera | A0A088AST6 |
| Apis mellifera | A0A088AC97 |
| Apis mellifera | A0A088ANX3 |
| Apis mellifera | A0A087ZV55 |

| | |
|---|---|
| Apis mellifera | A0A088AG76 |
| Apostasia she | A0A2I0ASJ4 |
| Apostasia she | A0A2H9ZXD3 |
| Apostasia she | A0A2I0B3M2 |
| Apostasia she | A0A2I0B5S4 |
| Apostasia she | A0A2I0B7B3 |
| Apostasia she | A0A2H9ZQW2 |
| Apostasia she | A0A2I0ADE7 |
| Apostasia she | A0A2I0AA30 |
| Apostasia she | A0A2I0B169 |
| Apostasia she | A0A2I0ABX1 |
| Apostasia she | A0A2I0AA27 |
| Apostasia she | A0A2I0B4N6 |
| Apostasia she | A0A2I0AQ30 |
| Apostasia she | A0A2I0AAT8 |
| Apostasia she | A0A2I0B7A3 |
| Apostasia she | A0A2I0B7B6 |
| Apostasia she | A0A2I0BFH8 |
| Apostasia she | A0A2I0A8W9 |
| Apostasia she | A0A2I0ACE4 |
| Apostasia she | A0A2I0A6S9 |
| Apostasia she | A0A2I0AA33 |
| Apostasia she | A0A2H9ZQW3 |
| Apostasia she | A0A2I0B912 |
| Apostasia she | A0A2H9ZYQ7 |
| Apostasia she | A0A2H9ZXB8 |
| Apostasia she | A0A2I0B2V6 |
| Apostasia she | A0A2H9ZX95 |
| Apostasia she | A0A2I0A9R7 |
| Aptenodytes | A0A087QH18 |
| Aptenodytes | A0A087QYC5 |
| Aptenodytes | A0A087QH19 |
| Aptenodytes | A0A087RE68 |
| Aquamicrobiu | A0A1I3QH19 |
| Aquamicrobiu | A0A1I3RZC0 |
| Aquamicrobiu | A0A0H1AF09 |
| Aquilegia coe | A0A2G5DNG3 |
| Aquilegia coe | A0A2G5EQ76 |
| Aquilegia coe | A0A2G5EQJ1 |
| Aquilegia coe | A0A2G5DCP1 |
| Aquilegia coe | A0A2G5C542 |

| | |
|---|---|
| Aquilegia coe | A0A2G5E5T2 |
| Aquilegia coe | A0A2G5EYF5 |
| Aquilegia coe | A0A2G5CKR8 |
| Aquilegia coe | A0A2G5D4L1 |
| Aquilegia coe | A0A2G5C571 |
| Aquilegia coe | A0A2G5CRQ3 |
| Aquilegia coe | A0A2G5EDS8 |
| Aquilegia coe | A0A2G5E7L7 |
| Aquilegia coe | A0A2G5E6D6 |
| Aquilegia coe | A0A2G5CRL5 |
| Aquilegia coe | A0A2G5C563 |
| Aquilegia coe | A0A2G5D4L6 |
| Aquilegia coe | A0A2G5E7H8 |
| Aquilegia coe | A0A2G5C847 |
| Aquilegia coe | A0A2G5ESB6 |
| Aquilegia coe | A0A2G5D4N4 |
| Aquilegia coe | A0A2G5CZF9 |
| Aquilegia coe | A0A2G5E7K7 |
| Aquilegia coe | A0A2G5DFA2 |
| Aquilegia coe | A0A2G5DIL6 |
| Aquilegia coe | A0A2G5ET28 |
| Aquilegia coe | A0A2G5D4L8 |
| Aquilegia coe | A0A2G5CH03 |
| Aquilegia coe | A0A2G5CZE7 |
| Aquilegia coe | A0A2G5DF97 |
| Aquilegia coe | A0A2G5CZE9 |
| Aquilegia coe | A0A2G5CRT5 |
| Aquilegia coe | A0A2G5EGS4 |
| Aquilegia coe | A0A2G5D562 |
| Aquilegia coe | A0A2G5EYR8 |
| Aquilegia coe | A0A2G5C581 |
| Aquilegia coe | A0A2G5DEI8 |
| Aquilegia coe | A0A2G5EYJ4 |
| Aquilegia coe | A0A2G5EYQ4 |
| Aquilegia coe | A0A2G5EYD9 |
| Aquilegia coe | A0A2G5D4M2 |
| Aquilegia coe | A0A2G5EYH4 |
| Aquilegia coe | A0A2G5EQJ0 |
| Aquilegia coe | A0A2G5D4M8 |
| Aquilegia coe | A0A2G5CRN7 |
| Aquilegia coe | A0A2G5ES32 |

| | |
|---|---|
| Aquilegia coe | A0A2G5CRM7 |
| Aquilegia coe | A0A2G5E7Q3 |
| Aquilegia coe | A0A2G5DFX8 |
| Aquilegia coe | A0A2G5CMW1 |
| Aquilegia coe | A0A2G5CRX1 |
| Aquilegia coe | A0A2G5CRI1 |
| Aquilegia coe | A0A2G5DG44 |
| Aquilegia coe | A0A2G5EYI8 |
| Aquilegia coe | A0A2G5EYI1 |
| Aquilegia coe | A0A2G5EYH5 |
| Aquilegia coe | A0A2G5EYG4 |
| Aquilegia coe | A0A2G5CRL0 |
| Aquilegia coe | A0A2G5D4K8 |
| Aquilegia coe | A0A2G5CGV5 |
| Aquilegia coe | A0A2G5DCI3 |
| Aquilegia coe | A0A2G5DCM8 |
| Aquilegia coe | A0A2G5E5V6 |
| Aquilegia coe | A0A2G5EYI0 |
| Aquilegia coe | A0A2G5D8X1 |
| Aquilegia coe | A0A2G5D4K5 |
| Aquilegia coe | A0A2G5CRR1 |
| Aquilegia coe | A0A2G5CRM4 |
| Aquilegia coe | A0A2G5E7J1 |
| Aquilegia coe | A0A2G5D5G9 |
| Aquilegia coe | A0A2G5E7K3 |
| Aquilegia coe | A0A2G5DE67 |
| Aquilegia coe | A0A2G5EYE0 |
| Aquilegia coe | A0A2G5DCF9 |
| Aquimonas v | A0A1G6XXE2 |
| Arabidopsis a | V5J679 |
| Arabidopsis a | V5J697 |
| Arabidopsis a | V5J681 |
| Arabidopsis a | V5J6E2 |
| Arabidopsis a | A0A3S9XJJ1 |
| Arabidopsis a | A0A451ES05 |
| Arabidopsis a | A0A451ES06 |
| Arabidopsis a | A0A3S5XGT4 |
| Arabidopsis a | V5J6V5 |
| Arabidopsis a | A0A3S5XGT3 |
| Arabidopsis a | A0A451ES07 |
| Arabidopsis a | V5J691 |

| | |
|---|---|
| Arabidopsis a | V5J6A0 |
| Arabidopsis a | V5J6E6 |
| Arabidopsis a | V5J6V1 |
| Arabidopsis a | V5J6X1 |
| Arabidopsis a | A0A3S5XGT1 |
| Arabidopsis a | A0A3Q9JP76 |
| Arabidopsis a | A0A3S5XGT0 |
| Arabidopsis a | V5J698 |
| Arabidopsis a | V5J688 |
| Arabidopsis a | V5J689 |
| Arabidopsis a | A0A3Q9JP70 |
| Arabidopsis a | A0A3Q9JP72 |
| Arabidopsis a | V5J682 |
| Arabidopsis a | V5J699 |
| Arabidopsis a | V5J6E4 |
| Arabidopsis a | A0A3Q9JP73 |
| Arabidopsis a | V5J692 |
| Arabidopsis a | V5J6E9 |
| Arabidopsis a | V5J6F1 |
| Arabidopsis a | V5J6W1 |
| Arabidopsis a | V5J690 |
| Arabidopsis a | V5J683 |
| Arabidopsis a | A0A451ES04 |
| Arabidopsis a | A0A3Q9JP74 |
| Arabidopsis a | A0A451ES08 |
| Arabidopsis a | V5J6W5 |
| Arabidopsis a | A0A3S5XGT2 |
| Arabidopsis a | A0A3Q9JP77 |
| Arabidopsis a | A0A451ES03 |
| Arabidopsis a | V5J680 |
| Arabidopsis h | I7EWH2 |
| Arabidopsis h | I7FWN1 |
| Arabidopsis h | D6NU58 |
| Arabidopsis h | I7FWN4 |
| Arabidopsis h | B2Y4N4 |
| Arabidopsis h | B1P3G7 |
| Arabidopsis h | D0EXX6 |
| Arabidopsis h | I4IY52 |
| Arabidopsis h | D0EXW2 |
| Arabidopsis h | B1P3E5 |
| Arabidopsis h | D0EXX4 |

| | |
|---|---|
| Arabidopsis h | B1P3K8 |
| Arabidopsis h | D0EXY1 |
| Arabidopsis h | A7XE46 |
| Arabidopsis h | B1P3H0 |
| Arabidopsis h | A0A0C5JYN3 |
| Arabidopsis h | D0EXW0 |
| Arabidopsis h | D0EXW6 |
| Arabidopsis h | A7XE66 |
| Arabidopsis h | B1P3I4 |
| Arabidopsis h | A7XE70 |
| Arabidopsis h | A0A068CIR0 |
| Arabidopsis h | A7XE48 |
| Arabidopsis h | A0A068CIP9 |
| Arabidopsis h | A0A0C5K603 |
| Arabidopsis h | A0A0C5K8H5 |
| Arabidopsis h | A0A0C5K8B3 |
| Arabidopsis h | D0EXY8 |
| Arabidopsis h | D0EXX7 |
| Arabidopsis h | A0A068CQG1 |
| Arabidopsis h | Q19MC1 |
| Arabidopsis h | B1P3I7 |
| Arabidopsis h | A0A068CM44 |
| Arabidopsis h | A0A0C5KK99 |
| Arabidopsis h | D0EXY2 |
| Arabidopsis h | B1P3G3 |
| Arabidopsis h | A0A0C5KK72 |
| Arabidopsis h | A7XE76 |
| Arabidopsis h | B1P3I5 |
| Arabidopsis h | Q19MC0 |
| Arabidopsis h | B5TD10 |
| Arabidopsis h | B1P3I6 |
| Arabidopsis h | A0A068CM61 |
| Arabidopsis h | D0EXX0 |
| Arabidopsis h | A7XE72 |
| Arabidopsis h | B1P3G6 |
| Arabidopsis h | A0A0C5K5V8 |
| Arabidopsis h | D0EXY3 |
| Arabidopsis h | A7XE85 |
| Arabidopsis h | D0EXW8 |
| Arabidopsis h | A7XE56 |
| Arabidopsis h | B5TD17 |

| | |
|---|---|
| Arabidopsis h | B5TD13 |
| Arabidopsis h | B5TD14 |
| Arabidopsis h | A7XE74 |
| Arabidopsis h | A7XE52 |
| Arabidopsis h | A7XE63 |
| Arabidopsis h | A7XE50 |
| Arabidopsis h | D0EXY7 |
| Arabidopsis h | A7XE68 |
| Arabidopsis h | B1P3K0 |
| Arabidopsis h | A0A068CM37 |
| Arabidopsis h | D0EXY0 |
| Arabidopsis h | D0EXW5 |
| Arabidopsis h | B1P3K4 |
| Arabidopsis h | A7XE58 |
| Arabidopsis h | B1P3H3 |
| Arabidopsis h | A7XE83 |
| Arabidopsis h | A0A0C5K8F4 |
| Arabidopsis h | A7XE60 |
| Arabidopsis h | A7XE64 |
| Arabidopsis h | A0A0C5K5X4 |
| Arabidopsis h | D0EXV9 |
| Arabidopsis h | D0EXY6 |
| Arabidopsis h | A0A0C5K8D6 |
| Arabidopsis h | B5TD16 |
| Arabidopsis h | D0EXX1 |
| Arabidopsis h | A7XE81 |
| Arabidopsis h | A7XE79 |
| Arabidopsis h | D0EXW1 |
| Arabidopsis h | A7XE54 |
| Arabidopsis h | B5TD11 |
| Arabidopsis h | B5TD12 |
| Arabidopsis h | I4IY53 |
| Arabidopsis h | B5TD15 |
| Arabidopsis h | B1P3J6 |
| Arabidopsis h | A0A068CQH3 |
| Arabidopsis h | A0A068CM51 |
| Arabidopsis h | I4IY51 |
| Arabidopsis h | B1P3E8 |
| Arabidopsis h | B1P3F7 |
| Arabidopsis h | D0EXW7 |
| Arabidopsis h | B1P3G8 |

| | |
|---|---|
| Arabidopsis k | I7FNZ1 |
| Arabidopsis k | I7FXW5 |
| Arabidopsis k | I7EWH7 |
| Arabidopsis k | I7FWN7 |
| Arabidopsis k | I7FNZ6 |
| Arabidopsis k | I7FXW2 |
| Arabidopsis k | I7EWG7 |
| Arabidopsis k | I7F535 |
| Arabidopsis l | D7KVV7 |
| Arabidopsis l | D7KW17 |
| Arabidopsis l | D7KW16 |
| Arabidopsis l | D7KVK3 |
| Arabidopsis l | D7MEU5 |
| Arabidopsis l | D7KMD6 |
| Arabidopsis l | D7KZN2 |
| Arabidopsis l | D7M0B8 |
| Arabidopsis l | D7MEH3 |
| Arabidopsis l | D7KSZ5 |
| Arabidopsis l | D7KVV6 |
| Arabidopsis l | D7LFY7 |
| Arabidopsis l | D7KV44 |
| Arabidopsis l | D7KW05 |
| Arabidopsis l | D7M4K4 |
| Arabidopsis l | D7KSZ4 |
| Arabidopsis l | D7KW07 |
| Arabidopsis l | D7KVI7 |
| Arabidopsis l | D7KMD5 |
| Arabidopsis l | D7KMC8 |
| Arabidopsis l | D7KW12 |
| Arabidopsis l | D7KW14 |
| Arabidopsis l | D7MWF8 |
| Arabidopsis l | D7MJM6 |
| Arabidopsis l | D7KW03 |
| Arabidopsis l | D7KEK1 |
| Arabidopsis l | D7KVJ8 |
| Arabidopsis l | D7LTX6 |
| Arabidopsis l | D7MUF3 |
| Arabidopsis l | D7KW18 |
| Arabidopsis l | D7KYK2 |
| Arabidopsis l | D7L500 |
| Arabidopsis l | D7KMD4 |

| | |
|---|---|
| Arabidopsis l | D7KW15 |
| Arabidopsis l | D7MJN8 |
| Arabidopsis l | D7MEU2 |
| Arabidopsis l | D7KW04 |
| Arabidopsis l | D7MWF7 |
| Arabidopsis l | D7LHZ3 |
| Arabidopsis l | D7M9K1 |
| Arabidopsis l | D7KMD7 |
| Arabidopsis l | D7MEU3 |
| Arabidopsis l | D7M1U9 |
| Arabidopsis l | D7M2R0 |
| Arabidopsis l | D7LIA6 |
| Arabidopsis l | D7KMD1 |
| Arabidopsis l | D7KU49 |
| Arabidopsis l | D7MEH4 |
| Arabidopsis l | D7KME3 |
| Arabidopsis l | D7L1X6 |
| Arabidopsis l | D7LTL2 |
| Arabidopsis l | D7KMC5 |
| Arabidopsis l | D7M5Q3 |
| Arabidopsis l | D7MJG2 |
| Arabidopsis l | V5J6F9 |
| Arabidopsis l | V5J6F7 |
| Arabidopsis l | V5J693 |
| Arabidopsis l | V5J6B0 |
| Arabidopsis l | A0A060D3V0 |
| Arabidopsis l | V5J695 |
| Arabidopsis l | V5J6Y3 |
| Arabidopsis l | V5J6X7 |
| Arabidopsis l | V5J6F3 |
| Arabidopsis l | V5J686 |
| Arabidopsis l | V5J6A1 |
| Arabidopsis l | V5J6F5 |
| Arabidopsis l | V5J6A8 |
| Arabidopsis l | V5J6Z0 |
| Arabidopsis l | V5J6Z6 |
| Arabidopsis l | V5J6A3 |
| Arabidopsis l | V5J696 |
| Arabidopsis l | V5J684 |
| Arabidopsis l | V5J6A6 |
| Arabidopsis l | V5J694 |

| | |
|---|---|
| Arabidopsis l | D0EXX3 |
| Arabidopsis l | D0EXZ4 |
| Arabidopsis l | G4WHA0 |
| Arabidopsis l | B5TD25 |
| Arabidopsis l | L7UVM3 |
| Arabidopsis l | Q84XJ6 |
| Arabidopsis l | G4WH68 |
| Arabidopsis l | A0A068CIS1 |
| Arabidopsis l | A0A451ERZ8 |
| Arabidopsis l | A0A451ES00 |
| Arabidopsis l | Q9ATK3 |
| Arabidopsis l | Q19MB7 |
| Arabidopsis l | A0A068CL41 |
| Arabidopsis l | Q19MB2 |
| Arabidopsis l | Q9ATK0 |
| Arabidopsis l | Q9ATK4 |
| Arabidopsis l | G4WH69 |
| Arabidopsis l | Q84XI9 |
| Arabidopsis l | Q84XK2 |
| Arabidopsis l | C3VIF0 |
| Arabidopsis l | Q27JI1 |
| Arabidopsis l | A0A068CQL8 |
| Arabidopsis l | Q9ATK2 |
| Arabidopsis l | A0A3Q9JP26 |
| Arabidopsis l | R9S9V2 |
| Arabidopsis l | Q19MB5 |
| Arabidopsis l | B5TD23 |
| Arabidopsis l | A0A068CIT0 |
| Arabidopsis l | A0A068CIT5 |
| Arabidopsis l | Q19MB6 |
| Arabidopsis l | A0A3Q9JP81 |
| Arabidopsis l | Q84XK1 |
| Arabidopsis l | A0A068CLV9 |
| Arabidopsis l | B5TD21 |
| Arabidopsis l | Q9ATJ7 |
| Arabidopsis l | Q19MB4 |
| Arabidopsis l | Q84XK3 |
| Arabidopsis l | G4WH91 |
| Arabidopsis l | Q19MB0 |
| Arabidopsis l | Q84XJ0 |
| Arabidopsis l | D0EXX2 |

| | |
|---|---|
| Arabidopsis l | Q9AVE1 |
| Arabidopsis l | G4WH89 |
| Arabidopsis l | Q19MB8 |
| Arabidopsis l | Q84XJ2 |
| Arabidopsis l | Q84XJ5 |
| Arabidopsis l | G4WH79 |
| Arabidopsis l | Q84XJ4 |
| Arabidopsis l | A0A1J0GTD5 |
| Arabidopsis l | Q9ATJ5 |
| Arabidopsis l | B5TD20 |
| Arabidopsis l | G4WH80 |
| Arabidopsis l | A0A451ERZ9 |
| Arabidopsis l | A0A451ES01 |
| Arabidopsis l | G4WH99 |
| Arabidopsis l | C7FE07 |
| Arabidopsis l | A0A068CLU4 |
| Arabidopsis l | A0A3S9XJG2 |
| Arabidopsis l | A0A068CM84 |
| Arabidopsis l | A0A451ES02 |
| Arabidopsis l | A0A068CL61 |
| Arabidopsis l | Q84XJ9 |
| Arabidopsis l | A0A3Q9JP24 |
| Arabidopsis l | Q84XJ1 |
| Arabidopsis l | A0A3Q9JP42 |
| Arabidopsis l | B5TD24 |
| Arabidopsis l | B5TD18 |
| Arabidopsis l | A0A068CQJ3 |
| Arabidopsis l | B5TD19 |
| Arabidopsis l | A0A3S5XGS9 |
| Arabidopsis l | B5TD27 |
| Arabidopsis l | Q19MB3 |
| Arabidopsis l | G4WH92 |
| Arabidopsis l | A0A3S5XGS8 |
| Arabidopsis l | A8IKA8 |
| Arabidopsis l | Q9ATJ4 |
| Arabidopsis l | G4WHA1 |
| Arabidopsis l | Q9ATJ6 |
| Arabidopsis l | Q9ATK1 |
| Arabidopsis l | Q84XI8 |
| Arabidopsis l | B5TD26 |
| Arabidopsis l | Q84XI4 |

| | |
|---|---|
| Arabidopsis l | Q9ATJ9 |
| Arabidopsis l | Q84XI7 |
| Arabidopsis l | Q84XJ3 |
| Arabidopsis l | D0EXV6 |
| Arabidopsis l | Q84XI6 |
| Arabidopsis l | G4WH81 |
| Arabidopsis l | Q84XJ7 |
| Arabidopsis l | B5TD22 |
| Arabidopsis l | Q9ATJ8 |
| Arabidopsis l | C7FE06 |
| Arabidopsis l | R9QT99 |
| Arabidopsis l | Q84XJ8 |
| Arabidopsis l | Q84XI5 |
| Arabidopsis l | D0EXV5 |
| Arabidopsis l | Q9AVE0 |
| Arabidopsis l | Q27JF2 |
| Arabidopsis l | Q27JG5 |
| Arabidopsis t | Q9LPZ9 |
| Arabidopsis t | P0DH86 |
| Arabidopsis t | Q9LW83 |
| Arabidopsis t | Q9S972 |
| Arabidopsis t | O64782 |
| Arabidopsis t | Q39086 |
| Arabidopsis t | O81905 |
| Arabidopsis t | Q39203 |
| Arabidopsis t | O81833 |
| Arabidopsis t | Q39202 |
| Arabidopsis t | Q9ZR08 |
| Arabidopsis t | O64477 |
| Arabidopsis t | Q9LZR8 |
| Arabidopsis t | Q9SXB3 |
| Arabidopsis t | Q9SXB4 |
| Arabidopsis t | O64784 |
| Arabidopsis t | Q9LPZ3 |
| Arabidopsis t | O64778 |
| Arabidopsis t | O64776 |
| Arabidopsis t | O64771 |
| Arabidopsis t | Q9ZVA1 |
| Arabidopsis t | Q9ZVA4 |
| Arabidopsis t | Q9ZVA5 |
| Arabidopsis t | O81906 |

Arabidopsis t P0DH87
Arabidopsis t Q9SXB8
Arabidopsis t Q9SXB5
Arabidopsis t O64783
Arabidopsis t O64774
Arabidopsis t Q9SY95
Arabidopsis t O64780
Arabidopsis t O64770
Arabidopsis t O64781
Arabidopsis t Q9SY89
Arabidopsis t Q9SYA0
Arabidopsis t O64793
Arabidopsis t O64777
Arabidopsis t Q9ZT07
Arabidopsis t P93756
Arabidopsis t Q9T058
Arabidopsis t O81832
Arabidopsis t A0A1P8AN24
Arabidopsis t A0A1P8B6N6
Arabidopsis t A0A1P8AM95
Arabidopsis t Q9SCT7
Arabidopsis t A0A1P8B6M9
Arabidopsis t A0A1P8AS04
Arabidopsis t A0A1P8AVJ2
Arabidopsis t Q9LHM1
Arabidopsis t A0A178VV00
Arabidopsis t A0A1P8B912
Arabidopsis t A0A178V611
Arabidopsis t A0A1P8B838
Arabidopsis t K9LQT0
Arabidopsis t A0A178WKT1
Arabidopsis t A0A178W8X6
Arabidopsis t A0A1P8AM91
Arabidopsis t A0A178WAM2
Arabidopsis t A0A178W4F8
Arabidopsis t A0A178WMJ8
Arabidopsis t A0A178WCF7
Arabidopsis t A0A384KCW1
Arabidopsis t A0A1P8AN12
Arabidopsis t K9LQP6
Arabidopsis t A0A178V4X9

Arabidopsis t A0A178WEA5
Arabidopsis t A0A178V3N4
Arabidopsis t A0A178V2J5
Arabidopsis t A0A1P8B6N5
Arabidopsis t A0A178WN39
Arabidopsis t A0A178UWV5
Arabidopsis t A0A178WFA7
Arabidopsis t A0A1P8AU51
Arabidopsis t F4I7F8
Arabidopsis t A0A1P8AS39
Arabidopsis t A0A1P8AVQ7
Arabidopsis t Q9FVW1
Arabidopsis t A0A1P8B837
Arabidopsis t A0A1P8B840
Arabidopsis t A0A384KEE5
Arabidopsis t A0A1P8AN14
Arabidopsis t A0A178UYZ8
Arabidopsis t A0A1I9LPL1
Arabidopsis t A0A1P8B600
Arabidopsis t L7USG8
Arabidopsis t A0A178V6V9
Arabidopsis t A0A1P8ATF5
Arabidopsis t A0A178VFP7
Arabidopsis t A0A178WA58
Arabidopsis t A0A1P8AWX0
Arabidopsis t A0A178WDF4
Arabidopsis t A0A178WC58
Arabidopsis t A0A178WA81
Arabidopsis t A0A178WI76
Arabidopsis t A0A178VYR4
Arabidopsis t A0A1P8ARZ8
Arabidopsis t A0A1P8AMD4
Arabidopsis t A0A1P8B859
Arabidopsis t A0A1I9LPL5
Arabidopsis t A0A1P8AMY7
Arabidopsis t A0A178WJN4
Arabidopsis t A0A178WJI0
Arabidopsis t A0A1P8ATC6
Arabidopsis t A0A1P8ARD8
Arabidopsis t A0A1P8AU37
Arabidopsis t A0A178WFP0

| | |
|---|---|
| Arabidopsis t | A0A178V0J5 |
| Arabidopsis t | A0A178VNJ9 |
| Arabidopsis t | A0A1P8ASG9 |
| Arabidopsis t | A0A1P8ASF1 |
| Arabidopsis t | A0A178WB58 |
| Arabidopsis t | A0A178W2U0 |
| Arabidopsis t | A0A1P8AMT2 |
| Arabidopsis t | A0A178W9H1 |
| Arabidopsis t | A0A1P8AM94 |
| Arabidopsis t | Q41272 |
| Arabidopsis t | A0A178WDZ2 |
| Arabidopsis t | F4I8U6 |
| Arabidopsis t | A0A1P8AVL2 |
| Arabidopsis t | A0A1P8AWT8 |
| Arabidopsis t | A0A1P8ASM5 |
| Arabidopsis t | A0A178UVF9 |
| Arabidopsis t | A0A1P8AN35 |
| Arabidopsis t | A0A178USE7 |
| Arabidopsis t | A0A1P8B5P5 |
| Arabidopsis t | A0A1I9LPL2 |
| Arabidopsis t | A0A1P8AMA7 |
| Arabidopsis t | A0A1P8B844 |
| Arabidopsis t | A0A1P8AM92 |
| Arabidopsis t | A0A1P8AQZ9 |
| Arabidopsis t | A0A1P8AMR1 |
| Arabidopsis t | A0A178WDY1 |
| Arabidopsis t | A0A1P8AMR8 |
| Arabidopsis t | A0A1P8AN06 |
| Arabidopsis t | A0A1P8AQN4 |
| Arabidopsis t | A7Y5W8 |
| Arabidopsis t | A0A1P8AMA0 |
| Arabidopsis t | A0A178W877 |
| Arabidopsis t | A0A178WD43 |
| Arabidopsis t | A0A1P8AN03 |
| Arabidopsis t | A0A1P8AMA3 |
| Arabidopsis t | A0A178UCA7 |
| Arabidopsis t | A0A178WK46 |
| Arabidopsis t | A0A1I9LPL3 |
| Arabidopsis t | A0A1P8B6M8 |
| Arabidopsis t | A0A1P8AVI8 |
| Arabidopsis t | A0A178WL67 |

| | |
|---|---|
| Arabidopsis t | A0A1I9LPL6 |
| Arabidopsis t | A0A178WAR7 |
| Arabidopsis t | A0A1P8B6N1 |
| Arabidopsis t | A0A1P8AW73 |
| Arabis alpina | A0A087GL67 |
| Arabis alpina | A0A087HF14 |
| Arabis alpina | A0A087G4N3 |
| Arabis alpina | A0A087HEZ2 |
| Arabis alpina | A0A087HEY3 |
| Arabis alpina | A0A087H0M1 |
| Arabis alpina | A0A087GH82 |
| Arabis alpina | A0A087GH84 |
| Arabis alpina | A0A087G682 |
| Arabis alpina | A0A087H3L7 |
| Arabis alpina | A0A087H3L8 |
| Arabis alpina | A0A087GFR5 |
| Arabis alpina | A0A087GFR4 |
| Arabis alpina | A0A087H8W0 |
| Arabis alpina | A0A087HEZ7 |
| Arabis alpina | A0A087HJG0 |
| Arabis alpina | A0A087GDA1 |
| Arabis alpina | A0A087HEZ3 |
| Arabis alpina | A0A087GH83 |
| Arabis alpina | A0A087HEZ6 |
| Arabis alpina | A0A087GX11 |
| Arabis alpina | A0A087H0M6 |
| Arabis alpina | A0A087GPZ0 |
| Arabis alpina | A0A087H0M5 |
| Arabis alpina | A0A087GSX0 |
| Arabis alpina | A0A087HCS1 |
| Arabis alpina | A0A087GDA2 |
| Arachis hypo | A0A445BB21 |
| Arachis hypo | A0A444YSR3 |
| Arachis hypo | A0A444XXG8 |
| Arachis hypo | A0A444Y4Z7 |
| Arachis hypo | A0A445C105 |
| Arachis hypo | A0A444WXX8 |
| Arachis hypo | A0A445DZ26 |
| Arachis hypo | A0A445C101 |
| Arachis hypo | A0A444XXT2 |
| Arachis hypo | A0A445BY59 |

| | |
|---|---|
| Arachis hypo{ | A0A445BY66 |
| Arachis hypo{ | A0A444XD73 |
| Arachis hypo{ | A0A445CY83 |
| Arachis hypo{ | A0A445BYC6 |
| Arachis hypo{ | A0A445BY70 |
| Arachis hypo{ | A0A445BYB8 |
| Arachis hypo{ | A0A445DYY7 |
| Arachis hypo{ | A0A445ECY2 |
| Arachis hypo{ | A0A445BYJ2 |
| Arachis hypo{ | A0A445CQB5 |
| Arachis hypo{ | A0A445BS77 |
| Arachis hypo{ | A0A445CI63 |
| Arachis hypo{ | A0A445BYE2 |
| Arachis hypo{ | A0A445CU59 |
| Arachis hypo{ | A0A445C0Z1 |
| Arachis hypo{ | A0A444Y4V5 |
| Arachis hypo{ | A0A445AGS4 |
| Arachis hypo{ | A0A444Y4X6 |
| Arachis hypo{ | A0A445BYN4 |
| Arachis hypo{ | A0A445AFB2 |
| Arachis hypo{ | A0A445DZ08 |
| Arachis hypo{ | A0A445BP40 |
| Arachis hypo{ | A0A445BB53 |
| Arachis hypo{ | A0A444Y4Z4 |
| Arachis hypo{ | A0A445BB59 |
| Arachis hypo{ | A0A445BB22 |
| Arachis hypo{ | A0A445C0Z7 |
| Arachis hypo{ | A0A445ECU2 |
| Arachis hypo{ | A0A444Y554 |
| Arachis hypo{ | A0A445BY98 |
| Arachis hypo{ | A0A444WNK3 |
| Arachis hypo{ | A0A445BB41 |
| Arachis hypo{ | A0A444Y534 |
| Arachis hypo{ | A0A445DYV4 |
| Arachis hypo{ | A0A444YP61 |
| Arachis hypo{ | A0A445BX15 |
| Arachis hypo{ | A0A444XXD3 |
| Arachis hypo{ | A0A445BYB1 |
| Arachis hypo{ | A0A444Y584 |
| Arachis hypo{ | A0A445DZ14 |
| Arachis hypo{ | A0A445BYB2 |

| | |
|---|---|
| Arachis hypo | A0A445A4A8 |
| Arachis hypo | A0A444WXZ0 |
| Arachis hypo | A0A445BYA3 |
| Arachis hypo | A0A444WNL4 |
| Arachis hypo | A0A445A406 |
| Arachis hypo | A0A445BX05 |
| Arachis hypo | A0A445ABG7 |
| Arachis hypo | A0A444WY41 |
| Arachis hypo | A0A444Y4V2 |
| Arachis hypo | A0A444YPC9 |
| Arachis hypo | A0A445DRJ0 |
| Arachis hypo | A0A445CHB6 |
| Arachis hypo | A0A444WY06 |
| Arachis hypo | A0A444Y6V2 |
| Arachis hypo | A0A444Z5W0 |
| Arachis hypo | A0A445DZ35 |
| Arachis hypo | A0A444XXE0 |
| Arachis hypo | A0A445DZ07 |
| Arachis hypo | A0A444WXX4 |
| Arachis hypo | A0A444Y4Y6 |
| Arachis hypo | A0A445BB20 |
| Arachis hypo | A0A444WXY1 |
| Arachis hypo | A0A445BYA9 |
| Arachis hypo | A0A445BY84 |
| Arachis hypo | A0A445C0Z5 |
| Arachis hypo | A0A445BYW2 |
| Arachis hypo | A0A445ED07 |
| Arachis hypo | A0A444WY99 |
| Arachis hypo | A0A445BB37 |
| Arachis hypo | A0A444Y4W1 |
| Arachis hypo | A0A444Y4Y7 |
| Arachis hypo | A0A445DYT5 |
| Arachis hypo | A0A445BY88 |
| Arachis hypo | A0A444Y526 |
| Arachis hypo | A0A444Y2G7 |
| Arachis hypo | A0A445DRJ8 |
| Arachis hypo | A0A445DRJ4 |
| Arachis hypo | A0A444Y562 |
| Arachis hypo | A0A445DZ00 |
| Arachis hypo | A0A444Y519 |
| Arachis hypo | A0A445BP74 |

| | |
|---|---|
| Arachis hypo | A0A444XBK0 |
| Arachis hypo | A0A444Y3V7 |
| Arachis hypo | A0A444Y6P6 |
| Arachis hypo | A0A445BY81 |
| Arachis hypo | A0A444WXZ1 |
| Arachis hypo | A0A445BYT7 |
| Arachis hypo | A0A445BY74 |
| Arachis hypo | A0A445BB43 |
| Arachis hypo | A0A444WXZ6 |
| Arachis hypo | A0A445CI88 |
| Arachis hypo | A0A445BYA0 |
| Arachis hypo | A0A445ECS9 |
| Arachis hypo | A0A445BB25 |
| Arachis hypo | A0A444Y532 |
| Arachis hypo | A0A445ECJ8 |
| Arachis hypo | A0A445BYD0 |
| Arachis hypo | A0A445CI74 |
| Arachis hypo | A0A445BB51 |
| Arachis hypo | A0A444YRD4 |
| Arachis hypo | A0A444Y514 |
| Arachis hypo | A0A444Y537 |
| Arachis hypo | A0A444XDC3 |
| Arachis hypo | A0A445A439 |
| Arachis hypo | A0A445EIW0 |
| Arachis hypo | A0A444Y4Y9 |
| Arachis hypo | A0A444Y6R2 |
| Arachis hypo | A0A444WY60 |
| Arachis hypo | A0A445A5E9 |
| Arachis hypo | A0A445BYB4 |
| Arachis hypo | A0A444Y6M3 |
| Arachis hypo | A0A444Y4X4 |
| Arachis hypo | A0A445C0Z4 |
| Arachis hypo | A0A445BYX5 |
| Arachis hypo | A0A445BYX9 |
| Arachis hypo | A0A445BB31 |
| Arachis hypo | A0A445BY77 |
| Arachis hypo | A0A445BYA5 |
| Arachis hypo | A0A445CI77 |
| Arachis hypo | A0A445E8F6 |
| Arachis hypo | A0A445AII5 |
| Arachis hypo | A0A445BYX2 |

| | |
|---|---|
| Arachis hypo{ | A0A445BYE0 |
| Arachis hypo{ | A0A445CBB0 |
| Arachis hypo{ | A0A444Y566 |
| Arachis hypo{ | A0A445DRJ3 |
| Arachis hypo{ | A0A444XD58 |
| Arachis hypo{ | A0A445DRQ2 |
| Arachis hypo{ | A0A444WY33 |
| Arachis hypo{ | A0A445E825 |
| Arachis hypo{ | A0A445BS20 |
| Arachis hypo{ | A0A445C104 |
| Arachis hypo{ | A0A445BYZ1 |
| Arachis hypo{ | A0A445BB33 |
| Arachis hypo{ | A0A445BB67 |
| Arachis hypo{ | A0A444WUD9 |
| Arachis hypo{ | A0A445BP39 |
| Arachis hypo{ | A0A445DG52 |
| Arachis hypo{ | A0A445A429 |
| Arachis hypo{ | A0A445BB24 |
| Arachis hypo{ | A0A444Y4T8 |
| Arachis hypo{ | A0A445BYM5 |
| Arachis hypo{ | A0A444Y4Y5 |
| Arachis hypo{ | A0A445BYV9 |
| Arachis hypo{ | A0A444WY17 |
| Arachis hypo{ | A0A445BY96 |
| Arachis hypo{ | A0A445DRK7 |
| Arachis hypo{ | A0A444Y529 |
| Arachis hypo{ | A0A445BB45 |
| Arachis hypo{ | A0A444Y6M2 |
| Arachis hypo{ | A0A444Y6J2 |
| Arachis hypo{ | A0A444Y6K9 |
| Arachis hypo{ | A0A445BP47 |
| Arachis hypo{ | A0A444WY50 |
| Arachis hypo{ | A0A445AZM8 |
| Arachis hypo{ | A0A445BB36 |
| Arachis hypo{ | A0A445C8A4 |
| Arachis hypo{ | A0A444WY27 |
| Arachis hypo{ | A0A445A408 |
| Arachis hypo{ | A0A444XXR6 |
| Arachis hypo{ | A0A444Y4W8 |
| Arachis hypo{ | A0A444WY44 |
| Arachis hypo{ | A0A444Y6V9 |

| | |
|---|---|
| Arachis hypo | A0A444Y594 |
| Arachis hypo | A0A445BYW3 |
| Arachis hypo | A0A445BB13 |
| Arachis hypo | A0A445DRJ9 |
| Arachis hypo | A0A444WXY2 |
| Arachis hypo | A0A445A416 |
| Arachis hypo | A0A445BZ11 |
| Arachis hypo | A0A445A4D2 |
| Arachis hypo | A0A290GKF7 |
| Arachis hypo | A0A445BKE9 |
| Arachis hypo | A0A444Y540 |
| Arachis hypo | A0A445BB44 |
| Arachis hypo | A0A445A4A2 |
| Arachis hypo | A0A445AIK2 |
| Arachis hypo | A0A445BY95 |
| Arachis hypo | A0A445CSW5 |
| Arachis hypo | A0A445A418 |
| Arachis hypo | A0A445BB46 |
| Arachis hypo | A0A444XXR8 |
| Arachis hypo | A0A444Y4Q5 |
| Arachis hypo | A0A445BB60 |
| Arachis hypo | A0A444Z8T4 |
| Arachis hypo | A0A445DJU8 |
| Arachis hypo | A0A445BB42 |
| Arachis hypo | A0A444Y6I9 |
| Arachis hypo | A0A444WY31 |
| Arachis hypo | A0A445BZ05 |
| Arachis hypo | A0A445CIM3 |
| Arachis hypo | A0A444Y4Y0 |
| Arachis hypo | A0A445BYI2 |
| Arachis hypo | A0A444Y542 |
| Arachis hypo | A0A444Y535 |
| Arachis hypo | A0A444WXZ9 |
| Arachis hypo | A0A445E7X5 |
| Arachis hypo | A0A445E0F1 |
| Arachis hypo | A0A445BB48 |
| Arachis hypo | A0A444Y4W6 |
| Arachis hypo | A0A444Y4Y8 |
| Arachis hypo | A0A445BB50 |
| Arachis hypo | A0A445AII6 |
| Arachis hypo | A0A445BY65 |

| | |
|---|---|
| Arachis hypo{ | A0A445E0N0 |
| Arachis hypo{ | A0A445BY56 |
| Arachis hypo{ | A0A445CBC1 |
| Arachis hypo{ | A0A444Y553 |
| Arachis hypo{ | A0A445A499 |
| Arachis hypo{ | A0A445DYX4 |
| Arachis hypo{ | A0A444XXS1 |
| Arachis hypo{ | A0A445E8D0 |
| Arachis hypo{ | A0A444WY11 |
| Arachis hypo{ | A0A445BX01 |
| Araucaria cur | A0A0D6R0F5 |
| Araucaria cur | A0A0D6QTT4 |
| Araucaria cur | A0A0D6QT12 |
| Araucaria cur | A0A0D6QTN1 |
| Araucaria cur | A0A0D6R4V4 |
| Araucaria cur | A0A0D6QT17 |
| Araucaria cur | A0A0D6QRX4 |
| Araucaria cur | A0A0D6QRZ3 |
| Araucaria cur | A0A0D6QTH7 |
| Araucaria cur | A0A0D6QR37 |
| Araucaria cur | A0A0D6QRW0 |
| Araucaria cur | A0A0D6QSZ0 |
| Araucaria cur | A0A0D6QR91 |
| Araucaria cur | A0A0D6QR45 |
| Archangium s | A0A1Q3HVR3 |
| Arenimonas c | A0A2P6M7B1 |
| Arenimonas c | A0A091BFC1 |
| Arion vulgaris | A0A0B6XVM2 |
| Armadillidiun | A0A444SIQ6 |
| Armadillidiun | A0A444TRU3 |
| Armadillidiun | A0A444SZK2 |
| Armadillidiun | A0A444SL52 |
| Armadillidiun | A0A444SR30 |
| Armadillidiun | A0A444SMM6 |
| Artemisia anr | A0A2U1MNQ4 |
| Artemisia anr | A0A2U1MVH2 |
| Artemisia anr | A0A2U1QG88 |
| Artemisia anr | A0A2U1KI18 |
| Artemisia anr | A0A2U1QD59 |
| Artemisia anr | A0A2U1KPH1 |
| Artemisia anr | A0A2U1KAR9 |

| | |
|---|---|
| Artemisia anr | A0A2U1Q3R8 |
| Artemisia anr | A0A2U1PGN2 |
| Artemisia anr | A0A2U1MXM3 |
| Artemisia anr | A0A2U1Q692 |
| Artemisia anr | A0A2U1MYT5 |
| Artemisia anr | A0A2U1QB37 |
| Artemisia anr | A0A2U1PC96 |
| Artemisia anr | A0A2U1MCE3 |
| Artemisia anr | A0A2U1P9R0 |
| Artemisia anr | A0A2U1LBZ4 |
| Artemisia anr | A0A2U1PTC0 |
| Artemisia anr | A0A2U1NVX8 |
| Artemisia anr | A0A2U1MRL9 |
| Artemisia anr | A0A2U1MF47 |
| Artemisia anr | A0A2U1MSA6 |
| Artemisia anr | A0A2U1MA90 |
| Artemisia anr | A0A2U1L2N0 |
| Artemisia anr | A0A2U1K9W3 |
| Artemisia anr | A0A2U1Q5W4 |
| Artemisia anr | A0A2U1NCZ8 |
| Artemisia anr | A0A2U1PHB5 |
| Artemisia anr | A0A2U1P0Y9 |
| Artemisia anr | A0A2U1MK05 |
| Artemisia anr | A0A2U1LB20 |
| Artemisia anr | A0A2U1LCW0 |
| Artemisia anr | A0A2U1P3C5 |
| Artemisia anr | A0A2U1M6L0 |
| Artemisia anr | A0A2U1QB05 |
| Artemisia anr | A0A2U1MFU2 |
| Artemisia anr | A0A2U1MRH5 |
| Artemisia anr | A0A2U1L7Y8 |
| Artemisia anr | A0A2U1Q3Y3 |
| Artemisia anr | A0A2U1KHM7 |
| Artemisia anr | A0A2U1L9K1 |
| Artemisia anr | A0A2U1PVA8 |
| Artemisia anr | A0A2U1KTL0 |
| Artemisia anr | A0A2U1PT94 |
| Artemisia anr | A0A2U1MKG2 |
| Artemisia anr | A0A2U1MIH0 |
| Artemisia anr | A0A2U1QHC0 |
| Artemisia anr | A0A2U1QKQ6 |

| | |
|---|---|
| Artemisia anr | A0A2U1PGR0 |
| Artemisia anr | A0A2U1MK14 |
| Artemisia anr | A0A2U1KKI3 |
| Artemisia anr | A0A2U1L628 |
| Artemisia anr | A0A2U1LK69 |
| Artemisia anr | A0A2U1LX06 |
| Artemisia anr | A0A2U1M3I4 |
| Artemisia anr | A0A2U1LEW8 |
| Artemisia anr | A0A2U1KKC2 |
| Artemisia anr | A0A2U1MXT3 |
| Artemisia anr | A0A2U1PTA5 |
| Artemisia anr | A0A2U1KHY4 |
| Artemisia anr | A0A2U1MUK8 |
| Artemisia anr | A0A2U1L5S7 |
| Artemisia anr | A0A2U1N8E3 |
| Artemisia anr | A0A2U1KIY8 |
| Artemisia anr | A0A2U1NEM6 |
| Artemisia anr | A0A2U1P0M6 |
| Artemisia anr | A0A2U1PDI3 |
| Artemisia anr | A0A2U1MSU5 |
| Artemisia anr | A0A2U1LJ61 |
| Artemisia anr | A0A2U1PQZ4 |
| Artemisia anr | A0A2U1PD36 |
| Artemisia anr | A0A2U1LEY7 |
| Artemisia anr | A0A2U1MJY2 |
| Artemisia anr | A0A2U1KDS1 |
| Artemisia anr | A0A2U1KXK7 |
| Artemisia anr | A0A2U1KWP0 |
| Artemisia anr | A0A2U1LG99 |
| Artemisia anr | A0A2U1NQJ3 |
| Artemisia anr | A0A2U1P450 |
| Artemisia anr | A0A2U1N3I9 |
| Artemisia anr | A0A2U1LNH3 |
| Artemisia anr | A0A2U1PZ37 |
| Artemisia anr | A0A2U1MK01 |
| Artemisia anr | A0A2U1P169 |
| Artemisia anr | A0A2U1L3H4 |
| Artemisia anr | A0A2U1NUG1 |
| Artemisia anr | A0A2U1LPS2 |
| Artemisia anr | A0A2U1PT69 |
| Artemisia anr | A0A2U1LZ92 |

| | |
|---|---|
| Artemisia anr | A0A2U1LEV4 |
| Artemisia anr | A0A2U1KQB6 |
| Artemisia anr | A0A2U1MFI2 |
| Artemisia anr | A0A2U1MWD6 |
| Artemisia anr | A0A2U1QDH1 |
| Artemisia anr | A0A2U1PD98 |
| Artemisia anr | A0A2U1KFT4 |
| Artemisia anr | A0A2U1PG62 |
| Artemisia anr | A0A2U1M3F8 |
| Artemisia anr | A0A2U1KBU3 |
| Artemisia anr | A0A2U1MMK0 |
| Artemisia anr | A0A2U1KFP7 |
| Artemisia anr | A0A2U1NQJ1 |
| Artemisia anr | A0A2U1P8A2 |
| Artemisia anr | A0A2U1N8W6 |
| Artemisia anr | A0A2U1MCQ6 |
| Artemisia anr | A0A2U1QB20 |
| Artemisia anr | A0A2U1MSB3 |
| Artemisia anr | A0A2U1MKZ0 |
| Artemisia anr | A0A2U1MY98 |
| Artemisia anr | A0A2U1PDB5 |
| Artemisia anr | A0A2U1MGQ2 |
| Artemisia anr | A0A2U1KLB8 |
| Artemisia anr | A0A2U1LNA4 |
| Artemisia anr | A0A2U1PIG5 |
| Artemisia anr | A0A2U1PG29 |
| Arthrobotrys | G1XPN8 |
| Arthrobotrys | G1XSL9 |
| Arthrobotrys | G1XGI6 |
| Arthrobotrys | G1XG00 |
| Arthrobotrys | G1XSA2 |
| Arthroderma | D4ARU1 |
| Arthroderma | E4UUF0 |
| Arundo dona | A0A0A9HRX8 |
| Arundo dona | A0A0A9TEG4 |
| Arundo dona | A0A0A9DPX1 |
| Arundo dona | A0A0A8YWY2 |
| Arundo dona | A0A0A9F543 |
| Arundo dona | A0A0A9E2Y2 |
| Arundo dona | A0A0A9E7P7 |
| Arundo dona | A0A0A9QLR5 |

| | |
|---|---|
| Arundo dona | A0A0A9DFU5 |
| Arundo dona | A0A0A8YEX4 |
| Arundo dona | A0A0A9NPY6 |
| Arundo dona | A0A0A9GL28 |
| Arundo dona | A0A0A9DM30 |
| Arundo dona | A0A0A9S6Y8 |
| Arundo dona | A0A0A9DF09 |
| Arundo dona | A0A0A9BMR2 |
| Arundo dona | A0A0A9LN34 |
| Arundo dona | A0A0A9TYC3 |
| Arundo dona | A0A0A9QZ24 |
| Arundo dona | A0A0A9DW73 |
| Arundo dona | A0A0A9H3W6 |
| Ascaris lumbr | A0A0M3IPJ6 |
| Ascaris lumbr | A0A0M3HP86 |
| Ascaris lumbr | A0A0M3IT96 |
| Ascaris lumbr | A0A0M3IBY8 |
| Ascaris lumbr | A0A0M3HND4 |
| Ascaris lumbr | A0A0M3HU83 |
| Ascaris lumbr | A0A0M3HEY4 |
| Ascaris lumbr | A0A0M3IH35 |
| Ascaris lumbr | A0A0M3ILR8 |
| Ascaris lumbr | A0A0M3IVC1 |
| Ascaris lumbr | A0A0M3IRC3 |
| Ascaris lumbr | A0A0M3IAG3 |
| Ascaris lumbr | A0A0M3IPG4 |
| Ascaris lumbr | A0A0M3HG49 |
| Ascaris lumbr | A0A0M3IRJ0 |
| Ascaris lumbr | A0A0M3HQ47 |
| Ascaris lumbr | A0A0M3I750 |
| Ascaris lumbr | A0A0M3ITR0 |
| Ascaris suum | F1L0P3 |
| Ascaris suum | F1L0G7 |
| Ascaris suum | F1LC48 |
| Ascaris suum | F1KWX7 |
| Ascaris suum | F1KVZ1 |
| Ascaris suum | F1KRY6 |
| Ascaris suum | F1L7L8 |
| Ascidiaceihak | A0A2R8B9F5 |
| Ascobolus im | A0A3N4IAA5 |
| Ascobolus im | A0A3N4IAW4 |

| | |
|---|---|
| Ascosphaera | A0A167YNV3 |
| Asparagus of | A0A1R3L6N1 |
| Asparagus of | A0A1R3L7T1 |
| Aspergillus ar | A0A2G7FEN3 |
| Aspergillus ar | A0A2G7G4M6 |
| Aspergillus bo | A0A1F7ZRV8 |
| Aspergillus bo | A0A1F8A4B8 |
| Aspergillus bo | A0A1F7ZUM9 |
| Aspergillus ca | A0A0U5FUR8 |
| Aspergillus ca | A0A0U5GHF6 |
| Aspergillus ca | A0A0U5FUB2 |
| Aspergillus ca | A0A0U5CCI5 |
| Aspergillus ca | A0A2I1DHJ1 |
| Aspergillus ca | A0A2I1D6S1 |
| Aspergillus ca | A0A2I2FIZ7 |
| Aspergillus ca | A0A2I2EZ59 |
| Aspergillus cl | A1CCE8 |
| Aspergillus cl | A1CBV4 |
| Aspergillus cr | A0A1E3BSI6 |
| Aspergillus el | A0A319DW04 |
| Aspergillus eu | A0A317WGQ7 |
| Aspergillus fla | B8NFC9 |
| Aspergillus fla | B8NFD0 |
| Aspergillus fla | B8NDN6 |
| Aspergillus fla | A0A2P2HFU4 |
| Aspergillus fla | A0A2P2HLC7 |
| Aspergillus fla | A0A2P2H4K6 |
| Aspergillus fla | A0A2P2HM09 |
| Aspergillus fla | A0A2P2HFT0 |
| Aspergillus fla | A0A2P2H9K4 |
| Aspergillus fla | A0A2P2HPX0 |
| Aspergillus fla | A0A3M7JRD9 |
| Aspergillus fla | A0A3M7JTE8 |
| Aspergillus fla | A0A3M7K264 |
| Aspergillus fla | A0A364MKB4 |
| Aspergillus fla | A0A364M579 |
| Aspergillus fu | A0A0J5Q2X8 |
| Aspergillus fu | A0A0J5PMA7 |
| Aspergillus gl | A0A1L9VP26 |
| Aspergillus gl | A0A1L9VC48 |
| Aspergillus gl | A0A1L9VP33 |

| | |
|---|---|
| Aspergillus gl | A0A1L9VNY3 |
| Aspergillus gl | A0A1L9VRE9 |
| Aspergillus he | A0A317WAR2 |
| Aspergillus ho | A0A395HFU3 |
| Aspergillus ib | A0A395GPB8 |
| Aspergillus ka | G7XTE0 |
| Aspergillus la | A0A319AAV7 |
| Aspergillus la | A0A319A7A0 |
| Aspergillus le | A0A0S7DE04 |
| Aspergillus le | A0A0S7E252 |
| Aspergillus le | A0A0S7EBW1 |
| Aspergillus lu | A0A1M3TFU6 |
| Aspergillus lu | A0A1M3T8Q8 |
| Aspergillus lu | A0A1M3TTZ5 |
| Aspergillus lu | A0A146F4B8 |
| Aspergillus m | A0A3D8T616 |
| Aspergillus m | A0A3D8S6I6 |
| Aspergillus m | A0A3D8T3A9 |
| Aspergillus m | A0A3D8QMJ3 |
| Aspergillus ne | A0A318YC13 |
| Aspergillus ne | A0A318Y1N8 |
| Aspergillus ne | A0A318YAK2 |
| Aspergillus ni | G3XMA4 |
| Aspergillus ni | A2QYV6 |
| Aspergillus ni | A2QTC1 |
| Aspergillus ni | A0A370BU00 |
| Aspergillus ni | A0A370BW53 |
| Aspergillus ni | A0A3F3R4A3 |
| Aspergillus ni | A0A117E3X4 |
| Aspergillus no | A0A0L1J937 |
| Aspergillus no | A0A2I1C107 |
| Aspergillus no | A0A2I1BTK7 |
| Aspergillus no | A0A2I1CJW4 |
| Aspergillus no | A0A2I1CCR0 |
| Aspergillus no | A0A2I1BTA2 |
| Aspergillus oc | A0A2T5M095 |
| Aspergillus oc | A0A2T5LRQ5 |
| Aspergillus oc | A0A0F8UID4 |
| Aspergillus or | I8U2J0 |
| Aspergillus or | I8A614 |
| Aspergillus or | I8U453 |

| | |
|---|---|
| Aspergillus or | Q2U7K1 |
| Aspergillus or | Q2UP96 |
| Aspergillus or | Q2U8A9 |
| Aspergillus or | Q2U2F0 |
| Aspergillus or | A0A1S9DBB6 |
| Aspergillus or | A0A1S9DIX8 |
| Aspergillus or | A0A1S9DJT8 |
| Aspergillus pa | A0A0F0IBG5 |
| Aspergillus pa | A0A0F0ICT4 |
| Aspergillus pl | A0A370PJK5 |
| Aspergillus pl | A0A370PHC2 |
| Aspergillus ra | A0A0F8X4Q4 |
| Aspergillus ru | A0A017SC51 |
| Aspergillus ru | A0A017SDA8 |
| Aspergillus sa | A0A318ZXC8 |
| Aspergillus sa | A0A318Z9C0 |
| Aspergillus sa | A0A318Z979 |
| Aspergillus sc | A0A3A2ZMA4 |
| Aspergillus sc | A0A319EXR3 |
| Aspergillus sc | A0A319EAA1 |
| Aspergillus sc | A0A319ECG3 |
| Aspergillus sc | A0A317VSX8 |
| Aspergillus st | A0A2I2GSR7 |
| Aspergillus ta | A0A2J5HZ31 |
| Aspergillus ta | A0A2J5HG69 |
| Aspergillus ta | A0A2J5IA66 |
| Aspergillus te | Q0CRW2 |
| Aspergillus th | A0A397GTA6 |
| Aspergillus tu | A0A1L9NER7 |
| Aspergillus tu | A0A1L9NHC1 |
| Aspergillus tu | A0A397I0S7 |
| Aspergillus tu | A0A3R7FTI7 |
| Aspergillus tu | A0A421CZV9 |
| Aspergillus tu | A0A397FZA9 |
| Aspergillus tu | A0A397H5H6 |
| Aspergillus tu | A0A397I608 |
| Aspergillus u | A0A0K8LIL4 |
| Aspergillus u | A0A319CFJ0 |
| Aspergillus u | A0A319C2C5 |
| Aspergillus u | A0A319CGR1 |
| Aspergillus va | A0A319B527 |

| | |
|---|---|
| Aspergillus vi | A0A2V5HBT7 |
| Aspergillus w | A0A1L9R4F3 |
| Aspergillus w | A0A1L9RJD3 |
| Aspergillus w | A0A1L9RI70 |
| Astatotilapia | A0A3P8R5U0 |
| Astatotilapia | A0A3P8QWI7 |
| Astatotilapia | A0A3P8NAK1 |
| Astatotilapia | A0A3P8NAL8 |
| Astatotilapia | A0A3P8NKJ4 |
| Astatotilapia | A0A3P8PUS6 |
| Astatotilapia | A0A3P8NAX7 |
| Asterias rube | A0A3R5SPP4 |
| Astyanax me: | W5LAJ7 |
| Astyanax me: | A0A3B1IZ91 |
| Astyanax me: | W5JZP1 |
| Astyanax me: | W5LF06 |
| Astyanax me: | A0A3B1JB59 |
| Astyanax me: | W5LF07 |
| Astyanax me: | W5KKK6 |
| Atta cephalot | A0A158NQ95 |
| Atta cephalot | A0A158NRJ0 |
| Atta cephalot | A0A158NUD3 |
| Atta cephalot | A0A158NCF5 |
| Atta cephalot | A0A158NFW1 |
| Atta cephalot | A0A158P387 |
| Atta colombi | A0A195BUK4 |
| Atta colombi | A0A195B440 |
| Atta colombi | A0A151I1P9 |
| Atta colombi | A0A195BWB4 |
| Atta colombi | A0A195BVH1 |
| Atta colombi | A0A195B228 |
| Atta colombi | A0A195BU26 |
| Atta colombi | A0A195BWF1 |
| Aurantimona | A0A1X0T3J5 |
| Aureimonas a | A0A1M6C5Z3 |
| Aureimonas a | A0A0B1Q6J5 |
| Aureimonas a | A0A0B1Q3S9 |
| Aureimonas j | A0A1H0C5Q6 |
| Aureimonas p | A0A1I2INE0 |
| Aureimonas p | A0A1I2J8H4 |
| Aureimonas s | A0A0S2EVY8 |

| | |
|---|---|
| Aureimonas s | A0A0Q6CLW5 |
| Aureimonas s | A0A0Q6D7V7 |
| Aureimonas s | A0A0Q6CNY5 |
| Aureimonas s | A0A0Q6DNE7 |
| Aureimonas s | A0A0Q6E7R1 |
| Aureimonas u | A0A175RVT2 |
| Aureimonas u | A0A175RB81 |
| Aureimonas u | A0A175RYG2 |
| Aureobasidiu | A0A074VR01 |
| Aureobasidiu | A0A074W6P0 |
| Aureobasidiu | A0A074VFW8 |
| Aureobasidiu | A0A074VY00 |
| Aureobasidiu | A0A074VWR6 |
| Aureobasidiu | A0A074VMM6 |
| Aureobasidiu | A0A074VQQ6 |
| Aureobasidiu | A0A074W863 |
| Aureobasidiu | A0A074VGQ4 |
| Aureobasidiu | A0A074WSI2 |
| Aureobasidiu | A0A074WT58 |
| Aureobasidiu | A0A074W896 |
| Aureobasidiu | A0A074WDM8 |
| Aureobasidiu | A0A074X5F4 |
| Aureobasidiu | A0A074WD87 |
| Aureobasidiu | A0A074WP99 |
| Aureobasidiu | A0A074WYP5 |
| Aureobasidiu | A0A074X7D6 |
| Aureobasidiu | A0A074X9M1 |
| Aureobasidiu | A0A074XKI4 |
| Aureobasidiu | A0A074XMC5 |
| Aureobasidiu | A0A074XW73 |
| Aureobasidiu | A0A074XUY6 |
| Aureobasidiu | A0A074XV26 |
| Aureobasidiu | A0A1A7MGL9 |
| Aureobasidiu | A0A1A7MIQ4 |
| Aureobasidiu | A0A1A7MHG3 |
| Aureobasidiu | A0A1A7MCT8 |
| Aureobasidiu | A0A1A7MLB2 |
| Aureobasidiu | A0A1A7MP46 |
| Aureobasidiu | A0A1A7MAV7 |
| Aureobasidiu | A0A1A7MR78 |
| Aureobasidiu | A0A1A7MEM4 |

| | |
|---|---|
| Aureobasidiu | A0A1A7MN21 |
| Aureobasidiu | A0A1A7MAJ0 |
| Aureobasidiu | A0A1A7MFI3 |
| Aureobasidiu | A0A1A7MS81 |
| Aureobasidiu | A0A1A7MDG6 |
| Aureobasidiu | A0A074Z9D4 |
| Aureobasidiu | A0A074XZQ2 |
| Aureobasidiu | A0A074Y4C9 |
| Aureobasidiu | A0A074YXY8 |
| Aureobasidiu | A0A074Y4Q8 |
| Aureobasidiu | A0A074YRD0 |
| Aureobasidiu | A0A074YLW2 |
| Aureobasidiu | A0A074YDL6 |
| Aureobasidiu | A0A074YCF2 |
| Aureobasidiu | A0A074YUS8 |
| Aureobasidiu | A0A074Y8S5 |
| Aureobasidiu | A0A074YQX7 |
| Aureobasidiu | A0A074YG47 |
| Aureobasidiu | A0A074XZ34 |
| Aureobasidiu | A0A074Z895 |
| Aureococcus | F0YIS4 |
| Austrofundul | A0A2I4AVX2 |
| Austrofundul | A0A2I4CU49 |
| Austrofundul | A0A2I4CCC7 |
| Austrofundul | A0A2I4D285 |
| Austrofundul | A0A2I4D8H5 |
| Austrofundul | A0A2I4CKP4 |
| Austrofundul | A0A2I4AVX1 |
| Austrofundul | A0A2I4BJ83 |
| Austrofundul | A0A2I4AVX5 |
| Austrofundul | A0A2I4CU47 |
| Auxenochlore | A0A1D2AC95 |
| Auxenochlore | A0A1D2ADW2 |
| Auxenochlore | A0A1D2A527 |
| Auxenochlore | A0A087SNA2 |
| Auxenochlore | A0A1D2AA88 |
| Auxenochlore | A0A3M7KY55 |
| Auxenochlore | A0A3M7L506 |
| Auxenochlore | A0A087S9Y1 |
| Auxenochlore | A0A3M7L688 |
| Auxenochlore | A0A1D2A2T0 |

| | |
|---|---|
| Auxenochlor€ | A0A1D1ZN49 |
| Auxenochlor€ | A0A1D2AGQ5 |
| Auxenochlor€ | A0A087SA78 |
| bacterium F1 | A0A202DKR6 |
| Bactrocera d( | A0A034V924 |
| Bactrocera d( | A0A034V101 |
| Bactrocera d( | A0A034VV04 |
| Bactrocera d( | A0A034VBU4 |
| Bactrocera d( | A0A034VP05 |
| Bactrocera d( | A0A034V269 |
| Bactrocera d( | A0A034VU87 |
| Bactrocera d( | A0A034VB68 |
| Bactrocera d( | A0A034WDU0 |
| Bactrocera d( | A0A034VQT0 |
| Bactrocera d( | A0A034VAC8 |
| Bactrocera d( | A0A034VA94 |
| Bactrocera la | A0A0K8WFY5 |
| Bactrocera la | A0A0K8V187 |
| Bactrocera la | A0A0K8TYA5 |
| Bactrocera la | A0A0K8U2Z1 |
| Bactrocera la | A0A0K8W143 |
| Bactrocera la | A0A0K8W107 |
| Bactrocera la | A0A0K8WAG5 |
| Bactrocera la | A0A0K8UYR3 |
| Bactrocera la | A0A0K8UC22 |
| Bactrocera la | A0A0K8W0F9 |
| Bactrocera la | A0A0K8URT7 |
| Bactrocera la | A0A0K8W7I2 |
| Bactrocera la | A0A0K8U7H8 |
| Bactrocera la | A0A0K8VH83 |
| Bactrocera la | A0A0K8VDD1 |
| Balaenoptera | A0A383ZG27 |
| Balaenoptera | A0A383ZV45 |
| Balaenoptera | A0A383Z1M8 |
| Balaenoptera | A0A383ZVD0 |
| Balaenoptera | A0A384AFX7 |
| Balaenoptera | A0A384AFQ0 |
| Balearica reg | A0A087VQ86 |
| Balearica reg | A0A087VHF0 |
| Balneola sp..› | A0A2E9UHU0 |
| Bambusicola | A0A2P4SWX5 |

| | |
|---|---|
| Bambusicola | A0A2P4SJS3 |
| Bandra mega | A0A2K9V7N6 |
| Bandra mega | A0A2K9V7Z4 |
| Bandra mega | A0A2K9V7Z1 |
| Bartonella ap | A0A1R0FC62 |
| Bartonella ap | A0A1U9MF28 |
| Basidiobolus | A0A1Y1WQS1 |
| Bathycoccus | K8EB66 |
| Bathycoccus | K8FDI0 |
| Bathycoccus | K8F4B2 |
| Bathycoccus | K8EQC3 |
| Bathycoccus | K8EJ82 |
| Bathycoccus | K8EAB0 |
| Bathycoccus | K8F6I2 |
| Bathycoccus : | E5ESL6 |
| Batrachochyt | F4NSE3 |
| Batrachochyt | F4NSY5 |
| Batrachochyt | A0A177WA49 |
| Batrachochyt | A0A177WAP2 |
| Batrachochyt | A0A1S8VW60 |
| Batrachochyt | A0A1S8VHV0 |
| Baudoinia pa | M2LEM6 |
| Baudoinia pa | M2MRY7 |
| Baudoinia pa | M2LJL5 |
| Baudoinia pa | M2N072 |
| Bauldia litora | A0A1G6CN88 |
| Bauldia litora | A0A1G6CP43 |
| Bdellovibrion | A0A2E7GCJ6 |
| Bdellovibrion | A0A2E7G7G0 |
| Bdellovibrion | A0A2E8HW08 |
| Beauveria ba: | J5JQG2 |
| Beauveria ba: | J4WGF4 |
| Beauveria ba: | J4KLZ6 |
| Beauveria ba: | A0A0A2VAI1 |
| Beauveria ba: | A0A0A2VQE8 |
| Beauveria ba: | A0A2N6NCG5 |
| Beauveria ba: | A0A2N6P0V4 |
| Beauveria ba: | A0A2N6NH65 |
| Beauveria ba: | A0A2S7Y029 |
| Beauveria ba: | A0A2S7XWT7 |
| Beauveria bro | A0A167DY06 |

| | |
|---|---|
| Beauveria bro | A0A162JT95 |
| Beijerinckia ir | B2IFP4 |
| Besnoitia bes | A0A2A9MK42 |
| Besnoitia bes | A0A2A9MEK6 |
| Besnoitia bes | A0A2A9MHF7 |
| Besnoitia bes | A0A2A9M2B2 |
| Besnoitia bes | A0A2A9MP64 |
| Besnoitia bes | A0A2A9M947 |
| Besnoitia bes | A0A2A9MH57 |
| Besnoitia bes | A0A2A9MHN4 |
| Besnoitia bes | A0A2A9M1K1 |
| Besnoitia bes | A0A2A9MK31 |
| Besnoitia bes | A0A2A9MLG1 |
| Besnoitia bes | A0A2A9M8Y8 |
| Besnoitia bes | A0A2A9MGA0 |
| Besnoitia bes | A0A2A9M957 |
| Besnoitia bes | A0A2A9M7H0 |
| Besnoitia bes | A0A2A9LZF4 |
| Besnoitia bes | A0A2A9MGC4 |
| Besnoitia bes | A0A2A9MDT6 |
| Besnoitia bes | A0A2A9MEJ6 |
| Besnoitia bes | A0A2A9MEY2 |
| Besnoitia bes | A0A2A9MBU9 |
| Besnoitia bes | A0A2A9MDP9 |
| Beta vulgaris | A0A0J8B6J9 |
| Beta vulgaris | A0A0J8B9T3 |
| Beta vulgaris | A0A0J8DZK2 |
| Beta vulgaris | A0A0J8B8G6 |
| Beta vulgaris | A0A0J8B6H3 |
| Beta vulgaris | A0A0J8B5L4 |
| Beta vulgaris | A0A0J8B9Q2 |
| Beta vulgaris | A0A0J8E0I7 |
| Biomphalaria | A0A2C9M6E5 |
| Biomphalaria | A0A2C9M2K3 |
| Biomphalaria | A0A2C9L085 |
| Biomphalaria | A0A2C9LUG0 |
| Biomphalaria | A0A2C9KVY0 |
| Biomphalaria | A0A2C9JZZ3 |
| Biomphalaria | A0A2C9LGM4 |
| Biomphalaria | A0A2C9MAU4 |
| Biomphalaria | A0A2C9KU12 |

Biomphalaria A0A2C9L1D2
Biomphalaria A0A2C9M004
Biomphalaria A0A2C9LZ70
Biomphalaria A0A2C9L2E0
Biomphalaria A0A2C9LWZ3
Biomphalaria A0A2C9LD29
Biomphalaria A0A2C9KJS2
Biomphalaria A0A2C9LP23
Biomphalaria A0A2C9KNW3
Biomphalaria A0A2C9LTI5
Biomphalaria A0A2C9M9Z8
Biomphalaria A0A2C9KM74
Biomphalaria A0A2C9KKC9
Biomphalaria A0A2C9JPS7
Biomphalaria A0A2C9M6Q8
Biomphalaria A0A2C9KBM1
Biomphalaria A0A2C9JPW5
Biomphalaria A0A2C9LE46
Biomphalaria A0A2C9LTU9
Biomphalaria A0A2C9JNC8
Biomphalaria A0A2C9LNH2
Biomphalaria A0A2C9LXF9
Biomphalaria A0A182YU34
Biomphalaria A0A2C9JRC5
Biomphalaria A0A2C9JBX8
Biomphalaria A0A2C9LL38
Biomphalaria A0A2C9KHG2
Biomphalaria A0A2C9M7I1
Biomphalaria A0A2C9KDN6
Biomphalaria A0A2C9LWA8
Biomphalaria A0A2C9K7K3
Biomphalaria A0A2C9KKA8
Biomphalaria A0A2C9L3Z3
Biomphalaria A0A2C9LBA6
Biomphalaria A0A2C9KJ30
Biomphalaria A0A2C9M3Q1
Biomphalaria A0A2C9M449
Biomphalaria A0A2C9KT20
Biomphalaria A0A2C9KXA4
Bipolaris oryz W6Z007
Bipolaris oryz W6ZCF9

| | |
|---|---|
| Bipolaris oryz | W6Z4B8 |
| Bipolaris oryz | W6ZJU0 |
| Bipolaris oryz | W6ZED0 |
| Bipolaris oryz | W6ZMD3 |
| Bipolaris oryz | W6ZA29 |
| Bipolaris oryz | W6Z9A5 |
| Bipolaris oryz | W6Z1B7 |
| Bipolaris victo | W7DVV8 |
| Bipolaris victo | W7EBL2 |
| Bipolaris victo | W7ER18 |
| Bipolaris victo | W7EF46 |
| Bipolaris victo | W7EPL2 |
| Bipolaris victo | W7ESE4 |
| Bipolaris victo | W7EV06 |
| Bipolaris victo | W7DVB6 |
| Bipolaris victo | W7E897 |
| Bipolaris zeic | W6YTV6 |
| Bipolaris zeic | W6Y989 |
| Bipolaris zeic | W6YAZ9 |
| Bipolaris zeic | W6XJ34 |
| Bipolaris zeic | W6Y7X4 |
| Bipolaris zeic | W6XU68 |
| Bipolaris zeic | W6Y6Y8 |
| Bipolaris zeic | W6Y442 |
| Biscutella neu | W6EIN7 |
| Biscutella neu | W6EMA3 |
| Biscutella neu | W6EMA1 |
| Biscutella neu | W6EIL6 |
| Biscutella neu | W6EK17 |
| Biscutella neu | W6EMB1 |
| Biscutella neu | W6EVT2 |
| Biscutella neu | W6EVS5 |
| Biscutella neu | W6EVU1 |
| Biscutella neu | W6EIM1 |
| Biscutella neu | W6EIS1 |
| Biscutella neu | W6EMA7 |
| Biscutella neu | W6EIT3 |
| Biscutella neu | W6EVS9 |
| Biscutella neu | W6EK05 |
| Biscutella neu | W6EMA5 |
| Biscutella neu | W6EK24 |

| | |
|---|---|
| Biscutella neu | W6EIM6 |
| Biscutella neu | W6EK09 |
| Biscutella neu | W6EK13 |
| Biscutella neu | W6EIP2 |
| Biscutella neu | W6EIS9 |
| Biscutella neu | W6EVT6 |
| Biscutella neu | W6EIS6 |
| Biscutella neu | W6EIU2 |
| Biscutella neu | W6EMA9 |
| Blastocatellia | A0A352FPY6 |
| Blastomonas | A0A2D6H9Y1 |
| Blastomyces | A0A1J9RE20 |
| Blattella gern | A0A2P8XY43 |
| Blattella gern | A0A2P8YQ78 |
| Blattella gern | A0A2P8YMN5 |
| Blattella gern | A0A2P8YR42 |
| Blattella gern | A0A2P8YJK3 |
| Blattella gern | A0A2P8YZ76 |
| Blattella gern | A0A2P8YR97 |
| Blattella gern | A0A2P8Z5Y9 |
| Bodo saltans | A0A2H4UUU3 |
| Bodo saltans | A0A2H4UU38 |
| Bodo saltans. | A0A0S4J9J2 |
| Bodo saltans. | A0A0S4J936 |
| Bodo saltans. | A0A0S4JQX2 |
| Bombyx mori | H9IXB3 |
| Bombyx mori | H9JA26 |
| Bombyx mori | H9JJP0 |
| Bombyx mori | H9JJP1 |
| Bombyx mori | H9JE81 |
| Bombyx mori | H9JE80 |
| Bos mutus.xls | L8IUV1 |
| Bos mutus.xls | L8IA26 |
| Bos mutus.xls | L8IEV1 |
| Bos mutus.xls | L8IY98 |
| Bos mutus.xls | L8IC35 |
| Bos taurus.xls | P06868 |
| Bos taurus.xls | F1MZD6 |
| Bos taurus.xls | A0A1L3G6L5 |
| Bos taurus.xls | Q6PZ62 |
| Bos taurus.xls | Q6Q0I7 |

| | |
|---|---|
| Bos taurus.xls | Q2KJB4 |
| Bos taurus.xls | E1B726 |
| Bos taurus.xls | E1BDW7 |
| Bos taurus.xls | A0A3Q1LSS6 |
| Bos taurus.xls | A0A3Q1LSX3 |
| Bos taurus.xls | A0A3Q1LT55 |
| Bos taurus.xls | F1MUT4 |
| Bos taurus.xls | Q5NTB3 |
| Bos taurus.xls | Q2KJ63 |
| Bos taurus.xls | Q24K22 |
| Bos taurus.xls | Q76BS1 |
| Bos taurus.xls | A7E350 |
| Bosea sp. RC/ | A0A370L6R0 |
| Botryosphaer | R1GKS9 |
| Botryosphaer | R1G4N0 |
| Botryotinia fu | A0A384J9B7 |
| Botryotinia fu | A0A384JR75 |
| Botryotinia fu | M7UKG0 |
| Botryotinia fu | M7U138 |
| Botryotinia fu | G2YNK3 |
| Botryotinia fu | G2XPN8 |
| Brachionus pl | A0A3M7RDQ8 |
| Brachionus pl | A0A3M7T8X2 |
| Brachionus pl | A0A3M7RNS1 |
| Brachionus pl | A0A3M7SMW6 |
| Brachionus pl | A0A3M7RME1 |
| Brachionus pl | A0A3M7RDY0 |
| Brachionus pl | A0A3M7P8R6 |
| Brachionus pl | A0A3M7PKU0 |
| Brachionus pl | A0A3M7RGA3 |
| Brachionus pl | A0A3M7PJ96 |
| Brachionus pl | A0A3M7SM38 |
| Brachionus pl | A0A3M7SFE0 |
| Brachionus pl | A0A3M7RAU7 |
| Brachionus pl | A0A3M7S8F6 |
| Brachionus pl | A0A3M7RN83 |
| Brachionus pl | A0A3M7T9E3 |
| Brachionus pl | A0A3M7SAU8 |
| Brachionus pl | A0A3M7RQA8 |
| Brachionus pl | A0A3M7PUQ9 |
| Brachionus pl | A0A3M7P3V5 |

| | |
|---|---|
| Brachionus pl | A0A3M7P7Y9 |
| Brachionus pl | A0A3M7QIW1 |
| Brachionus pl | A0A3M7REF6 |
| Brachionus pl | A0A3M7SUT2 |
| Brachionus pl | A0A3M7S822 |
| Brachionus pl | A0A3M7QVJ8 |
| Brachionus pl | A0A3M7P6F2 |
| Brachypodiur | I1HU53 |
| Brachypodiur | A0A2K2CXA4 |
| Brachypodiur | A0A2K2CWA4 |
| Brachypodiur | A0A0Q3GP11 |
| Brachypodiur | I1J295 |
| Brachypodiur | A0A2K2CIR5 |
| Brachypodiur | I1J2C4 |
| Brachypodiur | A0A2K2CGC5 |
| Brachypodiur | I1INT0 |
| Brachypodiur | I1J2S3 |
| Brachypodiur | A0A0Q3EAI5 |
| Brachypodiur | I1GMG6 |
| Brachypodiur | A0A2K2CIQ8 |
| Brachypodiur | A0A2K2CIR1 |
| Brachypodiur | A0A0Q3NIJ9 |
| Brachypodiur | A0A2K2CIS7 |
| Brachypodiur | I1IA55 |
| Brachypodiur | A0A2K2CST1 |
| Brachypodiur | A0A0Q3EAD2 |
| Brachypodiur | A0A0Q3JAA8 |
| Brachypodiur | A0A2K2CGA7 |
| Brachypodiur | I1J292 |
| Brachypodiur | A0A2K2CIR0 |
| Brachypodiur | I1GYM6 |
| Brachypodiur | A0A0Q3FVF0 |
| Brachypodiur | I1I349 |
| Brachypodiur | A0A0Q3ED56 |
| Brachypodiur | A0A2K2CIS4 |
| Brachypodiur | A0A0Q3QSF6 |
| Brachypodiur | I1HJD9 |
| Brachypodiur | A0A0Q3EYJ6 |
| Brachypodiur | A0A0Q3IF41 |
| Brachypodiur | A0A2K2D2Y3 |
| Brachypodiur | I1HHX6 |

| | |
|---|---|
| Brachypodiur | A0A0Q3GIB6 |
| Brachypodiur | A0A2K2DI89 |
| Brachypodiur | I1IXN9 |
| Brachypodiur | A0A2K2D7E6 |
| Brachypodiur | A0A0Q3GM35 |
| Brachypodiur | I1INT1 |
| Brachypodiur | I1IUS5 |
| Brachypodiur | I1HW64 |
| Brachypodiur | A0A0Q3R1L3 |
| Brachypodiur | I1HSB4 |
| Brachypodiur | I1HSB6 |
| Brachypodiur | I1GTM1 |
| Brachypodiur | A0A0Q3HD16 |
| Brachypodiur | I1IEU4 |
| Brachypodiur | A0A2K2CZG2 |
| Brachypodiur | A0A0Q3HM63 |
| Brachypodiur | I1HU55 |
| Brachypodiur | A0A2K2DFH8 |
| Brachypodiur | A0A2K2CGL7 |
| Brachypodiur | A0A2K2CIU7 |
| Brachypodiur | A0A0Q3JCV1 |
| Brachypodiur | A0A0Q3JHU3 |
| Brachypodiur | A0A2K2CGB5 |
| Brachypodiur | I1IU59 |
| Brachypodiur | A0A2K2CIT3 |
| Brachypodiur | A0A2K2DFD8 |
| Brachypodiur | A0A2K2DFF0 |
| Brachypodiur | I1GTK9 |
| Brachypodiur | A0A0Q3NYQ7 |
| Brachypodiur | A0A2K2CIR9 |
| Brachypodiur | A0A0Q3RAK5 |
| Brachypodiur | I1GL44 |
| Brachypodiur | I1HAY1 |
| Brachypodiur | I1IXN8 |
| Brachypodiur | A0A0Q3HI45 |
| Brachypodiur | A0A0Q3INA5 |
| Brachypodiur | A0A2K2CIS0 |
| Brachypodiur | A0A0Q3RM98 |
| Brachypodiur | A0A0Q3KXH5 |
| Brachypodiur | I1IFY2 |
| Brachypodiur | I1HSA3 |

| | |
|---|---|
| Brachypodiur | A0A0Q3H363 |
| Brachypodiur | I1GTM2 |
| Brachypodiur | A0A2K2CIS1 |
| Brachypodiur | A0A0Q3LFD5 |
| Brachypodiur | A0A2K2CIT7 |
| Brachypodiur | I1HW63 |
| Brachypodiur | A0A0Q3QIL7 |
| Brachypodiur | A0A0Q3JZ94 |
| Brachypodiur | A0A0Q3QBI3 |
| Brachypodiur | A0A2K2DFD4 |
| Brachypodiur | I1HDE8 |
| Brachypodiur | A0A0Q3IWS7 |
| Brachypodiur | A0A2K2CIP9 |
| Brachypodiur | A0A2K2DFE9 |
| Brachypodiur | A0A2K2CIT2 |
| Brachypodiur | A0A0Q3I905 |
| Brachypodiur | A0A0Q3EAH6 |
| Brachypodiur | A0A0Q3EAI4 |
| Brachypodiur | I1GXK1 |
| Brachypodiur | A0A0Q3IBU0 |
| Brachypodiur | A0A0Q3GCR1 |
| Brachypodiur | I1J2A0 |
| Brachypodiur | I1HEJ3 |
| Brachypodiur | A0A0Q3HSS0 |
| Brachypodiur | A0A0Q3NJG4 |
| Brachypodiur | A0A2K2CIR6 |
| Brachypodiur | I1HDE9 |
| Brachypodiur | A0A0Q3EAE1 |
| Brachypodiur | A0A0Q3HLT2 |
| Brachypodiur | A0A2K2CIQ9 |
| Brachypodiur | A0A2K2DFC3 |
| Brachypodiur | A0A2K2DFC5 |
| Brachypodiur | A0A0Q3LK38 |
| Brachypodiur | A0A2K2DI88 |
| Bradymonas | A0A2Z4FIU7 |
| Bradyrhizobia | A0A257KNM4 |
| Bradyrhizobia | F7QHU6 |
| Bradyrhizobia | A0A3S0G3E0 |
| Bradyrhizobia | A0A363TXD2 |
| Bradyrhizobia | A0A3S0H313 |
| Bradyrhizobia | A0A431M7G0 |

| | |
|---|---|
| Bradyrhizobia | A0A431QX01 |
| Bradyrhizobia | A0A3S0EKZ0 |
| Bradyrhizobiu | A0A1I7FU56 |
| Bradyrhizobiu | A0A1H1Y2P1 |
| Bradyrhizobiu | A0A1X3FPA0 |
| Bradyrhizobiu | A0A1X3GN34 |
| Bradyrhizobiu | A0A1X3GRS6 |
| Bradyrhizobiu | Q89V82 |
| Bradyrhizobiu | A0A2A6N5F9 |
| Bradyrhizobiu | A0A0E4BN53 |
| Bradyrhizobiu | A0A0D1MCK2 |
| Bradyrhizobiu | A0A1E3EC36 |
| Bradyrhizobiu | A0A1M5KHK6 |
| Bradyrhizobiu | A0A1N6F532 |
| Bradyrhizobiu | A0A1M7TM76 |
| Bradyrhizobiu | A0A1M5HCZ9 |
| Bradyrhizobiu | A0A1H4ZSB0 |
| Bradyrhizobiu | A0A410VES2 |
| Bradyrhizobiu | A0A1B1UPJ0 |
| Bradyrhizobiu | A0A023XWN2 |
| Bradyrhizobiu | G7DMB7 |
| Bradyrhizobiu | A0A1L3FRA1 |
| Bradyrhizobiu | A0A0A3XV48 |
| Bradyrhizobiu | A0A1Y2JI88 |
| Bradyrhizobiu | A0A0R3LBX3 |
| Bradyrhizobiu | A0A1M7AMK6 |
| Bradyrhizobiu | A0A1M6WVP9 |
| Bradyrhizobiu | A0A0R3M8G1 |
| Bradyrhizobiu | A0A2H9V6F8 |
| Bradyrhizobiu | A0A151FH31 |
| Bradyrhizobiu | K5CWF7 |
| Bradyrhizobiu | A0A0R3E5M9 |
| Bradyrhizobiu | A0A176YZA1 |
| Bradyrhizobiu | M4Z0Y5 |
| Bradyrhizobiu | M4ZBW7 |
| Bradyrhizobiu | A0A2U8P6V6 |
| Bradyrhizobiu | A0A0R3DMJ3 |
| Bradyrhizobiu | A0A1B9ZC66 |
| Bradyrhizobiu | A0A0R3MRW5 |
| Bradyrhizobiu | A0A1C3WWF3 |
| Bradyrhizobiu | A5EA89 |

| | |
|---|---|
| Bradyrhizobiu | A4YQL0 |
| Bradyrhizobiu | A4Z2E2 |
| Bradyrhizobiu | H0SMB8 |
| Bradyrhizobiu | H0SCU6 |
| Bradyrhizobiu | H0SBZ7 |
| Bradyrhizobiu | A0A2U8PT28 |
| Bradyrhizobiu | A0A2U8PN93 |
| Bradyrhizobiu | A0A2U8PT94 |
| Bradyrhizobiu | A0A2U8Q528 |
| Bradyrhizobiu | A0A2S6M8B5 |
| Bradyrhizobiu | A0A1Q5R4X0 |
| Bradyrhizobiu | A0A150UDS9 |
| Bradyrhizobiu | A0A176Z2D2 |
| Bradyrhizobiu | A0A109JGJ9 |
| Bradyrhizobiu | A0A1V5EYR3 |
| Bradyrhizobiu | A0A2A6NCZ9 |
| Bradyrhizobiu | A0A410W142 |
| Bradyrhizobiu | A0A109QD69 |
| Bradyrhizobiu | A0A1I3S3F5 |
| Bradyrhizobiu | A0A1I3J718 |
| Bradyrhizobiu | U1H1Q2 |
| Bradyrhizobiu | A0A150ZHL9 |
| Bradyrhizobiu | A0A0S6UQ35 |
| Bradyrhizobiu | A0A160UQP7 |
| Bradyrhizobiu | A0A1I3V8E7 |
| Bradyrhizobiu | A0A1I5BXB3 |
| Bradyrhizobiu | A0A2M8RAB9 |
| Bradyrhizobiu | A0A0Q6A3Y8 |
| Bradyrhizobiu | A0A1B9Z3Q7 |
| Bradyrhizobiu | A0A0D7NXE7 |
| Bradyrhizobiu | A0A0D7N7J5 |
| Bradyrhizobiu | A0A0D7Q5V0 |
| Bradyrhizobiu | A0A430MFG0 |
| Bradyrhizobiu | A0A2T2QQE6 |
| Bradyrhizobiu | A0A2T2PZ36 |
| Bradyrhizobiu | A0A2T2PSY2 |
| Bradyrhizobiu | A0A2T2Q0F5 |
| Bradyrhizobiu | A0A2T2PT25 |
| Bradyrhizobiu | A0A2T2PSY6 |
| Bradyrhizobiu | A0A2T2Q4P1 |
| Bradyrhizobiu | A0A1Q5SAH3 |

| | |
|---|---|
| Bradyrhizobiu | A0A1Q5R9S2 |
| Bradyrhizobiu | A0A1I4H0I3 |
| Bradyrhizobiu | A0A1H8BXF8 |
| Bradyrhizobiu | A0A1Y6KJQ2 |
| Bradyrhizobiu | A0A1Y6KP04 |
| Bradyrhizobiu | A0A2U3PS34 |
| Bradyrhizobiu | A0A1G7GWB1 |
| Bradyrhizobiu | A0A1G8B4K1 |
| Bradyrhizobiu | A0A1I4URV1 |
| Bradyrhizobiu | A0A3N6PAA8 |
| Bradyrhizobiu | I0FZQ2 |
| Bradyrhizobiu | A0A2K8Y6Q4 |
| Bradyrhizobiu | H0STR9 |
| Bradyrhizobiu | H0TBX2 |
| Bradyrhizobiu | H0TGS9 |
| Bradyrhizobiu | A0A2M6UE20 |
| Bradyrhizobiu | A0A1R1QEF7 |
| Bradyrhizobiu | A0A2A2VLQ2 |
| Bradyrhizobiu | I2QLW7 |
| Bradyrhizobiu | H5YCF8 |
| Bradyrhizobiu | A0A2A6PYD0 |
| Bradyrhizobiu | J2W3F2 |
| Bradyrhizobiu | J3I044 |
| Bradyrhizobiu | A0A160UAV3 |
| Bradyrhizobiu | A0A0R3KWP9 |
| Bradyrhizobiu | A0A0R3CA11 |
| Bradysia odor | A0A385GLI2 |
| Bradysia odor | A0A385GLJ0 |
| Branchiostom | C3YDL6 |
| Branchiostom | C3Z071 |
| Branchiostom | C3YAZ6 |
| Branchiostom | C3ZYT2 |
| Branchiostom | C3ZFD3 |
| Branchiostom | C3ZHY1 |
| Branchiostom | C3YYM3 |
| Branchiostom | C3ZAW6 |
| Branchiostom | C3XQ58 |
| Branchiostom | C3XQ51 |
| Branchiostom | C3Y7N4 |
| Branchiostom | C3YP82 |
| Branchiostom | C3ZLM9 |

| | |
|---|---|
| Branchiostom | C3Z0V1 |
| Branchiostom | C3XPW8 |
| Branchiostom | C3Y305 |
| Branchiostom | C3ZYG9 |
| Branchiostom | C3ZMW2 |
| Branchiostom | C3ZMB3 |
| Branchiostom | C3Y3P1 |
| Branchiostom | C3ZGD7 |
| Branchiostom | C3Y566 |
| Branchiostom | C3Z0R2 |
| Branchiostom | C3ZJ84 |
| Branchiostom | C3YU46 |
| Branchiostom | C3XY24 |
| Branchiostom | C3YC74 |
| Branchiostom | C3ZFT9 |
| Branchiostom | C3ZHS3 |
| Branchiostom | C3Y7E8 |
| Branchiostom | C3XW29 |
| Branchiostom | C3YDL9 |
| Branchiostom | C3ZT44 |
| Branchiostom | C3XUJ8 |
| Branchiostom | C3YP09 |
| Branchiostom | C3ZDY0 |
| Branchiostom | C3YS46 |
| Branchiostom | C3YDM0 |
| Branchiostom | C3ZZJ9 |
| Branchiostom | C3YCI7 |
| Branchiostom | C3YU47 |
| Branchiostom | C3YNF1 |
| Branchiostom | C3ZHW8 |
| Branchiostom | C3ZG87 |
| Branchiostom | C3XY53 |
| Branchiostom | C3YWE1 |
| Branchiostom | C3XW28 |
| Branchiostom | C3YMU3 |
| Branchiostom | C3ZK10 |
| Brassica barr | Q8LP65 |
| Brassica camp | Q84KY8 |
| Brassica camp | A0A397YWD8 |
| Brassica camp | Q84KX9 |
| Brassica camp | A0A3P5Z2T0 |

| | |
|---|---|
| Brassica cam | Q1XIH3 |
| Brassica cam | A8QWH1 |
| Brassica cam | A0A398A0U2 |
| Brassica cam | A0A1L7NT06 |
| Brassica cam | O23856 |
| Brassica cam | A0A397YDG2 |
| Brassica cam | B5U9A7 |
| Brassica cam | A0A397YVY9 |
| Brassica cam | A0A397YF45 |
| Brassica cam | O23848 |
| Brassica cam | A0A3P6C3Y6 |
| Brassica cam | A0A3P5ZPK2 |
| Brassica cam | A0A3P5XWY0 |
| Brassica cam | A0A397YV83 |
| Brassica cam | A0A397YFB7 |
| Brassica cam | A0A3P5Z9J5 |
| Brassica cam | A0A3P6CY69 |
| Brassica cam | A0A3P6D969 |
| Brassica cam | A0A3P5YJ21 |
| Brassica cam | A0A397YBC1 |
| Brassica cam | Q2MHJ1 |
| Brassica cam | A0A3P6CB57 |
| Brassica cam | Q948X2 |
| Brassica cam | A0A397Z250 |
| Brassica cam | A0A397YV95 |
| Brassica cam | A0A397XSU5 |
| Brassica cam | A8QZG2 |
| Brassica cam | B5U9A8 |
| Brassica cam | Q948X1 |
| Brassica cam | A0A397YA29 |
| Brassica cam | A0A397YPP3 |
| Brassica cam | Q9AYP7 |
| Brassica cam | O23850 |
| Brassica cam | O23855 |
| Brassica cam | Q6SVA0 |
| Brassica cam | A0A3P6A0C0 |
| Brassica cam | Q7DN95 |
| Brassica cam | A0A3P6ASY7 |
| Brassica cam | A0A398A9P6 |
| Brassica cam | A0A3P5ZV33 |
| Brassica cam | A0A3P6BWB7 |

| | |
|---|---|
| Brassica camp | A0A3P5YSC4 |
| Brassica camp | A0A397ZSN3 |
| Brassica camp | Q2EMZ8 |
| Brassica camp | Q6SVA3 |
| Brassica camp | A0A3P6A3C2 |
| Brassica camp | A8QZC5 |
| Brassica camp | A0A397XRI1 |
| Brassica camp | A0A3P5Z8N8 |
| Brassica camp | Q39279 |
| Brassica camp | A0A3P5YE62 |
| Brassica camp | A0A3P5XWD4 |
| Brassica camp | Q84KX7 |
| Brassica camp | A0A398AQ76 |
| Brassica camp | O23859 |
| Brassica camp | Q84KX6 |
| Brassica camp | Q39276 |
| Brassica camp | A0A3P5YCU6 |
| Brassica camp | A0A398AQ52 |
| Brassica camp | Q39280 |
| Brassica camp | Q18PE5 |
| Brassica camp | Q84KZ0 |
| Brassica camp | A0A3P6C0F7 |
| Brassica camp | O23851 |
| Brassica camp | A0A398A6C5 |
| Brassica camp | O23745 |
| Brassica camp | Q84KY9 |
| Brassica camp | A0A3P6CGF5 |
| Brassica camp | O23847 |
| Brassica camp | A0A398ARS1 |
| Brassica camp | A4UWM4 |
| Brassica camp | A0A3P6C098 |
| Brassica camp | A0A398AF65 |
| Brassica camp | A0A397YVY3 |
| Brassica camp | O23860 |
| Brassica camp | A0A3P5ZGB7 |
| Brassica camp | O23849 |
| Brassica camp | Q04034 |
| Brassica camp | O23861 |
| Brassica camp | Q84KY2 |
| Brassica camp | A0A397YCG0 |
| Brassica camp | Q84KY4 |

| | |
|---|---|
| Brassica camp | A8QWH4 |
| Brassica camp | A0A3P5Z9S3 |
| Brassica camp | A0A398AMH2 |
| Brassica camp | Q84KY1 |
| Brassica camp | A0A3P6A681 |
| Brassica camp | A0A397Y539 |
| Brassica camp | Q2MHJ2 |
| Brassica camp | E2IV49 |
| Brassica camp | A0A3P5Z8U6 |
| Brassica camp | Q84KY6 |
| Brassica camp | A0A1L7NT17 |
| Brassica camp | A0A397Y192 |
| Brassica camp | A0A397ZWZ8 |
| Brassica camp | A0A1L7NT09 |
| Brassica camp | O23852 |
| Brassica camp | O23854 |
| Brassica camp | A0A397KYV2 |
| Brassica camp | A0A398AKR7 |
| Brassica camp | Q39277 |
| Brassica camp | A0A3P5ZXM2 |
| Brassica camp | A0A397XTY3 |
| Brassica camp | A0A3P5ZIB6 |
| Brassica camp | Q84KY3 |
| Brassica camp | Q84KY0 |
| Brassica camp | Q41222 |
| Brassica camp | A0A3P6A1E3 |
| Brassica camp | A8QZG7 |
| Brassica camp | A0A3P6BVP4 |
| Brassica camp | Q8W407 |
| Brassica camp | Q84KY7 |
| Brassica camp | G3EXH7 |
| Brassica camp | A0A397YV80 |
| Brassica camp | Q9AYP5 |
| Brassica camp | Q948W9 |
| Brassica camp | A0A397Y1U1 |
| Brassica camp | A0A397XVY9 |
| Brassica camp | Q948X0 |
| Brassica camp | A0A398A3C5 |
| Brassica camp | A0A397ZLT6 |
| Brassica camp | A0A3P5ZRW3 |
| Brassica camp | A8QZC9 |

| | |
|---|---|
| Brassica camp | A0A398ASE3 |
| Brassica camp | A0A397KY95 |
| Brassica camp | A8QZF8 |
| Brassica camp | A0A398ATI7 |
| Brassica camp | Q9AYP2 |
| Brassica camp | A0A3P6ACK6 |
| Brassica camp | A0A3P5YRC2 |
| Brassica camp | A0A397YRH6 |
| Brassica camp | A0A397KZ86 |
| Brassica camp | Q8S9B0 |
| Brassica camp | B5U9A6 |
| Brassica camp | A0A3P6BCU6 |
| Brassica camp | Q6SV99 |
| Brassica camp | A0A397YN43 |
| Brassica camp | A0A397YFK7 |
| Brassica camp | A0A397YPJ3 |
| Brassica camp | Q6SV98 |
| Brassica camp | A0A1L7NT04 |
| Brassica camp | E2IV46 |
| Brassica camp | A0A397Z248 |
| Brassica camp | Q27IZ4 |
| Brassica camp | A0A1L7NT22 |
| Brassica camp | A0A397YJM2 |
| Brassica camp | A0A1L7NT10 |
| Brassica camp | A8QZG5 |
| Brassica camp | A0A398A3A1 |
| Brassica camp | A0A3P6C7S0 |
| Brassica camp | A0A397L944 |
| Brassica camp | Q84N09 |
| Brassica camp | Q1XIH1 |
| Brassica camp | A0A397XUJ7 |
| Brassica camp | Q1XHT4 |
| Brassica camp | O80430 |
| Brassica camp | A0A397XT43 |
| Brassica camp | A0A397YZX3 |
| Brassica camp | A8QWG5 |
| Brassica camp | O80344 |
| Brassica camp | A0A397ZXN6 |
| Brassica camp | A0A397YGY4 |
| Brassica camp | A0A3P5ZWE1 |
| Brassica camp | A0A397XUI4 |

| | |
|---|---|
| Brassica camp | A0A397Y2G3 |
| Brassica camp | E2IV47 |
| Brassica camp | A0A398AKX8 |
| Brassica camp | A0A3P6C0A8 |
| Brassica camp | O23857 |
| Brassica camp | A0A3P6C053 |
| Brassica camp | A0A397Z461 |
| Brassica camp | A0A397ZIG8 |
| Brassica camp | A0A1L7NT25 |
| Brassica camp | Q9ZNW9 |
| Brassica camp | A0A1L7NT13 |
| Brassica camp | A0A397Y4Y8 |
| Brassica camp | Q9SXK6 |
| Brassica camp | A0A3P5YMA1 |
| Brassica camp | A0A398A4W5 |
| Brassica camp | O23853 |
| Brassica camp | Q9AYP6 |
| Brassica camp | A0A397YQ80 |
| Brassica camp | A0A3P5ZPZ5 |
| Brassica camp | Q84KX8 |
| Brassica camp | A0A3P6CIH1 |
| Brassica camp | O23858 |
| Brassica camp | A0A3P5YA39 |
| Brassica camp | A0A3P5ZWS7 |
| Brassica camp | A0A397XQX1 |
| Brassica camp | A0A397L1V4 |
| Brassica camp | A0A3P6DPZ9 |
| Brassica camp | A0A398A9U1 |
| Brassica camp | A0A397ZGZ8 |
| Brassica camp | A0A397YV89 |
| Brassica camp | A0A3P5YQL4 |
| Brassica camp | A0A3P5Y7E1 |
| Brassica camp | Q1XIH2 |
| Brassica camp | A0A397YNB1 |
| Brassica camp | A0A3P6BQI0 |
| Brassica camp | A8QZD1 |
| Brassica camp | A0A3P6A4U7 |
| Brassica camp | A0A397YA55 |
| Brassica camp | Q84KY5 |
| Brassica camp | Q9SAZ7 |
| Brassica camp | A0A397XZ92 |

| | |
|---|---|
| Brassica camp | Q1XHT5 |
| Brassica camp | O80343 |
| Brassica camp | Q39278 |
| Brassica camp | A0A3P6B4Q3 |
| Brassica camp | A0A3P6C2F5 |
| Brassica camp | A0A397YX37 |
| Brassica camp | Q6SVA2 |
| Brassica camp | O48512 |
| Brassica camp | Q948X3 |
| Brassica camp | Q9AYP8 |
| Brassica camp | A0A3P5YM72 |
| Brassica camp | A0A3P5YDK4 |
| Brassica camp | A0A3P5ZQK6 |
| Brassica camp | A0A1L7NT14 |
| Brassica camp | A0A3P5ZUK5 |
| Brassica camp | E2IV48 |
| Brassica camp | Q9SXK7 |
| Brassica camp | Q18PE6 |
| Brassica camp | A0A397ZIB8 |
| Brassica camp | A0A397Z263 |
| Brassica camp | Q2PML8 |
| Brassica camp | Q39282 |
| Brassica camp | Q84N10 |
| Brassica camp | Q9ZNW6 |
| Brassica camp | A0A397ZR25 |
| Brassica camp | Q04168 |
| Brassica creti | A0A3N6S4L1 |
| Brassica creti | A0A3N6QXX1 |
| Brassica creti | A0A3N6QKJ3 |
| Brassica creti | A0A3N6TM80 |
| Brassica creti | A0A3N6QYA9 |
| Brassica creti | A0A3N6QM77 |
| Brassica creti | A0A3N6R7H2 |
| Brassica creti | A0A3N6QNP4 |
| Brassica creti | A0A3N6PZG5 |
| Brassica creti | A0A3N6QQN7 |
| Brassica creti | A0A3N6UQ88 |
| Brassica creti | A0A3N6RPS9 |
| Brassica creti | A0A3N6T5B6 |
| Brassica creti | A0A3N6S057 |
| Brassica creti | A0A3N6Q5N7 |

| | |
|---|---|
| Brassica creti | A0A3N6S1R2 |
| Brassica creti | A0A3N6U997 |
| Brassica creti | A0A3N6T699 |
| Brassica creti | A0A3N6SZL5 |
| Brassica creti | A0A3N6RNN4 |
| Brassica creti | A0A3N6UJ64 |
| Brassica creti | A0A3N6QYV1 |
| Brassica creti | A0A3N6UHA9 |
| Brassica creti | A0A3N6RZC7 |
| Brassica creti | A0A3N6S1W1 |
| Brassica creti | A0A3N6QNP5 |
| Brassica creti | A0A3N6RFD8 |
| Brassica creti | A0A3N6SMY5 |
| Brassica creti | A0A3N6R014 |
| Brassica creti | A0A3N6R511 |
| Brassica creti | A0A3N6QTL5 |
| Brassica creti | A0A3N6QXV3 |
| Brassica defle | Q8LP64 |
| Brassica insul | Q9SBK7 |
| Brassica mau | Q8LP62 |
| Brassica napu | Q01963 |
| Brassica napu | Q9M4E9 |
| Brassica napu | Q7FPE1 |
| Brassica napu | A0A078FY82 |
| Brassica napu | A0A078JHP5 |
| Brassica napu | Q43393 |
| Brassica napu | A0A078IH30 |
| Brassica napu | A0A078JPB3 |
| Brassica napu | A0A078GD60 |
| Brassica napu | A0A078G3D0 |
| Brassica napu | A0A078FY93 |
| Brassica napu | A0A078JQF9 |
| Brassica napu | A0A078HBI2 |
| Brassica napu | Q08700 |
| Brassica napu | A0A078I0E8 |
| Brassica napu | A0A078HA89 |
| Brassica napu | Q6SV95 |
| Brassica napu | A0A078HDD4 |
| Brassica napu | Q6SV97 |
| Brassica napu | A0A078HAC3 |
| Brassica napu | A0A078HEK1 |

| | |
|---|---|
| Brassica napu | A0A078IMG1 |
| Brassica napu | A0A078HR13 |
| Brassica napu | A0A078IFP5 |
| Brassica napu | A0A078HVX7 |
| Brassica napu | A0A078IAX0 |
| Brassica napu | A0A078HN20 |
| Brassica napu | A0A078F3B5 |
| Brassica napu | Q6SV92 |
| Brassica napu | A0A078GIA3 |
| Brassica napu | Q6SV96 |
| Brassica napu | Q39356 |
| Brassica napu | A0A078GM80 |
| Brassica napu | A0A078JYB1 |
| Brassica napu | A0A078H803 |
| Brassica napu | A0A078JCU5 |
| Brassica napu | A0A078HR81 |
| Brassica napu | A0A078J7X9 |
| Brassica napu | A0A078IDY2 |
| Brassica napu | A0A078GN24 |
| Brassica napu | A0A078HD30 |
| Brassica napu | A0A078H8L5 |
| Brassica napu | A0A078G7I9 |
| Brassica napu | A0A078HAD2 |
| Brassica napu | A0A078F337 |
| Brassica napu | A0A078I9I0 |
| Brassica napu | A0A078ILK5 |
| Brassica napu | A0A078H3H5 |
| Brassica napu | A0A078JMJ4 |
| Brassica napu | A0A078HEM0 |
| Brassica napu | A4UWL9 |
| Brassica napu | A0A078GS40 |
| Brassica napu | A0A078JKE7 |
| Brassica napu | A0A078FML4 |
| Brassica napu | A0A078FHJ4 |
| Brassica napu | A0A078FPN5 |
| Brassica napu | Q39363 |
| Brassica napu | A0A078J557 |
| Brassica napu | A0A078II36 |
| Brassica napu | A0A078IIQ7 |
| Brassica napu | A0A078HDC2 |
| Brassica napu | A0A078JFN3 |

| | |
|---|---|
| Brassica napu | A0A078FRP9 |
| Brassica napu | A0A078FIU1 |
| Brassica napu | A6P327 |
| Brassica napu | A0A078JBM2 |
| Brassica napu | A0A078GVV6 |
| Brassica napu | A0A078IDX4 |
| Brassica napu | Q39354 |
| Brassica napu | A0A078FI59 |
| Brassica napu | A0A078HT86 |
| Brassica napu | A0A078G2V4 |
| Brassica napu | A0A078JE37 |
| Brassica napu | A0A078FQ97 |
| Brassica napu | A0A078I9I3 |
| Brassica napu | A0A078FEU1 |
| Brassica napu | A0A078FKQ4 |
| Brassica napu | A0A078JIW9 |
| Brassica napu | A0A078G2W5 |
| Brassica napu | A0A078GU68 |
| Brassica napu | A0A078J1N7 |
| Brassica napu | A0A078G8J1 |
| Brassica napu | A0A078JWL9 |
| Brassica napu | A0A078J032 |
| Brassica napu | A0A078IYH8 |
| Brassica napu | A0A078JVW6 |
| Brassica napu | A0A078G3S4 |
| Brassica napu | A0A078FWL9 |
| Brassica napu | A0A078GJI3 |
| Brassica napu | A0A078IFM7 |
| Brassica napu | A0A078FQ82 |
| Brassica napu | A6P326 |
| Brassica napu | A0A078G3D4 |
| Brassica napu | A0A078FPH2 |
| Brassica napu | A0A078HAQ2 |
| Brassica napu | A0A078GYC5 |
| Brassica napu | A0A078HCX8 |
| Brassica napu | A0A078INX3 |
| Brassica napu | Q6SV94 |
| Brassica napu | A0A078JB85 |
| Brassica napu | A0A078HPD1 |
| Brassica napu | A0A078FIU7 |
| Brassica napu | A0A078GDP5 |

| | |
|---|---|
| Brassica napu | A0A078JQK2 |
| Brassica napu | A0A078EZE7 |
| Brassica napu | Q7DMS5 |
| Brassica napu | A0A078G200 |
| Brassica napu | A0A078IKU0 |
| Brassica napu | A0A078H3H1 |
| Brassica napu | A6P328 |
| Brassica napu | A0A078FC62 |
| Brassica napu | A0A078FE33 |
| Brassica napu | A4UWL8 |
| Brassica napu | A4UWL4 |
| Brassica napu | A4UWM3 |
| Brassica napu | A4UWL3 |
| Brassica napu | Q39357 |
| Brassica napu | A4UWL5 |
| Brassica napu | A4UWL6 |
| Brassica napu | Q04808 |
| Brassica nigra | Q8LP61 |
| Brassica olera | P22552 |
| Brassica olera | P22553 |
| Brassica olera | P22551 |
| Brassica olera | A0A0D3C9A8 |
| Brassica olera | A0A0D3BLV5 |
| Brassica olera | A0A0D2ZVD2 |
| Brassica olera | A0A0D3AAQ0 |
| Brassica olera | A0A0D3E522 |
| Brassica olera | A0A0D3DXZ3 |
| Brassica olera | A0A0D3A6W5 |
| Brassica olera | A0A0D3AAP8 |
| Brassica olera | A0A0D3CYG2 |
| Brassica olera | A0A0D3E1N6 |
| Brassica olera | A0A0D3E528 |
| Brassica olera | A0A0D3DA71 |
| Brassica olera | A0A0D3DP33 |
| Brassica olera | A0A0D2ZU01 |
| Brassica olera | A0A0D3AAR4 |
| Brassica olera | A0A0D3CJQ9 |
| Brassica olera | A0A0D2ZVF6 |
| Brassica olera | A0A0D3BVD8 |
| Brassica olera | A0A0D3DXZ2 |
| Brassica olera | A0A0D3BVD9 |

| | |
|---|---|
| Brassica olera | A0A0D3CM95 |
| Brassica olera | A0A0D3CY96 |
| Brassica olera | A0A0D3BFE7 |
| Brassica olera | A0A0D3E524 |
| Brassica olera | A0A0D3C5Q0 |
| Brassica olera | A0A0D3BXW1 |
| Brassica olera | A0A0D3BCW1 |
| Brassica olera | A0A0D3C227 |
| Brassica olera | A0A0D3E3G0 |
| Brassica olera | A0A0D3C9A3 |
| Brassica olera | A0A0D3DGD4 |
| Brassica olera | A0A0D3C9A0 |
| Brassica olera | A0A0D3DDG1 |
| Brassica olera | A0A0D3BW89 |
| Brassica olera | A0A0D3DE12 |
| Brassica olera | A0A0D3E3F9 |
| Brassica olera | A0A0D3AD84 |
| Brassica olera | A0A0D3CY99 |
| Brassica olera | A0A0D3DC95 |
| Brassica olera | A0A0D3AAR5 |
| Brassica olera | A0A0D3DMB4 |
| Brassica olera | A0A0D3AAP9 |
| Brassica olera | A0A0D3BYM9 |
| Brassica olera | A0A0D3EAU1 |
| Brassica olera | A0A0D3CY95 |
| Brassica olera | A0A0D3D1P0 |
| Brassica olera | A0A0D3C9A9 |
| Brassica olera | A0A0D2ZVS8 |
| Brassica olera | A0A0D3DVG2 |
| Brassica olera | A0A0D3BT69 |
| Brassica olera | A0A0D3A4U3 |
| Brassica olera | A0A0D3DXV5 |
| Brassica olera | A0A0D3B6D1 |
| Brassica olera | A0A0D2ZWS7 |
| Brassica olera | A0A0D3C538 |
| Brassica olera | A0A0D3C9C2 |
| Brassica olera | A0A0D3DXZ4 |
| Brassica olera | A0A0D3BVD6 |
| Brassica olera | A0A0D3C9A2 |
| Brassica olera | A0A0D3C5D3 |
| Brassica olera | A0A0D3DZL7 |

| | |
|---|---|
| Brassica olera | A0A0D3A4U2 |
| Brassica olera | A0A0D3ASG5 |
| Brassica olera | A0A0D3ASA4 |
| Brassica olera | A0A0D3DP20 |
| Brassica olera | A0A0D3D3Q8 |
| Brassica olera | A0A0D3CVY3 |
| Brassica olera | A0A0D3BAW1 |
| Brassica olera | A0A0D3AYW3 |
| Brassica olera | A0A0D3E523 |
| Brassica olera | A0A0D3BVG5 |
| Brassica olera | A0A0D3A6W4 |
| Brassica olera | A0A0D3DP40 |
| Brassica olera | A0A0D3E5Z4 |
| Brassica olera | A0A0D3EI38 |
| Brassica olera | A0A0D3DDG2 |
| Brassica olera | A0A0D3CEW1 |
| Brassica olera | A0A0D3B529 |
| Brassica olera | A0A0D3C9B0 |
| Brassica olera | A0A0D3C999 |
| Brassica olera | A0A0D3DP24 |
| Brassica olera | A0A0D3A2M9 |
| Brassica olera | A8VP90 |
| Brassica olera | Q09092 |
| Brassica olera | P17840 |
| Brassica olera | P17841 |
| Brassica olera | P07761 |
| Brassica olera | Q9SXH4 |
| Brassica olera | Q93VX9 |
| Brassica olera | O23743 |
| Brassica olera | Q93VR1 |
| Brassica olera | Q93VY7 |
| Brassica olera | Q93V96 |
| Brassica olera | A0A3P6DR72 |
| Brassica olera | O04383 |
| Brassica olera | Q9MB87 |
| Brassica olera | O23832 |
| Brassica olera | Q84KU6 |
| Brassica olera | Q8S9B2 |
| Brassica olera | A0A3P6EAU3 |
| Brassica olera | Q84KV5 |
| Brassica olera | Q1ET35 |

| | |
|---|---|
| Brassica olera | A0A3P6AZG2 |
| Brassica olera | A0A3P6DSC9 |
| Brassica olera | Q9SXH9 |
| Brassica olera | Q84KV2 |
| Brassica olera | Q39392 |
| Brassica olera | Q84KV1 |
| Brassica olera | Q93YJ8 |
| Brassica olera | A0A3P6E9L0 |
| Brassica olera | Q84KX3 |
| Brassica olera | A0A3P6CXY8 |
| Brassica olera | A0A3P6DPD4 |
| Brassica olera | A0A3P6BST4 |
| Brassica olera | A0A3P6CG45 |
| Brassica olera | A0A3P6CJK8 |
| Brassica olera | Q9ZQU7 |
| Brassica olera | B5U9A4 |
| Brassica olera | Q9T0M1 |
| Brassica olera | A0A3P6G216 |
| Brassica olera | Q84KW1 |
| Brassica olera | Q84KW3 |
| Brassica olera | A0A3P6DQ69 |
| Brassica olera | A0A3P6DM94 |
| Brassica olera | Q93YK2 |
| Brassica olera | A0A3P6H645 |
| Brassica olera | A0A3P6DJE8 |
| Brassica olera | O23835 |
| Brassica olera | A0A3P6EPZ7 |
| Brassica olera | O23744 |
| Brassica olera | Q9ZQU6 |
| Brassica olera | A0A3P6GT52 |
| Brassica olera | Q7DLP6 |
| Brassica olera | Q84KX5 |
| Brassica olera | Q93YJ7 |
| Brassica olera | Q84KW2 |
| Brassica olera | A0A3P6GDI9 |
| Brassica olera | Q39390 |
| Brassica olera | A0A3P6CMU3 |
| Brassica olera | O23844 |
| Brassica olera | A0A3P6FD20 |
| Brassica olera | A0A3P6EHB3 |
| Brassica olera | A0A3P6DJA8 |

| | |
|---|---|
| Brassica olera | A0A3P6EN76 |
| Brassica olera | A0A3P6G8V9 |
| Brassica olera | A0A3P6AM61 |
| Brassica olera | Q84KX2 |
| Brassica olera | Q93YK0 |
| Brassica olera | O23836 |
| Brassica olera | O23838 |
| Brassica olera | Q84KV7 |
| Brassica olera | Q84KW7 |
| Brassica olera | A0A3P6DCU3 |
| Brassica olera | A0A3P6GW39 |
| Brassica olera | A0A3P6FM07 |
| Brassica olera | A0A3P6CYJ8 |
| Brassica olera | A8QZJ0 |
| Brassica olera | O23837 |
| Brassica olera | Q84KW0 |
| Brassica olera | A0A3P6ECQ4 |
| Brassica olera | A0A3P6CA48 |
| Brassica olera | Q1ET30 |
| Brassica olera | A0A3P6BMB0 |
| Brassica olera | Q9MB88 |
| Brassica olera | Q84KX4 |
| Brassica olera | O23842 |
| Brassica olera | O04384 |
| Brassica olera | Q9SXH6 |
| Brassica olera | Q93VN5 |
| Brassica olera | O23841 |
| Brassica olera | A0A3P6FB97 |
| Brassica olera | A8QZI2 |
| Brassica olera | A0A3P6GBD6 |
| Brassica olera | A0A3P6G6E7 |
| Brassica olera | Q8W408 |
| Brassica olera | A0A3P6BHJ6 |
| Brassica olera | A0A3P6F1R2 |
| Brassica olera | A0A3P6FS98 |
| Brassica olera | A0A3P6DWV5 |
| Brassica olera | Q1XIG9 |
| Brassica olera | O23740 |
| Brassica olera | Q9SXH7 |
| Brassica olera | O23839 |
| Brassica olera | Q84KW4 |

| | |
|---|---|
| Brassica olera | Q9SXH5 |
| Brassica olera | T1Q059 |
| Brassica olera | A8QZI1 |
| Brassica olera | Q39393 |
| Brassica olera | Q84KV0 |
| Brassica olera | Q84KV3 |
| Brassica olera | O23840 |
| Brassica olera | A0A3P6F1Q3 |
| Brassica olera | A0A3P6GBB9 |
| Brassica olera | Q84KU5 |
| Brassica olera | O23863 |
| Brassica olera | A0A3P6EDW2 |
| Brassica olera | Q84KV4 |
| Brassica olera | O23895 |
| Brassica olera | Q84KU7 |
| Brassica olera | Q43410 |
| Brassica olera | A0A3P6GFP0 |
| Brassica olera | Q84KV9 |
| Brassica olera | A0A3P6CQM2 |
| Brassica olera | A0A3P6FF23 |
| Brassica olera | A0A3P6EV23 |
| Brassica olera | Q1ET32 |
| Brassica olera | A0A3P6DR77 |
| Brassica olera | A0A3P6AP88 |
| Brassica olera | Q84KU9 |
| Brassica olera | A0A3P6F267 |
| Brassica olera | P93068 |
| Brassica olera | A0A3P6F441 |
| Brassica olera | Q9SAN1 |
| Brassica olera | Q9SXH8 |
| Brassica olera | Q9T0M0 |
| Brassica olera | Q84KW5 |
| Brassica olera | Q84KW8 |
| Brassica olera | Q39389 |
| Brassica olera | Q84KU8 |
| Brassica olera | Q84KV6 |
| Brassica olera | O23834 |
| Brassica olera | Q8W409 |
| Brassica olera | O23833 |
| Brassica olera | Q39391 |
| Brassica olera | A0A3P6C6P4 |

| | |
|---|---|
| Brassica olera | A0A3P6E4S9 |
| Brassica olera | O23843 |
| Brassica olera | A0A3P6FEZ1 |
| Brassica olera | Q84KW6 |
| Brassica olera | A0A3P6G9V7 |
| Brassica olera | O23846 |
| Brassica olera | Q03349 |
| Brassica olera | T1Q7V5 |
| Brassica olera | Q84KW9 |
| Brassica olera | A0A3P6GQQ4 |
| Brassica olera | A0A3P6EI65 |
| Brassica olera | Q93YK1 |
| Brassica olera | A0A3P6EAA2 |
| Brassica olera | Q43411 |
| Brassica olera | A8QZH0 |
| Brassica olera | Q84KV8 |
| Brassica olera | A0A3P6CK21 |
| Brassica olera | O23862 |
| Brassica olera | Q9XFW5 |
| Brassica olera | Q1ET31 |
| Brassica olera | Q41177 |
| Brassica olera | A0A3P6FAU8 |
| Brassica olera | A0A3P6FN43 |
| Brassica olera | A0A3P6FGK6 |
| Brassica olera | A8QZH1 |
| Brassica olera | O23845 |
| Brassica olera | Q84KX1 |
| Brassica olera | Q93YJ9 |
| Brassica olera | Q84KX0 |
| Brassica olera | Q84KU4 |
| Brassica olera | A0A3P6B1L3 |
| Brassica olera | A0A3P6BNH0 |
| Brassica olera | A0A3P6GIA4 |
| Brassica oxyr | Q8LP60 |
| Brassica rapa | M4DRB6 |
| Brassica rapa | M4F1X0 |
| Brassica rapa | M4CIZ8 |
| Brassica rapa | M4DTL2 |
| Brassica rapa | M4FBV1 |
| Brassica rapa | M4EZL4 |
| Brassica rapa | M4CBQ0 |

Brassica rapa M4D8N1
Brassica rapa M4ESL3
Brassica rapa M4EE76
Brassica rapa M4D938
Brassica rapa M4CHF2
Brassica rapa M4ERK3
Brassica rapa M4F4F3
Brassica rapa M4DTM0
Brassica rapa M4EE77
Brassica rapa M4ECM7
Brassica rapa M4DE63
Brassica rapa M4FBV0
Brassica rapa M4ERK7
Brassica rapa M4DK16
Brassica rapa M4DTL3
Brassica rapa M4F8C2
Brassica rapa M4DTM3
Brassica rapa M4DK08
Brassica rapa M4DTL1
Brassica rapa M4D939
Brassica rapa M4CIU6
Brassica rapa M4EC54
Brassica rapa M4ESI7
Brassica rapa M4FHG6
Brassica rapa M4DAL3
Brassica rapa M4DTL9
Brassica rapa M4CIZ5
Brassica rapa M4DKI6
Brassica rapa M4ERK8
Brassica rapa M4FC19
Brassica rapa M4C7T3
Brassica rapa M4D4D3
Brassica rapa M4CU98
Brassica rapa M4EE78
Brassica rapa M4E2N1
Brassica rapa M4DTM1
Brassica rapa M4DK77
Brassica rapa M4DTK0
Brassica rapa M4FBW3
Brassica rapa M4D3D6
Brassica rapa M4DK14

| | |
|---|---|
| Brassica rapa | M4CZ69 |
| Brassica rapa | M4CCH3 |
| Brassica rapa | M4EEI3 |
| Brassica rapa | M4FCE8 |
| Brassica rapa | M4ESI6 |
| Brassica rapa | M4EXD7 |
| Brassica rapa | M4CIU7 |
| Brassica rapa | M4EC53 |
| Brassica rapa | M4CFP8 |
| Brassica rapa | M4EAX3 |
| Brassica rapa | M4EDR6 |
| Brassica rapa | M4CB53 |
| Brassica rapa | M4EE74 |
| Brassica rapa | M4FIP3 |
| Brassica rapa | M4FHL1 |
| Brassica rapa | M4DK11 |
| Brassica rapa | M4CIU5 |
| Brassica rapa | M4ESI5 |
| Brassica rapa | M4EE73 |
| Brassica rapa | M4DTM4 |
| Brassica rapa | M4DAL2 |
| Brassica rapa | M4EUB0 |
| Brassica souli | Q8LP66 |
| Brassica tour | Q8LP59 |
| Brassica villos | Q8LP58 |
| Breoghania c | A0A2T5VC63 |
| Breoghania s | A0A346R2I9 |
| Breoghania s | A0A346R164 |
| Brucellaceae | A0A3P1UMN2 |
| Brugia malayi | A0A0H5SIF6 |
| Brugia malayi | A0A0H5S642 |
| Brugia malayi | A0A0K0J6N5 |
| Brugia malayi | A0A0J9XW65 |
| Brugia malayi | A0A0K0J6N4 |
| Brugia malayi | A0A0H5S3W5 |
| Brugia malayi | A0A0K0JDL5 |
| Brugia malayi | A0A0H5S9G2 |
| Brugia malayi | A0A0H5S1D5 |
| Brugia malayi | A0A0H5SAY8 |
| Brugia malayi | A0A0H5SR05 |
| Brugia malayi | A0A1I9G1H5 |

| | |
|---|---|
| Brugia malayi | A0A158PX07 |
| Brugia malayi | A0A158PZU7 |
| Brugia malayi | A0A158PY61 |
| Brugia malayi | A0A0J9Y7A3 |
| Brugia malayi | A0A1I9G322 |
| Brugia malayi | A0A158PYI4 |
| Brugia malayi | A0A1I9G2Y4 |
| Brugia malayi | A0A0K0JDU5 |
| Brugia malayi | A0A0I9R304 |
| Brugia malayi | A0A0K0J681 |
| Brugia malayi | A0A0K0J644 |
| Brugia malayi | A0A158PX08 |
| Brugia malayi | A0A0K0JDU6 |
| Brugia malayi | A0A1I9GCN9 |
| Brugia malayi | A0A158PY62 |
| Brugia malayi | A0A0K0JU89 |
| Brugia malayi | A0A0K0IMD7 |
| Brugia malayi | A0A0J9XZT0 |
| Brugia malayi | A0A0J9Y9P8 |
| Brugia malayi | A0A0K0IQ14 |
| Brugia malayi | A0A0K0ISE1 |
| Brugia malayi | A0A0K0JJL0 |
| Brugia malayi | A0A0J9Y0R4 |
| Brugia malayi | A0A0J9YCN5 |
| Brugia pahan | A0A0N4TNB0 |
| Brugia pahan | A0A158PRG8 |
| Brugia pahan | A0A0N4TMV2 |
| Brugia pahan | A0A0N4TSS0 |
| Brugia pahan | A0A158PSR5 |
| Brugia pahan | A0A158PQG8 |
| Brugia pahan | A0A0N4SZC0 |
| Brugia pahan | A0A0N4T3W4 |
| Brugia pahan | A0A0N4TFB9 |
| Brugia pahan | A0A0N4TYT6 |
| Brugia pahan | A0A0N4T2A3 |
| Brugia pahan | A0A0N4TLH9 |
| Brugia pahan | A0A0N4SYT2 |
| Brugia pahan | A0A0N4TPM2 |
| Brugia pahan | A0A0N4TR02 |
| Brugia pahan | A0A0N4SWY6 |
| Brugia pahan | A0A0N4TNG8 |

| | |
|---|---|
| Brugia pahan | A0A0N4TVC8 |
| Brugia pahan | A0A0N4SZ35 |
| Brugia timori | A0A3P7TPN6 |
| Brugia timori | A0A158PT29 |
| Brugia timori | A0A0R3Q302 |
| Brugia timori | A0A3P7SG12 |
| Brugia timori | A0A0R3QBN5 |
| Brugia timori | A0A0R3QQ91 |
| Brugia timori | A0A0R3QX15 |
| Brugia timori | A0A0R3R813 |
| Brugia timori | A0A0R3QCX7 |
| Brugia timori | A0A0R3R1C6 |
| Brugia timori | A0A0R3R9R2 |
| Brugia timori | A0A0R3R309 |
| Brugia timori | A0A0R3QNJ6 |
| Brugia timori | A0A0R3QRA9 |
| Brugia timori | A0A0R3Q4B1 |
| Brugia timori | A0A3P7T0Q5 |
| Buceros rhinc | A0A091GKM8 |
| Buceros rhinc | A0A091I300 |
| Buceros rhinc | A0A091HAC8 |
| Buceros rhinc | A0A091HG60 |
| Bursaphelenc | A0A1I7S3Z5 |
| Bursaphelenc | A0A1I7RNK5 |
| Bursaphelenc | A0A1I7RQH8 |
| Bursaphelenc | A0A1I7SVU8 |
| Bursaphelenc | A0A1I7RV17 |
| Bursaphelenc | A0A1I7S9K1 |
| Bursaphelenc | A0A1I7SLP2 |
| Bursaphelenc | A0A1I7RIJ0 |
| Bursaphelenc | A0A1I7SAF2 |
| Bursaphelenc | A0A1I7RLM5 |
| Bursaphelenc | A0A1I7SQH5 |
| Bursaphelenc | A0A1I7S7L5 |
| Bursaphelenc | A0A1I7SX51 |
| Bursaphelenc | A0A1I7RIX3 |
| Bursaphelenc | A0A1I7S4P8 |
| Bursaphelenc | A0A1I7SLS0 |
| Bursaphelenc | A0A1I7SB83 |
| Bursaphelenc | A0A1I7SDI4 |
| Bursaphelenc | A0A1I7SER8 |

| | |
|---|---|
| Bursaphelenc | A0A1I7SST9 |
| Bursaphelenc | A0A1I7RHA2 |
| Bursaphelenc | A0A1I7RN79 |
| Bursaphelenc | A0A1I7RHH1 |
| Cadophora sp | A0A2V1BVR2 |
| Cadophora sp | A0A2V1BNQ3 |
| Cadophora sp | A0A2V1B6F2 |
| Cadophora sp | A0A2V1BD27 |
| Cadophora sp | A0A2V1BP60 |
| Cadophora sp | A0A2V1C858 |
| Cadophora sp | A0A2V1BJ06 |
| Cadophora sp | A0A2V1B5B9 |
| Cadophora sp | A0A2V1BWQ8 |
| Cadophora sp | A0A2V1CUT5 |
| Cadophora sp | A0A2V1CF77 |
| Cadophora sp | A0A2V1BSL5 |
| Caenorhabdit | B6VC06 |
| Caenorhabdit | G0NPZ3 |
| Caenorhabdit | G0MVC6 |
| Caenorhabdit | G0N8P6 |
| Caenorhabdit | G0M8F1 |
| Caenorhabdit | G0PGC1 |
| Caenorhabdit | G0M6R7 |
| Caenorhabdit | G0NE96 |
| Caenorhabdit | G0NN92 |
| Caenorhabdit | G0N7Q3 |
| Caenorhabdit | G0MCH5 |
| Caenorhabdit | G0MRH4 |
| Caenorhabdit | G0MHL2 |
| Caenorhabdit | G0PDT9 |
| Caenorhabdit | G0MVD5 |
| Caenorhabdit | G0M8F2 |
| Caenorhabdit | G0NLF9 |
| Caenorhabdit | G0P5N7 |
| Caenorhabdit | G0NU28 |
| Caenorhabdit | G0NII5 |
| Caenorhabdit | G0PEQ2 |
| Caenorhabdit | G0NE92 |
| Caenorhabdit | G0MHP5 |
| Caenorhabdit | G0NGT2 |
| Caenorhabdit | G0P5E1 |

| | |
|---|---|
| Caenorhabdit | B0K0A0 |
| Caenorhabdit | A8XM04 |
| Caenorhabdit | A8XU34 |
| Caenorhabdit | A8X9M4 |
| Caenorhabdit | E3CU51 |
| Caenorhabdit | A8WN02 |
| Caenorhabdit | A8XZL8 |
| Caenorhabdit | A8WT52 |
| Caenorhabdit | A8Y2W2 |
| Caenorhabdit | A8X5P5 |
| Caenorhabdit | A8XZR6 |
| Caenorhabdit | A8WPA1 |
| Caenorhabdit | A8XDB5 |
| Caenorhabdit | A8XK91 |
| Caenorhabdit | A8Y0T3 |
| Caenorhabdit | A8X0Q4 |
| Caenorhabdit | A8XQU5 |
| Caenorhabdit | A8WVZ7 |
| Caenorhabdit | A8WRC5 |
| Caenorhabdit | A8X0Q5 |
| Caenorhabdit | A8WXE5 |
| Caenorhabdit | A8Y4J3 |
| Caenorhabdit | Q27394 |
| Caenorhabdit | Q10125 |
| Caenorhabdit | Q09271 |
| Caenorhabdit | Q10952 |
| Caenorhabdit | Q17800 |
| Caenorhabdit | H9G2Z5 |
| Caenorhabdit | H2KZ73 |
| Caenorhabdit | B6EU70 |
| Caenorhabdit | A0A2C9C2X5 |
| Caenorhabdit | A0A2C9C344 |
| Caenorhabdit | Q7YX17 |
| Caenorhabdit | Q22631 |
| Caenorhabdit | C1P662 |
| Caenorhabdit | D0VWP5 |
| Caenorhabdit | Q21782 |
| Caenorhabdit | Q22214 |
| Caenorhabdit | A0A2C9C2R3 |
| Caenorhabdit | O61834 |
| Caenorhabdit | Q9XUE5 |

| | |
|---|---|
| Caenorhabdit | Q22815 |
| Caenorhabdit | Q94248 |
| Caenorhabdit | O17347 |
| Caenorhabdit | Q23327 |
| Caenorhabdit | O62201 |
| Caenorhabdit | H8W3Y1 |
| Caenorhabdit | Q18298 |
| Caenorhabdit | G5EBR9 |
| Caenorhabdit | G5ECB3 |
| Caenorhabdit | G5EDB1 |
| Caenorhabdit | Q20167 |
| Caenorhabdit | Q23328 |
| Caenorhabdit | Q22902 |
| Caenorhabdit | O45277 |
| Caenorhabdit | G5ECB4 |
| Caenorhabdit | Q8T3B7 |
| Caenorhabdit | K7GW62 |
| Caenorhabdit | K7HND9 |
| Caenorhabdit | K7GW65 |
| Caenorhabdit | A0A2H2ICP9 |
| Caenorhabdit | A0A2H2I0G9 |
| Caenorhabdit | K7GW61 |
| Caenorhabdit | K7GW66 |
| Caenorhabdit | K7H567 |
| Caenorhabdit | A0A2H2HYS4 |
| Caenorhabdit | K7H566 |
| Caenorhabdit | K7H1H4 |
| Caenorhabdit | A0A2H2I0E1 |
| Caenorhabdit | A0A2H2J0L9 |
| Caenorhabdit | K7GW60 |
| Caenorhabdit | H2WB38 |
| Caenorhabdit | A0A2H2IE65 |
| Caenorhabdit | A0A2H2IWD6 |
| Caenorhabdit | A0A2H2IAG5 |
| Caenorhabdit | A0A2H2IAK0 |
| Caenorhabdit | A0A2H2I6U0 |
| Caenorhabdit | A0A2H2HUZ7 |
| Caenorhabdit | K7GTV4 |
| Caenorhabdit | K7HVV9 |
| Caenorhabdit | A0A2H2I7Z8 |
| Caenorhabdit | K7H652 |

| | |
|---|---|
| Caenorhabdit | K7H407 |
| Caenorhabdit | A0A2H2JF03 |
| Caenorhabdit | A0A2H2HU97 |
| Caenorhabdit | A0A2H2IJJ4 |
| Caenorhabdit | K7H1H3 |
| Caenorhabdit | K7H408 |
| Caenorhabdit | K7H6J2 |
| Caenorhabdit | K7HAQ8 |
| Caenorhabdit | A0A2H2I481 |
| Caenorhabdit | K7GTV5 |
| Caenorhabdit | K7H6I4 |
| Caenorhabdit | K7HAQ7 |
| Caenorhabdit | K7HND8 |
| Caenorhabdit | A0A2H2I4H8 |
| Caenorhabdit | K7H6J3 |
| Caenorhabdit | K7HVW0 |
| Caenorhabdit | K7H6I3 |
| Caenorhabdit | A0A261ACA4 |
| Caenorhabdit | A0A261AH64 |
| Caenorhabdit | A0A261CPY0 |
| Caenorhabdit | A0A261CSP6 |
| Caenorhabdit | A0A261AU21 |
| Caenorhabdit | A0A261CNX8 |
| Caenorhabdit | A0A261CR66 |
| Caenorhabdit | A0A261BZ73 |
| Caenorhabdit | A0A261CE90 |
| Caenorhabdit | A0A260ZQN6 |
| Caenorhabdit | A0A261CNC8 |
| Caenorhabdit | A0A261CA21 |
| Caenorhabdit | A0A261CK25 |
| Caenorhabdit | A0A261CGY1 |
| Caenorhabdit | A0A261CP70 |
| Caenorhabdit | A0A261CFW0 |
| Caenorhabdit | A0A261BUT5 |
| Caenorhabdit | A0A261B5J5 |
| Caenorhabdit | A0A261ANV5 |
| Caenorhabdit | A0A260ZQ86 |
| Caenorhabdit | A0A261APP7 |
| Caenorhabdit | A0A261BID9 |
| Caenorhabdit | A0A260ZF97 |
| Caenorhabdit | A0A260ZPE3 |

| | |
|---|---|
| Caenorhabdit | A0A2G5T5T2 |
| Caenorhabdit | A0A2G5SYL6 |
| Caenorhabdit | A0A2G5U9C7 |
| Caenorhabdit | A0A2G5U9K5 |
| Caenorhabdit | A0A2G5ULQ0 |
| Caenorhabdit | A0A2G5SZG6 |
| Caenorhabdit | A0A2G5SPU6 |
| Caenorhabdit | A0A2G5UM21 |
| Caenorhabdit | A0A2G5U9C6 |
| Caenorhabdit | A0A2G5V3A7 |
| Caenorhabdit | A0A2G5U4H9 |
| Caenorhabdit | A0A2G5U0G8 |
| Caenorhabdit | A0A2G5T662 |
| Caenorhabdit | A0A2G5SPS0 |
| Caenorhabdit | A0A2G5SIZ1 |
| Caenorhabdit | A0A2G5V2K3 |
| Caenorhabdit | A0A2G5SQ73 |
| Caenorhabdit | A0A2G5U9E1 |
| Caenorhabdit | A0A2G5TVG4 |
| Caenorhabdit | A0A2G5U4R2 |
| Caenorhabdit | A0A2G5VRD3 |
| Caenorhabdit | A0A2G5U9B1 |
| Caenorhabdit | A0A2G5UKX7 |
| Caenorhabdit | A0A2G5TNF2 |
| Caenorhabdit | A0A2G5TY68 |
| Caenorhabdit | A0A2G5UG40 |
| Caenorhabdit | A0A2G5U9E5 |
| Caenorhabdit | A0A2G5U784 |
| Caenorhabdit | A0A2G5V2K1 |
| Caenorhabdit | A0A2G5UH11 |
| Caenorhabdit | A0A2G5SPN4 |
| Caenorhabdit | A0A2G5STL2 |
| Caenorhabdit | A0A2G5UV42 |
| Caenorhabdit | A0A2G5SYP7 |
| Caenorhabdit | A0A2G5THD7 |
| Caenorhabdit | A0A2G5TVG8 |
| Caenorhabdit | A0A2G5SPC7 |
| Caenorhabdit | A0A2G5TNG8 |
| Caenorhabdit | A0A2G5VF01 |
| Caenorhabdit | A0A2G5U9U1 |
| Caenorhabdit | A0A2G5U799 |

| | |
|---|---|
| Caenorhabdit | A0A2G5STT6 |
| Caenorhabdit | A0A2G5U4J0 |
| Caenorhabdit | A0A2G5VR02 |
| Caenorhabdit | A0A2G5VEG5 |
| Caenorhabdit | A0A2G5UKS0 |
| Caenorhabdit | A0A2G5STB6 |
| Caenorhabdit | A0A2G5SNI9 |
| Caenorhabdit | A0A2G5U534 |
| Caenorhabdit | A0A2G5U9C2 |
| Caenorhabdit | A0A2G5TI38 |
| Caenorhabdit | A0A2G5U9A0 |
| Caenorhabdit | A0A2G5TNA2 |
| Caenorhabdit | A0A2G5U4J4 |
| Caenorhabdit | A0A2G5TYA1 |
| Caenorhabdit | A0A2G5U787 |
| Caenorhabdit | E3N801 |
| Caenorhabdit | A0A2P4WK92 |
| Caenorhabdit | A0A2P4W3J2 |
| Caenorhabdit | E3M888 |
| Caenorhabdit | E3NWP9 |
| Caenorhabdit | E3M6Q4 |
| Caenorhabdit | E3M5X6 |
| Caenorhabdit | A0A2P4W7N0 |
| Caenorhabdit | A0A261BAQ3 |
| Caenorhabdit | A0A2P4WCC6 |
| Caenorhabdit | A0A261AJI9 |
| Caenorhabdit | A0A260ZJI1 |
| Caenorhabdit | A0A2P4V2Y6 |
| Caenorhabdit | E3M244 |
| Caenorhabdit | A0A2P4WFX4 |
| Caenorhabdit | A0A261AK10 |
| Caenorhabdit | E3NPV1 |
| Caenorhabdit | E3NQB6 |
| Caenorhabdit | E3M5X4 |
| Caenorhabdit | A0A261ATB5 |
| Caenorhabdit | A0A261AI47 |
| Caenorhabdit | A0A2P4VKF5 |
| Caenorhabdit | A0A260Z8D5 |
| Caenorhabdit | A0A261ALS6 |
| Caenorhabdit | A0A2P4WL44 |
| Caenorhabdit | E3M2F3 |

| | |
|---|---|
| Caenorhabdit | A0A261B0J8 |
| Caenorhabdit | E3LE81 |
| Caenorhabdit | E3MSS4 |
| Caenorhabdit | A0A2P4V2Z2 |
| Caenorhabdit | A0A2P4W7H8 |
| Caenorhabdit | A0A261BE63 |
| Caenorhabdit | A0A2P4VDA9 |
| Caenorhabdit | A0A2P4VDA1 |
| Caenorhabdit | A0A261AIL2 |
| Caenorhabdit | A0A2P4WLP1 |
| Caenorhabdit | A0A2P4W0U2 |
| Caenorhabdit | A0A2P4VIC1 |
| Caenorhabdit | E3N395 |
| Caenorhabdit | A0A2P4WBE7 |
| Caenorhabdit | E3MUD1 |
| Caenorhabdit | A0A261BE25 |
| Caenorhabdit | E3MQF4 |
| Caenorhabdit | A0A261A2K8 |
| Caenorhabdit | E3LY63 |
| Caenorhabdit | A0A2P4V2Y4 |
| Caenorhabdit | A0A2P4W7D8 |
| Caenorhabdit | E3NF96 |
| Caenorhabdit | E3MSS5 |
| Caenorhabdit | E3NAC3 |
| Caenorhabdit | A0A2P4WMV8 |
| Caenorhabdit | E3NS03 |
| Caenorhabdit | A0A2P4WTN2 |
| Caenorhabdit | A0A260ZUR0 |
| Caenorhabdit | A0A261B7H7 |
| Caenorhabdit | A0A261ASY1 |
| Caenorhabdit | E3LFD4 |
| Caenorhabdit | A0A261BCB7 |
| Caenorhabdit | E3LE82 |
| Caenorhabdit | E3MA09 |
| Caenorhabdit | A0A2P4VZW1 |
| Caenorhabdit | E3MK54 |
| Caenorhabdit | E3NE41 |
| Caenorhabdit | A0A261A2U8 |
| Caenorhabdit | A0A261AY11 |
| Caenorhabdit | E3ND68 |
| Caenorhabdit | A0A261A7H9 |

| | |
|---|---|
| Caenorhabdit | A0A2P4WIH5 |
| Caenorhabdit | A0A261AUN6 |
| Caenorhabdit | A0A260Z4Y7 |
| Caenorhabdit | A0A261ADY1 |
| Caenorhabdit | E3M221 |
| Caenorhabdit | E3LFC5 |
| Caenorhabdit | E3LZN5 |
| Caenorhabdit | A0A1I7TRC8 |
| Caenorhabdit | A0A1I7UFP5 |
| Caenorhabdit | A0A1I7TBP9 |
| Caenorhabdit | A0A1I7TB17 |
| Caenorhabdit | A0A1I7U139 |
| Caenorhabdit | A0A1I7UEG5 |
| Caenorhabdit | A0A1I7U2M5 |
| Caenorhabdit | A0A1I7U140 |
| Caenorhabdit | A0A1I7U2M6 |
| Caenorhabdit | A0A1I7T3H6 |
| Caenorhabdit | A0A1I7UDC8 |
| Caenorhabdit | A0A1I7V193 |
| Caenorhabdit | A0A1I7T400 |
| Caenorhabdit | A0A1I7U9T6 |
| Caenorhabdit | A0A1I7UTC6 |
| Caenorhabdit | A0A1I7TB16 |
| Caenorhabdit | A0A1I7V489 |
| Caenorhabdit | A0A1I7U190 |
| Caenorhabdit | A0A1I7TL14 |
| Caenorhabdit | A0A1I7U7E4 |
| Caenorhabdit | A0A1I7ULB1 |
| Caenorhabdit | A0A1I7U7E3 |
| Caenorhabdit | A0A1I7V194 |
| Caenorhabdit | A0A1I7T081 |
| Caenorhabdit | A0A1I7T3Z9 |
| Caenorhabdit | A0A1I7U7E6 |
| Caenorhabdit | A0A1I7U9T5 |
| Caenorhabdit | A0A1I7UTC7 |
| Caenorhabdit | A0A1I7UEG4 |
| Caenorhabdit | A0A1I7TBP8 |
| Cajanus cajar | A0A151SMC5 |
| Cajanus cajar | A0A151SPY9 |
| Cajanus cajar | A0A151UBP4 |
| Cajanus cajar | A0A151SVK7 |

Cajanus cajar A0A151S8Z9
Cajanus cajar A0A151UG46
Cajanus cajar A0A151TPQ2
Cajanus cajar A0A151S200
Cajanus cajar A0A151TU08
Cajanus cajar A0A151RB11
Cajanus cajar A0A151RFX6
Cajanus cajar A0A151TV23
Cajanus cajar A0A151SVI8
Cajanus cajar A0A151QZL4
Cajanus cajar A0A151R1S2
Cajanus cajar A0A151RF37
Cajanus cajar A0A151RBF8
Cajanus cajar A0A151R092
Cajanus cajar A0A151SY98
Cajanus cajar A0A151T3Q5
Cajanus cajar A0A151RFX8
Cajanus cajar A0A151S5X0
Cajanus cajar A0A151RKY9
Cajanus cajar A0A151SMF7
Cajanus cajar A0A151SVJ6
Cajanus cajar A0A151S231
Cajanus cajar A0A151SMD4
Cajanus cajar A0A151S219
Cajanus cajar A0A151S5A6
Cajanus cajar A0A151RBK0
Cajanus cajar A0A151RQ08
Cajanus cajar A0A151S2F9
Cajanus cajar A0A151QYZ7
Cajanus cajar A0A151RPY9
Cajanus cajar A0A151TBW7
Cajanus cajar A0A151SJN1
Cajanus cajar A0A151UDQ9
Cajanus cajar A0A151RDZ2
Cajanus cajar A0A151SH91
Cajanus cajar A0A151S126
Cajanus cajar A0A151RPW2
Cajanus cajar A0A151SMF2
Cajanus cajar A0A151S5R9
Cajanus cajar A0A151RKS3
Cajanus cajar A0A151T1R0

| | |
|---|---|
| Cajanus cajar | A0A151R0Z5 |
| Cajanus cajar | A0A151RPX4 |
| Cajanus cajar | A0A151RGD5 |
| Cajanus cajar | A0A151RFV9 |
| Cajanus cajar | A0A151RP79 |
| Cajanus cajar | A0A151RFY6 |
| Cajanus cajar | A0A151SJP5 |
| Cajanus cajar | A0A151RPZ1 |
| Cajanus cajar | A0A151S982 |
| Cajanus cajar | A0A151RXG6 |
| Cajanus cajar | A0A151RNK8 |
| Cajanus cajar | A0A151SM99 |
| Cajanus cajar | A0A151QZP1 |
| Cajanus cajar | A0A151RP46 |
| Cajanus cajar | A0A151RQ07 |
| Cajanus cajar | A0A151RGA5 |
| Cajanus cajar | A0A151UH94 |
| Cajanus cajar | A0A151T1Q6 |
| Cajanus cajar | A0A151SVE9 |
| Cajanus cajar | A0A151RPX7 |
| Cajanus cajar | A0A151S658 |
| Cajanus cajar | A0A151RL70 |
| Cajanus cajar | A0A151RE53 |
| Cajanus cajar | A0A151RL12 |
| Cajanus cajar | A0A151RQF6 |
| Cajanus cajar | A0A151TTP7 |
| Cajanus cajar | A0A151RBC2 |
| Cajanus cajar | A0A151RKT0 |
| Cajanus cajar | A0A151SWX4 |
| Callipepla squ | A0A226MM08 |
| Callipepla squ | A0A226MNE8 |
| Callipepla squ | A0A226MN47 |
| Callithrix jacc | F6WH76 |
| Callithrix jacc | F6S5L7 |
| Callithrix jacc | F7EDL5 |
| Callithrix jacc | F6ZZG6 |
| Callithrix jacc | F7I8A8 |
| Callithrix jacc | F6VX63 |
| Callithrix jacc | F7AA32 |
| Callithrix jacc | F6X343 |
| Callithrix jacc | F7GKJ7 |

| | |
|---|---|
| Callorhinchus | V9K997 |
| Callorhinchus | V9KGH2 |
| Callorhinchus | V9KIE5 |
| Callorhinus u | A0A3Q7P3N0 |
| Callorhinus u | A0A3Q7QI46 |
| Callorhinus u | A0A3Q7NAT6 |
| Callorhinus u | A0A3Q7RMW6 |
| Callorhinus u | A0A3Q7Q6J8 |
| Callorhinus u | A0A3Q7NFH9 |
| Callorhinus u | A0A3Q7Q880 |
| Callorhinus u | A0A3Q7QI91 |
| Callorhinus u | A0A3Q7PEL9 |
| Calypte anna | A0A091HYP2 |
| Calypte anna | A0A091HU29 |
| Camelus dror | A0A1C9EFM3 |
| Camelus feru | T0MC86 |
| Camelus feru | S9YNR1 |
| Camponotus | E2AYS0 |
| Camponotus | E2AW54 |
| Camponotus | E2AIC2 |
| Camponotus | E1ZYY0 |
| Camponotus | E2A6H3 |
| Camponotus | E2AEN9 |
| candidate div | A0A2G6E0Y9 |
| candidate div | A0A0G0Y802 |
| Candidatus D | A0A0T5ZUG2 |
| Candidatus Fi | A0A1F5VTJ5 |
| Candidatus La | A0A1F6GNK9 |
| Candidatus Le | A0A1F6IW58 |
| Candidatus Le | A0A1F6IW51 |
| Candidatus M | A0A2E6AVH4 |
| Candidatus M | A0A2E1QA46 |
| Candidatus M | A0A2E8VEV6 |
| Candidatus M | A0A081BMC8 |
| Candidatus N | A0A3E1D9A6 |
| Candidatus P | A0A2H0KSE9 |
| Candidatus R | A0A0J9E870 |
| Candidatus Tl | A0A2K8U7S7 |
| Candidatus V | A0A0S6WC51 |
| Canis lupus fa | Q867B7 |
| Canis lupus fa | F1PNV5 |

| | |
|---|---|
| Canis lupus fa | F1Q421 |
| Canis lupus fa | J9P7N9 |
| Canis lupus fa | U6C2S1 |
| Canis lupus fa | B5LX43 |
| Canis lupus fa | F1PPK8 |
| Canis lupus fa | E2QX19 |
| Capitella tele | X1ZK48 |
| Capitella tele | X2A7L8 |
| Capitella tele | R7T8C6 |
| Capitella tele | R7V393 |
| Capitella tele | R7U1A9 |
| Capitella tele | R7TZY2 |
| Capitella tele | R7V723 |
| Capitella tele | R7UC69 |
| Capitella tele | R7V5X1 |
| Capitella tele | R7TYC5 |
| Capitella tele | R7UCC3 |
| Capitella tele | R7TUV0 |
| Capitella tele | R7U099 |
| Capitella tele | R7UD86 |
| Capitella tele | R7VAV0 |
| Capitella tele | R7U4R0 |
| Capitella tele | R7V1P5 |
| Capitella tele | R7U5C7 |
| Capitella tele | R7VIS4 |
| Capitella tele | R7UZN9 |
| Capitella tele | R7UPR5 |
| Capitella tele | R7TN37 |
| Capitella tele | R7TTF7 |
| Capitella tele | N1PB13 |
| Capitella tele | R7T7C0 |
| Capitella tele | R7TW97 |
| Capitella tele | R7U2J7 |
| Capitella tele | R7V1H6 |
| Capitella tele | R7TQC9 |
| Capitella tele | R7VEV6 |
| Capitella tele | R7UHQ9 |
| Capitella tele | R7TNK1 |
| Capitella tele | R7U9W1 |
| Capitella tele | R7UFI6 |
| Capitella tele | R7TTM3 |

Capitella tele R7TL37
Capitella tele R7VDK3
Capitella tele N1PB94
Capitella tele R7UZW7
Capitella tele R7UR22
Capitella tele R7TMW2
Capitella tele R7TC17
Capitella tele R7U7K0
Capitella tele R7T380
Capitella tele R7TB71
Capitella tele R7TTG1
Capitella tele R7TV63
Capitella tele R7TDJ4
Capitella tele R7UDB7
Capitella tele R7TUJ8
Capitella tele R7TA27
Capitella tele R7T436
Capitella tele R7USY4
Capitella tele R7TG23
Capitella tele R7THU4
Capitella tele R7V2I5
Capitella tele R7TW72
Capitella tele R7U6P1
Capitella tele R7V1M6
Capitella tele R7U4L4
Capitella tele R7UEY2
Capitella tele R7TFT8
Capitella tele R7VJV9
Capitella tele R7TX09
Capitella tele R7UIB6
Capitella tele R7U426
Capitella tele R7VEG9
Capitella tele R7TMY6
Capitella tele R7ULX6
Capitella tele R7U7Q9
Capitella tele R7UCW6
Capitella tele R7UUK4
Capitella tele R7U544
Capitella tele R7UUF8
Capitella tele R7TA51
Capitella tele R7UFR7

| | |
|---|---|
| Capitella tele | R7T4S3 |
| Capitella tele | R7VFQ5 |
| Capitella tele | R7UDP1 |
| Capitella tele | R7UH53 |
| Capitella tele | R7TBV9 |
| Capitella tele | R7VFH4 |
| Capitella tele | R7V8L1 |
| Capitella tele | R7UGV2 |
| Capitella tele | R7VGL6 |
| Capitella tele | R7TAP8 |
| Capitella tele | R7TYU2 |
| Capitella tele | R7V4I7 |
| Capitella tele | R7TIG7 |
| Capitella tele | R7V0P1 |
| Capitella tele | R7T7U1 |
| Capitella tele | R7V6R4 |
| Capitella tele | R7U6D2 |
| Capitella tele | R7V5V9 |
| Capitella tele | R7TI76 |
| Capitella tele | R7VF14 |
| Capitella tele | R7UVT0 |
| Capitella tele | R7VG70 |
| Capra hircus.: | A0A452E0U1 |
| Capra hircus.: | A0A452E5W4 |
| Capra hircus.: | A0A452EN11 |
| Capra hircus.: | A0A452E0T2 |
| Capra hircus.: | A0A452E0U8 |
| Capra hircus.: | A0A452FN28 |
| Capra hircus.: | A0A452E7A0 |
| Capra hircus.: | A0A452E0H7 |
| Capronia cor | W9YD76 |
| Capronia epi | W9YRK9 |
| Capsella gran | Q15B70 |
| Capsella gran | Q15B72 |
| Capsella gran | Q15B73 |
| Capsella gran | B9VWJ3 |
| Capsella gran | C3USC5 |
| Capsella gran | C3USB9 |
| Capsella gran | C3USC3 |
| Capsella gran | B9VWJ2 |
| Capsella gran | A0A330PLR6 |

| | |
|---|---|
| Capsella gran | A0A330PLQ4 |
| Capsella gran | Q159E8 |
| Capsella gran | D6PR15 |
| Capsella gran | C3US88 |
| Capsella gran | Q159E6 |
| Capsella gran | D6PR17 |
| Capsella gran | Q159E2 |
| Capsella gran | B9VWJ4 |
| Capsella gran | D6PR14 |
| Capsella gran | C3US92 |
| Capsella gran | D6PR16 |
| Capsella gran | Q159E1 |
| Capsella gran | Q15B71 |
| Capsella gran | C3US83 |
| Capsella gran | A0A330PI91 |
| Capsella gran | D6PR65 |
| Capsella gran | C3USB8 |
| Capsella gran | A5A5D8 |
| Capsella gran | A0A330PL92 |
| Capsella gran | Q15B68 |
| Capsella gran | C3USC0 |
| Capsella gran | C3US90 |
| Capsella gran | C3USC9 |
| Capsella gran | C3USD0 |
| Capsella gran | C3US91 |
| Capsella gran | C3USC1 |
| Capsella gran | A0A330PL82 |
| Capsella gran | Q159E5 |
| Capsella gran | D6PR13 |
| Capsella gran | C3USC8 |
| Capsella gran | Q15B69 |
| Capsella gran | Q159E7 |
| Capsella gran | B9VWJ5 |
| Capsella gran | A0A330PKY6 |
| Capsella gran | C3USC6 |
| Capsella gran | Q159E4 |
| Capsella gran | C3USC4 |
| Capsella gran | C3US86 |
| Capsella gran | C3USC2 |
| Capsella gran | C3US84 |
| Capsella gran | Q159E3 |

| | |
|---|---|
| Capsella gran | C3US87 |
| Capsella gran | C3US82 |
| Capsella gran | D6PR63 |
| Capsella gran | D6PR62 |
| Capsella gran | C3US81 |
| Capsella gran | C3USC7 |
| Capsella rube | R0I6W8 |
| Capsella rube | R0GTT4 |
| Capsella rube | R0H9B0 |
| Capsella rube | R0GNE2 |
| Capsella rube | R0FI89 |
| Capsella rube | R0HTN8 |
| Capsella rube | R0GUQ7 |
| Capsella rube | R0HUY1 |
| Capsella rube | R0H0H3 |
| Capsella rube | R0I9M8 |
| Capsella rube | G4WHB0 |
| Capsella rube | R0IR13 |
| Capsella rube | R0GLW7 |
| Capsella rube | R0GG20 |
| Capsella rube | R0HTH2 |
| Capsella rube | R0HWX1 |
| Capsella rube | R0HH21 |
| Capsella rube | R0F138 |
| Capsella rube | R0G349 |
| Capsella rube | R0G9X0 |
| Capsella rube | R0H4Q7 |
| Capsella rube | G4WHA9 |
| Capsella rube | R0H3X2 |
| Capsella rube | R0IQY0 |
| Capsella rube | R0HDW3 |
| Capsella rube | R0GEJ4 |
| Capsella rube | R0I6F3 |
| Capsella rube | Q159E9 |
| Capsella rube | R0HEM0 |
| Capsella rube | R0GB39 |
| Capsella rube | R0EV33 |
| Capsella rube | R0IB39 |
| Capsella rube | R0GFC4 |
| Capsella rube | R0GDL4 |
| Capsella rube | R0H6A7 |

| | |
|---|---|
| Capsella rube | R0G917 |
| Capsella rube | R0GC32 |
| Capsella rube | R0FRL5 |
| Capsella rube | R0GGH9 |
| Capsella rube | R0GPB0 |
| Capsella rube | R0IKR4 |
| Capsella rube | R0I6F0 |
| Capsella rube | G3LPX2 |
| Capsella rube | C3US76 |
| Capsella rube | C3US93 |
| Capsella rube | G3LPQ9 |
| Capsella rube | Q159F0 |
| Capsicum anr | A0A2G2Z3C2 |
| Capsicum anr | A0A1U8HJK3 |
| Capsicum anr | A0A1U8FLD1 |
| Capsicum anr | A0A1U8G176 |
| Capsicum anr | A0A1U8FMT3 |
| Capsicum anr | A0A1U8G0B3 |
| Capsicum anr | A0A2G2YXS1 |
| Capsicum anr | A0A2G2Z497 |
| Capsicum anr | A0A2G2ZR65 |
| Capsicum anr | A0A2G3A8T5 |
| Capsicum anr | A0A1U8E006 |
| Capsicum anr | A0A1U8GC95 |
| Capsicum anr | A0A1U8FKZ6 |
| Capsicum anr | A0A1U8EHH7 |
| Capsicum anr | A0A1U8FWF1 |
| Capsicum anr | A0A2G2YVW6 |
| Capsicum anr | A0A1U8HID5 |
| Capsicum anr | A0A1U8HB94 |
| Capsicum anr | A0A2G2XXB9 |
| Capsicum anr | A0A1U8FUE7 |
| Capsicum anr | A0A1U8FH35 |
| Capsicum anr | A0A2G2YHC9 |
| Capsicum anr | A0A2G3A4Y4 |
| Capsicum anr | A0A1U8DSV0 |
| Capsicum anr | A0A2G2ZR66 |
| Capsicum anr | A0A2G3ACA4 |
| Capsicum anr | A0A1U8GJN2 |
| Capsicum anr | A0A2G2Z443 |
| Capsicum anr | A0A1U8GC84 |

| | |
|---|---|
| Capsicum anr | A0A2G2Z3R4 |
| Capsicum anr | A0A1U8G424 |
| Capsicum anr | A0A2G3A4P0 |
| Capsicum anr | A0A2G2XZF1 |
| Capsicum anr | A0A1U8HJC3 |
| Capsicum anr | A0A2G2Z3M0 |
| Capsicum anr | A0A1U8E541 |
| Capsicum anr | A0A2G2Y426 |
| Capsicum anr | A0A2G2XXC9 |
| Capsicum anr | A0A2G2Z5E8 |
| Capsicum anr | A0A2G2YRI6 |
| Capsicum anr | A0A1U8FFY8 |
| Capsicum anr | A0A2G3A8R0 |
| Capsicum anr | A0A2G3A8S2 |
| Capsicum anr | A0A2G2Z7W9 |
| Capsicum anr | A0A1U8F3X5 |
| Capsicum anr | A0A1U8FMU1 |
| Capsicum anr | A0A2G2Z4E1 |
| Capsicum anr | A0A2G2ZGY9 |
| Capsicum anr | A0A1U8EBB4 |
| Capsicum anr | A0A1U8GJN8 |
| Capsicum anr | A0A1U8F5I0 |
| Capsicum anr | A0A2G2ZR70 |
| Capsicum anr | A0A1U8FLJ4 |
| Capsicum anr | A0A2G2Z477 |
| Capsicum anr | A0A1U8FUF3 |
| Capsicum anr | A0A1U8H8D5 |
| Capsicum anr | A0A2G2ZGX9 |
| Capsicum anr | A0A1U8HCZ6 |
| Capsicum anr | A0A1U8FGU3 |
| Capsicum anr | A0A2G3AHC7 |
| Capsicum anr | A0A2G2ZRA6 |
| Capsicum anr | A0A2G2YBW0 |
| Capsicum anr | A0A1U8HJK7 |
| Capsicum anr | A0A2G2ZR53 |
| Capsicum anr | A0A2G2ZR72 |
| Capsicum anr | A0A1U8HHR9 |
| Capsicum anr | A0A1U8EQX3 |
| Capsicum anr | A0A1U8G097 |
| Capsicum anr | A0A2G2ZGX6 |
| Capsicum anr | A0A1U8HFC1 |

| | |
|---|---|
| Capsicum anı | A0A2G2Z069 |
| Capsicum anı | A0A1U8HJC5 |
| Capsicum anı | A0A1U8HF58 |
| Capsicum anı | A0A2G3AHK5 |
| Capsicum anı | A0A1U8GC83 |
| Capsicum anı | A0A2G2Z4E6 |
| Capsicum anı | A0A2G2Z592 |
| Capsicum anı | A0A1U8FWF6 |
| Capsicum anı | A0A1U8FKF0 |
| Capsicum anı | A0A1U8HBY4 |
| Capsicum anı | A0A1U8GC91 |
| Capsicum anı | A0A1U8FI52 |
| Capsicum anı | A0A2G2ZR54 |
| Capsicum anı | A0A1U8GLG7 |
| Capsicum anı | A0A2G2Z4F9 |
| Capsicum anı | A0A2G2XZG6 |
| Capsicum anı | A0A2G2Z588 |
| Capsicum anı | A0A1U8GKK0 |
| Capsicum anı | A0A1U8FMT6 |
| Capsicum anı | A0A1U8E509 |
| Capsicum anı | A0A2G3A8W8 |
| Capsicum anı | A0A1U8GL39 |
| Capsicum anı | A0A1U8H760 |
| Capsicum anı | A0A1U8HFP2 |
| Capsicum anı | A0A2G2Z4D1 |
| Capsicum anı | A0A1U8HC90 |
| Capsicum anı | A0A1U8FCB4 |
| Capsicum anı | A0A2G2Z5A7 |
| Capsicum anı | A0A2G2XTN5 |
| Capsicum anı | A0A2G3A3F6 |
| Capsicum anı | A0A2G2XJB3 |
| Capsicum anı | A0A1U8HJL6 |
| Capsicum anı | A0A2G3A8Y5 |
| Capsicum anı | A0A1U8FU12 |
| Capsicum anı | A0A2G2ZNM4 |
| Capsicum anı | A0A2G2YUR9 |
| Capsicum anı | A0A1U8FUG6 |
| Capsicum anı | A0A1U8FLF2 |
| Capsicum anı | A0A2G2Y4I0 |
| Capsicum anı | A0A1U8F5U4 |
| Capsicum anı | A0A2G2Y412 |

| | |
|---|---|
| Capsicum anr | A0A1U8FUE2 |
| Capsicum anr | A0A1U8HHS4 |
| Capsicum anr | A0A1U8FMU9 |
| Capsicum anr | A0A2G2ZR57 |
| Capsicum anr | A0A1U8HD42 |
| Capsicum anr | A0A1U8FL04 |
| Capsicum anr | A0A1U8HD34 |
| Capsicum anr | A0A2G2ZR87 |
| Capsicum anr | A0A2G2ZRA9 |
| Capsicum anr | A0A1U8HHK2 |
| Capsicum anr | A0A2G2Z3A1 |
| Capsicum anr | A0A2G2Z488 |
| Capsicum anr | A0A2G2ZZ34 |
| Capsicum anr | A0A2G2YUF5 |
| Capsicum anr | A0A1U8HD39 |
| Capsicum anr | A0A1U8H0K4 |
| Capsicum anr | A0A1U8GC89 |
| Capsicum anr | A0A2G3A8R4 |
| Capsicum anr | A0A1U8FMF8 |
| Capsicum anr | A0A1U8GCX8 |
| Capsicum anr | A0A2G2YQD6 |
| Capsicum anr | A0A1U8EA38 |
| Capsicum anr | A0A2G2Z3H8 |
| Capsicum anr | A0A1U8FQA6 |
| Capsicum anr | A0A2G2ZR51 |
| Capsicum anr | A0A2G2Z5Q1 |
| Capsicum anr | A0A1U8HD36 |
| Capsicum anr | A0A1U8FUF9 |
| Capsicum anr | A0A2G3A8R3 |
| Capsicum anr | A0A1U8FLD6 |
| Capsicum anr | A0A2G2ZR86 |
| Capsicum anr | A0A1U8FLE0 |
| Capsicum anr | A0A2G3A8S0 |
| Capsicum anr | A0A1U8GCY4 |
| Capsicum anr | A0A2G2ZR97 |
| Capsicum anr | A0A2G3A8U5 |
| Capsicum anr | A0A2G2Z5B6 |
| Capsicum anr | A0A2G2XZG5 |
| Capsicum anr | A0A2G3A8V8 |
| Capsicum anr | A0A1U8FGL4 |
| Capsicum anr | A0A1U8HJB8 |

| | |
|---|---|
| Capsicum anı | A0A2G2ZZZ3 |
| Capsicum anı | A0A2G3A8A0 |
| Capsicum anı | A0A2G2YVF7 |
| Capsicum anı | A0A2G2ZR77 |
| Capsicum anı | A0A2G2Z5J5 |
| Capsicum anı | A0A2G2ZR62 |
| Capsicum anı | A0A2G2Z449 |
| Capsicum anı | A0A2G2Z595 |
| Capsicum anı | A0A1U8GJM6 |
| Capsicum anı | A0A2G2Z4K3 |
| Capsicum anı | A0A1U8FLD5 |
| Capsicum anı | A0A1U8FFC8 |
| Capsicum anı | A0A1U8FEC5 |
| Capsicum anı | A0A1U8GMK6 |
| Capsicum anı | A0A2G3AFV5 |
| Capsicum anı | A0A2G2Z5L0 |
| Capsicum anı | A0A2G2Z584 |
| Capsicum anı | A0A2G2ZY67 |
| Capsicum anı | A0A1U8G841 |
| Capsicum anı | A0A2G2Z100 |
| Capsicum anı | A0A2G3A8V4 |
| Capsicum anı | A0A2G2YUT1 |
| Capsicum anı | A0A1U8GGN2 |
| Capsicum anı | A0A2G3A8X5 |
| Capsicum anı | A0A1U8GK83 |
| Capsicum anı | A0A1U8E139 |
| Capsicum anı | A0A1U8HLA8 |
| Capsicum anı | A0A1U8EB93 |
| Capsicum anı | A0A2G2Z3E9 |
| Capsicum bac | A0A2G2WEE9 |
| Capsicum bac | A0A2G2WVY4 |
| Capsicum bac | A0A2G2WW96 |
| Capsicum bac | A0A2G2VY63 |
| Capsicum bac | A0A2G2VQY9 |
| Capsicum bac | A0A2G2VAD8 |
| Capsicum bac | A0A2G2WPR1 |
| Capsicum bac | A0A2G2WDF9 |
| Capsicum bac | A0A2G2WEK5 |
| Capsicum bac | A0A2G2VR35 |
| Capsicum bac | A0A2G2VQY2 |
| Capsicum bac | A0A2G2WEJ2 |

| | |
|---|---|
| Capsicum bac | A0A2G2WG78 |
| Capsicum bac | A0A2G2WZR9 |
| Capsicum bac | A0A2G2X0B4 |
| Capsicum bac | A0A2G2WGJ3 |
| Capsicum bac | A0A2G2VBZ9 |
| Capsicum bac | A0A2G2XNZ0 |
| Capsicum bac | A0A2G2VX13 |
| Capsicum bac | A0A2G2WZV8 |
| Capsicum bac | A0A2G2WG79 |
| Capsicum bac | A0A2G2XMD7 |
| Capsicum bac | A0A2G2VAJ1 |
| Capsicum bac | A0A2G2XEE7 |
| Capsicum bac | A0A2G2WEJ3 |
| Capsicum bac | A0A2G2WDD7 |
| Capsicum bac | A0A2G2X0L9 |
| Capsicum bac | A0A2G2VAF3 |
| Capsicum bac | A0A2G2WEF4 |
| Capsicum bac | A0A2G2WG85 |
| Capsicum bac | A0A2G2VBR2 |
| Capsicum bac | A0A2G2WZY5 |
| Capsicum bac | A0A2G2XMN2 |
| Capsicum bac | A0A2G2WG76 |
| Capsicum bac | A0A2G2VAE7 |
| Capsicum bac | A0A2G2XBY4 |
| Capsicum bac | A0A2G2WRP5 |
| Capsicum bac | A0A2G2VAE4 |
| Capsicum bac | A0A2G2WEK2 |
| Capsicum bac | A0A2G2XEX2 |
| Capsicum bac | A0A2G2WZR1 |
| Capsicum bac | A0A2G2X0F7 |
| Capsicum bac | A0A2G2WED8 |
| Capsicum bac | A0A2G2X1N6 |
| Capsicum bac | A0A2G2VAK1 |
| Capsicum bac | A0A2G2WPT7 |
| Capsicum bac | A0A2G2XMC0 |
| Capsicum bac | A0A2G2WG69 |
| Capsicum bac | A0A2G2WG90 |
| Capsicum bac | A0A2G2WG92 |
| Capsicum bac | A0A2G2VRB8 |
| Capsicum bac | A0A2G2WED9 |
| Capsicum bac | A0A2G2WX08 |

| | |
|---|---|
| Capsicum bac | A0A2G2XCD8 |
| Capsicum bac | A0A2G2XLJ2 |
| Capsicum bac | A0A2G2XMA3 |
| Capsicum bac | A0A2G2XHT8 |
| Capsicum bac | A0A2G2X346 |
| Capsicum bac | A0A2G2X0C9 |
| Capsicum bac | A0A2G2VAE9 |
| Capsicum bac | A0A2G2WGA3 |
| Capsicum bac | A0A2G2VAD6 |
| Capsicum bac | A0A2G2WEI6 |
| Capsicum bac | A0A2G2VAD0 |
| Capsicum bac | A0A2G2W6E0 |
| Capsicum bac | A0A2G2WD99 |
| Capsicum bac | A0A2G2WEF8 |
| Capsicum bac | A0A2G2WZR7 |
| Capsicum bac | A0A2G2VAL5 |
| Capsicum bac | A0A2G2X3S1 |
| Capsicum bac | A0A2G2VCQ1 |
| Capsicum bac | A0A2G2WDF4 |
| Capsicum bac | A0A2G2VAC5 |
| Capsicum bac | A0A2G2VKF1 |
| Capsicum bac | A0A2G2VRK8 |
| Capsicum bac | A0A2G2X0B3 |
| Capsicum bac | A0A2G2WW24 |
| Capsicum bac | A0A2G2WG68 |
| Capsicum bac | A0A2G2VAF6 |
| Capsicum bac | A0A2G2UYX7 |
| Capsicum bac | A0A2G2WVX3 |
| Capsicum bac | A0A2G2X5R0 |
| Capsicum bac | A0A2G2VPU4 |
| Capsicum bac | A0A2G2WDS9 |
| Capsicum bac | A0A2G2WDD3 |
| Capsicum bac | A0A2G2XFD6 |
| Capsicum bac | A0A2G2VAF2 |
| Capsicum bac | A0A2G2X0H2 |
| Capsicum bac | A0A2G2WGA2 |
| Capsicum chi | A0A2G3D6E9 |
| Capsicum chi | A0A2G3C1F1 |
| Capsicum chi | A0A2G3C1E0 |
| Capsicum chi | A0A2G3C1G5 |
| Capsicum chi | A0A2G3C2Z3 |

Capsicum chi A0A2G3CPI0
Capsicum chi A0A2G3DCY1
Capsicum chi A0A2G3C316
Capsicum chi A0A2G3D2F1
Capsicum chi A0A2G3C2Z9
Capsicum chi A0A2G3C1E1
Capsicum chi A0A2G3D6B8
Capsicum chi A0A2G3CPH7
Capsicum chi A0A2G3BYB2
Capsicum chi A0A2G3CJH8
Capsicum chi A0A2G3C300
Capsicum chi A0A2G3D9N7
Capsicum chi A0A2G3CKE6
Capsicum chi A0A2G3C305
Capsicum chi A0A2G3C0I4
Capsicum chi A0A2G3BJQ2
Capsicum chi A0A2G3D6E1
Capsicum chi A0A2G3C1I7
Capsicum chi A0A2G3C0J7
Capsicum chi A0A2G3D6F2
Capsicum chi A0A2G3C0I0
Capsicum chi A0A2G3C307
Capsicum chi A0A2G3CK88
Capsicum chi A0A2G3D6G2
Capsicum chi A0A2G3D6D1
Capsicum chi A0A2G3C0J0
Capsicum chi A0A2G3C304
Capsicum chi A0A2G3B5S2
Capsicum chi A0A2G3CF44
Capsicum chi A0A2G3CKB8
Capsicum chi A0A2G3C314
Capsicum chi A0A2G3CPI6
Capsicum chi A0A2G3CPF9
Capsicum chi A0A2G3D2F8
Capsicum chi A0A2G3BYR9
Capsicum chi A0A2G3BIS4
Capsicum chi A0A2G3CPH6
Capsicum chi A0A2G3BNQ8
Capsicum chi A0A2G3D5W0
Capsicum chi A0A2G3DEU0
Capsicum chi A0A2G3D034

Capsicum chi A0A2G3C0M5
Capsicum chi A0A2G3D6E2
Capsicum chi A0A2G3CPH2
Capsicum chi A0A2G3C1F2
Capsicum chi A0A2G3D4Y8
Capsicum chi A0A2G3CUB9
Capsicum chi A0A2G3BNC4
Capsicum chi A0A2G3D6F6
Capsicum chi A0A2G3C2X9
Capsicum chi A0A2G3BKG5
Capsicum chi A0A2G3D6D5
Capsicum chi A0A2G3C180
Capsicum chi A0A2G3C355
Capsicum chi A0A2G3D6D6
Capsicum chi A0A2G3C0G2
Capsicum chi A0A2G3CPJ8
Capsicum chi A0A2G3CPG3
Capsicum chi A0A2G3C353
Capsicum chi A0A2G3CPG9
Capsicum chi A0A2G3AYH3
Capsicum chi A0A2G3DCW4
Capsicum chi A0A2G3C265
Capsicum chi A0A2G3CZ75
Capsicum chi A0A2G3C313
Capsicum chi A0A2G3CPH9
Capsicum chi A0A2G3BAY7
Capsicum chi A0A2G3D2T3
Capsicum chi A0A2G3BSY6
Capsicum chi A0A2G3ATR3
Capsicum chi A0A2G3CPI5
Capsicum chi A0A2G3BXI5
Capsicum chi A0A2G3D6G1
Capsicum chi A0A2G3CL76
Capsicum chi A0A2G3DD18
Capsicum chi A0A2G3D5A9
Capsicum chi A0A2G3D6C0
Capsicum chi A0A2G3CF81
Capsicum chi A0A2G3CPV5
Capsicum chi A0A2G3BN91
Capsicum chi A0A2G3D5Y6
Caragana kor M9RTK5

| | |
|---|---|
| Carassius aur | A0A0U2QU59 |
| Cariama crist | A0A091LRX5 |
| Cariama crist | A0A091LPV4 |
| Carios mimor | A0A147B803 |
| Castor canad | A0A250YCG5 |
| Castor fiber.x | A0A0K2A1W0 |
| Catenaria ang | A0A1Y2HTZ2 |
| Catenaria ang | A0A1Y2HS76 |
| Cathartes aur | A0A091LCZ8 |
| Cathartes aur | A0A091LCI0 |
| Cathartes aur | A0A091LH38 |
| Cathartes aur | A0A091LE22 |
| Catovirus CT\ | A0A1V0SBF1 |
| Caulobactera | A0A1B1AEB3 |
| Cavia porcell | H0UU59 |
| Cavia porcell | H0V4A7 |
| Cavia porcell | H0VJ51 |
| Cavia porcell | H0VJK2 |
| Cavia porcell | H0VMN8 |
| Cavia porcell | A0A1S6L629 |
| Cebus capuci | A0A2K5R7R9 |
| Cebus capuci | A0A2K5S2Q5 |
| Cebus capuci | A0A2K5S2P6 |
| Cebus capuci | A0A2K5RX96 |
| Cebus capuci | A0A2K5S9C9 |
| Cebus capuci | A0A2K5S2F6 |
| Cebus capuci | A0A2K5S9D7 |
| Cebus capuci | A0A2K5S2N2 |
| Celeribacter l | A0A2R4M5G7 |
| Celeribacter l | A0A1I3X2Y8 |
| Cephalotus fo | A0A1Q3CUV5 |
| Cephalotus fo | A0A1Q3BH96 |
| Cephalotus fo | A0A1Q3CMR9 |
| Cephalotus fo | A0A1Q3DDN1 |
| Cephalotus fo | A0A1Q3C1X7 |
| Cephalotus fo | A0A1Q3DAG8 |
| Cephalotus fo | A0A1Q3C304 |
| Cephalotus fo | A0A1Q3BHR2 |
| Cephalotus fo | A0A1Q3DDT5 |
| Cephalotus fo | A0A1Q3D9K6 |
| Cephalotus fo | A0A1Q3ATB9 |

| | |
|---|---|
| Cephalotus fo | A0A1Q3CIF3 |
| Cephalotus fo | A0A1Q3B9L6 |
| Cephalotus fo | A0A1Q3BTA5 |
| Ceraceosorus | A0A0P1BME7 |
| Ceraceosorus | A0A316VR68 |
| Ceraceosorus | A0A316VS87 |
| Ceraceosorus | A0A316VQI0 |
| Ceratitis capi | W8C6U2 |
| Ceratitis capi | W8C174 |
| Ceratitis capi | W8C1F8 |
| Ceratitis capi | W8ATI6 |
| Ceratitis capi | W8BF27 |
| Ceratitis capi | W8B238 |
| Ceratitis capi | W8BBW0 |
| Ceratitis capi | W8BCC6 |
| Cercocebus a | A0A2K5MGN8 |
| Cercocebus a | A0A2K5M9B1 |
| Cercocebus a | A0A2K5N9N6 |
| Cercocebus a | A0A2K5NA06 |
| Cercocebus a | A0A2K5MGR7 |
| Cercocebus a | A0A2K5N874 |
| Cercocebus a | A0A2K5N9U0 |
| Cercocebus a | A0A2K5LD51 |
| Cercocebus a | A0A2K5N9Z6 |
| Cercospora b | A0A2S6CCY8 |
| Cercospora b | A0A2S6BQS9 |
| Cercospora b | A0A2S6CDU3 |
| Cercospora b | A0A2S6BW05 |
| Cercospora b | A0A2S6BSE0 |
| Cercospora b | A0A2S6CLP9 |
| Cercospora b | A0A2S6C3L0 |
| Cercospora b | A0A2S6BTN4 |
| Cercospora b | A0A2S6C4G5 |
| Cercospora b | A0A2S6C470 |
| Cercospora b | A0A2S6BY54 |
| Cercospora b | A0A2S6C2K4 |
| Cercospora b | A0A2G5HUI6 |
| Cercospora b | A0A2G5IDK6 |
| Cercospora b | A0A2G5IA94 |
| Cercospora b | A0A2G5HDH1 |
| Cercospora b | A0A2G5HEC5 |

| | |
|---|---|
| Cercospora b | A0A2G5HCY6 |
| Cercospora b | A0A2G5HUJ2 |
| Cercospora b | A0A2G5I2T2 |
| Cercospora b | A0A2G5HEX4 |
| Cercospora b | A0A2G5HP23 |
| Cercospora b | A0A2G5HTF1 |
| Cercospora b | A0A2G5I1N9 |
| Cercospora b | A0A2G5HCN3 |
| Cercospora b | A0A2G5HGT9 |
| Cercospora b | A0A2G5IDY8 |
| Cercospora b | A0A2G5HL32 |
| Cercospora b | A0A2G5HYX7 |
| Cercospora b | A0A2G5I599 |
| Cercospora z | A0A2I0RLP9 |
| Cercospora z | A0A2I0RUR2 |
| Cercospora z | A0A2I0S1B2 |
| Cercospora z | A0A2I0RXR6 |
| Cercospora z | A0A2I0RX04 |
| Cercospora z | A0A2I0RUR7 |
| Cercospora z | A0A2I0RNL9 |
| Cercospora z | A0A2I0RI80 |
| Cercospora z | A0A2I0RRK8 |
| Cereibacter c | A0A2T4JY52 |
| Cereibacter c | A0A2W7S7A2 |
| Cervus elaph | A0A212C8S4 |
| Cervus elaph | A0A212C7P2 |
| Cervus elaph | A0A212CKS8 |
| Cervus elaph | A0A212C307 |
| Cervus elaph | A0A212C2K8 |
| Cervus elaph | A0A212C346 |
| Chaetomium | G0S7E3 |
| Chaetothyria | A0A3M7NZG2 |
| Chaetothyria | A0A3M7NLC0 |
| Chara braunii | A0A388K5P6 |
| Chara braunii | A0A388K2B3 |
| Chara braunii | A0A388LIL5 |
| Charadrius vc | A0A0A0A1C3 |
| Charadrius vc | A0A0A0ACH9 |
| Charadrius vc | A0A0A0A4L1 |
| Charadrius vc | A0A0A0AEL2 |
| Chelativorans | Q11FM9 |

| | |
|---|---|
| Chelativorans | Q11LR2 |
| Chelatococcu | A0A2W6Y8B9 |
| Chelonia myc | M7B6M9 |
| Chelonia myc | M7B4A9 |
| Chilo suppres | A0A437B6K6 |
| Chilo suppres | A0A437BD02 |
| Chilo suppres | A0A3S2P060 |
| Chilo suppres | A0A3S2NRB4 |
| Chilo suppres | A0A3S2TGF1 |
| Chiloscyllium | A0A401S8F4 |
| Chiloscyllium | A0A401SUD6 |
| Chiloscyllium | A0A401T7K4 |
| Chiloscyllium | A0A401S098 |
| Chlamydomo | A0A250WPW7 |
| Chlamydomo | A0A250WQA5 |
| Chlamydomo | A0A250WYW3 |
| Chlamydomo | A0A2K3DSA5 |
| Chlamydomo | A0A2K3E6A5 |
| Chlamydomo | A8HZX7 |
| Chlamydomo | A0A2K3DB77 |
| Chlamydomo | A8HQ30 |
| Chlamydomo | A8IBD2 |
| Chlamydomo | A0A2K3DBP7 |
| Chlamydomo | A0A2K3D8X6 |
| Chlamydomo | A0A2K3E6A3 |
| Chlamydomo | A0A2K3DB83 |
| Chlamydomo | A0A2K3CTH7 |
| Chlamydomo | A0A2K3CSI6 |
| Chlamydomo | A8IFR1 |
| Chlamydomo | A8IMR7 |
| Chlamydomo | A8HQN7 |
| Chlamydomo | A0A2K3E6U0 |
| Chlamydomo | A0A2K3DBQ5 |
| Chlamydomo | A0A2K3DBU7 |
| Chlamydomo | A8J8N4 |
| Chlamydomo | A0A2K3D2L0 |
| Chlamydomo | A0A2K3DS95 |
| Chlamydomo | A0A2K3DBP8 |
| Chlamydomo | A0A2K3DMT1 |
| Chlamydomo | A8J1H4 |
| Chlamydomo | A0A2K3D8R5 |

| | |
|---|---|
| Chlamydomo | A0A2K3DWJ6 |
| Chlamydomo | A0A2K3DCU6 |
| Chlamydomo | A8IKN5 |
| Chlamydomo | A0A2K3E6C5 |
| Chlamydomo | A0A2K3D9V3 |
| Chlamydomo | A8HX99 |
| Chlamydomo | A0A2K3D9V6 |
| Chlamydomo | A8JBG8 |
| Chlamydomo | A8IKN3 |
| Chlamydotis | A0A091L5H5 |
| Chlamydotis | A0A091L0G7 |
| Chlamydotis | A0A091KL34 |
| Chloebia gou | A0A3L8SH23 |
| Chloebia gou | A0A3L8S1X4 |
| Chloebia gou | A0A3L8SV11 |
| Chloebia gou | A0A3L8S925 |
| Chlorella sorc | A0A2P6TKL9 |
| Chlorella sorc | A0A2P6TF11 |
| Chlorella sorc | A0A2P6U331 |
| Chlorella sorc | A0A2P6TIU0 |
| Chlorella sorc | A0A2P6TGU7 |
| Chlorella sorc | A0A2P6TIQ9 |
| Chlorella sorc | A0A2P6TGF4 |
| Chlorella sorc | A0A2P6TFU2 |
| Chlorella sorc | A0A2P6TLH4 |
| Chlorella sorc | A0A2P6TJF6 |
| Chlorella sorc | A0A2P6U011 |
| Chlorella sorc | A0A2P6TXL2 |
| Chlorella sorc | A0A2P6U051 |
| Chlorella sorc | A0A2P6TCK0 |
| Chlorella sorc | A0A2P6TW22 |
| Chlorella sorc | A0A2P6TZN7 |
| Chlorella sorc | A0A2P6TCJ6 |
| Chlorella sorc | A0A2P6TUR6 |
| Chlorella sorc | A0A2P6THR0 |
| Chlorella sorc | A0A2P6TIQ5 |
| Chlorella sorc | A0A2P6TMR9 |
| Chlorella sorc | A0A2P6TR31 |
| Chlorella sorc | A0A2P6TH81 |
| Chlorella sorc | A0A2P6TP21 |
| Chlorella sorc | A0A2P6TVR3 |

| | |
|---|---|
| Chlorella sorc | A0A2P6TES6 |
| Chlorella sorc | A0A2P6TXZ7 |
| Chlorella sorc | A0A2P6TMQ7 |
| Chlorella sorc | A0A2P6TKW1 |
| Chlorella sorc | A0A2P6TQS8 |
| Chlorella sorc | A0A2P6TIZ9 |
| Chlorella sorc | A0A2P6TF15 |
| Chlorella sorc | A0A2P6U012 |
| Chlorella sorc | A0A2P6TMT2 |
| Chlorella sorc | A0A2P6U316 |
| Chlorella vari | E1ZPG0 |
| Chlorella vari | E1ZRL2 |
| Chlorella vari | E1Z6K2 |
| Chlorella vari | E1ZHG6 |
| Chlorella vari | E1ZJC2 |
| Chlorella vari | E1ZUC6 |
| Chlorella vari | E1ZHR7 |
| Chlorella vari | E1ZF04 |
| Chlorella vari | E1ZHR6 |
| Chlorella vari | E1Z7H7 |
| Chlorella vari | E1Z291 |
| Chlorella vari | E1Z6Q1 |
| Chlorella vari | E1ZJG5 |
| Chlorella vari | E1ZM59 |
| Chlorella vari | E1ZG06 |
| Chlorella vari | E1ZAC6 |
| Chlorella vari | E1Z965 |
| Chlorella vari | E1ZBZ7 |
| Chlorella vari | E1Z7U0 |
| Chlorocebus : | A0A0D9RIP2 |
| Chlorocebus : | A0A0D9S2M3 |
| Chlorocebus : | A0A0D9RHM4 |
| Chlorocebus : | A0A0D9S2M6 |
| Chlorocebus : | A0A0D9RQW6 |
| Chloroflexi ba | A0A2E1ZYY8 |
| Chondromyce | A0A017T0L6 |
| Chondromyce | A0A017T6P0 |
| Chromera vel | A0A0G4GGC1 |
| Chromera vel | A0A0G4G2T2 |
| Chromera vel | A0A0G4HXM5 |
| Chromera vel | A0A0G4HH89 |

| | |
|---|---|
| Chromera vel | A0A0G4GSE8 |
| Chromera vel | A0A0G4IBR5 |
| Chromera vel | A0A0G4H481 |
| Chromera vel | A0A0G4I991 |
| Chromera vel | A0A0G4FQB4 |
| Chromera vel | A0A0G4GMQ5 |
| Chromera vel | A0A0G4GZC4 |
| Chromera vel | A0A0G4HGC9 |
| Chromera vel | A0A0G4FVU0 |
| Chromera vel | A0A0G4F227 |
| Chromera vel | A0A0G4FYS2 |
| Chromera vel | A0A0G4FGP6 |
| Chromera vel | A0A0G4FEZ3 |
| Chromera vel | A0A0G4I700 |
| Chromera vel | A0A0G4GGB9 |
| Chromera vel | A0A0G4F023 |
| Chromera vel | A0A0G4FZM7 |
| Chromera vel | A0A0G4HZX5 |
| Chromera vel | A0A0G4HKE3 |
| Chromera vel | A0A0G4HYV2 |
| Chrysochrom | A0A0N9QJ81 |
| Chrysochrom | A0A0N9R3K7 |
| Chrysochrom | A0A0M0JLK2 |
| Chrysochrom | A0A0M0JFA1 |
| Chrysochrom | A0A0M0J9M7 |
| Chrysochrom | A0A0M0J502 |
| Chrysochrom | A0A0M0JW38 |
| Chrysochrom | A0A0M0JSY5 |
| Chrysochrom | A0A0M0JH75 |
| Cicer arietinu | A0A1S2Z201 |
| Cicer arietinu | A0A1S2Y6R3 |
| Cicer arietinu | A0A1S2YVX3 |
| Cicer arietinu | A0A1S2Y702 |
| Cicer arietinu | A0A1S2Z2P8 |
| Cicer arietinu | A0A1S2XCS8 |
| Cicer arietinu | A0A1S2XT54 |
| Cicer arietinu | A0A1S2XFS7 |
| Cicer arietinu | A0A1S2YU61 |
| Cicer arietinu | A0A1S3E0S5 |
| Cicer arietinu | A0A1S2XB68 |
| Cicer arietinu | A0A3Q7YCB7 |

| | |
|---|---|
| Cicer arietinu | A0A1S2YU62 |
| Cicer arietinu | A0A1S2XDH3 |
| Cicer arietinu | A0A1S3E9I2 |
| Cicer arietinu | A0A1S3EJ26 |
| Cicer arietinu | A0A1S3ED37 |
| Cicer arietinu | A0A1S2XP43 |
| Cicer arietinu | A0A3Q7XHP2 |
| Cicer arietinu | A0A1S2XCS7 |
| Cicer arietinu | A0A1S2X9K5 |
| Cicer arietinu | A0A1S2XP07 |
| Cicer arietinu | A0A3Q7YC42 |
| Cicer arietinu | A0A3Q7Y1H8 |
| Cicer arietinu | A0A1S3EDN3 |
| Cicer arietinu | A0A1S2XBI0 |
| Cicer arietinu | A0A3Q7YAU0 |
| Cicer arietinu | A0A3Q7Y285 |
| Cicer arietinu | A0A3Q7Y6L8 |
| Cicer arietinu | A0A1S2Z7A8 |
| Cicer arietinu | A0A3Q7XQV7 |
| Cicer arietinu | A0A3Q7YG81 |
| Cicer arietinu | A0A3Q7Y0F7 |
| Cicer arietinu | A0A1S2Z2N4 |
| Cicer arietinu | A0A3Q7XQV2 |
| Cicer arietinu | A0A1S2Y575 |
| Cicer arietinu | A0A1S2Z2N5 |
| Cicer arietinu | A0A3Q7WWB2 |
| Cicer arietinu | A0A1S2Y5U4 |
| Cicer arietinu | A0A1S3E9A9 |
| Cicer arietinu | A0A1S2X9L3 |
| Cicer arietinu | A0A3Q7XBV1 |
| Cicer arietinu | A0A1S2XRR7 |
| Cicer arietinu | A0A1S3E9B2 |
| Cicer arietinu | A0A3Q7XAI7 |
| Cicer arietinu | A0A3Q7YAZ7 |
| Cicer arietinu | A0A3Q7Y008 |
| Cicer arietinu | A0A3Q7YFJ7 |
| Cicer arietinu | A0A1S2Z200 |
| Cicer arietinu | A0A1S3E6X4 |
| Cicer arietinu | A0A1S2YU56 |
| Cicer arietinu | A0A1S2Y443 |
| Cicer arietinu | A0A1S3E803 |

| | |
|---|---|
| Cichorium int | Q84V84 |
| Cinnamomun | A0A443PRY7 |
| Cinnamomun | A0A443NX35 |
| Cinnamomun | A0A3S3NIB5 |
| Cinnamomun | A0A443P5K6 |
| Cinnamomun | A0A443PBN9 |
| Cinnamomun | A0A443PBP3 |
| Cinnamomun | A0A443P0S9 |
| Cinnamomun | A0A443PS04 |
| Cinnamomun | A0A443PW89 |
| Cinnamomun | A0A443PS03 |
| Cinnamomun | A0A3S3MEI8 |
| Cinnamomun | A0A443PN61 |
| Cinnamomun | A0A3S3N3K9 |
| Cinnamomun | A0A443NLN9 |
| Cinnamomun | A0A3S3MJQ9 |
| Cinnamomun | A0A3S3NA45 |
| Cinnamomun | A0A443P5V0 |
| Cinnamomun | A0A443P2S3 |
| Cinnamomun | A0A3S3PD76 |
| Cinnamomun | A0A3S3MXI8 |
| Cinnamomun | A0A443PCA0 |
| Cinnamomun | A0A443P2G4 |
| Cinnamomun | A0A443PBN2 |
| Cinnamomun | A0A443PS75 |
| Cinnamomun | A0A3S3MXL1 |
| Cinnamomun | A0A443PBP0 |
| Cinnamomun | A0A3S3MR36 |
| Cinnamomun | A0A3S3NVL5 |
| Cinnamomun | A0A443PBQ1 |
| Cinnamomun | A0A3S3P012 |
| Cinnamomun | A0A3S3NGN2 |
| Cinnamomun | A0A443PBP1 |
| Cinnamomun | A0A3S3QL46 |
| Cinnamomun | A0A443PBQ8 |
| Cinnamomun | A0A3S3MJ71 |
| Cinnamomun | A0A443PBP5 |
| Cinnamomun | A0A3S3PSG0 |
| Cinnamomun | A0A443PBN3 |
| Cinnamomun | A0A443PDZ2 |
| Cinnamomun | A0A3S3N939 |

| | |
|---|---|
| Cinnamomun | A0A3S3PEW5 |
| Cinnamomun | A0A3S4PKF1 |
| Cinnamomun | A0A3S3NWW4 |
| Cinnamomun | A0A3S3MXN2 |
| Cinnamomun | A0A3S3QIN5 |
| Cinnamomun | A0A3S3QJN6 |
| Cinnamomun | A0A3S3NA33 |
| Cinnamomun | A0A3S3NPA9 |
| Cinnamomun | A0A443NX39 |
| Cinnamomun | A0A443N5D8 |
| Cinnamomun | A0A3S3N3I2 |
| Cinnamomun | A0A443PBQ2 |
| Cinnamomun | A0A3S3MLU2 |
| Cinnamomun | A0A3S3MTZ8 |
| Cinnamomun | A0A3S3MVZ3 |
| Cinnamomun | A0A443PBN5 |
| Cinnamomun | A0A443PS14 |
| Cinnamomun | A0A3S3NH01 |
| Cinnamomun | A0A3S3MY34 |
| Cinnamomun | A0A3S3QYD7 |
| Cinnamomun | A0A3S3NIC6 |
| Cinnamomun | A0A443PMN8 |
| Cinnamomun | A0A3S3N4I9 |
| Cinnamomun | A0A443PBM7 |
| Cinnamomun | A0A443P2M2 |
| Cinnamomun | A0A3S4PQK3 |
| Cinnamomun | A0A3S3QQQ3 |
| Cinnamomun | A0A3S3QP22 |
| Cinnamomun | A0A443PLP3 |
| Cinnamomun | A0A3S3PD65 |
| Cinnamomun | A0A443PBN0 |
| Cinnamomun | A0A443NF22 |
| Cinnamomun | A0A443PLH6 |
| Cinnamomun | A0A443PS49 |
| Cinnamomun | A0A3S3NDM0 |
| Cinnamomun | A0A3S3PD86 |
| Cinnamomun | A0A443PJM2 |
| Cinnamomun | A0A3S3NJS6 |
| Cinnamomun | A0A443PLJ7 |
| Cinnamomun | A0A443PBQ0 |
| Cinnamomun | A0A3S3MXK7 |

| | |
|---|---|
| Cinnamomun | A0A443P0T7 |
| Ciona intestir | Q69HN0 |
| Ciona intestir | A0A1W5BT82 |
| Ciona intestir | A0A3Q0K068 |
| Ciona intestir | A0A3Q0JVM7 |
| Ciona intestir | A0A1W2WKK1 |
| Ciona intestir | A0A3Q0JM87 |
| Ciona intestir | A0A1W5BJJ9 |
| Ciona intestir | A0A3Q0K9Z4 |
| Ciona intestir | F6PY33 |
| Ciona intestir | F6UI41 |
| Ciona intestir | A0A1W5B4S5 |
| Ciona intestir | F6RQT3 |
| Ciona intestir | A0A1W5BDQ8 |
| Ciona intestir | A0A3Q0JQ58 |
| Ciona savigny | H2YF24 |
| Ciona savigny | H2ZPV7 |
| Citreicella sp. | I1ATJ2 |
| Citreicella sp. | D0D4L1 |
| Citreicella sp. | A0A355QKE1 |
| Citreicella sp. | A0A357RS17 |
| Citreimonas s | A0A1H3M0F3 |
| Citrus clemer | V4UM33 |
| Citrus clemer | V4S782 |
| Citrus clemer | V4WFC0 |
| Citrus clemer | V4US31 |
| Citrus clemer | V4UM62 |
| Citrus clemer | V4W3X3 |
| Citrus clemer | V4UM47 |
| Citrus clemer | V4UGX9 |
| Citrus clemer | V4TXJ0 |
| Citrus clemer | V4SX72 |
| Citrus clemer | V4T0F1 |
| Citrus clemer | V4UGU5 |
| Citrus clemer | V4TXM4 |
| Citrus clemer | V4UGV0 |
| Citrus clemer | V4WFE7 |
| Citrus clemer | V4UGU3 |
| Citrus clemer | V4UG60 |
| Citrus clemer | V4U8W0 |
| Citrus clemer | V4TPH0 |

Citrus clemer V4TXH2
Citrus clemer V4ULH8
Citrus clemer V4UM07
Citrus clemer V4UAV8
Citrus clemer V4WFD4
Citrus clemer V4UM16
Citrus clemer V4US56
Citrus clemer V4TXI0
Citrus clemer V4WFB7
Citrus clemer V4TXL6
Citrus clemer V4US24
Citrus clemer V4UGR6
Citrus clemer V4RYV2
Citrus clemer V4SQ65
Citrus clemer V4UM23
Citrus clemer V4UR15
Citrus clemer V4WFE2
Citrus clemer V4TQP7
Citrus clemer V4U0T7
Citrus clemer V4US39
Citrus clemer V4SZW8
Citrus clemer V4UGT3
Citrus clemer V4W877
Citrus clemer V4SUW1
Citrus clemer V4UGW8
Citrus clemer V4WFB2
Citrus clemer V4S160
Citrus clemer V4WFC6
Citrus clemer V4STW6
Citrus clemer V4T683
Citrus clemer V4UM36
Citrus clemer V4UEI5
Citrus clemer V4UM71
Citrus clemer V4USA4
Citrus clemer V4T894
Citrus clemer V4SP31
Citrus clemer V4TXG2
Citrus clemer V4W5E2
Citrus clemer V4SHZ9
Citrus clemer V4U0T3
Citrus clemer V4SDM6

Citrus clemer V4UGS3
Citrus clemer V4SNM6
Citrus clemer V4TXJ3
Citrus clemer V4TVE6
Citrus clemer V4RYU8
Citrus clemer V4W3W8
Citrus clemer V4US66
Citrus clemer V4W3G2
Citrus clemer V4UCD6
Citrus clemer V4WFN6
Citrus clemer V4TLL6
Citrus clemer V4US48
Citrus clemer V4U017
Citrus clemer V4US05
Citrus clemer V4TXJ7
Citrus clemer V4UM67
Citrus clemer V4US34
Citrus clemer V4US51
Citrus clemer V4US13
Citrus clemer V4ULH4
Citrus clemer V4TXK0
Citrus clemer V4UM52
Citrus clemer V4SQ60
Citrus clemer V4US61
Citrus clemer V4U986
Citrus clemer V4U5T3
Citrus clemer V4WFD7
Citrus clemer V4W873
Citrus clemer V4TXM0
Citrus clemer V4SV65
Citrus clemer V4UM41
Citrus clemer V4U452
Citrus clemer V4STW9
Citrus clemer V4STF2
Citrus clemer V4STL9
Citrus clemer V4UGV5
Citrus clemer V4U9H6
Citrus clemer V4W3W6
Citrus clemer V4U5T6
Citrus clemer V4SMY3
Citrus clemer V4SSX8

Citrus clemer V4UGT8
Citrus clemer V4S7S5
Citrus clemer V4T687
Citrus clemer V4UGS8
Citrus clemer V4ULH0
Citrus clemer V4UM27
Citrus limon.› B2CDF6
Citrus sinensi A0A067GG52
Citrus sinensi A0A067G6T4
Citrus sinensi A0A067GFX5
Citrus sinensi A0A067CZ86
Citrus sinensi A0A067GG63
Citrus sinensi A0A067GFU2
Citrus sinensi A0A067G437
Citrus sinensi A0A067G408
Citrus sinensi A0A067GAK8
Citrus sinensi A0A067GG46
Citrus sinensi A0A067ERZ0
Citrus sinensi A0A067G3Z5
Citrus sinensi A0A067G447
Citrus sinensi A0A067DNX9
Citrus sinensi A0A067ESK3
Citrus sinensi A0A067GFW5
Citrus sinensi A0A067EC69
Citrus sinensi A0A067DC17
Citrus sinensi A0A067FDW2
Citrus sinensi A0A067GFW9
Citrus sinensi A0A067DFP7
Citrus sinensi A0A067DJF5
Citrus sinensi A0A067G402
Citrus sinensi A0A067D6A0
Citrus sinensi A0A067DEZ1
Citrus sinensi A0A067D709
Citrus sinensi A0A067E075
Citrus sinensi A0A067ESU0
Citrus sinensi A0A067ERZ4
Citrus sinensi A0A067GG54
Citrus sinensi A0A067G3X5
Citrus sinensi A0A067GFU7
Citrus sinensi A0A067F300
Citrus sinensi A0A067G6V7

Citrus sinensi A0A067G416
Citrus sinensi A0A067F0B3
Citrus sinensi A0A067G7T6
Citrus sinensi A0A067GG67
Citrus sinensi A0A067FCY6
Citrus sinensi A0A067GFY6
Citrus sinensi A0A067G429
Citrus sinensi A0A067EZ26
Citrus sinensi A0A067G407
Citrus sinensi A0A067G444
Citrus sinensi A0A067DHM7
Citrus sinensi A0A067GG72
Citrus sinensi A0A067G7U1
Citrus sinensi A0A067DGG9
Citrus sinensi A0A067G3W6
Citrus sinensi A0A067F108
Citrus sinensi A0A067GG42
Citrus sinensi A0A067EXK6
Citrus sinensi A0A067DI35
Citrus sinensi A0A067G3X9
Citrus sinensi A0A067F5Z7
Citrus sinensi A0A067G419
Citrus sinensi A0A067DP49
Citrus sinensi A0A067D963
Citrus sinensi A0A067ESK4
Citrus sinensi A0A067GAK4
Citrus sinensi A0A067G6T1
Citrus sinensi A0A067G3Y0
Citrus sinensi A0A067DXE4
Citrus sinensi A0A067GFZ4
Citrus sinensi A0A067GG78
Citrus sinensi A0A067G404
Citrus sinensi A0A067DEU1
Citrus sinensi A0A067F642
Citrus sinensi A0A067DP44
Citrus sinensi A0A067EP43
Citrus sinensi A0A067ECN7
Citrus sinensi A0A067EFL5
Citrus sinensi A0A067G3Y8
Citrus sinensi A0A067ERY8
Citrus sinensi A0A067G3Y3

Citrus sinensi A0A067GFX8
Citrus sinensi A0A067EMR0
Citrus sinensi A0A067GG48
Citrus sinensi A0A067G7Q9
Citrus sinensi A0A067E705
Citrus sinensi A0A067G451
Citrus sinensi A0A067G6R7
Citrus sinensi A0A067DMY8
Citrus sinensi A0A067ECA5
Citrus sinensi A0A067EVI9
Citrus sinensi A0A067DT16
Citrus sinensi A0A067DFK5
Citrus sinensi A0A067DBK4
Citrus sinensi A0A067G3Z3
Citrus sinensi A0A067DLD4
Citrus sinensi A0A067EI62
Citrus sinensi A0A067EFX3
Citrus sinensi A0A067ENV7
Citrus sinensi A0A067EJB2
Citrus sinensi A0A067GG38
Citrus sinensi A0A067G3V1
Citrus sinensi A0A067GG58
Citrus sinensi A0A067DJW6
Citrus sinensi A0A067DRX6
Citrus sinensi A0A067GFV3
Citrus sinensi A0A067G6U4
Citrus sinensi A0A067G6W5
Citrus sinensi A0A067G6X4
Citrus sinensi A0A067DFT8
Citrus sinensi A0A067EP41
Citrus sinensi A0A067EIF8
Citrus sinensi A0A067G6Z0
Citrus sinensi A0A067FHX6
Citrus sinensi A0A067DIS5
Citrus sinensi A0A067DDY5
Citrus sinensi A0A067EMP5
Citrus sinensi A0A067G6S1
Citrus sinensi A0A067GG04
Citrus sinensi A0A067GFZ1
Citrus sinensi A0A067DJF0
Citrus sinensi A0A067F0Y8

Citrus sinensi A0A067G412
Citrus sinensi A0A067G6V2
Citrus sinensi A0A067G6U9
Citrus sinensi A0A067G3Z1
Citrus sinensi A0A067G420
Citrus sinensi A0A067G3V7
Citrus sinensi A0A067GK13
Citrus sinensi A0A067G7V2
Citrus sinensi A0A067GFY1
Citrus sinensi A0A067GK25
Citrus unshiu A0A2H5PKY8
Citrus unshiu A0A2H5PKZ4
Citrus unshiu A0A2H5Q9C5
Citrus unshiu A0A2H5PSJ2
Citrus unshiu A0A2H5PSH0
Citrus unshiu A0A2H5QJF9
Citrus unshiu A0A2H5PKR2
Citrus unshiu A0A2H5PFQ6
Citrus unshiu A0A2H5PSI2
Citrus unshiu A0A2H5PSG3
Citrus unshiu A0A2H5PSI8
Citrus unshiu A0A2H5PSM9
Citrus unshiu A0A2H5PSK6
Citrus unshiu A0A2H5PKW4
Citrus unshiu A0A2H5PKV3
Citrus unshiu A0A2H5PKZ1
Citrus unshiu A0A2H5PSG8
Citrus unshiu A0A2H5PSJ9
Citrus unshiu A0A2H5PSH4
Citrus unshiu A0A2H5PFR7
Citrus unshiu A0A2H5PL00
Citrus unshiu A0A2H5PLJ0
Citrus unshiu A0A2H5PKR5
Citrus unshiu A0A2H5P0E0
Citrus unshiu A0A2H5PYT1
Citrus unshiu A0A2H5Q8A5
Citrus unshiu A0A2H5PSK1
Citrus unshiu A0A2H5PLK0
Citrus unshiu A0A2H5QSC3
Citrus unshiu A0A2H5N494
Citrus unshiu A0A2H5NKK7

Citrus unshiu A0A2H5PKY5
Citrus unshiu A0A2H5MW58
Citrus unshiu A0A2H5PSJ1
Citrus unshiu A0A2H5QY31
Citrus unshiu A0A2H5PKT9
Citrus unshiu A0A2H5N543
Citrus unshiu A0A2H5PFR2
Citrus unshiu A0A2H5QX37
Citrus unshiu A0A2H5NYH0
Citrus unshiu A0A2H5PKT8
Citrus unshiu A0A2H5PKS3
Citrus unshiu A0A2H5PSI7
Citrus unshiu A0A2H5PKV8
Citrus unshiu A0A2H5PKW8
Citrus unshiu A0A2H5QGS6
Citrus unshiu A0A2H5PLH7
Citrus unshiu A0A2H5PKW9
Citrus unshiu A0A2H5N106
Citrus unshiu A0A2H5PSG4
Citrus unshiu A0A2H5PBR2
Citrus unshiu A0A2H5PIJ5
Citrus unshiu A0A2H5QGL4
Citrus unshiu A0A2H5PSL1
Citrus unshiu A0A2H5PSF5
Citrus unshiu A0A2H5QJI9
Citrus unshiu A0A2H5PWC1
Citrus unshiu A0A2H5PLG3
Citrus unshiu A0A2H5PSL8
Citrus unshiu A0A2H5PKU0
Citrus unshiu A0A2H5PKT6
Citrus unshiu A0A2H5QX47
Citrus unshiu A0A2H5PXG7
Citrus unshiu A0A2H5QUK2
Citrus unshiu A0A2H5PTF6
Citrus unshiu A0A2H5PCI5
Citrus unshiu A0A2H5PHZ3
Citrus unshiu A0A2H5QGN3
Citrus unshiu A0A2H5PKS2
Citrus unshiu A0A2H5PHW4
Citrus unshiu A0A2H5PSI9
Citrus unshiu A0A2H5QTN3

Citrus unshiu A0A2H5QDF2
Citrus unshiu A0A2H5PKW6
Citrus unshiu A0A2H5PKX5
Citrus unshiu A0A2H5P8A0
Citrus unshiu A0A2H5PKW0
Citrus unshiu A0A2H5PKV5
Citrus unshiu A0A2H5PKY2
Citrus unshiu A0A2H5PFQ1
Citrus unshiu A0A2H5Q8X1
Citrus unshiu A0A2H5PIK0
Citrus unshiu A0A2H5PSK7
Citrus unshiu A0A2H5PSH7
Citrus unshiu A0A2H5PTD6
Citrus unshiu A0A2H5PHY6
Citrus unshiu A0A2H5MVV6
Citrus unshiu A0A2H5PKT3
Citrus unshiu A0A2H5N2Y6
Citrus unshiu A0A2H5PKW2
Citrus unshiu A0A2H5PLM9
Citrus unshiu A0A2H5PKW5
Citrus unshiu A0A2H5PLK9
Citrus unshiu A0A2H5MW15
Citrus unshiu A0A2H5PKV2
Citrus unshiu A0A2H5MXC0
Citrus unshiu A0A2H5N221
Citrus unshiu A0A2H5QJF2
Citrus unshiu A0A2H5N2E3
Citrus unshiu A0A2H5PTC7
Citrus unshiu A0A2H5PSI1
Citrus unshiu A0A2H5PSH6
Citrus unshiu A0A2H5PSF9
Citrus unshiu A0A2H5PIL0
Citrus unshiu A0A2H5PSJ5
Citrus unshiu A0A2H5QDF6
Citrus unshiu A0A2H5PYF9
Citrus unshiu A0A2H5PFR4
Citrus unshiu A0A2H5PSM2
Citrus unshiu A0A2H5PLL9
Citrus unshiu A0A2H5QGR8
Citrus unshiu A0A2H5QSP4
Citrus unshiu A0A2H5PTB7

Citrus unshiu A0A2H5MXP2
Citrus unshiu A0A2H5QDF7
Citrus unshiu A0A2H5PW39
Citrus unshiu A0A2H5PKU4
Citrus unshiu A0A2H5PKV9
Citrus unshiu A0A2H5PKT1
Citrus unshiu A0A2H5PL05
Citrus unshiu A0A2H5MWP9
Citrus unshiu A0A2H5PKS5
Citrus unshiu A0A2H5PKY1
Citrus unshiu A0A2H5PKZ0
Citrus unshiu A0A2H5PKX3
Citrus unshiu A0A2H5PLN9
Citrus unshiu A0A2H5PSN3
Citrus unshiu A0A2H5PKY6
Citrus unshiu A0A2H5PSI0
Citrus unshiu A0A2H5PKU2
Citrus unshiu A0A2H5MY89
Citrus unshiu A0A2H5PKZ5
Citrus unshiu A0A2H5N248
Citrus unshiu A0A2H5MVR2
Citrus unshiu A0A2H5PC29
Citrus unshiu A0A2H5PSH8
Citrus unshiu A0A2H5PSJ6
Citrus unshiu A0A2H5PKZ9
Citrus unshiu A0A2H5PKV0
Citrus unshiu A0A2H5PKU3
Citrus unshiu A0A2H5QX61
Citrus unshiu A0A2H5PKU8
Citrus unshiu A0A2H5PXK1
Citrus unshiu A0A2H5PKS9
Citrus unshiu A0A2H5PSK2
Citrus unshiu A0A2H5PXH5
Citrus unshiu A0A2H5PKZ7
Cladophialop A0A0D2EPW4
Cladophialop A0A0D2H5D0
Cladophialop A0A0D2D5M4
Cladophialop A0A0D1ZW52
Cladophialop A0A0D2D9C0
Cladophialop W9VHA5
Clastoptera a A0A1B6E728

| | |
|---|---|
| Clastoptera a | A0A1B6DP44 |
| Clastoptera a | A0A1B6CQM5 |
| Clastoptera a | A0A1B6CRN7 |
| Clastoptera a | A0A1B6D4A1 |
| Clastoptera a | A0A1B6CPV4 |
| Clastoptera a | A0A1B6CH46 |
| Claviceps pur | M1VXP1 |
| Clohesyomyc | A0A1Y1ZIY5 |
| Clohesyomyc | A0A1Y2A3K8 |
| Clohesyomyc | A0A1Y1Y9H3 |
| Clohesyomyc | A0A1Y1ZK83 |
| Clohesyomyc | A0A1Y1Z9L6 |
| Clonorchis sir | A0A419PQI0 |
| Clonorchis sir | A0A3R7EPZ0 |
| Clonorchis sir | A0A419QAI4 |
| Clonorchis sir | A0A419PSE2 |
| Clonorchis sir | G7YA90 |
| Clonorchis sir | G7YBT5 |
| Clonorchis sir | A0A3R7DBB2 |
| Clonorchis sir | A0A3R7CQZ5 |
| Clonorchis sir | A0A419PZ32 |
| Clonorchis sir | G7YEU8 |
| Clonorchis sir | G7YWK9 |
| Clonorchis sir | G7Y624 |
| Clunio marinu | A0A1J1IGE1 |
| Clunio marinu | A0A1J1HN04 |
| Clunio marinu | A0A1J1IXY3 |
| Clunio marinu | A0A1J1IRI6 |
| Clunio marinu | A0A1J1I6X9 |
| Clunio marinu | A0A1J1HMP6 |
| Coccidioides | J3K384 |
| Coccidioides | J3K388 |
| Coccidioides | A0A0J8USD8 |
| Coccidioides | A0A0J8S0H2 |
| Coccidioides | A0A0J6Y7D1 |
| Coccidioides | A0A0J6Y7C8 |
| Coccidioides | A0A0J8QTA0 |
| Coccidioides | C5P676 |
| Coccidioides | C5P674 |
| Coccidioides | E9DHQ1 |
| Coccidioides | E9DIX1 |

| | |
|---|---|
| Coccidioides | E9DIM5 |
| Coccidioides | A0A0J6I586 |
| Coccidioides | A0A0J6I580 |
| Coccomyxa su | I0YJN3 |
| Coccomyxa su | I0YQC5 |
| Coccomyxa su | I0YZJ2 |
| Coccomyxa su | I0YQC4 |
| Coccomyxa su | I0Z909 |
| Coccomyxa su | I0YXG6 |
| Cochliobolus | N4WSB7 |
| Cochliobolus | N4XA49 |
| Cochliobolus | N4X4B7 |
| Cochliobolus | N4XWV7 |
| Cochliobolus | N4XAK1 |
| Cochliobolus | N4XNR5 |
| Cochliobolus | N4WIS8 |
| Cochliobolus | N4XE50 |
| Cochliobolus | N4WKJ0 |
| Cochliobolus | M2UDR7 |
| Cochliobolus | M2TCH5 |
| Cochliobolus | M2UPP8 |
| Cochliobolus | M2TUB1 |
| Cochliobolus | M2UZK2 |
| Cochliobolus | M2UC59 |
| Cochliobolus | M2TM44 |
| Cochliobolus | M2SS15 |
| Cochliobolus | M2TNS5 |
| Cochliobolus | M2SEH1 |
| Cochliobolus | M2TCD4 |
| Cochliobolus | M2REQ9 |
| Cochliobolus | M2T729 |
| Cochliobolus | M2QVU0 |
| Cochliobolus | M2SRC8 |
| Cochliobolus | M2SXY7 |
| Cochliobolus | M2SRR6 |
| Coffea canep | A0A068V1N7 |
| Coffea canep | A0A068TTW9 |
| Coffea canep | A0A068U5I0 |
| Coffea canep | A0A068UBR8 |
| Coffea canep | A0A068UMF4 |
| Coffea canep | A0A068VIN5 |

Coffea canep A0A068VAH2
Coffea canep A0A068VH78
Coffea canep A0A068U0Z7
Coffea canep A0A068VD61
Coffea canep A0A068UDX9
Coffea canep A0A068UF61
Coffea canep A0A068UEE7
Coffea canep A0A068TT72
Coffea canep A0A068TUQ8
Coffea canep A0A068U920
Coffea canep A0A068UBA7
Coffea canep A0A068UTC4
Coffea canep A0A068TTY9
Coffea canep A0A068V6K4
Coffea canep A0A068UCT7
Coffea canep A0A068VAV0
Coffea canep A0A068UAW2
Coffea canep A0A068TTU4
Coffea canep A0A068TUK9
Coffea canep A0A068UB95
Coffea canep A0A068U7P3
Coffea canep A0A068UAV7
Coffea canep A0A068U9M3
Coffea canep A0A068TR02
Coffea canep A0A068TUN3
Coffea canep A0A068UC35
Coffea canep A0A068VIA2
Coffea canep A0A068V9X1
Coffea canep A0A068TWF3
Coffea canep A0A068V7U6
Coffea canep A0A068U9A9
Coffea canep A0A068V658
Coffea canep A0A068VDF8
Coffea canep A0A068TUN8
Coffea canep A0A068URU0
Coffea canep A0A068TWI8
Coffea canep A0A068TPT8
Coffea canep A0A068UCP1
Coffea canep A0A068UB32
Coffea canep A0A068UU33
Coffea canep A0A068VDL2

| | |
|---|---|
| Coffea canep | A0A068UUD5 |
| Coffea canep | A0A068V600 |
| Coffea canep | A0A068U9B6 |
| Coffea canep | A0A068U735 |
| Coffea canep | A0A068UDJ3 |
| Coffea canep | A0A068VL89 |
| Coffea canep | A0A068V636 |
| Coffea canep | A0A068TTS6 |
| Coffea canep | A0A068VAU6 |
| Coffea canep | A0A068UAW8 |
| Coffea canep | A0A068TY94 |
| Coffea canep | A0A068V341 |
| Coffea canep | A0A068UDX1 |
| Coffea canep | A0A068U7P7 |
| Coffea canep | A0A068TRE6 |
| Coffea canep | A0A068U9C3 |
| Coffea canep | A0A068U9L9 |
| Coffea canep | A0A068UCP7 |
| Coffea canep | A0A068V5D0 |
| Coffea canep | A0A068U6Z2 |
| Coffea canep | A0A068UAL8 |
| Coffea canep | A0A068TUH5 |
| Coffea canep | A0A068UVZ3 |
| Cognatiyooni | A0A1I0RVR9 |
| Cohaesibacte | A0A285PJ03 |
| Cohaesibacte | A0A1I4ZNL6 |
| Cohaesibacte | A0A285M669 |
| Cohaesibacte | A0A2N5XUY7 |
| Coleophoma | A0A3D8Q658 |
| Coleophoma | A0A3D8S2L7 |
| Coleophoma | A0A3D8Q9J9 |
| Coleophoma | A0A3D8Q983 |
| Coleophoma | A0A3D8Q9G4 |
| Coleophoma | A0A3D8R5V5 |
| Coleophoma | A0A3D8STW8 |
| Coleophoma | A0A3D8Q6N2 |
| Coleophoma | A0A3D8Q4Q0 |
| Coleophoma | A0A3D8QNJ8 |
| Colinus virgin | A0A226PF24 |
| Colinus virgin | A0A226PDM2 |
| Colinus virgin | A0A226PAM3 |

| | |
|---|---|
| Colinus virgin | A0A226NT22 |
| Colius striatu | A0A091L082 |
| Colius striatu | A0A091KA09 |
| Colius striatu | A0A091K6S3 |
| Colletotrichu | A0A1Q8RWV1 |
| Colletotrichu | A0A1Q8RVP1 |
| Colletotrichu | A0A1Q8S2N5 |
| Colletotrichu | A0A1Q8S5V0 |
| Colletotrichu | A0A010QTI5 |
| Colletotrichu | A0A010SLX6 |
| Colletotrichu | A0A010RTR1 |
| Colletotrichu | L2FSS4 |
| Colletotrichu | L2GE62 |
| Colletotrichu | L2G3T0 |
| Colletotrichu | L2FT65 |
| Colletotrichu | L2FRD2 |
| Colletotrichu | L2GC03 |
| Colletotrichu | L2FYL0 |
| Colletotrichu | T0KBX5 |
| Colletotrichu | T0KBL3 |
| Colletotrichu | T0LNX5 |
| Colletotrichu | T0K3L5 |
| Colletotrichu | T0M6Z0 |
| Colletotrichu | T0K282 |
| Colletotrichu | T0KJU1 |
| Colletotrichu | T0KX95 |
| Colletotrichu | T0KV89 |
| Colletotrichu | E3QKF4 |
| Colletotrichu | E3Q9X3 |
| Colletotrichu | E3QHU7 |
| Colletotrichu | E3R0R1 |
| Colletotrichu | H1VPP1 |
| Colletotrichu | H1VE61 |
| Colletotrichu | A0A1B7XYH5 |
| Colletotrichu | H1VEI3 |
| Colletotrichu | A0A1B7XYR0 |
| Colletotrichu | A0A1B7Y121 |
| Colletotrichu | H1W3P5 |
| Colletotrichu | A0A162PMT3 |
| Colletotrichu | A0A166LHG7 |
| Colletotrichu | A0A1S1W2W6 |

| | |
|---|---|
| Colletotrichu | A0A1S1VWS2 |
| Colletotrichu | A0A162NJN8 |
| Colletotrichu | A0A162PCP5 |
| Colletotrichu | A0A135SZV4 |
| Colletotrichu | A0A135UT38 |
| Colletotrichu | A0A135T629 |
| Colletotrichu | A0A135UX69 |
| Colletotrichu | N4UZE4 |
| Colletotrichu | N4UZ75 |
| Colletotrichu | N4VFT2 |
| Colletotrichu | N4W403 |
| Colletotrichu | N4V0X7 |
| Colletotrichu | A0A1G4B117 |
| Colletotrichu | A0A1G4BBR1 |
| Colletotrichu | A0A1G4ATI1 |
| Colletotrichu | A0A135V7W1 |
| Colletotrichu | A0A135S139 |
| Colletotrichu | A0A135URM5 |
| Colletotrichu | A0A135V7C4 |
| Colletotrichu | A0A135RXN4 |
| Colletotrichu | A0A135RR28 |
| Colletotrichu | A0A135T1G0 |
| Colletotrichu | A0A135T432 |
| Colletotrichu | A0A135RPI7 |
| Colletotrichu | A0A066XMA0 |
| Colletotrichu | A0A066XY08 |
| Colletotrichu | A0A166RN27 |
| Colletotrichu | A0A166S4H7 |
| Colobus angc | A0A2K5JR89 |
| Colobus angc | A0A2K5JX41 |
| Colobus angc | A0A2K5JX37 |
| Colobus angc | A0A2K5JX55 |
| Colobus angc | A0A2K5JK04 |
| Colobus angc | A0A2K5HKG1 |
| Colobus angc | A0A2K5I747 |
| Colobus angc | A0A2K5I7F4 |
| Columba livia | A0A2I0MN33 |
| Columba livia | A0A2I0MN31 |
| Columba livia | A0A2I0MV52 |
| Columba livia | A0A2I0MV53 |
| Columba livia | A0A2I0MH63 |

| | |
|---|---|
| Columba livia | A0A2I0MN36 |
| Columba livia | A0A2I0MN41 |
| compost met | A0A3R4B871 |
| compost met | A0A407GJQ8 |
| compost met | A0A408KDV5 |
| compost met | A0A409SM37 |
| compost met | A0A3R2WYC1 |
| compost met | A0A3R4Z561 |
| compost met | A0A3R1NI64 |
| compost met | A0A405Z9C4 |
| compost met | A0A3R4NNR5 |
| Coniella lustr | A0A2T2ZVH0 |
| Coniella lustr | A0A2T2ZTN0 |
| Coniella lustr | A0A2T2ZW89 |
| Coniella lustr | A0A2T2ZSF1 |
| Coniochaeta | A0A1J7IVY8 |
| Coniochaeta | A0A420Y1M1 |
| Coniochaeta | A0A420Y084 |
| Coniosporiun | R7Z4G2 |
| Coniosporiun | R7YJ26 |
| Coniosporiun | R7YYV7 |
| Coniosporiun | R7Z2B6 |
| Coniosporiun | R7YKC6 |
| Coniosporiun | R7YYF5 |
| Consotaella s | A0A1T4MT25 |
| Corallococcus | H8MJT7 |
| Corchorus ca | A0A1R3IQG2 |
| Corchorus ca | A0A1R3JAU6 |
| Corchorus ca | A0A1R3JH31 |
| Corchorus ca | A0A1R3ICX9 |
| Corchorus ca | A0A1R3KDQ4 |
| Corchorus ca | A0A1R3IXI1 |
| Corchorus ca | A0A1R3I223 |
| Corchorus ca | A0A1R3I0S0 |
| Corchorus ca | A0A1R3JCQ7 |
| Corchorus ca | A0A1R3IHP6 |
| Corchorus ca | A0A1R3I8B4 |
| Corchorus ca | A0A1R3HWR3 |
| Corchorus ca | A0A1R3I8E3 |
| Corchorus ca | A0A1R3JH09 |
| Corchorus ca | A0A1R3IIR2 |

Corchorus ca A0A1R3I8B1
Corchorus ca A0A1R3GUC6
Corchorus ca A0A1R3G8B2
Corchorus ca A0A1R3IQM1
Corchorus ca A0A1R3GC79
Corchorus ca A0A1R3GXE2
Corchorus ca A0A1R3JH02
Corchorus ca A0A1R3K5M1
Corchorus ca A0A1R3JYE5
Corchorus ca A0A1R3IQK9
Corchorus ca A0A1R3IES9
Corchorus ca A0A1R3I214
Corchorus ca A0A1R3IQG7
Corchorus ca A0A1R3IQE6
Corchorus ca A0A1R3I248
Corchorus ca A0A1R3K9G1
Corchorus ca A0A1R3H0C3
Corchorus ca A0A1R3GS32
Corchorus ca A0A1R3J3W4
Corchorus ca A0A1R3H636
Corchorus ca A0A1R3I8F2
Corchorus ca A0A1R3FYR8
Corchorus oli A0A1R3KLM1
Corchorus oli A0A1R3K8S9
Corchorus oli A0A1R3KU50
Corchorus oli A0A1R3J5L5
Corchorus oli A0A1R3KBY3
Corchorus oli A0A1R3JYF5
Corchorus oli A0A1R3GIW4
Corchorus oli A0A1R3JDE5
Corchorus oli A0A1R3KM12
Corchorus oli A0A1R3HMG0
Corchorus oli A0A1R3I4C7
Corchorus oli A0A1R3KA03
Corchorus oli A0A1R3H436
Corchorus oli A0A1R3HD45
Corchorus oli A0A1R3KF17
Corchorus oli A0A1R3ICZ6
Corchorus oli A0A1R3JDF2
Corchorus oli A0A1R3FZ78
Corchorus oli A0A1R3K8S8

Corchorus oli A0A1R3J7N3
Corchorus oli A0A1R3H3T9
Corchorus oli A0A1R3GAX3
Corchorus oli A0A1R3IDM3
Corchorus oli A0A1R3JCH0
Corchorus oli A0A1R3H3T1
Corchorus oli A0A1R3JZ07
Corchorus oli A0A1R3KC06
Corchorus oli A0A1R3HV73
Corchorus oli A0A1R3JYS9
Corchorus oli A0A1R3KRY3
Corchorus oli A0A1R3H3Q5
Corchorus oli A0A1R3KBU9
Corchorus oli A0A1R3H4T5
Corchorus oli A0A1R3KBT7
Corchorus oli A0A1R3JYI7
Corchorus oli A0A1R3KBW6
Corchorus oli A0A1R3HEM2
Corchorus oli A0A1R3KBS9
Corchorus oli A0A1R3KBZ0
Corchorus oli A0A1R3I4A2
Corchorus oli A0A1R3H934
Corchorus oli A0A1R3K8R8
Corchorus oli A0A1R3KBZ3
Corchorus oli A0A1R3JYI0
Corchorus oli A0A1R3JYB7
Corchorus oli A0A1R3H3N1
Corchorus oli A0A1R3H913
Corchorus oli A0A1R3JDG2
Corchorus oli A0A1R3KBV2
Corchorus oli A0A1R3H8W0
Corchorus oli A0A1R3KU57
Corchorus oli A0A1R3I4C0
Corchorus oli A0A1R3H3R9
Corchorus oli A0A1R3JID2
Corchorus oli A0A1R3KBV6
Corchorus oli A0A1R3H3T2
Corchorus oli A0A1R3GSE3
Corchorus oli A0A1R3JYJ4
Corchorus oli A0A1R3KBX8
Corchorus oli A0A1R3KU24

| | |
|---|---|
| Corchorus oli | A0A1R3HPV5 |
| Corchorus oli | A0A1R3KGT6 |
| Corchorus oli | A0A1R3GJ03 |
| Corchorus oli | A0A1R3H3P6 |
| Corchorus oli | A0A1R3K9H9 |
| Corchorus oli | A0A1R3HSJ9 |
| Corchorus oli | A0A1R3KYZ9 |
| Cordyceps co | A0A162K7A9 |
| Cordyceps co | A0A168IRL7 |
| Cordyceps co | A0A179IKV7 |
| Cordyceps co | A0A179IAQ3 |
| Cordyceps fu | A0A167WMS9 |
| Cordyceps mi | G3J8R3 |
| Cordyceps mi | G3J375 |
| Cordyceps mi | G3JSZ4 |
| Cordyceps mi | G3JUF8 |
| Cordyceps mi | A0A2H4SK15 |
| Cordyceps mi | A0A2H4S7F4 |
| Cordyceps sp | A0A2C6AEQ6 |
| Cordyceps sp | A0A2C5ZXP2 |
| Corvus brach | A0A091ET11 |
| Corvus brach | A0A091EJ05 |
| Corvus brach | A0A091EWY0 |
| Corvus brach | A0A091F8A0 |
| Corvus brach | A0A091E9Y9 |
| Corynespora | A0A2T2N6K3 |
| Corynespora | A0A2T2NU07 |
| Corynespora | A0A2T2NYB4 |
| Corynespora | A0A2T2N5T0 |
| Corynespora | A0A2T2NUY0 |
| Corynespora | A0A2T2N6K9 |
| Crambe kralik | Q8LP57 |
| Crassostrea g | K1QBT9 |
| Crassostrea g | K1QP59 |
| Crassostrea g | K1Q5J6 |
| Crassostrea g | K1RL61 |
| Crassostrea g | K1PV69 |
| Crassostrea g | K1Q202 |
| Crassostrea g | K1RCS2 |
| Crassostrea g | K1R451 |
| Crassostrea g | K1QEX9 |

Crassostrea g K1RFL1
Crassostrea g K1QFL8
Crassostrea g K1R782
Crassostrea g K1PMX4
Crassostrea g K1QE58
Crassostrea g K1QC63
Crassostrea g K1Q2X3
Crassostrea g K1Q0K1
Crassostrea g K1RU37
Crassostrea g K1QRX2
Crassostrea g K1Q0F5
Crassostrea g K1PMX3
Crassostrea g K1QH76
Crassostrea g K1R7M2
Crassostrea g K1PKQ8
Crassostrea g K1QEU5
Crassostrea g K1QEV1
Crassostrea g K1Q3J0
Crassostrea g K1PJ39
Crassostrea g K1R0X1
Crassostrea g K1R8Y0
Crassostrea g K1QI12
Crassostrea g K1PUC3
Crassostrea g K1PZY8
Crassostrea g K1QNX1
Crassostrea g K1PEE6
Crassostrea g K1PQ16
Crassostrea g K1RKN1
Crassostrea g K1P151
Crassostrea g K1PCZ9
Crassostrea g K1P8Q7
Crassostrea g K1RDU0
Crassostrea g K1R0V6
Crassostrea g K1RNM0
Crassostrea g K1QGI9
Crassostrea g K1R168
Crassostrea g K1QEL7
Crassostrea g K1RBU1
Crassostrea g K1QRJ4
Crassostrea g K1R5R6
Crassostrea g K1QPK8

| | |
|---|---|
| Crassostrea g | K1QPU6 |
| Crassostrea g | K1QNB3 |
| Crassostrea g | K1RFB4 |
| Crassostrea g | K1P8F6 |
| Crassostrea g | K1QVB6 |
| Crassostrea g | K1QI33 |
| Crassostrea g | K1QDS6 |
| Crassostrea g | K1QVX2 |
| Crassostrea g | K1QKJ9 |
| Crassostrea g | K1QYN7 |
| Crassostrea g | K1R234 |
| Crassostrea g | K1Q9S5 |
| Crassostrea g | K1PXX6 |
| Crassostrea g | K1QRJ6 |
| Crassostrea g | K1RGH6 |
| Crassostrea g | K1PM36 |
| Crassostrea g | K1R3S9 |
| Crassostrea g | K1Q4S4 |
| Crassostrea g | K1R2E6 |
| Crassostrea g | K1Q9I9 |
| Crassostrea g | K1Q9I2 |
| Crassostrea g | K1QQI2 |
| Crassostrea g | K1P0S6 |
| Crassostrea g | K1RPI0 |
| Crassostrea g | K1QP97 |
| Crassostrea g | K1Q132 |
| Crassostrea g | K1PMA2 |
| Crassostrea g | K1P7M9 |
| Crassostrea g | K1PQG5 |
| Crassostrea g | K1QIV6 |
| Crassostrea g | K1PN45 |
| Crassostrea g | K1R926 |
| Crassostrea g | K1Q7G5 |
| Crassostrea g | K1QUA0 |
| Crassostrea g | K1PUE1 |
| Crassostrea g | K1PZF9 |
| Crassostrea g | K1QZS4 |
| Crassostrea g | K1QYB3 |
| Crassostrea g | K1QYH7 |
| Crassostrea g | K1PQV5 |
| Crenothrix pc | A0A1R4H3L7 |

| | |
|---|---|
| Cribrihabitan | A0A1H6XYT5 |
| Cricetulus gri | A0A3L7I648 |
| Cricetulus gri | G3HGQ6 |
| Cricetulus gri | A0A3L7IJG9 |
| Cricetulus gri | A0A3L7IG64 |
| Cricetulus gri | A0A061IMW5 |
| Cricetulus gri | A0A3L7HP47 |
| Cricetulus gri | A0A3L7IIX0 |
| Cricetulus gri | A0A061IRF7 |
| Cricetulus gri | G3HPQ5 |
| Cryptosporid | A0A1J4MK57 |
| Cryptosporid | A0A1J4MVW0 |
| Cryptosporid | A0A1J4MU07 |
| Cryptosporid | A0A1J4MTL5 |
| Cryptosporid | A0A0S4TCD4 |
| Cryptosporid | A0A0S4TBC4 |
| Cryptosporid | A0A0S4TAQ1 |
| Cryptosporid | A0A0S4TDE6 |
| Cryptosporid | A0A2P4YWR1 |
| Cryptosporid | A0A2P4Z358 |
| Cryptosporid | A0A2P4Z5F0 |
| Cryptosporid | A0A2P4YWR9 |
| Cryptosporid | B6AEN1 |
| Cryptosporid | B6AJC3 |
| Cryptosporid | B6A964 |
| Cryptosporid | Q5CSA4 |
| Cryptosporid | Q5CTT8 |
| Cryptosporid | Q5CSA0 |
| Cryptosporid | Q5CV27 |
| Cryptosporid | Q86PQ7 |
| Cryptosporid | A0A1J4MKT5 |
| Cryptosporid | A0A1J4MIN1 |
| Cryptosporid | A0A1J4MFD1 |
| Cryptotermes | A0A2J7PDR3 |
| Cryptotermes | A0A2J7QPV9 |
| Cryptotermes | A0A2J7Q2V5 |
| Cryptotermes | A0A2J7R433 |
| Cryptotermes | A0A2J7R431 |
| Cryptotermes | A0A2J7PHG6 |
| Cryptotermes | A0A2J7QD69 |
| Cryptotermes | A0A2J7R420 |

| | |
|---|---|
| Cryptotermes | A0A2J7PDS5 |
| Cryptotermes | A0A2J7QD68 |
| Cryptotermes | A0A2J7QD73 |
| Cryptotermes | A0A2J7R430 |
| Cryptotermes | A0A2J7RS16 |
| Cryptotermes | A0A2J7R424 |
| Cryptotermes | A0A2J7PWJ5 |
| Cryptotermes | A0A2J7R441 |
| Cryptotermes | A0A2J7R451 |
| Cryptotermes | A0A2J7R454 |
| Ctenopharyn | W8FRI5 |
| Cuculus cano | A0A091FW85 |
| Cuculus cano | A0A091GEV6 |
| Cuculus cano | A0A091GKK8 |
| Cuculus cano | A0A091GI80 |
| Cucumis melo | A0A1S4DST8 |
| Cucumis melo | A0A1S3B409 |
| Cucumis melo | A0A1S3B3R3 |
| Cucumis melo | A0A1S4DSQ5 |
| Cucumis melo | A0A1S4DSX1 |
| Cucumis melo | A0A1S3B440 |
| Cucumis melo | A0A1S3B585 |
| Cucumis melo | A0A1S3B4W7 |
| Cucumis melo | A0A1S3CSP7 |
| Cucumis melo | A0A1S3AZX8 |
| Cucumis melo | A0A1S3AZY6 |
| Cucumis melo | A0A1S3BZ12 |
| Cucumis melo | A0A1S4DSR7 |
| Cucumis melo | A0A1S4DSP8 |
| Cucumis melo | A0A1S4E4E3 |
| Cucumis melo | A0A1S3B4H2 |
| Cucumis melo | A0A1S3B453 |
| Cucumis melo | A0A1S3B3R8 |
| Cucumis melo | A0A1S3B5C7 |
| Cucumis melo | A0A1S4DTJ6 |
| Cucumis melo | A0A1S4DSS7 |
| Cucumis melo | A0A1S3B4D0 |
| Cucumis melo | A0A1S3B536 |
| Cucumis melo | A0A1S3B5U5 |
| Cucumis melo | A0A1S3B6L4 |
| Cucumis melo | A0A1S3B4A8 |

| | |
|---|---|
| Cucumis melo | A0A1S3BTL3 |
| Cucumis melo | A0A1S4DST5 |
| Cucumis melo | A0A1S3B4B3 |
| Cucumis melo | A0A1S3B516 |
| Cucumis melo | A0A1S3B4F1 |
| Cucumis melo | A0A1S4DZ25 |
| Cucumis melo | A0A1S3C0Q9 |
| Cucumis melo | A0A1S4DSE8 |
| Cucumis melo | A0A1S3CSN1 |
| Cucumis melo | A0A1S3BC50 |
| Cucumis melo | A0A1S4DSS6 |
| Cucumis melo | A0A1S3B7B8 |
| Cucumis melo | A0A1S3CF32 |
| Cucumis melo | A0A1S3C207 |
| Cucumis melo | A0A1S3B530 |
| Cucumis melo | A0A1S3BTI5 |
| Cucumis melo | A0A1S4DSE6 |
| Cucumis melo | A0A1S4DSS8 |
| Cucumis melo | A0A1S3CR70 |
| Cucumis melo | A0A1S3B575 |
| Cucumis sativ | A0A0A0LS73 |
| Cucumis sativ | A0A0A0LS68 |
| Cucumis sativ | A0A0A0LUY8 |
| Cucumis sativ | A0A0A0LX93 |
| Cucumis sativ | A0A0A0LUE6 |
| Cucumis sativ | A0A0A0LRR4 |
| Cucumis sativ | A0A0A0KRF4 |
| Cucumis sativ | A0A0A0K287 |
| Cucumis sativ | A0A0A0LEC1 |
| Cucumis sativ | A0A0A0LUZ3 |
| Cucumis sativ | A0A0A0LDL3 |
| Cucumis sativ | A0A0A0LUF1 |
| Cucumis sativ | A0A0A0LAF2 |
| Cucumis sativ | A0A0A0LRR1 |
| Cucumis sativ | A0A0A0KK10 |
| Cucumis sativ | A0A0A0LBA0 |
| Cucumis sativ | A0A0A0LX89 |
| Cucumis sativ | A0A0A0KC43 |
| Cucumis sativ | A0A0A0L6H7 |
| Cucumis sativ | A0A0A0K3N8 |
| Cucumis sativ | A0A0A0LUZ5 |

| | |
|---|---|
| Cucumis sativ | A0A0A0LFV8 |
| Cucumis sativ | A0A0A0LUY4 |
| Cucumis sativ | A0A0A0LII6 |
| Cucumis sativ | A0A0A0LXA0 |
| Cucumis sativ | A0A0A0LD19 |
| Cucumis sativ | A0A0A0LX86 |
| Cucumis sativ | A0A0A0KC37 |
| Cucumis sativ | A0A0A0LRR8 |
| Cucumis sativ | A0A0A0K8W5 |
| Cucumis sativ | A0A0A0L7U1 |
| Cucumis sativ | A0A0A0LRS1 |
| Cucumis sativ | A0A0A0KWZ1 |
| Cucumis sativ | A0A0A0LS76 |
| Cucumis sativ | A0A0A0LUD8 |
| Cucumis sativ | A0A0A0KSR7 |
| Cucumis sativ | A0A0A0LZJ7 |
| Cucumis sativ | A0A0A0K7B0 |
| Cuerna arida. | A0A1B6H4Y5 |
| Cuerna arida. | A0A1B6G4G8 |
| Cuerna arida. | A0A1B6EYW8 |
| Cuerna arida. | A0A1B6FLH2 |
| Cuerna arida. | A0A1B6FYJ9 |
| Cuerna arida. | A0A1B6GHV9 |
| Cuerna arida. | A0A1B6H2T7 |
| Cuerna arida. | A0A1B6FML8 |
| Cuerna arida. | A0A1B6GMC1 |
| Cuerna arida. | A0A1B6FMH3 |
| Cuerna arida. | A0A1B6FYA0 |
| Cuerna arida. | A0A1B6H1R8 |
| Culex quinqu | B0VZB8 |
| Culex quinqu | B0WCW6 |
| Culex quinqu | B0XHR5 |
| Culex quinqu | B0X861 |
| Culex quinqu | B0WCW5 |
| Culex quinqu | B0X859 |
| Culex quinqu | B0WBN2 |
| Culex quinqu | B0W214 |
| Culex quinqu | A0A1S4J279 |
| Culex quinqu | A0A1S4JBK4 |
| Culex quinqu | B0WCW7 |
| Culex tarsalis | A0A1Q3FSK0 |

| | |
|---|---|
| Culex tarsalis | A0A1Q3FS43 |
| Culicoides so | A0A336M006 |
| Culicoides so | A0A336LTK1 |
| Culicoides so | A0A336LRS2 |
| Culicoides so | A0A336N522 |
| Culicoides so | A0A336LVN5 |
| Culicoides so | A0A336MHF2 |
| Culicoides so | A0A336MGD3 |
| Culicoides so | A0A336MI32 |
| Culicoides so | A0A336MYJ1 |
| Culicoides so | A0A336LUN7 |
| Culicoides so | A0A336MT58 |
| Culicoides so | A0A336MRX6 |
| Culicoides so | A0A336LTL3 |
| Culicoides so | A0A336LMK2 |
| Cuscuta austr | A0A328DVR3 |
| Cuscuta austr | A0A328D129 |
| Cuscuta austr | A0A328D3T7 |
| Cuscuta austr | A0A328D453 |
| Cuscuta austr | A0A328DBU8 |
| Cuscuta austr | A0A328D973 |
| Cuscuta austr | A0A328DIK3 |
| Cuscuta austr | A0A328DIS6 |
| Cuspidothrix | A0A2S6CT90 |
| Cyanosarcina | A0A2T1F9F0 |
| Cyanothece s | A0A3B8Y4A2 |
| Cyclospora ca | A0A1D3CT75 |
| Cyclospora ca | A0A1D3D2G3 |
| Cyclospora ca | A0A1D3CZG8 |
| Cyclospora ca | A0A1D3D562 |
| Cyclospora ca | A0A1D3CXV2 |
| Cyclospora ca | A0A1D3CZF4 |
| Cyclospora ca | A0A1D3D4P8 |
| Cyclospora ca | A0A1D3D927 |
| Cyclospora ca | A0A1D3D792 |
| Cyclospora ca | A0A1D3CRX6 |
| Cyclospora ca | A0A1D3CZB5 |
| Cyclospora ca | A0A1D3CRI2 |
| Cyclospora ca | A0A1D3D067 |
| Cylicostephar | A0A3P6QZS2 |
| Cylicostephar | A0A3P6QGH9 |

| | |
|---|---|
| Cylicostephar | A0A3P6T344 |
| Cylicostephar | A0A3P6R246 |
| Cylicostephar | A0A3P6S9T1 |
| Cynara cardu | A0A103YIH9 |
| Cynara cardu | A0A124PXP6 |
| Cynara cardu | A0A103XS92 |
| Cynara cardu | A0A103XY72 |
| Cynara cardu | A0A103Y5E6 |
| Cynara cardu | A0A103XZ10 |
| Cynara cardu | A0A103XC99 |
| Cynara cardu | A0A124S064 |
| Cynara cardu | A0A103Y815 |
| Cynara cardu | A0A103XCM9 |
| Cynara cardu | A0A118JT41 |
| Cynara cardu | A0A118JSJ8 |
| Cynara cardu | A0A103XF71 |
| Cynara cardu | A0A118JU86 |
| Cynara cardu | A0A124SCT9 |
| Cynara cardu | A0A124SHJ7 |
| Cynara cardu | A0A103YJR9 |
| Cynara cardu | A0A103XQ49 |
| Cynara cardu | A0A118K316 |
| Cynara cardu | A0A103XC93 |
| Cynara cardu | A0A103XYY7 |
| Cynara cardu | A0A103Y467 |
| Cynara cardu | A0A118K2V5 |
| Cynara cardu | A0A124SHS0 |
| Cynara cardu | A0A103Y6D6 |
| Cynara cardu | A0A118HIH7 |
| Cynara cardu | A0A103XU24 |
| Cynara cardu | A0A103Y212 |
| Cynara cardu | A0A103Y1Z5 |
| Cynara cardu | A0A103YD45 |
| Cynara cardu | A0A103YJV2 |
| Cynara cardu | A0A103Y7Z6 |
| Cynara cardu | A0A124SHR9 |
| Cynara cardu | A0A103XQ54 |
| Cynara cardu | A0A124SDG2 |
| Cynara cardu | A0A103XWT3 |
| Cynara cardu | A0A103YIJ1 |
| Cynara cardu | A0A124SHK2 |

| | |
|---|---|
| Cynara cardu | A0A103XF58 |
| Cynara cardu | A0A118GC55 |
| Cynara cardu | A0A103XQI2 |
| Cynara cardu | A0A103Y7Z7 |
| Cynara cardu | A0A103XBG4 |
| Cynara cardu | A0A103XU62 |
| Cynara cardu | A0A118JTB4 |
| Cynara cardu | A0A103XZ33 |
| Cynara cardu | A0A103MM20 |
| Cynara cardu | A0A103Y9H4 |
| Cynara cardu | A0A103Y652 |
| Cynara cardu | A0A103Y6D3 |
| Cynara cardu | A0A118JTV8 |
| Cynara cardu | A0A103YDC6 |
| Cynara cardu | A0A103Y6C0 |
| Cynara cardu | A0A103VJ56 |
| Cynara cardu | A0A103Y9I0 |
| Cynara cardu | A0A103Y446 |
| Cynara cardu | A0A103XWW8 |
| Cynara cardu | A0A103YCS4 |
| Cynara cardu | A0A103XIZ2 |
| Cynara cardu | A0A103XDQ3 |
| Cynoglossus s | A0A3P8VG89 |
| Cynoglossus s | A0A3P8UTK9 |
| Cynoglossus s | A0A3P8VDI4 |
| Cynoglossus s | A0A3P8X2L6 |
| Cynoglossus s | A0A3P8X460 |
| Cynoglossus s | A0A3P8V360 |
| Cynoglossus s | A0A3P8VG76 |
| Cynoglossus s | A0A3P8X4M6 |
| Cyphellophor | W2RU83 |
| Cyphomyrme | A0A195CFB6 |
| Cyphomyrme | A0A151INZ7 |
| Cyphomyrme | A0A195CUA0 |
| Cyphomyrme | A0A195CJ60 |
| Cyphomyrme | A0A151IBX2 |
| Cyphomyrme | A0A195CDF0 |
| Cyprinodon v | A0A3Q2D697 |
| Cyprinodon v | A0A3Q2FWU5 |
| Cyprinodon v | A0A3Q2D220 |
| Cyprinodon v | A0A3Q2FWW5 |

| | |
|---|---|
| Cyprinodon v | A0A3Q2E8M8 |
| Cyprinodon v | A0A3Q2D1Q5 |
| Cyprinodon v | A0A3Q2D1Y0 |
| Cyprinodon v | A0A3Q2E737 |
| Cyprinodon v | A0A3Q2D1Q1 |
| Cyprinodon v | A0A3Q2FR25 |
| Cyprinodon v | A0A3Q2DW28 |
| Cyprinus carp | A0A0U2S2V6 |
| Cystobacter f | A0A250JIW8 |
| Cystoisospora | A0A2C6KJH5 |
| Cystoisospora | A0A2C6KQQ7 |
| Cystoisospora | A0A2C6KGJ2 |
| Cystoisospora | A0A2C6KJZ8 |
| Cystoisospora | A0A2C6KRS7 |
| Cystoisospora | A0A2C6KBF6 |
| Cystoisospora | A0A2C6KGB3 |
| Cystoisospora | A0A2C6L9R7 |
| Cystoisospora | A0A2C6KLD1 |
| Cystoisospora | A0A2C6LDQ6 |
| Cystoisospora | A0A2C6KWX7 |
| Cystoisospora | A0A2C6KJN3 |
| Cystoisospora | A0A2C6KC22 |
| Cystoisospora | A0A2C6KHJ6 |
| Cystoisospora | A0A2C6KXQ4 |
| Cystoisospora | A0A2C6LH90 |
| Cystoisospora | A0A2C6KQM3 |
| Cystoisospora | A0A2C6KV00 |
| Cystoisospora | A0A2C6KD09 |
| Cystoisospora | A0A2C6KJE6 |
| Cystoisospora | A0A2C6KGE6 |
| Cystoisospora | A0A2C6KQL0 |
| Cystoisospora | A0A2C6KFU7 |
| Cystoisospora | A0A2C6KW92 |
| Cystoisospora | A0A2C6KGC7 |
| Cystoisospora | A0A2C6KGB5 |
| Cystoisospora | A0A2C6L6S4 |
| Cystoisospora | A0A2C6L490 |
| Cytospora leu | A0A423VS70 |
| Cytospora leu | A0A423W0P4 |
| Cytospora leu | A0A423WTP3 |
| Cytospora leu | A0A423X084 |

| | |
|---|---|
| Daboia russel | A0A223PK27 |
| Dactylellina h | S8C318 |
| Dactylellina h | S8ASF1 |
| Dactylellina h | S8BMH2 |
| Dactylellina h | S8A7Q6 |
| Dactylellina h | S8A4U1 |
| Dactylellina h | S8BSD4 |
| Daldinia sp. E | A0A1Y2XBG1 |
| Daldinia sp. E | A0A1Y2WWT8 |
| Daldinia sp. E | A0A1Y2XCF1 |
| Daldinia sp. E | A0A1Y2WXC0 |
| Danaus plexi | A0A212F9C1 |
| Danaus plexi | A0A212FKU3 |
| Danaus plexi | A0A212FKV4 |
| Danaus plexi | A0A212F3I1 |
| Danaus plexi | A0A212EVI3 |
| Danaus plexi | A0A212FAB7 |
| Danio rerio.xl | Q6JAN0 |
| Danio rerio.xl | F1REK1 |
| Danio rerio.xl | F1Q890 |
| Danio rerio.xl | F6PA78 |
| Danio rerio.xl | A4IGA7 |
| Danio rerio.xl | Q5XFY1 |
| Danio rerio.xl | Q90ZN6 |
| Danio rerio.xl | E9QGF2 |
| Danio rerio.xl | Q5RGG3 |
| Danio rerio.xl | Q6PBA6 |
| Danio rerio.xl | F1QEZ1 |
| Daphnia mag | A0A0P5WPN5 |
| Daphnia mag | A0A0P5HGT8 |
| Daphnia mag | A0A0P5T1U8 |
| Daphnia mag | A0A0N8E7S9 |
| Daphnia mag | A0A0P6AL09 |
| Daphnia mag | A0A0P5Y462 |
| Daphnia mag | A0A0P5L4W8 |
| Daphnia mag | A0A0P5S420 |
| Daphnia mag | A0A0P5TKN5 |
| Daphnia mag | A0A0P6A674 |
| Daphnia mag | A0A0P5YJR7 |
| Daphnia mag | A0A0P5BMJ7 |
| Daphnia mag | A0A0P5U7V1 |

Daphnia mag A0A0P6C7R9
Daphnia mag A0A0P5M729
Daphnia mag A0A0P5DS66
Daphnia mag A0A0P5C4X4
Daphnia mag A0A0P5D7A2
Daphnia mag A0A0P6ATK6
Daphnia mag A0A0P5EBG6
Daphnia mag A0A0P5XUR1
Daphnia mag A0A0P6CTV5
Daphnia mag A0A0P5NLP7
Daphnia mag A0A0P5E9S4
Daphnia mag A0A0P5JUZ4
Daphnia mag A0A0P6CI84
Daphnia mag A0A0P6C183
Daphnia mag A0A0P5TSU1
Daphnia mag A0A0P6J986
Daphnia mag A0A0P5TJC9
Daphnia mag A0A0P5ZKH9
Daphnia mag A0A0N8CLP0
Daphnia mag A0A0P5BT62
Daphnia mag A0A0P6CKZ9
Daphnia mag A0A0P5RCJ4
Daphnia mag A0A0P6HJG1
Daphnia mag A0A0P4XKS6
Daphnia mag A0A0P5XUU9
Daphnia mag A0A0P5LNA4
Daphnia mag A0A0P5NMM3
Daphnia mag A0A0P6D731
Daphnia mag A0A0N8BA13
Daphnia mag A0A0P6B472
Daphnia mag A0A0N7ZWW0
Daphnia mag A0A0N8DMK8
Daphnia mag A0A0P5S7M2
Daphnia mag A0A0P5CWX9
Daphnia mag A0A0P5NCC6
Daphnia mag A0A0P6GJS2
Daphnia mag A0A0P5PR06
Daphnia mag A0A0P6H6H4
Daphnia mag A0A0P6GE29
Daphnia mag A0A0N8EJE4
Daphnia mag A0A0P5DGB0

Daphnia mag A0A0P5XV46
Daphnia mag A0A0P5F055
Daphnia mag A0A0P6I548
Daphnia mag A0A0P6IIP3
Daphnia mag A0A0P5T6R7
Daphnia mag A0A0P6HQE1
Daphnia mag A0A0P4YGM2
Daphnia mag A0A0P5BIZ1
Daphnia mag A0A0N8DGI8
Daphnia mag A0A0N8CBW6
Daphnia mag A0A0P5DBP6
Daphnia mag A0A0P5HLY6
Daphnia mag A0A0P5T4T8
Daphnia mag A0A0P5TZY1
Daphnia mag A0A0P6E4B8
Daphnia mag A0A0P6CID3
Daphnia mag A0A0P5REG3
Daphnia mag A0A0N8CB13
Daphnia mag A0A0N8CAL2
Daphnia mag A0A0P5SEK0
Daphnia mag A0A0N8AN73
Daphnia mag A0A0P5VJT5
Daphnia mag A0A0P5H9B6
Daphnia mag A0A0P4ZH74
Daphnia mag A0A0P5HXM7
Daphnia mag A0A0P6FTT5
Daphnia mag A0A0P5VB25
Daphnia mag A0A0P5Q0M1
Daphnia mag A0A0P5PUQ2
Daphnia mag A0A0P5Y2H7
Daphnia mag A0A0P5CZZ0
Daphnia mag A0A0P5BX45
Daphnia mag A0A0P5N855
Daphnia mag A0A0P5HF26
Daphnia mag A0A0P5K8Q2
Daphnia mag A0A0N8AQK2
Daphnia mag A0A0P5KWZ5
Daphnia mag A0A0P5TVC1
Daphnia mag A0A0P6GSX1
Daphnia mag A0A162QLH9
Daphnia mag A0A0P5WUS7

Daphnia mag A0A0P5ID96
Daphnia mag A0A0P5F0V1
Daphnia mag A0A0P5TUD8
Daphnia mag A0A0P5JTA5
Daphnia mag A0A0P5GZR4
Daphnia mag A0A0P6G736
Daphnia mag A0A0P5RD98
Daphnia mag A0A0P5Y4G8
Daphnia mag A0A0P6DIA9
Daphnia mag A0A0P4Z167
Daphnia mag A0A0N8DVY1
Daphnia mag A0A0P5ZUY7
Daphnia mag A0A0P4XS98
Daphnia mag A0A0P5WA39
Daphnia mag A0A0P5LT30
Daphnia mag A0A0P5PEL1
Daphnia mag A0A0P5YIS8
Daphnia mag A0A0P6J1P8
Daphnia mag A0A0P4XVA4
Daphnia mag A0A0P5GQ84
Daphnia mag A0A0P5VSK6
Daphnia mag A0A0N8E8I5
Daphnia mag A0A0P5Q8K8
Daphnia mag A0A0P4YLV2
Daphnia mag A0A0P5CFI1
Daphnia mag A0A0P5FCE1
Daphnia mag A0A0P5J8K4
Daphnia mag A0A0P5N364
Daphnia mag A0A0P5LSH1
Daphnia mag A0A0P5JKT7
Daphnia mag A0A0P5T0H1
Daphnia mag A0A0P5YD20
Daphnia mag A0A0P4XEQ9
Daphnia mag A0A0P5D8R1
Daphnia mag A0A0P5HZD6
Daphnia mag A0A0P5BFL0
Daphnia mag A0A0P5X5U9
Daphnia mag A0A0P5NCN9
Daphnia mag A0A0P4ZBT1
Daphnia mag A0A0P5C3E7
Daphnia mag A0A0P5J8A5

Daphnia mag A0A0P5N689
Daphnia mag A0A0N8A5F9
Daphnia mag A0A0P5U0B1
Daphnia mag A0A0P4XKJ3
Daphnia mag A0A0N8A4V3
Daphnia mag A0A0P5XZ51
Daphnia mag A0A0P4YXW8
Daphnia mag A0A0P5AIQ7
Daphnia mag A0A0P6GA05
Daphnia mag A0A0P6CWH0
Daphnia mag A0A0P5VVF0
Daphnia mag A0A0N8BSP2
Daphnia mag A0A0P4XN82
Daphnia mag A0A0P5AEQ6
Daphnia mag A0A0P4ZZ55
Daphnia mag A0A0P5EPQ8
Daphnia mag A0A0P4ZPB2
Daphnia mag A0A0N8ATY9
Daphnia mag A0A0P5TTT7
Daphnia mag A0A0P5RHI5
Daphnia mag A0A0P5UDV1
Daphnia mag A0A0N8EHV2
Daphnia mag A0A0P6DW53
Daphnia mag A0A0P4Y8F4
Daphnia mag A0A0P5CPG0
Daphnia mag A0A0P5SGL3
Daphnia mag A0A0P4YDX7
Daphnia mag A0A0P5C3L8
Daphnia mag A0A0P5HRA4
Daphnia mag A0A0P5QFT9
Daphnia mag A0A0P6EVG4
Daphnia mag A0A0P6AEB8
Daphnia mag A0A0P6GFW0
Daphnia mag A0A0P5BTL7
Daphnia mag A0A0N8E1L0
Daphnia mag A0A0P5YUR1
Daphnia mag A0A0P4YWK0
Daphnia mag A0A0P5PES0
Daphnia mag A0A162RSS1
Daphnia mag A0A0N7ZQ97
Daphnia mag A0A0P5H5A9

Daphnia mag A0A0P5W725
Daphnia mag A0A0P5PKS1
Daphnia mag A0A0P6IBH1
Daphnia mag A0A0P5KMN8
Daphnia mag A0A0P5N3S5
Daphnia mag A0A0N8EJ07
Daphnia mag A0A0P5MQE4
Daphnia mag A0A0P5PNX2
Daphnia mag A0A0P5RVR9
Daphnia mag A0A0P6I9Y3
Daphnia mag A0A0P6APJ6
Daphnia mag A0A0P6CI42
Daphnia mag A0A0P5AX29
Daphnia mag A0A0P6BQC9
Daphnia mag A0A0P6E3F3
Daphnia mag A0A0P6CUC0
Daphnia mag A0A0P5RW36
Daphnia mag A0A0N7ZLQ2
Daphnia mag A0A0P5TUC3
Daphnia mag A0A164NEQ9
Daphnia mag A0A0P5YU72
Daphnia mag A0A0P5MRU3
Daphnia mag A0A0P4XHH6
Daphnia mag A0A0P5SP99
Daphnia mag A0A0P5EW58
Daphnia mag A0A0P5LF22
Daphnia mag A0A0P5QI00
Daphnia mag A0A165ADR2
Daphnia mag A0A0N8BQK6
Daphnia mag A0A0P5N9W7
Daphnia mag A0A0P5CL62
Daphnia mag A0A0P5X9Q5
Daphnia mag A0A0P5NF07
Daphnia mag A0A0P5NQH2
Daphnia mag A0A0N8ACN6
Daphnia mag A0A0P5UDC3
Daphnia mag A0A0P5IA39
Daphnia mag A0A0P5YW13
Daphnia mag A0A0P5D6K9
Daphnia mag A0A0P5WU76
Daphnia mag A0A0P5ADU9

Daphnia mag A0A0P5HLP5
Daphnia mag A0A0P5PX75
Daphnia mag A0A0P5WMQ3
Daphnia mag A0A0P5WK98
Daphnia mag A0A0P6DWY3
Daphnia mag A0A0N8CYE6
Daphnia mag A0A0N8DAF5
Daphnia mag A0A0P5U6W6
Daphnia mag A0A0P5UHU0
Daphnia mag A0A0P5P0S3
Daphnia mag A0A164WWC9
Daphnia mag A0A0P5TQ92
Daphnia mag A0A0P5YKC6
Daphnia mag A0A0P5ECY0
Daphnia mag A0A0N8DNH8
Daphnia mag A0A0P6EUA0
Daphnia mag A0A0P5RUF3
Daphnia mag A0A0P5YP87
Daphnia mag A0A0P4XUA1
Daphnia mag A0A0P6FIA5
Daphnia mag A0A0P5TWW9
Daphnia mag A0A0N8BSD6
Daphnia mag A0A0P4Z326
Daphnia mag A0A0P5B454
Daphnia mag A0A164XEY4
Daphnia mag A0A0P5WQJ7
Daphnia mag A0A0P4YXX0
Daphnia mag A0A0P5RVL4
Daphnia mag A0A0N8AUG9
Daphnia mag A0A0P5ZPM0
Daphnia mag A0A0P4YEX3
Daphnia mag A0A0P5BTY4
Daphnia mag A0A0P6CM30
Daphnia mag A0A0P4WR04
Daphnia mag A0A0P5BT21
Daphnia mag A0A0P6GTG9
Daphnia mag A0A0P6H7K9
Daphnia mag A0A0P5DXB5
Daphnia mag A0A0P5D1S4
Daphnia mag A0A0P4Y9G9
Daphnia mag A0A164ZHL4

Daphnia mag A0A0P6ETE1
Daphnia mag A0A0P5ZR54
Daphnia mag A0A0P5G8Z5
Daphnia mag A0A0P5PKW0
Daphnia mag A0A0P5DCI3
Daphnia mag A0A0P6DUY5
Daphnia mag A0A0P5LVB4
Daphnia mag A0A0P6I8Z3
Daphnia mag A0A0P5QEZ7
Daphnia mag A0A0P5CK22
Daphnia mag A0A0P5U9K2
Daphnia mag A0A0P5E2Z0
Daphnia mag A0A0P4YWG8
Daphnia mag A0A164VXV0
Daphnia mag A0A0P5N8V1
Daphnia mag A0A0P5YVP4
Daphnia mag A0A0P4WYW8
Daphnia mag A0A0P4Y5I5
Daphnia mag A0A164XQP6
Daphnia mag A0A0P6HVX9
Daphnia mag A0A0P5UDG3
Daphnia mag A0A0P6DTI4
Daphnia mag A0A0N8AR21
Daphnia mag A0A0P5HVL3
Daphnia mag A0A0P4ZTX3
Daphnia mag A0A0P5NV81
Daphnia mag A0A0N8AXC3
Daphnia mag A0A0N8BWQ2
Daphnia mag A0A0P4Y6W6
Daphnia mag A0A0P5CI46
Daphnia mag A0A0P5GCD4
Daphnia mag A0A0P5DAL0
Daphnia mag A0A0N8DA56
Daphnia mag A0A0P6AXE9
Daphnia mag A0A0P5Q6F7
Daphnia mag A0A0P5Q4A7
Daphnia mag A0A0P5ARZ3
Daphnia mag A0A0P5IZW7
Daphnia mag A0A0P5V3V8
Daphnia mag A0A0P5N7S0
Daphnia mag A0A0N8A3D1

Daphnia mag A0A0P5WMW8
Daphnia mag A0A0P5IZ39
Daphnia mag A0A0P5MIM1
Daphnia mag A0A0P5RMT5
Daphnia mag A0A0P5DZF1
Daphnia mag A0A0P5LWH8
Daphnia mag A0A0P5C3K9
Daphnia mag A0A0N8CEH7
Daphnia mag A0A0P5DSL3
Daphnia mag A0A0P6EMY0
Daphnia mag A0A0P5YSJ8
Daphnia mag A0A0P5WJZ4
Daphnia mag A0A0P5WSP3
Daphnia mag A0A0P6F1X9
Daphnia mag A0A0N8AZ22
Daphnia mag A0A0P6IMA8
Daphnia mag A0A0P5E5X7
Daphnia mag A0A0P6EFJ2
Daphnia mag A0A0P5UL87
Daphnia mag A0A0P4Y7F0
Daphnia mag A0A0P5HFI7
Daphnia mag A0A0P5DIM6
Daphnia mag A0A0P5PZV2
Daphnia mag A0A0N8BZ24
Daphnia mag A0A0P6IB26
Daphnia mag A0A0N8EFF2
Daphnia mag A0A0P5WLD1
Daphnia mag A0A0P5U1N0
Daphnia mag A0A0P5NZP1
Daphnia mag A0A0N8BVC5
Daphnia mag A0A0P5LU81
Daphnia mag A0A0P4Z4U7
Daphnia mag A0A0P5TAQ3
Daphnia mag A0A0P5R736
Daphnia mag A0A162RDZ5
Daphnia mag A0A0P5G8N3
Daphnia mag A0A0N8B914
Daphnia mag A0A0N8CTU0
Daphnia mag A0A0P5W8C4
Daphnia mag A0A0P6I7V1
Daphnia mag A0A0P5Y7U6

Daphnia mag A0A0P6GKX2
Daphnia mag A0A0P6CI97
Daphnia mag A0A0P6HZU7
Daphnia mag A0A0P5YP46
Daphnia mag A0A0P5MTF8
Daphnia mag A0A0P5W4R4
Daphnia mag A0A0P6EJL6
Daphnia mag A0A0P5WKU1
Daphnia mag A0A0P5WJY7
Daphnia mag A0A0P5Y923
Daphnia mag A0A0P6CL50
Daphnia mag A0A0P5YP12
Daphnia mag A0A0P6JTG3
Daphnia mag A0A0N8AZB0
Daphnia mag A0A0P5RFE3
Daphnia mag A0A0P5MR11
Daphnia mag A0A0P5T4E7
Daphnia pule E9FZF1
Daphnia pule E9GUF9
Daphnia pule E9HG92
Daphnia pule E9H7K0
Daphnia pule E9FZT3
Daphnia pule E9H7J5
Daphnia pule E9GC79
Daphnia pule E9HGF3
Daphnia pule E9GXM8
Daphnia pule E9HXN3
Daphnia pule E9HL86
Daphnia pule E9HHQ3
Daphnia pule E9GB65
Daphnia pule E9GDU0
Daphnia pule E9FZB7
Daphnia pule E9FXU0
Daphnia pule E9HD48
Daphnia pule E9HD47
Daphnia pule E9GMC8
Daphnia pule E9GGB7
Daucus carot A0A164VCM7
Daucus carot A0A161YC10
Daucus carot A0A166G9N2
Daucus carot A0A161X5W6

Daucus carot A0A166F2H9
Daucus carot A0A166IMD7
Daucus carot A0A161ZJQ8
Daucus carot A0A165ZEG6
Daucus carot A0A164SJH2
Daucus carot A0A162AHQ4
Daucus carot A0A175YMP0
Daucus carot A0A165Z528
Daucus carot A0A166EUZ9
Daucus carot A0A175YJR0
Daucus carot A0A175YJB0
Daucus carot A0A166HZ24
Daucus carot A0A162AF38
Daucus carot A0A165WAR5
Daucus carot A0A164ZED3
Daucus carot A0A164X3K8
Daucus carot A0A165ZWM7
Daucus carot A0A164T5C8
Daucus carot A0A166GYV2
Daucus carot A0A164ZG59
Daucus carot A0A162ADM0
Daucus carot A0A166IUK9
Daucus carot A0A175YL17
Daucus carot A0A162AE67
Daucus carot A0A164Y320
Daucus carot A0A164TNL8
Daucus carot A0A175YLD1
Daucus carot A0A166IME5
Daucus carot A0A166AH33
Daucus carot A0A166I4U3
Daucus carot A0A164TNM5
Daucus carot A0A165Z541
Daucus carot A0A164SJF4
Daucus carot A0A165WAT6
Daucus carot A0A175YLF9
Daucus carot A0A166GYR4
Daucus carot A0A175YLB5
Daucus carot A0A175YLR0
Daucus carot A0A166IUK0
Daucus carot A0A175YLW6
Daucus carot A0A161ZVN8

Daucus carot A0A161ZTW6
Daucus carot A0A166GYN3
Daucus carot A0A166GYM4
Daucus carot A0A161ZKU6
Daucus carot A0A175YNJ8
Daucus carot A0A164TNT1
Daucus carot A0A166GV23
Daucus carot A0A164VUU7
Daucus carot A0A175YQ77
Daucus carot A0A166G9M3
Daucus carot A0A175YK13
Daucus carot A0A166GBE6
Daucus carot A0A162ADK4
Daucus carot A0A166I4V5
Daucus carot A0A175YNL1
Daucus carot A0A175YN80
Daucus carot A0A162B3S5
Daucus carot A0A162AD34
Daucus carot A0A175YKE6
Daucus carot A0A165Z6J7
Daucus carot A0A164W8A9
Daucus carot A0A175YLN5
Daucus carot A0A166AAK1
Daucus carot A0A164SJM1
Daucus carot A0A164ZI33
Daucus carot A0A165ZED0
Daucus carot A0A166JCX2
Daucus carot A0A166F2G3
Daucus carot A0A175YMV8
Daucus carot A0A166AH48
Daucus carot A0A175YM77
Daucus carot A0A166F590
Daucus carot A0A175YN69
Daucus carot A0A165ZEF7
Daucus carot A0A161ZHE5
Daucus carot A0A165Z522
Daucus carot A0A161Y240
Daucus carot A0A166IX90
Daucus carot A0A166CK63
Daucus carot A0A166AH68
Daucus carot A0A175YM10

| | |
|---|---|
| Daucus carot | A0A164WIW8 |
| Daucus carot | A0A175YMW5 |
| Daucus carot | A0A164W8B8 |
| Daucus carot | A0A166GYQ3 |
| Daucus carot | A0A166AH60 |
| Daucus carot | Q6I6Y6 |
| Deferribacter | D3P9U3 |
| Defluviimona | A0A159Z548 |
| Defluviimona | A0A2R8BKZ6 |
| Defluviimona | A0A2S8S490 |
| Delftia acidov | A0A2W5GAR1 |
| Delphinapter | A0A2Y9M5R2 |
| Delphinapter | A0A2Y9N707 |
| Delphinapter | A0A2Y9N2B0 |
| Delphinapter | A0A2Y9LKQ9 |
| Delphinapter | A0A2Y9MXX5 |
| Delphinapter | A0A2Y9M328 |
| Delphinapter | A0A2Y9LK73 |
| Deltaproteob | A0A2M7PDD6 |
| Deltaproteob | A0A2M7JKL0 |
| Deltaproteob | A0A2H0AN95 |
| Deltaproteob | A0A0T6AER9 |
| Deltaproteob | A0A2N2GCD1 |
| Deltaproteob | A0A1F9CSM5 |
| Deltaproteob | A0A2E7UQW9 |
| Deltaproteob | A0A2D5SNG4 |
| Deltaproteob | A0A2D6MP68 |
| Deltaproteob | A0A359C885 |
| Dendrobium | A0A2I0WMU7 |
| Dendrobium | A0A2I0XFC1 |
| Dendrobium | A0A2I0VZ05 |
| Dendrobium | A0A2I0VZ03 |
| Dendrobium | A0A2I0WD68 |
| Dendrobium | A0A2I0VCZ8 |
| Dendrobium | A0A2I0VY27 |
| Dendrobium | A0A2I0WAD1 |
| Dendrobium | A0A2I0VN57 |
| Dendrobium | A0A2I0VZ11 |
| Dendrobium | A0A2I0WY89 |
| Dendrobium | A0A2I0WQI1 |
| Dendrobium | A0A2I0WKE7 |

| | |
|---|---|
| Dendrobium | A0A2I0VZ09 |
| Dendrobium | A0A2I0XCU2 |
| Dendrobium | A0A2I0WR68 |
| Dendrobium | A0A2I0X2L0 |
| Dendrobium | A0A2I0VTF8 |
| Dendrobium | A0A2I0WD76 |
| Dendrobium | A0A2I0WMM2 |
| Dendrobium | A0A2I0VDJ3 |
| Dendrobium | A0A2I0V982 |
| Dendrobium | A0A2I0V8Q2 |
| Dendrobium | A0A2I0XIC7 |
| Dendrobium | A0A2I0VPC9 |
| Dendrobium | A0A2I0WAG8 |
| Dendrobium | A0A2I0WMK7 |
| Dendrobium | A0A2I0WUZ0 |
| Dendrobium | A0A2I0VZ10 |
| Dendrobium | A0A2I0V9B4 |
| Dendroctonu | U4UF05 |
| Dendroctonu | U4UJ61 |
| Dendroctonu | U4UHH1 |
| Dendroctonu | U4UAS2 |
| Dendroctonu | U4UHR8 |
| Dendroctonu | U4U368 |
| Dendroctonu | U4UMJ6 |
| Dendroctonu | U4UC62 |
| Dendroctonu | N6TCY6 |
| Dendroctonu | N6T5M3 |
| Dendroctonu | U4U5U9 |
| Dendroctonu | N6TNG4 |
| Dendroctonu | N6TYW1 |
| Dendroctonu | N6SW87 |
| Dendroctonu | N6SZ63 |
| Desulfobacte | A0A1J5E4F1 |
| Desulfobacte | A0A351CCD8 |
| Desulfobacte | A0A3N5QE56 |
| Desulfocapsa | M1NBW9 |
| Devosia croci | A0A1I7N2V7 |
| Devosia epid | A0A0F5QJQ8 |
| Devosia insul | A0A1E5XPL7 |
| Devosia insul | A0A1E5XM33 |
| Devosia limi [ | A0A0F5LGC2 |

| | |
|---|---|
| Devosia psycl | A0A0F5Q359 |
| Devosia ribof | A0A087M5P3 |
| Devosia ribof | A0A087M6Z6 |
| Devosia soli.x | A0A0F5L306 |
| Devosia sp. 1 | A0A087LR20 |
| Devosia sp. 6 | A0A1M3KU89 |
| Devosia sp. 6 | A0A1Q3Z4X7 |
| Devosia sp. D | A0A024KDE1 |
| Devosia sp. D | A0A024KC27 |
| Devosia sp. H | A0A0K0TS75 |
| Devosia sp. I5 | A0A2L1S3I6 |
| Devosia sp. L( | A0A085EUA2 |
| Devosia sp. L( | A0A085FE05 |
| Devosia sp. L | A0A0Q4PK00 |
| Devosia sp. M | A0A447I8R2 |
| Devosia sp. R | A0A0Q6PUQ7 |
| Devosia sp. R | A0A0Q7E2E7 |
| Devosia sp. R | A0A0Q7I7H2 |
| Devosia sp. R | A0A0T2MN01 |
| Devosia sp. R | A0A0Q8IC41 |
| Devosia sp. Y | A0A1H9CCU2 |
| Devosia sp. Y | A0A1H8ZG73 |
| Diaphorina ci | A0A3Q0JEG8 |
| Diaphorina ci | A0A3Q0J9Q0 |
| Diaphorina ci | A0A3Q0JCD1 |
| Diaphorina ci | A0A3Q0ISE2 |
| Diaphorina ci | A0A3Q0J8K4 |
| Diaphorina ci | A0A1S3D587 |
| Diaporthe am | A0A0G2HYC6 |
| Diaporthe am | A0A0G2FQS8 |
| Diaporthe am | A0A0G2HKP0 |
| Diaporthe am | A0A0G2FYA3 |
| Diaporthe am | A0A0G2FNE9 |
| Diaporthe am | A0A0G2HTH8 |
| Diaporthe he | A0A2P5HKQ7 |
| Diaporthe he | A0A2P5HFP5 |
| Diaporthe he | A0A2P5HKH9 |
| Diaporthe he | A0A2P5HW67 |
| Diaporthe he | A0A2P5HMQ7 |
| Diaporthe he | A0A2P5HTN7 |
| Diaporthe he | A0A2P5HMX3 |

| | |
|---|---|
| Diaporthe he | A0A2P5I8D4 |
| Dichantheliur | A0A1E5V5N8 |
| Dichantheliur | A0A1E5WE31 |
| Dichantheliur | A0A1E5VW96 |
| Dichantheliur | A0A1E5VZA9 |
| Dichantheliur | A0A1E5VG01 |
| Dichantheliur | A0A1E5VSM3 |
| Dichantheliur | A0A1E5W1E8 |
| Dichantheliur | A0A1E5VU03 |
| Dichantheliur | A0A1E5UWE4 |
| Dichantheliur | A0A1E5VYJ8 |
| Dichantheliur | A0A1E5VM23 |
| Dichantheliur | A0A1E5URK1 |
| Dichantheliur | A0A1E5WF62 |
| Dichantheliur | A0A1E5VK53 |
| Dichantheliur | A0A1E5UP39 |
| Dichantheliur | A0A1E5VK38 |
| Dichantheliur | A0A1E5V886 |
| Dichantheliur | A0A1E5W3D1 |
| Dichantheliur | A0A1E5WE45 |
| Dichantheliur | A0A1E5VDA3 |
| Dichantheliur | A0A1E5WHY6 |
| Dichantheliur | A0A1E5VK85 |
| Dichantheliur | A0A1E5UVP0 |
| Dichantheliur | A0A1E5UR82 |
| Dichantheliur | A0A1E5ULJ0 |
| Dichantheliur | A0A1E5VWU8 |
| Dichantheliur | A0A1E5V9T7 |
| Dichantheliur | A0A1E5V139 |
| Dichantheliur | A0A1E5ULP8 |
| Dichantheliur | A0A1E5VY77 |
| Dichantheliur | A0A1E5WKA8 |
| Dichantheliur | A0A1E5VIX5 |
| Dichantheliur | A0A1E5VIW0 |
| Dichantheliur | A0A1E5V1A3 |
| Dichantheliur | A0A1E5VVS7 |
| Dichantheliur | A0A1E5UY38 |
| Dichantheliur | A0A1E5VM04 |
| Dichantheliur | A0A1E5V8G9 |
| Dichantheliur | A0A1E5VVN3 |
| Dichantheliur | A0A1E5VKB6 |

| | |
|---|---|
| Dichantheliur | A0A1E5WFT2 |
| Dichantheliur | A0A1E5W4R6 |
| Dichantheliur | A0A1E5WAP0 |
| Dichantheliur | A0A1E5VM26 |
| Dichantheliur | A0A1E5WME9 |
| Dichantheliur | A0A1E5UVF0 |
| Dichantheliur | A0A1E5VMM0 |
| Dichantheliur | A0A1E5VU80 |
| Dichantheliur | A0A1E5VS42 |
| Dichantheliur | A0A1E5UV74 |
| Dichantheliur | A0A1E5VIT8 |
| Dichantheliur | A0A1E5VU74 |
| Dichantheliur | A0A1E5VIF0 |
| Dichantheliur | A0A1E5VW42 |
| Dichantheliur | A0A1E5W8Y3 |
| Dichantheliur | A0A1E5VS52 |
| Dichantheliur | A0A1E5WIF4 |
| Dichantheliur | A0A1E5ULC1 |
| Dichantheliur | A0A1E5UQP5 |
| Dichantheliur | A0A1E5VIJ8 |
| Dichantheliur | A0A1E5WCL7 |
| Dichantheliur | A0A1E5UW57 |
| Dictyocaulus | A0A0D8XPX3 |
| Dictyocaulus | A0A0D8X9K2 |
| Dictyocaulus | A0A0D8XPW4 |
| Dictyocaulus | A0A0D8XME1 |
| Dictyocaulus | A0A0D8X7W5 |
| Dictyocaulus | A0A0D8Y1F7 |
| Dictyocaulus | A0A0D8YBC7 |
| Dictyocaulus | A0A0D8XDB9 |
| Dictyocaulus | A0A0D8XCM5 |
| Dictyocaulus | A0A0D8XNV8 |
| Dictyocaulus | A0A0D8XSU8 |
| Dictyocaulus | A0A0D8Y821 |
| Dictyocaulus | A0A0D8Y118 |
| Dictyocaulus | A0A0D8XQ45 |
| Dictyocaulus | A0A0D8Y7C0 |
| Dictyocaulus | A0A0D8Y6I0 |
| Dictyocaulus | A0A0D8XYW9 |
| Dictyocaulus | A0A0D8Y6A7 |
| Dictyocaulus | A0A0D8XWZ8 |

| | |
|---|---|
| Dictyocaulus | A0A0D8XEJ7 |
| Dictyocaulus | A0A0D8XTU3 |
| Dictyostelium | Q54TD6 |
| Dictyostelium | F0ZKZ0 |
| Didymella ral | A0A163FB68 |
| Didymella ral | A0A163C076 |
| Didymella ral | A0A162ZDC7 |
| Didymella ral | A0A162ZPJ8 |
| Didymella ral | A0A163F3P6 |
| Didymella ral | A0A162WAZ1 |
| Didymella ral | A0A163J5K2 |
| Dinoroseobac | A0A2D8TLG9 |
| Dinothrombiu | A0A443QH82 |
| Dinothrombiu | A0A443QVN9 |
| Dinothrombiu | A0A3S4RFD9 |
| Dinothrombiu | A0A443Q9A2 |
| Dinothrombiu | A0A3S4Q503 |
| Dinothrombiu | A0A443Q760 |
| Dinothrombiu | A0A3S3QB39 |
| Dinothrombiu | A0A443RFM8 |
| Dinothrombiu | A0A3S3S9I9 |
| Dinothrombiu | A0A443R8P4 |
| Diplocarpon r | A0A2H3FTD0 |
| Diplocarpon r | A0A2H3F312 |
| Diplodia corti | A0A1J9S7L6 |
| Diplodia corti | A0A1J9R5G2 |
| Diplodia corti | A0A1J9RDT5 |
| Diplodia corti | A0A1J9RSV2 |
| Diplodia corti | A0A1J9R6F7 |
| Diplodia corti | A0A1J9QKG2 |
| Diplodia corti | A0A1J9SJL7 |
| Diplodia seria | A0A0G2GEA1 |
| Diplodia seria | A0A0G2E8Z2 |
| Diplodia seria | A0A0G2DVZ6 |
| Diplodia seria | A0A0G2GU77 |
| Diplodia seria | A0A1S8BAS3 |
| Diploscapter | A0A2A2JU22 |
| Diploscapter | A0A2A2L628 |
| Diploscapter | A0A2A2JTT9 |
| Diploscapter | A0A2A2L657 |
| Diploscapter | A0A2A2J1X0 |

Diploscapter A0A2A2K5D7
Diploscapter A0A2A2JJI2
Diploscapter A0A2A2JIU6
Diploscapter A0A2A2K5L0
Diploscapter A0A2A2KWG8
Diploscapter A0A2A2JS95
Diploscapter A0A2A2LN52
Diploscapter A0A2A2L620
Diploscapter A0A2A2J5B9
Diploscapter A0A2A2KVU0
Diploscapter A0A2A2L6B7
Diploscapter A0A2A2J211
Diploscapter A0A2A2LYR2
Diploscapter A0A2A2L669
Diploscapter A0A2A2L5U1
Diploscapter A0A2A2K927
Diploscapter A0A2A2L660
Diploscapter A0A2A2KCF6
Diploscapter A0A2A2KWT1
Diploscapter A0A2A2JR35
Diploscapter A0A2A2JRM6
Diploscapter A0A2A2K944
Diploscapter A0A2A2L6G5
Diploscapter A0A2A2JTJ2
Diploscapter A0A2A2J269
Diploscapter A0A2A2L631
Diploscapter A0A2A2JJ18
Diploscapter A0A2A2JW46
Diploscapter A0A2A2KN03
Diploscapter A0A2A2J7C6
Diploscapter A0A2A2LFJ4
Diploscapter A0A2A2JK27
Diploscapter A0A2A2LSI3
Diploscapter A0A2A2K984
Diploscapter A0A2A2J737
Diploscapter A0A2A2JVG0
Diploscapter A0A2A2LHE2
Diploscapter A0A2A2J9M8
Diploscapter A0A2A2LZT7
Diploscapter A0A2A2LK42
Diploscapter A0A2A2L5A9

| | |
|---|---|
| Diploscapter | A0A2A2L5Z3 |
| Diploscapter | A0A2A2K562 |
| Diploscapter | A0A2A2JIY3 |
| Diploscapter | A0A2A2JB04 |
| Diploscapter | A0A2A2JVQ1 |
| Diploscapter | A0A2A2M1Y0 |
| Diploscapter | A0A2A2L2G3 |
| Diploscapter | A0A2A2L2E8 |
| Diploscapter | A0A2A2L668 |
| Diploscapter | A0A2A2KEI4 |
| Diplotaxis eru | Q8LP56 |
| Diplotaxis siif | Q8LP55 |
| Dipodomys o | A0A1S3FYB6 |
| Dipodomys o | A0A1S3FZS6 |
| Dipodomys o | A0A1S3G699 |
| Dipodomys o | A0A1S3FYB4 |
| Dipodomys o | A0A1S3FW29 |
| Dipodomys o | A0A1S3G403 |
| Dipodomys o | A0A1S3FU82 |
| Dipodomys o | A0A1S3FX05 |
| Donghicola sp | A0A1M4MWK1 |
| Dontostemor | M1WLK5 |
| Dontostemor | M1WM21 |
| Dontostemor | M1WJ15 |
| Dontostemor | M1WKU1 |
| Dontostemor | M1WQZ1 |
| Dontostemor | M1WJ16 |
| Dorcoceras h | A0A2Z7CXC8 |
| Dorcoceras h | A0A2Z7D6Z2 |
| Dorcoceras h | A0A2Z7AFH7 |
| Dorcoceras h | A0A2Z7D8Q9 |
| Dorcoceras h | A0A2Z7CER6 |
| Dorcoceras h | A0A2Z7B0U4 |
| Dorcoceras h | A0A2Z7CJB8 |
| Dorcoceras h | A0A2Z7A354 |
| Dorcoceras h | A0A2Z7BCF0 |
| Dorcoceras h | A0A2Z6ZZX4 |
| Dorcoceras h | A0A2Z7D973 |
| Dorcoceras h | A0A2Z7AG07 |
| Dorcoceras h | A0A2Z7B1U4 |
| Dorcoceras h | A0A2Z7B0H7 |

| | |
|---|---|
| Dorcoceras h | A0A2Z7CYS7 |
| Dorcoceras h | A0A2Z7A3I7 |
| Dothistroma | M2YM64 |
| Dothistroma | N1PXS3 |
| Dothistroma | N1PBL6 |
| Dothistroma | M2WMG3 |
| Dothistroma | N1PFX2 |
| Dracunculus | A0A3P7PPB3 |
| Dracunculus | A0A3P7Q930 |
| Dracunculus | A0A0N4U4X9 |
| Dracunculus | A0A3P7PQ65 |
| Dracunculus | A0A0N4UGI5 |
| Dracunculus | A0A0N4UGV7 |
| Dracunculus | A0A0N4UKQ2 |
| Dracunculus | A0A0N4U0P8 |
| Dracunculus | A0A0N4UIB0 |
| Dracunculus | A0A0N4UGG1 |
| Dracunculus | A0A0N4UDY3 |
| Dracunculus | A0A0N4UNZ5 |
| Dracunculus | A0A0N4UEW0 |
| Dracunculus | A0A0N4U2K8 |
| Dracunculus | A0A0N4U723 |
| Dracunculus | A0A0N4U3H3 |
| Dracunculus | A0A158Q4C2 |
| Dracunculus | A0A0N4U9B2 |
| Dracunculus | A0A0N4ULW1 |
| Drechmeria c | A0A151GIG1 |
| Drechslerella | W7I2Z9 |
| Drosophila ar | B3MYN7 |
| Drosophila ar | B3MF92 |
| Drosophila ar | B3MFW4 |
| Drosophila ar | B3LWY2 |
| Drosophila ar | A0A0P8Y1T9 |
| Drosophila ar | B3M3A3 |
| Drosophila ar | B3M3A4 |
| Drosophila ar | B3LV56 |
| Drosophila bu | A0A0M4ESS7 |
| Drosophila bu | A0A0M4EKR6 |
| Drosophila bu | A0A0M4EV08 |
| Drosophila bu | A0A0M5J364 |
| Drosophila bu | A0A0M4EXF3 |

| | |
|---|---|
| Drosophila bu | A0A0M4EEP6 |
| Drosophila bu | A0A0M4EME8 |
| Drosophila bu | A0A0M4ECI2 |
| Drosophila er | B3P534 |
| Drosophila er | B3NSI7 |
| Drosophila er | A0A0Q5WP35 |
| Drosophila er | A0A0Q5WL19 |
| Drosophila er | B3N8M9 |
| Drosophila er | B3P5C3 |
| Drosophila er | B3P535 |
| Drosophila er | B3P944 |
| Drosophila er | B3P7G9 |
| Drosophila fic | A0A1W4V338 |
| Drosophila fic | A0A1W4UYM8 |
| Drosophila fic | A0A1W4W708 |
| Drosophila fic | A0A1W4UGH8 |
| Drosophila fic | A0A1W4UYB4 |
| Drosophila fic | A0A1W4U5N1 |
| Drosophila fic | A0A1W4UGL8 |
| Drosophila fic | A0A1W4U500 |
| Drosophila fic | A0A1W4V9D3 |
| Drosophila fic | A0A1W4VW08 |
| Drosophila fic | A0A1W4UR70 |
| Drosophila gr | B4JTG4 |
| Drosophila gr | B4JSG6 |
| Drosophila gr | B4J4M5 |
| Drosophila gr | B4J8W1 |
| Drosophila gr | B4JRW8 |
| Drosophila gr | B4JMU5 |
| Drosophila gr | B4J4M4 |
| Drosophila gr | B4JRW7 |
| Drosophila gr | B4JG50 |
| Drosophila gu | A0A3B0KC09 |
| Drosophila gu | A0A3B0KS22 |
| Drosophila gu | A0A3B0JG58 |
| Drosophila gu | A0A3B0JKN7 |
| Drosophila gu | A0A3B0K9B5 |
| Drosophila gu | A0A3B0KB54 |
| Drosophila gu | A0A3B0JFB9 |
| Drosophila gu | A0A3B0K860 |
| Drosophila gu | A0A3B0JT97 |

| | |
|---|---|
| Drosophila gu | A0A3B0JFL0 |
| Drosophila gu | A0A3B0JKK1 |
| Drosophila m | Q9BMD4 |
| Drosophila m | Q86P35 |
| Drosophila m | A0A0B4LF37 |
| Drosophila m | A0A0B4LEX9 |
| Drosophila m | Q9BMD5 |
| Drosophila m | Q8IGS2 |
| Drosophila m | A1Z7G0 |
| Drosophila m | Q1WWD1 |
| Drosophila m | B6IDK6 |
| Drosophila m | Q9W5Z0 |
| Drosophila m | B5X535 |
| Drosophila m | Q32KF1 |
| Drosophila m | A1Z7G1 |
| Drosophila m | Q8MS37 |
| Drosophila m | Q9VCR4 |
| Drosophila m | Q9VAG2 |
| Drosophila m | Q9V9X0 |
| Drosophila m | Q9V9X1 |
| Drosophila m | A1Z8J3 |
| Drosophila m | A1Z8J4 |
| Drosophila m | A0A0Q9X7G2 |
| Drosophila m | B4KR75 |
| Drosophila m | A0A0Q9X8J5 |
| Drosophila m | B4K8A2 |
| Drosophila m | B4KT11 |
| Drosophila m | B4K8A0 |
| Drosophila m | B4K4N1 |
| Drosophila m | B4KT12 |
| Drosophila m | B4KMN4 |
| Drosophila m | B4L2D3 |
| Drosophila pe | B4GBV0 |
| Drosophila pe | B4GP76 |
| Drosophila pe | B4GYU7 |
| Drosophila pe | B4GYU9 |
| Drosophila pe | B4GGH2 |
| Drosophila pe | B4GBV1 |
| Drosophila pe | B4GNR1 |
| Drosophila pe | B4H315 |
| Drosophila ps | B5E0A1 |

| | |
|---|---|
| Drosophila ps | Q29BN3 |
| Drosophila ps | B5DYT0 |
| Drosophila ps | B5DYS9 |
| Drosophila ps | A0A0R3NKK5 |
| Drosophila ps | Q293D9 |
| Drosophila ps | B5DKG8 |
| Drosophila se | B4HRZ3 |
| Drosophila se | B4IM31 |
| Drosophila se | B4HNF3 |
| Drosophila se | B4HRZ2 |
| Drosophila se | B4HZF8 |
| Drosophila se | B4IJ01 |
| Drosophila se | B4I033 |
| Drosophila se | B4HFA6 |
| Drosophila si | B4QFU2 |
| Drosophila si | A0A0J9TYZ5 |
| Drosophila si | A0A0J9TWC5 |
| Drosophila si | B4QSN2 |
| Drosophila si | B4QB78 |
| Drosophila si | B4NSA9 |
| Drosophila si | B4QSN1 |
| Drosophila si | B4R0T6 |
| Drosophila si | A0A0J9R8B2 |
| Drosophila si | B4R722 |
| Drosophila si | A0A0J9RAB3 |
| Drosophila vi | B4LMY0 |
| Drosophila vi | B4MEP6 |
| Drosophila vi | A0A0Q9WGQ1 |
| Drosophila vi | B4LZT0 |
| Drosophila vi | B4LMX9 |
| Drosophila vi | B4LX01 |
| Drosophila vi | B4LMZ8 |
| Drosophila vi | B4MEC9 |
| Drosophila vi | B4LX02 |
| Drosophila w | B4NIE3 |
| Drosophila w | B4NMW3 |
| Drosophila w | B4NA23 |
| Drosophila w | B4MYB6 |
| Drosophila w | B4N9U9 |
| Drosophila w | B4NMW2 |
| Drosophila w | B4NIE6 |

| | |
|---|---|
| Drosophila w | B4NCY5 |
| Drosophila ya | B4P7G4 |
| Drosophila ya | B4PPJ2 |
| Drosophila ya | A0A0R1E8R0 |
| Drosophila ya | B4PWJ2 |
| Drosophila ya | B4P2V3 |
| Drosophila ya | B4PNE6 |
| Drosophila ya | B4PNE5 |
| Drosophila ya | A0A0R1E9F3 |
| Drosophila ya | B4PN38 |
| Drosophila ya | A0A0R1DN71 |
| Drosophila ya | A0A0R1EAK2 |
| Drosophila ya | A0A0R1E956 |
| Dryobates pu | A0A093GPD4 |
| Dryobates pu | A0A093GWQ7 |
| Duddingtonia | A0A437ACB8 |
| Duddingtonia | A0A436ZUE2 |
| Duddingtonia | A0A437A9N2 |
| Dufourea nov | A0A154NWA8 |
| Dufourea nov | A0A154PP16 |
| Dufourea nov | A0A154PDD4 |
| Dufourea nov | A0A154P8R1 |
| Dufourea nov | A0A154PGX2 |
| Dufourea nov | A0A154PF39 |
| Echinaster se | A0A2I6QP38 |
| Echinococcus | W6UHM1 |
| Echinococcus | A0A068WP17 |
| Echinococcus | W6UV63 |
| Echinococcus | W6UEB0 |
| Echinococcus | A0A068WL63 |
| Echinococcus | A0A068WDY5 |
| Echinococcus | A0A068X3L0 |
| Echinococcus | A0A068Y252 |
| Echinococcus | A0A068Y2J7 |
| Echinococcus | A0A068Y2T5 |
| Echinococcus | A0A087VY50 |
| Echinostoma | A0A183AP36 |
| Ectocarpus si | D8LQ47 |
| Ectocarpus si | D7FKT5 |
| Egretta garze | A0A091JLU0 |
| Egretta garze | A0A091JD63 |

| | |
|---|---|
| Egretta garze | A0A091J4H6 |
| Egretta garze | A0A091JSL3 |
| Eimeria acerv | U6GJ11 |
| Eimeria acerv | U6GCD3 |
| Eimeria acerv | U6GPI2 |
| Eimeria acerv | U6GEH8 |
| Eimeria acerv | U6GV22 |
| Eimeria acerv | U6GQW4 |
| Eimeria brun | U6LNL3 |
| Eimeria brun | U6M125 |
| Eimeria brun | U6LQZ6 |
| Eimeria brun | U6LPX3 |
| Eimeria brun | U6LG02 |
| Eimeria brun | U6LEP7 |
| Eimeria maxi | U6M8Q0 |
| Eimeria maxi | G0LEU7 |
| Eimeria maxi | U6M1Y7 |
| Eimeria maxi | U6MAY8 |
| Eimeria maxi | U6LYQ2 |
| Eimeria mitis | U6KE00 |
| Eimeria necat | U6N3N3 |
| Eimeria necat | U6N256 |
| Eimeria necat | U6MUL8 |
| Eimeria necat | U6MIM2 |
| Eimeria necat | U6MTA8 |
| Eimeria necat | U6MUE5 |
| Eimeria necat | U6MZX5 |
| Eimeria necat | B0FLX7 |
| Eimeria necat | U6N5K1 |
| Eimeria praec | U6H1N4 |
| Eimeria praec | U6GYZ4 |
| Eimeria stied | A0A1S6JVG4 |
| Eimeria tenel | U6KPE6 |
| Eimeria tenel | U6L5A4 |
| Eimeria tenel | U6KPH4 |
| Eimeria tenel | A9Q278 |
| Eimeria tenel | U6KZX4 |
| Eimeria tenel | U6KR47 |
| Eimeria tenel | U6L0S8 |
| Eimeria tenel | H9B9Q2 |
| Eimeria tenel | Q9U966 |

| | |
|---|---|
| Elaeophora e | A0A0R3S3W6 |
| Elaeophora e | A0A0R3RMT7 |
| Elaeophora e | A0A0R3S175 |
| Elaeophora e | A0A158Q7L5 |
| Elaeophora e | A0A0R3S6Y6 |
| Elaeophora e | A0A0R3RSG8 |
| Elaeophora e | A0A0R3RQS7 |
| Elaeophora e | A0A158Q8Q1 |
| Elaeophora e | A0A0R3RJF9 |
| Elaeophora e | A0A0R3S0T9 |
| Elaeophora e | A0A0R3RLA6 |
| Elaeophora e | A0A158Q7Z5 |
| Elaeophora e | A0A0R3RNM3 |
| Elaeophora e | A0A0R3RQF1 |
| Elaeophora e | A0A0R3RND4 |
| Elsinoe austr | A0A2P7ZU16 |
| Elsinoe austr | A0A2P8A0K2 |
| Elsinoe austr | A0A2P7ZTR3 |
| Elsinoe austr | A0A2P7Z2I6 |
| Elsinoe austr | A0A2P8A4U8 |
| Elsinoe austr | A0A2P8AJH4 |
| Elsinoe austr | A0A2P8A0V9 |
| Elsinoe austr | A0A2P7ZE80 |
| Elsinoe austr | A0A2P7ZEA0 |
| Elsinoe austr | A0A2P7YG32 |
| Elsinoe austr | A0A2P8A7V0 |
| Elsinoe austr | A0A2P7ZAA3 |
| Elsinoe austr | A0A2P7Z3V7 |
| Elsinoe austr | A0A2P8AIG7 |
| Elsinoe austr | A0A2P8A7U5 |
| Elsinoe austr | A0A2P7Z292 |
| Elsinoe austr | A0A2P8ABP9 |
| Elsinoe austr | A0A2P7ZCS8 |
| Elsinoe austr | A0A2P7ZQ48 |
| Elsinoe austr | A0A2P7Z2J1 |
| Elsinoe austr | A0A2P8AE22 |
| Elsinoe austr | A0A2P8A8C3 |
| Elsinoe austr | A0A2P7ZQ26 |
| Elysia chlorot | A0A433T326 |
| Elysia chlorot | A0A3S1BH73 |
| Elysia chlorot | A0A3S0ZKA2 |

| | |
|---|---|
| Elysia chlorot | A0A433TIN4 |
| Elysia chlorot | A0A3S0ZE69 |
| Elysia chlorot | A0A3S1BVS8 |
| Elysia chlorot | A0A433SIG5 |
| Elysia chlorot | A0A433U408 |
| Elysia chlorot | A0A433TGA1 |
| Elysia chlorot | A0A433TX18 |
| Elysia chlorot | A0A433U537 |
| Elysia chlorot | A0A433TXC9 |
| Elysia chlorot | A0A3S1HNN2 |
| Elysia chlorot | A0A3S1BNP3 |
| Elysia chlorot | A0A3S1BSL7 |
| Elysia chlorot | A0A3S1BG98 |
| Elysia chlorot | A0A3S1AGJ4 |
| Elysia chlorot | A0A433UC51 |
| Elysia chlorot | A0A3S1CCH7 |
| Emiliania hux | G8DGE3 |
| Emiliania hux | R1EI28 |
| Emiliania hux | R1BAM4 |
| Emiliania hux | R1B683 |
| Emiliania hux | R1C548 |
| Emiliania hux | R1C7G4 |
| Emiliania hux | R1CY05 |
| Emiliania hux | R1DNH1 |
| Emiliania hux | R1F3X8 |
| Emiliania hux | R1BID0 |
| Emiliania hux | R1B9P9 |
| Emiliania hux | R1FUQ2 |
| Emiliania hux | R1E328 |
| Emiliania hux | R1BF00 |
| Emiliania hux | R1D562 |
| Emiliania hux | A0A0D3I431 |
| Emiliania hux | R1DXZ7 |
| Emiliania hux | R1FC05 |
| Emiliania hux | R1G197 |
| Emiliania hux | R1F3B4 |
| Emiliania hux | R1BP77 |
| Emiliania hux | R1EXU6 |
| Emiliania hux | R1FN73 |
| Emiliania hux | R1EEU0 |
| Emiliania hux | R1C4W0 |

| | |
|---|---|
| Emiliania hux | R1D1K2 |
| Endocarpon p | U1GDJ7 |
| Endocarpon p | U1HN00 |
| Endocarpon p | U1HVD3 |
| Enhydra lutri: | A0A2Y9J0Y1 |
| Enhydra lutri: | A0A2Y9J0V1 |
| Enhydra lutri: | A0A2Y9J9R1 |
| Enhydra lutri: | A0A2Y9KTY8 |
| Enhydra lutri: | A0A2Y9KI03 |
| Enhydra lutri: | A0A2Y9KPG6 |
| Enhydra lutri: | A0A2Y9IZ91 |
| Enhydra lutri: | A0A2Y9J4D9 |
| Ensete ventri | A0A444ETX7 |
| Ensete ventri | A0A444FD73 |
| Ensete ventri | A0A427BAD6 |
| Ensete ventri | A0A444E0U3 |
| Ensete ventri | A0A445MMM3 |
| Ensete ventri | A0A444DKE0 |
| Ensete ventri | A0A426Y944 |
| Ensete ventri | A0A444DNP9 |
| Ensete ventri | A0A444CQP6 |
| Ensete ventri | A0A426ZAP5 |
| Ensete ventri | A0A444DGM2 |
| Ensete ventri | A0A426XMX6 |
| Ensete ventri | A0A444DJ88 |
| Ensete ventri | A0A444FQB9 |
| Ensete ventri | A0A444EPL5 |
| Ensete ventri | A0A444DYW9 |
| Ensete ventri | A0A426Z8Y2 |
| Ensete ventri | A0A427A7Z6 |
| Ensete ventri | A0A427ADX1 |
| Ensete ventri | A0A427A375 |
| Ensete ventri | A0A426ZAK1 |
| Ensete ventri | A0A426Y7Y5 |
| Ensete ventri | A0A444F0S5 |
| Ensete ventri | A0A426Z0W7 |
| Ensete ventri | A0A426XTU0 |
| Ensete ventri | A0A426X112 |
| Ensete ventri | A0A445MI72 |
| Ensete ventri | A0A426Y1Q2 |
| Ensete ventri | A0A426XZ16 |

| | |
|---|---|
| Ensete ventri | A0A444FSB7 |
| Ensete ventri | A0A444DA88 |
| Ensete ventri | A0A444CS48 |
| Ensete ventri | A0A426YR82 |
| Ensete ventri | A0A426YBE7 |
| Ensete ventri | A0A426WZY4 |
| Ensete ventri | A0A444EBR5 |
| Ensete ventri | A0A444G7H0 |
| Ensete ventri | A0A444FF33 |
| Ensete ventri | A0A427BBY0 |
| Ensete ventri | A0A426YCT9 |
| Ensete ventri | A0A426X001 |
| Ensete ventri | A0A426XKV3 |
| Ensete ventri | A0A426YHE0 |
| Ensete ventri | A0A444FQX5 |
| Ensete ventri | A0A444FXT6 |
| Ensete ventri | A0A444G8Y1 |
| Ensete ventri | A0A444E0K6 |
| Ensete ventri | A0A444FSD3 |
| Ensete ventri | A0A444C1A8 |
| Ensete ventri | A0A444EZB8 |
| Ensete ventri | A0A444EYD8 |
| Ensete ventri | A0A444FT20 |
| Ensete ventri | A0A426YPX6 |
| Ensete ventri | A0A444E275 |
| Ensete ventri | A0A444FI43 |
| Ensete ventri | A0A426YFG1 |
| Ensete ventri | A0A427B9L9 |
| Ensete ventri | A0A445MGF6 |
| Ensete ventri | A0A426Z6T2 |
| Ensete ventri | A0A426XGC5 |
| Ensete ventri | A0A444GI94 |
| Ensete ventri | A0A444C1E0 |
| Ensete ventri | A0A444GAN1 |
| Ensete ventri | A0A444C1T9 |
| Ensete ventri | A0A444GIB6 |
| Ensete ventri | A0A444G7E9 |
| Ensifer adhae | W8IAH2 |
| Ensifer adhae | A0A0L8BYH5 |
| Ensifer adhae | A0A2V4WJQ1 |
| Ensifer adhae | A0A285XA72 |

| | |
|---|---|
| Ensifer adhae | D1CSF5 |
| Ensifer alkalis | A0A1E3VH17 |
| Ensifer glycin | A0A178XXR5 |
| Ensifer sojae | A0A249PID2 |
| Ensifer sp. AF | A0A2V4ZAK9 |
| Ensifer sp. AF | A0A2V4ZCS8 |
| Ensifer sp. LC | A0A1C0SLC3 |
| Ensifer sp. LC | A0A1C0SWS9 |
| Ensifer sp. LC | A0A1S1T063 |
| Ensifer sp. M | A0A395L2C9 |
| Ensifer sp. M | A0A2N8JDI4 |
| Ensifer sp. NI | A0A2T3DA21 |
| Ensifer sp. Rc | A0A0Q9EJQ9 |
| Ensifer sp. YR | A0A1G9QSV3 |
| Enterobius ve | A0A0N4UYF4 |
| Enterobius ve | A0A0N4VB29 |
| Enterobius ve | A0A0N4VKQ5 |
| Enterobius ve | A0A158Q9W4 |
| Enterobius ve | A0A0N4V3E1 |
| Enterobius ve | A0A0N4V9C5 |
| Enterobius ve | A0A0N4VBZ0 |
| Enterobius ve | A0A0N4V6Q2 |
| Enterobius ve | A0A0N4V4P5 |
| Enterobius ve | A0A0N4USF4 |
| Enterobius ve | A0A0N4UVL6 |
| Enterobius ve | A0A0N4UTY5 |
| Enterobius ve | A0A0N4VH10 |
| Enterobius ve | A0A0N4V8H6 |
| Enterobius ve | A0A0N4UVC2 |
| Enterobius ve | A0A158QAV3 |
| Enterobius ve | A0A158QB01 |
| Enterobius ve | A0A0N4VNF8 |
| Enterobius ve | A0A0N4VFH7 |
| Enterobius ve | A0A0N4UVC1 |
| Enterobius ve | A0A0N4V445 |
| Enterobius ve | A0A3P6INW5 |
| Epibacterium | A0A1H2U8J0 |
| Epicoccum ni | A0A1Y2LTI9 |
| Epicoccum ni | A0A1Y2M3K0 |
| Epicoccum ni | A0A1Y2LUZ8 |
| Epicoccum ni | A0A1Y2LKS8 |

| | |
|---|---|
| Epicoccum ni | A0A1Y2LT57 |
| Epicoccum ni | A0A1Y2M6T7 |
| Epicoccum ni | A0A1Y2M0F2 |
| Epicoccum ni | A0A1Y2LRR1 |
| Epicoccum ni | A0A1Y2M3W5 |
| Epicoccum ni | A0A1Y2LUZ6 |
| Epicoccum ni | A0A1Y2M927 |
| Epicoccum ni | A0A1Y2LUE0 |
| Epicoccum ni | A0A1Y2LJN4 |
| Epicoccum ni | A0A1Y2LMN7 |
| Epinephelus ( | G4WG37 |
| Equus caballu | F6USP9 |
| Equus caballu | F6ZWI6 |
| Equus caballu | F6PR90 |
| Equus caballu | F7BK76 |
| Equus caballu | A0A3Q2H7K1 |
| Equus caballu | A0A3Q2HUE1 |
| Equus caballu | F6T3E4 |
| Equus caballu | A0A3Q2L7R0 |
| Equus caballu | A0A3Q2L1V8 |
| Erinaceus eur | A0A1S3WHJ7 |
| Erinaceus eur | A0A1S2ZDK0 |
| Erinaceus eur | A0A1S3WHE5 |
| Erinaceus eur | A0A1S3A071 |
| Erinaceus eur | A0A1S3W4L1 |
| Erinaceus eur | A0A1S3W4J1 |
| Erinaceus eur | A0A1S2ZMK4 |
| Erinaceus eur | A0A1S3A1S4 |
| Erinaceus eur | A0A1S2ZDJ8 |
| Erinaceus eur | Q29485 |
| Eruca vesicar | Q8LP54 |
| Eruca vesicar | Q8LP53 |
| Erucastrum a | Q8LP52 |
| Erucastrum g | Q8LP51 |
| Erysimum ch | O80355 |
| Erythranthe g | A0A022QTQ7 |
| Erythranthe g | A0A022PT68 |
| Erythranthe g | A0A022R2X9 |
| Erythranthe g | A0A022QF32 |
| Erythranthe g | A0A022R014 |
| Erythranthe g | A0A022RNY5 |

| | |
|---|---|
| Erythranthe g | A0A022QYC6 |
| Erythranthe g | A0A022QY85 |
| Erythranthe g | A0A022RTU8 |
| Erythranthe g | A0A022QN22 |
| Erythranthe g | A0A022QIR8 |
| Erythranthe g | A0A022QN16 |
| Erythranthe g | A0A022R8V5 |
| Erythranthe g | A0A022Q9S0 |
| Erythranthe g | A0A022QRE8 |
| Erythranthe g | A0A022PWI2 |
| Erythranthe g | A0A022QGP2 |
| Erythranthe g | A0A022QI21 |
| Erythranthe g | A0A022R1Z8 |
| Erythranthe g | A0A022QJW6 |
| Erythranthe g | A0A022RSP1 |
| Erythranthe g | A0A022R428 |
| Erythranthe g | A0A022QNQ9 |
| Erythranthe g | A0A022QUF4 |
| Erythranthe g | A0A022R0W8 |
| Erythranthe g | A0A022QJX3 |
| Erythranthe g | A0A022QY81 |
| Erythranthe g | A0A022QT32 |
| Erythranthe g | A0A022QZT5 |
| Erythranthe g | A0A022QZQ9 |
| Erythranthe g | A0A022QW47 |
| Erythranthe g | A0A022RTP0 |
| Erythranthe g | A0A022R1U4 |
| Erythranthe g | A0A022QEX0 |
| Erythranthe g | A0A022R559 |
| Erythranthe g | A0A022R2X4 |
| Erythranthe g | A0A022QPW6 |
| Erythranthe g | A0A022R1U7 |
| Erythranthe g | A0A022R4G1 |
| Erythranthe g | A0A022RYE8 |
| Erythranthe g | A0A022RS75 |
| Erythranthe g | A0A022QX25 |
| Erythranthe g | A0A022QZ29 |
| Erythranthe g | A0A022QGM9 |
| Erythranthe g | A0A022QFN7 |
| Erythranthe g | A0A022R018 |
| Erythranthe g | A0A022QKT8 |

| | |
|---|---|
| Erythranthe g | A0A022R8M0 |
| Erythrobacte | A0A074M1L8 |
| Erythrobacte | A0A0N8GK13 |
| Eschrichtius r | A0A2F0AWT5 |
| Esox lucius.xl | A0A3P9A6E8 |
| Esox lucius.xl | A0A3P9AKF4 |
| Esox lucius.xl | A0A3P8ZCM4 |
| Esox lucius.xl | A0A3P8YY57 |
| Esox lucius.xl | A0A3P8YIZ1 |
| Esox lucius.xl | A0A3P8YIY1 |
| Esox lucius.xl | A0A3P8Z4D8 |
| Esox lucius.xl | A0A3P8XUT0 |
| Esox lucius.xl | A0A3P8X7I2 |
| Eucalyptus gr | A0A058ZUL1 |
| Eucalyptus gr | A0A059C9G5 |
| Eucalyptus gr | A0A059C6C8 |
| Eucalyptus gr | A0A058ZUX1 |
| Eucalyptus gr | A0A059CAD5 |
| Eucalyptus gr | A0A059C8D0 |
| Eucalyptus gr | A0A059C8W2 |
| Eucalyptus gr | A0A059BMB1 |
| Eucalyptus gr | A0A059CF25 |
| Eucalyptus gr | A0A059BJE7 |
| Eucalyptus gr | A0A059A1V7 |
| Eucalyptus gr | A0A059CAF8 |
| Eucalyptus gr | A0A059D2A0 |
| Eucalyptus gr | A0A058ZTJ3 |
| Eucalyptus gr | A0A058ZS00 |
| Eucalyptus gr | A0A059BMY5 |
| Eucalyptus gr | A0A059C9N1 |
| Eucalyptus gr | A0A059CAG3 |
| Eucalyptus gr | A0A059BJZ0 |
| Eucalyptus gr | A0A059BVD6 |
| Eucalyptus gr | A0A059CD96 |
| Eucalyptus gr | A0A059D0P0 |
| Eucalyptus gr | A0A059CDS6 |
| Eucalyptus gr | A0A059C900 |
| Eucalyptus gr | A0A059CDJ8 |
| Eucalyptus gr | A0A059C9K6 |
| Eucalyptus gr | A0A059BCC2 |
| Eucalyptus gr | A0A059C904 |

Eucalyptus gr A0A059C2G3
Eucalyptus gr A0A059C908
Eucalyptus gr A0A059CF20
Eucalyptus gr A0A058ZUY0
Eucalyptus gr A0A059BX54
Eucalyptus gr A0A059BMJ5
Eucalyptus gr A0A059DHG7
Eucalyptus gr A0A059C975
Eucalyptus gr A0A059DG99
Eucalyptus gr A0A059CL14
Eucalyptus gr A0A059CDT6
Eucalyptus gr A0A059BNH3
Eucalyptus gr A0A059C9N8
Eucalyptus gr A0A058ZSM4
Eucalyptus gr A0A059CDK9
Eucalyptus gr A0A059BLH2
Eucalyptus gr A0A059DCZ0
Eucalyptus gr A0A059CZX4
Eucalyptus gr A0A059BAQ1
Eucalyptus gr A0A059CAB5
Eucalyptus gr A0A059CAX2
Eucalyptus gr A0A059BVE0
Eucalyptus gr A0A059A785
Eucalyptus gr A0A059C1G2
Eucalyptus gr A0A059A615
Eucalyptus gr A0A059A075
Eucalyptus gr A0A059C9A3
Eucalyptus gr A0A059C8U8
Eucalyptus gr A0A059B9B7
Eucalyptus gr A0A059BMI0
Eucalyptus gr A0A059BRV2
Eucalyptus gr A0A059BKX0
Eucalyptus gr A0A059BNH7
Eucalyptus gr A0A059A6I8
Eucalyptus gr A0A059C945
Eucalyptus gr A0A059C6U7
Eucalyptus gr A0A059BVV6
Eucalyptus gr A0A059AXJ7
Eucalyptus gr A0A059DF87
Eucalyptus gr A0A059C9T6
Eucalyptus gr A0A059DFY6

Eucalyptus gr A0A059BN16
Eucalyptus gr A0A059CF13
Eucalyptus gr A0A058ZSA5
Eucalyptus gr A0A059D643
Eucalyptus gr A0A059D1S0
Eucalyptus gr A0A059D249
Eucalyptus gr A0A059BVR8
Eucalyptus gr A0A058ZTY8
Eucalyptus gr A0A059BTY0
Eucalyptus gr A0A059BVQ8
Eucalyptus gr A0A059CDF2
Eucalyptus gr A0A059BWS9
Eucalyptus gr A0A059DGY3
Eucalyptus gr A0A058ZUE3
Eucalyptus gr A0A059BR54
Eucalyptus gr A0A059CDJ4
Eucalyptus gr A0A059CE26
Eucalyptus gr A0A059BLY4
Eucalyptus gr A0A059CE51
Eucalyptus gr A0A059BB89
Eucalyptus gr A0A059DGF5
Eucalyptus gr A0A059BUD2
Eucalyptus gr A0A059BLZ5
Eucalyptus gr A0A059CUD3
Eucalyptus gr A0A059C982
Eucalyptus gr A0A059C9H8
Eucalyptus gr A0A059C958
Eucalyptus gr A0A059BB94
Eucalyptus gr A0A059CAH7
Eucalyptus gr A0A059CXX9
Eucalyptus gr A0A059C8V4
Eucalyptus gr A0A059CE41
Eucalyptus gr A0A059BLY9
Eucalyptus gr A0A059CDR9
Eucalyptus gr A0A059CAB0
Eucalyptus gr A0A059DGX5
Eucalyptus gr A0A059BJD9
Eucalyptus gr A0A059BMJ2
Eucalyptus gr A0A059BNG0
Eucalyptus gr A0A059BM31
Eucalyptus gr A0A058ZQZ8

Eucalyptus gr A0A059CPC6
Eucalyptus gr A0A059CF29
Eucalyptus gr A0A058ZSW8
Eucalyptus gr A0A059CEE1
Eucalyptus gr A0A059BM22
Eucalyptus gr A0A059CAC4
Eucalyptus gr A0A059A163
Eucalyptus gr A0A059A230
Eucalyptus gr A0A059BVD2
Eucalyptus gr A0A059CEG0
Eucalyptus gr A0A059CBT7
Eucalyptus gr A0A059C9B7
Eucalyptus gr A0A059C1I4
Eucalyptus gr A0A059BUV4
Eucalyptus gr A0A059D2V1
Eucalyptus gr A0A059D2V0
Eucalyptus gr A0A059C631
Eucalyptus gr A0A059CD83
Eucalyptus gr A0A059CDT5
Eucalyptus gr A0A059B8N1
Eucalyptus gr A0A059C5L8
Eucalyptus gr A0A059BMX2
Eucalyptus gr A0A059A0X2
Eucalyptus gr A0A059BMX6
Eucalyptus gr A0A059AAQ7
Eucalyptus gr A0A059CDH3
Eucalyptus gr A0A059CE67
Eucalyptus gr A0A059BMA5
Eucalyptus gr A0A059DGF0
Eucalyptus gr A0A059BMG9
Eucalyptus gr A0A059AB79
Eucalyptus gr A0A059CSN4
Eucalyptus gr A0A059CED3
Eucalyptus gr A0A059C9F2
Eucalyptus gr A0A059CI09
Eucalyptus gr A0A059BMW7
Eucalyptus gr A0A059BLI8
Eucalyptus gr A0A059CE37
Eucalyptus gr A0A059A790
Eucalyptus gr A0A058ZU59
Eucalyptus gr A0A059BSQ0

Eucalyptus gr A0A059D582
Eucalyptus gr A0A059CEP6
Eucalyptus gr A0A059C965
Eucalyptus gr A0A059CB94
Eucalyptus gr A0A059BEM4
Eucalyptus gr A0A059CD78
Eucalyptus gr A0A059CE46
Eucalyptus gr A0A059BLG8
Eucalyptus gr A0A059DG00
Eucalyptus gr A0A059CF15
Eucalyptus gr A0A059DDF4
Eucalyptus gr A0A059BJQ8
Eucalyptus gr A0A059B892
Eucalyptus gr A0A059BUD7
Eucalyptus gr A0A059BAR2
Eucalyptus gr A0A059BMY1
Eucalyptus gr A0A059AWY3
Eucalyptus gr A0A059B897
Eucalyptus gr A0A059BJX1
Eucalyptus gr A0A058ZSV0
Eucalyptus gr A0A059BNF8
Eucalyptus gr A0A059A0R6
Eucalyptus gr A0A059A823
Eucalyptus gr A0A059BKW4
Eucalyptus gr A0A059BDD1
Eucalyptus gr A0A059C6B4
Eucalyptus gr A0A059DFF0
Eucalyptus gr A0A059DFE3
Eucalyptus gr A0A059BMB6
Eucalyptus gr A0A059BBN7
Eucalyptus gr A0A059D2I7
Eucalyptus gr A0A059C9R7
Eucalyptus gr A0A059BTK0
Eucalyptus gr A0A059BBA1
Eucalyptus gr A0A058ZW23
Eucalyptus gr A0A059A1Z3
Eucalyptus gr A0A059BM37
Eucalyptus gr A0A059C9W0
Eucalyptus gr A0A059CF08
Eucalyptus gr A0A059BMH9
Eucalyptus gr A0A059A6J2

Eucalyptus gr A0A059BM34
Eucalyptus gr A0A059BLW8
Eucalyptus gr A0A059AKP5
Eucalyptus gr A0A059CAE7
Eucalyptus gr A0A058ZV14
Eucalyptus gr A0A059DFZ6
Eucalyptus gr A0A059C153
Eucalyptus gr A0A059B9A1
Eucalyptus gr A0A059DFZ4
Eucalyptus gr A0A059A113
Eucalyptus gr A0A059A235
Eucalyptus gr A0A059C9M3
Eucalyptus gr A0A059BMI4
Eucalyptus gr A0A059DGY9
Eucalyptus gr A0A058ZVI2
Eucalyptus gr A0A059BMB9
Eucalyptus gr A0A059CB42
Eucalyptus gr A0A059BLZ9
Eucalyptus gr A0A059BUZ3
Eucalyptus gr A0A059C536
Eucalyptus gr A0A059CDU0
Eucalyptus gr A0A059C8X4
Eucalyptus gr A0A059B232
Eucalyptus gr A0A059AB02
Eucalyptus gr A0A058ZZN8
Eucalyptus gr A0A059CBU2
Eucalyptus gr A0A059C1M6
Eucalyptus gr A0A059DGG0
Eucalyptus gr A0A059D5Y0
Eucalyptus gr A0A059CAC9
Eucalyptus gr A0A059C9Q5
Eucalyptus gr A0A059C9M7
Eucalyptus gr A0A059BVV1
Eucalyptus gr A0A058ZZL0
Eucalyptus gr A0A059DG97
Eucalyptus gr A0A059CDS9
Eucalyptus gr A0A059BJX7
Eucalyptus gr A0A059CDG7
Eucalyptus gr A0A059CAI8
Eucalyptus gr A0A059CDS1
Eucalyptus gr A0A059BLH9

Eucalyptus gr A0A059C8X9
Eucalyptus gr A0A059C976
Eucalyptus gr A0A059CEE5
Eucalyptus gr A0A059CAJ3
Eucalyptus gr A0A059BNG6
Eucalyptus gr A0A058ZV47
Eucalyptus gr A0A059BR56
Eucalyptus gr A0A059BVR3
Eucalyptus gr A0A059BUQ9
Eucalyptus gr A0A059BVR0
Eucalyptus gr A0A059C7M3
Eucalyptus gr A0A059BB21
Eucalyptus gr A0A059DFD9
Eucalyptus gr A0A059BRU7
Eucalyptus gr A0A059C951
Eucalyptus gr A0A059CEQ0
Eucalyptus gr A0A059A7T8
Eucalyptus gr A0A059CAX8
Eucalyptus gr A0A058ZQT7
Eucalyptus gr A0A059C0W1
Eucalyptus gr A0A059C9H1
Eucalyptus gr A0A059D1S5
Eucalyptus gr A0A058ZXR1
Eucalyptus gr A0A059CF34
Eucalyptus gr A0A059BNE6
Eucalyptus gr A0A059DGE5
Eucalyptus gr A0A059C970
Eucalyptus gr A0A059CDU4
Eucalyptus gr A0A058ZVT6
Eucalyptus gr A0A059D0Q2
Eucalyptus gr A0A059C9F8
Eucalyptus gr A0A059DG91
Eucalyptus gr A0A059CA18
Eucalyptus gr A0A059BLV9
Eucalyptus gr A0A059CAP5
Eucalyptus gr A0A059A0T8
Eucalyptus gr A0A058ZRN7
Eucalyptus gr A0A059BB75
Eucalyptus gr A0A059CDK1
Eucalyptus gr A0A059BVR4
Eucalyptus gr A0A059CAE0

| | |
|---|---|
| Eucalyptus gr | A0A059BM19 |
| Eucalyptus gr | A0A058ZSW3 |
| Eucalyptus gr | A0A059DG93 |
| Eucalyptus gr | A0A059CE63 |
| Eucalyptus gr | A0A058ZVN1 |
| Eucalyptus gr | A0A059D240 |
| Eucalyptus gr | A0A059CAE9 |
| Eucalyptus gr | A0A059BCB2 |
| Eucalyptus gr | A0A059CE56 |
| Eucalyptus gr | A0A059BWP5 |
| Eucalyptus gr | A0A059C9H4 |
| Eucalyptus gr | A0A059BRU8 |
| Eucalyptus gr | A0A059BAZ9 |
| Eucalyptus gr | A0A059CDH8 |
| Eucalyptus gr | A0A059C9M0 |
| Eucalyptus gr | A0A059BC92 |
| Eucalyptus gr | A0A059CED7 |
| Eucalyptus gr | A0A058ZTK4 |
| Eucalyptus gr | A0A059CDF8 |
| Eucalyptus gr | A0A059BMZ2 |
| Eucalyptus gr | A0A059CFL3 |
| Eucalyptus gr | A0A059C947 |
| Eufriesea me: | A0A310S483 |
| Eufriesea me: | A0A310SBN6 |
| Eufriesea me: | A0A310SWL1 |
| Eufriesea me: | A0A310SLW1 |
| Euroglyphus | A0A1Y3B584 |
| Euroglyphus | A0A1Y3AU99 |
| Euroglyphus | A0A1Y3AXF6 |
| Euroglyphus | A0A1Y3B164 |
| Euroglyphus | A0A1Y3BLQ2 |
| Euroglyphus | A0A1Y3AUF8 |
| Euroglyphus | A0A1Y3BLR8 |
| Euroglyphus | A0A1Y3AZZ7 |
| Euroglyphus | A0A1Y3AWF6 |
| Euroglyphus | A0A1Y3BKE0 |
| Euryarchaeot | A0A2E4CYV5 |
| Eurypyga heli | A0A093IZQ0 |
| Eutrema sals | V4MUM3 |
| Eutrema sals | V4L7H3 |
| Eutrema sals | V4KWJ2 |

| | |
|---|---|
| Eutrema sals | V4KQS5 |
| Eutrema sals | V4KY09 |
| Eutrema sals | V4MJV0 |
| Eutrema sals | V4LNJ7 |
| Eutrema sals | V4KDR5 |
| Eutrema sals | V4L028 |
| Eutrema sals | V4MC52 |
| Eutrema sals | V4L095 |
| Eutrema sals | V4L2I0 |
| Eutrema sals | V4KJ04 |
| Eutrema sals | V4KNB2 |
| Eutrema sals | V4KHG6 |
| Eutrema sals | V4M7D2 |
| Eutrema sals | V4MED4 |
| Eutrema sals | V4MBM7 |
| Eutrema sals | V4LX42 |
| Eutrema sals | V4LB40 |
| Eutrema sals | V4MDH2 |
| Eutrema sals | V4K896 |
| Eutrema sals | V4KDS9 |
| Eutrema sals | V4P511 |
| Eutrema sals | V4LVA5 |
| Eutrema sals | V4KWJ7 |
| Eutrema sals | V4LD36 |
| Eutrema sals | V4KC43 |
| Eutrema sals | V4KHG0 |
| Eutrema sals | V4L090 |
| Eutrema sals | V4KDS4 |
| Eutrema sals | V4L8J2 |
| Eutrema sals | V4KPI9 |
| Eutrema sals | V4JV72 |
| Eutrema sals | V4MIC9 |
| Eutrema sals | V4KPJ9 |
| Eutrema sals | V4JTX9 |
| Eutrema sals | V4NUT8 |
| Eutrema sals | V4NAM5 |
| Eutrema sals | V4MH01 |
| Eutrema sals | V4KFK6 |
| Exophiala der | H6C4W5 |
| Exophiala me | A0A0D1Y6D0 |
| Exophiala me | A0A438NHT6 |

| | |
|---|---|
| Exophiala olig | A0A0D2AIX8 |
| Exophiala olig | A0A0D2DN60 |
| Exophiala sid | A0A0D1X4X0 |
| Exophiala sid | A0A0D1YE49 |
| Exophiala sid | A0A0D1WNZ7 |
| Exophiala sid | A0A0D1YQP5 |
| Exophiala spi | A0A0D1Z1V9 |
| Exophiala spi | A0A0D2BF00 |
| Exophiala xer | A0A0D2CZ88 |
| Exophiala xer | A0A0D2CUE6 |
| Exophiala xer | A0A0D2D105 |
| Fagus sylvatic | A0A2N9F471 |
| Fagus sylvatic | A0A2N9G5U5 |
| Fagus sylvatic | A0A2N9ID93 |
| Fagus sylvatic | A0A2N9I6Q1 |
| Fagus sylvatic | A0A2N9IVV1 |
| Fagus sylvatic | A0A2N9F4G1 |
| Fagus sylvatic | A0A2N9F6W7 |
| Fagus sylvatic | A0A2N9F3H1 |
| Fagus sylvatic | A0A2N9FT24 |
| Fagus sylvatic | A0A2N9ERT8 |
| Fagus sylvatic | A0A2N9G5U0 |
| Fagus sylvatic | A0A2N9EZK0 |
| Fagus sylvatic | A0A2N9I7P8 |
| Fagus sylvatic | A0A2N9EYQ2 |
| Fagus sylvatic | A0A2N9HI25 |
| Fagus sylvatic | A0A2N9I8I7 |
| Fagus sylvatic | A0A2N9H741 |
| Fagus sylvatic | A0A2N9H8W4 |
| Fagus sylvatic | A0A2N9F7R0 |
| Fagus sylvatic | A0A2N9H6F3 |
| Fagus sylvatic | A0A2N9H2X2 |
| Fagus sylvatic | A0A2N9J533 |
| Fagus sylvatic | A0A2N9G8Z2 |
| Fagus sylvatic | A0A2N9HS19 |
| Fagus sylvatic | A0A2N9HL38 |
| Fagus sylvatic | A0A2N9IBZ3 |
| Fagus sylvatic | A0A2N9G5W6 |
| Fagus sylvatic | A0A2N9HCP4 |
| Fagus sylvatic | A0A2N9H5Z2 |
| Fagus sylvatic | A0A2N9FDE0 |

| | |
|---|---|
| Fagus sylvatic | A0A2N9F3U1 |
| Fagus sylvatic | A0A2N9IVM7 |
| Fagus sylvatic | A0A2N9FCL3 |
| Fagus sylvatic | A0A2N9HX85 |
| Fagus sylvatic | A0A2N9FNS6 |
| Fagus sylvatic | A0A2N9FA80 |
| Fagus sylvatic | A0A2N9F6Y2 |
| Fagus sylvatic | A0A2N9EZI6 |
| Fagus sylvatic | A0A2N9HDA4 |
| Fagus sylvatic | A0A2N9I2U2 |
| Fagus sylvatic | A0A2N9IY45 |
| Fagus sylvatic | A0A2N9EYP9 |
| Fagus sylvatic | A0A2N9HL99 |
| Fagus sylvatic | A0A2N9H4B2 |
| Fagus sylvatic | A0A2N9HRU4 |
| Fagus sylvatic | A0A2N9HI04 |
| Fagus sylvatic | A0A2N9FE92 |
| Fagus sylvatic | A0A2N9HPG2 |
| Fagus sylvatic | A0A2N9FX35 |
| Fagus sylvatic | A0A2N9H9S5 |
| Fagus sylvatic | A0A2N9IUZ4 |
| Fagus sylvatic | A0A2N9IV03 |
| Fagus sylvatic | A0A2N9I632 |
| Fagus sylvatic | A0A2N9J6T9 |
| Fagus sylvatic | A0A2N9EYX2 |
| Fagus sylvatic | A0A2N9FCP0 |
| Fagus sylvatic | A0A2N9F9Y2 |
| Fagus sylvatic | A0A2N9HTG9 |
| Fagus sylvatic | A0A2N9H672 |
| Fagus sylvatic | A0A2N9GTW2 |
| Fagus sylvatic | A0A2N9IZ71 |
| Fagus sylvatic | A0A2N9HNC1 |
| Fagus sylvatic | A0A2N9HE29 |
| Fagus sylvatic | A0A2N9FRE3 |
| Fagus sylvatic | A0A2N9HZG0 |
| Fagus sylvatic | A0A2N9FDT8 |
| Fagus sylvatic | A0A2N9GA03 |
| Fagus sylvatic | A0A2N9HIJ0 |
| Fagus sylvatic | A0A2N9ICZ8 |
| Fagus sylvatic | A0A2N9GJI4 |
| Fagus sylvatic | A0A2N9IM33 |

| | |
|---|---|
| Fagus sylvatic | A0A2N9HTB4 |
| Fagus sylvatic | A0A2N9GJM5 |
| Fagus sylvatic | A0A2N9HL23 |
| Fagus sylvatic | A0A2N9FCP6 |
| Fagus sylvatic | A0A2N9G696 |
| Fagus sylvatic | A0A2N9IPQ1 |
| Fagus sylvatic | A0A2N9GAM8 |
| Fagus sylvatic | A0A2N9J640 |
| Fagus sylvatic | A0A2N9FCQ2 |
| Fagus sylvatic | A0A2N9EM72 |
| Fagus sylvatic | A0A2N9EE92 |
| Fagus sylvatic | A0A2N9ED67 |
| Fagus sylvatic | A0A2N9G5P0 |
| Fagus sylvatic | A0A2N9I9D1 |
| Fagus sylvatic | A0A2N9HYD5 |
| Fagus sylvatic | A0A2N9F021 |
| Fagus sylvatic | A0A2N9H5L7 |
| Fagus sylvatic | A0A2N9IG02 |
| Fagus sylvatic | A0A2N9GLI0 |
| Fagus sylvatic | A0A2N9ELE2 |
| Fagus sylvatic | A0A2N9ER54 |
| Fagus sylvatic | A0A2N9GLH4 |
| Fagus sylvatic | A0A2N9F7N7 |
| Fagus sylvatic | A0A2N9HKW5 |
| Fagus sylvatic | A0A2N9I6X0 |
| Fagus sylvatic | A0A2N9I1I5 |
| Fagus sylvatic | A0A2N9HHV9 |
| Fagus sylvatic | A0A2N9I2Z7 |
| Fagus sylvatic | A0A2N9EM92 |
| Fagus sylvatic | A0A2N9G9P1 |
| Fagus sylvatic | A0A2N9EYP4 |
| Fagus sylvatic | A0A2N9H6F1 |
| Fagus sylvatic | A0A2N9GWJ5 |
| Fagus sylvatic | A0A2N9HS70 |
| Fagus sylvatic | A0A2N9HAB2 |
| Fagus sylvatic | A0A2N9FAP6 |
| Fagus sylvatic | A0A2N9HI47 |
| Fagus sylvatic | A0A2N9EZA0 |
| Fagus sylvatic | A0A2N9H5Z0 |
| Fagus sylvatic | A0A2N9FCU8 |
| Fagus sylvatic | A0A2N9GXA5 |

| | |
|---|---|
| Fagus sylvatic | A0A2N9FZ00 |
| Fagus sylvatic | A0A2N9ER32 |
| Fagus sylvatic | A0A2N9H0J5 |
| Fagus sylvatic | A0A2N9IT13 |
| Fagus sylvatic | A0A2N9HLN1 |
| Fagus sylvatic | A0A2N9GJ59 |
| Fagus sylvatic | A0A2N9F736 |
| Fagus sylvatic | A0A2N9EI38 |
| Fagus sylvatic | A0A2N9HRI8 |
| Fagus sylvatic | A0A2N9FLN9 |
| Fagus sylvatic | A0A2N9I3I6 |
| Fagus sylvatic | A0A2N9G737 |
| Fagus sylvatic | A0A2N9J704 |
| Fagus sylvatic | A0A2N9I596 |
| Fagus sylvatic | A0A2N9FCQ0 |
| Fagus sylvatic | A0A2N9HWH5 |
| Fagus sylvatic | A0A2N9GWS3 |
| Fagus sylvatic | A0A2N9IMS6 |
| Fagus sylvatic | A0A2N9I1K9 |
| Fagus sylvatic | A0A2N9J9T5 |
| Fagus sylvatic | A0A2N9HRW8 |
| Fagus sylvatic | A0A2N9HQ34 |
| Fagus sylvatic | A0A2N9GA49 |
| Fagus sylvatic | A0A2N9GID3 |
| Fagus sylvatic | A0A2N9EE63 |
| Fagus sylvatic | A0A2N9HNV5 |
| Fagus sylvatic | A0A2N9IY81 |
| Fagus sylvatic | A0A2N9EYQ0 |
| Fagus sylvatic | A0A2N9IVN9 |
| Fagus sylvatic | A0A2N9EME6 |
| Fagus sylvatic | A0A2N9H6H0 |
| Fagus sylvatic | A0A2N9ERJ4 |
| Fagus sylvatic | A0A2N9F311 |
| Fagus sylvatic | A0A2N9IBN6 |
| Fagus sylvatic | A0A2N9HKN1 |
| Fagus sylvatic | A0A2N9EYQ9 |
| Fagus sylvatic | A0A2N9IGP9 |
| Fagus sylvatic | A0A2N9H563 |
| Fagus sylvatic | A0A2N9I8M7 |
| Fagus sylvatic | A0A2N9I612 |
| Fagus sylvatic | A0A2N9HB53 |

| | |
|---|---|
| Fagus sylvatic | A0A2N9HHE3 |
| Fagus sylvatic | A0A2N9FNV9 |
| Fagus sylvatic | A0A2N9GFL6 |
| Fagus sylvatic | A0A2N9GUQ4 |
| Fagus sylvatic | A0A2N9EYQ8 |
| Fagus sylvatic | A0A2N9HFF6 |
| Fagus sylvatic | A0A2N9GAZ3 |
| Fagus sylvatic | A0A2N9GZD3 |
| Fagus sylvatic | A0A2N9IE26 |
| Fagus sylvatic | A0A2N9HWE4 |
| Fagus sylvatic | A0A2N9EZ06 |
| Fagus sylvatic | A0A2N9EE70 |
| Fagus sylvatic | A0A2N9IH59 |
| Fagus sylvatic | A0A2N9G6C0 |
| Fagus sylvatic | A0A2N9IC01 |
| Fagus sylvatic | A0A2N9HTA6 |
| Fagus sylvatic | A0A2N9HEY7 |
| Fagus sylvatic | A0A2N9EM85 |
| Fagus sylvatic | A0A2N9FTC8 |
| Fagus sylvatic | A0A2N9J971 |
| Fagus sylvatic | A0A2N9FSD1 |
| Fagus sylvatic | A0A2N9HLB3 |
| Fagus sylvatic | A0A2N9HHG3 |
| Fagus sylvatic | A0A2N9I338 |
| Fagus sylvatic | A0A2N9FD43 |
| Fagus sylvatic | A0A2N9HYX6 |
| Fagus sylvatic | A0A2N9HM72 |
| Fagus sylvatic | A0A2N9IGI8 |
| Fagus sylvatic | A0A2N9IVH2 |
| Fagus sylvatic | A0A2N9HDT5 |
| Fagus sylvatic | A0A2N9FXK4 |
| Fagus sylvatic | A0A2N9GE58 |
| Fagus sylvatic | A0A2N9F4K3 |
| Fagus sylvatic | A0A2N9G9R4 |
| Fagus sylvatic | A0A2N9FW96 |
| Fagus sylvatic | A0A2N9GXP3 |
| Fagus sylvatic | A0A2N9FYD6 |
| Fagus sylvatic | A0A2N9IAR8 |
| Fagus sylvatic | A0A2N9F9M6 |
| Fagus sylvatic | A0A2N9GCZ4 |
| Fagus sylvatic | A0A2N9G2I6 |

| | |
|---|---|
| Fagus sylvatic | A0A2N9EYU2 |
| Fagus sylvatic | A0A2N9G9Q1 |
| Fagus sylvatic | A0A2N9IX44 |
| Fagus sylvatic | A0A2N9IHR5 |
| Fagus sylvatic | A0A2N9H6L7 |
| Fagus sylvatic | A0A2N9HT54 |
| Fagus sylvatic | A0A2N9EEA4 |
| Fagus sylvatic | A0A2N9I6X5 |
| Fagus sylvatic | A0A2N9GG82 |
| Fagus sylvatic | A0A2N9FAN8 |
| Fagus sylvatic | A0A2N9G0E5 |
| Fagus sylvatic | A0A2N9HDU0 |
| Fagus sylvatic | A0A2N9IY90 |
| Fagus sylvatic | A0A2N9JBS8 |
| Fagus sylvatic | A0A2N9HW38 |
| Fagus sylvatic | A0A2N9HRB8 |
| Fagus sylvatic | A0A2N9GE76 |
| Fagus sylvatic | A0A2N9HGR5 |
| Fagus sylvatic | A0A2N9HQP2 |
| Fagus sylvatic | A0A2N9F979 |
| Fagus sylvatic | A0A2N9JBR8 |
| Fagus sylvatic | A0A2N9I7X6 |
| Fagus sylvatic | A0A2N9IAJ3 |
| Fagus sylvatic | A0A2N9EZS8 |
| Fagus sylvatic | A0A2N9H640 |
| Fagus sylvatic | A0A2N9G6V6 |
| Fagus sylvatic | A0A2N9HI57 |
| Fagus sylvatic | A0A2N9H894 |
| Fagus sylvatic | A0A2N9IAS3 |
| Fagus sylvatic | A0A2N9F3L2 |
| Fagus sylvatic | A0A2N9HEM6 |
| Fagus sylvatic | A0A2N9G7C0 |
| Fagus sylvatic | A0A2N9F1Q0 |
| Fagus sylvatic | A0A2N9EYP0 |
| Fagus sylvatic | A0A2N9J7F4 |
| Fagus sylvatic | A0A2N9H1C2 |
| Fagus sylvatic | A0A2N9ERP4 |
| Fagus sylvatic | A0A2N9HZ89 |
| Fagus sylvatic | A0A2N9IA84 |
| Fagus sylvatic | A0A2N9IH48 |
| Fagus sylvatic | A0A2N9FXU7 |

| | |
|---|---|
| Fagus sylvatic | A0A2N9H6E0 |
| Fagus sylvatic | A0A2N9I4V8 |
| Fagus sylvatic | A0A2N9IER8 |
| Fagus sylvatic | A0A2N9IVP2 |
| Fagus sylvatic | A0A2N9IHP1 |
| Fagus sylvatic | A0A2N9I4B6 |
| Fagus sylvatic | A0A2N9H6F8 |
| Fagus sylvatic | A0A2N9IGJ1 |
| Fagus sylvatic | A0A2N9IFD7 |
| Fagus sylvatic | A0A2N9J390 |
| Fagus sylvatic | A0A2N9HZI8 |
| Fagus sylvatic | A0A2N9IGJ3 |
| Fagus sylvatic | A0A2N9FUZ8 |
| Fagus sylvatic | A0A2N9G2R4 |
| Fagus sylvatic | A0A2N9ERG2 |
| Fagus sylvatic | A0A2N9GAN0 |
| Falsochrobac | A0A364JW94 |
| Fasciola hepa | A0A2H1C6Y1 |
| Fasciola hepa | A0A2H1C2I6 |
| Felis catus.xls | Q9BH09 |
| Felis catus.xls | A0A2I2UAP2 |
| Felis catus.xls | M3WZJ1 |
| Felis catus.xls | M3VWV2 |
| Felis catus.xls | A0A2K5TIV6 |
| Felis catus.xls | M3X3T9 |
| Ficedula albic | U3K7I1 |
| Ficedula albic | U3K2Y8 |
| Ficedula albic | U3KBU0 |
| Ficedula albic | U3KBT4 |
| Ficedula albic | U3KCE5 |
| Flavimaricola | A0A238LAN8 |
| Flavimaricola | A0A238LBM3 |
| Flavobacteria | A0A2A4M1B2 |
| Flavobacteria | A0A2D5LP88 |
| Flexilinea floc | A0A0S7BYF0 |
| Flexilinea floc | A0A0K8PA51 |
| Folsomia can | A0A226E742 |
| Folsomia can | A0A226EIX0 |
| Folsomia can | A0A226EJL6 |
| Folsomia can | A0A226DUU9 |
| Folsomia can | A0A226ERX6 |

| | |
|---|---|
| Folsomia can | A0A226EKR3 |
| Folsomia can | A0A226EXA3 |
| Folsomia can | A0A226CZ90 |
| Folsomia can | A0A226E675 |
| Folsomia can | A0A226D8L3 |
| Folsomia can | A0A226DA03 |
| Folsomia can | A0A226E8X0 |
| Folsomia can | A0A226ES37 |
| Folsomia can | A0A226E4B6 |
| Folsomia can | A0A226DYP9 |
| Folsomia can | A0A226F3C7 |
| Folsomia can | A0A226DMW5 |
| Fonsecaea m | A0A177EZP3 |
| Fonsecaea m | A0A0D2JU71 |
| Fonsecaea m | A0A0D2JHM0 |
| Fonsecaea m | A0A178BR81 |
| Fonsecaea m | A0A178BZ24 |
| Fonsecaea m | A0A178BL33 |
| Fonsecaea nu | A0A178CTQ1 |
| Fonsecaea pe | A0A0D2E424 |
| Fopius arisan | A0A0C9PLF2 |
| Fopius arisan | A0A0C9Q7Z9 |
| Fopius arisan | A0A0C9Q6E7 |
| Fopius arisan | A0A0C9RT62 |
| Fopius arisan | A0A0C9RVW4 |
| Fopius arisan | A0A0C9QIV9 |
| Fopius arisan | A0A0C9RYR6 |
| Fopius arisan | A0A0C9QF47 |
| Fopius arisan | A0A0C9RAM8 |
| Fopius arisan | A0A0C9R7C8 |
| Fragaria x ana | W8PA20 |
| Fukomys dam | A0A091DZJ9 |
| Fukomys dam | A0A091DXE1 |
| Fukomys dam | A0A091DVM4 |
| Fukomys dam | A0A091CMD3 |
| Fulmarus glac | A0A093I9I3 |
| Fulmarus glac | A0A093ILG6 |
| Fulmarus glac | A0A093IZP1 |
| Fulvimarina r | A0A1W2EWK8 |
| Fulvimarina r | A0A1W2D4E0 |
| Fulvimarina p | Q0G412 |

| | |
|---|---|
| Fulvimarina s | A0A2E1H9K8 |
| Fundulus het | A0A146MU99 |
| Fundulus het | A0A146YM67 |
| Fundulus het | A0A146XF35 |
| Fundulus het | A0A146SKJ1 |
| Fundulus het | A0A146NDC0 |
| Fundulus het | A0A146SAR9 |
| Fundulus het | A0A3Q2P288 |
| Fundulus het | A0A3Q2U5Q0 |
| Fundulus het | A0A146UV90 |
| Fundulus het | A0A3Q2PXJ3 |
| Fundulus het | A0A147AF21 |
| Fundulus het | A0A146NFQ6 |
| Fundulus het | A0A146MVK4 |
| Fundulus het | A0A146NM15 |
| Fundulus het | A0A3Q2NV28 |
| Fundulus het | A0A146NKH6 |
| Fundulus het | A0A3Q2QG44 |
| Fundulus het | A0A3Q2NUN0 |
| Fundulus het | A0A146MX16 |
| Fundulus het | A0A146NV94 |
| Fundulus het | A0A146MUA7 |
| Fundulus het | A0A3Q2NRI0 |
| Fundulus het | A0A146UVF0 |
| Fundulus het | A0A146ML14 |
| Fundulus het | A0A3Q2P3A5 |
| Fundulus het | A0A146NEU9 |
| Fundulus het | A0A146NDC3 |
| Fundulus het | A0A146Y0F0 |
| Fundulus het | A0A146MMW4 |
| Fundulus het | A0A146R9E1 |
| Fundulus het | A0A146YKS9 |
| Fundulus het | A0A146NEK3 |
| Fundulus het | A0A147AX70 |
| Fundulus het | A0A3Q2P247 |
| Fundulus het | A0A146YWT4 |
| Fundulus het | A0A146NEL7 |
| Fundulus het | A0A3Q2STI2 |
| Fundulus het | A0A146YKR1 |
| Fundulus het | A0A146ML01 |
| Fundulus het | A0A3Q2NUL5 |

| | |
|---|---|
| fungal sp. No | A0A0S6XK74 |
| fungal sp. No | A0A0S6X4V7 |
| fungal sp. No | A0A0C9MAW8 |
| fungal sp. No | A0A0S6XHR7 |
| fungal sp. No | A0A0S6XII5 |
| fungal sp. No | A0A0S6XQ97 |
| fungal sp. No | A0A0S6XL07 |
| fungal sp. No | A0A0C9LMV7 |
| fungal sp. No | A0A0S6XI06 |
| fungal sp. No | A0A0S6XTD5 |
| fungal sp. No | A0A0S6XM18 |
| fungal sp. No | A0A0S6XLT8 |
| fungal sp. No | A0A0S6XMF5 |
| fungal sp. No | A0A0C9ME04 |
| fungal sp. No | A0A0S6XI16 |
| fungal sp. No | A0A0S6XNW0 |
| fungal sp. No | A0A1V1T8U0 |
| fungal sp. No | A0A1V1T9B7 |
| fungal sp. No | A0A1V1TBD7 |
| Fusarium aml | A0A428UEU2 |
| Fusarium aml | A0A428TVT7 |
| Fusarium aml | A0A428UJD4 |
| Fusarium aml | A0A428UW00 |
| Fusarium aml | A0A428U053 |
| Fusarium aml | A0A428U9S6 |
| Fusarium aml | A0A428TW22 |
| Fusarium aml | A0A428T8Z1 |
| Fusarium aml | A0A428USM7 |
| Fusarium aml | A0A428RYB6 |
| Fusarium aml | A0A428U1S8 |
| Fusarium aml | A0A428TDI1 |
| Fusarium aml | A0A428U0L8 |
| Fusarium aml | A0A428TDN9 |
| Fusarium aml | A0A428TDI8 |
| Fusarium aml | A0A428UAR9 |
| Fusarium cul | A0A2T4GF04 |
| Fusarium cul | A0A2T4GEI2 |
| Fusarium cul | A0A2T4GM20 |
| Fusarium cul | A0A2T4GQ39 |
| Fusarium cul | A0A2T4GZI7 |
| Fusarium cul | A0A2T4HAV6 |

| | |
|---|---|
| Fusarium culı | A0A2T4H2K6 |
| Fusarium culı | A0A2T4GU55 |
| Fusarium culı | A0A2T4GS35 |
| Fusarium culı | A0A2T4GCH5 |
| Fusarium culı | A0A2T4GFN6 |
| Fusarium culı | A0A2T4GL46 |
| Fusarium culı | A0A2T4GYZ7 |
| Fusarium culı | A0A2T4GVC4 |
| Fusarium culı | A0A2T4GS29 |
| Fusarium culı | A0A2T4GYX5 |
| Fusarium culı | A0A2T4GET1 |
| Fusarium culı | A0A2T4H3Y6 |
| Fusarium culı | A0A2T4GRL8 |
| Fusarium euv | A0A430M7X9 |
| Fusarium euv | A0A430KY02 |
| Fusarium euv | A0A430LYZ3 |
| Fusarium euv | A0A430L5I8 |
| Fusarium euv | A0A430LMK1 |
| Fusarium euv | A0A430LAE0 |
| Fusarium euv | A0A430M9V2 |
| Fusarium euv | A0A430LTZ7 |
| Fusarium euv | A0A430LXM6 |
| Fusarium euv | A0A430LXM7 |
| Fusarium euv | A0A430L0Q7 |
| Fusarium euv | A0A430LT59 |
| Fusarium euv | A0A430L7B2 |
| Fusarium euv | A0A430LDB8 |
| Fusarium euv | A0A430LIS3 |
| Fusarium euv | A0A430M2H2 |
| Fusarium euv | A0A430LBT6 |
| Fusarium euv | A0A430M5A4 |
| Fusarium euv | A0A430LG97 |
| Fusarium kur | A0A3M2RQX3 |
| Fusarium kur | A0A3M2SJ30 |
| Fusarium kur | A0A3M2S4P6 |
| Fusarium kur | A0A3M2RRC6 |
| Fusarium kur | A0A3M2RNA0 |
| Fusarium kur | A0A3M2S004 |
| Fusarium kur | A0A3M2SGY5 |
| Fusarium kur | A0A3M2RVM1 |
| Fusarium kur | A0A3M2RP84 |

| | |
|---|---|
| Fusarium kur | A0A3M2SAI4 |
| Fusarium kur | A0A3M2RVN5 |
| Fusarium kur | A0A3M2SMU0 |
| Fusarium kur | A0A3M2SDA7 |
| Fusarium kur | A0A3M2SDH3 |
| Fusarium lan | A0A0N0V5S0 |
| Fusarium lan | A0A0M9EXB1 |
| Fusarium lan | A0A0N0DCT1 |
| Fusarium lan | A0A0N0V8P4 |
| Fusarium lan | A0A0M9ETX5 |
| Fusarium lan | A0A0M9ERF9 |
| Fusarium lan | A0A0M9EKX4 |
| Fusarium lan | A0A0M9ETU9 |
| Fusarium lan | A0A0N0DBP2 |
| Fusarium lan | A0A0M9EQI9 |
| Fusarium lan | A0A0M9F2P3 |
| Fusarium lon | A0A395SMB0 |
| Fusarium lon | A0A395SU16 |
| Fusarium lon | A0A395T394 |
| Fusarium lon | A0A395TAS6 |
| Fusarium lon | A0A395RLH4 |
| Fusarium lon | A0A395TAR6 |
| Fusarium lon | A0A395RR81 |
| Fusarium lon | A0A395T1Z9 |
| Fusarium lon | A0A395T8X0 |
| Fusarium lon | A0A395T8W6 |
| Fusarium lon | A0A395T7Q5 |
| Fusarium lon | A0A395RWV1 |
| Fusarium lon | A0A395T6G5 |
| Fusarium lon | A0A395SJK3 |
| Fusarium lon | A0A395T9N8 |
| Fusarium lon | A0A395TBU9 |
| Fusarium lon | A0A395SWW9 |
| Fusarium lon | A0A395SJJ1 |
| Fusarium lon | A0A395T993 |
| Fusarium lon | A0A395RYQ6 |
| Fusarium lon | A0A395TAA6 |
| Fusarium lon | A0A395RI56 |
| Fusarium ma | A0A1L7T250 |
| Fusarium ma | A0A1L7SNI5 |
| Fusarium ma | A0A1L7SY16 |

| | |
|---|---|
| Fusarium ma | A0A1L7U4Q3 |
| Fusarium ma | A0A1L7SQ32 |
| Fusarium ma | A0A1L7TU10 |
| Fusarium ma | A0A1L7UDI1 |
| Fusarium ma | A0A1L7UAA5 |
| Fusarium ma | A0A1L7TG43 |
| Fusarium ma | A0A1L7TZT8 |
| Fusarium ma | A0A1L7SRG8 |
| Fusarium ma | A0A1L7UDC6 |
| Fusarium ma | A0A1L7TI59 |
| Fusarium ma | A0A1L7TU11 |
| Fusarium ma | A0A1L7ST20 |
| Fusarium ma | A0A1L7TSY6 |
| Fusarium ma | A0A1L7T2L2 |
| Fusarium ma | A0A1L7TN71 |
| Fusarium ma | A0A1L7UJW9 |
| Fusarium ma | A0A1L7UHL6 |
| Fusarium ma | A0A1L7STR4 |
| Fusarium oxy | F9F0W5 |
| Fusarium oxy | F9FQB6 |
| Fusarium oxy | F9FN55 |
| Fusarium oxy | F9G9P5 |
| Fusarium oxy | F9FRR0 |
| Fusarium oxy | F9FNV4 |
| Fusarium oxy | F9F3M4 |
| Fusarium oxy | F9F4N5 |
| Fusarium oxy | F9G352 |
| Fusarium oxy | F9G5P5 |
| Fusarium oxy | F9G3D6 |
| Fusarium oxy | F9F676 |
| Fusarium oxy | F9GDQ2 |
| Fusarium oxy | F9FWK6 |
| Fusarium oxy | F9FLK6 |
| Fusarium oxy | F9G1R4 |
| Fusarium oxy | F9G1R3 |
| Fusarium oxy | F9GBZ7 |
| Fusarium oxy | A0A3L6P746 |
| Fusarium oxy | A0A3L6NN57 |
| Fusarium oxy | A0A3L6NQQ4 |
| Fusarium oxy | A0A3L6MW91 |
| Fusarium oxy | A0A3L6NQT5 |

Fusarium oxy A0A3L6NPD4
Fusarium oxy A0A3L6NTI7
Fusarium oxy A0A3L6NEX4
Fusarium oxy A0A3L6N5S7
Fusarium oxy A0A3L6N706
Fusarium oxy A0A3L6NWN1
Fusarium oxy A0A3L6NBP9
Fusarium oxy A0A3L6MWX0
Fusarium oxy A0A3L6N4Z1
Fusarium oxy A0A3L6NXP5
Fusarium oxy A0A3L6N0T9
Fusarium oxy A0A3L6NTQ6
Fusarium oxy A0A3L6NQ88
Fusarium oxy A0A3L6MUU1
Fusarium oxy X0HDS5
Fusarium oxy X0IS02
Fusarium oxy X0HCS8
Fusarium oxy X0HRY6
Fusarium oxy X0HVQ1
Fusarium oxy X0HNT3
Fusarium oxy X0HXU3
Fusarium oxy X0HEE6
Fusarium oxy X0HT53
Fusarium oxy X0HE86
Fusarium oxy X0IFF6
Fusarium oxy X0I6Z5
Fusarium oxy X0HQT8
Fusarium oxy X0HZM6
Fusarium oxy X0HS18
Fusarium oxy X0HBE0
Fusarium oxy X0GNF1
Fusarium oxy X0HVP9
Fusarium oxy X0HB55
Fusarium oxy X0H868
Fusarium oxy X0HRQ2
Fusarium oxy N4TVA0
Fusarium oxy N4UWB4
Fusarium oxy N4UQ87
Fusarium oxy N4TTM6
Fusarium oxy N4U5D0
Fusarium oxy N4U486

| | |
|---|---|
| Fusarium oxy | N4TXC1 |
| Fusarium oxy | N4TPR8 |
| Fusarium oxy | N4UXC7 |
| Fusarium oxy | N4UER4 |
| Fusarium oxy | N4U8I0 |
| Fusarium oxy | N4V0A9 |
| Fusarium oxy | N4UMR2 |
| Fusarium oxy | N4UFQ1 |
| Fusarium oxy | N4UFD6 |
| Fusarium oxy | N4U504 |
| Fusarium oxy | N4US39 |
| Fusarium oxy | N4TMJ2 |
| Fusarium oxy | N1S2M4 |
| Fusarium oxy | N1RYI2 |
| Fusarium oxy | N1RQ64 |
| Fusarium oxy | N1RQ00 |
| Fusarium oxy | N1S7G0 |
| Fusarium oxy | N1RYM7 |
| Fusarium oxy | N1SAV4 |
| Fusarium oxy | N1SAD8 |
| Fusarium oxy | N1RGZ6 |
| Fusarium oxy | N1S5R5 |
| Fusarium oxy | N1RB09 |
| Fusarium oxy | N1R8K0 |
| Fusarium oxy | N1SB76 |
| Fusarium oxy | N1RPT0 |
| Fusarium oxy | N1RPN7 |
| Fusarium oxy | N1S4B5 |
| Fusarium oxy | N1R9G0 |
| Fusarium oxy | N1RV49 |
| Fusarium oxy | N1RC32 |
| Fusarium oxy | N1S7H6 |
| Fusarium oxy | X0KKH1 |
| Fusarium oxy | X0JD64 |
| Fusarium oxy | X0J7Z6 |
| Fusarium oxy | X0JRR5 |
| Fusarium oxy | X0IXV2 |
| Fusarium oxy | X0K4Y4 |
| Fusarium oxy | X0LFY3 |
| Fusarium oxy | X0IRL6 |
| Fusarium oxy | X0K6S9 |

Fusarium oxy X0J8E8
Fusarium oxy X0J8L0
Fusarium oxy X0KS72
Fusarium oxy X0JS17
Fusarium oxy X0KGE6
Fusarium oxy X0J5F8
Fusarium oxy X0KBP3
Fusarium oxy X0IQZ7
Fusarium oxy X0LJG6
Fusarium oxy X0K565
Fusarium oxy X0K4R2
Fusarium oxy X0JMF5
Fusarium oxy A0A0D2XFR8
Fusarium oxy A0A0D2XY32
Fusarium oxy A0A0D2Y321
Fusarium oxy A0A0D2XC34
Fusarium oxy A0A0D2X9W2
Fusarium oxy A0A0D2XKS9
Fusarium oxy A0A0D2XG03
Fusarium oxy A0A0D2YBX0
Fusarium oxy A0A0C4DIY3
Fusarium oxy A0A0J9W4W2
Fusarium oxy A0A0D2Y0C7
Fusarium oxy A0A0J9WPH9
Fusarium oxy A0A0J9VTQ2
Fusarium oxy A0A0J9VIA4
Fusarium oxy A0A0J9UPV1
Fusarium oxy A0A0J9WAX7
Fusarium oxy A0A0J9WCE8
Fusarium oxy A0A0J9UUH4
Fusarium oxy A0A0D2Y340
Fusarium oxy A0A0J9W359
Fusarium oxy A0A0D2Y341
Fusarium oxy A0A0J9WQB9
Fusarium oxy A0A0D2YGJ8
Fusarium oxy A0A0J9VU72
Fusarium oxy X0AFU5
Fusarium oxy W9Z4V9
Fusarium oxy W9ZBD0
Fusarium oxy X0AIL8
Fusarium oxy X0ARG8

Fusarium oxy X0BEV5
Fusarium oxy X0ATQ4
Fusarium oxy X0AKC9
Fusarium oxy X0AP22
Fusarium oxy W9ZFM9
Fusarium oxy X0AH58
Fusarium oxy X0B6E5
Fusarium oxy X0AJE3
Fusarium oxy X0AC84
Fusarium oxy X0B5X6
Fusarium oxy W9ZBV2
Fusarium oxy X0A7A2
Fusarium oxy W9ZGB5
Fusarium oxy X0A725
Fusarium oxy W9NLV3
Fusarium oxy W9PM75
Fusarium oxy W9NVE9
Fusarium oxy W9PM48
Fusarium oxy W9NUL9
Fusarium oxy W9NXE1
Fusarium oxy W9NJT9
Fusarium oxy W9P038
Fusarium oxy W9PVA4
Fusarium oxy W9PIK8
Fusarium oxy W9NWZ0
Fusarium oxy W9NMU0
Fusarium oxy W9QB84
Fusarium oxy W9P2D0
Fusarium oxy W9PWG0
Fusarium oxy W9NHW9
Fusarium oxy W9Q8W3
Fusarium oxy W9NIE4
Fusarium oxy W9Q072
Fusarium oxy W9P3J9
Fusarium oxy W9PEW3
Fusarium oxy A0A2H3G3H1
Fusarium oxy A0A2H3GP16
Fusarium oxy A0A2H3GPR0
Fusarium oxy A0A2H3GI26
Fusarium oxy A0A2H3G3A7
Fusarium oxy A0A2H3GT41

Fusarium oxy A0A2H3HFX7
Fusarium oxy A0A2H3H582
Fusarium oxy A0A2H3HWB5
Fusarium oxy A0A2H3G8B5
Fusarium oxy A0A2H3H7J7
Fusarium oxy A0A2H3HDQ9
Fusarium oxy A0A2H3HBA8
Fusarium oxy A0A2H3HUG8
Fusarium oxy A0A2H3GTZ9
Fusarium oxy A0A2H3H1W4
Fusarium oxy A0A2H3H7B8
Fusarium oxy A0A2H3GR19
Fusarium oxy A0A2H3HIX3
Fusarium oxy X0CBT2
Fusarium oxy X0CJZ6
Fusarium oxy X0CA20
Fusarium oxy X0C0K3
Fusarium oxy X0BLK2
Fusarium oxy X0BYZ9
Fusarium oxy X0BHR3
Fusarium oxy X0D8W0
Fusarium oxy X0D9D6
Fusarium oxy X0BNE2
Fusarium oxy X0CCJ8
Fusarium oxy X0D7P5
Fusarium oxy X0C5I2
Fusarium oxy X0BWY5
Fusarium oxy X0BK94
Fusarium oxy X0C4F8
Fusarium oxy X0BU43
Fusarium oxy X0CH27
Fusarium oxy X0CCJ4
Fusarium oxy X0BT77
Fusarium oxy X0BGL4
Fusarium oxy X0BSM4
Fusarium oxy X0CBV1
Fusarium oxy X0KM84
Fusarium oxy X0KL93
Fusarium oxy X0LI06
Fusarium oxy X0LM24
Fusarium oxy X0L461

| | |
|---|---|
| Fusarium oxy | X0N5D7 |
| Fusarium oxy | X0MNF5 |
| Fusarium oxy | X0LVS2 |
| Fusarium oxy | X0LLL6 |
| Fusarium oxy | X0KZC8 |
| Fusarium oxy | X0MQ25 |
| Fusarium oxy | X0LJZ4 |
| Fusarium oxy | X0LYG6 |
| Fusarium oxy | X0MWI5 |
| Fusarium oxy | X0MGF6 |
| Fusarium oxy | X0MLD1 |
| Fusarium oxy | X0MQJ9 |
| Fusarium oxy | W9JZF8 |
| Fusarium oxy | W9K3B2 |
| Fusarium oxy | W9K2I6 |
| Fusarium oxy | W9KHK3 |
| Fusarium oxy | W9JKU5 |
| Fusarium oxy | W9KL95 |
| Fusarium oxy | W9KMV3 |
| Fusarium oxy | W9KB71 |
| Fusarium oxy | W9LDU1 |
| Fusarium oxy | W9K179 |
| Fusarium oxy | W9JPG0 |
| Fusarium oxy | W9KU91 |
| Fusarium oxy | W9KWA8 |
| Fusarium oxy | W9K2L5 |
| Fusarium oxy | W9JY72 |
| Fusarium oxy | W9JLD1 |
| Fusarium oxy | W9JH38 |
| Fusarium oxy | W9JK71 |
| Fusarium oxy | A0A420PSH8 |
| Fusarium oxy | A0A420PDU2 |
| Fusarium oxy | A0A2H3SXG2 |
| Fusarium oxy | A0A420PMW4 |
| Fusarium oxy | A0A2H3TNB9 |
| Fusarium oxy | A0A2H3TUT4 |
| Fusarium oxy | A0A2H3TT16 |
| Fusarium oxy | A0A2H3TGN6 |
| Fusarium oxy | A0A420MLC5 |
| Fusarium oxy | A0A420SEB3 |
| Fusarium oxy | A0A2H3TL83 |

Fusarium oxy A0A420S1Y6
Fusarium oxy A0A420QWS3
Fusarium oxy A0A420PGL0
Fusarium oxy A0A420NGZ2
Fusarium oxy A0A420R9X9
Fusarium oxy A0A420QHW6
Fusarium oxy A0A420S0P8
Fusarium oxy A0A420TXI9
Fusarium oxy A0A420UC75
Fusarium oxy A0A420MBT8
Fusarium oxy A0A420PHU3
Fusarium oxy A0A420QP35
Fusarium oxy A0A420P6T6
Fusarium oxy A0A420QIK4
Fusarium oxy A0A420PYP6
Fusarium oxy A0A2H3T198
Fusarium oxy A0A420QQV1
Fusarium oxy A0A420NBX5
Fusarium oxy A0A420NZN8
Fusarium oxy A0A420QQT6
Fusarium oxy A0A420RGE4
Fusarium oxy A0A420PWD7
Fusarium oxy A0A2H3TZQ8
Fusarium oxy A0A420NEW3
Fusarium oxy A0A2H3SMM6
Fusarium oxy A0A420RVH4
Fusarium oxy A0A420NNM5
Fusarium oxy A0A2H3T9F1
Fusarium oxy A0A2H3TSH4
Fusarium oxy A0A420RN85
Fusarium oxy A0A420N964
Fusarium oxy A0A420MDX2
Fusarium oxy A0A420NZR8
Fusarium oxy A0A2H3TND6
Fusarium oxy A0A2H3SPH1
Fusarium oxy A0A2H3TLK4
Fusarium oxy A0A420R6D6
Fusarium oxy A0A420QGY8
Fusarium oxy A0A420P5A3
Fusarium oxy A0A420NJK8
Fusarium oxy A0A2H3TLA9

| | |
|---|---|
| Fusarium oxy | A0A420MLJ4 |
| Fusarium oxy | A0A2H3U2W3 |
| Fusarium oxy | A0A420R1M2 |
| Fusarium oxy | A0A420UEN2 |
| Fusarium oxy | A0A420SA71 |
| Fusarium oxy | A0A420PG81 |
| Fusarium oxy | A0A420NEU8 |
| Fusarium oxy | A0A2H3T2A7 |
| Fusarium oxy | A0A420U4W6 |
| Fusarium oxy | A0A420PC63 |
| Fusarium oxy | A0A420Q8J1 |
| Fusarium oxy | A0A420U9L0 |
| Fusarium oxy | A0A420N211 |
| Fusarium oxy | A0A2H3T2B5 |
| Fusarium oxy | A0A420SKN9 |
| Fusarium oxy | A0A420RIQ3 |
| Fusarium oxy | A0A2H3T2H0 |
| Fusarium oxy | A0A420MQK5 |
| Fusarium oxy | A0A420S1Y5 |
| Fusarium oxy | A0A2H3SKQ3 |
| Fusarium oxy | A0A420RSV8 |
| Fusarium oxy | A0A420QNM7 |
| Fusarium oxy | A0A420U9X9 |
| Fusarium oxy | A0A420QBG3 |
| Fusarium poa | A0A1B8AI36 |
| Fusarium poa | A0A1B8AIV6 |
| Fusarium poa | A0A1B8AQ51 |
| Fusarium poa | A0A1B8AHP8 |
| Fusarium poa | A0A1B8B148 |
| Fusarium poa | A0A1B8AZG1 |
| Fusarium poa | A0A1B8AZ43 |
| Fusarium poa | A0A1B8B687 |
| Fusarium poa | A0A1B8AHJ0 |
| Fusarium poa | A0A1B8ARR3 |
| Fusarium poa | A0A1B8B037 |
| Fusarium poa | A0A1B8ARV5 |
| Fusarium poa | A0A1B8AQS5 |
| Fusarium poa | A0A1B8ADY5 |
| Fusarium poa | A0A1B8AE77 |
| Fusarium poa | A0A1B8AH23 |
| Fusarium pro | A0A1L7VUI1 |

Fusarium pro A0A1L7VQT1
Fusarium pro A0A1L7VTN8
Fusarium pro A0A1L7VQ64
Fusarium pro A0A1L7VRM1
Fusarium pro A0A1L7W234
Fusarium pro A0A1L7W7F8
Fusarium pro A0A1L7W3Q1
Fusarium pro A0A1L7VZE6
Fusarium pro A0A1L7W7T0
Fusarium pro A0A1L7W068
Fusarium pro A0A1L7VBG9
Fusarium pro A0A1L7VJ89
Fusarium pro A0A1L7V6L5
Fusarium pro A0A1L7VLJ1
Fusarium pro A0A1L7W6T5
Fusarium pro A0A1L7W2B6
Fusarium pro A0A1L7W3Q7
Fusarium pro A0A1L7VAZ6
Fusarium pro A0A1L7W9S2
Fusarium pro A0A1L7VG50
Fusarium pro A0A1L7V558
Fusarium pro A0A1L7VE71
Fusarium pro A0A1L7VBJ5
Fusarium pse K3V6C2
Fusarium pse K3VNW8
Fusarium pse K3UVA9
Fusarium pse K3VI86
Fusarium pse K3VD86
Fusarium pse K3VSF8
Fusarium pse K3VZG0
Fusarium pse K3W149
Fusarium pse K3UVV0
Fusarium pse K3VF03
Fusarium pse K3VSH8
Fusarium pse K3V8J7
Fusarium pse K3W1U8
Fusarium pse K3UBY1
Fusarium pse K3VV23
Fusarium pse K3V722
Fusarium pse K3UTP7
Fusarium pse K3VPR6

| | |
|---|---|
| Fusarium pse | K3VW92 |
| Fusarium pse | K3VVL6 |
| Fusarium pse | A0A096PED2 |
| Fusarium sp. | A0A428RK34 |
| Fusarium sp. | A0A428SD27 |
| Fusarium sp. | A0A428RDV5 |
| Fusarium sp. | A0A428RQ68 |
| Fusarium sp. | A0A428SN67 |
| Fusarium sp. | A0A428Q5F5 |
| Fusarium sp. | A0A428SNN1 |
| Fusarium sp. | A0A428S0J4 |
| Fusarium sp. | A0A428RZR3 |
| Fusarium sp. | A0A428RSR8 |
| Fusarium sp. | A0A428RXQ1 |
| Fusarium sp. | A0A428RX08 |
| Fusarium sp. | A0A428RZJ5 |
| Fusarium sp. | A0A428SEZ3 |
| Fusarium sp. | A0A428RPS7 |
| Fusarium sp. | A0A428RNL1 |
| Fusarium sp. | A0A428RDW7 |
| Fusarium sp. | A0A428UIA3 |
| Fusarium sp. | A0A428SWL5 |
| Fusarium sp. | A0A428U3R5 |
| Fusarium sp. | A0A428T156 |
| Fusarium sp. | A0A428U3Z4 |
| Fusarium sp. | A0A428TXX9 |
| Fusarium sp. | A0A428SFD7 |
| Fusarium sp. | A0A428TWR6 |
| Fusarium sp. | A0A428UIJ1 |
| Fusarium sp. | A0A428SQQ5 |
| Fusarium sp. | A0A428U227 |
| Fusarium sp. | A0A428UCR8 |
| Fusarium sp. | A0A428THD3 |
| Fusarium sp. | A0A428S8Z0 |
| Fusarium sp. | A0A428UDE6 |
| Fusarium sp. | A0A428SKX5 |
| Fusarium sp. | A0A428PKQ2 |
| Fusarium sp. | A0A428QUF4 |
| Fusarium sp. | A0A428P7J0 |
| Fusarium sp. | A0A428P7H3 |
| Fusarium sp. | A0A428NDT7 |

Fusarium sp. A0A428PH59
Fusarium sp. A0A428Q7Z0
Fusarium sp. A0A428QDG1
Fusarium sp. A0A428RAS8
Fusarium sp. A0A428NYA2
Fusarium sp. A0A428NM38
Fusarium sp. A0A428Q0Y4
Fusarium sp. A0A428PKP7
Fusarium sp. A0A428PTI2
Fusarium sp. A0A428NUP9
Fusarium sp. A0A428PR63
Fusarium sp. A0A428P8A4
Fusarium sp. A0A428R4H3
Fusarium sp. A0A428NIV0
Fusarium sp. A0A428NZ95
Fusarium sp. A0A428PCW5
Fusarium sp. A0A428Q8W1
Fusarium sp. A0A428NR66
Fusarium sp. A0A428NN65
Fusarium sp. A0A428QD28
Fusarium sp. A0A428QWP2
Fusarium sp. A0A428PBY7
Fusarium sp. A0A428PA97
Fusarium sp. A0A428QKE9
Fusarium sp. A0A428P7T0
Fusarium sp. A0A428NXE6
Fusarium sp. A0A428P4U5
Fusarium sp. A0A428QQ80
Fusarium sp. A0A428PCY9
Fusarium sp. A0A428NSA9
Fusarium sp. A0A428PX74
Fusarium sp. A0A428NS70
Fusarium sp. A0A395MGQ2
Fusarium sp. A0A395MGA8
Fusarium sp. A0A395M7J9
Fusarium sp. A0A395MAG0
Fusarium sp. A0A395MLX9
Fusarium sp. A0A395N4U8
Fusarium sp. A0A395MHX2
Fusarium sp. A0A395N159
Fusarium sp. A0A395M5H5

| | |
|---|---|
| Fusarium sp. | A0A395MHI4 |
| Fusarium sp. | A0A395MIF4 |
| Fusarium sp. | A0A395MBX0 |
| Fusarium sp. | A0A395N2D5 |
| Fusarium sp. | A0A395N4Y5 |
| Fusarium sp. | A0A395N3C1 |
| Fusarium sp. | A0A395MVX4 |
| Fusarium sp. | A0A395MI62 |
| Fusarium sp. | A0A395MVQ6 |
| Fusarium sp. | A0A395MCF6 |
| Fusarium sp. | A0A395N1D8 |
| Fusarium sp. | A0A395N4X6 |
| Fusarium sp. | A0A395MZD0 |
| Fusarium sp. | A0A366S714 |
| Fusarium sp. | A0A366QWA4 |
| Fusarium sp. | A0A366QY56 |
| Fusarium sp. | A0A366S516 |
| Fusarium sp. | A0A366R509 |
| Fusarium sp. | A0A366S9E0 |
| Fusarium sp. | A0A366S648 |
| Fusarium sp. | A0A366S7J1 |
| Fusarium sp. | A0A366S6E2 |
| Fusarium sp. | A0A366RU34 |
| Fusarium sp. | A0A366SAJ5 |
| Fusarium sp. | A0A366S5H5 |
| Fusarium sp. | A0A366RLX0 |
| Fusarium sp. | A0A366RZB1 |
| Fusarium sp. | A0A366RSR3 |
| Fusarium sp. | A0A366S6U1 |
| Fusarium sp. | A0A366SA36 |
| Fusarium sp. | A0A366S608 |
| Fusarium sp. | A0A366R259 |
| Fusarium sp. | A0A090N5T3 |
| Fusarium sp. | A0A090MBD1 |
| Fusarium sp. | A0A090N5N0 |
| Fusarium sp. | W9I3C2 |
| Fusarium sp. | W9HY89 |
| Fusarium sp. | W9IHK5 |
| Fusarium sp. | W9HQZ2 |
| Fusarium sp. | W9IYB6 |
| Fusarium sp. | W9I981 |

| | |
|---|---|
| Fusarium sp. | W9IXH4 |
| Fusarium sp. | W9IFQ1 |
| Fusarium sp. | W9HPW8 |
| Fusarium sp. | W9HU54 |
| Fusarium sp. | W9HPA2 |
| Fusarium sp. | W9J633 |
| Fusarium sp. | W9IFB4 |
| Fusarium sp. | W9J1V2 |
| Fusarium sp. | W9IC02 |
| Fusarium sp. | W9HM81 |
| Fusarium spo | A0A395SCF9 |
| Fusarium spo | A0A395SN15 |
| Fusarium spo | A0A395SBE5 |
| Fusarium spo | A0A395S6D9 |
| Fusarium spo | A0A395SH08 |
| Fusarium spo | A0A395SBA6 |
| Fusarium spo | A0A395RVT2 |
| Fusarium spo | A0A395RRP8 |
| Fusarium spo | A0A395SJC5 |
| Fusarium spo | A0A395SIC0 |
| Fusarium spo | A0A395RMJ7 |
| Fusarium spo | A0A395RYB3 |
| Fusarium spo | A0A395RVS6 |
| Fusarium spo | A0A395SNH5 |
| Fusarium spo | A0A395SDT1 |
| Fusarium spo | A0A395S6D1 |
| Fusarium spo | A0A395SDH6 |
| Fusarium ven | A0A2L2U374 |
| Fusarium ven | A0A2L2TJ64 |
| Fusarium ven | A0A2L2U3G9 |
| Fusarium ven | A0A2L2TQW0 |
| Fusarium ven | A0A2L2TFB0 |
| Fusarium ven | A0A2L2TKB7 |
| Fusarium ven | A0A2L2TJF0 |
| Fusarium ven | A0A2L2TLG9 |
| Fusarium ven | A0A2L2TPW8 |
| Fusarium ven | A0A2L2U3E5 |
| Fusarium ven | A0A2L2T6R1 |
| Fusarium ven | A0A2L2TZ15 |
| Fusarium ven | A0A2L2TR51 |
| Fusarium ven | A0A2L2T472 |

| | |
|---|---|
| Fusarium ven | A0A2L2SR66 |
| Fusarium ven | A0A2L2TU58 |
| Fusarium ven | A0A2L2T569 |
| Fusarium ven | A0A2L2U3K6 |
| Fusarium ven | A0A2L2SSM8 |
| Fusarium ven | A0A2L2TMX3 |
| Fusarium ven | A0A2L2TCF7 |
| Gaeavirus sp. | A0A3G4ZYX2 |
| Gaeumannor | J3PC95 |
| Gallionellales | A0A2M7N8H7 |
| Gallus gallus. | Q90978 |
| Gallus gallus. | Q90865 |
| Gallus gallus. | A0A1D5PAR6 |
| Gallus gallus. | F1NWX6 |
| Gallus gallus. | R4GMH5 |
| Gallus gallus. | E1C8A1 |
| Gallus gallus. | F1NUN5 |
| Gallus gallus. | A0A3Q2U5D1 |
| Gallus gallus. | Q788Q2 |
| Gambusia aff | A0A315UVV8 |
| Gambusia aff | A0A315UQL6 |
| Gambusia aff | A0A315VVX5 |
| Gambusia aff | A0A315UYF8 |
| Gambusia aff | A0A315UVQ5 |
| Gammaprote | A0A1G0KYN1 |
| Gammaprote | A0A2D5I3I9 |
| Gammaprote | A0A2D5PM94 |
| Gammaprote | A0A2E1JMT7 |
| Gasterosteus | G3N792 |
| Gasterosteus | G3NR28 |
| Gasterosteus | G3Q8D9 |
| Gasterosteus | G3NR40 |
| Gasterosteus | G3NX83 |
| Gasterosteus | G3NR48 |
| Gasterosteus | G3NX96 |
| Gasterosteus | G3N8R7 |
| Gasterosteus | G3PGI4 |
| Gavia stellata | A0A093F083 |
| Gemmobacte | A0A1H7ZTR7 |
| Gemmobacte | A0A2T6B2Z8 |
| Gemmobacte | A0A3S3UUA3 |

| | |
|---|---|
| Gemmobacte | A0A451GG88 |
| Gemmobacte | A0A1N7QCJ4 |
| Gemmobacte | A0A3P3DPT3 |
| Gemmobacte | A0A398BPI3 |
| Gemmobacte | A0A398BSF3 |
| Gemmobacte | A0A358B3Q1 |
| Genlisea aure | S8CSG8 |
| Genlisea aure | S8BXM1 |
| Genlisea aure | S8ECB0 |
| Geotrichum c | A0A0J9XCZ4 |
| Gibberella fuj | S0E9M1 |
| Gibberella fuj | S0EAW3 |
| Gibberella fuj | S0E050 |
| Gibberella fuj | S0DSB2 |
| Gibberella fuj | S0E1P0 |
| Gibberella fuj | S0DJL6 |
| Gibberella fuj | S0ECM0 |
| Gibberella fuj | S0DSY7 |
| Gibberella fuj | S0E5X7 |
| Gibberella fuj | S0EAP9 |
| Gibberella fuj | S0DSC6 |
| Gibberella fuj | S0EMX3 |
| Gibberella fuj | S0E357 |
| Gibberella fuj | S0E2Q5 |
| Gibberella fuj | S0ECL2 |
| Gibberella fuj | S0E6U9 |
| Gibberella fuj | S0DJP2 |
| Gibberella fuj | S0DZF4 |
| Gibberella fuj | S0EFD2 |
| Gibberella fuj | S0E497 |
| Gibberella fuj | A0A0I9Z138 |
| Gibberella fuj | A0A0I9YAH3 |
| Gibberella fuj | A0A0I9XHA9 |
| Gibberella fuj | A0A0I9Y8Q9 |
| Gibberella fuj | A0A0I9YM17 |
| Gibberella fuj | A0A0I9XFP3 |
| Gibberella fuj | A0A0J0BUY0 |
| Gibberella fuj | A0A2H3RKI8 |
| Gibberella fuj | A0A2H3SN28 |
| Gibberella fuj | A0A0J0CHN8 |
| Gibberella fuj | A0A2H3RR38 |

| | |
|---|---|
| Gibberella fuj | A0A2H3SWB6 |
| Gibberella fuj | A0A0I9X760 |
| Gibberella fuj | A0A2H3S7W2 |
| Gibberella fuj | A0A0J0CQW4 |
| Gibberella fuj | A0A2H3S359 |
| Gibberella fuj | A0A2H3SM63 |
| Gibberella fuj | A0A2H3RNJ8 |
| Gibberella fuj | A0A2H3RMB7 |
| Gibberella fuj | A0A2H3SWM1 |
| Gibberella fuj | A0A0J0CT57 |
| Gibberella fuj | A0A2H3RRY9 |
| Gibberella fuj | A0A0I9Y942 |
| Gibberella fuj | A0A2H3RGR6 |
| Gibberella fuj | A0A0I9XEV3 |
| Gibberella fuj | A0A2H3RUA2 |
| Gibberella fuj | A0A0I9XAZ3 |
| Gibberella fuj | A0A0J0BYF6 |
| Gibberella fuj | A0A2H3RVF2 |
| Gibberella fuj | A0A2H3RW12 |
| Gibberella fuj | A0A0I9XY78 |
| Gibberella fuj | A0A0I9YHM8 |
| Gibberella fuj | A0A2H3RR57 |
| Gibberella fuj | A0A0I9X911 |
| Gibberella fuj | A0A0I9XJR0 |
| Gibberella fuj | A0A0J0C013 |
| Gibberella fuj | A0A2H3RVE0 |
| Gibberella fuj | A0A2H3SQE6 |
| Gibberella fuj | A0A0I9XAJ6 |
| Gibberella fuj | A0A2H3SHG6 |
| Gibberella fuj | A0A2H3R980 |
| Gibberella fuj | A0A0I9XDN8 |
| Gibberella int | A0A420SS68 |
| Gibberella int | A0A420TP29 |
| Gibberella int | A0A420UB76 |
| Gibberella int | A0A365NIV2 |
| Gibberella int | A0A365N6M1 |
| Gibberella int | A0A365MMV3 |
| Gibberella int | A0A365N4F2 |
| Gibberella int | A0A365NNN5 |
| Gibberella int | A0A365ND98 |
| Gibberella int | A0A420T6U9 |

| | |
|---|---|
| Gibberella int | A0A365NJW8 |
| Gibberella int | A0A420SEE3 |
| Gibberella int | A0A365N416 |
| Gibberella int | A0A365NIH4 |
| Gibberella int | A0A420SZ33 |
| Gibberella int | A0A365MQF6 |
| Gibberella int | A0A420UA77 |
| Gibberella int | A0A365MQ14 |
| Gibberella int | A0A365MMJ0 |
| Gibberella int | A0A420SDB1 |
| Gibberella int | A0A365MPJ1 |
| Gibberella int | A0A365N099 |
| Gibberella int | A0A365NDF6 |
| Gibberella int | A0A420SSE2 |
| Gibberella int | A0A420T6Q6 |
| Gibberella int | A0A420U9Y0 |
| Gibberella int | A0A365N1D1 |
| Gibberella int | A0A420T9A6 |
| Gibberella int | A0A365N275 |
| Gibberella int | A0A365MYB1 |
| Gibberella int | A0A420TU20 |
| Gibberella int | A0A420TTT9 |
| Gibberella int | A0A365MRF5 |
| Gibberella int | A0A365MKN4 |
| Gibberella int | A0A420SGE2 |
| Gibberella int | A0A420TM01 |
| Gibberella int | A0A420SQ44 |
| Gibberella int | A0A365N043 |
| Gibberella int | A0A365NGM1 |
| Gibberella int | A0A420SEZ4 |
| Gibberella int | A0A420SPD6 |
| Gibberella int | A0A420SWC9 |
| Gibberella int | A0A365MMK9 |
| Gibberella mo | W7M0V8 |
| Gibberella mo | W7MQW8 |
| Gibberella mo | W7LIK3 |
| Gibberella mo | W7N541 |
| Gibberella mo | A0A139YBX2 |
| Gibberella mo | W7MTU5 |
| Gibberella mo | W7NCD1 |
| Gibberella mo | W7N6Q6 |

| | |
|---|---|
| Gibberella m | W7MUN6 |
| Gibberella m | W7ME54 |
| Gibberella m | W7MSX0 |
| Gibberella m | W7MR98 |
| Gibberella m | W7MZZ1 |
| Gibberella m | W7N269 |
| Gibberella m | W7N008 |
| Gibberella m | W7LSV5 |
| Gibberella m | W7N837 |
| Gibberella m | W7LEM9 |
| Gibberella m | W7MXJ6 |
| Gibberella m | A0A139YBM8 |
| Gibberella m | W7M9D1 |
| Gibberella m | W7MZW8 |
| Gibberella m | A0A366Q8Q7 |
| Gibberella m | A0A366Q909 |
| Gibberella m | A0A366R737 |
| Gibberella m | A0A366S001 |
| Gibberella m | A0A366QHY9 |
| Gibberella m | A0A366PSC3 |
| Gibberella m | A0A366R685 |
| Gibberella m | A0A366NS06 |
| Gibberella m | A0A366NR83 |
| Gibberella m | A0A366RE72 |
| Gibberella m | A0A366P026 |
| Gibberella m | A0A366P298 |
| Gibberella m | A0A366Q4J4 |
| Gibberella m | A0A366QZP0 |
| Gibberella m | A0A366Q9V0 |
| Gibberella m | A0A366PGF9 |
| Gibberella m | A0A366RYF1 |
| Gibberella m | A0A366RSA3 |
| Gibberella m | A0A366QJB0 |
| Gibberella m | A0A366RE50 |
| Gibberella m | A0A366QC02 |
| Gibberella m | A0A366PJS9 |
| Gibberella m | A0A366PTB2 |
| Gibberella m | A0A366Q4I7 |
| Gibberella m | A0A366QQ61 |
| Gibberella m | A0A366Q3H7 |
| Gibberella m | A0A366Q2F5 |

| | |
|---|---|
| Gibberella m( | A0A366RV94 |
| Gibberella m( | A0A366RKF4 |
| Gibberella m( | A0A366PYT4 |
| Gibberella m( | A0A366PSP3 |
| Gibberella m( | A0A366R173 |
| Gibberella m( | A0A366RE58 |
| Gibberella m( | A0A366QPK3 |
| Gibberella m( | A0A366PGR0 |
| Gibberella m( | A0A366R225 |
| Gibberella ny | A0A2K0W5D0 |
| Gibberella ny | A0A2K0UTI6 |
| Gibberella ny | A0A2K0W931 |
| Gibberella ny | A0A2K0WC09 |
| Gibberella ny | A0A2K0UTH2 |
| Gibberella ny | A0A2K0WV95 |
| Gibberella ny | A0A2K0WIM1 |
| Gibberella ny | A0A2K0W2B0 |
| Gibberella ny | A0A2K0W2K6 |
| Gibberella ny | A0A2K0VVG2 |
| Gibberella ny | A0A2K0W1N5 |
| Gibberella ny | A0A2K0UTG7 |
| Gibberella ny | A0A2K0W8T1 |
| Gibberella ny | A0A2K0W365 |
| Gibberella ny | A0A2K0VVT0 |
| Gibberella ny | A0A2K0WAU9 |
| Gibberella ny | A0A2K0W3Y3 |
| Gibberella ze | I1S305 |
| Gibberella ze | A0A098DHT5 |
| Gibberella ze | A0A098DJ50 |
| Gibberella ze | I1RTL3 |
| Gibberella ze | I1S3P5 |
| Gibberella ze | A0A098DZ54 |
| Gibberella ze | I1RLE4 |
| Gibberella ze | I1S5T9 |
| Gibberella ze | I1RTD2 |
| Gibberella ze | I1RV98 |
| Gibberella ze | I1S3P4 |
| Gibberella ze | A0A098D7I4 |
| Gibberella ze | A0A098DJX2 |
| Gibberella ze | A0A098DZM5 |
| Gibberella ze | I1RA55 |

| | |
|---|---|
| Gibberella ze | I1RXM0 |
| Gibberella ze | I1R9J0 |
| Gibberella ze | I1RKL8 |
| Gibberella ze | I1S2Z6 |
| Gibberella ze | A0A1C3YKN6 |
| Gibberella ze | A0A098E223 |
| Gibberella ze | A0A1C3YHH6 |
| Gibberella ze | A0A1C3YKD6 |
| Gibberella ze | A0A1C3YKB1 |
| Gibberella ze | A0A2H3HMC5 |
| Gibberella ze | A0A2H3GRJ6 |
| Gibberella ze | A0A2H3HSA2 |
| Gibberella ze | A0A2H3GYF5 |
| Gibberella ze | A0A2H3FZY4 |
| Gibberella ze | A0A2H3HDH3 |
| Gibberella ze | A0A2H3G941 |
| Gibberella ze | A0A2H3H4N8 |
| Gibberella ze | A0A2H3GNB0 |
| Gibberella ze | A0A2H3GIG1 |
| Gibberella ze | A0A2H3FML8 |
| Gibberella ze | A0A2H3G768 |
| Gibberella ze | A0A2H3H9N3 |
| Gibberella ze | A0A2H3HFP0 |
| Gibberella ze | A0A2H3GGM8 |
| Gibberella ze | A0A2H3FHF7 |
| Gibberella ze | A0A2H3GHH8 |
| Gigaspora ros | A0A397VR36 |
| Gigaspora ros | A0A397W4J7 |
| Gigaspora ros | A0A397VYK2 |
| Glarea lozoye | S3D022 |
| Glarea lozoye | S3CIG6 |
| Glarea lozoye | S3D8B4 |
| Glarea lozoye | S3DNA8 |
| Glarea lozoye | H0EQA9 |
| Globisporang | K3WFA2 |
| Globisporang | K3WFA4 |
| Globisporang | K3WN51 |
| Globisporang | K3WWN0 |
| Globisporang | K3WCW4 |
| Globisporang | K3W5W5 |
| Globisporang | K3WZR8 |

| | |
|---|---|
| Globisporang | K3WJD0 |
| Globisporang | K3WPX5 |
| Globisporang | K3WPX2 |
| Globisporang | K3WZD7 |
| Globisporang | K3WD79 |
| Globisporang | K3XBK6 |
| Globisporang | K3WN48 |
| Globisporang | K3WWN3 |
| Globisporang | K3WI68 |
| Globisporang | K3WRX0 |
| Globisporang | K3WI73 |
| Globisporang | K3WCV9 |
| Globisporang | K3WY68 |
| Globisporang | K3WCW1 |
| Globisporang | K3WPX3 |
| Globisporang | K3X667 |
| Globisporang | K3WHY8 |
| Globisporang | K3WGZ7 |
| Globisporang | K3WCW6 |
| Globisporang | K3WI69 |
| Globisporang | K3WI70 |
| Globisporang | K3WRY9 |
| Globisporang | K3XDL0 |
| Globisporang | K3WCV8 |
| Globisporang | K3WP93 |
| Globisporang | K3WPX4 |
| Globisporang | K3WGZ2 |
| Globisporang | K3WCW3 |
| Globisporang | K3WCU2 |
| Globisporang | K3WCU9 |
| Globisporang | K3WK48 |
| Globisporang | K3WFA3 |
| Globodera pa | A0A183BI04 |
| Globodera pa | A0A183CL17 |
| Globodera pa | A0A183BNL9 |
| Globodera pa | A0A183CQJ0 |
| Globodera pa | A0A183CDV8 |
| Globodera pa | A0A183CRD4 |
| Globodera pa | A0A183CBM9 |
| Globodera pa | A0A183BUJ7 |
| Globodera pa | A0A183BWP9 |

| | |
|---|---|
| Globodera pa | A0A183CDW0 |
| Globodera pa | A0A183CBA0 |
| Globodera pa | A0A183BWW1 |
| Glossina aust | A0A1A9VE33 |
| Glossina aust | A0A1A9UM99 |
| Glossina aust | A0A1A9UMI6 |
| Glossina aust | A0A1A9UK14 |
| Glossina aust | A0A1A9UT06 |
| Glossina aust | A0A1A9UM92 |
| Glossina aust | A0A1A9UV32 |
| Glossina brev | A0A1A9WZX2 |
| Glossina brev | A0A1A9WYV2 |
| Glossina brev | A0A1A9X167 |
| Glossina brev | A0A1A9X2J2 |
| Glossina brev | A0A1A9WP94 |
| Glossina brev | A0A1A9X5N1 |
| Glossina brev | A0A1A9X170 |
| Glossina brev | A0A1A9W7X4 |
| Glossina fusc | A0A1A9XI74 |
| Glossina fusc | A0A1A9XI75 |
| Glossina fusc | A0A1A9YJ04 |
| Glossina fusc | A0A1A9XI83 |
| Glossina fusc | A0A1A9XZJ7 |
| Glossina fusc | A0A1A9XDF9 |
| Glossina fusc | A0A1A9XLY9 |
| Glossina fusc | A0A1A9Y8K0 |
| Glossina mor | A0A1B0FQZ0 |
| Glossina mor | A0A1B0G5Z0 |
| Glossina mor | A0A1B0FMH8 |
| Glossina mor | A0A1B0F9A8 |
| Glossina mor | A0A1B0FN27 |
| Glossina mor | A0A1B0FJ31 |
| Glossina mor | A0A1B0FF31 |
| Glossina palli | A0A1B0ACN4 |
| Glossina palli | A0A1A9ZN42 |
| Glossina palli | A0A1B0AFV3 |
| Glossina palli | A0A1A9Z7T1 |
| Glossina palli | A0A1B0ACN9 |
| Glossina palli | A0A1A9Z0F1 |
| Glossina palli | A0A1B0ACN0 |
| Glossina palli | A0A1B0AAP1 |

| | |
|---|---|
| Glossina palp | A0A1B0AYB5 |
| Glossina palp | A0A1B0B913 |
| Glossina palp | A0A1B0BQV5 |
| Glossina palp | A0A1B0BRF1 |
| Glossina palp | A0A1B0AMM5 |
| Glossina palp | A0A1B0B615 |
| Glossina palp | A0A1B0BSL7 |
| Glycine max.) | C6ZRQ3 |
| Glycine max.) | C6ZS29 |
| Glycine max.) | K7L2L4 |
| Glycine max.) | I1L742 |
| Glycine max.) | A0A0R0EVN8 |
| Glycine max.) | I1KE73 |
| Glycine max.) | I1LSR9 |
| Glycine max.) | I1LSR6 |
| Glycine max.) | A0A0R0KG24 |
| Glycine max.) | K7M8Z4 |
| Glycine max.) | K7KXE1 |
| Glycine max.) | I1LSX3 |
| Glycine max.) | I1MIG9 |
| Glycine max.) | A0A0R0HEP2 |
| Glycine max.) | K7KXI5 |
| Glycine max.) | A0A0R0H4D6 |
| Glycine max.) | I1ME78 |
| Glycine max.) | K7KXH0 |
| Glycine max.) | K7LCV5 |
| Glycine max.) | I1MI78 |
| Glycine max.) | K7M2T9 |
| Glycine max.) | I1LST1 |
| Glycine max.) | A0A0R0L9K9 |
| Glycine max.) | K7M1S0 |
| Glycine max.) | K7K3A5 |
| Glycine max.) | I1KZA4 |
| Glycine max.) | K7M1R9 |
| Glycine max.) | K7KBW2 |
| Glycine max.) | A0A0R0FWY9 |
| Glycine max.) | K7LU18 |
| Glycine max.) | A0A0R0IPR8 |
| Glycine max.) | I1KE80 |
| Glycine max.) | A0A0R0JLU1 |
| Glycine max.) | K7K3A6 |

| | |
|---|---|
| Glycine max.) | I1K7Y3 |
| Glycine max.) | K7LU19 |
| Glycine max.) | K7LUX6 |
| Glycine max.) | I1M2F1 |
| Glycine max.) | A0A0R0IHT4 |
| Glycine max.) | K7K9A9 |
| Glycine max.) | I1KEB9 |
| Glycine max.) | A0A0R0GW83 |
| Glycine max.) | A0A0R0JNK4 |
| Glycine max.) | K7LUW1 |
| Glycine max.) | K7KXF2 |
| Glycine max.) | I1LSY1 |
| Glycine max.) | A0A0R0KJ29 |
| Glycine max.) | K7KXI4 |
| Glycine max.) | I1LGT5 |
| Glycine max.) | K7MZ69 |
| Glycine max.) | A0A0R0HDR1 |
| Glycine max.) | I1KEC5 |
| Glycine max.) | I1KE67 |
| Glycine max.) | K7M2F6 |
| Glycine max.) | C6ZRW7 |
| Glycine max.) | A0A0R0IRG6 |
| Glycine max.) | K7M2E9 |
| Glycine max.) | A0A0R0H8A8 |
| Glycine max.) | I1KE60 |
| Glycine max.) | A0A0R0JLS9 |
| Glycine max.) | K7M8T0 |
| Glycine max.) | K7MCU5 |
| Glycine max.) | K7KXH6 |
| Glycine max.) | A0A0R0HHK1 |
| Glycine max.) | A0A0R0H240 |
| Glycine max.) | K7L050 |
| Glycine max.) | I1M2E7 |
| Glycine max.) | K7LAQ9 |
| Glycine max.) | A0A0R0HUZ3 |
| Glycine max.) | A0A0R0HHN2 |
| Glycine max.) | A0A0R0H5B6 |
| Glycine max.) | K7LAS7 |
| Glycine max.) | I1KEA9 |
| Glycine max.) | A0A0R0HHS7 |
| Glycine max.) | A0A0R0KYQ5 |

| | |
|---|---|
| Glycine max.) | A0A0R0JT19 |
| Glycine max.) | K7LUV4 |
| Glycine max.) | K7KXC8 |
| Glycine max.) | A0A0R0HFB3 |
| Glycine max.) | K7M046 |
| Glycine max.) | A0A0R0HDU4 |
| Glycine max.) | I1MKV1 |
| Glycine max.) | A0A0R0JM62 |
| Glycine max.) | A0A0R0JM46 |
| Glycine max.) | K7M1R8 |
| Glycine max.) | I1MI79 |
| Glycine max.) | K7M8Z5 |
| Glycine max.) | A0A0R0H590 |
| Glycine max.) | A0A0R0EG65 |
| Glycine max.) | K7MCU6 |
| Glycine max.) | A0A0R0J530 |
| Glycine max.) | A0A0R0HEH1 |
| Glycine max.) | A0A0R0JRN4 |
| Glycine max.) | K7LUW3 |
| Glycine max.) | I1JTI1 |
| Glycine max.) | K7MCU7 |
| Glycine max.) | A0A0R0JRK8 |
| Glycine max.) | K7M2F1 |
| Glycine max.) | A0A0R0INR7 |
| Glycine max.) | A0A0R0JLL7 |
| Glycine max.) | K7K3A1 |
| Glycine max.) | A0A0R0FWT7 |
| Glycine max.) | A0A0R0H6A2 |
| Glycine max.) | K7KXD7 |
| Glycine max.) | K7LQJ4 |
| Glycine max.) | I1KZA6 |
| Glycine max.) | A0A0R0EHM4 |
| Glycine max.) | K7LCV6 |
| Glycine max.) | A0A0R0JLQ4 |
| Glycine max.) | A0A0R0FXR9 |
| Glycine max.) | I1LSY2 |
| Glycine max.) | K7KXH7 |
| Glycine max.) | A0A0R0H7L6 |
| Glycine max.) | K7LUX4 |
| Glycine max.) | I1M2F0 |
| Glycine max.) | A0A0R0H5I9 |

| | |
|---|---|
| Glycine max.) | A0A0R0K7P2 |
| Glycine max.) | A0A0R0GTK5 |
| Glycine max.) | K7KBW3 |
| Glycine max.) | I1KE90 |
| Glycine max.) | K7KXI2 |
| Glycine max.) | A0A0R0H043 |
| Glycine max.) | K7M1R3 |
| Glycine max.) | I1KF87 |
| Glycine max.) | K7KDF6 |
| Glycine max.) | K7KXH2 |
| Glycine max.) | A0A0R0JLC9 |
| Glycine max.) | I1M3F1 |
| Glycine max.) | A0A0R0JVL9 |
| Glycine max.) | A0A0R0HEE2 |
| Glycine max.) | A0A0R0HFQ6 |
| Glycine max.) | A0A0R0G425 |
| Glycine max.) | K7KXF3 |
| Glycine max.) | A0A0R0H539 |
| Glycine max.) | I1KE88 |
| Glycine max.) | I1KE68 |
| Glycine max.) | A0A0R0J260 |
| Glycine max.) | I1LSW7 |
| Glycine max.) | I1M5U0 |
| Glycine max.) | K7KI11 |
| Glycine max.) | I1KE70 |
| Glycine max.) | I1LAB6 |
| Glycine max.) | A0A0R0J0N7 |
| Glycine max.) | K7LAS8 |
| Glycine max.) | I1KKX7 |
| Glycine max.) | K7KXE2 |
| Glycine max.) | A0A0R0KD14 |
| Glycine max.) | K7MCT9 |
| Glycine max.) | I1LSX8 |
| Glycine max.) | K7LAQ6 |
| Glycine max.) | A0A0R0G5B0 |
| Glycine max.) | K7KXH9 |
| Glycine max.) | A0A0R0JLZ5 |
| Glycine max.) | A0A0R0H884 |
| Glycine max.) | K7L8G9 |
| Glycine max.) | A0A0R0FXA7 |
| Glycine max.) | I1KZ76 |

| | |
|---|---|
| Glycine max.) | K7LUV3 |
| Glycine max.) | A0A0R0IC73 |
| Glycine max.) | K7LUY2 |
| Glycine max.) | K7KXH5 |
| Glycine max.) | K7LVZ6 |
| Glycine max.) | I1KZ77 |
| Glycine max.) | K7M2F0 |
| Glycine max.) | I1JNE6 |
| Glycine max.) | A0A0R0EWT5 |
| Glycine max.) | K7K3A2 |
| Glycine max.) | I1MNZ4 |
| Glycine max.) | K7M1R5 |
| Glycine max.) | A0A0R0JTU4 |
| Glycine max.) | I1M2E9 |
| Glycine max.) | K7MYD3 |
| Glycine max.) | I1JPX6 |
| Glycine max.) | K7LVZ4 |
| Glycine max.) | A0A0R0H5T0 |
| Glycine max.) | I1LSR5 |
| Glycine max.) | I1KEB0 |
| Glycine max.) | K7LUW9 |
| Glycine max.) | K7M1R6 |
| Glycine max.) | I1KQR8 |
| Glycine max.) | A0A0R0H5Y2 |
| Glycine max.) | I1M2E5 |
| Glycine max.) | K7LVZ3 |
| Glycine max.) | K7LUS0 |
| Glycine max.) | I1LLA6 |
| Glycine max.) | K7KXD3 |
| Glycine max.) | A0A0R0JLM5 |
| Glycine max.) | K7KD19 |
| Glycine max.) | C6ZRU8 |
| Glycine max.) | K7MG77 |
| Glycine max.) | I1LSY4 |
| Glycine max.) | A0A0R0GV60 |
| Glycine max.) | I1KEB1 |
| Glycine max.) | A0A0R0H7N3 |
| Glycine max.) | A0A0R0G5X7 |
| Glycine max.) | A0A0R0H529 |
| Glycine max.) | A0A0R0HEQ7 |
| Glycine max.) | K7LUV6 |

| | |
|---|---|
| Glycine max.› | A0A0R0JRU4 |
| Glycine max.› | I1KEB7 |
| Glycine max.› | A0A0R0H5H4 |
| Glycine max.› | A0A0R0L9J9 |
| Glycine max.› | A0A0R0H871 |
| Glycine max.› | A0A0R0FNK4 |
| Glycine max.› | K7LUV7 |
| Glycine max.› | A0A0R0HHZ0 |
| Glycine max.› | A0A0R0JLR9 |
| Glycine max.› | K7KD21 |
| Glycine max.› | K7M1R7 |
| Glycine max.› | A0A0R0GTN5 |
| Glycine max.› | K7KXI0 |
| Glycine max.› | K7MCU9 |
| Glycine max.› | K7M1R2 |
| Glycine max.› | I1KE64 |
| Glycine max.› | A0A0R0LIX3 |
| Glycine max.› | K7KT18 |
| Glycine max.› | K7LUW8 |
| Glycine max.› | I1KEA7 |
| Glycine max.› | A0A0R0H5K4 |
| Glycine max.› | A0A0R0HFT7 |
| Glycine max.› | A0A0R0JT32 |
| Glycine max.› | A0A368UHR2 |
| Glycine max.› | K7KXH1 |
| Glycine soja.› | A0A445IKH7 |
| Glycine soja.› | A0A445K4J5 |
| Glycine soja.› | A0A445IBB4 |
| Glycine soja.› | A0A445IC32 |
| Glycine soja.› | A0A445IBG5 |
| Glycine soja.› | A0A445KEL5 |
| Glycine soja.› | A0A445I9N1 |
| Glycine soja.› | A0A445IA22 |
| Glycine soja.› | A0A445KEC4 |
| Glycine soja.› | A0A0B2R8A6 |
| Glycine soja.› | A0A0B2RTJ3 |
| Glycine soja.› | A0A0B2R410 |
| Glycine soja.› | A0A0B2RYE8 |
| Glycine soja.› | A0A0B2R3Z9 |
| Glycine soja.› | A0A0B2NQG4 |
| Glycine soja.› | A0A0B2SR89 |

Glycine soja.› A0A0B2SHQ9
Glycine soja.› A0A0B2Q0P0
Glycine soja.› A0A0B2QFX4
Glycine soja.› A0A445LDU1
Glycine soja.› A0A0B2PJD4
Glycine soja.› A0A0B2PLC0
Glycine soja.› A0A0B2R8Z3
Glycine soja.› A0A0B2P2R6
Glycine soja.› A0A0B2SNF0
Glycine soja.› A0A0B2QYM0
Glycine soja.› A0A445KEJ3
Glycine soja.› A0A0B2SG46
Glycine soja.› A0A0B2SPZ0
Glycine soja.› A0A0B2NXR2
Glycine soja.› A0A0B2S035
Glycine soja.› A0A0B2S336
Glycine soja.› A0A0B2SSB0
Glycine soja.› A0A445GWF0
Glycine soja.› A0A445IFF3
Glycine soja.› A0A445HSY4
Glycine soja.› A0A445IB72
Glycine soja.› A0A0B2RZB0
Glycine soja.› A0A445IBT0
Glycine soja.› A0A0B2SIU2
Glycine soja.› A0A0B2PEM2
Glycine soja.› A0A0B2PA07
Glycine soja.› A0A0B2RQ81
Glycine soja.› A0A0B2QDD9
Glycine soja.› A0A0B2NSG5
Glycine soja.› A0A0B2PTL8
Glycine soja.› A0A0B2PCY8
Glycine soja.› A0A0B2QIS7
Glycine soja.› A0A0B2PHY5
Glycine soja.› A0A0B2QXQ7
Glycine soja.› A0A0B2SRB5
Glycine soja.› A0A0B2QH24
Glycine soja.› A0A0B2QEU6
Glycine soja.› A0A0B2SLN0
Glycine soja.› A0A0B2NSK2
Glycine soja.› A0A0B2PEM8
Glycine soja.› A0A0B2PD35

Glycine soja.› A0A0B2R1I6
Glycine soja.› A0A445L056
Glycine soja.› A0A445IBD0
Glycine soja.› A0A445KEK2
Glycine soja.› A0A0B2R2T4
Glycine soja.› A0A445JAL8
Glycine soja.› A0A445HQ61
Glycine soja.› A0A445JPE3
Glycine soja.› A0A445FGF3
Glycine soja.› A0A445M1W2
Glycine soja.› A0A445HN76
Glycine soja.› A0A445IA45
Glycine soja.› A0A445HQC5
Glycine soja.› A0A445HQE3
Glycine soja.› A0A445JV37
Glycine soja.› A0A445HPY3
Glycine soja.› A0A445KEI0
Glycine soja.› A0A445IB85
Glycine soja.› A0A445KE92
Glycine soja.› A0A445HPU3
Glycine soja.› A0A445HQB7
Glycine soja.› A0A445KEQ7
Glycine soja.› A0A445KVH1
Glycine soja.› A0A445KEH0
Glycine soja.› A0A0B2RLN3
Glycine soja.› A0A445KEG9
Glycine soja.› A0A445KEI1
Glycine soja.› A0A445L055
Glycine soja.› A0A445I607
Glycine soja.› A0A445HPX9
Glycine soja.› A0A445JAP1
Glycine soja.› A0A0B2QGK7
Glycine soja.› A0A445KF02
Glycine soja.› A0A445GWE9
Glycine soja.› A0A445JAK5
Glycine soja.› A0A445KV70
Glycine soja.› A0A445GGC1
Glycine soja.› A0A445L8H0
Glycine soja.› A0A445HSS9
Glycine soja.› A0A445KZL6
Glycine soja.› A0A445IZ56

| | |
|---|---|
| Glycine soja.› | A0A445KF53 |
| Glycine soja.› | A0A0B2PCH0 |
| Glycine soja.› | A0A0B2QFX9 |
| Glycine soja.› | A0A0B2P0P0 |
| Glycine soja.› | A0A0B2RPE1 |
| Glycine soja.› | A0A0B2NX34 |
| Glycine soja.› | A0A0B2P9T3 |
| Glycine soja.› | A0A0B2S2G1 |
| Glycine soja.› | A0A445I3H0 |
| Glycine soja.› | A0A445HXA0 |
| Glycine soja.› | A0A445KEJ2 |
| Glycine soja.› | A0A0B2RKG5 |
| Glycine soja.› | A0A0B2QGL4 |
| Glycine soja.› | A0A0B2P4X3 |
| Glycine soja.› | A0A445IZ50 |
| Glycine soja.› | A0A445IZ45 |
| Glycine soja.› | A0A445HPU8 |
| Glycine soja.› | A0A445IA11 |
| Glycine soja.› | A0A445KED2 |
| Glycine soja.› | A0A445KEI5 |
| Glycine soja.› | A0A445I9L6 |
| Glycine soja.› | A0A445IBA1 |
| Glycine soja.› | A0A445L054 |
| Glycine soja.› | A0A445GQ74 |
| Glycine soja.› | A0A445HQD2 |
| Glycine soja.› | A0A445GQ03 |
| Glycine soja.› | A0A445HQD8 |
| Glycine soja.› | A0A445HQC1 |
| Glycine soja.› | A0A445HQK7 |
| Glycine soja.› | A0A445HST5 |
| Glycine soja.› | A0A445KF01 |
| Glycine soja.› | A0A445HPX5 |
| Glycine soja.› | A0A445JPF3 |
| Glycine soja.› | A0A445KEN4 |
| Glycine soja.› | A0A445KEU4 |
| Glycine soja.› | A0A445IB73 |
| Glycine soja.› | A0A445FIR5 |
| Glycine soja.› | A0A445IBH5 |
| Glycine soja.› | A0A445GWZ7 |
| Glycine soja.› | A0A445I9N6 |
| Glycine soja.› | A0A445JPH7 |

| | |
|---|---|
| Glycine soja.› | A0A445HQ82 |
| Glycine soja.› | A0A445KET6 |
| Glycine soja.› | A0A445KVD1 |
| Glycine soja.› | A0A445L5S4 |
| Glycine soja.› | A0A445IBB3 |
| Glycine soja.› | A0A445LQP4 |
| Glycine soja.› | A0A0B2PDS8 |
| Glycine soja.› | A0A0B2QBU1 |
| Glycine soja.› | A0A445KEM0 |
| Glycine soja.› | A0A445HQA5 |
| Glycine soja.› | A0A445K4C6 |
| Glycine soja.› | A0A445I9M8 |
| Glycine soja.› | A0A445HQG0 |
| Glycine soja.› | A0A445JPL0 |
| Glycine soja.› | A0A445JAK4 |
| Glycine soja.› | A0A445KEF7 |
| Glycine soja.› | A0A445JPB5 |
| Glycine soja.› | A0A0B2RX25 |
| Glycine soja.› | A0A0B2SJF3 |
| Glycine soja.› | A0A0B2SL58 |
| Glycine soja.› | A0A0B2QH31 |
| Glycine soja.› | A0A445FNQ5 |
| Glycine soja.› | A0A445HPS5 |
| Glycine soja.› | A0A445KEW3 |
| Glycine soja.› | A0A445I3X8 |
| Glycine soja.› | A0A445FNS9 |
| Glycine soja.› | A0A445LBW9 |
| Glycine soja.› | A0A445L5L3 |
| Glycine soja.› | A0A445HQ49 |
| Glycine soja.› | A0A445HQ40 |
| Glycine soja.› | A0A445HPZ2 |
| Glycine soja.› | A0A445KED1 |
| Glycine soja.› | A0A445HQK9 |
| Glycine soja.› | A0A445HSU7 |
| Glycine soja.› | A0A445IB74 |
| Glycine soja.› | A0A445GQ81 |
| Glycine soja.› | A0A445KEK1 |
| Glycine soja.› | A0A445KEA4 |
| Glycine soja.› | A0A445HSU4 |
| Glycine soja.› | A0A445HN83 |
| Glycine soja.› | A0A0B2R9H1 |

| | |
|---|---|
| Glycine soja.› | A0A0B2PX39 |
| Glycine soja.› | A0A0B2PQ09 |
| Glycine soja.› | A0A0B2R9G6 |
| Glycine soja.› | A0A0B2NXL9 |
| Glycine soja.› | A0A0B2RS26 |
| Glycine soja.› | A0A0B2QYI7 |
| Glycine soja.› | A0A0B2QVJ9 |
| Glycine soja.› | A0A0B2NXT7 |
| Glycine soja.› | A0A0B2QCR2 |
| Glycine soja.› | A0A0B2PX80 |
| Glycine soja.› | A0A445KE89 |
| Glycine soja.› | A0A445HQ13 |
| Glycine soja.› | A0A445GQ70 |
| Glycine soja.› | A0A445KES8 |
| Glycine soja.› | A0A445HQG2 |
| Glycine soja.› | A0A0B2SMR2 |
| Glycine soja.› | A0A0B2R2T0 |
| Glycine soja.› | A0A445JAN3 |
| Glycine soja.› | A0A445KED5 |
| Glycine soja.› | A0A0B2RR49 |
| Glycine soja.› | A0A445JPM6 |
| Glycine soja.› | A0A445HQ19 |
| Glycine soja.› | A0A445IC34 |
| Glycine soja.› | A0A445M1S7 |
| Glycine soja.› | A0A445I3D7 |
| Glycine soja.› | A0A445JJ41 |
| Glycine soja.› | A0A0B2PNA6 |
| Glycine soja.› | A0A0B2R151 |
| Glycine soja.› | A0A445HT18 |
| Glycine soja.› | A0A445JPC2 |
| Glycine soja.› | A0A445IC64 |
| Glycine soja.› | A0A445FF01 |
| Glycine soja.› | A0A0B2QCR6 |
| Glycine soja.› | A0A445IB88 |
| Glycine soja.› | A0A445KEM6 |
| Glycine soja.› | A0A445KEJ0 |
| Glycine soja.› | A0A445F251 |
| Glycine soja.› | A0A445GJ58 |
| Glycine soja.› | A0A445HSW1 |
| Glycine soja.› | A0A445HPQ4 |
| Glycine soja.› | A0A445JPA6 |

Glycine soja.› A0A445HSU3
Glycine soja.› A0A445L7J1
Glycine soja.› A0A0B2PE82
Glycine soja.› A0A445HPU5
Glycine soja.› A0A445J754
Glycine soja.› A0A445HPW8
Glycine soja.› A0A445KEB3
Glycine soja.› A0A445GQ56
Glycine soja.› A0A445HPX8
Glycine soja.› A0A445HSU0
Glycine soja.› A0A445I9M0
Glycine soja.› A0A445HPV7
Glycine soja.› A0A445HQK5
Glycine soja.› A0A445HGW0
Glycine soja.› A0A445HQ03
Glycine soja.› A0A445K7Z2
Glycine soja.› A0A445KF30
Glycine soja.› A0A445I9P9
Glycine soja.› A0A445HPY4
Glycine soja.› A0A445JP99
Glycine soja.› A0A445JYL2
Glycine soja.› A0A445KEJ4
Glycine soja.› A0A445I9P3
Glycine soja.› A0A445L7J0
Glycine soja.› A0A445JB06
Glycine soja.› A0A445KEB9
Glycine soja.› A0A445IB86
Glycine soja.› A0A445HQF8
Glycine soja.› A0A445GWL4
Glycine soja.› A0A445L9S1
Glycine soja.› A0A445HSV5
Glycine soja.› A0A445KEE3
Glycine soja.› A0A445KEX3
Glycine soja.› A0A445JTF8
Glycine soja.› A0A445JB16
Glycine soja.› A0A445KET4
Glycine soja.› A0A445JXZ8
Glycine soja.› A0A0B2SV99
Glycine soja.› A0A0B2PTA3
Glycine soja.› A0A445GQ60
Glycine soja.› A0A445HPZ9

| | |
|---|---|
| Glycine soja.› | A0A445HQE5 |
| Glycine soja.› | A0A445JPB6 |
| Glycine soja.› | A0A445JPB2 |
| Glycine soja.› | A0A445KGH5 |
| Glycine soja.› | A0A445HQD5 |
| Glycine soja.› | A0A445JYL7 |
| Glycine soja.› | A0A445KEK3 |
| Glycine soja.› | A0A0B2RV84 |
| Glycine soja.› | A0A445IZ62 |
| Glycine soja.› | A0A445HQ72 |
| Glycine soja.› | A0A0B2NU47 |
| Glycine soja.› | A0A445M242 |
| Glycine soja.› | A0A0B2PI23 |
| Glycine soja.› | A0A0B2NVK2 |
| Glycine soja.› | A0A0B2NXP7 |
| Glycine soja.› | A0A445JPA5 |
| Glycine soja.› | A0A445KEY2 |
| Glycine soja.› | A0A445GWI7 |
| Glycine soja.› | A0A0B2RRS1 |
| Glycine soja.› | A0A445GWF9 |
| Glycine soja.› | A0A445KEJ8 |
| Glycine soja.› | A0A0B2QVY8 |
| Glycine soja.› | A0A445HPR4 |
| Glycine soja.› | A0A0B2PU27 |
| Glycine soja.› | A0A445M229 |
| Glycine soja.› | A0A0B2SLA2 |
| Glycine soja.› | A0A445GM55 |
| Glycine soja.› | A0A0B2PGJ9 |
| Glycine soja.› | A0A445GMK7 |
| Glycine soja.› | A0A0B2R815 |
| Glycine soja.› | A0A445GWF8 |
| Glycine soja.› | A0A445IC59 |
| Glycine soja.› | A0A445GM53 |
| Glycine soja.› | A0A445KEG8 |
| Glycine soja.› | A0A445HQK6 |
| Glycine soja.› | A0A445I9S8 |
| Glycine soja.› | A0A0B2R405 |
| Glycine soja.› | A0A445KEG0 |
| Gonapodya p | A0A139AN84 |
| Gonapodya p | A0A139AWG3 |
| Gongylonema | A0A3P7LQ98 |

| | |
|---|---|
| Gongylonema | A0A3P6SB28 |
| Gongylonema | A0A183E3M5 |
| Gongylonema | A0A183DSQ9 |
| Gongylonema | A0A183D9Z8 |
| Gongylonema | A0A3P6S9V6 |
| Gongylonema | A0A3P7MR12 |
| Gongylonema | A0A183D9N7 |
| Gongylonema | A0A183CUV3 |
| Gongylonema | A0A183CZ86 |
| Gongylonema | A0A183CVQ5 |
| Gongylonema | A0A183E7Y0 |
| Gongylonema | A0A183D9A1 |
| Gongylonema | A0A183CUC2 |
| Gongylonema | A0A183DYS1 |
| Gongylonema | A0A183D4T0 |
| Gongylonema | A0A183CZ38 |
| Gongylonema | A0A183EWR6 |
| Gongylonema | A0A183D6N3 |
| Gongylonema | A0A183DDG5 |
| Gongylonema | A0A183DJR2 |
| Gongylonema | A0A183D2S8 |
| Gonium pect | A0A150G4C6 |
| Gonium pect | A0A150G869 |
| Gonium pect | A0A150GQC4 |
| Gonium pect | A0A150GM43 |
| Gonium pect | A0A150GGI9 |
| Gonium pect | A0A150FY45 |
| Gonium pect | A0A150G7V9 |
| Gonium pect | A0A150G6Z0 |
| Gonium pect | A0A150GXZ1 |
| Gonium pect | A0A150GXR5 |
| Gonium pect | A0A150GAR7 |
| Gonium pect | A0A150G2N9 |
| Gonium pect | A0A150G968 |
| Gonium pect | A0A150FX17 |
| Gonium pect | A0A150G2Z2 |
| Gonium pect | A0A150GY18 |
| Gonium pect | A0A150FX43 |
| Gonium pect | A0A150G717 |
| Gonium pect | A0A150GGC8 |
| Gonium pect | A0A150GQP5 |

| | |
|---|---|
| Gonium pect | A0A150GGE4 |
| Gonium pect | A0A150G565 |
| Gonium pect | A0A150G8B8 |
| Gonium pect | A0A150FU74 |
| Gopherus aga | A0A452H3F3 |
| Gopherus aga | A0A452H966 |
| Gopherus aga | A0A452H952 |
| Gorilla gorilla | G3RXF7 |
| Gorilla gorilla | G3R4W6 |
| Gorilla gorilla | G3RNU9 |
| Gorilla gorilla | G3R2X1 |
| Gorilla gorilla | A0A2I2ZIP3 |
| Gorilla gorilla | A0A2I2YSL6 |
| Gorilla gorilla | G3RZ73 |
| Gorilla gorilla | A0A2I2ZYB7 |
| Gorilla gorilla | G3QLL7 |
| Gossypium ar | A0A0B0N1R7 |
| Gossypium ar | A0A0B0PPF2 |
| Gossypium ar | A0A0B0NU75 |
| Gossypium ar | A0A0B0MYX5 |
| Gossypium ar | A0A0B0MTX0 |
| Gossypium ar | A0A0B0MXU6 |
| Gossypium ar | A0A0B0MZL4 |
| Gossypium ar | A0A0B0NP12 |
| Gossypium ar | A0A0B0N0U6 |
| Gossypium ar | A0A0B0N855 |
| Gossypium ar | A0A0B0PYF7 |
| Gossypium ar | A0A0B0N5G9 |
| Gossypium ba | A0A2P5QME1 |
| Gossypium ba | A0A2P5WSV9 |
| Gossypium ba | A0A2P5R530 |
| Gossypium ba | A0A2P5QAQ1 |
| Gossypium ba | A0A2P5RJB5 |
| Gossypium ba | A0A2P5PX52 |
| Gossypium ba | A0A2P5R799 |
| Gossypium ba | A0A2P5XQ00 |
| Gossypium ba | A0A2P5R525 |
| Gossypium ba | A0A2P5YJR7 |
| Gossypium ba | A0A2P5WUP4 |
| Gossypium ba | A0A2P5XQE1 |
| Gossypium ba | A0A2P5W537 |

| | |
|---|---|
| Gossypium b: | A0A2P5QYR4 |
| Gossypium b: | A0A2P5RJD9 |
| Gossypium b: | A0A2P5SAL6 |
| Gossypium b: | A0A2P5R4Z5 |
| Gossypium b: | A0A2P5RMX0 |
| Gossypium b: | A0A2P5Y4V6 |
| Gossypium b: | A0A2P5XRT3 |
| Gossypium b: | A0A2P5RJC7 |
| Gossypium b: | A0A2P5XLL9 |
| Gossypium b: | A0A2P5QP15 |
| Gossypium b: | A0A2P5YQK4 |
| Gossypium b: | A0A2P5Q2I3 |
| Gossypium b: | A0A2P5W0S5 |
| Gossypium b: | A0A2P5RX37 |
| Gossypium b: | A0A2P5RFB1 |
| Gossypium b: | A0A2P5XRP7 |
| Gossypium b: | A0A2P5YMC7 |
| Gossypium b: | A0A2P5XPY8 |
| Gossypium b: | A0A2P5W117 |
| Gossypium b: | A0A2P5Q5R1 |
| Gossypium b: | A0A2P5QZB7 |
| Gossypium b: | A0A2P5Y5I5 |
| Gossypium b: | A0A2P5SRC9 |
| Gossypium b: | A0A2P5RY44 |
| Gossypium b: | A0A2P5WP31 |
| Gossypium b: | A0A2P5RWS6 |
| Gossypium b: | A0A2P5RMT2 |
| Gossypium b: | A0A2P5Y6U2 |
| Gossypium b: | A0A2P5R4X6 |
| Gossypium b: | A0A2P5YX52 |
| Gossypium b: | A0A2P5XNK6 |
| Gossypium b: | A0A2P5WUK6 |
| Gossypium b: | A0A2P5XT89 |
| Gossypium b: | A0A2P5SBC1 |
| Gossypium b: | A0A2P5SP55 |
| Gossypium b: | A0A2P5W510 |
| Gossypium b: | A0A2P5RJE1 |
| Gossypium b: | A0A2P5QIM8 |
| Gossypium b: | A0A2P5SJY1 |
| Gossypium b: | A0A2P5QJX4 |
| Gossypium b: | A0A2P5S011 |

| | |
|---|---|
| Gossypium b: | A0A2P5QSI7 |
| Gossypium b: | A0A2P5WKV8 |
| Gossypium b: | A0A2P5RN97 |
| Gossypium b: | A0A2P5RZU3 |
| Gossypium b: | A0A2P5XM03 |
| Gossypium b: | A0A2P5S017 |
| Gossypium b: | A0A2P5WUK4 |
| Gossypium b: | A0A2P5QAP8 |
| Gossypium b: | A0A2P5W0S1 |
| Gossypium b: | A0A2P5RMG9 |
| Gossypium b: | A0A2P5VYB6 |
| Gossypium b: | A0A2P5Y509 |
| Gossypium b: | A0A2P5W461 |
| Gossypium b: | A0A2P5Y5H9 |
| Gossypium b: | A0A2P5X2H7 |
| Gossypium b: | A0A2P5XQF5 |
| Gossypium b: | A0A2P5QC40 |
| Gossypium b: | A0A2P5WKV0 |
| Gossypium b: | A0A2P5R7M3 |
| Gossypium b: | A0A2P5WII7 |
| Gossypium b: | A0A2P5XRR5 |
| Gossypium b: | A0A2P5S3Y3 |
| Gossypium b: | A0A2P5PX26 |
| Gossypium b: | A0A2P5XT09 |
| Gossypium b: | A0A2P5Y4V2 |
| Gossypium b: | A0A2P5XYY9 |
| Gossypium b: | A0A2P5YJS7 |
| Gossypium b: | A0A2P5RWY7 |
| Gossypium b: | A0A2P5SP77 |
| Gossypium b: | A0A2P5RWV6 |
| Gossypium b: | A0A2P5R4Z6 |
| Gossypium b: | A0A2P5R7C6 |
| Gossypium b: | A0A2P5WUM4 |
| Gossypium b: | A0A2P5R7B5 |
| Gossypium b: | A0A2P5YX45 |
| Gossypium b: | A0A2P5R500 |
| Gossypium b: | A0A2P5W2X4 |
| Gossypium b: | A0A2P5QPE5 |
| Gossypium b: | A0A2P5SP93 |
| Gossypium b: | A0A2P5RMJ4 |
| Gossypium b: | A0A2P5Q5Q4 |

| | |
|---|---|
| Gossypium b | A0A2P5Y5L5 |
| Gossypium b | A0A2P5R4Y1 |
| Gossypium b | A0A2P5Y9P7 |
| Gossypium b | A0A2P5RZT0 |
| Gossypium b | A0A2P5XRP9 |
| Gossypium b | A0A2P5RUA4 |
| Gossypium b | A0A2P5Y109 |
| Gossypium b | A0A2P5R7B6 |
| Gossypium b | A0A2P5QAB2 |
| Gossypium b | A0A2P5QJY8 |
| Gossypium b | A0A2P5SV42 |
| Gossypium b | A0A2P5WSV7 |
| Gossypium b | A0A2P5RKW8 |
| Gossypium b | A0A2P5WKX2 |
| Gossypium b | A0A2P5QJM8 |
| Gossypium b | A0A2P5SE83 |
| Gossypium b | A0A2P5SP86 |
| Gossypium b | A0A2P5R7B3 |
| Gossypium b | A0A2P5Q4C5 |
| Gossypium b | A0A2P5SE70 |
| Gossypium b | A0A2P5RMS7 |
| Gossypium b | A0A2P5YQL9 |
| Gossypium b | A0A2P5WKU3 |
| Gossypium b | A0A2P5R5J1 |
| Gossypium b | A0A2P5RN93 |
| Gossypium b | A0A2P5SP50 |
| Gossypium b | A0A2P5XRU3 |
| Gossypium b | A0A2P5VNQ9 |
| Gossypium b | A0A2P5PXQ4 |
| Gossypium b | A0A2P5R7A8 |
| Gossypium b | A0A2P5RMU3 |
| Gossypium b | A0A2P5RU57 |
| Gossypium b | A0A2P5RLV9 |
| Gossypium b | A0A2P5XRW6 |
| Gossypium b | A0A2P5XRW3 |
| Gossypium b | A0A2P5R7B4 |
| Gossypium b | A0A2P5WKW0 |
| Gossypium b | A0A2P5VXE2 |
| Gossypium b | A0A2P5VQP7 |
| Gossypium b | A0A2P5QAS2 |
| Gossypium hi | A0A1U8LTN4 |

| | |
|---|---|
| Gossypium hi | A0A1U8IY86 |
| Gossypium hi | A0A1U8JWT7 |
| Gossypium hi | A0A1U8MSL2 |
| Gossypium hi | A0A1U8NIT3 |
| Gossypium hi | A0A1U8JW83 |
| Gossypium hi | A0A1U8M088 |
| Gossypium hi | A0A1U8PII4 |
| Gossypium hi | A0A1U8MW63 |
| Gossypium hi | A0A1U8PIK8 |
| Gossypium hi | A0A1U8PIH7 |
| Gossypium hi | A0A1U8NKW8 |
| Gossypium hi | A0A1U8KUL9 |
| Gossypium hi | A0A1U8MSI7 |
| Gossypium hi | A0A1U8MQ96 |
| Gossypium hi | A0A1U8MJ01 |
| Gossypium hi | A0A1U8INS0 |
| Gossypium hi | A0A1U8J3Z6 |
| Gossypium hi | A0A1U8MZ75 |
| Gossypium hi | A0A1U8J8V8 |
| Gossypium hi | A0A1U8K7I4 |
| Gossypium hi | A0A1U8HRC6 |
| Gossypium hi | A0A1U8JD70 |
| Gossypium hi | A0A1U8K7G9 |
| Gossypium hi | A0A1U8MW53 |
| Gossypium hi | A0A1U8M3G9 |
| Gossypium hi | A0A1U8PS02 |
| Gossypium hi | A0A1U8MWK7 |
| Gossypium hi | A0A1U8IQI9 |
| Gossypium hi | A0A1U8IQ79 |
| Gossypium hi | A0A1U8J5M7 |
| Gossypium hi | A0A1U8PL10 |
| Gossypium hi | A0A1U8PJ03 |
| Gossypium hi | A0A1U8KPI2 |
| Gossypium hi | A0A1U8KDH7 |
| Gossypium hi | A0A1U8KPN3 |
| Gossypium hi | A0A1U8NGI9 |
| Gossypium hi | A0A1U8MQB6 |
| Gossypium hi | A0A1U8N4R6 |
| Gossypium hi | A0A1U8KIK7 |
| Gossypium hi | A0A1U8NU95 |
| Gossypium hi | A0A1U8K568 |

| | |
|---|---|
| Gossypium hi | A0A1U8JUG2 |
| Gossypium hi | A0A1U8KP21 |
| Gossypium hi | A0A1U8HJH7 |
| Gossypium hi | A0A1U8MU48 |
| Gossypium hi | A0A1U8J417 |
| Gossypium hi | A0A1U8MTB7 |
| Gossypium hi | A0A1U8IZ76 |
| Gossypium hi | A0A1U8MSL8 |
| Gossypium hi | A0A1U8KFJ8 |
| Gossypium hi | A0A1U8IIC3 |
| Gossypium hi | A0A1U8PRU7 |
| Gossypium hi | I0B675 |
| Gossypium hi | A0A1U8NVM4 |
| Gossypium hi | A0A1U8N3Z5 |
| Gossypium hi | A0A1U8J436 |
| Gossypium hi | A0A1U8MWV1 |
| Gossypium hi | A0A1U8L9V7 |
| Gossypium hi | A0A1U8MU65 |
| Gossypium hi | A0A1U8K7H3 |
| Gossypium hi | A0A1U8J5E0 |
| Gossypium hi | A0A1U8NTW1 |
| Gossypium hi | A0A1U8J456 |
| Gossypium hi | A0A1U8LQA8 |
| Gossypium hi | A0A1U8LU33 |
| Gossypium hi | A0A1U8MJ39 |
| Gossypium hi | A0A1U8KJM4 |
| Gossypium hi | A0A1U8N710 |
| Gossypium hi | A0A1U8IIU3 |
| Gossypium hi | A0A1U8HZZ8 |
| Gossypium hi | A0A1U8JME3 |
| Gossypium hi | A0A1U8MSV8 |
| Gossypium hi | A0A1U8LQ71 |
| Gossypium hi | A0A1U8KI27 |
| Gossypium hi | A0A1U8N0I2 |
| Gossypium hi | A0A1U8NVW2 |
| Gossypium hi | A0A1U8JTJ3 |
| Gossypium hi | A0A1U8MU44 |
| Gossypium hi | A0A1U8HXT9 |
| Gossypium hi | A0A1U8KN26 |
| Gossypium hi | A0A1U8NAH6 |
| Gossypium hi | A0A1U8MPV7 |

| | |
|---|---|
| Gossypium hi | A0A1U8NWF2 |
| Gossypium hi | A0A1U8JA35 |
| Gossypium hi | A0A1U8MWB9 |
| Gossypium hi | A0A1U8NWR2 |
| Gossypium hi | A0A1U8MU69 |
| Gossypium hi | A0A1U8N0D3 |
| Gossypium hi | A0A1U8MXQ1 |
| Gossypium hi | A0A1U8KM96 |
| Gossypium hi | A0A1U8NTW9 |
| Gossypium hi | A0A1U8KM40 |
| Gossypium hi | A0A1U8KUG4 |
| Gossypium hi | A0A1U8MSS6 |
| Gossypium hi | A0A1U8HR08 |
| Gossypium hi | A0A2P0JA21 |
| Gossypium hi | A0A1U8N932 |
| Gossypium hi | A0A1U8LW85 |
| Gossypium hi | A0A1U8NVG7 |
| Gossypium hi | A0A1U8MT98 |
| Gossypium hi | A0A1U8L9U6 |
| Gossypium hi | A0A1U8NWQ7 |
| Gossypium hi | A0A1U8JLD2 |
| Gossypium hi | A0A1U8NWF7 |
| Gossypium hi | A0A1U8JPV5 |
| Gossypium hi | A0A1U8PJ85 |
| Gossypium hi | A0A1U8IIP2 |
| Gossypium hi | A0A1U8NU86 |
| Gossypium hi | A0A1U8IJ22 |
| Gossypium ra | A0A0D2NHR6 |
| Gossypium ra | A0A0D2S9W7 |
| Gossypium ra | A0A0D2SHQ6 |
| Gossypium ra | A0A0D2R293 |
| Gossypium ra | A0A0D2UMW3 |
| Gossypium ra | A0A0D2U925 |
| Gossypium ra | A0A0D2UF09 |
| Gossypium ra | A0A0D2R2A3 |
| Gossypium ra | A0A0D2N3S2 |
| Gossypium ra | A0A0D2Q7B3 |
| Gossypium ra | A0A0D2NJM3 |
| Gossypium ra | A0A0D2Q7C6 |
| Gossypium ra | A0A0D2U4N2 |
| Gossypium ra | A0A0D2U7M3 |

| | |
|---|---|
| Gossypium ra | A0A0D2VH44 |
| Gossypium ra | A0A0D2R1G8 |
| Gossypium ra | A0A0D2U3X9 |
| Gossypium ra | A0A0D2NYP5 |
| Gossypium ra | A0A0D2NJL3 |
| Gossypium ra | A0A0D2S4R5 |
| Gossypium ra | A0A0D2RIS5 |
| Gossypium ra | A0A0D2SLX3 |
| Gossypium ra | A0A0D2W2H3 |
| Gossypium ra | A0A0D2Q066 |
| Gossypium ra | A0A0D2R018 |
| Gossypium ra | A0A0D2VAH9 |
| Gossypium ra | A0A0D2NJN0 |
| Gossypium ra | A0A0D2VJL8 |
| Gossypium ra | A0A0D2T102 |
| Gossypium ra | A0A0D2Q7A2 |
| Gossypium ra | A0A0D2U3Z0 |
| Gossypium ra | A0A0D2VAL8 |
| Gossypium ra | A0A0D2V8Y9 |
| Gossypium ra | A0A0D2Q2C4 |
| Gossypium ra | A0A0D2TKF7 |
| Gossypium ra | A0A0D2NHP9 |
| Gossypium ra | A0A0D2S2B9 |
| Gossypium ra | A0A0D2TEH9 |
| Gossypium ra | A0A0D2ST81 |
| Gossypium ra | A0A0D2PWM1 |
| Gossypium ra | A0A0D2SZD9 |
| Gossypium ra | A0A0D2M695 |
| Gossypium ra | A0A0D2RIT8 |
| Gossypium ra | A0A0D2Q9G7 |
| Gossypium ra | A0A0D2QW76 |
| Gossypium ra | A0A0D2NG46 |
| Gossypium ra | A0A0D2RFL5 |
| Gossypium ra | A0A0D2U430 |
| Gossypium ra | A0A0D2Q063 |
| Gossypium ra | A0A0D2R2E2 |
| Gossypium ra | A0A0D2QHX8 |
| Gossypium ra | A0A0D2V1I8 |
| Gossypium ra | A0A0D2PWL4 |
| Gossypium ra | A0A0D2NHQ3 |
| Gossypium ra | A0A0D2R874 |

| | |
|---|---|
| Gossypium ra | A0A0D2SKU2 |
| Gossypium ra | A0A0D2NJR3 |
| Gossypium ra | A0A0D2NHR3 |
| Gossypium ra | A0A0D2S5T3 |
| Gossypium ra | A0A0D2ST84 |
| Gossypium ra | A0A0D2VA22 |
| Gossypium ra | A0A0D2T0Z5 |
| Gossypium ra | A0A0D2R008 |
| Gossypium ra | A0A0D2Q078 |
| Gossypium ra | A0A0D2NHS1 |
| Gossypium ra | A0A0D2TRL9 |
| Gossypium ra | A0A0D2R2W6 |
| Gossypium ra | A0A0D2TF92 |
| Gossypium ra | A0A0D2RIT4 |
| Gossypium ra | A0A0D2M871 |
| Gossypium ra | A0A0D2T212 |
| Gossypium ra | A0A0D2NTC5 |
| Gossypium ra | A0A0D2R013 |
| Gossypium ra | A0A0D2R877 |
| Gossypium ra | A0A0D2TUH1 |
| Gossypium ra | A0A0D2TTI3 |
| Gossypium ra | A0A0D2Q2G3 |
| Gossypium ra | A0A0D2RSA1 |
| Gossypium ra | A0A0D2TQ92 |
| Gossypium ra | A0A0D2VUT2 |
| Gossypium ra | A0A0D2M7Q0 |
| Gossypium ra | A0A0D2TWT0 |
| Gossypium ra | A0A0D2RAR6 |
| Gossypium ra | A0A0D2Q9E3 |
| Gossypium ra | A0A0D2U0Y6 |
| Gossypium ra | A0A0D2M841 |
| Gossypium ra | A0A0D2UGZ9 |
| Gossypium ra | A0A0D2Q059 |
| Gossypium ra | A0A0D2PQT1 |
| Gossypium ra | A0A0D2SUB8 |
| Gossypium ra | A0A0D2UNZ7 |
| Gossypium ra | A0A0D2V1J1 |
| Gossypium ra | A0A0D2Q081 |
| Gossypium ra | A0A0D2R033 |
| Gossypium ra | A0A0D2RQB0 |
| Gossypium ra | A0A0D2M672 |

| | |
|---|---|
| Gossypium ra | A0A0D2UF05 |
| Gossypium ra | A0A0D2UUZ6 |
| Gossypium ra | A0A0D2W712 |
| Gossypium ra | A0A0D2UPR4 |
| Gossypium ra | A0A0D2UF15 |
| Gossypium ra | A0A0D2SYV9 |
| Gossypium ra | A0A0D2V121 |
| Gossypium ra | A0A0D2Q2D6 |
| Gossypium ra | A0A0D2NIU8 |
| Gossypium ra | A0A0D2UCR1 |
| Gossypium ra | A0A0D2QA06 |
| Gossypium ra | A0A0D2Q2G4 |
| Gossypium ra | A0A0D2RIU4 |
| Graphocepha | A0A1B6MBH5 |
| Graphocepha | A0A1B6MT83 |
| Graphocepha | A0A1B6KHB7 |
| Graphocepha | A0A1B6LPX0 |
| Graphocepha | A0A1B6MRL1 |
| Graphocepha | A0A1B6KWB9 |
| Graphocepha | A0A1B6KTZ0 |
| Gregarina nip | A0A023AXR7 |
| Gregarina nip | A0A023AYU7 |
| Gregarina nip | A0A023B052 |
| Gregarina nip | A0A023BAV3 |
| Gregarina nip | A0A023B0X7 |
| Gregarina nip | A0A023B381 |
| Gregarina nip | A0A023B0X6 |
| Gregarina nip | A0A023AXP8 |
| Gregarina nip | A0A023B1C3 |
| Gregarina nip | A0A023B0P6 |
| Guillardia the | L1JJD2 |
| Guillardia the | L1JJX0 |
| Guillardia the | L1JTS3 |
| Guillardia the | L1JKL7 |
| Guillardia the | L1JCN2 |
| Guillardia the | L1IX89 |
| Guillardia the | L1K113 |
| Guillardia the | L1IWJ0 |
| Guillardia the | L1IIZ2 |
| Guillardia the | L1JCP1 |
| Gulo gulo.xlsx | A0A3P4KMI5 |

| | |
|---|---|
| Gulo gulo.xls: | A0A3P4RU50 |
| Gulo gulo.xls: | A0A3P4RW44 |
| Gulo gulo.xls: | A0A3P4RAU2 |
| Gulo gulo.xls: | A0A3P4RWP0 |
| Habropoda la | A0A0L7QLL2 |
| Habropoda la | A0A0L7R1I8 |
| Habropoda la | A0A0L7RF61 |
| Habropoda la | A0A0L7R6V2 |
| Habropoda la | A0A0L7R538 |
| Habropoda la | A0A0L7RFI2 |
| Haematobact | A0A212ACR4 |
| Haematobact | A0A086XX93 |
| Haematobact | A0A212AJW7 |
| Haemonchus | W6NGT9 |
| Haemonchus | W6NNJ2 |
| Haemonchus | A0A158QM82 |
| Haemonchus | A0A0N4W5U3 |
| Haemonchus | A0A158QK55 |
| Haemonchus | A0A3P7UET5 |
| Haemonchus | A0A3P7TV91 |
| Haemonchus | A0A3P7XBC6 |
| Haemonchus | A0A3P7TQB2 |
| Haemonchus | A0A0N4WUQ0 |
| Haemonchus | A0A0N4W8T5 |
| Haemonchus | A0A158QNJ5 |
| Haemonchus | A0A0N4WA48 |
| Haemonchus | A0A0N4X5R8 |
| Haemonchus | A0A0N4WPF3 |
| Haemonchus | A0A0N4X266 |
| Haemonchus | A0A0N4WAZ2 |
| Haemonchus | A0A158QLJ4 |
| Haemonchus | A0A158QPL6 |
| Haemonchus | A0A158QR36 |
| Haemonchus | A0A158QKJ2 |
| Haemonchus | A0A0N4WU40 |
| Haemonchus | A0A0N4WJV0 |
| Haemonchus | A0A0N4WKR9 |
| Haemonchus | A0A0N4WC06 |
| Haemonchus | A0A158QRG0 |
| Haemonchus | A0A0N4W2R6 |
| Haemonchus | A0A0N4WAT5 |

| | |
|---|---|
| Haemonchus | A0A0N4WNL0 |
| Haemonchus | A0A0N4WCR5 |
| Haemonchus | A0A0N4WS57 |
| Haemonchus | A0A0N4WAG3 |
| Haliaeetus all | A0A091PRS1 |
| Haliaeetus all | A0A091PH99 |
| Haliaeetus all | A0A091PGF6 |
| Haliaeetus all | A0A091PJK8 |
| Halocynthiiba | A0A126UY30 |
| Halomonas c | A0A1M7I344 |
| Halomonas h | W1NCY1 |
| Hammondia l | A0A074SYS7 |
| Hammondia l | A0A074SUJ0 |
| Hammondia l | A0A074TKE0 |
| Hammondia l | A0A074SSB4 |
| Hammondia l | A0A074T9T4 |
| Hammondia l | A0A074SY06 |
| Hammondia l | A0A074SSP2 |
| Hammondia l | A0A074SRW4 |
| Hammondia l | A0A074T002 |
| Hammondia l | A0A074SV30 |
| Hammondia l | A0A074SUT3 |
| Hammondia l | A0A074SME0 |
| Hammondia l | A0A074T415 |
| Hammondia l | A0A074T0Q6 |
| Hammondia l | A0A074TPN2 |
| Hammondia l | A0A074SMC6 |
| Handroanthu | A0A2G9HBA5 |
| Handroanthu | A0A2G9I607 |
| Handroanthu | A0A2G9GS19 |
| Handroanthu | A0A2G9HFW2 |
| Handroanthu | A0A2G9G3H0 |
| Handroanthu | A0A2G9GBC9 |
| Handroanthu | A0A2G9H9N3 |
| Handroanthu | A0A2G9FWW5 |
| Handroanthu | A0A2G9G020 |
| Handroanthu | A0A2G9H900 |
| Handroanthu | A0A2G9G8U4 |
| Handroanthu | A0A2G9H5E2 |
| Handroanthu | A0A2G9G0T8 |
| Handroanthu | A0A2G9G7H6 |

| | |
|---|---|
| Handroanthu | A0A2G9G7F0 |
| Handroanthu | A0A2G9HQH7 |
| Handroanthu | A0A2G9GLL5 |
| Handroanthu | A0A2G9I602 |
| Handroanthu | A0A2G9HMW9 |
| Handroanthu | A0A2G9H7Y4 |
| Handroanthu | A0A2G9HVS1 |
| Handroanthu | A0A2G9H5H4 |
| Handroanthu | A0A2G9FW99 |
| Handroanthu | A0A2G9G9M9 |
| Handroanthu | A0A2G9FX93 |
| Handroanthu | A0A2G9G0T2 |
| Handroanthu | A0A2G9GFB5 |
| Handroanthu | A0A2G9HQJ0 |
| Handroanthu | A0A2G9I4A0 |
| Handroanthu | A0A2G9G0G1 |
| Handroanthu | A0A2G9HNZ9 |
| Handroanthu | A0A2G9GUR0 |
| Handroanthu | A0A2G9H3N1 |
| Handroanthu | A0A2G9FY38 |
| Handroanthu | A0A2G9G8A6 |
| Handroanthu | A0A2G9GBD2 |
| Handroanthu | A0A2G9HQ39 |
| Handroanthu | A0A2G9HP33 |
| Handroanthu | A0A2G9G948 |
| Handroanthu | A0A2G9G8Y1 |
| Handroanthu | A0A2G9GK44 |
| Handroanthu | A0A2G9G7D7 |
| Handroanthu | A0A2G9IBJ2 |
| Handroanthu | A0A2G9G5U6 |
| Handroanthu | A0A2G9HBA6 |
| Handroanthu | A0A2G9G7K3 |
| Handroanthu | A0A2G9G7Z8 |
| Handroanthu | A0A2G9G482 |
| Handroanthu | A0A2G9GNC7 |
| Handroanthu | A0A2G9HBD0 |
| Handroanthu | A0A2G9GA17 |
| Handroanthu | A0A2G9GNR7 |
| Handroanthu | A0A2G9GGQ5 |
| Handroanthu | A0A2G9I619 |
| Handroanthu | A0A2G9G874 |

| | |
|---|---|
| Handroanthu | A0A2G9GY40 |
| Handroanthu | A0A2G9HZV9 |
| Handroanthu | A0A2G9G7A6 |
| Handroanthu | A0A2G9GZU4 |
| Handroanthu | A0A2G9GC79 |
| Handroanthu | A0A2G9G877 |
| Handroanthu | A0A2G9G5N8 |
| Handroanthu | A0A2G9I606 |
| Handroanthu | A0A2G9HM51 |
| Handroanthu | A0A2G9GUP3 |
| Handroanthu | A0A2G9GEQ2 |
| Handroanthu | A0A2G9HNL0 |
| Handroanthu | A0A2G9H7Q3 |
| Handroanthu | A0A2G9G4X6 |
| Handroanthu | A0A2G9IBI1 |
| Handroanthu | A0A2G9GJX1 |
| Handroanthu | A0A2G9I6C9 |
| Handroanthu | A0A2G9G5B6 |
| Handroanthu | A0A2G9GB96 |
| Handroanthu | A0A2G9HHD9 |
| Handroanthu | A0A2G9GFY4 |
| Handroanthu | A0A2G9H5D1 |
| Handroanthu | A0A2G9GFA7 |
| Handroanthu | A0A2G9GBC7 |
| Handroanthu | A0A2G9H719 |
| Handroanthu | A0A2G9H716 |
| Handroanthu | A0A2G9IAY0 |
| Handroanthu | A0A2G9GX58 |
| Handroanthu | A0A2G9GA47 |
| Handroanthu | A0A2G9GKN2 |
| Handroanthu | A0A2G9FXC4 |
| Handroanthu | A0A2G9FY42 |
| Handroanthu | A0A2G9FXI3 |
| Handroanthu | A0A2G9H718 |
| Handroanthu | A0A2G9HP44 |
| Handroanthu | A0A2G9GMC6 |
| Handroanthu | A0A2G9HMZ3 |
| Handroanthu | A0A2G9I5Y5 |
| Handroanthu | A0A2G9G1J5 |
| Handroanthu | A0A2G9H762 |
| Handroanthu | A0A2G9GKG3 |

| | |
|---|---|
| Handroanthu | A0A2G9IBT8 |
| Handroanthu | A0A2G9GT09 |
| Handroanthu | A0A2G9G6Z8 |
| Handroanthu | A0A2G9I585 |
| Handroanthu | A0A2G9HHH7 |
| Handroanthu | A0A2G9HTZ3 |
| Handroanthu | A0A2G9HHL8 |
| Handroanthu | A0A2G9H5I0 |
| Handroanthu | A0A2G9I2J4 |
| Handroanthu | A0A2G9HP00 |
| Handroanthu | A0A2G9GSG6 |
| Handroanthu | A0A2G9FYE8 |
| Handroanthu | A0A2G9H932 |
| Handroanthu | A0A2G9G8L4 |
| Haplochromis | A0A3Q2WKE2 |
| Haplochromis | A0A3Q2VWA3 |
| Haplochromis | A0A3Q2VMX2 |
| Haplochromis | A0A3Q2W714 |
| Haplochromis | A0A3Q2V9X5 |
| Haplochromis | A0A3Q3BK39 |
| Haplochromis | A0A3Q3CJW9 |
| Harpegnatho | E2BZU8 |
| Harpegnatho | E2BV16 |
| Harpegnatho | E2C0I8 |
| Harpegnatho | E2C2K1 |
| Harpegnatho | E2B4Q6 |
| Harpegnatho | E2B676 |
| Hartmanniba | A0A2C9D802 |
| Helianthus ar | A0A251VNS3 |
| Helianthus ar | A0A251T9D7 |
| Helianthus ar | A0A251S5U8 |
| Helianthus ar | A0A251RZC7 |
| Helianthus ar | A0A251URM3 |
| Helianthus ar | A0A251U4N7 |
| Helianthus ar | A0A251SPB8 |
| Helianthus ar | A0A251U419 |
| Helianthus ar | A0A251TB44 |
| Helianthus ar | A0A251VQC9 |
| Helianthus ar | A0A251TTA7 |
| Helianthus ar | A0A251T9Z3 |
| Helianthus ar | A0A251T8T6 |

Helianthus ar A0A251S5S5
Helianthus ar A0A251USV7
Helianthus ar A0A251V9P5
Helianthus ar A0A251SNM0
Helianthus ar A0A251RSS9
Helianthus ar A0A251ST77
Helianthus ar A0A251U4U7
Helianthus ar A0A251URK0
Helianthus ar A0A251V8P7
Helianthus ar A0A251V4L5
Helianthus ar A0A251T8Z6
Helianthus ar A0A251V3F7
Helianthus ar A0A251SGZ7
Helianthus ar A0A251UQR4
Helianthus ar A0A251S8P2
Helianthus ar A0A251SC29
Helianthus ar A0A251ULP0
Helianthus ar A0A251T9K9
Helianthus ar A0A251T9M7
Helianthus ar A0A251TAJ6
Helianthus ar A0A251T967
Helianthus ar A0A251U5M7
Helianthus ar A0A251UIR2
Helianthus ar A0A251RT61
Helianthus ar A0A251TNE9
Helianthus ar A0A251T9J2
Helianthus ar A0A251THT3
Helianthus ar A0A251TZ72
Helianthus ar A0A251U550
Helianthus ar A0A251U473
Helianthus ar A0A251TNN4
Helianthus ar A0A251UQL0
Helianthus ar A0A251RSW7
Helianthus ar A0A251U760
Helianthus ar A0A251VDL9
Helianthus ar A0A251UDJ8
Helianthus ar A0A251U3Z2
Helianthus ar A0A251T939
Helianthus ar A0A251V5W8
Helianthus ar A0A251TDT0
Helianthus ar A0A251SDU7

Helianthus ar A0A251T4J5
Helianthus ar A0A251SMJ5
Helianthus ar A0A251TA85
Helianthus ar A0A251S960
Helianthus ar A0A251T8Z8
Helianthus ar A0A251UJM0
Helianthus ar A0A251T4T1
Helianthus ar A0A251VPZ1
Helianthus ar A0A251TCA5
Helianthus ar A0A251TAB3
Helianthus ar A0A251SW79
Helianthus ar A0A251RXK0
Helianthus ar A0A251S8G7
Helianthus ar A0A251TSR7
Helianthus ar A0A251U418
Helianthus ar A0A251T909
Helianthus ar A0A251TAA8
Helianthus ar A0A1Y3BZF9
Helianthus ar A0A251T8Y8
Helianthus ar A0A251RSP8
Helianthus ar A0A251SSD8
Helianthus ar A0A251T949
Helianthus ar A0A251RMY8
Helianthus ar A0A251TA21
Helianthus ar A0A251T2S5
Helianthus ar A0A251RN23
Helianthus ar A0A251T906
Helianthus ar A0A251UL16
Helianthus ar A0A251RV73
Helianthus ar A0A251RTF5
Helianthus ar A0A251S990
Helianthus ar A0A251STX9
Helianthus ar A0A251S8Y4
Helianthus ar A0A251URH5
Helianthus ar A0A251S0K1
Helianthus ar A0A251UEQ1
Helianthus ar A0A251RNY5
Helianthus ar A0A251TL03
Helianthus ar A0A251SS56
Helianthus ar A0A251TN10
Helianthus ar A0A251THY3

Helianthus ar A0A251S995
Helianthus ar A0A251TA25
Helianthus ar A0A251T5L3
Helianthus ar A0A251V227
Helianthus ar A0A251T959
Helianthus ar A0A251TL38
Helianthus ar A0A251TAD2
Helianthus ar A0A251SXL9
Helianthus ar A0A251U5K9
Helianthus ar A0A251T266
Helianthus ar A0A251U403
Helianthus ar A0A251V611
Helianthus ar A0A251THH7
Helianthus ar A0A251T9X4
Helianthus ar A0A251TV70
Helianthus ar A0A251TAX5
Helianthus ar A0A251RS35
Helianthus ar A0A251RS51
Helianthus ar A0A251T7B5
Helianthus ar A0A251T1U1
Helianthus ar A0A251RNF0
Helianthus ar A0A251VBF9
Helianthus ar A0A251UHX2
Helianthus ar A0A251U6B1
Helianthus ar A0A251ULD7
Helianthus ar A0A251T3N2
Helianthus ar A0A251TA13
Helianthus ar A0A251T908
Helianthus ar A0A251U4V0
Helianthus ar A0A251UAK0
Helianthus ar A0A251TAL5
Helianthus ar A0A251VCQ1
Helianthus ar A0A251SMN4
Helianthus ar A0A251U509
Helianthus ar A0A251U5G8
Helianthus ar A0A251SMR2
Helianthus ar A0A251UHY6
Helianthus ar A0A251TY69
Helianthus ar A0A251TCD7
Helianthus ar A0A251U4T1
Helianthus ar A0A251TJD6

| | |
|---|---|
| Helianthus ar | A0A251U476 |
| Helianthus ar | A0A251UPA1 |
| Helianthus ar | A0A251T447 |
| Helianthus ar | A0A251T2R7 |
| Helianthus ar | A0A251U4S3 |
| Helianthus ar | A0A251VPX0 |
| Helianthus ar | A0A251T8U9 |
| Helianthus ar | A0A251T1W6 |
| Helianthus ar | A0A251T929 |
| Helianthus ar | A0A251T9Q8 |
| Helianthus ar | A0A251T2N3 |
| Helianthus ar | A0A251ULK0 |
| Helianthus ar | A0A251TA08 |
| Helianthus ar | A0A251T9E6 |
| Helianthus ar | A0A251U5H8 |
| Helianthus ar | A0A251T585 |
| Helianthus ar | A0A251UNM9 |
| Helianthus ar | A0A251SVM1 |
| Helianthus ar | A0A251S0B8 |
| Helianthus ar | A0A251TAB8 |
| Helianthus ar | A0A251U4T7 |
| Helianthus ar | A0A251VUB1 |
| Helianthus ar | A0A251U519 |
| Helianthus ar | A0A251UJH6 |
| Helianthus ar | A0A251VLZ7 |
| Helianthus ar | A0A251SM06 |
| Helianthus ar | A0A251T1Y3 |
| Helianthus ar | A0A251T2F8 |
| Helianthus ar | A0A251SA55 |
| Helianthus ar | A0A251TAM7 |
| Helianthus ar | A0A251RTF9 |
| Helianthus ar | A0A251RTR7 |
| Helianthus ar | A0A251U5H1 |
| Helianthus ar | A0A251UTN4 |
| Helianthus ar | A0A251UTW0 |
| Helianthus ar | A0A251SU98 |
| Helianthus ar | A0A251UQN1 |
| Helianthus ar | A0A251SNN8 |
| Helianthus ar | A0A251U3Z3 |
| Helianthus ar | A0A251USS8 |
| Helianthus ar | A0A251TMP9 |

| | |
|---|---|
| Helianthus ar | A0A251UTL3 |
| Helianthus ar | A0A251SS58 |
| Helianthus ar | A0A251S4B6 |
| Helianthus ar | A0A251SKT0 |
| Helianthus ar | A0A251T8U8 |
| Helianthus ar | A0A251U463 |
| Helianthus ar | A0A251U4V7 |
| Helianthus ar | A0A251U422 |
| Helianthus ar | A0A251TAS0 |
| Helianthus ar | A0A251RMW7 |
| Helianthus ar | A0A251TAE1 |
| Helianthus ar | A0A251U8J7 |
| Helianthus ar | A0A251VPQ8 |
| Helianthus ar | A0A251U8C6 |
| Helianthus ar | A0A251SMR8 |
| Helianthus ar | A0A251TA09 |
| Helianthus ar | A0A251SS64 |
| Helianthus ar | A0A251VRM9 |
| Helianthus ar | A0A251U6A1 |
| Helianthus ar | A0A251VHM6 |
| Helianthus ar | A0A251U4T8 |
| Helianthus ar | A0A251T3G0 |
| Helicocarpus | A0A2B7WMP5 |
| Helicocarpus | A0A2B7XNZ2 |
| Helicosporidi | A0A059LJS3 |
| Helicosporidi | A0A059LQC7 |
| Heligmosomo | A0A183GCJ4 |
| Heligmosomo | A0A183FDS9 |
| Heligmosomo | A0A183GQL5 |
| Heligmosomo | A0A183FNG0 |
| Heligmosomo | A0A183F5N0 |
| Heligmosomo | A0A183F1S4 |
| Heligmosomo | A0A183G9U4 |
| Heligmosomo | A0A183FAM8 |
| Heligmosomo | A0A183GCA0 |
| Heligmosomo | A0A183GSN8 |
| Heligmosomo | A0A183FC10 |
| Heligmosomo | A0A183FLN4 |
| Heligmosomo | A0A183GR87 |
| Heligmosomo | A0A183FV70 |
| Heligmosomo | A0A183FLD7 |

| | |
|---|---|
| Heligmosomo | A0A183GDB3 |
| Heligmosomo | A0A183FE94 |
| Heligmosomo | A0A183FLN5 |
| Heligmosomo | A0A183GEW6 |
| Heligmosomo | A0A3P7TAF8 |
| Heligmosomo | A0A3P7XS44 |
| Heligmosomo | A0A3P7UBD8 |
| Heligmosomo | A0A3P7YBC3 |
| Heligmosomo | A0A3P8CJY4 |
| Heligmosomo | A0A3P8B899 |
| Heligmosomo | A0A3P8BPW5 |
| Heligmosomo | A0A3P7XWA8 |
| Heligmosomo | A0A3P8FFK8 |
| Heligmosomo | A0A3P8BG03 |
| Heligmosomo | A0A3P7Y3L6 |
| Heligmosomo | A0A3P8AS32 |
| Heligmosomo | A0A3P8DWK6 |
| Heligmosomo | A0A3P8CGZ0 |
| Heligmosomo | A0A3P7YEL9 |
| Heligmosomo | A0A3P8A3U8 |
| Heligmosomo | A0A3P7XYB2 |
| Heligmosomo | A0A3P8C2W2 |
| Heligmosomo | A0A3P8BGP5 |
| Heligmosomo | A0A3P8BSP4 |
| Heligmosomo | A0A3P8E3J7 |
| Heligmosomo | A0A3P7X235 |
| Heligmosomo | A0A3P8DNT7 |
| Heligmosomo | A0A3P7WBJ4 |
| Heliothis vire | A0A2A4K0R0 |
| Heliothis vire | A0A2A4J5Y3 |
| Heliothis vire | A0A2A4J736 |
| Heliothis vire | A0A2A4IUN6 |
| Heliothis vire | A0A2A4JF45 |
| Heliothis vire | A0A2A4JFF2 |
| Heliothis vire | A0A2A4K3D6 |
| Heliothis vire | A0A2A4JED9 |
| Heliothis vire | A0A2A4IXP5 |
| Helobdella ro | T1FFM4 |
| Helobdella ro | T1F4T5 |
| Helobdella ro | T1FEH2 |
| Helobdella ro | T1F969 |

| | |
|---|---|
| Helobdella ro | T1FKL0 |
| Helobdella ro | T1FFM5 |
| Helobdella ro | T1EQR6 |
| Helobdella ro | T1F4U5 |
| Helobdella ro | T1F4U0 |
| Helobdella ro | T1FB09 |
| Helobdella ro | T1EQR5 |
| Helobdella ro | T1F1E0 |
| Helobdella ro | T1FEJ2 |
| Helobdella ro | T1FUN0 |
| Helobdella ro | T1F4U7 |
| Helobdella ro | T1FJS5 |
| Helobdella ro | T1FFA4 |
| Helobdella ro | T1F854 |
| Helobdella ro | T1FVJ6 |
| Helobdella ro | T1F4U6 |
| Helobdella ro | T1F4U3 |
| Helobdella ro | T1FVK2 |
| Helobdella ro | T1F5M2 |
| Helobdella ro | T1FEF5 |
| Helobdella ro | T1FPY2 |
| Helobdella ro | T1FUT1 |
| Helobdella ro | T1ETQ4 |
| Helobdella ro | T1FVK1 |
| Helobdella ro | T1F970 |
| Helobdella ro | T1EVQ0 |
| Helobdella ro | T1F9B8 |
| Helobdella ro | T1FR64 |
| Helobdella ro | T1FBN8 |
| Helobdella ro | T1FCJ6 |
| Helobdella ro | T1FJA3 |
| Helobdella ro | T1F4T4 |
| Helobdella ro | T1FVK0 |
| Helobdella ro | T1F4T9 |
| Helobdella ro | T1FDP6 |
| Helobdella ro | T1EQR7 |
| Helobdella ro | T1FUN1 |
| Helobdella ro | T1EP89 |
| Helobdella ro | T1ET71 |
| Helobdella ro | T1FFD2 |
| Helobdella ro | T1FDN8 |

| | |
|---|---|
| Helobdella ro | T1FEJ8 |
| Helobdella ro | T1F4T6 |
| Helobdella ro | T1EQP1 |
| Helobdella ro | T1F4U4 |
| Helobdella ro | T1FPT1 |
| Helobdella ro | T1F960 |
| Helobdella ro | T1FLN5 |
| Helobdella ro | T1FB69 |
| Helobdella ro | T1F954 |
| Helobdella ro | T1EZ95 |
| Helobdella ro | T1EQP5 |
| Helobdella ro | T1FV32 |
| Helobdella ro | T1FKK7 |
| Helobdella ro | T1F659 |
| Helobdella ro | T1FLC6 |
| Helobdella ro | T1FJL9 |
| Helobdella ro | T1FDA4 |
| Helobdella ro | T1F4U8 |
| Helobdella ro | T1F916 |
| Helobdella ro | T1F973 |
| Helobdella ro | T1FHL6 |
| Helobdella ro | T1FJN5 |
| Helobdella ro | T1FUI3 |
| Helobdella ro | T1FLN6 |
| Helobdella ro | T1FJM1 |
| Helobdella ro | T1EQP4 |
| Helobdella ro | T1F4T3 |
| Helobdella ro | T1FBM8 |
| Helobdella ro | T1FS89 |
| Helobdella ro | T1EZ93 |
| Helobdella ro | T1FPY5 |
| Helobdella ro | T1FF45 |
| Helobdella ro | T1FEG8 |
| Helobdella ro | T1FPZ1 |
| Helobdella ro | T1EQN9 |
| Helobdella ro | T1FVV8 |
| Helobdella ro | T1EPB3 |
| Helobdella ro | T1FLR7 |
| Helobdella ro | T1F975 |
| Helobdella ro | T1FFM6 |
| Helobdella ro | T1FFG1 |

| | |
|---|---|
| Helobdella ro | T1FBA6 |
| Helobdella ro | T1FDR4 |
| Helobdella ro | T1FBP2 |
| Helobdella ro | T1F682 |
| Helobdella ro | T1FPY3 |
| Helobdella ro | T1EQP3 |
| Helobdella ro | T1F4U2 |
| Helobdella ro | T1FBN9 |
| Helobdella ro | T1F974 |
| Helobdella ro | T1FG81 |
| Helobdella ro | T1F4V1 |
| Helobdella ro | T1FBM7 |
| Helobdella ro | T1F4T7 |
| Helobdella ro | T1FG80 |
| Helobdella ro | T1FH28 |
| Helobdella ro | T1FUI8 |
| Helobdella ro | T1FB01 |
| Helobdella ro | T1ETA4 |
| Helobdella ro | T1FB02 |
| Helobdella ro | T1F4T8 |
| Helobdella ro | T1EQR0 |
| Helobdella ro | T1F174 |
| Henriciella al | A0A399RHT4 |
| Henriciella sp | A0A399RLR9 |
| Henriciella sp | A0A399RSL7 |
| Henriciella sp | A0A399RMI5 |
| Henriciella sp | A0A2D8YFK7 |
| Heterocepha | A0A0P6JB42 |
| Heterocepha | G5BZW9 |
| Heterocepha | G5BNV2 |
| Heterocepha | G5BNV1 |
| Heterocepha | G5B0R1 |
| Heterocepha | A0A0P6IYV4 |
| Heterocepha | A0A0P6JIW8 |
| Heterocepha | A0A0P6J2P0 |
| Heterocepha | G5BC53 |
| Heterorhabdi | A0A1I7W716 |
| Heterorhabdi | A0A1I7XH77 |
| Heterorhabdi | A0A1I7WWV5 |
| Heterorhabdi | A0A1I7XSR7 |
| Heterorhabdi | A0A1I7XAF8 |

| | |
|---|---|
| Heterorhabdi | A0A1I7XHQ1 |
| Heterorhabdi | A0A1I7X146 |
| Heterorhabdi | A0A1I7XM37 |
| Heterorhabdi | A0A1I7X745 |
| Heterorhabdi | A0A1I7XHZ8 |
| Heterorhabdi | A0A1I7XII8 |
| Heterorhabdi | A0A1I7X5P0 |
| Heterorhabdi | A0A1I7WM55 |
| Hippea marit | F2LUA4 |
| Hippocampus | A0A3Q2XCJ9 |
| Hippocampus | A0A3Q3D5K9 |
| Hippocampus | A0A3Q2YZJ7 |
| Hippocampus | A0A3Q2YUT9 |
| Hippocampus | A0A3Q2YLY3 |
| Hirondellea g | A0A2P2ICY2 |
| Hirschfeldia i | Q9SEB5 |
| Hirschfeldia i | Q9SEB2 |
| Hirschfeldia i | Q9SEB3 |
| Hirschfeldia i | Q9SEB6 |
| Hirschfeldia i | Q9SEB4 |
| Hirschfeldia i | Q9SEC5 |
| Hirsutella mi | A0A0F7ZU04 |
| Hirudovirus s | V5L2N5 |
| Hirudovirus s | V5L3N5 |
| Hirudovirus s | V5L3U0 |
| Hirundo rusti | A0A3M0JH56 |
| Hirundo rusti | A0A3M0KB79 |
| Hirundo rusti | A0A3M0KPT5 |
| Hoeflea halo | A0A286IFF5 |
| Hoeflea mari | A0A317PT30 |
| Hoeflea phot | A9D968 |
| Hoeflea sp. B | A0A098TDW6 |
| Hoeflea sp. B | A0A0F2PZU5 |
| Hoeflea sp. II | A0A0F7PKQ6 |
| Hoeflea sp..xl | A0A2G2DJB6 |
| Homalodisca | A0A1B6IND1 |
| Homalodisca | A0A1B6JBA6 |
| Homalodisca | A0A1B6IQ37 |
| Homalodisca | A0A1B6HRP0 |
| Homo sapien | P03951 |
| Homo sapien | P00747 |

Homo sapien P03952
Homo sapien P26927
Homo sapien P14210
Homo sapien Q02325
Homo sapien Q15195
Homo sapien Q2TV78
Homo sapien C9WSJ4
Homo sapien G3XAK1
Homo sapien B7Z3P3
Homo sapien A0A0F7G8J1
Homo sapien H0Y596
Homo sapien B4DMX2
Homo sapien C9J075
Homo sapien A8K6K9
Homo sapien C9JS80
Homo sapien B2R7F8
Homo sapien Q59H59
Homo sapien Q9UEG0
Homo sapien C9JCT1
Homo sapien C9JDP4
Homo sapien F8WCD6
Homo sapien Q9UE70
Homo sapien Q53QL7
Homo sapien E9PBC5
Homo sapien B4DPN1
Homo sapien B4DPQ8
Homo sapien A0A2R8YEI1
Homo sapien A6PVI2
Homo sapien B7Z538
Homo sapien A8K9A9
Homo sapien H0YAC1
Homo sapien D6RB32
Homo sapien Q5TEH5
Homo sapien C9WSJ3
Homo sapien B7Z250
Homo sapien Q13208
Homo sapien Q53GN8
Hondaea ferr A0A2R5GHI4
Hondaea ferr A0A2R5GZ90
Hondaea ferr A0A2R5G055
Hondaea ferr A0A2R5G2W7

| | |
|---|---|
| Hondaea ferr | A0A2R5GHT1 |
| Hondaea ferr | A0A2R5GPJ4 |
| Hondaea ferr | A0A2R5GQ72 |
| Hondaea ferr | A0A2R5GFH2 |
| Hondaea ferr | A0A2R5GZ49 |
| Hondaea ferr | A0A2R5GMR6 |
| Hondaea ferr | A0A2R5FCZ9 |
| Hondaea ferr | A0A2R5GLX6 |
| Hondaea ferr | A0A2R5GNL7 |
| Hondaea ferr | A0A2R5GI37 |
| Hondaea ferr | A0A2R5GR48 |
| Hondaea ferr | A0A2R5GP36 |
| Hondaea ferr | A0A2R5GCC8 |
| Hondaea ferr | A0A2R5GTZ6 |
| Hondaea ferr | A0A2R5GIQ6 |
| Hondaea ferr | A0A2R5GBG8 |
| Hondaea ferr | A0A2R5GIT4 |
| Hondaea ferr | A0A2R5G3T7 |
| Hondaea ferr | A0A2R5GCH5 |
| Hondaea ferr | A0A2R5GJ87 |
| Hondaea ferr | A0A2R5GGB6 |
| Hondaea ferr | A0A2R5G7Y5 |
| Hondaea ferr | A0A2R5GWX3 |
| Hondaea ferr | A0A2R5GQR5 |
| Hondaea ferr | A0A2R5GUR1 |
| Hondaea ferr | A0A2R5GIX0 |
| Hondaea ferr | A0A2R5GVR7 |
| Hondaea ferr | A0A2R5G8F7 |
| Hondaea ferr | A0A2R5GQH2 |
| Hondaea ferr | A0A2R5GRQ0 |
| Hondaea ferr | A0A2R5GDB9 |
| Hordeum vul | F2CSY2 |
| Hordeum vul | M0XR46 |
| Hordeum vul | A0A287JPN5 |
| Hordeum vul | F2CQZ9 |
| Hordeum vul | M0Z8K2 |
| Hordeum vul | A0A287JKX2 |
| Hordeum vul | A0A287IUF4 |
| Hordeum vul | F2CZD5 |
| Hordeum vul | A0A287KLF9 |
| Hordeum vul | M0YWF3 |

| | |
|---|---|
| Hordeum vul | M0Y2Z8 |
| Hordeum vul | A0A287JLH6 |
| Hordeum vul | A0A287N5Q1 |
| Hordeum vul | A0A287Q5F5 |
| Hordeum vul | M0X4U6 |
| Hordeum vul | A0A287JL69 |
| Hordeum vul | A0A287LS65 |
| Hordeum vul | M0XR48 |
| Hordeum vul | A0A287JKB3 |
| Hordeum vul | A0A287V5S3 |
| Hordeum vul | F2E6E0 |
| Hordeum vul | A0A287VXI1 |
| Hordeum vul | A0A287KLC6 |
| Hordeum vul | F2E5G0 |
| Hordeum vul | A0A287VXR9 |
| Hordeum vul | A0A287KLB4 |
| Hordeum vul | A0A287JL74 |
| Hordeum vul | A0A287JLD4 |
| Hordeum vul | M0X5T3 |
| Hordeum vul | A0A287JKR9 |
| Hordeum vul | M0X4U1 |
| Hordeum vul | M0WTQ2 |
| Hordeum vul | A0A287JL26 |
| Hordeum vul | A0A287S531 |
| Hordeum vul | M0X4U2 |
| Hordeum vul | A0A287LSB4 |
| Hordeum vul | M0XH34 |
| Hordeum vul | A0A287N5X8 |
| Hordeum vul | A0A287DZN4 |
| Hordeum vul | M0UVT7 |
| Hordeum vul | M0Y3I5 |
| Hordeum vul | A0A287JKM2 |
| Hordeum vul | A0A287SWW6 |
| Hordeum vul | M0Y2Q7 |
| Hordeum vul | A0A287JKJ1 |
| Hordeum vul | M0Y9R6 |
| Hordeum vul | A0A287MK73 |
| Hordeum vul | M0W648 |
| Hordeum vul | M0X5T4 |
| Hordeum vul | F2DE34 |
| Hordeum vul | A0A287KLB0 |

| | |
|---|---|
| Hordeum vul | A0A287JKS4 |
| Hordeum vul | A0A287LS98 |
| Hordeum vul | A0A287S544 |
| Hordeum vul | M0X4U8 |
| Hordeum vul | A0A287XZA8 |
| Hordeum vul | M0VMR6 |
| Hordeum vul | A0A287JPW4 |
| Hordeum vul | A0A287KLD9 |
| Hordeum vul | M0V1J8 |
| Hordeum vul | A0A287JPJ2 |
| Hordeum vul | A0A287JPN9 |
| Hordeum vul | A0A287JKU9 |
| Hordeum vul | M0YRS9 |
| Hordeum vul | M0W0I1 |
| Hordeum vul | A0A287Q137 |
| Hordeum vul | M0X645 |
| Hordeum vul | M0VB75 |
| Hordeum vul | A0A287XIX9 |
| Hordeum vul | A0A287JVX3 |
| Hordeum vul | M0VXQ3 |
| Hordeum vul | M0X4U9 |
| Hordeum vul | A0A287S793 |
| Hordeum vul | A0A287KLA1 |
| Hordeum vul | A0A287HTP8 |
| Hordeum vul | A0A287JKA6 |
| Hordeum vul | A0A287XAT4 |
| Hordeum vul | A0A287LSA0 |
| Hordeum vul | A0A287HU15 |
| Hordeum vul | A0A287LS94 |
| Hordeum vul | A0A287ULQ4 |
| Hordeum vul | A0A287G0I2 |
| Hordeum vul | A0A287I9B0 |
| Hordeum vul | A0A287X4W8 |
| Hordeum vul | A0A287V603 |
| Hordeum vul | A0A287JKW2 |
| Hordeum vul | A0A287VXJ1 |
| Hordeum vul | A0A287S581 |
| Hordeum vul | M0USE4 |
| Hordeum vul | A0A287G0I0 |
| Hordeum vul | M0YWF2 |
| Hordeum vul | M0YRS8 |

| | |
|---|---|
| Hordeum vul | A0A287EG31 |
| Hordeum vul | A0A287G0I1 |
| Hordeum vul | A0A287JPP7 |
| Hordeum vul | A0A287X536 |
| Hordeum vul | A0A287Q497 |
| Hordeum vul | A0A287JPV5 |
| Hordeum vul | F2DPF7 |
| Hordeum vul | A0A287JKP7 |
| Hordeum vul | A0A287H0I3 |
| Hordeum vul | F2E9C6 |
| Hordeum vul | A0A287SY82 |
| Hordeum vul | A0A287H0K6 |
| Hordeum vul | A0A287JKX4 |
| Hordeum vul | A0A287LSB2 |
| Hordeum vul | A0A287JPY8 |
| Hordeum vul | A0A287G0J3 |
| Hordeum vul | A0A287VSN4 |
| Hordeum vul | A0A287JKP4 |
| Hordeum vul | F2E736 |
| Hordeum vul | A0A287E1S1 |
| Hordeum vul | A0A287JKX3 |
| Hordeum vul | A0A287JVM6 |
| Hordeum vul | A0A287IUE8 |
| Hordeum vul | A0A287NM49 |
| Hordeum vul | M0WIJ2 |
| Hordeum vul | F2D8U5 |
| Hordeum vul | A0A287MNZ7 |
| Hordeum vul | A0A2R9IZN9 |
| Hordeum vul | F2CR46 |
| Hordeum vul | A0A287IUJ8 |
| Hordeum vul | A0A287S537 |
| Hordeum vul | A0A287S520 |
| Hordeum vul | A0A287VS95 |
| Hordeum vul | A0A287XZG6 |
| Hordeum vul | A0A287JKV6 |
| Hordeum vul | A0A287JK91 |
| Hordeum vul | M0UW74 |
| Hordeum vul | A0A287H0K5 |
| Hordeum vul | A0A287XMV2 |
| Hordeum vul | A0A287ETX1 |
| Hordeum vul | A0A287EJ56 |

| | |
|---|---|
| Hordeum vul | M0VKE6 |
| Hordeum vul | A0A287Q400 |
| Hordeum vul | A0A287IUH9 |
| Hordeum vul | A0A287H787 |
| Hordeum vul | A0A287Q1D6 |
| Hordeum vul | A0A287VSA5 |
| Hordeum vul | A0A287S509 |
| Hordeum vul | A0A287H719 |
| Hordeum vul | A0A287JKP2 |
| Hordeum vul | F2DZW7 |
| Hordeum vul | A0A287SK29 |
| Hordeum vul | A0A287JL87 |
| Hordeum vul | A0A287LSA4 |
| Hordeum vul | A0A287IGH2 |
| Hordeum vul | A0A287EE31 |
| Hordeum vul | F2DVH7 |
| Hordeum vul | A0A287XS45 |
| Hordeum vul | A0A287XIR7 |
| Hordeum vul | A0A287IGL1 |
| Hordeum vul | A0A287I9A0 |
| Hordeum vul | A0A287DZP5 |
| Hordeum vul | A0A287EG67 |
| Hordeum vul | M0XKB6 |
| Hordeum vul | A0A287MK90 |
| Hordeum vul | A0A287JPN4 |
| Hordeum vul | A0A287XIJ9 |
| Hordeum vul | A0A287JKI2 |
| Hordeum vul | A0A287JKU7 |
| Hordeum vul | A0A287X8T7 |
| Hordeum vul | A0A287JKL9 |
| Hordeum vul | M0US74 |
| Hordeum vul | A0A287H0Q6 |
| Hordeum vul | A0A287HTT3 |
| Hordeum vul | A0A287XAZ9 |
| Hordeum vul | F2DVP4 |
| Hordeum vul | A0A287IGC1 |
| Hordeum vul | A0A287LSA9 |
| Hordeum vul | A0A287JKZ7 |
| Hordeum vul | A0A287JDY8 |
| Hordeum vul | A0A287EXW8 |
| Hordeum vul | A0A287JPK4 |

| | |
|---|---|
| Hordeum vul | A0A287XJ32 |
| Hordeum vul | A0A287XS09 |
| Hordeum vul | A0A287H0I2 |
| Hordeum vul | A0A287GWW0 |
| Hordeum vul | A0A287JKA0 |
| Hordeum vul | A0A287GYF4 |
| Hordeum vul | A0A287JKJ3 |
| Hordeum vul | A0A287HU10 |
| Hordeum vul | A0A287JKV4 |
| Hordeum vul | A0A287JLG8 |
| Hordeum vul | A0A287JDY3 |
| Hordeum vul | M0YMH8 |
| Hordeum vul | A0A287JKH8 |
| Hordeum vul | A0A287HTI0 |
| Hordeum vul | M0WAI6 |
| Hordeum vul | A0A287I9D8 |
| Hordeum vul | A0A287LSE2 |
| Hordeum vul | A0A287LSE8 |
| Hordeum vul | F2DDM1 |
| Hordeum vul | A0A287Q5G8 |
| Hordeum vul | A0A287JKQ6 |
| Hordeum vul | A0A287LSE1 |
| Hordeum vul | A0A287LSD9 |
| Hordeum vul | A0A287S577 |
| Hordeum vul | A0A287JDM9 |
| Hordeum vul | A0A287XIY2 |
| Hordeum vul | M0W0I0 |
| Hordeum vul | A0A287V653 |
| Hordeum vul | A0A287JKI8 |
| Hordeum vul | A0A287EJ83 |
| Hordeum vul | A0A287JKY5 |
| Hordeum vul | A0A287JKY7 |
| Hordeum vul | A0A287H0I8 |
| Hordeum vul | A0A287JL01 |
| Hordeum vul | A0A287N0H0 |
| Hordeum vul | M0W0I5 |
| Hordeum vul | F2DZS5 |
| Hordeum vul | M0UIL6 |
| Hordeum vul | A0A287JL57 |
| Hordeum vul | A0A287EDZ4 |
| Hordeum vul | M0UVT6 |

| | |
|---|---|
| Hordeum vul | A0A287LSB9 |
| Hordeum vul | A0A287H0H7 |
| Hordeum vul | A0A287LS90 |
| Hordeum vul | A0A287LSA1 |
| Hordeum vul | A0A287JKZ5 |
| Hordeum vul | A0A287JKQ5 |
| Hordeum vul | A0A287IGK1 |
| Hordeum vul | M0UDY0 |
| Hordeum vul | M0WBF9 |
| Hordeum vul | A0A287JL20 |
| Hordeum vul | A0A287XIT5 |
| Hordeum vul | A0A287H0J8 |
| Hordeum vul | A0A287MK96 |
| Hordeum vul | A0A287GX15 |
| Hordeum vul | A0A287JKZ9 |
| Hordeum vul | A0A287L9W8 |
| Hordeum vul | A0A287G0I8 |
| Hordeum vul | A0A287LSA5 |
| Hordeum vul | A0A287Q632 |
| Hordeum vul | A0A287JKZ4 |
| Hordeum vul | A0A287KQ24 |
| Hordeum vul | A0A287M8A3 |
| Hordeum vul | A0A287LS92 |
| Hordeum vul | A0A287S515 |
| Hordeum vul | A0A287JDV3 |
| Hordeum vul | A0A287HU21 |
| Hordeum vul | A0A287XS50 |
| Hordeum vul | A0A287E1N3 |
| Hordeum vul | A0A287N0G0 |
| Hordeum vul | A0A287M896 |
| Hordeum vul | A0A287JKZ2 |
| Hordeum vul | M0XZ56 |
| Hordeum vul | A0A287JKM9 |
| Hordeum vul | A0A287V7H0 |
| Hordeum vul | A0A287JKR1 |
| Hordeum vul | A0A287L9U3 |
| Hordeum vul | A0A287V5K1 |
| Hordeum vul | A0A287HU00 |
| Hordeum vul | F2DV22 |
| Hordeum vul | A0A287S4V9 |
| Hordeum vul | A0A287JL54 |

| | |
|---|---|
| Hordeum vul | A0A287NLW9 |
| Hordeum vul | A0A287LS76 |
| Hordeum vul | A0A287EGH8 |
| Hordeum vul | A0A287JKU5 |
| Hordeum vul | A0A287XIJ8 |
| Hordeum vul | A0A287EG37 |
| Hordeum vul | A0A287E1N0 |
| Hordeum vul | F2DZG8 |
| Hordeum vul | F2DWF4 |
| Hordeum vul | A0A287JDQ7 |
| Hordeum vul | A0A287KH06 |
| Hordeum vul | A0A287VXV7 |
| Hordeum vul | A0A287GFZ2 |
| Hordeum vul | A0A287MK60 |
| Hordeum vul | A0A287H0J7 |
| Hordeum vul | A0A287H0H8 |
| Hordeum vul | A0A287E1M5 |
| Hordeum vul | A0A287EJ40 |
| Hordeum vul | A0A287G0I5 |
| Hordeum vul | A0A287I9D4 |
| Hordeum vul | F2D0V2 |
| Hordeum vul | A0A287HTP3 |
| Hordeum vul | A0A287XIZ1 |
| Hordeum vul | F2E792 |
| Hordeum vul | A0A287H0N6 |
| Hordeum vul | A0A287JPZ0 |
| Hordeum vul | A0A287S554 |
| Hordeum vul | A0A287JLI4 |
| Hordeum vul | A0A287LS59 |
| Hordeum vul | A0A287U4M0 |
| Hordeum vul | A0A287MK62 |
| Hordeum vul | M0X8T3 |
| Hordeum vul | F2D0M2 |
| Hordeum vul | A0A287ETW5 |
| Hordeum vul | A0A287JKR4 |
| Hordeum vul | A0A287G0I7 |
| Hordeum vul | A0A287EG04 |
| Hordeum vul | A0A287S504 |
| Hordeum vul | A0A287XAP8 |
| Hordeum vul | M0UQL7 |
| Hordeum vul | M0X1P9 |

| | |
|---|---|
| Hordeum vul | A0A287JKQ4 |
| Hordeum vul | A0A287JKG2 |
| Hordeum vul | F2DJ24 |
| Hordeum vul | A0A287JKD6 |
| Hordeum vul | A0A287T4N7 |
| Hordeum vul | F2D6A9 |
| Hordeum vul | A0A287S550 |
| Hordeum vul | A0A287S507 |
| Hordeum vul | A0A287JKY8 |
| Hordeum vul | F2D857 |
| Hordeum vul | F2E0N1 |
| Hordeum vul | F2D0S6 |
| Hordeum vul | A0A287HTS3 |
| Hordeum vul | A0A287HU22 |
| Hordeum vul | A0A287N5T2 |
| Hordeum vul | F2E071 |
| Hordeum vul | A0A287LSD5 |
| Hordeum vul | A0A287SWR5 |
| Hordeum vul | F2DYG0 |
| Hordeum vul | A0A287E1M9 |
| Hordeum vul | A0A287TTW0 |
| Hordeum vul | F2E424 |
| Hordeum vul | A0A287T1C1 |
| Hordeum vul | A0A287IGI8 |
| Hordeum vul | A0A287LSC1 |
| Hordeum vul | F2DGX1 |
| Hordeum vul | A0A287N5Y3 |
| Hordeum vul | A0A287LSD3 |
| Hordeum vul | A0A287XS04 |
| Hordeum vul | A0A287HTT9 |
| Hordeum vul | A0A287S5B0 |
| Hordeum vul | A0A287LSI8 |
| Hordeum vul | A0A287JKK0 |
| Hordeum vul | A0A287TU03 |
| Hordeum vul | F2DJ90 |
| Hordeum vul | A0A287S5A2 |
| Hordeum vul | A0A287JE19 |
| Hordeum vul | A0A287JKS0 |
| Hordeum vul | A0A287H785 |
| Hordeum vul | A0A287G0H8 |
| Hordeum vul | F2EEQ7 |

| | |
|---|---|
| Hordeum vul | M0X5F1 |
| Hordeum vul | A0A287LSD4 |
| Hordeum vul | F2CSL7 |
| Hordeum vul | A0A287LSF0 |
| Hordeum vul | A0A287N5T8 |
| Hordeum vul | A0A287XZF8 |
| Hordeum vul | M0Y1V6 |
| Hordeum vul | A0A287G0H7 |
| Hordeum vul | A0A287JK96 |
| Hordeum vul | A0A287T4Y6 |
| Hordeum vul | A0A287G0I6 |
| Hordeum vul | A0A287XSE7 |
| Hordeum vul | F2DBV1 |
| Hordeum vul | A0A287LSC8 |
| Hordeum vul | A0A287JKV9 |
| Hordeum vul | A0A287KLG4 |
| Hordeum vul | A0A287JPP4 |
| Hordeum vul | A0A287JKI4 |
| Hordeum vul | A0A287JLH1 |
| Hordeum vul | A0A287JL30 |
| Hordeum vul | A0A287LS79 |
| Hordeum vul | A0A287JKX1 |
| Hordeum vul | A0A287SWV1 |
| Hordeum vul | A0A287XMW2 |
| Hordeum vul | A0A287LS85 |
| Hordeum vul | A0A287SX07 |
| Hordeum vul | A0A287VXG8 |
| Hordeum vul | A0A287V5Z3 |
| Hordeum vul | A0A287JGN6 |
| Hordeum vul | A0A287S4X9 |
| Hordeum vul | A0A287JKT0 |
| Hordeum vul | A0A287XIU0 |
| Hordeum vul | A0A287IGE2 |
| Hordeum vul | A0A287JPV4 |
| Hordeum vul | A0A287JL84 |
| Hordeum vul | A0A287H0R0 |
| Hordeum vul | F2DSN1 |
| Hordeum vul | A0A287IGB2 |
| Hordeum vul | A0A287SYA0 |
| Hordeum vul | A0A287L9W9 |
| Hordeum vul | A0A287SWS0 |

| | |
|---|---|
| Hordeum vul | A0A287VDJ7 |
| Hordeum vul | M0V508 |
| Hordeum vul | A0A287XAQ3 |
| Hordeum vul | F2ED78 |
| Hordeum vul | A0A287JLE3 |
| Hordeum vul | A0A287JKK2 |
| Hordeum vul | A0A287H0L7 |
| Hordeum vul | A0A287XIX4 |
| Hordeum vul | A0A287JKY3 |
| Hordeum vul | A0A287V5G7 |
| Hordeum vul | A0A287NLN4 |
| Hordeum vul | A0A287JVK7 |
| Hordeum vul | M0UR24 |
| Hordeum vul | A0A287E1L3 |
| Hordeum vul | F2D4F6 |
| Hordeum vul | F2DZY6 |
| Hordeum vul | A0A287JKH3 |
| Hordeum vul | M0V5N5 |
| Hordeum vul | M0XKB5 |
| Hordeum vul | A0A287H0P5 |
| Hordeum vul | A0A287S580 |
| Hordeum vul | A0A287JK95 |
| Hordeum vul | M0YRS5 |
| Hordeum vul | A0A287S4W4 |
| Hordeum vul | A0A287H0I7 |
| Hordeum vul | M0YDW1 |
| Hordeum vul | F2E6M1 |
| Hordeum vul | A0A2R9IZN3 |
| Hordeum vul | F2CXU1 |
| Hordeum vul | M0UFX9 |
| Hordeum vul | A0A287MK66 |
| Hordeum vul | A0A287KLC4 |
| Hordeum vul | A0A287FZA9 |
| Hordeum vul | A0A287H0L1 |
| Hordeum vul | A0A287L9W2 |
| Hordeum vul | A0A287N0F8 |
| Hordeum vul | A0A287M890 |
| Hordeum vul | A0A287JKR0 |
| Hordeum vul | A0A287S5E8 |
| Hordeum vul | A0A287JL81 |
| Hordeum vul | A0A287JL40 |

| | |
|---|---|
| Hordeum vul | A0A287SY15 |
| Hordeum vul | A0A287V5G1 |
| Hordeum vul | A0A287Q1Y2 |
| Hordeum vul | A0A287I9D7 |
| Hordeum vul | A0A287JKD0 |
| Hordeum vul | A0A287JKQ0 |
| Hordeum vul | A0A287EJ45 |
| Hordeum vul | A0A287JGA6 |
| Hordeum vul | A0A287HTT1 |
| Hordeum vul | M0YMI0 |
| Hordeum vul | A0A287IGH4 |
| Hordeum vul | A0A287HTR9 |
| Hordeum vul | A0A287XIR2 |
| Hordeum vul | A0A287IUF8 |
| Hordeum vul | A0A287XS55 |
| Hordeum vul | A0A287X4U2 |
| Hordeum vul | A0A287KGI7 |
| Hordeum vul | A0A287JKN7 |
| Hordeum vul | A0A287LS81 |
| Hordeum vul | A0A287V5I8 |
| Hordeum vul | A0A287Q176 |
| Hordeum vul | A0A287JKE5 |
| Hordeum vul | A0A287GG05 |
| Hordeum vul | A0A287JKL1 |
| Hordeum vul | A0A287IUE9 |
| Hordeum vul | A0A287IUL0 |
| Hordeum vul | M0YMH9 |
| Hordeum vul | A0A287JL90 |
| Hordeum vul | A0A287XAW7 |
| Hordeum vul | A0A287H0P0 |
| Hordeum vul | A0A287JKK7 |
| Hordeum vul | F2DQ07 |
| Hordeum vul | F2DVF8 |
| Hordeum vul | A0A287EXV5 |
| Hordeum vul | A0A287EXX5 |
| Hordeum vul | A0A287EG25 |
| Hordeum vul | A0A287XZG3 |
| Hordeum vul | A0A287XBL1 |
| Hordeum vul | A0A287JKV7 |
| Hordeum vul | A0A287G0I4 |
| Hordeum vul | A0A287V5J4 |

| | |
|---|---|
| Hordeum vul | A0A287X4U7 |
| Hordeum vul | A0A287S5B5 |
| Hordeum vul | A0A287EG62 |
| Hordeum vul | A0A287JDT8 |
| Hordeum vul | A0A287JPY5 |
| Hordeum vul | A0A287EG71 |
| Hordeum vul | A0A287JKX7 |
| Hordeum vul | A0A287JLI6 |
| Hordeum vul | A0A287N0F4 |
| Hordeum vul | A0A287HU27 |
| Hordeum vul | A0A287G0J5 |
| Hordeum vul | A0A287MK69 |
| Hordeum vul | F2DP44 |
| Hordeum vul | A0A287Q173 |
| Hordeum vul | A0A287I9A6 |
| Hordeum vul | A0A287ULB9 |
| Hordeum vul | A0A287XSC3 |
| Hordeum vul | A0A287JKF5 |
| Hordeum vul | A0A287XRW5 |
| Hordeum vul | A0A287N5W5 |
| Hordeum vul | A0A287HU05 |
| Hordeum vul | A0A287Q638 |
| Hordeum vul | F2DDU9 |
| Hordeum vul | F2EJV9 |
| Hordeum vul | A0A287UVQ5 |
| Hordeum vul | A0A287LS87 |
| Hordeum vul | A0A287LS64 |
| Hordeum vul | A0A287E1P0 |
| Hordeum vul | A0A287I9C6 |
| Hordeum vul | A0A287KLC0 |
| Hordeum vul | A0A287MUT0 |
| Hordeum vul | F2DUT1 |
| Hordeum vul | M0W647 |
| Hordeum vul | A0A287JGC0 |
| Hordeum vul | M0X5T2 |
| Hordeum vul | A0A287KLF4 |
| Hordeum vul | F2DGU3 |
| Hordeum vul | M0XSP1 |
| Hordeum vul | M0W3T9 |
| Hordeum vul | A0A287I9C1 |
| Hordeum vul | A0A287JL89 |

| | |
|---|---|
| Hordeum vul | A0A287H0J6 |
| Hordeum vul | A0A287EFT8 |
| Hordeum vul | A0A287K4Z4 |
| Hordeum vul | A0A287FZB8 |
| Hordeum vul | A0A287XIW0 |
| Hordeum vul | A0A287KGP2 |
| Hordeum vul | A0A287NLR1 |
| Hordeum vul | A0A287SYB4 |
| Hordeum vul | A0A287H0L5 |
| Hordeum vul | A0A287HU06 |
| Hordeum vul | A0A287HTJ9 |
| Hordeum vul | A0A287KLD6 |
| Hordeum vul | A0A287V5F6 |
| Hordeum vul | A0A287LSC4 |
| Hordeum vul | A0A287EG21 |
| Hordeum vul | A0A287VST9 |
| Hordeum vul | A0A287LSA6 |
| Hordeum vul | A0A287S505 |
| Hordeum vul | A0A287H0K2 |
| Hordeum vul | A0A287JKK1 |
| Hordeum vul | A0A287HU32 |
| Hordeum vul | A0A287IUK4 |
| Hordeum vul | A0A287JGI4 |
| Hordeum vul | M0YSV2 |
| Hordeum vul | A0A287G0I9 |
| Hordeum vul | A0A287GY94 |
| Hordeum vul | A0A287JK87 |
| Hordeum vul | A0A287JKX9 |
| Hordeum vul | M0XCH9 |
| Hordeum vul | A0A287LS99 |
| Hordeum vul | A0A287S571 |
| Hortaea werr | A0A1Z5T7T5 |
| Hortaea werr | A0A1Z5T1G6 |
| Hortaea werr | A0A1Z5SZU9 |
| Hortaea werr | A0A1Z5TRS5 |
| Hortaea werr | A0A1Z5SNU5 |
| Hortaea werr | A0A1Z5T2F6 |
| Hortaea werr | A0A1Z5TKS3 |
| Hortaea werr | A0A1Z5T220 |
| Hortaea werr | A0A1Z5T860 |
| Hortaea werr | A0A3M6XEF2 |

Hortaea werr A0A3M6YRP2
Hortaea werr A0A3M7B8I9
Hortaea werr A0A3M7BD57
Hortaea werr A0A3M7EN81
Hortaea werr A0A3M7IHZ9
Hortaea werr A0A3M7F5J7
Hortaea werr A0A3M7CY13
Hortaea werr A0A3M7IF39
Hortaea werr A0A3M7IKN2
Hortaea werr A0A3M7F3Z2
Hortaea werr A0A3M7J989
Hortaea werr A0A3M7DTD4
Hortaea werr A0A3M6X1R5
Hortaea werr A0A3M7B9I1
Hortaea werr A0A3M7HRW2
Hortaea werr A0A3M6ZPM6
Hortaea werr A0A3M7A6V9
Hortaea werr A0A3M7CW34
Hortaea werr A0A3M7FE79
Hortaea werr A0A3M6Z7L5
Hortaea werr A0A3M6WLT3
Hortaea werr A0A3M7IN62
Hortaea werr A0A3M7AEZ4
Hortaea werr A0A3M7D1M2
Hortaea werr A0A3M7GWG3
Hortaea werr A0A3M6YTF6
Hortaea werr A0A3M7FEQ6
Hortaea werr A0A3M7GDI6
Hortaea werr A0A3M7GKN4
Hortaea werr A0A3M7F5S1
Hortaea werr A0A3M7H9H8
Hortaea werr A0A3M7E7K4
Hortaea werr A0A3M7BMV2
Hortaea werr A0A3M6YFN9
Hortaea werr A0A3M7HTY7
Hortaea werr A0A3M7C6P1
Hortaea werr A0A3M6Y8Q0
Hortaea werr A0A3M7GBS4
Hortaea werr A0A3M7FVY9
Hortaea werr A0A3M7AI77
Hortaea werr A0A3M7H9D0

Hortaea werr A0A3M7F5D2
Hortaea werr A0A3M7EZS7
Hortaea werr A0A3M6Y2F1
Hortaea werr A0A3M7EXQ6
Hortaea werr A0A3M7CWK5
Hortaea werr A0A3M7FQH4
Hortaea werr A0A3M7FKN6
Hortaea werr A0A3M7CZJ1
Hortaea werr A0A3M7HXS6
Hortaea werr A0A3M7ITW6
Hortaea werr A0A3M7DCL5
Hortaea werr A0A3M7FY57
Hortaea werr A0A3M7AY00
Hortaea werr A0A3M7HF76
Hortaea werr A0A3M6XNG0
Hortaea werr A0A3M7IR78
Hortaea werr A0A3M7D168
Hortaea werr A0A3M7EZ41
Hortaea werr A0A3M7E391
Hortaea werr A0A3M7GVU9
Hortaea werr A0A3M7FYN0
Hortaea werr A0A3M7BRN4
Hortaea werr A0A3M6X571
Hortaea werr A0A3M6XIS4
Hortaea werr A0A3M7IC91
Hortaea werr A0A3M7F1K5
Hortaea werr A0A3M6WR01
Hortaea werr A0A3M6WED0
Hortaea werr A0A3M7BI24
Hortaea werr A0A3M7BYP9
Hortaea werr A0A3M7EB66
Hortaea werr A0A3M7ETI4
Hortaea werr A0A3M7HLN2
Hortaea werr A0A3M7DQ18
Hortaea werr A0A3M6X7W9
Hortaea werr A0A3M7C0U4
Hortaea werr A0A3M7AI42
Hortaea werr A0A3M7DXT2
Hortaea werr A0A3M7CVC3
Hortaea werr A0A3M7EYQ9
Hortaea werr A0A3M7DQ84

| | |
|---|---|
| Hortaea werr | A0A3M7I235 |
| Hortaea werr | A0A3M7IA13 |
| Hyalangium r | A0A085W504 |
| Hyaloperono: | M4B6K3 |
| Hyaloperono: | M4B6K2 |
| Hyaloscypha | A0A2J6RSW5 |
| Hyaloscypha | A0A2J6RMZ2 |
| Hyaloscypha | A0A2J6RSV8 |
| Hyaloscypha | A0A2J6RGD2 |
| Hyaloscypha | A0A2J6RVZ8 |
| Hyaloscypha | A0A2J6S4F4 |
| Hyaloscypha | A0A2J6R4R6 |
| Hyaloscypha | A0A2J6RYD0 |
| Hydatigena ta | A0A0R3WZ01 |
| Hydractinia s | A8S0U4 |
| Hydrogenoph | A0A2M7IBC9 |
| hydrotherma | A0A3B0T141 |
| hydrotherma | A0A170PTA1 |
| hydrotherma | A0A3B0T4U7 |
| hydrotherma | A0A3B0S6G0 |
| hydrotherma | A0A3B0YDW4 |
| hydrotherma | A0A3B0RX13 |
| Hymenolepis | A0A158QEB2 |
| Hymenolepis | A0A3P7BEC2 |
| Hymenolepis | A0A158QBS7 |
| Hyphomicrob | A0A1I7NHV8 |
| Hyphomicrob | A0A2P5KXM8 |
| Hyphomicrob | A0A2P5KRL9 |
| Hyphomicrob | A0A2S5PBC1 |
| Hyphomicrob | A0A2S5PC10 |
| Hyphomicrob | A0A2S5P4Z9 |
| Hyphomonac | A0A0F2QD64 |
| Hyphomonac | A0A1Z9YBG4 |
| Hyphomonac | A0A2D8ZSJ2 |
| Hyphomonac | A0A2E3AXJ3 |
| Hyphomonac | A0A2E4S0J7 |
| Hyphomonac | A0A2D8ZV35 |
| Hyphomonas | A0A069E6T4 |
| Hyphomonas | A0A3B9H2G8 |
| Hyphomonas | A0A059DX53 |
| Hyphomonas | A0A3B9KYM9 |

| | |
|---|---|
| Hyphomonas | A0A062U7B1 |
| Hyphomonas | A0A062UCV7 |
| Hyphomonas | A0A059FLM4 |
| Hyphomonas | A0A059G005 |
| Hyphomonas | A0A059F7I7 |
| Hyphomonas | A0A059FSS0 |
| Hyphomonas | Q0BYL9 |
| Hyphomonas | Q0C462 |
| Hyphomonas | A0A059G8G7 |
| Hyphomonas | A0A062VJB5 |
| Hyphomonas | A0A258BI69 |
| Hyphomonas | A0A259LR84 |
| Hyphomonas | A0A0F2RG41 |
| Hyphomonas | A0A328JPZ2 |
| Hyphomonas | A0A344WD66 |
| Hyphomonas | A0A059DSX7 |
| Hyphomonas | A0A059EES6 |
| Hyphomonas | A0A062TZP2 |
| Hyphomonas | A0A2D8W5J1 |
| Hyphomonas | A0A3D2LBT6 |
| Hyphomonas | A0A350ZTY8 |
| Hyphomonas | A0A2D8ER08 |
| Hyphomonas | A0A2E7B2U2 |
| Hyphomonas | A0A2E3JXR2 |
| Hypocrea atr | G9P8Z6 |
| Hypocrea atr | G9NQ37 |
| Hypocrea jec | A0A024SGL6 |
| Hypocrea jec | A0A024SLI6 |
| Hypocrea jec | G0RDV7 |
| Hypocrea jec | G0RFA4 |
| Hypocrea vire | G9MYE6 |
| Hypocrea vire | G9NAA5 |
| Hypocrea vire | G9NBF0 |
| Hypocrea vire | G9MT46 |
| Hypoxylon sp | A0A1Y2W191 |
| Hypoxylon sp | A0A1Y2VQD0 |
| Hypoxylon sp | A0A1Y2VLV2 |
| Hypoxylon sp | A0A1Y2UVW5 |
| Hypoxylon sp | A0A1Y2V838 |
| Hypoxylon sp | A0A1Y2V1I2 |
| Hypoxylon sp | A0A1Y2TBN0 |

| | |
|---|---|
| Hypoxylon sp | A0A1Y2U492 |
| Hypoxylon sp | A0A1Y2TUW0 |
| Hypoxylon sp | A0A1Y2TMZ5 |
| Hypsibius duj | A0A1W0X097 |
| Hypsibius duj | A0A1W0WIH9 |
| Hypsibius duj | A0A1W0WWV0 |
| Hypsibius duj | A0A1W0X6K9 |
| Hypsibius duj | A0A1W0X4X0 |
| Hypsibius duj | A0A1W0WI98 |
| Hypsibius duj | A0A1W0WIM8 |
| Hypsibius duj | A0A1W0WIA3 |
| Hypsibius duj | A0A1W0WQ69 |
| Hypsibius duj | A0A1W0WPS6 |
| Hypsibius duj | A0A1W0WNF7 |
| Hypsibius duj | A0A1W0X6A8 |
| Hypsibius duj | A0A1W0X3K6 |
| Hypsibius duj | A0A1W0WF75 |
| Hypsibius duj | A0A1W0XCI5 |
| Hypsibius duj | A0A1W0WSL7 |
| Hypsibius duj | A0A1W0X0I9 |
| Hypsibius duj | A0A1W0WBY5 |
| Hypsibius duj | A0A1W0X277 |
| Hypsibius duj | A0A1W0X910 |
| Hypsibius duj | A0A1W0WDI1 |
| Hypsibius duj | A0A1W0WXQ4 |
| Hypsibius duj | A0A1W0X8W3 |
| Hypsibius duj | A0A1W0WR27 |
| Hypsibius duj | A0A1W0WFY4 |
| Hypsibius duj | A0A1W0WII5 |
| Hypsibius duj | A0A1W0WII2 |
| Hypsibius duj | A0A1W0WLH6 |
| Hypsibius duj | A0A1W0WG72 |
| Hypsibius duj | A0A1W0WBT5 |
| Iconisemion s | A0A1A7YAK1 |
| Iconisemion s | A0A1A7WJK9 |
| Ictalurus pun | A0A2D0S0Y9 |
| Ictalurus pun | W5UJ51 |
| Ictalurus pun | A0A2D0PYF3 |
| Ictalurus pun | A0A2D0PVZ8 |
| Ictalurus pun | A0A2D0PZW9 |
| Ictalurus pun | A0A2D0S0B2 |

| | |
|---|---|
| Ictalurus pun | A0A2D0PX86 |
| Ictalurus pun | A0A2D0PN50 |
| Ictalurus pun | A0A2D0PUK1 |
| Ictalurus pun | A0A2D0PXQ6 |
| Ictalurus pun | A0A2D0PYM6 |
| Ictalurus pun | A0A2D0PW18 |
| Ictalurus pun | W5UM75 |
| Ictalurus pun | W5UH86 |
| Ictidomys tric | I3ME70 |
| Ictidomys tric | I3NFV2 |
| Ictidomys tric | I3M1N9 |
| Ictidomys tric | I3MLI1 |
| Ictidomys tric | I3ME62 |
| Imhoffiella pu | W9VHP2 |
| Intoshia linei. | A0A177B984 |
| Intoshia linei. | A0A177B686 |
| Ipomoea trifi | Q40096 |
| Ipomoea trifi | Q40100 |
| Ipomoea trifi | Q40099 |
| Ipomoea trifi | Q40098 |
| Ixodes ricinus | V5HBD1 |
| Ixodes ricinus | V5H4U1 |
| Ixodes ricinus | V5HJ86 |
| Ixodes ricinus | A0A147BGK7 |
| Ixodes ricinus | A0A131YAQ2 |
| Ixodes ricinus | A0A131XWE4 |
| Ixodes ricinus | A0A1C8FEE7 |
| Ixodes ricinus | V5HEL7 |
| Ixodes ricinus | A0A147BMG4 |
| Ixodes ricinus | V5ICT5 |
| Ixodes scapul | B7QCV3 |
| Ixodes scapul | B7Q2N4 |
| Ixodes scapul | B7QM85 |
| Ixodes scapul | B7P1S6 |
| Ixodes scapul | B7P6R1 |
| Ixodes scapul | B7QDU0 |
| Ixodes scapul | B7PWP4 |
| Ixodes scapul | B7PQI9 |
| Ixodes scapul | B7Q7Z0 |
| Ixodes scapul | B7QM84 |
| Ixodes scapul | B7QM83 |

| | |
|---|---|
| Ixodes scapul | B7PM04 |
| Ixodes scapul | B7PQI8 |
| Jaminaea ros | A0A316V1R6 |
| Jannaschia pc | A0A1I3UGX8 |
| Jannaschia se | A0A2Y9B448 |
| Jannaschia sp | Q28VT1 |
| Jannaschia sp | A0A1B6YPR7 |
| Jannaschia sp | A0A1B6Z0H5 |
| Jatropha curc | A0A067LHY6 |
| Jatropha curc | A0A067KUJ8 |
| Jatropha curc | A0A067L0Z2 |
| Jatropha curc | A0A067L5M7 |
| Jatropha curc | A0A067JDL7 |
| Jatropha curc | A0A067L394 |
| Jatropha curc | A0A067JQ57 |
| Jatropha curc | A0A067JDL1 |
| Jatropha curc | A0A067LB03 |
| Jatropha curc | A0A067KQJ4 |
| Jatropha curc | A0A067JMW2 |
| Jatropha curc | A0A067K6D5 |
| Jatropha curc | A0A067KR73 |
| Jatropha curc | A0A067JMV5 |
| Jatropha curc | A0A067KZS2 |
| Jatropha curc | A0A067KVG4 |
| Jatropha curc | A0A067KS71 |
| Jatropha curc | A0A067KMS7 |
| Jatropha curc | A0A067JLG1 |
| Jatropha curc | A0A067KWS2 |
| Jatropha curc | A0A067JLG5 |
| Jatropha curc | A0A067KVF4 |
| Jatropha curc | A0A067JMV9 |
| Jatropha curc | A0A067L3A3 |
| Jatropha curc | A0A067LB09 |
| Jatropha curc | A0A067KXT3 |
| Jatropha curc | A0A067JDL4 |
| Jatropha curc | A0A067JFW9 |
| Jatropha curc | A0A067KRX7 |
| Jatropha curc | A0A067LDD5 |
| Jatropha curc | A0A067L3A8 |
| Jatropha curc | A0A067KRY8 |
| Jatropha curc | A0A067JLG8 |

| | |
|---|---|
| Jatropha curc | A0A067L4G0 |
| Jatropha curc | A0A067JAL4 |
| Jatropha curc | A0A067JQM4 |
| Jatropha curc | A0A067KEY5 |
| Jatropha curc | A0A067KZQ0 |
| Jatropha curc | A0A067JQ92 |
| Jatropha curc | A0A067JII4 |
| Jatropha curc | A0A067K362 |
| Jatropha curc | A0A067JAK6 |
| Jatropha curc | A0A067L4F6 |
| Jatropha curc | A0A067L4H1 |
| Jatropha curc | A0A067KVG0 |
| Jatropha curc | A0A067JAQ2 |
| Jatropha curc | A0A067L398 |
| Jatropha curc | A0A067L3B7 |
| Jatropha curc | A0A067LDW4 |
| Jatropha curc | A0A067JAQ7 |
| Jatropha curc | A0A067JWU9 |
| Jatropha curc | A0A067L392 |
| Jatropha curc | A0A067K7M8 |
| Jatropha curc | A0A067KS66 |
| Jatropha curc | A0A067KRX3 |
| Jatropha curc | A0A067KVG9 |
| Jatropha curc | A0A067KB60 |
| Jatropha curc | A0A067LBK7 |
| Jatropha curc | A0A067KS82 |
| Jatropha curc | A0A067K9W6 |
| Jatropha curc | A0A067KF77 |
| Jeongeupia sp | A0A1D9BCC6 |
| Juglans regia. | A0A2I4GK62 |
| Juglans regia. | A0A2I4G4M2 |
| Juglans regia. | A0A2I4H823 |
| Juglans regia. | A0A2I4FLZ3 |
| Juglans regia. | A0A2I4FU95 |
| Juglans regia. | A0A2I4DI82 |
| Juglans regia. | A0A2I4EPU3 |
| Juglans regia. | A0A2I4E0M1 |
| Juglans regia. | A0A2I4HI12 |
| Juglans regia. | A0A2I4H7B3 |
| Juglans regia. | A0A2I4GAB4 |
| Juglans regia. | A0A2I4E0L2 |

Juglans regia. A0A2I4FZE8
Juglans regia. A0A2I4HNL1
Juglans regia. A0A2I4FFM6
Juglans regia. A0A2I4EKA5
Juglans regia. A0A2I4ERC6
Juglans regia. A0A2I4H782
Juglans regia. A0A2I4H784
Juglans regia. A0A2I4HSV7
Juglans regia. A0A2I4EKB9
Juglans regia. A0A2I4G4H1
Juglans regia. A0A2I4HV68
Juglans regia. A0A2I4H7D7
Juglans regia. A0A2I4EPT2
Juglans regia. A0A2I4DYI9
Juglans regia. A0A2I4DYZ0
Juglans regia. A0A2I4GFL8
Juglans regia. A0A2I4E0L5
Juglans regia. A0A2I4GBN2
Juglans regia. A0A2I4FA92
Juglans regia. A0A2I4DU39
Juglans regia. A0A2I4E0L6
Juglans regia. A0A2I4FU89
Juglans regia. A0A2I4GBS2
Juglans regia. A0A2I4HLF8
Juglans regia. A0A2I4DYZ5
Juglans regia. A0A2I4EGC9
Juglans regia. A0A2I4DU26
Juglans regia. A0A2I4DGG7
Juglans regia. A0A2I4GV20
Juglans regia. A0A2I4E0K7
Juglans regia. A0A2I4H7H3
Juglans regia. A0A2I4HHT8
Juglans regia. A0A2I4DYJ2
Juglans regia. A0A2I4FLN8
Juglans regia. A0A2I4H7E7
Juglans regia. A0A2I4GV14
Juglans regia. A0A2I4GT05
Juglans regia. A0A2I4DGF8
Juglans regia. A0A2I4FFQ2
Juglans regia. A0A2I4DKP2
Juglans regia. A0A2I4GV12

Juglans regia. A0A2I4FFG4
Juglans regia. A0A2I4FIE0
Juglans regia. A0A2I4GGL6
Juglans regia. A0A2I4FZE7
Juglans regia. A0A2I4ERD8
Juglans regia. A0A2I4GT07
Juglans regia. A0A2I4DKP0
Juglans regia. A0A2I4FJE4
Juglans regia. A0A2I4HTL0
Juglans regia. A0A2I4HHJ4
Juglans regia. A0A2I4GV23
Juglans regia. A0A2I4EKK6
Juglans regia. A0A2I4HNL3
Juglans regia. A0A2I4GA68
Juglans regia. A0A2I4HEF2
Juglans regia. A0A2I4HLF5
Juglans regia. A0A2I4H4Y0
Juglans regia. A0A2I4GAY6
Juglans regia. A0A2I4ERB9
Juglans regia. A0A2I4FU46
Juglans regia. A0A2I4HTX2
Juglans regia. A0A2I4DQG1
Juglans regia. A0A2I4ERD4
Juglans regia. A0A2I4H6J2
Juglans regia. A0A2I4HP53
Juglans regia. A0A2I4GAU2
Juglans regia. A0A2I4EGB5
Juglans regia. A0A2I4DEN2
Juglans regia. A0A2I4ERC8
Juglans regia. A0A2I4FIB5
Juglans regia. A0A2I4FLY6
Juglans regia. A0A2I4G9T9
Juglans regia. A0A2I4ERD7
Juglans regia. A0A2I4H7C7
Juglans regia. A0A2I4FFP8
Juglans regia. A0A2I4HHI9
Juglans regia. A0A2I4FIE8
Juglans regia. A0A2I4H4L2
Juglans regia. A0A2I4GG21
Juglans regia. A0A2I4G912
Juglans regia. A0A2I4GBS9

Juglans regia. A0A2I4HPJ3
Juglans regia. A0A2I4GAA0
Juglans regia. A0A2I4EKJ5
Juglans regia. A0A2I4HSV3
Juglans regia. A0A2I4HTK6
Juglans regia. A0A2I4FIL5
Juglans regia. A0A2I4EK97
Juglans regia. A0A2I4FFQ0
Juglans regia. A0A2I4GV28
Juglans regia. A0A2I4HIN1
Juglans regia. A0A2I4EMX6
Juglans regia. A0A2I4HV91
Juglans regia. A0A2I4ERD1
Juglans regia. A0A2I4DU21
Juglans regia. A0A2I4EKU1
Juglans regia. A0A2I4E0L9
Juglans regia. A0A2I4GBS6
Juglans regia. A0A2I4HP69
Juglans regia. A0A2I4GWB0
Juglans regia. A0A2I4GAW7
Juglans regia. A0A2I4DET7
Juglans regia. A0A2I4G4J0
Juglans regia. A0A2I4GV30
Juglans regia. A0A2I4HV71
Juglans regia. A0A2I4FKE4
Juglans regia. A0A2I4E7G7
Juglans regia. A0A2I4GBT4
Juglans regia. A0A2I4DGG1
Juglans regia. A0A2I4DU29
Juglans regia. A0A2I4FFP3
Juglans regia. A0A2I4DN46
Juglans regia. A0A2I4HAL5
Juglans regia. A0A2I4FLV5
Juglans regia. A0A2I4FID4
Juglans regia. A0A2I4HV84
Juglans regia. A0A2I4GBV7
Juglans regia. A0A2I4E8G1
Juglans regia. A0A2I4E8H5
Juglans regia. A0A2I4E6A6
Juglans regia. A0A2I4HNL4
Juglans regia. A0A2I4HEG9

| | |
|---|---|
| Juglans regia. | A0A2I4E3B8 |
| Juglans regia. | A0A2I4EWD4 |
| Juglans regia. | A0A2I4GBR4 |
| Juglans regia. | A0A2I4EG75 |
| Juglans regia. | A0A2I4GAA7 |
| Juglans regia. | A0A2I4EG66 |
| Juglans regia. | A0A2I4HG78 |
| Juglans regia. | A0A2I4HSV9 |
| Juglans regia. | A0A2I4DTA7 |
| Juglans regia. | A0A2I4HNR2 |
| Juglans regia. | A0A2I4H926 |
| Juglans regia. | A0A2I4EG64 |
| Juglans regia. | A0A2I4DGG9 |
| Juglans regia. | A0A2I4DEM3 |
| Juglans regia. | A0A2I4EN07 |
| Juglans regia. | A0A2I4FU87 |
| Juglans regia. | A0A2I4FIJ8 |
| Juglans regia. | A0A2I4GBT2 |
| Juglans regia. | A0A2I4EGC4 |
| Kaistia algaru | A0A2S5U2E7 |
| Kaistia soli DS | A0A1M4XZE6 |
| Kaistia sp. SC | A0A1E4CVK7 |
| Kaistia sp. SC | A0A1E4C319 |
| Klebsormidiu | A0A1Y1I6D8 |
| Klebsormidiu | A0A1Y1IA49 |
| Klosneuvirus | A0A1V0SHW7 |
| Kocuria rhizo | A0A2W5CL52 |
| Kryptolebias | A0A3Q2Z9K7 |
| Kryptolebias | A0A3Q3B1Z9 |
| Kryptolebias | A0A3Q3B902 |
| Kryptolebias | A0A3Q3F291 |
| Kryptolebias | A0A3Q3BH26 |
| Kryptolebias | A0A3Q3B198 |
| Kwoniella ma | A0A1B9IXV1 |
| Labrenzia agg | A0NVX3 |
| Labrenzia agg | A0NMA0 |
| Labrenzia agg | A0A0M6Y1A7 |
| Labrenzia agg | A0A0M6Y878 |
| Labrenzia alb | A0A0M7AJT2 |
| Labrenzia alb | A0A0M6YXQ5 |
| Labrenzia ale | A0A0M6ZWY2 |

| | |
|---|---|
| Labrenzia ma | A0A2S3UK78 |
| Labrenzia sp. | A0A2T7H985 |
| Labrenzia sp. | U7G8P7 |
| Labrenzia sp. | U7GAT6 |
| Labrenzia sp. | A0A163UN31 |
| Labrenzia sp. | A0A163WYR8 |
| Labrenzia sp. | A0A222F1C4 |
| Labrenzia sp. | A0A222FCV3 |
| Labrenzia sua | A0A1M7CSY2 |
| Labrus bergyl | A0A3Q3N8L8 |
| Labrus bergyl | A0A3Q3GBJ6 |
| Labrus bergyl | A0A3Q3FDA4 |
| Labrus bergyl | A0A3Q3GEZ8 |
| Labrus bergyl | A0A3Q3GFE0 |
| Labrus bergyl | A0A3Q3N8X5 |
| Labrus bergyl | A0A3Q3GFJ4 |
| Labrus bergyl | A0A3Q3N8N2 |
| Labrus bergyl | A0A3Q3GFZ5 |
| Labrus bergyl | A0A3Q3GB54 |
| Labrus bergyl | A0A3Q3GFB9 |
| Lactuca sativa | A0A2J6K9H8 |
| Lactuca sativa | A0A2J6M9S4 |
| Lactuca sativa | A0A2J6LQZ4 |
| Lactuca sativa | A0A2J6JXN1 |
| Lactuca sativa | A0A2J6KN62 |
| Lactuca sativa | A0A2J6KKT3 |
| Lactuca sativa | A0A2J6KC77 |
| Lactuca sativa | A0A2J6MH90 |
| Lactuca sativa | A0A2J6LXZ5 |
| Lactuca sativa | A0A2J6JZ56 |
| Lactuca sativa | A0A2J6LVI0 |
| Lactuca sativa | A0A2J6K3Q9 |
| Lactuca sativa | A0A2J6L847 |
| Lactuca sativa | A0A2J6L7A5 |
| Lactuca sativa | A0A2J6K925 |
| Lactuca sativa | A0A2J6K8G7 |
| Lactuca sativa | A0A2J6MDS4 |
| Lactuca sativa | A0A2J6M338 |
| Lactuca sativa | A0A2J6LCC7 |
| Lactuca sativa | A0A2J6MGQ2 |
| Lactuca sativa | A0A2J6M6H7 |

| | |
|---|---|
| Lactuca sativa | A0A2J6KQ34 |
| Lactuca sativa | A0A2J6MB10 |
| Lactuca sativa | A0A2J6KSB6 |
| Lactuca sativa | A0A2J6M2X5 |
| Lactuca sativa | A0A2J6KQ62 |
| Lactuca sativa | A0A2J6KD61 |
| Lactuca sativa | A0A2J6KDN2 |
| Lactuca sativa | A0A2J6K1B8 |
| Lactuca sativa | A0A2J6K3R7 |
| Lactuca sativa | A0A2J6MI63 |
| Lactuca sativa | A0A2J6MGT2 |
| Lactuca sativa | A0A2J6JXJ5 |
| Lactuca sativa | A0A2J6M5A7 |
| Lactuca sativa | A0A2J6KQC0 |
| Lactuca sativa | A0A2J6JL93 |
| Lactuca sativa | A0A2J6KKU2 |
| Lactuca sativa | A0A2J6JT61 |
| Lactuca sativa | A0A2J6JYI5 |
| Lactuca sativa | A0A2J6KK13 |
| Lactuca sativa | A0A2J6M2T7 |
| Lactuca sativa | A0A2J6KQK0 |
| Lactuca sativa | A0A2J6KS85 |
| Lactuca sativa | A0A2J6LUT0 |
| Lactuca sativa | A0A2J6KKG9 |
| Lactuca sativa | A0A2J6JXL9 |
| Lactuca sativa | A0A2J6LRN0 |
| Lactuca sativa | A0A2J6M2U7 |
| Lactuca sativa | A0A2J6K992 |
| Lactuca sativa | A0A2J6JM53 |
| Lactuca sativa | A0A2J6KI92 |
| Lactuca sativa | A0A2J6MGS8 |
| Lactuca sativa | A0A2J6MH59 |
| Lactuca sativa | A0A2J6JU53 |
| Lactuca sativa | A0A2J6KQ13 |
| Lactuca sativa | A0A2J6MH39 |
| Lactuca sativa | A0A2J6KDQ7 |
| Lactuca sativa | A0A2J6JXP1 |
| Lactuca sativa | A0A2J6JXM0 |
| Lactuca sativa | A0A2J6K937 |
| Lactuca sativa | A0A2J6JXM4 |
| Lactuca sativa | A0A2J6LRI2 |

| | |
|---|---|
| Lactuca sativa | A0A2J6K9T5 |
| Lactuca sativa | A0A2J6JU38 |
| Lactuca sativa | A0A2J6L6J2 |
| Lactuca sativa | A0A2J6LFL5 |
| Lactuca sativa | A0A2J6KI16 |
| Lactuca sativa | A0A2J6KQR3 |
| Lactuca sativa | A0A2J6KQ10 |
| Lactuca sativa | A0A2J6LRK6 |
| Lactuca sativa | A0A2J6M2U5 |
| Lactuca sativa | A0A2J6MFF9 |
| Lactuca sativa | A0A2J6MFK8 |
| Lactuca sativa | A0A2J6LRQ0 |
| Lactuca sativa | A0A2J6JHP7 |
| Lactuca sativa | A0A2J6L1L1 |
| Lactuca sativa | A0A2J6KKP5 |
| Lactuca sativa | A0A2J6LL57 |
| Lactuca sativa | A0A2J6K8G6 |
| Lactuca sativa | A0A2J6L780 |
| Lactuca sativa | A0A2J6K927 |
| Lactuca sativa | A0A2J6JX67 |
| Lactuca sativa | A0A2J6JXK7 |
| Lactuca sativa | A0A2J6JPF2 |
| Lactuca sativa | A0A2J6LV17 |
| Lactuca sativa | A0A2J6LRI5 |
| Lactuca sativa | A0A2J6M2Y3 |
| Lactuca sativa | A0A2J6MGY5 |
| Lactuca sativa | A0A2J6L1K0 |
| Lactuca sativa | A0A2J6LVM5 |
| Lactuca sativa | A0A2J6M6F0 |
| Lactuca sativa | A0A2J6LUM7 |
| Lactuca sativa | A0A2J6MGS4 |
| Lactuca sativa | A0A2J6KZL2 |
| Lactuca sativa | A0A2J6M316 |
| Lactuca sativa | A0A2J6JXN5 |
| Lactuca sativa | A0A2J6JXL3 |
| Lactuca sativa | A0A2J6KFI6 |
| Lactuca sativa | A0A2J6KDB6 |
| Lactuca sativa | A0A2J6LRT0 |
| Lactuca sativa | A0A2J6JM37 |
| Lactuca sativa | A0A2J6KQI6 |
| Lactuca sativa | A0A2J6L1H5 |

| | |
|---|---|
| Lactuca sativa | A0A2J6JXA5 |
| Lactuca sativa | A0A2J6JXK1 |
| Lactuca sativa | A0A2J6KJY7 |
| Lactuca sativa | A0A2J6JW82 |
| Lactuca sativa | A0A2J6KQH9 |
| Lactuca sativa | A0A2J6M327 |
| Lactuca sativa | A0A2J6M2S7 |
| Lactuca sativa | A0A2J6K9D7 |
| Lactuca sativa | A0A2J6K917 |
| Lactuca sativa | A0A2J6KQ59 |
| Lactuca sativa | A0A2J6KJX6 |
| Lactuca sativa | A0A2J6KI82 |
| Lactuca sativa | A0A2J6L1I8 |
| Lactuca sativa | A0A2J6L7M7 |
| Lactuca sativa | A0A2J6KJV7 |
| Lactuca sativa | A0A2J6MGS5 |
| Lactuca sativa | A0A2J6M8C4 |
| Lactuca sativa | A0A2J6M2U1 |
| Lactuca sativa | A0A2J6L6Y1 |
| Lactuca sativa | A0A2J6KQ32 |
| Lactuca sativa | A0A2J6L9N2 |
| Lagenidium g | W0G9U1 |
| Lagenidium g | W0G5H4 |
| Larimichthys | A0A0F8CNM7 |
| Larimichthys | A0A0F8ASU5 |
| Lasius niger.x | A0A0J7NTZ9 |
| Lasius niger.x | A0A0J7KV86 |
| Lasius niger.x | A0A0J7KKG3 |
| Lasius niger.x | A0A0J7NL95 |
| Lasius niger.x | A0A0J7NBH3 |
| Lasius niger.x | A0A0J7KUZ9 |
| Lasius niger.x | A0A0J7KRY0 |
| Lasius niger.x | A0A0J7L2K3 |
| Latimeria cha | M3XJC9 |
| Latimeria cha | M3XGZ2 |
| Latimeria cha | H3ACN6 |
| Latimeria cha | H3AXI3 |
| Latimeria cha | M3XKU7 |
| Latrodectus h | E7D1R2 |
| Leavenworth | B0ZRK1 |
| Leavenworth | B0ZRN7 |

Leavenworth B0ZRK3
Leavenworth B0ZRK7
Leavenworth B0ZRK0
Leavenworth B0ZRK2
Leavenworth B0ZRJ7
Leavenworth B0ZRM2
Leavenworth B0ZRM5
Leavenworth B0ZRI5
Leavenworth B0ZRP4
Leavenworth B0ZRI9
Leavenworth B0ZRK4
Leavenworth B0ZRP3
Leavenworth B0ZRM6
Leavenworth I1V1S2
Leavenworth B0ZRL1
Leavenworth B0ZRM7
Leavenworth B0ZRL9
Leavenworth A0A2Z4HJD9
Leavenworth B0ZRI6
Leavenworth R9S7T9
Leavenworth B0ZRJ2
Leavenworth B0ZRM8
Leavenworth B0ZRI4
Leavenworth B0ZRK6
Leavenworth B0ZRJ6
Leavenworth B0ZRN9
Leavenworth B0ZRN2
Leavenworth B0ZRJ1
Leavenworth B0ZRM9
Leavenworth B0ZRN6
Leavenworth B0ZRK9
Leavenworth B0ZRM1
Leavenworth B0ZRJ3
Leavenworth B0ZRL7
Leavenworth B0ZRI2
Leavenworth R9S7P5
Leavenworth B0ZRI8
Leavenworth B0ZRL6
Leavenworth B0ZRL5
Leavenworth B0ZRN3
Leavenworth R9S7H9

| | |
|---|---|
| Leavenworth | B0ZRN5 |
| Leavenworth | B0ZRL0 |
| Leavenworth | R9S9P4 |
| Leavenworth | B0ZRJ4 |
| Leavenworth | B0ZRI3 |
| Leavenworth | B0ZRL8 |
| Leavenworth | B0ZRI7 |
| Leavenworth | B0ZRN4 |
| Leavenworth | B0ZRL4 |
| Leavenworth | B0ZRL3 |
| Leavenworth | B0ZRJ8 |
| Leavenworth | B0ZRM4 |
| Leavenworth | B0ZRK8 |
| Leavenworth | I3QHF6 |
| Leavenworth | B0ZRJ5 |
| Leavenworth | B0ZRN0 |
| Leavenworth | B0ZRK5 |
| Leavenworth | R9SA46 |
| Leavenworth | B0ZRM0 |
| Leavenworth | B0ZRP0 |
| Leavenworth | A0A2Z4HJM8 |
| Leavenworth | R9S9V7 |
| Leavenworth | B0ZRJ9 |
| Leavenworth | B0ZRN1 |
| Leersia perrie | A0A0D9W6W8 |
| Leersia perrie | A0A0D9XWL1 |
| Leersia perrie | A0A0D9W161 |
| Leersia perrie | A0A0D9WRU9 |
| Leersia perrie | A0A0D9V8W7 |
| Leersia perrie | A0A0D9XNS0 |
| Leersia perrie | A0A0D9WA06 |
| Leersia perrie | A0A0D9V3Y6 |
| Leersia perrie | A0A0D9W9Z1 |
| Leersia perrie | A0A0D9V8T3 |
| Leersia perrie | A0A0D9V8U6 |
| Leersia perrie | A0A0D9V8Q3 |
| Leersia perrie | A0A0D9XGF3 |
| Leersia perrie | A0A0D9WXV1 |
| Leersia perrie | A0A0D9W4W1 |
| Leersia perrie | A0A0D9W9Y8 |
| Leersia perrie | A0A0D9X052 |

| | |
|---|---|
| Leersia perrie | A0A0D9XGI1 |
| Leersia perrie | A0A0D9WD80 |
| Leersia perrie | A0A0D9X057 |
| Leersia perrie | A0A0D9W9Z0 |
| Leersia perrie | A0A0D9WTD1 |
| Leersia perrie | A0A0D9VVN7 |
| Leersia perrie | A0A0D9V8T2 |
| Leersia perrie | A0A0D9XB99 |
| Leersia perrie | A0A0D9W2T7 |
| Leersia perrie | A0A0D9X065 |
| Leersia perrie | A0A0D9W9Z8 |
| Leersia perrie | A0A0D9UY17 |
| Leersia perrie | A0A0D9XZW4 |
| Leersia perrie | A0A0D9XGF7 |
| Leersia perrie | A0A0D9XIR5 |
| Leersia perrie | A0A0D9WA00 |
| Leersia perrie | A0A0D9W4V1 |
| Leersia perrie | A0A0D9WR41 |
| Leersia perrie | A0A0D9XGF4 |
| Leersia perrie | A0A0D9W9Y9 |
| Leersia perrie | A0A0D9V3F7 |
| Leersia perrie | A0A0D9WA08 |
| Leersia perrie | A0A0D9VVD3 |
| Leersia perrie | A0A0D9WRG7 |
| Leersia perrie | A0A0D9X056 |
| Leersia perrie | A0A0D9WA07 |
| Leersia perrie | A0A0D9WA13 |
| Leersia perrie | A0A0D9V6J6 |
| Leersia perrie | A0A0D9WIB2 |
| Leersia perrie | A0A0D9UWX5 |
| Leersia perrie | A0A0D9XGF8 |
| Leersia perrie | A0A0D9WGJ8 |
| Leersia perrie | A0A0D9XWK7 |
| Leersia perrie | A0A0D9V3Y5 |
| Leersia perrie | A0A0D9XWK5 |
| Leersia perrie | A0A0D9W4V4 |
| Leersia perrie | A0A0D9XGI2 |
| Leersia perrie | A0A0D9V3F8 |
| Leersia perrie | A0A0D9XZV4 |
| Leersia perrie | A0A0D9W1R5 |
| Leersia perrie | A0A0D9V6K0 |

| | |
|---|---|
| Leersia perrie | A0A0D9W4V9 |
| Leersia perrie | A0A0D9V6J7 |
| Leersia perrie | A0A0D9UYF7 |
| Leersia perrie | A0A0D9VRC5 |
| Leersia perrie | A0A0D9XGF5 |
| Leersia perrie | A0A0D9W4V2 |
| Leersia perrie | A0A0D9XNR9 |
| Leersia perrie | A0A0D9V6J2 |
| Leersia perrie | A0A0D9V3Y7 |
| Leersia perrie | A0A0D9W4V3 |
| Leersia perrie | A0A0D9W3A9 |
| Leersia perrie | A0A0D9WA14 |
| Leersia perrie | A0A0D9W4V7 |
| Leersia perrie | A0A0D9V7B0 |
| Leersia perrie | A0A0D9V3Y8 |
| Leersia perrie | A0A0D9V1T5 |
| Leersia perrie | A0A0D9WQ05 |
| Leersia perrie | A0A0D9VAI1 |
| Leersia perrie | A0A0D9UYF9 |
| Leersia perrie | A0A0D9V6J9 |
| Leersia perrie | A0A0D9W3B3 |
| Leersia perrie | A0A0D9W9Z3 |
| Leersia perrie | A0A0D9WD66 |
| Leersia perrie | A0A0D9W2T8 |
| Leersia perrie | A0A0D9V8T6 |
| Leersia perrie | A0A0D9WRU8 |
| Leersia perrie | A0A0D9UYF8 |
| Leersia perrie | A0A0D9WA03 |
| Leersia perrie | A0A0D9WAH9 |
| Leersia perrie | A0A0D9XZV3 |
| Leersia perrie | A0A0D9XQ93 |
| Leersia perrie | A0A0D9WA05 |
| Leersia perrie | A0A0D9WR42 |
| Leersia perrie | A0A0D9X053 |
| Leersia perrie | A0A0D9W9Z9 |
| Leersia perrie | A0A0D9W9Z6 |
| Leersia perrie | A0A0D9WGJ7 |
| Leersia perrie | A0A0D9XH63 |
| Leersia perrie | A0A0D9V6J3 |
| Leersia perrie | A0A0D9W9Y7 |
| Leersia perrie | A0A0D9X503 |

| | |
|---|---|
| Leersia perrie | A0A0D9W3B1 |
| Leersia perrie | A0A0D9W4V6 |
| Leersia perrie | A0A0D9W4V5 |
| Leishmania b | A0A3P3YXB7 |
| Leishmania b | A0A3P3YXB3 |
| Leishmania b | A4H3V1 |
| Leishmania d | A0A3S5H596 |
| Leishmania g | A0A1E1INT0 |
| Leishmania ir | A0A381M993 |
| Leishmania m | E9AK14 |
| Leishmania p | A0A088RHV5 |
| Leisingera aq | A0A0P1H8K4 |
| Leisingera me | V9VZT4 |
| Lentibacter a | A0A1H3GY41 |
| Lepeophtheir | A0A0K2TCD8 |
| Lepeophtheir | A0A0K2V009 |
| Lepeophtheir | A0A0K2T1Y7 |
| Lepeophtheir | A0A0K2TPD6 |
| Lepeophtheir | A0A0K2UTX9 |
| Lepeophtheir | A0A0K2TTX1 |
| Lepeophtheir | A0A0K2U3D2 |
| Lepeophtheir | A0A0K2T870 |
| Lepeophtheir | A0A0K2U346 |
| Lepeophtheir | A0A0K2T1X1 |
| Lepeophtheir | A0A0K2SXW0 |
| Lepeophtheir | A0A0K2UNM9 |
| Lepeophtheir | A0A0K2TDI0 |
| Lepeophtheir | A0A0K2TFU3 |
| Lepeophtheir | A0A0K2TBQ4 |
| Lepeophtheir | A0A0K2UUX8 |
| Lepeophtheir | A0A0K2SZ08 |
| Lepeophtheir | A0A0K2U2F7 |
| Lepeophtheir | A0A0K2UVK3 |
| Lepeophtheir | A0A0K2TZM7 |
| Lepeophtheir | A0A0K2UP87 |
| Lepeophtheir | A0A0K2USH8 |
| Lepeophtheir | A0A0K2U0I6 |
| Lepeophtheir | A0A0K2UUV6 |
| Lepeophtheir | A0A0K2UUZ9 |
| Lepeophtheir | A0A0K2UTJ0 |
| Lepeophtheir | A0A0K2TZZ8 |

| | |
|---|---|
| Lepeophtheir | A0A0K2UVW1 |
| Lepeophtheir | A0A0K2UU23 |
| Lepeophtheir | A0A0K2UVG2 |
| Lepeophtheir | A0A0K2UWI6 |
| Lepisosteus c | W5NER0 |
| Lepisosteus c | W5M199 |
| Lepisosteus c | W5NEQ8 |
| Lepisosteus c | W5NFF8 |
| Lepisosteus c | W5NAB0 |
| Leptolyngbya | A0A3M9ZBN0 |
| Leptonychote | A0A2U3YDU7 |
| Leptonychote | A0A2U3X6Q7 |
| Leptonychote | A0A2U3YLJ6 |
| Leptonychote | A0A2U3YE66 |
| Leptonychote | A0A2U3YDS5 |
| Leptonychote | A0A2U3YDS8 |
| Leptonychote | A0A2U3XSF8 |
| Leptonychote | A0A2U3XSB8 |
| Leptonychote | A0A2U3X6P8 |
| Leptosomus ( | A0A091PSC7 |
| Leptosomus ( | A0A091PQK1 |
| Leptosomus ( | A0A091Q2P9 |
| Leptosphaeri | E4ZGL2 |
| Leptosphaeri | E5AD62 |
| Leptosphaeri | E5A991 |
| Leptotrombic | A0A443SF82 |
| Leptotrombic | A0A443RUF8 |
| Leptotrombic | A0A443SUS0 |
| Leptotrombic | A0A443SE45 |
| Leptotrombic | A0A443RVB4 |
| Leptotrombic | A0A443SM33 |
| Leptotrombic | A0A443SQ21 |
| Leptotrombic | A0A443SF48 |
| Leptotrombic | A0A443SHP2 |
| Leptotrombic | A0A443SQI1 |
| Leptotrombic | A0A443S6A4 |
| Leptotrombic | A0A443SH42 |
| Leptotrombic | A0A443SI69 |
| Leptotrombic | A0A443SN48 |
| Lethenteron ( | A0A2Z5XDQ4 |
| Lethenteron ( | A0A2Z5XDQ7 |

| | |
|---|---|
| Leucosporidiu | A0A1Y2FYL8 |
| Limosa lappo | A0A2I0UCB4 |
| Limosa lappo | A0A2I0U654 |
| Limosa lappo | A0A2I0THH2 |
| Limosa lappo | A0A2I0U125 |
| Lingula ungui | A0A1S3KCR3 |
| Lingula ungui | A0A1S3I3B4 |
| Lingula ungui | A0A2R2MTN1 |
| Lingula ungui | A0A1S3JBH9 |
| Lingula ungui | A0A1S3HNX8 |
| Lingula ungui | A0A1S3HFI9 |
| Lingula ungui | A0A1S3I1V9 |
| Lingula ungui | A0A1S3JR18 |
| Lingula ungui | A0A1S3IEH2 |
| Lingula ungui | A0A1S3J1T8 |
| Lingula ungui | A0A1S3H9T3 |
| Lingula ungui | A0A1S3HBY0 |
| Lingula ungui | A0A1S3K773 |
| Lingula ungui | A0A2R2MTZ4 |
| Lingula ungui | A0A1S3INU8 |
| Lingula ungui | A0A1S3IE57 |
| Lingula ungui | A0A1S3JU66 |
| Lingula ungui | A0A1S3I9N3 |
| Lingula ungui | A0A1S3JZ08 |
| Lingula ungui | A0A1S3K775 |
| Lingula ungui | A0A1S3JZ64 |
| Lingula ungui | A0A1S3JQD4 |
| Lingula ungui | A0A1S3J5V4 |
| Lingula ungui | A0A1S3GZQ5 |
| Lingula ungui | A0A1S3IAW8 |
| Lingula ungui | A0A1S3K770 |
| Lingula ungui | A0A1S3HKV5 |
| Lingula ungui | A0A1S3I1A6 |
| Lingula ungui | A0A1S3JFE9 |
| Lingula ungui | A0A1S3I1Y2 |
| Lingula ungui | A0A1S3IQU7 |
| Lingula ungui | A0A1S3KBV0 |
| Lingula ungui | A0A1S3KC90 |
| Lingula ungui | A0A1S3INP9 |
| Lingula ungui | A0A1S3KEU6 |
| Lingula ungui | A0A1S3H081 |

Lingula ungui A0A1S3K6K4
Lingula ungui A0A1S3K0C7
Lingula ungui A0A1S3I3C2
Lingula ungui A0A1S3KFD1
Lingula ungui A0A1S3KEE4
Lingula ungui A0A1S3KBC6
Lingula ungui A0A1S3H208
Lingula ungui A0A2R2MSR5
Lingula ungui A0A1S3K4Q7
Lingula ungui A0A1S3JEN3
Lingula ungui A0A1S3GZU4
Lingula ungui A0A1S3I1W2
Lingula ungui A0A1S3JBK0
Lingula ungui A0A1S3KHB8
Lingula ungui A0A1S3INY3
Lingula ungui A0A2R2MLU4
Lingula ungui A0A1S3H7Z1
Lingula ungui A0A1S3HI84
Lingula ungui A0A1S3HZL4
Lingula ungui A0A1S3KCC7
Lingula ungui A0A1S3ISS0
Lingula ungui A0A1S3IMR2
Lingula ungui A0A1S3HER8
Lingula ungui A0A2R2MJK4
Lingula ungui A0A1S3KEG7
Lingula ungui A0A1S3HEB9
Lingula ungui A0A1S3K901
Lingula ungui A0A1S3K0Q7
Lingula ungui A0A1S3K7Q8
Lingula ungui A0A1S3KA03
Lingula ungui A0A1S3JR32
Lingula ungui A0A1S3H2K1
Lingula ungui A0A1S3I4L8
Lingula ungui A0A1S3HD07
Lingula ungui A0A1S3KAP7
Lingula ungui A0A1S3J0J2
Lingula ungui A0A1S3J4U5
Lingula ungui A0A1S3JX74
Lingula ungui A0A1S3K1K5
Lingula ungui A0A1S3IC24
Lingula ungui A0A1S3JXG0

Lingula ungui A0A1S3K6B9
Lingula ungui A0A1S3JI63
Lingula ungui A0A2R2MRD9
Lingula ungui A0A1S3HR32
Lingula ungui A0A1S3JYB9
Lingula ungui A0A1S3KBX3
Lingula ungui A0A1S3JYD2
Lingula ungui A0A1S3HYX9
Lingula ungui A0A1S3KFK1
Lingula ungui A0A1S3I1T9
Lingula ungui A0A1S3ILB6
Lingula ungui A0A1S3K0U4
Lingula ungui A0A1S3JHR9
Lingula ungui A0A1S3JBP0
Lingula ungui A0A1S3JSJ3
Lingula ungui A0A1S3ICN0
Lingula ungui A0A1S3JR04
Lingula ungui A0A1S3HBM0
Lingula ungui A0A1S3J6C4
Lingula ungui A0A1S3ID77
Lingula ungui A0A1S3JZB8
Lingula ungui A0A1S3IZ38
Lingula ungui A0A1S3JW52
Lingula ungui A0A1S3J693
Lingula ungui A0A2R2MIB3
Lingula ungui A0A1S3I2T0
Lingula ungui A0A1S3K221
Lingula ungui A0A1S3KBT6
Lingula ungui A0A1S3HVT4
Lingula ungui A0A1S3IHR4
Lingula ungui A0A1S3KD35
Lingula ungui A0A1S3GZL9
Lingula ungui A0A2R2MJM7
Lingula ungui A0A1S3K756
Lingula ungui A0A2R2MRG8
Lingula ungui A0A1S3IIG5
Lingula ungui A0A1S3JYC8
Lingula ungui A0A1S3H9K2
Lingula ungui A0A1S3JQN5
Lingula ungui A0A1S3JZA1
Lingula ungui A0A2R2MNI0

| | |
|---|---|
| Lingula ungui | A0A1S3KB21 |
| Lipotes vexilli | A0A340XF42 |
| Lipotes vexilli | A0A340XZU6 |
| Lipotes vexilli | A0A340WGH7 |
| Lipotes vexilli | A0A340Y6H1 |
| Lipotes vexilli | A0A340Y5U1 |
| Lipotes vexilli | A0A340X8C1 |
| Lipotes vexilli | A0A340X7S9 |
| Lipotes vexilli | A0A340X8B6 |
| Lipotes vexilli | A0A340XEK1 |
| Lipotes vexilli | A0A340XC28 |
| Lipotes vexilli | A0A340XC32 |
| Litomosoides | A0A3P7K4H3 |
| Litomosoides | A0A3P6TYE8 |
| Litomosoides | A0A3P7M9S7 |
| Litomosoides | A0A3P7M8L9 |
| Litomosoides | A0A3P6TRE5 |
| Litomosoides | A0A3P6V4Q9 |
| Litomosoides | A0A3P6S6Z6 |
| Litomosoides | A0A3P6T8W6 |
| Litomosoides | A0A3P6USV6 |
| Litomosoides | A0A3P6UER0 |
| Litomosoides | A0A3P6U602 |
| Litomosoides | A0A3P6U3U8 |
| Litomosoides | A0A3P6UAP6 |
| Litomosoides | A0A3P6SQ54 |
| Litomosoides | A0A3P6SHG7 |
| Litomosoides | A0A3P6V945 |
| Litomosoides | A0A3P6RZY6 |
| Litoreibacter | A0A1H2WL46 |
| Litoreibacter | S9QDQ2 |
| Litoreibacter | A0A1M5BZJ4 |
| Litoreibacter | A0A1I6G0Y4 |
| Litorimicrobi | A0A1H9DXX5 |
| Litorimonas t | A0A420WEA0 |
| Litorimonas t | A0A420WFF1 |
| Litorimonas t | A0A420WKP0 |
| Littorina litto | A0A0A7RPZ1 |
| Littorina litto | A0A0A7RPW5 |
| Littorina litto | A0A0A7RPU1 |
| Loa loa.xlsx | A0A1I7VDG7 |

| | |
|---|---|
| Loa loa.xlsx | A0A1I7VW75 |
| Loa loa.xlsx | A0A1S0U4V4 |
| Loa loa.xlsx | A0A1I7VW66 |
| Loa loa.xlsx | A0A1I7W211 |
| Loa loa.xlsx | A0A1I7VX34 |
| Loa loa.xlsx | A0A1S0U2W5 |
| Loa loa.xlsx | A0A1S0U290 |
| Loa loa.xlsx | A0A1S0TQ84 |
| Loa loa.xlsx | A0A1S0U4B4 |
| Loa loa.xlsx | A0A1I7V595 |
| Loa loa.xlsx | A0A1S0TXG2 |
| Loa loa.xlsx | A0A1S0TGL3 |
| Loa loa.xlsx | A0A1I7VEP2 |
| Loa loa.xlsx | A0A1I7W1P7 |
| Loa loa.xlsx | A0A1I7VM68 |
| Loa loa.xlsx | A0A1S0U513 |
| Loa loa.xlsx | A0A1I7V7Z8 |
| Loa loa.xlsx | A0A1S0TYR1 |
| Loa loa.xlsx | A0A1I7VP24 |
| Loa loa.xlsx | A0A1I7W1K7 |
| Loa loa.xlsx | A0A1S0U521 |
| Loa loa.xlsx | A0A1I7VEI3 |
| Loa loa.xlsx | A0A1S0TM76 |
| Loa loa.xlsx | A0A1S0TVK0 |
| Loa loa.xlsx | A0A1I7V913 |
| Loa loa.xlsx | A0A1S0U7X7 |
| Loa loa.xlsx | A0A1S0TNT1 |
| Loa loa.xlsx | A0A1S0TXD9 |
| Loa loa.xlsx | A0A1S0U5P5 |
| Loa loa.xlsx | A0A1I7VER6 |
| Loa loa.xlsx | A0A1I7VJ23 |
| Loa loa.xlsx | A0A1S0TNL5 |
| Loa loa.xlsx | A0A1S0TP23 |
| Loktanella sp | A0A366W4W6 |
| Lomentospor | A0A2N3N2R9 |
| Lonchura stri | A0A218UQI0 |
| Lonchura stri | A0A218V7U2 |
| Longibacter s | A0A2A8CVC7 |
| Lophiomus se | B3XXB9 |
| Lottia gigante | V4ASX7 |
| Lottia gigante | V4CNX9 |

| | |
|---|---|
| Lottia gigante | V4AQ34 |
| Lottia gigante | V4A085 |
| Lottia gigante | V3ZGG1 |
| Lottia gigante | V3ZBJ5 |
| Lottia gigante | V4AP13 |
| Lottia gigante | V4A568 |
| Lottia gigante | V4B215 |
| Lottia gigante | V3ZQU5 |
| Lottia gigante | V3ZUA7 |
| Lottia gigante | V4AXR6 |
| Lottia gigante | V4AG04 |
| Lottia gigante | V4BSL1 |
| Lottia gigante | V4A7Y1 |
| Lottia gigante | V4BB69 |
| Lottia gigante | V4BDZ0 |
| Lottia gigante | V4A5J1 |
| Lottia gigante | V3ZJU6 |
| Lottia gigante | V4B1H4 |
| Lottia gigante | V4AGQ9 |
| Lottia gigante | V4BAA4 |
| Lottia gigante | V4BQJ5 |
| Lottia gigante | V4AWC8 |
| Lottia gigante | V4CGE2 |
| Lottia gigante | V4A396 |
| Lottia gigante | V3ZYP3 |
| Lottia gigante | V3ZKE9 |
| Lottia gigante | V4BQJ9 |
| Lottia gigante | V4B0J4 |
| Lottia gigante | V4CE61 |
| Lottia gigante | V3ZTQ2 |
| Lottia gigante | V4AZ96 |
| Lottia gigante | V4ACW6 |
| Lottia gigante | V3ZKU5 |
| Lottia gigante | V4ADH8 |
| Lottia gigante | V4BC73 |
| Lottia gigante | V4A6L5 |
| Lottia gigante | V3ZFI8 |
| Lottia gigante | V4ANB3 |
| Lottia gigante | V3ZGM4 |
| Lottia gigante | V4C4P7 |
| Lottia gigante | V4AFG6 |

| | |
|---|---|
| Lottia gigante | V4AAJ8 |
| Lottia gigante | V3ZJQ9 |
| Lottia gigante | V4B893 |
| Lottia gigante | V4A878 |
| Lottia gigante | V4AAG6 |
| Lottia gigante | V3ZVU7 |
| Lottia gigante | V4A5C0 |
| Lottia gigante | V4AF63 |
| Lottia gigante | V3ZP38 |
| Lottia gigante | V4ANZ3 |
| Lottia gigante | V4BCE9 |
| Lottia gigante | V4A5K9 |
| Lottia gigante | V4BEG6 |
| Lottia gigante | V4ADA5 |
| Lottia gigante | V4BFS5 |
| Lottia gigante | V4ABU2 |
| Lottia gigante | V4AGS3 |
| Lottia gigante | V4BI89 |
| Lottia gigante | V4A0F5 |
| Lottia gigante | V4BZ59 |
| Lottia gigante | V4C8K6 |
| Lottia gigante | V4ANZ0 |
| Lottia gigante | V4AAL1 |
| Lottia gigante | V4BV31 |
| Lottia gigante | V4AM58 |
| Lottia gigante | V4BSL7 |
| Lottia gigante | V4AJ20 |
| Lottia gigante | V3ZA19 |
| Lottia gigante | V4BF63 |
| Lottia gigante | V4AA83 |
| Lottia gigante | V3ZC53 |
| Lottia gigante | V4AZU1 |
| Lottia gigante | V4A6E5 |
| Lottia gigante | V4A9B0 |
| Lottia gigante | V4AZF9 |
| Lottia gigante | V3ZRI2 |
| Lottia gigante | V3ZE05 |
| Lottia gigante | V4BHI1 |
| Lottia gigante | V4CFZ0 |
| Lottia gigante | V4AAD3 |
| Lottia gigante | V4BND1 |

| | |
|---|---|
| Lottia gigante | V4A8U5 |
| Lottia gigante | V4AAP8 |
| Lottia gigante | V4A8U0 |
| Lottia gigante | V4AQF9 |
| Lottia gigante | V3Z3B9 |
| Lottia gigante | V3Z9D7 |
| Lottia gigante | V4BXB3 |
| Lottia gigante | V3ZKT4 |
| Lottia gigante | V4B8Q8 |
| Lottia gigante | V4A2F6 |
| Lottia gigante | V4BGT4 |
| Lottia gigante | V3ZJF4 |
| Lottia gigante | V4ATM4 |
| Lottia gigante | V4BAE3 |
| Lottia gigante | V4ARJ8 |
| Lottia gigante | V4BB58 |
| Lottia gigante | V4AZ26 |
| Lottia gigante | V4AJG9 |
| Lottia gigante | V4B9R7 |
| Lottia gigante | V3Z371 |
| Lottia gigante | V4AGU9 |
| Lottia gigante | V4ALG6 |
| Lottia gigante | V3Z819 |
| Lottia gigante | V4AXG5 |
| Lottia gigante | V3ZP70 |
| Lottia gigante | V4A0V8 |
| Lottia gigante | V4AZ89 |
| Lottia gigante | V4BDK1 |
| Lottia gigante | V3ZNC3 |
| Lottia gigante | V4A9Q6 |
| Lottia gigante | V3ZHP8 |
| Lottia gigante | V4AWY7 |
| Lottia gigante | V4C4C1 |
| Lottia gigante | V4B8W2 |
| Lottia gigante | V4B493 |
| Lottia gigante | V4BY76 |
| Lottia gigante | V3ZM12 |
| Lottia gigante | V4C7E0 |
| Lottia gigante | V4AUH2 |
| Lottia gigante | V4CL71 |
| Lottia gigante | V3ZP26 |

| | |
|---|---|
| Lottia gigante | V3ZV91 |
| Lotus japonic | Q70I28 |
| Lotus japonic | B0BLA2 |
| Lotus japonic | Q70I30 |
| Lotus japonic | Q8GSP1 |
| Lotus japonic | B0BLA4 |
| Lotus japonic | Q70I29 |
| Lotus japonic | Q8GSP2 |
| Lotus japonic | I3SEG1 |
| Lotus japonic | Q8GSP0 |
| Loxodonta af | G3TJ67 |
| Loxodonta af | G3SSI7 |
| Loxodonta af | G3U701 |
| Loxodonta af | G3TJ74 |
| Loxodonta af | G3U7P9 |
| Lucilia cuprin | A0A0L0CF27 |
| Lucilia cuprin | A0A0L0BZD3 |
| Lucilia cuprin | A0A0L0CF53 |
| Lucilia cuprin | A0A0L0CB36 |
| Lucilia cuprin | A0A0L0CRG8 |
| Lucilia cuprin | A0A0L0C513 |
| Lucilia cuprin | A0A0L0BPA5 |
| Lucilia cuprin | A0A0L0CB71 |
| Lucilia cuprin | A0A0L0BLS4 |
| Lunaria annu | Q8LP50 |
| Lupinus angu | A0A1J7H6R4 |
| Lupinus angu | A0A1J7GA64 |
| Lupinus angu | A0A1J7HQD3 |
| Lupinus angu | A0A394DIC4 |
| Lupinus angu | A0A1J7I619 |
| Lupinus angu | A0A1J7H3V7 |
| Lupinus angu | A0A1J7GFK8 |
| Lupinus angu | A0A1J7GZ77 |
| Lupinus angu | A0A1J7HH30 |
| Lupinus angu | A0A1J7HEQ0 |
| Lupinus angu | A0A1J7GDB9 |
| Lupinus angu | A0A1J7GNM1 |
| Lupinus angu | A0A1J7I6S6 |
| Lupinus angu | A0A1J7HD03 |
| Lupinus angu | A0A1J7FWA6 |
| Lutibaculum | V4RQY9 |

| | |
|---|---|
| Lutimaribacte | A0A1H0GW80 |
| Lutzomyia lor | A0A1B0CHS8 |
| Lutzomyia lor | A0A1B0CDT4 |
| Lutzomyia lor | A0A1B0CQS2 |
| Lutzomyia lor | A0A1B0CJ91 |
| Lutzomyia lor | A0A1B0CLB1 |
| Lygus hesper | A0A146LQ91 |
| Lygus hesper | A0A146M6Y4 |
| Lygus hesper | A0A146KQT6 |
| Lygus hesper | A0A0K8SEA4 |
| Lygus hesper | A0A0A9ZGD8 |
| Lygus hesper | A0A0A9XMZ2 |
| Lygus hesper | A0A0A9X4C5 |
| Lygus hesper | A0A0A9ZB53 |
| Lygus hesper | A0A0A9VYR8 |
| Lygus hesper | A0A0A9YIK1 |
| Lygus hesper | A0A0K8T345 |
| Lygus hesper | A0A0A9VV93 |
| Lygus hesper | A0A0A9X899 |
| Lygus hesper | A0A0A9XR44 |
| Lygus hesper | A0A0K8SWZ6 |
| Lygus hesper | A0A0K8SQ51 |
| Lygus hesper | A0A146L1N9 |
| Lygus hesper | A0A146LN03 |
| Lygus hesper | A0A0A9YAH9 |
| Lysobacter gu | A0A0S2G5W4 |
| Macaca fascic | A0A2K5VVK2 |
| Macaca fascic | G7PKF4 |
| Macaca fascic | A0A2K5WCZ4 |
| Macaca fascic | G7PKF5 |
| Macaca fascic | A0A2K5WGT6 |
| Macaca fascic | G7P478 |
| Macaca fascic | A0A2K5VTJ9 |
| Macaca fascic | C9E9X5 |
| Macaca fascic | A2V9Z1 |
| Macaca fascic | A0A2K5WGT9 |
| Macaca fascic | A0A2K5TJJ3 |
| Macaca fascic | A0A2K5WH02 |
| Macaca fascic | A0A2K5VTQ7 |
| Macaca fascic | C9E9X6 |
| Macaca fascic | G8F4N5 |

| | |
|---|---|
| Macaca fasci | A0A2K5TJJ6 |
| Macaca mula | F7A5G3 |
| Macaca mula | F6PJA8 |
| Macaca mula | F6PJA1 |
| Macaca mula | G7MSF8 |
| Macaca mula | F7CC91 |
| Macaca mula | A0A1D5RGB5 |
| Macaca mula | H9FEQ2 |
| Macaca mula | G7ML90 |
| Macaca mula | G7MSF7 |
| Macaca mula | F6PJ95 |
| Macaca mula | F7A5F6 |
| Macaca mula | A0A1D5Q7U7 |
| Macaca mula | G7MM01 |
| Macaca mula | P12545 |
| Macaca nem | A0A2K6DQD1 |
| Macaca nem | A0A2K6DDC3 |
| Macaca nem | A0A2K6DDA5 |
| Macaca nem | A0A2K6DD94 |
| Macaca nem | A0A2K6DCN9 |
| Macaca nem | A0A2K6DH51 |
| Macaca nem | A0A2K6E3X7 |
| Macaca nem | A0A2K6E3W9 |
| Macaca nem | A0A2K6DQE6 |
| Macleaya cor | A0A200QXH0 |
| Macleaya cor | A0A200RDT1 |
| Macleaya cor | A0A200Q4Y2 |
| Macleaya cor | A0A200QXM5 |
| Macleaya cor | A0A200Q8X0 |
| Macleaya cor | A0A200QRV5 |
| Macleaya cor | A0A200QQN4 |
| Macleaya cor | A0A200Q130 |
| Macleaya cor | A0A200QBX7 |
| Macleaya cor | A0A200R1B1 |
| Macleaya cor | A0A200QTR7 |
| Macleaya cor | A0A200QXK1 |
| Macleaya cor | A0A200QXI1 |
| Macleaya cor | A0A200QQN7 |
| Macleaya cor | A0A200Q8Y1 |
| Macleaya cor | A0A200Q8J1 |
| Macleaya cor | A0A200Q0W5 |

| | |
|---|---|
| Macleaya cor | A0A200QXH4 |
| Macleaya cor | A0A200QIT5 |
| Macleaya cor | A0A200Q0X7 |
| Macleaya cor | A0A200QXF5 |
| Macleaya cor | A0A200QRS4 |
| Macleaya cor | A0A200Q8V5 |
| Macleaya cor | A0A200Q2G3 |
| Macleaya cor | A0A200Q0W0 |
| Macleaya cor | A0A200QXP0 |
| Macrophomi | K2RH51 |
| Macrophomi | K2RS30 |
| Macrophomi | K2RCW8 |
| Macrophomi | K2RNM7 |
| Macrophomi | K2S8A1 |
| Macrophomi | K2RUQ8 |
| Macropus eu | O18783 |
| Macrostomu | A0A1I8GHF8 |
| Macrostomu | A0A1I8FVR7 |
| Macrostomu | A0A1I8H365 |
| Macrostomu | A0A1I8J167 |
| Macrostomu | A0A267EWK4 |
| Macrostomu | A0A1I8IBQ6 |
| Macrostomu | A0A1I8JEV8 |
| Macrostomu | A0A1I8FXH1 |
| Macrostomu | A0A1I8GNZ2 |
| Macrostomu | A0A267GQH1 |
| Macrostomu | A0A267GGW3 |
| Macrostomu | A0A1I8GL54 |
| Macrostomu | A0A1I8HG15 |
| Macrostomu | A0A1I8IV72 |
| Macrostomu | A0A267G2E7 |
| Macrostomu | A0A1I8G1P9 |
| Macrostomu | A0A267EA31 |
| Macrostomu | A0A267GUN8 |
| Macrostomu | A0A1I8IW00 |
| Macrostomu | A0A1I8IVY1 |
| Macrostomu | A0A1I8IRT6 |
| Macrostomu | A0A1I8H8D5 |
| Macrostomu | A0A1I8G8D6 |
| Macrostomu | A0A1I8HC33 |
| Macrostomu | A0A1I8IMI9 |

| | |
|---|---|
| Macrostomu | A0A267FXX8 |
| Macrostomu | A0A1I8J0U1 |
| Macrostomu | A0A1I8GHG0 |
| Macrostomu | A0A1I8HE61 |
| Macrostomu | A0A1I8J1U2 |
| Macrostomu | A0A1I8JJX3 |
| Macrostomu | A0A1I8GHR2 |
| Macrostomu | A0A1I8JKN1 |
| Macrostomu | A0A1I8FX03 |
| Macrostomu | A0A1I8J0K0 |
| Macrostomu | A0A1I8GPQ5 |
| Macrostomu | A0A267FEP6 |
| Macrostomu | A0A1I8GLQ8 |
| Macrostomu | A0A1I8J5S5 |
| Macrostomu | A0A1I8GNC3 |
| Macrostomu | A0A1I8GF89 |
| Macrostomu | A0A267DMG9 |
| Macrostomu | A0A267H2N9 |
| Macrostomu | A0A1I8JF80 |
| Macrostomu | A0A1I8GE13 |
| Macrostomu | A0A267DXV5 |
| Macrostomu | A0A1I8H2N3 |
| Macrostomu | A0A1I8FHG8 |
| Macrostomu | A0A1I8GCM3 |
| Macrostomu | A0A1I8FWJ3 |
| Macrostomu | A0A1I8GNW6 |
| Macrostomu | A0A1I8IZ00 |
| Macrostomu | A0A267F306 |
| Macrostomu | A0A267EXC8 |
| Macrostomu | A0A267GX64 |
| Macrostomu | A0A1I8GFR2 |
| Macrostomu | A0A1I8I316 |
| Macrostomu | A0A1I8ID15 |
| Macrostomu | A0A1I8G103 |
| Macrostomu | A0A267E9Q2 |
| Macrostomu | A0A267DNT5 |
| Macrostomu | A0A1I8G9H2 |
| Macrostomu | A0A1I8J5P8 |
| Macrostomu | A0A267E0M5 |
| Macrostomu | A0A267G2S7 |
| Macrostomu | A0A1I8FV01 |

| | |
|---|---|
| Macrostomu | A0A1I8HV10 |
| Macrostomu | A0A1I8J8F5 |
| Macrostomu | A0A1I8GJX9 |
| Macrostomu | A0A1I8H4R7 |
| Macrostomu | A0A1I8J9R2 |
| Macrostomu | A0A1I8JG79 |
| Macrostomu | A0A1I8GZN4 |
| Macrostomu | A0A1I8J542 |
| Macrostomu | A0A1I8JFP1 |
| Macrostomu | A0A1I8HVW8 |
| Macrostomu | A0A1I8J222 |
| Macrostomu | A0A1I8G6K2 |
| Macrostomu | A0A1I8GEP5 |
| Macrostomu | A0A1I8G513 |
| Macrostomu | A0A1I8I3M0 |
| Macrostomu | A0A1I8J5N9 |
| Macrostomu | A0A1I8FTM3 |
| Macrostomu | A0A1I8HQT8 |
| Macrostomu | A0A1I8GI38 |
| Macrostomu | A0A1I8GEN9 |
| Macrostomu | A0A1I8G9F7 |
| Macrostomu | A0A1I8IHH8 |
| Macrostomu | A0A267H9R8 |
| Macrostomu | A0A267GB32 |
| Macrostomu | A0A267GHE2 |
| Macrostomu | A0A267EFJ6 |
| Macrostomu | A0A1I8FYE4 |
| Macrostomu | A0A267F4N8 |
| Macrostomu | A0A1I8FY63 |
| Macrostomu | A0A1I8FMH1 |
| Macrostomu | A0A1I8GYH3 |
| Macrostomu | A0A1I8IU21 |
| Macrostomu | A0A1I8GI06 |
| Macrostomu | A0A1I8F694 |
| Macrostomu | A0A1I8GHZ2 |
| Macrostomu | A0A1I8GIV2 |
| Macrostomu | A0A1I8FT25 |
| Macrostomu | A0A267F4Y9 |
| Macrostomu | A0A1I8JK67 |
| Macrostomu | A0A1I8GKV5 |
| Macrostomu | A0A1I8GC08 |

| | |
|---|---|
| Macrostomu | A0A1I8GBL5 |
| Macrostomu | A0A1I8GKD7 |
| Macrostomu | A0A1I8HSV8 |
| Macrostomu | A0A267FJ10 |
| Macrostomu | A0A1I8J628 |
| Macrostomu | A0A1I8HKD5 |
| Macrostomu | A0A1I8IZJ9 |
| Macrostomu | A0A1I8HM08 |
| Macrostomu | A0A267GBV5 |
| Macrostomu | A0A1I8J8D6 |
| Macrostomu | A0A267GVU8 |
| Macrostomu | A0A267GP16 |
| Macrostomu | A0A1I8JFH6 |
| Macrostomu | A0A1I8J7E2 |
| Macrostomu | A0A1I8GC74 |
| Macrostomu | A0A1I8H460 |
| Macrostomu | A0A267E6R9 |
| Macrostomu | A0A1I8FRS9 |
| Macrostomu | A0A267G961 |
| Macrostomu | A0A267DRT5 |
| Macrostomu | A0A1I8HUT3 |
| Macrostomu | A0A1I8FU86 |
| Macrostomu | A0A267EFS8 |
| Macrostomu | A0A1I8HYI4 |
| Macrostomu | A0A1I8IQU1 |
| Macrostomu | A0A267ETV5 |
| Macrostomu | A0A267E615 |
| Macrostomu | A0A1I8FYK6 |
| Macrostomu | A0A1I8FWD9 |
| Macrostomu | A0A267ETH5 |
| Macrostomu | A0A1I8J0A4 |
| Macrostomu | A0A1I8FYP1 |
| Macrostomu | A0A1I8GUN9 |
| Macrostomu | A0A267FDY5 |
| Macrostomu | A0A1I8IHJ7 |
| Madurella my | A0A175WH97 |
| Madurella my | A0A175W7N1 |
| Madurella my | A0A175WB07 |
| Magnaporthe | G4MZI7 |
| Magnaporthe | Q2KGY0 |
| Magnaporthe | G4MXA4 |

| | |
|---|---|
| Magnaporthe | G4MZU5 |
| Magnaporthe | L7J520 |
| Magnaporthe | L7JDZ1 |
| Magnaporthe | L7JG00 |
| Magnaporthe | L7HS16 |
| Magnaporthe | L7I490 |
| Magnaporthi | A0A0C4EBJ2 |
| Magnetovibri | A0A2E6X7J5 |
| Magnetovibri | A0A2E6X7K3 |
| Maliponia aq | A0A238JQ45 |
| Mameliella al | A0A0B3RUU6 |
| Mameliella al | A0A0B3S7C3 |
| Mameliella sp | A0A2E9I414 |
| Mameliella sp | A0A2E9I2Q1 |
| Mameliella sp | A0A2E9I2P9 |
| Manacus vite | A0A093QGB8 |
| Manacus vite | A0A093Q5H9 |
| Mandrillus le | A0A2K5ZUK4 |
| Mandrillus le | A0A2K5Z4M0 |
| Mandrillus le | A0A2K6AE30 |
| Mandrillus le | A0A2K5Z4L5 |
| Mandrillus le | A0A2K5Z4N9 |
| Mandrillus le | A0A2K6ADW4 |
| Mandrillus le | A0A2K5YTU2 |
| Mandrillus le | A0A2K5Y8K1 |
| Mandrillus le | A0A2K5ZUK1 |
| Mandrillus le | A0A2K5Z4U2 |
| Manihot escu | A0A2C9UZL6 |
| Manihot escu | A0A2C9UQY7 |
| Manihot escu | A0A2C9U9R2 |
| Manihot escu | A0A2C9W6L4 |
| Manihot escu | A0A2C9WFX9 |
| Manihot escu | A0A2C9VN71 |
| Manihot escu | A0A2C9V805 |
| Manihot escu | A0A199UBF6 |
| Manihot escu | A0A2C9U5I8 |
| Manihot escu | A0A2C9U8X9 |
| Manihot escu | A0A2C9U847 |
| Manihot escu | A0A2C9V7G1 |
| Manihot escu | A0A2C9VAX0 |
| Manihot escu | A0A2C9VBF0 |

| | |
|---|---|
| Manihot escu | A0A2C9VNX9 |
| Manihot escu | A0A2C9VLQ8 |
| Manihot escu | A0A2C9U446 |
| Manihot escu | A0A2C9VAL0 |
| Manihot escu | A0A2C9U4D2 |
| Manihot escu | A0A2C9U801 |
| Manihot escu | A0A2C9U4I0 |
| Manihot escu | A0A2C9U805 |
| Manihot escu | A0A2C9U7Y6 |
| Manihot escu | A0A199UAQ8 |
| Manihot escu | A0A2C9UYI9 |
| Manihot escu | A0A199UBH6 |
| Manihot escu | A0A2C9U8Y9 |
| Manihot escu | A0A2C9UPX0 |
| Manihot escu | A0A2C9UFH3 |
| Manihot escu | A0A2C9U3X3 |
| Manihot escu | A0A2C9W8T4 |
| Manihot escu | A0A2C9U4C4 |
| Manihot escu | A0A2C9UF19 |
| Manihot escu | A0A2C9W6W4 |
| Manihot escu | A0A2C9U642 |
| Manihot escu | A0A2C9VNX3 |
| Manihot escu | A0A2C9UK30 |
| Manihot escu | A0A2C9VP96 |
| Manihot escu | A0A2C9W6N6 |
| Manihot escu | A0A199U9M8 |
| Manihot escu | A0A2C9UUL6 |
| Manihot escu | A0A2C9URP0 |
| Manihot escu | A0A2C9UZM3 |
| Manihot escu | A0A2C9WC49 |
| Manihot escu | A0A2C9UX56 |
| Manihot escu | A0A2C9U928 |
| Manihot escu | A0A2C9UMD6 |
| Manihot escu | A0A2C9UFX9 |
| Manihot escu | A0A2C9U9Q2 |
| Manihot escu | A0A2C9W8S6 |
| Manihot escu | A0A199UC06 |
| Manihot escu | A0A2C9UF31 |
| Manihot escu | A0A2C9U4L0 |
| Manihot escu | A0A2C9U5B5 |
| Manihot escu | A0A2C9U487 |

| | |
|---|---|
| Manihot escu | A0A2C9UDV6 |
| Manihot escu | A0A199UC16 |
| Manihot escu | A0A2C9VLR9 |
| Manihot escu | A0A2C9U0H2 |
| Manihot escu | A0A2C9VN68 |
| Manihot escu | A0A2C9UPX3 |
| Manihot escu | A0A2C9UPE4 |
| Manihot escu | A0A2C9UY52 |
| Manihot escu | A0A199UBJ1 |
| Manihot escu | A0A2C9W426 |
| Manihot escu | A0A2C9WEY1 |
| Manihot escu | A0A2C9W6P6 |
| Manihot escu | A0A2C9W2Y5 |
| Manihot escu | A0A2C9W6Q6 |
| Manihot escu | A0A2C9VLT1 |
| Manihot escu | A0A2C9U478 |
| Manihot escu | A0A2C9U869 |
| Manihot escu | A0A2C9UZ81 |
| Manihot escu | A0A2C9WFM5 |
| Manihot escu | A0A2C9W6P5 |
| Manihot escu | A0A2C9U8V8 |
| Manihot escu | A0A2C9U4K0 |
| Manihot escu | A0A2C9UFD3 |
| Manihot escu | A0A2C9UUW9 |
| Manihot escu | A0A2C9VNC9 |
| Manihot escu | A0A2C9VF19 |
| Manihot escu | A0A2C9UEZ1 |
| Manihot escu | A0A2C9UAW2 |
| Manihot escu | A0A2C9UYH0 |
| Manihot escu | A0A199UBF1 |
| Manihot escu | A0A2C9VN80 |
| Manihot escu | A0A2C9U9K7 |
| Manihot escu | A0A2C9VTQ5 |
| Manihot escu | A0A2C9U5A6 |
| Manihot escu | A0A2C9UPQ0 |
| Manihot escu | A0A2C9U8W8 |
| Manihot escu | A0A199UAZ0 |
| Manihot escu | A0A2C9VNE5 |
| Manihot escu | A0A2C9UPZ5 |
| Manihot escu | A0A2C9U839 |
| Manihot escu | A0A2C9UFW8 |

| | |
|---|---|
| Manihot escu | A0A2C9VB28 |
| Manihot escu | A0A2C9U816 |
| Manihot escu | A0A2C9U5B6 |
| Manihot escu | A0A2C9ULZ6 |
| Manihot escu | A0A2C9W6L3 |
| Manihot escu | A0A2C9VLU2 |
| Manihot escu | A0A199UBJ8 |
| Manihot escu | A0A2C9U935 |
| Manihot escu | A0A2C9VAT4 |
| Manihot escu | A0A2C9U7Q9 |
| Manihot escu | A0A2C9VLX7 |
| Manihot escu | A0A2C9W6M3 |
| Manihot escu | A0A2C9U632 |
| Manihot escu | A0A2C9VMC6 |
| Manihot escu | A0A2C9UEM3 |
| Manihot escu | A0A2C9UFH5 |
| Manihot escu | A0A2C9U0N6 |
| Manihot escu | A0A199UBH1 |
| Manihot escu | A0A199UC12 |
| Manihot escu | A0A2C9VLU8 |
| Manihot escu | A0A2C9VLS2 |
| Manihot escu | A0A2C9VP90 |
| Manihot escu | A0A2C9WEW1 |
| Manihot escu | A0A2C9U537 |
| Manihot escu | A0A2C9U4I6 |
| Manihot escu | A0A2C9VE98 |
| Manihot escu | A0A251IUP5 |
| Manihot escu | A0A199UBE6 |
| Manihot escu | A0A251JEQ5 |
| Marchantia p | A0A176VVC5 |
| Marchantia p | A0A176VSU9 |
| Marchantia p | A0A2R6WIU0 |
| Marchantia p | A0A2R6WIQ4 |
| Maricaulis ma | Q0AR81 |
| Maricaulis sa | A0A1G9Q3S7 |
| Maricaulis sp | A0A1Q8FK24 |
| Maricaulis sp | A0A2E2XTA6 |
| Maricaulis sp | A0A2D7Y7C9 |
| Marichromat | A0A161Q3D4 |
| Marichromat | W0E1A6 |
| Marichromat | A0A3M8NI56 |

| | |
|---|---|
| marine sedim | A0A0F9HEY4 |
| Marinibacteri | A0A225NEP4 |
| Marinicauda | A0A2U2BRH1 |
| Marinicauda | A0A2U2BSN0 |
| Marinovum a | A0A0H4LDC5 |
| Maritimibact | A3VM18 |
| Maritimibact | A0A1X7NU32 |
| Maritimibact | A0A2D9YV70 |
| Maritimibact | A0A2D8PDZ1 |
| Marssonina b | K1XNZ5 |
| Marssonina b | K1WYE2 |
| Marssonina b | K1XJ48 |
| Marssonina c | A0A218YS99 |
| Marssonina c | A0A218Z724 |
| Martelella m | A0A2T0JV19 |
| Mastacembe | A0A3Q3MRJ9 |
| Mastacembe | A0A3Q3LF07 |
| Mastacembe | A0A3Q3MRQ7 |
| Mastacembe | A0A3Q3MWE6 |
| Mastacembe | A0A3Q3S4P9 |
| Mastacembe | A0A3Q3LIA0 |
| Mastacembe | A0A3Q3MRT6 |
| Mastacembe | A0A3Q3MXL6 |
| Mastacembe | A0A3Q3SHD3 |
| Mastacembe | A0A3Q3RXI1 |
| Maylandia ze | A0A3P9CUC6 |
| Maylandia ze | A0A3P9CVH7 |
| Maylandia ze | A0A3P9CXJ8 |
| Maylandia ze | A0A3P9BY20 |
| Maylandia ze | A0A3P9BCB7 |
| Maylandia ze | A0A3P9CDB0 |
| Maylandia ze | A0A3P9CW19 |
| Medicago tru | G7ITJ8 |
| Medicago tru | G7IK65 |
| Medicago tru | A0A072TZ18 |
| Medicago tru | A0A072UMY1 |
| Medicago tru | A0A072TMX7 |
| Medicago tru | G7LFV0 |
| Medicago tru | G7IK59 |
| Medicago tru | G7LFU9 |
| Medicago tru | A0A072UY36 |

| | |
|---|---|
| Medicago tru | A0A072ULK1 |
| Medicago tru | A0A072UP57 |
| Medicago tru | G7LFV1 |
| Medicago tru | G7JF50 |
| Medicago tru | G7JSW6 |
| Medicago tru | G7IK51 |
| Medicago tru | G7JF44 |
| Medicago tru | G7LFU8 |
| Medicago tru | G7IK53 |
| Medicago tru | G7IVB0 |
| Medicago tru | A0A072VB47 |
| Medicago tru | G7IRL5 |
| Medicago tru | G7IK61 |
| Medicago tru | A0A072UA49 |
| Medicago tru | A0A072ULW3 |
| Medicago tru | A0A072V563 |
| Medicago tru | G7LFU6 |
| Medicago tru | A0A072UZ61 |
| Medicago tru | G7L503 |
| Medicago tru | G7K610 |
| Medicago tru | G7J0P4 |
| Medicago tru | A0A072V398 |
| Medicago tru | G7IK50 |
| Medicago tru | A0A072TZS6 |
| Medicago tru | A0A072UUL6 |
| Medicago tru | A0A072UTW4 |
| Medicago tru | G7IZ17 |
| Medicago tru | G7K613 |
| Medicago tru | A0A072UFT0 |
| Medicago tru | G7IK52 |
| Medicago tru | A0A072U488 |
| Medicago tru | A0A072UVG0 |
| Medicago tru | A0A072UN27 |
| Medicago tru | G7IVA0 |
| Medicago tru | A0A072TZ03 |
| Medicago tru | G7IZ18 |
| Medicago tru | G7IK54 |
| Medicago tru | G7J0Q4 |
| Medicago tru | A0A072VB61 |
| Medicago tru | A0A072VAM1 |
| Medicago tru | Q2HTU4 |

| | |
|---|---|
| Medicago tru | G7JF49 |
| Medicago tru | G7J0Q7 |
| Medicago tru | A0A072TPH7 |
| Medicago tru | G7L990 |
| Medicago tru | G7LDQ6 |
| Medicago tru | G7LA23 |
| Medicago tru | A0A072UA25 |
| Medicago tru | A0A072UUL9 |
| Medicago tru | A0A072TYW2 |
| Medicago tru | A0A072U0S7 |
| Medicago tru | G7IYY4 |
| Medicago tru | A0A072TZR6 |
| Medicago tru | G7L9B9 |
| Medicago tru | A0A072THU9 |
| Medicago tru | A0A072UA87 |
| Medicago tru | A0A072TZY0 |
| Medicago tru | G7J708 |
| Medicago tru | G7L199 |
| Medicago tru | A0A072TQM4 |
| Medicago tru | G7IYX7 |
| Medicago tru | A0A072TQ44 |
| Medicago tru | A0A072TZ15 |
| Medicago tru | G7LA28 |
| Medicago tru | G7IH64 |
| Medicago tru | A0A072UA68 |
| Medicago tru | G7IYX3 |
| Medicago tru | A0A072UNR6 |
| Medicago tru | G7KZ40 |
| Medicago tru | G7LA34 |
| Medicago tru | G7L986 |
| Medicago tru | A0A072UA82 |
| Medicago tru | G7IYY1 |
| Medicago tru | A0A072TZ68 |
| Medicago tru | A0A072UR75 |
| Medicago tru | A0A072THC7 |
| Medicago tru | A0A072TZY9 |
| Medicago tru | G7JZD9 |
| Medicago tru | A0A072V9X1 |
| Medicago tru | G7J0Q3 |
| Medicago tru | A0A072U1F5 |
| Medicago tru | A0A072TPK7 |

| | |
|---|---|
| Medicago tru | A0A072TYZ8 |
| Medicago tru | A0A072V4F9 |
| Medicago tru | A0A072TQ73 |
| Medicago tru | G7LBP9 |
| Medicago tru | A0A072TLA3 |
| Medicago tru | G7ITK1 |
| Medicago tru | A0A072V4F3 |
| Medicago tru | A0A072UN78 |
| Medicago tru | A0A072VL26 |
| Medicago tru | G7IYX6 |
| Medicago tru | A0A072U1H2 |
| Medicago tru | A0A072V3X0 |
| Medicago tru | A0A072U1E3 |
| Medicago tru | A0A072TS26 |
| Medicago tru | A0A072U0S1 |
| Medicago tru | G7KKY8 |
| Medicago tru | G7LA08 |
| Medicago tru | A0A072TZ74 |
| Medicago tru | A0A072U0N7 |
| Medicago tru | A0A072UP51 |
| Medicago tru | A0A072VBN5 |
| Medicago tru | A0A072TTD3 |
| Medicago tru | G7J0Q1 |
| Medicago tru | A0A072UP20 |
| Medicago tru | A0A072TZ09 |
| Medicago tru | A0A072TZ31 |
| Medicago tru | G7IRK8 |
| Medicago tru | A0A072UVG3 |
| Medicago tru | A0A072VAL5 |
| Medicago tru | G7IK58 |
| Medicago tru | A0A072TQ29 |
| Medicago tru | G7JXF4 |
| Medicago tru | G7IZ14 |
| Medicago tru | A0A072UTW1 |
| Medicago tru | A0A072TQF1 |
| Medicago tru | A0A072UTA1 |
| Medicago tru | G7LA16 |
| Medicago tru | G7L9C7 |
| Medicago tru | A0A072TQ50 |
| Medicago tru | G7JXG0 |
| Medicago tru | A0A072TZ10 |

| | |
|---|---|
| Medicago tru | A0A072UTV6 |
| Medicago tru | G7JSW4 |
| Medicago tru | G7JSV8 |
| Medicago tru | A0A072UKC3 |
| Medicago tru | A0A072UN82 |
| Medicago tru | A0A072V8H5 |
| Medicago tru | A0A396GBY0 |
| Medicago tru | A0A396IB06 |
| Medicago tru | A0A396I8P5 |
| Medicago tru | A0A396I8N0 |
| Medicago tru | A0A396II14 |
| Medicago tru | A0A396GIR1 |
| Medicago tru | A0A396IJL1 |
| Medicago tru | A0A396IAC3 |
| Medicago tru | A0A396GKB0 |
| Medicago tru | A0A396GKM1 |
| Medicago tru | A0A396GID7 |
| Medicago tru | A0A396GXQ7 |
| Medicago tru | A0A396IPQ6 |
| Medicago tru | A0A396H464 |
| Medicago tru | A0A396J1V4 |
| Medicago tru | A0A396GHQ0 |
| Medicago tru | A0A396IJS7 |
| Medicago tru | A0A396GXP2 |
| Medicago tru | A0A396IJV7 |
| Medicago tru | A0A396IFP3 |
| Medicago tru | A0A396GZH7 |
| Medicago tru | A0A396J1V8 |
| Medicago tru | A0A396GIN5 |
| Medicago tru | A0A396IKZ3 |
| Medicago tru | A0A396IEA5 |
| Medicago tru | A0A396GIL7 |
| Medicago tru | A0A396GJK4 |
| Medicago tru | A0A396H410 |
| Medicago tru | A0A396J0V6 |
| Medicago tru | A0A396GXZ2 |
| Medicago tru | A0A396H073 |
| Medicago tru | A0A396IH94 |
| Medicago tru | A0A396IJQ8 |
| Medicago tru | A0A396JG02 |
| Medicago tru | A0A396ICL0 |

| | |
|---|---|
| Medicago tru | A0A396H452 |
| Medicago tru | A0A396IB36 |
| Medicago tru | A0A396J5K8 |
| Medicago tru | A0A396H4I4 |
| Medicago tru | A0A396GKI2 |
| Medicago tru | A0A396GAD8 |
| Medicago tru | A0A396JBU6 |
| Medicago tru | A0A396I3J8 |
| Medicago tru | A0A396GIN1 |
| Medicago tru | A0A396IQ10 |
| Medicago tru | A0A396I8N1 |
| Medicago tru | A0A396GC28 |
| Medicago tru | A0A396GHT6 |
| Medicago tru | A0A396H0A3 |
| Medicago tru | A0A396IJM8 |
| Medicago tru | A0A396IDT3 |
| Medicago tru | A0A396H2F4 |
| Medicago tru | A0A396J9F7 |
| Medicago tru | A0A396J198 |
| Medicago tru | A0A396IGJ9 |
| Medicago tru | A0A396IMY4 |
| Medicago tru | A0A396HUI6 |
| Medicago tru | A0A396GXR8 |
| Medicago tru | A0A396GJX9 |
| Medicago tru | A0A396GMB1 |
| Medicago tru | A0A396IQZ7 |
| Medicago tru | A0A396JBT7 |
| Medicago tru | A0A072TFN9 |
| Medicago tru | A2Q3L3 |
| Medicago tru | G7JSW9 |
| Medicago tru | G7JF45 |
| Medicago tru | A0A072TZ95 |
| Medicago tru | G7L1A0 |
| Medicago tru | A0A072TZR2 |
| Medicago tru | A0A072TLV0 |
| Medicago tru | G7L2A0 |
| Medicago tru | A0A072UDX2 |
| Medicago tru | G7IXM3 |
| Medicago tru | G7L299 |
| Medicago tru | G7L994 |
| Medicago tru | G7IRM2 |

| | |
|---|---|
| Medicago tru | A0A072UTA5 |
| Medicago tru | G7JZE3 |
| Medicago tru | A0A072U7C9 |
| Medicago tru | G7IVB2 |
| Medicago tru | A0A072TQ06 |
| Medicago tru | A0A072UQ11 |
| Medicago tru | G7L9B8 |
| Medicago tru | A0A072VJ29 |
| Medicago tru | G7J0P5 |
| Medicago tru | A0A072ULW9 |
| Medicago tru | A0A072TR17 |
| Medicago tru | A0A072U021 |
| Medicago tru | G7J0P6 |
| Medicago tru | A0A072USP0 |
| Medicago tru | G7JSX3 |
| Medicago tru | A0A072UF01 |
| Medicago tru | G7L2A2 |
| Medicago tru | A0A072U1F1 |
| Medicago tru | G7IYX8 |
| Medicago tru | G7IK56 |
| Medicago tru | G7IZ19 |
| Medicago tru | G7IK62 |
| Medicago tru | G7ITK2 |
| Medicago tru | G7JSX1 |
| Medicago tru | G7JSW5 |
| Medicago tru | A0A072UM50 |
| Medicago tru | A0A072VE90 |
| Medicago tru | A0A072TGW7 |
| Medicago tru | A0A072VAH2 |
| Medicago tru | A0A072VGD0 |
| Medicago tru | A0A396HUX6 |
| Medicago tru | A0A396GF10 |
| Medicago tru | A0A396J9G3 |
| Medicago tru | A0A396GLE7 |
| Medicago tru | A0A396J1B2 |
| Medicago tru | A0A396GK01 |
| Medicago tru | A0A396II24 |
| Medicago tru | A0A396H0B4 |
| Medicago tru | A0A396IAC8 |
| Medicago tru | A0A396GKN1 |
| Medicago tru | A0A396H9W1 |

| | |
|---|---|
| Medicago tru | A0A396GKK2 |
| Medicago tru | A0A396ICL7 |
| Medicago tru | A0A396GQP4 |
| Medicago tru | A0A396GIS2 |
| Medicago tru | A0A396GZH2 |
| Medicago tru | A0A396GZ32 |
| Medicago tru | A0A396J4V5 |
| Medicago tru | A0A396IIB1 |
| Medicago tru | A0A396GY25 |
| Medicago tru | A0A396IPU4 |
| Megaselia sca | T1GV96 |
| Megaselia sca | T1GCN7 |
| Megavirus ch | G5CS05 |
| Megavirus ch | G5CSX9 |
| Megavirus ch | G5CSY0 |
| Megavirus co | K7YVH7 |
| Megavirus co | K7Y8W4 |
| Megavirus co | K7YGR5 |
| Megavirus co | H2EAU8 |
| Megavirus co | H2EAV0 |
| Megavirus co | H2EAE1 |
| Megavirus lba | L7Y445 |
| Megavirus lba | L7Y5I8 |
| Megavirus lba | L7XXX5 |
| Megavirus vit | A0A2P1EIL3 |
| Megavirus vit | A0A2P1EIK7 |
| Megavirus vit | A0A2P1EI82 |
| Melanaphis s | A0A2H8TNX4 |
| Melanaphis s | A0A2H8TQD0 |
| Meleagris gal | G1N219 |
| Meleagris gal | G3UPW8 |
| Meleagris gal | G3USW9 |
| Meleagris gal | G1NG08 |
| Meleagris gal | G1NIF9 |
| Meleagris gal | G1MVV6 |
| Meliniomyces | A0A2J6SGV1 |
| Meliniomyces | A0A2J6SNC2 |
| Meliniomyces | A0A2J6T0X6 |
| Meliniomyces | A0A2J6TAR8 |
| Melipona qua | A0A0M8ZTA1 |
| Melipona qua | A0A0N1IT07 |

| | |
|---|---|
| Melipona qua | A0A0M9AA61 |
| Melipona qua | A0A0N0BEK1 |
| Melipona qua | A0A0N0BFS3 |
| Melipona qua | A0A0N1IU30 |
| Melipona qua | A0A0N0BKD6 |
| Melittangium | A0A250ING7 |
| Meloidogyne | A0A1I8B4B8 |
| Meloidogyne | A0A1I8BK90 |
| Meloidogyne | A0A1I8AXB8 |
| Meloidogyne | A0A1I8AXZ1 |
| Meloidogyne | A0A1I8C0R8 |
| Meloidogyne | A0A1I8BY25 |
| Meloidogyne | A0A1I8BIP0 |
| Meloidogyne | A0A1I8B5R6 |
| Meloidogyne | A0A1I8BYM1 |
| Meloidogyne | A0A1I8BQ11 |
| Meloidogyne | A0A1I8B4D5 |
| Meloidogyne | A0A1I8BG52 |
| Meloidogyne | A0A1I8AY49 |
| Meloidogyne | A0A1I8BQG6 |
| Meloidogyne | A0A1I8BBQ9 |
| Meloidogyne | A0A1I8BCA3 |
| Merops nubi | A0A091QTC3 |
| Merops nubi | A0A091QC15 |
| Merops nubi | A0A091QSV6 |
| Merops nubi | A0A091QSR7 |
| Mesitornis ur | A0A091QUC8 |
| Mesitornis ur | A0A091RMA7 |
| Mesocestoid | A0A0R3U559 |
| Mesocricetus | A0A1U7Q8Z9 |
| Mesocricetus | A0A1U7R772 |
| Mesocricetus | A0A1U8C9R2 |
| Mesocricetus | A0A3Q0CLL1 |
| Mesocricetus | A0A1U7QLW9 |
| Mesocricetus | A0A3Q0CJX2 |
| Mesocricetus | A0A1U7R4K2 |
| Mesocricetus | A0A3Q0CDV3 |
| Mesocricetus | A0A1U8BGC2 |
| Mesocricetus | A0A1U8BFL9 |
| Mesocricetus | A0A1U8BRY1 |
| Mesorhizobiu | A0A1I3VIU9 |

| | |
|---|---|
| Mesorhizobiu | A0A1I4EVC7 |
| Mesorhizobiu | H0HTQ8 |
| Mesorhizobiu | A0A212DNS9 |
| Mesorhizobiu | G6YBH1 |
| Mesorhizobiu | A0A2W5AVW2 |
| Mesorhizobiu | A0A2W4ZIG5 |
| Mesorhizobiu | L0KNN6 |
| Mesorhizobiu | A0A1X7PA47 |
| Mesorhizobiu | A0A1X7PI50 |
| Mesorhizobiu | E8TE24 |
| Mesorhizobiu | A0A143NC21 |
| Mesorhizobiu | A0A2P9AXN5 |
| Mesorhizobiu | A0A2P7SR23 |
| Mesorhizobiu | A0A330I171 |
| Mesorhizobiu | A0A330H340 |
| Mesorhizobiu | A0A2T4IQ86 |
| Mesorhizobiu | A0A068DAZ6 |
| Mesorhizobiu | A0A068D8X3 |
| Mesorhizobiu | Q982A2 |
| Mesorhizobiu | Q982A0 |
| Mesorhizobiu | Q983Q7 |
| Mesorhizobiu | Q98K29 |
| Mesorhizobiu | Q982A3 |
| Mesorhizobiu | A0A3M9XHZ6 |
| Mesorhizobiu | A0A3A5KV12 |
| Mesorhizobiu | A0A2W7C7Z2 |
| Mesorhizobiu | A0A2W7CT51 |
| Mesorhizobiu | A0A271L788 |
| Mesorhizobiu | M5F7L2 |
| Mesorhizobiu | A0A1G9IZK6 |
| Mesorhizobiu | A0A1L3SUG4 |
| Mesorhizobiu | F7Y3J3 |
| Mesorhizobiu | F7YDM6 |
| Mesorhizobiu | A0A090DAC7 |
| Mesorhizobiu | A0A090GF48 |
| Mesorhizobiu | A0A090GJ86 |
| Mesorhizobiu | A0A090EAQ0 |
| Mesorhizobiu | A0A0K2VPY4 |
| Mesorhizobiu | A0A1R3VHN0 |
| Mesorhizobiu | A0A1R3VG59 |
| Mesorhizobiu | A0A1G5YRE1 |

| | |
|---|---|
| Mesorhizobiu | A0A2A6F891 |
| Mesorhizobiu | A0A2P7SMF1 |
| Mesorhizobiu | A0A1Q3LTY8 |
| Mesorhizobiu | A0A1Q4AJJ6 |
| Mesorhizobiu | A0A1A5T9T0 |
| Mesorhizobiu | A0A395KH22 |
| Mesorhizobiu | A0A3R9Y594 |
| Mesorhizobiu | A0A2N7SIF6 |
| Mesorhizobiu | A6N7X9 |
| Mesorhizobiu | A6N7X6 |
| Mesorhizobiu | A6N7X7 |
| Mesorhizobiu | A6N7X5 |
| Mesorhizobiu | A0A330GWG4 |
| Mesorhizobiu | A0A3A3HUH9 |
| Mesorhizobiu | X6KLP5 |
| Mesorhizobiu | X6K883 |
| Mesorhizobiu | X6KA10 |
| Mesorhizobiu | V7HDL0 |
| Mesorhizobiu | X6JHB4 |
| Mesorhizobiu | X6JBU3 |
| Mesorhizobiu | X6IKL6 |
| Mesorhizobiu | X6HND0 |
| Mesorhizobiu | V7H8P5 |
| Mesorhizobiu | X6FM84 |
| Mesorhizobiu | X6GLA2 |
| Mesorhizobiu | A0A371XGB0 |
| Mesorhizobiu | A0A1S1SGU3 |
| Mesorhizobiu | A0A1S1S606 |
| Mesorhizobiu | X6EAG6 |
| Mesorhizobiu | X6DBJ5 |
| Mesorhizobiu | X6DP76 |
| Mesorhizobiu | X6EWU4 |
| Mesorhizobiu | X6F6P8 |
| Mesorhizobiu | X6EVU3 |
| Mesorhizobiu | X6CWC0 |
| Mesorhizobiu | X6C4J0 |
| Mesorhizobiu | X6DDI2 |
| Mesorhizobiu | V7G7Z9 |
| Mesorhizobiu | X5Y7Z1 |
| Mesorhizobiu | A0A0E2NHH7 |
| Mesorhizobiu | V7FGS0 |

| | |
|---|---|
| Mesorhizobiu | X5V9Q4 |
| Mesorhizobiu | X5V974 |
| Mesorhizobiu | X5UDZ6 |
| Mesorhizobiu | X5RA25 |
| Mesorhizobiu | X5PRZ7 |
| Mesorhizobiu | X5QT36 |
| Mesorhizobiu | X5QMX6 |
| Mesorhizobiu | X5QIF0 |
| Mesorhizobiu | A0A436RRF8 |
| Mesorhizobiu | A0A3S2R3M1 |
| Mesorhizobiu | A0A437Z7D9 |
| Mesorhizobiu | A0A3S2YUX2 |
| Mesorhizobiu | A0A3S3C863 |
| Mesorhizobiu | A0A3S3YBG7 |
| Mesorhizobiu | A0A437YW26 |
| Mesorhizobiu | A0A3S3A539 |
| Mesorhizobiu | A0A437ZX63 |
| Mesorhizobiu | A0A437VVF2 |
| Mesorhizobiu | A0A3S2YVR6 |
| Mesorhizobiu | A0A437WTZ3 |
| Mesorhizobiu | A0A437VGM9 |
| Mesorhizobiu | A0A434THQ7 |
| Mesorhizobiu | A0A434SHC2 |
| Mesorhizobiu | A0A434NS16 |
| Mesorhizobiu | A0A434P1D2 |
| Mesorhizobiu | A0A434MM88 |
| Mesorhizobiu | A0A3S1ILE2 |
| Mesorhizobiu | A0A434PMQ0 |
| Mesorhizobiu | A0A434L3Y6 |
| Mesorhizobiu | A0A3Q8YPJ5 |
| Mesorhizobiu | A0A3S9DBX0 |
| Mesorhizobiu | A0A3S9DC85 |
| Mesorhizobiu | A0A3S9DBY6 |
| Mesorhizobiu | A0A3S9D2V1 |
| Mesorhizobiu | A0A3S2KY76 |
| Mesorhizobiu | A0A437WND3 |
| Mesorhizobiu | A0A3S9FK01 |
| Mesorhizobiu | A0A3Q9AI45 |
| Mesorhizobiu | A0A3S3LC86 |
| Mesorhizobiu | A0A443GJA2 |
| Mesorhizobiu | A0A3S2I1J4 |

| | |
|---|---|
| Mesorhizobiu | A0A436SS41 |
| Mesorhizobiu | A0A3Q8YI66 |
| Mesorhizobiu | A0A3S1KJ31 |
| Mesorhizobiu | A0A434SWL3 |
| Mesorhizobiu | A0A3S2KH58 |
| Mesorhizobiu | A0A3S1NX24 |
| Mesorhizobiu | A0A444KCU0 |
| Mesorhizobiu | A0A3S3R3Y8 |
| Mesorhizobiu | A0A444K8W2 |
| Mesorhizobiu | A0A435D6W3 |
| Mesorhizobiu | A0A3S1QJG7 |
| Mesorhizobiu | A0A3S9BTR5 |
| Mesorhizobiu | A0A436QSC2 |
| Mesorhizobiu | A0A436SLU0 |
| Mesorhizobiu | A0A436SBV8 |
| Mesorhizobiu | A0A3S9DJ20 |
| Mesorhizobiu | A0A3Q8YYA6 |
| Mesorhizobiu | A0A3S9DRS6 |
| Mesorhizobiu | A0A434VEP9 |
| Mesorhizobiu | A0A3S1E0H8 |
| Mesorhizobiu | A0A436Y6Z9 |
| Mesorhizobiu | A0A436VBS5 |
| Mesorhizobiu | A0A437ZLF5 |
| Mesorhizobiu | A0A436P9R0 |
| Mesorhizobiu | A0A436PHM8 |
| Mesorhizobiu | A0A3S2IWX9 |
| Mesorhizobiu | A0A3S2YQZ9 |
| Mesorhizobiu | A0A437X9T7 |
| Mesorhizobiu | A0A437X6J9 |
| Mesorhizobiu | A0A436WL48 |
| Mesorhizobiu | A0A436SR86 |
| Mesorhizobiu | A0A3S2IF88 |
| Mesorhizobiu | A0A3S2KLT7 |
| Mesorhizobiu | A0A436SRB7 |
| Mesorhizobiu | A0A436YYK0 |
| Mesorhizobiu | A0A435B6A1 |
| Mesorhizobiu | A0A3S2QL44 |
| Mesorhizobiu | A0A434RXG0 |
| Mesorhizobiu | A0A436UW61 |
| Mesorhizobiu | A0A436WQQ7 |
| Mesorhizobiu | A0A3S2LXT0 |

| | |
|---|---|
| Mesorhizobiu | A0A3S1MLP3 |
| Mesorhizobiu | A0A3Q8ZQ16 |
| Mesorhizobiu | A0A436R115 |
| Mesorhizobiu | A0A3S3QWG9 |
| Mesorhizobiu | A0A434MRM8 |
| Mesorhizobiu | A0A3S1HQV7 |
| Mesorhizobiu | A0A434MG50 |
| Mesorhizobiu | A0A434JX21 |
| Mesorhizobiu | A0A3S1HYB4 |
| Mesorhizobiu | A0A434EHK3 |
| Mesorhizobiu | A0A434CMJ8 |
| Mesorhizobiu | A0A454WRI7 |
| Mesorhizobiu | A0A3S2FRB0 |
| Mesorhizobiu | A0A436KR71 |
| Mesorhizobiu | A0A436KVN4 |
| Mesorhizobiu | A0A3S2F8X3 |
| Mesorhizobiu | A0A3S1VTV1 |
| Mesorhizobiu | A0A435QXV3 |
| Mesorhizobiu | A0A3S1V2H9 |
| Mesorhizobiu | A0A435NFG8 |
| Mesorhizobiu | A0A435K6G4 |
| Mesorhizobiu | A0A3S1V0K8 |
| Mesorhizobiu | A0A436HLD5 |
| Mesorhizobiu | A0A3S2ERE3 |
| Mesorhizobiu | A0A3S2AZW4 |
| Mesorhizobiu | A0A3S2LEV5 |
| Mesorhizobiu | A0A3S2DZD0 |
| Mesorhizobiu | A0A436BIE9 |
| Mesorhizobiu | A0A3S2FP85 |
| Mesorhizobiu | A0A435P576 |
| Mesorhizobiu | A0A3S1U647 |
| Mesorhizobiu | A0A3S1TF94 |
| Mesorhizobiu | A0A3S1UAZ5 |
| Mesorhizobiu | A0A3S2DDW8 |
| Mesorhizobiu | A0A436BS52 |
| Mesorhizobiu | A0A3S1VGD1 |
| Mesorhizobiu | A0A435KA11 |
| Mesorhizobiu | A0A3S1VM62 |
| Mesorhizobiu | A0A3S1T8V1 |
| Mesorhizobiu | A0A3S1TXY0 |
| Mesorhizobiu | A0A435UXY3 |

| | |
|---|---|
| Mesorhizobiu | A0A435U085 |
| Mesorhizobiu | A0A3S1X503 |
| Mesorhizobiu | A0A435QVB4 |
| Mesorhizobiu | A0A436GBI0 |
| Mesorhizobiu | A0A436GCM6 |
| Mesorhizobiu | A0A436HTL9 |
| Mesorhizobiu | A0A3S2HVT7 |
| Mesorhizobiu | A0A436KG24 |
| Mesorhizobiu | A0A435W6B2 |
| Mesorhizobiu | A0A3S1ZTB0 |
| Mesorhizobiu | A0A3S1U3E5 |
| Mesorhizobiu | A0A435BZ93 |
| Mesorhizobiu | A0A3S2DEI1 |
| Mesorhizobiu | A0A435APW1 |
| Mesorhizobiu | A0A3S1QT55 |
| Mesorhizobiu | A0A3S1WAI0 |
| Mesorhizobiu | A0A3S1VV68 |
| Mesorhizobiu | A0A3S2G7B5 |
| Mesorhizobiu | A0A435MSE4 |
| Mesorhizobiu | A0A3S2E2D1 |
| Mesorhizobiu | A0A3S2FII5 |
| Mesorhizobiu | A0A435FX83 |
| Mesorhizobiu | A0A3S2I3S2 |
| Mesorhizobiu | A0A436GZ56 |
| Mesorhizobiu | A0A435RZZ7 |
| Mesorhizobiu | A0A3S2PD80 |
| Mesorhizobiu | A0A3S2ASJ9 |
| Mesorhizobiu | A0A3S2CHW9 |
| Mesorhizobiu | A0A435H1J4 |
| Mesorhizobiu | A0A3S2K8R7 |
| Mesorhizobiu | A0A434HTC2 |
| Mesorhizobiu | A0A434L053 |
| Mesorhizobiu | A0A3S2B6C9 |
| Mesorhizobiu | A0A435W3M5 |
| Mesorhizobiu | A0A3S1RN73 |
| Mesorhizobiu | A0A3S1TEX9 |
| Mesorhizobiu | A0A435WUH2 |
| Mesorhizobiu | A0A3S1YYJ5 |
| Mesorhizobiu | A0A3S1Z5H9 |
| Mesorhizobiu | A0A3S1P7C5 |
| Mesorhizobiu | A0A3S1TWK1 |

| | |
|---|---|
| Mesorhizobiu | A0A3S1Y0D5 |
| Mesorhizobiu | A0A436CQA8 |
| Mesorhizobiu | A0A3S2H9U8 |
| Mesorhizobiu | A0A435PLS9 |
| Mesorhizobiu | A0A434UUY9 |
| Mesorhizobiu | A0A3S1LTZ1 |
| Mesorhizobiu | A0A3S1NPV1 |
| Mesorhizobiu | A0A3S1NFM4 |
| Mesorhizobiu | A0A3S1Q0N3 |
| Mesorhizobiu | A0A3S1PV14 |
| Mesorhizobiu | A0A3T0JEF6 |
| Mesorhizobiu | A0A434H439 |
| Mesorhizobiu | A0A3S1HEH3 |
| Mesorhizobiu | A0A434C2T4 |
| Mesorhizobiu | A0A3S1S2I8 |
| Mesorhizobiu | A0A3S2A461 |
| Mesorhizobiu | A0A3Q8ZK90 |
| Mesorhizobiu | A0A3Q8ZHB5 |
| Mesorhizobiu | A0A3Q8ZIL9 |
| Mesorhizobiu | A0A3S9EAD3 |
| Mesorhizobiu | A0A3S1MPI8 |
| Mesorhizobiu | A0A434UBI8 |
| Mesorhizobiu | A0A434XFT3 |
| Mesorhizobiu | A0A436XWS3 |
| Mesorhizobiu | A0A3S9F3K3 |
| Mesorhizobiu | A0A3S1LKL0 |
| Mesorhizobiu | A0A3Q8XZG9 |
| Mesorhizobiu | A0A1I5G550 |
| Mesorhizobiu | A0A090DJ28 |
| Mesorhizobiu | A0A090DM34 |
| Mesorhizobiu | A0A090DBY7 |
| Mesorhizobiu | A0A1S1TYY4 |
| Mesorhizobiu | A0A3S1DJG9 |
| Mesorhizobiu | A0A434DDZ6 |
| Mesorhizobiu | A0A0Q6P090 |
| Mesorhizobiu | A0A0Q8AAI4 |
| Mesorhizobiu | A0A0Q8AHI2 |
| Mesorhizobiu | A0A0Q8KRL3 |
| Mesorhizobiu | A0A0Q7WFD1 |
| Mesorhizobiu | A0A0Q7WHK9 |
| Mesorhizobiu | A0A1E2T1C8 |

| | |
|---|---|
| Mesorhizobiu | A0A090EMI5 |
| Mesorhizobiu | M5FEZ9 |
| Mesorhizobiu | A0A1C2DYW0 |
| Mesorhizobiu | A0A3S1CC63 |
| Mesorhizobiu | A0A1W6X7T5 |
| Mesorhizobiu | A0A2A3E108 |
| Mesorhizobiu | A0A2A3D8Z1 |
| Mesorhizobiu | A0A2A3DCN7 |
| Mesorhizobiu | A0A2A3CRF1 |
| Mesorhizobiu | A0A2A3B4T1 |
| Mesorhizobiu | A0A1A5S3H4 |
| Mesorhizobiu | A0A2A3C913 |
| Mesorhizobiu | A0A2A3BMA6 |
| Mesorhizobiu | A0A2A3BJT5 |
| Mesorhizobiu | A0A2A3BJX8 |
| Mesorhizobiu | A0A2A3BT00 |
| Mesorhizobiu | A0A2A3APY2 |
| Mesorhizobiu | A0A3F2UDB7 |
| Mesorhizobiu | A0A1I7B871 |
| Mesorhizobiu | A0A440WS30 |
| Mesorhizobiu | A0A441BU16 |
| Mesorhizobiu | A0A3S4GRJ1 |
| Mesorhizobiu | A0A441KJB7 |
| Mesorhizobiu | A0A441X4E6 |
| Mesorhizobiu | A0A440BP34 |
| Mesorhizobiu | A0A442P447 |
| Mesorhizobiu | A0A440DU90 |
| Mesorhizobiu | A0A440FRF7 |
| Mesorhizobiu | A0A439HIJ1 |
| Mesorhizobiu | A0A440UWQ5 |
| Mesorhizobiu | A0A440VCK8 |
| Mesorhizobiu | A0A3S4JKR1 |
| Mesorhizobiu | A0A439H2W3 |
| Mesorhizobiu | A0A442F213 |
| Mesorhizobiu | A0A3S3MKL6 |
| Mesorhizobiu | A0A440LVW5 |
| Mesorhizobiu | A0A439LX41 |
| Mesorhizobiu | A0A443D0J6 |
| Mesorhizobiu | A0A439SGK3 |
| Mesorhizobiu | A0A441U7J5 |
| Mesorhizobiu | A0A439Y445 |

| | |
|---|---|
| Mesorhizobiι | A0A3S3FIG9 |
| Mesorhizobiι | A0A442J3G7 |
| Mesorhizobiι | A0A439J7B3 |
| Mesorhizobiι | A0A3S3JNQ5 |
| Mesorhizobiι | A0A440S8T8 |
| Mesorhizobiι | A0A3S3FVM8 |
| Mesorhizobiι | A0A442U2E3 |
| Mesorhizobiι | A0A3S4EUE8 |
| Mesorhizobiι | A0A439VKN8 |
| Mesorhizobiι | A0A439QVQ7 |
| Mesorhizobiι | A0A442LAV1 |
| Mesorhizobiι | A0A3S3ENK4 |
| Mesorhizobiι | A0A440VEQ9 |
| Mesorhizobiι | A0A441HPM6 |
| Mesorhizobiι | A0A439V159 |
| Mesorhizobiι | A0A442JPW1 |
| Mesorhizobiι | A0A439P052 |
| Mesorhizobiι | A0A442WYU6 |
| Mesorhizobiι | A0A3S3DCE8 |
| Mesorhizobiι | A0A3S3F5U4 |
| Mesorhizobiι | A0A3S3DDR4 |
| Mesorhizobiι | A0A3S4EHK5 |
| Mesorhizobiι | A0A442ZIC7 |
| Mesorhizobiι | A0A442Y373 |
| Mesorhizobiι | A0A442Y1R2 |
| Mesorhizobiι | A0A440J505 |
| Mesorhizobiι | A0A442JQC4 |
| Mesorhizobiι | A0A442SPV3 |
| Mesorhizobiι | A0A442VU88 |
| Mesorhizobiι | A0A439PZL8 |
| Mesorhizobiι | A0A440GZD4 |
| Mesorhizobiι | A0A442S531 |
| Mesorhizobiι | A0A440KF60 |
| Mesorhizobiι | A0A442GQ66 |
| Mesorhizobiι | A0A442QWA3 |
| Mesorhizobiι | A0A441U254 |
| Mesorhizobiι | A0A440N3Y3 |
| Mesorhizobiι | A0A442MNF5 |
| Mesorhizobiι | A0A3S3F8A3 |
| Mesorhizobiι | A0A3S4BUS5 |
| Mesorhizobiι | A0A3S3FRU0 |

| | |
|---|---|
| Mesorhizobiu | A0A3S3K008 |
| Mesorhizobiu | A0A3S4KZN4 |
| Mesorhizobiu | A0A3S3JXW3 |
| Mesorhizobiu | A0A443AW77 |
| Mesorhizobiu | A0A439U373 |
| Mesorhizobiu | A0A442M396 |
| Mesorhizobiu | A0A440I076 |
| Mesorhizobiu | A0A443ER01 |
| Mesorhizobiu | A0A440CPT0 |
| Mesorhizobiu | A0A442G6N0 |
| Mesorhizobiu | A0A442JQB2 |
| Mesorhizobiu | A0A441T6H6 |
| Mesorhizobiu | A0A439IC62 |
| Mesorhizobiu | A0A439ZLB8 |
| Mesorhizobiu | A0A439FL51 |
| Mesorhizobiu | A0A440L807 |
| Mesorhizobiu | A0A442XI98 |
| Mesorhizobiu | A0A440NAA5 |
| Mesorhizobiu | A0A442XL36 |
| Mesorhizobiu | A0A442Q031 |
| Mesorhizobiu | A0A440T1P1 |
| Mesorhizobiu | A0A441CKQ3 |
| Mesorhizobiu | A0A439V3E1 |
| Mesorhizobiu | A0A441SU59 |
| Mesorhizobiu | A0A442YNW8 |
| Mesorhizobiu | A0A442HP09 |
| Mesorhizobiu | A0A439GTQ9 |
| Mesorhizobiu | A0A439Z2H0 |
| Mesorhizobiu | A0A440JZN6 |
| Mesorhizobiu | A0A442YRX4 |
| Mesorhizobiu | A0A442GXF5 |
| Mesorhizobiu | A0A439MT07 |
| Mesorhizobiu | A0A443G159 |
| Mesorhizobiu | A0A442NGT0 |
| Mesorhizobiu | A0A442JPV2 |
| Mesorhizobiu | A0A439IIB9 |
| Mesorhizobiu | A0A443A4B0 |
| Mesorhizobiu | A0A439LUJ8 |
| Mesorhizobiu | A0A440CVJ3 |
| Mesorhizobiu | A0A440GVE4 |
| Mesorhizobiu | A0A440JFB0 |

| | |
|---|---|
| Mesorhizobiu | A0A441TTA0 |
| Mesorhizobiu | A0A443EMA7 |
| Mesorhizobiu | A0A442IY47 |
| Mesorhizobiu | A0A3S3EEL7 |
| Mesorhizobiu | A0A443APX6 |
| Mesorhizobiu | A0A441V9P3 |
| Mesorhizobiu | A0A442J2Z3 |
| Mesorhizobiu | A0A442UTB2 |
| Mesorhizobiu | A0A440IBK9 |
| Mesorhizobiu | A0A440EGN0 |
| Mesorhizobiu | A0A439XJM6 |
| Mesorhizobiu | A0A443G1J9 |
| Mesorhizobiu | A0A442JQC7 |
| Mesorhizobiu | A0A442WB73 |
| Mesorhizobiu | A0A442ZRR9 |
| Mesorhizobiu | A0A440EB87 |
| Mesorhizobiu | A0A442I508 |
| Mesorhizobiu | A0A442B2D2 |
| Mesorhizobiu | A0A439RGV3 |
| Mesorhizobiu | A0A442XE59 |
| Mesorhizobiu | A0A441XQ43 |
| Mesorhizobiu | A0A439KBP8 |
| Mesorhizobiu | A0A443FHR0 |
| Mesorhizobiu | A0A442V753 |
| Mesorhizobiu | A0A439XT51 |
| Mesorhizobiu | A0A439KGC3 |
| Mesorhizobiu | A0A439FUH3 |
| Mesorhizobiu | A0A442VDM9 |
| Mesorhizobiu | A0A440FQR3 |
| Mesorhizobiu | A0A443CHW7 |
| Mesorhizobiu | A0A443CDR9 |
| Mesorhizobiu | A0A443E755 |
| Mesorhizobiu | A0A442YE60 |
| Mesorhizobiu | A0A442QCW7 |
| Mesorhizobiu | A0A443E034 |
| Mesorhizobiu | A0A443BMA7 |
| Mesorhizobiu | A0A440JYY7 |
| Mesorhizobiu | A0A442NIN0 |
| Mesorhizobiu | A0A3P3G863 |
| Mesorhizobiu | A0A271LPN7 |
| Mesorhizobiu | A0A3A5KAS4 |

| | |
|---|---|
| Mesorhizobiu | A0A3A5L844 |
| Mesorhizobiu | A0A271KG65 |
| Metarhizium | E9DV60 |
| Metarhizium | A0A0B2WJX9 |
| Metarhizium | A0A0B4GVG0 |
| Metarhizium | A0A0B4GKK8 |
| Metarhizium | A0A0B4IAF8 |
| Metarhizium | A0A0B4G2X5 |
| Metarhizium | A0A167AMX9 |
| Metarhizium | A0A014N1B6 |
| Methanoregu | A0A3N5N557 |
| Methanosaet | A0A1V4V301 |
| Methanosaet | A0A1V5AGG2 |
| Methanosaet | A0A1V5AWY6 |
| Methanospiri | Q2FN86 |
| Methanospiri | A0A2V2MVB0 |
| Methylarcula | A0A3R7NX32 |
| Methylobacte | A0A2R4WGM2 |
| Methylobrevi | A0A1E3GZU6 |
| Methylobrevi | A0A1E3H0I6 |
| Methylocella | B8EJN0 |
| Methylocysta | A0A395CUD5 |
| Methylosinus | A0A177P9W2 |
| Methylosinus | A0A2U1SNI8 |
| Methylosinus | A0A2D2CXA0 |
| Micractinium | A0A2P6V6K0 |
| Micractinium | A0A2P6VA87 |
| Micractinium | A0A2P6VCG7 |
| Micractinium | A0A2P6V942 |
| Micractinium | A0A2P6VI36 |
| Micractinium | A0A2P6VHH0 |
| Micractinium | A0A2P6VA39 |
| Micractinium | A0A2P6V3M9 |
| Micractinium | A0A2P6V2A4 |
| Micractinium | A0A2P6VB87 |
| Micractinium | A0A2P6VES9 |
| Micractinium | A0A2P6VPE2 |
| Micractinium | A0A2P6VH52 |
| Micractinium | A0A2P6VES2 |
| Micractinium | A0A2P6VML2 |
| Micractinium | A0A2P6V2M3 |

| | |
|---|---|
| Micractinium | A0A2P6VH58 |
| Micractinium | A0A2P6VK95 |
| Micractinium | A0A2P6VMI8 |
| Micractinium | A0A2P6VBH1 |
| Micractinium | A0A2P6V9Q5 |
| Micractinium | A0A2P6VI19 |
| Micractinium | A0A2P6VGM6 |
| Microbotryur | A0A238FKR1 |
| Microcoleace | A0A349JEI0 |
| Microdochiur | A0A136J612 |
| Microdochiur | A0A136IJV1 |
| Microdochiur | A0A136J606 |
| Microdochiur | A0A136J3L2 |
| Microdochiur | A0A136ITG3 |
| Micromonas | C1EAS9 |
| Micromonas | C1E8G6 |
| Micromonas | C1E8N1 |
| Micromonas | C1FJX3 |
| Micromonas | C1E192 |
| Micromonas | C1MHF5 |
| Micromonas | C1MZS3 |
| Micromonas | C1MT37 |
| Microvirga sp | A0A086MKH3 |
| Mimivirus Bo | A0A159ZQX3 |
| Mimivirus Bo | A0A159ZSK6 |
| Mimivirus Bo | A0A165XJE2 |
| Mimivirus sp. | A0A3S8UXA9 |
| Mimivirus sp. | A0A3Q8U814 |
| Mimivirus sp. | A0A3S8UYS9 |
| Minicystis ros | A0A1L6LTM2 |
| Mitsuaria noc | A0A246H318 |
| Mitsuaria sp. | A0A2G9CDV5 |
| Mitsuaria sp. | A0A1I6IYG6 |
| Mizuhopecte | A0A210Q014 |
| Mizuhopecte | A0A210Q865 |
| Mizuhopecte | A0A210QCM8 |
| Mizuhopecte | A0A210QRE0 |
| Mizuhopecte | A0A210PFV8 |
| Mizuhopecte | A0A210Q3Z0 |
| Mizuhopecte | A0A210QIL1 |
| Mizuhopecte | A0A210QAM1 |

| | |
|---|---|
| Mizuhopecte | A0A210R204 |
| Mizuhopecte | A0A210Q475 |
| Mizuhopecte | A0A210PED4 |
| Mizuhopecte | A0A210Q439 |
| Mizuhopecte | A0A210QTF2 |
| Mizuhopecte | A0A210R6V4 |
| Mizuhopecte | A0A210PET2 |
| Mizuhopecte | A0A210QAL4 |
| Mizuhopecte | A0A210QH12 |
| Mizuhopecte | A0A210PTJ6 |
| Mizuhopecte | A0A210QDU9 |
| Mizuhopecte | A0A210PKW3 |
| Mizuhopecte | A0A210PYD7 |
| Mizuhopecte | A0A210PEI4 |
| Mizuhopecte | A0A210Q2X6 |
| Mizuhopecte | A0A210QH27 |
| Mizuhopecte | A0A210Q9G7 |
| Mizuhopecte | A0A210QMZ3 |
| Mizuhopecte | A0A210PNY2 |
| Mizuhopecte | A0A210PKZ8 |
| Mizuhopecte | A0A210QAL2 |
| Mizuhopecte | A0A210QQF1 |
| Mizuhopecte | A0A210Q9Y6 |
| Mola mola.xl: | A0A3Q3W1J6 |
| Mola mola.xl: | A0A3Q3WBY9 |
| Mola mola.xl: | A0A3Q3W970 |
| Mola mola.xl: | A0A3Q4B8Q9 |
| Mola mola.xl: | A0A3Q3WI21 |
| Mola mola.xl: | A0A3Q3XDI1 |
| Mola mola.xl: | A0A3Q3VSX4 |
| Monilinia fru | A0A395J3G5 |
| Monilinia fru | A0A395J2C4 |
| Moniliophtho | E2L5Y0 |
| Moniliophtho | V2YJ73 |
| Moniliophtho | A0A0W0G933 |
| Monodelphis | F7G1K1 |
| Monodelphis | F7F243 |
| Monodelphis | F7FKH6 |
| Monodelphis | F6Y0F0 |
| Monodelphis | F7FL16 |
| Monopterus | A0A3Q3IFF7 |

| | |
|---|---|
| Monopterus | A0A3Q3Q2I1 |
| Monopterus | A0A3Q3Q2G6 |
| Monopterus | A0A3Q3QBW9 |
| Monopterus | A0A3Q3JMK3 |
| Monopterus | A0A3Q3ILB0 |
| Monopterus | A0A3Q3IFK2 |
| Monopterus | A0A3Q3KJG2 |
| Monoraphidi | A0A0D2MPA4 |
| Monoraphidi | A0A0D2LGW3 |
| Monoraphidi | A0A0D2N2N7 |
| Monoraphidi | A0A0D2J9H2 |
| Monoraphidi | A0A0D2M8B4 |
| Monosiga bre | A9V705 |
| Monosiga bre | A9V3K4 |
| Monosiga bre | A9V0V7 |
| Monosiga bre | A9VCQ3 |
| Monosiga bre | A9VCI2 |
| Monosiga bre | A9V610 |
| Monosiga bre | A9VDX2 |
| Monosiga bre | A9UWN2 |
| Monosiga bre | A9VAX1 |
| Monosiga bre | A9UWV1 |
| Monosiga bre | A9UYM0 |
| Monosiga bre | A9V068 |
| Monosiga bre | A9V646 |
| Monosiga bre | A9V0Z5 |
| Monosiga bre | A9UNM7 |
| Monosiga bre | A9UTM0 |
| Monosiga bre | A9UVN3 |
| Moricandia a | Q8LP49 |
| Morus notabi | W9S1W6 |
| Morus notabi | W9SC56 |
| Morus notabi | W9S2J3 |
| Morus notabi | W9RMV6 |
| Morus notabi | W9R2F2 |
| Morus notabi | W9SEU6 |
| Morus notabi | W9RS41 |
| Morus notabi | W9RZT6 |
| Morus notabi | W9RGD7 |
| Morus notabi | W9SVD8 |
| Morus notabi | W9S9U6 |

| | |
|---|---|
| Morus notabi | W9SAT2 |
| Morus notabi | W9SGF4 |
| Morus notabi | W9SFY6 |
| Morus notabi | W9R8I4 |
| Morus notabi | W9QN88 |
| Morus notabi | W9RRQ7 |
| Morus notabi | W9S089 |
| Morus notabi | W9QYJ9 |
| Morus notabi | W9SJF4 |
| Morus notabi | W9R8B3 |
| Morus notabi | W9RV46 |
| Morus notabi | W9SEU1 |
| Morus notabi | W9SHC4 |
| Morus notabi | W9QI76 |
| Morus notabi | W9RLJ1 |
| Morus notabi | W9RUW2 |
| Morus notabi | W9S558 |
| Morus notabi | W9S6F9 |
| Morus notabi | W9QMK7 |
| Morus notabi | W9S5M0 |
| Morus notabi | W9S6T4 |
| Morus notabi | W9RG28 |
| Morus notabi | W9SB99 |
| Morus notabi | W9RS35 |
| Morus notabi | W9SIC7 |
| Morus notabi | W9S774 |
| Morus notabi | W9R087 |
| Morus notabi | W9S4J8 |
| Morus notabi | W9QCV1 |
| Morus notabi | W9QMB2 |
| Morus notabi | W9QMB6 |
| Morus notabi | W9RTS6 |
| Morus notabi | W9QJS9 |
| Morus notabi | W9RP60 |
| Morus notabi | W9S6W6 |
| Morus notabi | W9R543 |
| Moumouviru | A0A2P1EL58 |
| Moumouviru | A0A2P1ELG5 |
| Moumouviru | A0A2P1ELE8 |
| Moumouviru | M1NNC1 |
| Moumouviru | M1PN76 |

Moumouviru M1PXF7
Moumouviru H2EEU0
Moumouviru H2EF15
Moumouviru H2EET9
Mucuna prur A0A371HKR4
Mucuna prur A0A371E8L2
Mucuna prur A0A371HQJ5
Mucuna prur A0A371F8L2
Mucuna prur A0A371HH60
Mucuna prur A0A371E8Q5
Mucuna prur A0A371GAP6
Mucuna prur A0A371I1M6
Mucuna prur A0A371HCN9
Mucuna prur A0A371HKR5
Mucuna prur A0A371GHE4
Mucuna prur A0A371FRB0
Mucuna prur A0A371H143
Mucuna prur A0A371IEP8
Mucuna prur A0A371HMC9
Mucuna prur A0A371G0L4
Mucuna prur A0A371H7G4
Mucuna prur A0A371F5G3
Mucuna prur A0A371EQX7
Mucuna prur A0A371HK74
Mucuna prur A0A371HCI3
Mucuna prur A0A371F802
Mucuna prur A0A371GT85
Mucuna prur A0A371E352
Mucuna prur A0A371H7E8
Mucuna prur A0A371EKZ3
Mucuna prur A0A371EVC7
Mucuna prur A0A371HGU5
Mucuna prur A0A371GDQ1
Mucuna prur A0A371H1M0
Mucuna prur A0A371G1Y0
Mucuna prur A0A371EKI3
Mucuna prur A0A371HKT8
Mucuna prur A0A371HDB0
Mucuna prur A0A371GTF3
Mucuna prur A0A371G807
Mucuna prur A0A371G1X9

Mucuna prur A0A371F145
Mucuna prur A0A371GCZ5
Mucuna prur A0A371HJ92
Mucuna prur A0A371HMD7
Mucuna prur A0A371GXN7
Mucuna prur A0A371FRU0
Mucuna prur A0A371HB49
Mucuna prur A0A371HDD5
Mucuna prur A0A371HKS1
Mucuna prur A0A371IER3
Mucuna prur A0A371EZ33
Mucuna prur A0A371HH99
Mucuna prur A0A371F945
Mucuna prur A0A371IEM8
Mucuna prur A0A371E4N2
Mucuna prur A0A371EL02
Mucuna prur A0A371HBI0
Mucuna prur A0A371F936
Mucuna prur A0A371HME2
Mucuna prur A0A371FJ78
Mucuna prur A0A371GQ72
Mucuna prur A0A371HCE8
Mucuna prur A0A371EN60
Mucuna prur A0A371GC08
Mucuna prur A0A371E6H1
Mucuna prur A0A371HKB0
Mucuna prur A0A371FWJ5
Mucuna prur A0A371HMF4
Mucuna prur A0A371GXU7
Mucuna prur A0A371HMY2
Mucuna prur A0A371HME4
Mucuna prur A0A371ECI5
Mucuna prur A0A371EGW9
Mucuna prur A0A371HMM7
Mucuna prur A0A371HK49
Mucuna prur A0A371H7E5
Mucuna prur A0A371EBC1
Mucuna prur A0A371EB07
Mucuna prur A0A371ECG7
Mucuna prur A0A371FNR9
Mucuna prur A0A371HAZ0

| | |
|---|---|
| Mucuna prur | A0A371EUQ1 |
| Mucuna prur | A0A371E529 |
| Mucuna prur | A0A371HCF5 |
| Mucuna prur | A0A371EGY1 |
| Mucuna prur | A0A371ET34 |
| Mucuna prur | A0A371FNL4 |
| Mucuna prur | A0A371E3B1 |
| Mucuna prur | A0A371EG56 |
| Mucuna prur | A0A371HB25 |
| Mucuna prur | A0A371HB07 |
| Mucuna prur | A0A371I1M7 |
| Mucuna prur | A0A371ER06 |
| Mus musculu | P20918 |
| Mus musculu | Q08048 |
| Mus musculu | P26928 |
| Mus musculu | P26262 |
| Mus musculu | Q3V1T9 |
| Mus musculu | Q6TCI0 |
| Mus musculu | Q8C4E2 |
| Mus musculu | E0CXN0 |
| Mus musculu | H9H9R5 |
| Mus musculu | Q3UZ05 |
| Mus musculu | Q91XG8 |
| Mus musculu | Q8C9G5 |
| Mus musculu | Q91Y47 |
| Musa acumin | M0SRJ9 |
| Musa acumin | M0SUC6 |
| Musa acumin | M0U4G9 |
| Musa acumin | M0RY71 |
| Musa acumin | M0SRJ7 |
| Musa acumin | M0TM00 |
| Musa acumin | M0S383 |
| Musa acumin | M0RH24 |
| Musa acumin | M0TF78 |
| Musa acumin | M0TDY2 |
| Musa acumin | M0U372 |
| Musa acumin | M0TB77 |
| Musa acumin | M0T6Z8 |
| Musa acumin | M0SRJ8 |
| Musa acumin | M0RVH0 |
| Musa acumin | M0TDY4 |

| | |
|---|---|
| Musa acumin | M0T1W0 |
| Musa acumin | M0S810 |
| Musa acumin | M0U587 |
| Musa acumin | M0TCZ2 |
| Musa acumin | M0RY70 |
| Musa acumin | M0TNE6 |
| Musa acumin | M0UAY4 |
| Musa acumin | M0SP19 |
| Musa acumin | M0TVR6 |
| Musa acumin | M0S240 |
| Musa acumin | M0S243 |
| Musa acumin | M0TZM0 |
| Musa acumin | M0T308 |
| Musa acumin | M0U8W4 |
| Musa acumin | M0TLZ6 |
| Musca dome: | A0A1I8MY63 |
| Musca dome: | A0A1I8M6T0 |
| Musca dome: | A0A1I8M8N5 |
| Musca dome: | A0A1I8N6V2 |
| Musca dome: | A0A1I8NAG5 |
| Musca dome: | A0A1I8MV57 |
| Musca dome: | A0A1I8NEV1 |
| Mustela puto | G9K488 |
| Mustela puto | M3YFX5 |
| Mustela puto | M3Z3L5 |
| Mustela puto | M3YFW6 |
| Mustela puto | M3YQ99 |
| Mustela puto | M3XZU6 |
| Myceliophth( | G2QGY9 |
| Mycoplana di | A0A2T5B5R9 |
| Mycoplana di | A0A2T5BAW0 |
| Myotis brand | S7PRA1 |
| Myotis brand | S7QDN3 |
| Myotis brand | S7MS05 |
| Myotis brand | S7NFT0 |
| Myotis davidi | L5LR27 |
| Myotis davidi | L5LGA9 |
| Myotis davidi | L5M976 |
| Myotis lucifu; | G1PGD5 |
| Myotis lucifu; | G1PSR3 |
| Myotis lucifu; | G1Q6A3 |

| | |
|---|---|
| Mytilus gallo| A0A409V789 |
| Mytilus gallo| A0A3L5TVE3 |
| Mytilus gallo| A0A3L5TQ46 |
| Mytilus gallo| A0A3R5Q2Z7 |
| Myxococcale | A0A2E8RW71 |
| Myxococcale | A0A2D5EZ81 |
| Myxococcus | A0A0F7E4G9 |
| Myxococcus | A0A1I0BCH0 |
| Myxococcus | L7U0L6 |
| Nannochloris | Q68BK5 |
| Nasonia vitri | K7J454 |
| Nasonia vitri | K7IRC8 |
| Nasonia vitri | K7J148 |
| Nasonia vitri | K7J2B0 |
| Nasonia vitri | K7J4P8 |
| Necator ame | W2TTT0 |
| Necator ame | W2TUD4 |
| Necator ame | W2TVZ2 |
| Necator ame | W2TY26 |
| Necator ame | W2SM98 |
| Necator ame | W2TYP1 |
| Necator ame | W2U1I9 |
| Necator ame | W2TDU4 |
| Necator ame | W2TCG9 |
| Necator ame | W2T3P4 |
| Necator ame | W2TVN9 |
| Necator ame | W2TK17 |
| Necator ame | W2SPF8 |
| Necator ame | W2TZT5 |
| Necator ame | W2SJP5 |
| Necator ame | W2T4Y4 |
| Necator ame | W2TP33 |
| Necator ame | W2TVL3 |
| Necator ame | W2TNE7 |
| Necator ame | W2T4H9 |
| Necator ame | W2T7H4 |
| Necator ame | W2T2S3 |
| Necator ame | W2TRM4 |
| Nectria haem | C7YKV9 |
| Nectria haem | C7ZLU7 |
| Nectria haem | C7YHL5 |

| | |
|---|---|
| Nectria haem | C7ZII7 |
| Nectria haem | C7ZNF6 |
| Nectria haem | C7Z7I1 |
| Nectria haem | C7YTH9 |
| Nectria haem | C7ZD94 |
| Nectria haem | C7ZLD5 |
| Nectria haem | C7ZBG8 |
| Nectria haem | C7YSG4 |
| Nectria haem | C7ZEX1 |
| Nectria haem | C7Z173 |
| Nectria haem | C7ZLD0 |
| Nectria haem | C7YXE5 |
| Nelumbo nuc | A0A1U8Q9H4 |
| Nelumbo nuc | A0A1U7ZHR0 |
| Nelumbo nuc | A0A1U8AZI5 |
| Nelumbo nuc | A0A1U8QBE6 |
| Nelumbo nuc | A0A1U7ZR51 |
| Nelumbo nuc | A0A1U7ZR95 |
| Nelumbo nuc | A0A1U7YTF2 |
| Nelumbo nuc | A0A1U8PYZ2 |
| Nelumbo nuc | A0A1U7ZWW8 |
| Nelumbo nuc | A0A1U8A5W5 |
| Nelumbo nuc | A0A1U8B247 |
| Nelumbo nuc | A0A1U8A090 |
| Nelumbo nuc | A0A1U8AYK2 |
| Nelumbo nuc | A0A1U8Q8H4 |
| Nelumbo nuc | A0A1U8B4B8 |
| Nelumbo nuc | A0A1U8A4R2 |
| Nelumbo nuc | A0A1U7Z5Y2 |
| Nelumbo nuc | A0A1U7Z9J8 |
| Nelumbo nuc | A0A1U7ZJI1 |
| Nelumbo nuc | A0A1U8A7C2 |
| Nelumbo nuc | A0A1U8B095 |
| Nelumbo nuc | A0A1U7ZYN8 |
| Nelumbo nuc | A0A1U8QBD9 |
| Nelumbo nuc | A0A1U7ZYY7 |
| Nelumbo nuc | A0A1U8Q7Y9 |
| Nelumbo nuc | A0A1U7ZSA9 |
| Nelumbo nuc | A0A1U8ARE9 |
| Nelumbo nuc | A0A1U8Q7X5 |
| Nelumbo nuc | A0A1U8BCK6 |

| | |
|---|---|
| Nelumbo nuc | A0A1U7ZLL7 |
| Nelumbo nuc | A0A1U7Z475 |
| Nelumbo nuc | A0A1U8QCS1 |
| Nelumbo nuc | A0A1U7ZID0 |
| Nelumbo nuc | A0A1U8B0A1 |
| Nelumbo nuc | A0A1U7YTS6 |
| Nelumbo nuc | A0A1U8AM32 |
| Nelumbo nuc | A0A1U8AN45 |
| Nelumbo nuc | A0A1U8B9J8 |
| Nelumbo nuc | A0A1U8AIZ5 |
| Nelumbo nuc | A0A1U8Q0W3 |
| Nelumbo nuc | A0A1U7ZY78 |
| Nelumbo nuc | A0A1U8AZI0 |
| Nelumbo nuc | A0A1U8BI59 |
| Nelumbo nuc | A0A1U8ABD3 |
| Nelumbo nuc | A0A1U8AY95 |
| Nelumbo nuc | A0A1U7ZHQ5 |
| Nelumbo nuc | A0A1U8QAY8 |
| Nelumbo nuc | A0A1U7ZFF0 |
| Nelumbo nuc | A0A1U7ZRU9 |
| Nelumbo nuc | A0A1U7Z1L5 |
| Nelumbo nuc | A0A1U8A603 |
| Nelumbo nuc | A0A1U7Z5Y7 |
| Nelumbo nuc | A0A1U8ARI0 |
| Nelumbo nuc | A0A1U8Q9G1 |
| Nelumbo nuc | A0A1U8ALH3 |
| Nelumbo nuc | A0A1U7Z9W7 |
| Nelumbo nuc | A0A1U8Q6V5 |
| Nelumbo nuc | A0A1U8BCG2 |
| Nelumbo nuc | A0A1U7Z2V6 |
| Nelumbo nuc | A0A1U7Z4C3 |
| Nelumbo nuc | A0A1U7Z3J0 |
| Nelumbo nuc | A0A1U8A7U0 |
| Nelumbo nuc | A0A1U8AN31 |
| Nelumbo nuc | A0A1U8Q7F4 |
| Nelumbo nuc | A0A1U8Q349 |
| Nelumbo nuc | A0A1U8BA38 |
| Nelumbo nuc | A0A1U8BGB5 |
| Nelumbo nuc | A0A1U8AM80 |
| Nelumbo nuc | A0A1U7ZY31 |
| Nematostella | A7SZH1 |

Nematostella A7SIZ3
Nematostella A7SRX6
Nematostella A7SBL4
Nematostella A7RSM4
Nematostella A7RNE4
Nematostella A7RMZ7
Nematostella A7S8E8
Nematostella A7RX77
Nematostella A7RFP1
Nematostella A7RRU0
Nematostella A7SBM9
Nematostella A7RQ47
Nematostella A7SHY9
Nematostella A7S6R4
Nematostella A8DUZ5
Nematostella A7RU88
Nematostella A7T3J8
Nematostella A7SHH8
Nematostella A7S968
Nematostella A7SRE8
Nematostella A7RPS0
Nematostella A7SM86
Nematostella A7SEE3
Nematostella A7S366
Nematostella A7T6F6
Nematostella A7RWW1
Nematostella A7S6Y4
Nematostella A7S6U4
Nematostella A7SAJ9
Nematostella A7S7H3
Nematostella A7SFL5
Nematostella A7S6T9
Nematostella A7RNV4
Nematostella A7SDI1
Nematostella A7S941
Nematostella A7RKH7
Nematostella A7S6Y5
Nematostella A7T2L8
Nematostella A7SEM5
Nematostella A7SGJ1
Nematostella A7SRX5

Nematostella A7RU08
Nematostella A7S942
Nematostella A7SRE3
Nematostella A7T7G4
Nematostella A7SCA6
Nematostella A7RIK4
Nematostella A7RWB2
Nematostella A7T1F4
Nematostella A7RQ57
Nematostella A7SI46
Nematostella A7SBD5
Nematostella A7RHX3
Nematostella A7RQH4
Nematostella A7T3Q6
Nematostella A7SFU7
Nematostella A7RSX6
Nematostella A7RWB4
Nematostella A7RUJ4
Nematostella A7SVL2
Nematostella A7RVC4
Nematostella A7SXQ3
Nematostella A7SIB1
Nematostella A7SCH3
Nematostella A7S3Q4
Nematostella A7RWZ4
Nematostella A7S293
Nematostella A7SX86
Nematostella A7SSU6
Nematostella A7S547
Nematostella A7SR11
Nematostella A7S6D1
Nematostella A7SDG5
Nematostella A7RJD0
Nematostella A7RQE2
Nematostella A7RP92
Nematostella A7S3L4
Nematostella A7T9M2
Nematostella A7T780
Nematostella A7T8U0
Nematostella A7SE24
Nematostella A7RYZ8

| | |
|---|---|
| Nematostella | A7SD38 |
| Nematostella | A7SC13 |
| Nematostella | A7S6L2 |
| Nematostella | A7SJ07 |
| Nematostella | A7SN79 |
| Nematostella | A7TCJ6 |
| Nematostella | A7RTC0 |
| Nematostella | A7T1Z2 |
| Nematostella | A7RWV8 |
| Nematostella | A7S6U0 |
| Nematostella | A7SL04 |
| Nematostella | A7RJN3 |
| Nematostella | A7SNB6 |
| Nematostella | A7RM74 |
| Nematostella | A7SK70 |
| Nematostella | A7S6Y3 |
| Neocallimasti | A0A1Y2BK59 |
| Neocallimasti | A0A1Y2A5T7 |
| Neolamprolo | A0A3Q4MLG0 |
| Neolamprolo | A0A3Q4I4H3 |
| Neolamprolo | A0A3Q4I627 |
| Neolamprolo | A0A3Q4H1X3 |
| Neolamprolo | A0A3Q4HU08 |
| Neolamprolo | A0A3Q4H1Q8 |
| Neolamprolo | A0A3Q4GWT9 |
| Neolamprolo | A0A3Q4IEG5 |
| Neolecta irre | A0A1U7LM76 |
| Neomonachu | A0A2Y9H1Q2 |
| Neomonachu | A0A2Y9HDN4 |
| Neomonachu | A0A2Y9HJV5 |
| Neomonachu | A0A2Y9HGI1 |
| Neomonachu | A0A2Y9HK72 |
| Neomonachu | A0A2Y9HH66 |
| Neomonachu | A0A2Y9GTJ0 |
| Neomonachu | A0A2Y9HJV1 |
| Neomonachu | A0A2Y9H6M8 |
| Neonectria di | A0A0P7B6P7 |
| Neonectria di | A0A0P7BBW4 |
| Neonectria di | A0A0P7BTD6 |
| Neonectria di | A0A0N8H811 |
| Neonectria di | A0A0P7BG58 |

| | |
|---|---|
| Neonectria di | A0A0P7BD71 |
| Neonectria di | A0A0P7ALI4 |
| Neonectria di | A0A0P7BIY5 |
| Neonectria di | A0A0P7B5T9 |
| Neonectria di | A0A0P7B6I9 |
| Neonectria di | A0A0P7BLF3 |
| Neonectria di | A0A0P7AUR2 |
| Neonectria di | A0A0P7AKI4 |
| Neonectria di | A0A0P7BL89 |
| Neonectria di | A0A0P7BSA0 |
| Neophocaena | A0A341BPU6 |
| Neophocaena | A0A341BNE4 |
| Neophocaena | A0A341C1A0 |
| Neophocaena | A0A341BM03 |
| Neophocaena | A0A341ATW5 |
| Neophocaena | A0A341C3X0 |
| Neophocaena | A0A341BNF2 |
| Neophocaena | A0A341CM99 |
| Neophocaena | A0A341BLW0 |
| Neorhizobium | A0A068TC39 |
| Neorhizobium | A0A0T7H3C9 |
| Neorhizobium | A0A0T7FB36 |
| Neorhizobium | A0A0T7FXT8 |
| Neorhizobium | A0A068SW47 |
| Neorhizobium | A0A0T7HIV6 |
| Neorhizobium | A0A386E2E6 |
| Neorhizobium | A0A346YQL0 |
| Neosartorya | A1DN01 |
| Neosartorya | Q4WBJ3 |
| Neosartorya | B0YAB7 |
| Neosartorya | B0XSS6 |
| Neosartorya | A0A229XX59 |
| Neosartorya | A0A229X3Y5 |
| Neospora car | F0V833 |
| Neospora car | F0V892 |
| Neospora car | F0VKE6 |
| Neospora car | F0VBW4 |
| Neospora car | F0V9H8 |
| Neospora car | F0JAV0 |
| Neospora car | F0VJW9 |
| Neospora car | F0VDC1 |

| | |
|---|---|
| Neospora car | F0V7W6 |
| Neospora car | F0VJW8 |
| Neospora car | F0VI90 |
| Neospora car | F0VKF4 |
| Neospora car | A0A0F7UGU6 |
| Neospora car | A0A0F7UM27 |
| Neospora car | F0VKE8 |
| Neospora car | A0A0F7UH55 |
| Neospora car | F0VQW3 |
| Neospora car | F0V8T4 |
| Neospora car | A0A0F7UIB5 |
| Neospora car | A0A0F7U8D9 |
| Neospora car | A0A0F7U4M4 |
| Neospora car | F0V8K8 |
| Neospora car | F0VKE7 |
| Neospora car | A0A0F7UQW1 |
| Neospora car | F0VCD3 |
| Neospora car | F0VJW7 |
| Neospora car | A0A0H4LVB4 |
| Neospora car | A0A0H4LYS9 |
| Neospora car | A0A0H4MAT4 |
| Neospora car | Q2KHJ3 |
| Neotoma lep | A0A1A6FUU5 |
| Neotoma lep | A0A1A6FTR8 |
| Neotoma lep | A0A1A6GY88 |
| Neotoma lep | A0A1A6HIS2 |
| Neotoma lep | A0A1A6FW21 |
| Neovison visc | U6CR68 |
| Nephila clavi | A0A2P6L9E4 |
| Nephila clavi | A0A2P6L9F6 |
| Nephila clavi | A0A2P6LI56 |
| Nephila clavi | A0A2P6L9G3 |
| Nephila clavi | A0A2P6KM95 |
| Nephila clavi | A0A2P6K685 |
| Nephila clavi | A0A2P6L9F0 |
| Nephila clavi | A0A2P6K678 |
| Nephila clavi | A0A2P6KT77 |
| Nephila clavi | A0A2P6K695 |
| Neslia panicu | D6PR18 |
| Neslia panicu | D6PR67 |
| Nestor notab | A0A091S6P5 |

| | |
|---|---|
| Nestor notab | A0A091SBL5 |
| Nestor notab | A0A091T5V5 |
| Nestor notab | A0A091S1J9 |
| Neurospora c | A7UVW4 |
| Neurospora c | Q7S109 |
| Neurospora c | V5ILM8 |
| Neurospora c | Q7SFE8 |
| Neurospora c | V5ILY1 |
| Neurospora c | A0A0B0DW84 |
| Neurospora c | Q96U11 |
| Neurospora c | A0A0B0E396 |
| Neurospora c | A0A0B0E9R9 |
| Neurospora c | A0A0B0E9P7 |
| Neurospora t | F8N082 |
| Neurospora t | F8N4S9 |
| Neurospora t | F8MEC8 |
| Neurospora t | F8MC30 |
| Neurospora t | G4U7X4 |
| Neurospora t | G4UFH6 |
| Neurospora t | G4UD81 |
| Neurospora t | G4U9Q3 |
| Nicotiana att | A0A1J6HWP7 |
| Nicotiana att | A0A314L2N5 |
| Nicotiana att | A0A314KMT1 |
| Nicotiana att | A0A1J6IQT9 |
| Nicotiana att | A0A314KMZ2 |
| Nicotiana att | A0A314KWS7 |
| Nicotiana att | A0A314LDT0 |
| Nicotiana att | A0A1J6J3K8 |
| Nicotiana att | A0A1J6KDN9 |
| Nicotiana att | A0A1J6ISD8 |
| Nicotiana att | A0A1J6I486 |
| Nicotiana att | A0A1J6ICK1 |
| Nicotiana att | A0A1J6KII4 |
| Nicotiana att | A0A1J6JDG5 |
| Nicotiana att | A0A314KM80 |
| Nicotiana att | A0A1J6IQM4 |
| Nicotiana att | A0A314L5L5 |
| Nicotiana att | A0A314KNT5 |
| Nicotiana att | A0A314L814 |
| Nicotiana att | A0A1J6I9A0 |

| | |
|---|---|
| Nicotiana att | A0A314L5W9 |
| Nicotiana att | A0A1J6INQ4 |
| Nicotiana att | A0A1J6IB07 |
| Nicotiana att | A0A1J6IG63 |
| Nicotiana att | A0A1J6JYU0 |
| Nicotiana att | A0A1J6KUR6 |
| Nicotiana att | A0A314L5M3 |
| Nicotiana att | A0A1J6I9G8 |
| Nicotiana att | A0A1J6KXS8 |
| Nicotiana att | A0A1J6L9X1 |
| Nicotiana att | A0A1J6IAJ9 |
| Nicotiana att | A0A314KLH0 |
| Nicotiana att | A0A314KHP6 |
| Nicotiana att | A0A1J6K8T5 |
| Nicotiana att | A0A1J6K0Y6 |
| Nicotiana att | A0A1J6KY06 |
| Nicotiana att | A0A1J6I3J3 |
| Nicotiana att | A0A1J6I7I0 |
| Nicotiana att | A0A1J6J380 |
| Nicotiana att | A0A314L474 |
| Nicotiana att | A0A1J6HU55 |
| Nicotiana att | A0A1J6KMP9 |
| Nicotiana att | A0A1J6JV58 |
| Nicotiana att | A0A314L438 |
| Nicotiana att | A0A1J6IJN2 |
| Nicotiana att | A0A314L6V5 |
| Nicotiana att | G4XXY5 |
| Nicotiana att | A0A1J6J9K7 |
| Nicotiana att | A0A1J6KSS8 |
| Nicotiana att | A0A1J6IF17 |
| Nicotiana att | A0A1J6ID41 |
| Nicotiana att | A0A314L7K3 |
| Nicotiana att | A0A1J6L0B3 |
| Nicotiana att | A0A1J6IWL3 |
| Nicotiana att | A0A1J6IGU9 |
| Nicotiana ber | B9A9C5 |
| Nicotiana glu | B6EB06 |
| Nicotiana syl | A0A1U7VDG6 |
| Nicotiana syl | A0A1U7VUJ3 |
| Nicotiana syl | A0A1U7X9V3 |
| Nicotiana syl | A0A1U7WCZ7 |

| | |
|---|---|
| Nicotiana sylv | A0A1U7XDD8 |
| Nicotiana sylv | A0A1U7VEB0 |
| Nicotiana sylv | A0A1U7Y2A8 |
| Nicotiana sylv | A0A1U7XQK4 |
| Nicotiana sylv | A0A1U7WXX2 |
| Nicotiana sylv | A0A1U7WDB0 |
| Nicotiana sylv | A0A1U7YLM9 |
| Nicotiana sylv | A0A1U7WKL4 |
| Nicotiana sylv | A0A1U7WLL2 |
| Nicotiana sylv | A0A1U7XQZ3 |
| Nicotiana sylv | A0A1U7W2G6 |
| Nicotiana sylv | A0A1U7WAK5 |
| Nicotiana sylv | A0A1U7W505 |
| Nicotiana sylv | A0A1U7WKF6 |
| Nicotiana sylv | A0A1U7VZG2 |
| Nicotiana sylv | A0A1U7WH46 |
| Nicotiana sylv | A0A1U7Y9T1 |
| Nicotiana sylv | A0A1U7WED3 |
| Nicotiana sylv | A0A1U7WV83 |
| Nicotiana sylv | A0A1U7VL33 |
| Nicotiana sylv | A0A1U7XGW4 |
| Nicotiana sylv | A0A1U7VQE3 |
| Nicotiana sylv | A0A1U7YRY2 |
| Nicotiana sylv | A0A1U7W1W1 |
| Nicotiana sylv | A0A1U7XXL7 |
| Nicotiana sylv | A0A1U7Z1C5 |
| Nicotiana sylv | A0A1U7YL89 |
| Nicotiana sylv | A0A1U7WMH9 |
| Nicotiana sylv | A0A1U7VPX9 |
| Nicotiana sylv | A0A1U7X9V8 |
| Nicotiana sylv | A0A1U7V5P9 |
| Nicotiana sylv | A0A1U7XLP8 |
| Nicotiana sylv | A0A1U7WT92 |
| Nicotiana sylv | A0A1U7V2M1 |
| Nicotiana sylv | A0A1U7WD36 |
| Nicotiana sylv | A0A1U7VZ90 |
| Nicotiana sylv | A0A1U7XDE3 |
| Nicotiana sylv | A0A1U7V541 |
| Nicotiana sylv | A0A1U7VJP5 |
| Nicotiana sylv | A0A1U7W2Y1 |
| Nicotiana sylv | A0A1U7VKV2 |

| | |
|---|---|
| Nicotiana sylv | A0A1U7W5R6 |
| Nicotiana sylv | A0A1U7WW15 |
| Nicotiana sylv | A0A1U7W0I9 |
| Nicotiana sylv | A0A1U7Y7Q9 |
| Nicotiana sylv | A0A1U7VN54 |
| Nicotiana sylv | A0A1U7VUC4 |
| Nicotiana sylv | A0A1U7WBT7 |
| Nicotiana sylv | A0A1U7XL68 |
| Nicotiana sylv | A0A1U7W973 |
| Nicotiana sylv | A0A1U7X5P3 |
| Nicotiana sylv | A0A1U7YK15 |
| Nicotiana sylv | A0A1U7XTF4 |
| Nicotiana sylv | A0A1U7WPB7 |
| Nicotiana sylv | A0A1U7UUN4 |
| Nicotiana sylv | A0A1U7WCX4 |
| Nicotiana sylv | A0A1U7XPV3 |
| Nicotiana sylv | A0A1U7Y3E7 |
| Nicotiana sylv | A0A1U7XGV9 |
| Nicotiana sylv | A0A1U7Y6J9 |
| Nicotiana sylv | A0A1U7VQF3 |
| Nicotiana sylv | A0A1U7XPM9 |
| Nicotiana sylv | A0A1U7WVB8 |
| Nicotiana sylv | A0A1U7W731 |
| Nicotiana sylv | A0A1U7XYW4 |
| Nicotiana sylv | A0A1U7VPS9 |
| Nicotiana sylv | A0A1U7WLD5 |
| Nicotiana sylv | A0A1U7XC49 |
| Nicotiana sylv | A0A1U7XLK4 |
| Nicotiana sylv | A0A1U7W910 |
| Nicotiana sylv | A0A1U7WKM8 |
| Nicotiana sylv | A0A1U7Y8N9 |
| Nicotiana sylv | A0A1U7VET8 |
| Nicotiana sylv | A0A1U7WMI3 |
| Nicotiana sylv | A0A1U7VC33 |
| Nicotiana sylv | A0A1U7XCN9 |
| Nicotiana sylv | A0A1U7W9Z4 |
| Nicotiana sylv | A0A1U7X4E1 |
| Nicotiana sylv | A0A1U7VW07 |
| Nicotiana sylv | A0A1U7W8N5 |
| Nicotiana sylv | A0A1U7VNJ5 |
| Nicotiana sylv | A0A1U7VUJ7 |

| | |
|---|---|
| Nicotiana syl | A0A1U7XS99 |
| Nicotiana tab | A0A1S4A8G6 |
| Nicotiana tab | A0A1S3Z9W1 |
| Nicotiana tab | A0A1S3ZKW9 |
| Nicotiana tab | A0A1S3YVQ2 |
| Nicotiana tab | A0A1S3ZKG6 |
| Nicotiana tab | A0A1S3ZPQ4 |
| Nicotiana tab | A0A1S3XZW1 |
| Nicotiana tab | A0A1S4AIL3 |
| Nicotiana tab | A0A1S4AMR9 |
| Nicotiana tab | A0A1S4DLY3 |
| Nicotiana tab | A0A1S4AQV3 |
| Nicotiana tab | A0A1S4A3C8 |
| Nicotiana tab | A0A1S4BVH7 |
| Nicotiana tab | A0A1S3YT81 |
| Nicotiana tab | A0A1S4DJR1 |
| Nicotiana tab | A0A1S4BG39 |
| Nicotiana tab | A0A1S4AAV5 |
| Nicotiana tab | A0A1S3YCL1 |
| Nicotiana tab | A0A1S3YGV9 |
| Nicotiana tab | A0A1S4AS71 |
| Nicotiana tab | A0A1S4CBP5 |
| Nicotiana tab | A0A1S4AIV1 |
| Nicotiana tab | A0A1S3YGK9 |
| Nicotiana tab | A0A1S4C8A5 |
| Nicotiana tab | A0A1S3ZLY6 |
| Nicotiana tab | A0A1S3XKI4 |
| Nicotiana tab | A0A1S4BVI6 |
| Nicotiana tab | A0A1S3YS52 |
| Nicotiana tab | A0A1S3XGG9 |
| Nicotiana tab | A0A1S4C2V4 |
| Nicotiana tab | A0A1S4C7V6 |
| Nicotiana tab | A0A1S3YX32 |
| Nicotiana tab | A0A1S3X9D4 |
| Nicotiana tab | A0A1S4D768 |
| Nicotiana tab | A0A1S4CCG0 |
| Nicotiana tab | A0A1S4C7R5 |
| Nicotiana tab | A0A1S4AAW3 |
| Nicotiana tab | A0A1S3ZFI4 |
| Nicotiana tab | A0A1S4C0I7 |
| Nicotiana tab | A0A1S3Z379 |

| | |
|---|---|
| Nicotiana tab | A0A1S4CT07 |
| Nicotiana tab | A0A1S4BZ07 |
| Nicotiana tab | A0A1S4CBP9 |
| Nicotiana tab | A0A1S3YCE1 |
| Nicotiana tab | A0A1S3Y0A5 |
| Nicotiana tab | A0A1S4DI92 |
| Nicotiana tab | A0A1S4ASV5 |
| Nicotiana tab | A0A1S3XFC6 |
| Nicotiana tab | A0A1S3X1W3 |
| Nicotiana tab | A0A1S4D7F2 |
| Nicotiana tab | A0A1S4D8R5 |
| Nicotiana tab | A0A1S4C899 |
| Nicotiana tab | A0A1S4AB13 |
| Nicotiana tab | A0A1S3YI37 |
| Nicotiana tab | A0A1S4BCW1 |
| Nicotiana tab | A0A1S3ZBM9 |
| Nicotiana tab | A0A1S3ZWG5 |
| Nicotiana tab | A0A1S3YH70 |
| Nicotiana tab | A0A1S4DEP7 |
| Nicotiana tab | A0A1S3YCZ3 |
| Nicotiana tab | A0A1S4BQF7 |
| Nicotiana tab | A0A1S3Z2M2 |
| Nicotiana tab | A0A1S4DM05 |
| Nicotiana tab | A0A1S4DN45 |
| Nicotiana tab | A0A1S4BD11 |
| Nicotiana tab | A0A1S3Z7R3 |
| Nicotiana tab | A0A1S3ZF88 |
| Nicotiana tab | A0A1S3ZBH4 |
| Nicotiana tab | A0A1S3ZK91 |
| Nicotiana tab | A0A1S3XKN5 |
| Nicotiana tab | A0A1S4DEY0 |
| Nicotiana tab | A0A1S4DPZ0 |
| Nicotiana tab | A0A1S3YCZ7 |
| Nicotiana tab | A0A1S4BJE9 |
| Nicotiana tab | A0A1S4B6P0 |
| Nicotiana tab | A0A1S4DER9 |
| Nicotiana tab | A0A1S4A3U1 |
| Nicotiana tab | A0A1S3ZG58 |
| Nicotiana tab | A0A1S4BX98 |
| Nicotiana tab | A0A1S3YZW8 |
| Nicotiana tab | A0A1S4AAQ4 |

| | |
|---|---|
| Nicotiana tab | A0A1S3Y6Q2 |
| Nicotiana tab | A0A1S3Y621 |
| Nicotiana tab | A0A1S4BVJ0 |
| Nicotiana tab | A0A1S4DI53 |
| Nicotiana tab | A0A1S4AGM7 |
| Nicotiana tab | A0A1S3ZWE8 |
| Nicotiana tab | A0A1S4C7S1 |
| Nicotiana tab | A0A1S4C2N1 |
| Nicotiana tab | A0A1S4BVM4 |
| Nicotiana tab | A0A1S4AHG2 |
| Nicotiana tab | A0A1S4BIZ9 |
| Nicotiana tab | A0A1S3Z1I1 |
| Nicotiana tab | A0A1S4BD90 |
| Nicotiana tab | A0A1S3ZKX2 |
| Nicotiana tab | A0A1S4B4B4 |
| Nicotiana tab | A0A1S3Z7T1 |
| Nicotiana tab | A0A1S3ZXB1 |
| Nicotiana tab | A0A1S3YHU7 |
| Nicotiana tab | B9A9C4 |
| Nicotiana tab | A0A1S4BF74 |
| Nicotiana tab | A0A1S4A7H4 |
| Nicotiana tab | A0A1S3YH89 |
| Nicotiana tab | A0A1S3ZFQ4 |
| Nicotiana tab | A0A1S3XJQ9 |
| Nicotiana tab | A0A1S4AV00 |
| Nicotiana tab | A0A1S3ZWC2 |
| Nicotiana tab | A0A1S4AJE1 |
| Nicotiana tab | A0A1S4C3P0 |
| Nicotiana tab | A0A1S3Z2H5 |
| Nicotiana tab | A0A1S3Z761 |
| Nicotiana tab | A0A1S3ZGF4 |
| Nicotiana tab | A0A1S4ARU2 |
| Nicotiana tab | A0A1S4AAC7 |
| Nicotiana tab | A0A1S3ZBB3 |
| Nicotiana tab | A0A1S3ZWF8 |
| Nicotiana tab | A0A1S3ZIS3 |
| Nicotiana tab | A0A1S4BU26 |
| Nicotiana tab | A0A1S3ZIZ8 |
| Nicotiana tab | A0A1S4CTL2 |
| Nicotiana tab | A0A1S3Y6I1 |
| Nicotiana tab | A0A1S4BXG7 |

| | |
|---|---|
| Nicotiana tab | A0A1S3WZV5 |
| Nicotiana tab | A0A1S4BFA7 |
| Nicotiana tab | A0A1S3YUE5 |
| Nicotiana tab | A0A1S3ZFP6 |
| Nicotiana tab | A0A1S4BBN7 |
| Nicotiana tab | A0A1S4AAQ8 |
| Nicotiana tab | A0A1S4A7H2 |
| Nicotiana tab | A0A1S3YHW7 |
| Nicotiana tab | A0A1S4BF06 |
| Nicotiana tab | A0A1S4B7P0 |
| Nicotiana tab | A0A1S4A3X6 |
| Nicotiana tab | A0A1S4BF18 |
| Nicotiana tab | A0A1S4DQJ7 |
| Nicotiana tab | A0A1S4C7S6 |
| Nicotiana tab | A0A1S4DI64 |
| Nicotiana tab | A0A1S3YNY7 |
| Nicotiana tab | A0A1S3YZ96 |
| Nicotiana tab | E5D7E5 |
| Nicotiana tab | A0A1S4BTQ9 |
| Nicotiana tab | A0A1S4AAV8 |
| Nicotiana tab | A0A1S4BVJ1 |
| Nicotiana tab | A0A1S4CI20 |
| Nicotiana tab | A0A1S3YL50 |
| Nicotiana tab | A0A1S4AHU4 |
| Nicotiana tab | A0A1S4AGG2 |
| Nicotiana tab | A0A1S4C7S0 |
| Nicotiana tab | A0A1S4AGE4 |
| Nicotiana tab | A0A1S4C7G1 |
| Nicotiana tab | A0A1S4C1I5 |
| Nicotiana tab | A0A1S4C7F1 |
| Nicotiana tab | A0A1S4DEL0 |
| Nicotiana tab | A0A1S4C4Q3 |
| Nicotiana tab | A0A1S4BTX0 |
| Nicotiana tab | A0A1S4CUC0 |
| Niemeyer viru | A0A0U2UIV4 |
| Niemeyer viru | A0A0U2TUD0 |
| Niemeyer viru | A0A0U2K3G9 |
| Nipponia nipp | A0A091VZY0 |
| Nipponia nipp | A0A091W0K6 |
| Nipponia nipp | A0A091UP08 |
| Nipponia nipp | A0A091ULT1 |

| | |
|---|---|
| Nippostrongy | A0A0N4XE48 |
| Nippostrongy | A0A3P7CNB1 |
| Nippostrongy | A0A0N4YIX3 |
| Nippostrongy | A0A3P7A1Y9 |
| Nippostrongy | A0A3P7AQW4 |
| Nippostrongy | A0A0N4Y5X9 |
| Nippostrongy | A0A158R0D4 |
| Nippostrongy | A0A0N4XTQ9 |
| Nippostrongy | A0A3P7B0Z6 |
| Nippostrongy | A0A0N4YDL0 |
| Nippostrongy | A0A0N4Y521 |
| Nippostrongy | A0A0N4XTZ4 |
| Nippostrongy | A0A0N4YX44 |
| Nippostrongy | A0A158QWT4 |
| Nippostrongy | A0A158R0F4 |
| Nippostrongy | A0A0N4YRN0 |
| Nippostrongy | A0A0N4XCL2 |
| Nippostrongy | A0A158R0A2 |
| Nippostrongy | A0A0N4XP57 |
| Nippostrongy | A0A0N4YJU3 |
| Nippostrongy | A0A158QZV0 |
| Nippostrongy | A0A0N4XMB6 |
| Nippostrongy | A0A158R2Z3 |
| Nippostrongy | A0A0N4YT44 |
| Nippostrongy | A0A0N4XW83 |
| Nippostrongy | A0A158R099 |
| Nippostrongy | A0A0N4XDZ8 |
| Nippostrongy | A0A0N4XLC5 |
| Nippostrongy | A0A0N4YJQ3 |
| Nippostrongy | A0A0N4YT73 |
| Nippostrongy | A0A0N4XKU9 |
| Nippostrongy | A0A158QZH2 |
| Nippostrongy | A0A158QX22 |
| Nippostrongy | A0A0N4YVS7 |
| Nippostrongy | A0A0N4XWH8 |
| Nippostrongy | A0A0N4XTR1 |
| Nippostrongy | A0A0N4YUM8 |
| Nippostrongy | A0A0N4YSD3 |
| Nitratireduct | K2PNL4 |
| Nitratireduct | K2P3G0 |
| Nitratireduct | K2LLY9 |

| | |
|---|---|
| Nitratireduct | A0A1H4K6F6 |
| Nitratireduct | A0A2P8ND91 |
| Nitratireduct | A0A2E0JHX2 |
| Nitrobacter h | Q1QRJ6 |
| Nitrobacter s | A0A1Q3JTV4 |
| Nitrobacter s | A0A1M3C7Q1 |
| Nitrobacter s | A3WSN9 |
| Nitrobacter v | A0A1V4I335 |
| Nitrobacter w | Q3SMU6 |
| Nitrosomona | A0A1H8BAS7 |
| Nitrosomona | A0A1I0D0D5 |
| Nitrosomona | A0A1H9S965 |
| Nitrosomona | A0A358JLT2 |
| Nitrospira sp. | A0A2N6HD53 |
| Noccaea caer | A0A1J3FP03 |
| Noccaea caer | A0A1J3HMF5 |
| Noccaea caer | A0A1J3HW34 |
| Noccaea caer | A0A1J3GB39 |
| Noccaea caer | A0A1J3JNB8 |
| Noccaea caer | A0A1J3JZS2 |
| Noccaea caer | A0A1J3ISG7 |
| Noccaea caer | A0A1J3DXH5 |
| Noccaea caer | A0A1J3E580 |
| Noccaea caer | A0A1J3GBL6 |
| Noccaea caer | A0A1J3K818 |
| Noccaea caer | A0A1J3H057 |
| Noccaea caer | A0A1J3GQN5 |
| Noccaea caer | A0A1J3HRL3 |
| Noccaea caer | A0A1J3DRG8 |
| Noccaea caer | A0A1J3GNA1 |
| Noccaea caer | A0A1J3J3Z7 |
| Noccaea caer | A0A1J3JRE5 |
| Noccaea caer | A0A1J3FL37 |
| Noccaea caer | A0A1J3JQL6 |
| Noccaea caer | A0A1J3GRF7 |
| Noccaea caer | A0A1J3JX24 |
| Noccaea caer | A0A1J3GGC1 |
| Noccaea caer | A0A1J3E7G3 |
| Noccaea caer | A0A1J3F3A5 |
| Noccaea caer | A0A1J3JPT9 |
| Noccaea caer | A0A1J3DU93 |

Noccaea caer A0A1J3DTR9
Noccaea caer A0A1J3CXR2
Noccaea caer A0A1J3JBY9
Noccaea caer A0A1J3E9F3
Noccaea caer A0A1J3FSR4
Noccaea caer A0A1J3JJ88
Noccaea caer A0A1J3H7F2
Noccaea caer A0A1J3JTS5
Noccaea caer A0A1J3FCK6
Noccaea caer A0A1J3FKJ0
Noccaea caer A0A1J3DSR5
Noccaea caer A0A1J3GCV7
Noccaea caer A0A1J3FI92
Noccaea caer A0A1J3G1G3
Noccaea caer A0A1J3HR35
Noccaea caer A0A1J3DHA0
Noccaea caer A0A1J3E845
Noccaea caer A0A1J3FS72
Noccaea caer A0A1J3G1T7
Noccaea caer A0A1J3H5P9
Noccaea caer A0A1J3CQE8
Noccaea caer A0A1J3IWX7
Noccaea caer A0A1J3G0P7
Noccaea caer A0A1J3IM74
Noccaea caer A0A1J3DTR0
Noccaea caer A0A1J3JQL3
Noccaea caer A0A1J3E7J0
Noccaea caer A0A1J3DTZ1
Noccaea caer A0A1J3J3Q6
Noccaea caer A0A1J3CCB9
Noccaea caer A0A1J3FIV7
Noccaea caer A0A1J3EAN5
Noccaea caer A0A1J3IVU3
Noccaea caer A0A1J3GZI8
Noccaea caer A0A1J3HK52
Noccaea caer A0A1J3HAN9
Noccaea caer A0A1J3JM42
Noccaea caer A0A1J3DI79
Noccaea caer A0A1J3DSA9
Noccaea caer A0A1J3DX70
Noccaea caer A0A1J3HPH3

Noccaea caer A0A1J3DW17
Noccaea caer A0A1J3DSG0
Noccaea caer A0A1J3FWI3
Noccaea caer A0A1J3IK70
Noccaea caer A0A1J3IIK4
Noccaea caer A0A1J3JKH6
Noccaea caer A0A1J3CFM4
Noccaea caer A0A1J3EV57
Noccaea caer A0A1J3IX87
Noccaea caer A0A1J3I886
Noccaea caer A0A1J3J875
Noccaea caer A0A1J3EEU1
Noccaea caer A0A1J3ES29
Noccaea caer A0A1J3JTY3
Noccaea caer A0A1J3I7Z8
Noccaea caer A0A1J3EFY5
Noccaea caer A0A1J3CL94
Noccaea caer A0A1J3I0U3
Noccaea caer A0A1J3IVB8
Noccaea caer A0A1J3D311
Noccaea caer A0A1J3EEH9
Noccaea caer A0A1J3DS04
Noccaea caer A0A1J3D5T0
Noccaea caer A0A1J3DME5
Noccaea caer A0A1J3IH96
Noccaea caer A0A1J3F2S3
Noccaea caer A0A1J3JRD9
Noccaea caer A0A1J3GZM1
Noccaea caer A0A1J3FB28
Noccaea caer A0A1J3EN22
Noccaea caer A0A1J3HDE8
Noccaea caer A0A1J3CMT3
Noccaea caer A0A1J3EEN0
Noccaea caer A0A1J3FFP7
Noccaea caer A0A1J3JUT5
Noccaea caer A0A1J3D2I7
Noccaea caer A0A1J3H709
Noccaea caer A0A1J3DF21
Noccaea caer A0A1J3J0E4
Noccaea caer A0A1J3H499
Noccaea caer A0A1J3GMB9

| | |
|---|---|
| Noccaea caer | A0A1J3GVE5 |
| Noccaea caer | A0A1J3DWJ7 |
| Noccaea caer | A0A1J3JHE1 |
| Noccaea caer | A0A1J3DMM0 |
| Noccaea caer | A0A1J3JS61 |
| Noccaea caer | A0A1J3IP86 |
| Noccaea caer | A0A1J3FSR0 |
| Noccaea caer | A0A1J3J1W0 |
| Noccaea caer | A0A1J3GG27 |
| Noccaea caer | A0A1J3EMF4 |
| Noccaea caer | A0A1J3G4X8 |
| Noccaea caer | A0A1J3ELH1 |
| Noccaea caer | A0A1J3EPX9 |
| Noccaea caer | A0A1J3GZP2 |
| Noccaea caer | A0A1J3F8K9 |
| Nomascus leu | A0A2I3GPI4 |
| Nomascus leu | G1RKX8 |
| Nomascus leu | M3ZBA9 |
| Nomascus leu | G1RLC0 |
| Nomascus leu | A0A2I3G9T1 |
| Nomascus leu | G1RXL8 |
| Nomascus leu | A0A2I3HH53 |
| Nomascus leu | G1RLE8 |
| Nomascus leu | G1R5A0 |
| Nostoc comm | A0A2R5FFP8 |
| Nostoc sp. 'P | A0A2D0H9I4 |
| Nostoc sp. 33 | A0A318AZE0 |
| Nothobranch | A0A1A8UHT3 |
| Nothobranch | A0A1A8U2N8 |
| Nothobranch | A0A1A8UGE6 |
| Nothobranch | A0A1A8AMP3 |
| Nothobranch | A0A1A8UYV9 |
| Nothobranch | A0A1A8DKN8 |
| Nothobranch | A0A1A8CX61 |
| Nothobranch | A0A1A8F858 |
| Nothobranch | A0A1A8KU64 |
| Nothobranch | A0A1A8J1X1 |
| Nothobranch | A0A1A8HP15 |
| Nothobranch | A0A1A8Q0T0 |
| Nothobranch | A0A1A8M3J2 |
| Nothobranch | A0A1A8N5R8 |

| | |
|---|---|
| Nothobranch | A0A1A8S3A7 |
| Novosphingo | A0A3D1P0S1 |
| Novosphingo | A0A3D1P0P9 |
| Oceanicaulis | A0A2T4ZM68 |
| Oceanicaulis | A0A2D9MLX3 |
| Oceanicaulis | A0A2E5UVY4 |
| Oceanicola gı | Q2CK54 |
| Oceanicola lit | A0A1N6H695 |
| Oceanicola sp | A0A254R8C3 |
| Oceaniovalib | K2I7J2 |
| Ochrobactrur | A6X4J9 |
| Ochrobactrur | A0A011UJA9 |
| Ochrobactrur | A0A137XJ12 |
| Ochrobactrur | A0A3R8ZDT7 |
| Ochrobactrur | A0A1J6HPX3 |
| Ochrobactrur | A0A256FNR5 |
| Ochrobactrur | A0A397M124 |
| Ochrobactrur | A0A2P9HHH8 |
| Ochrobactrur | U4V6A2 |
| Ochrobactrur | C4WPE7 |
| Ochrobactrur | M5JM46 |
| Ochrobactrur | A0A256GVM1 |
| Ochrobactrur | A0A2S7J3N1 |
| Ochrobactrur | A0A256G407 |
| Ochrobactrur | A0A1A9FPL5 |
| Ochrobactrur | A0A256F053 |
| Ochrobactrur | A0A2M9SAC5 |
| Ochrobactrur | A0A2M9RTW1 |
| Ochrobactrur | A0A2M9RPN7 |
| Ochrobactrur | A0A248UL68 |
| Ochrobactrur | A0A248UDK1 |
| Ochrobactrur | A0A421BXV1 |
| Ochrobactrur | U1XL45 |
| Ochrobactrur | A0A3N0BIY1 |
| Ochrobactrur | A0A2S8YAJ0 |
| Ochrobactrur | A0A2S8Y7R3 |
| Ochrobactrur | A0A2S8TZD4 |
| Ochrobactrur | A0A1V3YWR0 |
| Ochrobactrur | A0A317LE59 |
| Ochrobactrur | A0A292GL07 |
| Ochrobactrur | A0A256FB23 |

| | |
|---|---|
| Octopus bima | A0A0L8HR65 |
| Octopus bima | A0A0L8G008 |
| Octopus bima | A0A0L8GIC5 |
| Octopus bima | A0A0L8G5H6 |
| Octopus bima | A0A0L8HWE5 |
| Octopus bima | A0A0L8HQP4 |
| Octopus bima | A0A0L8HYY9 |
| Octopus bima | A0A0L8G081 |
| Octopus bima | A0A0L8HPA3 |
| Octopus bima | A0A0L8HXI7 |
| Octopus bima | A0A0L8HS26 |
| Octopus bima | A0A0L8FY82 |
| Octopus bima | A0A0L8HWF0 |
| Octopus bima | A0A0L8HQ38 |
| Octopus bima | A0A0L8G444 |
| Octopus bima | A0A0L8IDR0 |
| Octopus bima | A0A0L8G3A8 |
| Octopus bima | A0A0L8HMG8 |
| Octopus bima | A0A0L8HW99 |
| Octopus bima | A0A0L8HWE2 |
| Octopus bima | A0A0L8H6T5 |
| Octopus bima | A0A0L8HQN8 |
| Octopus bima | A0A0L8HP85 |
| Octopus bima | A0A0L8HDL0 |
| Octopus bima | A0A0L8FNP5 |
| Octopus bima | A0A0L8FJ32 |
| Octopus bima | A0A0L8HW19 |
| Octopus bima | A0A0L8GXG5 |
| Octopus bima | A0A0L8GT95 |
| Octopus bima | A0A0L8GXD9 |
| Octopus bima | A0A0L8HWB9 |
| Octopus bima | A0A0L8GSV1 |
| Octopus bima | A0A0L8GYH4 |
| Octopus bima | A0A0L8FUE7 |
| Octopus bima | A0A0L8HNN0 |
| Octopus bima | A0A0L8H7N3 |
| Octopus bima | A0A0L8GWJ9 |
| Octopus bima | A0A0L8G1Z8 |
| Octopus bima | A0A0L8HP77 |
| Octopus bima | A0A0L8G183 |
| Octopus bima | A0A0L8FRC1 |

| | |
|---|---|
| Octopus bima | A0A0L8I283 |
| Octopus bima | A0A0L8HWA4 |
| Octopus bima | A0A0L8I485 |
| Octopus bima | A0A0L8GWC4 |
| Octopus bima | A0A0L8H6J5 |
| Octopus bima | A0A0L8HZ03 |
| Octopus bima | A0A0L8HPE9 |
| Octopus bima | A0A0L8FRF9 |
| Odobenus ro | A0A2U3VG05 |
| Odobenus ro | A0A2U3VGE7 |
| Odobenus ro | A0A2U3VVF2 |
| Odobenus ro | A0A2U3X506 |
| Odobenus ro | A0A2U3W015 |
| Odobenus ro | A0A2U3VZS1 |
| Odobenus ro | A0A2U3ZHH0 |
| Odobenus ro | A0A2U3X5D6 |
| Odobenus ro | A0A2U3X502 |
| Odobenus ro | A0A2U3X508 |
| Oesophagost | A0A0B1SMS5 |
| Oesophagost | A0A0B1SZF3 |
| Oesophagost | A0A0B1T7G4 |
| Oesophagost | A0A0B1TNZ7 |
| Oesophagost | A0A0B1SVQ8 |
| Oesophagost | A0A0B1TSF3 |
| Oesophagost | A0A0B1THL2 |
| Oesophagost | A0A0B1RPJ3 |
| Oesophagost | A0A0B1TVI4 |
| Oesophagost | A0A0B1T7X3 |
| Oesophagost | A0A0B1TLL5 |
| Oesophagost | A0A0B1SJV9 |
| Oesophagost | A0A0B1SZQ3 |
| Oesophagost | A0A0B1TJJ8 |
| Oesophagost | A0A0B1TQU4 |
| Oesophagost | A0A0B1SPE9 |
| Oesophagost | A0A0B1TRM5 |
| Oesophagost | A0A0B1TM49 |
| Oesophagost | A0A0B1TP05 |
| Oesophagost | A0A0B1TLV5 |
| Oesophagost | A0A0B1SLD5 |
| Oesophagost | A0A0B1T699 |
| Oesophagost | A0A0B1TQQ2 |

| | |
|---|---|
| Oesophagost | A0A0B1SIU2 |
| Oidiodendror | A0A0C3GVM9 |
| Oidiodendror | A0A0C3GHZ3 |
| Oidiodendror | A0A0C3GYJ2 |
| Oidiodendror | A0A0C3C6A0 |
| Oidiodendror | A0A0C3GWZ0 |
| Oikopleura di | E4YAT7 |
| Oikopleura di | E4X175 |
| Oikopleura di | E4Z726 |
| Oikopleura di | E4XPL0 |
| Oikopleura di | E4YKT2 |
| Oikopleura di | E4WX62 |
| Oikopleura di | E4WXA9 |
| Oikopleura di | E4X236 |
| Oikopleura di | E4XM60 |
| Oikopleura di | E4X025 |
| Oikopleura di | E4YGB1 |
| Oikopleura di | E4YEZ4 |
| Oikopleura di | E4Y3D2 |
| Okeania hirsu | A0A3N6PYR2 |
| Olea europae | D7RJY6 |
| Olea europae | D7RJY4 |
| Oligotropha c | B6JDB0 |
| Onchocerca f | A0A238BQ40 |
| Onchocerca f | A0A183HYH0 |
| Onchocerca f | A0A183HKI7 |
| Onchocerca f | A0A183H9Q1 |
| Onchocerca f | A0A183HZR7 |
| Onchocerca f | A0A183HM98 |
| Onchocerca f | A0A183I5Y8 |
| Onchocerca f | A0A183H621 |
| Onchocerca f | A0A183HW79 |
| Onchocerca f | A0A183HZQ7 |
| Onchocerca f | A0A183HBN7 |
| Onchocerca f | A0A183HGE2 |
| Onchocerca f | A0A183I2D1 |
| Onchocerca f | A0A183I087 |
| Onchocerca f | A0A183GZD3 |
| Onchocerca f | A0A238C1L1 |
| Onchocerca f | A0A238C0I5 |
| Onchocerca f | A0A238BZT1 |

| | |
|---|---|
| Onchocerca f | A0A238C1R8 |
| Onchocerca f | A0A3P7VC50 |
| Onchocerca f | A0A183HF33 |
| Onchocerca f | A0A238BHJ2 |
| Onchocerca f | A0A238C1C5 |
| Onchocerca f | A0A238BUI0 |
| Onchocerca f | A0A238BPJ5 |
| Onchocerca c | A0A3P7KGB4 |
| Onchocerca c | A0A182ERB1 |
| Onchocerca c | A0A3P7JTR2 |
| Onchocerca c | A0A182EUC0 |
| Onchocerca c | A0A182E9R2 |
| Onchocerca c | A0A182E294 |
| Onchocerca c | A0A3P6SEL8 |
| Onchocerca c | A0A3P6STP6 |
| Onchocerca c | A0A182EK23 |
| Onchocerca c | A0A182EGU8 |
| Onchocerca c | A0A182EBS9 |
| Onchocerca c | A0A182ELR8 |
| Onchocerca c | A0A182E8N8 |
| Onchocerca c | A0A182E853 |
| Onchocerca c | A0A182E148 |
| Onchocerca c | A0A182E7K9 |
| Onchocerca c | A0A182DWJ9 |
| Onchocerca c | A0A182E5V3 |
| Onchocerca v | A0A044TXH1 |
| Onchocerca v | A0A044TB24 |
| Onchocerca v | A0A044V6J3 |
| Onchocerca v | A0A044SAK8 |
| Onchocerca v | A0A044T6C9 |
| Onchocerca v | A0A044TQE7 |
| Onchocerca v | A0A044VG28 |
| Onchocerca v | A0A044TXN9 |
| Onchocerca v | A0A044QW86 |
| Onchocerca v | A0A044TQV4 |
| Onchocerca v | A0A044TS10 |
| Onchocerca v | A0A044VIE3 |
| Onchocerca v | A0A044SXT1 |
| Onchocerca v | A0A2K6W0F3 |
| Onchocerca v | A0A2K6VJT9 |
| Oncopeltus fa | A0A2R7VUY6 |

| | |
|---|---|
| Oncopeltus fa | A0A2R7WMN5 |
| Oncopeltus fa | A0A2R7X3S8 |
| Oncopeltus fa | A0A2R7VTV5 |
| Oncopeltus fa | A0A2R7WIU5 |
| Oncopeltus fa | A0A2R7WIV9 |
| Oncopeltus fa | A0A2R7WJA5 |
| Oncorhynchu | A0A060X7U1 |
| Oncorhynchu | A0A060XRA8 |
| Oncorhynchu | A0A060XME5 |
| Oncorhynchu | A0A060XFN8 |
| Oncorhynchu | A0A060W8B1 |
| Oncorhynchu | A0A060WHF0 |
| Oncorhynchu | Q5DVP8 |
| Oncorhynchu | A0A060XU84 |
| Ooceraea bir | A0A026W7N3 |
| Ooceraea bir | A0A3L8E197 |
| Ooceraea bir | A0A026VZ91 |
| Ooceraea bir | A0A026WBB7 |
| Ooceraea bir | A0A3L8DVC2 |
| Ooceraea bir | A0A3L8DB52 |
| Ooceraea bir | A0A026W4C2 |
| Ooceraea bir | A0A3L8DPD8 |
| Ooceraea bir | A0A026X1V9 |
| Ooceraea bir | A0A3L8E1U8 |
| Ooceraea bir | A0A3L8DP99 |
| Ooceraea bir | A0A026W2Z5 |
| Ooceraea bir | A0A3L8DFM8 |
| Operophtera | A0A0L7LTQ8 |
| Operophtera | A0A0L7L2P1 |
| Operophtera | A0A0L7LM72 |
| Operophtera | A0A0L7L0G5 |
| Operophtera | A0A0L7L335 |
| Operophtera | A0A0L7L2H0 |
| Operophtera | A0A0L7LA69 |
| Ophiocordyce | T5ADX7 |
| Ophiocordyce | T5AAZ9 |
| Ophiocordyce | A0A2A9PMS4 |
| Ophiocordyce | A0A2A9PLL3 |
| Ophiophagus | V8NK09 |
| Ophiophagus | V8PHB4 |
| Ophiostoma | | S3D177 |

| | |
|---|---|
| Ophiostoma p | S3C2I7 |
| Opisthocomu | A0A091WHX0 |
| Opisthocomu | A0A091XY89 |
| Opisthocomu | A0A091WJ34 |
| Opisthocomu | A0A091WK45 |
| Opisthorchis | A0A1S8WUG3 |
| Opisthorchis | A0A074ZV38 |
| Opisthorchis | A0A075A3I4 |
| Opisthorchis | A0A1S8X8S4 |
| Opisthorchis | A0A1S8WQZ1 |
| Opisthorchis | A0A074Z2H8 |
| Opisthorchis | A0A074Z9I0 |
| Opisthorchis | A0A1S8WPP4 |
| Opisthorchis | A0A074ZMS3 |
| Opitutae bact | A0A2E9RXB9 |
| Orchesella cir | A0A1D2NBH8 |
| Orchesella cir | A0A1D2MZY1 |
| Orchesella cir | A0A1D2NJJ1 |
| Orchesella cir | A0A1D2MYU3 |
| Orchesella cir | A0A1D2ND60 |
| Oreochromis | I3K037 |
| Oreochromis | I3KFD6 |
| Oreochromis | I3K036 |
| Oreochromis | I3KFB8 |
| Oreochromis | I3KCE9 |
| Oreochromis | I3JAQ5 |
| Oreochromis | I3JAR3 |
| Oreochromis | I3KCF0 |
| Oreochromis | I3KFB7 |
| Oreochromis | I3J849 |
| Oreochromis | I3J1D2 |
| Ornatilinea a | A0A0P6YFR3 |
| Ornithodoros | A0A293M3U3 |
| Ornithodoros | A0A293LHY5 |
| Ornithorhync | F7CM56 |
| Ornithorhync | F7BQ99 |
| Ornithorhync | F7FSH7 |
| Ornithorhync | F7CM40 |
| Ornithorhync | F7FSI1 |
| Orychophrag | O80354 |
| Oryctes borb | A0A0T6BGW1 |

| | |
|---|---|
| Oryctes borb | A0A0T6B7H3 |
| Oryctes borb | A0A0T6B598 |
| Oryctolagus c | Q95ME7 |
| Oryctolagus c | D0PTV6 |
| Oryctolagus c | G1U9Q9 |
| Oryctolagus c | G1SSQ6 |
| Oryctolagus c | G1T127 |
| Oryctolagus c | G1SMZ3 |
| Oryctolagus c | G1SFS6 |
| Oryctolagus c | D0PPI8 |
| Oryctolagus c | C9E9X4 |
| Oryza alta.xls | E0CW66 |
| Oryza alta.xls | E0CW57 |
| Oryza austral | B9V0G8 |
| Oryza barthii. | A0A0D3HR23 |
| Oryza barthii. | A0A0D3HAE3 |
| Oryza barthii. | A0A0D3EL74 |
| Oryza barthii. | A0A0D3G020 |
| Oryza barthii. | A0A0D3HR22 |
| Oryza barthii. | A0A0D3EX91 |
| Oryza barthii. | A0A0D3H9S4 |
| Oryza barthii. | A0A0D3GD84 |
| Oryza barthii. | A0A0D3ESA3 |
| Oryza barthii. | A0A0D3EUU8 |
| Oryza barthii. | A0A0D3FJY4 |
| Oryza barthii. | A0A0D3GS78 |
| Oryza barthii. | A0A0D3EUU6 |
| Oryza barthii. | A0A0D3G016 |
| Oryza barthii. | A0A0D3EUU9 |
| Oryza barthii. | A0A0D3GS89 |
| Oryza barthii. | A0A0D3FKE2 |
| Oryza barthii. | A0A0D3ESB1 |
| Oryza barthii. | A0A0D3H9S5 |
| Oryza barthii. | A0A0D3FU00 |
| Oryza barthii. | A0A0D3EUU5 |
| Oryza barthii. | A0A0D3GD82 |
| Oryza barthii. | A0A0D3EUU4 |
| Oryza barthii. | A0A0D3EUV0 |
| Oryza barthii. | A0A0D3FU04 |
| Oryza barthii. | A0A0D3ESA2 |
| Oryza barthii. | A0A0D3EWT3 |

| | |
|---|---|
| Oryza barthii | A0A0D3GI36 |
| Oryza barthii | A0A0D3HI98 |
| Oryza barthii | A0A0D3EQN3 |
| Oryza barthii | A0A0D3FR49 |
| Oryza barthii | A0A0D3EL79 |
| Oryza barthii | A0A0D3FXQ4 |
| Oryza barthii | A0A0D3G018 |
| Oryza barthii | A0A0D3H9R8 |
| Oryza barthii | A0A0D3FTZ3 |
| Oryza barthii | A0A0D3G022 |
| Oryza barthii | A0A0D3GHQ2 |
| Oryza barthii | A0A0D3EX86 |
| Oryza barthii | A0A0D3EWT4 |
| Oryza barthii | A0A0D3FVT7 |
| Oryza barthii | A0A0D3HHT7 |
| Oryza barthii | A0A0D3GS79 |
| Oryza barthii | A0A0D3G017 |
| Oryza barthii | A0A0D3G132 |
| Oryza barthii | A0A0D3EL82 |
| Oryza barthii | A0A0D3FVU0 |
| Oryza barthii | A0A0D3H9S1 |
| Oryza barthii | A0A0D3FVT5 |
| Oryza barthii | A0A0D3HCH4 |
| Oryza barthii | A0A0D3EL84 |
| Oryza barthii | A0A0D3FU01 |
| Oryza barthii | A0A0D3EUU3 |
| Oryza barthii | A0A0D3HS10 |
| Oryza barthii | A0A0D3HV71 |
| Oryza barthii | A0A0D3G021 |
| Oryza barthii | A0A0D3H9R9 |
| Oryza barthii | A0A0D3EUV2 |
| Oryza barthii | A0A0D3ESB4 |
| Oryza barthii | A0A0D3FFK8 |
| Oryza barthii | A0A0D3EX49 |
| Oryza barthii | A0A0D3HSZ8 |
| Oryza barthii | A0A0D3HX90 |
| Oryza barthii | A0A0D3H9S2 |
| Oryza barthii | A0A0D3GIF2 |
| Oryza barthii | A0A0D3G9H3 |
| Oryza barthii | A0A0D3HJF3 |
| Oryza barthii | A0A0D3G019 |

| | |
|---|---|
| Oryza barthii | A0A0D3FVT4 |
| Oryza barthii | A0A0D3G3N6 |
| Oryza barthii | A0A0D3FUK4 |
| Oryza barthii | A0A0D3HHT5 |
| Oryza barthii | A0A0D3G011 |
| Oryza barthii | A0A0D3GS82 |
| Oryza barthii | A0A0D3GS90 |
| Oryza barthii | A0A0D3GI35 |
| Oryza barthii | A0A0D3G012 |
| Oryza barthii | A0A0D3G7P0 |
| Oryza barthii | A0A0D3G013 |
| Oryza barthii | A0A0D3G550 |
| Oryza barthii | A0A0D3G024 |
| Oryza barthii | A0A0D3FVS9 |
| Oryza barthii | A0A0D3EL80 |
| Oryza barthii | A0A0D3HCH7 |
| Oryza barthii | A0A0D3EX87 |
| Oryza barthii | A0A0D3FVT0 |
| Oryza barthii | A0A0D3GI32 |
| Oryza barthii | A0A0D3HHT9 |
| Oryza barthii | A0A0D3G3N8 |
| Oryza barthii | A0A0D3FRT4 |
| Oryza barthii | A0A0D3ESB0 |
| Oryza barthii | A0A0D3FT93 |
| Oryza barthii | A0A0D3HHU2 |
| Oryza barthii | A0A0D3G023 |
| Oryza barthii | A0A0D3FUM6 |
| Oryza barthii | A0A0D3ESA7 |
| Oryza barthii | A0A0D3GS81 |
| Oryza barthii | A0A0D3FVS8 |
| Oryza barthii | A0A0D3HJF4 |
| Oryza barthii | A0A0D3G3N7 |
| Oryza barthii | A0A0D3G3P0 |
| Oryza barthii | A0A0D3EJR8 |
| Oryza barthii | A0A0D3G131 |
| Oryza barthii | A0A0D3EL81 |
| Oryza barthii | A0A0D3ERR0 |
| Oryza barthii | A0A0D3G9G2 |
| Oryza barthii | A0A0D3FAT2 |
| Oryza barthii | A0A0D3EUU7 |
| Oryza barthii | A0A0D3HV70 |

| | |
|---|---|
| Oryza barthii | A0A0D3EL83 |
| Oryza barthii | A0A0D3FU02 |
| Oryza brachy | J3L4R2 |
| Oryza brachy | J3L0V7 |
| Oryza brachy | J3MLV8 |
| Oryza brachy | J3LUL2 |
| Oryza brachy | J3L5D6 |
| Oryza brachy | J3LLH5 |
| Oryza brachy | B9V0I5 |
| Oryza brachy | J3MZX1 |
| Oryza brachy | J3LPV8 |
| Oryza brachy | J3L2Q4 |
| Oryza brachy | J3NBU7 |
| Oryza brachy | A0A0U1WXR2 |
| Oryza brachy | J3M4A9 |
| Oryza brachy | G2XM38 |
| Oryza brachy | J3L2R2 |
| Oryza brachy | J3MFK0 |
| Oryza brachy | J3N1W7 |
| Oryza brachy | J3L2A1 |
| Oryza brachy | J3LX34 |
| Oryza brachy | J3KXU0 |
| Oryza brachy | J3M263 |
| Oryza brachy | J3MZW5 |
| Oryza brachy | J3M1Q6 |
| Oryza brachy | J3MLV6 |
| Oryza brachy | J3LQ96 |
| Oryza brachy | J3MZW8 |
| Oryza brachy | J3NB41 |
| Oryza brachy | J3KXT4 |
| Oryza brachy | J3L6M2 |
| Oryza brachy | J3MQB8 |
| Oryza brachy | J3KWJ2 |
| Oryza brachy | J3LXT3 |
| Oryza brachy | J3M4A7 |
| Oryza brachy | J3MID2 |
| Oryza brachy | J3M1Q3 |
| Oryza brachy | J3L4R1 |
| Oryza brachy | J3KXU3 |
| Oryza brachy | J3L2Q7 |
| Oryza brachy | J3MEX9 |

| | |
|---|---|
| Oryza brachy | J3MLV9 |
| Oryza brachy | J3M4A8 |
| Oryza brachy | J3MF85 |
| Oryza brachy | G2XM40 |
| Oryza brachy | J3LVJ9 |
| Oryza brachy | J3LXT4 |
| Oryza brachy | J3L6J2 |
| Oryza brachy | J3LWL4 |
| Oryza brachy | J3MQW0 |
| Oryza brachy | J3KUK1 |
| Oryza brachy | J3M1Q9 |
| Oryza brachy | J3M265 |
| Oryza brachy | J3M1Q4 |
| Oryza brachy | J3MEX8 |
| Oryza brachy | J3LXT1 |
| Oryza brachy | J3KW69 |
| Oryza brachy | J3L2Q8 |
| Oryza brachy | J3L809 |
| Oryza brachy | J3M264 |
| Oryza brachy | J3NFE2 |
| Oryza brachy | J3MF84 |
| Oryza brachy | J3L4Q9 |
| Oryza brachy | J3LXT5 |
| Oryza brachy | J3LW49 |
| Oryza brachy | J3KXU2 |
| Oryza brachy | J3LWL5 |
| Oryza brachy | J3L2Q3 |
| Oryza brachy | J3MLW9 |
| Oryza brachy | J3L2Q5 |
| Oryza brachy | J3M780 |
| Oryza brachy | J3M7G4 |
| Oryza brachy | J3MGY9 |
| Oryza brachy | J3LXT6 |
| Oryza brachy | J3N0F9 |
| Oryza glaberr | I1NQI2 |
| Oryza glaberr | I1NQI4 |
| Oryza glaberr | I1PZU2 |
| Oryza glaberr | I1NKC7 |
| Oryza glaberr | I1PL71 |
| Oryza glaberr | I1QTK2 |
| Oryza glaberr | I1NU21 |

| | |
|---|---|
| Oryza glaberr | I1P904 |
| Oryza glaberr | I1R3R0 |
| Oryza glaberr | I1NTS1 |
| Oryza glaberr | I1NQI7 |
| Oryza glaberr | I1PL76 |
| Oryza glaberr | I1NQI3 |
| Oryza glaberr | I1R3R2 |
| Oryza glaberr | I1R4I3 |
| Oryza glaberr | I1PCQ7 |
| Oryza glaberr | I1QS23 |
| Oryza glaberr | G2XMM6 |
| Oryza glaberr | I1PL69 |
| Oryza glaberr | I1Q381 |
| Oryza glaberr | I1PJJ4 |
| Oryza glaberr | G2XMM7 |
| Oryza glaberr | G2XMM5 |
| Oryza glaberr | I1PPM3 |
| Oryza glaberr | I1QR22 |
| Oryza glaberr | I1PTT5 |
| Oryza glaberr | I1R3R1 |
| Oryza glaberr | G2XMM4 |
| Oryza glaberr | I1PPM4 |
| Oryza glaberr | I1PIJ8 |
| Oryza glaberr | I1QR27 |
| Oryza glaberr | I1PL68 |
| Oryza glaberr | I1QR20 |
| Oryza glaberr | I1PQE8 |
| Oryza glaberr | I1PL73 |
| Oryza glaberr | I1PST0 |
| Oryza glaberr | I1NU52 |
| Oryza glaberr | I1PL72 |
| Oryza glaberr | I1QR24 |
| Oryza glaberr | I1PX09 |
| Oryza glaberr | I1PZT8 |
| Oryza glaberr | I1PPM7 |
| Oryza glaberr | I1NQI6 |
| Oryza glaberr | G2XMV1 |
| Oryza glaberr | I1PST2 |
| Oryza glaberr | I1PMV0 |
| Oryza glaberr | I1QR21 |
| Oryza glaberr | I1R8A7 |

| | |
|---|---|
| Oryza glaberr | I1R3Q9 |
| Oryza glaberr | I1PL77 |
| Oryza glaberr | I1NTS0 |
| Oryza glaberr | I1QYI5 |
| Oryza glaberr | I1NLF0 |
| Oryza glaberr | I1QR28 |
| Oryza glaberr | I1PPM6 |
| Oryza glaberr | I1NS91 |
| Oryza glaberr | I1PK04 |
| Oryza glaberr | I1QXD1 |
| Oryza glaberr | I1PL79 |
| Oryza glaberr | I1P0W9 |
| Oryza glaberr | I1PCD1 |
| Oryza glaberr | I1P4I6 |
| Oryza glaberr | I1PPL9 |
| Oryza glaberr | I1NU57 |
| Oryza glaberr | I1PK06 |
| Oryza glaberr | I1Q383 |
| Oryza glaberr | I1R5B8 |
| Oryza glaberr | I1NQI9 |
| Oryza glaberr | I1NLF3 |
| Oryza glaberr | I1QXD4 |
| Oryza glaberr | I1PK03 |
| Oryza glaberr | I1QBG0 |
| Oryza glaberr | I1QBF9 |
| Oryza glaberr | I1QR19 |
| Oryza glaberr | I1PPM5 |
| Oryza glaberr | I1QBG7 |
| Oryza glaberr | I1PH64 |
| Oryza glaberr | I1QBF7 |
| Oryza glaberr | I1QYI6 |
| Oryza glaberr | I1NQI5 |
| Oryza glaberr | G2XLC0 |
| Oryza glaberr | I1PPM0 |
| Oryza glaberr | I1Q380 |
| Oryza glaberr | I1PQE7 |
| Oryza glaberr | I1PIJ7 |
| Oryza glaberr | I1PSR4 |
| Oryza glaberr | I1PL75 |
| Oryza glaberr | I1NLE2 |
| Oryza glaberr | I1PPM2 |

| | |
|---|---|
| Oryza glaberr | I1PL74 |
| Oryza glaberr | I1PSS6 |
| Oryza glaberr | I1PSS8 |
| Oryza glaberr | I1NTS2 |
| Oryza glaberr | I1QTK4 |
| Oryza glaberr | I1Q3F8 |
| Oryza glaberr | I1NLF2 |
| Oryza glaberr | I1NS88 |
| Oryza glumip | A0A0D9YK84 |
| Oryza glumip | A0A0D9YCX9 |
| Oryza glumip | A0A0D9ZG94 |
| Oryza glumip | A0A0D9ZKN7 |
| Oryza glumip | A0A0D9ZJI1 |
| Oryza glumip | A0A0D9YIH2 |
| Oryza glumip | A0A0D9ZKN8 |
| Oryza glumip | A0A0D9ZR94 |
| Oryza glumip | A0A0E0B677 |
| Oryza glumip | A0A0D9YFW0 |
| Oryza glumip | A0A0D9Y5E5 |
| Oryza glumip | A0A0E0AL88 |
| Oryza glumip | A0A0D9YI07 |
| Oryza glumip | A0A0E0ARK9 |
| Oryza glumip | A0A0D9Y5D1 |
| Oryza glumip | A0A0D9ZMW8 |
| Oryza glumip | A0A0D9Y5E7 |
| Oryza glumip | A0A0D9Z436 |
| Oryza glumip | A0A0D9ZR74 |
| Oryza glumip | A0A0E0AL85 |
| Oryza glumip | A0A0D9ZIM8 |
| Oryza glumip | A0A0D9YFW3 |
| Oryza glumip | A0A0E0A5I8 |
| Oryza glumip | A0A0D9Y5C8 |
| Oryza glumip | A0A0D9ZUP3 |
| Oryza glumip | A0A0E0BEX9 |
| Oryza glumip | A0A0E0AL72 |
| Oryza glumip | A0A0E0AL76 |
| Oryza glumip | A0A0D9ZR97 |
| Oryza glumip | A0A0E0BFC3 |
| Oryza glumip | A0A0E0B678 |
| Oryza glumip | A0A0D9ZIN0 |
| Oryza glumip | A0A0D9ZFJ1 |

Oryza glumip A0A0D9ZRA5
Oryza glumip A0A0D9YFW4
Oryza glumip A0A0D9ZRA4
Oryza glumip A0A0D9ZR98
Oryza glumip A0A0D9ZRA7
Oryza glumip A0A0D9ZR81
Oryza glumip A0A0E0AIR6
Oryza glumip A0A0D9YA73
Oryza glumip A0A0E0BNH0
Oryza glumip A0A0D9YIC5
Oryza glumip A0A0E0BNG7
Oryza glumip A0A0D9ZG93
Oryza glumip A0A0D9ZZ82
Oryza glumip A0A0D9ZJF8
Oryza glumip A0A0E0A5J0
Oryza glumip A0A0D9Z9I2
Oryza glumip A0A0D9ZRS7
Oryza glumip A0A0D9Y5C9
Oryza glumip A0A0D9ZIM7
Oryza glumip A0A0E0AL74
Oryza glumip A0A0D9YC92
Oryza glumip A0A0D9Y4Z1
Oryza glumip A0A0D9YCY7
Oryza glumip A0A0D9ZRA6
Oryza glumip A0A0D9ZR95
Oryza glumip A0A0D9Y5E4
Oryza glumip A0A0D9Y5E6
Oryza glumip A0A0D9ZR83
Oryza glumip A0A0D9Y3F4
Oryza glumip A0A0E0BGJ9
Oryza glumip A0A0E0BR07
Oryza glumip A0A0D9YFX1
Oryza glumip A0A0D9ZR90
Oryza glumip A0A0D9YCY3
Oryza glumip A0A0D9YI02
Oryza glumip A0A0D9ZIN2
Oryza glumip A0A0E0B670
Oryza glumip A0A0E0B668
Oryza glumip A0A0D9ZR80
Oryza glumip A0A0E0BNG4
Oryza glumip A0A0D9YYI7

Oryza glumip A0A0E0A5J1
Oryza glumip A0A0E0BR04
Oryza glumip A0A0E0B667
Oryza glumip A0A0E0B671
Oryza glumip A0A0D9YI05
Oryza glumip A0A0D9YCY4
Oryza glumip A0A0D9ZUP5
Oryza glumip A0A0D9Y5D0
Oryza glumip A0A0D9ZR79
Oryza glumip A0A0D9YIG5
Oryza glumip A0A0E0ARL0
Oryza glumip A0A0E0ABJ9
Oryza glumip A0A0D9ZKP7
Oryza glumip A0A0E0BPR9
Oryza glumip A0A0D9Z920
Oryza glumip A0A0D9YIH3
Oryza glumip A0A0D9Y5E8
Oryza glumip A0A0E0B705
Oryza glumip A0A0D9Y5D3
Oryza glumip A0A0D9YFW1
Oryza glumip A0A0E0ARL1
Oryza glumip A0A0E0BVT0
Oryza glumip A0A0E0A194
Oryza glumip A0A0D9ZRA3
Oryza glumip A0A0D9YFW2
Oryza glumip A0A0D9ZUP4
Oryza glumip A0A0D9ZR96
Oryza glumip A0A0D9Y5D2
Oryza glumip A0A0E0B985
Oryza glumip A0A0E0B664
Oryza glumip A0A0D9ZRS6
Oryza glumip A0A0D9ZKN9
Oryza glumip A0A0E0B982
Oryza glumip A0A0D9YT30
Oryza glumip A0A0D9YCY1
Oryza glumip A0A0D9YFX0
Oryza glumip A0A0D9YIG8
Oryza glumip A0A0D9ZG92
Oryza glumip A0A0D9ZR82
Oryza glumip A0A0D9YFW8
Oryza glumip A0A0E0ARK8

| | |
|---|---|
| Oryza glumip | A0A0D9ZR78 |
| Oryza glumip | A0A0E0AIE6 |
| Oryza glumip | A0A0D9ZIM6 |
| Oryza glumip | A0A0D9YFW7 |
| Oryza glumip | A0A0D9ZKP8 |
| Oryza glumip | A0A0D9YI03 |
| Oryza glumip | A0A0E0BTG8 |
| Oryza glumip | A0A0D9ZRA2 |
| Oryza meridi | A0A0E0DCW6 |
| Oryza meridi | A0A0E0C8L4 |
| Oryza meridi | A0A0E0DFW2 |
| Oryza meridi | A0A0E0EUL2 |
| Oryza meridi | A0A0E0DE62 |
| Oryza meridi | A0A0E0CAW3 |
| Oryza meridi | A0A0E0DNX0 |
| Oryza meridi | A0A0E0EUM2 |
| Oryza meridi | A0A0E0CLE9 |
| Oryza meridi | A0A0E0EUK8 |
| Oryza meridi | A0A0E0BXB3 |
| Oryza meridi | A0A0E0DE59 |
| Oryza meridi | A0A0E0DC72 |
| Oryza meridi | A0A0E0EUK7 |
| Oryza meridi | A0A0E0DIT6 |
| Oryza meridi | A0A0E0C5U5 |
| Oryza meridi | A0A0E0ECX2 |
| Oryza meridi | A0A0E0DXL5 |
| Oryza meridi | A0A0E0DIT9 |
| Oryza meridi | A0A0E0DSY1 |
| Oryza meridi | A0A0E0EWS8 |
| Oryza meridi | A0A0E0EUL1 |
| Oryza meridi | A0A0E0FCQ8 |
| Oryza meridi | A0A0E0FA77 |
| Oryza meridi | A0A0E0EUL0 |
| Oryza meridi | A0A0E0EUM1 |
| Oryza meridi | A0A0E0C5U3 |
| Oryza meridi | A0A0E0CBD2 |
| Oryza meridi | A0A0E0DE49 |
| Oryza meridi | A0A0E0DE58 |
| Oryza meridi | A0A0E0CBC9 |
| Oryza meridi | A0A0E0E351 |
| Oryza meridi | A0A0E0CL32 |

| | |
|---|---|
| Oryza meridi | A0A0E0F268 |
| Oryza meridi | A0A0E0CAW4 |
| Oryza meridi | A0A0E0DC71 |
| Oryza meridi | A0A0E0C599 |
| Oryza meridi | A0A0E0DIU3 |
| Oryza meridi | A0A0E0E354 |
| Oryza meridi | A0A0E0DC70 |
| Oryza meridi | A0A0E0ECX0 |
| Oryza meridi | A0A0E0BZ76 |
| Oryza meridi | A0A0E0FAG5 |
| Oryza meridi | A0A0E0F2L6 |
| Oryza meridi | A0A0E0DE64 |
| Oryza meridi | A0A0E0C5U8 |
| Oryza meridi | A0A0E0DXL9 |
| Oryza meridi | A0A0E0F269 |
| Oryza meridi | A0A0E0CB81 |
| Oryza meridi | A0A0E0E2S8 |
| Oryza meridi | A0A0E0EWS6 |
| Oryza meridi | A0A0E0C5U2 |
| Oryza meridi | A0A0E0C5U7 |
| Oryza meridi | A0A0E0DML9 |
| Oryza meridi | A0A0E0DC67 |
| Oryza meridi | A0A0E0DCZ3 |
| Oryza meridi | A0A0E0E3L6 |
| Oryza meridi | A0A0E0DE54 |
| Oryza meridi | A0A0E0FA73 |
| Oryza meridi | A0A0E0D9W7 |
| Oryza meridi | A0A0E0BZ80 |
| Oryza meridi | A0A0E0DIS8 |
| Oryza meridi | A0A0E0FA68 |
| Oryza meridi | A0A0E0BZ68 |
| Oryza meridi | A0A0E0DC68 |
| Oryza meridi | A0A0E0C3C6 |
| Oryza meridi | A0A0E0D9B1 |
| Oryza meridi | A0A0E0DBN8 |
| Oryza meridi | A0A0E0ECW5 |
| Oryza meridi | A0A0E0EUK6 |
| Oryza meridi | A0A0E0D2Q4 |
| Oryza meridi | A0A0E0F3G5 |
| Oryza meridi | A0A0E0FAZ3 |
| Oryza meridi | A0A0E0DIU0 |

| | |
|---|---|
| Oryza meridi | A0A0E0DIS6 |
| Oryza meridi | A0A0E0DE56 |
| Oryza meridi | A0A0E0FA37 |
| Oryza meridi | A0A0E0DJC4 |
| Oryza meridi | A0A0E0FA69 |
| Oryza meridi | A0A0E0ECX4 |
| Oryza meridi | A0A0E0EIN0 |
| Oryza meridi | A0A0E0DIS7 |
| Oryza meridi | A0A0E0DE65 |
| Oryza meridi | A0A0E0DML8 |
| Oryza meridi | A0A0E0EA66 |
| Oryza meridi | A0A0E0BZ79 |
| Oryza meridi | A0A0E0FDQ6 |
| Oryza meridi | A0A0E0DIS5 |
| Oryza meridi | A0A0E0C8L7 |
| Oryza meridi | A0A0E0ECW7 |
| Oryza meridi | A0A0E0E264 |
| Oryza meridi | A0A0E0FDQ5 |
| Oryza meyeri | B9V0L1 |
| Oryza minuta | E0CW78 |
| Oryza nivara. | A0A0E0GKI4 |
| Oryza nivara. | A0A0E0FIN1 |
| Oryza nivara. | A0A0E0FIP8 |
| Oryza nivara. | A0A0U1WYS9 |
| Oryza nivara. | A0A0E0HGH4 |
| Oryza nivara. | A0A0E0J8N9 |
| Oryza nivara. | A0A0E0INR6 |
| Oryza nivara. | A0A0E0H0N8 |
| Oryza nivara. | A0A0E0FQS6 |
| Oryza nivara. | A0A0E0GKI6 |
| Oryza nivara. | A0A0E0GKI2 |
| Oryza nivara. | A0A0E0HXK8 |
| Oryza nivara. | A0A0E0J697 |
| Oryza nivara. | A0A0E0GX03 |
| Oryza nivara. | A0A0E0FQR8 |
| Oryza nivara. | A0A0E0HSU5 |
| Oryza nivara. | A0A0E0FQ49 |
| Oryza nivara. | A0A0E0H0P1 |
| Oryza nivara. | A0A0E0GX02 |
| Oryza nivara. | A0A0E0HJW4 |
| Oryza nivara. | A0A0E0HS89 |

Oryza nivara. A0A0E0GKI7
Oryza nivara. A0A0E0I7U3
Oryza nivara. A0A0E0H0P8
Oryza nivara. A0A0E0FGF6
Oryza nivara. A0A0E0J8N8
Oryza nivara. A0A0E0INB5
Oryza nivara. A0A0E0HUM6
Oryza nivara. A0A0E0H0N9
Oryza nivara. A0A0E0FWA9
Oryza nivara. A0A0E0GDA8
Oryza nivara. B9V0Q1
Oryza nivara. A0A0E0H2X3
Oryza nivara. A0A0E0HY24
Oryza nivara. A0A0E0G7M5
Oryza nivara. A0A0E0FIM7
Oryza nivara. A0A0E0FQR3
Oryza nivara. A0A0E0H6W8
Oryza nivara. A0A0E0GXL4
Oryza nivara. A0A0E0IXT2
Oryza nivara. A0A0E0FWU1
Oryza nivara. A0A0E0FQR6
Oryza nivara. A0A0E0H0P7
Oryza nivara. A0A0E0JB34
Oryza nivara. A0A0E0GW59
Oryza nivara. A0A0E0HGH5
Oryza nivara. A0A0E0IZH0
Oryza nivara. A0A0E0H0P6
Oryza nivara. A0A0E0INB6
Oryza nivara. A0A0E0FIN2
Oryza nivara. A0A0E0HR40
Oryza nivara. A0A0E0GKJ1
Oryza nivara. A0A0E0FIP9
Oryza nivara. A0A0E0INC5
Oryza nivara. A0A0E0H0N5
Oryza nivara. A0A0E0HY06
Oryza nivara. A0A0E0FQR7
Oryza nivara. A0A0E0GKJ2
Oryza nivara. A0A0E0GKH3
Oryza nivara. A0A0E0I273
Oryza nivara. A0A0E0INB4
Oryza nivara. A0A0E0IZH1

Oryza nivara. A0A0E0FIN0
Oryza nivara. A0A0E0HGG1
Oryza nivara. A0A0E0HS95
Oryza nivara. A0A0E0GKI5
Oryza nivara. A0A0E0H0P0
Oryza nivara. A0A0E0J2D2
Oryza nivara. A0A0E0GXK3
Oryza nivara. A0A0E0J696
Oryza nivara. A0A0E0I250
Oryza nivara. A0A0E0GKI3
Oryza nivara. A0A0E0FQS1
Oryza nivara. A0A0E0HLM1
Oryza nivara. A0A0E0GXR3
Oryza nivara. A0A0E0FWQ0
Oryza nivara. A0A0E0J8N4
Oryza nivara. A0A0E0HS91
Oryza nivara. A0A0E0HLM4
Oryza nivara. A0A0E0IZG9
Oryza nivara. A0A0E0I254
Oryza nivara. A0A0E0INC4
Oryza nivara. A0A0E0FU79
Oryza nivara. A0A0E0GYT0
Oryza nivara. A0A0E0J9E6
Oryza nivara. A0A0E0FLY3
Oryza nivara. A0A0E0FIP7
Oryza nivara. A0A0E0GPL9
Oryza nivara. A0A0E0HST7
Oryza nivara. A0A0E0J7G5
Oryza nivara. A0A0E0FI93
Oryza nivara. A0A0E0FWT5
Oryza nivara. A0A0E0I249
Oryza nivara. A0A0E0I0E2
Oryza nivara. A0A0E0H0N6
Oryza nivara. A0A0E0GP69
Oryza nivara. A0A0E0H0N7
Oryza nivara. A0A0E0IGG9
Oryza nivara. A0A0E0IXT5
Oryza nivara. A0A0E0IXT1
Oryza nivara. A0A0E0I252
Oryza nivara. A0A0E0FU81
Oryza nivara. A0A0E0GKJ0

Oryza nivara. A0A0E0JCW6
Oryza nivara. A0A0E0JB33
Oryza nivara. A0A0E0GP70
Oryza nivara. A0A0E0GKI9
Oryza nivara. A0A0E0HSU3
Oryza nivara. A0A0E0FIN9
Oryza nivara. A0A0E0J698
Oryza nivara. A0A0E0H6W7
Oryza nivara. A0A0E0FNQ7
Oryza nivara. A0A0E0GKJ3
Oryza nivara. A0A0E0FWT8
Oryza nivara. A0A0E0J2D0
Oryza nivara. A0A0E0HT73
Oryza nivara. A0A0E0HS88
Oryza nivara. A0A0E0FIN8
Oryza nivara. A0A0E0INB8
Oryza nivara. A0A0E0FU76
Oryza nivara. A0A0E0FIN7
Oryza nivara. B9V0Q0
Oryza nivara. A0A0E0FIN4
Oryza puncta A0A0E0KQI0
Oryza puncta A0A0E0KF64
Oryza puncta A0A0E0KVP1
Oryza puncta A0A0E0MCH2
Oryza puncta A0A0E0M481
Oryza puncta A0A0E0KM14
Oryza puncta A0A0E0LRX0
Oryza puncta A0A0E0JS11
Oryza puncta A0A0E0JFS6
Oryza puncta A0A0E0KQJ0
Oryza puncta A0A0E0KVP4
Oryza puncta A0A0E0M493
Oryza puncta B9V0S7
Oryza puncta A0A0E0KNA2
Oryza puncta A0A0E0LM11
Oryza puncta A0A0E0KVP5
Oryza puncta A0A0E0LRW9
Oryza puncta A0A0E0JPL8
Oryza puncta A0A0E0KYW2
Oryza puncta A0A0E0JFS5
Oryza puncta A0A0E0MC44

Oryza puncta A0A0E0JPM2
Oryza puncta A0A0E0LCW7
Oryza puncta A0A0E0JQG4
Oryza puncta A0A0E0LDK4
Oryza puncta A0A0E0MJ94
Oryza puncta A0A0E0KVP2
Oryza puncta A0A0E0JPL6
Oryza puncta A0A0E0LHZ7
Oryza puncta A0A0E0LBV1
Oryza puncta A0A0E0L4C9
Oryza puncta A0A0E0JPM1
Oryza puncta A0A0E0KQI6
Oryza puncta A0A0E0M482
Oryza puncta A0A0E0M4V9
Oryza puncta A0A0E0LZP4
Oryza puncta A0A0E0JFQ8
Oryza puncta A0A0E0LYC7
Oryza puncta A0A0E0MJ93
Oryza puncta A0A0E0MQ50
Oryza puncta A0A0E0M487
Oryza puncta A0A0E0LRW7
Oryza puncta A0A0E0KML7
Oryza puncta A0A0E0LM19
Oryza puncta A0A0E0KW47
Oryza puncta A0A0E0KVN9
Oryza puncta A0A0E0M6P1
Oryza puncta A0A0E0M6P2
Oryza puncta A0A0U1WXQ2
Oryza puncta A0A0E0LZM8
Oryza puncta A0A0E0KSK1
Oryza puncta A0A0E0KQI4
Oryza puncta A0A0E0JE78
Oryza puncta A0A0E0K0K1
Oryza puncta A0A0E0KAM8
Oryza puncta A0A0E0MKA3
Oryza puncta A0A0E0KQI5
Oryza puncta A0A0E0L2K7
Oryza puncta A0A0E0JFS2
Oryza puncta A0A0E0JRX9
Oryza puncta A0A0E0KFK8
Oryza puncta A0A0E0KVN4

Oryza puncta A0A0E0KQI8
Oryza puncta A0A0E0JM75
Oryza puncta A0A0E0LCW4
Oryza puncta A0A0E0KQI3
Oryza puncta A0A0E0LD93
Oryza puncta A0A0E0LCX3
Oryza puncta A0A0E0M492
Oryza puncta A0A0E0KNT6
Oryza puncta A0A0E0K5I5
Oryza puncta A0A0E0JLN9
Oryza puncta A0A0E0KVP3
Oryza puncta A0A0E0KRU6
Oryza puncta A0A0E0JM78
Oryza puncta A0A0E0KYV9
Oryza puncta A0A0E0KLA8
Oryza puncta A0A0E0JS10
Oryza puncta A0A0E0LM06
Oryza puncta A0A0E0MC39
Oryza puncta A0A0E0KNA3
Oryza puncta A0A0E0JRK2
Oryza puncta A0A0E0KYY4
Oryza puncta A0A0E0LM08
Oryza puncta A0A0E0JRK4
Oryza puncta A0A0E0JLN5
Oryza puncta A0A0E0L8B2
Oryza puncta A0A0E0MDQ4
Oryza puncta A0A0E0KVN6
Oryza puncta A0A0E0JM73
Oryza puncta A0A0E0KQI2
Oryza puncta A0A0E0JPL9
Oryza puncta A0A0E0JM82
Oryza puncta A0A0E0LYG7
Oryza puncta A0A0E0KNT9
Oryza puncta A0A0E0LET8
Oryza puncta A0A0E0JS09
Oryza puncta A0A0E0LZM7
Oryza puncta A0A0E0KNU2
Oryza puncta A0A0E0JS12
Oryza puncta A0A0E0LRW8
Oryza puncta A0A0E0JM79
Oryza puncta A0A0E0JRX7

| | |
|---|---|
| Oryza puncta | A0A0E0KYY5 |
| Oryza puncta | A0A0E0LZM9 |
| Oryza puncta | A0A0E0JTU7 |
| Oryza ridleyi. | E0CWC2 |
| Oryza rufipog | A0A0E0P929 |
| Oryza rufipog | A0A0E0QUS7 |
| Oryza rufipog | A0A0E0Q9Q2 |
| Oryza rufipog | A0A0E0R5M3 |
| Oryza rufipog | A0A0E0R5M2 |
| Oryza rufipog | A0A0E0N0I2 |
| Oryza rufipog | A0A0E0PPL5 |
| Oryza rufipog | A0A0E0N0I7 |
| Oryza rufipog | A0A0E0N0I1 |
| Oryza rufipog | A0A0E0PEJ2 |
| Oryza rufipog | A0A0E0QUS6 |
| Oryza rufipog | A0A0E0PEJ4 |
| Oryza rufipog | A0A0E0PY75 |
| Oryza rufipog | A0A0E0R492 |
| Oryza rufipog | A0A0E0N0I4 |
| Oryza rufipog | A0A0E0PI15 |
| Oryza rufipog | A0A0E0NWN9 |
| Oryza rufipog | A0A0E0QF46 |
| Oryza rufipog | A0A0E0R5M4 |
| Oryza rufipog | A0A0E0PTZ3 |
| Oryza rufipog | A0A0E0MRI6 |
| Oryza rufipog | A0A0E0PEI3 |
| Oryza rufipog | A0A0E0PYZ1 |
| Oryza rufipog | A0A0E0R5M1 |
| Oryza rufipog | A0A0E0PEJ6 |
| Oryza rufipog | A0A0E0P930 |
| Oryza rufipog | A0A0E0R3S3 |
| Oryza rufipog | A0A0E0RKL1 |
| Oryza rufipog | A0A0E0Q9N8 |
| Oryza rufipog | A0A0E0MYM4 |
| Oryza rufipog | A0A0E0RI41 |
| Oryza rufipog | A0A0E0N0H4 |
| Oryza rufipog | A0A0E0Q9P1 |
| Oryza rufipog | A0A0E0PZV4 |
| Oryza rufipog | A0A0E0PEJ5 |
| Oryza rufipog | A0A0E0PZG7 |
| Oryza rufipog | A0A0E0QF44 |

| | |
|---|---|
| Oryza rufipog | A0A0E0P3N3 |
| Oryza rufipog | A0A0E0P717 |
| Oryza rufipog | A0A0E0PZ15 |
| Oryza rufipog | A0A0E0Q9Q1 |
| Oryza rufipog | A0A0E0P724 |
| Oryza rufipog | A0A0E0P936 |
| Oryza rufipog | A0A0E0REI6 |
| Oryza rufipog | A0A0E0N628 |
| Oryza rufipog | A0A0E0NRR4 |
| Oryza rufipog | A0A0E0QUS8 |
| Oryza rufipog | A0A0E0MS90 |
| Oryza rufipog | A0A0E0QVH0 |
| Oryza rufipog | A0A0E0RDA8 |
| Oryza rufipog | A0A0E0P672 |
| Oryza rufipog | A0A0E0N0H5 |
| Oryza rufipog | A0A0E0PEI8 |
| Oryza rufipog | A0A0E0PEJ3 |
| Oryza rufipog | A0A0E0N633 |
| Oryza rufipog | A0A0E0MT99 |
| Oryza rufipog | A0A0E0QF47 |
| Oryza rufipog | A0A0E0QF43 |
| Oryza rufipog | A0A0E0PZG2 |
| Oryza rufipog | A0A0E0P571 |
| Oryza rufipog | A0A0E0PF17 |
| Oryza rufipog | A0A0E0MT98 |
| Oryza rufipog | A0A0E0PEI6 |
| Oryza rufipog | A0A0E0NL99 |
| Oryza rufipog | A0A0E0QXV0 |
| Oryza rufipog | A0A0E0MTA1 |
| Oryza rufipog | A0A0E0N631 |
| Oryza rufipog | A0A0E0N5M2 |
| Oryza rufipog | A0A0E0N3D9 |
| Oryza rufipog | A0A0E0Q505 |
| Oryza rufipog | A0A0E0PI17 |
| Oryza rufipog | A0A0E0P556 |
| Oryza rufipog | A0A0E0P7R6 |
| Oryza rufipog | A0A0E0Q6Z5 |
| Oryza rufipog | A0A0E0P4C6 |
| Oryza rufipog | A0A0E0PF18 |
| Oryza rufipog | A0A0E0QUT5 |
| Oryza rufipog | A0A0E0N3E2 |

| | |
|---|---|
| Oryza rufipog | A0A0E0P934 |
| Oryza rufipog | A0A0E0P723 |
| Oryza rufipog | A0A0E0MSU8 |
| Oryza rufipog | A0A0E0RDA9 |
| Oryza rufipog | A0A0E0N0J0 |
| Oryza rufipog | A0A0E0N5M3 |
| Oryza rufipog | A0A0E0QUT6 |
| Oryza rufipog | A0A0E0P935 |
| Oryza rufipog | A0A0E0NFS2 |
| Oryza rufipog | A0A0E0QXU8 |
| Oryza rufipog | A0A0E0MT87 |
| Oryza rufipog | A0A0E0P933 |
| Oryza rufipog | A0A0E0N5Z2 |
| Oryza rufipog | A0A0E0RFL8 |
| Oryza rufipog | A0A0E0PZG5 |
| Oryza rufipog | A0A0E0PEJ1 |
| Oryza rufipog | A0A0E0P726 |
| Oryza rufipog | A0A0E0PBB3 |
| Oryza rufipog | A0A0E0NX59 |
| Oryza rufipog | A0A0E0PTY5 |
| Oryza rufipog | A0A0E0Q9N9 |
| Oryza rufipog | A0A0E0N3D6 |
| Oryza sativa s | A2YMH7 |
| Oryza sativa s | B8AP81 |
| Oryza sativa s | B8A9Q7 |
| Oryza sativa s | B8B3W2 |
| Oryza sativa s | A2YDY8 |
| Oryza sativa s | A2XYE9 |
| Oryza sativa s | B8AUV8 |
| Oryza sativa s | B8BFX5 |
| Oryza sativa s | A2XPF2 |
| Oryza sativa s | B8B7X0 |
| Oryza sativa s | B8AUV4 |
| Oryza sativa s | B8AAG4 |
| Oryza sativa s | A2XRP3 |
| Oryza sativa s | B8B3K4 |
| Oryza sativa s | B8AAN9 |
| Oryza sativa s | B8A6L2 |
| Oryza sativa s | B8BIX3 |
| Oryza sativa s | B8B784 |
| Oryza sativa s | B8A6Z2 |

| | |
|---|---|
| Oryza sativa s | B8BE63 |
| Oryza sativa s | A2X9W0 |
| Oryza sativa s | B8BBD1 |
| Oryza sativa s | B8AYF8 |
| Oryza sativa s | A2Z666 |
| Oryza sativa s | A2Y9R2 |
| Oryza sativa s | B8A7D5 |
| Oryza sativa s | B8ASQ4 |
| Oryza sativa s | B8A779 |
| Oryza sativa s | A2ZIL9 |
| Oryza sativa s | A2XRP6 |
| Oryza sativa s | B8AUV7 |
| Oryza sativa s | A2YEC0 |
| Oryza sativa s | B8AAP0 |
| Oryza sativa s | A2WVT4 |
| Oryza sativa s | B8BM83 |
| Oryza sativa s | A2XTD8 |
| Oryza sativa s | B8B2E5 |
| Oryza sativa s | A2ZBG9 |
| Oryza sativa s | B8AUV2 |
| Oryza sativa s | B8AS67 |
| Oryza sativa s | B8BEE8 |
| Oryza sativa s | A2YMH8 |
| Oryza sativa s | A2Y2G0 |
| Oryza sativa s | A2YMG6 |
| Oryza sativa s | B8ASF5 |
| Oryza sativa s | A2ZCH9 |
| Oryza sativa s | A2XTE2 |
| Oryza sativa s | B8ATQ3 |
| Oryza sativa s | B8BE64 |
| Oryza sativa s | A2XTD2 |
| Oryza sativa s | B8ATQ2 |
| Oryza sativa s | B8AS68 |
| Oryza sativa s | B8ATQ5 |
| Oryza sativa s | B8A780 |
| Oryza sativa s | A2WM87 |
| Oryza sativa s | B8BLW7 |
| Oryza sativa s | B8AK50 |
| Oryza sativa s | A2YMG7 |
| Oryza sativa s | A2XTE5 |
| Oryza sativa s | B8AUZ5 |

| | |
|---|---|
| Oryza sativa s | A2WTM1 |
| Oryza sativa s | B8AUV3 |
| Oryza sativa s | A2Z664 |
| Oryza sativa s | A2Z3T6 |
| Oryza sativa s | A2WXE7 |
| Oryza sativa s | B8AB24 |
| Oryza sativa s | A2YER5 |
| Oryza sativa s | B8AAN5 |
| Oryza sativa s | A2WXT2 |
| Oryza sativa s | B8AUW7 |
| Oryza sativa s | B8BE67 |
| Oryza sativa s | A2WVT7 |
| Oryza sativa s | A2XVC1 |
| Oryza sativa s | A2YTP0 |
| Oryza sativa s | B8BLW6 |
| Oryza sativa s | B8A783 |
| Oryza sativa s | A2XTD1 |
| Oryza sativa s | A2XTD0 |
| Oryza sativa s | B9V0N4 |
| Oryza sativa s | B8ATQ4 |
| Oryza sativa s | A2ZB52 |
| Oryza sativa s | B8BEE9 |
| Oryza sativa s | A5JS87 |
| Oryza sativa s | A2ZB38 |
| Oryza sativa s | A2WTM3 |
| Oryza sativa s | A2Y9Q7 |
| Oryza sativa s | B8AZQ7 |
| Oryza sativa s | B8A782 |
| Oryza sativa s | B8AUW5 |
| Oryza sativa s | A2XE17 |
| Oryza sativa s | A2ZN71 |
| Oryza sativa s | A2WRY3 |
| Oryza sativa s | A2WKV0 |
| Oryza sativa s | A2XTD4 |
| Oryza sativa s | B8A7B9 |
| Oryza sativa s | A2Y0R8 |
| Oryza sativa s | B8AUV5 |
| Oryza sativa s | B8ATY7 |
| Oryza sativa s | A2XII0 |
| Oryza sativa s | A2XTE4 |
| Oryza sativa s | A2X5J7 |

| | |
|---|---|
| Oryza sativa s | A2Y0Q5 |
| Oryza sativa s | B8BE62 |
| Oryza sativa s | B8AVL9 |
| Oryza sativa s | A2ZB35 |
| Oryza sativa s | A2WT48 |
| Oryza sativa s | A2XTE3 |
| Oryza sativa s | B8AUV6 |
| Oryza sativa s | A2XNU0 |
| Oryza sativa s | A2XXZ2 |
| Oryza sativa s | B8AUW6 |
| Oryza sativa s | B8BE66 |
| Oryza sativa s | B8AUV1 |
| Oryza sativa s | B8B3W0 |
| Oryza sativa s | B8BP34 |
| Oryza sativa s | A2ZHT0 |
| Oryza sativa s | A2ZCI0 |
| Oryza sativa s | Q0JDY0 |
| Oryza sativa s | Q69MP3 |
| Oryza sativa s | B9FEC4 |
| Oryza sativa s | B9FBW5 |
| Oryza sativa s | B9EYI9 |
| Oryza sativa s | Q0JD96 |
| Oryza sativa s | B9FZD1 |
| Oryza sativa s | B9FCW6 |
| Oryza sativa s | B9GBP9 |
| Oryza sativa s | Q5N835 |
| Oryza sativa s | A0A0P0Y6H8 |
| Oryza sativa s | B9G4Y0 |
| Oryza sativa s | B9FMN5 |
| Oryza sativa s | A0A0P0WFW5 |
| Oryza sativa s | B9ETD6 |
| Oryza sativa s | Q60D67 |
| Oryza sativa s | A3CFG6 |
| Oryza sativa s | A0A0P0UZX5 |
| Oryza sativa s | A0A0P0XQJ7 |
| Oryza sativa s | Q0J9S4 |
| Oryza sativa s | A0A0P0WIC4 |
| Oryza sativa s | Q5Z607 |
| Oryza sativa s | Q0J9S9 |
| Oryza sativa s | A0A0P0WSZ1 |
| Oryza sativa s | Q2R920 |

| | |
|---|---|
| Oryza sativa s | A0A0N7KR84 |
| Oryza sativa s | A0A0P0XQY3 |
| Oryza sativa s | A0A0P0V6A7 |
| Oryza sativa s | Q5NB60 |
| Oryza sativa s | Q0J9S8 |
| Oryza sativa s | Q5QNL1 |
| Oryza sativa s | Q69MN9 |
| Oryza sativa s | A0A0N7KJ20 |
| Oryza sativa s | A3C195 |
| Oryza sativa s | Q7X6C7 |
| Oryza sativa s | Q0JEB3 |
| Oryza sativa s | A0A0P0Y0I8 |
| Oryza sativa s | B9FCQ2 |
| Oryza sativa s | Q2QY66 |
| Oryza sativa s | B9FMM3 |
| Oryza sativa s | B9FEC2 |
| Oryza sativa s | B9FCW7 |
| Oryza sativa s | A0A0P0XQ09 |
| Oryza sativa s | A0A0P0XQ04 |
| Oryza sativa s | A0A0P0WXZ2 |
| Oryza sativa s | B9FCQ0 |
| Oryza sativa s | C7J132 |
| Oryza sativa s | A0A0P0XRC3 |
| Oryza sativa s | Q7XQT3 |
| Oryza sativa s | Q0JH14 |
| Oryza sativa s | A3BL04 |
| Oryza sativa s | C7J1M6 |
| Oryza sativa s | Q2RAY8 |
| Oryza sativa s | Q5QLM8 |
| Oryza sativa s | A3AVE3 |
| Oryza sativa s | A3BCZ7 |
| Oryza sativa s | A0A0P0Y6V4 |
| Oryza sativa s | A3AXR8 |
| Oryza sativa s | Q5JQW1 |
| Oryza sativa s | A0A0P0WF90 |
| Oryza sativa s | A0A0P0WF87 |
| Oryza sativa s | B9F816 |
| Oryza sativa s | B9FCQ5 |
| Oryza sativa s | A0A0P0XYU6 |
| Oryza sativa s | C7J3E6 |
| Oryza sativa s | B9FXT7 |

| | |
|---|---|
| Oryza sativa s | Q0IZT3 |
| Oryza sativa s | A0A0P0Y7N1 |
| Oryza sativa s | A0A0P0X5S1 |
| Oryza sativa s | A0A0P0VAV4 |
| Oryza sativa s | A0A0P0WG14 |
| Oryza sativa s | A0A0P0XQ32 |
| Oryza sativa s | A0A0P0WIA4 |
| Oryza sativa s | A0A0P0WAJ2 |
| Oryza sativa s | A2ZYH9 |
| Oryza sativa s | Q0J9T9 |
| Oryza sativa s | Q0JEB1 |
| Oryza sativa s | Q2QWY4 |
| Oryza sativa s | Q7XL24 |
| Oryza sativa s | B9FEC1 |
| Oryza sativa s | B9GE14 |
| Oryza sativa s | A3AS58 |
| Oryza sativa s | A0A0P0VBI4 |
| Oryza sativa s | B9G4X2 |
| Oryza sativa s | A0A0P0V6F2 |
| Oryza sativa s | A0A0P0W6Q5 |
| Oryza sativa s | Q0J9T4 |
| Oryza sativa s | B9FCP9 |
| Oryza sativa s | A0A0P0XYD6 |
| Oryza sativa s | B9EUW0 |
| Oryza sativa s | Q0J9S6 |
| Oryza sativa s | Q8H353 |
| Oryza sativa s | A0A0P0V6E9 |
| Oryza sativa s | B9FCP8 |
| Oryza sativa s | Q7XQU2 |
| Oryza sativa s | A0A0P0VBE8 |
| Oryza sativa s | B9EU56 |
| Oryza sativa s | A0A0N7KHG5 |
| Oryza sativa s | Q7XMH7 |
| Oryza sativa s | B9F999 |
| Oryza sativa s | Q0J9T3 |
| Oryza sativa s | Q0ITY0 |
| Oryza sativa s | B9G985 |
| Oryza sativa s | Q0E0R8 |
| Oryza sativa s | Q7XL27 |
| Oryza sativa s | B9FCQ4 |
| Oryza sativa s | B9EU61 |

| | |
|---|---|
| Oryza sativa s | A3A0E3 |
| Oryza sativa s | B9EUW2 |
| Oryza sativa s | B9EVA3 |
| Oryza sativa s | A0A0P0Y700 |
| Oryza sativa s | A0A0P0XPZ9 |
| Oryza sativa s | A0A0P0W9H2 |
| Oryza sativa s | A0A0P0WF77 |
| Oryza sativa s | A0A0P0WF71 |
| Oryza sativa s | A0A0N7KDE3 |
| Oryza sativa s | A0A0N7KJS0 |
| Oryza sativa s | Q0JEP1 |
| Oryza sativa s | Q7XQU6 |
| Oryza sativa s | B9FKZ6 |
| Oryza sativa s | A0A0P0X7Q1 |
| Oryza sativa s | A0A0P0WAE2 |
| Oryza sativa s | A0A0P0XRC1 |
| Oryza sativa s | B9EYJ0 |
| Oryza sativa s | B9FCQ1 |
| Oryza sativa s | B9ETD5 |
| Oryza sativa s | B9GCM6 |
| Oryza sativa s | A3B8R2 |
| Oryza sativa s | Q6ATL4 |
| Oryza sativa s | Q7XQU0 |
| Oryza sativa s | A3C9L3 |
| Oryza sativa s | B9FD08 |
| Oryza sativa s | B9G4X9 |
| Oryza sativa s | A3A014 |
| Oryza sativa s | B9FCQ3 |
| Oryza sativa s | B9FMN4 |
| Oryza sativa s | B9G4X5 |
| Oryza sativa s | A3CEN4 |
| Oryza sativa s | B9FF59 |
| Oryza sativa s | A0A0P0WFH6 |
| Oryza sativa s | Q7XJ40 |
| Oryza sativa s | A0A0P0W6K3 |
| Oryza sativa s | A0A0P0WAN9 |
| Oryza sativa s | B9FXT6 |
| Oryza sativa s | B9FF58 |
| Oryza sativa s | A0A0P0WF66 |
| Oryza sativa s | Q7XNW1 |
| Oryza sativa s | A0A0P0WF95 |

| | |
|---|---|
| Oryza sativa s | A3A0E0 |
| Oryza sativa s | Q7XQT7 |
| Oryza sativa s | Q8W054 |
| Oryza sativa s | Q69MN2 |
| Oryza sativa s | Q7XI25 |
| Oryza sativa s | Q5NAQ9 |
| Oryza sativa s | Q8LMB0 |
| Oryza sativa s | Q2QPI9 |
| Oryza sativa s | Q7XHV3 |
| Oryza sativa s | Q2R917 |
| Oryza sativa s | Q5QLR7 |
| Oryza sativa s | Q5QLN0 |
| Oryza sativa s | Q9LE23 |
| Oryza sativa s | Q5ZAL3 |
| Oryza sativa s | Q0JH38 |
| Oryza sativa s | Q53PX8 |
| Oryza sativa s | Q0IUV4 |
| Oryza sativa s | Q8W3C9 |
| Oryza sativa s | Q5N9Y5 |
| Oryza sativa s | Q7XMR2 |
| Oryza sativa s | Q0JKJ3 |
| Oryza sativa s | Q7XXJ5 |
| Oryza sativa s | Q5ZAK8 |
| Oryza sativa s | Q7XQU3 |
| Oryza sativa s | Q2QLI8 |
| Oryza sativa s | Q851M7 |
| Oryza sativa s | Q60D69 |
| Oryza sativa s | Q7EZ32 |
| Oryza sativa s | Q60EJ2 |
| Oryza sativa s | Q5ZBQ1 |
| Oryza sativa s | Q8H821 |
| Oryza sativa s | Q6ZF83 |
| Oryza sativa s | Q8S0R2 |
| Oryza sativa s | Q7XQT8 |
| Oryza sativa s | Q5Z604 |
| Oryza sativa s | Q7XTD6 |
| Oryza sativa s | Q7X6I1 |
| Oryza sativa s | Q2RAY6 |
| Oryza sativa s | Q5VRR9 |
| Oryza sativa s | Q84MF7 |
| Oryza sativa s | Q6Z6H3 |

| | |
|---|---|
| Oryza sativa s | Q5Z601 |
| Oryza sativa s | Q7XTD7 |
| Oryza sativa s | Q9LDT8 |
| Oryza sativa s | Q2RAY2 |
| Oryza sativa s | Q7XMH4 |
| Oryza sativa s | Q5Z932 |
| Oryza sativa s | Q69XK0 |
| Oryza sativa s | Q2QUP5 |
| Oryza sativa s | Q0J9T7 |
| Oryza sativa s | Q69MM4 |
| Oryza sativa s | Q0JD95 |
| Oryza sativa s | Q2QY70 |
| Oryza sativa s | Q5SMQ2 |
| Oryza sativa s | Q2QY68 |
| Oryza sativa s | Q7XMG8 |
| Oryza sativa s | Q7XMG6 |
| Oryza sativa s | Q2RAY4 |
| Oryza sativa s | Q2R918 |
| Oryza sativa s | Q0D733 |
| Oryza sativa s | Q10JE9 |
| Oryza sativa s | Q5NB61 |
| Oryza sativa s | Q6H6Q2 |
| Oryza sativa s | Q8LMB2 |
| Oryza sativa s | Q7XQT9 |
| Oryza sativa s | B7F353 |
| Oryza sativa s | Q7XMH8 |
| Oryza sativa s | Q7XMG5 |
| Oryza sativa s | Q7XMC0 |
| Oryza sativa s | Q5QLM9 |
| Oryza sativa s | Q0J9T2 |
| Oryza sativa s | Q7EZ33 |
| Oryza sativa s | A2ZWE1 |
| Oryza sativa s | Q8LQN6 |
| Oryza sativa s | Q8W059 |
| Oryza sativa s | Q8LQH1 |
| Oryza sativa s | B9FEC3 |
| Oryza sativa s | Q60D78 |
| Oryza sativa s | Q60D68 |
| Oryza sativa s | Q0JIQ1 |
| Oryza sativa s | Q5ZAV5 |
| Oryza sativa s | B9FJP0 |

| | |
|---|---|
| Oryza sativa s | Q7Y1P1 |
| Oryza sativa s | Q2QY71 |
| Oryza sativa s | Q943Q1 |
| Oryza sativa s | Q7XMH0 |
| Oryza sativa s | Q7XUM4 |
| Oryza sativa s | Q7XQX5 |
| Oryza sativa s | Q69MM9 |
| Oryza sativa s | Q0JPG7 |
| Oryza sativa s | Q7XSF1 |
| Oryza sativa s | Q7XMH5 |
| Oryza sativa s | Q7XMH1 |
| Oryza sativa.) | Q25A54 |
| Oryza sativa.) | Q01KU2 |
| Oryza sativa.) | Q01J05 |
| Oryza sativa.) | Q25A55 |
| Oryza sativa.) | Q8LLI2 |
| Oryza sativa.) | Q25A21 |
| Oryza sativa.) | Q01MD3 |
| Oryza sativa.) | Q8LLI3 |
| Oryza sativa.) | Q01K00 |
| Oryza sativa.) | Q25AB4 |
| Oryza sativa.) | Q01KM2 |
| Oryza sativa.) | Q01MD6 |
| Oryza sativa.) | Q25A60 |
| Oryza sativa.) | Q8LLI4 |
| Oryza sativa.) | Q01K01 |
| Oryza sativa.) | Q01K07 |
| Oryza sativa.) | Q9LLN5 |
| Oryza sativa.) | Q01MD7 |
| Oryza sativa.) | Q01LZ8 |
| Oryza sativa.) | Q01N33 |
| Oryza sativa.) | Q01K06 |
| Oryza sativa.) | Q01N32 |
| Oryza sativa.) | Q25A58 |
| Oryza sativa.) | Q25A57 |
| Oryza sativa.) | Q01MD8 |
| Oryza sativa.) | Q01K02 |
| Oryza sativa.) | K0JBC6 |
| Oryza sativa.) | Q25AB6 |
| Oryza sativa.) | Q01M86 |
| Oryza sativa.) | Q8LLI1 |

| | |
|---|---|
| Oryza sativa.: | Q25A00 |
| Oryza sativa.: | Q01M00 |
| Oryza sativa.: | Q259Z7 |
| Oryza sativa.: | Q25A01 |
| Oryzias javan | A0A437CU31 |
| Oryzias javan | A0A3S2U5L5 |
| Oryzias javan | A0A3S2MCZ5 |
| Oryzias javan | A0A437DAI2 |
| Oryzias javan | A0A3S2MFY0 |
| Oryzias javan | A0A437DEQ2 |
| Oryzias latipe | A0A3P9ICR8 |
| Oryzias latipe | Q50LG6 |
| Oryzias latipe | A0A3B3I1H2 |
| Oryzias latipe | A0A3P9MP48 |
| Oryzias latipe | A0A3P9K011 |
| Oryzias latipe | A0A3P9I2I7 |
| Oryzias latipe | H2LF02 |
| Oryzias latipe | A0A3P9JAV4 |
| Oryzias latipe | A0A3P9LI60 |
| Oryzias latipe | A0A3P9J614 |
| Oryzias latipe | H2MEZ4 |
| Oryzias latipe | A0A3P9MAE3 |
| Oryzias latipe | A0A3P9LHP9 |
| Oryzias latipe | H2MEU2 |
| Oryzias latipe | H2M0V9 |
| Oryzias latipe | A0A3P9HMN6 |
| Oryzias latipe | A0A3P9K000 |
| Oryzias latipe | A0A3P9JJS7 |
| Oryzias latipe | A0A3P9LIA1 |
| Oryzias mela: | A0A3B3D7P0 |
| Oryzias mela: | I1SRG1 |
| Oryzias mela: | A0A3B3D5Q6 |
| Oryzias mela: | A0A3B3BZR1 |
| Oryzias mela: | A0A3B3CM08 |
| Oryzias mela: | I1SRH8 |
| Oryzias mela: | A0A3B3DHX8 |
| Osmerus moı | C1BLZ6 |
| Ostreococcus | A4S6G3 |
| Ostreococcus | A0A1Y5IBM2 |
| Ostreococcus | A0A096P7Z7 |
| Ostreococcus | A0A090N4R5 |

| | |
|---|---|
| Ostreococcus | A0A1Y5I2L6 |
| Ostreococcus | A0A1Y5IDE4 |
| Ostreococcus | A0A1Y5I7I8 |
| Ostreococcus | A0A090LXV4 |
| Ostreococcus | A0A096P9I0 |
| Ostreococcus | A0A1Y5I593 |
| Ostreococcus | A0A096P7B4 |
| Ostreococcus | A0A090MCI5 |
| Otolemur gar | H0XRY6 |
| Otolemur gar | H0WTA2 |
| Otolemur gar | H0WJV6 |
| Otolemur gar | H0X5J0 |
| Otolemur gar | H0WJV3 |
| Ovis aries.xls: | W5PEM0 |
| Ovis aries.xls: | W5P3R3 |
| Ovis aries.xls: | W5PEI4 |
| Ovis aries.xls: | W5PUG1 |
| Ovis aries.xls: | W5Q5H9 |
| Pacmanvirus | A0A1X6WFX5 |
| Pacmanvirus | A0A1X6WF83 |
| Paenirhodob: | A0A086XR69 |
| Pan paniscus. | A0A2R9CEQ2 |
| Pan paniscus. | A0A2R9B5R5 |
| Pan paniscus. | A0A2R9CAN4 |
| Pan paniscus. | A0A2R9B8J6 |
| Pan paniscus. | A0A2R9B1Y7 |
| Pan paniscus. | A0A2R9CER0 |
| Pan paniscus. | A0A2R9CES0 |
| Pan paniscus. | A0A2R9CL99 |
| Pan paniscus. | A0A2R9BBJ0 |
| Pan troglodyt | A0A2J8L7X0 |
| Pan troglodyt | A0A2J8JZZ2 |
| Pan troglodyt | A0A2J8L7Y1 |
| Pan troglodyt | A0A2J8L7X6 |
| Pan troglodyt | A0A2J8L807 |
| Pan troglodyt | A0A2I3RJ89 |
| Pan troglodyt | A0A2I3S7A8 |
| Pan troglodyt | A0A2J8L7W7 |
| Pan troglodyt | A0A2J8L7X2 |
| Pan troglodyt | A0A2J8LGB0 |
| Pan troglodyt | A0A2I3S7L0 |

| | |
|---|---|
| Pan troglodyt | A0A2I3SM29 |
| Pan troglodyt | A0A2J8JZY8 |
| Pan troglodyt | H2QIA7 |
| Pan troglodyt | A0A2J8K5V2 |
| Pan troglodyt | A0A2J8LGB6 |
| Pan troglodyt | A0A2J8LGB4 |
| Pan troglodyt | A0A2I3S5X5 |
| Pan troglodyt | K7CWX4 |
| Pan troglodyt | A0A2J8L7V9 |
| Pan troglodyt | H2QUU8 |
| Pan troglodyt | A0A2J8K020 |
| Pan troglodyt | H2QU06 |
| Pan troglodyt | A0A2I3TJS6 |
| Pan troglodyt | A0A2I3T423 |
| Pan troglodyt | A0A2I3REN2 |
| Pan troglodyt | H2QQJ4 |
| Pan troglodyt | A0A2I3TKS5 |
| Pan troglodyt | A0A2I3SRY8 |
| Pan troglodyt | K7BAC9 |
| Pan troglodyt | K7AMD5 |
| Pan troglodyt | A0A2J8K5U6 |
| Pan troglodyt | H2R8A3 |
| Panaeolus cy | A0A409YP06 |
| Panaeolus cy | A0A409YU19 |
| Panicum halli | A0A2T7D0I5 |
| Panicum halli | A0A2T7EVK0 |
| Panicum halli | A0A2T7EDS2 |
| Panicum halli | A0A2T7E6P0 |
| Panicum halli | A0A2T7DT33 |
| Panicum halli | A0A2T7CS41 |
| Panicum halli | A0A2T7CTI1 |
| Panicum halli | A0A2T7E553 |
| Panicum halli | A0A2T7E643 |
| Panicum halli | A0A2T7ET21 |
| Panicum halli | A0A2T7ER75 |
| Panicum halli | A0A2T7DIN6 |
| Panicum halli | A0A2T7DDD3 |
| Panicum halli | A0A2T7CBD2 |
| Panicum halli | A0A2T7EVJ0 |
| Panicum halli | A0A2T7D0G2 |
| Panicum halli | A0A2T7CT36 |

Panicum halli A0A2T7E6C1
Panicum halli A0A2T7C4S3
Panicum halli A0A2T7DFU5
Panicum halli A0A2T7DFU8
Panicum halli A0A2T7CMR2
Panicum halli A0A2T7EF41
Panicum halli A0A2T7EWK1
Panicum halli A0A2T7BYK9
Panicum halli A0A2T7DLQ8
Panicum halli A0A2T7DJ77
Panicum halli A0A2T7D0E9
Panicum halli A0A2T7C3A3
Panicum halli A0A2T7DDF0
Panicum halli A0A2T7EW50
Panicum halli A0A2T7CBC7
Panicum halli A0A2T7CL36
Panicum halli A0A2T7EX72
Panicum halli A0A2T7EIN3
Panicum halli A0A2T7D0E7
Panicum halli A0A2T7D509
Panicum halli A0A2T7CFW6
Panicum halli A0A2T7DT37
Panicum halli A0A2T7CUW6
Panicum halli A0A2T7D0D5
Panicum halli A0A2T7DLD3
Panicum halli A0A2T7F513
Panicum halli A0A2T7E7E6
Panicum halli A0A2T7F551
Panicum halli A0A2T7F5F3
Panicum halli A0A2T7D0D9
Panicum halli A0A2T7F5F1
Panicum halli A0A2T7CUW0
Panicum halli A0A2T7D494
Panicum halli A0A2T7EBW9
Panicum halli A0A2T7D0E0
Panicum halli A0A2T7E7B7
Panicum halli A0A2T7DFW0
Panicum halli A0A2T7F436
Panicum halli A0A2T7EVJ2
Panicum halli A0A2T7CUW5
Panicum halli A0A2T7CUZ3

Panicum halli A0A2T7CMN6
Panicum halli A0A2T7E7C7
Panicum halli A0A2T7CB30
Panicum halli A0A2T7EX73
Panicum halli A0A2T7EX64
Panicum halli A0A2T7ER93
Panicum halli A0A2T7D0F4
Panicum halli A0A2T7EVJ5
Panicum halli A0A2T7ERX6
Panicum halli A0A2T7EX66
Panicum halli A0A2T7DDD0
Panicum halli A0A2T7D0E4
Panicum halli A0A2T7CBK9
Panicum halli A0A2T7E0A7
Panicum halli A0A2T7D0F3
Panicum halli A0A2T7D0D8
Panicum halli A0A2T7DV28
Panicum halli A0A2T7DT28
Panicum halli A0A2T7DIM9
Panicum halli A0A2T7DX83
Panicum halli A0A2T7DIM2
Panicum halli A0A2T7F574
Panicum halli A0A2T7F5H7
Panicum halli A0A2T7D0J0
Panicum halli A0A2T7DFV2
Panicum halli A0A2T7EVK4
Panicum halli A0A2T7CUW4
Panicum halli A0A2T7DX94
Panicum halli A0A2T7D0Q9
Panicum halli A0A2T7EVI9
Panicum halli A0A2T7EJU9
Panicum halli A0A2T7EBW4
Panicum halli A0A2T7CB26
Panicum halli A0A2T7EBW3
Panicum halli A0A2T7EVI4
Panicum halli A0A2T7DFU4
Panicum halli A0A2T7C4G3
Panicum halli A0A2T7DJ67
Panicum halli A0A2T7D0F2
Panicum halli A0A2T7DFW9
Panicum halli A0A2T7E6H9

Panicum halli A0A2T7DLC8
Panicum halli A0A2T7BYV2
Panicum halli A0A2T7ENB4
Panicum halli A0A2T7CUV7
Panicum halli A0A2T7D0Y3
Panicum halli A0A2T7DLE0
Panicum halli A0A2T7CUW3
Panicum halli A0A2T7DRA7
Panicum halli A0A2T7DDA9
Panicum halli A0A2T7D0K1
Panicum halli A0A2T7DLD0
Panicum halli A0A2T7D0Q0
Panicum halli A0A2T7EBV8
Panicum halli A0A2T7D0H3
Panicum halli A0A2T7E7B6
Panicum halli A0A270R6Y3
Panicum halli A0A2S3HXW3
Panicum halli A0A2T8IPC6
Panicum halli A0A2T8KJ78
Panicum halli A0A2T8ICA6
Panicum halli A0A2S3HTA8
Panicum halli A0A2S3IAP2
Panicum halli A0A2T8I4D3
Panicum halli A0A2T8KV97
Panicum halli A0A2S3H0F9
Panicum halli A0A2S3IAP8
Panicum halli A0A2S3H1N8
Panicum halli A0A2S3HIG5
Panicum halli A0A2S3IAR5
Panicum halli A0A2T8KQB7
Panicum halli A0A2S3I5P1
Panicum halli A0A2T8IE12
Panicum halli A0A2T8KRJ5
Panicum halli A0A2S3H2E7
Panicum halli A0A2S3HGY6
Panicum halli A0A2S3IKM6
Panicum halli A0A2S3IAM3
Panicum halli A0A2S3H186
Panicum halli A0A2S3I0N4
Panicum halli A0A2T8I4D7
Panicum halli A0A2S3H205

Panicum halli A0A2S3H7F5
Panicum halli A0A2T8IJZ0
Panicum halli A0A2S3H1U7
Panicum halli A0A2T8KUQ8
Panicum halli A0A2T8IC99
Panicum halli A0A2T8I0T0
Panicum halli A0A270R6C5
Panicum halli A0A2T8IE65
Panicum halli A0A2S3HTE7
Panicum halli A0A2T8KRK0
Panicum halli A0A2T8IE16
Panicum halli A0A2S3HAQ3
Panicum halli A0A2S3IGG3
Panicum halli A0A2S3HPQ6
Panicum halli A0A2S3HR11
Panicum halli A0A2T8IDZ9
Panicum halli A0A2S3ID39
Panicum halli A0A2T8IE34
Panicum halli A0A2S3H3A8
Panicum halli A0A2S3GXJ4
Panicum halli A0A2S3H396
Panicum halli A0A2T8KM77
Panicum halli A0A2S3HUX0
Panicum halli A0A2S3IAS7
Panicum halli A0A2S3H762
Panicum halli A0A2T8IE17
Panicum halli A0A2T8IE14
Panicum halli A0A2S3IAP0
Panicum halli A0A2T8KS60
Panicum halli A0A2S3IS50
Panicum halli A0A2S3HP70
Panicum halli A0A2S3HK05
Panicum halli A0A2S3IQN4
Panicum halli A0A2S3IPM5
Panicum halli A0A2S3HAQ6
Panicum halli A0A2S3HR06
Panicum halli A0A2T8KNP2
Panicum halli A0A2T8KRH6
Panicum halli A0A2S3IGI2
Panicum halli A0A2S3HR12
Panicum halli A0A2S3HD01

| | |
|---|---|
| Panicum halli | A0A2S3H5Z0 |
| Panicum halli | A0A2S3IE86 |
| Panicum halli | A0A2S3HT43 |
| Panicum halli | A0A2S3IQM7 |
| Panicum halli | A0A2T8IJ38 |
| Panicum halli | A0A2S3I1E5 |
| Panicum halli | A0A2S3IB91 |
| Panicum halli | A0A2T8JCF9 |
| Panicum halli | A0A2S3IAQ3 |
| Panicum halli | A0A2T8IE28 |
| Panicum halli | A0A2S3I6M3 |
| Panicum halli | A0A2T8KV35 |
| Panicum halli | A0A2S3IBD2 |
| Panicum halli | A0A2S3IPQ8 |
| Panicum halli | A0A2T8KHH4 |
| Panicum halli | A0A2S3H6Z9 |
| Panicum halli | A0A2S3IAP1 |
| Panicum halli | A0A2S3HV18 |
| Panicum halli | A0A2S3HBZ4 |
| Panicum halli | A0A2S3HRI0 |
| Panicum halli | A0A2S3I6M7 |
| Panicum halli | A0A2S3GP95 |
| Panicum halli | A0A2S3HUW3 |
| Panicum halli | A0A2T8KRL1 |
| Panicum halli | A0A2S3HJZ4 |
| Panicum halli | A0A2S3HP69 |
| Panicum halli | A0A2T8I1B4 |
| Panicum halli | A0A2S3IAN4 |
| Panicum halli | A0A2T8IAS4 |
| Panicum halli | A0A2T8I1S3 |
| Panicum halli | A0A2T8IDZ2 |
| Panicum halli | A0A2S3H370 |
| Panicum halli | A0A2S3HW47 |
| Panicum halli | A0A2S3IAP4 |
| Panicum halli | A0A2T8IE05 |
| Panicum halli | A0A2T8KRJ0 |
| Panicum halli | A0A2S3HGB8 |
| Panicum halli | A0A2S3HZ22 |
| Panicum halli | A0A2S3GNM4 |
| Panicum halli | A0A2S3HFW8 |
| Panicum halli | A0A2S3H7E9 |

| | |
|---|---|
| Panicum halli | A0A2S3HIN5 |
| Panicum halli | A0A2T8KLZ9 |
| Panicum halli | A0A2S3I6H4 |
| Panicum halli | A0A2S3IAN6 |
| Panicum halli | A0A2T8I8I3 |
| Panicum halli | A0A2T8IBV6 |
| Panicum halli | A0A2S3I506 |
| Panicum halli | A0A2T8IJW5 |
| Panicum halli | A0A2S3IAP7 |
| Panicum halli | A0A2S3HAP8 |
| Panicum halli | A0A2T8KHP9 |
| Panicum halli | A0A2S3I6I3 |
| Panicum halli | A0A2S3GTT4 |
| Panicum halli | A0A2T8I1Q5 |
| Panicum halli | A0A2S3IST7 |
| Panicum halli | A0A2T8KVD3 |
| Panicum halli | A0A2T8IE22 |
| Panicum halli | A0A2T8IJV0 |
| Panicum halli | A0A2T8IE03 |
| Panicum halli | A0A2S3GYI1 |
| Panicum halli | A0A2S3IAT7 |
| Panicum halli | A0A2S3HZR6 |
| Panicum halli | A0A2S3IAN7 |
| Panicum halli | A0A2T8IKX9 |
| Panicum milia | A0A3L6TCR6 |
| Panicum milia | A0A3L6PZS7 |
| Panicum milia | A0A3L6S5J8 |
| Panicum milia | A0A3L6PDC5 |
| Panicum milia | A0A3L6QHJ2 |
| Panicum milia | A0A3L6PX19 |
| Panicum milia | A0A3L6PCT4 |
| Panicum milia | A0A3L6PL39 |
| Panicum milia | A0A3L6PX91 |
| Panicum milia | A0A3L6S5N0 |
| Panicum milia | A0A3L6QA61 |
| Panicum milia | A0A3L6PG20 |
| Panicum milia | A0A3L6SUD9 |
| Panicum milia | A0A3L6RDD5 |
| Panicum milia | A0A3L6Q4M6 |
| Panicum milia | A0A3L6PCG6 |
| Panicum milia | A0A3L6SC35 |

| | |
|---|---|
| Panicum mili | A0A3L6QNE5 |
| Panicum mili | A0A3L6PWM5 |
| Panicum mili | A0A3L6SS66 |
| Panicum mili | A0A3L6S1T6 |
| Panicum mili | A0A3L6SHN4 |
| Panicum mili | A0A3L6PTZ7 |
| Panicum mili | A0A3L6TP99 |
| Panicum mili | A0A3L6SU86 |
| Panicum mili | A0A3L6R9E9 |
| Panicum mili | A0A3L6SL66 |
| Panicum mili | A0A3L6QSF1 |
| Panicum mili | A0A3L6Q3C3 |
| Panicum mili | A0A3L6TLT5 |
| Panicum mili | A0A3L6PTZ2 |
| Panicum mili | A0A3L6Q8B0 |
| Panicum mili | A0A3L6QWW8 |
| Panicum mili | A0A3L6RTV3 |
| Panicum mili | A0A3L6SY93 |
| Panicum mili | A0A3L6THN0 |
| Panicum mili | A0A3L6S034 |
| Panicum mili | A0A3L6SRC3 |
| Panicum mili | A0A3L6QXJ3 |
| Panicum mili | A0A3L6T0D2 |
| Panicum mili | A0A3L6T329 |
| Panicum mili | A0A3L6R3K6 |
| Panicum mili | A0A3L6RCL0 |
| Panicum mili | A0A3L6T8M5 |
| Panicum mili | A0A3L6QUJ7 |
| Panicum mili | A0A3L6SPW4 |
| Panicum mili | A0A3L6T7S6 |
| Panicum mili | A0A3L6QCA5 |
| Panicum mili | A0A3L6RXE5 |
| Panicum mili | A0A3L6T8Z5 |
| Panicum mili | A0A3L6RSY0 |
| Panicum mili | A0A3L6SVG3 |
| Panicum mili | A0A3L6R496 |
| Panicum mili | A0A3L6RL96 |
| Panicum mili | A0A3L6PWI7 |
| Panicum mili | A0A3L6T589 |
| Panicum mili | A0A3L6T3C1 |
| Panicum mili | A0A3L6R910 |

| | |
|---|---|
| Panicum mili | A0A3L6QPB7 |
| Panicum mili | A0A3L6SAN9 |
| Panicum mili | A0A3L6Q7C9 |
| Panicum mili | A0A3L6T2B5 |
| Panicum mili | A0A3L6SZ69 |
| Panicum mili | A0A3L6R088 |
| Panicum mili | A0A3L6PV23 |
| Panicum mili | A0A3L6R2W0 |
| Panicum mili | A0A3L6TPL9 |
| Panicum mili | A0A3L6S8F1 |
| Panicum mili | A0A3L6QW70 |
| Panicum mili | A0A3L6PE95 |
| Panicum mili | A0A3L6R0F3 |
| Panicum mili | A0A3L6RH98 |
| Panicum mili | A0A3L6SDA3 |
| Panicum mili | A0A3L6P9T3 |
| Panicum mili | A0A3L6PX52 |
| Panicum mili | A0A3L6RQD8 |
| Panicum mili | A0A3L6S172 |
| Panicum mili | A0A3L6TFE8 |
| Panicum mili | A0A3L6SSN5 |
| Panicum mili | A0A3L6S7G8 |
| Panicum mili | A0A3L6SBX7 |
| Panicum mili | A0A3L6QI59 |
| Panicum mili | A0A3L6SGX8 |
| Panicum mili | A0A3L6PSV3 |
| Panicum mili | A0A3L6PB20 |
| Panicum mili | A0A3L6SW57 |
| Panicum mili | A0A3L6PS89 |
| Panicum mili | A0A3L6T996 |
| Panicum mili | A0A3L6QZB1 |
| Panicum mili | A0A3L6Q8B5 |
| Panicum mili | A0A3L6RFQ9 |
| Panicum mili | A0A3L6R701 |
| Panicum mili | A0A3L6QDT6 |
| Panicum mili | A0A3L6SN21 |
| Panicum mili | A0A3L6T804 |
| Panicum mili | A0A3L6QYH1 |
| Panicum mili | A0A3L6SXL6 |
| Panicum mili | A0A3L6TUW6 |
| Panicum mili | A0A3L6STY3 |

| | |
|---|---|
| Panicum milia | A0A3L6TQD8 |
| Panicum milia | A0A3L6QWJ0 |
| Panicum milia | A0A3L6T346 |
| Pannonibacte | A0A0K6HYJ7 |
| Pannonibacte | A0A378ZUH8 |
| Pannonibacte | A0A0U3PCH3 |
| Panstrongylu | A0A224XG17 |
| Panstrongylu | A0A224XJC5 |
| Panstrongylu | A0A224XLL6 |
| Panstrongylu | A0A224XI83 |
| Papilio macha | A0A194RNJ8 |
| Papilio macha | A0A194R138 |
| Papilio macha | A0A0N1PJM0 |
| Papilio macha | A0A194RDJ8 |
| Papilio macha | A0A194R2I6 |
| Papilio polyte | I4DM08 |
| Papilio xuthu | A0A0N1I4I2 |
| Papilio xuthu | A0A194PVK1 |
| Papilio xuthu | A0A194PQ35 |
| Papilio xuthu | I4DIJ2 |
| Papilio xuthu | A0A194QJF1 |
| Papilio xuthu | A0A194PTA1 |
| Papilio xuthu | A0A194PXM5 |
| Papio anubis. | A0A2I3MER4 |
| Papio anubis. | A0A096NQZ2 |
| Papio anubis. | A0A2I3MEH4 |
| Papio anubis. | A0A096MT27 |
| Papio anubis. | A0A096NZY9 |
| Papio anubis. | A0A2I3LKX1 |
| Papio anubis. | A0A096MR85 |
| Papio anubis. | A0A2I3M8D7 |
| Papio anubis. | A0A096NC95 |
| Paracoccus al | A0A1H6MTW0 |
| Paracoccus ai | S5XSK4 |
| Paracoccus b | A0A3D9XH37 |
| Paracoccus cl | A0A1G9LDL1 |
| Paracoccus co | A0A1W6CWQ5 |
| Paracoccus d | A1B4L0 |
| Paracoccus d | A0A2W5A1K7 |
| Paracoccus h | A0A099F544 |
| Paracoccus lu | A0A368YRU1 |

| | |
|---|---|
| Paracoccus p | A0A1K1YUK1 |
| Paracoccus p | A0A454NRD8 |
| Paracoccus p | A0A1I5KJ26 |
| Paracoccus sa | A0A2A2GLC7 |
| Paracoccus sa | A0A1N7M497 |
| Paracoccus sa | A0A099GIU4 |
| Paracoccus sa | A0A1H2UIH5 |
| Paracoccus sp | A0A418SQD8 |
| Paracoccus sp | A0A2K9F7T9 |
| Paracoccus sp | A0A3N7HKP6 |
| Paracoccus sp | A0A1X7K5P3 |
| Paracoccus sp | A0A3S4DYQ8 |
| Paracoccus sp | A0A0V0Q0U8 |
| Paracoccus th | A0A1N6QLJ3 |
| Paracoccus v | A0A3E0CAT2 |
| Paracoccus v | A0A369U330 |
| Paracoccus y | A0A386UNW4 |
| Paracoccus y | A0A2D2C4W0 |
| Paracoccus y | A0A1V0GXG8 |
| Paracoccus zh | A0A2H5F0H5 |
| Paralichthys ( | D1M702 |
| Paramesorhiz | A0A135HYP7 |
| Paramormyrc | A0A3B3T1V7 |
| Paramormyrc | A0A3B3T2D5 |
| Paramormyrc | A0A3B3QTI7 |
| Paramormyrc | A0A3B3T216 |
| Paramormyrc | A0A3B3R816 |
| Paramormyrc | A0A3B3T3Q0 |
| Paramormyrc | A0A3B3QMB5 |
| Paramormyrc | A0A3B3T389 |
| Paramormyrc | A0A3B3T3H8 |
| Paramormyrc | A0A3B3T3U8 |
| Paraphaeosp | A0A177CZE2 |
| Paraphaeosp | A0A177CRA0 |
| Paraphaeosp | A0A177CUG9 |
| Paraphaeosp | A0A177C382 |
| Paraphaeosp | A0A177C6N6 |
| Paraphaeosp | A0A177CID1 |
| Pararhizobiur | A0A1C7P136 |
| Parasponia ar | A0A2P5BJA4 |
| Parasponia ar | A0A2P5BW50 |

| | |
|---|---|
| Parasponia aı | A0A2P5B4R0 |
| Parasponia aı | A0A2P5BW53 |
| Parasponia aı | A0A2P5BW51 |
| Parasponia aı | A0A2P5BM82 |
| Parasponia aı | A0A2P5A6J9 |
| Parasponia aı | A0A2P5BW67 |
| Parasponia aı | A0A2P5E4M6 |
| Parasponia aı | A0A2P5D441 |
| Parasponia aı | A0A2P5BW21 |
| Parasponia aı | A0A2P5BKD8 |
| Parasponia aı | A0A2P5E4L5 |
| Parasponia aı | A0A2P5BMA8 |
| Parasponia aı | A0A2P5BKI9 |
| Parasponia aı | A0A2P5AA39 |
| Parasponia aı | A0A2P5BMB4 |
| Parasponia aı | A0A2P5B4Q4 |
| Parasponia aı | A0A2P5BM80 |
| Parasponia aı | A0A2P5DYR7 |
| Parasponia aı | A0A2P5AHJ9 |
| Parasponia aı | A0A2P5E4N0 |
| Parasponia aı | A0A2P5D258 |
| Parasponia aı | A0A2P5BW45 |
| Parasponia aı | A0A2P5BKH5 |
| Parasponia aı | A0A2P5E4Q5 |
| Parasponia aı | A0A2P5A6G1 |
| Parasponia aı | A0A2P5BM96 |
| Parasponia aı | A0A2P5B4Q6 |
| Parasponia aı | A0A2P5BW66 |
| Parasponia aı | A0A2P5BM92 |
| Parasponia aı | A0A2P5BN11 |
| Parasponia aı | A0A2P5BMD0 |
| Parasponia aı | A0A2P5CS31 |
| Parasponia aı | A0A2P5BMA5 |
| Parasponia aı | A0A2P5E4N1 |
| Parasponia aı | A0A2P5B4P1 |
| Parasponia aı | A0A2P5CSJ2 |
| Parasponia aı | A0A2P5BM72 |
| Parasponia aı | A0A2P5E4Q3 |
| Parasponia aı | A0A2P5BH32 |
| Parasponia aı | A0A2P5B4S4 |
| Parasponia aı | A0A2P5CFJ1 |

| | |
|---|---|
| Parasponia aı | A0A2P5E4M4 |
| Parasponia aı | A0A2P5DP24 |
| Parasponia aı | A0A2P5BIW1 |
| Parasponia aı | A0A2P5CC51 |
| Parasponia aı | A0A2P5BM84 |
| Parasponia aı | A0A2P5BM71 |
| Parasponia aı | A0A2P5BJC8 |
| Parasponia aı | A0A2P5B4T3 |
| Parasponia aı | A0A2P5BKJ2 |
| Parasponia aı | A0A2P5AA49 |
| Parasponia aı | A0A2P5BW30 |
| Parasponia aı | A0A2P5BW41 |
| Parasponia aı | A0A2P5BW35 |
| Parasponia aı | A0A2P5B1T6 |
| Parasponia aı | A0A2P5E4R8 |
| Parasponia aı | A0A2P5B1R8 |
| Parasponia aı | A0A2P5DGX2 |
| Parasponia aı | A0A2P5B4R2 |
| Parasponia aı | A0A2P5B4Q1 |
| Parasponia aı | A0A2P5B4S3 |
| Parasponia aı | A0A2P5BJA2 |
| Parasponia aı | A0A2P5BW77 |
| Parasponia aı | A0A2P5BW40 |
| Parasponia aı | A0A2P5B4Q8 |
| Parasponia aı | A0A2P5DYQ3 |
| Parasponia aı | A0A2P5CA29 |
| Parasponia aı | A0A2P5AIJ3 |
| Parasponia aı | A0A2P5BM74 |
| Parasponia aı | A0A2P5E4J2 |
| Parasponia aı | A0A2P5BJC9 |
| Parasponia aı | A0A2P5BM50 |
| Parasponia aı | A0A2P5A6F1 |
| Parasponia aı | A0A2P5BW89 |
| Parasponia aı | A0A2P5BW37 |
| Parasponia aı | A0A2P5BW20 |
| Parasponia aı | A0A2P5E4P8 |
| Parasponia aı | A0A2P5BW58 |
| Parasponia aı | A0A2P5E4M9 |
| Parasponia aı | A0A2P5E4P4 |
| Parasponia aı | A0A2P5E4N5 |
| Parasponia aı | A0A2P5B4R9 |

| | |
|---|---|
| Parasponia ar | A0A2P5A6G8 |
| Parasponia ar | A0A2P5DYT0 |
| Parasponia ar | A0A2P5BW31 |
| Parasponia ar | A0A2P5AEM1 |
| Parasponia ar | A0A2P5D7Y5 |
| Parasponia ar | A0A2P5BW71 |
| Parasponia ar | A0A2P5BMA2 |
| Parasponia ar | A0A2P5AH20 |
| Parasponia ar | A0A2P5B4S8 |
| Parasponia ar | A0A2P5BMC0 |
| Parasponia ar | A0A2P5BXQ1 |
| Parasponia ar | A0A2P5BVZ5 |
| Parasponia ar | A0A2P5BKL3 |
| Parasteatoda | A0A2L2YD42 |
| Parasteatoda | A0A2L2Z1W7 |
| Parasteatoda | A0A2L2Z8H1 |
| Parasteatoda | A0A2L2ZA53 |
| Parasteatoda | A0A2L2Z530 |
| Parasteatoda | A0A2L2YWS1 |
| Parastrongylo | A0A0N4ZHI4 |
| Parastrongylo | A0A0N5A0H4 |
| Parastrongylo | A0A0N4ZPS9 |
| Parastrongylo | A0A0N5A3M1 |
| Parastrongylo | A0A0N4ZRF2 |
| Parastrongylo | A0A0N4Z938 |
| Parastrongylo | A0A0N4Z4J1 |
| Parastrongylo | A0A0N4ZEA2 |
| Parastrongylo | A0A0N4Z1R8 |
| Parastrongylo | A0A0N4Z6U2 |
| Parastrongylo | A0A0N4ZP70 |
| Parastrongylo | A0A0N5A5G9 |
| Parastrongylo | A0A0N4ZHI3 |
| Parastrongylo | A0A0N4ZNC4 |
| Parastrongylo | A0A0N4ZKC3 |
| Parastrongylo | A0A0N4ZPV9 |
| Parastrongylo | A0A0N4Z312 |
| Parastrongylo | A0A0N4ZIU0 |
| Parastrongylo | A0A0N4Z5S0 |
| Parastrongylo | A0A0N4ZS52 |
| Paroedura pi | A0A402ESB7 |
| Paroedura pi | A0A402FAU3 |

| | |
|---|---|
| Paroedura pi | A0A402FMT0 |
| Paroedura pi | A0A402EEM0 |
| Parvibaculum | A0A2D8XC50 |
| Patagioenas f | A0A1V4K051 |
| Patagioenas f | A0A1V4JKY3 |
| Patagioenas f | A0A1V4JG98 |
| Patagioenas f | A0A1V4JKX2 |
| Patagioenas f | A0A1V4K8H7 |
| Patagioenas f | A0A1V4K1P3 |
| Patagioenas f | A0A1V4K8P3 |
| Pectinophora | A0A1E1WSI0 |
| Pectinophora | A0A1E1WED2 |
| Pediculus hur | E0VCL9 |
| Pediculus hur | E0VHE3 |
| Pediculus hur | E0VC84 |
| Pediculus hur | E0VX62 |
| Pediculus hur | E0VX61 |
| Pediculus hur | E0VJE4 |
| Pelagibaca ab | A0A1P8UUY1 |
| Pelagibaca be | Q0FI61 |
| Pelagibaca sp | A0A2D6BAV5 |
| Pelagibacteri | A0A1E4DW35 |
| Pelagibacteri | A0A1E4FC43 |
| Pelagibacteri | A0A1E4GBY0 |
| Pelagibacteri | A0A1E4GLE6 |
| Pelagicola lito | A0A1Y5RQR3 |
| Pelagicola sp. | A0A2U2CX27 |
| Pelagimonas | A0A238KIZ4 |
| Pelagivirga se | A0A2T7G6G5 |
| Pelecanus cri | A0A091SM44 |
| Pelecanus cri | A0A091TMT4 |
| Pelecanus cri | A0A091TWU6 |
| Pelecanus cri | A0A091STZ2 |
| Pelodiscus sir | E5RLR1 |
| Pelodiscus sir | K7GFR6 |
| Pelodiscus sir | K7GGY8 |
| Pelodiscus sir | K7FLF2 |
| Pelolinea sub | A0A347ZT03 |
| Penaeus vanr | A0A3R7LS22 |
| Penaeus vanr | A0A423SJ34 |
| Penaeus vanr | A0A423TTP3 |

| | |
|---|---|
| Penaeus vanr | A0A3R7M4Q0 |
| Penaeus vanr | A0A423SF89 |
| Penaeus vanr | A0A3R7MK76 |
| Penaeus vanr | A0A3R7P3K5 |
| Penaeus vanr | A0A423U6F6 |
| Penaeus vanr | A0A423TME0 |
| Penaeus vanr | A0A423TTP4 |
| Penaeus vanr | A0A3R7M498 |
| Penaeus vanr | A0A3R7MRL0 |
| Penaeus vanr | A0A3R7P0E8 |
| Penaeus vanr | A0A3R7PNR3 |
| Penaeus vanr | A0A3R7PF83 |
| Penicilliopsis | A0A1L9S4X6 |
| Penicilliopsis | A0A1L9SBG1 |
| Penicillium ar | A0A1V6PCL2 |
| Penicillium ar | A0A1F5LG55 |
| Penicillium ar | A0A1F5LFH2 |
| Penicillium ar | A0A1F5LBV1 |
| Penicillium ar | A0A1F5L7N0 |
| Penicillium ar | A0A1F5LFH7 |
| Penicillium ar | A0A1F5L4M3 |
| Penicillium ca | A0A0G4PQR1 |
| Penicillium co | A0A1V6UT76 |
| Penicillium co | A0A1V6URX4 |
| Penicillium co | A0A1V6USL6 |
| Penicillium de | A0A1V6PHF0 |
| Penicillium ex | A0A0A2ITH3 |
| Penicillium fla | A0A1V6SGC7 |
| Penicillium ita | A0A0A2KKT0 |
| Penicillium na | A0A1V6WU75 |
| Penicillium na | A0A1V6Z413 |
| Penicillium ox | S8B8Z7 |
| Penicillium ox | S7ZRP3 |
| Penicillium ox | S7ZGN1 |
| Penicillium ox | S7ZMP6 |
| Penicillium pa | A0A135LCH0 |
| Penicillium pa | A0A135LCH3 |
| Penicillium pa | A0A135LCQ1 |
| Penicillium ro | W6Q172 |
| Penicillium ru | B6HSU8 |
| Penicillium sc | A0A1V6QSF1 |

| | |
|---|---|
| Penicillium sc | A0A1V6QSE8 |
| Penicillium sp | A0A2H3III4 |
| Penicillium sp | A0A2H3IC68 |
| Penicillium st | A0A1V6SRD0 |
| Penicillium su | A0A1Q5T8E6 |
| Penicillium su | A0A1Q5TYZ6 |
| Penicillium vu | A0A1V6RF18 |
| Penicillium vu | A0A1V6RE75 |
| Penicillium vu | A0A1V6RG25 |
| Penicillium vu | A0A1V6S597 |
| Penicillium vu | A0A1V6RFI7 |
| Perca flavesc | F1C726 |
| Perca flavesc | F1C7D0 |
| Periconia ma | A0A2V1DWV5 |
| Periconia ma | A0A2V1D2D5 |
| Periconia ma | A0A2V1DIZ4 |
| Periconia ma | A0A2V1DJW4 |
| Periconia ma | A0A2V1DLD9 |
| Periconia ma | A0A2V1D8T0 |
| Periophthalm | A0A3B3ZPQ5 |
| Periophthalm | A0A3B3ZY43 |
| Periophthalm | A0A3B3ZXI2 |
| Periophthalm | A0A3B3ZP74 |
| Periophthalm | A0A3B3ZYU4 |
| Periophthalm | A0A3B4AGU9 |
| Periophthalm | A0A3B3ZP80 |
| Periophthalm | A0A3B3ZYS8 |
| Perkinsela sp | A0A0L1L095 |
| Peronospora | A0A3M6VL27 |
| Peronospora | A0A3M6VAW8 |
| Peronospora | A0A425CJM8 |
| Peronospora | A0A3R7Y325 |
| Peronospora | A0A3M6V6Z7 |
| Pestalotiopsis | W3XET2 |
| Pestalotiopsis | W3X3J5 |
| Pestalotiopsis | W3XMF5 |
| Pestalotiopsis | W3XPT6 |
| Pestalotiopsis | W3XC43 |
| Pestalotiopsis | W3WRT1 |
| Pestalotiopsis | W3XNX9 |
| Pezoloma eri | A0A2J6QKC3 |

| | |
|---|---|
| Phaeobacter | A0A366X020 |
| Phaeobacter | A0A1B0ZTK3 |
| Phaeobacter | A0A0P1GVW2 |
| Phaeobacter | A0A0P1IHK9 |
| Phaeodactylu | B7G5U0 |
| Phaeomoniel | A0A0G2E5S8 |
| Phaeosphaer | Q0UMP0 |
| Phaeosphaer | Q0U636 |
| Phaeosphaer | Q0V168 |
| Phaeosphaer | Q0V5G3 |
| Phaeosphaer | Q0V4N1 |
| Phaeosphaer | Q0TZX6 |
| Phaeosphaer | Q0UPL0 |
| Phaeosphaer | Q0UWY3 |
| Phaeosphaer | Q0UP20 |
| Phaethon lep | A0A091ULD4 |
| Phaethon lep | A0A091T5G0 |
| Phaethon lep | A0A091TAK8 |
| Phaethon lep | A0A091TBV2 |
| Phalacrocora | A0A093QYY4 |
| Phalacrocora | A0A093TDY9 |
| Phalacrocora | A0A093QPS2 |
| Phalacrocora | A0A093QSI2 |
| Phaseolus an | A0A0L9TKR2 |
| Phaseolus an | A0A0L9VGP0 |
| Phaseolus an | A0A0L9UFV5 |
| Phaseolus an | A0A0L9ULD5 |
| Phaseolus an | A0A0L9TCV5 |
| Phaseolus an | A0A0L9V3Z2 |
| Phaseolus an | A0A0L9UKZ9 |
| Phaseolus an | A0A0L9U8P2 |
| Phaseolus an | A0A0L9U553 |
| Phaseolus an | A0A0L9TLV1 |
| Phaseolus an | A0A0L9U6Q9 |
| Phaseolus an | A0A0L9UIH5 |
| Phaseolus an | A0A0L9ULI3 |
| Phaseolus an | A0A0L9UM53 |
| Phaseolus an | A0A0L9U552 |
| Phaseolus an | A0A0L9V319 |
| Phaseolus an | A0A0L9UN60 |
| Phaseolus an | A0A0L9TLV8 |

| | |
|---|---|
| Phaseolus an | A0A0L9ULD1 |
| Phaseolus an | A0A0L9V362 |
| Phaseolus an | A0A0L9V8X3 |
| Phaseolus an | A0A0L9V600 |
| Phaseolus an | A0A0L9ULS9 |
| Phaseolus an | A0A0L9T851 |
| Phaseolus an | A0A0L9VUB8 |
| Phaseolus an | A0A0L9TGQ7 |
| Phaseolus an | A0A0L9U4V0 |
| Phaseolus an | A0A0L9THP8 |
| Phaseolus an | A0A0L9VUS9 |
| Phaseolus an | A0A0L9V9B9 |
| Phaseolus an | A0A0L9UIP6 |
| Phaseolus an | A0A0L9UPU8 |
| Phaseolus an | A0A0L9TKM5 |
| Phaseolus an | A0A0L9V913 |
| Phaseolus an | A0A0L9VTA1 |
| Phaseolus an | A0A0L9V3F3 |
| Phaseolus an | A0A0L9V376 |
| Phaseolus an | A0A0L9UN55 |
| Phaseolus an | A0A0L9V4V3 |
| Phaseolus an | A0A0L9V1W9 |
| Phaseolus an | A0A0L9V3H0 |
| Phaseolus an | A0A0L9VFN3 |
| Phaseolus an | A0A0L9UTN1 |
| Phaseolus an | A0A0L9TGU5 |
| Phaseolus an | A0A0L9V3V3 |
| Phaseolus an | A0A0L9UPN2 |
| Phaseolus an | A0A0L9TGG9 |
| Phaseolus an | A0A0L9V486 |
| Phaseolus an | A0A0L9UMB7 |
| Phaseolus an | A0A0L9VG17 |
| Phaseolus an | A0A0L9UJE5 |
| Phaseolus an | A0A0L9TUI5 |
| Phaseolus an | A0A0L9V464 |
| Phaseolus an | A0A0L9VFU0 |
| Phaseolus an | A0A0L9U5F8 |
| Phaseolus an | A0A0L9U882 |
| Phaseolus an | A0A0L9UJ28 |
| Phaseolus an | A0A0L9VSL2 |
| Phaseolus an | A0A0L9UMD9 |

| | |
|---|---|
| Phaseolus an | A0A0L9UM48 |
| Phaseolus an | A0A0L9VFN1 |
| Phaseolus an | A0A0L9V5Z3 |
| Phaseolus vu | V7BQJ3 |
| Phaseolus vu | V7CQM2 |
| Phaseolus vu | V7AIN2 |
| Phaseolus vu | V7BX92 |
| Phaseolus vu | V7B6U7 |
| Phaseolus vu | V7AIF8 |
| Phaseolus vu | V7BXZ6 |
| Phaseolus vu | V7ALS3 |
| Phaseolus vu | V7B4W4 |
| Phaseolus vu | V7AHY7 |
| Phaseolus vu | V7AGA5 |
| Phaseolus vu | V7AIM9 |
| Phaseolus vu | V7AIL8 |
| Phaseolus vu | V7CPR4 |
| Phaseolus vu | V7BXT0 |
| Phaseolus vu | V7AIF2 |
| Phaseolus vu | V7AHM7 |
| Phaseolus vu | V7BUN3 |
| Phaseolus vu | V7ALT3 |
| Phaseolus vu | V7AMQ9 |
| Phaseolus vu | V7AIH2 |
| Phaseolus vu | V7AJU3 |
| Phaseolus vu | V7APL9 |
| Phaseolus vu | V7B7X1 |
| Phaseolus vu | V7ALI5 |
| Phaseolus vu | V7AHN7 |
| Phaseolus vu | V7BVD1 |
| Phaseolus vu | V7AHQ4 |
| Phaseolus vu | V7AMQ6 |
| Phaseolus vu | V7AJR9 |
| Phaseolus vu | V7BRW9 |
| Phaseolus vu | V7AE52 |
| Phaseolus vu | V7ALQ7 |
| Phaseolus vu | V7BV89 |
| Phaseolus vu | V7AJK3 |
| Phaseolus vu | V7D2F6 |
| Phaseolus vu | V7BZ49 |
| Phaseolus vu | V7AHK3 |

Phaseolus vu V7CPG4
Phaseolus vu V7B5J8
Phaseolus vu V7AID3
Phaseolus vu V7AIM3
Phaseolus vu V7BYK2
Phaseolus vu V7B3V1
Phaseolus vu V7AH65
Phaseolus vu V7BXM2
Phaseolus vu V7AJP5
Phaseolus vu V7BYJ9
Phaseolus vu V7BQM9
Phaseolus vu V7BZ42
Phaseolus vu V7B3X6
Phaseolus vu V7AJE9
Phaseolus vu V7AJS5
Phaseolus vu V7ATU1
Phaseolus vu V7CPG0
Phaseolus vu V7CM72
Phaseolus vu V7AST9
Phaseolus vu V7ALL6
Phaseolus vu V7CZ14
Phaseolus vu V7BTA1
Phaseolus vu V7AKL3
Phaseolus vu V7BXZ1
Phaseolus vu V7AHE0
Phaseolus vu V7ALR2
Phaseolus vu V7ALS5
Phaseolus vu V7AKL1
Phaseolus vu V7AVN9
Phaseolus vu V7C143
Phaseolus vu V7C7H2
Phaseolus vu V7CA41
Phaseolus vu V7ALS8
Phaseolus vu V7B451
Phaseolus vu V7D003
Phaseolus vu V7ANE1
Phaseolus vu V7AVN4
Phaseolus vu V7BUI7
Phaseolus vu V7AMN5
Phaseolus vu V7AHR0
Phaseolus vu V7BTA2

| | |
|---|---|
| Phaseolus vu | V7AMM7 |
| Phaseolus vu | V7AHK9 |
| Phaseolus vu | V7AM09 |
| Phaseolus vu | V7ALF8 |
| Phaseolus vu | V7BUL6 |
| Phaseolus vu | V7AGA9 |
| Phaseolus vu | V7AHL4 |
| Phaseolus vu | V7BWF6 |
| Phialocephala | A0A194X242 |
| Phialocephala | A0A194XBS6 |
| Phialocephala | A0A132BBB6 |
| Phialocephala | A0A194WTN9 |
| Phialocephala | A0A132B592 |
| Phialocephala | A0A194X7W7 |
| Phialocephala | A0A194X856 |
| Phialocephala | A0A194XR54 |
| Phialocephala | A0A194WSB8 |
| Phialocephala | A0A194WSX5 |
| Phialocephala | A0A194WTK6 |
| Phialocephala | A0A194XX66 |
| Phialocephala | A0A132BCG7 |
| Phialocephala | A0A132B964 |
| Phialocephala | A0A1L7XLE9 |
| Phialocephala | A0A1L7X859 |
| Phialocephala | A0A1L7WF44 |
| Phialocephala | A0A1L7XLL5 |
| Phialocephala | A0A1L7XTN8 |
| Phialocephala | A0A1L7X9R0 |
| Phialocephala | A0A1L7WMU3 |
| Phialophora a | A0A0N0NQR5 |
| Phialophora a | A0A0N1HGJ5 |
| Phialophora a | A0A0N0NQ51 |
| Phialophora a | A0A0N1GX61 |
| Phialophora a | A0A0N1H4B9 |
| Phialophora a | A0A0N1NZ06 |
| Phialophora a | A0A0N1HG31 |
| Phialophora c | A0A370TCU2 |
| Phialophora c | A0A370TYQ2 |
| Phialophora c | A0A370TJW0 |
| Phialophora c | A0A370TQF0 |
| Phlebotomus | A0A1B0GNH6 |

| | |
|---|---|
| Phlebotomus | A0A1B0D9P3 |
| Phlebotomus | A0A1B0DGU9 |
| Phlebotomus | A0A1B0DI86 |
| Phlebotomus | A0A1B0GP61 |
| Phoenicopter | A0A091U239 |
| Phoenicopter | A0A091UAN5 |
| Phoenix dact | A0A3Q0HVZ9 |
| Phoenix dact | A0A3Q0HMY7 |
| Phoenix dact | A0A3Q0I1K4 |
| Phoenix dact | A0A2H3ZIZ6 |
| Phoenix dact | A0A2H3ZAB6 |
| Phoenix dact | A0A3Q0I914 |
| Phoenix dact | A0A2H3X918 |
| Phoenix dact | A0A3Q0HT42 |
| Phoenix dact | A0A2H3ZS58 |
| Phoenix dact | A0A3Q0IFF5 |
| Phoenix dact | A0A3Q0HQF8 |
| Phoenix dact | A0A3Q0HSM9 |
| Phoenix dact | A0A3Q0HVI7 |
| Phoenix dact | A0A3Q0HTF0 |
| Phoenix dact | A0A2H3XGF9 |
| Phoenix dact | A0A2H3X9J5 |
| Phoenix dact | A0A2H3X0Q6 |
| Phoenix dact | A0A2H3X3J0 |
| Phoenix dact | A0A2H3YZ59 |
| Phoenix dact | A0A2H3Y805 |
| Phoenix dact | A0A2H3X8K1 |
| Phoenix dact | A0A2H3XA64 |
| Phoenix dact | A0A2H3WXV9 |
| Phoenix dact | A0A2H3X7L6 |
| Phoenix dact | A0A2H3ZCS3 |
| Phoenix dact | A0A2H3ZIK9 |
| Phoenix dact | A0A3Q0HR72 |
| Phoenix dact | A0A2H3XQV7 |
| Phoenix dact | A0A2H4A1V3 |
| Phoenix dact | A0A2H3ZJF3 |
| Phoenix dact | A0A3Q0HQX3 |
| Phoenix dact | A0A3Q0HSL1 |
| Phoenix dact | A0A2H3X0N3 |
| Phoenix dact | A0A2H3YKV9 |
| Phoenix dact | A0A2H3Z3E8 |

| | |
|---|---|
| Phoenix dact | A0A3Q0HVZ4 |
| Phoenix dact | A0A3Q0HTK9 |
| Phoenix dact | A0A2H3XJR1 |
| Phoenix dact | A0A3Q0I0R7 |
| Phoenix dact | A0A2H3YI52 |
| Phoenix dact | A0A2H3X0N6 |
| Phoenix dact | A0A3Q0I4E3 |
| Phoenix dact | A0A2H3Y849 |
| Phoenix dact | A0A2H3ZQ89 |
| Phoenix dact | A0A2H3Y8E0 |
| Phoenix dact | A0A2H3ZT49 |
| Phoenix dact | A0A3Q0IGU8 |
| Phoenix dact | A0A3Q0HRL0 |
| Phoenix dact | A0A2H3YUK0 |
| Phoenix dact | A0A2H3X6E4 |
| Phoenix dact | A0A3Q0IBE3 |
| Phoenix dact | A0A3Q0I4N2 |
| Phoenix dact | A0A2H3Z4R9 |
| Phoenix dact | A0A2H3Y059 |
| Phoenix dact | A0A2H3WYX2 |
| Phoenix dact | A0A3Q0HQU7 |
| Phoenix dact | A0A2H3ZI64 |
| Phoenix dact | A0A2H3XQR6 |
| Phoenix dact | A0A2H3ZW91 |
| Photinus pyra | A0A1Y1KHG4 |
| Photinus pyra | A0A1Y1MWD5 |
| Photinus pyra | A0A1Y1N456 |
| Photinus pyra | A0A1Y1LYJ5 |
| Photinus pyra | A0A1Y1K6Q9 |
| Photinus pyra | A0A1Y1KVK9 |
| Photinus pyra | A0A1Y1NDL0 |
| Photinus pyra | A0A1Y1MWC5 |
| Photinus pyra | A0A1Y1KDA6 |
| Photinus pyra | A0A1Y1N145 |
| Photinus pyra | A0A1Y1N670 |
| Photinus pyra | A0A1Y1MXX0 |
| Phyllobacteri | A0A2E1XX07 |
| Phyllobacteri | A0A368ZAV4 |
| Phyllobacteri | A0A368Z5S6 |
| Phyllobacteri | A0A2P7BX27 |
| Phyllobacteri | A0A2P7B2B9 |

| | |
|---|---|
| Phyllobacteri | A0A2P7B199 |
| Phyllobacteri | A0A318STX6 |
| Phyllobacteri | A0A2S9JB27 |
| Phyllobacteri | A0A368K297 |
| Phyllobacteri | A0A2P7BBS7 |
| Phyllobacteri | A0A2S9IZ33 |
| Phyllobacteri | A0A1H0QMP3 |
| Phyllobacteri | J3CCV4 |
| Phyllobacteri | A0A1H0UPJ4 |
| Phyllobacteri | A0A2N9VRL8 |
| Physcomitrel | A9SNW9 |
| Physcomitrel | A0A2K1K665 |
| Physcomitrel | A0A2K1KEJ6 |
| Physcomitrel | A0A2K1K664 |
| Physcomitrel | A0A2K1IS53 |
| Physcomitrel | A9T7Y0 |
| Physcomitrel | A0A2K1JCS2 |
| Physcomitrel | A0A2K1ISB0 |
| Physcomitrel | A9TEC1 |
| Physcomitrel | A9SP81 |
| Physeter mac | A0A2Y9FNN2 |
| Physeter mac | A0A2Y9FMM8 |
| Physeter mac | A0A2Y9S5B0 |
| Physeter mac | A0A2Y9FL53 |
| Physeter mac | A0A2Y9FNH7 |
| Physeter mac | A0A2Y9FVN1 |
| Physeter mac | A0A2Y9RZG7 |
| Phytophthora | A0A329R7W9 |
| Phytophthora | A0A329R9F3 |
| Phytophthora | A0A329RYD9 |
| Phytophthora | A0A329SSU7 |
| Phytophthora | A0A329RKN5 |
| Phytophthora | A0A329S4Z6 |
| Phytophthora | A0A329T4J1 |
| Phytophthora | A0A329SHI1 |
| Phytophthora | A0A329S9V0 |
| Phytophthora | A0A329SRY9 |
| Phytophthora | A0A329RNK1 |
| Phytophthora | A0A329SNY9 |
| Phytophthora | A0A329S7M0 |
| Phytophthora | A0A329S1Z2 |

| | |
|---|---|
| Phytophthora | A0A329SRE1 |
| Phytophthora | A0A329S1D7 |
| Phytophthora | A0A329SWT6 |
| Phytophthora | A0A329S4B3 |
| Phytophthora | A0A329SPJ8 |
| Phytophthora | A0A329RIU8 |
| Phytophthora | A0A329SPU9 |
| Phytophthora | A0A329RT73 |
| Phytophthora | D0MY50 |
| Phytophthora | D0NX36 |
| Phytophthora | D0NX35 |
| Phytophthora | D0NTA4 |
| Phytophthora | D0MY47 |
| Phytophthora | D0P016 |
| Phytophthora | D0NEY9 |
| Phytophthora | D0NYL6 |
| Phytophthora | D0NTB0 |
| Phytophthora | D0P1E8 |
| Phytophthora | D0N2D7 |
| Phytophthora | D0N7R8 |
| Phytophthora | D0MT29 |
| Phytophthora | D0NRT7 |
| Phytophthora | D0MRR0 |
| Phytophthora | D0MY49 |
| Phytophthora | D0P0T9 |
| Phytophthora | D0NUN5 |
| Phytophthora | D0NQ00 |
| Phytophthora | D0NZF4 |
| Phytophthora | D0N386 |
| Phytophthora | D0P028 |
| Phytophthora | Q2M441 |
| Phytophthora | B9VUU8 |
| Phytophthora | A0A421GDD8 |
| Phytophthora | A0A3R7K3Z6 |
| Phytophthora | A0A3F2RDE5 |
| Phytophthora | A0A421FHH8 |
| Phytophthora | A0A3R7K0A4 |
| Phytophthora | A0A3R7HDZ2 |
| Phytophthora | A0A421ETD0 |
| Phytophthora | A0A421FA97 |
| Phytophthora | A0A3F2RJX1 |

| | |
|---|---|
| Phytophthora | A0A3F2RE35 |
| Phytophthora | A0A3F2RF52 |
| Phytophthora | A0A3R7IQA6 |
| Phytophthora | A0A3F2RY02 |
| Phytophthora | A0A3F2RER3 |
| Phytophthora | A0A3F2RZC8 |
| Phytophthora | A0A3F2RR68 |
| Phytophthora | A0A3R7KUW5 |
| Phytophthora | A0A3R7JT10 |
| Phytophthora | A0A3R7J7R7 |
| Phytophthora | A0A3F2RF49 |
| Phytophthora | A0A3R7G004 |
| Phytophthora | A0A3R7HP88 |
| Phytophthora | A0A225WAC9 |
| Phytophthora | A0A225WNV7 |
| Phytophthora | A0A225VPW0 |
| Phytophthora | A0A225VX31 |
| Phytophthora | A0A225V0S2 |
| Phytophthora | A0A225WNJ4 |
| Phytophthora | A0A225VUT8 |
| Phytophthora | A0A225V861 |
| Phytophthora | A0A225VY03 |
| Phytophthora | A0A225VWU9 |
| Phytophthora | A0A225WU58 |
| Phytophthora | A0A225UTR0 |
| Phytophthora | A0A225US74 |
| Phytophthora | A0A225VHE2 |
| Phytophthora | A0A225WUN7 |
| Phytophthora | A0A225VTE7 |
| Phytophthora | A0A225WIB3 |
| Phytophthora | A0A225WKQ4 |
| Phytophthora | A0A0W8C7T1 |
| Phytophthora | A0A0W8CDF7 |
| Phytophthora | A0A0W8DMY0 |
| Phytophthora | A0A0W8DQS0 |
| Phytophthora | A0A0W8C6H3 |
| Phytophthora | A0A0W8DC43 |
| Phytophthora | A0A0W8CY53 |
| Phytophthora | A0A0W8C244 |
| Phytophthora | A0A0W8DC46 |
| Phytophthora | A0A0W8BQT9 |

| | |
|---|---|
| Phytophthora | A0A0W8BN24 |
| Phytophthora | A0A0W8C8H2 |
| Phytophthora | A0A0W8CP70 |
| Phytophthora | A0A0W8C0L1 |
| Phytophthora | A0A0W8CT29 |
| Phytophthora | A0A0W8DME7 |
| Phytophthora | A0A0W8D9R0 |
| Phytophthora | A0A0W8C2P6 |
| Phytophthora | A0A0W8CY27 |
| Phytophthora | A0A0W8CIP5 |
| Phytophthora | A0A0W8B357 |
| Phytophthora | A0A0W8AZV8 |
| Phytophthora | A0A2P4YEQ1 |
| Phytophthora | A0A2P4YER5 |
| Phytophthora | A0A2P4Y0Y6 |
| Phytophthora | A0A2P4XL73 |
| Phytophthora | A0A2P4XH46 |
| Phytophthora | A0A2P4XJK4 |
| Phytophthora | A0A2P4XVK1 |
| Phytophthora | A0A2P4Y6J7 |
| Phytophthora | A0A2P4X3D0 |
| Phytophthora | A0A2P4XLE8 |
| Phytophthora | A0A2P4X3F5 |
| Phytophthora | A0A2P4YFR7 |
| Phytophthora | A0A2P4Y2R6 |
| Phytophthora | A0A2P4YH90 |
| Phytophthora | A0A2P4XUW4 |
| Phytophthora | W2Q467 |
| Phytophthora | W2RHF8 |
| Phytophthora | W2QVF4 |
| Phytophthora | W2Q4M7 |
| Phytophthora | W2PJM9 |
| Phytophthora | W2PLH6 |
| Phytophthora | W2Q1I5 |
| Phytophthora | W2PKG5 |
| Phytophthora | W2QD00 |
| Phytophthora | W2QZF4 |
| Phytophthora | W2QTN8 |
| Phytophthora | W2Q634 |
| Phytophthora | W2Q9Y7 |
| Phytophthora | W2PJB5 |

| | |
|---|---|
| Phytophthora | W2R209 |
| Phytophthora | W2PFP4 |
| Phytophthora | W2PU68 |
| Phytophthora | W2QCZ4 |
| Phytophthora | W2REW3 |
| Phytophthora | W2PEG7 |
| Phytophthora | W2Q870 |
| Phytophthora | W2WEV6 |
| Phytophthora | W2X913 |
| Phytophthora | W2X1M9 |
| Phytophthora | W2X5D1 |
| Phytophthora | W2VX79 |
| Phytophthora | W2WUZ2 |
| Phytophthora | W2X348 |
| Phytophthora | W2XWF0 |
| Phytophthora | W2VZ24 |
| Phytophthora | W2WSU4 |
| Phytophthora | W2W607 |
| Phytophthora | W2W0N8 |
| Phytophthora | W2VXA4 |
| Phytophthora | W2XT69 |
| Phytophthora | W2WTF3 |
| Phytophthora | W2W2G5 |
| Phytophthora | W2XAA7 |
| Phytophthora | W2WHH5 |
| Phytophthora | W2VUZ5 |
| Phytophthora | W2XAB2 |
| Phytophthora | W3A0G8 |
| Phytophthora | W2Y5P0 |
| Phytophthora | W2ZM34 |
| Phytophthora | W3A6G4 |
| Phytophthora | W2ZDS8 |
| Phytophthora | W2ZHI4 |
| Phytophthora | W2YS73 |
| Phytophthora | W2Z3U6 |
| Phytophthora | W2YNN5 |
| Phytophthora | W2Y9I4 |
| Phytophthora | W2Z3S0 |
| Phytophthora | W2ZBQ5 |
| Phytophthora | W2YBW1 |
| Phytophthora | W2ZD75 |

| | |
|---|---|
| Phytophthora | W2ZHQ3 |
| Phytophthora | W2YGM9 |
| Phytophthora | W2Y8A6 |
| Phytophthora | W2YD26 |
| Phytophthora | W2Z356 |
| Phytophthora | W2YBY5 |
| Phytophthora | V9EY76 |
| Phytophthora | V9E3H4 |
| Phytophthora | V9DZZ5 |
| Phytophthora | V9FZZ4 |
| Phytophthora | V9FE93 |
| Phytophthora | V9E8N5 |
| Phytophthora | V9EY37 |
| Phytophthora | V9EX27 |
| Phytophthora | V9F9H0 |
| Phytophthora | V9E3Y2 |
| Phytophthora | V9ENS5 |
| Phytophthora | V9F7A9 |
| Phytophthora | V9F7C6 |
| Phytophthora | V9E3S0 |
| Phytophthora | V9FXZ9 |
| Phytophthora | V9FC98 |
| Phytophthora | V9EKG8 |
| Phytophthora | V9EI96 |
| Phytophthora | V9FDK5 |
| Phytophthora | V9EC42 |
| Phytophthora | V9E1H4 |
| Phytophthora | A0A081AYV0 |
| Phytophthora | A0A081A0S9 |
| Phytophthora | A0A081A8Z3 |
| Phytophthora | A0A080ZLK1 |
| Phytophthora | A0A081AB04 |
| Phytophthora | A0A081B3V1 |
| Phytophthora | A0A080Z747 |
| Phytophthora | A0A081A0T1 |
| Phytophthora | A0A080Z748 |
| Phytophthora | A0A080Z738 |
| Phytophthora | A0A080ZP81 |
| Phytophthora | A0A081AYU8 |
| Phytophthora | A0A080Z4Y8 |
| Phytophthora | A0A081AYU4 |

| | |
|---|---|
| Phytophthora | A0A081A1R2 |
| Phytophthora | A0A080Z2Z3 |
| Phytophthora | A0A081AB01 |
| Phytophthora | A0A081AH92 |
| Phytophthora | A0A080ZDR9 |
| Phytophthora | A0A080ZAI3 |
| Phytophthora | W2LFS8 |
| Phytophthora | W2HKN0 |
| Phytophthora | W2L0R5 |
| Phytophthora | W2P1G7 |
| Phytophthora | W2FR98 |
| Phytophthora | W2K4B4 |
| Phytophthora | W2J7J1 |
| Phytophthora | W2KMQ1 |
| Phytophthora | W2FPQ8 |
| Phytophthora | W2K6H5 |
| Phytophthora | W2K5X3 |
| Phytophthora | W2P694 |
| Phytophthora | W2L728 |
| Phytophthora | W2K2G5 |
| Phytophthora | W2MHW4 |
| Phytophthora | W2NET7 |
| Phytophthora | W2GUQ1 |
| Phytophthora | W2J9V7 |
| Phytophthora | W2GNC4 |
| Phytophthora | W2H193 |
| Phytophthora | O42830 |
| Phytophthora | W2HY18 |
| Phytophthora | W2K697 |
| Phytophthora | W2MTG5 |
| Phytophthora | W2MC44 |
| Phytophthora | W2KBB4 |
| Phytophthora | W2G7E9 |
| Phytophthora | W2IG72 |
| Phytophthora | W2J2Z4 |
| Phytophthora | W2J2Z6 |
| Phytophthora | W2MA47 |
| Phytophthora | W2FRI0 |
| Phytophthora | W2M865 |
| Phytophthora | W2GWC8 |
| Phytophthora | W2GWD7 |

| | |
|---|---|
| Phytophthora | W2L8Q2 |
| Phytophthora | W2I021 |
| Phytophthora | W2M2S0 |
| Phytophthora | W2N786 |
| Phytophthora | W2N694 |
| Phytophthora | W2G037 |
| Phytophthora | W2G9R5 |
| Phytophthora | W2JQ56 |
| Phytophthora | W2I5H9 |
| Phytophthora | W2HZQ8 |
| Phytophthora | W2MA56 |
| Phytophthora | W2ISN0 |
| Phytophthora | W2H2J5 |
| Phytophthora | W2NEV2 |
| Phytophthora | W2HNT1 |
| Phytophthora | W2JB43 |
| Phytophthora | W2I254 |
| Phytophthora | W2MW48 |
| Phytophthora | W2NEU1 |
| Phytophthora | H3G593 |
| Phytophthora | H3H1Z2 |
| Phytophthora | H3GT31 |
| Phytophthora | H3HDS4 |
| Phytophthora | H3H1Y9 |
| Phytophthora | H3H2Y9 |
| Phytophthora | H3H2Y8 |
| Phytophthora | H3G5T4 |
| Phytophthora | H3H1Y8 |
| Phytophthora | B2ZU37 |
| Phytophthora | B2ZU68 |
| Phytophthora | B2ZU62 |
| Phytophthora | H3GQA8 |
| Phytophthora | B2ZU49 |
| Phytophthora | H3H1Y6 |
| Phytophthora | H3H1U8 |
| Phytophthora | H3GLB2 |
| Phytophthora | H3G5P9 |
| Phytophthora | H3H1Z1 |
| Phytophthora | B2ZU72 |
| Phytophthora | H3GQA7 |
| Phytophthora | B2ZU45 |

| | |
|---|---|
| Phytophthora | H3H2Z0 |
| Phytophthora | H3GT32 |
| Phytophthora | H3G5B0 |
| Phytophthora | H3H2Y7 |
| Phytophthora | H3GDG9 |
| Phytophthora | B2ZU71 |
| Phytophthora | H3H1Z0 |
| Phytophthora | B2ZU70 |
| Phytophthora | B2ZU32 |
| Phytophthora | A0A088CB98 |
| Phytophthora | B2ZU23 |
| Phytophthora | H3GVW9 |
| Phytophthora | B2ZU31 |
| Phytophthora | A0A2L0WUS4 |
| Phytophthora | A1EAQ0 |
| Phytophthora | G4Z5J6 |
| Phytophthora | G4ZN72 |
| Phytophthora | G4ZGF9 |
| Phytophthora | G4YIR5 |
| Phytophthora | G5ABJ2 |
| Phytophthora | G4Z8P9 |
| Phytophthora | G5AEI8 |
| Phytophthora | G4ZGF6 |
| Phytophthora | G4YRJ3 |
| Phytophthora | G4Z802 |
| Phytophthora | G5A0U5 |
| Phytophthora | G4YU93 |
| Phytophthora | G4YIR6 |
| Phytophthora | G4YTG2 |
| Phytophthora | G5AEI6 |
| Phytophthora | G4Z8P8 |
| Phytophthora | G4ZRE5 |
| Phytophthora | G4YIR7 |
| Phytophthora | G4YXF1 |
| Phytophthora | G4Z8Q8 |
| Phytophthora | G4Z5J7 |
| Phytophthora | G4YRE7 |
| Phytophthora | G4ZT51 |
| Phytophthora | G4Z5J8 |
| Phytophthora | G4Z8Q0 |
| Phytophthora | G5AEI5 |

| | |
|---|---|
| Phytophthora | G4YW39 |
| Phytophthora | G4ZTF9 |
| Picea sitchen | B8LRA5 |
| Picea sitchen | B8LRG3 |
| Pinctada fuca | A0A194ANQ8 |
| Pipistrellus al | L0N3W3 |
| Piromyces fin | A0A1Y1VF86 |
| Piromyces fin | A0A1Y1V4N4 |
| Piromyces sp | A0A1Y3NGF1 |
| Planctomycet | A0A2D9PRI4 |
| Planoprotost | A0A2P6NN51 |
| Planoprotost | A0A2P6NBD6 |
| Planoprotost | A0A2P6NS16 |
| Planoprotost | A0A2P6NFX0 |
| Planoprotost | A0A2P6NBF7 |
| Planoprotost | A0A2P6NM78 |
| Planoprotost | A0A2P6NZX6 |
| Planoprotost | A0A2P6NQW3 |
| Planoprotost | A0A2P6NJ93 |
| Planoprotost | A0A2P6NKK7 |
| Planoprotost | A0A2P6NXQ1 |
| Planoprotost | A0A2P6N084 |
| Planoprotost | A0A2P6N8A1 |
| Planoprotost | A0A2P6MQS5 |
| Planoprotost | A0A2P6NWD1 |
| Planoprotost | A0A2P6N6L8 |
| Planoprotost | A0A2P6NUU8 |
| Planoprotost | A0A2P6NYV0 |
| Planoprotost | A0A2P6MVG3 |
| Planoprotost | A0A2P6N8R1 |
| Planoprotost | A0A2P6N8J2 |
| Planoprotost | A0A2P6NNF7 |
| Planoprotost | A0A2P6N1U3 |
| Planoprotost | A0A2P6NZG2 |
| Planoprotost | A0A2P6NJU9 |
| Planoprotost | A0A2P6NU31 |
| Planoprotost | A0A2P6NNX6 |
| Planoprotost | A0A2P6MNV7 |
| Planoprotost | A0A2P6N4J7 |
| Planoprotost | A0A2P6P052 |
| Planoprotost | A0A2P6NPL0 |

| | |
|---|---|
| Planoprotost | A0A2P6N398 |
| Planoprotost | A0A2P6NVG2 |
| Planoprotost | A0A2P6MWF9 |
| Planoprotost | A0A2P6NLH5 |
| Planoprotost | A0A2P6N891 |
| Planoprotost | A0A2P6NKS8 |
| Planoprotost | A0A2P6NRU2 |
| Planoprotost | A0A2P6NI63 |
| Planoprotost | A0A2P6NPR4 |
| Planoprotost | A0A2P6NBF0 |
| Planoprotost | A0A2P6NL20 |
| Planoprotost | A0A2P6NRH4 |
| Planoprotost | A0A2P6NWM6 |
| Planoprotost | A0A2P6MUC2 |
| Planoprotost | A0A2P6MQR6 |
| Planoprotost | A0A2P6NL12 |
| Planoprotost | A0A2P6NVA7 |
| Planoprotost | A0A2P6NKZ3 |
| Planoprotost | A0A2P6MZC6 |
| Planoprotost | A0A2P6NII8 |
| Planoprotost | A0A2P6P0D1 |
| Planoprotost | A0A2P6NUJ5 |
| Planoprotost | A0A2P6NMA2 |
| Planoprotost | A0A2P6NGP7 |
| Planoprotost | A0A2P6MT35 |
| Planoprotost | A0A2P6MSH3 |
| Planoprotost | A0A2P6NK96 |
| Planoprotost | A0A2P6NR97 |
| Plasmopara h | A0A0P1B1W3 |
| Plasmopara h | A0A0P1A668 |
| Plasmopara h | A0A0P1B457 |
| Plasmopara h | A0A0P1ALX1 |
| Platycephalus | D0VX36 |
| Platycephalus | D0VX37 |
| Platynereis d | A0A411F668 |
| Pleomorphor | A0A2G9WWN9 |
| Pleomorphor | A0A1I4U8D5 |
| Pleomorphor | A0A2Z6HWP3 |
| Pleomorphor | A0A2Z6I0M2 |
| Pochonia chla | A0A179F4S7 |
| Pochonia chla | A0A179F441 |

| | |
|---|---|
| Pocillopora d | A0A3M6TS79 |
| Pocillopora d | A0A3M6USF2 |
| Pocillopora d | A0A3M6U8L5 |
| Pocillopora d | A0A3M6T8B8 |
| Pocillopora d | A0A3M6UTQ7 |
| Pocillopora d | A0A3M6V1M2 |
| Pocillopora d | A0A3M6TMM1 |
| Pocillopora d | A0A3M6T9H4 |
| Pocillopora d | A0A3M6UUH4 |
| Pocillopora d | A0A3M6UCX4 |
| Pocillopora d | A0A3M6TR63 |
| Pocillopora d | A0A3M6TQV1 |
| Pocillopora d | A0A3M6UUS3 |
| Pocillopora d | A0A3M6T9L3 |
| Pocillopora d | A0A3M6UH21 |
| Pocillopora d | A0A3M6UFJ0 |
| Pocillopora d | A0A3M6TLA8 |
| Pocillopora d | A0A3M6TXJ3 |
| Pocillopora d | A0A3M6UNB9 |
| Pocillopora d | A0A3M6TV38 |
| Pocillopora d | A0A3M6UY98 |
| Pocillopora d | A0A3M6UGR8 |
| Pocillopora d | A0A3M6UHN8 |
| Pocillopora d | A0A3M6TQC2 |
| Pocillopora d | A0A3M6UZT4 |
| Pocillopora d | A0A3M6TG14 |
| Pocillopora d | A0A3M6UAH9 |
| Pocillopora d | A0A3M6U389 |
| Pocillopora d | A0A3M6T9B4 |
| Pocillopora d | A0A3M6TK43 |
| Pocillopora d | A0A3M6TKR9 |
| Pocillopora d | A0A3M6UW14 |
| Pocillopora d | A0A3M6USD4 |
| Pocillopora d | A0A3M6TZG9 |
| Pocillopora d | A0A3M6UIH2 |
| Pocillopora d | A0A3M6UHX1 |
| Pocillopora d | A0A3M6TJU1 |
| Pocillopora d | A0A3M6TLI2 |
| Pocillopora d | A0A3M6UQ55 |
| Pocillopora d | A0A3M6UM67 |
| Pocillopora d | A0A3M6T7C9 |

| | |
|---|---|
| Pocillopora d | A0A3M6T879 |
| Pocillopora d | A0A3M6TMQ8 |
| Pocillopora d | A0A3M6U799 |
| Pocillopora d | A0A3M6UYP3 |
| Pocillopora d | A0A3M6UY17 |
| Pocillopora d | A0A3M6V3Y0 |
| Pocillopora d | A0A3M6UUQ2 |
| Pocillopora d | A0A3M6TKU3 |
| Pocillopora d | A0A3M6UYG1 |
| Pocillopora d | A0A3M6TTJ5 |
| Pocillopora d | A0A3M6TY63 |
| Pocillopora d | A0A3M6TFR8 |
| Pocillopora d | A0A3M6U3C3 |
| Pocillopora d | A0A3M6U1C9 |
| Pocillopora d | A0A3M6U041 |
| Pocillopora d | A0A3M6U1S3 |
| Pocillopora d | A0A3M6TKF4 |
| Pocillopora d | A0A3M6TTS0 |
| Pocillopora d | A0A3M6U9P4 |
| Pocillopora d | A0A3M6TU17 |
| Pocillopora d | A0A3M6V0G4 |
| Pocillopora d | A0A3M6UKJ5 |
| Pocillopora d | A0A3M6U9D2 |
| Pocillopora d | A0A3M6UMI0 |
| Pocillopora d | A0A3M6V1P4 |
| Pocillopora d | A0A3M6TZE9 |
| Pocillopora d | A0A3M6U9C4 |
| Pocillopora d | A0A3M6UK43 |
| Pocillopora d | A0A3M6TDQ0 |
| Pocillopora d | A0A3M6UQ00 |
| Pocillopora d | A0A3M6T9N0 |
| Pocillopora d | A0A3M6TL38 |
| Pocillopora d | A0A3M6UGQ2 |
| Pocillopora d | A0A3M6T5E6 |
| Pocillopora d | A0A3M6TYN4 |
| Pocillopora d | A0A3M6UZA1 |
| Pocillopora d | A0A3M6V499 |
| Pocillopora d | A0A3M6TZ45 |
| Pocillopora d | A0A3M6T7Z7 |
| Pocillopora d | A0A3M6UQN2 |
| Pocillopora d | A0A3M6V0A1 |

| | |
|---|---|
| Pocillopora d | A0A3M6UAD8 |
| Pocillopora d | A0A3M6TA22 |
| Pocillopora d | A0A3M6UL69 |
| Pocillopora d | A0A3M6V562 |
| Pocillopora d | A0A3M6UHQ1 |
| Pocillopora d | A0A3M6V6G0 |
| Pocillopora d | A0A3M6U1J0 |
| Pocillopora d | A0A3M6ULX9 |
| Pocillopora d | A0A3M6TX33 |
| Pocillopora d | A0A3M6V3R3 |
| Pocillopora d | A0A3M6UG70 |
| Pocillopora d | A0A3M6TFV1 |
| Pocillopora d | A0A3M6TCE3 |
| Pocillopora d | A0A3M6U1L3 |
| Pocillopora d | A0A3M6USJ5 |
| Pocillopora d | A0A3M6UV09 |
| Pocillopora d | A0A3M6THX6 |
| Pocillopora d | A0A3M6TSD4 |
| Pocillopora d | A0A3M6UHD5 |
| Pocillopora d | A0A3M6TIM9 |
| Pocillopora d | A0A3M6V609 |
| Pocillopora d | A0A3M6UMN1 |
| Pocillopora d | A0A3M6T7T9 |
| Pocillopora d | A0A3M6TRU4 |
| Pocillopora d | A0A3M6UPF6 |
| Pocillopora d | A0A3M6TA00 |
| Pocillopora d | A0A3M6TTM9 |
| Pocillopora d | A0A3M6T9S3 |
| Pocillopora d | A0A3M6V4P3 |
| Pocillopora d | A0A3M6URI8 |
| Pocillopora d | A0A3M6T6W2 |
| Pocillopora d | A0A3M6TGI7 |
| Pocillopora d | A0A3M6U3G3 |
| Pocillopora d | A0A3M6UL17 |
| Pocillopora d | A0A3M6UCE1 |
| Pocillopora d | A0A3M6USZ6 |
| Pocillopora d | A0A3M6UMF9 |
| Pocillopora d | A0A3M6TET4 |
| Pocillopora d | A0A3M6TG44 |
| Pocillopora d | A0A3M6U3J1 |
| Pocillopora d | A0A3M6UXV8 |

Pocillopora d A0A3M6TWK8
Pocillopora d A0A3M6UV46
Pocillopora d A0A3M6UL16
Pocillopora d A0A3M6ULU8
Pocillopora d A0A3M6T6A8
Pocillopora d A0A3M6U377
Pocillopora d A0A3M6TAL1
Pocillopora d A0A3M6T5X9
Pocillopora d A0A3M6V0S5
Pocillopora d A0A3M6UZX3
Pocillopora d A0A3M6V292
Pocillopora d A0A3M6USE0
Pocillopora d A0A3M6V1C4
Pocillopora d A0A3M6TRT6
Pocillopora d A0A3M6TGP2
Pocillopora d A0A3M6TNZ6
Pocillopora d A0A3M6U6B3
Pocillopora d A0A3M6TYI9
Pocillopora d A0A3M6U5N1
Pocillopora d A0A3M6TZH8
Pocillopora d A0A3M6TZB0
Pocillopora d A0A3M6U1T5
Pocillopora d A0A3M6UTP6
Pocillopora d A0A3M6V3G9
Pocillopora d A0A3M6UAU9
Pocillopora d A0A3M6UVP6
Pocillopora d A0A3M6UUS4
Pocillopora d A0A3M6U1B0
Pocillopora d A0A3M6U9H5
Pocillopora d A0A3M6U252
Pocillopora d A0A3M6T542
Pocillopora d A0A3M6UJV7
Pocillopora d A0A3M6TA55
Pocillopora d A0A3M6T9T7
Pocillopora d A0A3M6TWX2
Pocillopora d A0A3M6V721
Pocillopora d A0A3M6U9I9
Pocillopora d A0A3M6U1G7
Pocillopora d A0A3M6U527
Pocillopora d A0A3M6U1R3
Pocillopora d A0A3M6V1Z1

Pocillopora d A0A3M6UQ36
Pocillopora d A0A3M6TI45
Pocillopora d A0A3M6UIZ7
Pocillopora d A0A3M6UMS9
Pocillopora d A0A3M6UY19
Pocillopora d A0A3M6UPE3
Pocillopora d A0A3M6TNQ1
Pocillopora d A0A3M6UMU0
Pocillopora d A0A3M6TGA7
Pocillopora d A0A3M6TX72
Pocillopora d A0A3M6V494
Pocillopora d A0A3M6UG64
Pocillopora d A0A3M6UK86
Pocillopora d A0A3M6UYI3
Pocillopora d A0A3M6V0I9
Pocillopora d A0A3M6T8D1
Pocillopora d A0A3M6TZS0
Pocillopora d A0A3M6TCK3
Pocillopora d A0A3M6UF72
Pocillopora d A0A3M6URU5
Pocillopora d A0A3M6U5L9
Pocillopora d A0A3M6UQ23
Pocillopora d A0A3M6U9G1
Pocillopora d A0A3M6UGB6
Pocillopora d A0A3M6UUP1
Pocillopora d A0A3M6TGX0
Pocillopora d A0A3M6UKV3
Pocillopora d A0A3M6U1N8
Pocillopora d A0A3M6V463
Pocillopora d A0A3M6UKM5
Pocillopora d A0A3M6TYF4
Pocillopora d A0A3M6U0N0
Pocillopora d A0A3M6UMN0
Pocillopora d A0A3M6U2G5
Pocillopora d A0A3M6T9Q3
Pocillopora d A0A3M6TY60
Pocillopora d A0A3M6UCM1
Pocillopora d A0A3M6TVJ0
Pocillopora d A0A3M6U3I0
Pocillopora d A0A3M6VBM0
Pocillopora d A0A3M6TTN4

| | |
|---|---|
| Pocillopora d | A0A3M6TST7 |
| Pocillopora d | A0A3M6V143 |
| Pocillopora d | A0A3M6TTD9 |
| Pocillopora d | A0A3M6U218 |
| Pocillopora d | A0A3M6U1Q8 |
| Pocillopora d | A0A3M6T410 |
| Pocillopora d | A0A3M6UKS4 |
| Pocillopora d | A0A3M6T9Q5 |
| Pocillopora d | A0A3M6US86 |
| Pocillopora d | A0A3M6TYB9 |
| Pocillopora d | A0A3M6TTW0 |
| Pocillopora d | A0A3M6TCD6 |
| Pocillopora d | A0A3M6UMC9 |
| Pocillopora d | A0A3M6USH7 |
| Pocillopora d | A0A3M6TD69 |
| Pocillopora d | A0A3M6UK11 |
| Pocillopora d | A0A3M6UVR5 |
| Pocillopora d | A0A3M6U2Q5 |
| Pocillopora d | A0A3M6V3V5 |
| Pocillopora d | A0A3M6UBX9 |
| Pocillopora d | A0A3M6UK88 |
| Pocillopora d | A0A3M6UUI1 |
| Pocillopora d | A0A3M6TKL2 |
| Pocillopora d | A0A3M6V010 |
| Pocillopora d | A0A3M6TK69 |
| Pocillopora d | A0A3M6TDY5 |
| Pocillopora d | A0A3M6TIX2 |
| Pocillopora d | A0A3M6TYN3 |
| Podiceps cris | A0A094L7R4 |
| Podiceps cris | A0A094NEK3 |
| Podiceps cris | A0A094KG51 |
| Podiceps cris | A0A094LT96 |
| Podospora ar | B2B3F1 |
| Podospora ar | B2AEW5 |
| Podospora ar | B2ASY6 |
| Podospora ar | B2B631 |
| Podospora ar | A0A090CJ60 |
| Podospora cc | A0A447CBI5 |
| Podospora cc | A0A447CFK1 |
| Podospora cc | A0A447C1B4 |
| Podospora cc | A0A447CHJ2 |

| | |
|---|---|
| Poecilia form | A0A087YC08 |
| Poecilia form | A0A087YDJ9 |
| Poecilia form | A0A096M9X7 |
| Poecilia form | A0A096MHY9 |
| Poecilia form | A0A096LUY1 |
| Poecilia form | A0A087XAX0 |
| Poecilia form | A0A087YK42 |
| Poecilia form | A0A096M7R2 |
| Poecilia form | A0A087YBW9 |
| Poecilia latipi | A0A3B3V3Z1 |
| Poecilia latipi | A0A3B3TJZ9 |
| Poecilia latipi | A0A3B3V421 |
| Poecilia latipi | A0A3B3V1M0 |
| Poecilia latipi | A0A3B3UUE6 |
| Poecilia latipi | A0A3B3W2N5 |
| Poecilia latipi | A0A3B3VXR8 |
| Poecilia latipi | A0A3B3TK34 |
| Poecilia latipi | A0A3B3USD0 |
| Poecilia latipi | A0A3B3TX33 |
| Poecilia latipi | A0A3B3VD97 |
| Poecilia latipi | A0A3B3V427 |
| Poecilia latipi | A0A3B3TZ12 |
| Poecilia mexi | A0A3B3XYA2 |
| Poecilia mexi | A0A3B3YGN7 |
| Poecilia mexi | A0A3B3YU59 |
| Poecilia mexi | A0A3B3YUM0 |
| Poecilia mexi | A0A3B3XXC4 |
| Poecilia mexi | A0A3B3XJP2 |
| Poecilia mexi | A0A3B3XXG3 |
| Poecilia mexi | A0A3B3X0F3 |
| Poecilia mexi | A0A3B3XY03 |
| Poecilia mexi | A0A3B3YUR5 |
| Poecilia mexi | A0A3B3XRS1 |
| Poecilia mexi | A0A3B3Z3V6 |
| Poecilia mexi | A0A3B3WRD1 |
| Poecilia mexi | A0A3B3YVC7 |
| Poecilia retic | A0A3P9Q3Q9 |
| Poecilia retic | A0A3P9Q2T6 |
| Poecilia retic | A0A3P9NA69 |
| Poecilia retic | A0A3P9Q2U6 |
| Poecilia retic | A0A3P9Q2Z7 |

| | |
|---|---|
| Poecilia reticı | A0A3P9Q320 |
| Poecilia reticı | A0A3P9Q335 |
| Poecilia reticı | A0A3P9Q431 |
| Poecilia reticı | A0A3P9Q3Q0 |
| Poecilia reticı | A0A3P9Q4J8 |
| Poecilia reticı | A0A3P9Q3N5 |
| Poecilia reticı | A0A3P9PH03 |
| Poecilia reticı | A0A3P9Q356 |
| Poeciliopsis p | A0A0S7K5B4 |
| Poeciliopsis p | A0A0S7J5R7 |
| Poeciliopsis p | A0A0S7K6Q0 |
| Poeciliopsis p | A0A0S7KT15 |
| Poeciliopsis p | A0A0S7K5P7 |
| Poeciliopsis p | A0A0S7M8P1 |
| Poeciliopsis p | A0A0S7L542 |
| Poeciliopsis p | A0A0S7ELW6 |
| Poeciliopsis p | A0A0S7K6K9 |
| Poeciliopsis p | A0A0S7K5V9 |
| Poeciliopsis p | A0A0S7M7Y8 |
| Poeciliopsis p | A0A0S7EN80 |
| Poeciliopsis p | A0A0S7EV87 |
| Pohlia nutans | A0A125SL75 |
| Pohlia nutans | A0A125SL77 |
| Polyandrocar | Q9Y1V3 |
| Polymorphun | F2IW28 |
| Polytolypa hy | A0A2B7YVT1 |
| Polytolypa hy | A0A2B7Y009 |
| Polytolypa hy | A0A2B7XRG7 |
| Pomacea can | A0A2T7PUF8 |
| Pomacea can | A0A2T7P4A6 |
| Pomacea can | A0A2T7P2C2 |
| Pomacea can | A0A2T7NW21 |
| Pomacea can | A0A2T7NVT2 |
| Pomacea can | A0A2T7P293 |
| Pomacea can | A0A2T7NRV8 |
| Pomacea can | A0A2T7P283 |
| Pomacea can | A0A2T7NUX7 |
| Pomacea can | A0A2T7NZW8 |
| Pomacea can | A0A2T7P282 |
| Pomacea can | A0A2T7NPR7 |
| Pomacea can | A0A2T7NP19 |

| | |
|---|---|
| Pomacea can | A0A2T7PDK6 |
| Pomacea can | A0A2T7PBM0 |
| Pomacea can | A0A2T7NH27 |
| Pomacea can | A0A2T7NGZ5 |
| Pomacea can | A0A2T7PVC4 |
| Pomacea can | A0A2T7P296 |
| Pomacea can | A0A2T7NPP6 |
| Pomacea can | A0A2T7PIV1 |
| Pomacea can | A0A2T7NT09 |
| Pomacea can | A0A2T7P290 |
| Pomacea can | A0A2T7P2A0 |
| Pomacea can | A0A2T7NT13 |
| Pomacea can | A0A2T7PXQ6 |
| Pomacea can | A0A2T7P2A4 |
| Pomacea can | A0A2T7P9Q0 |
| Pomacea can | A0A2T7NDL4 |
| Pomacea can | A0A2T7NMC9 |
| Pomacea can | A0A2T7PII0 |
| Pomacea can | A0A2T7Q143 |
| Pomacea can | A0A2T7P231 |
| Pomacea can | A0A2T7NAQ2 |
| Pomacea can | A0A2T7NUW9 |
| Pomacea can | A0A2T7PVQ3 |
| Pomacea can | A0A2T7PQ77 |
| Pongo abelii.: | A0A2J8T957 |
| Pongo abelii.: | K7EVW6 |
| Pongo abelii.: | A0A2J8VLI2 |
| Pongo abelii.: | A0A2J8VLJ4 |
| Pongo abelii.: | A0A2J8T3S9 |
| Pongo abelii.: | A0A2J8VLH8 |
| Pongo abelii.: | A0A2J8T3R6 |
| Pongo abelii.: | H2PKT3 |
| Pongo abelii.: | A0A2J8T3Q1 |
| Pongo abelii.: | A0A2J8VLJ0 |
| Pongo abelii.: | A0A2J8TGX6 |
| Pongo abelii.: | A0A2J8T3Q9 |
| Pongo abelii.: | H2PAR3 |
| Pongo abelii.: | A0A2J8T3Q0 |
| Pongo abelii.: | A0A2J8VLK6 |
| Pongo abelii.: | A0A2J8VLJ5 |
| Pongo abelii.: | A0A2J8TGY3 |

| | |
|---|---|
| Pongo abelii.: | H2PEX4 |
| Pongo abelii.: | A0A2J8VLK7 |
| Pongo abelii.: | A0A2J8T3Q5 |
| Pongo abelii.: | A0A2J8T3T1 |
| Pongo abelii.: | Q5R976 |
| Pongo abelii.: | A0A2J8T3R5 |
| Pongo abelii.: | Q5R502 |
| Pongo abelii.: | H2PEX7 |
| Pongo abelii.: | H2PN16 |
| Pongo abelii.: | A0A2J8T950 |
| Pongo abelii.: | Q5R8X6 |
| Pontibaca me | A0A1R3WSM0 |
| Ponticaulis sp | A0A2D9RCA7 |
| Ponticaulis sp | A0A2D6GQ64 |
| Ponticaulis sp | A0A2D7IW65 |
| Ponticaulis sp | A0A2D9RE05 |
| Ponticoccus r | A0A0W7WIT4 |
| Ponticoccus r | A0A0W7WNN3 |
| Ponticoccus r | A0A0W7WKJ9 |
| Ponticoccus r | A0A0W7WED2 |
| Populus tome | A0A385JHS5 |
| Populus tome | A0A3S9LX70 |
| Populus tome | A0A385JID9 |
| Populus tome | A0A385JIC9 |
| Populus tome | A0A385JIW5 |
| Populus tome | A0A3S9LX83 |
| Populus tome | A0A3S9LX72 |
| Populus tome | A0A385JIF5 |
| Populus tome | A0A3S9LX97 |
| Populus tome | A0A1L6K4F6 |
| Populus tome | A0A3S9LX71 |
| Populus tome | A0A3S9LX61 |
| Populus tome | A0A3S9LXA8 |
| Populus tome | A0A3S9LX98 |
| Populus tome | A0A3S9LXB3 |
| Populus tome | A0A3S9LX57 |
| Populus tome | A0A3S9LXA3 |
| Populus trich | A9PJX6 |
| Populus trich | A0A3N7FJX8 |
| Populus trich | B9HYR9 |
| Populus trich | A0A3N7FA41 |

Populus trich A0A2K1Y5Z8
Populus trich U5FEZ9
Populus trich U7E2S0
Populus trich A0A3N7FPW1
Populus trich A0A2K2CBM7
Populus trich A0A3N7H0W6
Populus trich A0A3N7FQG9
Populus trich B9I033
Populus trich A0A2K1YMT8
Populus trich A0A2K2CD15
Populus trich A0A2K1YMD4
Populus trich A0A2K1YF78
Populus trich A0A2K1YF67
Populus trich B9HYQ3
Populus trich A0A2K2AP05
Populus trich A0A2K2CBR0
Populus trich A0A3N7G2Z5
Populus trich U5GW32
Populus trich A0A2K1YMF9
Populus trich A0A2K1YMF8
Populus trich A0A2K2CBV8
Populus trich U7DXY3
Populus trich A0A2K1YLY5
Populus trich A0A3N7H7G1
Populus trich B9HYS3
Populus trich A0A2K1YJB9
Populus trich U7E151
Populus trich A0A2K1R9J9
Populus trich A0A2K2CBS5
Populus trich A0A2K1XDN9
Populus trich A0A3N7FS13
Populus trich U5G9S9
Populus trich A0A2K1YF84
Populus trich B9I035
Populus trich B9NGV9
Populus trich A0A3N7EEZ9
Populus trich A0A3N7ETB6
Populus trich A0A3N7GAS2
Populus trich U7DV48
Populus trich A0A3N7EEW7
Populus trich A0A3N7G946

Populus trich A0A2K1YMS1
Populus trich A0A2K2AP11
Populus trich A0A3N7GLP6
Populus trich A0A3N7FPY6
Populus trich A0A3N7FJT6
Populus trich U7DVW0
Populus trich A0A2K2BAE9
Populus trich A0A3N7FRW4
Populus trich A0A3N7GIT0
Populus trich A0A2K1YF54
Populus trich A9PFH0
Populus trich A0A3N7FPB6
Populus trich A0A2K2CBN3
Populus trich A0A2K1YMF6
Populus trich A0A2K1YF72
Populus trich A0A2K1YF37
Populus trich A0A3N7FK42
Populus trich A0A2K1YJI6
Populus trich A0A2K2CBU0
Populus trich A0A3N7FQ27
Populus trich A0A2K1YMF7
Populus trich A0A2K1YMK1
Populus trich A0A2K1R9K1
Populus trich U7DZT8
Populus trich A0A2K2CBV2
Populus trich A0A2K1YRU7
Populus trich U5FQT0
Populus trich A0A2K1R435
Populus trich A0A2K2CBU5
Populus trich A0A3N7FKB6
Populus trich A0A3N7EYY1
Populus trich A0A3N7F931
Populus trich A0A2K1R9M9
Populus trich A0A3N7HDJ2
Populus trich A0A3N7H721
Populus trich A0A2K2CBR5
Populus trich A0A2K1YMD5
Populus trich U7DWR5
Populus trich U7E0R0
Populus trich A0A3N7G378
Populus trich A0A3N7G0M4

Populus trich A0A2K2CBS7
Populus trich U5GSZ3
Populus trich A0A2K1YF20
Populus trich A0A2K1YCZ6
Populus trich A0A2K2AP00
Populus trich A0A2K2CBS4
Populus trich A0A2K1R9L1
Populus trich A0A2K1R9K8
Populus trich A0A2K1YF88
Populus trich A0A2K2CBR3
Populus trich A0A3N7EYL9
Populus trich A0A3N7FPG8
Populus trich U7DYK7
Populus trich A0A3N7EJM8
Populus trich A0A2K1YMF0
Populus trich B9IFY4
Populus trich A0A2K1R9K3
Populus trich A0A2K1R9L4
Populus trich U7E198
Populus trich A0A3N7FBU8
Populus trich A0A3N7HE18
Populus trich U7E213
Populus trich A0A3N7FRQ4
Populus trich A0A2K1R7A4
Populus trich U7DZU3
Populus trich A0A3N7FTM4
Populus trich A0A2K1YJJ2
Populus trich A0A2K2CAQ6
Populus trich A0A3N7F920
Populus trich A0A3N7FQE7
Populus trich A0A2K2AAA5
Populus trich B9HAE5
Populus trich A0A2K1XB50
Populus trich A0A2K1YMH4
Populus trich A0A2K1YJA7
Populus trich A0A2K1YMF3
Populus trich U5FZN7
Populus trich A0A2K1WRN2
Populus trich A0A3N7EEZ8
Populus trich A0A2K1R9I9
Populus trich B9I048

Populus trich A0A2K1R9K2
Populus trich A0A2K2CBW8
Populus trich A0A3N7FBT5
Populus trich A0A2K1YF32
Populus trich U5FP35
Populus trich A0A3N7H774
Populus trich A0A3N7FUN1
Populus trich A9PF61
Populus trich A0A2K1YJJ1
Populus trich A0A2K2AQM5
Populus trich A0A2K1YF42
Populus trich A0A2K1R9J8
Populus trich A0A3N7ERB7
Populus trich A0A2K1ZA48
Populus trich A0A3N7FRY4
Populus trich A0A2K2AAA9
Populus trich A0A3N7G2Z7
Populus trich A0A2K1YF57
Populus trich A0A3N7EBP4
Populus trich A0A2K1R5I9
Populus trich A0A3N7EXE8
Populus trich B9INT4
Populus trich B9MU91
Populus trich A0A3N7EII0
Populus trich A0A3N7HDF5
Populus trich A0A2K1YF71
Populus trich A0A2K2AAC8
Populus trich A0A2K2CBV0
Populus trich A0A2K1R9L6
Populus trich A0A2K1YF62
Populus trich A0A3N7EMG0
Populus trich A0A3N7GEG5
Populus trich A0A3N7FKU0
Populus trich A0A2K2CC26
Populus trich A0A2K1R9K7
Populus trich A0A2K1R9L9
Populus trich A0A3N7FQE2
Populus trich A0A3N7FE30
Populus trich B9IQL6
Populus trich B9HYQ6
Populus trich A0A2K2CBS0

Populus trich A0A2K1Y5X3
Populus trich A0A2K2CBN5
Populus trich A0A3N7I1Z5
Populus trich A0A2K1R9K4
Populus trich U7E275
Populus trich A0A2K2CBV3
Populus trich A0A3N7GT81
Populus trich A0A3N7FK00
Populus trich A0A2K1YJB7
Populus trich A0A3N7GCP1
Populus trich A0A3N7FXT6
Populus trich A0A3N7FWX1
Populus trich A0A2K2CBR2
Populus trich U7E2B5
Populus trich A0A2K1YJJ4
Populus trich A0A2K2CBQ6
Populus trich A0A2K1Y5Y0
Populus trich U7DTV7
Populus trich A0A2K1R9J0
Populus trich A0A2K1YJC0
Populus trich A0A3N7EN77
Populus trich A0A3N7HDH1
Populus trich A0A2K2CBV5
Populus trich A0A2K1YMS3
Populus trich A0A3N7H760
Populus trich A0A2K2CBU2
Populus trich A0A3N7FMC7
Populus trich A0A3N7FFN1
Populus trich A0A2K1YII9
Populus trich A0A2K1YFA5
Populus trich A0A2K1YMG3
Populus trich Q0ZCC5
Populus trich A0A3N7F1A3
Populus trich A0A2K1YMC5
Populus trich A0A2K1R9L8
Populus trich A0A2K2CBS9
Populus trich A0A2K1YF89
Populus trich A0A2K1YJB0
Populus trich A0A3N7G5C0
Populus trich B9I819
Populus trich A0A3N7EZT8

Populus trich B9H1V2
Populus trich A0A2K1YF22
Populus trich A0A2K1YKS5
Populus trich A0A3N7FQ08
Populus trich A0A2K2CBS8
Populus trich A0A2K2AC56
Populus trich A0A2K1YF85
Populus trich A0A2K2AA68
Populus trich A0A2K2CBS3
Populus trich A0A2K1YF29
Populus trich A0A2K1YF09
Populus trich U7DZK5
Populus trich A0A2K2AA71
Populus trich A0A2K2CBQ2
Populus trich A0A3N7F8U4
Populus trich B9INU0
Populus trich A0A2K2CD09
Populus trich A0A2K2BQD8
Populus trich A0A2K2CBR1
Populus trich A0A2K1YME5
Populus trich A0A3N7FHR5
Populus trich A0A2K1YIJ7
Populus trich U7DSM0
Populus trich B9MUS9
Populus trich B9H1U4
Populus trich A0A3N7FQ25
Populus trich A0A2K1YF75
Populus trich A0A2K1YME2
Populus trich A0A2K1YF38
Populus trich A0A3N7G9D1
Populus trich B9H9X2
Populus trich B9I050
Populus trich A0A2K1R9I3
Populus trich A0A3N7F8V7
Populus trich A0A2K1WUY0
Populus trich A0A2K2AP04
Populus trich A0A3N7FUP4
Populus trich A0A2K2AP06
Populus trich A0A3N7FF46
Populus trich A0A3N7FTN5
Populus trich A0A2K1YF58

| | |
|---|---|
| Porphyra um | A0A1X6NVT8 |
| Porphyra um | A0A1X6NQM4 |
| Porphyra um | A0A1X6PFD2 |
| Porphyra um | A0A1X6P8C6 |
| Porphyra um | A0A1X6NNT9 |
| Porphyra um | A0A1X6P5P1 |
| Porphyrobact | A0A0N0JQI8 |
| Powai lake m | A0A167RBV0 |
| Powai lake m | A0A167R7I2 |
| Powai lake m | A0A167RBU3 |
| Primorskyiba | A0A2R8AUK0 |
| Pristionchus | H3FLN2 |
| Pristionchus | H3FU64 |
| Pristionchus | A0A2A6CLV0 |
| Pristionchus | H3G269 |
| Pristionchus | H3DYL1 |
| Pristionchus | A0A2A6BKN6 |
| Pristionchus | A0A2A6CL96 |
| Pristionchus | H3F912 |
| Pristionchus | A0A2A6CKS7 |
| Pristionchus | H3DXL2 |
| Pristionchus | H3EPT2 |
| Pristionchus | A0A2A6BKX0 |
| Pristionchus | A0A2A6BRD4 |
| Pristionchus | H3EMR6 |
| Pristionchus | H3F239 |
| Pristionchus | H3FWN3 |
| Pristionchus | H3EMS5 |
| Pristionchus | H3E6Q3 |
| Pristionchus | A0A2A6CW20 |
| Pristionchus | H3G484 |
| Pristionchus | A0A2A6CWJ1 |
| Pristionchus | A0A2A6BQE5 |
| Pristionchus | H3G483 |
| Pristionchus | A0A2A6C8D4 |
| Pristionchus | A0A2A6CWL7 |
| Pristionchus | H3F6C9 |
| Pristionchus | A0A2A6BXL8 |
| Pristionchus | A0A2A6BTB7 |
| Pristionchus | H3E9C6 |
| Pristionchus | H3F0T6 |

Pristionchus | A0A2A6CRJ5
Pristionchus | H3FU41
Pristionchus | A0A2A6CC90
Pristionchus | A0A2A6CXB5
Pristionchus | H3EUD8
Pristionchus | H3EUB5
Pristionchus | A0A2A6CVS9
Pristionchus | A0A2A6C5D7
Pristionchus | H3EYA4
Pristionchus | A0A2A6CNU0
Pristionchus | A0A2A6CPM8
Pristionchus | H3FXU2
Pristionchus | H3FXP1
Pristionchus | H3FH55
Pristionchus | H3FDU0
Pristionchus | H3DRR5
Pristionchus | H3F6D1
Pristionchus | A0A2A6C0F3
Pristionchus | A0A2A6CJP0
Pristionchus | H3FUP4
Pristionchus | H3EC28
Pristionchus | A0A2A6C3M2
Pristionchus | H3E6Q4
Pristionchus | A0A2A6BWE1
Pristionchus | A0A2A6CV10
Pristionchus | A0A2A6C2I9
Pristionchus | A0A2A6BTN5
Pristionchus | H3FDT3
Pristionchus | A0A2A6BRT5
Pristionchus | H3EUB4
Pristionchus | A0A2A6BRY8
Pristionchus | H3EIL7
Pristionchus | A0A2A6B687
Pristionchus | H3F237
Pristionchus | A0A2A6B8K1
Pristionchus | A0A2A6CBV8
Pristionchus | A0A2A6BIV2
Pristionchus | A0A2A6BTB0
Pristionchus | H3EA22
Pristionchus | A0A2A6CPA6
Pristionchus | A0A2A6D2U8

| | |
|---|---|
| Pristionchus | A0A2A6CLQ3 |
| Pristionchus | H3G4B7 |
| Pristionchus | A0A2A6BP16 |
| Pristionchus | H3FP67 |
| Pristionchus | A0A2A6B6P2 |
| Pristionchus | A0A454XRJ9 |
| Pristionchus | A0A2A6CS22 |
| Pristionchus | A0A2A6BMJ9 |
| Pristionchus | A0A2A6CVF2 |
| Pristionchus | A0A2A6C5B7 |
| Pristionchus | H3FKW8 |
| Pristionchus | H3E9C4 |
| Pristionchus | H3FLR8 |
| Pristionchus | A0A2A6CL26 |
| Pristionchus | A0A2A6CAV6 |
| Pristionchus | A0A2A6C0V0 |
| Pristionchus | H3DT54 |
| Pristionchus | A0A2A6C9M6 |
| Pristionchus | H3F624 |
| Pristionchus | A0A2A6CIU4 |
| Pristionchus | H3G421 |
| Pristionchus | H3EKC8 |
| Pristionchus | A0A2A6CYT9 |
| Pristionchus | A0A2A6CJH0 |
| Pristionchus | A0A2A6CTB6 |
| Pristionchus | H3FFI6 |
| Pristionchus | A0A2A6BY19 |
| Pristionchus | A0A2A6BF62 |
| Pristionchus | A0A2A6C2U7 |
| Pristionchus | A0A2A6C8U6 |
| Pristionchus | A0A2A6CM39 |
| Pristionchus | A0A2A6BTW0 |
| Pristionchus | A0A2A6B724 |
| Pristionchus | H3EWG7 |
| Pristionchus | A0A2A6CGU1 |
| Pristionchus | A0A2A6C3N2 |
| Pristionchus | A0A2A6BRI6 |
| Propithecus c | A0A2K6FYF0 |
| Propithecus c | A0A2K6FWB1 |
| Propithecus c | A0A2K6FWE4 |
| Propithecus c | A0A2K6GMV5 |

| | |
|---|---|
| Propithecus c | A0A2K6FWC7 |
| Propithecus c | A0A2K6GU92 |
| Propithecus c | A0A2K6FYE7 |
| Propithecus c | A0A2K6EIZ2 |
| Prosthecomic | A0A0P6W8Z7 |
| Proteobacter | A0A1W9IFG2 |
| Proteobacter | A0A1W9I8T4 |
| Proteobacter | A0A2W4P8I4 |
| Proteobacter | A0A2W4P3J5 |
| Proteobacter | A0A3A0F8E5 |
| Proteobacter | A0A317HTN1 |
| Protopolystor | A0A448WDT9 |
| Protopolystor | A0A3S5A576 |
| Prunus cerasi | E6YCZ6 |
| Prunus cerasi | E6YCZ1 |
| Prunus persic | M5WXQ1 |
| Prunus persic | A0A251PZP6 |
| Prunus persic | A0A251Q7E1 |
| Prunus persic | A0A251PF29 |
| Prunus persic | A0A251PTU7 |
| Prunus persic | M5WCL9 |
| Prunus persic | A0A251Q445 |
| Prunus persic | M5VZY3 |
| Prunus persic | A0A251PKI1 |
| Prunus persic | A0A251Q478 |
| Prunus persic | A0A251P735 |
| Prunus persic | A0A251PF35 |
| Prunus persic | M5WJ95 |
| Prunus persic | A0A251Q7C8 |
| Prunus persic | A0A251PKC2 |
| Prunus persic | M5WMF2 |
| Prunus persic | Q6DU55 |
| Prunus persic | M5X5D7 |
| Prunus persic | M5WWL5 |
| Prunus persic | M5VUG0 |
| Prunus persic | A0A251PN29 |
| Prunus persic | A0A251PMZ2 |
| Prunus persic | A0A251RBW4 |
| Prunus persic | M5WM53 |
| Prunus persic | M5WZG6 |
| Prunus persic | A0A251Q443 |

| | |
|---|---|
| Prunus persic | M5XEL7 |
| Prunus persic | M5X9Q1 |
| Prunus persic | A0A251PMZ1 |
| Prunus persic | A0A251PN38 |
| Prunus persic | A0A251REA2 |
| Prunus persic | A0A251Q484 |
| Prunus persic | A0A251RGM7 |
| Prunus persic | M5X9P7 |
| Prunus persic | A0A251PF27 |
| Prunus persic | M5VTG3 |
| Prunus persic | M5WZA7 |
| Prunus persic | M5WZT3 |
| Prunus persic | A0A251PF21 |
| Prunus persic | M5WK87 |
| Prunus persic | A0A251PZD2 |
| Prunus persic | A0A251PLL1 |
| Prunus persic | A0A251NRG7 |
| Prunus persic | A0A251PID0 |
| Prunus persic | M5WZ40 |
| Prunus persic | A0A251NHZ5 |
| Prunus persic | A0A251PWJ2 |
| Prunus persic | M5WPH8 |
| Prunus persic | M5VZT1 |
| Prunus persic | A0A251PKJ4 |
| Prunus persic | A0A251Q4B5 |
| Prunus persic | A0A251PLJ6 |
| Prunus persic | A0A251PGK7 |
| Prunus persic | M5X1J1 |
| Prunus persic | M5WG45 |
| Prunus persic | M5WD24 |
| Prunus persic | A0A251PF18 |
| Prunus persic | A0A251PN12 |
| Prunus persic | A0A251PLY0 |
| Prunus persic | A0A251PMY4 |
| Prunus persic | A0A251Q434 |
| Prunus persic | M5X8S1 |
| Prunus persic | M5X455 |
| Prunus persic | A0A251PYL5 |
| Prunus persic | M5XQT3 |
| Prunus persic | M5X887 |
| Prunus persic | M5X3N3 |

| | |
|---|---|
| Prunus persic | M5Y1R6 |
| Prunus persic | M5W6E4 |
| Prunus persic | A0A251P734 |
| Prunus persic | M5WH13 |
| Prunus persic | M5Y3U1 |
| Prunus persic | M5X159 |
| Prunus persic | M5WK95 |
| Prunus persic | A0A251PF34 |
| Prunus persic | A0A251Q485 |
| Prunus persic | M5WPP2 |
| Prunus persic | A0A251NRF5 |
| Prunus persic | A0A251PF25 |
| Prunus persic | M5WLB2 |
| Prunus persic | A0A251Q454 |
| Prunus persic | M5XAQ5 |
| Prunus persic | A0A251PN32 |
| Prunus persic | M5WPS0 |
| Prunus persic | M5X3A2 |
| Prunus persic | A0A251PN76 |
| Prunus persic | A0A251PKI0 |
| Prunus persic | A0A251PQ97 |
| Prunus persic | M5WXJ1 |
| Prunus persic | A0A251PN13 |
| Prunus persic | A0A251PF23 |
| Prunus persic | M5WKA3 |
| Prunus persic | M5WM79 |
| Prunus persic | A0A251Q5C4 |
| Prunus persic | A0A251Q4D5 |
| Prunus persic | A0A251RGX7 |
| Prunus persic | M5X9S4 |
| Prunus persic | A0A251NRE5 |
| Prunus persic | M5WXU6 |
| Prunus persic | A0A251REE6 |
| Prunus persic | A0A251PN41 |
| Prunus persic | M5XL06 |
| Prunus persic | A0A251Q458 |
| Prunus persic | M5X216 |
| Prunus persic | M5WH09 |
| Prunus persic | A0A251Q463 |
| Prunus persic | M5X8Y5 |
| Prunus persic | A0A251Q479 |

| | |
|---|---|
| Prunus persic | M5XMQ7 |
| Prunus persic | M5XHF0 |
| Prunus persic | M5WQ28 |
| Prunus persic | M5X7U6 |
| Prunus persic | M5WVR7 |
| Prunus persic | M5WSH4 |
| Prunus persic | A0A251NDA6 |
| Prunus persic | A0A251PN25 |
| Prunus persic | A0A251PN53 |
| Prunus persic | A0A251Q7F0 |
| Prunus persic | A0A251Q464 |
| Prunus persic | A0A251NT57 |
| Prunus persic | M5X6Z5 |
| Prunus persic | A0A251PRA6 |
| Prunus persic | A0A251PLN9 |
| Prunus persic | M5WPY5 |
| Prunus persic | M5WPV3 |
| Prunus persic | A0A251NIW0 |
| Prunus persic | M5WJJ1 |
| Prunus persic | M5X3N0 |
| Prunus persic | A0A251Q190 |
| Prunus persic | M5X3I1 |
| Prunus persic | A0A251N2H7 |
| Prunus persic | A0A251PK13 |
| Prunus persic | M5WNI3 |
| Prunus persic | A0A251PN15 |
| Prunus persic | A0A251PPI2 |
| Prunus persic | A0A251PWI5 |
| Prunus persic | A0A251PEY5 |
| Prunus persic | A0A251NFQ2 |
| Prunus persic | A0A251Q5B2 |
| Prunus persic | A0A251PF13 |
| Prunus persic | A0A251PN44 |
| Prunus persic | A0A251Q436 |
| Prunus persic | A0A251PEZ5 |
| Prunus persic | M5WVG0 |
| Prunus persic | A0A251PN23 |
| Prunus persic | A0A251PF11 |
| Prunus persic | M5WPG0 |
| Prunus persic | A0A251PPF4 |
| Prunus persic | A0A251PWI0 |

| | |
|---|---|
| Prunus persic | A0A251Q435 |
| Prunus persic | M5WN54 |
| Prunus persic | M5WDJ6 |
| Prunus persic | M5X2T2 |
| Prunus persic | A0A251REB1 |
| Prunus persic | A0A251PPD9 |
| Prunus persic | M5X9G3 |
| Prunus persic | A0A251NKN9 |
| Prunus persic | A0A251PWL4 |
| Prunus persic | A0A251Q4A4 |
| Prunus persic | M5WZT2 |
| Prunus persic | M5W812 |
| Prunus persic | M5WX54 |
| Prunus persic | A0A251PKK4 |
| Prunus persic | M5X978 |
| Prunus persic | M5WZV5 |
| Prunus persic | M5WPY0 |
| Prunus persic | A0A251PF28 |
| Prunus persic | A0A251PN85 |
| Prunus persic | M5WWQ5 |
| Prunus persic | A0A251PN40 |
| Prunus persic | A0A251PWJ6 |
| Prunus persic | M5XIJ4 |
| Prunus persic | A0A251QKG7 |
| Prunus persic | M5WNK5 |
| Prunus persic | M5WM02 |
| Prunus persic | A0A251Q470 |
| Prunus persic | A0A251Q456 |
| Prunus persic | M5VVP2 |
| Prunus persic | A0A251PF31 |
| Prunus persic | A0A251PEZ9 |
| Prunus persic | A0A251PN33 |
| Prunus persic | M5WWU8 |
| Prunus persic | A0A251PEC6 |
| Prunus persic | A0A251PKE3 |
| Prunus persic | M5WPH5 |
| Prunus yedoe | A0A314U9M1 |
| Prunus yedoe | A0A314Y373 |
| Prunus yedoe | A0A314Z0A8 |
| Prunus yedoe | A0A314ZKH3 |
| Prunus yedoe | A0A314XG48 |

| | |
|---|---|
| Prunus yedoe | A0A314YPW4 |
| Prunus yedoe | A0A314XMH1 |
| Prunus yedoe | A0A314YT22 |
| Prunus yedoe | A0A314XP08 |
| Prunus yedoe | A0A314XQJ9 |
| Prunus yedoe | A0A314Y542 |
| Prunus yedoe | A0A314UQL0 |
| Prunus yedoe | A0A314YBU3 |
| Prunus yedoe | A0A314YQ99 |
| Prunus yedoe | A0A314UY39 |
| Prunus yedoe | A0A314YQW7 |
| Prunus yedoe | A0A314YYP7 |
| Prunus yedoe | A0A314YVF9 |
| Prunus yedoe | A0A314XGA1 |
| Prunus yedoe | A0A314UX31 |
| Prunus yedoe | A0A314Z5K6 |
| Prunus yedoe | A0A314Z001 |
| Prunus yedoe | A0A314YRN2 |
| Prunus yedoe | A0A314YT57 |
| Prunus yedoe | A0A314YBR0 |
| Prunus yedoe | A0A314XP39 |
| Prunus yedoe | A0A314Z720 |
| Prunus yedoe | A0A314Y5X9 |
| Prunus yedoe | A0A314UHG7 |
| Prunus yedoe | A0A314Z8Q0 |
| Prunus yedoe | A0A314Y699 |
| Prunus yedoe | A0A314YB32 |
| Prunus yedoe | A0A314UTP8 |
| Prunus yedoe | A0A314UPQ4 |
| Prunus yedoe | A0A314V0I2 |
| Prunus yedoe | A0A314ZFK4 |
| Prunus yedoe | A0A314YL52 |
| Prunus yedoe | A0A315AX60 |
| Prunus yedoe | A0A314YAA4 |
| Prunus yedoe | A0A314Z0M5 |
| Prunus yedoe | A0A314Z3J4 |
| Prunus yedoe | A0A314Y7M3 |
| Prunus yedoe | A0A314V0X4 |
| Prunus yedoe | A0A314YXT4 |
| Prunus yedoe | A0A314YUF3 |
| Prunus yedoe | A0A314UJA2 |

| | |
|---|---|
| Prunus yedoe | A0A315AGJ7 |
| Prunus yedoe | A0A314ZPW0 |
| Prunus yedoe | A0A314Y1L0 |
| Prunus yedoe | A0A314XZH9 |
| Prunus yedoe | A0A314ZSU2 |
| Prunus yedoe | A0A314UXP5 |
| Prunus yedoe | A0A314Y0Z2 |
| Prunus yedoe | A0A314YAA6 |
| Prunus yedoe | A0A314Y818 |
| Prunus yedoe | A0A314Y3J8 |
| Prunus yedoe | A0A314ZB68 |
| Prunus yedoe | A0A314ZP76 |
| Prunus yedoe | A0A314YUG3 |
| Prunus yedoe | A0A314Z802 |
| Prunus yedoe | A0A314UEM7 |
| Prunus yedoe | A0A314YRH3 |
| Prunus yedoe | A0A314Y8A2 |
| Prunus yedoe | A0A314UWI0 |
| Prunus yedoe | A0A315ARL6 |
| Prunus yedoe | A0A315A6R0 |
| Prunus yedoe | A0A314UJ27 |
| Prunus yedoe | A0A314YAF3 |
| Prunus yedoe | A0A314UEA8 |
| Prunus yedoe | A0A314Y3B8 |
| Prunus yedoe | A0A314ZHU3 |
| Prunus yedoe | A0A314YJP3 |
| Prunus yedoe | A0A314XSE1 |
| Prunus yedoe | A0A314YFP3 |
| Prunus yedoe | A0A314Y3T4 |
| Prunus yedoe | A0A314Y829 |
| Prunus yedoe | A0A314Z9C8 |
| Prunus yedoe | A0A314ZVV1 |
| Prunus yedoe | A0A314YLI7 |
| Prunus yedoe | A0A314ZJ97 |
| Prunus yedoe | A0A314YUI6 |
| Prunus yedoe | A0A314YBY2 |
| Prunus yedoe | A0A314XVQ6 |
| Prunus yedoe | A0A314XIP0 |
| Prunus yedoe | A0A314XP62 |
| Prunus yedoe | A0A314UP42 |
| Prunus yedoe | A0A314UV53 |

| | |
|---|---|
| Prunus yedoe | A0A314XIQ0 |
| Prunus yedoe | A0A314Z5Z0 |
| Prunus yedoe | A0A314YBQ1 |
| Prunus yedoe | A0A314YSN1 |
| Prunus yedoe | A0A314YKD5 |
| Prunus yedoe | A0A314UCK4 |
| Prunus yedoe | A0A314YAE2 |
| Prunus yedoe | A0A314YCB1 |
| Prunus yedoe | A0A314XVQ9 |
| Prunus yedoe | A0A314UXN9 |
| Pseudaestuar | A0A0L1JPX5 |
| Pseudaminob | A0A1V8RTC8 |
| Pseudaminob | A0A1V8RVS0 |
| Pseudaminob | A0A316C775 |
| Pseudaminob | A0A432VCJ6 |
| Pseudaminob | A0A432UZB1 |
| Pseudanabae | K9SMW0 |
| Pseudo-nitzsc | A0A448Z8L7 |
| Pseudoaltero | A0A167HEY0 |
| Pseudoaltero | A0A0F6AB37 |
| Pseudoaltero | A0A161Z1S3 |
| Pseudoaltero | A0A1C0TWN0 |
| Pseudobacte | A0A1Y6CLA4 |
| Pseudobacte | A0A1Y6C4J4 |
| Pseudobacte | A0A1Y6CJZ7 |
| Pseudocercos | A0A139HR58 |
| Pseudocercos | A0A139H9G6 |
| Pseudocercos | A0A139HU89 |
| Pseudocercos | A0A139HUE9 |
| Pseudocercos | A0A139H0C0 |
| Pseudocercos | A0A139HE34 |
| Pseudocercos | A0A139HF24 |
| Pseudocercos | M3AJY5 |
| Pseudocercos | M3ABI7 |
| Pseudocercos | N1QAP8 |
| Pseudocercos | M3AM66 |
| Pseudocercos | A0A139IDZ4 |
| Pseudocercos | A0A139ISD7 |
| Pseudocercos | A0A139IAH9 |
| Pseudocercos | A0A139I9V6 |
| Pseudocercos | A0A139IQ32 |

| | |
|---|---|
| Pseudocercos | A0A139HLG1 |
| Pseudocercos | A0A139I5S0 |
| Pseudocercos | A0A139IE78 |
| Pseudocercos | A0A139IRW9 |
| Pseudocercos | A0A139I5G1 |
| Pseudocercos | A0A139I8T4 |
| Pseudochrob | A0A366DTY3 |
| Pseudocohnil | A0A0V0Q930 |
| Pseudogymn | L8FX08 |
| Pseudogymn | A0A177A5U9 |
| Pseudogymn | A0A177AEB2 |
| Pseudogymn | A0A1B8FQ74 |
| Pseudogymn | A0A1B8FAM6 |
| Pseudogymn | A0A1B8FAB7 |
| Pseudogymn | A0A1B8EVM3 |
| Pseudogymn | A0A1B8DV39 |
| Pseudogymn | A0A1B8EGD8 |
| Pseudogymn | A0A1B8DP42 |
| Pseudogymn | A0A1B8DS74 |
| Pseudogymn | A0A1B8E4V2 |
| Pseudogymn | A0A1B8DQT9 |
| Pseudogymn | A0A1B8D360 |
| Pseudogymn | A0A1B8DGK9 |
| Pseudogymn | A0A094EGR9 |
| Pseudogymn | A0A094DWV3 |
| Pseudogymn | A0A094E8W1 |
| Pseudogymn | A0A094ECR8 |
| Pseudogymn | A0A094F5X7 |
| Pseudogymn | A0A093X015 |
| Pseudogymn | A0A093Y9E1 |
| Pseudogymn | A0A093ZSP4 |
| Pseudogymn | A0A093XA42 |
| Pseudogymn | A0A094AHX5 |
| Pseudogymn | A0A093YEN8 |
| Pseudogymn | A0A093YI19 |
| Pseudogymn | A0A093ZRK7 |
| Pseudogymn | A0A093ZUL1 |
| Pseudogymn | A0A093YA26 |
| Pseudogymn | A0A093Y8L0 |
| Pseudogymn | A0A093XLF2 |
| Pseudogymn | A0A093XX47 |

| | |
|---|---|
| Pseudogymn | A0A093XB82 |
| Pseudogymn | A0A093ZC45 |
| Pseudogymn | A0A093ZL44 |
| Pseudogymn | A0A093Z902 |
| Pseudogymn | A0A094BF38 |
| Pseudogymn | A0A094B572 |
| Pseudogymn | A0A094B804 |
| Pseudogymn | A0A094AQ42 |
| Pseudogymn | A0A094AF07 |
| Pseudogymn | A0A094CGQ0 |
| Pseudogymn | A0A094AJK3 |
| Pseudogymn | A0A094C0V4 |
| Pseudogymn | A0A094AXJ9 |
| Pseudogymn | A0A094C5H2 |
| Pseudogymn | A0A094CCB3 |
| Pseudogymn | A0A094DRK2 |
| Pseudogymn | A0A094D2R0 |
| Pseudogymn | A0A094DJC7 |
| Pseudogymn | A0A094D600 |
| Pseudogymn | A0A094CUY7 |
| Pseudogymn | A0A094D6Z9 |
| Pseudogymn | A0A094CLP7 |
| Pseudogymn | A0A094FL50 |
| Pseudogymn | A0A094DH72 |
| Pseudogymn | A0A094FUT8 |
| Pseudogymn | A0A094GG55 |
| Pseudogymn | A0A094FUK7 |
| Pseudogymn | A0A094G2V6 |
| Pseudogymn | A0A094G2T3 |
| Pseudogymn | A0A094F380 |
| Pseudogymn | A0A094HAD8 |
| Pseudogymn | A0A094FXE5 |
| Pseudogymn | A0A094F3W2 |
| Pseudogymn | A0A094IUK4 |
| Pseudogymn | A0A094FUX7 |
| Pseudogymn | A0A094GE65 |
| Pseudogymn | A0A094IFS5 |
| Pseudogymn | A0A094FC36 |
| Pseudogymn | A0A094HXC7 |
| Pseudogymn | A0A094HR56 |
| Pseudogymn | A0A094JG43 |

| | |
|---|---|
| Pseudogymn | A0A094HUW8 |
| Pseudogymn | A0A094H2I0 |
| Pseudogymn | A0A094GL04 |
| Pseudogymn | A0A094IWA8 |
| Pseudogymn | A0A094HPW1 |
| Pseudogymn | A0A094I5T1 |
| Pseudogymn | A0A094K6B1 |
| Pseudogymn | A0A094H1N9 |
| Pseudogymn | A0A094KPW5 |
| Pseudogymn | A0A094HXM1 |
| Pseudogymn | A0A094IA09 |
| Pseudogymn | A0A094I941 |
| Pseudogymn | A0A094HH70 |
| Pseudogymn | A0A094H7Z8 |
| Pseudogymn | A0A094HW58 |
| Pseudogymn | A0A1B8CEM6 |
| Pseudogymn | A0A1B8C167 |
| Pseudogymn | A0A1B8BVH8 |
| Pseudogymn | A0A1B8BVN1 |
| Pseudogymn | A0A1B8CLH8 |
| Pseudogymn | A0A1B8GFJ5 |
| Pseudogymn | A0A1B8GIK4 |
| Pseudogymn | A0A1B8G6U4 |
| Pseudogymn | A0A1B8G8P5 |
| Pseudogymn | A0A1B8GXU4 |
| Pseudolabrys | A0A371BDI6 |
| Pseudolabrys | A0A0Q7TDU5 |
| Pseudolabrys | A0A345ZXK6 |
| Pseudomassa | A0A1Y2DW08 |
| Pseudomassa | A0A1Y2D6M0 |
| Pseudomassa | A0A1Y2E0V4 |
| Pseudomassa | A0A1Y2DSV9 |
| Pseudomassa | A0A1Y2D8N4 |
| Pseudomassa | A0A1Y2E3R2 |
| Pseudomassa | A0A1Y2E4X6 |
| Pseudomassa | A0A1Y2E8Q9 |
| Pseudomona | A0A2V4SND8 |
| Pseudomona | A0A2T5HEC1 |
| Pseudoocean | A0A0A0EH55 |
| Pseudoocean | A3U263 |
| Pseudoocean | A0A1X6ZN84 |

| | |
|---|---|
| Pseudoocean | A0A1I1NTG6 |
| Pseudoocean | A0A399J8R4 |
| Pseudoocean | A0A2E3H3U4 |
| Pseudoocean | A0A2E2PAW3 |
| Pseudopelagi | A0A1M6RS66 |
| Pseudorhizob | A0A081MMN6 |
| Pseudorhodo | A0A1H8C883 |
| Pseudorhodo | A0A433S634 |
| Pseudorhodo | A0A2M8MZT2 |
| Pseudorhodo | A0A257GRH4 |
| Pseudorhodo | A0A1W6ZZH9 |
| Pseudoruege | A0A2T0RJD8 |
| Pseudoruege | A0A1G8QA31 |
| Pseudoxanth | A0A1M7Z7T5 |
| Pseudozyma | A0A081CEQ5 |
| Pterocles gut | A0A093DR50 |
| Pteropus alec | L5JNS7 |
| Pteropus alec | L5KBX5 |
| Pteropus alec | L5KUG3 |
| Pundamilia n | A0A3B4GGZ1 |
| Pundamilia n | A0A3B4ER40 |
| Pundamilia n | A0A3B4EPW7 |
| Pundamilia n | A0A3B4EUN9 |
| Pundamilia n | A0A3B4G794 |
| Pundamilia n | A0A3B4GLL4 |
| Pundamilia n | A0A3B4F895 |
| Punica granat | A0A218X271 |
| Punica granat | A0A2I0HMY7 |
| Punica granat | A0A2I0J4H2 |
| Punica granat | A0A218X161 |
| Punica granat | A0A2I0IJU8 |
| Punica granat | A0A2I0J4I9 |
| Punica granat | A0A218Y2C4 |
| Punica granat | A0A2I0HIQ9 |
| Punica granat | A0A2I0J9K2 |
| Punica granat | A0A218Y341 |
| Punica granat | A0A2I0IQT3 |
| Punica granat | A0A2I0KMH9 |
| Punica granat | A0A2I0IJ58 |
| Punica granat | A0A2I0KZ71 |
| Punica granat | A0A218X0P0 |

| | |
|---|---|
| Punica granat | A0A218WZC1 |
| Punica granat | A0A218WYH1 |
| Punica granat | A0A218WXK1 |
| Punica granat | A0A2I0JQJ0 |
| Punica granat | A0A218WZ09 |
| Punica granat | A0A2I0HYS2 |
| Punica granat | A0A218WXG4 |
| Punica granat | A0A218WUD1 |
| Punica granat | A0A2I0HAP0 |
| Punica granat | A0A218XP01 |
| Punica granat | A0A218WER6 |
| Punica granat | A0A2I0IQZ3 |
| Punica granat | A0A2I0HSV7 |
| Punica granat | A0A218Y426 |
| Punica granat | A0A2I0HST7 |
| Punica granat | A0A218Y3D0 |
| Punica granat | A0A2I0HPH7 |
| Punica granat | A0A218Y2B4 |
| Punica granat | A0A218WC26 |
| Punica granat | A0A218VVJ2 |
| Punica granat | A0A2I0KRY6 |
| Punica granat | A0A218Y294 |
| Punica granat | A0A2I0I1Y2 |
| Punica granat | A0A218WXR5 |
| Punica granat | A0A218WY26 |
| Punica granat | A0A2I0HEN9 |
| Punica granat | A0A2I0KGT9 |
| Punica granat | A0A218X1M6 |
| Punica granat | A0A218X8E1 |
| Punica granat | A0A2I0I1U5 |
| Punica granat | A0A2I0K6N7 |
| Punica granat | A0A218X1Y4 |
| Punica granat | A0A218Y0Y4 |
| Punica granat | A0A2I0HHU4 |
| Punica granat | A0A2I0KWU4 |
| Punica granat | A0A2I0IB26 |
| Punica granat | A0A2I0KD33 |
| Punica granat | A0A2I0JRD0 |
| Punica granat | A0A218Y212 |
| Punica granat | A0A218VZL6 |
| Punica granat | A0A218X0N1 |

| | |
|---|---|
| Punica grana | A0A218Y398 |
| Punica grana | A0A218X6V7 |
| Punica grana | A0A218W3Q4 |
| Punica grana | A0A218X0A3 |
| Punica grana | A0A2I0IHT5 |
| Punica grana | A0A218VUN0 |
| Punica grana | A0A218Y175 |
| Punica grana | A0A218WY07 |
| Punica grana | A0A2I0I1X9 |
| Punica grana | A0A218Y1X1 |
| Punica grana | A0A218X956 |
| Punica grana | A0A218Y478 |
| Punica grana | A0A218XUG7 |
| Punica grana | A0A2I0I3E5 |
| Punica grana | A0A218WYF1 |
| Punica grana | A0A2I0I1U6 |
| Punica grana | A0A2I0KZ81 |
| Punica grana | A0A218Y494 |
| Punica grana | A0A218W9L6 |
| Punica grana | A0A218XS02 |
| Punica grana | A0A218Y227 |
| Punica grana | A0A218X264 |
| Punica grana | A0A218WXQ6 |
| Punica grana | A0A2I0L0F4 |
| Punica grana | A0A2I0J4I3 |
| Punica grana | A0A2I0HKJ3 |
| Punica grana | A0A2I0KZ88 |
| Punica grana | A0A2I0KZ67 |
| Punica grana | A0A2I0KYL9 |
| Punica grana | A0A218X7K9 |
| Punica grana | A0A218WXT2 |
| Punica grana | A0A2I0KZ70 |
| Punica grana | A0A218WY64 |
| Punica grana | A0A2I0KW29 |
| Punica grana | A0A2I0KZA1 |
| Punica grana | A0A2I0J932 |
| Punica grana | A0A2I0IJ71 |
| Punica grana | A0A2I0KG46 |
| Punica grana | A0A2I0I6U2 |
| Punica grana | A0A2I0HSU7 |
| Punica grana | A0A218WXY7 |

| | |
|---|---|
| Punica granat | A0A218XFG8 |
| Punica granat | A0A218XFR5 |
| Punica granat | A0A2I0KN21 |
| Punica granat | A0A2I0HKH6 |
| Punica granat | A0A2I0I2C3 |
| Punica granat | A0A218X1C5 |
| Punica granat | A0A218Y271 |
| Punica granat | A0A2I0KET6 |
| Punica granat | A0A218X305 |
| Punica granat | A0A218Y234 |
| Punica granat | A0A218Y351 |
| Punica granat | A0A2I0KX43 |
| Punica granat | A0A218X2P8 |
| Punica granat | A0A218Y2C0 |
| Punica granat | A0A2I0IHV8 |
| Punica granat | A0A2I0IPE4 |
| Punica granat | A0A218WYC8 |
| Punica granat | A0A2I0KFP7 |
| Punica granat | A0A2I0IHT9 |
| Punica granat | A0A218XW17 |
| Punica granat | A0A218XFS5 |
| Punica granat | A0A218Y0Y2 |
| Punica granat | A0A2I0IQG0 |
| Punica granat | A0A2I0KAW6 |
| Punica granat | A0A2I0KC11 |
| Punica granat | A0A2I0JYX5 |
| Punica granat | A0A218W566 |
| Punica granat | A0A218WF52 |
| Punica granat | A0A218WZS3 |
| Punica granat | A0A218Y3E1 |
| Punica granat | A0A2I0HIE5 |
| Punica granat | A0A2I0KWT0 |
| Punica granat | A0A2I0HT88 |
| Punica granat | A0A218X7K7 |
| Punica granat | A0A218WXZ7 |
| Punica granat | A0A2I0L4U1 |
| Punica granat | A0A218WY16 |
| Punica granat | A0A218WYD8 |
| Punica granat | A0A218XUR4 |
| Punica granat | A0A218WY96 |
| Punica granat | A0A2I0KZ61 |

| | |
|---|---|
| Punica granat | A0A218X3M6 |
| Punica granat | A0A218XAX6 |
| Punica granat | A0A218XDE2 |
| Puniceibacte | A0A2G8RAW8 |
| Puniceibacte | A0A238VTL4 |
| Puniceibacte | A0A0J5QAI6 |
| Purpureocilli | A0A179GCR8 |
| Purpureocilli | A0A2U3DTP8 |
| Purpureocilli | A0A2U3DSM1 |
| Purpureocilli | A0A179G1G3 |
| Purpureocilli | A0A2U3ECC8 |
| Purpureocilli | A0A179HQR1 |
| Purpureocilli | A0A179GK49 |
| Pygocentrus | A0A3B4BXA6 |
| Pygocentrus | A0A3B4DRA6 |
| Pygocentrus | A0A3B4BXD6 |
| Pygocentrus | A0A3B4D157 |
| Pygocentrus | A0A3B4BXJ5 |
| Pygocentrus | A0A3B4C326 |
| Pygocentrus | A0A3B4C1E6 |
| Pygocentrus | A0A3B4C446 |
| Pygocentrus | A0A3B4DN28 |
| Pygoscelis ad | A0A093P1B8 |
| Pygoscelis ad | A0A093RLN1 |
| Pygoscelis ad | A0A093QRK6 |
| Pygoscelis ad | A0A093N8L5 |
| Pyrenochaeta | A0A178E6Z5 |
| Pyrenochaeta | A0A178EA49 |
| Pyrenochaeta | A0A178EEX5 |
| Pyrenochaeta | A0A178DK72 |
| Pyrenochaeta | A0A178DY86 |
| Pyrenophora | A0A3M7M5G6 |
| Pyrenophora | A0A3M7MFK4 |
| Pyrenophora | A0A3M7MG51 |
| Pyrenophora | A0A3M7M128 |
| Pyrenophora | A0A3M7MCE0 |
| Pyrenophora | E3S6M1 |
| Pyrenophora | E3S7C0 |
| Pyrenophora | E3RU61 |
| Pyrenophora | E3RPP7 |
| Pyrenophora | B2WLR3 |

| | |
|---|---|
| Pyrenophora | B2VUM0 |
| Pyrenophora | B2WMT4 |
| Pyrenophora | B2WBS3 |
| Pyrenophora | A0A2W1GRF2 |
| Pyrenophora | A0A2W1HFA3 |
| Pyrenophora | A0A2W1HWY2 |
| Pyrenophora | A0A2W1EGN9 |
| Pyrenophora | A0A2W1ELB0 |
| Pythium insic | A0A2D4C6Q4 |
| Pythium insic | A0A2D4BUE8 |
| Pythium insic | A0A2D4CGH4 |
| Pythium insic | A0A2D4C0E0 |
| Pythium insic | A0A2D4C4K6 |
| Pythium insic | A0A2D4C7F2 |
| Pythium insic | A0A2D4BJS7 |
| Pythium insic | A0A2D4BHR4 |
| Pythium insic | A0A2D4BT34 |
| Pythium insic | A0A2D4BXK6 |
| Pythium insic | A0A2D4BRG2 |
| Pythium insic | A0A2D4BI32 |
| Pythium insic | A0A2D4C690 |
| Pythium insic | A0A2D4BGL9 |
| Pythium insic | A0A2D4C6A5 |
| Pythium insic | A0A2D4C3B2 |
| Pythium insic | A0A2D4C8R4 |
| Pythium insic | A0A2D4C6I3 |
| Rachicladosp | A0A1V8SZ61 |
| Rachicladosp | A0A1V8TRB4 |
| Rachicladosp | A0A1V8T5T1 |
| Rachicladosp | A0A1V8TF18 |
| Rachicladosp | A0A1V8TEM5 |
| Rachicladosp | A0A1V8SMS7 |
| Rachicladosp | A0A1V8TN34 |
| Rachicladosp | A0A1V8SPQ2 |
| Rachicladosp | A0A1V8TVE0 |
| Rachicladosp | A0A1V8ST60 |
| Rachicladosp | A0A1V8TZH6 |
| Rachicladosp | A0A1V8UTC1 |
| Rachicladosp | A0A1V8V8Q1 |
| Rachicladosp | A0A1V8UBK3 |
| Rachicladosp | A0A1V8UW76 |

| | |
|---|---|
| Rachicladosp | A0A1V8U0N8 |
| Rachicladosp | A0A1V8U6W6 |
| Rachicladosp | A0A1V8TZA8 |
| Rachicladosp | A0A1V8U585 |
| Ramazzottius | A0A1D1VG42 |
| Ramazzottius | A0A1D1UP37 |
| Ramazzottius | A0A1D1UTW6 |
| Ramazzottius | A0A1D1VPW1 |
| Ramazzottius | A0A1D1UMW3 |
| Ramazzottius | A0A1D1V436 |
| Ramazzottius | A0A1D1W957 |
| Ramazzottius | A0A1D1UIH8 |
| Ramazzottius | A0A1D1UU11 |
| Ramazzottius | A0A1D1W7T4 |
| Ramazzottius | A0A1D1VJV1 |
| Ramazzottius | A0A1D1VJT6 |
| Ramazzottius | A0A1D1VAI5 |
| Ramularia co | A0A2D3V2X9 |
| Ramularia co | A0A2D3ULC3 |
| Ramularia co | A0A2D3UWU9 |
| Ramularia co | A0A2D3V3D6 |
| Ramularia co | A0A2D3V0K0 |
| Ramularia co | A0A2D3UT01 |
| Ramularia co | A0A2D3V4J3 |
| Raphanus rap | Q9SBP5 |
| Raphanus rap | Q9SBP1 |
| Raphanus rap | Q9SBP4 |
| Raphanus rap | Q9SBP3 |
| Raphanus rap | Q9SBP2 |
| Raphanus rap | Q9SBP0 |
| Raphanus rap | A0A0D5XMK2 |
| Raphanus rap | A0A0D5XNG1 |
| Raphanus rap | Q8LP63 |
| Raphanus rap | A0A0D5XNJ2 |
| Raphanus sat | A0A387IHF6 |
| Raphanus sat | O80347 |
| Raphanus sat | O80351 |
| Raphanus sat | Q6L8R0 |
| Raphanus sat | A0A2D1CRR0 |
| Raphanus sat | A0A3B1EZU5 |
| Raphanus sat | O80350 |

Raphanus sat A0A387IHI8
Raphanus sat O80346
Raphanus sat Q5QHE9
Raphanus sat A0A387IHE2
Raphanus sat D2I902
Raphanus sat A0A387IHI2
Raphanus sat A0A387IHE1
Raphanus sat O80349
Raphanus sat A0A3B1EZV1
Raphanus sat Q6L8R6
Raphanus sat Q5QHE7
Raphanus sat Q940W8
Raphanus sat A0A387IHI7
Raphanus sat Q001P7
Raphanus sat Q6L8R1
Raphanus sat G4XN45
Raphanus sat G4XN49
Raphanus sat A0A387IHI6
Raphanus sat Q6L8R4
Raphanus sat A0A387IHH3
Raphanus sat G4XN46
Raphanus sat Q6L8Q9
Raphanus sat Q6L8R2
Raphanus sat A0A3B1EZU3
Raphanus sat Q6L8R3
Raphanus sat O80356
Raphanus sat A0A387IHI4
Raphanus sat Q940W6
Raphanus sat A0A387IHH9
Raphanus sat O80352
Raphanus sat Q940X0
Raphanus sat Q5QHE8
Raphanus sat G4XN47
Raphanus sat Q5QHE6
Raphanus sat Q6L8R7
Raphanus sat Q5QH04
Raphanus sat A0A387IHF0
Raphanus sat O80353
Raphanus sat A0A2D1CRR9
Raphanus sat O80348
Raphanus sat G4XN48

| | |
|---|---|
| Raphanus sat | Q6Q2Z5 |
| Raphanus sat | A0FLG6 |
| Raphanus sat | A0A387IHE8 |
| Raphanus sat | A0A3B1EZU4 |
| Raphanus sat | Q6L8R5 |
| Raphanus sat | Q940W9 |
| Raphanus sat | P93511 |
| Raphanus sat | Q940W7 |
| Raphidocelis | A0A2V0NKU4 |
| Raphidocelis | A0A2V0PBY3 |
| Raphidocelis | A0A2V0PFW8 |
| Raphidocelis | A0A2V0P2D6 |
| Raphidocelis | A0A2V0PQQ0 |
| Raphidocelis | A0A2V0NRX2 |
| Raphidocelis | A0A2V0NLP1 |
| Raphidocelis | A0A2V0NLF4 |
| Rattus norve | Q01177 |
| Rattus norve | P17945 |
| Rattus norve | P14272 |
| Rattus norve | A0A0G2K4I9 |
| Rattus norve | F7FMS0 |
| Rattus norve | P70521 |
| Rattus norve | Q5FVS2 |
| Rattus norve | Q6TUF8 |
| Reticuliterme | A0A1V1FYM3 |
| Reticuliterme | A0A1V1FQA1 |
| Rhabditopha | A0A1I8CMM6 |
| Rhabditopha | A0A1I8C8R0 |
| Rhabditopha | A0A1I8CK66 |
| Rhabditopha | A0A1I8CA14 |
| Rhabditopha | A0A1I8C8S3 |
| Rhabditopha | A0A1I8D6L7 |
| Rhabditopha | A0A1I8CRQ9 |
| Rhabditopha | A0A1I8C3E9 |
| Rhabditopha | A0A1I8CQ74 |
| Rhabditopha | A0A1I8CUY7 |
| Rhabditopha | A0A1I8CVM8 |
| Rhabditopha | A0A1I8D1P2 |
| Rhabditopha | A0A1I8CH38 |
| Rhabditopha | A0A1I8CBE6 |
| Rhabditopha | A0A1I8CKM7 |

| | |
|---|---|
| Rhabditophar | A0A1I8CF77 |
| Rhabditophar | A0A1I8CLA6 |
| Rhabditophar | A0A1I8CV32 |
| Rhinopithecu | A0A2K6L170 |
| Rhinopithecu | A0A2K6MRU0 |
| Rhinopithecu | A0A2K6MWT8 |
| Rhinopithecu | A0A2K6N2E4 |
| Rhinopithecu | A0A2K6MN57 |
| Rhinopithecu | A0A2K6MWU2 |
| Rhinopithecu | A0A2K6MWT4 |
| Rhinopithecu | A0A2K6N2J7 |
| Rhinopithecu | A0A2K6L173 |
| Rhinopithecu | A0A2K6PRN4 |
| Rhinopithecu | A0A2K6NWB1 |
| Rhinopithecu | A0A2K6N799 |
| Rhinopithecu | A0A2K6RCW8 |
| Rhinopithecu | A0A2K6NTA1 |
| Rhinopithecu | A0A2K6N7A2 |
| Rhinopithecu | A0A2K6N796 |
| Rhinopithecu | A0A2K6NWB3 |
| Rhinopithecu | A0A2K6NWB7 |
| Rhinopithecu | A0A2K6PRM9 |
| Rhinopithecu | A0A2K6NTA8 |
| Rhipicephalus | Q8I6X5 |
| Rhizobiaceae | A0A2D8K1E4 |
| Rhizobiales b | A0A1Q4C6M3 |
| Rhizobiales b | A0A1Q4DD97 |
| Rhizobiales b | A0A1M2Z0A0 |
| Rhizobiales b | A0A1Q3KY06 |
| Rhizobiales b | A0A2E4Z0V6 |
| Rhizobiales b | A0A2E2MCQ3 |
| Rhizobiales b | A0A2E2M8M6 |
| Rhizobiales b | A0A2N6AXD7 |
| Rhizobiales b | A0A2N6B0J3 |
| Rhizobiales b | A0A2A5G4Z8 |
| Rhizobiales b | A0A2N6B7F0 |
| Rhizobiales b | A0A2A4P7M7 |
| Rhizobiales b | A0A3A0EE51 |
| Rhizobiales b | A0A2A4R9T6 |
| Rhizobiales b | A0A349QK41 |
| Rhizobiales b | A0A2A4NQQ8 |

| | |
|---|---|
| Rhizobium ac | A0A0N1DQF6 |
| Rhizobium ac | A0A0N0LCX8 |
| Rhizobium ae | A0A1C3Y3G1 |
| Rhizobium all | A0A2U2DQR8 |
| Rhizobium alt | A0A109K3K2 |
| Rhizobium ar | A0A3S0SF88 |
| Rhizobium es | A0A246DZ15 |
| Rhizobium et | Q2K005 |
| Rhizobium et | B3Q100 |
| Rhizobium et | B3PUG4 |
| Rhizobium et | A0A1L5PF86 |
| Rhizobium et | S5S373 |
| Rhizobium et | A0A0A8GIJ8 |
| Rhizobium et | F2AFE1 |
| Rhizobium et | A0A327YDD0 |
| Rhizobium fa | W6RLA5 |
| Rhizobium fre | G9AGQ7 |
| Rhizobium fre | A0A2L0H058 |
| Rhizobium fre | A0A2A6LWB4 |
| Rhizobium fre | A0A2L0HFL1 |
| Rhizobium fre | A0A2A6LVY4 |
| Rhizobium fre | N6V882 |
| Rhizobium ga | A0A0B4XEY4 |
| Rhizobium ga | A0A1L5NUL5 |
| Rhizobium gr | S3H7T7 |
| Rhizobium gr | A0A370KJW2 |
| Rhizobium ha | A0A1C3WED0 |
| Rhizobium hi | A0A2A6KF27 |
| Rhizobium ja | A0A387FJY0 |
| Rhizobium la | A0A1S9GBZ1 |
| Rhizobium la | A0A1S9GD17 |
| Rhizobium le | A0A072BW40 |
| Rhizobium le | C6B7V3 |
| Rhizobium le | C6B6X0 |
| Rhizobium le | B6A442 |
| Rhizobium le | W0IT72 |
| Rhizobium le | J0KL73 |
| Rhizobium le | J0L6R4 |
| Rhizobium le | I9N4J7 |
| Rhizobium le | I9NL44 |
| Rhizobium le | A0A1C9HR08 |

| | |
|---|---|
| Rhizobium le | A0A1C9HXI1 |
| Rhizobium le | A0A3E1B354 |
| Rhizobium le | A0A3E1AZA5 |
| Rhizobium le | A0A3E1BI76 |
| Rhizobium le | Q1M687 |
| Rhizobium le | Q1M724 |
| Rhizobium le | J0K300 |
| Rhizobium le | J0VE20 |
| Rhizobium le | A0A1Q8H6V5 |
| Rhizobium le | A0A2L1CRP2 |
| Rhizobium le | A0A1Q8HGY6 |
| Rhizobium le | A0A222U8G4 |
| Rhizobium le | A0A222U658 |
| Rhizobium le | A0A444NFR8 |
| Rhizobium le | A0A179BUW9 |
| Rhizobium le | A0A1B1CLA0 |
| Rhizobium le | A0A154IRG5 |
| Rhizobium le | A0A1L3ZLB7 |
| Rhizobium le | A0A3S4Z848 |
| Rhizobium le | A0A154IPB2 |
| Rhizobium le | A0A444HN98 |
| Rhizobium le | A0A444IGJ5 |
| Rhizobium le | A0A444NC52 |
| Rhizobium le | A0A3S3WH50 |
| Rhizobium le | A0A2K9ZEW6 |
| Rhizobium le | A0A2K9ZE91 |
| Rhizobium le | A0A2Z4YEZ8 |
| Rhizobium le | A0A3S4ZBC5 |
| Rhizobium le | A0A2Z4YX29 |
| Rhizobium lo | A0A1G4TJ35 |
| Rhizobium lo | A0A1B4YZE1 |
| Rhizobium lo | A0A1B4YZG6 |
| Rhizobium lo | A0A1B4YZE6 |
| Rhizobium lo | A0A2N5AWF1 |
| Rhizobium lo | A0A1B4YGK0 |
| Rhizobium lo | A0A316CJ14 |
| Rhizobium lo | A0A117N3F6 |
| Rhizobium lo | A0A101KVC7 |
| Rhizobium lo | A0A1B4YE29 |
| Rhizobium lo | A0A1A5HXY1 |
| Rhizobium lo | A0A1A5QH69 |

| | |
|---|---|
| Rhizobium lo | A0A1A5R3P7 |
| Rhizobium lu | A0A1C3W6S0 |
| Rhizobium m | Q92VA6 |
| Rhizobium m | A0A2J0Z7E8 |
| Rhizobium m | A0A437F0L4 |
| Rhizobium m | A0A222GTF2 |
| Rhizobium m | A0A437G4H5 |
| Rhizobium m | A0A222K2F7 |
| Rhizobium m | A0A3S2TJR4 |
| Rhizobium m | A0A437E078 |
| Rhizobium m | K0Q667 |
| Rhizobium m | K0PTJ5 |
| Rhizobium m | A0A1C3U1J6 |
| Rhizobium m | A0A1C3WEX5 |
| Rhizobium na | A0A376ASS4 |
| Rhizobium na | A0A376AR20 |
| Rhizobium ne | A0A0D8KAH4 |
| Rhizobium or | A0A1X7CLE5 |
| Rhizobium or | A0A1Q9AYC7 |
| Rhizobium or | A0A1Q8ZX36 |
| Rhizobium ph | A0A1W6GK72 |
| Rhizobium ph | A0A1W6GQX6 |
| Rhizobium ph | A0A0M3GJ47 |
| Rhizobium ph | A0A0M3GFH7 |
| Rhizobium ph | A0A192TDW1 |
| Rhizobium ph | A0A192TJG2 |
| Rhizobium ph | A0A2U3CMZ0 |
| Rhizobium pi | A0A3R9AP92 |
| Rhizobium pu | A0A1L9CN61 |
| Rhizobium pu | A0A1S9ESQ4 |
| Rhizobium ra | A0A176XJZ8 |
| Rhizobium ra | A0A0X8IY53 |
| Rhizobium ra | A0A0D0J656 |
| Rhizobium ra | A0A176X4G7 |
| Rhizobium ra | A0A1B9SXL8 |
| Rhizobium ra | A0A0D0KZ04 |
| Rhizobium ra | A0A1V2AH49 |
| Rhizobium ra | A0A1B9UR06 |
| Rhizobium ra | A0A2T7W9Z1 |
| Rhizobium ra | A0A2L2LHE9 |
| Rhizobium ra | A0A083ZGC6 |

| | |
|---|---|
| Rhizobium ra | A0A2C5Z3N8 |
| Rhizobium ra | A0A109VT16 |
| Rhizobium ra | A0A3G2DPM2 |
| Rhizobium ra | A0A420FYH9 |
| Rhizobium ra | A0A024IWY1 |
| Rhizobium ra | A0A3B8F8H3 |
| Rhizobium ra | A0A2T7W547 |
| Rhizobium rh | A0A1Q9ALQ8 |
| Rhizobium rh | A0A1Q9AGL9 |
| Rhizobium se | A0A376ABJ2 |
| Rhizobium so | A0A3R9CXJ4 |
| Rhizobium so | A0A2A5KUS5 |
| Rhizobium sp | A0A248W2A8 |
| Rhizobium sp | A0A444LBM9 |
| Rhizobium sp | A0A1L8PLX4 |
| Rhizobium sp | A0A1M3P8H7 |
| Rhizobium sp | A0A1M2YPN5 |
| Rhizobium sp | A0A143ZFH2 |
| Rhizobium sp | A0A0N1AWY9 |
| Rhizobium sp | A0A0N1A5V6 |
| Rhizobium sp | A0A1B9RQK2 |
| Rhizobium sp | A0A1B9S8M0 |
| Rhizobium sp | A0A292ECM0 |
| Rhizobium sp | A0A292EB32 |
| Rhizobium sp | A0A420CH60 |
| Rhizobium sp | A0A420CND4 |
| Rhizobium sp | A0A285Y1D3 |
| Rhizobium sp | J2DBP0 |
| Rhizobium sp | J2WMP8 |
| Rhizobium sp | A0A432P6G8 |
| Rhizobium sp | A0A432NWE3 |
| Rhizobium sp | A0A3S0Q0E5 |
| Rhizobium sp | A0A2A6JF54 |
| Rhizobium sp | J5QAT9 |
| Rhizobium sp | A0A387GV30 |
| Rhizobium sp | W6WJL6 |
| Rhizobium sp | J2IQH9 |
| Rhizobium sp | J1T8K3 |
| Rhizobium sp | A0A1W6KW30 |
| Rhizobium sp | A0A2A6R2N3 |
| Rhizobium sp | A0A098RMU8 |

Rhizobium sp A0A060I507
Rhizobium sp U4Q372
Rhizobium sp A0A2A6K1I0
Rhizobium sp A0A2T3EJJ0
Rhizobium sp A0A2S6BBW7
Rhizobium sp A0A2A6ICH9
Rhizobium sp A0A2A6I5L3
Rhizobium sp A0A2A6HV73
Rhizobium sp A0A2A6HMS6
Rhizobium sp A0A2A6I066
Rhizobium sp A0A2A6KX61
Rhizobium sp A0A0M3B778
Rhizobium sp A0A0M3B5N6
Rhizobium sp A0A1S1T9A9
Rhizobium sp A0A1S1TD64
Rhizobium sp A0A0Q4WAW7
Rhizobium sp A0A0Q5D0H9
Rhizobium sp A0A0Q5E1T4
Rhizobium sp A0A0Q5W5J8
Rhizobium sp A0A0Q5WYK9
Rhizobium sp A0A0Q6H5F7
Rhizobium sp A0A2A6LEH7
Rhizobium sp A0A192M5Y9
Rhizobium sp A0A235AY41
Rhizobium sp A0A235AJ79
Rhizobium sp A0A235AZH9
Rhizobium sp A0A1G5HB02
Rhizobium sp A0A1H9WX89
Rhizobium sp A0A1I0ZNA8
Rhizobium sp A0A1H6GJQ1
Rhizobium sp A0A329AJB1
Rhizobium sp L0NJT2
Rhizobium sp A0A1W6PV42
Rhizobium sp A0A1W6PZX1
Rhizobium sp A0A2L0W608
Rhizobium sp A0A2L0W9E8
Rhizobium sp A0A1G8WJR3
Rhizobium sp H4EZX2
Rhizobium sp K0VNQ6
Rhizobium sp A0A2V4UN27
Rhizobium sp A0A2V4VIK9

| | |
|---|---|
| Rhizobium sp | A0A2V4UL45 |
| Rhizobium sp | A0A246SBW7 |
| Rhizobium sp | A0A246T4R1 |
| Rhizobium sp | A0A246T354 |
| Rhizobium sp | A0A246TWX7 |
| Rhizobium sp | A0A437NBS0 |
| Rhizobium sp | A0A0Q6QM12 |
| Rhizobium sp | A0A0Q6RH95 |
| Rhizobium sp | A0A0Q6S6H3 |
| Rhizobium sp | A0A0Q6VUN6 |
| Rhizobium sp | A0A0Q6UVB6 |
| Rhizobium sp | A0A0Q6ZH68 |
| Rhizobium sp | A0A0Q7XTZ6 |
| Rhizobium sp | A0A0T1WTZ2 |
| Rhizobium sp | A0A0Q7NJK2 |
| Rhizobium sp | A0A0Q7PCQ9 |
| Rhizobium sp | A0A0Q8C8S6 |
| Rhizobium sp | A0A0Q8BTC4 |
| Rhizobium sp | A0A0Q8GJX7 |
| Rhizobium sp | A0A0Q8NRH2 |
| Rhizobium sp | A0A1S1PS94 |
| Rhizobium sp | A0A1N6XKQ2 |
| Rhizobium sp | A0A1N6UY85 |
| Rhizobium sp | A0A1W2DVC3 |
| Rhizobium sp | A0A2T3GU97 |
| Rhizobium sp | A0A2N4XYT7 |
| Rhizobium sp | A0A2V5CDS1 |
| Rhizobium sp | A0A198YKG9 |
| Rhizobium sp | A0A198YI97 |
| Rhizobium sp | A0A2K0ZQK3 |
| Rhizobium sp | A0A2K0ZP10 |
| Rhizobium sp | A0A1E4YFI2 |
| Rhizobium sp | A0A095UEN9 |
| Rhizobium sp | A0A3D3BI31 |
| Rhizobium sp | A0A2W6XJX2 |
| Rhizobium sp | A0A2M8U1J2 |
| Rhizobium sp | A0A353G6F7 |
| Rhizobium su | A0A285U5F1 |
| Rhizobium su | A0A2N0DBB1 |
| Rhizobium ta | A0A1Q9A5B5 |
| Rhizobium til | A0A1H8WJ94 |

| | |
|---|---|
| Rhizobium tib | A0A1H8IHA7 |
| Rhizobium tr | L0LF24 |
| Rhizobium tr | A0A329Y7X4 |
| Rhizobium tr | A0A2T0P999 |
| Rhizobium tu | A0A2W4CTD1 |
| Rhizobium tu | A0A2W4B084 |
| Rhizobium va | A0A432PRA9 |
| Rhizoclosmat | A0A1Y2CKU5 |
| Rhizoclosmat | A0A1Y2CE17 |
| Rhizoclosmat | A0A1Y2B783 |
| Rhizoclosmat | A0A1Y2AMS3 |
| Rhizoclosmat | A0A1Y2C375 |
| Rhizoclosmat | A0A1Y2C103 |
| Rhizoclosmat | A0A1Y2CFU8 |
| Rhizoclosmat | A0A1Y2BGJ3 |
| Rhizoclosmat | A0A1Y2CP39 |
| Rhizoclosmat | A0A1Y2BQK0 |
| Rhizoclosmat | A0A1Y2CED3 |
| Rhizoclosmat | A0A1Y2AE69 |
| Rhizoclosmat | A0A1Y2CJV3 |
| Rhizoclosmat | A0A1Y2CDM5 |
| Rhizoclosmat | A0A1Y2C0V7 |
| Rhizoclosmat | A0A1Y2BFL8 |
| Rhizoclosmat | A0A1Y2CJS8 |
| Rhizoclosmat | A0A1Y2AQ77 |
| Rhizoclosmat | A0A1Y2BHM1 |
| Rhizoclosmat | A0A1Y2CL98 |
| Rhizophora m | A0A2P2IMF7 |
| Rhizophora m | A0A2P2IIC6 |
| Rhizophora m | A0A2P2JRF0 |
| Rhizophora m | A0A2P2II72 |
| Rhizophora m | A0A2P2MSG9 |
| Rhizophora m | A0A2P2MSG6 |
| Rhizophora m | A0A2P2MQG3 |
| Rhizophora m | A0A2P2MSF5 |
| Rhizophora m | A0A2P2K922 |
| Rhodanobact | A0A1V3P122 |
| Rhodnius pro | T1HW46 |
| Rhodnius pro | T1HKB1 |
| Rhodnius pro | T1HVY5 |
| Rhodnius pro | T1H815 |

| | |
|---|---|
| Rhodnius pro | T1I1K7 |
| Rhodnius pro | T1HB28 |
| Rhodobacter | A0A1N7LNA9 |
| Rhodobacter | A0A2T5JUG7 |
| Rhodobacter | A0A2T4JF26 |
| Rhodobacter | D5AST4 |
| Rhodobacter | A0A0E2PIW9 |
| Rhodobacter | A0A1G7E3U5 |
| Rhodobacter | A0A330HGS9 |
| Rhodobacter | A0A285SYE0 |
| Rhodobacter | A0A239GLZ7 |
| Rhodobacter | A0A285CQ45 |
| Rhodobacter | A0A366K6J3 |
| Rhodobacter | A0A1H4S7Q6 |
| Rhodobacter | A0A421BL93 |
| Rhodobacter | C8RWS1 |
| Rhodobacter | A0A3D6CUW1 |
| Rhodobacter | Q3IY48 |
| Rhodobacter | A4WNG2 |
| Rhodobacter | A3PFM8 |
| Rhodobacter | B9KQL4 |
| Rhodobacter | A0A370XYR7 |
| Rhodobacter | A0A2W5U3H5 |
| Rhodobacter | A0A3G6W6P9 |
| Rhodobacter | A0A2T4J571 |
| Rhodobacter | A0A2T4JJD8 |
| Rhodobacter | A0A1N7KZF5 |
| Rhodobacter | A0A318U1P8 |
| Rhodobacter | A0A2G1CH11 |
| Rhodobacter | A0A372EZL6 |
| Rhodobacter | A0A327M0R4 |
| Rhodobacter | A0A3N2R174 |
| Rhodobacter | A0A347UC92 |
| Rhodobacter | A0A3S3T2P1 |
| Rhodobacter | A0A2M7EPC2 |
| Rhodobacter | A0A1B6YLV4 |
| Rhodobacter | A0A0P7W120 |
| Rhodobacter | B6ATT1 |
| Rhodobacter | A3JM87 |
| Rhodobacter | A0A418SAV9 |
| Rhodobacter | A0A3L9Y7X3 |

| | |
|---|---|
| Rhodobacter | A0A257F196 |
| Rhodobacter | A0A0B4E3A1 |
| Rhodobacter | A0A372IHY4 |
| Rhodobacter | A0A3D4DUJ2 |
| Rhodobacter | A0A3D5PSG7 |
| Rhodobacter | A0A2V5FZ51 |
| Rhodobacter | A0A2D4V2U9 |
| Rhodobacter | A0A2A4NME6 |
| Rhodobacter | A0A2E3PJV4 |
| Rhodobacter | A0A2E3FHN5 |
| Rhodobacter | A0A356V427 |
| Rhodobacter | A0A2E3PH72 |
| Rhodobacter | A0A350AHD8 |
| Rhodobacter | A0A2E3PP51 |
| Rhodobacter | A0A2A4ZFU9 |
| Rhodobacter | A0A2D5J967 |
| Rhodobacter | A0A2V5G5E0 |
| Rhodobacter | A0A2D5IXH4 |
| Rhodobacter | A0A259DA28 |
| Rhodobacter | A0A259LR21 |
| Rhodobacter | A0A1Q4CVQ4 |
| Rhodobacter | A0A1G3GSI9 |
| Rhodobacter | B6B7T2 |
| Rhodobacter | A0A2G6ISQ3 |
| Rhodobacter | A0A2G6CFH3 |
| Rhodobacter | A0A2G6I8H1 |
| Rhodobacter | A0A2G6II81 |
| Rhodobacter | A0A2G6IV12 |
| Rhodobiacea | A0A2A4S6F2 |
| Rhodobiacea | A0A355T6E9 |
| Rhodobiacea | A0A2Z4UG21 |
| Rhodobiacea | A0A2E5BRN7 |
| Rhodobiacea | A0A2A5EMY3 |
| Rhodobium c | A0A327JMD5 |
| Rhodoplanes | A0A327KJ78 |
| Rhodoplanes | A0A327K3R9 |
| Rhodoplanes | A0A3S4B6R7 |
| Rhodoplanes | A0A327L3N5 |
| Rhodoplanes | A0A127F2Y1 |
| Rhodopseudo | A0A318TNH4 |
| Rhodopseudo | Q6NBK5 |

| | |
|---|---|
| Rhodopseudo | Q07TU8 |
| Rhodopseudo | Q212S6 |
| Rhodopseudo | Q21BD4 |
| Rhodopseudo | Q13CZ0 |
| Rhodopseudo | E6VBP2 |
| Rhodopseudo | Q2IR81 |
| Rhodopseudo | B3QEG9 |
| Rhodopseudo | A0A0D7EX63 |
| Rhodopseudo | A0A0D7F517 |
| Rhodopseudo | A0A323UFY3 |
| Rhodopseudo | A0A2R4GPL3 |
| Rhodopseudo | A0A418UYX9 |
| Rhodopseudo | A0A336JVI2 |
| Rhodopseudo | A0A1H8TZ49 |
| Rhodopseudo | A0A0N1C8P1 |
| Rhodosalinus | A0A365U4P5 |
| Rhodospirilla | A0A2E7W3G6 |
| Rhodovulum | A0A2T5U026 |
| Rhodovulum | A0A428K1H3 |
| Rhodovulum | A0A366C229 |
| Rhodovulum | A0A1V0I275 |
| Rhodovulum | J5PI78 |
| Rhodovulum | A0A2E2JWZ9 |
| Rhodovulum | A0A1Q9QUC6 |
| Rhynchospor | A0A1E1KPQ4 |
| Rhynchospor | A0A1E1LMW4 |
| Rhynchospor | A0A1E1KU29 |
| Rhynchospor | A0A1E1KM25 |
| Rhynchospor | A0A1E1JTK9 |
| Rhynchospor | A0A1E1LGC2 |
| Rhynchospor | A0A1E1K9E7 |
| Rhynchospor | A0A1E1KXI0 |
| Rhynchospor | A0A1E1KAU5 |
| Rhynchospor | A0A1E1LP64 |
| Rhynchospor | A0A1E1MDX2 |
| Rhynchospor | A0A1E1M5W1 |
| Rhynchospor | A0A1E1MFU2 |
| Rhynchospor | A0A1E1M9F8 |
| Rhynchospor | A0A1E1MTL8 |
| Rhynchospor | A0A1E1MD61 |
| Rhynchospor | A0A1E1M215 |

| | |
|---|---|
| Ricinus comm | B9SSB2 |
| Ricinus comm | B9STG0 |
| Ricinus comm | B9SEN3 |
| Ricinus comm | B9RXY2 |
| Ricinus comm | B9SXC2 |
| Ricinus comm | B9RQP9 |
| Ricinus comm | B9RFM4 |
| Ricinus comm | B9SIG2 |
| Ricinus comm | B9SSB7 |
| Ricinus comm | B9TAY3 |
| Ricinus comm | B9R758 |
| Ricinus comm | B9RHL2 |
| Ricinus comm | B9SJQ1 |
| Ricinus comm | B9T444 |
| Ricinus comm | B9SV42 |
| Ricinus comm | B9S7P6 |
| Ricinus comm | B9SIQ7 |
| Ricinus comm | B9SSB5 |
| Ricinus comm | B9T7U7 |
| Ricinus comm | B9RLS4 |
| Ricinus comm | B9SIR0 |
| Ricinus comm | B9R757 |
| Ricinus comm | B9R755 |
| Ricinus comm | B9SSA1 |
| Ricinus comm | B9RXX5 |
| Ricinus comm | B9R760 |
| Ricinus comm | B9SNG5 |
| Ricinus comm | B9RQP5 |
| Ricinus comm | B9SSC0 |
| Ricinus comm | B9R761 |
| Ricinus comm | B9S0R9 |
| Ricinus comm | B9SRH2 |
| Ricinus comm | B9SFF3 |
| Ricinus comm | B9RI65 |
| Ricinus comm | B9S9P6 |
| Ricinus comm | B9SFF2 |
| Ricinus comm | B9SFX3 |
| Ricinus comm | B9RXY0 |
| Ricinus comm | B9RXY1 |
| Ricinus comm | B9RVX8 |
| Ricinus comm | B9SW97 |

| | |
|---|---|
| Ricinus comm | B9SFX5 |
| Ricinus comm | B9SFF5 |
| Ricinus comm | B9SIR1 |
| Ricinus comm | B9SZ96 |
| Ricinus comm | B9RXX9 |
| Ricinus comm | B9SFF1 |
| Ricinus comm | B9RFM6 |
| Ricinus comm | B9S858 |
| Ricinus comm | B9SSB8 |
| Ricinus comm | B9RV63 |
| Ricinus comm | B9RLS2 |
| Ricinus comm | B9T7U8 |
| Robiginitoma | A0A2G2GCP5 |
| Robiginitoma | A0A2G2R0P0 |
| Robiginitoma | A0A2G1YWK1 |
| Robiginitoma | A0A2G2HQX6 |
| Robiginitoma | A0A2G2GH02 |
| Rosa chinens | A0A2P6Q8G4 |
| Rosa chinens | A0A2P6SHN7 |
| Rosa chinens | A0A2P6SCY4 |
| Rosa chinens | A0A2P6Q365 |
| Rosa chinens | A0A2P6QUU6 |
| Rosa chinens | A0A2P6Q342 |
| Rosa chinens | A0A2P6S2Q9 |
| Rosa chinens | A0A2P6QU66 |
| Rosa chinens | A0A2P6PXU3 |
| Rosa chinens | A0A2P6Q3D1 |
| Rosa chinens | A0A2P6S898 |
| Rosa chinens | A0A2P6SAU9 |
| Rosa chinens | A0A2P6RJH7 |
| Rosa chinens | A0A2P6Q908 |
| Rosa chinens | A0A2P6QQ00 |
| Rosa chinens | A0A2P6QVT9 |
| Rosa chinens | A0A2P6QJ32 |
| Rosa chinens | A0A2P6Q399 |
| Rosa chinens | A0A2P6S8C4 |
| Rosa chinens | A0A2P6Q341 |
| Rosa chinens | A0A2P6RSA0 |
| Rosa chinens | A0A2P6PFN5 |
| Rosa chinens | A0A2P6Q3C0 |
| Rosa chinens | A0A2P6RCT1 |

Rosa chinens A0A2P6QB94
Rosa chinens A0A2P6Q3C5
Rosa chinens A0A2P6Q391
Rosa chinens A0A2P6QHF8
Rosa chinens A0A2P6Q8G0
Rosa chinens A0A2P6S2T7
Rosa chinens A0A2P6S756
Rosa chinens A0A2P6RTY8
Rosa chinens A0A2P6Q351
Rosa chinens A0A2P6QBC3
Rosa chinens A0A2P6QBG0
Rosa chinens A0A2P6PXY7
Rosa chinens A0A2P6QBF8
Rosa chinens A0A2P6RCR1
Rosa chinens A0A2P6QAY4
Rosa chinens A0A2P6Q3C2
Rosa chinens A0A2P6Q3E7
Rosa chinens A0A2P6R3M6
Rosa chinens A0A2P6Q0T1
Rosa chinens A0A2P6RX36
Rosa chinens A0A2P6SAB0
Rosa chinens A0A2P6Q3A9
Rosa chinens A0A2P6Q390
Rosa chinens A0A2P6QB71
Rosa chinens A0A2P6S6W6
Rosa chinens A0A2P6S765
Rosa chinens A0A2P6Q3D5
Rosa chinens A0A2P6Q8T0
Rosa chinens A0A2P6QBH7
Rosa chinens A0A2P6Q3A8
Rosa chinens A0A2P6Q3C6
Rosa chinens A0A2P6Q8Z8
Rosa chinens A0A2P6Q367
Rosa chinens A0A2P6RX44
Rosa chinens A0A2P6QU56
Rosa chinens A0A2P6QE17
Rosa chinens A0A2P6QE07
Rosa chinens A0A2P6QVC9
Rosa chinens A0A2P6QAX3
Rosa chinens A0A2P6Q355
Rosa chinens A0A2P6Q396

Rosa chinens A0A2P6RRU2
Rosa chinens A0A2P6Q2N8
Rosa chinens A0A2P6RUA2
Rosa chinens A0A2P6Q3A7
Rosa chinens A0A2P6RRQ5
Rosa chinens A0A2P6SCB0
Rosa chinens A0A2P6RU56
Rosa chinens A0A2P6RS89
Rosa chinens A0A2P6Q372
Rosa chinens A0A2P6QM19
Rosa chinens A0A2P6PX37
Rosa chinens A0A2P6Q374
Rosa chinens A0A2P6Q0Q2
Rosa chinens A0A2P6SGJ1
Rosa chinens A0A2P6QR39
Rosa chinens A0A2P6Q381
Rosa chinens A0A2P6Q362
Rosa chinens A0A2P6QBG3
Rosa chinens A0A2P6S2K8
Rosa chinens A0A2P6S2R0
Rosa chinens A0A2P6Q3B2
Rosa chinens A0A2P6QBH3
Rosa chinens A0A2P6RBI2
Rosa chinens A0A2P6S6X4
Rosa chinens A0A2P6PD71
Rosa chinens A0A2P6SD05
Rosa chinens A0A2P6S6W4
Rosa chinens A0A2P6RST1
Rosa chinens A0A2P6QBL5
Rosa chinens A0A2P6PXZ1
Rosa chinens A0A2P6Q348
Rosa chinens A0A2P6RVC0
Rosa chinens A0A2P6RRR7
Rosa chinens A0A2P6S2Q6
Rosa chinens A0A2P6S2L9
Rosa chinens A0A2P6Q397
Rosa chinens A0A2P6Q8F1
Rosa chinens A0A2P6S2P5
Rosa chinens A0A2P6QBJ8
Rosa chinens A0A2P6QVN2
Rosa chinens A0A2P6Q8K4

Rosa chinens A0A2P6QBA5
Rosa chinens A0A2P6RU59
Rosa chinens A0A2P6Q8F3
Rosa chinens A0A2P6QR92
Rosa chinens A0A2P6P691
Rosa chinens A0A2P6Q8G5
Rosa chinens A0A2P6PDR1
Rosa chinens A0A2P6QB88
Rosa chinens A0A2P6RRU7
Rosa chinens A0A2P6Q394
Rosa chinens A0A2P6SAD4
Rosa chinens A0A2P6SAC0
Rosa chinens A0A2P6Q3A3
Rosa chinens A0A2P6Q8H0
Rosa chinens A0A2P6S8D3
Rosa chinens A0A2P6Q8G3
Rosa chinens A0A2P6PDQ4
Rosa chinens A0A2P6QM80
Rosa chinens A0A2P6RCQ5
Rosa chinens A0A2P6Q8U1
Rosa chinens A0A2P6Q0P6
Rosa chinens A0A2P6RVC4
Rosa chinens A0A2P6Q3B1
Rosa chinens A0A2P6SAY2
Rosa chinens A0A2P6RUW8
Rosa chinens A0A2P6Q8U4
Rosa chinens A0A2P6QW50
Rosa chinens A0A2P6R760
Rosa chinens A0A2P6RCX2
Rosa chinens A0A2P6RRU6
Rosa chinens A0A2P6REX4
Rosa chinens A0A2P6PDQ7
Rosa chinens A0A2P6SLZ0
Rosa chinens A0A2P6Q385
Rosa chinens A0A2P6S6Y9
Rosa chinens A0A2P6Q8L4
Rosa chinens A0A2P6QQ26
Rosa chinens A0A2P6RCS7
Rosa chinens A0A2P6SD19
Rosa chinens A0A2P6RU70
Rosa chinens A0A2P6R9I7

Rosa chinens A0A2P6S2N8
Rosa chinens A0A2P6Q8V1
Rosa chinens A0A2P6PXZ2
Rosa chinens A0A2P6S8C8
Rosa chinens A0A2P6QM44
Rosa chinens A0A2P6QBI9
Rosa chinens A0A2P6PAH1
Rosa chinens A0A2P6QBF9
Rosa chinens A0A2P6QBI2
Rosa chinens A0A2P6QBK6
Rosa chinens A0A2P6Q387
Rosa chinens A0A2P6Q392
Rosa chinens A0A2P6QBI6
Rosa chinens A0A2P6REY3
Rosa chinens A0A2P6Q8I8
Rosa chinens A0A2P6QVT7
Rosa chinens A0A2P6QA23
Rosa chinens A0A2P6S790
Rosa chinens A0A2P6QL63
Rosa chinens A0A2P6RW66
Rosa chinens A0A2P6S8G7
Rosa chinens A0A2P6QR51
Rosa chinens A0A2P6Q3C7
Rosa chinens A0A2P6PXY8
Rosa chinens A0A2P6Q3C3
Rosa chinens A0A2P6Q3D6
Rosa chinens A0A2P6Q398
Rosa chinens A0A2P6QB80
Rosa chinens A0A2P6S2F9
Rosa chinens A0A2P6S2M5
Rosa chinens A0A2P6PEP6
Rosa chinens A0A2P6QBC9
Rosa chinens A0A2P6RS63
Rosa chinens A0A2P6RRT9
Rosa chinens A0A2P6QBC1
Rosa chinens A0A2P6Q388
Rosa chinens A0A2P6S745
Rosa chinens A0A2P6PXY5
Rosa chinens A0A2P6RS64
Rosa chinens A0A2P6Q8Y2
Rosa chinens A0A2P6QP61

| | |
|---|---|
| Rosa chinens | A0A2P6PD75 |
| Rosa chinens | A0A2P6S2K2 |
| Rosa chinens | A0A2P6QBA3 |
| Rosa chinens | A0A2P6R9L9 |
| Rosa chinens | A0A2P6RWS3 |
| Rosa chinens | A0A2P6S2N2 |
| Rosa chinens | A0A2P6S729 |
| Rosa chinens | A0A2P6Q8F7 |
| Rosa chinens | A0A2P6QBK3 |
| Rosa chinens | A0A2P6S742 |
| Rosa chinens | A0A2P6RS56 |
| Rosa chinens | A0A2P6PDH3 |
| Rosa chinens | A0A2P6Q393 |
| Rosa chinens | A0A2P6QBH0 |
| Rosa chinens | A0A2P6Q8Y0 |
| Rosa chinens | A0A2P6Q378 |
| Rosa chinens | A0A2P6PY16 |
| Rosa chinens | A0A2P6S8D1 |
| Rosa chinens | A0A2P6S8F6 |
| Rosa chinens | A0A2P6Q8U6 |
| Rosa chinens | A0A2P6Q3J3 |
| Rosa chinens | A0A2P6PXZ6 |
| Rosa chinens | A0A2P6R9M2 |
| Rosa chinens | A0A2P6SB38 |
| Rosa chinens | A0A2P6QBK4 |
| Rosa chinens | A0A2P6QB92 |
| Rosa chinens | A0A2P6QBL2 |
| Rosa chinens | A0A2P6SCZ3 |
| Rosa chinens | A0A2P6QUV7 |
| Rosa chinens | A0A2P6QWZ3 |
| Rosa chinens | A0A2P6Q389 |
| Rosa chinens | A0A2P6QA32 |
| Roseibaca cal | A0A0P7W3J3 |
| Roseibaca ek | A0A3B0MGU5 |
| Roseibacteriu | W8RZV3 |
| Roseibium sp | E2CPR9 |
| Roseicyclus n | A0A316GM60 |
| Roseisalinus a | A0A1Y5S619 |
| Roseivivax isc | X7F4L9 |
| Roseivivax lei | A0A1N7JUR1 |
| Roseivivax pa | A0A2T6B3V4 |

| | |
|---|---|
| Roseivivax se | A0A1I1U481 |
| Rosellinia nec | A0A1W2TSD6 |
| Rosellinia nec | A0A1W2TQM6 |
| Roseobacter | B7RMG1 |
| Roseobacter | A3X6K1 |
| Roseobacter | A4ET50 |
| Roseovarius a | A0A1X7BTA5 |
| Roseovarius a | A0A1X6YTQ2 |
| Roseovarius a | A0A0T5NYW0 |
| Roseovarius i | A0A0T5PEA0 |
| Roseovarius l | A0A1I4YX20 |
| Roseovarius r | A0A1N7E9E7 |
| Roseovarius r | A3SKX6 |
| Roseovarius r | A0A348WF88 |
| Roseovarius s | A0A176F0X4 |
| Roseovarius s | A0A176FFA7 |
| Roseovarius s | A0A3A8B3W9 |
| Roseovarius s | A0A365TZC0 |
| Roseovarius s | A0A2E2S3M6 |
| Roseovarius s | A0A2D9CQ95 |
| Roseovarius t | A0A1H8EWM5 |
| Rubellimicrol | A0A017HNB8 |
| Ruegeria litor | A0A1Y5RCZ8 |
| Ruegeria mar | A0A1G7AEP1 |
| Ruegeria mar | A0A0X3TTM9 |
| Ruegeria mec | A0A2R8C330 |
| Ruegeria pon | Q5LWT4 |
| Ruegeria sp. / | A0A346SKI6 |
| Ruegeria sp. I | A0A2V3VSP3 |
| Ruegeria sp. I | A0A1E3D7V4 |
| Ruegeria sp. I | A0A1E3DCB0 |
| Rutstroemia s | A0A2S7PS49 |
| Rutstroemia s | A0A2S7PDR4 |
| Rutstroemia s | A0A2S7Q765 |
| Saccharum hy | A0A2R4QNS3 |
| Sagittula sp. I | A0A2K8WAW8 |
| Sagittula stell | A3K2N2 |
| Saimiri bolivi | A0A2K6UT93 |
| Saimiri bolivi | A0A2K6SSN8 |
| Saimiri bolivi | A0A2K6S9K0 |
| Saimiri bolivi | A0A2K6S9I8 |

| | |
|---|---|
| Saimiri bolivi | A0A2K6UT83 |
| Saimiri bolivi | A0A2K6SSN3 |
| Saimiri bolivi | A0A2K6SSP3 |
| Saimiri bolivi | A0A2K6UTP6 |
| Saimiri bolivi | A0A2K6UYV6 |
| Saimiri bolivi | A0A2K6UYX4 |
| Saitoella com | A0A0E9NMM4 |
| Salinihabitans | A0A1H8QTL5 |
| Salipiger aest | A0A327Y1D3 |
| Salipiger mar | A0A1G8I7T9 |
| Salipiger muc | S9QTS7 |
| Salipiger prof | A0A1U7D1X3 |
| Salipiger thio | A0A1G7KG72 |
| Salmo salar.x | A0A1S3LE46 |
| Salmo salar.x | A0A1S3S8H0 |
| Salmo salar.x | A0A1S3SH05 |
| Salmo salar.x | A0A1S3MAB2 |
| Salmo salar.x | A0A1S3SH07 |
| Salmo salar.x | A0A1S3SH04 |
| Salmo salar.x | A0A1S3MLI3 |
| Salmo salar.x | A0A1S3SH06 |
| Salmo salar.x | A0A1S3N1I0 |
| Salmo salar.x | A0A1S3SH08 |
| Salmo salar.x | A0A1S3SIB8 |
| Salmo salar.x | A0A1S3SIC4 |
| Salmo salar.x | A0A1S3KP78 |
| Salmo salar.x | A0A1S3SH12 |
| Salmo salar.x | A0A1S3LE48 |
| Salmo salar.x | A0A1S3P228 |
| Salpingoeca r | F2TWZ6 |
| Salpingoeca r | F2UF68 |
| Salpingoeca r | F2TWR8 |
| Salpingoeca r | F2UG85 |
| Salpingoeca r | F2UPS4 |
| Salpingoeca r | F2USW3 |
| Salpingoeca r | F2U618 |
| Salpingoeca r | F2UA60 |
| Salpingoeca r | F2UFK9 |
| Salpingoeca r | F2U2Y5 |
| Salpingoeca r | F2UGI4 |
| Salpingoeca r | F2U386 |

| | |
|---|---|
| Salpingoeca r | F2UIR6 |
| Salpingoeca r | F2UHB7 |
| Salpingoeca r | F2TVR9 |
| Salpingoeca r | F2UJ78 |
| Salpingoeca r | F2UIC5 |
| Salpingoeca r | F2UDJ0 |
| Salpingoeca r | F2U6I3 |
| Samba virus.› | A0A140E138 |
| Samba virus.› | W6GGU3 |
| Samba virus.› | A0A140E0U4 |
| Sandaracinus | A0A2E0TIG6 |
| Saprolegnia c | T0RAA8 |
| Saprolegnia c | T0RYA1 |
| Saprolegnia c | T0QTS1 |
| Saprolegnia c | T0Q639 |
| Saprolegnia c | T0RYL1 |
| Saprolegnia c | T0PVB7 |
| Saprolegnia c | T0RQN3 |
| Saprolegnia c | T0R976 |
| Saprolegnia c | T0RRW1 |
| Saprolegnia c | T0QUW1 |
| Saprolegnia c | T0QNZ2 |
| Saprolegnia c | T0RMV6 |
| Saprolegnia c | T0PNV2 |
| Saprolegnia c | T0RAF6 |
| Saprolegnia c | T0QYA6 |
| Saprolegnia c | T0S243 |
| Saprolegnia c | T0RG82 |
| Saprolegnia c | T0QXL6 |
| Saprolegnia c | T0RI62 |
| Saprolegnia c | T0RPR9 |
| Saprolegnia c | T0R5M4 |
| Saprolegnia c | T0RIQ1 |
| Saprolegnia c | T0RBY8 |
| Saprolegnia c | T0QL36 |
| Saprolegnia c | T0PWZ1 |
| Saprolegnia c | T0QQ07 |
| Saprolegnia c | T0RBY3 |
| Saprolegnia c | T0RH69 |
| Saprolegnia c | T0QGJ1 |
| Saprolegnia c | T0RJM1 |

| | |
|---|---|
| Saprolegnia c | T0PH14 |
| Saprolegnia c | T0R5A9 |
| Saprolegnia c | T0Q529 |
| Saprolegnia c | T0PVB2 |
| Saprolegnia c | T0RYU4 |
| Saprolegnia c | T0PU14 |
| Saprolegnia c | T0RJM4 |
| Saprolegnia c | T0QZF9 |
| Saprolegnia c | T0R762 |
| Saprolegnia c | T0S1C6 |
| Saprolegnia c | T0R8A8 |
| Saprolegnia c | T0Q4I6 |
| Saprolegnia c | T0QAA3 |
| Saprolegnia c | T0Q2C6 |
| Saprolegnia c | T0QMW7 |
| Saprolegnia c | T0QN46 |
| Saprolegnia c | T0RIN6 |
| Saprolegnia c | T0QI45 |
| Saprolegnia c | T0QJB1 |
| Saprolegnia c | T0QUH0 |
| Saprolegnia c | T0RQM9 |
| Saprolegnia c | T0Q4C3 |
| Saprolegnia p | A0A067BU55 |
| Saprolegnia p | A0A067BYE3 |
| Saprolegnia p | A0A067BYA7 |
| Saprolegnia p | A0A067BJE2 |
| Saprolegnia p | A0A067C8S2 |
| Saprolegnia p | A0A067CBB6 |
| Saprolegnia p | A0A067C0P6 |
| Saprolegnia p | A0A067BSE2 |
| Saprolegnia p | A0A067BIR1 |
| Saprolegnia p | A0A067C156 |
| Saprolegnia p | A0A067BJJ2 |
| Saprolegnia p | A0A067CR05 |
| Saprolegnia p | A0A067CK80 |
| Saprolegnia p | A0A067C126 |
| Saprolegnia p | A0A067CKM6 |
| Saprolegnia p | A0A067C128 |
| Saprolegnia p | A0A067BMR9 |
| Saprolegnia p | A0A067CL67 |
| Saprolegnia p | A0A067BJT0 |

| | |
|---|---|
| Saprolegnia p | A0A067CFL9 |
| Saprolegnia p | A0A067BLZ4 |
| Saprolegnia p | A0A067BU12 |
| Saprolegnia p | A0A067BRT0 |
| Saprolegnia p | A0A067CCE3 |
| Saprolegnia p | A0A067CGD2 |
| Saprolegnia p | A0A067BEG5 |
| Saprolegnia p | A0A067CYL5 |
| Saprolegnia p | A0A067CF87 |
| Saprolegnia p | A0A067C4Y2 |
| Saprolegnia p | A0A067C8W4 |
| Saprolegnia p | A0A067BUH1 |
| Saprolegnia p | A0A067D5R2 |
| Saprolegnia p | A0A067CQB6 |
| Saprolegnia p | A0A067CG93 |
| Saprolegnia p | A0A067CKH3 |
| Saprolegnia p | A0A067C9Y0 |
| Saprolegnia p | A0A067BWR1 |
| Saprolegnia p | A0A067BRT6 |
| Saprolegnia p | A0A067CC90 |
| Saprolegnia p | A0A067BYB2 |
| Saprolegnia p | A0A067C8G3 |
| Saprolegnia p | A0A067C2R2 |
| Saprolegnia p | A0A067D5R8 |
| Saprolegnia p | A0A067CDQ0 |
| Saprolegnia p | A0A067BZY9 |
| Saprolegnia p | A0A067C9M8 |
| Saprolegnia p | A0A067C9D7 |
| Saprolegnia p | A0A067C6I2 |
| Saprolegnia p | A0A067CHH5 |
| Saprolegnia p | A0A067BRS1 |
| Saprolegnia p | A0A067BXJ9 |
| Saprolegnia p | A0A067CJ02 |
| SAR86 cluster | J5KAB5 |
| SAR86 cluster | A0A368BEY8 |
| Sarcocystis m | Q08668 |
| Sarcocystis m | P81860 |
| Sarcocystis m | Q26539 |
| Sarcophilus h | G3WZ37 |
| Sarcophilus h | G3VTC0 |
| Sarcophilus h | G3WZ36 |

| | |
|---|---|
| Sarcophilus h | G3VQX8 |
| Sarcophilus h | G3VAC2 |
| Sarcophilus h | G3VAC3 |
| Sarcophilus h | G3WT27 |
| Sarcophilus h | G3VQX9 |
| Sarcoptes sca | A0A131ZZD7 |
| Sarcoptes sca | A0A132A4Q6 |
| Sarcoptes sca | A0A132A395 |
| Sarcoptes sca | A0A132A5Y6 |
| Sarcoptes sca | A0A132A5S6 |
| Sarcoptes sca | A0A131ZTE1 |
| Sarcoptes sca | A0A131ZSZ0 |
| Saudi moumo | A0A1S5V1F4 |
| Saudi moumo | A0A1S5V1F6 |
| Saudi moumo | A0A1S5V127 |
| Scedosporiun | A0A084G238 |
| Scedosporiun | A0A084G6H4 |
| Scedosporiun | A0A084G8P5 |
| Schistocepha | A0A0X3PVM5 |
| Schistosoma | A0A430QM43 |
| Schistosoma | A0A430QQG0 |
| Schistosoma | A0A183JN66 |
| Schistosoma | A0A183JYW3 |
| Schistosoma | A0A095C245 |
| Schistosoma | A0A094ZN30 |
| Schistosoma | Q5DBB4 |
| Schistosoma | A0A3Q0KQV0 |
| Schistosoma | A0A183MF64 |
| Schistosoma | A0A183NRF4 |
| Schistosoma | A0A3P8BBP8 |
| Schistosoma | A0A183R1Q6 |
| Schizaphis gr | A0A2S2N7W5 |
| Schizaphis gr | A0A2S2NT68 |
| Schizaphis gr | A0A2S2P2Z5 |
| Schizaphis gr | A0A2S2NBX7 |
| Schizaphis gr | A0A2S2NWJ0 |
| Schizaphis gr | A0A2S2PP89 |
| Schizaphis gr | A0A2S2NVK1 |
| Schizaphis gr | A0A2S2NBX8 |
| Schizaphis gr | A0A2S2NQG9 |
| Schizaphis gr | A0A2S2NP55 |

| | |
|---|---|
| Scleropages f | A0A1W5APA3 |
| Scleropages f | A0A0P7VVQ1 |
| Scleropages f | A0A1W5APF2 |
| Scleropages f | A0A0P7XEA4 |
| Scleropages f | A0A0P7ZBC4 |
| Scleropages f | A0A0P7UFY3 |
| Scleropages f | A0A1W4Y2F3 |
| Scleropages f | A0A0P7ULB2 |
| Scleropages f | A0A1W4YBZ6 |
| Scleropages f | A0A1W4YNQ9 |
| Scleropages f | A0A1W5A615 |
| Scleropages f | A0A0P7UJJ9 |
| Scleropages f | A0A1W4ZAZ3 |
| Sclerotinia bo | W9CB98 |
| Sclerotinia bo | W9C6T6 |
| Sclerotinia sc | A0A1D9Q902 |
| Sclerotinia sc | A0A1D9PTT0 |
| Sclerotinia sc | A0A1D9PV26 |
| Sclerotinia sc | A7E7Z9 |
| Sclerotinia sc | A7E800 |
| Sclerotinia sc | A7F252 |
| Scophthalmu | A0A2U9B2H8 |
| Scophthalmu | A0A2U9BYQ0 |
| Scophthalmu | A0A2U9CUU6 |
| Scophthalmu | A0A2U9CZF4 |
| Scophthalmu | A0A2U9BGH2 |
| Scophthalmu | A0A2U9CZ16 |
| Scophthalmu | A0A2U9CYT4 |
| Scophthalmu | A0A2U9D1N9 |
| Scophthalmu | A0A2U9CUM9 |
| Scophthalmu | A0A2U9CYZ6 |
| Scyliorhinus t | A0A401NHP4 |
| Scyliorhinus t | A0A401NSN8 |
| Scyliorhinus t | A0A401NY63 |
| Scyliorhinus t | A0A401PJU3 |
| Scyliorhinus t | A0A401P1Q0 |
| Scyliorhinus t | A0A2Z5XDQ5 |
| Scylla olivace | A0A0P4WNK8 |
| Scylla olivace | A0A0P4WFH3 |
| Scylla olivace | A0A0P4WR24 |
| Scylla olivace | A0A0N7ZDF3 |

| | |
|---|---|
| Scylla olivace | A0A0P4WF65 |
| Scylla olivace | A0A0P4WMW9 |
| Scytalidium li | A0A3E2GVD1 |
| Sedimentitale | A0A1I6XD87 |
| Sedimentitale | A0A3T0N402 |
| Sedimentitale | A0A3T0N638 |
| Selaginella m | D8R4F7 |
| Selaginella m | D8SC04 |
| Selaginella m | D8QZ24 |
| Selaginella m | D8TB11 |
| Selaginella m | D8T7A8 |
| Selaginella m | D8TF62 |
| Selaginella m | D8S1J3 |
| Selaginella m | D8SDZ7 |
| Selaginella m | D8QTB9 |
| Selaginella m | D8RST4 |
| Selaginella m | D8S4Y9 |
| Selaginella m | D8QZK2 |
| Selaginella m | D8T4B9 |
| Selaginella m | D8SRG6 |
| Selaginella m | D8RA35 |
| Selaginella m | D8RHS8 |
| Selaginella m | D8S2E7 |
| Selaginella m | D8TG77 |
| Selaginella m | D8RWA6 |
| Selaginella m | D8SX02 |
| Selaginella m | D8SNE6 |
| Selaginella m | D8SC01 |
| Selaginella m | D8QTB8 |
| Selaginella m | D8S6U2 |
| Selaginella m | D8R007 |
| Selaginella m | D8R008 |
| Selaginella m | D8RVP2 |
| Senecio squa | Q5NJB1 |
| Sepioteuthis | R4G2C9 |
| Sepioteuthis | R4G7D8 |
| Seriola dume | A0A3B4TE38 |
| Seriola dume | A0A3B4T255 |
| Seriola dume | A0A3B4THQ6 |
| Seriola dume | A0A3B4TCE1 |
| Seriola dume | A0A3B4UM85 |

| | |
|---|---|
| Seriola laland | A0A3B4X7F7 |
| Seriola laland | A0A3B4WXD7 |
| Seriola laland | A0A3B4WCE8 |
| Seriola laland | A0A3B4WX70 |
| Seriola laland | A0A3B4X514 |
| Seriola laland | A0A3B4Y8L3 |
| Setaria italica | K4A2F5 |
| Setaria italica | K3Z304 |
| Setaria italica | A0A368R0J1 |
| Setaria italica | A0A368REB7 |
| Setaria italica | K3XQN4 |
| Setaria italica | A0A368RTU8 |
| Setaria italica | A0A368RZT1 |
| Setaria italica | A0A368RZL0 |
| Setaria italica | A0A368PG49 |
| Setaria italica | A0A368SB62 |
| Setaria italica | A0A368RZH4 |
| Setaria italica | K3XPE2 |
| Setaria italica | K3XSM1 |
| Setaria italica | K3Z530 |
| Setaria italica | A0A368R959 |
| Setaria italica | A0A368R9A6 |
| Setaria italica | K3ZF16 |
| Setaria italica | A0A368QBK5 |
| Setaria italica | A0A368QCH9 |
| Setaria italica | A0A368Q4U7 |
| Setaria italica | A0A368Q482 |
| Setaria italica | K3ZPX7 |
| Setaria italica | A0A368RTX5 |
| Setaria italica | A0A368RTT9 |
| Setaria italica | K3XPE3 |
| Setaria italica | K3YDB8 |
| Setaria italica | A0A368RS21 |
| Setaria italica | K3YDQ0 |
| Setaria italica | A0A368RU04 |
| Setaria italica | A0A368QK95 |
| Setaria italica | A0A368RIC1 |
| Setaria italica | A0A368QBY1 |
| Setaria italica | A0A368R0I2 |
| Setaria italica | A0A368QB50 |
| Setaria italica | A0A368R8H3 |

| | |
|---|---|
| Setaria italica | K4A740 |
| Setaria italica | A0A368RSJ9 |
| Setaria italica | A0A368QB49 |
| Setaria italica | A0A368QCI5 |
| Setaria italica | A0A368R1M3 |
| Setaria italica | A0A368Q1T4 |
| Setaria italica | A0A368RZS3 |
| Setaria italica | K3YDS3 |
| Setaria italica | K3Y1Y9 |
| Setaria italica | A0A368Q4R1 |
| Setaria italica | K3ZCG7 |
| Setaria italica | K3XFB5 |
| Setaria italica | A0A368RZL6 |
| Setaria italica | A0A368RZL1 |
| Setaria italica | A0A368RZK1 |
| Setaria italica | K4A6H6 |
| Setaria italica | A0A368RZK0 |
| Setaria italica | K3YPU6 |
| Setaria italica | K3YEW1 |
| Setaria italica | K3YGJ6 |
| Setaria italica | K3XSH2 |
| Setaria italica | A0A368Q490 |
| Setaria italica | A0A368SI17 |
| Setaria italica | A0A368RZG9 |
| Setaria italica | A0A368RZG3 |
| Setaria italica | K3XEF3 |
| Setaria italica | K3YD20 |
| Setaria italica | A0A368RBT8 |
| Setaria italica | K3ZQI9 |
| Setaria italica | K3Y585 |
| Setaria italica | A0A368Q289 |
| Setaria italica | A0A368R1M5 |
| Setaria italica | A0A368R0L6 |
| Setaria italica | K3Y129 |
| Setaria italica | A0A368SAN2 |
| Setaria italica | A0A368Q6K5 |
| Setaria italica | A0A368Q6G7 |
| Setaria italica | A0A368Q569 |
| Setaria italica | K3Y505 |
| Setaria italica | K3ZZR5 |
| Setaria italica | K3ZRF9 |

Setaria italica A0A368RZI0
Setaria italica A0A368RZJ2
Setaria italica A0A368Q4J9
Setaria italica A0A368QR38
Setaria italica A0A368QXB9
Setaria italica A0A368Q621
Setaria italica A0A368PRU5
Setaria italica A0A368S195
Setaria italica A0A368RZM3
Setaria italica K3YD40
Setaria italica A0A368RZK9
Setaria italica A0A368R030
Setaria italica A0A368PK67
Setaria italica A0A368Q4N4
Setaria italica A0A368RU08
Setaria italica A0A368RZE2
Setaria italica A0A368RBV9
Setaria italica A0A368RZK6
Setaria italica K3XVB7
Setaria italica K3XIC1
Setaria italica A0A368R502
Setaria italica K3XEI2
Setaria italica K4A357
Setaria italica A0A368R9C0
Setaria italica A0A368QYB1
Setaria italica A0A368Q4M4
Setaria italica A0A368QBF6
Setaria italica K3Y2F6
Setaria italica A0A368S2P0
Setaria italica A0A368R035
Setaria italica A0A368RVM9
Setaria italica A0A368RZF9
Setaria italica K3XEM4
Setaria italica K3Y572
Setaria italica A0A368SR44
Setaria italica A0A368QMF4
Setaria italica K3YCM4
Setaria italica K4ALT0
Setaria italica K4A323
Setaria italica A0A368QPX0
Setaria italica A0A368RZI2

| | |
|---|---|
| Setaria italica | A0A368Q570 |
| Setaria italica | K4A115 |
| Setaria italica | K4AMY1 |
| Setaria italica | A0A368R970 |
| Setaria italica | K3Y5N0 |
| Setaria italica | A0A368RBZ6 |
| Setaria italica | A0A368RE63 |
| Setaria italica | A0A368SH77 |
| Setaria italica | A0A368RZR9 |
| Setaria italica | A0A368Q474 |
| Setaria italica | K3YEX8 |
| Setaria italica | A0A368R046 |
| Setaria italica | K4A086 |
| Setaria italica | A0A368RZF3 |
| Setaria italica | A0A368RH05 |
| Setaria italica | K3ZLB6 |
| Setaria italica | A0A368RRV9 |
| Setaria italica | K3Y545 |
| Setaria italica | K3Y5K2 |
| Setaria italica | A0A368Q6F6 |
| Setaria italica | K3YDM6 |
| Setaria italica | K4A5Z2 |
| Setaria italica | A0A368QBG1 |
| Setaria italica | K3XVB8 |
| Setaria italica | A0A368QJJ5 |
| Setaria italica | A0A368RZN4 |
| Setaria italica | K3YCU3 |
| Setaria italica | K3Z037 |
| Setaria italica | A0A0Q3PQD3 |
| Setaria italica | A0A368RFZ5 |
| Setaria italica | A0A368SP74 |
| Setaria italica | A0A368R3U7 |
| Setaria italica | A0A368RZU1 |
| Setaria italica | A0A368SII8 |
| Setaria italica | A0A368QW32 |
| Setaria italica | A0A368S227 |
| Setaria italica | A0A368RG75 |
| Setaria italica | A0A368RAQ1 |
| Setaria italica | A0A368Q480 |
| Setaria italica | A0A368RZI6 |
| Setaria italica | A0A368QC37 |

| | |
|---|---|
| Setaria italica | K3Z3T5 |
| Setaria italica | A0A368STV8 |
| Setaria italica | A0A368RZM9 |
| Setaria italica | A0A368QCP0 |
| Setaria italica | A0A368SC73 |
| Setaria italica | A0A368R950 |
| Setaria italica | K4A5S5 |
| Setaria italica | K3XQB1 |
| Setaria italica | A0A368SB67 |
| Setaria italica | K3ZQM2 |
| Setaria italica | K3YDM0 |
| Setaria italica | A0A368RZM0 |
| Setaria italica | A0A368R4P0 |
| Setaria italica | A0A368SBE7 |
| Setaria italica | K3YDB1 |
| Setaria italica | K4A1K7 |
| Setaria italica | K3ZQP3 |
| Setaria italica | K3Y559 |
| Setaria italica | A0A368SP77 |
| Setaria italica | K3XEG7 |
| Setaria italica | A0A368R690 |
| Setaria italica | K3Z3U1 |
| Setaria italica | K3XEE4 |
| Setaria italica | K3Y247 |
| Setaria italica | K3YC50 |
| Setaria italica | K4A5Y3 |
| Setaria italica | A0A368R8Y6 |
| Setaria italica | K3Y179 |
| Setaria italica | K3ZQP4 |
| Setaria italica | K4A5Y7 |
| Setosphaeria | R0IH14 |
| Setosphaeria | R0JVZ7 |
| Setosphaeria | R0K2V0 |
| Shimia abyssi | A0A2P8FBJ9 |
| Shimia aestu | A0A1I4K541 |
| Shimia haliot | A0A1I4DH54 |
| Shimia marin | A0A0P1FE41 |
| Shimia sp. SK | A0A0N0V3A4 |
| Shinella sp. D | A0A021XGD0 |
| Shinella sp. H | A0A1A9FVP4 |
| Shinella sp. W | A0A372NA60 |

| | |
|---|---|
| Shinella sp. W | A0A372NIR6 |
| Shinella sp..xl | A0A2W6YNN0 |
| Shinella sp..xl | A0A2W6XTN2 |
| Shinella sp..xl | A0A2W6TGS4 |
| Silicibacter sp | C9D0Y9 |
| Sinapis alba.x | U3M974 |
| Sinapis alba.x | Q8LP48 |
| Sinapis arven | Q8LP47 |
| Sinapis arven | Q9SEC7 |
| Sinapis arven | Q8LP46 |
| Sinapis arven | Q9SEB7 |
| Sinapis arven | Q9SEC6 |
| Sinirhodobac | A0A443L654 |
| Sinirhodobac | A0A443KHG8 |
| Sinirhodobac | A0A443IWD5 |
| Sinirhodobac | A0A443LU18 |
| Sinirhodobac | A0A443KBX3 |
| Sinirhodobac | A0A443JST8 |
| Sinobacterium | A0A3N2DQJ3 |
| Sinorhizobium | A0A1L3LEX8 |
| Sinorhizobium | A0A1L3KYZ3 |
| Sinorhizobium | A0A2S3YKP1 |
| Sinorhizobium | A0A2S3YS27 |
| Sinorhizobium | A0A1L3LHN3 |
| Sinorhizobium | A0A1L3LX24 |
| Sinorhizobium | D1CSK6 |
| Sinorhizobium | C3KRM2 |
| Sinorhizobium | A0A2Z3G701 |
| Sinorhizobium | A0A2U8GC90 |
| Sinorhizobium | A0A249Q4C1 |
| Sinorhizobium | A0A0T6ZYX4 |
| Sinorhizobium | I3X127 |
| Sinorhizobium | I3XBU9 |
| Sinorhizobium | A0A0E0UPB7 |
| Sinorhizobium | F7XGE8 |
| Sinorhizobium | H0G729 |
| Sinorhizobium | A0A178Y5H3 |
| Sinorhizobium | A0A178YA01 |
| Sinorhizobium | A0A1V3R0M3 |
| Sinorhizobium | A0A2A6PA40 |
| Sinorhizobium | A0A249P2H9 |

| | |
|---|---|
| Sinorhizobiur | A0A249NZ41 |
| Sinorhizobiur | A0A0T6ZY03 |
| Sinorhizobiur | A0A0T6YWU1 |
| Sinorhizobiur | A0A226CRW2 |
| Sinorhizobiur | A0A2N8JWB1 |
| Sinorhizobiur | A0A1G5WW04 |
| Sinorhizobiur | A0A2A6MGF3 |
| Sinorhizobiur | A0A2A6MEJ3 |
| Sinorhizobiur | A0A1B3MF54 |
| Sinorhizobiur | A0A1B3MIS4 |
| Sinorhizobiur | A0A0T6XVR8 |
| Sinorhizobiur | A0A0T6Z662 |
| Sipha flava.xl | A0A2S2R1U8 |
| Sipha flava.xl | A0A2S2R590 |
| Sipha flava.xl | A0A2S2PZK0 |
| Sipha flava.xl | A0A2S2PXW4 |
| Sipha flava.xl | A0A2S2R5W8 |
| Sipha flava.xl | A0A2S2PY34 |
| Sipha flava.xl | A0A2S2PWP9 |
| Sipha flava.xl | A0A2S2Q0X7 |
| Sisymbrium i | R9S9Z3 |
| Sisymbrium i | R9SAI4 |
| Sisymbrium i | R9S9I9 |
| Soboliphyme | A0A183IUB1 |
| Soboliphyme | A0A183IUA7 |
| Soboliphyme | A0A183IL94 |
| Soboliphyme | A0A183J2V0 |
| Soboliphyme | A0A183IH80 |
| Soboliphyme | A0A183IL56 |
| Soboliphyme | A0A183ISK1 |
| Soboliphyme | A0A183J3L0 |
| Soboliphyme | A0A183IHX2 |
| Soboliphyme | A0A183IAW3 |
| Soboliphyme | A0A183IZI2 |
| Soboliphyme | A0A183IV49 |
| Soboliphyme | A0A183IMR6 |
| Soboliphyme | A0A183IBD8 |
| Soboliphyme | A0A183IAM1 |
| Soboliphyme | A0A183IN87 |
| Soboliphyme | A0A183IPU9 |
| Soboliphyme | A0A183IMI5 |

| | |
|---|---|
| Soboliphyme | A0A183J9K8 |
| Soboliphyme | A0A183ITM5 |
| Soboliphyme | A0A3P7ZKR8 |
| Soboliphyme | A0A183IID0 |
| Solanum bulk | Q17U57 |
| Solanum chac | A0A0V0GU91 |
| Solanum chac | A0A0V0IAL3 |
| Solanum chac | A0A0V0IL88 |
| Solanum chac | A0A0V0IT75 |
| Solanum chac | A0A0V0H401 |
| Solanum chac | A0A0V0GP59 |
| Solanum chac | A0A0V0IZC1 |
| Solanum chac | A0A0V0ITT0 |
| Solanum chac | A0A0V0ISB1 |
| Solanum chac | A0A0V0HK80 |
| Solanum chac | A0A0V0HB28 |
| Solanum chac | A0A0V0IWF8 |
| Solanum chac | A0A0V0I770 |
| Solanum chac | A0A0V0GWH3 |
| Solanum chac | A0A0V0HTT1 |
| Solanum chac | A0A0V0IQW6 |
| Solanum chac | A0A0V0ISZ6 |
| Solanum chac | A0A0V0HEP1 |
| Solanum chac | A0A0V0IW22 |
| Solanum chac | A0A0V0IQJ4 |
| Solanum chac | A0A0V0ITB2 |
| Solanum lyco | A0A3Q7FRL4 |
| Solanum lyco | A0A3Q7JRJ5 |
| Solanum lyco | A0A3Q7IBI2 |
| Solanum lyco | A0A3Q7GDZ4 |
| Solanum lyco | A0A3Q7F4P0 |
| Solanum lyco | K4CH55 |
| Solanum lyco | K4B9J0 |
| Solanum lyco | A0A3Q7FY38 |
| Solanum lyco | A0A3Q7HF37 |
| Solanum lyco | A0A3Q7HDB6 |
| Solanum lyco | A0A3Q7G892 |
| Solanum lyco | A0A3Q7G5Y6 |
| Solanum lyco | A0A3Q7F4L9 |
| Solanum lyco | A0A3Q7F595 |
| Solanum lyco | A0A3Q7G6E0 |

| | |
|---|---|
| Solanum lyco | A0A3Q7IAB2 |
| Solanum lyco | A0A3Q7G8E1 |
| Solanum lyco | A0A3Q7FWM1 |
| Solanum lyco | A0A3Q7EL09 |
| Solanum lyco | A0A3Q7J270 |
| Solanum lyco | A0A3Q7F4Q9 |
| Solanum lyco | A0A3Q7HBS1 |
| Solanum lyco | A0A3Q7HEV3 |
| Solanum lyco | A0A3Q7EWM9 |
| Solanum lyco | A0A3Q7F4N0 |
| Solanum lyco | A0A3Q7IAP8 |
| Solanum lyco | A0A3Q7F4K4 |
| Solanum lyco | A0A3Q7HYY7 |
| Solanum lyco | A0A3Q7G5X9 |
| Solanum lyco | A0A3Q7HTG3 |
| Solanum lyco | A0A3Q7FFS6 |
| Solanum lyco | A0A3Q7G8V9 |
| Solanum lyco | A0A3Q7HF02 |
| Solanum lyco | A0A3Q7GP81 |
| Solanum lyco | A0A3Q7F2Q7 |
| Solanum lyco | A0A3Q7HHX3 |
| Solanum lyco | A0A3Q7FRL0 |
| Solanum lyco | A0A3Q7HHD3 |
| Solanum lyco | A0A3Q7FRP6 |
| Solanum lyco | A0A3Q7F582 |
| Solanum lyco | A0A3Q7G5Z5 |
| Solanum lyco | A0A3Q7I091 |
| Solanum lyco | K4BE95 |
| Solanum lyco | A0A3Q7I1N1 |
| Solanum lyco | A0A3Q7IBL7 |
| Solanum lyco | A0A3Q7HQY4 |
| Solanum lyco | A0A3Q7INM2 |
| Solanum lyco | A0A3Q7G218 |
| Solanum lyco | A0A3Q7F3P0 |
| Solanum lyco | A0A3Q7I8J1 |
| Solanum lyco | A0A3Q7IBE5 |
| Solanum lyco | A0A3Q7I8I7 |
| Solanum pen | A0A0D4BTD1 |
| Solanum pen | A0A0D4BT35 |
| Solanum pen | A0A0D4BTY2 |
| Solanum tub | M1C323 |

| | |
|---|---|
| Solanum tube | M1BNV7 |
| Solanum tube | M1A4E2 |
| Solanum tube | M1A1M0 |
| Solanum tube | M1C8X1 |
| Solanum tube | M0ZGV4 |
| Solanum tube | M1CGZ5 |
| Solanum tube | M1B352 |
| Solanum tube | M1C1G6 |
| Solanum tube | M1DW58 |
| Solanum tube | M1BYZ9 |
| Solanum tube | M1BIT6 |
| Solanum tube | M1D080 |
| Solanum tube | M1B369 |
| Solanum tube | M1CJN9 |
| Solanum tube | M1BGH7 |
| Solanum tube | M0ZKJ6 |
| Solanum tube | M1A1J1 |
| Solanum tube | M1BZ01 |
| Solanum tube | M1BGB2 |
| Solanum tube | M1B373 |
| Solanum tube | M1BNW8 |
| Solanum tube | M1CVL9 |
| Solanum tube | M1BP27 |
| Solanum tube | M1BGH6 |
| Solanum tube | M1BP21 |
| Solanum tube | M1DRI9 |
| Solanum tube | M0ZGV5 |
| Solanum tube | M1A1J0 |
| Solanum tube | M1AFX4 |
| Solanum tube | A0A0M3STW8 |
| Solanum tube | M1BIU3 |
| Solanum tube | M1CRP0 |
| Solanum tube | M1CVM0 |
| Solanum tube | M1CGZ4 |
| Solanum tube | M1BNV6 |
| Solanum tube | M1AFZ1 |
| Solanum tube | M1D081 |
| Solanum tube | M1CH47 |
| Solanum tube | M0ZUR7 |
| Solanum tube | M1A1I5 |
| Solanum tube | M1BGH3 |

| | |
|---|---|
| Solanum tube | M1CVL4 |
| Solanum tube | M1C1G5 |
| Solanum tube | M1ASH7 |
| Solanum tube | M1BGG2 |
| Solanum tube | M1BNW9 |
| Solanum tube | M1A1M2 |
| Solanum tube | M1A1I7 |
| Solanum tube | M1C1G7 |
| Solanum tube | M1BP20 |
| Solanum tube | M1BNV8 |
| Solanum tube | M1BZ00 |
| Solanum tube | M1BNX0 |
| Solanum tube | M1A1I8 |
| Solanum tube | M1BGB0 |
| Solanum tube | M1DRP4 |
| Solanum tube | M1CGZ2 |
| Solanum tube | M1A226 |
| Solanum tube | M1CGZ3 |
| Solanum tube | M1A1I9 |
| Solanum tube | M1AJH1 |
| Solanum tube | M1B367 |
| Solanum tube | M1B363 |
| Solanum tube | M1B357 |
| Solanum tube | M1BIU2 |
| Solanum tube | M1BGA3 |
| Solanum tube | M1CVM1 |
| Solanum tube | M1AJH2 |
| Solanum tube | M1BNV5 |
| Solanum tube | M1BIU4 |
| Solanum tube | M1B356 |
| Solanum tube | M1AFY9 |
| Solanum tube | M1BGA2 |
| Solanum tube | M1C322 |
| Solanum tube | M1CGZ7 |
| Solanum tube | M1AFZ0 |
| Solanum tube | M1B366 |
| Solanum tube | M1DZ63 |
| Solanum tube | M1BGA7 |
| Solanum tube | M1BGA4 |
| Solanum tube | M1BGG5 |
| Solanum tube | M1C8X2 |

| | |
|---|---|
| Solanum tube | M1CC72 |
| Solanum tube | M1BNV4 |
| Solanum tube | M1BNW1 |
| Solanum tube | M1CJP1 |
| Solanum tube | M1BIT7 |
| Solanum tube | M1CVL6 |
| Solanum tube | M1A4E5 |
| Solanum tube | M1CH44 |
| Solanum tube | M1B371 |
| Solenopsis in | E9J8M2 |
| Solenopsis in | E9IHS5 |
| Solenopsis in | E9IN88 |
| Solenopsis in | E9J669 |
| Solenopsis in | E9J909 |
| Solenopsis in | E9J910 |
| Solenopsis in | E9IHS6 |
| Solenopsis in | E9IN89 |
| Sorangium ce | A0A150STA1 |
| Sordaria mac | F7VYZ6 |
| Sordaria mac | F7VR72 |
| Sordaria mac | F7W2U3 |
| Sorghum bicc | A0A1Z5R405 |
| Sorghum bicc | A0A1W0W663 |
| Sorghum bicc | A0A1Z5RKG3 |
| Sorghum bicc | A0A1Z5R8K9 |
| Sorghum bicc | C5YTR3 |
| Sorghum bicc | A0A1B6Q1T4 |
| Sorghum bicc | A0A1B6P7F5 |
| Sorghum bicc | A0A1W0W011 |
| Sorghum bicc | C5XIK1 |
| Sorghum bicc | C5X8J9 |
| Sorghum bicc | A0A1W0W6A4 |
| Sorghum bicc | A0A1B6QDZ9 |
| Sorghum bicc | A0A1Z5R8L3 |
| Sorghum bicc | C5XNJ5 |
| Sorghum bicc | A0A1B6PQ00 |
| Sorghum bicc | A0A1B6Q6I4 |
| Sorghum bicc | A0A1W0W4G9 |
| Sorghum bicc | C5YRQ1 |
| Sorghum bicc | A0A1Z5RF80 |
| Sorghum bicc | C5XKC4 |

| | |
|---|---|
| Sorghum bicc | A0A1B6QBX0 |
| Sorghum bicc | C5XG13 |
| Sorghum bicc | A0A1B6QJV5 |
| Sorghum bicc | A0A194YPP4 |
| Sorghum bicc | C5YI80 |
| Sorghum bicc | A0A1Z5S8X7 |
| Sorghum bicc | A0A1B6PN86 |
| Sorghum bicc | A0A1B6Q0T3 |
| Sorghum bicc | A0A1Z5RF61 |
| Sorghum bicc | A0A1B6P674 |
| Sorghum bicc | A0A1W0VTK0 |
| Sorghum bicc | A0A194YKI9 |
| Sorghum bicc | A0A1B6QP87 |
| Sorghum bicc | A0A1B6QBJ7 |
| Sorghum bicc | A0A1B6PNF9 |
| Sorghum bicc | A0A194YKJ5 |
| Sorghum bicc | C5Z879 |
| Sorghum bicc | A0A1Z5RCU0 |
| Sorghum bicc | A0A1B6QDZ0 |
| Sorghum bicc | A0A1W0W4H9 |
| Sorghum bicc | C5YFX5 |
| Sorghum bicc | A0A1B6QEU8 |
| Sorghum bicc | C5YZF4 |
| Sorghum bicc | A0A1B6QE05 |
| Sorghum bicc | C5XAW9 |
| Sorghum bicc | A0A1B6QPL1 |
| Sorghum bicc | C5YGN7 |
| Sorghum bicc | A0A1Z5RCV0 |
| Sorghum bicc | C5YGM5 |
| Sorghum bicc | A0A1Z5R1H2 |
| Sorghum bicc | A0A1Z5RGB4 |
| Sorghum bicc | C5XAX0 |
| Sorghum bicc | A0A1W0VYV0 |
| Sorghum bicc | A0A1B6P8A7 |
| Sorghum bicc | A0A1Z5RF79 |
| Sorghum bicc | C5XNJ6 |
| Sorghum bicc | C5XL73 |
| Sorghum bicc | A0A1Z5RF74 |
| Sorghum bicc | C5X9Y1 |
| Sorghum bicc | A0A1B6Q255 |
| Sorghum bicc | A0A1Z5RFW8 |

| | |
|---|---|
| Sorghum bicc | A0A1B6Q7G8 |
| Sorghum bicc | A0A1Z5RIL9 |
| Sorghum bicc | C5WQK6 |
| Sorghum bicc | A0A1Z5RKG6 |
| Sorghum bicc | A0A1W0W6M4 |
| Sorghum bicc | A0A194YHF9 |
| Sorghum bicc | A0A1B6P5Y4 |
| Sorghum bicc | A0A1B6PNE9 |
| Sorghum bicc | A0A1W0VTM8 |
| Sorghum bicc | A0A1W0VTM2 |
| Sorghum bicc | C5XLS4 |
| Sorghum bicc | A0A1Z5RFC4 |
| Sorghum bicc | A0A1Z5R535 |
| Sorghum bicc | C5YT42 |
| Sorghum bicc | A0A1B6PNG4 |
| Sorghum bicc | C5Y4M3 |
| Sorghum bicc | C5X494 |
| Sorghum bicc | C5YV85 |
| Sorghum bicc | C5WV97 |
| Sorghum bicc | A0A1Z5RF78 |
| Sorghum bicc | C5XSW0 |
| Sorghum bicc | C5X2A2 |
| Sorghum bicc | C5YGP6 |
| Sorghum bicc | A0A1B6QE00 |
| Sorghum bicc | A0A1W0W6E0 |
| Sorghum bicc | A0A1B6QBX3 |
| Sorghum bicc | A0A1B6PNF5 |
| Sorghum bicc | A0A1Z5S3X9 |
| Sorghum bicc | A0A1W0W6U5 |
| Sorghum bicc | A0A1W0W672 |
| Sorghum bicc | A0A1B6Q6J1 |
| Sorghum bicc | C5YGP5 |
| Sorghum bicc | A0A1W0VSW8 |
| Sorghum bicc | A0A1B6QBW8 |
| Sorghum bicc | A0A1B6QBW6 |
| Sorghum bicc | C5YI84 |
| Sorghum bicc | Q8LJZ0 |
| Sphaceloma ı | A0A2K1QVG0 |
| Sphaceloma ı | A0A2K1QHN6 |
| Sphaceloma ı | A0A2K1QKQ3 |
| Sphaceloma ı | A0A2K1QK63 |

| | |
|---|---|
| Sphaceloma r | A0A2K1QSX4 |
| Sphaceloma r | A0A2K1QFH6 |
| Sphaceloma r | A0A2K1QKW0 |
| Sphaceloma r | A0A2K1QK44 |
| Sphaceloma r | A0A2K1QZH3 |
| Sphaceloma r | A0A2K1QY48 |
| Sphaceloma r | A0A2K1QX05 |
| Sphaceloma r | A0A2K1QSH2 |
| Sphaceloma r | A0A2K1R2J1 |
| Sphaceloma r | A0A2K1QRG3 |
| Sphaceloma r | A0A2K1QKB9 |
| Sphaceloma r | A0A2K1QGJ2 |
| Sphaceloma r | A0A2K1R3T9 |
| Sphaerulina r | M3AV53 |
| Sphaerulina r | M3BZM1 |
| Sphingobacte | A0A349QUZ7 |
| Sphingobacte | A0A354KFP2 |
| Sphingomona | A0A2T7WNY4 |
| Sphingomona | A0A2T7WSG0 |
| Sphingopyxis | A0A1L3JC18 |
| Sphingopyxis | A0A1L3JC23 |
| Sphingopyxis | A0A1L3JC30 |
| Sphingorhabo | A0A371B272 |
| Sphingorhabo | A0A371BHY4 |
| Sphingorhabo | A0A371BHT8 |
| Spinacia olera | A0A0K9RR85 |
| Spinacia olera | A0A0K9RDP3 |
| Spinacia olera | A0A0K9QHY2 |
| Spinacia olera | A0A0K9QR37 |
| Spinacia olera | A0A0K9R7Z2 |
| Spinacia olera | A0A0K9Q8S7 |
| Spinacia olera | A0A0K9S0X2 |
| Spinacia olera | A0A0K9QPL4 |
| Spinacia olera | A0A0K9RF56 |
| Spinacia olera | A0A0K9S213 |
| Spinacia olera | A0A0K9RH17 |
| Spinacia olera | A0A0K9RT23 |
| Spinacia olera | A0A0K9RB42 |
| Spinacia olera | A0A0K9QZ62 |
| Spinacia olera | A0A0K9RVT8 |
| Spinacia olera | A0A0K9R8M3 |

| | |
|---|---|
| Spinacia olera | A0A0K9Q888 |
| Spinacia olera | A0A0K9RMQ3 |
| Spinacia olera | A0A0K9QW25 |
| Spinacia olera | A0A0K9S082 |
| Spinacia olera | A0A0K9R4K7 |
| Spinacia olera | A0A0K9QRC1 |
| Spizellomyces | A0A0L0HA63 |
| Spizellomyces | A0A0L0H8G6 |
| Spodoptera f | A0A2H1WV32 |
| Spodoptera f | A0A2H1VNI1 |
| Spodoptera f | A0A2H1VQJ4 |
| Spodoptera f | A0A2H1WDX9 |
| Spodoptera f | A0A2H1W4L3 |
| Spongospora | A0A0H5QFI6 |
| Spongospora | A0A0H5QHL0 |
| Sporothrix br | A0A0C2ER25 |
| Sporothrix in | A0A162MRJ9 |
| Sporothrix in | A0A167PWC0 |
| Sporothrix sc | U7Q5P1 |
| Sporothrix sc | A0A0F2LUQ6 |
| Stachybotrys | A0A084B8D0 |
| Stachybotrys | A0A084AUC4 |
| Stachybotrys | A0A084AH70 |
| Stachybotrys | A0A084AW91 |
| Stachybotrys | A0A084R5E6 |
| Stachybotrys | A0A084R699 |
| Stachybotrys | A0A084RSS0 |
| Stachybotrys | A0A084R0N3 |
| Stachybotrys | A0A084QS85 |
| Stachybotrys | A0A084QWE9 |
| Stachybotrys | A0A084QR64 |
| Stagonospora | A0A178AM66 |
| Stagonospora | A0A178B4D4 |
| Stagonospora | A0A178B8B6 |
| Stagonospora | A0A178AC45 |
| Stagonospora | A0A178AYG8 |
| Stagonospora | A0A178B8W6 |
| Stagonospora | A0A178AV13 |
| Stagonospora | A0A178B2D4 |
| Stappia indica | A0A285S9J4 |
| Stappia sp. 22 | A0A1Y1SPA5 |

| | |
|---|---|
| Stappia sp. AI | A0A420AAV3 |
| Stappia sp. ES | A0A1H2FDZ7 |
| Stappia sp..xl | A0A2E6Y7I2 |
| Stegastes par | A0A3B4ZKK7 |
| Stegastes par | A0A3B5B6X3 |
| Stegastes par | A0A3B4ZPZ3 |
| Stegastes par | A0A3B4ZIA4 |
| Stegastes par | A0A3B5ADV9 |
| Stegastes par | A0A3B4ZH28 |
| Stegastes par | A0A3B4ZII7 |
| Stegodyphus | A0A087T2K8 |
| Stegodyphus | A0A087U8M8 |
| Stegodyphus | A0A087UVI0 |
| Stegodyphus | A0A087UVI1 |
| Stegodyphus | A0A087UVH9 |
| Stegodyphus | A0A087T2K7 |
| Stegodyphus | A0A087TCV3 |
| Stegodyphus | A0A087SWB4 |
| Stegodyphus | A0A087UKG6 |
| Stegodyphus | A0A087SVJ4 |
| Stegodyphus | A0A087UKG5 |
| Stegodyphus | A0A087UBD7 |
| Stegodyphus | A0A087UAP8 |
| Stegodyphus | A0A087TMG3 |
| Stegodyphus | A0A087T9G3 |
| Stegodyphus | A0A087T7G6 |
| Stegodyphus | A0A087T2C9 |
| Stegodyphus | A0A087T0G3 |
| Stegodyphus | A0A087TEI6 |
| Stegodyphus | A0A087UVX7 |
| Stegodyphus | A0A087THX0 |
| Steinernema | A0A1I8AEG0 |
| Steinernema | A0A1I7Y5R0 |
| Steinernema | A0A1I8AKS5 |
| Steinernema | A0A1I8AL78 |
| Steinernema | A0A1I7Y544 |
| Steinernema | A0A1I8A6X3 |
| Steinernema | A0A1I8ATW3 |
| Steinernema | A0A1I8AU60 |
| Steinernema | A0A1I7YBL2 |
| Steinernema | A0A1I8A4V1 |

| | |
|---|---|
| Steinernema | A0A1I7Z6J2 |
| Steinernema | A0A1I8A5G4 |
| Steinernema | A0A1I7YSK9 |
| Steinernema | A0A1I7YBL9 |
| Steinernema | A0A1I7Y0N0 |
| Steinernema | A0A1I7ZHS2 |
| Steinernema | A0A1I7Y6B7 |
| Steinernema | A0A1I7YB77 |
| Steinernema | A0A1I7ZQD0 |
| Steinernema | A0A1I7ZLK9 |
| Steinernema | A0A1I7ZW10 |
| Steinernema | A0A1I7Y539 |
| Steinernema | A0A1I7Z2T9 |
| Steinernema | A0A1I7YQG9 |
| Steinernema | A0A1I7ZVA4 |
| Steinernema | A0A1I7YYH6 |
| Steinernema | A0A1I8AAT9 |
| Steinernema | A0A1I8AAI7 |
| Steinernema | A0A1I7Z483 |
| Steinernema | A0A1I8ARD3 |
| Stemphylium | A0A364NG07 |
| Stemphylium | A0A364N8Y3 |
| Stemphylium | A0A364NBE1 |
| Stichopus jap | A0A2G8KGD5 |
| Stichopus jap | A0A2G8KPK3 |
| Stichopus jap | A0A2G8KY26 |
| Stichopus jap | A0A2G8JWF4 |
| Stichopus jap | A0A2G8KY13 |
| Stichopus jap | A0A2G8KMR0 |
| Stichopus jap | A0A2G8K0N1 |
| Stichopus jap | A0A2G8LND1 |
| Stichopus jap | A0A2G8LNU1 |
| Stichopus jap | A0A2G8JX12 |
| Stichopus jap | A0A2G8KPX2 |
| Stichopus jap | A0A2G8KPE7 |
| Stichopus jap | A0A2G8K4I4 |
| Stichopus jap | A0A2G8K4G4 |
| Stichopus jap | A0A2G8JWF7 |
| Stichopus jap | A0A2G8L1A0 |
| Stichopus jap | A0A2G8KPD9 |
| Stichopus jap | A0A2G8KPE6 |

| | |
|---|---|
| Stichopus jap | A0A2G8L837 |
| Stichopus jap | A0A2G8LH26 |
| Stigmatella a | E3FLQ8 |
| Stigmatella a | Q098U7 |
| Stomoxys cal | A0A1I8PQ32 |
| Stomoxys cal | A0A1I8QFD3 |
| Stomoxys cal | A0A1I8PLI1 |
| Stomoxys cal | A0A1I8NR90 |
| Stomoxys cal | A0A1I8QDM2 |
| Stomoxys cal | A0A1I8PQE0 |
| Stomoxys cal | A0A1I8QFE6 |
| Stomoxys cal | A0A1I8QFB9 |
| Stomoxys cal | A0A1I8NPE1 |
| Strigamia ma | T1JBA9 |
| Strigamia ma | T1ILR6 |
| Strigamia ma | T1IZB1 |
| Strigamia ma | T1IYR6 |
| Strigamia ma | T1J1D9 |
| Strigamia ma | T1J1D8 |
| Strigamia ma | T1J1E1 |
| Strigamia ma | T1J2Z9 |
| Strigamia ma | T1IYR5 |
| Strigamia ma | T1IUF2 |
| Strigamia ma | T1IWK2 |
| Strongylocen | W4YFT8 |
| Strongylocen | W4XEH1 |
| Strongylocen | W4Y9R3 |
| Strongylocen | W4Y7C8 |
| Strongylocen | W4Y7X2 |
| Strongylocen | W4ZAY8 |
| Strongylocen | W4YX16 |
| Strongylocen | W4ZEP3 |
| Strongylocen | W4Z4Q9 |
| Strongylocen | W4Y1M2 |
| Strongylocen | W4Z490 |
| Strongylocen | W4XV31 |
| Strongylocen | W4XT52 |
| Strongylocen | W4Y4J9 |
| Strongylocen | W4YJJ7 |
| Strongylocen | W4Y4H6 |
| Strongylocen | W4XV55 |

Strongylocen W4XRN9
Strongylocen W4YFP9
Strongylocen W4ZK20
Strongylocen W4YB28
Strongylocen W4ZB03
Strongylocen W4YM62
Strongylocen W4ZIR0
Strongylocen W4YUE3
Strongylocen W4Z6A9
Strongylocen W4Y9R4
Strongylocen W4Z5T2
Strongylocen W4ZD80
Strongylocen W4XET4
Strongylocen W4XV33
Strongylocen W4Y748
Strongylocen W4YH25
Strongylocen W4YVK8
Strongylocen W4Y2F2
Strongylocen W4YXL4
Strongylocen W4Z1A9
Strongylocen W4YQ09
Strongylocen W4XPT9
Strongylocen W4ZG23
Strongylocen W4Y9H1
Strongylocen W4XN66
Strongylocen W4Z197
Strongylocen W4ZKJ1
Strongylocen W4YMZ9
Strongylocen W4Z0S4
Strongylocen W4YI87
Strongylocen W4Y1N4
Strongylocen W4Z525
Strongylocen W4XT71
Strongylocen W4XAZ6
Strongylocen W4Y432
Strongylocen W4Z5T3
Strongylocen W4XGA6
Strongylocen W4Z430
Strongylocen W4XDS2
Strongylocen W4YJJ8
Strongylocen W4Y9R5

| | |
|---|---|
| Strongylocen | W4XX39 |
| Strongylocen | W4Z280 |
| Strongyloides | A0A0N5BIA2 |
| Strongyloides | A0A0N5CCT3 |
| Strongyloides | A0A0N5CGC3 |
| Strongyloides | A0A0N5BMA1 |
| Strongyloides | A0A0N5B5D0 |
| Strongyloides | A0A0N5CGC2 |
| Strongyloides | A0A0N5BQ06 |
| Strongyloides | A0A0N5B5E6 |
| Strongyloides | A0A0N5BCK0 |
| Strongyloides | A0A0N5B3M3 |
| Strongyloides | A0A0N5C548 |
| Strongyloides | A0A0N5C796 |
| Strongyloides | A0A0N5BRL1 |
| Strongyloides | A0A0N5BF41 |
| Strongyloides | A0A0N5C8M1 |
| Strongyloides | A0A0N5C2G3 |
| Strongyloides | A0A0N5C549 |
| Strongyloides | A0A0N5C5W7 |
| Strongyloides | A0A0N5BTC5 |
| Strongyloides | A0A090MZF3 |
| Strongyloides | A0A090LAM2 |
| Strongyloides | A0A090KSC6 |
| Strongyloides | A0A090KTL6 |
| Strongyloides | A0A090MTX8 |
| Strongyloides | A0A090L5S9 |
| Strongyloides | A0A090KXK0 |
| Strongyloides | A0A090KWR4 |
| Strongyloides | A0A090KR38 |
| Strongyloides | A0A090LBK0 |
| Strongyloides | A0A090KZM2 |
| Strongyloides | A0A090LMM7 |
| Strongyloides | A0A090LMB5 |
| Strongyloides | A0A090L0U9 |
| Strongyloides | A0A090L8Q0 |
| Strongyloides | A0A090LEE0 |
| Strongyloides | A0A090KR99 |
| Strongyloides | A0A090LS81 |
| Strongyloides | A0A090LH59 |
| Strongyloides | A0A090L2B1 |

| | |
|---|---|
| Strongyloides | A0A090KZU3 |
| Strongyloides | A0A0K0EQ10 |
| Strongyloides | A0A0K0EN25 |
| Strongyloides | A0A0K0ENZ3 |
| Strongyloides | A0A0K0EPC1 |
| Strongyloides | A0A0K0E3P4 |
| Strongyloides | A0A0K0E4B4 |
| Strongyloides | A0A0K0EFB6 |
| Strongyloides | A0A0K0ES05 |
| Strongyloides | A0A0K0EEE0 |
| Strongyloides | A0A0K0ESE2 |
| Strongyloides | A0A0K0E6Z7 |
| Strongyloides | A0A0K0DU24 |
| Strongyloides | A0A0K0E1P0 |
| Strongyloides | A0A0K0EGC5 |
| Strongyloides | A0A0K0EE54 |
| Strongyloides | A0A0K0DX88 |
| Strongyloides | A0A0K0DTQ3 |
| Strongyloides | A0A0K0EGC4 |
| Strongyloides | A0A0K0E8Q0 |
| Strongyloides | A0A0K0EJ96 |
| Strongyloides | A0A0K0E4B5 |
| Strongyloides | A0A0K0FAY7 |
| Strongyloides | A0A0K0FYJ7 |
| Strongyloides | A0A0K0F449 |
| Strongyloides | A0A0K0F450 |
| Strongyloides | A0A0K0F9B8 |
| Strongyloides | A0A0K0FA09 |
| Strongyloides | A0A0K0F2N2 |
| Strongyloides | A0A0K0G164 |
| Strongyloides | A0A0K0FJN2 |
| Strongyloides | A0A0K0FCQ4 |
| Strongyloides | A0A0K0G4U8 |
| Strongyloides | A0A0K0EWF5 |
| Strongyloides | A0A0K0F5J9 |
| Strongyloides | A0A0K0F3F0 |
| Strongyloides | A0A0K0EUB2 |
| Strongyloides | A0A0K0F008 |
| Strongyloides | A0A0K0FCQ5 |
| Strongyloides | A0A0K0EZ99 |
| Strongyloides | A0A0K0F0L9 |

| | |
|---|---|
| Strongyloides | A0A0K0FEV0 |
| Strongylus vu | A0A3P7ILC4 |
| Strongylus vu | A0A3P7I9T4 |
| Strongylus vu | A0A3P7JE85 |
| Strongylus vu | A0A3P7IEU2 |
| Strongylus vu | A0A3P7KH13 |
| Strongylus vu | A0A3P7IJW3 |
| Strongylus vu | A0A3P7IZH6 |
| Strongylus vu | A0A3P7I491 |
| Strongylus vu | A0A3P7LAH1 |
| Strongylus vu | A0A3P7J5F1 |
| Strongylus vu | A0A3P7IRM1 |
| Strongylus vu | A0A3P7JJX8 |
| Strongylus vu | A0A3P7J7H7 |
| Strongylus vu | A0A3P7IN45 |
| Strongylus vu | A0A3P7JMN8 |
| Strongylus vu | A0A3P7INY8 |
| Strongylus vu | A0A3P7LGM1 |
| Strongylus vu | A0A3P7J2L6 |
| Strongylus vu | A0A3P7JB01 |
| Strongylus vu | A0A3P7JA70 |
| Strongylus vu | A0A3P7L1J1 |
| Struthio came | A0A093KD36 |
| Struthio came | A0A093HHU1 |
| Struthio came | A0A093HGM9 |
| Stylophora pi | A0A2B4SM39 |
| Stylophora pi | A0A2B4RRZ6 |
| Stylophora pi | A0A2B4SJV0 |
| Stylophora pi | A0A2B4SAG5 |
| Stylophora pi | A0A2B4SES4 |
| Stylophora pi | A0A2B4RBI1 |
| Stylophora pi | A0A2B4S2H8 |
| Stylophora pi | A0A2B4S3A5 |
| Stylophora pi | A0A2B4SNH9 |
| Stylophora pi | A0A2B4S207 |
| Stylophora pi | A0A2B4S6F9 |
| Stylophora pi | A0A2B4R7J5 |
| Stylophora pi | A0A2B4SFG4 |
| Stylophora pi | A0A2B4S741 |
| Stylophora pi | A0A2B4SJX2 |
| Stylophora pi | A0A2B4RAJ9 |

Stylophora pi A0A2B4SUX3
Stylophora pi A0A2B4SH64
Stylophora pi A0A2B4SU12
Stylophora pi A0A2B4R4B9
Stylophora pi A0A2B4S3B6
Stylophora pi A0A2B4R832
Stylophora pi A0A2B4SXX8
Stylophora pi A0A2B4SUD0
Stylophora pi A0A2B4R2R9
Stylophora pi A0A2B4RFK9
Stylophora pi A0A2B4RMT1
Stylophora pi A0A2B4R623
Stylophora pi A0A2B4R7E0
Stylophora pi A0A2B4RD03
Stylophora pi A0A2B4S9T7
Stylophora pi A0A2B4RFP1
Stylophora pi A0A2B4RPK6
Stylophora pi A0A2B4R401
Stylophora pi A0A2B4RAC0
Stylophora pi A0A2B4RS71
Stylophora pi A0A2B4RXZ8
Stylophora pi A0A2B4T1E8
Stylophora pi A0A2B4RAX9
Stylophora pi A0A2B4R5W3
Stylophora pi A0A2B4REV2
Stylophora pi A0A2B4RBF8
Stylophora pi A0A2B4SUP0
Stylophora pi A0A2B4RLC2
Stylophora pi A0A2B4T0K3
Stylophora pi A0A2B4RYT3
Stylophora pi A0A2B4RW39
Stylophora pi A0A2B4RR37
Stylophora pi A0A2B4RK16
Stylophora pi A0A2B4SDB6
Stylophora pi A0A2B4SD97
Stylophora pi A0A2B4R0H8
Stylophora pi A0A2B4SY37
Stylophora pi A0A2B4RMM4
Stylophora pi A0A2B4R9P6
Stylophora pi A0A2B4SAP0
Stylophora pi A0A2B4S7C1

Stylophora pi A0A2B4T076
Stylophora pi A0A2B4RQ27
Stylophora pi A0A2B4RZ31
Stylophora pi A0A2B4SW57
Stylophora pi A0A2B4RQZ6
Stylophora pi A0A2B4SW29
Stylophora pi A0A2B4SXJ7
Stylophora pi A0A2B4R394
Stylophora pi A0A2B4REL3
Stylophora pi A0A2B4SKC3
Stylophora pi A0A2B4RCQ4
Stylophora pi A0A2B4RCF9
Stylophora pi A0A2B4SGL1
Stylophora pi A0A2B4SKU4
Stylophora pi A0A2B4S352
Stylophora pi A0A2B4RR27
Stylophora pi A0A2B4R5T8
Stylophora pi A0A2B4RIF0
Stylophora pi A0A2B4SI30
Stylophora pi A0A2B4RF56
Stylophora pi A0A2B4S048
Stylophora pi A0A2B4S8S0
Stylophora pi A0A2B4RCW0
Stylophora pi A0A2B4S421
Stylophora pi A0A2B4S3Y9
Stylophora pi A0A2B4SWB5
Stylophora pi A0A2B4RNZ9
Stylophora pi A0A2B4RKY5
Stylophora pi A0A2B4SYK2
Stylophora pi A0A2B4RCU9
Stylophora pi A0A2B4RG52
Stylophora pi A0A2B4SQ47
Stylophora pi A0A2B4SFY8
Stylophora pi A0A2B4SEZ0
Stylophora pi A0A2B4RQ71
Stylophora pi A0A2B4RI82
Stylophora pi A0A2B4RXE4
Stylophora pi A0A2B4R8V7
Stylophora pi A0A2B4SNN3
Stylophora pi A0A2B4S8F7
Stylophora pi A0A2B4R5T1

Stylophora pi A0A2B4RA51
Stylophora pi A0A2B4RNS9
Stylophora pi A0A2B4RAH7
Stylophora pi A0A2B4RTJ2
Stylophora pi A0A2B4SKD6
Stylophora pi A0A2B4RCL9
Stylophora pi A0A2B4S0E1
Stylophora pi A0A2B4S0Y6
Stylophora pi A0A2B4RJW9
Stylophora pi A0A2B4SZU0
Stylophora pi A0A2B4SRH6
Stylophora pi A0A2B4T229
Stylophora pi A0A2B4S0G9
Stylophora pi A0A2B4S040
Stylophora pi A0A2B4RDG4
Stylophora pi A0A2B4R9R0
Stylophora pi A0A2B4R5S5
Stylophora pi A0A2B4SPN4
Stylophora pi A0A2B4S8L9
Stylophora pi A0A2B4SBF7
Stylophora pi A0A2B4R907
Stylophora pi A0A2B4RDE8
Stylophora pi A0A2B4RFA7
Stylophora pi A0A2B4S2N4
Stylophora pi A0A2B4SVB4
Stylophora pi A0A2B4SXM9
Stylophora pi A0A2B4RIY2
Stylophora pi A0A2B4RLZ3
Stylophora pi A0A2B4S0I5
Stylophora pi A0A2B4S7S6
Stylophora pi A0A2B4RYB4
Stylophora pi A0A2B4SE06
Stylophora pi A0A2B4S470
Stylophora pi A0A2B4R8S2
Stylophora pi A0A2B4SYR5
Stylophora pi A0A2B4RI65
Stylophora pi A0A2B4RE37
Stylophora pi A0A2B4RJT3
Stylophora pi A0A2B4RHB2
Stylophora pi A0A2B4S982
Stylophora pi A0A2B4SI73

| | |
|---|---|
| Stylophora pi | A0A2B4RI05 |
| Stylophora pi | A0A2B4RX94 |
| Stylophora pi | A0A2B4SNE8 |
| Stylophora pi | A0A2B4RJP3 |
| Stylophora pi | A0A2B4SBI8 |
| Stylophora pi | A0A2B4R7I8 |
| Stylophora pi | A0A2B4RED5 |
| Stylophora pi | A0A2B4R819 |
| Stylophora pi | A0A2B4RAT8 |
| Stylophora pi | A0A2B4RDB8 |
| Stylophora pi | A0A2B4RM65 |
| Stylophora pi | A0A2B4SMY1 |
| Stylophora pi | A0A2B4RXN2 |
| Stylophora pi | A0A2B4SDQ3 |
| Stylophora pi | A0A2B4S5V1 |
| Sulfitobacter | A0A1I1VVN6 |
| Sulfitobacter | A0A073ILQ0 |
| Sulfitobacter | A0A196QUR2 |
| Sulfitobacter | A0A420DSB1 |
| Sulfitobacter | A0A061SV07 |
| Sulfitobacter | A0A2T6CEN1 |
| Sulfitobacter | A0A221K513 |
| Sulfitobacter | A0A1J0WMJ3 |
| Sulfitobacter | A0A196P088 |
| Sulfitobacter | A0A345Q6D2 |
| Sus scrofa.xls | P06867 |
| Sus scrofa.xls | F1SPS6 |
| Sus scrofa.xls | F1SB93 |
| Sus scrofa.xls | F1RZN7 |
| Sus scrofa.xls | A0A286ZVK1 |
| Sus scrofa.xls | O97506 |
| Sus scrofa.xls | A0A286ZR66 |
| Sus scrofa.xls | F1SB81 |
| Symbiodiniur | A0A1Q9ESG7 |
| Symbiodiniur | A0A1Q9CQI2 |
| Symbiodiniur | A0A1Q9C6C8 |
| Symbiodiniur | A0A1Q9CCQ0 |
| Symbiodiniur | A0A1Q9EEK4 |
| Symbiodiniur | A0A1Q9D1E5 |
| Symbiodiniur | A0A1Q9DKU8 |
| Symbiodiniur | A0A1Q9DRV2 |

| | |
|---|---|
| Symbiodiniur | A0A1Q9F029 |
| Symbiodiniur | A0A1Q9ESU8 |
| Symbiodiniur | A0A1Q9EQ68 |
| Symbiodiniur | A0A1Q9D2R1 |
| Symbiodiniur | A0A1Q9E4J1 |
| Symbiodiniur | A0A1Q9CW84 |
| Symbiodiniur | A0A1Q9CMK9 |
| Symbiodiniur | A0A1Q9CF00 |
| Symbiodiniur | A0A1Q9C5U3 |
| Symbiodiniur | A0A1Q9E8L2 |
| Symbiodiniur | A0A1Q9D4N5 |
| Symbiodiniur | A0A1Q9EKZ9 |
| Symbiodiniur | A0A1Q9C5H9 |
| Symbiodiniur | A0A1Q9E9H4 |
| Symbiodiniur | A0A1Q9C600 |
| Symbiodiniur | A0A1Q9CGB9 |
| Symbiodiniur | A0A1Q9BXP7 |
| Symbiodiniur | A0A1Q9CQY3 |
| Symbiodiniur | A0A1Q9CR00 |
| Symbiodiniur | A0A1Q9CAA1 |
| Symbiodiniur | A0A1Q9EMW5 |
| Symbiodiniur | A0A1Q9DB15 |
| Symbiodiniur | A0A1Q9CW09 |
| Symbiodiniur | A0A1Q9CR59 |
| Symbiodiniur | A0A1Q9ED26 |
| Symbiodiniur | A0A1Q9ERZ2 |
| Symbiodiniur | A0A1Q9CQZ9 |
| Symbiodiniur | A0A1Q9CZK1 |
| Symbiodiniur | A0A1Q9DNS4 |
| Symbiodiniur | A0A1Q9F1U4 |
| Symbiodiniur | A0A1Q9CT81 |
| Symbiodiniur | A0A1Q9D3Z6 |
| Symbiodiniur | A0A1Q9CQ66 |
| Symbiodiniur | A0A1Q9CSN6 |
| Symbiodiniur | A0A1Q9ERZ8 |
| Symbiodiniur | A0A1Q9D1P7 |
| Symbiodiniur | A0A1Q9ES05 |
| Symbiodiniur | A0A1Q9CUS2 |
| Symbiodiniur | A0A1Q9DZF4 |
| Symbiodiniur | A0A1Q9CAP7 |
| Symbiodiniur | A0A1Q9DAV4 |

| | |
|---|---|
| Symbiodiniur | A0A1Q9DVU6 |
| Symbiodiniur | A0A1Q9BZ14 |
| Symbiodiniur | A0A1Q9DA38 |
| Symbiodiniur | A0A1Q9EUE8 |
| Symbiodiniur | A0A1Q9DT95 |
| Symbiodiniur | A0A1Q9EUA8 |
| Symbiodiniur | A0A1Q9DLH6 |
| Symbiodiniur | A0A1Q9D262 |
| Symbiodiniur | A0A1Q9D697 |
| Symbiodiniur | A0A1Q9DPD5 |
| Symbiodiniur | A0A1Q9EBY1 |
| Symbiodiniur | A0A1Q9D3F0 |
| Symbiodiniur | A0A1Q9CMG7 |
| Symbiodiniur | A0A1Q9EUA6 |
| Symbiodiniur | A0A1Q9F599 |
| Synechococci | A0A1Z9KW32 |
| Synechococci | A0A1Z9L6H2 |
| Syntrophoba | A0A2M7BBY6 |
| Syntrophus sp | A0A1V4Y3U4 |
| Syphacia mur | A0A0N5AQ74 |
| Syphacia mur | A0A0N5AVN4 |
| Syphacia mur | A0A0N5AWJ9 |
| Syphacia mur | A0A0N5AKP3 |
| Syphacia mur | A0A0N5ASE0 |
| Syphacia mur | A0A0N5AQE8 |
| Syphacia mur | A0A0N5AV92 |
| Syphacia mur | A0A0N5ABP9 |
| Syphacia mur | A0A0N5AMT9 |
| Syphacia mur | A0A158R5G1 |
| Syphacia mur | A0A0N5AYU3 |
| Syphacia mur | A0A0N5AT40 |
| Syphacia mur | A0A0N5A8Y8 |
| Syphacia mur | A0A0N5AUH4 |
| Syphacia mur | A0A158R6C4 |
| Syphacia mur | A0A158R5U4 |
| Syphacia mur | A0A0N5AX21 |
| Syphacia mur | A0A0N5AAC3 |
| Syphacia mur | A0A0N5AES2 |
| Syphacia mur | A0A0N5AR84 |
| Syphacia mur | A0A0N5AWX5 |
| Tabrizicola sp | A0A411Z4S3 |

| | |
|---|---|
| Tabrizicola sp | A0A3S8U2P2 |
| Tabrizicola sp | A0A2N4YG23 |
| Taenia asiatic | A0A0R3VTQ5 |
| Taenia asiatic | A0A3P6NAE9 |
| Taenia asiatic | A0A158R765 |
| Taeniopygia g | H0ZG36 |
| Taeniopygia g | H0YWJ9 |
| Taeniopygia g | H0ZY21 |
| Taeniopygia g | H0ZML2 |
| Taeniopygia g | H0Z9B7 |
| Taeniopygia g | H0Z415 |
| Takifugu rubr | A0A3B5KNQ7 |
| Takifugu rubr | A0A0S3P5S3 |
| Takifugu rubr | A0A3B5JVN8 |
| Takifugu rubr | H2U253 |
| Takifugu rubr | A0A3B5K7U1 |
| Takifugu rubr | H2UIX2 |
| Takifugu rubr | A0A3B5K478 |
| Takifugu rubr | H2U252 |
| Takifugu rubr | A0A3B5K8C0 |
| Takifugu rubr | H2UB48 |
| Takifugu rubr | A0A3B5KSI5 |
| Takifugu rubr | H2UB51 |
| Talaromyces | A0A364L110 |
| Talaromyces | A0A1Q5QBL9 |
| Talaromyces | A0A0U1LQB1 |
| Talaromyces | B6QAW5 |
| Talaromyces | B6Q7P1 |
| Talaromyces | B6QVR0 |
| Talaromyces | B8M4W0 |
| Talaromyces | B8MRL4 |
| Taphrina defo | R4X8Q8 |
| Tardiphaga ro | A0A163ZG59 |
| Tardiphaga sp | A0A370QWB2 |
| Tardiphaga sp | A0A1H6PBW2 |
| Tardiphaga sp | A0A239NTK9 |
| Tardiphaga sp | A0A2U0TH97 |
| Tardiphaga sp | A0A420ATD9 |
| Tarsius syrich | A0A3Q0DMW9 |
| Tarsius syrich | A0A1U7SYT3 |
| Tarsius syrich | A0A1U7U5V8 |

| | |
|---|---|
| Tarsius syrich | A0A1U7U7M3 |
| Tarsius syrich | A0A1U7TY18 |
| Tarsius syrich | A0A3Q0DKG9 |
| Tarsius syrich | A0A1U7SMG8 |
| Tarsius syrich | A0A1U7UC10 |
| Tarsius syrich | A0A3Q0DG51 |
| Tarsius syrich | A0A3Q0DM20 |
| Tateyamaria | A0A1P8MZ84 |
| Tateyamaria | A0A0B4C503 |
| Tauraco erytl | A0A093CH68 |
| Tauraco erytl | A0A093CND7 |
| Tauraco erytl | A0A093CVC3 |
| Tauraco erytl | A0A093CQR8 |
| Teladorsagia | A0A2G9TXP9 |
| Teladorsagia | A0A2G9U3X5 |
| Teladorsagia | A0A2G9UGS1 |
| Teladorsagia | A0A2G9UVF5 |
| Teladorsagia | A0A2G9UD74 |
| Teladorsagia | A0A2G9TB90 |
| Teladorsagia | A0A2G9UAM5 |
| Teladorsagia | A0A2G9T9Y6 |
| Teladorsagia | A0A2G9U041 |
| Teladorsagia | A0A2G9UK14 |
| Teladorsagia | A0A2G9U3J8 |
| Teladorsagia | A0A2G9U5G6 |
| Teladorsagia | A0A2G9U6U5 |
| Teladorsagia | A0A2G9UCE1 |
| Teladorsagia | A0A2G9V4H0 |
| Teladorsagia | A0A2G9UF90 |
| Teladorsagia | A0A2G9U0T0 |
| Teladorsagia | A0A2G9UAE0 |
| Teladorsagia | A0A2G9U9D4 |
| Teladorsagia | A0A2G9UTY8 |
| Teladorsagia | A0A2G9UH07 |
| Tenacibaculu | A0A3C1S3E6 |
| Tenacibaculu | A0A3C1S4B8 |
| Tepidicaulis r | A0A081BAR4 |
| Teredinibacte | C5BI67 |
| Terrestrivirus | A0A3G4ZLM1 |
| Tetrabaena s | A0A2J7ZPE6 |
| Tetrabaena s | A0A2J7ZJB7 |

| | |
|---|---|
| Tetrabaena s | A0A2J8AH70 |
| Tetrabaena s | A0A2J8AIZ8 |
| Tetrabaena s | A0A2J8A820 |
| Tetrabaena s | A0A2J8A4H3 |
| Tetrabaena s | A0A2J8AFD0 |
| Tetrabaena s | A0A2J7ZQC3 |
| Tetrabaena s | A0A2J8AFF5 |
| Tetrabaena s | A0A2J7ZX95 |
| Tetrabaena s | A0A2J8A7Q4 |
| Tetrabaena s | A0A2J7ZFF2 |
| Tetradesmus | A0A383W609 |
| Tetradesmus | A0A383W2V6 |
| Tetradesmus | A0A383VA02 |
| Tetradesmus | A0A383VTL8 |
| Tetradesmus | A0A383VEC8 |
| Tetradesmus | A0A383W0C6 |
| Tetradesmus | A0A383VRV3 |
| Tetradesmus | A0A383VVE2 |
| Tetradesmus | A0A383VV90 |
| Tetradesmus | A0A383WH61 |
| Tetradesmus | A0A383VIT8 |
| Tetradesmus | A0A383VX83 |
| Tetradesmus | A0A383V476 |
| Tetradesmus | A0A383WKH6 |
| Tetradesmus | A0A383VAP4 |
| Tetradesmus | A0A383VSI6 |
| Tetrahymena | Q236L9 |
| Tetrahymena | Q2I2L4 |
| Tetranychus t | A0A3G5ANY0 |
| Tetranychus u | T1L2L9 |
| Tetranychus u | T1JTQ3 |
| Tetranychus u | T1JZ41 |
| Tetranychus u | T1JZ38 |
| Tetranychus u | T1KP84 |
| Tetranychus u | T1L6E3 |
| Tetranychus u | T1KXB8 |
| Tetranychus u | T1KLX8 |
| Tetranychus u | T1K1X0 |
| Tetranychus u | T1KJV3 |
| Tetranychus u | T1JQK0 |
| Tetranychus u | T1K882 |

| | |
|---|---|
| Tetranychus ( | T1KJA7 |
| Tetranychus ( | T1L2F0 |
| Tetranychus ( | T1JY66 |
| Tetranychus ( | T1KEM9 |
| Tetranychus ( | T1L0H5 |
| Tetranychus ( | T1KY75 |
| Tetranychus ( | T1KVC5 |
| Tetranychus ( | T1KP75 |
| Tetranychus ( | T1L6B7 |
| Tetranychus ( | T1L4K2 |
| Tetranychus ( | T1L0U8 |
| Tetranychus ( | T1K0C2 |
| Tetranychus ( | T1JY60 |
| Tetranychus ( | T1KXD7 |
| Tetranychus ( | T1K0A5 |
| Tetranychus ( | T1KP31 |
| Tetranychus ( | T1KNH0 |
| Tetranychus ( | T1L2M1 |
| Tetranychus ( | T1KVC7 |
| Tetranychus ( | T1K0C1 |
| Tetranychus ( | A0A158P574 |
| Tetranychus ( | T1L0H6 |
| Tetranychus ( | T1K0C0 |
| Tetranychus ( | T1K6H1 |
| Tetranychus ( | T1K0A4 |
| Tetranychus ( | T1L1K0 |
| Tetranychus ( | T1JXS1 |
| Tetranychus ( | T1K414 |
| Tetranychus ( | A0A158P575 |
| Tetranychus ( | T1KVB7 |
| Tetranychus ( | T1K276 |
| Tetranychus ( | T1K0A1 |
| Tetranychus ( | T1KW32 |
| Tetranychus ( | T1L1J9 |
| Tetranychus ( | T1KP56 |
| Tetranychus ( | T1L4K1 |
| Tetranychus ( | T1KP60 |
| Tetranychus ( | T1K0A3 |
| Tetranychus ( | T1K411 |
| Tetranychus ( | T1L2E5 |
| Tetranychus ( | T1KVC8 |

| | |
|---|---|
| Tetranychus ( | T1K0A9 |
| Tetranychus ( | T1KF99 |
| Tetranychus ( | T1KP59 |
| Tetranychus ( | T1JXR7 |
| Tetranychus ( | T1JXS5 |
| Tetranychus ( | T1K0A7 |
| Tetranychus ( | T1K0E3 |
| Tetranychus ( | T1JV21 |
| Tetranychus ( | T1KP64 |
| Tetranychus ( | T1K7E7 |
| Tetraodon ni | Q4RF08 |
| Tetraodon ni | H3CM48 |
| Tetraodon ni | Q4RX91 |
| Tetraodon ni | H3CWI4 |
| Tetraodon ni | H3DBA6 |
| Tetraodon ni | H3D0B7 |
| Tetraodon ni | Q4RF09 |
| Tetraodon ni | Q4SIS6 |
| Tetraodon ni | Q4SBT4 |
| Tetraodon ni | H3C6P0 |
| Tetraodon ni | H3DPE4 |
| Tetraodon ni | H3CV10 |
| Tetraodon ni | H3CWH8 |
| Tetraselmis s | A0A061RM82 |
| Tetraselmis s | A0A061QU17 |
| Tetraselmis s | A0A061SFA1 |
| Tetraselmis s | A0A061S0K3 |
| Tetraselmis s | A0A061R340 |
| Tetraselmis s | A0A061S6B9 |
| Tetraselmis s | A0A061RKH4 |
| Tetraselmis s | A0A061SLP2 |
| Tetraselmis s | A0A061QSN4 |
| Tetraselmis s | A0A061S9I3 |
| Tetraselmis s | A0A061R7K6 |
| Tetraselmis s | A0A061QSD4 |
| Tetraselmis s | A0A061R2B1 |
| Tetraselmis s | A0A061R0X8 |
| Tetraselmis s | A0A061QSK8 |
| Tetraselmis s | A0A061SA98 |
| Tetraselmis v | A0A2P0VNG3 |
| Tetraselmis v | A0A2P0VNJ5 |

| | |
|---|---|
| Thalassiosira | K0R003 |
| Thalassiosira | K0T569 |
| Thalassiosira | K0R8M5 |
| Thalassiosira | K0R4V6 |
| Thalassiosira | K0TI40 |
| Thalassiosira | K0S3W9 |
| Thalassiosira | K0R053 |
| Thalassiosira | K0RJ23 |
| Thalassiosira | K0R2N3 |
| Thalassobius | A0A0P1FYC3 |
| Thalassobius | A0A0P1FXN8 |
| Thalassococc | A0A369TVA4 |
| Thalassococc | A0A369TQK7 |
| Thalassomon | A0A0D8D6E0 |
| Theileria orie | J4C7F0 |
| Theileria orie | A0A2T7I917 |
| Theileria orie | A0A2T7IN49 |
| Thelazia calli | A0A0N5D4A3 |
| Thelazia calli | A0A0N5DAH6 |
| Thelazia calli | A0A0N5D0V9 |
| Thelazia calli | A0A0N5CNH4 |
| Thelazia calli | A0A0N5D8L1 |
| Thelazia calli | A0A0N5CN06 |
| Thelazia calli | A0A0N5D5Z9 |
| Thelazia calli | A0A158RB75 |
| Thelazia calli | A0A158RB18 |
| Thelazia calli | A0A158RCJ6 |
| Thelazia calli | A0A0N5CL57 |
| Thelazia calli | A0A0N5CMM8 |
| Thelazia calli | A0A0N5D941 |
| Thelazia calli | A0A0N5CU49 |
| Thelazia calli | A0A0N5CLL0 |
| Thelazia calli | A0A0N5D6J3 |
| Thelazia calli | A0A0N5CU48 |
| Thelazia calli | A0A0N5D3M6 |
| Thelazia calli | A0A0N5D0H3 |
| Thelazia calli | A0A0N5CU50 |
| Thelohanellu | A0A0C2N594 |
| Theobroma c | A0A061FES3 |
| Theobroma c | A0A061F6M2 |
| Theobroma c | A0A061F6Q9 |

Theobroma c A0A061F858
Theobroma c A0A061F7U2
Theobroma c A0A061E4J3
Theobroma c A0A061F7S0
Theobroma c A0A061DSY4
Theobroma c A0A061F7R6
Theobroma c A0A061F5X2
Theobroma c A0A061F5W4
Theobroma c A0A061DK79
Theobroma c A0A061GFZ4
Theobroma c A0A061FE28
Theobroma c A0A061FDY8
Theobroma c A0A061GMY9
Theobroma c A0A061DT57
Theobroma c A0A061FE35
Theobroma c A0A061F6S4
Theobroma c A0A061ESM6
Theobroma c A0A061F7P5
Theobroma c A0A061FE32
Theobroma c A0A061GN00
Theobroma c A0A061DJB7
Theobroma c A0A061F829
Theobroma c A0A061FAS2
Theobroma c A0A061GJY1
Theobroma c A0A061GJ76
Theobroma c A0A061E2S8
Theobroma c A0A061FXH6
Theobroma c A0A061FQL2
Theobroma c A0A061GGE3
Theobroma c A0A061GSK0
Theobroma c A0A061F5V9
Theobroma c A0A061GGC0
Theobroma c A0A061F6T1
Theobroma c A0A061FEW2
Theobroma c A0A061GMZ5
Theobroma c A0A061FA86
Theobroma c A0A061E2J3
Theobroma c A0A061GF26
Theobroma c A0A061F5X6
Theobroma c A0A061F6R7
Theobroma c A0A061FII3

Theobroma c A0A061GA93
Theobroma c A0A061GGC6
Theobroma c A0A061FBL4
Theobroma c A0A061ETG0
Theobroma c A0A061F7N9
Theobroma c A0A061EAF9
Theobroma c A0A061GG11
Theobroma c A0A061F862
Theobroma c A0A061GF19
Theobroma c A0A061F9V0
Theobroma c A0A061F5Y4
Theobroma c A0A061GS10
Theobroma c A0A061F798
Theobroma c A0A061DLF5
Theobroma c A0A061GH37
Theobroma c A0A061FEJ0
Theobroma c A0A061F7L9
Theobroma c A0A061ERV1
Theobroma c A0A061GGD8
Theobroma c A0A061GFZ9
Theobroma c A0A061F6A9
Theobroma c A0A061F7T9
Theobroma c A0A061F463
Theobroma c S1RWJ4
Theobroma c A0A061F7A3
Theobroma c A0A061E4H9
Theobroma c A0A061F772
Theobroma c A0A061F7R1
Theobroma c A0A061F7T8
Theobroma c A0A061E2S4
Theobroma c A0A061F771
Theobroma c A0A061FDW5
Theobroma c A0A061FE49
Theobroma c A0A061E2U3
Theobroma c A0A061F6S1
Theobroma c A0A061F7S7
Theobroma c A0A061E4H6
Theobroma c A0A061E3P7
Theobroma c A0A061F784
Theobroma c A0A061F5W9
Theobroma c A0A061F6J2

| | |
|---|---|
| Theobroma c | A0A061FFC7 |
| Theobroma c | A0A061E9M0 |
| Theobroma c | A0A061GG05 |
| Theobroma c | A0A061F6R3 |
| Theobroma c | A0A061GF31 |
| Theobroma c | A0A061EQQ6 |
| Theobroma c | A0A061E3Q5 |
| Theobroma c | A0A061FDY1 |
| Theobroma c | A0A061FGI5 |
| Theobroma c | A0A061GH52 |
| Theobroma c | A0A061G8B1 |
| Theobroma c | A0A061FE26 |
| Theobroma c | A0A061EZN1 |
| Theobroma c | A0A061ERB8 |
| Theobroma c | A0A061EAF4 |
| Theobroma c | A0A061E3Q2 |
| Theobroma c | A0A061GN44 |
| Theobroma c | A0A061ETH9 |
| Theobroma c | A0A061GGD2 |
| Theobroma c | A0A061FC33 |
| Theobroma c | A0A061GBN6 |
| Thermodesul | A0A177E7S8 |
| Thermosulfid | A0A0S3QUQ3 |
| Thermosulfu | A0A179D652 |
| Thielavia terr | G2QZQ3 |
| Thielavia terr | A0A3S4ATD0 |
| Thiocapsa ma | F9UF48 |
| Thiocapsa ros | A0A1H2T5K1 |
| Thioclava ind | A0A074JZA8 |
| Thiocystis vio | I3YA25 |
| Thioflavicocc | L0H1B6 |
| Thiorhodococ | G2DZW1 |
| Thiotrichales | A0A1H6F9F4 |
| Thraustothec | A0A0A7CLS4 |
| Thraustothec | A0A0A7CM27 |
| Thraustothec | A0A0A7CMA6 |
| Thraustothec | A0A1V9ZCI3 |
| Thraustothec | A0A0A7CLY0 |
| Thraustothec | A0A1V9Y7R8 |
| Thraustothec | A0A1V9ZQ59 |
| Thraustothec | A0A1V9ZX23 |

| | |
|---|---|
| Thraustothec | A0A0A7CM73 |
| Thraustothec | A0A1V9ZWR3 |
| Thraustothec | A0A0A7CLP3 |
| Thraustothec | A0A1V9ZWD5 |
| Thraustothec | A0A1W0A8G8 |
| Thraustothec | A0A1V9Z6L2 |
| Thraustothec | A0A0A7CMR6 |
| Thraustothec | A0A1W0A9L7 |
| Thraustothec | A0A1V9Y3Z2 |
| Thraustothec | A0A0A7CMJ9 |
| Thraustothec | A0A0A7CLZ4 |
| Thraustothec | A0A0A7CLK9 |
| Thraustothec | A0A1V9Y7V8 |
| Thraustothec | A0A0A7CLD0 |
| Thraustothec | A0A0A7CM37 |
| Thraustothec | A0A1V9YW96 |
| Thraustothec | A0A0A7CMD3 |
| Thraustothec | A0A1V9ZV18 |
| Thraustothec | A0A1V9ZQU5 |
| Thraustothec | A0A1W0A6N9 |
| Thraustothec | A0A0A7CM15 |
| Thraustothec | A0A1V9ZX55 |
| Thraustothec | A0A1V9YS19 |
| Thraustothec | A0A1W0A3D9 |
| Thraustothec | A0A1W0A1E0 |
| Thraustothec | A0A0A7CLV9 |
| Thraustothec | A0A1V9YHU4 |
| Thraustothec | A0A1V9Y7T4 |
| Thraustothec | A0A1V9YS21 |
| Thraustothec | A0A1W0A6T9 |
| Thraustothec | A0A1W0A2R4 |
| Thraustothec | A0A1V9ZRW9 |
| Thraustothec | A0A0A7CMA3 |
| Thraustothec | A0A0A7CM84 |
| Thraustothec | A0A0A7CM21 |
| Thraustothec | A0A0A7CM83 |
| Tilletia caries | A0A177U9L0 |
| Tilletia contro | A0A177VNW5 |
| Tilletiaria anc | A0A066V6E8 |
| Tinamus gutt | A0A099ZKM4 |
| Tinamus gutt | A0A099YWG8 |

| | |
|---|---|
| Tinamus gutt | A0A099ZHV8 |
| Tinamus gutt | A0A099ZGB5 |
| Togninia mini | R8BMT1 |
| Togninia mini | R8BT17 |
| Tolypocladiur | A0A2S4KXQ7 |
| Tolypocladiur | A0A2S4KU31 |
| Torrubiella he | A0A0A1T7V7 |
| Toxocara can | A0A183V6M5 |
| Toxocara can | A0A183UMU0 |
| Toxocara can | A0A183V0Y5 |
| Toxocara can | A0A183VH40 |
| Toxocara can | A0A183URQ0 |
| Toxocara can | A0A183UCS5 |
| Toxocara can | A0A183UGS8 |
| Toxocara can | A0A183VGT9 |
| Toxocara can | A0A183UGB4 |
| Toxocara can | A0A183UJW2 |
| Toxocara can | A0A183VFR8 |
| Toxocara can | A0A183V6I3 |
| Toxocara can | A0A183V8F0 |
| Toxocara can | A0A183UKJ7 |
| Toxocara can | A0A183UZI3 |
| Toxocara can | A0A183UCS4 |
| Toxocara can | A0A183TZ03 |
| Toxocara can | A0A183UXD4 |
| Toxocara can | A0A183UPE8 |
| Toxocara can | A0A183VHD0 |
| Toxocara can | A0A183V3X8 |
| Toxocara can | A0A183UN76 |
| Toxocara can | A0A183V0R7 |
| Toxocara can | A0A183UZ80 |
| Toxocara can | A0A0B2V222 |
| Toxocara can | A0A0B2W1F4 |
| Toxocara can | A0A0B2VYQ4 |
| Toxocara can | A0A0B2VKQ9 |
| Toxocara can | A0A0B2VKX7 |
| Toxocara can | A0A0B2VCC1 |
| Toxocara can | A0A0B2VBP0 |
| Toxocara can | A0A0B2VDF6 |
| Toxocara can | A0A0B2VTH8 |
| Toxocara can | A0A0B2VE70 |

| | |
|---|---|
| Toxocara can | A0A183V0Y4 |
| Toxocara can | A0A183TW29 |
| Toxocara can | A0A3P7IEE1 |
| Toxocara can | A0A0B2V8X8 |
| Toxocara can | A0A0B2V3M1 |
| Toxocara can | A0A183U4G9 |
| Toxocara can | A0A3P7EW77 |
| Toxocara can | A0A0B2VCY3 |
| Toxocara can | A0A0B2VUZ8 |
| Toxocara can | A0A0B2VRS5 |
| Toxocara can | A0A0B2VS27 |
| Toxocara can | A0A0B2V756 |
| Toxocara can | A0A0B2V7G5 |
| Toxocara can | A0A0B2VLY9 |
| Toxocara can | A0A0B2V1Y2 |
| Toxocara can | A0A0B2UVF7 |
| Toxocara can | A0A0B2UUB7 |
| Toxocara can | A0A3P7F9F7 |
| Toxocara can | A0A0B2VJJ4 |
| Toxocara can | A0A183UFM8 |
| Toxocara can | A0A0B2UVS9 |
| Toxocara can | A0A0B2W5N5 |
| Toxoplasma g | A0A125YNB1 |
| Toxoplasma g | S8EWI3 |
| Toxoplasma g | A0A125YP11 |
| Toxoplasma g | S8EXK9 |
| Toxoplasma g | S8FBN1 |
| Toxoplasma g | S8GJR8 |
| Toxoplasma g | S8GTW7 |
| Toxoplasma g | S8GU33 |
| Toxoplasma g | A0A125YN33 |
| Toxoplasma g | S8GE96 |
| Toxoplasma g | S8EWJ0 |
| Toxoplasma g | S8G8W7 |
| Toxoplasma g | S8EUX1 |
| Toxoplasma g | S8F2N4 |
| Toxoplasma g | S8EVK9 |
| Toxoplasma g | S8GIC9 |
| Toxoplasma g | S8FBK4 |
| Toxoplasma g | S8F0R5 |
| Toxoplasma g | S8EZX1 |

| | |
|---|---|
| Toxoplasma g | S7UGM9 |
| Toxoplasma g | S7UVS3 |
| Toxoplasma g | S7W921 |
| Toxoplasma g | S7VS18 |
| Toxoplasma g | S7ULA3 |
| Toxoplasma g | S7W1D2 |
| Toxoplasma g | A0A125YUY4 |
| Toxoplasma g | S7VZR7 |
| Toxoplasma g | S7V4Y3 |
| Toxoplasma g | S7VNS6 |
| Toxoplasma g | S7VUK4 |
| Toxoplasma g | S7UNU6 |
| Toxoplasma g | S7URJ3 |
| Toxoplasma g | S7WIY4 |
| Toxoplasma g | S7UL27 |
| Toxoplasma g | S7UID9 |
| Toxoplasma g | S7UMV2 |
| Toxoplasma g | S7UFC4 |
| Toxoplasma g | A0A125YP10 |
| Toxoplasma g | S7V4X6 |
| Toxoplasma g | A0A125YMJ2 |
| Toxoplasma g | B6KPA5 |
| Toxoplasma g | B9QJ65 |
| Toxoplasma g | V5BN10 |
| Toxoplasma g | B9PXJ3 |
| Toxoplasma g | A0A0F7UY76 |
| Toxoplasma g | A0A0F7V6F0 |
| Toxoplasma g | A0A0F7UZC2 |
| Toxoplasma g | V4ZN28 |
| Toxoplasma g | V5B2C7 |
| Toxoplasma g | B9QHA2 |
| Toxoplasma g | A0A125YYV2 |
| Toxoplasma g | V4ZE44 |
| Toxoplasma g | V4ZEU8 |
| Toxoplasma g | V4Z8E3 |
| Toxoplasma g | V4Z683 |
| Toxoplasma g | A0A0F7UY36 |
| Toxoplasma g | A0A0F7VB90 |
| Toxoplasma g | A0A0F7V4E0 |
| Toxoplasma g | V4YRM6 |
| Toxoplasma g | B9QHI1 |

| | |
|---|---|
| Toxoplasma g | A0A0F7USH6 |
| Toxoplasma g | A0A0F7UP20 |
| Toxoplasma g | A0A0F7UTL5 |
| Toxoplasma g | A0A0F7V2Y2 |
| Toxoplasma g | A0A125YH93 |
| Toxoplasma g | V4ZAX9 |
| Toxoplasma g | V4ZEU3 |
| Toxoplasma g | V4ZQP4 |
| Toxoplasma g | A0A0F7V7R9 |
| Toxoplasma g | V4Z2B5 |
| Toxoplasma g | A0A0F7V2N5 |
| Toxoplasma g | V4ZSH6 |
| Toxoplasma g | A0A139Y8B9 |
| Toxoplasma g | A0A139XR75 |
| Toxoplasma g | A0A139XT94 |
| Toxoplasma g | A0A139XR22 |
| Toxoplasma g | A0A139XUT5 |
| Toxoplasma g | A0A139XV46 |
| Toxoplasma g | A0A139Y523 |
| Toxoplasma g | A0A139XII1 |
| Toxoplasma g | A0A139XUW2 |
| Toxoplasma g | A0A139XRH6 |
| Toxoplasma g | A0A139XI69 |
| Toxoplasma g | A0A139XIG5 |
| Toxoplasma g | A0A139XQZ9 |
| Toxoplasma g | A0A139XIW7 |
| Toxoplasma g | A0A139XYI0 |
| Toxoplasma g | A0A139XIH6 |
| Toxoplasma g | A0A139YAJ0 |
| Toxoplasma g | A0A139XI83 |
| Toxoplasma g | A0A139XUB1 |
| Toxoplasma g | A0A139XIA3 |
| Toxoplasma g | A0A425HTR7 |
| Toxoplasma g | A0A425HLG8 |
| Toxoplasma g | A0A3R8FZK1 |
| Toxoplasma g | A0A3R7YPK8 |
| Toxoplasma g | A0A3R8BM20 |
| Toxoplasma g | A0A3R8AJM0 |
| Toxoplasma g | A0A425HTJ6 |
| Toxoplasma g | A0A3R7Z551 |
| Toxoplasma g | A0A3R7ZDF1 |

| | |
|---|---|
| Toxoplasma g | A0A3R7ZER8 |
| Toxoplasma g | A0A3R8GAD6 |
| Toxoplasma g | A0A425HLE6 |
| Toxoplasma g | A0A425HRD9 |
| Toxoplasma g | A0A425HTP5 |
| Toxoplasma g | A0A425HNT2 |
| Toxoplasma g | A0A425I1A5 |
| Toxoplasma g | A0A425HTB8 |
| Toxoplasma g | A0A425HNI2 |
| Toxoplasma g | A0A3R8AFY2 |
| Toxoplasma g | A0A2G8XSQ7 |
| Toxoplasma g | A0A2G8XUD9 |
| Toxoplasma g | A0A2G8Y8M6 |
| Toxoplasma g | A0A2G8XUK9 |
| Toxoplasma g | A0A2G8XV07 |
| Toxoplasma g | A0A2G8XU33 |
| Toxoplasma g | A0A2G8XVA9 |
| Toxoplasma g | A0A2G8XLM8 |
| Toxoplasma g | A0A2G8XM15 |
| Toxoplasma g | A0A2G8Y7C4 |
| Toxoplasma g | A0A2G8XYW5 |
| Toxoplasma g | A0A2G8XU35 |
| Toxoplasma g | A0A2G8YB46 |
| Toxoplasma g | A0A2G8XVB5 |
| Toxoplasma g | A0A2G8XLP9 |
| Toxoplasma g | A0A2G8XM02 |
| Toxoplasma g | A0A2G8XWV2 |
| Toxoplasma g | A0A086KZL5 |
| Toxoplasma g | A0A086L8S1 |
| Toxoplasma g | A0A086KXX3 |
| Toxoplasma g | A0A086JKK2 |
| Toxoplasma g | A0A086LDQ0 |
| Toxoplasma g | A0A086JHJ8 |
| Toxoplasma g | A0A086K039 |
| Toxoplasma g | A0A086K045 |
| Toxoplasma g | A0A086JSN5 |
| Toxoplasma g | A0A086LF74 |
| Toxoplasma g | A0A086KUA3 |
| Toxoplasma g | A0A086LGR4 |
| Toxoplasma g | A0A086KUA2 |
| Toxoplasma g | A0A086LC13 |

| | |
|---|---|
| Toxoplasma g | A0A086KT20 |
| Toxoplasma g | A0A086KZ32 |
| Toxoplasma g | A0A086KBL4 |
| Toxoplasma g | A0A086KJB6 |
| Toxoplasma g | A0A086KPR8 |
| Toxoplasma g | A0A086JRW4 |
| Toxoplasma g | A0A086JAN6 |
| Toxoplasma g | A0A086JTG7 |
| Toxoplasma g | A0A086JAS2 |
| Toxoplasma g | A0A086JRJ6 |
| Toxoplasma g | A0A086KBL3 |
| Toxoplasma g | A0A086KAV4 |
| Toxoplasma g | A0A086J9Y0 |
| Toxoplasma g | A0A086K6N1 |
| Toxoplasma g | A0A086JBU3 |
| Toxoplasma g | A0A086KWD4 |
| Toxoplasma g | A0A086JSD4 |
| Toxoplasma g | A0A086KNW2 |
| Toxoplasma g | A0A086JH56 |
| Toxoplasma g | A0A086QPX5 |
| Toxoplasma g | A0A086QPX6 |
| Toxoplasma g | A0A086QS28 |
| Toxoplasma g | A0A086QR16 |
| Toxoplasma g | A0A086QSB4 |
| Toxoplasma g | A0A086QZQ0 |
| Toxoplasma g | A0A086QXJ7 |
| Toxoplasma g | A0A086PMI4 |
| Toxoplasma g | A0A086PJV3 |
| Toxoplasma g | A0A086QPJ5 |
| Toxoplasma g | A0A086Q7X9 |
| Toxoplasma g | A0A086QSK0 |
| Toxoplasma g | A0A086QDJ1 |
| Toxoplasma g | A0A086Q5C2 |
| Toxoplasma g | A0A086PJS5 |
| Toxoplasma g | A0A086PJV4 |
| Toxoplasma g | A0A086PP13 |
| Toxoplasma g | A0A086JYT0 |
| Toxoplasma g | A0A086JYV7 |
| Toxoplasma g | A0A086KQT7 |
| Toxoplasma g | A0A086JUX0 |
| Toxoplasma g | A0A086JEK0 |

| | |
|---|---|
| Toxoplasma g | A0A086K9B0 |
| Toxoplasma g | A0A086J6M2 |
| Toxoplasma g | A0A086L3H4 |
| Toxoplasma g | A0A086JYV2 |
| Toxoplasma g | A0A086J6A5 |
| Toxoplasma g | A0A086JAT8 |
| Toxoplasma g | A0A086JW34 |
| Toxoplasma g | A0A086KKD3 |
| Toxoplasma g | A0A086J6E7 |
| Toxoplasma g | A0A086KW68 |
| Toxoplasma g | A0A086JUW9 |
| Toxoplasma g | A0A086JYW7 |
| Toxoplasma g | A0A086LN03 |
| Toxoplasma g | A0A086LVF2 |
| Toxoplasma g | A0A086LIM6 |
| Toxoplasma g | A0A086LJ27 |
| Toxoplasma g | A0A086LIY4 |
| Toxoplasma g | A0A086LWU1 |
| Toxoplasma g | A0A086MAA6 |
| Toxoplasma g | A0A086LX43 |
| Toxoplasma g | A0A086LX44 |
| Toxoplasma g | A0A086LTJ4 |
| Toxoplasma g | A0A086M0U2 |
| Toxoplasma g | A0A086M4H2 |
| Toxoplasma g | A0A086LIS3 |
| Toxoplasma g | A0A086M306 |
| Toxoplasma g | A0A086LUX9 |
| Toxoplasma g | A0A086LV63 |
| Toxoplasma g | A0A086M612 |
| Toxoplasma g | A0A086M0U7 |
| Toxoplasma g | A0A086LJF7 |
| Toxoplasma g | A0A086LJF6 |
| Toxoplasma g | A0A2T6IZJ0 |
| Toxoplasma g | A0A2T6II53 |
| Toxoplasma g | A0A2T6IQ36 |
| Toxoplasma g | A0A2T6IQ28 |
| Toxoplasma g | A0A2T6IUI9 |
| Toxoplasma g | A0A2T6IIA3 |
| Toxoplasma g | A0A2T6II49 |
| Toxoplasma g | A0A2T6ID59 |
| Toxoplasma g | A0A2T6II52 |

| | |
|---|---|
| Toxoplasma g | A0A2T6IRS2 |
| Toxoplasma g | A0A2T6J4Y3 |
| Toxoplasma g | A0A2T6IDB2 |
| Toxoplasma g | A0A2T6ISI1 |
| Toxoplasma g | A0A2T6J2S6 |
| Toxoplasma g | A0A2T6IS88 |
| Toxoplasma g | A0A2T6IQD1 |
| Toxoplasma g | A0A151GZ57 |
| Toxoplasma g | A0A151HHT1 |
| Toxoplasma g | A0A151HQ98 |
| Toxoplasma g | A0A151GZ62 |
| Toxoplasma g | A0A151HIC2 |
| Toxoplasma g | A0A151HH74 |
| Toxoplasma g | A0A151HFL2 |
| Toxoplasma g | A0A151H2M3 |
| Toxoplasma g | A0A151H4P4 |
| Toxoplasma g | A0A151HDF6 |
| Toxoplasma g | A0A151GYR0 |
| Toxoplasma g | A0A151HIU0 |
| Toxoplasma g | A0A151HH38 |
| Toxoplasma g | A0A151HIP1 |
| Toxoplasma g | A0A151HGR3 |
| Toxoplasma g | A0A151HL78 |
| Toxoplasma g | A0A151HGN6 |
| Toxoplasma g | A0A086PTH3 |
| Toxoplasma g | A0A086PPL1 |
| Toxoplasma g | A0A086PIW8 |
| Toxoplasma g | A0A086PGB6 |
| Toxoplasma g | A0A086QHG9 |
| Toxoplasma g | A0A086PYL8 |
| Toxoplasma g | A0A086PIX2 |
| Toxoplasma g | A0A086PGB7 |
| Toxoplasma g | A0A086PG37 |
| Toxoplasma g | A0A086PFY3 |
| Toxoplasma g | A0A086PSX1 |
| Toxoplasma g | A0A086PFS3 |
| Toxoplasma g | A0A086Q6Y4 |
| Toxoplasma g | A0A086PYK3 |
| Toxoplasma g | A0A086PZC2 |
| Toxoplasma g | A0A086PIX4 |
| Toxoplasma g | A0A086Q258 |

| | |
|---|---|
| Toxoplasma g | A0A086PSX2 |
| Toxoplasma g | A0A086PYV6 |
| Toxoplasma g | A0A086QF78 |
| Toxoplasma g | Q1JSE8 |
| Toxoplasma g | D8L561 |
| Toxoplasma g | D8L553 |
| Toxoplasma g | D8L555 |
| Toxoplasma g | D8L551 |
| Toxoplasma g | D8L554 |
| Toxoplasma g | D8L556 |
| Toxoplasma g | D8L552 |
| Toxoplasma g | D8L559 |
| Toxoplasma g | Q1JSK6 |
| Toxoplasma g | D8L558 |
| Toxoplasma g | D8L550 |
| Toxoplasma g | Q9XZH7 |
| Toxoplasma g | D8L557 |
| Trachemys sc | W8VX31 |
| Trachymyrme | A0A195E6P4 |
| Trachymyrme | A0A195DN17 |
| Trachymyrme | A0A195DN14 |
| Trachymyrme | A0A151JQX2 |
| Trachymyrme | A0A151J1A2 |
| Trachymyrme | A0A195DID1 |
| Trachymyrme | A0A151J003 |
| Trachymyrme | A0A151J024 |
| Trachymyrme | A0A195F9C1 |
| Trachymyrme | A0A195FXK7 |
| Trachymyrme | A0A195FSC2 |
| Trachymyrme | A0A195F1L6 |
| Trachymyrme | A0A195EZ81 |
| Trachymyrme | A0A195FP32 |
| Trachymyrme | A0A151WZ08 |
| Trachymyrme | A0A151X5L2 |
| Trachymyrme | A0A151XCA5 |
| Trachymyrme | A0A151WZC1 |
| Trachymyrme | A0A151WMX4 |
| Trachymyrme | A0A151WFX6 |
| Tranquillimor | A0A438AHQ8 |
| Trema orient | A0A2P5EEL5 |
| Trema orient | A0A2P5E9P3 |

Trema orient A0A2P5E9P1
Trema orient A0A2P5D0W9
Trema orient A0A2P5D0Z2
Trema orient A0A2P5D0Y8
Trema orient A0A2P5F2U6
Trema orient A0A2P5F1R2
Trema orient A0A2P5EHK8
Trema orient A0A2P5D0X0
Trema orient A0A2P5F9V6
Trema orient A0A2P5FFA8
Trema orient A0A2P5BR45
Trema orient A0A2P5E9N6
Trema orient A0A2P5BDV9
Trema orient A0A2P5E9S9
Trema orient A0A2P5E300
Trema orient A0A2P5BB99
Trema orient A0A2P5D0W8
Trema orient A0A2P5AU86
Trema orient A0A2P5ELI9
Trema orient A0A2P5BDW2
Trema orient A0A2P5BXZ5
Trema orient A0A2P5CM83
Trema orient A0A2P5BDX9
Trema orient A0A2P5EJU3
Trema orient A0A2P5E5Q8
Trema orient A0A2P5EJY0
Trema orient A0A2P5E9N1
Trema orient A0A2P5E7C6
Trema orient A0A2P5DNM0
Trema orient A0A2P5D111
Trema orient A0A2P5CM53
Trema orient A0A2P5EJU9
Trema orient A0A2P5EGL2
Trema orient A0A2P5D0X3
Trema orient A0A2P5DNJ7
Trema orient A0A2P5E9Q9
Trema orient A0A2P5BR70
Trema orient A0A2P5E7F0
Trema orient A0A2P5EM89
Trema orient A0A2P5CM78
Trema orient A0A2P5E7F7

Trema orient A0A2P5EM81
Trema orient A0A2P5EM56
Trema orient A0A2P5E7E7
Trema orient A0A2P5E7E4
Trema orient A0A2P5BDZ8
Trema orient A0A2P5D0Y7
Trema orient A0A2P5E7F8
Trema orient A0A2P5D0W7
Trema orient A0A2P5AU55
Trema orient A0A2P5E7E0
Trema orient A0A2P5E9N7
Trema orient A0A2P5EGQ1
Trema orient A0A2P5E7C3
Trema orient A0A2P5AVW6
Trema orient A0A2P5E7E1
Trema orient A0A2P5D0X7
Trema orient A0A2P5E7E9
Trema orient A0A2P5CM65
Trema orient A0A2P5D0Y2
Trema orient A0A2P5E9K7
Trema orient A0A2P5EK01
Trema orient A0A2P5E796
Trema orient A0A2P5CM46
Trema orient A0A2P5AU80
Trema orient A0A2P5DKI1
Trema orient A0A2P5EEU6
Trema orient A0A2P5AW26
Trema orient A0A2P5E7I1
Trema orient A0A2P5AU99
Trema orient A0A2P5E7K1
Trema orient A0A2P5E9P7
Trema orient A0A2P5EMF7
Trema orient A0A2P5CM64
Trema orient A0A2P5E7F2
Trema orient A0A2P5DUH7
Trema orient A0A2P5EGL5
Trema orient A0A2P5EJX1
Trema orient A0A2P5E329
Trema orient A0A2P5E7G0
Trema orient A0A2P5EMG7
Trema orient A0A2P5E305

| | |
|---|---|
| Trema orient | A0A2P5ARA6 |
| Trema orient | A0A2P5C4S2 |
| Trema orient | A0A2P5EJT3 |
| Trema orient | A0A2P5D0X8 |
| Trema orient | A0A2P5E322 |
| Trema orient | A0A2P5EM74 |
| Trema orient | A0A2P5BDW0 |
| Trema orient | A0A2P5B3V3 |
| Trema orient | A0A2P5BCJ5 |
| Trema orient | A0A2P5EJU7 |
| Trema orient | A0A2P5BCJ8 |
| Trema orient | A0A2P5B9F2 |
| Trema orient | A0A2P5E7C7 |
| Trema orient | A0A2P5E9R8 |
| Trema orient | A0A2P5EGP2 |
| Trema orient | A0A2P5B3Y1 |
| Trema orient | A0A2P5F9B9 |
| Trema orient | A0A2P5EJU0 |
| Trema orient | A0A2P5E9R2 |
| Trema orient | A0A2P5EGN4 |
| Triatoma infe | A0A161M4M9 |
| Triatoma infe | A0A023EZN7 |
| Triatoma infe | A0A023F552 |
| Tribolium cas | D6X332 |
| Tribolium cas | D6WSE9 |
| Tribolium cas | D2A5T8 |
| Tribolium cas | D6WCQ0 |
| Tribolium cas | D6WEC2 |
| Tribolium cas | A0A139WM22 |
| Tribolium cas | A0A139WEU2 |
| Tribolium cas | A0A139WF13 |
| Trichechus m | A0A2Y9D6R9 |
| Trichechus m | A0A2Y9RN69 |
| Trichechus m | A0A2Y9D6V5 |
| Trichechus m | A0A2Y9DUI6 |
| Trichechus m | A0A2Y9REY6 |
| Trichechus m | A0A2Y9DUP3 |
| Trichechus m | A0A2Y9RJN7 |
| Trichechus m | A0A2Y9QP02 |
| Trichechus m | A0A2Y9D6R6 |
| Trichechus m | A0A2Y9R8V0 |

Trichinella br A0A0V1CBM7
Trichinella br A0A0V1CL20
Trichinella br A0A0V1CF23
Trichinella br A0A0V1D321
Trichinella br A0A0V1DE70
Trichinella br A0A0V1CNZ7
Trichinella br A0A0V1DDL1
Trichinella br A0A0V1CYA0
Trichinella br A0A0V1D3X8
Trichinella br A0A0V1D312
Trichinella br A0A0V1D2Z4
Trichinella br A0A0V1DGI6
Trichinella br A0A0V1DIR9
Trichinella br A0A0V1DGQ0
Trichinella br A0A0V1D9K9
Trichinella br A0A0V1DCG4
Trichinella br A0A0V1D9Q7
Trichinella br A0A0V1CAI1
Trichinella br A0A0V1CF35
Trichinella br A0A0V1DGL4
Trichinella br A0A0V1CBT7
Trichinella br A0A0V1CQV4
Trichinella br A0A0V1CL47
Trichinella br A0A0V1D4Y2
Trichinella br A0A0V1CL07
Trichinella br A0A0V1CJ57
Trichinella br A0A0V1CQS1
Trichinella br A0A0V1CPE6
Trichinella br A0A0V1D2G1
Trichinella br A0A0V1DCQ6
Trichinella br A0A0V1CGZ2
Trichinella br A0A0V1CKH2
Trichinella br A0A0V1C8F4
Trichinella br A0A0V1D5M4
Trichinella br A0A0V1D5G8
Trichinella br A0A0V1CY65
Trichinella br A0A0V1CIY6
Trichinella br A0A0V1CS09
Trichinella br A0A0V1DDG4
Trichinella br A0A0V1D628
Trichinella br A0A0V1CL51

| | |
|---|---|
| Trichinella br | A0A0V1D3L4 |
| Trichinella br | A0A0V1D9P6 |
| Trichinella br | A0A0V1CF47 |
| Trichinella br | A0A0V1CEP1 |
| Trichinella br | A0A0V1CR67 |
| Trichinella br | A0A0V1D2S7 |
| Trichinella br | A0A0V1CCF9 |
| Trichinella br | A0A0V1CI86 |
| Trichinella br | A0A0V1D4Z1 |
| Trichinella br | A0A0V1D8K4 |
| Trichinella br | A0A0V1CY85 |
| Trichinella br | A0A0V1CIZ3 |
| Trichinella br | A0A0V1CLE7 |
| Trichinella br | A0A0V1D8Y2 |
| Trichinella br | A0A0V1CIU3 |
| Trichinella br | A0A0V1CLV0 |
| Trichinella br | A0A0V1D4H1 |
| Trichinella br | A0A0V1D4T5 |
| Trichinella br | A0A0V1CLT4 |
| Trichinella br | A0A0V1CC34 |
| Trichinella br | A0A0V1CQM7 |
| Trichinella br | A0A0V1CIG0 |
| Trichinella br | A0A0V1C615 |
| Trichinella br | A0A0V1CYG1 |
| Trichinella br | A0A0V1CL59 |
| Trichinella br | A0A0V1D176 |
| Trichinella br | A0A0V1CLC2 |
| Trichinella br | A0A0V1DDF7 |
| Trichinella br | A0A0V1CJ14 |
| Trichinella mu | A0A0V0U668 |
| Trichinella mu | A0A0V0TU85 |
| Trichinella mu | A0A0V0U6A2 |
| Trichinella mu | A0A0V0UGN9 |
| Trichinella mu | A0A0V0T888 |
| Trichinella mu | A0A0V0T7J2 |
| Trichinella mu | A0A0V0TGR2 |
| Trichinella mu | A0A0V0TE11 |
| Trichinella mu | A0A0V0TLQ5 |
| Trichinella mu | A0A0V0TLA0 |
| Trichinella mu | A0A0V0UAM2 |
| Trichinella mu | A0A0V0UI40 |

| | |
|---|---|
| Trichinella m | A0A0V0TLP7 |
| Trichinella m | A0A0V0TBB8 |
| Trichinella m | A0A0V0U7B1 |
| Trichinella m | A0A0V0TSM0 |
| Trichinella m | A0A0V0TU80 |
| Trichinella m | A0A0V0UAP6 |
| Trichinella m | A0A0V0TWW5 |
| Trichinella m | A0A0V0UI96 |
| Trichinella m | A0A0V0T000 |
| Trichinella m | A0A0V0TLE6 |
| Trichinella m | A0A0V0U6B0 |
| Trichinella m | A0A0V0UIZ5 |
| Trichinella m | A0A0V0TKS4 |
| Trichinella m | A0A0V0U6A5 |
| Trichinella m | A0A0V0TUK2 |
| Trichinella m | A0A0V0U664 |
| Trichinella m | A0A0V0TGD8 |
| Trichinella m | A0A0V0U6L1 |
| Trichinella m | A0A0V0TW68 |
| Trichinella m | A0A0V0TZG6 |
| Trichinella m | A0A0V0TZB3 |
| Trichinella m | A0A0V0U3H5 |
| Trichinella m | A0A0V0TR48 |
| Trichinella m | A0A0V0UAV4 |
| Trichinella m | A0A0V0TSU0 |
| Trichinella m | A0A0V0U6J8 |
| Trichinella m | A0A0V0TQX3 |
| Trichinella m | A0A0V0TCV2 |
| Trichinella m | A0A0V0T869 |
| Trichinella m | A0A0V0T5X4 |
| Trichinella m | A0A0V0T304 |
| Trichinella m | A0A0V0U0F0 |
| Trichinella m | A0A0V0T3U0 |
| Trichinella m | A0A0V0TSV2 |
| Trichinella m | A0A0V0T8Y2 |
| Trichinella m | A0A0V0TLJ2 |
| Trichinella m | A0A0V0TTX4 |
| Trichinella m | A0A0V0THH3 |
| Trichinella m | A0A0V0TCQ5 |
| Trichinella m | A0A0V0UGL3 |
| Trichinella m | A0A0V0T7W1 |

| | |
|---|---|
| Trichinella m | A0A0V0TGF1 |
| Trichinella m | A0A0V0TT12 |
| Trichinella m | A0A0V0TV07 |
| Trichinella m | A0A0V0UJZ1 |
| Trichinella m | A0A0V0T591 |
| Trichinella m | A0A0V0TFQ9 |
| Trichinella m | A0A0V0TW29 |
| Trichinella m | A0A0V0U516 |
| Trichinella m | A0A0V0TPM2 |
| Trichinella na | A0A0V1KNK0 |
| Trichinella na | A0A0V1KLF7 |
| Trichinella na | A0A0V1KYR3 |
| Trichinella na | A0A0V1KYC8 |
| Trichinella na | A0A0V1LNJ6 |
| Trichinella na | A0A0V1LAI7 |
| Trichinella na | A0A0V1KPV0 |
| Trichinella na | A0A0V1KPB9 |
| Trichinella na | A0A0V1KKU9 |
| Trichinella na | A0A0V1L1F4 |
| Trichinella na | A0A1Y3EXK1 |
| Trichinella na | A0A0V1LJJ4 |
| Trichinella na | A0A0V1LAS4 |
| Trichinella na | A0A0V1KMH3 |
| Trichinella na | A0A1Y3E8Z7 |
| Trichinella na | A0A1Y3E6K9 |
| Trichinella na | A0A1Y3EHJ6 |
| Trichinella na | A0A0V1LIU2 |
| Trichinella na | A0A1Y3E6I2 |
| Trichinella na | A0A1Y3EHG2 |
| Trichinella na | A0A1Y3EE55 |
| Trichinella na | A0A0V1KPJ6 |
| Trichinella na | A0A0V1L0L9 |
| Trichinella na | A0A0V1LEN4 |
| Trichinella na | A0A0V1LMK8 |
| Trichinella na | A0A1Y3EMP1 |
| Trichinella na | A0A0V1KMD9 |
| Trichinella na | A0A1Y3E8I5 |
| Trichinella na | A0A0V1L1M0 |
| Trichinella na | A0A1Y3E9F8 |
| Trichinella na | A0A0V1LIH7 |
| Trichinella na | A0A0V1L1N0 |

Trichinella na A0A1Y3EK45
Trichinella na A0A0V1L5K7
Trichinella na A0A0V1LAY9
Trichinella na A0A0V1L1G4
Trichinella na A0A1Y3E3J3
Trichinella na A0A0V1KM43
Trichinella na A0A0V1LBC3
Trichinella na A0A1Y3E9Q0
Trichinella na A0A0V1KYR0
Trichinella na A0A0V1L1E8
Trichinella na A0A0V1KP52
Trichinella na A0A0V1L0I9
Trichinella na A0A0V1LEV0
Trichinella na A0A0V1LAP5
Trichinella na A0A0V1L1M7
Trichinella na A0A0V1LE33
Trichinella na A0A1Y3EH27
Trichinella na A0A1Y3E603
Trichinella na A0A0V1LJH2
Trichinella na A0A0V1LMI4
Trichinella na A0A0V1KYB0
Trichinella na A0A1Y3ET15
Trichinella na A0A0V1L1E6
Trichinella na A0A0V1KR57
Trichinella na A0A0V1L136
Trichinella na A0A0V1LT84
Trichinella na A0A0V1KLJ0
Trichinella na A0A1Y3EP57
Trichinella na A0A1Y3EUE7
Trichinella na A0A0V1KXX6
Trichinella na A0A1Y3EF82
Trichinella na A0A1Y3EZI1
Trichinella na A0A0V1KXY2
Trichinella na A0A1Y3E7T0
Trichinella na A0A0V1KPL5
Trichinella na A0A1Y3E8Z4
Trichinella na A0A0V1KX10
Trichinella na A0A0V1LFN7
Trichinella na A0A1Y3ENN2
Trichinella na A0A1Y3EJ87
Trichinella na A0A1Y3F0N2

| | |
|---|---|
| Trichinella na | A0A1Y3EDK3 |
| Trichinella na | A0A1Y3E9V6 |
| Trichinella na | A0A0V1KPY5 |
| Trichinella na | A0A1Y3EJ46 |
| Trichinella na | A0A0V1LE29 |
| Trichinella na | A0A0V1LED3 |
| Trichinella na | A0A0V1LLN6 |
| Trichinella na | A0A1Y3EUS4 |
| Trichinella na | A0A0V1L2N9 |
| Trichinella na | A0A0V1L3H8 |
| Trichinella na | A0A0V1KR45 |
| Trichinella na | A0A0V1LAL0 |
| Trichinella na | A0A0V1KS99 |
| Trichinella na | A0A1Y3EGT9 |
| Trichinella na | A0A0V1LCN5 |
| Trichinella na | A0A1Y3ETI7 |
| Trichinella na | A0A1Y3EAN3 |
| Trichinella na | A0A1Y3EH07 |
| Trichinella na | A0A0V1LU40 |
| Trichinella na | A0A1Y3E7K6 |
| Trichinella na | A0A0V1KSI2 |
| Trichinella na | A0A0V1LES1 |
| Trichinella na | A0A1Y3E7U1 |
| Trichinella na | A0A1Y3E9F9 |
| Trichinella na | A0A0V1LB56 |
| Trichinella na | A0A0V1LNE5 |
| Trichinella na | A0A0V1L7E6 |
| Trichinella na | A0A0V1L6Z9 |
| Trichinella ne | A0A0V0RYN5 |
| Trichinella ne | A0A0V0SCW7 |
| Trichinella ne | A0A0V0RIU1 |
| Trichinella ne | A0A0V0RN79 |
| Trichinella ne | A0A0V0SH81 |
| Trichinella ne | A0A0V0SCZ0 |
| Trichinella ne | A0A0V0RV95 |
| Trichinella ne | A0A0V0S538 |
| Trichinella ne | A0A0V0RM32 |
| Trichinella ne | A0A0V0SCM0 |
| Trichinella ne | A0A0V0RFG5 |
| Trichinella ne | A0A0V0RRH9 |
| Trichinella ne | A0A0V0RRB5 |

| | |
|---|---|
| Trichinella ne | A0A0V0RG56 |
| Trichinella ne | A0A0V0S806 |
| Trichinella ne | A0A0V0RZJ1 |
| Trichinella ne | A0A0V0RDH2 |
| Trichinella ne | A0A0V0RGM5 |
| Trichinella ne | A0A0V0SNF7 |
| Trichinella ne | A0A0V0S2V9 |
| Trichinella ne | A0A0V0RYH3 |
| Trichinella ne | A0A0V0RGW1 |
| Trichinella ne | A0A0V0SCX3 |
| Trichinella ne | A0A0V0REU3 |
| Trichinella ne | A0A0V0RZP8 |
| Trichinella ne | A0A0V0RT14 |
| Trichinella ne | A0A0V0S6I1 |
| Trichinella ne | A0A0V0RJS2 |
| Trichinella ne | A0A0V0SEE5 |
| Trichinella ne | A0A0V0RZB8 |
| Trichinella ne | A0A0V0RK34 |
| Trichinella ne | A0A0V0S5T4 |
| Trichinella ne | A0A0V0RZM6 |
| Trichinella ne | A0A0V0SMX2 |
| Trichinella ne | A0A0V0RX40 |
| Trichinella ne | A0A0V0S9B4 |
| Trichinella ne | A0A0V0RFF5 |
| Trichinella ne | A0A0V0RYQ4 |
| Trichinella ne | A0A0V0S5G1 |
| Trichinella ne | A0A0V0REL5 |
| Trichinella ne | A0A0V0RSM5 |
| Trichinella ne | A0A0V0RYQ1 |
| Trichinella ne | A0A0V0SCP1 |
| Trichinella ne | A0A0V0SCK5 |
| Trichinella ne | A0A0V0RF35 |
| Trichinella ne | A0A0V0RFT3 |
| Trichinella ne | A0A0V0RPN2 |
| Trichinella ne | A0A0V0RQB5 |
| Trichinella ne | A0A0V0RHM2 |
| Trichinella ne | A0A0V0RZD1 |
| Trichinella ne | A0A0V0REM5 |
| Trichinella ne | A0A0V0S4Z2 |
| Trichinella ne | A0A0V0RZT2 |
| Trichinella ne | A0A0V0RS35 |

| | |
|---|---|
| Trichinella ne | A0A0V0S652 |
| Trichinella ne | A0A0V0RG20 |
| Trichinella ne | A0A0V0RM92 |
| Trichinella ne | A0A0V0S584 |
| Trichinella ne | A0A0V0S4X7 |
| Trichinella ne | A0A0V0RDD2 |
| Trichinella ne | A0A0V0S541 |
| Trichinella ne | A0A0V0REJ1 |
| Trichinella ne | A0A0V0S0D1 |
| Trichinella ne | A0A0V0RK03 |
| Trichinella ne | A0A0V0S3G7 |
| Trichinella ne | A0A0V0RX96 |
| Trichinella ne | A0A0V0RHC1 |
| Trichinella ne | A0A0V0RSZ9 |
| Trichinella ne | A0A0V0RIE7 |
| Trichinella ne | A0A0V0S9G5 |
| Trichinella ne | A0A0V0RTR4 |
| Trichinella ne | A0A0V0RIE5 |
| Trichinella ne | A0A0V0RYP8 |
| Trichinella ne | A0A0V0RSK8 |
| Trichinella ne | A0A0V0RFI6 |
| Trichinella ne | A0A0V0S2H6 |
| Trichinella ne | A0A0V0S5G8 |
| Trichinella ne | A0A0V0S0F9 |
| Trichinella ne | A0A0V0S5R1 |
| Trichinella pa | A0A0V1MVW8 |
| Trichinella pa | A0A0V1MAI7 |
| Trichinella pa | A0A0V1MYP5 |
| Trichinella pa | A0A0V1N2V7 |
| Trichinella pa | A0A0V1MUG7 |
| Trichinella pa | A0A0V1MUG4 |
| Trichinella pa | A0A0V1MYQ2 |
| Trichinella pa | A0A0V1MGC0 |
| Trichinella pa | A0A0V1N6W8 |
| Trichinella pa | A0A0V1N6W7 |
| Trichinella pa | A0A0V1MF20 |
| Trichinella pa | A0A0V1MNC5 |
| Trichinella pa | A0A0V1MJU5 |
| Trichinella pa | A0A0V1MYR0 |
| Trichinella pa | A0A0V1MZ66 |
| Trichinella pa | A0A0V1MAV6 |

| | |
|---|---|
| Trichinella pa | A0A0V1MYQ0 |
| Trichinella pa | A0A0V1N2Z9 |
| Trichinella pa | A0A0V1MH28 |
| Trichinella pa | A0A0V1MVV2 |
| Trichinella pa | A0A0V1N167 |
| Trichinella pa | A0A0V1MYJ7 |
| Trichinella pa | A0A0V1MT15 |
| Trichinella pa | A0A0V1MH70 |
| Trichinella pa | A0A0V1M2G6 |
| Trichinella pa | A0A0V1MUW3 |
| Trichinella pa | A0A0V1M9J0 |
| Trichinella pa | A0A0V1MHZ6 |
| Trichinella pa | A0A0V1MTF5 |
| Trichinella pa | A0A0V1MGN0 |
| Trichinella pa | A0A0V1MZ68 |
| Trichinella pa | A0A0V1N6Q2 |
| Trichinella pa | A0A0V1MGD7 |
| Trichinella pa | A0A0V1MIW0 |
| Trichinella pa | A0A0V1N5S4 |
| Trichinella pa | A0A0V1MYM3 |
| Trichinella pa | A0A0V1MI03 |
| Trichinella pa | A0A0V1MYL8 |
| Trichinella pa | A0A0V1MSA4 |
| Trichinella pa | A0A0V1MZA7 |
| Trichinella pa | A0A0V1MZ29 |
| Trichinella pa | A0A0V1MUK2 |
| Trichinella pa | A0A0V1MZ89 |
| Trichinella pa | A0A0V1MUM5 |
| Trichinella pa | A0A0V1MIL5 |
| Trichinella pa | A0A0V1MZ88 |
| Trichinella pa | A0A0V1MTC8 |
| Trichinella pa | A0A0V1M8W6 |
| Trichinella pa | A0A0V1M0N9 |
| Trichinella pa | A0A0V1N9C6 |
| Trichinella pa | A0A0V1MN65 |
| Trichinella pa | A0A0V1MNB1 |
| Trichinella pa | A0A0V1MWG2 |
| Trichinella pa | A0A0V1N8S1 |
| Trichinella pa | A0A0V1MYJ6 |
| Trichinella pa | A0A0V1N750 |
| Trichinella pa | A0A0V1MH06 |

| | |
|---|---|
| Trichinella pa | A0A0V1MYI7 |
| Trichinella pa | A0A0V1MYQ1 |
| Trichinella pa | A0A0V0ZT98 |
| Trichinella pa | A0A0V1AAJ2 |
| Trichinella pa | A0A0V0ZDL2 |
| Trichinella pa | A0A0V1AFC5 |
| Trichinella pa | A0A0V0ZMA2 |
| Trichinella pa | A0A0V0ZI13 |
| Trichinella pa | A0A0V1ADL5 |
| Trichinella pa | A0A0V0ZA63 |
| Trichinella pa | A0A0V0ZJI5 |
| Trichinella pa | A0A0V1AGR2 |
| Trichinella pa | A0A0V0ZEN6 |
| Trichinella pa | A0A0V0ZJY6 |
| Trichinella pa | A0A0V0ZI71 |
| Trichinella pa | A0A0V0ZJV3 |
| Trichinella pa | A0A0V1A9E4 |
| Trichinella pa | A0A0V0ZKA1 |
| Trichinella pa | A0A0V1A5G3 |
| Trichinella pa | A0A0V1AH40 |
| Trichinella pa | A0A0V0Z9W9 |
| Trichinella pa | A0A0V0ZMW0 |
| Trichinella pa | A0A0V0ZPP6 |
| Trichinella pa | A0A0V0ZJ05 |
| Trichinella pa | A0A0V1A350 |
| Trichinella pa | A0A0V0Z9K2 |
| Trichinella pa | A0A0V0ZQB1 |
| Trichinella pa | A0A0V1A5H0 |
| Trichinella pa | A0A0V0ZQQ6 |
| Trichinella pa | A0A0V1A5B9 |
| Trichinella pa | A0A0V0ZCY3 |
| Trichinella pa | A0A0V1ACU1 |
| Trichinella pa | A0A0V0ZID7 |
| Trichinella pa | A0A0V0ZJE6 |
| Trichinella pa | A0A0V0ZA86 |
| Trichinella pa | A0A0V1AEG0 |
| Trichinella pa | A0A0V0ZHM8 |
| Trichinella pa | A0A0V0ZU82 |
| Trichinella pa | A0A0V0ZXE1 |
| Trichinella pa | A0A0V1A735 |
| Trichinella pa | A0A0V0ZBC9 |

| | |
|---|---|
| Trichinella pa | A0A0V1A2Y6 |
| Trichinella pa | A0A0V1ADC4 |
| Trichinella pa | A0A0V1A315 |
| Trichinella pa | A0A0V0ZG93 |
| Trichinella pa | A0A0V0ZG08 |
| Trichinella pa | A0A0V0ZBZ0 |
| Trichinella pa | A0A0V0ZC36 |
| Trichinella pa | A0A0V0ZBF0 |
| Trichinella pa | A0A0V1A3P0 |
| Trichinella pa | A0A0V0ZBK7 |
| Trichinella pa | A0A0V1ACP2 |
| Trichinella pa | A0A0V0ZUN7 |
| Trichinella pa | A0A0V1ACR5 |
| Trichinella pa | A0A0V1A6I5 |
| Trichinella pa | A0A0V0ZRK6 |
| Trichinella pa | A0A0V0ZDM2 |
| Trichinella pa | A0A0V0ZC11 |
| Trichinella pa | A0A0V0ZSY6 |
| Trichinella pa | A0A0V0ZAI4 |
| Trichinella pa | A0A0V1AGP3 |
| Trichinella pa | A0A0V0ZK36 |
| Trichinella pa | A0A0V1ABN4 |
| Trichinella pa | A0A0V0ZK58 |
| Trichinella pa | A0A0V0Z425 |
| Trichinella ps | A0A0V1K9D8 |
| Trichinella ps | A0A0V1JL75 |
| Trichinella ps | A0A0V1KA64 |
| Trichinella ps | A0A0V1IA74 |
| Trichinella ps | A0A0V1ECJ0 |
| Trichinella ps | A0A0V1ERM8 |
| Trichinella ps | A0A0V1EJT7 |
| Trichinella ps | A0A0V1E276 |
| Trichinella ps | A0A0V1K9L3 |
| Trichinella ps | A0A0V1IPI2 |
| Trichinella ps | A0A0V1IKD9 |
| Trichinella ps | A0A0V1IKA3 |
| Trichinella ps | A0A0V1IPX2 |
| Trichinella ps | A0A0V1KD92 |
| Trichinella ps | A0A0V1JZQ7 |
| Trichinella ps | A0A0V0YKR7 |
| Trichinella ps | A0A0V1JCJ6 |

Trichinella ps A0A0V1JE87
Trichinella ps A0A0V1J8T7
Trichinella ps A0A0V1JS72
Trichinella ps A0A0V1IJ16
Trichinella ps A0A0V0YL79
Trichinella ps A0A0V1K592
Trichinella ps A0A0V1IE69
Trichinella ps A0A0V1K9L2
Trichinella ps A0A0V0XEJ1
Trichinella ps A0A0V1JC56
Trichinella ps A0A0V1I359
Trichinella ps A0A0V1K7S5
Trichinella ps A0A0V1IQ50
Trichinella ps A0A0V1J492
Trichinella ps A0A0V0YE74
Trichinella ps A0A0V0Y8V0
Trichinella ps A0A0V1EVN0
Trichinella ps A0A0V1K9K9
Trichinella ps A0A0V1KD78
Trichinella ps A0A0V1G4T5
Trichinella ps A0A0V1IAL1
Trichinella ps A0A0V1FH81
Trichinella ps A0A0V1IDD8
Trichinella ps A0A0V1JC58
Trichinella ps A0A0V1J0K8
Trichinella ps A0A0V0Y705
Trichinella ps A0A0V1EVL8
Trichinella ps A0A0V1K9B4
Trichinella ps A0A0V0XNP3
Trichinella ps A0A0V1FRY9
Trichinella ps A0A0V1IAQ7
Trichinella ps A0A0V0XX86
Trichinella ps A0A0V1IQJ7
Trichinella ps A0A0V1JE40
Trichinella ps A0A0V1FHE6
Trichinella ps A0A0V1K4X2
Trichinella ps A0A0V1H366
Trichinella ps A0A0V1FQ76
Trichinella ps A0A0V1EVA7
Trichinella ps A0A0V0XW16
Trichinella ps A0A0V1K5I2

Trichinella ps A0A0V1EUZ6
Trichinella ps A0A0V0YGT7
Trichinella ps A0A0V1IAS5
Trichinella ps A0A0V1E2U6
Trichinella ps A0A0V1E1V7
Trichinella ps A0A0V1G2W2
Trichinella ps A0A0V1JI52
Trichinella ps A0A0V1G298
Trichinella ps A0A0V1EV41
Trichinella ps A0A0V1EMN5
Trichinella ps A0A0V1JHF0
Trichinella ps A0A0V1KDC2
Trichinella ps A0A0V1K5A6
Trichinella ps A0A0V1JJD4
Trichinella ps A0A0V1K9V2
Trichinella ps A0A0V1FLB8
Trichinella ps A0A0V0XY64
Trichinella ps A0A0V1FYN9
Trichinella ps A0A0V1J0M9
Trichinella ps A0A0V1J4L5
Trichinella ps A0A0V0YMV1
Trichinella ps A0A0V1FQM2
Trichinella ps A0A0V1GQ57
Trichinella ps A0A0V1KEH4
Trichinella ps A0A0V0YGT1
Trichinella ps A0A0V0XY09
Trichinella ps A0A0V1EUS6
Trichinella ps A0A0V1K589
Trichinella ps A0A0V0XTS1
Trichinella ps A0A0V1ETL0
Trichinella ps A0A0V1FFS0
Trichinella ps A0A0V1JL92
Trichinella ps A0A0V1ED64
Trichinella ps A0A0V1ETL9
Trichinella ps A0A0V1EV55
Trichinella ps A0A0V1J0J8
Trichinella ps A0A0V1G235
Trichinella ps A0A0V1KDB0
Trichinella ps A0A0V1K3Q7
Trichinella ps A0A0V1EEX1
Trichinella ps A0A0V0XTR8

Trichinella ps A0A0V1FEX1
Trichinella ps A0A0V0Y6W9
Trichinella ps A0A0V0XKW1
Trichinella ps A0A0V0XKX5
Trichinella ps A0A0V1K9J3
Trichinella ps A0A0V0XKY0
Trichinella ps A0A0V1EMK3
Trichinella ps A0A0V1FRL6
Trichinella ps A0A0V1F4T6
Trichinella ps A0A0V1KDD7
Trichinella ps A0A0V1FBJ2
Trichinella ps A0A0V1K421
Trichinella ps A0A0V1FBW8
Trichinella ps A0A0V1EVA1
Trichinella ps A0A0V0XU04
Trichinella ps A0A0V1IRZ9
Trichinella ps A0A0V1EV45
Trichinella ps A0A0V0XNW4
Trichinella ps A0A0V1EVN2
Trichinella ps A0A0V0YMZ8
Trichinella ps A0A0V0XJ62
Trichinella ps A0A0V0YFK6
Trichinella ps A0A0V1JPJ3
Trichinella ps A0A0V1J0J3
Trichinella ps A0A0V0Y715
Trichinella ps A0A0V1FMV7
Trichinella ps A0A0V1JCP9
Trichinella ps A0A0V1E9M7
Trichinella ps A0A0V1KDC1
Trichinella ps A0A0V1FEU7
Trichinella ps A0A0V1K0T0
Trichinella ps A0A0V1DST8
Trichinella ps A0A0V1IHH3
Trichinella ps A0A0V1J0I8
Trichinella ps A0A0V1G0P4
Trichinella ps A0A0V0XL17
Trichinella ps A0A0V1ESY5
Trichinella ps A0A0V1K9H0
Trichinella ps A0A0V0XUM5
Trichinella ps A0A0V1EVN4
Trichinella ps A0A0V1EU35

Trichinella ps A0A0V1DXH2
Trichinella ps A0A0V1JCS1
Trichinella ps A0A0V0XNL7
Trichinella ps A0A0V1E5G0
Trichinella ps A0A0V1G1X6
Trichinella ps A0A0V1EWX6
Trichinella ps A0A0V1K6H8
Trichinella ps A0A0V1FAF9
Trichinella ps A0A0V0YNX4
Trichinella ps A0A0V1K3Q0
Trichinella ps A0A0V1JE80
Trichinella ps A0A0V1HB36
Trichinella ps A0A0V1IYZ4
Trichinella ps A0A0V1KEG0
Trichinella ps A0A0V0XUB1
Trichinella ps A0A0V0YN83
Trichinella ps A0A0V1IIS1
Trichinella ps A0A0V1K9N4
Trichinella ps A0A0V1EV51
Trichinella ps A0A0V1K5S7
Trichinella ps A0A0V0XS62
Trichinella ps A0A0V1IVQ1
Trichinella ps A0A0V1FQU3
Trichinella ps A0A0V1K9E0
Trichinella ps A0A0V1JPK7
Trichinella ps A0A0V1ETF5
Trichinella ps A0A0V1FAR0
Trichinella ps A0A0V1ENV6
Trichinella ps A0A0V1FMQ4
Trichinella ps A0A0V1JPN2
Trichinella ps A0A0V1J380
Trichinella ps A0A0V0XJI2
Trichinella ps A0A0V1KDE3
Trichinella ps A0A0V1DXF5
Trichinella ps A0A0V0YE73
Trichinella ps A0A0V1F7L0
Trichinella ps A0A0V1JEB3
Trichinella ps A0A0V1F5X2
Trichinella ps A0A0V0YMM2
Trichinella ps A0A0V1FBB6
Trichinella ps A0A0V1ESS6

Trichinella ps A0A0V1G0K7
Trichinella ps A0A0V1EVU6
Trichinella ps A0A0V1JBG7
Trichinella ps A0A0V1G4S3
Trichinella ps A0A0V0YMZ0
Trichinella ps A0A0V1KDD9
Trichinella ps A0A0V1IVW1
Trichinella ps A0A0V0YGE0
Trichinella ps A0A0V1JZL9
Trichinella ps A0A0V0XXD7
Trichinella ps A0A0V0XEA7
Trichinella ps A0A0V1JE69
Trichinella ps A0A0V1J0L5
Trichinella ps A0A0V0XFS5
Trichinella ps A0A0V1K9R0
Trichinella ps A0A0V1G5W1
Trichinella ps A0A0V1JCL5
Trichinella ps A0A0V0XUK2
Trichinella ps A0A0V1DX07
Trichinella ps A0A0V0XZB1
Trichinella ps A0A0V1IZ01
Trichinella ps A0A0V1ET77
Trichinella ps A0A0V1F060
Trichinella ps A0A0V1JEA8
Trichinella ps A0A0V1FY33
Trichinella ps A0A0V1FQA4
Trichinella ps A0A0V1JHS7
Trichinella ps A0A0V1ED73
Trichinella ps A0A0V1K5M6
Trichinella ps A0A0V0Y1Z1
Trichinella ps A0A0V1JCM7
Trichinella ps A0A0V1IHF8
Trichinella ps A0A0V1J0Q3
Trichinella ps A0A0V1IVX3
Trichinella ps A0A0V1ETT3
Trichinella ps A0A0V1EU49
Trichinella ps A0A0V1E0G2
Trichinella ps A0A0V1KD83
Trichinella ps A0A0V1FAL4
Trichinella ps A0A0V1ET37
Trichinella ps A0A0V0XNN7

Trichinella ps A0A0V1K5K3
Trichinella ps A0A0V0XRQ1
Trichinella ps A0A0V1EKP8
Trichinella ps A0A0V1G1T4
Trichinella ps A0A0V1JJ61
Trichinella ps A0A0V1IHH9
Trichinella ps A0A0V1EVH0
Trichinella ps A0A0V1ETP5
Trichinella ps A0A0V0XUH1
Trichinella ps A0A0V0XLI9
Trichinella ps A0A0V1J0K0
Trichinella ps A0A0V1IYY2
Trichinella ps A0A0V1JPL8
Trichinella ps A0A0V0XNN5
Trichinella ps A0A0V0Y6W4
Trichinella ps A0A0V1G0P5
Trichinella sp A0A0V0WI90
Trichinella sp A0A0V0WS36
Trichinella sp A0A0V0X5U7
Trichinella sp A0A0V0WZP8
Trichinella sp A0A0V0WF21
Trichinella sp A0A0V0WMF7
Trichinella sp A0A0V0WJN0
Trichinella sp A0A0V0X647
Trichinella sp A0A0V0WMC0
Trichinella sp A0A0V0X8U0
Trichinella sp A0A0V0WGF2
Trichinella sp A0A0V0WGK4
Trichinella sp A0A0V0WEM4
Trichinella sp A0A0V0X8Q3
Trichinella sp A0A0V0WH09
Trichinella sp A0A0V0WML0
Trichinella sp A0A0V0X6B4
Trichinella sp A0A0V0WZT2
Trichinella sp A0A0V0WS86
Trichinella sp A0A0V0X7G3
Trichinella sp A0A0V0WYZ7
Trichinella sp A0A0V0W8L3
Trichinella sp A0A0V0W0I6
Trichinella sp A0A0V0WGI7
Trichinella sp A0A0V0WYQ7

Trichinella sp A0A0V0WX01
Trichinella sp A0A0V0WF38
Trichinella sp A0A0V0W1G0
Trichinella sp A0A0V0W776
Trichinella sp A0A0V0WXK0
Trichinella sp A0A0V0WA24
Trichinella sp A0A0V0X335
Trichinella sp A0A0V0WS82
Trichinella sp A0A0V0W1K5
Trichinella sp A0A0V0W1Z3
Trichinella sp A0A0V0W1S4
Trichinella sp A0A0V0WAI4
Trichinella sp A0A0V0WHC9
Trichinella sp A0A0V0V0P6
Trichinella sp A0A0V0WHA1
Trichinella sp A0A0V0WM77
Trichinella sp A0A0V0WJ02
Trichinella sp A0A0V0X0M5
Trichinella sp A0A0V0X5Y4
Trichinella sp A0A0V0WF91
Trichinella sp A0A0V0X004
Trichinella sp A0A0V0X092
Trichinella sp A0A0V0WZR3
Trichinella sp A0A0V0X0N1
Trichinella sp A0A0V0X081
Trichinella sp A0A0V0WDP4
Trichinella sp A0A0V0WD03
Trichinella sp A0A0V0WHG0
Trichinella sp A0A0V0W0M1
Trichinella sp A0A0V0ULH6
Trichinella sp A0A0V0WX65
Trichinella sp A0A0V0WJM1
Trichinella sp A0A0V0V1L2
Trichinella sp A0A0V0WKK3
Trichinella sp A0A0V0WJS6
Trichinella sp A0A0V0W1F0
Trichinella sp A0A0V0WET2
Trichinella sp A0A0V0W1R2
Trichinella sp A0A0V0WK58
Trichinella sp A0A0V0W771
Trichinella sp A0A0V0WAQ3

Trichinella sp A0A0V0WJZ4
Trichinella sp A0A0V0V151
Trichinella sp A0A0V0WWY8
Trichinella sp A0A0V0VUH1
Trichinella sp A0A0V0X4G5
Trichinella sp A0A0V0WXW5
Trichinella sp A0A0V0WHJ7
Trichinella sp A0A0V0WJM5
Trichinella sp A0A0V0WF23
Trichinella sp A0A0V0UYA2
Trichinella sp A0A0V0W7A7
Trichinella sp A0A0V0VH07
Trichinella sp A0A0V0W798
Trichinella sp A0A0V0WXH6
Trichinella sp A0A0V0W6U1
Trichinella sp A0A0V0X369
Trichinella sp A0A0V0W1Y2
Trichinella sp A0A0V0W1I5
Trichinella sp A0A0V0WJZ8
Trichinella sp A0A0V0W0F4
Trichinella sp A0A0V0W1K6
Trichinella sp A0A0V0VYK2
Trichinella sp A0A0V0WAX9
Trichinella sp A0A0V0X4A4
Trichinella sp A0A0V0WY03
Trichinella sp A0A0V0WHZ9
Trichinella sp A0A0V0W4Y7
Trichinella sp A0A0V0WM81
Trichinella sp A0A0V1P5U0
Trichinella sp A0A0V1NES3
Trichinella sp A0A0V1P2I1
Trichinella sp A0A0V1PI68
Trichinella sp A0A0V1P853
Trichinella sp A0A0V1NF35
Trichinella sp A0A0V1PCG3
Trichinella sp A0A0V1NXA6
Trichinella sp A0A0V1NL85
Trichinella sp A0A0V1NGD5
Trichinella sp A0A0V1PF26
Trichinella sp A0A0V1PK08
Trichinella sp A0A0V1NHJ3

Trichinella sp A0A0V1NLL0
Trichinella sp A0A0V1PLE3
Trichinella sp A0A0V1P795
Trichinella sp A0A0V1PJS4
Trichinella sp A0A0V1NY22
Trichinella sp A0A0V1P7K0
Trichinella sp A0A0V1P5F5
Trichinella sp A0A0V1NLV9
Trichinella sp A0A0V1NG13
Trichinella sp A0A0V1P9L4
Trichinella sp A0A0V1P7L8
Trichinella sp A0A0V1NEN6
Trichinella sp A0A0V1P771
Trichinella sp A0A0V1NM54
Trichinella sp A0A0V1P5T0
Trichinella sp A0A0V1P8Q0
Trichinella sp A0A0V1NE86
Trichinella sp A0A0V1P280
Trichinella sp A0A0V1NS27
Trichinella sp A0A0V1P770
Trichinella sp A0A0V1PI77
Trichinella sp A0A0V1P7C0
Trichinella sp A0A0V1PFJ0
Trichinella sp A0A0V1P721
Trichinella sp A0A0V1NYC7
Trichinella sp A0A0V1P6T8
Trichinella sp A0A0V1NLN2
Trichinella sp A0A0V1NLK1
Trichinella sp A0A0V1PAZ0
Trichinella sp A0A0V1P2H3
Trichinella sp A0A0V1NXE2
Trichinella sp A0A0V1P6D6
Trichinella sp A0A0V1NVS2
Trichinella sp A0A0V1NVB0
Trichinella sp A0A0V1P6Q0
Trichinella sp A0A0V1PF39
Trichinella sp A0A0V1PEY2
Trichinella sp A0A0V1P7H3
Trichinella sp A0A0V1P9E1
Trichinella sp A0A0V1NX80
Trichinella sp A0A0V1NTT6

Trichinella sp A0A0V1NYS7
Trichinella sp A0A0V1P6M8
Trichinella sp A0A0V1NGF7
Trichinella sp A0A0V1P4Z0
Trichinella sp A0A0V0VJK1
Trichinella sp A0A0V0VR93
Trichinella sp A0A0V0UTV5
Trichinella sp A0A0V0UQR6
Trichinella sp A0A0V0UM00
Trichinella sp A0A0V0V174
Trichinella sp A0A0V0V426
Trichinella sp A0A0V0UTZ7
Trichinella sp A0A0V0VRE1
Trichinella sp A0A0V0VRI1
Trichinella sp A0A0V0UNQ3
Trichinella sp A0A0V0VRC8
Trichinella sp A0A0V0VE39
Trichinella sp A0A0V0VFJ5
Trichinella sp A0A0V0UQ81
Trichinella sp A0A0V0VFQ4
Trichinella sp A0A0V0VFH2
Trichinella sp A0A0V0VPC7
Trichinella sp A0A0V0UUY9
Trichinella sp A0A0V0VRE0
Trichinella sp A0A0V0VP79
Trichinella sp A0A0V0V552
Trichinella sp A0A0V0VFW1
Trichinella sp A0A0V0VS73
Trichinella sp A0A0V0VWE9
Trichinella sp A0A0V0V155
Trichinella sp A0A0V0VCB7
Trichinella sp A0A0V0VMC3
Trichinella sp A0A0V0V7D6
Trichinella sp A0A0V0UPR0
Trichinella sp A0A0V0UUV8
Trichinella sp A0A0V0V210
Trichinella sp A0A0V0VE27
Trichinella sp A0A0V0VRQ3
Trichinella sp A0A0V0VEC8
Trichinella sp A0A0V0UQH4
Trichinella sp A0A0V0VFX1

Trichinella sp A0A0V0VSL6
Trichinella sp A0A0V0VRE9
Trichinella sp A0A0V0V6V2
Trichinella sp A0A0V0VP70
Trichinella sp A0A0V0VSK7
Trichinella sp A0A0V0UUX7
Trichinella sp A0A0V0VED2
Trichinella sp A0A0V0VFN7
Trichinella sp A0A0V0V499
Trichinella sp A0A0V0V3J5
Trichinella sp A0A0V0V587
Trichinella sp A0A0V0VW86
Trichinella sp A0A0V0VQ95
Trichinella sp A0A0V0VCJ2
Trichinella sp A0A0V0VSS5
Trichinella sp A0A0V0UJV4
Trichinella sp A0A0V0UPU4
Trichinella sp A0A0V0VGC0
Trichinella sp A0A0V0U3P2
Trichinella sp A0A0V0VE47
Trichinella sp A0A0V0UTV0
Trichinella sp A0A0V0UMK8
Trichinella sp A0A0V0T3U1
Trichinella sp A0A0V0VPC1
Trichinella sp A0A0V0V159
Trichinella sp A0A0V0VRW7
Trichinella sp A0A0V0VTE1
Trichinella sp A0A0V0VJ88
Trichinella sp A0A0V0UVL3
Trichinella sp A0A0V0UUP5
Trichinella sp A0A0V0VE17
Trichinella sp A0A0V0V119
Trichinella sp A0A0V0UNR0
Trichinella sp A0A0V0VFK5
Trichinella sp A0A0V0VLU4
Trichinella sp A0A0V0VDL9
Trichinella sp A0A0V0VW85
Trichinella sp A0A0V0VRG0
Trichinella sp A0A0V0UQQ3
Trichinella sp A0A0V0VEJ2
Trichinella sp A0A0V0VDY8

Trichinella sp A0A0V1BV44
Trichinella sp A0A0V1ASI9
Trichinella sp A0A0V1AWR7
Trichinella sp A0A0V1C327
Trichinella sp A0A0V1ARX7
Trichinella sp A0A0V1ASM2
Trichinella sp A0A0V1BM31
Trichinella sp A0A0V1ASM0
Trichinella sp A0A0V1BY88
Trichinella sp A0A0V1BSS1
Trichinella sp A0A0V1BU64
Trichinella sp A0A0V1BYB4
Trichinella sp A0A0V1B1M0
Trichinella sp A0A0V1BFH0
Trichinella sp A0A0V1BTA4
Trichinella sp A0A0V1BU83
Trichinella sp A0A0V1BYZ5
Trichinella sp A0A0V1BYW3
Trichinella sp A0A0V1ASI6
Trichinella sp A0A0V1B1C8
Trichinella sp A0A0V1BUM6
Trichinella sp A0A0V1BY31
Trichinella sp A0A0V1AWB4
Trichinella sp A0A0V1AUR8
Trichinella sp A0A0V1BTZ8
Trichinella sp A0A0V1AZ75
Trichinella sp A0A0V1BF39
Trichinella sp A0A0V0Z9Z1
Trichinella sp A0A0V1B1K9
Trichinella sp A0A0V1BF98
Trichinella sp A0A0V1AUN2
Trichinella sp A0A0V1BG87
Trichinella sp A0A0V1AQK3
Trichinella sp A0A0V1BXK5
Trichinella sp A0A0V1BGU0
Trichinella sp A0A0V1BV17
Trichinella sp A0A0V1BY17
Trichinella sp A1DYF2
Trichinella sp A0A0V1C087
Trichinella sp A0A0V1AW82
Trichinella sp A0A0V1ARD6

Trichinella sp A0A0V1BYW4
Trichinella sp A0A0V1AQ90
Trichinella sp A0A0V1ARB8
Trichinella sp A0A0V1BHN0
Trichinella sp A0A0V1AQK5
Trichinella sp A0A0V1BSR9
Trichinella sp A0A0V1AV98
Trichinella sp E5S9D6
Trichinella sp A0A0V1ASN4
Trichinella sp A0A0V1BZS3
Trichinella sp A0A0V1BYF9
Trichinella sp A0A0V1BY68
Trichinella sp A0A0V1AWE1
Trichinella sp A0A0V1C324
Trichinella sp A0A0V1BV78
Trichinella sp A0A0V1BIN6
Trichinella sp A0A0V1BV40
Trichinella sp A0A0V1BZG3
Trichinella sp A0A0V1AT94
Trichinella sp A0A0V1AZ10
Trichinella sp A0A0V1AUX5
Trichinella sp A0A0V1AU60
Trichinella sp A0A0V1AQI4
Trichinella sp A0A0V1BG93
Trichinella sp A0A0V1BGM2
Trichinella sp A0A0V1BT93
Trichinella sp A0A0V1BYD9
Trichinella sp A0A0V1C2F3
Trichinella sp A0A0V1ALY2
Trichinella sp A0A0V1BY24
Trichinella sp A0A0V1BY86
Trichinella sp A0A0V1BUE7
Trichinella sp E5SVP5
Trichinella sp A0A0V1ARU4
Trichinella sp A0A0V1BMR3
Trichinella sp A0A0V1ARE7
Trichinella zir A0A0V1HCU6
Trichinella zir A0A0V1HT58
Trichinella zir A0A0V1HB44
Trichinella zir A0A0V1HUT8
Trichinella zir A0A0V1I1I8

Trichinella zir A0A0V1HNV1
Trichinella zir A0A0V1GP67
Trichinella zir A0A0V1H9N6
Trichinella zir A0A0V1HMQ6
Trichinella zir A0A0V1H7J5
Trichinella zir A0A0V1I1W3
Trichinella zir A0A0V1HAB7
Trichinella zir A0A0V1HXL0
Trichinella zir A0A0V1HW85
Trichinella zir A0A0V1H4C6
Trichinella zir A0A0V1I5B5
Trichinella zir A0A0V1HR65
Trichinella zir A0A0V1HBV3
Trichinella zir A0A0V1I259
Trichinella zir A0A0V1I3Y0
Trichinella zir A0A0V1HLA0
Trichinella zir A0A0V1HW02
Trichinella zir A0A0V1HLD1
Trichinella zir A0A0V1HVP2
Trichinella zir A0A0V1HLM5
Trichinella zir A0A0V1I5C9
Trichinella zir A0A0V1I289
Trichinella zir A0A0V1HBL5
Trichinella zir A0A0V1H4G7
Trichinella zir A0A0V1H7I9
Trichinella zir A0A0V1HYK5
Trichinella zir A0A0V1I3Y7
Trichinella zir A0A0V1GSV2
Trichinella zir A0A0V1HE40
Trichinella zir A0A0V1H7R5
Trichinella zir A0A0V1I5B7
Trichinella zir A0A0V1HAK7
Trichinella zir A0A0V1HBJ8
Trichinella zir A0A0V1HX30
Trichinella zir A0A0V1HDT3
Trichinella zir A0A0V1I4J6
Trichinella zir A0A0V1H559
Trichinella zir A0A0V1HQU5
Trichinella zir A0A0V1HAU6
Trichinella zir A0A0V1HAX6
Trichinella zir A0A0V1HCF7

| | |
|---|---|
| Trichinella zir | A0A0V1HEP2 |
| Trichinella zir | A0A0V1HCA8 |
| Trichinella zir | A0A0V1HEI2 |
| Trichinella zir | A0A0V1HTQ9 |
| Trichinella zir | A0A0V1H6F1 |
| Trichobilharz | A0A183VXQ2 |
| Trichobilharz | A0A183VLN0 |
| Trichobilharz | A0A3P8HFW6 |
| Trichoderma | A0A395NMG6 |
| Trichoderma | A0A395NGU7 |
| Trichoderma | A0A395NHQ3 |
| Trichoderma | A0A2T3ZPF6 |
| Trichoderma | A0A2T3ZAJ6 |
| Trichoderma | A0A2T3YU56 |
| Trichoderma | A0A2T3YQC3 |
| Trichoderma | A0A2T3YQB8 |
| Trichoderma | A0A2T3ZA54 |
| Trichoderma | A0A2T3YQC0 |
| Trichoderma | A0A2T4B6T4 |
| Trichoderma | A0A2T4B5A6 |
| Trichoderma | A0A2K0TAN5 |
| Trichoderma | A0A1T3CP57 |
| Trichoderma | A0A1T3CIF4 |
| Trichoderma | A0A2T3ZRT3 |
| Trichoderma | A0A2T3ZXU3 |
| Trichoderma | A0A2T4ANA8 |
| Trichoderma | A0A2T3ZX63 |
| Trichoderma | A0A2K0TYA4 |
| Trichoderma | A0A2N1LZN2 |
| Trichoderma | A0A0F9ZNQ8 |
| Trichoderma | A0A2N1L2X0 |
| Trichoderma | A0A2N1LKG2 |
| Trichoderma | A0A2K0U895 |
| Trichoderma | A0A2N1LM61 |
| Trichoderma | A0A0F9X8N3 |
| Trichoderma | A0A2T4CJY3 |
| Trichoderma | A0A2T4CIH3 |
| Trichoderma | A0A2H2Z6Q9 |
| Trichoderma | A0A2H2Z2T5 |
| Trichomalops | A0A232F3J2 |
| Trichomalops | A0A232FIF8 |

| | |
|---|---|
| Trichomalops | A0A232F941 |
| Trichomalops | A0A232F4N8 |
| Trichomalops | A0A232F7T9 |
| Trichophyton | F2SMU2 |
| Trichophyton | A0A022W2V9 |
| Trichophyton | A0A178F2Y8 |
| Trichophyton | A0A022XTQ1 |
| Trichophyton | A0A178FP64 |
| Trichoplax sp | A0A369RWP4 |
| Trichuris mur | A0A0N5E1P0 |
| Trichuris mur | A0A0N5DL77 |
| Trichuris mur | A0A0N5DPD5 |
| Trichuris mur | A0A0N5DH91 |
| Trichuris mur | A0A0N5DXR0 |
| Trichuris mur | A0A0N5DPD6 |
| Trichuris mur | A0A0N5DRN3 |
| Trichuris mur | A0A0N5DYA9 |
| Trichuris mur | A0A0N5DYK6 |
| Trichuris mur | A0A0N5DUR4 |
| Trichuris mur | A0A0N5DEG2 |
| Trichuris mur | A0A0N5DVP1 |
| Trichuris mur | A0A0N5DR95 |
| Trichuris mur | A0A0N5DR76 |
| Trichuris mur | A0A0N5E6W2 |
| Trichuris mur | A0A0N5DHF1 |
| Trichuris mur | A0A0N5DRT2 |
| Trichuris mur | A0A0N5DD95 |
| Trichuris mur | A0A0N5DM82 |
| Trichuris suis | A0A085NT68 |
| Trichuris suis | A0A085N873 |
| Trichuris suis | A0A085MF87 |
| Trichuris suis | A0A085MJD4 |
| Trichuris suis | A0A0B1PQG5 |
| Trichuris suis | A0A085M562 |
| Trichuris suis | A0A085LN72 |
| Trichuris suis | A0A085N482 |
| Trichuris suis | A0A0B1PME0 |
| Trichuris suis | A0A0B1PKA9 |
| Trichuris suis | A0A085MJ81 |
| Trichuris suis | A0A085LWZ9 |
| Trichuris suis | A0A085NTB9 |

| | |
|---|---|
| Trichuris suis | A0A0B1PI26 |
| Trichuris suis | A0A0B1PIN9 |
| Trichuris suis | A0A085MPH9 |
| Trichuris suis | A0A085LXR5 |
| Trichuris suis | A0A085NL76 |
| Trichuris suis | A0A085NGB0 |
| Trichuris suis | A0A085M826 |
| Trichuris suis | A0A0B1PP49 |
| Trichuris suis | A0A085MVC4 |
| Trichuris suis | A0A085M330 |
| Trichuris suis | A0A085LIA0 |
| Trichuris suis | A0A085N5V9 |
| Trichuris suis | A0A085MK20 |
| Trichuris suis | A0A0B1PRQ3 |
| Trichuris suis | A0A085NN81 |
| Trichuris suis | A0A085LIK5 |
| Trichuris suis | A0A085M8H3 |
| Trichuris suis | A0A085MX09 |
| Trichuris suis | A0A085N8R6 |
| Trichuris suis | A0A085M2H7 |
| Trichuris suis | A0A085NKE8 |
| Trichuris suis | A0A085LQT6 |
| Trichuris trich | A0A077YW89 |
| Trichuris trich | A0A077YZL9 |
| Trichuris trich | A0A077Z1E9 |
| Trichuris trich | A0A077ZBA3 |
| Trichuris trich | A0A077Z7J7 |
| Trichuris trich | A0A077Z0B4 |
| Trichuris trich | A0A077ZF09 |
| Trichuris trich | A0A077Z0L9 |
| Trichuris trich | A0A077Z8C3 |
| Trichuris trich | A0A077Z0L5 |
| Trichuris trich | A0A077YVM8 |
| Trichuris trich | A0A077ZAR2 |
| Trichuris trich | A0A077ZB27 |
| Trichuris trich | A0A077Z4C4 |
| Trichuris trich | A0A077ZFL2 |
| Trichuris trich | A0A077YY03 |
| Trichuris trich | A0A077ZGT3 |
| Trichuris trich | A0A077Z244 |
| Trichuris trich | A0A077ZE86 |

| | |
|---|---|
| Trifolium me( | A0A392PT57 |
| Trifolium me( | A0A392M1H5 |
| Trifolium me( | A0A392TWG0 |
| Trifolium me( | A0A392PBX8 |
| Trifolium me( | A0A392N1T7 |
| Trifolium me( | A0A392Q1K1 |
| Trifolium me( | A0A392NG93 |
| Trifolium me( | A0A392NDB2 |
| Trifolium me( | A0A392RWA2 |
| Trifolium me( | A0A392NAQ6 |
| Trifolium me( | A0A392NVW7 |
| Trifolium me( | A0A392PE82 |
| Trifolium me( | A0A392PGQ4 |
| Trifolium me( | A0A392N2X5 |
| Trifolium me( | A0A392PG95 |
| Trifolium me( | A0A392N4E7 |
| Trifolium me( | A0A392PA83 |
| Trifolium me( | A0A392NWT8 |
| Trifolium me( | A0A392NQL4 |
| Trifolium me( | A0A392MBY4 |
| Trifolium me( | A0A392Q8A5 |
| Trifolium me( | A0A392PMK5 |
| Trifolium me( | A0A392MI51 |
| Trifolium me( | A0A392NQF2 |
| Trifolium me( | A0A392Q3T9 |
| Trifolium me( | A0A392NX73 |
| Trifolium me( | A0A392N067 |
| Trifolium me( | A0A392P4S9 |
| Trifolium me( | A0A392NR91 |
| Trifolium me( | A0A392MH64 |
| Trifolium me( | A0A392NFX7 |
| Trifolium pra | A0A2K3N545 |
| Trifolium pra | A0A2K3NIL3 |
| Trifolium pra | A0A2K3NQL2 |
| Trifolium pra | A0A2K3PPK5 |
| Trifolium pra | A0A2K3PAC8 |
| Trifolium pra | A0A2K3N5G0 |
| Trifolium pra | A0A2K3PGR8 |
| Trifolium pra | A0A2K3PAJ7 |
| Trifolium pra | A0A2K3PNI2 |
| Trifolium pra | A0A2K3L2K4 |

| | |
|---|---|
| Trifolium pra | A0A2K3NKF3 |
| Trifolium pra | A0A2K3MWB2 |
| Trifolium pra | A0A2K3MNA4 |
| Trifolium pra | A0A2K3PFA1 |
| Trifolium pra | A0A2K3LYQ1 |
| Trifolium pra | A0A2K3NPR5 |
| Trifolium pra | A0A2K3PKJ0 |
| Trifolium pra | A0A2K3KY21 |
| Trifolium pra | A0A2K3P821 |
| Trifolium pra | A0A2K3N0U2 |
| Trifolium pra | A0A2K3NIX8 |
| Trifolium pra | A0A2K3PCX5 |
| Trifolium pra | A0A2K3MTX0 |
| Trifolium pra | A0A2K3P0N9 |
| Trifolium pra | A0A2K3K2X8 |
| Trifolium pra | A0A2K3K1G0 |
| Trifolium pra | A0A2K3P4N6 |
| Trifolium pra | A0A2K3MMY0 |
| Trifolium pra | A0A2K3N390 |
| Trifolium pra | A0A2K3PRU0 |
| Trifolium pra | A0A2K3NHE9 |
| Trifolium pra | A0A2K3MP42 |
| Trifolium pra | A0A2K3L7A3 |
| Trifolium pra | A0A2K3KDT6 |
| Trifolium pra | A0A2K3L793 |
| Trifolium pra | A0A2K3NJM8 |
| Trifolium pra | A0A2K3NHS7 |
| Trifolium pra | A0A2K3M7W6 |
| Trifolium pra | A0A2K3P4W6 |
| Trifolium pra | A0A2K3NNI7 |
| Trifolium pra | A0A2K3L7X0 |
| Trifolium pra | A0A2K3LKJ0 |
| Trifolium pra | A0A2K3MQU1 |
| Trifolium pra | A0A2K3JN77 |
| Trifolium pra | A0A2K3PRC7 |
| Trifolium pra | A0A2K3LSP2 |
| Trifolium pra | A0A2K3LQR6 |
| Trifolium pra | A0A2K3NAV2 |
| Trifolium pra | A0A2K3N5F0 |
| Trifolium pra | A0A2K3P2Q7 |
| Trifolium pra | A0A2K3NIV7 |

| | |
|---|---|
| Trifolium pra | A0A2K3MTU5 |
| Trifolium pra | A0A2K3MZS0 |
| Trifolium pra | A0A2K3JM43 |
| Trifolium pra | A0A2K3LPU5 |
| Trifolium pra | A0A2K3N4M8 |
| Trifolium pra | A0A2K3PJS0 |
| Trifolium pra | A0A2K3NB85 |
| Trifolium pra | A0A2K3M7I7 |
| Trifolium pra | A0A2K3NDE7 |
| Trifolium pra | A0A2K3KSU5 |
| Trifolium pra | A0A2K3PLE0 |
| Trifolium pra | A0A2K3M3U1 |
| Trifolium pra | A0A2K3L147 |
| Trifolium pra | A0A2K3P033 |
| Trifolium pra | A0A2K3LVH3 |
| Trifolium pra | A0A2K3N391 |
| Trifolium pra | A0A2K3NHE1 |
| Trifolium pra | A0A2K3NPS4 |
| Trifolium pra | A0A2K3L0P8 |
| Trifolium pra | A0A2K3MRR9 |
| Trifolium pra | A0A2K3P8M4 |
| Trifolium pra | A0A2K3M9G4 |
| Trifolium pra | A0A2K3JVE5 |
| Trifolium pra | A0A2K3LWR5 |
| Trifolium pra | A0A2K3KZ94 |
| Trifolium pra | A0A2K3KSU6 |
| Trifolium pra | A0A2K3LB77 |
| Trifolium pra | A0A2K3NP95 |
| Trifolium pra | A0A2K3N177 |
| Trifolium pra | A0A2K3P0P9 |
| Trifolium pra | A0A2K3LRR4 |
| Trifolium pra | A0A2K3PL10 |
| Trifolium pra | A0A2K3PJV4 |
| Trifolium pra | A0A2K3NSR0 |
| Trifolium pra | A0A2K3KLD8 |
| Trifolium pra | A0A2K3LCR5 |
| Trifolium pra | A0A2K3NTH2 |
| Trifolium pra | A0A2K3PKC0 |
| Trifolium pra | A0A2K3MMJ1 |
| Trifolium sub | A0A2Z6LMC9 |
| Trifolium sub | A0A2Z6MW96 |

Trifolium sub A0A2Z6M7P9
Trifolium sub A0A2Z6NDV6
Trifolium sub A0A2Z6MC05
Trifolium sub A0A2Z6MAQ7
Trifolium sub A0A2Z6N0I3
Trifolium sub A0A2Z6LQ21
Trifolium sub A0A2Z6MXN7
Trifolium sub A0A2Z6M5F5
Trifolium sub A0A2Z6MXE3
Trifolium sub A0A2Z6P679
Trifolium sub A0A2Z6LR55
Trifolium sub A0A2Z6MQ73
Trifolium sub A0A2Z6MDJ9
Trifolium sub A0A2Z6NUW8
Trifolium sub A0A2Z6MKZ5
Trifolium sub A0A2Z6NWU0
Trifolium sub A0A2Z6ME77
Trifolium sub A0A2Z6MPN7
Trifolium sub A0A2Z6NJ42
Trifolium sub A0A2Z6LL70
Trifolium sub A0A2Z6LM12
Trifolium sub A0A2Z6P1H7
Trifolium sub A0A2Z6LMM9
Trifolium sub A0A2Z6MPK7
Trifolium sub A0A2Z6N5W6
Trifolium sub A0A2Z6MAP8
Trifolium sub A0A2Z6N1K5
Trifolium sub A0A2Z6NTN7
Trifolium sub A0A2Z6LQG0
Trifolium sub A0A2Z6LQW7
Trifolium sub A0A2Z6LPI7
Trifolium sub A0A2Z6MQ16
Trifolium sub A0A2Z6MDD8
Trifolium sub A0A2Z6P268
Trifolium sub A0A2Z6LPS2
Trifolium sub A0A2Z6P3R5
Trifolium sub A0A2Z6MYS4
Trifolium sub A0A2Z6LPQ7
Trifolium sub A0A2Z6NJK2
Trifolium sub A0A2Z6P441
Trifolium sub A0A2Z6NAG1

Trifolium sub A0A2Z6MMM3
Trifolium sub A0A2Z6M4K0
Trifolium sub A0A2Z6MVN2
Trifolium sub A0A2Z6LPR4
Trifolium sub A0A2Z6LTH3
Trifolium sub A0A2Z6PUS7
Trifolium sub A0A2Z6MQE3
Trifolium sub A0A2Z6NF42
Trifolium sub A0A2Z6MF49
Trifolium sub A0A2Z6MQ83
Trifolium sub A0A2Z6MRQ0
Trifolium sub A0A2Z6MFP6
Trifolium sub A0A2Z6NXT5
Trifolium sub A0A2Z6N7M7
Trifolium sub A0A2Z6P962
Trifolium sub A0A2Z6LPY4
Trifolium sub A0A2Z6NL64
Trifolium sub A0A2Z6M8M8
Trifolium sub A0A2Z6P518
Trifolium sub A0A2Z6P2F4
Trifolium sub A0A2Z6MYB5
Trifolium sub A0A2Z6LTQ0
Trifolium sub A0A2Z6LV78
Trifolium sub A0A2Z6MVE1
Trifolium sub A0A2Z6LKE4
Trifolium sub A0A2Z6PDH2
Trifolium sub A0A2Z6M528
Trifolium sub A0A2Z6MVF6
Triticum aest A0A3B6NPP9
Triticum aest A0A3B6IPA0
Triticum aest A0A3B6GXA5
Triticum aest A0A3B6CFN8
Triticum aest A0A3B6GVI4
Triticum aest A0A3B6AVA7
Triticum aest A0A3B6ITK6
Triticum aest A0A3B6DM60
Triticum aest A0A3B6AVU8
Triticum aest A0A3B6EJV5
Triticum aest A0A3B6DAX4
Triticum aest A0A3B6I2Q0
Triticum aest A0A3B6FV60

Triticum aest A0A077RVZ3
Triticum aest A0A3B6GWT1
Triticum aest A0A3B6FV65
Triticum aest A0A3B5Y188
Triticum aest A0A3B6FQR7
Triticum aest A0A3B5Y1P0
Triticum aest A0A3B6CIE0
Triticum aest A0A3B5YYY7
Triticum aest A0A3B6GVI9
Triticum aest A0A3B6DAD8
Triticum aest A0A3B6C3X4
Triticum aest A0A3B6MW88
Triticum aest A0A3B6IKZ5
Triticum aest A0A3B6B802
Triticum aest A0A3B6C476
Triticum aest A0A3B5XTG1
Triticum aest A0A3B6CDR2
Triticum aest A0A3B6CFN2
Triticum aest A0A3B6DQD8
Triticum aest A0A3B6ML17
Triticum aest A0A3B6D535
Triticum aest A0A3B6TPW9
Triticum aest A0A3B6DN59
Triticum aest A0A3B6HWP0
Triticum aest A0A3B5Z3I2
Triticum aest A0A3B6KT61
Triticum aest A0A3B6FWV1
Triticum aest A0A3B5ZXT0
Triticum aest A0A3B6SIG2
Triticum aest A0A3B6CFN4
Triticum aest A0A3B6DQC5
Triticum aest A0A2X0SW72
Triticum aest A0A3B6FER1
Triticum aest A0A3B6CFL5
Triticum aest A0A3B6CDW1
Triticum aest A0A3B6QD29
Triticum aest A0A3B6U554
Triticum aest A0A3B6MWW1
Triticum aest A0A3B6B6Z0
Triticum aest A0A3B6IXZ8
Triticum aest A0A3B6C118

Triticum aest A0A3B6MWI7
Triticum aest A0A3B6RA84
Triticum aest A0A3B6DM95
Triticum aest A0A3B6KNX5
Triticum aest A0A3B5XTI8
Triticum aest A0A3B6RP60
Triticum aest A0A0R8CAF3
Triticum aest A0A3B6B844
Triticum aest A0A3B5YZH3
Triticum aest A0A3B5ZZY7
Triticum aest A0A3B5ZYM3
Triticum aest A0A3B6DM69
Triticum aest A0A3B6D841
Triticum aest A0A3B6IWH6
Triticum aest A0A3B6NXU2
Triticum aest A0A3B6D593
Triticum aest A0A3B6C633
Triticum aest A0A3B6UAV8
Triticum aest A0A3B6CDW2
Triticum aest A0A3B6RDT9
Triticum aest A0A3B6CDV2
Triticum aest A0A3B6LUF7
Triticum aest A0A3B6DQ92
Triticum aest A0A3B6SL80
Triticum aest A0A3B6N0N1
Triticum aest A0A3B6DLD6
Triticum aest A0A2X0SND0
Triticum aest A0A3B5ZZF6
Triticum aest A0A3B6B2F3
Triticum aest A0A3B6MYX9
Triticum aest A0A3B5Z1X9
Triticum aest A0A3B6DMT7
Triticum aest A0A3B6KN82
Triticum aest A0A3B6NWS4
Triticum aest A0A3B5XT21
Triticum aest A0A3B6C8K8
Triticum aest A0A3B6DMB1
Triticum aest A0A3B6JIP6
Triticum aest A0A3B6G0F8
Triticum aest A0A3B6CFH5
Triticum aest A0A3B6C728

Triticum aest A0A3B6QP32
Triticum aest A0A3B6RMY0
Triticum aest A0A3B6DML0
Triticum aest A0A3B6RPD1
Triticum aest A0A3B6SA65
Triticum aest A0A3B6DQD5
Triticum aest A0A3B6MP47
Triticum aest A0A3B6DL71
Triticum aest A0A3B6IV36
Triticum aest A0A3B6DJM6
Triticum aest A0A3B6MWH8
Triticum aest A0A3B6B8U6
Triticum aest A0A3B6TNK8
Triticum aest A0A3B6MWJ6
Triticum aest A0A3B6CFR7
Triticum aest A0A3B6GPP5
Triticum aest A0A2X0TGT9
Triticum aest A0A2X0S0A4
Triticum aest A0A3B6B6Z5
Triticum aest A0A3B6AR12
Triticum aest A0A3B6DDD0
Triticum aest A0A3B6TFJ0
Triticum aest A0A3B6CI24
Triticum aest A0A3B6TPC6
Triticum aest A0A3B6B568
Triticum aest A0A3B6IWJ1
Triticum aest A0A3B6SEP2
Triticum aest A0A3B6KLI6
Triticum aest A0A3B6EKY9
Triticum aest A0A3B6CG16
Triticum aest A0A3B6B8T6
Triticum aest A0A3B6RPA4
Triticum aest A0A3B6TT98
Triticum aest A0A3B6H304
Triticum aest A0A3B6EH46
Triticum aest A0A3B6FUN9
Triticum aest A0A3B6CFS0
Triticum aest A0A3B6KRY5
Triticum aest A0A3B6B571
Triticum aest A0A3B6TQ03
Triticum aest A0A3B6GVH4

Triticum aest A0A3B6CDR7
Triticum aest A0A3B6I3N0
Triticum aest A0A3B6B7D6
Triticum aest A0A3B6CFM9
Triticum aest A0A3B6LRY8
Triticum aest A0A3B6DJM2
Triticum aest A0A3B6MV81
Triticum aest A0A3B5ZNL3
Triticum aest A0A3B6A183
Triticum aest A0A3B6QP21
Triticum aest A0A3B6BZH0
Triticum aest A0A3B6J0A9
Triticum aest A0A3B6B640
Triticum aest A0A3B6DL82
Triticum aest A0A3B6QNH4
Triticum aest A0A3B6U4W3
Triticum aest A0A3B5Z2P0
Triticum aest A0A3B6CDX1
Triticum aest A0A3B6JP90
Triticum aest A0A3B6S8I2
Triticum aest A0A3B6MV89
Triticum aest A0A3B6LXS1
Triticum aest A0A3B6QP12
Triticum aest A0A3B6B6T4
Triticum aest A0A3B5XT90
Triticum aest A0A3B6DQE1
Triticum aest A0A3B6CH00
Triticum aest A0A3B6KN77
Triticum aest A0A077RVZ1
Triticum aest A0A3B5Z285
Triticum aest A0A3B6AZK9
Triticum aest A0A3B6B840
Triticum aest A0A3B6CFZ6
Triticum aest A0A3B6SBZ7
Triticum aest A0A3B6PVE3
Triticum aest A0A3B6LDI3
Triticum aest A0A3B6N2M7
Triticum aest A0A3B6C6J6
Triticum aest A0A3B6DN39
Triticum aest A0A3B6DLB0
Triticum aest A0A3B6B710

Triticum aest A0A3B6DN68
Triticum aest A0A3B6LQP4
Triticum aest A0A3B5XVZ4
Triticum aest A0A3B6TQM2
Triticum aest A0A3B6I168
Triticum aest A0A3B6CFM3
Triticum aest A0A3B6LXQ4
Triticum aest A0A3B6CDS2
Triticum aest A0A3B5Z158
Triticum aest A0A3B6TXD5
Triticum aest A0A3B6UAM6
Triticum aest A0A2X0SCN5
Triticum aest A0A3B6UC57
Triticum aest A0A3B6SRM5
Triticum aest A0A3B6TW06
Triticum aest A0A2X0SNF0
Triticum aest A0A3B6MII3
Triticum aest A0A3B6CEN4
Triticum aest A0A3B6DJN6
Triticum aest A0A3B6B7S8
Triticum aest A0A3B6KNC2
Triticum aest A0A3B6CDU3
Triticum aest A0A3B6C714
Triticum aest A0A3B6MXU5
Triticum aest A0A3B6QKM8
Triticum aest A0A3B6FN69
Triticum aest A0A3B5ZME3
Triticum aest A0A3B6N370
Triticum aest A0A3B6B8R0
Triticum aest A0A3B6LRZ5
Triticum aest A0A3B6LY19
Triticum aest A0A3B6B6X0
Triticum aest A0A3B6GTI5
Triticum aest A0A3B6HX59
Triticum aest A0A3B6LR65
Triticum aest A0A3B6KLI0
Triticum aest A0A3B6B7J7
Triticum aest A0A3B5Y5C6
Triticum aest A0A3B6SDH9
Triticum aest A0A3B6SPZ5
Triticum aest A0A3B6D4X4

Triticum aest A0A3B6DQM1
Triticum aest A0A3B6DDV3
Triticum aest A0A3B6TAX9
Triticum aest A0A3B6KKP1
Triticum aest A0A3B6AVA1
Triticum aest A0A3B6DLC1
Triticum aest A0A3B6EKT7
Triticum aest A0A3B6SQQ6
Triticum aest A0A3B6C001
Triticum aest A0A3B6H0E6
Triticum aest A0A3B6LEV2
Triticum aest A0A3B6LS79
Triticum aest A0A3B6B644
Triticum aest A0A3B6DE47
Triticum aest A0A3B6DL94
Triticum aest A0A2X0SNW6
Triticum aest A0A3B6AX84
Triticum aest A0A3B6BXA3
Triticum aest A0A3B6TT93
Triticum aest A0A3B5XUG7
Triticum aest A0A3B6PV04
Triticum aest A0A3B6CDW7
Triticum aest A0A3B6ARF6
Triticum aest A0A3B6B7Q6
Triticum aest A0A3B6A030
Triticum aest A0A3B6JCZ0
Triticum aest A0A3B6KKN0
Triticum aest A0A3B5Y464
Triticum aest A0A3B6B7H2
Triticum aest A0A3B6CDT7
Triticum aest A0A3B5Z410
Triticum aest A0A3B6BZG5
Triticum aest A0A3B6CFP5
Triticum aest A0A3B6SKQ8
Triticum aest A0A3B6G1T3
Triticum aest A0A3B6LP90
Triticum aest A0A3B6MPM0
Triticum aest A0A3B5ZQN2
Triticum aest A0A3B6MWW7
Triticum aest A0A3B6FI27
Triticum aest A0A3B6GSI9

Triticum aest A0A3B5Z165
Triticum aest A0A3B6CEJ7
Triticum aest A0A3B6H694
Triticum aest A0A3B5Z1E9
Triticum aest A0A3B6CFP7
Triticum aest A0A3B6HT43
Triticum aest A0A3B6RSP0
Triticum aest A0A3B5YT59
Triticum aest A0A3B6CFN5
Triticum aest A0A3B6KNX0
Triticum aest A0A3B6TK76
Triticum aest A0A3B6SIF3
Triticum aest A0A3B6PST4
Triticum aest A0A3B6CC02
Triticum aest A0A3B6TUP1
Triticum aest A0A3B6B0K5
Triticum aest A0A3B6CHY5
Triticum aest A0A3B6B7R1
Triticum aest A0A3B6B826
Triticum aest A0A1D5UGG3
Triticum aest A0A3B5ZZV1
Triticum aest A0A3B6B022
Triticum aest A0A3B6KD22
Triticum aest A0A3B6KRL2
Triticum aest A0A3B6U115
Triticum aest A0A3B6GNZ2
Triticum aest A0A3B6JDU7
Triticum aest A0A3B5Y527
Triticum aest A0A2X0SBL5
Triticum aest A0A3B6CDT0
Triticum aest A0A3B6C3Z6
Triticum aest A0A3B5Y3Q9
Triticum aest A0A3B6QL00
Triticum aest A0A3B6SJY6
Triticum aest A0A3B6ESF4
Triticum aest A0A3B6DJP0
Triticum aest A0A3B6ER67
Triticum aest A0A3B6RJV3
Triticum aest A0A3B6CFJ4
Triticum aest A0A3B5Y5D0
Triticum aest A0A3B6CDT5

Triticum aest A0A3B6JF83
Triticum aest A0A3B6SDH4
Triticum aest A0A3B6DL99
Triticum aest A0A3B6AQB5
Triticum aest A0A3B6H388
Triticum aest A0A3B6PSS9
Triticum aest A0A3B6FRH5
Triticum aest A0A3B6NSZ0
Triticum aest A0A3B6CFN7
Triticum aest A0A3B6C3W9
Triticum aest A0A3B6ENW4
Triticum aest A0A3B6B7N8
Triticum aest A0A3B6DGA7
Triticum aest A0A3B6D4L4
Triticum aest A0A3B6MXW0
Triticum aest A0A2X0SNE0
Triticum aest A0A3B6DNS5
Triticum aest A0A3B6B820
Triticum aest A0A3B6DL80
Triticum aest A0A3B6LS69
Triticum aest A0A3B6CI19
Triticum aest A0A3B6TNR8
Triticum aest A0A3B5ZVB3
Triticum aest A0A3B6R9Q9
Triticum aest A0A3B6CG09
Triticum aest A0A3B5ZME6
Triticum aest A0A3B6IMW8
Triticum aest A0A3B6SLK7
Triticum aest A0A3B6H2G6
Triticum aest A0A3B6KNM0
Triticum aest A0A3B6DAE3
Triticum aest A0A3B6SLZ3
Triticum aest A0A3B6IR46
Triticum aest A0A3B6CDX8
Triticum aest A0A3B6RMR3
Triticum aest A0A3B6JK14
Triticum aest A0A3B6LP53
Triticum aest A0A3B6SQQ3
Triticum aest A0A3B6LSC8
Triticum aest R9UC29
Triticum aest A0A2X0SUV4

Triticum aest A0A3B6B733
Triticum aest A0A3B6GZL0
Triticum aest A0A2X0SCV8
Triticum aest A0A3B6BXQ7
Triticum aest A0A3B6RPT7
Triticum aest Q56DH9
Triticum aest A0A3B6TE76
Triticum aest A0A3B6SNL4
Triticum aest A0A3B6B7Z9
Triticum aest A0A3B6IZM9
Triticum aest A0A3B6CG90
Triticum aest A0A3B6TPR0
Triticum aest A0A3B6AWA3
Triticum aest A0A3B6B7V1
Triticum aest I0JTU1
Triticum aest A0A3B6GWS6
Triticum aest A0A3B5Z166
Triticum aest A0A3B6EEA5
Triticum aest D8L9S5
Triticum aest A0A3B6AYR6
Triticum aest A0A3B6CFM6
Triticum aest A0A3B6SHS1
Triticum aest A0A3B6KSI0
Triticum aest A0A3B6CFN3
Triticum aest A0A3B6NLJ3
Triticum aest A0A3B6CFN9
Triticum aest A0A3B5Z2J0
Triticum aest A0A2X0SBH9
Triticum aest A0A2X0U0Z3
Triticum aest A0A3B6MXW5
Triticum aest A0A3B6LQP0
Triticum aest A0A3B6JI87
Triticum aest A0A3B6REN4
Triticum aest A0A3B6RMR6
Triticum aest A0A3B6BZR8
Triticum aest A0A3B6B5R5
Triticum aest A0A3B6CI12
Triticum aest A0A3B6DHA3
Triticum aest A0A3B6KAL1
Triticum aest A0A3B6B715
Triticum aest A0A2X0SPL7

Triticum aest A0A3B5ZMU9
Triticum aest A0A3B6H6M7
Triticum aest A0A3B6AWQ0
Triticum aest A0A3B6TF32
Triticum aest A0A3B6HPJ6
Triticum aest A0A3B6RTD5
Triticum aest A0A3B6DKA3
Triticum aest A0A3B6LJM9
Triticum aest A0A3B6ESW5
Triticum aest A0A3B6DM74
Triticum aest A0A3B6TW12
Triticum aest A0A3B6B6D0
Triticum aest A0A3B6LRZ0
Triticum aest A0A3B6MXV5
Triticum aest A0A3B6RB18
Triticum aest A0A3B6NWL7
Triticum aest A0A3B6AVX3
Triticum aest A0A3B6C9J4
Triticum aest A0A3B6SQF7
Triticum aest A0A2X0U1E9
Triticum aest A0A3B6B7V7
Triticum aest A0A3B6RPG4
Triticum aest A0A3B6CF85
Triticum aest A0A3B6D812
Triticum aest A0A3B6LM48
Triticum aest A0A3B6TLB6
Triticum aest A0A3B6AVW3
Triticum aest A0A3B6EHS5
Triticum aest A0A3B6IK35
Triticum aest A0A3B6B8F8
Triticum aest A0A2X0SCT9
Triticum aest A0A3B6U7R0
Triticum aest A0A3B5ZP26
Triticum aest A0A3B5XTI4
Triticum aest A0A3B6U6Q9
Triticum aest A0A3B5ZRQ8
Triticum aest A0A3B6DPF6
Triticum aest A0A3B6RR09
Triticum aest R9UFG9
Triticum aest A0A3B6QL55
Triticum aest A0A3B6EM35

Triticum aest A0A3B6DAS8
Triticum aest A0A3B6EM42
Triticum aest A0A3B6EJL4
Triticum aest A0A3B6B738
Triticum aest A0A3B6FRT0
Triticum aest A0A3B6GQZ4
Triticum aest A0A3B6NVE3
Triticum aest A0A3B5Y4W4
Triticum aest A0A3B6HT50
Triticum aest A0A3B6RQ82
Triticum aest A0A3B6JK06
Triticum aest A0A2X0SCX3
Triticum aest A0A3B6DAS6
Triticum aest A0A3B6B0L1
Triticum aest A0A3B6U6W6
Triticum aest A0A3B6PSW1
Triticum aest A0A3B6MI22
Triticum aest A0A3B6EM39
Triticum aest A0A3B6FUK0
Triticum aest A0A3B6EJW4
Triticum aest A0A3B6C439
Triticum aest A0A3B5ZYD7
Triticum aest A0A3B6FUJ5
Triticum aest A0A3B6C3V9
Triticum aest A0A3B6DAW9
Triticum aest A0A3B6C3Y1
Triticum aest A0A3B6H012
Triticum aest A0A3B6NIA4
Triticum aest A0A2X0RZC4
Triticum aest A0A3B6DE52
Triticum aest A0A3B5ZYE5
Triticum aest A0A2X0U219
Triticum aest A0A3B6EH34
Triticum aest A0A3B6AVB1
Triticum aest A0A3B6GXA9
Triticum aest A0A3B6EJV9
Triticum aest A0A3B6DN34
Triticum aest A0A3B6KN67
Triticum aest A0A3B6C6J2
Triticum aest A0A3B6AX65
Triticum aest A0A3B6AX70

Triticum aest A0A3B6A137
Triticum aest A0A3B6CFK5
Triticum aest A0A3B6FRS5
Triticum aest A0A3B6NX53
Triticum aest A0A3B6DKK6
Triticum aest A0A3B6QCT9
Triticum aest A0A3B6LXE2
Triticum aest A0A3B6GXB4
Triticum aest A0A3B6C471
Triticum aest A0A3B6AWQ3
Triticum aest A0A3B6DNI2
Triticum aest A0A3B6A1F9
Triticum aest A0A3B6AVW8
Triticum aest A0A3B6EJK8
Triticum aest A0A3B6FUK5
Triticum aest A0A3B6JE30
Triticum aest A0A3B6GWT6
Triticum aest A0A3B6CDY1
Triticum aest A0A3B6DKS5
Triticum aest A0A3B6AWA8
Triticum aest A0A3B6C434
Triticum aest A0A3B6GVH9
Triticum aest A0A3B6B5S4
Triticum aest A0A3B6DLD1
Triticum aest A0A3B5XW78
Triticum aest A0A3B6H0F8
Triticum aest A0A3B6N1N1
Triticum aest A0A3B6AZ76
Triticum aest A0A3B6MWV6
Triticum aest A0A3B5ZZF3
Triticum aest A0A3B6CG64
Triticum aest A0A3B6H5X7
Triticum aest A0A2X0SDG7
Triticum aest A0A3B6TPX5
Triticum aest A0A3B6BXA2
Triticum aest A0A3B5XTC5
Triticum aest A0A3B6CFT5
Triticum aest A0A3B6DM84
Triticum aest A0A3B6BWH3
Triticum aest A0A3B6HTC9
Triticum aest A0A3B6PUX9

Triticum aest A0A3B5YQ73
Triticum aest A0A3B6DBG8
Triticum aest A0A3B6SK28
Triticum aest A0A3B6MWX7
Triticum aest A0A3B6CDU5
Triticum aest A0A3B6DMH8
Triticum aest A0A3B6LUS0
Triticum aest A0A3B6DM89
Triticum aest A0A3B6LRZ3
Triticum aest A0A3B6DJM9
Triticum aest A0A3B5Z2V7
Triticum aest A0A3B6B883
Triticum aest A0A3B6B6C5
Triticum aest A0A3B5Z2G7
Triticum aest A0A3B6B8V7
Triticum aest A0A3B6DMH3
Triticum aest A0A3B6EF95
Triticum aest A0A3B5YQF7
Triticum aest A0A3B6CHZ0
Triticum aest A0A3B5XWA5
Triticum aest A0A3B5XTJ9
Triticum aest A0A3B6LRZ4
Triticum aest A0A3B6HYQ9
Triticum aest A0A3B6RP18
Triticum aest A0A3B6GZZ9
Triticum aest A0A3B6AQC9
Triticum aest A0A3B6B980
Triticum aest A0A3B6LRX9
Triticum aest A0A3B6DLC6
Triticum aest A0A3B6NXN1
Triticum aest A0A3B6SLK3
Triticum aest A0A3B6B7T6
Triticum aest A0A2X0SCN7
Triticum aest A0A3B6CDX6
Triticum aest A0A3B6DMA2
Triticum aest A0A3B6B036
Triticum aest A0A3B6B576
Triticum aest A0A3B6C3Z0
Triticum aest A0A3B6ASJ6
Triticum aest A0A3B6HW93
Triticum aest A0A2X0RZB1

Triticum aest A0A3B6CFS5
Triticum aest A0A3B6DBG3
Triticum aest A0A3B6KSQ6
Triticum aest A0A3B6C6A4
Triticum aest A0A2X0TS56
Triticum aest A0A3B6PLI1
Triticum aest A0A3B6B719
Triticum aest A0A3B6CFR6
Triticum aest A0A3B6H004
Triticum aest A0A3B6C787
Triticum aest A0A2X0TYQ5
Triticum aest A0A3B5ZXN5
Triticum aest A0A3B6CG60
Triticum aest A0A3B6DLA4
Triticum aest A0A3B6DQ26
Triticum aest A0A3B6SF92
Triticum aest A0A2X0SBJ0
Triticum aest A0A3B6DLN1
Triticum aest A0A3B6E8J0
Triticum aest A0A3B6GLR2
Triticum aest A0A3B6KUH0
Triticum aest A0A3B5YQG9
Triticum aest A0A3B6CFK3
Triticum aest A0A3B6B8P2
Triticum aest A0A3B6DMG8
Triticum aest A0A2X0SBA4
Triticum aest A0A3B6TUB4
Triticum aest A0A3B6EDF4
Triticum aest A0A3B6RPB1
Triticum aest A0A3B6DMF8
Triticum aest A0A3B6UCL0
Triticum aest A0A3B6PUV2
Triticum aest A0A3B6H0E9
Triticum aest A0A3B5Y7C0
Triticum aest A0A3B6GN82
Triticum aest A0A3B6AX75
Triticum aest A0A3B6LQN7
Triticum aest A0A3B6DLB6
Triticum aest A0A3B6DKS9
Triticum aest A0A3B6LRY4
Triticum aest A0A3B6DKI6

Triticum aest A0A3B6GX38
Triticum aest A0A3B5ZMG1
Triticum aest A0A3B5Z1I1
Triticum aest A0A2X0SCS8
Triticum aest A0A3B6TI56
Triticum aest A0A3B6CFI4
Triticum aest A0A3B6AQU8
Triticum aest A0A3B6C782
Triticum aest A0A3B5ZZT3
Triticum aest A0A3B6KL84
Triticum aest A0A3B6RL94
Triticum aest A0A3B6TQ21
Triticum aest A0A3B6LRY9
Triticum aest A0A3B6QN56
Triticum aest A0A3B5YRB3
Triticum aest A0A3B6UAH4
Triticum aest A0A3B6AR71
Triticum aest A0A3B6KED9
Triticum aest A0A3B6TWJ5
Triticum aest A0A3B6ERL9
Triticum aest A0A3B6B6E3
Triticum aest A0A3B6H1E1
Triticum aest A0A3B6DN47
Triticum aest A0A3B6FXS0
Triticum aest A0A3B6MX31
Triticum aest A0A3B6FMU4
Triticum aest A0A3B6APH5
Triticum aest A0A2X0RZY7
Triticum aest A0A3B6DNR6
Triticum aest A0A3B6RL87
Triticum aest A0A3B6TMT8
Triticum aest A0A3B6RQ86
Triticum aest A0A3B6EJJ5
Triticum aest A0A3B6B232
Triticum aest A0A3B6RQ77
Triticum aest A0A2X0TMP3
Triticum aest A0A1D6RPH0
Triticum aest A0A2X0SCP5
Triticum aest A0A1D5UXF1
Triticum aest A0A077S3W5
Triticum aest W5D450

Triticum aest A0A1D5UN73
Triticum aest W5C0L4
Triticum aest A0A1D5UXD7
Triticum aest A0A077RQ16
Triticum aest A0A1D5UVZ2
Triticum aest A0A077RZC2
Triticum aest A0A077RQQ4
Triticum aest A0A077RFI1
Triticum aest A0A341ZF31
Triticum aest A0A1D5UKP6
Triticum aest A0A077S2M2
Triticum aest A0A1D5UK73
Triticum aest A0A1D5UY85
Triticum aest A0A077S0P5
Triticum turg A0A446MVQ3
Triticum turg A0A446LHZ4
Triticum turg A0A446REL1
Triticum turg A0A446MUR9
Triticum turg A0A446UJ83
Triticum turg A0A446YSZ1
Triticum turg A0A446TC95
Triticum turg A0A446SLK3
Triticum turg A0A446LI13
Triticum turg A0A446MV11
Triticum turg A0A446UJ84
Triticum turg A0A446NKA8
Triticum turg A0A446Q0W6
Triticum turg A0A446VU93
Triticum turg A0A446Y561
Triticum turg A0A446NRX1
Triticum turg A0A446SJZ5
Triticum turg A0A446L3Y0
Triticum turg A0A446MRJ4
Triticum turg A0A446KV21
Triticum turg A0A446LHU9
Triticum turg A0A446MRR5
Triticum turg A0A446PDR2
Triticum turg A0A446Q0X4
Triticum turg A0A446TCN2
Triticum turg A0A446LI10
Triticum turg A0A446MVF3

Triticum turg A0A446LHZ0
Triticum turg A0A446PK86
Triticum turg A0A446MWF9
Triticum turg A0A446K644
Triticum turg A0A446VWF4
Triticum turg A0A446MWD8
Triticum turg A0A446M560
Triticum turg A0A446W8K1
Triticum turg A0A446YTG2
Triticum turg A0A446LLE9
Triticum turg A0A446LLA9
Triticum turg A0A446MV52
Triticum turg A0A446MRN3
Triticum turg A0A446LHW9
Triticum turg A0A446NRW8
Triticum turg A0A446RQK6
Triticum turg A0A446RQM9
Triticum turg A0A446M588
Triticum turg A0A446UJ35
Triticum turg A0A446UJ06
Triticum turg A0A446VSN4
Triticum turg A0A446UJB3
Triticum turg A0A446LMD3
Triticum turg A0A446S8Q2
Triticum turg A0A446LI11
Triticum turg A0A446LHV2
Triticum turg A0A446REN0
Triticum turg A0A446YTE0
Triticum turg A0A446MRL7
Triticum turg A0A446LHV8
Triticum turg A0A446MRL2
Triticum turg A0A446UJ76
Triticum turg A0A446TC94
Triticum turg A0A446IP34
Triticum turg A0A446NRW5
Triticum turg A0A446XSM0
Triticum turg A0A446TCA5
Triticum turg A0A446KA03
Triticum turg A0A446YTD1
Triticum turg A0A446MW57
Triticum turg A0A446M577

Triticum turg A0A446UJ96
Triticum turg A0A446K654
Triticum turg A0A446LLJ7
Triticum turg A0A446K022
Triticum turg A0A446WQH4
Triticum turg A0A446KP30
Triticum turg A0A446MUV2
Triticum turg A0A446X5A7
Triticum turg A0A446NRU9
Triticum turg A0A446MVY3
Triticum turg A0A446MUV4
Triticum turg A0A446X576
Triticum turg A0A446NRV0
Triticum turg A0A446L422
Triticum turg A0A446LLB4
Triticum turg A0A446L9C3
Triticum turg A0A446YVN7
Triticum turg A0A446NRW9
Triticum turg A0A446XZM9
Triticum turg A0A446MV59
Triticum turg A0A446Y2M9
Triticum turg A0A446LL76
Triticum turg A0A446MV73
Triticum turg A0A446UJ87
Triticum turg A0A446L975
Triticum turg A0A446Y3N5
Triticum turg A0A446KV41
Triticum turg A0A446K617
Triticum turg A0A446QGE1
Triticum turg A0A446MV32
Triticum turg A0A446PK69
Triticum turg A0A446SK23
Triticum turg A0A446YZ67
Triticum turg A0A446MI06
Triticum turg A0A446LI16
Triticum turg A0A446XY56
Triticum turg A0A446YP66
Triticum turg A0A446LL77
Triticum turg A0A446REX2
Triticum turg A0A446NK72
Triticum turg A0A446UJA1

Triticum turg A0A446YLI9
Triticum turg A0A446K601
Triticum turg A0A446UJ43
Triticum turg A0A446NK74
Triticum turg A0A446LL83
Triticum turg A0A446XPQ6
Triticum turg A0A446VUA3
Triticum turg A0A446NRU8
Triticum turg A0A446KUV2
Triticum turg A0A446Q9Q4
Triticum turg A0A446K5J4
Triticum turg A0A446WSK7
Triticum turg A0A446NRZ1
Triticum turg A0A446M568
Triticum turg A0A446JML1
Triticum turg A0A446SLJ0
Triticum turg A0A446R9Q0
Triticum turg A0A446UK57
Triticum turg A0A446LI26
Triticum turg A0A446M587
Triticum turg A0A446XU41
Triticum turg A0A446UJ71
Triticum turg A0A446T6X8
Triticum turg A0A446KV32
Triticum turg A0A446LM41
Triticum turg A0A446L404
Triticum turg A0A446MV88
Triticum turg A0A446NRW1
Triticum turg A0A446J771
Triticum turg A0A446M551
Triticum turg A0A446KUW7
Triticum turg A0A446KUU3
Triticum turg A0A446MV70
Triticum turg A0A446LHW7
Triticum turg A0A446MV74
Triticum turg A0A446M5B2
Triticum turg A0A446LL46
Triticum turg A0A446MV68
Triticum turg A0A446PK92
Triticum turg A0A446MVS2
Triticum turg A0A446XVJ2

Triticum turg A0A446LHU2
Triticum turg A0A446UKD3
Triticum turg A0A446M590
Triticum turg A0A446K035
Triticum turg A0A446SLG8
Triticum turg A0A446LKQ3
Triticum turg A0A446QEY0
Triticum turg A0A446MUR6
Triticum turg A0A446IP50
Triticum turg A0A446P3S7
Triticum turg A0A446QNL4
Triticum turg A0A446R7C5
Triticum turg A0A446JML0
Triticum turg A0A446NRW7
Triticum turg A0A446MRN0
Triticum turg A0A446MRN9
Triticum turg A0A446MUS6
Triticum turg A0A446R1Y3
Triticum turg A0A446LJB5
Triticum turg A0A446KHJ9
Triticum turg A0A446YPI4
Triticum turg A0A446XPR5
Triticum turg A0A446LHT9
Triticum turg A0A446KUZ6
Triticum turg A0A446MVB0
Triticum turg A0A446K6J7
Triticum turg A0A446LHY6
Triticum turg A0A446Q9Y2
Triticum turg A0A446TBU5
Triticum turg A0A446L175
Triticum turg A0A446MRK5
Triticum turg A0A446Q9T2
Triticum turg A0A446UJ22
Triticum turg A0A446M562
Triticum turg A0A446MRI6
Triticum turg A0A446P3S9
Triticum turg A0A446SK01
Triticum turg A0A446YS27
Triticum turg A0A446MI08
Triticum turg A0A446MCY9
Triticum turg A0A446MRR0

Triticum turg A0A446MRK8
Triticum turg A0A446KV16
Triticum turg A0A446M570
Triticum turg A0A446L164
Triticum turg A0A446M582
Triticum turg A0A446REF2
Triticum turg A0A446JA06
Triticum turg A0A446MRP1
Triticum turg A0A446M580
Triticum turg A0A446M565
Triticum turg A0A446SLJ2
Triticum turg A0A446J9N6
Triticum turg A0A446MDG6
Triticum turg A0A446UJC2
Triticum turg A0A446UJ81
Triticum turg A0A446SLP8
Triticum turg A0A446LL86
Triticum turg A0A446Y563
Triticum turg A0A446MZY1
Triticum turg A0A446K620
Triticum turg A0A446MVV2
Triticum turg A0A446NB74
Triticum turg A0A446LLA2
Triticum turg A0A446MUS4
Triticum turg A0A446TT86
Triticum turg A0A446LRA6
Triticum turg A0A446L9G2
Triticum turg A0A446L414
Triticum turg A0A446K5H4
Triticum turg A0A446SXH2
Triticum turg A0A446MV89
Triticum turg A0A446MRJ7
Triticum turg A0A446QNN2
Triticum turg A0A446MRS6
Triticum turg A0A446L9B9
Triticum turg A0A446NRU1
Triticum turg A0A446MWC8
Triticum turg A0A446MVB4
Triticum turg A0A446UJ63
Triticum turg A0A446NRY5
Triticum turg A0A446K9J8

Triticum turg A0A446MW84
Triticum turg A0A446USM4
Triticum turg A0A446K5G2
Triticum turg A0A446MRT0
Triticum turg A0A446UQJ9
Triticum turg A0A446JML2
Triticum turg A0A446UJ99
Triticum turg A0A446MSW5
Triticum turg A0A446MUV5
Triticum turg A0A446L406
Triticum turg A0A446RS14
Triticum turg A0A446SAY9
Triticum turg A0A446YNY1
Triticum turg A0A446NS01
Triticum turg A0A446KV78
Triticum turg A0A446LL87
Triticum turg A0A446K036
Triticum turg A0A446L9A2
Triticum turg A0A446NRY3
Triticum turg A0A446MUH3
Triticum turg A0A446NRV5
Triticum turg A0A446L3X8
Triticum turg A0A446LL80
Triticum turg A0A446QLZ4
Triticum turg A0A446UJ94
Triticum turg A0A446MUY4
Triticum turg A0A446NAL7
Triticum turg A0A446VV46
Triticum turg A0A446MVA2
Triticum turg A0A446LM62
Triticum turg A0A446UJ59
Triticum turg A0A446MRJ9
Triticum turg A0A446MV20
Triticum turg A0A446LHR2
Triticum turg A0A446LL97
Triticum turg A0A446MRG0
Triticum turg A0A446LHX1
Triticum turg A0A446U9T5
Triticum turg A0A446XVM3
Triticum turg A0A446M558
Triticum turg A0A446R9M6

Triticum turg A0A446JGG2
Triticum turg A0A446UJ85
Triticum turg A0A446TQK5
Triticum turg A0A446KVB1
Triticum turg A0A446YTB0
Triticum turg A0A446MY43
Triticum turg A0A446K667
Triticum turg A0A446MVJ1
Triticum turg A0A446LL78
Triticum turg A0A446TR27
Triticum turg A0A446TQK4
Triticum turg A0A446REI4
Triticum turg A0A446JA28
Triticum turg A0A446Q0X1
Triticum turg A0A446K647
Triticum turg A0A446RLD6
Triticum turg A0A446RM11
Triticum turg A0A446LSA0
Triticum turg A0A446MUX5
Triticum turg A0A446MVA1
Triticum turg A0A446K6G4
Triticum turg A0A446UJ65
Triticum turg A0A446KGH3
Triticum turg A0A446KV82
Triticum turg A0A446KV63
Triticum turg A0A446MV50
Triticum turg A0A446NK01
Triticum turg A0A446IHX8
Triticum turg A0A446MRT7
Triticum turg A0A446K639
Triticum turg A0A446MRH0
Triticum turg A0A446MV40
Triticum turg A0A446R1L1
Triticum turg A0A446RQM3
Triticum turg A0A446K5K0
Triticum turg A0A446MD27
Triticum turg A0A446MW47
Triticum turg A0A446LHX2
Triticum turg A0A446TQK1
Triticum turg A0A446L9F6
Triticum turg A0A446RDR5

Triticum turg A0A446XSM3
Triticum turg A0A446KV34
Triticum turg A0A446MV21
Triticum turg A0A446M526
Triticum turg A0A446R9J1
Triticum turg A0A446QA33
Triticum turg A0A446NS73
Triticum turg A0A446Q9P9
Triticum turg A0A446TT87
Triticum turg A0A446SMN3
Triticum turg A0A446P4B8
Triticum turg A0A446K5Z2
Triticum turg A0A446MV84
Triticum turg A0A446LHW3
Triticum turg A0A446UIZ6
Triticum turg A0A446XTI5
Triticum turg A0A446RZS7
Triticum turg A0A446LJA8
Triticum turg A0A446KUZ3
Triticum turg A0A446MRI0
Triticum turg A0A446QA44
Triticum turg A0A446MUT9
Triticum turg A0A446LLE5
Triticum turg A0A446VUA1
Triticum turg A0A446M5A4
Triticum turg A0A446LHT8
Triticum turg A0A446KGV8
Triticum turg A0A446J7A1
Triticum turg A0A446M579
Triticum turg A0A446RQK2
Triticum turg A0A446LR36
Triticum turg A0A446NRY7
Triticum turg A0A446NK41
Triticum turg A0A446NRV3
Triticum turg A0A446XPD9
Triticum turg A0A446K031
Triticum turg A0A446N0X5
Triticum turg A0A446P3S3
Triticum turg A0A446Q9R0
Triticum turg A0A446UJ70
Triticum turg A0A446TBW3

Triticum turg A0A446UJC6
Triticum turg A0A446S8Q6
Triticum turg A0A446X3F4
Triticum turg A0A446MVC2
Triticum turg A0A446WSC8
Triticum turg A0A446RZ79
Triticum turg A0A446Y556
Triticum turg A0A446SJZ8
Triticum turg A0A446Q9P3
Triticum turg A0A446Y367
Triticum turg A0A446II08
Triticum turg A0A446WMX9
Triticum turg A0A446Q9Q0
Triticum turg A0A446L3Z8
Triticum turg A0A446LJJ7
Triticum turg A0A446J765
Triticum turg A0A446UJ32
Triticum turg A0A446K938
Triticum turg A0A446NRY8
Triticum turg A0A446UJ23
Triticum turg A0A446N128
Triticum turg A0A446UJ69
Triticum turg A0A446LL91
Triticum turg A0A446JP82
Triticum turg A0A446MRL0
Triticum turg A0A446Q9P6
Triticum turg A0A446SLQ8
Triticum turg A0A446LHT2
Triticum turg A0A446LR56
Triticum turg A0A446MVD0
Triticum turg A0A446NRX7
Triticum turg A0A446K9M4
Triticum turg A0A446MVX3
Triticum turg A0A446K1Q8
Triticum turg A0A446NAD8
Triticum turg A0A446N9Y9
Triticum turg A0A446UJ82
Triticum turg A0A446NK52
Triticum turg A0A446YTG9
Triticum turg A0A446MRM3
Triticum turg A0A446UJ89

Triticum turg A0A446NS53
Triticum turg A0A446KFS7
Triticum turg A0A446Q8P2
Triticum turg A0A446TIF8
Triticum turg A0A446PVA8
Triticum turg A0A446WSF5
Triticum turg A0A446K631
Triticum turg A0A446NAK8
Triticum turg A0A446TBV6
Triticum turg A0A446L425
Triticum turg A0A446MW67
Triticum turg A0A446XZR0
Triticum turg A0A446NK46
Triticum turg A0A446U2C0
Triticum turg A0A446JGI5
Triticum turg A0A446P474
Triticum turg A0A446K605
Triticum turg A0A446YZ88
Triticum turg A0A446M589
Triticum turg A0A446PV83
Triticum turg A0A446YAM4
Triticum turg A0A446LMA9
Triticum turg A0A446MWH4
Triticum turg A0A446QNY6
Triticum turg A0A446QH58
Triticum turg A0A446LHY0
Triticum turg A0A446KV42
Triticum turg A0A446M569
Triticum turg A0A446Q0X5
Triticum turg A0A446KV52
Triticum turg A0A446MVE3
Triticum turg A0A446MAN9
Triticum turg A0A446MUR3
Triticum turg A0A446M561
Triticum turg A0A446MVD1
Triticum turg A0A446X556
Triticum turg A0A446M5B3
Triticum turg A0A446R1T8
Triticum turg A0A446KUY6
Triticum turg A0A446LHX4
Triticum turg A0A446P114

Triticum turg A0A446MRM4
Triticum turg A0A446LHS8
Triticum turg A0A446UJA5
Triticum turg A0A446XNN9
Triticum turg A0A446NRV9
Triticum turg A0A446NRY6
Triticum turg A0A446LHY1
Triticum turg A0A446X3F2
Triticum turg A0A446NRX3
Triticum turg A0A446WT80
Triticum turg A0A446NRV2
Triticum turg A0A446TBX2
Triticum turg A0A446MRI3
Triticum turg A0A446NS43
Triticum turg A0A446MWZ7
Triticum turg A0A446WSF8
Triticum turg A0A446JGF3
Triticum turg A0A446X5C0
Triticum turg A0A446NYC9
Triticum turg A0A446WSC2
Triticum turg A0A446X5E5
Triticum turg A0A446MRJ3
Triticum turg A0A446MDA4
Triticum turg A0A446MRG9
Triticum turg A0A446Y558
Triticum turg A0A446UJ53
Triticum turg A0A446NRY0
Triticum turg A0A446KUV4
Triticum turg A0A446MV57
Triticum turg A0A446KUW8
Triticum turg A0A446LL74
Triticum turg A0A446LHW1
Triticum turg A0A446SWQ4
Triticum turg A0A446YTB1
Triticum turg A0A446YP03
Triticum turg R9UD00
Triticum turg A0A446M557
Triticum turg A0A446MRP0
Triticum turg A0A446NA00
Triticum turg A0A446L155
Triticum turg A0A446MVK1

Triticum turg A0A446MV76
Triticum turg A0A446LHY3
Triticum turg A0A446XU55
Triticum turg A0A446WTD2
Triticum turg A0A446WSD5
Triticum turg A0A446XS86
Triticum turg A0A446LI36
Triticum turg A0A446J7A5
Triticum turg A0A446LI37
Triticum turg A0A446YPB5
Triticum turg A0A446TQL2
Triticum turg A0A446NAE6
Triticum turg A0A446IP93
Triticum turg A0A446MD40
Triticum turg A0A446NRW2
Triticum turg A0A446MUS9
Triticum turg A0A446MV97
Triticum turg A0A446P468
Triticum turg A0A446USI9
Triticum turg A0A446MVI9
Triticum turg A0A446UJ90
Triticum turg A0A446J796
Triticum turg A0A446R1M6
Triticum turg A0A446KHK3
Triticum turg A0A446RQL8
Triticum turg A0A446VUB8
Triticum turg A0A446RQK5
Triticum turg A0A446IP14
Triticum turg A0A446KV31
Triticum turg A0A446LLA5
Triticum turg A0A446Y304
Triticum turg A0A446MUN3
Triticum turg A0A446MV33
Triticum turg A0A446KV62
Triticum turg A0A446KVC1
Triticum turg A0A446LL47
Triticum turg A0A446NRX4
Triticum turg A0A446L993
Triticum turg A0A446M585
Triticum turg A0A446IP36
Triticum turg A0A446UQG9

Triticum turg A0A446M527
Triticum turg A0A446VAR7
Triticum turg A0A446MUY8
Triticum turg A0A446L401
Triticum turg A0A446U9S4
Triticum turg A0A446Y557
Triticum turg A0A446M553
Triticum turg A0A446MV07
Triticum turg A0A446UWT0
Triticum turg A0A446NBI9
Triticum turg A0A446M528
Triticum turg A0A446LLF8
Triticum turg A0A446Q0V8
Triticum turg A0A446LI24
Triticum turg A0A446MUU3
Triticum turg A0A446YLJ5
Triticum turg A0A446MV00
Triticum turg A0A446MRJ2
Triticum turg A0A446TBW0
Triticum turg A0A446KUQ8
Triticum turg A0A446R9L4
Triticum turg A0A446MUL5
Triticum turg A0A446TBX1
Triticum turg A0A446IHX1
Triticum turg A0A446KV07
Triticum turg A0A446N0X4
Triticum turg A0A446K677
Triticum turg A0A446RZ10
Triticum turg A0A446MI12
Triticum turg A0A446V4H5
Triticum turg A0A446MV35
Triticum turg A0A446VQ63
Triticum turg A0A446MUM6
Triticum turg A0A446LLB3
Triticum turg A0A446QEZ2
Triticum turg A0A446PK75
Triticum turg A0A446XNN5
Triticum turg A0A446K6I0
Triticum turg A0A446TC97
Triticum turg A0A446M5A2
Triticum turg A0A446KV22

Triticum turg A0A446TRY9
Triticum turg A0A446NRW4
Triticum turg A0A446SK08
Triticum turg A0A446UWU1
Triticum turg A0A446MVP0
Triticum turg A0A446MUY6
Triticum turg A0A446L973
Triticum turg A0A446MW55
Triticum turg A0A446K6L6
Triticum turg A0A446XPB9
Triticum turg A0A446LHU8
Triticum turg A0A446J9R0
Triticum turg A0A446MD20
Triticum turg A0A446LI00
Triticum turg A0A446UZM1
Triticum turg A0A446MRL6
Triticum turg A0A446YRY6
Triticum turg A0A446MRN5
Triticum turg A0A446LHV3
Triticum turg A0A446R1U8
Triticum turg A0A446MW53
Triticum turg A0A446MVW1
Triticum turg A0A446N457
Triticum turg A0A446Q0X7
Triticum turg A0A446KUW1
Triticum turg A0A446LL55
Triticum turg A0A446L487
Triticum turg A0A446JGM8
Triticum turg A0A446QQX8
Triticum turg A0A446YZ82
Triticum turg A0A446Y148
Triticum turg A0A446WQF2
Triticum turg A0A446MIA1
Triticum turg A0A446NRU7
Triticum turg A0A446UJ72
Triticum turg A0A446XU56
Triticum turg A0A446L9A0
Triticum turg A0A446PV58
Triticum turg A0A446WSH4
Triticum turg A0A446SAW2
Triticum turg A0A446NS67

Triticum turg A0A446MRL5
Triticum turg A0A446MRK7
Triticum turg A0A446YTC2
Triticum turg A0A446RT60
Triticum turg A0A446YT98
Triticum turg A0A446WU68
Triticum turg A0A446UJ54
Triticum turg A0A446J7B3
Triticum turg A0A446VLN6
Triticum turg A0A446VUB4
Triticum turg A0A446Q9P0
Triticum turg A0A446LHV1
Triticum turg A0A446UJ60
Triticum turg A0A446MV39
Triticum turg A0A446MVB2
Triticum turg A0A446KV26
Triticum turg A0A446LHV4
Triticum turg A0A446KUW4
Triticum turg A0A446KV73
Triticum turg A0A446Q0W0
Triticum turg A0A446MV61
Triticum turg A0A446Y562
Triticum turg A0A446MV98
Triticum turg A0A446R7D5
Triticum turg A0A446MV30
Triticum turg A0A446KV89
Triticum turg A0A446IHW9
Triticum turg A0A446PV86
Triticum turg A0A446WVE1
Triticum turg A0A446MT54
Triticum turg A0A446L3Y7
Triticum turg A0A446MRL9
Triticum turg A0A446LLB2
Triticum turg A0A446MV41
Triticum turg A0A446RZ83
Triticum turg A0A446X395
Triticum turg A0A446U371
Triticum turg A0A446UW14
Triticum turg A0A446MUU4
Triticum turg A0A446QA53
Triticum turg A0A446SLI6

Triticum turg A0A446LM93
Triticum turg A0A446NRZ4
Triticum turg A0A446KVA1
Triticum turg A0A446PV26
Triticum turg A0A446L9A8
Triticum turg A0A446QLB6
Triticum turg A0A446MRN4
Triticum turg A0A446L4A2
Triticum turg A0A446UJ74
Triticum turg A0A446UW15
Triticum turg A0A446UJ12
Triticum turg A0A446WUB2
Triticum turg A0A446MRL4
Triticum turg A0A446MVE2
Triticum turg A0A446KUU1
Triticum turg A0A446SLN0
Triticum turg A0A446Y3A1
Triticum turg A0A446L430
Triticum turg A0A446Q8F8
Triticum turg A0A446SK49
Triticum turg A0A446KUU5
Triticum turg A0A446RQP4
Triticum turg A0A446Y2R3
Triticum turg A0A446LHV9
Triticum turg A0A446UJA0
Triticum turg A0A446NRV7
Triticum turg A0A446NRW6
Triticum turg A0A446UEC6
Triticum turg A0A446MRI7
Triticum turg A0A446MRJ5
Triticum turg A0A446SXG6
Triticum turg A0A446MRQ6
Triticum turg A0A446MZY9
Triticum turg A0A446P423
Triticum turg A0A446M576
Triticum turg A0A446Q9N8
Triticum turg A0A446LZ06
Triticum turg A0A446IP45
Triticum turg A0A446M592
Triticum turg A0A446X394
Triticum turg A0A446KUU6

Triticum turg A0A446LNU4
Triticum turg A0A446KUX7
Triticum turg A0A446M574
Triticum turg A0A446UJB1
Triticum turg A0A446MVG4
Triticum turg A0A446MVM9
Triticum turg A0A446LSA5
Triticum turg A0A446WSC9
Triticum turg A0A446YSZ5
Triticum turg A0A446VU91
Triticum turg A0A446MVE4
Triticum turg A0A446KV50
Triticum turg A0A446MUZ5
Triticum turg A0A446UJA3
Triticum turg A0A446N116
Triticum turg A0A446RDR4
Triticum turg A0A446L474
Triticum turg A0A446LLE4
Triticum turg A0A446MT19
Triticum turg A0A446UJA4
Triticum turg A0A446MRI4
Triticum turg A0A446LHS2
Triticum turg A0A446MVL2
Triticum turg A0A446Q9R4
Triticum turg A0A446YS15
Triticum turg A0A446MUR2
Triticum turg A0A446MRQ2
Triticum turg A0A446M578
Triticum turg A0A446KUZ2
Triticum turg A0A446RS84
Triticum turg A0A446QR39
Triticum turg A0A446UJ79
Triticum turg A0A446LHU0
Triticum turg A0A446MVA5
Triticum turg A0A446NRV8
Triticum turg A0A446R7D1
Triticum turg A0A446KUY2
Triticum turg A0A446LLE1
Triticum turg A0A446M599
Triticum turg A0A446UZL9
Triticum turg A0A446MV19

Triticum turg A0A446II24
Triticum turg A0A446MV85
Triticum urar M7YPH8
Triticum urar M7ZVA3
Triticum urar M7ZF50
Triticum urar M7YWV4
Triticum urar M7Z5B8
Triticum urar M7YN61
Triticum urar M8AE45
Triticum urar M7ZNB1
Triticum urar M7YXB0
Triticum urar M8AST1
Triticum urar M7YR24
Triticum urar M7ZA47
Triticum urar M7YT26
Triticum urar M7YJ40
Triticum urar M8ADD0
Triticum urar M7ZJA7
Triticum urar M7YQ54
Triticum urar M8AUM9
Triticum urar M8ANE0
Triticum urar M7ZXQ8
Triticum urar M8AUZ7
Triticum urar M7ZHG8
Triticum urar M7ZXN9
Triticum urar M7ZBE9
Triticum urar M7YEQ0
Triticum urar M7ZCG6
Triticum urar M8AAR3
Triticum urar M8A1P2
Triticum urar T1NTJ1
Triticum urar T1N2R3
Triticum urar T1LF75
Triticum urar T1N4J2
Triticum urar T1MJA6
Triticum urar T1M819
Triticum urar T1LC35
Triticum urar T1MQA5
Triticum urar T1LSX1
Triticum urar T1MIZ5
Triticum urar T1LNZ0

Triticum urar T1N671
Triticum urar T1MUZ3
Triticum urar T1NI12
Triticum urar T1NKJ6
Triticum urar T1NR99
Triticum urar T1LEM3
Triticum urar T1M0W0
Triticum urar T1NAZ0
Triticum urar T1MFP5
Triticum urar T1LZ01
Triticum urar T1NPH7
Triticum urar T1NHL1
Triticum urar T1MDZ0
Triticum urar T1NS03
Triticum urar T1M9E3
Triticum urar T1LM18
Triticum urar T1LAC9
Triticum urar T1LBZ0
Triticum urar T1MXQ5
Triticum urar T1MQD4
Triticum urar T1MPW5
Triticum urar T1LM09
Triticum urar T1NPT5
Triticum urar T1LM19
Triticum urar T1LU09
Triticum urar T1NK32
Triticum urar T1LUD5
Triticum urar T1NF00
Triticum urar T1NC78
Triticum urar T1NN82
Triticum urar T1MA35
Triticum urar T1L9J1
Triticum urar T1MBB1
Triticum urar T1LXJ4
Triticum urar T1LKE2
Triticum urar T1LGV7
Triticum urar T1MAS0
Triticum urar T1MPD3
Triticum urar T1MSD0
Triticum urar T1LRP0
Triticum urar T1MF28

Triticum urar T1L9J3
Triticum urar T1M1J0
Triticum urar T1NWB7
Triticum urar T1M5P2
Triticum urar T1NBE0
Triticum urar T1LGV4
Triticum urar T1LSN9
Triticum urar T1NGW2
Triticum urar T1LNY8
Triticum urar T1MDI5
Triticum urar T1NVC5
Triticum urar T1NHJ6
Triticum urar T1LSX0
Triticum urar T1L8X2
Triticum urar T1MZ83
Triticum urar T1LD28
Triticum urar T1LAN4
Triticum urar T1LAN5
Triticum urar T1NM62
Triticum urar T1LF74
Triticum urar T1LGV3
Triticum urar T1LF77
Triticum urar T1MQD3
Triticum urar T1LI56
Triticum urar T1NW29
Triticum urar T1N4G6
Triticum urar T1M6J5
Triticum urar T1LC36
Triticum urar T1M1I9
Triticum urar T1LHM4
Triticum urar T1MAK7
Triticum urar T1ME31
Triticum urar T1MKF8
Triticum urar T1LI57
Triticum urar T1LM22
Triticum urar T1LBJ3
Triticum urar T1LAN6
Triticum urar M7YNN7
Triticum urar M7Z646
Triticum urar M8A2P4
Tropicibacter A0A0P1GFE6

| | |
|---|---|
| Tropicibacter | A0A0P1GTJ8 |
| Tropicibacter | A0A238J853 |
| Tropicimonas | A0A239KUI1 |
| Tropilaelaps r | A0A1V9XN76 |
| Tropilaelaps r | A0A1V9XRJ6 |
| Tropilaelaps r | A0A1V9XWL9 |
| Tropilaelaps r | A0A1V9XIR5 |
| Tropilaelaps r | A0A1V9X1B6 |
| Tropilaelaps r | A0A1V9Y2Z8 |
| Tropilaelaps r | A0A1V9X236 |
| Tropilaelaps r | A0A1V9X4X2 |
| Tupaia belang | A0A144ZKR4 |
| Tupaia chiner | L9J9W1 |
| Tupaia chiner | L9J9Q4 |
| Tupaia chiner | L8Y0B6 |
| Tupanvirus d | A0A2K9L9B3 |
| Tupanvirus d | A0A2K9L3H7 |
| Tupanvirus d | A0A2K9L521 |
| Tupanvirus d | A0A2K9L700 |
| Tupanvirus d | A0A2K9L519 |
| Tupanvirus s | A0A2K9L1A9 |
| Tupanvirus s | A0A2K9L2G0 |
| Tupanvirus s | A0A2K9L0F4 |
| Tupanvirus s | A0A2K9L648 |
| Tupanvirus s | A0A2K9L6M7 |
| Tursiops trun | A0A2U4CNJ2 |
| Tursiops trun | A0A2U3V1A2 |
| Tursiops trun | A0A2U4CNJ3 |
| Tursiops trun | A0A2U4A725 |
| Tursiops trun | A0A2U4CNU7 |
| Tursiops trun | A0A2U4CNK8 |
| Tursiops trun | A0A2U4BS66 |
| Tursiops trun | A0A2U4A724 |
| Tyto alba.xlsx | A0A093FWK3 |
| Tyto alba.xlsx | A0A093FQ32 |
| Umbilicaria p | A0A1W5CXS0 |
| Umbilicaria p | A0A1W5CXL4 |
| uncultured ba | K2AG36 |
| uncultured Pl | A0A212KZK8 |
| Ursus americ | A0A452RK74 |
| Ursus americ | A0A452RW34 |

| | |
|---|---|
| Ursus americ | A0A452QNT0 |
| Ursus americ | A0A452RK69 |
| Ursus arctos | A0A3Q7VTE7 |
| Ursus arctos | A0A3Q7W7U3 |
| Ursus arctos | A0A3Q7WRV2 |
| Ursus arctos | A0A3Q7XCY5 |
| Ursus arctos | A0A3Q7WHD8 |
| Ursus arctos | A0A3Q7V5V4 |
| Ursus arctos | A0A3Q7X1Z0 |
| Ursus arctos | A0A3Q7WRV6 |
| Ursus arctos | A0A3Q7XCZ2 |
| Ursus arctos | A0A3Q7XNM7 |
| Ursus maritin | A0A384BVT0 |
| Ursus maritin | A0A384DPH5 |
| Ursus maritin | A0A384BP91 |
| Ursus maritin | A0A384D4U8 |
| Ursus maritin | A0A384BPA7 |
| Ursus maritin | A0A384DNT9 |
| Ursus maritin | A0A452VCB3 |
| Ursus maritin | A0A452U7G2 |
| Ursus maritin | A0A452VCC2 |
| Ursus maritin | A0A452U7G7 |
| Valsa mali va | A0A194V1F2 |
| Valsa mali va | A0A194UVJ5 |
| Valsa mali va | A0A194ULX3 |
| Valsa mali va | A0A194VCV2 |
| Valsa mali.xls | A0A194W0K5 |
| Valsa mali.xls | A0A194W6W9 |
| Valsa mali.xls | A0A194VR95 |
| Valsa mali.xls | A0A194W4H9 |
| Valsa malicol | A0A423VPM3 |
| Valsa malicol | A0A423VSL1 |
| Valsa malicol | A0A423WD94 |
| Valsa malicol | A0A423W9F7 |
| Valsa malicol | A0A423WZH0 |
| Valsa malicol | A0A423WV28 |
| Valsa sordida | A0A423WIT3 |
| Valsa sordida | A0A423WH16 |
| Valsa sordida | A0A423VR78 |
| Valsa sordida | A0A423VA80 |
| Variibacter g | A0A0S3PPN4 |

| | |
|---|---|
| Variovorax bo | A0A1E7U9P5 |
| Variovorax bo | A0A250DK61 |
| Variovorax pa | E6V8Z5 |
| Variovorax pa | A0A0P9AFP2 |
| Variovorax sp | A0A2G6X860 |
| Variovorax sp | A0A1I5PZ82 |
| Variovorax sp | A0A2N4SJ19 |
| Variovorax sp | A0A2E9C131 |
| Venturia inae | A0A370IXJ3 |
| Venturia inae | A0A370JCW3 |
| Verrucomicro | A0A2V5TIP4 |
| Verrucomicro | A0A2D5MT73 |
| Verruconis ga | A0A0D1XT13 |
| Verruconis ga | A0A0D1YG70 |
| Verticillium a | C9SWV4 |
| Verticillium d | G2XFJ0 |
| Verticillium d | A0A2J8BUS4 |
| Verticillium d | A0A444SAT6 |
| Verticillium d | A0A366Q651 |
| Verticillium lo | A0A0G4M299 |
| Verticillium lo | A0A0G4MAI0 |
| Verticillium lo | A0A0G4MID9 |
| Verticillium lo | A0A0G4MA18 |
| Verticillium n | A0A3M9Y2Y5 |
| Vigna angular | A0A0S3SI41 |
| Vigna angular | A0A0S3S9L1 |
| Vigna angular | A0A0S3SVK9 |
| Vigna angular | A0A0S3RUP5 |
| Vigna angular | A0A0S3RDA5 |
| Vigna angular | A0A0S3S2D1 |
| Vigna angular | A0A0S3SFB8 |
| Vigna angular | A0A0S3SI12 |
| Vigna angular | A0A0S3S983 |
| Vigna angular | A0A0S3SHB1 |
| Vigna angular | A0A0S3S2C4 |
| Vigna angular | A0A0S3RAQ9 |
| Vigna angular | A0A0S3SI52 |
| Vigna angular | A0A0S3SBH8 |
| Vigna angular | A0A0S3SHD6 |
| Vigna angular | A0A0S3RUY5 |
| Vigna angular | A0A0S3RV54 |

| | |
|---|---|
| Vigna angulaı | A0A0S3SWB4 |
| Vigna angulaı | A0A0S3SI31 |
| Vigna angulaı | A0A0S3SIJ7 |
| Vigna angulaı | A0A0S3S2E3 |
| Vigna angulaı | A0A0S3T100 |
| Vigna angulaı | A0A0S3SAD2 |
| Vigna angulaı | A0A0S3SFJ3 |
| Vigna angulaı | A0A0S3S9H7 |
| Vigna angulaı | A0A0S3QZQ6 |
| Vigna angulaı | A0A0S3SKU3 |
| Vigna angulaı | A0A0S3SHG6 |
| Vigna angulaı | A0A0S3S9C6 |
| Vigna angulaı | A0A0S3RUP4 |
| Vigna angulaı | A0A0S3SYJ7 |
| Vigna angulaı | A0A0S3RUT8 |
| Vigna angulaı | A0A0S3SII5 |
| Vigna angulaı | A0A0S3RTM0 |
| Vigna angulaı | A0A0S3QZS6 |
| Vigna angulaı | A0A0S3RQC6 |
| Vigna angulaı | A0A0S3SBP8 |
| Vigna angulaı | A0A0S3SKN8 |
| Vigna angulaı | A0A0S3S9A9 |
| Vigna angulaı | A0A0S3S9H4 |
| Vigna angulaı | A0A0S3S2C1 |
| Vigna angulaı | A0A0S3S9B5 |
| Vigna angulaı | A0A0S3RV98 |
| Vigna angulaı | A0A0S3RBJ5 |
| Vigna angulaı | A0A0S3QZN0 |
| Vigna angulaı | A0A0S3SHY7 |
| Vigna angulaı | A0A0S3SA88 |
| Vigna angulaı | A0A0S3SU71 |
| Vigna angulaı | A0A0S3SVC8 |
| Vigna angulaı | A0A0S3RUR6 |
| Vigna angulaı | A0A0S3SHH7 |
| Vigna angulaı | A0A0S3RUS8 |
| Vigna angulaı | A0A0S3SB63 |
| Vigna angulaı | A0A0S3T043 |
| Vigna angulaı | A0A0S3RZL4 |
| Vigna angulaı | A0A0S3S969 |
| Vigna angulaı | A0A0S3RZ03 |
| Vigna angulaı | A0A0S3SB96 |

| | |
|---|---|
| Vigna angulai | A0A0S3QZV8 |
| Vigna angulai | A0A0S3SHB5 |
| Vigna angulai | A0A0S3S9G1 |
| Vigna angulai | A0A0S3QZX3 |
| Vigna angulai | A0A0S3S2F1 |
| Vigna radiata | A0A3Q0FG34 |
| Vigna radiata | A0A1S3UY68 |
| Vigna radiata | A0A1S3URP2 |
| Vigna radiata | A0A1S3URH0 |
| Vigna radiata | A0A3Q0ET99 |
| Vigna radiata | A0A3Q0EYW5 |
| Vigna radiata | A0A1S3TL02 |
| Vigna radiata | A0A1S3W0Y9 |
| Vigna radiata | A0A1S3U2W6 |
| Vigna radiata | A0A1S3W0T2 |
| Vigna radiata | A0A1S3W0T7 |
| Vigna radiata | A0A1S3W144 |
| Vigna radiata | A0A1S3U4J5 |
| Vigna radiata | A0A1S3TNA8 |
| Vigna radiata | A0A1S3T9S3 |
| Vigna radiata | A0A1S3VJI6 |
| Vigna radiata | A0A1S3TMS8 |
| Vigna radiata | A0A3Q0ESJ5 |
| Vigna radiata | A0A1S3TLG9 |
| Vigna radiata | A0A1S3VUM7 |
| Vigna radiata | A0A1S3TUL8 |
| Vigna radiata | A0A1S3TSI3 |
| Vigna radiata | A0A1S3U0N3 |
| Vigna radiata | A0A3Q0EZS4 |
| Vigna radiata | A0A1S3VLI3 |
| Vigna radiata | A0A1S3U6F1 |
| Vigna radiata | A0A3Q0EVU4 |
| Vigna radiata | A0A3Q0ET57 |
| Vigna radiata | A0A1S3VJJ8 |
| Vigna radiata | A0A1S3T8I0 |
| Vigna radiata | A0A1S3W0Y5 |
| Vigna radiata | A0A3Q0F2X9 |
| Vigna radiata | A0A1S3TGW0 |
| Vigna radiata | A0A1S3VEP0 |
| Vigna radiata | A0A1S3VT28 |
| Vigna radiata | A0A1S3U039 |

Vigna radiata A0A3Q0FL44
Vigna radiata A0A1S3W0L1
Vigna radiata A0A1S3VU23
Vigna radiata A0A3Q0EIT8
Vigna radiata A0A3Q0EQK2
Vigna radiata A0A1S3TYK8
Vigna radiata A0A1S3U5X8
Vigna radiata A0A1S3TUD5
Vigna radiata A0A3Q0F6W5
Vigna radiata A0A1S3U6A4
Vigna radiata A0A1S3TSG7
Vigna radiata A0A3Q0ERZ1
Vigna radiata A0A3Q0FGU0
Vigna radiata A0A3Q0EP47
Vigna radiata A0A3Q0ELC1
Vigna radiata A0A1S3U943
Vigna radiata A0A3Q0EVT5
Vigna radiata A0A3Q0F3I1
Vigna radiata A0A1S3VE13
Vigna radiata A0A1S3VKA1
Vigna radiata A0A1S3TI08
Vigna radiata A0A3Q0FK37
Vigna radiata A0A1S3VLA4
Vigna radiata A0A3Q0F5J2
Vigna radiata A0A1S3U186
Vigna radiata A0A3Q0FKN7
Vigna radiata A0A3Q0FHK9
Vigna radiata A0A1S3TXM9
Vigna radiata A0A1S3U6A9
Vigna radiata A0A1S3TLE5
Vigna radiata A0A1S3W149
Vigna radiata A0A1S3U6B7
Vigna radiata A0A1S3VCH7
Vigna radiata A0A1S3TU21
Vigna radiata A0A1S3U951
Vigna radiata A0A3Q0F0J3
Vigna radiata A0A3Q0EUC6
Vigna radiata A0A3Q0EX51
Vigna radiata A0A1S3USU6
Vigna radiata A0A1S3VJQ9
Vigna radiata A0A1S3VSU2

| | |
|---|---|
| Vigna radiata | A0A3Q0FH19 |
| Vigna radiata | A0A3Q0EQ28 |
| Vigna radiata | A0A3Q0EWF0 |
| Vigna radiata | A0A3Q0F0F3 |
| Vigna radiata | A0A3Q0FGP8 |
| Vigna radiata | A0A1S3VBZ6 |
| Vitis vinifera. | A5C4W8 |
| Vitis vinifera. | A0A438HK40 |
| Vitis vinifera. | F6GZ95 |
| Vitis vinifera. | A0A438HSA8 |
| Vitis vinifera. | A0A438K4K8 |
| Vitis vinifera. | F6H2A3 |
| Vitis vinifera. | F6H2N4 |
| Vitis vinifera. | A0A438HPG9 |
| Vitis vinifera. | F6GXQ1 |
| Vitis vinifera. | F6GYD8 |
| Vitis vinifera. | F6HRT0 |
| Vitis vinifera. | A5BHL6 |
| Vitis vinifera. | A5B7V9 |
| Vitis vinifera. | A0A438IDB7 |
| Vitis vinifera. | A0A438JV19 |
| Vitis vinifera. | A0A438J4D4 |
| Vitis vinifera. | A0A438J4P8 |
| Vitis vinifera. | F6H2B3 |
| Vitis vinifera. | F6HL29 |
| Vitis vinifera. | A5BJG8 |
| Vitis vinifera. | A5AH51 |
| Vitis vinifera. | A5BYN3 |
| Vitis vinifera. | F6H549 |
| Vitis vinifera. | F6HD38 |
| Vitis vinifera. | F6HD40 |
| Vitis vinifera. | F6HL32 |
| Vitis vinifera. | A5BFJ9 |
| Vitis vinifera. | A5ALH9 |
| Vitis vinifera. | F6I0H1 |
| Vitis vinifera. | F6GZ96 |
| Vitis vinifera. | F6H296 |
| Vitis vinifera. | D7U4I0 |
| Vitis vinifera. | F6I7L4 |
| Vitis vinifera. | A5ACT5 |
| Vitis vinifera. | F6H5E3 |

| | |
|---|---|
| Vitis vinifera. | A5BH08 |
| Vitis vinifera. | F6H5E2 |
| Vitis vinifera. | A5BRV2 |
| Vitis vinifera. | A5AD43 |
| Vitis vinifera. | F6HW07 |
| Vitis vinifera. | F6HL33 |
| Vitis vinifera. | F6GYE5 |
| Vitis vinifera. | A0A438G9R9 |
| Vitis vinifera. | A0A438GDM0 |
| Vitis vinifera. | A0A438H0P0 |
| Vitis vinifera. | A5ASQ2 |
| Vitis vinifera. | F6GXQ3 |
| Vitis vinifera. | F6HJX6 |
| Vitis vinifera. | F6GZ98 |
| Vitis vinifera. | A5BY61 |
| Vitis vinifera. | A5B986 |
| Vitis vinifera. | D7T5C7 |
| Vitis vinifera. | A5C5N7 |
| Vitis vinifera. | E0CSX4 |
| Vitis vinifera. | F6H7Z9 |
| Vitis vinifera. | D7TJ25 |
| Vitis vinifera. | A5BIS9 |
| Vitis vinifera. | F6H543 |
| Vitis vinifera. | F6GZX7 |
| Vitis vinifera. | A5BXA4 |
| Vitis vinifera. | F6H1X1 |
| Vitis vinifera. | A5BQP9 |
| Vitis vinifera. | F6H2A5 |
| Vitis vinifera. | A5BH06 |
| Vitis vinifera. | D7TRQ9 |
| Vitis vinifera. | A0A438HNC4 |
| Vitis vinifera. | A5BKS7 |
| Vitis vinifera. | D7UAZ5 |
| Vitis vinifera. | D7U6R5 |
| Vitis vinifera. | A5ALK0 |
| Vitis vinifera. | F6HL31 |
| Vitis vinifera. | F6H2G3 |
| Vitis vinifera. | A5AQY5 |
| Vitis vinifera. | F6HYF4 |
| Vitis vinifera. | F6H2B6 |
| Vitis vinifera. | F6I2E8 |

| | |
|---|---|
| Vitis vinifera. | E0CSZ7 |
| Vitis vinifera. | F6H2G5 |
| Vitis vinifera. | F6HL30 |
| Vitis vinifera. | F6I1Y7 |
| Vitis vinifera. | A5BRU4 |
| Vitis vinifera. | A5BQQ0 |
| Vitis vinifera. | F6H298 |
| Vitis vinifera. | F6H2C0 |
| Vitis vinifera. | A5C1U2 |
| Vitis vinifera. | F6GZ97 |
| Vitis vinifera. | F6HH96 |
| Vitis vinifera. | F6H2A9 |
| Vitis vinifera. | F6HW06 |
| Vitis vinifera. | A5BXZ3 |
| Vitis vinifera. | A0A438EUR3 |
| Vitis vinifera. | A0A438DP60 |
| Vitis vinifera. | A0A438IDB3 |
| Vitis vinifera. | F6GYG6 |
| Vitis vinifera. | A0A438G921 |
| Vitis vinifera. | A0A438BSL5 |
| Vitis vinifera. | A0A438G9P6 |
| Vitis vinifera. | A0A438HN97 |
| Vitis vinifera. | A0A438GDR0 |
| Vitis vinifera. | A5AG42 |
| Vitis vinifera. | A0A438GF75 |
| Vitis vinifera. | A0A438HLK8 |
| Vitis vinifera. | A0A438G9V2 |
| Vitis vinifera. | A0A438G9Q8 |
| Vitis vinifera. | A0A438H0K0 |
| Vitis vinifera. | A0A438ID79 |
| Vitis vinifera. | A0A438HSB1 |
| Vitis vinifera. | A0A438HK35 |
| Vitis vinifera. | A0A438G270 |
| Vitis vinifera. | A0A438G9S5 |
| Vitis vinifera. | A0A438HAT9 |
| Vitis vinifera. | A5BHL5 |
| Vitis vinifera. | A5AUE6 |
| Vitis vinifera. | A0A438BPR1 |
| Vitis vinifera. | F6H559 |
| Vitis vinifera. | A0A438BSE3 |
| Vitis vinifera. | A0A438DBA0 |

| | |
|---|---|
| Vitis vinifera. | A0A438CR06 |
| Vitis vinifera. | A0A438BPQ2 |
| Vitis vinifera. | A0A438G973 |
| Vitis vinifera. | A0A438JAW2 |
| Vitis vinifera. | A0A438FCT8 |
| Vitis vinifera. | A0A438HSB6 |
| Vitis vinifera. | A0A438JS26 |
| Vitis vinifera. | A0A438EA03 |
| Vitis vinifera. | A0A438GV90 |
| Vitis vinifera. | A0A438CSM6 |
| Vitis vinifera. | A0A438HS93 |
| Vitis vinifera. | A0A438BPP3 |
| Vitis vinifera. | A0A438GVE3 |
| Vitis vinifera. | A0A438CK55 |
| Vitis vinifera. | A0A438G8Z3 |
| Vitis vinifera. | A0A438BSB7 |
| Vitis vinifera. | A0A438ITA4 |
| Vitis vinifera. | A0A438K4I2 |
| Vitis vinifera. | A0A438BSD0 |
| Vitis vinifera. | A0A438FQF7 |
| Vitis vinifera. | A0A438C0M0 |
| Vitis vinifera. | A0A438GDR3 |
| Vitis vinifera. | A0A438BSF9 |
| Vitis vinifera. | A0A438G3D9 |
| Vitis vinifera. | A0A438KNQ7 |
| Vitis vinifera. | A0A438G911 |
| Vitis vinifera. | A5BHL9 |
| Vitis vinifera. | A0A438EGN8 |
| Vitis vinifera. | F6H2A8 |
| Vitis vinifera. | A0A438J4V0 |
| Vitis vinifera. | A0A438K4L8 |
| Vitis vinifera. | A0A438GDM4 |
| Vitis vinifera. | A0A438DY93 |
| Vitis vinifera. | A0A438HN88 |
| Vitis vinifera. | A0A438HN57 |
| Vitis vinifera. | A0A438GVF0 |
| Vitis vinifera. | A0A438GDM2 |
| Vitis vinifera. | A0A438GVA2 |
| Vitis vinifera. | A0A438BS99 |
| Vitis vinifera. | A0A438DBE8 |
| Vitis vinifera. | A0A438JAQ1 |

| | |
|---|---|
| Vitis vinifera. | A0A438DP71 |
| Vitis vinifera. | A0A438BS80 |
| Vitis vinifera. | F6H2A0 |
| Vitis vinifera. | A0A438G9S9 |
| Vitis vinifera. | A0A438HU56 |
| Vitis vinifera. | A0A438BPN8 |
| Vitis vinifera. | A0A438HS63 |
| Vitis vinifera. | F6GXQ9 |
| Vitis vinifera. | A0A438GVC3 |
| Vitis vinifera. | A0A438FWU9 |
| Vitis vinifera. | A0A438GDN5 |
| Vitis vinifera. | A0A438GDN4 |
| Vitis vinifera. | A0A438GDQ3 |
| Vitis vinifera. | A0A438EUS9 |
| Vitis vinifera. | A0A438E5D3 |
| Vitis vinifera. | A0A438I408 |
| Vitis vinifera. | A0A438G271 |
| Vitis vinifera. | A0A438DE69 |
| Vitis vinifera. | F6GYD9 |
| Vitis vinifera. | A0A438HPI4 |
| Vitis vinifera. | A0A438BPY9 |
| Vitis vinifera. | A0A438BSI7 |
| Vitis vinifera. | A0A438I405 |
| Vitis vinifera. | A0A438BS96 |
| Vitis vinifera. | A0A438K4H2 |
| Vitis vinifera. | A0A438GDQ0 |
| Vitis vinifera. | A0A438C054 |
| Vitis vinifera. | A0A438H4L4 |
| Vitis vinifera. | A0A438G257 |
| Vitis vinifera. | A0A438BSC6 |
| Vitis vinifera. | A0A438G9Q4 |
| Vitis vinifera. | A0A438BPQ7 |
| Vitis vinifera. | A0A438HSC6 |
| Vitis vinifera. | A0A438GVB8 |
| Vitis vinifera. | A0A438IDD7 |
| Vitis vinifera. | A0A438HPK3 |
| Vitis vinifera. | A0A438G9Q3 |
| Vitis vinifera. | A0A438BSC1 |
| Vitis vinifera. | A0A438G5Q0 |
| Vitis vinifera. | A0A438BSM6 |
| Vitis vinifera. | A0A438BSC0 |

| | |
|---|---|
| Vitis vinifera. | A0A438DP46 |
| Vitis vinifera. | A0A438DGJ9 |
| Vitis vinifera. | F6GYE3 |
| Vitis vinifera. | A0A438JLB1 |
| Vitis vinifera. | A0A438IDA7 |
| Vitis vinifera. | A0A438GVD0 |
| Vitis vinifera. | A0A438GDQ6 |
| Vitis vinifera. | A0A438DF00 |
| Vitis vinifera. | A0A438DP50 |
| Vitis vinifera. | A0A438G9P1 |
| Vitis vinifera. | A0A438GBQ1 |
| Vitis vinifera. | A0A438JKA7 |
| Vitis vinifera. | A0A438KF31 |
| Vitis vinifera. | A0A438IDB0 |
| Vitis vinifera. | A0A438G907 |
| Vitis vinifera. | A0A438H6B4 |
| Vitis vinifera. | A0A438GV93 |
| Vitis vinifera. | A0A438G943 |
| Vitis vinifera. | A0A438CYC5 |
| Vitis vinifera. | A0A438BSE2 |
| Vitis vinifera. | A0A438BSD9 |
| Vitis vinifera. | A0A438DB91 |
| Vitis vinifera. | A0A438G9Q7 |
| Vitis vinifera. | A0A438BSA2 |
| Vitis vinifera. | A0A438GVA1 |
| Vitis vinifera. | A0A438GDN6 |
| Vitis vinifera. | A0A438FCS0 |
| Vitis vinifera. | A5AQ78 |
| Vitis vinifera. | A0A438C802 |
| Vitis vinifera. | A0A438HLD8 |
| Vitis vinifera. | A0A438HZ10 |
| Vitis vinifera. | A0A438GVD3 |
| Vitis vinifera. | A0A438G908 |
| Vitis vinifera. | A0A438GVB4 |
| Vitis vinifera. | A5BLU0 |
| Vitis vinifera. | A0A438HPJ7 |
| Vitis vinifera. | A0A438G5P6 |
| Vitis vinifera. | A0A438DE77 |
| Vitis vinifera. | A0A438C111 |
| Vitis vinifera. | A0A438HCT6 |
| Vitis vinifera. | D7SRN6 |

| | |
|---|---|
| Vitis vinifera. | F6GXQ6 |
| Vitis vinifera. | A5BKZ4 |
| Vitis vinifera. | A0A438BSD1 |
| Vitis vinifera. | F6H2G7 |
| Vitis vinifera. | F6GZX9 |
| Vitis vinifera. | F6GZD2 |
| Vitis vinifera. | F6H294 |
| Vitis vinifera. | F6I0I1 |
| Vitis vinifera. | A5ACT7 |
| Vitis vinifera. | A5BRU5 |
| Vitis vinifera. | A5BVG8 |
| Vitis vinifera. | F6H542 |
| Vitis vinifera. | F6HL34 |
| Vitis vinifera. | F6I2A7 |
| Vitis vinifera. | F6H291 |
| Vitis vinifera. | E0CT00 |
| Vitis vinifera. | A5AQI0 |
| Vitis vinifera. | A5C9P3 |
| Vitis vinifera. | F6HFB8 |
| Vitis vinifera. | F6GXQ0 |
| Vitis vinifera. | A5BGQ9 |
| Vitis vinifera. | E0CSJ3 |
| Vitis vinifera. | F6HRS4 |
| Vitis vinifera. | A5C0J8 |
| Vitis vinifera. | F6HD39 |
| Vitis vinifera. | F6H801 |
| Vitis vinifera. | F6I1Y9 |
| Vitis vinifera. | F6H2B7 |
| Vitis vinifera. | F6GZX6 |
| Vitis vinifera. | F6H2G6 |
| Vitis vinifera. | F6GYE4 |
| Vitis vinifera. | A5BEU8 |
| Vitis vinifera. | E0CT03 |
| Vitis vinifera. | F6H295 |
| Vitis vinifera. | F6I2E9 |
| Vitis vinifera. | A0A438GVB5 |
| Vitis vinifera. | A0A438DPA7 |
| Vitis vinifera. | A0A438DP66 |
| Vitis vinifera. | A0A438HAT4 |
| Vitis vinifera. | A0A438CQW8 |
| Vitis vinifera. | A0A438HLG1 |

| | |
|---|---|
| Vitis vinifera. | A5C437 |
| Vitis vinifera. | F6GZX1 |
| Vitis vinifera. | A0A438DBZ7 |
| Vitis vinifera. | A5BEK3 |
| Vitis vinifera. | D7TK53 |
| Vitis vinifera. | F6HYF3 |
| Vitis vinifera. | F6H329 |
| Vitis vinifera. | A5BIW6 |
| Vitis vinifera. | A0A438D3S7 |
| Vitis vinifera. | A5APY4 |
| Vitis vinifera. | F6HL08 |
| Vitis vinifera. | F6H2A2 |
| Vitis vinifera. | A5BHL4 |
| Vitis vinifera. | A0A438EA11 |
| Vitis vinifera. | A0A438CFB9 |
| Vitis vinifera. | A0A438G8X0 |
| Vitis vinifera. | A0A438EAE1 |
| Vitis vinifera. | A0A438GDP7 |
| Vitis vinifera. | A0A438BPQ9 |
| Vitis vinifera. | A0A438JP69 |
| Vitis vinifera. | A0A438HLN0 |
| Vitis vinifera. | A0A438G975 |
| Vitis vinifera. | A0A438HPK0 |
| Vitis vinifera. | A0A438BSA0 |
| Vitis vinifera. | A0A438FHJ0 |
| Vitis vinifera. | A0A438G906 |
| Vitis vinifera. | F6GXR2 |
| Vitis vinifera. | A0A438G919 |
| Vitis vinifera. | A0A438F254 |
| Vitis vinifera. | A0A438EGP7 |
| Vitis vinifera. | A0A438BS88 |
| Vitis vinifera. | A0A438HS95 |
| Vitis vinifera. | A0A438GVD6 |
| Vitis vinifera. | A0A438E9X1 |
| Vitis vinifera. | A5AWE1 |
| Vitis vinifera. | A0A438GVB1 |
| Vitis vinifera. | A0A438G952 |
| Vitis vinifera. | A0A438IDD5 |
| Vitis vinifera. | A0A438G9P4 |
| Vitis vinifera. | A0A438BSF2 |
| Vitis vinifera. | A0A438G5Q1 |

| | |
|---|---|
| Vitis vinifera. | A0A438HS43 |
| Vitis vinifera. | A0A438JK94 |
| Vitis vinifera. | A0A438JAS5 |
| Vitis vinifera. | A0A438ID88 |
| Vitis vinifera. | A0A438ITJ2 |
| Vitis vinifera. | A0A438HLS7 |
| Vitis vinifera. | A0A438DP63 |
| Vitis vinifera. | A0A438HPH4 |
| Vitis vinifera. | A0A438BPR7 |
| Vitis vinifera. | A0A438G918 |
| Vitis vinifera. | A0A438JAY5 |
| Vitis vinifera. | A0A438G9T4 |
| Vitis vinifera. | A5BFK0 |
| Vitis vinifera. | A0A438BSE1 |
| Vitis vinifera. | A5C187 |
| Vitis vinifera. | F6H290 |
| Vitis vinifera. | F6I1Y8 |
| Vitis vinifera. | A5AS83 |
| Vitis vinifera. | A5C1Y4 |
| Vitis vinifera. | F6H292 |
| Vitis vinifera. | A5AST3 |
| Vitis vinifera. | F6HCT4 |
| Vitis vinifera. | A0A438HS08 |
| Vitis vinifera. | A0A438IT97 |
| Vitrella brass | A0A0G4H1K6 |
| Vitrella brass | A0A0G4FPW0 |
| Vitrella brass | A0A0G4EVW8 |
| Vitrella brass | A0A0G4G2P6 |
| Vitrella brass | A0A0G4EX32 |
| Vitrella brass | A0A0G4EK41 |
| Vitrella brass | A0A0G4GV94 |
| Vitrella brass | A0A0G4FK32 |
| Vitrella brass | A0A0G4FSG0 |
| Vitrella brass | A0A0G4GBE1 |
| Vitrella brass | A0A0G4FEA5 |
| Vitrella brass | A0A0G4G667 |
| Vitrella brass | A0A0G4G5L3 |
| Vitrella brass | A0A0G4GUX0 |
| Vitrella brass | A0A0G4H7X7 |
| Vitrella brass | A0A0G4FED4 |
| Vitrella brass | A0A0G4GC70 |

| | |
|---|---|
| Vitrella brass | A0A0G4EL21 |
| Vitrella brass | A0A0G4EFA5 |
| Vitrella brass | A0A0G4FS69 |
| Vitrella brass | A0A0G4G6X5 |
| Vitrella brass | A0A0G4GSW5 |
| Vitrella brass | A0A0G4FB70 |
| Vitrella brass | A0A0G4FJV3 |
| Vitrella brass | A0A0G4ELW9 |
| Vitrella brass | A0A0G4F302 |
| Vitrella brass | A0A0G4H591 |
| Vitrella brass | A0A0G4GFR4 |
| Vitrella brass | A0A0G4FNT8 |
| Vitrella brass | A0A0G4H797 |
| Vitrella brass | A0A0G4FKH8 |
| Vitrella brass | A0A0G4GBQ5 |
| Vitrella brass | A0A0G4G8Z9 |
| Vitrella brass | A0A0G4FBG5 |
| Vitrella brass | A0A0G4FUC6 |
| Vitrella brass | A0A0G4EES0 |
| Vitrella brass | A0A0G4GY87 |
| Vitrella brass | A0A0G4FN79 |
| Vitrella brass | A0A0G4FG97 |
| Vitrella brass | A0A0G4GU04 |
| Vitrella brass | A0A0G4GAX4 |
| Vitrella brass | A0A0G4EKA7 |
| Vitrella brass | A0A0G4E9C4 |
| Vitrella brass | A0A0G4GA99 |
| Vitrella brass | A0A0G4GC21 |
| Vitrella brass | A0A0G4F4H5 |
| Vitrella brass | A0A0G4GTL7 |
| Vitrella brass | A0A0G4EC69 |
| Vitrella brass | A0A0G4EDQ0 |
| Vitrella brass | A0A0G4F1H0 |
| Vitrella brass | A0A0G4GX24 |
| Vitrella brass | A0A0G4GSZ4 |
| Vitrella brass | A0A0G4GQI0 |
| Vitrella brass | A0A0G4FAD2 |
| Vitrella brass | A0A0G4EYY7 |
| Vitrella brass | A0A0G4ECR3 |
| Vitrella brass | A0A0G4FRN1 |
| Vitrella brass | A0A0G4F3I1 |

| | |
|---|---|
| Vitrella brass | A0A0G4GJ27 |
| Vitrella brass | A0A0G4FWK0 |
| Vitrella brass | A0A0G4FT88 |
| Vitrella brass | A0A0G4ER04 |
| Vitrella brass | A0A0G4FH06 |
| Vitrella brass | A0A0G4GJ92 |
| Vitrella brass | A0A0G4EN10 |
| Vitrella brass | A0A0G4FC64 |
| Vitrella brass | A0A0G4G604 |
| Vitrella brass | A0A0G4H617 |
| Vitrella brass | A0A0G4EE30 |
| Vitrella brass | A0A0G4EMH2 |
| Vitrella brass | A0A0G4EXD3 |
| Vitrella brass | A0A0G4F467 |
| Vitrella brass | A0A0G4ESG9 |
| Vitrella brass | A0A0G4H5A7 |
| Vitrella brass | A0A0G4H147 |
| Vitrella brass | A0A0G4ELP5 |
| Vitrella brass | A0A0G4EW80 |
| Vitrella brass | A0A0G4EYN3 |
| Vitrella brass | A0A0G4EVX9 |
| Vitrella brass | A0A0G4G194 |
| Vitrella brass | A0A0G4FYF0 |
| Vitrella brass | A0A0G4ENT8 |
| Vitrella brass | A0A0G4E913 |
| Vitrella brass | A0A0G4EZ90 |
| Vitrella brass | A0A0G4GDZ1 |
| Vitrella brass | A0A0G4EV77 |
| Vitrella brass | A0A0G4F2L7 |
| Vitrella brass | A0A0G4FW07 |
| Vitrella brass | A0A0G4END2 |
| Vitrella brass | A0A0G4G025 |
| Vitrella brass | A0A0G4EJ16 |
| Vitrella brass | A0A0G4G3W1 |
| Vitrella brass | A0A0G4ED81 |
| Vitrella brass | A0A0G4EA18 |
| Vitrella brass | A0A0G4GZE5 |
| Vitrella brass | A0A0G4FZK6 |
| Vitrella brass | A0A0G4F356 |
| Vitrella brass | A0A0G4GY92 |
| Vitrella brass | A0A0G4EP61 |

| | |
|---|---|
| Vitrella brass | A0A0G4ERN6 |
| Vitrella brass | A0A0G4F257 |
| Vitrella brass | A0A0G4FG25 |
| Vitrella brass | A0A0G4GJ87 |
| Vitrella brass | A0A0G4GFW8 |
| Vitrella brass | A0A0G4GN70 |
| Vitrella brass | A0A0G4EGD8 |
| Vitrella brass | A0A0G4GV35 |
| Vitrella brass | A0A0G4EDK7 |
| Vitrella brass | A0A0G4FDV3 |
| Vitrella brass | A0A0G4EN36 |
| Vitrella brass | A0A0G4FU39 |
| Vitrella brass | A0A0G4G6I2 |
| Vitrella brass | A0A0G4FAX9 |
| Volvox carter | Q948Y6 |
| Volvox carter | D8U662 |
| Volvox carter | D8UDJ2 |
| Volvox carter | D8UHS7 |
| Volvox carter | D8UHS1 |
| Volvox carter | D8U663 |
| Volvox carter | D8TZF2 |
| Volvox carter | D8TWV3 |
| Volvox carter | D8TZP2 |
| Volvox carter | D8ULZ4 |
| Volvox carter | D8U0L1 |
| Volvox carter | D8THZ2 |
| Volvox carter | D8U661 |
| Volvox carter | D8U9Q0 |
| Volvox carter | D8TUD4 |
| Volvox carter | D8TWS3 |
| Volvox carter | D8U5V3 |
| Volvox carter | D8TWS6 |
| Volvox carter | D8U656 |
| Volvox carter | D8UHS6 |
| Volvox carter | D8U0K8 |
| Volvox carter | D8TWV6 |
| Volvox carter | D8U3S1 |
| Vulpes vulpes | A0A3Q7TWA6 |
| Vulpes vulpes | A0A3Q7T0P7 |
| Vulpes vulpes | A0A3Q7SNP7 |
| Vulpes vulpes | A0A3Q7V334 |

| | |
|---|---|
| Vulpes vulpes | A0A3Q7TKS7 |
| Vulpes vulpes | A0A3Q7TAD2 |
| Vulpes vulpes | A0A3Q7TW96 |
| Vulpes vulpes | A0A3Q7UEY3 |
| Wenxinia ma | A0A0D0QEL7 |
| Wenxinia ma | A0A0D0PB00 |
| Wenxinia sae | A0A1M6EL45 |
| Wollemia nol | A0A0C9S462 |
| Wollemia nol | A0A0C9QPG3 |
| Wollemia nol | A0A0C9RV88 |
| Wollemia nol | A0A0C9RSU0 |
| Wollemia nol | A0A0C9QPV3 |
| Wollemia nol | A0A0C9RSN7 |
| Wollemia nol | A0A0C9S6M8 |
| Wuchereria b | A0A1I8E9A3 |
| Wuchereria b | J9DND2 |
| Wuchereria b | A0A3P7DL77 |
| Wuchereria b | A0A1I8ELN8 |
| Wuchereria b | A0A3P7EWD2 |
| Wuchereria b | A0A1I8EZC2 |
| Wuchereria b | J9ETT3 |
| Wuchereria b | A0A1I8EY91 |
| Wuchereria b | A0A1I8EL26 |
| Wuchereria b | A0A1I8EZH7 |
| Wuchereria b | A0A3P7GM02 |
| Wuchereria b | J9DZA9 |
| Wuchereria b | A0A3P7FT60 |
| Wuchereria b | A0A3P7GCY4 |
| Wuchereria b | J9E1R8 |
| Wuchereria b | A0A1I8EFL2 |
| Wuchereria b | A0A1I8EF35 |
| Wuchereria b | A0A3P7EEQ9 |
| Wuchereria b | A0A3P7FCC9 |
| Wuchereria b | A0A3P7DVJ1 |
| Wuchereria b | A0A3P7DV86 |
| Wuchereria b | A0A1I8ET18 |
| Wuchereria b | A0A3P7DDB0 |
| Wuchereria b | A0A3P7DLR8 |
| Wuchereria b | A0A1I8F0Y1 |
| Wuchereria b | J9F728 |
| Wuchereria b | J9F5K7 |

| | |
|---|---|
| Wuchereria k | A0A1I8ELS4 |
| Wuchereria k | A0A3P7EE66 |
| Wuchereria k | A0A1I8EA12 |
| Wuchereria k | J9ELM2 |
| Wuchereria k | J9EVZ5 |
| Wuchereria k | J9E6Y8 |
| Wuchereria k | J9ESJ0 |
| Wuchereria k | J9AHN9 |
| Wuchereria k | A0A3P7GJ34 |
| Wuchereria k | A0A1I8ESS3 |
| Wuchereria k | J9E0L8 |
| Wuchereria k | J9EHN5 |
| Wuchereria k | A0A1I8ECU1 |
| Wuchereria k | J9FCK3 |
| Wuchereria k | A0A3P7FKH0 |
| Wuchereria k | A0A3P7EG90 |
| Wuchereria k | J9F005 |
| Wuchereria k | J9FHZ7 |
| Wuchereria k | J9F0L7 |
| Wuchereria k | J9EDU4 |
| Wuchereria k | A0A1I8EPI5 |
| Wuchereria k | A0A1I8ENJ3 |
| Wuchereria k | A0A1I8EHS2 |
| Wuchereria k | A0A3P7E3Y0 |
| Wuchereria k | A0A1I8ER89 |
| Wuchereria k | J9AZF1 |
| Wuchereria k | J9B8V2 |
| Xenopus laev | A0A1L8G2F6 |
| Xenopus laev | Q91691 |
| Xenopus laev | A0A1L8GHB0 |
| Xenopus laev | A0A1L8G2M6 |
| Xenopus laev | A0A1L8GP52 |
| Xenopus laev | Q7ZTN9 |
| Xenopus laev | A0A1L8GTY1 |
| Xenopus laev | A0A1L8GZD1 |
| Xenopus laev | Q6GP14 |
| Xenopus laev | Q6AZS7 |
| Xenopus laev | P70006 |
| Xenopus trop | F7EDW8 |
| Xenopus trop | F6YRB8 |
| Xenopus trop | F7AR31 |

| | |
|---|---|
| Xenopus.xlsx | Q91402 |
| Xinfangfangia | A0A3P5XFD2 |
| Xiphophorus | A0A3B5KRY2 |
| Xiphophorus | A0A3B5KXI9 |
| Xiphophorus | A0A3B5KR76 |
| Xiphophorus | A0A3B5MPU6 |
| Xiphophorus | A0A3B5M999 |
| Xiphophorus | A0A3B5KPF4 |
| Xiphophorus | A0A3B5KR79 |
| Xiphophorus | A0A3B5KQW4 |
| Xiphophorus | A0A3B5MUP9 |
| Xiphophorus | A0A3B5M3B3 |
| Xiphophorus | A0A3B5MDS7 |
| Xiphophorus | A0A3B5KJF0 |
| Xiphophorus | A0A3B5MW25 |
| Xiphophorus | M4ASA0 |
| Xiphophorus | A0A3B5R264 |
| Xiphophorus | M4AQQ5 |
| Xiphophorus | A0A3B5R3X9 |
| Xiphophorus | M4AAJ3 |
| Xiphophorus | M3ZXK8 |
| Xiphophorus | M3ZTS0 |
| Xiphophorus | M3ZXL1 |
| Xiphophorus | M3ZXJ9 |
| Xiphophorus | M3ZQC7 |
| Xiphophorus | A0A3B5PQ81 |
| Xylaria gramr | A0A439DFT7 |
| Xylaria gramr | A0A439CN67 |
| Xylaria gramr | A0A439CTF6 |
| Yangia sp..xls | A0A2E1D5Y3 |
| Yoonia sedim | A0A2T6KMQ7 |
| Yoonia tamle | A0A1I6G2U0 |
| Zea mays.xls› | A0A3L6D9L1 |
| Zea mays.xls› | A0A3L6E7F7 |
| Zea mays.xls› | A0A317YAC8 |
| Zea mays.xls› | A0A3L6DRV9 |
| Zea mays.xls› | A0A3L6E476 |
| Zea mays.xls› | A0A317YET9 |
| Zea mays.xls› | A0A3L6E218 |
| Zea mays.xls› | A0A317YKF8 |
| Zea mays.xls› | B6TY46 |

Zea mays.xls› A0A317Y521
Zea mays.xls› P17801
Zea mays.xls› A0A1D6KQI4
Zea mays.xls› C0P6F5
Zea mays.xls› A0A1D6KH09
Zea mays.xls› A0A096T7Z1
Zea mays.xls› A0A1D6M0L4
Zea mays.xls› A0A1D6LBB2
Zea mays.xls› A0A1D6DXI3
Zea mays.xls› A0A1D6F8W0
Zea mays.xls› A0A1D6N9S7
Zea mays.xls› A0A1D6DXH8
Zea mays.xls› A0A1D6H9Q0
Zea mays.xls› A0A1D6GN29
Zea mays.xls› K7UZN2
Zea mays.xls› A0A1D6IEQ3
Zea mays.xls› A0A1D6P507
Zea mays.xls› A0A1D6DXM7
Zea mays.xls› A0A1D6J6V6
Zea mays.xls› K7VD35
Zea mays.xls› A0A1D6IAX9
Zea mays.xls› C0P9N1
Zea mays.xls› A0A1D6JF78
Zea mays.xls› A0A1D6GA13
Zea mays.xls› A0A1D6FGJ5
Zea mays.xls› K7VRP4
Zea mays.xls› A0A1D6DXM5
Zea mays.xls› A0A1D6ML66
Zea mays.xls› A0A1D6GA09
Zea mays.xls› A0A1D6N9S3
Zea mays.xls› A0A1D6HLB4
Zea mays.xls› A0A1D6IER4
Zea mays.xls› A0A1D6N9Q9
Zea mays.xls› C0HFM8
Zea mays.xls› A0A1D6KIF9
Zea mays.xls› A0A1D6IEQ5
Zea mays.xls› A0A1D6GA12
Zea mays.xls› A0A1D6KAQ9
Zea mays.xls› A0A1D6IEI3
Zea mays.xls› A0A1D6FSS8
Zea mays.xls› A0A1D6IEI1

Zea mays.xls› A0A1D6N9R8
Zea mays.xls› A0A1D6MFC2
Zea mays.xls› A0A1D6M2E6
Zea mays.xls› A0A1D6J6V7
Zea mays.xls› A0A3L6E4L3
Zea mays.xls› A0A1D6DXH6
Zea mays.xls› A0A1D6J6V1
Zea mays.xls› A0A1D6J6V2
Zea mays.xls› A0A1D6M0N6
Zea mays.xls› A0A1D6EWT3
Zea mays.xls› B4G1J2
Zea mays.xls› A0A1D6ITQ0
Zea mays.xls› A0A1D6KQI3
Zea mays.xls› A0A1D6KL97
Zea mays.xls› A0A1D6N9R3
Zea mays.xls› K7UIF8
Zea mays.xls› A0A1D6M0N7
Zea mays.xls› A0A1D6EB48
Zea mays.xls› A0A1D6QAA8
Zea mays.xls› A0A1D6P2B1
Zea mays.xls› A0A1D6M0N8
Zea mays.xls› A0A3L6DU79
Zea mays.xls› A0A3L6EU45
Zea mays.xls› A0A317Y2S5
Zea mays.xls› B7ZXB0
Zea mays.xls› A0A3L6FU88
Zea mays.xls› A0A3L6DL74
Zea mays.xls› B6U2B7
Zea mays.xls› A0A3L6FAJ7
Zea mays.xls› A0A3L6DY96
Zea mays.xls› A0A1Q1B749
Zea mays.xls› A0A3L6ET80
Zea mays.xls› B6SWL2
Zea mays.xls› A0A3L6GBB9
Zea mays.xls› A0A3L6G7X6
Zea mays.xls› A0A3L6E4K1
Zea mays.xls› A0A3L6D8A7
Zea mays.xls› A0A3L6DCB9
Zea mays.xls› A0A3L6DDA9
Zea mays.xls› B4F952
Zea mays.xls› A0A3L6G155

Zea mays.xls› O24571
Zea mays.xls› A0A3L6G2G8
Zea mays.xls› B6ST43
Zea mays.xls› A0A3L6FWI8
Zea mays.xls› A0A3L6EU60
Zea mays.xls› B4FYC6
Zea mays.xls› A0A1V0E7H4
Zea mays.xls› B4F8X7
Zea mays.xls› A0A1D6M0M0
Zea mays.xls› A0A1D6KIH3
Zea mays.xls› A0A1D6N9S8
Zea mays.xls› A0A1D6G7A2
Zea mays.xls› A0A1D6KNU3
Zea mays.xls› A0A1D6P015
Zea mays.xls› K7TKD2
Zea mays.xls› C0P3V1
Zea mays.xls› A0A3L6G196
Zea mays.xls› A0A1D6DXM6
Zea mays.xls› A0A1D6HWK7
Zea mays.xls› A0A1D6DXF1
Zea mays.xls› A0A1D6IEQ4
Zea mays.xls› A0A1D6KII4
Zea mays.xls› B4FB05
Zea mays.xls› A0A1D6IBG7
Zea mays.xls› A0A1D6DXI2
Zea mays.xls› A0A1D6N9R2
Zea mays.xls› K7UC64
Zea mays.xls› A0A1D6G136
Zea mays.xls› A0A1D6LHG9
Zea mays.xls› A0A1D6Q5J8
Zea mays.xls› A0A1D6DXH9
Zea mays.xls› A0A1D6FSS9
Zea mays.xls› A0A1D6DXI7
Zea mays.xls› A0A1D6G137
Zea mays.xls› A0A1D6IEI2
Zea mays.xls› A0A1D6JTK4
Zea mays.xls› A0A1D6JF77
Zea mays.xls› K7U3E7
Zea mays.xls› A0A1D6QHL5
Zea mays.xls› A0A1D6HLB7
Zea mays.xls› A0A1D6I5R1

Zea mays.xls› A0A1D6GA10
Zea mays.xls› A0A1D6LGF6
Zea mays.xls› A0A1D6DXM4
Zea mays.xls› A0A1D6EUL5
Zea mays.xls› A0A317YAM4
Zea mays.xls› A0A3L6ETW0
Zea mays.xls› A0A3L6E3M3
Zea mays.xls› O49974
Zea mays.xls› A0A3L6FYL3
Zea mays.xls› A0A3L6EW75
Zea mays.xls› A0A317Y7G4
Zea mays.xls› A0A317Y3N4
Zea mays.xls› A0A3L6FI05
Zea mays.xls› A0A3L6DN41
Zea mays.xls› C0PE10
Zea mays.xls› A0A3L6G6F6
Zea mays.xls› A0A3L6G1E6
Zea mays.xls› A0A3L6DU70
Zea mays.xls› C0HEN9
Zea mays.xls› A0A3L6G8Y0
Zea mays.xls› A0A3L6FT98
Zea mays.xls› A0A3L6EX22
Zea mays.xls› A0A3L6EHP0
Zea mays.xls› O82103
Zea mays.xls› A0A3L6DLS3
Zea mays.xls› B7ZXT8
Zea mays.xls› A0A3L6F189
Zea mays.xls› A0A3L6FL65
Zea mays.xls› A0A3L6E7I6
Zea mays.xls› B7ZWT1
Zea mays.xls› A0A3L6G679
Zea mays.xls› A0A3L6DII2
Zea mays.xls› A0A3L6FF64
Zea mays.xls› A0A3L6DKE6
Zea mays.xls› A0A3L6FXI5
Zea mays.xls› A0A3L6GCE3
Zea mays.xls› A0A3L6FWN1
Zea mays.xls› A0A3L6FGU1
Zea mays.xls› A0A3L6GBY8
Zea mays.xls› A0A3L6EFC7
Zea mays.xls› A0A3L6FT74

| | |
|---|---|
| Zea mays.xls› | A0A3L6EBY7 |
| Zea mays.xls› | A0A317Y6J9 |
| Zea mays.xls› | C0P3U3 |
| Zeugodacus c | A0A0A1XIZ1 |
| Zeugodacus c | A0A0A1XS55 |
| Zeugodacus c | A0A0A1X650 |
| Zeugodacus c | A0A0A1XQ87 |
| Zeugodacus c | A0A0A1X1C0 |
| Zeugodacus c | A0A0A1XB00 |
| Zeugodacus c | A0A0A1X8W7 |
| Zeugodacus c | A0A0A1WII0 |
| Zeugodacus c | A0A0A1XIF3 |
| Zeugodacus c | A0A0A1WSF6 |
| Zhengella ma | A0A2G1QQ59 |
| Zootermopsis | A0A067RG78 |
| Zootermopsis | A0A067RN91 |
| Zootermopsis | A0A067R6A7 |
| Zootermopsis | A0A067RBZ2 |
| Zootermopsis | A0A067RIJ5 |
| Zostera marir | A0A0K9NV78 |
| Zostera marir | A0A0K9Q5Q1 |
| Zostera marir | A0A0K9PK14 |
| Zostera marir | A0A0K9P5Q8 |
| Zostera marir | A0A0K9NYD2 |
| Zostera marir | A0A0K9PEI3 |
| Zostera marir | A0A0K9PPY6 |
| Zostera marir | A0A0K9PXR7 |
| Zostera marir | A0A0K9PLY7 |
| Zostera marir | A0A0K9NWC4 |
| Zostera marir | A0A0K9PKC6 |
| Zostera marir | A0A0K9NQT5 |
| Zostera marir | A0A0K9P3S2 |
| Zostera marir | A0A0K9PQY0 |
| Zostera marir | A0A0K9PLW0 |
| Zostera marir | A0A0K9P3C4 |
| Zostera marir | A0A0K9PPJ9 |
| Zostera marir | A0A0K9P0C4 |
| Zostera marir | A0A0K9PY30 |
| Zostera marir | A0A0K9PNK6 |
| Zostera marir | A0A0K9PP88 |
| Zostera marir | A0A0K9P1W2 |

| | |
|---|---|
| Zostera marir | A0A0K9NS31 |
| Zostera marir | A0A0K9NWR2 |
| Zostera marir | A0A0K9NMN3 |
| Zostera marir | A0A0K9NJ97 |
| Zostera marir | A0A0K9PD87 |
| Zostera marir | A0A0K9NTS5 |
| Zostera marir | A0A0K9Q4R1 |
| Zostera marir | A0A0K9NIZ5 |
| Zostera marir | A0A0K9NQH7 |
| Zostera marir | A0A0K9PG70 |
| Zostera marir | A0A0K9NRX4 |
| Zostera marir | A0A0K9NV14 |
| Zostera marir | A0A0K9NLX9 |
| Zostera marir | A0A0K9P5H7 |
| Zostera marir | A0A0K9NPE2 |
| Zostera marir | A0A0K9P819 |
| Zostera marir | A0A0K9PRP8 |
| Zostera marir | A0A0K9P5Q0 |
| Zostera marir | A0A0K9PM56 |
| Zymoseptoria | A0A0F4GX70 |
| Zymoseptoria | A0A0F4GNN3 |
| Zymoseptoria | A0A0F4GDY3 |
| Zymoseptoria | A0A0F4G7R0 |
| Zymoseptoria | A0A0F4GJL4 |
| Zymoseptoria | A0A0F4G525 |
| Zymoseptoria | A0A0F4GKZ5 |
| Zymoseptoria | A0A0F4GK64 |
| Zymoseptoria | A0A0F4GK12 |
| Zymoseptoria | A0A0F4GVA7 |
| Zymoseptoria | A0A0F4GTQ5 |
| Zymoseptoria | A0A0F4GJJ0 |
| Zymoseptoria | A0A0F4GPQ0 |
| Zymoseptoria | A0A0F4GLI2 |
| Zymoseptoria | F9X4T1 |
| Zymoseptoria | F9XHT4 |
| Zymoseptoria | F9XH72 |
| Zymoseptoria | F9XJG6 |
| Zymoseptoria | F9XDZ3 |
| Zymoseptoria | F9X146 |
| Zymoseptoria | F9XK84 |
| Zymoseptoria | F9XCF6 |

| | |
|---|---|
| Zymoseptoria | F9XP87 |
| Zymoseptoria | F9X6W1 |
| Zymoseptoria | F9XFV9 |
| Zymoseptoria | F9XM39 |
| Zymoseptoria | F9XAK9 |
| Zymoseptoria | F9XNE3 |
| Zymoseptoria | A0A1Y6LP08 |
| Zymoseptoria | A0A1Y6LZ79 |
| Zymoseptoria | A0A1Y6LY30 |
| Zymoseptoria | A0A1Y6LT89 |
| Zymoseptoria | A0A1Y6L814 |
| Zymoseptoria | A0A1Y6LQH5 |
| Zymoseptoria | A0A1Y6LQU6 |
| Zymoseptoria | A0A1Y6L5Q9 |
| Zymoseptoria | A0A1Y6LXG6 |
| Zymoseptoria | A0A1Y6LN68 |
| Zymoseptoria | A0A1Y6LFZ2 |
| Zymoseptoria | A0A1Y6LBN0 |
| Zymoseptoria | A0A1Y6LDV9 |
| Zymoseptoria | A0A1Y6L8C9 |
| Zymoseptoria | A0A1Y6LRA7 |
| Zymoseptoria | A0A2H1FJ88 |
| Zymoseptoria | A0A2H1GP22 |
| Zymoseptoria | A0A2H1H488 |
| Zymoseptoria | A0A2H1FZK2 |
| Zymoseptoria | A0A2H1H533 |
| Zymoseptoria | A0A2H1FLY6 |
| Zymoseptoria | A0A2H1H4S8 |
| Zymoseptoria | A0A2H1GT42 |
| Zymoseptoria | A0A2H1GHG1 |
| Zymoseptoria | A0A2H1H5M8 |
| Zymoseptoria | A0A2H1GTE8 |
| Zymoseptoria | A0A2H1FQ02 |
| Zymoseptoria | A0A2H1G5G9 |
| Zymoseptoria | A0A2H1GXW6 |
| Zymoseptoria | A0A2H1GZK0 |
| Zymoseptoria | A0A2H1GNU0 |
| Zymoseptoria | A0A2H1GST3 |
| Zymoseptoria | A0A2H1H7P5 |
| Zymoseptoria | A0A1X7RYL6 |
| Zymoseptoria | A0A1X7RF14 |

| | |
|---|---|
| Zymoseptoria | A0A1X7RL97 |
| Zymoseptoria | A0A1X7RUL5 |
| Zymoseptoria | A0A1X7S239 |
| Zymoseptoria | A0A1X7RV33 |
| Zymoseptoria | A0A1X7RNL6 |
| Zymoseptoria | A0A1X7RY43 |
| Zymoseptoria | A0A1X7S9D0 |
| Zymoseptoria | A0A1X7S2M2 |
| Zymoseptoria | A0A1X7S7L9 |
| Zymoseptoria | A0A1X7RBY3 |
| Zymoseptoria | A0A1X7S0P9 |
| Zymoseptoria | A0A1X7RID6 |

**otein identification number (ID) and functional annotation of PAN domain-containing proteins.**

| Function |
|---|
| GON-Abd-1 |
| GON-Abd-2 |
| Uncharacterized protein MAMA_L838 |
| PAN/APPLE-like domain protein |
| PAN domain-containing protein |
| PAN domain-containing protein |
| PAN domain-containing protein |
| Putative PAN domain-containing protein |
| Putative PAN domain-containing protein |
| Uncharacterized protein |
| Uncharacterized protein |
| Glycosyl hydrolases family 32 superfamily protein |
| Uncharacterized protein |
| Uncharacterized protein |
| Uncharacterized protein |
| Uncharacterized protein |
| Uncharacterized protein |
| Uncharacterized protein |
| Uncharacterized protein |
| Uncharacterized protein |
| Pan domain-containing |
| Putative PAN domain-containing protein |
| Uncharacterized protein L719 |
| Putative PAN domain-containing protein |
| Uncharacterized protein L719 |
| Putative PAN domain-containing protein |
| Uncharacterized protein |
| Pan domain-containing |
| Putative PAN domain-containing protein L775 |
| Uncharacterized protein L719 |
| Putative PAN domain-containing protein R486 |
| Pan domain-containing |
| Uncharacterized protein |
| Putative PAN domain-containing protein |
| Putative PAN domain-containing protein |
| PAN domain-containing protein |
| Uncharacterized protein |
| PAN domain-containing protein |
| Plasminogen (Fragment) |

Plasminogen (Fragment)
Uncharacterized protein
Uncharacterized protein
Uncharacterized protein
Uncharacterized protein
Uncharacterized protein
Plasminogen
Uncharacterized protein
Uncharacterized protein
Secreted protein (Fragment)
Secreted protein
Secreted protein
Secreted protein
Carbohydrate-binding protein
Carbohydrate-binding protein
GPI anchor protein
Secreted protein
Secreted protein
Secreted protein
Secreted protein
Uncharacterized protein
Uncharacterized protein (Fragment)
Lysostaphin
Uncharacterized protein
Uncharacterized protein
Uncharacterized protein
Secreted protein
Uncharacterized protein
Secreted protein
Secreted protein
Uncharacterized protein
Secreted protein (Fragment)
Uncharacterized protein
Uncharacterized protein (Fragment)
Uncharacterized protein
Uncharacterized protein (Fragment)
Secreted protein
Secreted protein
Uncharacterized protein
Secreted protein (Fragment)
Uncharacterized protein

Uncharacterized protein (Fragment)
Uncharacterized protein
Secreted protein
Secreted protein
GPI anchor protein
Uncharacterized protein
Uncharacterized protein
Uncharacterized protein
Glucanase
Secreted protein
Uncharacterized protein
Carbohydrate-binding protein (Fragment)
Glucanase
Uncharacterized protein
Uncharacterized protein (Fragment)
Uncharacterized protein
Uncharacterized protein
Uncharacterized protein
Uncharacterized protein
Uncharacterized protein (Fragment)
Uncharacterized protein
Uncharacterized protein
Uncharacterized protein
Plasma kallikrein (Fragment)
Coagulation factor XI
Neurotrypsin
Uncharacterized protein
Uncharacterized protein
Cuticlin-1
Uncharacterized protein
Uncharacterized protein
Uncharacterized protein
Uncharacterized protein
G-type lectin S-receptor-like serine/threonine-protein kinase
G-type lectin S-receptor-like serine/threonine-protein kinase
Epidermis-specific secreted glycoprotein
G-type lectin S-receptor-like serine/threonine-protein kinase
G-type lectin S-receptor-like serine/threonine-protein kinase
G-type lectin S-receptor-like serine/threonine-protein kinase
G-type lectin S-receptor-like serine/threonine-protein kinase
G-type lectin S-receptor-like serine/threonine-protein kinase

G-type lectin S-receptor-like serine/threonine-protein kinase
G-type lectin S-receptor-like serine/threonine-protein kinase
PAN domain-containing protein
G-type lectin S-receptor-like serine/threonine-protein kinase
G-type lectin S-receptor-like serine/threonine-protein kinase
Receptor protein kinase
G-type lectin S-receptor-like serine/threonine-protein kinase
G-type lectin S-receptor-like serine/threonine-protein kinase
G-type lectin S-receptor-like serine/threonine-protein kinase
G-type lectin S-receptor-like serine/threonine-protein kinase
G-type lectin S-receptor-like serine/threonine-protein kinase
G-type lectin S-receptor-like serine/threonine-protein kinase
G-type lectin S-receptor-like serine/threonine-protein kinase
G-type lectin S-receptor-like serine/threonine-protein kinase
G-type lectin S-receptor-like serine/threonine-protein kinase
G-type lectin S-receptor-like serine/threonine-protein kinase
Receptor protein kinase
Epidermis-specific secreted glycoprotein
G-type lectin S-receptor-like serine/threonine-protein kinase
G-type lectin S-receptor-like serine/threonine-protein kinase
G-type lectin S-receptor-like serine/threonine-protein kinase
Receptor protein kinase
G-type lectin S-receptor-like serine/threonine-protein kinase
G-type lectin S-receptor-like serine/threonine-protein kinase
G-type lectin S-receptor-like serine/threonine-protein kinase
G-type lectin S-receptor-like serine/threonine-protein kinase
G-type lectin S-receptor-like serine/threonine-protein kinase
G-type lectin S-receptor-like serine/threonine-protein kinase
G-type lectin S-receptor-like serine/threonine-protein kinase (Fragment)
G-type lectin S-receptor-like serine/threonine-protein kinase
G-type lectin S-receptor-like serine/threonine-protein kinase
G-type lectin S-receptor-like serine/threonine-protein kinase (Fragment)
Receptor-like serine/threonine-protein kinase
G-type lectin S-receptor-like serine/threonine-protein kinase
G-type lectin S-receptor-like serine/threonine-protein kinase
G-type lectin S-receptor-like serine/threonine-protein kinase
G-type lectin S-receptor-like serine/threonine-protein kinase
Receptor-like serine/threonine-protein kinase
G-type lectin S-receptor-like serine/threonine-protein kinase (Fragment)
G-type lectin S-receptor-like serine/threonine-protein kinase
G-type lectin S-receptor-like serine/threonine-protein kinase

G-type lectin S-receptor-like serine/threonine-protein kinase
G-type lectin S-receptor-like serine/threonine-protein kinase
G-type lectin S-receptor-like serine/threonine-protein kinase
G-type lectin S-receptor-like serine/threonine-protein kinase
G-type lectin S-receptor-like serine/threonine-protein kinase (Fragment)
G-type lectin S-receptor-like serine/threonine-protein kinase
PAN domain-containing protein
G-type lectin S-receptor-like serine/threonine-protein kinase
G-type lectin S-receptor-like serine/threonine-protein kinase
G-type lectin S-receptor-like serine/threonine-protein kinase
G-type lectin S-receptor-like serine/threonine-protein kinase
G-type lectin S-receptor-like serine/threonine-protein kinase
G-type lectin S-receptor-like serine/threonine-protein kinase
Receptor-like serine/threonine-protein kinase
Uncharacterized protein
Uncharacterized protein
Uncharacterized protein
Uncharacterized protein
Uncharacterized protein
Uncharacterized protein
Uncharacterized protein
Uncharacterized protein
Uncharacterized protein
Endoxylanase-like protein
Uncharacterized protein
Uncharacterized protein
PAN/APPLE-like domain-containing protein (Fragment)
No-mechanoreceptor potential a
AAEL008647-PA
Uncharacterized protein
AAEL009594-PA
AAEL002066-PA
AAEL007579-PA
AAEL003278-PA
AAEL009592-PA
Uncharacterized protein
Uncharacterized protein
Uncharacterized protein
Uncharacterized protein
Uncharacterized protein
Uncharacterized protein

Uncharacterized protein
Uncharacterized protein
Uncharacterized protein
Uncharacterized protein
Uncharacterized protein
Uncharacterized protein
Uncharacterized protein
Uncharacterized protein
Uncharacterized protein
Uncharacterized protein
Uncharacterized protein
Uncharacterized protein
Uncharacterized protein
Uncharacterized protein
Uncharacterized protein
Uncharacterized protein
Uncharacterized protein
Uncharacterized protein
Uncharacterized protein
Uncharacterized protein
Uncharacterized protein
Uncharacterized protein
Uncharacterized protein
Uncharacterized protein
Uncharacterized protein
Uncharacterized protein
Uncharacterized protein
Uncharacterized protein
Uncharacterized protein
Uncharacterized protein
Uncharacterized protein
Uncharacterized protein
Uncharacterized protein
Uncharacterized protein
Uncharacterized protein
Uncharacterized protein
Uncharacterized protein
Uncharacterized protein
Uncharacterized protein
Uncharacterized protein
Uncharacterized protein

Uncharacterized protein
Uncharacterized protein
Uncharacterized protein
Uncharacterized protein
Uncharacterized protein
Uncharacterized protein
Uncharacterized protein
Uncharacterized protein
Uncharacterized protein
Uncharacterized protein
Uncharacterized protein
Uncharacterized protein
Uncharacterized protein
Uncharacterized protein
Uncharacterized protein
Uncharacterized protein
Uncharacterized protein
Uncharacterized protein
Uncharacterized protein
Uncharacterized protein
Uncharacterized protein
Uncharacterized protein
Uncharacterized protein
Uncharacterized protein
Uncharacterized protein
Uncharacterized protein
Uncharacterized protein
Uncharacterized protein
Uncharacterized protein
Uncharacterized protein
Uncharacterized protein
Uncharacterized protein
Uncharacterized protein
Uncharacterized protein
Uncharacterized protein
Uncharacterized protein
Uncharacterized protein
Uncharacterized protein
Uncharacterized protein
Uncharacterized protein
Uncharacterized protein

Uncharacterized protein
Uncharacterized protein
Uncharacterized protein
Uncharacterized protein
Uncharacterized protein
Uncharacterized protein
Uncharacterized protein
Uncharacterized protein
Uncharacterized protein
Uncharacterized protein
Uncharacterized protein
Uncharacterized protein
Uncharacterized protein
Uncharacterized protein
Uncharacterized protein
Uncharacterized protein
Uncharacterized protein
Uncharacterized protein
Uncharacterized protein
Uncharacterized protein
Uncharacterized protein
Uncharacterized protein
Uncharacterized protein
Uncharacterized protein
Uncharacterized protein
Uncharacterized protein
Uncharacterized protein
Uncharacterized protein
Uncharacterized protein
Uncharacterized protein
Uncharacterized protein
Uncharacterized protein
Uncharacterized protein
Uncharacterized protein
Uncharacterized protein
Uncharacterized protein
Uncharacterized protein
Uncharacterized protein
Uncharacterized protein
Uncharacterized protein
Uncharacterized protein

Uncharacterized protein
Uncharacterized protein
Uncharacterized protein
Uncharacterized protein
Uncharacterized protein
Uncharacterized protein
Uncharacterized protein
Uncharacterized protein
Uncharacterized protein
Uncharacterized protein
Uncharacterized protein
Uncharacterized protein
Uncharacterized protein
Uncharacterized protein
Uncharacterized protein
Uncharacterized protein
Uncharacterized protein
Uncharacterized protein
Uncharacterized protein
Uncharacterized protein
Uncharacterized protein
Uncharacterized protein
Uncharacterized protein
Uncharacterized protein
Uncharacterized protein
Uncharacterized protein
Uncharacterized protein
Uncharacterized protein
Uncharacterized protein
Uncharacterized protein
Uncharacterized protein
Uncharacterized protein
Uncharacterized protein
Uncharacterized protein
Uncharacterized protein
Uncharacterized protein
Uncharacterized protein
Uncharacterized protein
Uncharacterized protein
Uncharacterized protein
Uncharacterized protein

Uncharacterized protein
Uncharacterized protein
Uncharacterized protein
Uncharacterized protein
Uncharacterized protein
Uncharacterized protein
Uncharacterized protein
Uncharacterized protein
Uncharacterized protein
Uncharacterized protein
Uncharacterized protein
Uncharacterized protein
Uncharacterized protein
Uncharacterized protein
Uncharacterized protein
Uncharacterized protein
Uncharacterized protein
Uncharacterized protein
Uncharacterized protein
Uncharacterized protein
Uncharacterized protein
Uncharacterized protein
Uncharacterized protein
Uncharacterized protein
Uncharacterized protein
Uncharacterized protein
Uncharacterized protein
Uncharacterized protein
Uncharacterized protein
Uncharacterized protein
Uncharacterized protein
Uncharacterized protein
Uncharacterized protein
Uncharacterized protein
Uncharacterized protein
Uncharacterized protein
Uncharacterized protein
Uncharacterized protein
Uncharacterized protein
Uncharacterized protein
Uncharacterized protein

Uncharacterized protein
Uncharacterized protein
Uncharacterized protein
Uncharacterized protein
Uncharacterized protein
Uncharacterized protein
Uncharacterized protein
Uncharacterized protein
Uncharacterized protein
Uncharacterized protein
Uncharacterized protein
Uncharacterized protein
Uncharacterized protein
Uncharacterized protein
Uncharacterized protein
Uncharacterized protein
Uncharacterized protein
Uncharacterized protein
Uncharacterized protein
Uncharacterized protein
Uncharacterized protein
Uncharacterized protein
Uncharacterized protein
Uncharacterized protein
Uncharacterized protein
Uncharacterized protein
Uncharacterized protein
Uncharacterized protein
Uncharacterized protein
Uncharacterized protein
Uncharacterized protein
Uncharacterized protein
Uncharacterized protein
Uncharacterized protein
Uncharacterized protein
Uncharacterized protein
Uncharacterized protein
Uncharacterized protein
Uncharacterized protein
Uncharacterized protein
Uncharacterized protein

Uncharacterized protein
Uncharacterized protein
Uncharacterized protein
Uncharacterized protein
Uncharacterized protein
Uncharacterized protein
Uncharacterized protein
Uncharacterized protein
Uncharacterized protein
Uncharacterized protein
Uncharacterized protein
Uncharacterized protein
Uncharacterized protein
Uncharacterized protein
Uncharacterized protein
Uncharacterized protein
Uncharacterized protein
Uncharacterized protein
Uncharacterized protein
Uncharacterized protein
Uncharacterized protein
Uncharacterized protein
Uncharacterized protein
Uncharacterized protein
Uncharacterized protein
Uncharacterized protein
Uncharacterized protein
Uncharacterized protein
Uncharacterized protein
Uncharacterized protein
Uncharacterized protein
Uncharacterized protein
Uncharacterized protein
Uncharacterized protein
Uncharacterized protein
Uncharacterized protein
Uncharacterized protein
Uncharacterized protein
Uncharacterized protein
Uncharacterized protein
Uncharacterized protein

Uncharacterized protein
Uncharacterized protein
Uncharacterized protein
Uncharacterized protein
Uncharacterized protein
Uncharacterized protein
Uncharacterized protein
Uncharacterized protein
Uncharacterized protein
Uncharacterized protein
Uncharacterized protein
Uncharacterized protein
Uncharacterized protein
Uncharacterized protein
Uncharacterized protein
Uncharacterized protein
Uncharacterized protein
Uncharacterized protein
Uncharacterized protein
Uncharacterized protein
Uncharacterized protein
Uncharacterized protein
Uncharacterized protein
Uncharacterized protein
Uncharacterized protein
Uncharacterized protein
Uncharacterized protein
Uncharacterized protein
Uncharacterized protein
Uncharacterized protein
Uncharacterized protein
Uncharacterized protein
Uncharacterized protein
Uncharacterized protein
Uncharacterized protein
Uncharacterized protein
Uncharacterized protein
Uncharacterized protein
Uncharacterized protein
Uncharacterized protein
Uncharacterized protein

Uncharacterized protein
Uncharacterized protein
Uncharacterized protein
Uncharacterized protein
Uncharacterized protein
Uncharacterized protein
Uncharacterized protein
Uncharacterized protein
Uncharacterized protein
Uncharacterized protein
Uncharacterized protein
Uncharacterized protein
Uncharacterized protein
Uncharacterized protein
Uncharacterized protein
Uncharacterized protein
Uncharacterized protein
Uncharacterized protein
Uncharacterized protein
Uncharacterized protein
Uncharacterized protein
Uncharacterized protein
Uncharacterized protein
Uncharacterized protein
Uncharacterized protein
Uncharacterized protein
Uncharacterized protein
Uncharacterized protein
Uncharacterized protein
Uncharacterized protein
Uncharacterized protein
Uncharacterized protein
Uncharacterized protein
Uncharacterized protein
Uncharacterized protein
Uncharacterized protein
Uncharacterized protein
Uncharacterized protein
Uncharacterized protein
Uncharacterized protein
Uncharacterized protein

Uncharacterized protein
Uncharacterized protein
Uncharacterized protein
Uncharacterized protein
Uncharacterized protein
Uncharacterized protein
Uncharacterized protein
Uncharacterized protein
Uncharacterized protein
Uncharacterized protein
Uncharacterized protein
Uncharacterized protein
Uncharacterized protein
Uncharacterized protein
Uncharacterized protein
Uncharacterized protein
Uncharacterized protein
Uncharacterized protein
Uncharacterized protein
Uncharacterized protein
Uncharacterized protein
Uncharacterized protein
Uncharacterized protein
Uncharacterized protein
Uncharacterized protein
Uncharacterized protein
Uncharacterized protein
Uncharacterized protein
Uncharacterized protein
Uncharacterized protein
Uncharacterized protein
Uncharacterized protein
Uncharacterized protein
Uncharacterized protein
Uncharacterized protein
Uncharacterized protein
Uncharacterized protein
Uncharacterized protein
Uncharacterized protein
Uncharacterized protein
Uncharacterized protein

Uncharacterized protein
Uncharacterized protein
Uncharacterized protein
Uncharacterized protein
Uncharacterized protein
Uncharacterized protein
Uncharacterized protein
Uncharacterized protein
Uncharacterized protein
Uncharacterized protein
Uncharacterized protein
Uncharacterized protein
Uncharacterized protein
Uncharacterized protein
Uncharacterized protein
Uncharacterized protein
Uncharacterized protein
Uncharacterized protein
Uncharacterized protein
Uncharacterized protein
Uncharacterized protein
Uncharacterized protein
Uncharacterized protein
Uncharacterized protein
Uncharacterized protein
Uncharacterized protein
Uncharacterized protein
Uncharacterized protein
Uncharacterized protein
Uncharacterized protein
Uncharacterized protein
Uncharacterized protein
Uncharacterized protein
Uncharacterized protein
Uncharacterized protein
Uncharacterized protein
Uncharacterized protein
Uncharacterized protein
Uncharacterized protein
Uncharacterized protein
Uncharacterized protein

Uncharacterized protein
Uncharacterized protein
Uncharacterized protein
Uncharacterized protein
Uncharacterized protein
Uncharacterized protein
Uncharacterized protein
Uncharacterized protein
Uncharacterized protein
Uncharacterized protein
Uncharacterized protein
Uncharacterized protein
Uncharacterized protein
Uncharacterized protein
Uncharacterized protein
Uncharacterized protein
Uncharacterized protein
Uncharacterized protein
Uncharacterized protein
Uncharacterized protein
Uncharacterized protein
Uncharacterized protein
Uncharacterized protein
Uncharacterized protein
Uncharacterized protein
Uncharacterized protein
Uncharacterized protein
Uncharacterized protein
Uncharacterized protein
Uncharacterized protein
Uncharacterized protein
Uncharacterized protein
Uncharacterized protein
Uncharacterized protein
Uncharacterized protein
Uncharacterized protein
Uncharacterized protein
Uncharacterized protein
Uncharacterized protein
Uncharacterized protein
Uncharacterized protein

Uncharacterized protein
Uncharacterized protein
Uncharacterized protein
Uncharacterized protein
Uncharacterized protein
Uncharacterized protein
Uncharacterized protein
Uncharacterized protein
Uncharacterized protein
Uncharacterized protein
Uncharacterized protein
Uncharacterized protein
Uncharacterized protein
Uncharacterized protein
Uncharacterized protein
Uncharacterized protein
Uncharacterized protein
Uncharacterized protein
Uncharacterized protein
Uncharacterized protein
Uncharacterized protein
Uncharacterized protein
Uncharacterized protein
Uncharacterized protein
Uncharacterized protein
Uncharacterized protein
Uncharacterized protein
Uncharacterized protein
Uncharacterized protein
Uncharacterized protein
Uncharacterized protein
Uncharacterized protein
Uncharacterized protein
Uncharacterized protein
Uncharacterized protein
Uncharacterized protein
Uncharacterized protein
Uncharacterized protein
Uncharacterized protein
Uncharacterized protein
Uncharacterized protein

Uncharacterized protein
Uncharacterized protein
Uncharacterized protein
Uncharacterized protein
Putative serine/threonine-protein kinase receptor
Putative serine/threonine-protein kinase receptor
Putative serine/threonine-protein kinase receptor
Putative serine/threonine-protein kinase receptor
Putative receptor protein kinase ZmPK1
Putative serine/threonine-protein kinase receptor
Putative receptor protein kinase ZmPK1
S-locus-specific glycoprotein S6
Putative receptor protein kinase ZmPK1
S-locus-specific glycoprotein S13
Putative receptor protein kinase ZmPK1
Putative receptor protein kinase ZmPK1
Putative serine/threonine-protein kinase receptor
S-locus-specific glycoprotein S13
S-locus-specific glycoprotein S13
Putative serine/threonine-protein kinase receptor
Putative serine/threonine-protein kinase receptor
Putative serine/threonine-protein kinase receptor
Putative receptor protein kinase ZmPK1
Putative receptor protein kinase ZmPK1
Putative serine/threonine-protein kinase receptor
Putative receptor protein kinase ZmPK1
Putative receptor protein kinase ZmPK1
Putative receptor protein kinase ZmPK1
Putative receptor protein kinase ZmPK1
Putative serine/threonine-protein kinase receptor
Putative serine/threonine-protein kinase receptor
Putative serine/threonine-protein kinase receptor
Putative serine/threonine-protein kinase receptor
Putative serine/threonine-protein kinase receptor
Putative receptor protein kinase ZmPK1
Putative serine/threonine-protein kinase receptor
Putative receptor protein kinase ZmPK1
Putative receptor protein kinase ZmPK1
Putative serine/threonine-protein kinase receptor
Putative receptor protein kinase ZmPK1
S-locus-specific glycoprotein

Putative serine/threonine-protein kinase receptor
Putative serine/threonine-protein kinase receptor
Putative receptor protein kinase ZmPK1
Putative serine/threonine-protein kinase receptor
Putative serine/threonine-protein kinase receptor
Putative serine/threonine-protein kinase receptor
Putative serine/threonine-protein kinase receptor
Putative serine/threonine-protein kinase receptor
Putative serine/threonine-protein kinase receptor
Putative serine/threonine-protein kinase receptor
Putative serine/threonine-protein kinase
Cysteine-rich receptor-like protein kinase 10
Putative serine/threonine-protein kinase receptor
Putative serine/threonine-protein kinase receptor
Uncharacterized protein
Putative serine/threonine-protein kinase receptor
PAN domain-containing protein
Putative receptor protein kinase ZmPK1
Putative serine/threonine-protein kinase receptor
Uncharacterized protein
Putative serine/threonine-protein kinase receptor
Putative serine/threonine-protein kinase receptor
Cysteine-rich receptor-like protein kinase 10
Putative serine/threonine-protein kinase receptor
Putative receptor protein kinase ZmPK1
Putative serine/threonine-protein kinase receptor
Putative serine/threonine-protein kinase receptor
Putative receptor protein kinase ZmPK1
Putative serine/threonine-protein kinase receptor
S-locus-specific glycoprotein S13
Putative serine/threonine-protein kinase receptor
Putative serine/threonine-protein kinase receptor
Putative serine/threonine-protein kinase receptor
Putative receptor protein kinase ZmPK1
Putative serine/threonine-protein kinase receptor
Putative receptor protein kinase ZmPK1
Putative serine/threonine-protein kinase receptor
Putative receptor protein kinase ZmPK1
Putative serine/threonine-protein kinase receptor
Putative receptor protein kinase ZmPK1
Putative serine/threonine-protein kinase receptor

Putative receptor protein kinase ZmPK1
Putative serine/threonine-protein kinase receptor
Putative serine/threonine-protein kinase receptor
Putative serine/threonine-protein kinase receptor
Cysteine-rich receptor-like protein kinase 10
Putative serine/threonine-protein kinase receptor
Putative serine/threonine-protein kinase receptor
Putative serine/threonine-protein kinase receptor
Putative serine/threonine-protein kinase receptor
Putative serine/threonine-protein kinase receptor
Putative serine/threonine-protein kinase receptor
Putative serine/threonine-protein kinase receptor
Putative serine/threonine-protein kinase receptor
Putative serine/threonine-protein kinase receptor
Putative serine/threonine-protein kinase receptor
Putative serine/threonine-protein kinase receptor
S-locus-specific glycoprotein BS29-2
Putative serine/threonine-protein kinase receptor
Putative serine/threonine-protein kinase receptor
Putative serine/threonine-protein kinase receptor
Putative serine/threonine-protein kinase receptor
Putative serine/threonine-protein kinase receptor
Putative serine/threonine-protein kinase receptor
Putative serine/threonine-protein kinase receptor
Cysteine-rich receptor-like protein kinase 6
Putative serine/threonine-protein kinase receptor
Putative serine/threonine-protein kinase receptor
Putative serine/threonine-protein kinase receptor
Putative serine/threonine-protein kinase receptor
Putative serine/threonine-protein kinase receptor
Putative serine/threonine-protein kinase receptor
Putative serine/threonine-protein kinase receptor
Putative serine/threonine-protein kinase receptor
Putative serine/threonine-protein kinase receptor
Putative LRR receptor-like serine/threonine-protein kinase
Putative cysteine-rich receptor-like protein kinase 35
Putative receptor protein kinase ZmPK1
Putative serine/threonine-protein kinase receptor
Putative serine/threonine-protein kinase receptor
Putative receptor protein kinase ZmPK1
Putative serine/threonine-protein kinase receptor

Putative serine/threonine-protein kinase receptor
Putative serine/threonine-protein kinase receptor
S-locus-specific glycoprotein BS29-2
Putative receptor protein kinase ZmPK1
Putative serine/threonine-protein kinase receptor
Putative receptor protein kinase ZmPK1
Putative serine/threonine-protein kinase receptor
Uncharacterized protein
Putative serine/threonine-protein kinase receptor
Putative serine/threonine-protein kinase receptor
Putative receptor protein kinase ZmPK1
Putative serine/threonine-protein kinase receptor
Putative serine/threonine-protein kinase receptor
Putative serine/threonine-protein kinase receptor
Putative receptor protein kinase ZmPK1
Putative serine/threonine-protein kinase receptor
Cysteine-rich receptor-like protein kinase 10
Putative serine/threonine-protein kinase receptor
Putative serine/threonine-protein kinase receptor
Putative receptor protein kinase ZmPK1
Uncharacterized protein
Uncharacterized protein
PAN domain-containing protein
Uncharacterized protein
PAN domain protein
PAN domain
Apple domain-containing protein
Apple domain-containing protein
Apple domain-containing protein
Apple domain-containing protein
PAN domain-containing protein
Apple domain-containing protein
Apple domain-containing protein
Apple domain-containing protein
Envelope protein 164
I10L
uncharacterized protein LOC108741704
uncharacterized protein LOC108732347
uncharacterized protein LOC108734666
uncharacterized protein LOC108734669
Alpha-2-macroglobulin domain-containing protein 2

Membrane protein
Uncharacterized protein
Uncharacterized protein
Uncharacterized protein
Uncharacterized protein
Uncharacterized protein
Uncharacterized protein
Uncharacterized protein
Membrane protein
Membrane protein
Uncharacterized protein
Uncharacterized protein
Uncharacterized protein
Uncharacterized protein
Uncharacterized protein
Uncharacterized protein
Uncharacterized protein
Uncharacterized protein
Uncharacterized protein
Membrane protein
Uncharacterized protein
Uncharacterized protein
Serine protease
Uncharacterized protein
Uncharacterized protein
Membrane protein
Uncharacterized protein
Uncharacterized protein
Uncharacterized protein
Uncharacterized protein
Uncharacterized protein
Uncharacterized protein
Uncharacterized protein
Uncharacterized protein
Tetratricopeptide repeat protein
Uncharacterized protein
Uncharacterized protein
Uncharacterized protein
Uncharacterized protein
Membrane protein
Uncharacterized protein

Uncharacterized protein
Uncharacterized protein
MG2 domain protein
Uncharacterized protein
Membrane protein
Uncharacterized protein (Fragment)
Uncharacterized protein (Fragment)
Uncharacterized protein
Uncharacterized protein
Uncharacterized protein
Uncharacterized protein
Uncharacterized protein
Uncharacterized protein
Uncharacterized protein
Uncharacterized protein
Uncharacterized protein
Uncharacterized protein
Uncharacterized protein
Uncharacterized protein (Fragment)
Uncharacterized protein
Uncharacterized protein
VWA domain-containing protein
Uncharacterized protein
Uncharacterized protein
Uncharacterized protein
Uncharacterized protein
Coagulation factor XI
Uncharacterized protein
Kallikrein B1
Uncharacterized protein
Uncharacterized protein (Fragment)
Uncharacterized protein (Fragment)
Hepatocyte growth factor
Uncharacterized protein (Fragment)
Receptor-like serine/threonine-protein kinase SD1-8
Uncharacterized protein
Uncharacterized protein
Uncharacterized protein
Uncharacterized protein
AlNc14C96G5867 protein
Cellulose binding elicitor lectin (CBEL) putative

Cellulose binding elicitor lectin (CBEL) putative
Cellulose binding elicitor lectin (CBEL) putative
AlNc14C96G5853 protein
Uncharacterized protein
Plasminogen
Hepatocyte growth factor
Plasma kallikrein isoform A
Hepatocyte growth factor
Plasminogen
Hepatocyte growth factor-like protein
Plasma kallikrein isoform B
hepatocyte growth factor-like protein isoform X1
plasma kallikrein
Plasminogen
Hepatocyte growth factor
Alpha-2-macroglobulin domain protein
Uncharacterized protein
Uncharacterized protein
Uncharacterized protein
Macrophage stimulating 1 (Fragment)
Macrophage stimulating 1 (Fragment)
Macrophage stimulating 1 (Fragment)
Uncharacterized protein
Apple domain-containing protein
Uncharacterized protein (Fragment)
Alpha-2-macroglobulin
Uncharacterized protein
PAN domain-containing protein
PAN domain-containing protein
PAN domain-containing protein
Alpha-2-macroglobulin family protein
Alpha-2-macroglobulin family protein (Fragment)
Uncharacterized protein (Fragment)
Uncharacterized protein
Uncharacterized protein (Fragment)
Uncharacterized protein
Uncharacterized protein
Uncharacterized protein
Uncharacterized protein
Galactose oxidase
Uncharacterized protein (Fragment)

Uncharacterized protein
Uncharacterized protein
Uncharacterized protein
Uncharacterized protein (Fragment)
Hepatocyte growth factor
Coagulation factor XI-like protein
Hepatocyte growth factor-like protein-like isoform X2
Uncharacterized protein
Putative conserved secreted protein
Putative conserved secreted protein (Fragment)
Uncharacterized protein
Uncharacterized protein
Uncharacterized protein
Uncharacterized protein
Uncharacterized protein
Uncharacterized protein
Uncharacterized protein
Uncharacterized protein
Uncharacterized protein
Uncharacterized protein
Uncharacterized protein
Uncharacterized protein
Uncharacterized protein
Uncharacterized protein
Uncharacterized protein
Uncharacterized protein
Uncharacterized protein
Uncharacterized protein
Uncharacterized protein
Uncharacterized protein
Uncharacterized protein
Uncharacterized protein
Uncharacterized protein
Uncharacterized protein
PAN domain
Alpha-2-macroglobulin family N-terminal region
Membrane protein
Uncharacterized protein
Uncharacterized protein
Uncharacterized protein
Plasminogen

Macrophage stimulating 1
Plasminogen
Plasminogen
Plasminogen
Hepatocyte growth factor a
Plasminogen
Macrophage stimulating 1
Plasminogen
Macrophage stimulating 1
Plasminogen
Macrophage stimulating 1
Hepatocyte growth factor
Uncharacterized protein
Uncharacterized protein
Hepatocyte growth factor
Uncharacterized protein
Uncharacterized protein
Plasminogen
Uncharacterized protein
Hepatocyte growth factor a
Uncharacterized protein
Macrophage stimulating 1
Uncharacterized protein
Hepatocyte growth factor
Venom protein
Venom protein
PAN domain-containing protein
Uncharacterized protein
Macrophage stimulating 1
Plasminogen
Uncharacterized protein
Plasminogen
Uncharacterized protein
Hepatocyte growth factor a
Uncharacterized protein
Macrophage stimulating 1
Hepatocyte growth factor
Macrophage stimulating 1
Uncharacterized protein
Uncharacterized protein
G-type lectin S-receptor-like serine/threonine-protein kinase SD3-1

G-type lectin S-receptor-like serine/threonine-protein kinase
Putative receptor protein kinase ZmPK1
G-type lectin S-receptor-like serine/threonine-protein kinase SD2-5
Receptor-like serine/threonine-protein kinase SD1-8
Receptor-like serine/threonine-protein kinase SD1-8
Receptor-like serine/threonine-protein kinase SD1-8 (Fragment)
G-type lectin S-receptor-like serine/threonine-protein kinase RLK1
Receptor-like serine/threonine-protein kinase SD1-8 (Fragment)
G-type lectin S-receptor-like serine/threonine-protein kinase SD2-5
G-type lectin S-receptor-like serine/threonine-protein kinase
Receptor-like serine/threonine-protein kinase SD1-8
G-type lectin S-receptor-like serine/threonine-protein kinase
G-type lectin S-receptor-like serine/threonine-protein kinase
G-type lectin S-receptor-like serine/threonine-protein kinase
G-type lectin S-receptor-like serine/threonine-protein kinase SD2-2
Putative receptor protein kinase ZmPK1
G-type lectin S-receptor-like serine/threonine-protein kinase (Fragment)
G-type lectin S-receptor-like serine/threonine-protein kinase
G-type lectin S-receptor-like serine/threonine-protein kinase B120
G-type lectin S-receptor-like serine/threonine-protein kinase
Putative receptor protein kinase ZmPK1
G-type lectin S-receptor-like serine/threonine-protein kinase SD2-5
G-type lectin S-receptor-like serine/threonine-protein kinase (Fragment)
Receptor-like serine/threonine-protein kinase SD1-8
G-type lectin S-receptor-like serine/threonine-protein kinase
Plasminogen
Uncharacterized protein
Coagulation factor XI
Plasminogen (Fragment)
Hepatocyte growth factor (Fragment)
Plasma kallikrein (Fragment)
Plasminogen (Fragment)
PAN domain protein
PAN domain protein
PAN domain protein
Uncharacterized protein
PAN domain protein
PAN domain protein
PAN domain protein
PAN domain protein (Fragment)
PAN domain protein

PAN domain protein (Fragment)
PAN domain protein
PAN domain protein
PAN domain protein
PAN domain protein (Fragment)
PAN domain protein
Uncharacterized protein
PAN domain protein
PAN domain protein
PAN domain protein
PAN domain protein (Fragment)
PAN domain protein
PAN domain protein
PAN domain protein
PAN domain protein (Fragment)
PAN domain protein
PAN domain protein (Fragment)
Low-density lipoprotein receptor domain class A
PAN domain protein (Fragment)
PAN domain protein
PAN domain protein
Uncharacterized protein
PAN domain protein
PAN domain protein
Uncharacterized protein
PAN domain protein
Uncharacterized protein
Uncharacterized protein
Uncharacterized protein
Uncharacterized protein
PAN domain protein (Fragment)
Uncharacterized protein
Uncharacterized protein
Uncharacterized protein
Uncharacterized protein
Uncharacterized protein
Uncharacterized protein
Uncharacterized protein
PAN domain protein
Uncharacterized protein
Uncharacterized protein

Uncharacterized protein
PAN domain protein
Uncharacterized protein (Fragment)
Uncharacterized protein
Uncharacterized protein
Uncharacterized protein
Uncharacterized protein
PAN domain protein
Uncharacterized protein
PAN domain protein
Uncharacterized protein
Uncharacterized protein
Uncharacterized protein
Uncharacterized protein
Uncharacterized protein
Uncharacterized protein
Uncharacterized protein
Uncharacterized protein
Uncharacterized protein
Uncharacterized protein
Uncharacterized protein
Uncharacterized protein
Uncharacterized protein
Uncharacterized protein
Uncharacterized protein
Uncharacterized protein
PAN domain protein
Uncharacterized protein
Uncharacterized protein
Uncharacterized protein
Uncharacterized protein (Fragment)
Uncharacterized protein
Zona pellucida-like domain protein
Uncharacterized protein
Uncharacterized protein
Uncharacterized protein
Uncharacterized protein
Uncharacterized protein
Uncharacterized protein
Uncharacterized protein
Uncharacterized protein

PAN domain protein
Uncharacterized protein
Uncharacterized protein
Uncharacterized protein
Uncharacterized protein
Uncharacterized protein
Uncharacterized protein
Uncharacterized protein
Uncharacterized protein
PAN domain protein
Uncharacterized protein
Uncharacterized protein
Uncharacterized protein
Uncharacterized protein (Fragment)
Uncharacterized protein
PAN domain protein (Fragment)
Uncharacterized protein
Uncharacterized protein
Uncharacterized protein
Uncharacterized protein
Uncharacterized protein
Uncharacterized protein
Uncharacterized protein
Uncharacterized protein
Uncharacterized protein
Uncharacterized protein
Uncharacterized protein
Uncharacterized protein
Uncharacterized protein
Uncharacterized protein
Uncharacterized protein
Uncharacterized protein
Uncharacterized protein
PAN domain protein
PAN domain protein
Uncharacterized protein
Uncharacterized protein
Uncharacterized protein
PAN domain protein
Uncharacterized protein
Uncharacterized protein

Uncharacterized protein
PAN domain protein
Uncharacterized protein
PAN domain protein
Uncharacterized protein
Uncharacterized protein
PAN domain protein
PAN domain protein
PAN domain protein
PAN domain protein (Fragment)
PAN domain protein
PAN domain protein
PAN domain protein
PAN domain protein
PAN domain protein
PAN domain protein (Fragment)
Uncharacterized protein
PAN domain protein
PAN domain protein (Fragment)
Uncharacterized protein
PAN domain protein
PAN domain protein
PAN domain protein (Fragment)
PAN domain protein (Fragment)
PAN domain protein (Fragment)
PAN domain protein
PAN domain protein
PAN domain protein (Fragment)
PAN domain protein
PAN domain protein
PAN domain protein (Fragment)
PAN domain protein
PAN domain protein
PAN domain protein (Fragment)
PAN domain protein
PAN domain protein
Uncharacterized protein
Uncharacterized protein
Uncharacterized protein
Uncharacterized protein
Uncharacterized protein

Uncharacterized protein
Uncharacterized protein
Uncharacterized protein
Uncharacterized protein
Uncharacterized protein
Uncharacterized protein
Uncharacterized protein
Uncharacterized protein
Uncharacterized protein
Uncharacterized protein
Uncharacterized protein
Uncharacterized protein
Uncharacterized protein
Uncharacterized protein
Uncharacterized protein
Uncharacterized protein
Uncharacterized protein
Uncharacterized protein
Uncharacterized protein
Uncharacterized protein
Uncharacterized protein
Uncharacterized protein
Uncharacterized protein
Uncharacterized protein
Uncharacterized protein
Uncharacterized protein
Uncharacterized protein
Uncharacterized protein
Uncharacterized protein
Uncharacterized protein
Uncharacterized protein
Uncharacterized protein
Uncharacterized protein
Uncharacterized protein
Uncharacterized protein
Uncharacterized protein
Uncharacterized protein
Uncharacterized protein
Uncharacterized protein
Uncharacterized protein
Uncharacterized protein

Uncharacterized protein
Uncharacterized protein
Uncharacterized protein
Uncharacterized protein
Properdin (inferred by orthology to a human protein)
Uncharacterized protein
Uncharacterized protein
Uncharacterized protein
Uncharacterized protein
Uncharacterized protein
Uncharacterized protein
Uncharacterized protein
Uncharacterized protein
Properdin (inferred by orthology to a human protein)
Uncharacterized protein
Let-653 protein (inferred by orthology to a C. elegans protein)
Uncharacterized protein
Uncharacterized protein
Acrosin (inferred by orthology to a human protein)
Uncharacterized protein
Uncharacterized protein
Uncharacterized protein
Hepatocyte growth factor
Uncharacterized protein
Plasminogen
Uncharacterized protein
Hepatocyte growth factor
Uncharacterized protein
Uncharacterized protein
Uncharacterized protein
Uncharacterized protein
Uncharacterized protein
Uncharacterized protein
Uncharacterized protein
Uncharacterized protein
ENSANGG00000015172 protein (Fragment)
ENSANGG00000015172 protein (Fragment)
Uncharacterized protein
ENSANGG00000015172 protein (Fragment)
Uncharacterized protein
ENSANGG00000015172 protein (Fragment)

Uncharacterized protein
Uncharacterized protein
Uncharacterized protein
Uncharacterized protein
Uncharacterized protein
Uncharacterized protein
Uncharacterized protein
Uncharacterized protein
Uncharacterized protein
Uncharacterized protein
Uncharacterized protein (Fragment)
Uncharacterized protein
Uncharacterized protein
Uncharacterized protein
Uncharacterized protein
Uncharacterized protein
Uncharacterized protein
Uncharacterized protein
Uncharacterized protein
Uncharacterized protein
Uncharacterized protein
Uncharacterized protein
Uncharacterized protein
Uncharacterized protein
Uncharacterized protein
Uncharacterized protein
Uncharacterized protein
Uncharacterized protein
Trynity
Uncharacterized protein
No-mechanoreceptor potential a
Neyo
Uncharacterized protein
Uncharacterized protein
Uncharacterized protein
Uncharacterized protein
Uncharacterized protein
Uncharacterized protein
Uncharacterized protein
Uncharacterized protein
Uncharacterized protein

Uncharacterized protein
Uncharacterized protein
Uncharacterized protein
Uncharacterized protein
Uncharacterized protein
Uncharacterized protein
Uncharacterized protein
Uncharacterized protein
Uncharacterized protein
Uncharacterized protein
Uncharacterized protein
Uncharacterized protein
Uncharacterized protein
Uncharacterized protein
Uncharacterized protein
Uncharacterized protein
Uncharacterized protein
Uncharacterized protein
Uncharacterized protein
AGAP007100-PA
ENSANGG00000015172 protein (Fragment)
AGAP002316-PA
AGAP003012-PA
AGAP009003-PA (Fragment)
Uncharacterized protein
Uncharacterized protein
ENSANGG00000015172 protein (Fragment)
ENSANGG00000015172 protein (Fragment)
Uncharacterized protein
Uncharacterized protein
Uncharacterized protein
Uncharacterized protein
Uncharacterized protein
Uncharacterized protein
Uncharacterized protein
Uncharacterized protein
Uncharacterized protein
Uncharacterized protein
Uncharacterized protein
Uncharacterized protein
Uncharacterized protein

Uncharacterized protein
Uncharacterized protein
Uncharacterized protein
Uncharacterized protein
Uncharacterized protein
Uncharacterized protein
Uncharacterized protein
Uncharacterized protein
Uncharacterized protein
Uncharacterized protein
Uncharacterized protein
Uncharacterized protein
Uncharacterized protein
Uncharacterized protein
Uncharacterized protein
Uncharacterized protein
Uncharacterized protein
AGAP002316-PA-like protein
AGAP007100-PA-like protein
Uncharacterized protein
AGAP009003-PA-like protein
Uncharacterized protein
Uncharacterized protein
Uncharacterized protein
Uncharacterized protein
Uncharacterized protein
Uncharacterized protein
Uncharacterized protein
Uncharacterized protein
PAN domain-containing protein
Putative lipoprotein YfhM
PAN domain protein
Receptor-like serine/threonine-protein kinase SD1-8 (Fragment)
Receptor-like serine/threonine-protein kinase SD1-8 (Fragment)
Receptor-like serine/threonine-protein kinase SD1-8
PAN domain-containing protein At5g03700 (Fragment)
G-type lectin S-receptor-like serine/threonine-protein kinase At2g19130
Receptor-like serine/threonine-protein kinase SD1-8 (Fragment)
Receptor-like serine/threonine-protein kinase SD1-8 (Fragment)
G-type lectin S-receptor-like serine/threonine-protein kinase RLK1 (Fragment)
Receptor-like serine/threonine-protein kinase SD1-8 (Fragment)

G-type lectin S-receptor-like serine/threonine-protein kinase At1g34300
G-type lectin S-receptor-like serine/threonine-protein kinase At5g35370 (Fragment)
Receptor-like serine/threonine-protein kinase SD1-8 (Fragment)
G-type lectin S-receptor-like serine/threonine-protein kinase RKS1 (Fragment)
Receptor-like serine/threonine-protein kinase SD1-8 (Fragment)
G-type lectin S-receptor-like serine/threonine-protein kinase SD2-5 (Fragment)
Receptor-like serine/threonine-protein kinase SD1-8 (Fragment)
G-type lectin S-receptor-like serine/threonine-protein kinase B120 (Fragment)
Receptor-like serine/threonine-protein kinase SD1-8 (Fragment)
Receptor-like serine/threonine-protein kinase SD1-6 (Fragment)
G-type lectin S-receptor-like serine/threonine-protein kinase RKS1 (Fragment)
Receptor-like serine/threonine-protein kinase SD1-8 (Fragment)
G-type lectin S-receptor-like serine/threonine-protein kinase RKS1 (Fragment)
Receptor-like serine/threonine-protein kinase SD1-6 (Fragment)
G-type lectin S-receptor-like serine/threonine-protein kinase SD1-29
Receptor-like serine/threonine-protein kinase SD1-8 (Fragment)
G-type lectin S-receptor-like serine/threonine-protein kinase RKS1 (Fragment)
Receptor-like serine/threonine-protein kinase SD1-8 (Fragment)
G-type lectin S-receptor-like serine/threonine-protein kinase At2g19130 (Fragment)
G-type lectin S-receptor-like serine/threonine-protein kinase B120 (Fragment)
G-type lectin S-receptor-like serine/threonine-protein kinase At1g34300
G-type lectin S-receptor-like serine/threonine-protein kinase SD2-2 (Fragment)
G-type lectin S-receptor-like serine/threonine-protein kinase At2g19130 (Fragment)
G-type lectin S-receptor-like serine/threonine-protein kinase At2g19130
G-type lectin S-receptor-like serine/threonine-protein kinase SD1-13 (Fragment)
G-type lectin S-receptor-like serine/threonine-protein kinase At2g19130 (Fragment)
G-type lectin S-receptor-like serine/threonine-protein kinase At2g19130 (Fragment)
Receptor-like serine/threonine-protein kinase SD1-8 (Fragment)
G-type lectin S-receptor-like serine/threonine-protein kinase SD3-1
G-type lectin S-receptor-like serine/threonine-protein kinase At2g19130 (Fragment)
G-type lectin S-receptor-like serine/threonine-protein kinase At2g19130
G-type lectin S-receptor-like serine/threonine-protein kinase SD1-13 (Fragment)
Receptor-like serine/threonine-protein kinase SD1-8 (Fragment)
S-locus-specific glycoprotein S6 (Fragment)
Receptor-like serine/threonine-protein kinase SD1-8 (Fragment)
G-type lectin S-receptor-like serine/threonine-protein kinase At2g19130 (Fragment)
Receptor-like serine/threonine-protein kinase SD1-8 (Fragment)
G-type lectin S-receptor-like serine/threonine-protein kinase B120 (Fragment)
G-type lectin S-receptor-like serine/threonine-protein kinase SD3-1
Plasminogen (Fragment)
Plasminogen (Fragment)

Hepatocyte growth factor (Fragment)
Uncharacterized protein
Uncharacterized protein
Hepatocyte growth factor
Plasminogen
Uncharacterized protein
Uncharacterized protein
Uncharacterized protein
Uncharacterized protein
Hepatocyte growth factor (Fragment)
Uncharacterized protein
Uncharacterized protein
Uncharacterized protein
Uncharacterized protein
Uncharacterized protein
Uncharacterized protein
Uncharacterized protein
Uncharacterized protein (Fragment)
Uncharacterized protein
Uncharacterized protein
Uncharacterized protein
Uncharacterized protein
Uncharacterized protein (Fragment)
Uncharacterized protein
Uncharacterized protein
Uncharacterized protein (Fragment)
Uncharacterized protein
Uncharacterized protein
Uncharacterized protein
Uncharacterized protein
Uncharacterized protein (Fragment)
Uncharacterized protein
Uncharacterized protein
Uncharacterized protein
Uncharacterized protein
Uncharacterized protein (Fragment)
Uncharacterized protein (Fragment)
Uncharacterized protein (Fragment)
Uncharacterized protein
Uncharacterized protein
Uncharacterized protein (Fragment)

Uncharacterized protein (Fragment)
Uncharacterized protein
Uncharacterized protein
Uncharacterized protein
Uncharacterized protein
Uncharacterized protein
Uncharacterized protein
Uncharacterized protein
Uncharacterized protein
Uncharacterized protein
Uncharacterized protein
Uncharacterized protein
Uncharacterized protein
Uncharacterized protein
Uncharacterized protein
Uncharacterized protein
Uncharacterized protein
Uncharacterized protein
Uncharacterized protein
Uncharacterized protein
Uncharacterized protein
Uncharacterized protein
Uncharacterized protein
Uncharacterized protein
Uncharacterized protein (Fragment)
Uncharacterized protein
Uncharacterized protein
Uncharacterized protein (Fragment)
Uncharacterized protein
Uncharacterized protein
Uncharacterized protein
Uncharacterized protein
Uncharacterized protein
Uncharacterized protein
Uncharacterized protein
Uncharacterized protein
Uncharacterized protein
Uncharacterized protein
Uncharacterized protein (Fragment)
Uncharacterized protein
Uncharacterized protein

Uncharacterized protein
Uncharacterized protein
Uncharacterized protein
Uncharacterized protein
Uncharacterized protein (Fragment)
Uncharacterized protein (Fragment)
Uncharacterized protein (Fragment)
Uncharacterized protein
Uncharacterized protein (Fragment)
Uncharacterized protein
Uncharacterized protein
Uncharacterized protein (Fragment)
Uncharacterized protein (Fragment)
Uncharacterized protein
Uncharacterized protein (Fragment)
Uncharacterized protein
Uncharacterized protein
Uncharacterized protein
Uncharacterized protein (Fragment)
Uncharacterized protein (Fragment)
Uncharacterized protein
Uncharacterized protein
Uncharacterized protein (Fragment)
Uncharacterized protein
Uncharacterized protein (Fragment)
Uncharacterized protein
Uncharacterized protein
Uncharacterized protein
Uncharacterized protein
Uncharacterized protein (Fragment)
Uncharacterized protein
Uncharacterized protein
Uncharacterized protein
Uncharacterized protein (Fragment)
Uncharacterized protein
Uncharacterized protein (Fragment)
Uncharacterized protein
Uncharacterized protein
Uncharacterized protein (Fragment)
Uncharacterized protein
Uncharacterized protein

Uncharacterized protein
Uncharacterized protein
Uncharacterized protein
Uncharacterized protein
Uncharacterized protein
Uncharacterized protein
Uncharacterized protein (Fragment)
Uncharacterized protein (Fragment)
Uncharacterized protein
Uncharacterized protein (Fragment)
Uncharacterized protein (Fragment)
Uncharacterized protein (Fragment)
Uncharacterized protein
Uncharacterized protein
Uncharacterized protein
Uncharacterized protein (Fragment)
Uncharacterized protein
Uncharacterized protein
Uncharacterized protein
Uncharacterized protein
Uncharacterized protein
Uncharacterized protein
Uncharacterized protein
Uncharacterized protein
Uncharacterized protein
Uncharacterized protein
Uncharacterized protein
Uncharacterized protein
Uncharacterized protein
Uncharacterized protein
Uncharacterized protein (Fragment)
Uncharacterized protein
Uncharacterized protein
Uncharacterized protein
Uncharacterized protein
Uncharacterized protein
Uncharacterized protein
Uncharacterized protein
Uncharacterized protein
Uncharacterized protein
Uncharacterized protein

Uncharacterized protein
Uncharacterized protein
Uncharacterized protein
Uncharacterized protein
Uncharacterized protein
Uncharacterized protein
Uncharacterized protein
Uncharacterized protein
Uncharacterized protein
Uncharacterized protein
Uncharacterized protein (Fragment)
Uncharacterized protein
Uncharacterized protein
Uncharacterized protein
Uncharacterized protein
Uncharacterized protein (Fragment)
Uncharacterized protein
Uncharacterized protein (Fragment)
Uncharacterized protein
Uncharacterized protein
Uncharacterized protein (Fragment)
Uncharacterized protein (Fragment)
Uncharacterized protein
Uncharacterized protein
Uncharacterized protein
Uncharacterized protein
Uncharacterized protein
Uncharacterized protein
Uncharacterized protein
Uncharacterized protein (Fragment)
Uncharacterized protein (Fragment)
Uncharacterized protein
Uncharacterized protein
Uncharacterized protein
Uncharacterized protein
Uncharacterized protein
Uncharacterized protein
Uncharacterized protein
Uncharacterized protein (Fragment)
Uncharacterized protein
Uncharacterized protein

Uncharacterized protein
Uncharacterized protein
Uncharacterized protein
Uncharacterized protein (Fragment)
Uncharacterized protein
Uncharacterized protein (Fragment)
Uncharacterized protein
Uncharacterized protein (Fragment)
Uncharacterized protein
Uncharacterized protein
Uncharacterized protein
Uncharacterized protein
Uncharacterized protein
Uncharacterized protein (Fragment)
Uncharacterized protein
Uncharacterized protein
Uncharacterized protein
Uncharacterized protein (Fragment)
Uncharacterized protein
Uncharacterized protein
Uncharacterized protein (Fragment)
Uncharacterized protein
Uncharacterized protein
Uncharacterized protein
Uncharacterized protein
Uncharacterized protein (Fragment)
Uncharacterized protein
Uncharacterized protein
Uncharacterized protein
Uncharacterized protein
Uncharacterized protein
Uncharacterized protein
Uncharacterized protein
Uncharacterized protein (Fragment)
Uncharacterized protein
Uncharacterized protein
Uncharacterized protein (Fragment)
Uncharacterized protein
Uncharacterized protein (Fragment)
Uncharacterized protein (Fragment)
Uncharacterized protein

Uncharacterized protein
Uncharacterized protein
Uncharacterized protein
Uncharacterized protein (Fragment)
Uncharacterized protein
Uncharacterized protein (Fragment)
Uncharacterized protein
Uncharacterized protein
Uncharacterized protein (Fragment)
Uncharacterized protein
Uncharacterized protein
Uncharacterized protein (Fragment)
Uncharacterized protein
Uncharacterized protein
Uncharacterized protein
Uncharacterized protein
Uncharacterized protein
Uncharacterized protein
Uncharacterized protein
Uncharacterized protein (Fragment)
Uncharacterized protein (Fragment)
Uncharacterized protein
Uncharacterized protein
Uncharacterized protein (Fragment)
Uncharacterized protein
Uncharacterized protein (Fragment)
Uncharacterized protein
Uncharacterized protein
Uncharacterized protein
Uncharacterized protein
Uncharacterized protein
Uncharacterized protein
Uncharacterized protein (Fragment)
Uncharacterized protein
Uncharacterized protein
Uncharacterized protein
Uncharacterized protein
Uncharacterized protein (Fragment)
Uncharacterized protein
Uncharacterized protein
Uncharacterized protein

Uncharacterized protein
Uncharacterized protein
Uncharacterized protein (Fragment)
Uncharacterized protein (Fragment)
Uncharacterized protein
Uncharacterized protein
Uncharacterized protein
Uncharacterized protein (Fragment)
Uncharacterized protein
Uncharacterized protein (Fragment)
Uncharacterized protein
Uncharacterized protein
Uncharacterized protein
Uncharacterized protein (Fragment)
Uncharacterized protein (Fragment)
Uncharacterized protein
Uncharacterized protein (Fragment)
Uncharacterized protein (Fragment)
Uncharacterized protein
Uncharacterized protein
Uncharacterized protein
Uncharacterized protein
Uncharacterized protein
Uncharacterized protein (Fragment)
Uncharacterized protein (Fragment)
Uncharacterized protein (Fragment)
Uncharacterized protein
Uncharacterized protein (Fragment)
Uncharacterized protein
Uncharacterized protein (Fragment)
Uncharacterized protein
Uncharacterized protein
Uncharacterized protein (Fragment)
Uncharacterized protein
Uncharacterized protein
Uncharacterized protein
Uncharacterized protein
Uncharacterized protein
Uncharacterized protein
Uncharacterized protein (Fragment)
Uncharacterized protein (Fragment)

Uncharacterized protein
Uncharacterized protein
Uncharacterized protein (Fragment)
Uncharacterized protein (Fragment)
Uncharacterized protein
Uncharacterized protein
Uncharacterized protein
Uncharacterized protein
Uncharacterized protein
Uncharacterized protein
Uncharacterized protein
Uncharacterized protein
Uncharacterized protein (Fragment)
Uncharacterized protein
Uncharacterized protein
Uncharacterized protein
Uncharacterized protein
Uncharacterized protein (Fragment)
Uncharacterized protein
Uncharacterized protein
Uncharacterized protein (Fragment)
Uncharacterized protein
Uncharacterized protein (Fragment)
Uncharacterized protein
Uncharacterized protein
Uncharacterized protein (Fragment)
Uncharacterized protein (Fragment)
Uncharacterized protein (Fragment)
Uncharacterized protein
Uncharacterized protein
Uncharacterized protein
Uncharacterized protein
Uncharacterized protein
Uncharacterized protein (Fragment)
Uncharacterized protein
Uncharacterized protein
Uncharacterized protein (Fragment)
Uncharacterized protein
Uncharacterized protein
Uncharacterized protein
Uncharacterized protein

Uncharacterized protein
Uncharacterized protein
Uncharacterized protein
Uncharacterized protein
Uncharacterized protein
Uncharacterized protein (Fragment)
Uncharacterized protein
Uncharacterized protein
Uncharacterized protein
Uncharacterized protein
Uncharacterized protein
Uncharacterized protein
Uncharacterized protein
Uncharacterized protein
Uncharacterized protein
Uncharacterized protein
Uncharacterized protein
Uncharacterized protein
Uncharacterized protein
Uncharacterized protein (Fragment)
Uncharacterized protein
Uncharacterized protein
Uncharacterized protein (Fragment)
Uncharacterized protein
Uncharacterized protein (Fragment)
Uncharacterized protein
Uncharacterized protein (Fragment)
Uncharacterized protein (Fragment)
Uncharacterized protein (Fragment)
Uncharacterized protein
Uncharacterized protein
Uncharacterized protein
Uncharacterized protein
Uncharacterized protein
Uncharacterized protein (Fragment)
Uncharacterized protein
Uncharacterized protein
Uncharacterized protein
Uncharacterized protein (Fragment)
Uncharacterized protein (Fragment)
Uncharacterized protein

Uncharacterized protein (Fragment)
Uncharacterized protein
Uncharacterized protein (Fragment)
Uncharacterized protein
Uncharacterized protein
Uncharacterized protein
Uncharacterized protein (Fragment)
Uncharacterized protein
Uncharacterized protein
Uncharacterized protein (Fragment)
Uncharacterized protein
Uncharacterized protein (Fragment)
Uncharacterized protein
Uncharacterized protein
Uncharacterized protein (Fragment)
Uncharacterized protein
Uncharacterized protein
Uncharacterized protein
Uncharacterized protein
Uncharacterized protein
Uncharacterized protein
Uncharacterized protein
Uncharacterized protein
Uncharacterized protein
Uncharacterized protein
Uncharacterized protein
Uncharacterized protein
Uncharacterized protein
Uncharacterized protein
Uncharacterized protein
Uncharacterized protein (Fragment)
Uncharacterized protein
Uncharacterized protein
Uncharacterized protein
Uncharacterized protein (Fragment)
Uncharacterized protein
Uncharacterized protein
Uncharacterized protein
Uncharacterized protein
Uncharacterized protein
Uncharacterized protein

Uncharacterized protein (Fragment)
Uncharacterized protein (Fragment)
Uncharacterized protein
Uncharacterized protein
Uncharacterized protein (Fragment)
Uncharacterized protein (Fragment)
Uncharacterized protein
Uncharacterized protein
Uncharacterized protein
Uncharacterized protein
Uncharacterized protein
Uncharacterized protein (Fragment)
Uncharacterized protein
Uncharacterized protein
Uncharacterized protein
Uncharacterized protein
Uncharacterized protein
Uncharacterized protein
Uncharacterized protein
Uncharacterized protein
Uncharacterized protein
Uncharacterized protein (Fragment)
Uncharacterized protein
Uncharacterized protein (Fragment)
Uncharacterized protein
Uncharacterized protein (Fragment)
Uncharacterized protein
Uncharacterized protein
Uncharacterized protein
Uncharacterized protein
Uncharacterized protein
Uncharacterized protein
Uncharacterized protein
Uncharacterized protein (Fragment)
Uncharacterized protein
Uncharacterized protein (Fragment)
Uncharacterized protein (Fragment)
Uncharacterized protein
Uncharacterized protein
Uncharacterized protein
Uncharacterized protein

Uncharacterized protein
Uncharacterized protein
Uncharacterized protein
Uncharacterized protein
Uncharacterized protein
Uncharacterized protein (Fragment)
Uncharacterized protein
Uncharacterized protein
Uncharacterized protein
Uncharacterized protein
Uncharacterized protein
Uncharacterized protein
Uncharacterized protein
Uncharacterized protein
Uncharacterized protein
Uncharacterized protein
Uncharacterized protein
Uncharacterized protein
Uncharacterized protein
Uncharacterized protein
Uncharacterized protein
Uncharacterized protein
Uncharacterized protein
Uncharacterized protein (Fragment)
Uncharacterized protein
Uncharacterized protein
Uncharacterized protein
Uncharacterized protein (Fragment)
Cuticlin-1
Low-density lipoprotein receptor-related protein
Ras-related protein Rab-31
Uncharacterized protein
Neurotrypsin
Uncharacterized protein
Uncharacterized protein
Uncharacterized protein
Uncharacterized protein
Uncharacterized protein
Uncharacterized protein
Uncharacterized protein
Uncharacterized protein

Uncharacterized protein
G-type lectin S-receptor-like serine/threonine-protein kinase
Receptor-like serine/threonine-protein kinase SD1-8
PAN domain-containing protein
Receptor protein kinase ZmPK1
G-type lectin S-receptor-like serine/threonine-protein kinase B120
G-type lectin S-receptor-like serine/threonine-protein kinase
G-type lectin S-receptor-like serine/threonine-protein kinase SD3-1
G-type lectin S-receptor-like serine/threonine-protein kinase
G-type lectin S-receptor-like serine/threonine-protein kinase RLK1
Receptor-like serine/threonine-protein kinase SD1-8
G-type lectin S-receptor-like serine/threonine-protein kinase
G-type lectin S-receptor-like serine/threonine-protein kinase
G-type lectin S-receptor-like serine/threonine-protein kinase
Receptor-like serine/threonine-protein kinase SD1-8
G-type lectin S-receptor-like serine/threonine-protein kinase B120
G-type lectin S-receptor-like serine/threonine-protein kinase B120
G-type lectin S-receptor-like serine/threonine-protein kinase B120
Receptor protein kinase ZmPK1
G-type lectin S-receptor-like serine/threonine-protein kinase RLK1
G-type lectin S-receptor-like serine/threonine-protein kinase
G-type lectin S-receptor-like serine/threonine-protein kinase
Receptor-like serine/threonine-protein kinase SD1-8
PAN domain-containing protein
G-type lectin S-receptor-like serine/threonine-protein kinase SD2-2
Receptor-like serine/threonine-protein kinase SD1-8
G-type lectin S-receptor-like serine/threonine-protein kinase
G-type lectin S-receptor-like serine/threonine-protein kinase
G-type lectin S-receptor-like serine/threonine-protein kinase
Plasminogen (Fragment)
Plasma kallikrein (Fragment)
Plasminogen (Fragment)
Hepatocyte growth factor (Fragment)
Uncharacterized protein
PAN domain-containing protein
Membrane protein
Uncharacterized protein
Uncharacterized protein
Uncharacterized protein
Uncharacterized protein
Uncharacterized protein

Uncharacterized protein
Uncharacterized protein
Uncharacterized protein
Uncharacterized protein
Uncharacterized protein
Uncharacterized protein
Uncharacterized protein
Uncharacterized protein
Uncharacterized protein (Fragment)
Uncharacterized protein (Fragment)
Uncharacterized protein
Uncharacterized protein
Uncharacterized protein
Uncharacterized protein (Fragment)
Uncharacterized protein
Uncharacterized protein (Fragment)
Uncharacterized protein
Uncharacterized protein
Uncharacterized protein
Uncharacterized protein
Uncharacterized protein
Uncharacterized protein (Fragment)
Uncharacterized protein
Uncharacterized protein
Uncharacterized protein
Uncharacterized protein
Uncharacterized protein
Uncharacterized protein
Uncharacterized protein
Uncharacterized protein
Uncharacterized protein
Uncharacterized protein
Uncharacterized protein
Uncharacterized protein
Uncharacterized protein
Uncharacterized protein
Uncharacterized protein
Uncharacterized protein
Uncharacterized protein
Uncharacterized protein
Uncharacterized protein (Fragment)

Uncharacterized protein
Uncharacterized protein
Uncharacterized protein
Uncharacterized protein
Uncharacterized protein
Uncharacterized protein
Uncharacterized protein
Uncharacterized protein
Uncharacterized protein
Uncharacterized protein
Uncharacterized protein
Uncharacterized protein
Uncharacterized protein (Fragment)
Uncharacterized protein
Uncharacterized protein
Uncharacterized protein
Uncharacterized protein
Uncharacterized protein
Uncharacterized protein
Uncharacterized protein (Fragment)
Uncharacterized protein
Uncharacterized protein
Uncharacterized protein
Uncharacterized protein
Uncharacterized protein
Uncharacterized protein
Uncharacterized protein
Uncharacterized protein
PAN domain-containing protein
S-receptor kinase SRK03 isoform a (Fragment)
S-receptor kinase SRK09 (Fragment)
S-receptor kinase SRK17 (Fragment)
S-receptor kinase SRK06 (Fragment)
S-locus receptor kinase (Fragment)
S-locus receptor kinase (Fragment)
S-locus receptor kinase (Fragment)
S-locus receptor kinase (Fragment)
S-receptor kinase SRK14b isoform b (Fragment)
S-locus receptor kinase (Fragment)
S-locus receptor kinase (Fragment)
S-receptor kinase SRK65 (Fragment)

S-receptor kinase SRK72 isoform b (Fragment)
S-receptor kinase SRK21 (Fragment)
S-receptor kinase SRK08 (Fragment)
S-receptor kinase SRK75 (Fragment)
S-locus receptor kinase (Fragment)
S-locus receptor kinase (Fragment)
S-locus receptor kinase (Fragment)
S-receptor kinase SRK16 (Fragment)
S-receptor kinase SRK03 isoform b (Fragment)
S-receptor kinase SRK14 (Fragment)
S-locus receptor kinase (Fragment)
S-locus receptor kinase (Fragment)
S-receptor kinase SRK64 (Fragment)
S-receptor kinase SRK70 (Fragment)
S-receptor kinase SRK06 isoform a (Fragment)
S-locus receptor kinase (Fragment)
S-receptor kinase SRK69 (Fragment)
S-receptor kinase SRK71 (Fragment)
S-receptor kinase SRK74 (Fragment)
S-receptor kinase SRK62 (Fragment)
S-receptor kinase SRK18 (Fragment)
S-receptor kinase SRK73 (Fragment)
S-locus receptor kinase (Fragment)
S-locus receptor kinase (Fragment)
S-locus receptor kinase (Fragment)
S-receptor kinase SRK72 isoform a (Fragment)
S-locus receptor kinase (Fragment)
S-locus receptor kinase (Fragment)
S-locus receptor kinase (Fragment)
S-receptor kinase SRK11 (Fragment)
SRK (Fragment)
SRK
SRK
SRK
Putative S-locus lectin protein kinase family protein
ARK3 (Fragment)
S-receptor kinase (Fragment)
ARK3 protein
S-receptor kinase (Fragment)
S-related kinase 9 (Fragment)
S-receptor kinase (Fragment)

ARK3 (Fragment)
S-receptor kinase (Fragment)
S-receptor kinase (Fragment)
ARK3 (Fragment)
SRKp
S-receptor kinase (Fragment)
S-receptor kinase (Fragment)
S-receptor kinase (Fragment)
ARK3 (Fragment)
S-receptor kinase (Fragment)
SRKp
S-receptor kinase (Fragment)
At4g21390p-like protein
At4g21390p-like protein
SRKp
ARK3p
S-receptor kinase (Fragment)
S-receptor kinase (Fragment)
SRKp
S receptor kinase SRK04 (Fragment)
ARK3 (Fragment)
ARK3p
ARK3p
S-receptor kinase (Fragment)
S-related kinase 9 (Fragment)
SRKp
S-receptor kinase (Fragment)
ARK3 (Fragment)
S receptor kinase SRK10 (Fragment)
S receptor kinase (Fragment)
ARK3 (Fragment)
ARK3p
S-receptor kinase (Fragment)
S-receptor kinase (Fragment)
ARK3 (Fragment)
ARK3p
S-receptor kinase (Fragment)
S-receptor kinase (Fragment)
S-receptor kinase (Fragment)
S-receptor kinase (Fragment)
S receptor kinase (Fragment)

S receptor kinase (Fragment)
S receptor kinase (Fragment)
S-receptor kinase (Fragment)
S-receptor kinase (Fragment)
S-receptor kinase (Fragment)
S-receptor kinase (Fragment)
S-receptor kinase (Fragment)
S-receptor kinase (Fragment)
ARK3 (Fragment)
SRKp
S-receptor kinase (Fragment)
S-receptor kinase (Fragment)
ARK3 (Fragment)
S-receptor kinase (Fragment)
ARK3 (Fragment)
S-receptor kinase (Fragment)
SRKp
S-receptor kinase (Fragment)
S-receptor kinase (Fragment)
At4g21390p-like protein
S-receptor kinase (Fragment)
S-receptor kinase (Fragment)
At4g21390p-like protein
S receptor kinase (Fragment)
S-receptor kinase (Fragment)
S-receptor kinase (Fragment)
S-receptor kinase (Fragment)
S-receptor kinase (Fragment)
S-receptor kinase (Fragment)
S receptor kinase (Fragment)
S receptor kinase (Fragment)
Arabidopsis halleri haplotype Ah15 S-locus region genomic sequence, BAC library Aha_B_L406, BAC clor
S receptor kinase (Fragment)
ARK3 (Fragment)
At4g21390p-like protein
At4g21390p-like protein
SRK protein
S-related kinase 9 (Fragment)
S-related kinase 9 (Fragment)
S-receptor kinase (Fragment)
ARK3 (Fragment)

SRK
SRK
SRK
SRK
SRK
SRK (Fragment)
SRK
SRK
Uncharacterized protein
Predicted protein
Uncharacterized protein
Predicted protein
Predicted protein
S-locus lectin protein kinase family protein
Predicted protein
Predicted protein
Uncharacterized protein
Predicted protein
Uncharacterized protein
Predicted protein
Predicted protein
Uncharacterized protein
Uncharacterized protein
Uncharacterized protein
Uncharacterized protein
Predicted protein
Uncharacterized protein
Uncharacterized protein
S-locus lectin protein kinase family protein
Predicted protein
Uncharacterized protein
Predicted protein
Predicted protein
Sugar binding protein
Predicted protein
Uncharacterized protein
Uncharacterized protein
S-locus lectin protein kinase family protein
Uncharacterized protein
CES101
S-locus lectin protein kinase family protein

Predicted protein
Predicted protein
Uncharacterized protein
Uncharacterized protein
Predicted protein (Fragment)
Uncharacterized protein
Predicted protein
Predicted protein
At4g21380
Uncharacterized protein
S-locus lectin protein kinase family protein
Predicted protein
Predicted protein
Uncharacterized protein
Predicted protein
Uncharacterized protein
S-locus lectin protein kinase family protein
Uncharacterized protein
Predicted protein
Uncharacterized protein
Predicted protein
S-receptor kinase SRK67 (Fragment)
S-receptor kinase SRK64 (Fragment)
S-receptor kinase SRK06 isoform b (Fragment)
S-receptor kinase SRK69 (Fragment)
S-locus receptor kinase s-haplotype 1
S-receptor kinase SRK63 (Fragment)
S-receptor kinase SRK60 (Fragment)
S-receptor kinase SRK16 (Fragment)
S-receptor kinase SRK08 (Fragment)
S-receptor kinase SRK62 (Fragment)
S-receptor kinase SRK03 (Fragment)
S-receptor kinase SRK25 isoform b (Fragment)
S-receptor kinase SRK65 (Fragment)
S-receptor kinase SRK74 isoform b (Fragment)
S-receptor kinase SRK68 (Fragment)
S-receptor kinase SRK18 isoform a (Fragment)
S-receptor kinase SRK66 isoform a (Fragment)
S-receptor kinase SRK06 isoform a (Fragment)
S-receptor kinase SRK60 (Fragment)
S-receptor kinase SRK25 isoform a (Fragment)

S-receptor kinase (Fragment)
S-receptor kinase (Fragment)
Uncharacterized protein
S receptor kinase (Fragment)
S-receptor kinase (Fragment)
S-receptor kinase 13-7 (Fragment)
Uncharacterized protein
ARK3p
S-locus receptor kinase (Fragment)
S-locus receptor kinase (Fragment)
S-receptor kinase (Fragment)
S receptor kinase SRK09 (Fragment)
At4g21390p-like protein
S receptor kinase SRK31 (Fragment)
S-receptor kinase (Fragment)
S-receptor kinase (Fragment)
Uncharacterized protein
S-receptor kinase 13-17 (Fragment)
S-related kinase 9 (Fragment)
SRK (Fragment)
Self-incompatibility protein 8 (Fragment)
SRKp
S-receptor kinase (Fragment)
S-locus receptor kinase (Fragment)
S-domain receptor kinase (Fragment)
S receptor kinase SRK18 (Fragment)
S receptor kinase (Fragment)
SRKp
SRKp
S receptor kinase SRK16 (Fragment)
S-locus receptor kinase (Fragment)
S-related kinase 10.1A (Fragment)
At4g21390p-like protein
S receptor kinase (Fragment)
S-receptor kinase (Fragment)
S receptor kinase SRK22 (Fragment)
S-related kinase 8 (Fragment)
Uncharacterized protein
S receptor kinase SRK37 (Fragment)
S-receptor kinase 13-15 (Fragment)
S-receptor kinase (Fragment)

SRKa (Fragment)
Uncharacterized protein SRK
S receptor kinase SRK06 (Fragment)
S-receptor kinase 13-12 (Fragment)
S-receptor kinase 13-8 (Fragment)
Uncharacterized protein SRK
S-receptor kinase 13-10 (Fragment)
S-locus receptor kinase (Fragment)
S-receptor kinase (Fragment)
S receptor kinase (Fragment)
Uncharacterized protein ARK3
S-locus receptor kinase (Fragment)
S-locus receptor kinase (Fragment)
Uncharacterized protein SRK
S-locus receptor kinase 25
ARK3p
S-locus receptor kinase (Fragment)
ARK3p
S-locus receptor kinase (Fragment)
ARK3p
S-related kinase 14 (Fragment)
S-locus receptor kinase (Fragment)
S-receptor kinase 13-14 (Fragment)
S-locus receptor kinase (Fragment)
S receptor kinase (Fragment)
S receptor kinase (Fragment)
At4g21390p-like protein
S receptor kinase (Fragment)
S-locus receptor kinase (Fragment)
S receptor kinase (Fragment)
S receptor kinase SRK25 (Fragment)
Uncharacterized protein
S-locus receptor kinase (Fragment)
SRK protein (Fragment)
S-receptor kinase (Fragment)
Uncharacterized protein
S-receptor kinase (Fragment)
S-receptor kinase (Fragment)
S-receptor kinase 13-18 (Fragment)
S receptor kinase (Fragment)
S-receptor kinase 13-30 (Fragment)

S-receptor kinase (Fragment)
S-receptor kinase 13-25 (Fragment)
S-receptor kinase 13-11 (Fragment)
S-receptor kinase (Fragment)
S-receptor kinase 13-28 (Fragment)
Uncharacterized protein
S-receptor kinase 13-6 (Fragment)
S receptor kinase (Fragment)
S-receptor kinase (Fragment)
S-locus receptor kinase 6
S-locus receptor kinase 13-15 (Fragment)
S-receptor kinase 13-2 (Fragment)
S-receptor kinase 13-29 (Fragment)
S-receptor kinase (Fragment)
SRKb
Self-incompatibility protein 8 (Fragment)
Self-incompatibility protein 8 (Fragment)
G-type lectin S-receptor-like serine/threonine-protein kinase SD1-13
G-type lectin S-receptor-like serine/threonine-protein kinase SRK
G-type lectin S-receptor-like serine/threonine-protein kinase CES101
Receptor-like serine/threonine-protein kinase SD1-6
G-type lectin S-receptor-like serine/threonine-protein kinase SD1-29
Receptor-like serine/threonine-protein kinase SD1-7
Receptor-like serine/threonine-protein kinase SD1-8
G-type lectin S-receptor-like serine/threonine-protein kinase SD2-2
G-type lectin S-receptor-like serine/threonine-protein kinase SD1-1
G-type lectin S-receptor-like serine/threonine-protein kinase RLK1
G-type lectin S-receptor-like serine/threonine-protein kinase At4g03230
G-type lectin S-receptor-like serine/threonine-protein kinase At2g19130
PAN domain-containing protein At5g03700
G-type lectin S-receptor-like serine/threonine-protein kinase At1g11280
G-type lectin S-receptor-like serine/threonine-protein kinase At1g11300
G-type lectin S-receptor-like serine/threonine-protein kinase At1g61360
G-type lectin S-receptor-like serine/threonine-protein kinase At1g11410
G-type lectin S-receptor-like serine/threonine-protein kinase At1g61420
G-type lectin S-receptor-like serine/threonine-protein kinase At1g61440
G-type lectin S-receptor-like serine/threonine-protein kinase At1g61480
EP1-like glycoprotein 1
EP1-like glycoprotein 3
EP1-like glycoprotein 4
G-type lectin S-receptor-like serine/threonine-protein kinase B120

Putative inactive G-type lectin S-receptor-like serine/threonine-protein kinase SRK
G-type lectin S-receptor-like serine/threonine-protein kinase At1g11330
G-type lectin S-receptor-like serine/threonine-protein kinase At1g11303
G-type lectin S-receptor-like serine/threonine-protein kinase At1g61370
G-type lectin S-receptor-like serine/threonine-protein kinase At1g61460
G-type lectin S-receptor-like serine/threonine-protein kinase At1g61550
G-type lectin S-receptor-like serine/threonine-protein kinase At1g61400
G-type lectin S-receptor-like serine/threonine-protein kinase At1g61490
G-type lectin S-receptor-like serine/threonine-protein kinase At1g61390
Putative G-type lectin S-receptor-like serine/threonine-protein kinase At1g61610
G-type lectin S-receptor-like serine/threonine-protein kinase At1g61500
G-type lectin S-receptor-like serine/threonine-protein kinase At1g67520
G-type lectin S-receptor-like serine/threonine-protein kinase At1g61430
G-type lectin S-receptor-like serine/threonine-protein kinase RKS1
G-type lectin S-receptor-like serine/threonine-protein kinase SD3-1
G-type lectin S-receptor-like serine/threonine-protein kinase At4g11900
G-type lectin S-receptor-like serine/threonine-protein kinase At4g27290
Lectin protein kinase family protein
S-locus lectin protein kinase family protein
S-locus lectin protein kinase family protein
D-mannose binding lectin protein with Apple-like carbohydrate-binding domain-containing protein
S-locus lectin protein kinase family protein
S-locus lectin protein kinase family protein
S-locus lectin protein kinase family protein
S glycoprotein
Uncharacterized protein
Receptor-like protein kinase 4
Uncharacterized protein
G-type lectin S-receptor-like Serine/Threonine-kinase
S domain subfamily 1 receptor-like kinase
SD1-29
Uncharacterized protein
S-locus lectin protein kinase family protein
Uncharacterized protein
Uncharacterized protein
Uncharacterized protein
Uncharacterized protein
Uncharacterized protein
Lectin protein kinase family protein
S domain subfamily receptor-like kinase
Uncharacterized protein

Uncharacterized protein
Uncharacterized protein
RK3
S-locus lectin protein kinase family protein
Uncharacterized protein
Uncharacterized protein
Uncharacterized protein
S-locus lectin protein kinase family protein
S-locus lectin protein kinase family protein
S-locus lectin protein kinase family protein
S-locus lectin protein kinase family protein
NS1 glycoprotein, putative; 15463-17897
G-type lectin S-receptor-like Serine/Threonine-kinase
G-type lectin S-receptor-like Serine/Threonine-kinase
Uncharacterized protein
Lectin protein kinase family protein
Uncharacterized protein
Lectin protein kinase family protein
Receptor kinase 3
S-receptor kinase
Uncharacterized protein
G-type lectin S-receptor-like Serine/Threonine-kinase
RFO3
Uncharacterized protein
S-locus glycoprotein family protein
Uncharacterized protein
Uncharacterized protein
Uncharacterized protein
Uncharacterized protein
Uncharacterized protein
S-locus lectin protein kinase family protein
S-locus lectin protein kinase family protein
G-type lectin S-receptor-like Serine/Threonine-kinase
Lectin protein kinase family protein
S-locus lectin protein kinase family protein
Uncharacterized protein
Uncharacterized protein
G-type lectin S-receptor-like Serine/Threonine-kinase
S-locus lectin protein kinase family protein
S-locus lectin protein kinase family protein
Uncharacterized protein

S-locus lectin protein kinase family protein
Uncharacterized protein
S-locus lectin protein kinase family protein
G-type lectin S-receptor-like Serine/Threonine-kinase
Uncharacterized protein
Uncharacterized protein
S-locus lectin protein kinase family protein
Uncharacterized protein
S-locus lectin protein kinase family protein
AtS1 protein
Uncharacterized protein
S-locus lectin protein kinase family protein
S-locus lectin protein kinase family protein
S-locus lectin protein kinase family protein
S-locus lectin protein kinase family protein
Uncharacterized protein
Lectin protein kinase family protein
B120
S-locus lectin protein kinase family protein
Lectin protein kinase family protein
S-locus lectin protein kinase family protein
G-type lectin S-receptor-like Serine/Threonine-kinase
S-locus lectin protein kinase family protein
Receptor kinase 1
S-locus lectin protein kinase family protein
SD1-13
S-locus lectin protein kinase family protein
Lectin protein kinase family protein
S-domain-1 29
Truncated At4g21370
S-locus lectin protein kinase family protein
Uncharacterized protein
Uncharacterized protein
Lectin protein kinase family protein
S-locus lectin protein kinase family protein
Uncharacterized protein
Uncharacterized protein
Lectin protein kinase family protein
S-locus lectin protein kinase family protein
Receptor kinase 2
Uncharacterized protein

Lectin protein kinase family protein
RK1
S-locus lectin protein kinase family protein
S-locus lectin protein kinase family protein
Uncharacterized protein
Uncharacterized protein
Uncharacterized protein
Uncharacterized protein
Uncharacterized protein
Uncharacterized protein
Uncharacterized protein
Uncharacterized protein
Uncharacterized protein
Uncharacterized protein
Uncharacterized protein
Uncharacterized protein
Uncharacterized protein
Uncharacterized protein
Uncharacterized protein
Uncharacterized protein
Uncharacterized protein
Uncharacterized protein
Uncharacterized protein
Uncharacterized protein
Uncharacterized protein
Uncharacterized protein
Uncharacterized protein
Uncharacterized protein
Uncharacterized protein
Uncharacterized protein
Uncharacterized protein
Uncharacterized protein
Uncharacterized protein
Uncharacterized protein
Uncharacterized protein
Uncharacterized protein
Uncharacterized protein
Uncharacterized protein
Uncharacterized protein
Uncharacterized protein
Uncharacterized protein

Uncharacterized protein
Uncharacterized protein
Uncharacterized protein
Uncharacterized protein
Uncharacterized protein
Uncharacterized protein
Uncharacterized protein
Uncharacterized protein
Uncharacterized protein
Uncharacterized protein
Uncharacterized protein
Uncharacterized protein
Uncharacterized protein
Uncharacterized protein
Uncharacterized protein
Uncharacterized protein
Uncharacterized protein
Uncharacterized protein
Uncharacterized protein
Uncharacterized protein
Uncharacterized protein
Uncharacterized protein
Uncharacterized protein
Uncharacterized protein
Uncharacterized protein
Uncharacterized protein
Uncharacterized protein
Uncharacterized protein
Uncharacterized protein
Uncharacterized protein
Uncharacterized protein
Uncharacterized protein
Uncharacterized protein
Uncharacterized protein
Uncharacterized protein
Uncharacterized protein
Uncharacterized protein
Uncharacterized protein
Uncharacterized protein
Uncharacterized protein
Uncharacterized protein

Uncharacterized protein
Uncharacterized protein
Uncharacterized protein
Uncharacterized protein
Uncharacterized protein
Uncharacterized protein
Uncharacterized protein
Uncharacterized protein
Uncharacterized protein
Uncharacterized protein
Uncharacterized protein
Uncharacterized protein
Uncharacterized protein
Uncharacterized protein
Uncharacterized protein
Uncharacterized protein
Uncharacterized protein
Uncharacterized protein
Uncharacterized protein
Uncharacterized protein
Uncharacterized protein
Uncharacterized protein
Uncharacterized protein
Uncharacterized protein
Uncharacterized protein
Uncharacterized protein
Uncharacterized protein
Uncharacterized protein
Uncharacterized protein
Uncharacterized protein
Uncharacterized protein
Uncharacterized protein
Uncharacterized protein
Uncharacterized protein
Uncharacterized protein
Uncharacterized protein
Uncharacterized protein
Uncharacterized protein
Uncharacterized protein
Uncharacterized protein
Uncharacterized protein

Uncharacterized protein
Uncharacterized protein
Uncharacterized protein
Uncharacterized protein
Uncharacterized protein
Uncharacterized protein
Uncharacterized protein
Uncharacterized protein
Uncharacterized protein
Uncharacterized protein
Uncharacterized protein
Uncharacterized protein
Uncharacterized protein
Uncharacterized protein
Uncharacterized protein
Uncharacterized protein
Uncharacterized protein
Uncharacterized protein
Uncharacterized protein
Uncharacterized protein
Uncharacterized protein
Uncharacterized protein
Uncharacterized protein
Uncharacterized protein
Uncharacterized protein
Uncharacterized protein
Uncharacterized protein
Uncharacterized protein
Uncharacterized protein
Uncharacterized protein
Uncharacterized protein
Uncharacterized protein
Uncharacterized protein
Uncharacterized protein
Uncharacterized protein
Uncharacterized protein
Uncharacterized protein
Uncharacterized protein
Uncharacterized protein
Uncharacterized protein
Uncharacterized protein

Uncharacterized protein
Uncharacterized protein
Uncharacterized protein
Uncharacterized protein
Uncharacterized protein
Uncharacterized protein
Uncharacterized protein
Uncharacterized protein
Uncharacterized protein
Uncharacterized protein
Uncharacterized protein
Uncharacterized protein
Uncharacterized protein
Uncharacterized protein
Uncharacterized protein
Uncharacterized protein
Uncharacterized protein
Uncharacterized protein
Uncharacterized protein
Uncharacterized protein
Uncharacterized protein
Uncharacterized protein
Uncharacterized protein
Uncharacterized protein
Uncharacterized protein
Uncharacterized protein
Uncharacterized protein
Uncharacterized protein
Uncharacterized protein
Uncharacterized protein
Uncharacterized protein
Uncharacterized protein
Uncharacterized protein
Uncharacterized protein
Uncharacterized protein
Uncharacterized protein
Uncharacterized protein
Uncharacterized protein
Uncharacterized protein
Uncharacterized protein
Uncharacterized protein

Uncharacterized protein
Uncharacterized protein
Uncharacterized protein
Uncharacterized protein
Uncharacterized protein
Uncharacterized protein
Uncharacterized protein
Uncharacterized protein
Resistance protein (Fragment)
Uncharacterized protein
Uncharacterized protein
Uncharacterized protein
Uncharacterized protein
Uncharacterized protein
Uncharacterized protein
Uncharacterized protein
Uncharacterized protein
Uncharacterized protein
Uncharacterized protein
Uncharacterized protein
Uncharacterized protein
Uncharacterized protein
Uncharacterized protein
Uncharacterized protein
Uncharacterized protein
Uncharacterized protein
Uncharacterized protein
Uncharacterized protein
Uncharacterized protein
Uncharacterized protein
Uncharacterized protein
Uncharacterized protein
Uncharacterized protein
Uncharacterized protein
Uncharacterized protein
Uncharacterized protein
Uncharacterized protein
Uncharacterized protein
Uncharacterized protein
Uncharacterized protein
Uncharacterized protein

Uncharacterized protein
Uncharacterized protein
Uncharacterized protein
Uncharacterized protein
Uncharacterized protein
Uncharacterized protein
Uncharacterized protein
Uncharacterized protein
Uncharacterized protein
Uncharacterized protein
Uncharacterized protein
Uncharacterized protein
Uncharacterized protein
Uncharacterized protein
Uncharacterized protein
Uncharacterized protein
Uncharacterized protein
Uncharacterized protein
Uncharacterized protein
Uncharacterized protein
Uncharacterized protein
Uncharacterized protein
Uncharacterized protein
Uncharacterized protein
Uncharacterized protein
Uncharacterized protein
Uncharacterized protein
Uncharacterized protein (Fragment)
Uncharacterized protein
Papilin (Fragment)
Uncharacterized protein
Uncharacterized protein
Neurotrypsin (Fragment)
Uncharacterized protein (Fragment)
EGF-like domain-containing protein
S-locus glycoprotein domain-containing protein
Receptor-like protein kinase 4
Apple-like protein
S-locus glycoprotein domain-containing protein
Apple-like protein
Apple-like protein

Receptor protein kinase ZmPK1
Apple-like protein
Bulb-type lectin domain-containing protein
Apple-like protein
S-locus glycoprotein domain-containing protein
G-type lectin S-receptor-like serine/threonine-protein kinase
Apple-like protein
S-locus lectin protein kinase family protein
Curculin-like (Mannose-binding) lectin family protein / PAN domain-containing protein
S-locus glycoprotein domain-containing protein
S-locus glycoprotein domain-containing protein
S-locus glycoprotein domain-containing protein
S-locus glycoprotein domain-containing protein
EGF-like domain-containing protein
Uncharacterized protein
Uncharacterized protein
S-locus glycoprotein domain-containing protein
Apple-like protein
Apple-like protein
Bulb-type lectin domain-containing protein
GPCR kinase
S-locus glycoprotein domain-containing protein
Uncharacterized protein
S-locus glycoprotein domain-containing protein
Bulb-type lectin domain-containing protein
S-domain-1 29
S-locus glycoprotein domain-containing protein
G-type lectin S-receptor-like serine/threonine-protein kinase
Uncharacterized protein
EGF-like domain-containing protein
S-locus glycoprotein domain-containing protein
S-locus glycoprotein domain-containing protein
Apple-like protein
Bulb-type lectin domain-containing protein
S-locus glycoprotein domain-containing protein
Apple-like protein
S-locus glycoprotein domain-containing protein
S-locus glycoprotein domain-containing protein
Apple-like protein
EGF-like domain-containing protein
Receptor protein kinase ZmPK1

Apple-like protein
S-locus glycoprotein domain-containing protein
EGF-like domain-containing protein
Apple-like protein
Bulb-type lectin domain-containing protein
PAN domain-containing protein
Uncharacterized protein
S-locus lectin protein kinase family protein
Receptor-like protein kinase 4
Apple-like protein
S-locus glycoprotein domain-containing protein
Receptor kinase 3
Uncharacterized protein
Uncharacterized protein
G-type lectin S-receptor-like serine/threonine-protein kinase
Apple-like protein
S-locus glycoprotein domain, Bulb-type lectin domain, PAN/Apple domain protein
Uncharacterized protein
S-locus glycoprotein domain-containing protein
Apple-like protein
S-receptor kinase-like protein 1
G-type lectin S-receptor-like serine/threonine-protein kinase
S-locus glycoprotein domain-containing protein
Receptor kinase 3
S-locus glycoprotein domain-containing protein
Bulb-type lectin domain-containing protein
S-locus glycoprotein domain-containing protein
Apple-like protein
Apple-like protein
S-locus glycoprotein domain-containing protein
S-locus lectin protein kinase family protein
Apple-like protein
G-type lectin S-receptor-like serine/threonine-protein kinase
S-locus glycoprotein domain, Bulb-type lectin domain, PAN/Apple domain protein
S-locus glycoprotein domain-containing protein
S-locus glycoprotein domain-containing protein
S-locus glycoprotein domain-containing protein
G-type lectin S-receptor-like serine/threonine-protein kinase
S-locus lectin protein kinase family protein
S-locus glycoprotein domain-containing protein
Apple-like protein

Receptor kinase 3
Apple-like protein
Apple-like protein
Apple-like protein
S-locus glycoprotein domain-containing protein
S-locus glycoprotein domain-containing protein
S-locus glycoprotein domain-containing protein
Apple-like protein
Uncharacterized protein
Apple-like protein
Apple-like protein
Bulb-type lectin domain-containing protein
Apple-like protein
Bulb-type lectin domain-containing protein
Uncharacterized protein
Bulb-type lectin domain-containing protein
G-type lectin S-receptor-like serine/threonine-protein kinase
Lectin protein kinase family protein
S-locus glycoprotein domain-containing protein
S-locus glycoprotein domain-containing protein
S-locus glycoprotein domain-containing protein
S-locus glycoprotein domain-containing protein
DNA repair metallo-beta-lactamase family protein
Receptor kinase 3
S-locus glycoprotein domain, Bulb-type lectin domain, PAN/Apple domain protein
Bulb-type lectin domain-containing protein
Uncharacterized protein
Uncharacterized protein
Uncharacterized protein
Uncharacterized protein
Uncharacterized protein
Uncharacterized protein
Uncharacterized protein
Pk3
Uncharacterized protein
Uncharacterized protein
Pk3
Uncharacterized protein
Uncharacterized protein
Uncharacterized protein
Uncharacterized protein

Uncharacterized protein
Uncharacterized protein
Uncharacterized protein
Uncharacterized protein
Uncharacterized protein
Uncharacterized protein
Uncharacterized protein
Uncharacterized protein
Uncharacterized protein
Uncharacterized protein
Uncharacterized protein
Uncharacterized protein
Uncharacterized protein
Uncharacterized protein
Uncharacterized protein
Uncharacterized protein
Uncharacterized protein
Uncharacterized protein
Uncharacterized protein
Uncharacterized protein
Uncharacterized protein
Uncharacterized protein
Uncharacterized protein
Uncharacterized protein
Uncharacterized protein
Uncharacterized protein
Uncharacterized protein
Uncharacterized protein
Uncharacterized protein
Uncharacterized protein
Uncharacterized protein
Uncharacterized protein
Uncharacterized protein
Uncharacterized protein
Uncharacterized protein (Fragment)
Uncharacterized protein
Uncharacterized protein
Uncharacterized protein
Uncharacterized protein
Putative lipoprotein YfhM
Uncharacterized protein
Uncharacterized protein

Apple-like protein
Uncharacterized protein
Uncharacterized protein
Uncharacterized protein
Uncharacterized protein
Uncharacterized protein
Uncharacterized protein
Uncharacterized protein
Uncharacterized protein
Uncharacterized protein
Uncharacterized protein
Uncharacterized protein
Uncharacterized protein
Uncharacterized protein
Uncharacterized protein
Uncharacterized protein
Hypothetical low complexity protein
Uncharacterized protein
Uncharacterized protein
Uncharacterized protein
Uncharacterized protein
Uncharacterized protein
Uncharacterized protein
Uncharacterized protein
Uncharacterized protein
Uncharacterized protein
Uncharacterized protein
Uncharacterized protein
Uncharacterized protein
Uncharacterized protein
Uncharacterized protein
Uncharacterized protein
Uncharacterized protein
Uncharacterized protein
Uncharacterized protein
Uncharacterized protein
Uncharacterized protein
Uncharacterized protein
Uncharacterized protein
Uncharacterized protein
Uncharacterized protein

Uncharacterized protein
Uncharacterized protein
Uncharacterized protein
Uncharacterized protein
Uncharacterized protein
Glycosyltransferase family 31
Uncharacterized protein
Uncharacterized protein
Uncharacterized protein
Uncharacterized protein
Uncharacterized protein
Uncharacterized protein
Glycosyltransferase family 31 protein
Uncharacterized protein
Glycosyltransferase family 31
Uncharacterized protein
Uncharacterized protein
Uncharacterized protein
Uncharacterized protein
Uncharacterized protein
Family 31 glycosyltransferase
Uncharacterized protein
Uncharacterized protein
Uncharacterized protein
Aspergillus niger contig An09c0040, genomic contig
Uncharacterized protein
Uncharacterized protein
Uncharacterized protein
Glycosyltransferase family 31
Uncharacterized protein
Uncharacterized protein
Uncharacterized protein
Uncharacterized protein
Uncharacterized protein
Uncharacterized protein
Uncharacterized protein
Uncharacterized protein
Uncharacterized protein
Uncharacterized protein
Uncharacterized protein
Uncharacterized protein

Uncharacterized protein
Uncharacterized protein
Uncharacterized protein
Uncharacterized protein
Fringe-like
N/apple PAN domain protein
Fringe-like
Fringe-like protein
Uncharacterized protein
Uncharacterized protein
Uncharacterized protein
Uncharacterized protein
Uncharacterized protein
Uncharacterized protein
Uncharacterized protein
Uncharacterized protein
Uncharacterized protein
Uncharacterized protein
Uncharacterized protein
Uncharacterized protein
Uncharacterized protein
Uncharacterized protein
Uncharacterized protein
Uncharacterized protein
Uncharacterized protein
Uncharacterized protein
Uncharacterized protein
Uncharacterized protein
Uncharacterized protein
Uncharacterized protein (Fragment)
Uncharacterized protein
Uncharacterized protein
Uncharacterized protein
Uncharacterized protein
Uncharacterized protein
Uncharacterized protein
Reticulocyte-binding protein 2 homolog a
Uncharacterized protein
Uncharacterized protein
Uncharacterized protein
Uncharacterized protein

Uncharacterized protein
Uncharacterized protein
Uncharacterized protein
Uncharacterized protein
Hepatocyte growth factor
Uncharacterized protein
Plasminogen
Uncharacterized protein
Uncharacterized protein
Uncharacterized protein
Plasminogen
Coelomotrypsin
Hepatocyte growth factor
Hepatocyte growth factor
Hepatocyte growth factor
Plasminogen
Uncharacterized protein
Plasminogen
Uncharacterized protein
Uncharacterized protein
Uncharacterized protein
Uncharacterized protein
Uncharacterized protein
Uncharacterized protein
Uncharacterized protein
Uncharacterized protein
Ras-related protein RabJ
Neurotrypsin (Fragment)
Uncharacterized protein
Uncharacterized protein
Uncharacterized protein
Cuticlin-1 (Fragment)
Translation initiation factor IF-2 (Fragment)
Alpha-2-macroglobulin domain-containing protein
Uncharacterized protein
Uncharacterized protein
Uncharacterized protein
Uncharacterized protein
Uncharacterized protein
Uncharacterized protein
Uncharacterized protein

Uncharacterized protein
Uncharacterized protein
Uncharacterized protein
Uncharacterized protein
Uncharacterized protein
Uncharacterized protein
Uncharacterized protein
Uncharacterized protein
Uncharacterized protein
Uncharacterized protein
Uncharacterized protein
Uncharacterized protein
Uncharacterized protein
Uncharacterized protein
Uncharacterized protein
Uncharacterized protein
Uncharacterized protein (Fragment)
Uncharacterized protein (Fragment)
Uncharacterized protein
Uncharacterized protein
Uncharacterized protein
Uncharacterized protein
Uncharacterized protein
Uncharacterized protein (Fragment)
Uncharacterized protein (Fragment)
Uncharacterized protein
Uncharacterized protein (Fragment)
Uncharacterized protein
Uncharacterized protein
Uncharacterized protein (Fragment)
Uncharacterized protein
Uncharacterized protein
Uncharacterized protein
Putative beta-glucosidase
Uncharacterized protein
Uncharacterized protein
Putative cytosolic iron-sulfur protein assembly protein 1
Tyrosine--tRNA ligase
Uncharacterized protein
Uncharacterized protein
Uncharacterized protein

MFS general substrate transporter
Uncharacterized protein
Putative MFS monocarboxylate transporter
SRF-TF-domain-containing protein
Uncharacterized protein
Uncharacterized protein
Uncharacterized protein
Uncharacterized protein
Uncharacterized protein
Uncharacterized protein
Uncharacterized protein
Carbohydrate-binding module family 50 protein
Uncharacterized protein
Uncharacterized protein
Uncharacterized protein
Uncharacterized protein
Carbohydrate esterase family 1 protein
Uncharacterized protein
Uncharacterized protein
Uncharacterized protein
Uncharacterized protein
hepatocyte growth factor-like isoform X3
coagulation factor XI-like
thyroglobulin
hepatocyte growth factor-like protein
plasminogen-like
Plasminogen
hepatocyte growth factor-like isoform X2
Hepatocyte growth factor
hepatocyte growth factor-like isoform X1
coagulation factor XI-like
Uncharacterized protein (Fragment)
Uncharacterized protein (Fragment)
Uncharacterized protein
Peptidyl-prolyl cis-trans isomerase D
Uncharacterized protein
Uncharacterized protein
Uncharacterized protein
Uncharacterized protein
Uncharacterized protein
Peptidyl-prolyl cis-trans isomerase (Fragment)

Uncharacterized protein (Fragment)
Uncharacterized protein (Fragment)
Mannan endo-1,4-beta-mannosidase 4
Uncharacterized protein
Uncharacterized protein (Fragment)
Uncharacterized protein (Fragment)
Uncharacterized protein (Fragment)
Uncharacterized protein (Fragment)
Uncharacterized protein
Uncharacterized protein
Uncharacterized protein (Fragment)
Uncharacterized protein (Fragment)
Low-density lipoprotein receptor-related protein 11
Translation initiation factor IF-2 (Fragment)
Uncharacterized protein
Uncharacterized protein
Uncharacterized protein (Fragment)
Uncharacterized protein
EGF-like domain-containing protein 2
Uncharacterized protein
Uncharacterized protein
Translation initiation factor IF-2
Uncharacterized protein
Uncharacterized protein
Uncharacterized protein
Translation initiation factor IF-2
Uncharacterized protein
Uncharacterized protein
Uncharacterized protein
Uncharacterized protein
Uncharacterized protein
coagulation factor XI
Hepatocyte growth factor
hepatocyte growth factor-like protein
Hepatocyte growth factor
plasminogen isoform X2
Plasminogen
Plasminogen (Fragment)
Coagulation factor XI (Fragment)
Uncharacterized protein
Uncharacterized protein (Fragment)

Uncharacterized protein
Uncharacterized protein
Pan domain-containing protein (Fragment)
Pan domain-containing protein
PAN domain-containing protein
PAN domain-containing protein
Uncharacterized protein
Uncharacterized protein
Uncharacterized protein
Uncharacterized protein
Uncharacterized protein
Uncharacterized protein
Uncharacterized protein
Uncharacterized protein
Uncharacterized protein
Uncharacterized protein
Uncharacterized protein
Uncharacterized protein
Uncharacterized protein
Uncharacterized protein
Uncharacterized protein
Uncharacterized protein
Uncharacterized protein
Uncharacterized protein
Uncharacterized protein
PAN domain-containing protein
Uncharacterized protein
Uncharacterized protein
Uncharacterized protein (Fragment)
Uncharacterized protein
PAN domain containing protein
Uncharacterized protein
Uncharacterized protein
Uncharacterized protein
Uncharacterized protein
Methionine-rich protein
Uncharacterized protein
Uncharacterized protein
Uncharacterized protein
Uncharacterized protein
Apple-like protein

PAN domain containing protein
Alpha-2-macroglobulin domain protein
Uncharacterized protein
1,3(4)-beta-glucanase
Uncharacterized protein
Putative microneme protein
Uncharacterized protein
PAN domain-containing protein
PAN/Apple domain-containing protein
PAN domain-containing protein
PAN domain-containing protein
PAN domain-containing protein
Microneme protein MIC4
PAN domain-containing protein
Microneme protein MIC17B
Microneme protein MIC17A
Microneme protein MIC17A
Uncharacterized protein
PAN/Apple domain-containing protein
Microneme protein MIC17A
Uncharacterized protein
PAN domain-containing protein
Uncharacterized protein
Subtilisin SUB12
Uncharacterized protein
Uncharacterized protein
Uncharacterized protein
Uncharacterized protein
Uncharacterized protein
Uncharacterized protein
Uncharacterized protein
Uncharacterized protein
Uncharacterized protein
Uncharacterized protein
Uncharacterized protein
Uncharacterized protein
Uncharacterized protein
Uncharacterized protein
Uncharacterized protein
Uncharacterized protein
Uncharacterized protein

Uncharacterized protein
Uncharacterized protein
Uncharacterized protein
Uncharacterized protein
Uncharacterized protein
Uncharacterized protein
Uncharacterized protein
Uncharacterized protein
Uncharacterized protein
Uncharacterized protein
Uncharacterized protein
Uncharacterized protein
Uncharacterized protein
Uncharacterized protein
Uncharacterized protein
Uncharacterized protein
Uncharacterized protein
Uncharacterized protein
Uncharacterized protein
Uncharacterized protein
Uncharacterized protein
Uncharacterized protein
Uncharacterized protein
Uncharacterized protein
Uncharacterized protein
Uncharacterized protein
Uncharacterized protein
Uncharacterized protein
Uncharacterized protein
Uncharacterized protein
Uncharacterized protein
Uncharacterized protein
Uncharacterized protein
Uncharacterized protein
Uncharacterized protein
Uncharacterized protein
Uncharacterized protein
Uncharacterized protein
Uncharacterized protein
Copper radical oxidase
Copper radical oxidase

Copper radical oxidase
Uncharacterized protein
Uncharacterized protein (Fragment)
Uncharacterized protein (Fragment)
Uncharacterized protein
Uncharacterized protein (Fragment)
Uncharacterized protein
Copper radical oxidase
Uncharacterized protein
Uncharacterized protein (Fragment)
Copper radical oxidase
Carbohydrate-binding module family 18 protein
Uncharacterized protein (Fragment)
Uncharacterized protein
Uncharacterized protein
Uncharacterized protein
Uncharacterized protein
Uncharacterized protein
Uncharacterized protein (Fragment)
Uncharacterized protein
Uncharacterized protein
Uncharacterized protein (Fragment)
Copper radical oxidase
Copper radical oxidase
Aly8/ARK3-related protein (Fragment)
SRK-related protein (Fragment)
SRK-related protein (Fragment)
SRK-related protein (Fragment)
SRK-related protein (Fragment)
Aly327420-related protein (Fragment)
SRK-related protein (Fragment)
SRK-related protein (Fragment)
Aly8/ARK3-related protein (Fragment)
SRK-related protein (Fragment)
SRK-related protein (Fragment)
SRK-related protein (Fragment)
SRK-related protein (Fragment)
SRK-related protein (Fragment)
SRK-related protein (Fragment)
SRK-related protein (Fragment)
Aly327420-related protein (Fragment)

SRK-related protein (Fragment)
SRK-related protein (Fragment)
SRK-related protein (Fragment)
Aly327420-related protein (Fragment)
SRK-related protein (Fragment)
SRK-related protein (Fragment)
SRK-related protein (Fragment)
Aly327420-related protein (Fragment)
Aly8/ARK3-related protein (Fragment)
Uncharacterized protein
Uncharacterized protein
Uncharacterized protein
Uncharacterized protein (Fragment)
Uncharacterized protein
Uncharacterized protein
Uncharacterized protein
Uncharacterized protein
Uncharacterized protein (Fragment)
Uncharacterized protein
Uncharacterized protein
Pan domain containing protein
Uncharacterized protein
Beta-fructofuranosidase-like protein, putative (Fragment)
Glycoside hydrolase, putative
Protein kinase, putative
Uncharacterized protein
Uncharacterized protein
Uncharacterized protein
Uncharacterized protein
Uncharacterized protein
Uncharacterized protein
Hepatocyte growth factor-like protein
Plasminogen (Fragment)
Plasma kallikrein (Fragment)
Hepatocyte growth factor (Fragment)
Coagulation factor XI
Plasminogen
Hepatocyte growth factor
Hepatocyte growth factor
Coagulation factor XI (Fragment)
Blood coagulation factor XI (Fragment)

F11 protein
Plasminogen
Hepatocyte growth factor-like protein
Hepatocyte growth factor
Hepatocyte growth factor-like protein
Coagulation factor XI
Coagulation factor XI
Coagulation factor XI
Plasma kallikrein
Hepatocyte growth factor-like protein
Hepatocyte growth factor
PLG protein
Uncharacterized protein
Putative carbohydrate-binding wsc protein
Putative Apple protein
Uncharacterized protein
Uncharacterized protein
Uncharacterized protein
Putative pan domain containing protein
Uncharacterized protein
Uncharacterized protein
Antigen B membrane (Fragment)
Coagulation factor XI-like
SCP-like extracellular
Uncharacterized protein
Fibroblast growth factor receptor-like (Fragment)
Uncharacterized protein
Uncharacterized protein (Fragment)
Antigen B membrane
Antigen B membrane (Fragment)
Uncharacterized protein
Uncharacterized protein
Antigen B membrane
Uncharacterized protein (Fragment)
Uncharacterized protein
Uncharacterized protein
Cellulose binding elicitor
Golgi-associated plant pathogenesis-related 1-like
Fibroblast growth factor receptor-like
SCP-like extracellular
EF-hand domain-containing D1

Antigen B membrane
Fibroblast growth factor receptor-like (Fragment)
Uncharacterized protein
Uncharacterized protein
Cellulose binding elicitor
Antigen B membrane
Cysteine dioxygenase (Fragment)
Uncharacterized protein
Uncharacterized protein
Uncharacterized protein
Uncharacterized protein
Uncharacterized protein
Uncharacterized protein
Uncharacterized protein
Uncharacterized protein
Uncharacterized protein
Uncharacterized protein
Uncharacterized protein
Uncharacterized protein
Uncharacterized protein
Uncharacterized protein
Uncharacterized protein
Uncharacterized protein
Uncharacterized protein
Uncharacterized protein
Uncharacterized protein
Uncharacterized protein
Uncharacterized protein
Uncharacterized protein
Uncharacterized protein
Uncharacterized protein
Uncharacterized protein
Uncharacterized protein
Uncharacterized protein
Uncharacterized protein
Uncharacterized protein
Uncharacterized protein
Uncharacterized protein
Uncharacterized protein
Uncharacterized protein
Uncharacterized protein

Uncharacterized protein
Uncharacterized protein
Uncharacterized protein
Uncharacterized protein
Uncharacterized protein
Uncharacterized protein
Uncharacterized protein
Uncharacterized protein
Uncharacterized protein
Uncharacterized protein
Uncharacterized protein
Uncharacterized protein
Uncharacterized protein
Uncharacterized protein
Uncharacterized protein
Uncharacterized protein (Fragment)
Uncharacterized protein
Uncharacterized protein
Uncharacterized protein
Uncharacterized protein (Fragment)
Uncharacterized protein
Uncharacterized protein
Uncharacterized protein
Uncharacterized protein
Uncharacterized protein
Uncharacterized protein
Uncharacterized protein
Uncharacterized protein
Uncharacterized protein
Uncharacterized protein
Uncharacterized protein
Uncharacterized protein
Uncharacterized protein
Uncharacterized protein
Uncharacterized protein
Uncharacterized protein
Uncharacterized protein
Uncharacterized protein
Uncharacterized protein
Uncharacterized protein
Uncharacterized protein

Uncharacterized protein
Uncharacterized protein
Uncharacterized protein
Uncharacterized protein
Uncharacterized protein
Uncharacterized protein
Uncharacterized protein
Uncharacterized protein
Uncharacterized protein
Uncharacterized protein
Uncharacterized protein
Uncharacterized protein
Uncharacterized protein
Uncharacterized protein
Uncharacterized protein
Uncharacterized protein
Uncharacterized protein
Uncharacterized protein
Uncharacterized protein
Uncharacterized protein
Uncharacterized protein
Uncharacterized protein
Uncharacterized protein
Uncharacterized protein
Uncharacterized protein (Fragment)
Uncharacterized protein
Uncharacterized protein
Uncharacterized protein
Uncharacterized protein
Uncharacterized protein
Uncharacterized protein
Uncharacterized protein
Uncharacterized protein
Uncharacterized protein
Uncharacterized protein
Apple domain-containing protein
Uncharacterized protein
Apple domain-containing protein
Apple domain-containing protein
Apple domain-containing protein
Apple domain-containing protein

Apple domain-containing protein
Apple domain-containing protein
PAN domain-containing protein
PAN domain-containing protein
Uncharacterized protein
Uncharacterized protein
Apple domain-containing protein
Blr1165 protein
Apple domain-containing protein
Uncharacterized protein
Apple domain-containing protein
Apple domain-containing protein
PAN domain-containing protein
PAN domain-containing protein
PAN domain-containing protein
PAN domain-containing protein
PAN domain-containing protein
Apple domain-containing protein
Apple domain-containing protein
Uncharacterized protein
Uncharacterized protein
Apple domain-containing protein
Apple domain-containing protein
Apple domain-containing protein
Apple domain-containing protein
PAN domain-containing protein
PAN domain-containing protein
Apple domain-containing protein
PAN domain-containing protein
Apple domain-containing protein
Uncharacterized protein
Apple domain-containing protein
Apple domain-containing protein
Uncharacterized protein
Uncharacterized protein
Apple domain-containing protein
Apple domain-containing protein
Apple domain-containing protein
Apple domain-containing protein
PAN domain-containing protein
Uncharacterized protein

Uncharacterized protein
Uncharacterized protein
Uncharacterized protein
Uncharacterized protein
Uncharacterized protein
Uncharacterized protein
Apple domain-containing protein
Uncharacterized protein
Apple domain-containing protein
Apple domain-containing protein
Apple domain-containing protein
Apple domain-containing protein
Apple domain-containing protein
Apple domain-containing protein
Apple domain-containing protein
Apple domain-containing protein
Apple domain-containing protein
Apple domain-containing protein
PAN domain-containing protein
PAN domain-containing protein
Monoamine oxidase
Apple domain-containing protein
Uncharacterized protein
Uncharacterized protein
PAN domain-containing protein
PAN domain-containing protein
Apple domain-containing protein
Apple domain-containing protein
Apple domain-containing protein
Apple domain-containing protein
Apple domain-containing protein
Apple domain-containing protein
Apple domain-containing protein
Apple domain-containing protein
Uncharacterized protein
Uncharacterized protein
Uncharacterized protein
Uncharacterized protein
Uncharacterized protein
Apple domain-containing protein
Apple domain-containing protein

Apple domain-containing protein
PAN domain-containing protein
PAN domain-containing protein
Uncharacterized protein
Uncharacterized protein
Uncharacterized protein
PAN domain-containing protein
PAN domain-containing protein
PAN domain-containing protein
Apple domain-containing protein
Uncharacterized protein
Apple domain-containing protein
Uncharacterized protein
Uncharacterized protein
Uncharacterized protein
Apple domain-containing protein
Apple domain-containing protein
Apple domain-containing protein
PAN domain
PAN domain protein
Apple domain-containing protein
PAN domain PAN-like domain containing protein
PAN domain containing protein
Uncharacterized protein
Apple domain-containing protein
Apple domain-containing protein
Chitinase
Chitinase
Uncharacterized protein
Uncharacterized protein (Fragment)
Uncharacterized protein
Uncharacterized protein
Uncharacterized protein
Uncharacterized protein
Uncharacterized protein
Uncharacterized protein
Uncharacterized protein
Uncharacterized protein
Uncharacterized protein
Uncharacterized protein (Fragment)
Uncharacterized protein

Uncharacterized protein
Uncharacterized protein
Uncharacterized protein
Uncharacterized protein
Uncharacterized protein
Uncharacterized protein
Uncharacterized protein
Uncharacterized protein
Uncharacterized protein
Uncharacterized protein
Uncharacterized protein
Uncharacterized protein
Uncharacterized protein
Uncharacterized protein
Uncharacterized protein
Uncharacterized protein
Uncharacterized protein
Uncharacterized protein (Fragment)
Uncharacterized protein
Uncharacterized protein
Uncharacterized protein
Uncharacterized protein
Uncharacterized protein
Uncharacterized protein
Uncharacterized protein
Uncharacterized protein
Uncharacterized protein
Uncharacterized protein
Uncharacterized protein
Uncharacterized protein
Uncharacterized protein
Uncharacterized protein
Uncharacterized protein
Uncharacterized protein
Uncharacterized protein
Uncharacterized protein
S-locus-related I (Fragment)
S-locus receptor kinase (Fragment)
Uncharacterized protein
S-locus receptor kinase (Fragment)
Uncharacterized protein

BrSLGf2b protein
S receptor kinase
Uncharacterized protein
SRK like receptor kinase
S glycoprotein (Fragment)
Uncharacterized protein
S-locus glycoprotein (Fragment)
Uncharacterized protein
Uncharacterized protein
S glycoprotein (Fragment)
Uncharacterized protein
Uncharacterized protein
Uncharacterized protein
Uncharacterized protein
Uncharacterized protein
Uncharacterized protein (Fragment)
Uncharacterized protein
Uncharacterized protein
Uncharacterized protein
Uncharacterized protein
S receptor kinase 44 (Fragment)
Uncharacterized protein
Receptor kinase 5
Uncharacterized protein
Uncharacterized protein
Uncharacterized protein
S-locus receptor kinase (Fragment)
S-locus glycoprotein (Fragment)
Receptor kinase 6
Uncharacterized protein
Uncharacterized protein
S locus glycoprotein (Fragment)
S glycoprotein (Fragment)
S glycoprotein (Fragment)
S-locus glycoprotein (Fragment)
Uncharacterized protein
S-receptor kinase SRK9 (Fragment)
Uncharacterized protein
Uncharacterized protein
Uncharacterized protein
Uncharacterized protein

Uncharacterized protein
Uncharacterized protein
S-locus glycoprotein
S-locus receptor kinase
Uncharacterized protein
S receptor kinase-54
Uncharacterized protein
Uncharacterized protein
SLG8
Uncharacterized protein
Uncharacterized protein
S-locus receptor kinase (Fragment)
Uncharacterized protein
S glycoprotein (Fragment)
S-locus glycoprotein (Fragment)
Receptor protein kinase SRK8
Uncharacterized protein
Uncharacterized protein
B.campestris S-locus-specific glycoprotein (Fragment)
S receptor kinase (Fragment)
S-locus receptor kinase (Fragment)
Uncharacterized protein
S glycoprotein (Fragment)
Uncharacterized protein
S-receptor kinase
S-locus receptor kinase (Fragment)
Uncharacterized protein
S glycoprotein (Fragment)
Uncharacterized protein
S receptor kinase
Uncharacterized protein
Uncharacterized protein
Uncharacterized protein
S glycoprotein (Fragment)
Uncharacterized protein
S glycoprotein (Fragment)
NS1 glycoprotein (Fragment)
S glycoprotein (Fragment)
S-locus receptor kinase (Fragment)
Uncharacterized protein
S-locus receptor kinase (Fragment)

S locus glycoprotein
Uncharacterized protein
Uncharacterized protein
S-locus receptor kinase (Fragment)
Uncharacterized protein
Uncharacterized protein
S receptor kinase 40 (Fragment)
S locus protein 4 (Fragment)
Uncharacterized protein
S-locus receptor kinase (Fragment)
SRK like receptor kinase
Uncharacterized protein
Uncharacterized protein
SRK like receptor kinase
S glycoprotein (Fragment)
S glycoprotein (Fragment)
Uncharacterized protein
Uncharacterized protein
Receptor protein kinase SRK12
Uncharacterized protein
Uncharacterized protein
Uncharacterized protein
S-locus receptor kinase (Fragment)
S-locus receptor kinase (Fragment)
S-glycoprotein
Uncharacterized protein
S-locus receptor kinase (Fragment)
Uncharacterized protein
S receptor kinase 47 (Fragment)
S-locus receptor kinase (Fragment)
S locus protein 6
Uncharacterized protein
S locus receptor kinase (Fragment)
S-locus related 2-9B (Fragment)
Uncharacterized protein
Uncharacterized protein
S-locus related 2-9A (Fragment)
Uncharacterized protein
Uncharacterized protein
Uncharacterized protein
S locus glycoprotein-54

Uncharacterized protein
Uncharacterized protein (Fragment)
S-locus receptor kinase (Fragment)
Uncharacterized protein
S-locus glycoprotein (Fragment)
Uncharacterized protein
Uncharacterized protein
Uncharacterized protein
Uncharacterized protein
S receptor kinase (Fragment)
S-locus glycoprotein (Fragment)
Uncharacterized protein
S-locus receptor kinase
Uncharacterized protein
Uncharacterized protein
Uncharacterized protein
S-locus glycoprotein (Fragment)
SRK like receptor kinase
S locus protein 1 (Fragment)
Uncharacterized protein
S locus receptor kinase
SRK like receptor kinase
Uncharacterized protein
SRK like receptor kinase
S-locus receptor kinase (Fragment)
Uncharacterized protein
Uncharacterized protein
Uncharacterized protein
S-locus glycoprotein (Fragment)
BrSRK-f2 protein
Uncharacterized protein
S-receptor kinase
NS3-glycoprotein
Uncharacterized protein
Uncharacterized protein
S receptor kinase
SRK29
Uncharacterized protein
Uncharacterized protein
Uncharacterized protein
Uncharacterized protein

Uncharacterized protein
S locus protein 2 (Fragment)
Uncharacterized protein
Uncharacterized protein
S glycoprotein (Fragment)
Uncharacterized protein
Uncharacterized protein
Uncharacterized protein
SRK like receptor kinase
SRK45
SRK like receptor kinase
Uncharacterized protein
Glycoprotein
Uncharacterized protein (Fragment)
Uncharacterized protein
S glycoprotein (Fragment)
S locus glycoprotein
Uncharacterized protein
Uncharacterized protein
S-locus receptor kinase (Fragment)
Uncharacterized protein
S glycoprotein (Fragment)
Uncharacterized protein
Uncharacterized protein
Uncharacterized protein
Uncharacterized protein
Uncharacterized protein
Uncharacterized protein
Uncharacterized protein
Uncharacterized protein
Uncharacterized protein
Uncharacterized protein
BrSLGf2a protein
Uncharacterized protein (Fragment)
Uncharacterized protein
S-locus receptor kinase (Fragment)
Uncharacterized protein
Uncharacterized protein
S-locus receptor kinase (Fragment)
SLG45
Uncharacterized protein

S-receptor kinase
SLG29
SLG12
Uncharacterized protein
Uncharacterized protein
Uncharacterized protein
S-locus glycoprotein (Fragment)
Receptor kinase 1
Receptor kinase 3
S locus glycoprotein (Fragment)
Uncharacterized protein
Uncharacterized protein
Uncharacterized protein
SRK like receptor kinase
Uncharacterized protein
S locus protein 3 (Fragment)
Glycoprotein
S receptor kinase (Fragment)
Uncharacterized protein
Uncharacterized protein
S locus glycoprotein
S-locus glycoprotein of Brassica campestris S9-homozygote
S receptor kinase (Fragment)
S-locus receptor kinase
Uncharacterized protein
S-locus glycoprotein
Uncharacterized protein
Uncharacterized protein
Uncharacterized protein
Uncharacterized protein
Uncharacterized protein
Uncharacterized protein (Fragment)
Uncharacterized protein
Uncharacterized protein
Uncharacterized protein
Uncharacterized protein
Uncharacterized protein
Uncharacterized protein
Uncharacterized protein
Uncharacterized protein (Fragment)
Uncharacterized protein

Uncharacterized protein
Uncharacterized protein
Uncharacterized protein (Fragment)
Uncharacterized protein
Uncharacterized protein
Uncharacterized protein
Uncharacterized protein
Uncharacterized protein (Fragment)
Uncharacterized protein
Uncharacterized protein (Fragment)
Uncharacterized protein (Fragment)
Uncharacterized protein
Uncharacterized protein (Fragment)
Uncharacterized protein
Uncharacterized protein
Uncharacterized protein (Fragment)
Uncharacterized protein
S-locus-related I (Fragment)
S-locus related 1 (Fragment)
S-locus-related I (Fragment)
S-locus receptor kinase
S-locus glycoprotein
S-locus glycoprotein
BnaC05g08640D protein
BnaA09g51740D protein
S-receptor kinase
BnaC08g14740D protein
BnaAnng30410D protein
BnaA08g12250D protein
BnaA08g25350D protein
BnaC05g08740D protein
BnaCnng56910D protein
BnaA09g47650D protein
S-locus glycoprotein type II (Fragment)
BnaC02g16590D protein
BnaA03g19850D protein
S-locus receptor kinase
BnaA09g47640D protein
S-locus receptor kinase
BnaC08g42100D protein
BnaC06g27660D protein

BnaC08g14760D protein
BnaA04g10940D protein (Fragment)
BnaC01g43350D protein
BnaCnng09910D protein
BnaC04g33120D protein
BnaC06g27240D protein
BnaA07g25480D protein
S-locus glycoprotein (Fragment)
BnaA06g12270D protein
S-locus glycoprotein (Fragment)
SLR1 glycoprotein
BnaA03g60150D protein
BnaAnng33740D protein
BnaC08g42090D protein
BnaCnng40530D protein
BnaC06g27230D protein
BnaC01g41150D protein
BnaA01g22600D protein
BnaA09g46530D protein
BnaA04g14930D protein
BnaA09g47660D protein
BnaA08g25330D protein
BnaC08g42110D protein
BnaA07g25460D protein
BnaA01g22610D protein
BnaC08g14850D protein
BnaA03g60620D protein
BnaA03g57950D protein
BnaC06g27650D protein
S receptor kinase (Fragment)
BnaC01g19470D protein
BnaC03g72660D protein
BnaA10g07090D protein
BnaA09g40890D protein
BnaCnng04390D protein
S-locus glycoprotein
BnaA07g38400D protein
BnaA01g22750D protein
BnaC02g44050D protein
BnaA09g47440D protein
BnaA03g58780D protein

BnaC02g34450D protein
BnaA09g13770D protein
S-locus glycoprotein (Fragment)
BnaCnng38470D protein
BnaC01g19460D protein
BnaA01g22760D protein
S-locus glycoprotein
BnaA09g13780D protein
BnaCnng09940D protein
BnaC05g08660D protein
BnaC01g41140D protein
BnaCnng04150D protein
BnaA01g22660D protein
BnaA09g13740D protein
BnaCnng04180D protein
BnaAnng22710D protein
BnaC05g08760D protein
BnaC07g32470D protein
BnaC08g14920D protein
BnaA08g25370D protein
BnaCnng69790D protein
BnaC04g54970D protein
BnaA04g29170D protein
BnaAnng35500D protein (Fragment)
BnaC05g08670D protein
BnaA09g05490D protein
BnaC04g13630D protein
BnaC01g43340D protein
BnaCnng04170D protein
S-locus glycoprotein (Fragment)
BnaA08g25300D protein
BnaC04g47960D protein
BnaA09g47630D protein
BnaC05g37330D protein
BnaC08g42120D protein
BnaCnng21290D protein
S-locus glycoprotein (Fragment)
BnaCnng38150D protein
BnaC06g27220D protein
BnaA09g13720D protein
BnaA04g05920D protein

BnaCnng55880D protein
BnaA07g25440D protein
Serine/threonine kinase receptor
BnaC05g08650D protein
BnaC01g44280D protein
BnaA03g60570D protein
S-locus receptor kinase (Fragment)
BnaC05g23360D protein
BnaCnng02640D protein
S receptor kinase
S-locus glycoprotein (Fragment)
BnaC06g27610D protein
S-locus glycoprotein (Fragment)
Glycoprotein
S-locus glycoprotein (Fragment)
S-locus glycoprotein (Fragment)
Glycoprotein
S-locus-related I (Fragment)
S-locus-specific glycoprotein BS29-1
S-locus-specific glycoprotein BS29-2
S-locus-specific glycoprotein
Uncharacterized protein
Serine/threonine-protein kinase
Serine/threonine-protein kinase
Serine/threonine-protein kinase
Uncharacterized protein
Serine/threonine-protein kinase
Uncharacterized protein
Serine/threonine-protein kinase
Uncharacterized protein
Uncharacterized protein
Uncharacterized protein
Uncharacterized protein
Serine/threonine-protein kinase
Serine/threonine-protein kinase
Serine/threonine-protein kinase
Uncharacterized protein
Uncharacterized protein
Uncharacterized protein
Uncharacterized protein
Serine/threonine-protein kinase

Uncharacterized protein
Uncharacterized protein
Uncharacterized protein
Uncharacterized protein
Uncharacterized protein
Uncharacterized protein
Uncharacterized protein
Serine/threonine-protein kinase
Uncharacterized protein
Uncharacterized protein
Uncharacterized protein
Serine/threonine-protein kinase
Serine/threonine-protein kinase
Uncharacterized protein
Uncharacterized protein
Uncharacterized protein
Uncharacterized protein
Uncharacterized protein
Serine/threonine-protein kinase
Serine/threonine-protein kinase
Uncharacterized protein
Uncharacterized protein
Uncharacterized protein
Serine/threonine-protein kinase
Uncharacterized protein
Uncharacterized protein
Serine/threonine-protein kinase
Uncharacterized protein
Serine/threonine-protein kinase
Uncharacterized protein
Uncharacterized protein
Uncharacterized protein
Serine/threonine-protein kinase
Uncharacterized protein
Uncharacterized protein
Uncharacterized protein
Serine/threonine-protein kinase
Uncharacterized protein
Uncharacterized protein
Serine/threonine-protein kinase
Uncharacterized protein

Uncharacterized protein
Serine/threonine-protein kinase
Uncharacterized protein
Serine/threonine-protein kinase
Serine/threonine-protein kinase
Uncharacterized protein
Uncharacterized protein
Serine/threonine-protein kinase
Serine/threonine-protein kinase
Serine/threonine-protein kinase
Serine/threonine-protein kinase
Serine/threonine-protein kinase
Uncharacterized protein
Uncharacterized protein
Uncharacterized protein
Uncharacterized protein
Uncharacterized protein
Uncharacterized protein
Serine/threonine-protein kinase
Uncharacterized protein
Uncharacterized protein
S13-b receptor kinase
Putative serine/threonine-protein kinase receptor
S-locus-specific glycoprotein S13
S-locus-specific glycoprotein S14 (Fragment)
S-locus-specific glycoprotein S6
SRK13-b
SLGB protein (Fragment)
SFR1 protein
SLGB protein (Fragment)
SLGA protein (Fragment)
SLGA protein (Fragment)
Uncharacterized protein
Serine /threonine kinase
S60 S-locus receptor kinase
S glycoprotein (Fragment)
S-locus glycoprotein (Fragment)
S receptor kinase (Fragment)
Uncharacterized protein
S-locus receptor kinase (Fragment)
S-locus receptor kinase

Uncharacterized protein
Uncharacterized protein
SLG2-b
S-locus glycoprotein (Fragment)
Srk29 protein
S-locus glycoprotein (Fragment)
SRK protein (Fragment)
Uncharacterized protein (Fragment)
S-locus receptor kinase (Fragment)
Uncharacterized protein
Uncharacterized protein
Uncharacterized protein
Uncharacterized protein
Uncharacterized protein
SLG23Bol
S-locus glycoprotein (Fragment)
SRK5 protein (Fragment)
Uncharacterized protein
S-locus receptor kinase (Fragment)
S-locus receptor kinase (Fragment)
Uncharacterized protein (Fragment)
Uncharacterized protein
SLR2 protein (Fragment)
Uncharacterized protein
Uncharacterized protein
S glycoprotein (Fragment)
Uncharacterized protein
SFR3 protein
SRK23Bol (Fragment)
Uncharacterized protein (Fragment)
SLGB protein
S-locus receptor kinase (Fragment)
SRK protein (Fragment)
S-locus receptor kinase (Fragment)
Uncharacterized protein
SLR3-1 protein
Uncharacterized protein
S glycoprotein (Fragment)
Uncharacterized protein
Uncharacterized protein
Uncharacterized protein

Uncharacterized protein
Uncharacterized protein
Uncharacterized protein
S-locus receptor kinase (Fragment)
SRK protein (Fragment)
S glycoprotein (Fragment)
S glycoprotein (Fragment)
S-locus receptor kinase (Fragment)
S-locus receptor kinase (Fragment)
Uncharacterized protein
Uncharacterized protein
Uncharacterized protein
Uncharacterized protein
S-locus receptor kinase (Fragment)
S glycoprotein (Fragment)
S-locus receptor kinase (Fragment)
Uncharacterized protein
Uncharacterized protein
S-locus receptor kinase
Uncharacterized protein
S18 S-locus receptor kinase
S-locus receptor kinase (Fragment)
S glycoprotein (Fragment)
Serine/threonine kinase
SLG13-b (Fragment)
SRK protein (Fragment)
S glycoprotein (Fragment)
Uncharacterized protein
S-locus receptor kinase (Fragment)
Uncharacterized protein
Uncharacterized protein
S receptor kinase 32 (Fragment)
Uncharacterized protein
Uncharacterized protein
Uncharacterized protein
Uncharacterized protein
BoSRK-28 protein (Fragment)
SLG-Sc protein (Fragment)
SLG13 (Fragment)
S glycoprotein (Fragment)
S-locus receptor kinase (Fragment)

SRK13
S-locus receptor kinase (Fragment)
S-locus receptor kinase (Fragment)
S-receptor kinase-like protein
S-locus glycoprotein (Fragment)
S-locus glycoprotein (Fragment)
S blycoprotein (Fragment)
Uncharacterized protein (Fragment)
Uncharacterized protein
S-locus glycoprotein (Fragment)
S glycoprotein (Fragment)
Uncharacterized protein
S-locus glycoprotein (Fragment)
S glycoprotein (Fragment)
S-locus glycoprotein (Fragment)
S locus glycoprotein
Uncharacterized protein
S-locus receptor kinase (Fragment)
Uncharacterized protein
Uncharacterized protein
Uncharacterized protein
S-locus receptor kinase
Uncharacterized protein
Uncharacterized protein
S-locus glycoprotein (Fragment)
Uncharacterized protein
Receptor-like kinase
Uncharacterized protein
S glycoprotein (Fragment)
SRK2-b
SFR2 protein (Fragment)
S-locus receptor kinase (Fragment)
S-locus receptor kinase (Fragment)
SLG glycoprotein (Fragment)
S-locus glycoprotein (Fragment)
S-locus receptor kinase (Fragment)
S glycoprotein (Fragment)
S receptor kinase 12 (Fragment)
S glycoprotein (Fragment)
SLR3-2 protein
Uncharacterized protein

Uncharacterized protein
S glycoprotein (Fragment)
Uncharacterized protein
S-locus receptor kinase (Fragment)
Uncharacterized protein
S glycoprotein (Fragment)
Self-incompatibility locus specific glycoprotein
S-locus receptor kinase (Fragment)
S-locus receptor kinase (Fragment)
Uncharacterized protein
Uncharacterized protein
SRK protein (Fragment)
Uncharacterized protein
SRK3 gene
S-locus receptor kinase (Fragment)
S-locus receptor kinase (Fragment)
Uncharacterized protein
S glycoprotein (Fragment)
SRK15 protein (Fragment)
S-locus glycoprotein
S-locus-specific glycoprotein homolog (Fragment)
Uncharacterized protein
Uncharacterized protein (Fragment)
Uncharacterized protein (Fragment)
S-locus receptor kinase (Fragment)
S glycoprotein (Fragment)
S-locus receptor kinase (Fragment)
SRK protein (Fragment)
S-locus receptor kinase (Fragment)
S-locus glycoprotein (Fragment)
Uncharacterized protein
Uncharacterized protein
Uncharacterized protein
S-locus-related I (Fragment)
Uncharacterized protein
Uncharacterized protein
Uncharacterized protein
Uncharacterized protein
Uncharacterized protein
Uncharacterized protein
Uncharacterized protein

Uncharacterized protein
Uncharacterized protein
Uncharacterized protein
Uncharacterized protein
Uncharacterized protein
Uncharacterized protein
Uncharacterized protein
Uncharacterized protein
Uncharacterized protein
Uncharacterized protein
Uncharacterized protein
Uncharacterized protein
Uncharacterized protein
Uncharacterized protein
Uncharacterized protein
Uncharacterized protein
Uncharacterized protein
Uncharacterized protein
Uncharacterized protein
Uncharacterized protein
Uncharacterized protein
Uncharacterized protein
Uncharacterized protein
Uncharacterized protein
Uncharacterized protein
Uncharacterized protein
Uncharacterized protein
Uncharacterized protein
Uncharacterized protein
Uncharacterized protein
Uncharacterized protein
Uncharacterized protein
Uncharacterized protein
Uncharacterized protein
Uncharacterized protein
Uncharacterized protein
Uncharacterized protein
Uncharacterized protein
Uncharacterized protein
Uncharacterized protein
Uncharacterized protein

Uncharacterized protein
Uncharacterized protein
Uncharacterized protein
Uncharacterized protein
Uncharacterized protein
Uncharacterized protein
Uncharacterized protein
Uncharacterized protein
Uncharacterized protein
Uncharacterized protein
Uncharacterized protein
Uncharacterized protein
Uncharacterized protein
Uncharacterized protein
Uncharacterized protein
Uncharacterized protein
Uncharacterized protein
Uncharacterized protein
Uncharacterized protein
Uncharacterized protein
Uncharacterized protein
Uncharacterized protein
S-locus-related I (Fragment)
S-locus-related I (Fragment)
S-locus-related I (Fragment)
Uncharacterized protein
Uncharacterized protein
Uncharacterized protein
Uncharacterized protein
Bm2565
Bm6023
BMA-NOAH-1, isoform c
BMA-NOAH-2
BMA-NOAH-1, isoform a
Bm6373
Bm4198
Bm1777
BMA-CUTL-28
BMA-CUTL-17
Bm10928
Bm2629, isoform b

Cation-transporting ATPase
Bm4699, isoform f
Bm2255
Bm10200
Bm4699, isoform f
Uncharacterized protein
Bm4699, isoform c
Bm4248, isoform b
BMA-SRAP-1
Bm2491
Uncharacterized protein
Cation-transporting ATPase
Bm4248, isoform b
Bm2255
Bm2255
Bm7942
Uncharacterized protein
Bm4248, isoform a
Bm2491 (Fragment)
Uncharacterized protein
Uncharacterized protein
BMA-SRAP-1
Bm4248, isoform b
Bm7942 (Fragment)
Uncharacterized protein
Uncharacterized protein
Uncharacterized protein
Uncharacterized protein
Uncharacterized protein
Uncharacterized protein
Uncharacterized protein
Uncharacterized protein
Uncharacterized protein
Uncharacterized protein
Uncharacterized protein
Uncharacterized protein
Uncharacterized protein
Uncharacterized protein
Uncharacterized protein
Uncharacterized protein
Uncharacterized protein

Uncharacterized protein
Uncharacterized protein
Uncharacterized protein
Uncharacterized protein
Uncharacterized protein
Uncharacterized protein
Uncharacterized protein
Uncharacterized protein
Uncharacterized protein
Uncharacterized protein
Uncharacterized protein
Uncharacterized protein
Uncharacterized protein
Uncharacterized protein
Uncharacterized protein
Uncharacterized protein
Uncharacterized protein
Uncharacterized protein
Plasma kallikrein (Fragment)
Plasminogen (Fragment)
Hepatocyte growth factor (Fragment)
Plasminogen (Fragment)
Uncharacterized protein
Uncharacterized protein
Uncharacterized protein
Uncharacterized protein
Uncharacterized protein
Uncharacterized protein
Uncharacterized protein
Uncharacterized protein
Uncharacterized protein
Uncharacterized protein
Uncharacterized protein
Uncharacterized protein
Uncharacterized protein
Uncharacterized protein
Uncharacterized protein
Uncharacterized protein
Uncharacterized protein
Uncharacterized protein
Uncharacterized protein

Uncharacterized protein
Uncharacterized protein
Uncharacterized protein
Uncharacterized protein
Uncharacterized protein
Uncharacterized protein
Uncharacterized protein
Uncharacterized protein
Uncharacterized protein
Uncharacterized protein
Uncharacterized protein
Uncharacterized protein
Glycosyltransferase family 31 protein
Copper radical oxidase
Uncharacterized protein
Uncharacterized protein
Uncharacterized protein
Uncharacterized protein
Uncharacterized protein
CBN-CUTL-18 protein
Uncharacterized protein
Uncharacterized protein
Uncharacterized protein
Uncharacterized protein
Uncharacterized protein
Uncharacterized protein
Uncharacterized protein
Uncharacterized protein
Uncharacterized protein
Uncharacterized protein
Uncharacterized protein
Uncharacterized protein
Uncharacterized protein
Uncharacterized protein
Uncharacterized protein
Uncharacterized protein
CBN-CUTL-17 protein
Uncharacterized protein
Uncharacterized protein
Uncharacterized protein
Uncharacterized protein

Protein CBG17597
Protein CBG15350
Protein CBR-NOAH-2
Protein CBG09784
Protein CBG14590
Protein CBG00402
Protein CBG21204
Protein CBG02948
Protein CBG22791
Protein CBR-TAG-303
Protein CBG21281
Protein CBG00939
Protein CBG11308
Protein CBG14589
Protein CBG21662
Protein CBR-LET-653
Protein CBG17333
Protein CBR-NOAH-1
Protein CBR-TRY-2
Protein CBG06184
Protein CBG04379
Protein CBG23169
Protein let-653
Uncharacterized protein F52C9.5
Uncharacterized protein C34C12.7
Uncharacterized protein B0361.9
HGF/MSP/plasminogen-like protein
Uncharacterized protein
NOmpA Homolog (Drosophila nompA: no mechanoreceptor potential A)
Uncharacterized protein
Uncharacterized protein
Uncharacterized protein
Uncharacterized protein
Uncharacterized protein
Uncharacterized protein
Uncharacterized protein
Uncharacterized protein
Uncharacterized protein
Uncharacterized protein
CUTiclin-Like
NOmpA Homolog (Drosophila nompA: no mechanoreceptor potential A)

Uncharacterized protein
Uncharacterized protein
Uncharacterized protein
Uncharacterized protein
Uncharacterized protein
NOmpA Homolog (Drosophila nompA: no mechanoreceptor potential A)
CUTiclin-Like
CUTiclin-Like
Serine Rich Adhesion Protein-like
Serine Rich Adhesion Protein-like
CUTiclin-Like
Uncharacterized protein
Uncharacterized protein
Uncharacterized protein
Serine Rich Adhesion Protein-like
NOmpA Homolog (Drosophila nompA: no mechanoreceptor potential A)
Uncharacterized protein
Uncharacterized protein
Uncharacterized protein
Uncharacterized protein
Uncharacterized protein
Uncharacterized protein
Uncharacterized protein
Uncharacterized protein
Uncharacterized protein
Uncharacterized protein
Uncharacterized protein
Uncharacterized protein
Uncharacterized protein
Uncharacterized protein
Uncharacterized protein
Uncharacterized protein
Uncharacterized protein
Uncharacterized protein
Uncharacterized protein
Uncharacterized protein
Uncharacterized protein
Uncharacterized protein
Uncharacterized protein
Uncharacterized protein
Uncharacterized protein

Uncharacterized protein
Uncharacterized protein
Uncharacterized protein
Uncharacterized protein
Uncharacterized protein
Uncharacterized protein
Uncharacterized protein
Uncharacterized protein
Uncharacterized protein
Uncharacterized protein
Uncharacterized protein
Uncharacterized protein
Uncharacterized protein
Uncharacterized protein
Uncharacterized protein
Uncharacterized protein
Uncharacterized protein
Uncharacterized protein (Fragment)
Uncharacterized protein (Fragment)
Uncharacterized protein (Fragment)
Uncharacterized protein (Fragment)
Uncharacterized protein (Fragment)
Uncharacterized protein (Fragment)
Uncharacterized protein (Fragment)
Uncharacterized protein (Fragment)
Uncharacterized protein (Fragment)
Uncharacterized protein (Fragment)
Uncharacterized protein (Fragment)
Uncharacterized protein (Fragment)
Uncharacterized protein (Fragment)
Uncharacterized protein (Fragment)
Uncharacterized protein (Fragment)
Uncharacterized protein (Fragment)
Uncharacterized protein (Fragment)
Uncharacterized protein (Fragment)
Uncharacterized protein (Fragment)
Uncharacterized protein (Fragment)
Uncharacterized protein (Fragment)
Uncharacterized protein (Fragment)
Uncharacterized protein (Fragment)
Uncharacterized protein (Fragment)

Uncharacterized protein
Uncharacterized protein
Uncharacterized protein
Uncharacterized protein
Uncharacterized protein
Uncharacterized protein
Uncharacterized protein
Uncharacterized protein
Uncharacterized protein
Uncharacterized protein
Uncharacterized protein
Uncharacterized protein
Uncharacterized protein
Uncharacterized protein
Uncharacterized protein
Uncharacterized protein
Uncharacterized protein
Uncharacterized protein
Uncharacterized protein
Uncharacterized protein
Uncharacterized protein
Uncharacterized protein
Uncharacterized protein
Uncharacterized protein
Uncharacterized protein
Uncharacterized protein
Uncharacterized protein
Uncharacterized protein
Uncharacterized protein
Uncharacterized protein
Uncharacterized protein
Uncharacterized protein
Uncharacterized protein
Uncharacterized protein
Uncharacterized protein
Uncharacterized protein
Uncharacterized protein
Uncharacterized protein
Uncharacterized protein
Uncharacterized protein
Uncharacterized protein

Uncharacterized protein
Uncharacterized protein
Uncharacterized protein
Uncharacterized protein
Uncharacterized protein
Uncharacterized protein
Uncharacterized protein
Uncharacterized protein
Uncharacterized protein
Uncharacterized protein
Uncharacterized protein
Uncharacterized protein
Uncharacterized protein
Uncharacterized protein
Uncharacterized protein
CRE-NOAH-2 protein
Uncharacterized protein
Uncharacterized protein
Uncharacterized protein
Uncharacterized protein
Uncharacterized protein
Uncharacterized protein
Uncharacterized protein (Fragment)
Uncharacterized protein (Fragment)
Uncharacterized protein
Uncharacterized protein (Fragment)
Uncharacterized protein (Fragment)
Uncharacterized protein
Uncharacterized protein
Uncharacterized protein (Fragment)
Uncharacterized protein (Fragment)
Uncharacterized protein
Uncharacterized protein
Uncharacterized protein
Uncharacterized protein (Fragment)
Uncharacterized protein (Fragment)
Uncharacterized protein
Uncharacterized protein (Fragment)
Uncharacterized protein (Fragment)
Uncharacterized protein
Uncharacterized protein

Uncharacterized protein (Fragment)
Uncharacterized protein
CRE-CUTL-27 protein
Uncharacterized protein
Uncharacterized protein
Uncharacterized protein (Fragment)
Uncharacterized protein (Fragment)
Uncharacterized protein (Fragment)
Uncharacterized protein (Fragment)
Uncharacterized protein
Uncharacterized protein
Uncharacterized protein
Uncharacterized protein
Uncharacterized protein (Fragment)
Uncharacterized protein
Uncharacterized protein (Fragment)
Uncharacterized protein
Uncharacterized protein (Fragment)
CRE-NOAH-1 protein
Uncharacterized protein
Uncharacterized protein
CRE-CUTL-18 protein
CRE-LET-653 protein
Uncharacterized protein
Uncharacterized protein
Uncharacterized protein
Uncharacterized protein
Uncharacterized protein (Fragment)
Uncharacterized protein (Fragment)
Uncharacterized protein (Fragment)
Uncharacterized protein
Uncharacterized protein (Fragment)
Uncharacterized protein
Uncharacterized protein
Uncharacterized protein (Fragment)
Uncharacterized protein
Uncharacterized protein
Uncharacterized protein (Fragment)
Uncharacterized protein (Fragment)
Uncharacterized protein
Uncharacterized protein (Fragment)

Uncharacterized protein (Fragment)
Uncharacterized protein (Fragment)
Uncharacterized protein (Fragment)
Uncharacterized protein (Fragment)
Uncharacterized protein
Uncharacterized protein
Uncharacterized protein
Uncharacterized protein
Uncharacterized protein
Uncharacterized protein
Uncharacterized protein
Uncharacterized protein
Uncharacterized protein
Uncharacterized protein
Uncharacterized protein
Uncharacterized protein
Uncharacterized protein
Uncharacterized protein
Uncharacterized protein
Uncharacterized protein
Uncharacterized protein
Uncharacterized protein
Uncharacterized protein
Uncharacterized protein
Uncharacterized protein
Uncharacterized protein
Uncharacterized protein
Uncharacterized protein
Uncharacterized protein
Uncharacterized protein
Uncharacterized protein
Uncharacterized protein
Uncharacterized protein
Uncharacterized protein
Uncharacterized protein
Uncharacterized protein
Uncharacterized protein
Putative serine/threonine-protein kinase receptor
Putative receptor protein kinase ZmPK1
Putative receptor protein kinase ZmPK1
Putative serine/threonine-protein kinase receptor

Putative serine/threonine-protein kinase receptor
Putative serine/threonine-protein kinase receptor
Putative serine/threonine-protein kinase receptor
Cysteine-rich receptor-like protein kinase 10
Putative cysteine-rich receptor-like protein kinase 20
Cysteine-rich receptor-like protein kinase 10
Putative serine/threonine-protein kinase receptor
PAN domain-containing protein At5g03700 family
Putative serine/threonine-protein kinase receptor
Putative serine/threonine-protein kinase receptor
Putative serine/threonine-protein kinase receptor (Fragment)
Putative receptor protein kinase ZmPK1
Putative serine/threonine-protein kinase receptor
Putative serine/threonine-protein kinase receptor (Fragment)
Putative serine/threonine-protein kinase receptor
S-locus-specific glycoprotein S13
Putative serine/threonine-protein kinase receptor
Putative receptor protein kinase ZmPK1
Putative serine/threonine-protein kinase receptor
Putative serine/threonine-protein kinase receptor
Putative serine/threonine-protein kinase receptor
Putative serine/threonine-protein kinase receptor
Putative serine/threonine-protein kinase receptor
Putative serine/threonine-protein kinase receptor
Putative receptor protein kinase ZmPK1
Putative serine/threonine-protein kinase receptor
Putative serine/threonine-protein kinase receptor
S-locus-specific glycoprotein S6
Putative serine/threonine-protein kinase receptor
Putative serine/threonine-protein kinase receptor
Uncharacterized protein (Fragment)
Putative serine/threonine-protein kinase receptor (Fragment)
S-locus-specific glycoprotein BS29-2 (Fragment)
Cysteine-rich receptor-like protein kinase 10
Acidic endochitinase
Putative serine/threonine-protein kinase receptor
Putative serine/threonine-protein kinase receptor
Putative serine/threonine-protein kinase receptor
Putative receptor protein kinase ZmPK1
Putative serine/threonine-protein kinase receptor
Putative serine/threonine-protein kinase receptor

Cysteine-rich receptor-like protein kinase 10
Putative serine/threonine-protein kinase receptor
Putative serine/threonine-protein kinase receptor
Putative serine/threonine-protein kinase receptor
Cysteine-rich receptor-like protein kinase 6
Putative serine/threonine-protein kinase receptor
Putative serine/threonine-protein kinase receptor
Putative serine/threonine-protein kinase receptor
Epidermis-specific secreted glycoprotein EP1
Putative receptor protein kinase ZmPK1
Putative serine/threonine-protein kinase receptor
Putative serine/threonine-protein kinase receptor
Putative serine/threonine-protein kinase receptor
Cysteine-rich receptor-like protein kinase 10
Putative serine/threonine-protein kinase receptor
Putative serine/threonine-protein kinase receptor
Cysteine-rich receptor-like protein kinase 6
Putative serine/threonine-protein kinase receptor
Putative serine/threonine-protein kinase receptor
Putative serine/threonine-protein kinase receptor
Putative LRR receptor-like serine/threonine-protein kinase At1g56140 family
Putative cysteine-rich receptor-like protein kinase 12
Putative serine/threonine-protein kinase receptor
Putative serine/threonine-protein kinase receptor
Cysteine-rich receptor-like protein kinase 10
Putative receptor protein kinase ZmPK1
Putative serine/threonine-protein kinase receptor
Putative serine/threonine-protein kinase receptor
Putative serine/threonine-protein kinase receptor
Uncharacterized protein
Uncharacterized protein
Uncharacterized protein
Kallikrein B1
Uncharacterized protein
Hepatocyte growth factor
Coagulation factor XI
Hepatocyte growth factor
Coagulation factor XI
Plasminogen
Kallikrein B1
Hepatocyte growth factor

Hepatocyte growth factor
Plasminogen
Hepatocyte growth factor-like protein (Fragment)
Hepatocyte growth factor
plasma kallikrein isoform X2
Plasminogen
plasma kallikrein isoform X1
hepatocyte growth factor-like protein
Plasminogen
coagulation factor XI isoform X2
coagulation factor XI isoform X1
Hepatocyte growth factor
Hepatocyte growth factor (Fragment)
Coagulation factor XI (Fragment)
Hepatocyte growth factor (Fragment)
Macrophage stimulating 1-like protein
Uncharacterized protein (Fragment)
Uncharacterized protein
Uncharacterized protein
Uncharacterized protein
Uncharacterized protein
Neurotrypsin
Cuticlin-1
Uncharacterized protein
Heme chaperone HemW
Uncharacterized protein
Uncharacterized protein
Uncharacterized protein
Uncharacterized protein
Uncharacterized protein
Uncharacterized protein
Uncharacterized protein (Fragment)
Uncharacterized protein
Uncharacterized protein
PAN domain-containing protein
Uncharacterized protein (Fragment)
PAN domain protein
Alpha-2-macroglobulin
Uncharacterized protein
Hepatocyte growth factor
Kallikrein B1

Plasminogen
Uncharacterized protein
Prekallikrein
Factor XI
Coagulation factor XI
Uncharacterized protein
Uncharacterized protein
Uncharacterized protein
Uncharacterized protein
Uncharacterized protein
Uncharacterized protein
Uncharacterized protein
Uncharacterized protein
Uncharacterized protein
Uncharacterized protein
Uncharacterized protein
Uncharacterized protein
Uncharacterized protein
Uncharacterized protein
Uncharacterized protein
Uncharacterized protein
Uncharacterized protein
Uncharacterized protein
Uncharacterized protein
Uncharacterized protein
Uncharacterized protein (Fragment)
Uncharacterized protein
Uncharacterized protein
Uncharacterized protein
Uncharacterized protein
Uncharacterized protein
Uncharacterized protein
Uncharacterized protein
Uncharacterized protein
Uncharacterized protein
Uncharacterized protein (Fragment)
Uncharacterized protein
Uncharacterized protein
Uncharacterized protein
Uncharacterized protein
Uncharacterized protein (Fragment)
Uncharacterized protein

Uncharacterized protein (Fragment)
Uncharacterized protein
Uncharacterized protein
Uncharacterized protein
Uncharacterized protein
Uncharacterized protein
Uncharacterized protein
Uncharacterized protein
Uncharacterized protein
Uncharacterized protein
Uncharacterized protein
Uncharacterized protein
Uncharacterized protein
Uncharacterized protein
Uncharacterized protein
Uncharacterized protein
Uncharacterized protein
Uncharacterized protein
Uncharacterized protein
Uncharacterized protein
Uncharacterized protein
Uncharacterized protein
Uncharacterized protein
Uncharacterized protein
Uncharacterized protein
Uncharacterized protein (Fragment)
Uncharacterized protein
Uncharacterized protein (Fragment)
Uncharacterized protein
Uncharacterized protein
Uncharacterized protein
Uncharacterized protein
Uncharacterized protein
Uncharacterized protein
Uncharacterized protein
Uncharacterized protein
Uncharacterized protein
Uncharacterized protein
Uncharacterized protein
Uncharacterized protein
Uncharacterized protein

Uncharacterized protein
Uncharacterized protein
Uncharacterized protein
Uncharacterized protein
Uncharacterized protein
Uncharacterized protein
Uncharacterized protein
Uncharacterized protein (Fragment)
Uncharacterized protein
Uncharacterized protein
Uncharacterized protein
Uncharacterized protein
Uncharacterized protein
Uncharacterized protein
Uncharacterized protein
Uncharacterized protein
Uncharacterized protein
Uncharacterized protein
Uncharacterized protein
Uncharacterized protein
Uncharacterized protein
Uncharacterized protein
Hepatocyte growth factor
Uncharacterized protein
Uncharacterized protein
Uncharacterized protein
Hepatocyte growth factor
Uncharacterized protein
Plasminogen
Hepatocyte growth factor
Uncharacterized protein
Uncharacterized protein
S-receptor kinase (Fragment)
S-receptor kinase (Fragment)
S-receptor kinase (Fragment)
Putative S-receptor kinase (Fragment)
ARK3 (Fragment)
ARK3 (Fragment)
ARK3 (Fragment)
Putative S-receptor kinase (Fragment)
At421370-like protein

At421380-like protein
ARK3-like protein (Fragment)
AT4G21390-like protein (Fragment)
SRK (Fragment)
ARK3-like protein (Fragment)
AT4G21390-like protein (Fragment)
ARK3-like protein (Fragment)
Putative S-receptor kinase (Fragment)
AT4G21390-like protein (Fragment)
SRK (Fragment)
AT4G21390-like protein (Fragment)
ARK3-like protein (Fragment)
S-receptor kinase (Fragment)
SRK (Fragment)
At421370-like protein
AT4G27290-like protein (Fragment)
ARK3 (Fragment)
S-locus receptor kinase SRK7
At421380-like protein
S-receptor kinase (Fragment)
ARK3 (Fragment)
SRK (Fragment)
ARK3 (Fragment)
ARK3 (Fragment)
SRK (Fragment)
ARK3 (Fragment)
At421390-like protein
ARK3-like protein (Fragment)
AT4G21390-like protein (Fragment)
ARK3 (Fragment)
S-receptor kinase (Fragment)
ARK3-like protein (Fragment)
Putative S-receptor kinase (Fragment)
At421390-like protein
ARK3 (Fragment)
ARK3-like protein (Fragment)
ARK3 (Fragment)
SRK (Fragment)
ARK3 (Fragment)
SRK (Fragment)
ARK3-like protein (Fragment)

SRK (Fragment)
SRK (Fragment)
AT4G27290-like protein (Fragment)
AT4G27290-like protein (Fragment)
SRK (Fragment)
ARK3 (Fragment)
Uncharacterized protein
Uncharacterized protein
Uncharacterized protein
Uncharacterized protein
Uncharacterized protein
Uncharacterized protein
Uncharacterized protein
Uncharacterized protein
Protein DETOXIFICATION (Fragment)
Uncharacterized protein
Uncharacterized protein
Uncharacterized protein
Uncharacterized protein
Uncharacterized protein
Uncharacterized protein
Uncharacterized protein (Fragment)
Uncharacterized protein
Uncharacterized protein
Uncharacterized protein
Uncharacterized protein
Uncharacterized protein
Uncharacterized protein
Uncharacterized protein
Uncharacterized protein
Uncharacterized protein
Uncharacterized protein
Uncharacterized protein
ARK3-like protein (Fragment)
Uncharacterized protein
Uncharacterized protein
Uncharacterized protein
Uncharacterized protein
Uncharacterized protein
Uncharacterized protein
Uncharacterized protein

Uncharacterized protein
Uncharacterized protein
Uncharacterized protein
Uncharacterized protein
Uncharacterized protein (Fragment)
Uncharacterized protein (Fragment)
Uncharacterized protein
AT4G27290-like protein (Fragment)
SRK (Fragment)
ARK3 (Fragment)
AT4G21390-like protein (Fragment)
ARK3-like protein (Fragment)
G-type lectin S-receptor-like serine/threonine-protein kinase At2g19130
G-type lectin S-receptor-like serine/threonine-protein kinase At4g03230 isoform X1
G-type lectin S-receptor-like serine/threonine-protein kinase At1g11330 isoform X4
G-type lectin S-receptor-like serine/threonine-protein kinase SD3-1 isoform X1
G-type lectin S-receptor-like serine/threonine-protein kinase At1g11300 isoform X1
G-type lectin S-receptor-like serine/threonine-protein kinase At4g27290 isoform X2
G-type lectin S-receptor-like serine/threonine-protein kinase SD2-2
Uncharacterized protein
G-type lectin S-receptor-like serine/threonine-protein kinase
Uncharacterized protein
G-type lectin S-receptor-like serine/threonine-protein kinase RLK1
G-type lectin S-receptor-like serine/threonine-protein kinase At4g27290
G-type lectin S-receptor-like serine/threonine-protein kinase
G-type lectin S-receptor-like serine/threonine-protein kinase At4g27290
G-type lectin S-receptor-like serine/threonine-protein kinase At1g11300 isoform X6
Uncharacterized protein
G-type lectin S-receptor-like serine/threonine-protein kinase SD2-5
G-type lectin S-receptor-like serine/threonine-protein kinase At4g03230 isoform X2
G-type lectin S-receptor-like serine/threonine-protein kinase At4g03230 isoform X2
G-type lectin S-receptor-like serine/threonine-protein kinase At1g11300 isoform X2
G-type lectin S-receptor-like serine/threonine-protein kinase At4g27290
G-type lectin S-receptor-like serine/threonine-protein kinase
G-type lectin S-receptor-like serine/threonine-protein kinase At4g27290 isoform X1
G-type lectin S-receptor-like serine/threonine-protein kinase SD2-2
Uncharacterized protein
G-type lectin S-receptor-like serine/threonine-protein kinase
G-type lectin S-receptor-like serine/threonine-protein kinase
Uncharacterized protein
G-type lectin S-receptor-like serine/threonine-protein kinase At4g27290

G-type lectin S-receptor-like serine/threonine-protein kinase
G-type lectin S-receptor-like serine/threonine-protein kinase CES101
Uncharacterized protein
G-type lectin S-receptor-like serine/threonine-protein kinase
G-type lectin S-receptor-like serine/threonine-protein kinase
Uncharacterized protein
putative receptor protein kinase ZmPK1
Uncharacterized protein
Uncharacterized protein
G-type lectin S-receptor-like serine/threonine-protein kinase
Uncharacterized protein
G-type lectin S-receptor-like serine/threonine-protein kinase At1g61370 isoform X2
G-type lectin S-receptor-like serine/threonine-protein kinase RKS1
G-type lectin S-receptor-like serine/threonine-protein kinase
Uncharacterized protein
LOW QUALITY PROTEIN: G-type lectin S-receptor-like serine/threonine-protein kinase SD2-5
receptor-like serine/threonine-protein kinase SD1-8 isoform X1
Uncharacterized protein
Uncharacterized protein
G-type lectin S-receptor-like serine/threonine-protein kinase RLK1
G-type lectin S-receptor-like serine/threonine-protein kinase At4g27290 isoform X2
G-type lectin S-receptor-like serine/threonine-protein kinase At1g61370 isoform X2
G-type lectin S-receptor-like serine/threonine-protein kinase
G-type lectin S-receptor-like serine/threonine-protein kinase SD2-5
Uncharacterized protein
G-type lectin S-receptor-like serine/threonine-protein kinase At1g11330 isoform X5
LOW QUALITY PROTEIN: G-type lectin S-receptor-like serine/threonine-protein kinase RLK1
Uncharacterized protein
G-type lectin S-receptor-like serine/threonine-protein kinase At4g27290
G-type lectin S-receptor-like serine/threonine-protein kinase
Uncharacterized protein
Uncharacterized protein
Uncharacterized protein
G-type lectin S-receptor-like serine/threonine-protein kinase At4g03230 isoform X4
G-type lectin S-receptor-like serine/threonine-protein kinase At4g27290 isoform X1
Uncharacterized protein
G-type lectin S-receptor-like serine/threonine-protein kinase At1g11300
G-type lectin S-receptor-like serine/threonine-protein kinase At4g27290
Uncharacterized protein
Uncharacterized protein
G-type lectin S-receptor-like serine/threonine-protein kinase At4g03230

G-type lectin S-receptor-like serine/threonine-protein kinase
G-type lectin S-receptor-like serine/threonine-protein kinase At4g27290
uncharacterized protein LOC107876757
Uncharacterized protein
G-type lectin S-receptor-like serine/threonine-protein kinase At4g27290
Uncharacterized protein
Receptor-like serine/threonine-protein kinase SD1-6
G-type lectin S-receptor-like serine/threonine-protein kinase At1g11410
Uncharacterized protein
G-type lectin S-receptor-like serine/threonine-protein kinase SD2-5
G-type lectin S-receptor-like serine/threonine-protein kinase At4g27290 isoform X2
uncharacterized protein LOC107858220
G-type lectin S-receptor-like serine/threonine-protein kinase (Fragment)
G-type lectin S-receptor-like serine/threonine-protein kinase At4g27290 isoform X2
Uncharacterized protein
Uncharacterized protein
Uncharacterized protein
LOW QUALITY PROTEIN: G-type lectin S-receptor-like serine/threonine-protein kinase At1g67520
G-type lectin S-receptor-like serine/threonine-protein kinase At1g11300 isoform X3
PAN domain-containing protein At5g03700
G-type lectin S-receptor-like serine/threonine-protein kinase
putative receptor protein kinase ZmPK1
LOW QUALITY PROTEIN: uncharacterized protein LOC107876846
putative receptor protein kinase ZmPK1 isoform X1
Uncharacterized protein
G-type lectin S-receptor-like serine/threonine-protein kinase SD2-5
G-type lectin S-receptor-like serine/threonine-protein kinase At4g27290
Uncharacterized protein
Uncharacterized protein
Uncharacterized protein
Uncharacterized protein
G-type lectin S-receptor-like serine/threonine-protein kinase
G-type lectin S-receptor-like serine/threonine-protein kinase
G-type lectin S-receptor-like serine/threonine-protein kinase At4g03230 isoform X4
Uncharacterized protein (Fragment)
Uncharacterized protein
G-type lectin S-receptor-like serine/threonine-protein kinase At1g11330
Uncharacterized protein
Uncharacterized protein
G-type lectin S-receptor-like serine/threonine-protein kinase At1g61370 isoform X1
Uncharacterized protein

G-type lectin S-receptor-like serine/threonine-protein kinase At1g11330 isoform X2
G-type lectin S-receptor-like serine/threonine-protein kinase At4g03230 isoform X3
G-type lectin S-receptor-like serine/threonine-protein kinase At1g11410
Uncharacterized protein
G-type lectin S-receptor-like serine/threonine-protein kinase At4g27290 isoform X1
G-type lectin S-receptor-like serine/threonine-protein kinase At4g03230 isoform X3
G-type lectin S-receptor-like serine/threonine-protein kinase At4g27290
G-type lectin S-receptor-like serine/threonine-protein kinase
G-type lectin S-receptor-like serine/threonine-protein kinase At4g27290
putative receptor protein kinase ZmPK1 isoform X2
Uncharacterized protein
Uncharacterized protein
Uncharacterized protein
Uncharacterized protein
G-type lectin S-receptor-like serine/threonine-protein kinase At4g27290 isoform X2
G-type lectin S-receptor-like serine/threonine-protein kinase At1g11330
G-type lectin S-receptor-like serine/threonine-protein kinase At4g27290 isoform X1
Receptor-like serine/threonine-protein kinase SD1-8
G-type lectin S-receptor-like serine/threonine-protein kinase At4g03230 isoform X1
G-type lectin S-receptor-like serine/threonine-protein kinase
Uncharacterized protein
LOW QUALITY PROTEIN: uncharacterized protein LOC107844086
G-type lectin S-receptor-like serine/threonine-protein kinase
G-type lectin S-receptor-like serine/threonine-protein kinase At4g27290 isoform X1
Uncharacterized protein
G-type lectin S-receptor-like serine/threonine-protein kinase
G-type lectin S-receptor-like serine/threonine-protein kinase At4g27290
G-type lectin S-receptor-like serine/threonine-protein kinase B120
Uncharacterized protein
G-type lectin S-receptor-like serine/threonine-protein kinase At1g11330 isoform X1
G-type lectin S-receptor-like serine/threonine-protein kinase
G-type lectin S-receptor-like serine/threonine-protein kinase At4g27290
G-type lectin S-receptor-like serine/threonine-protein kinase
G-type lectin S-receptor-like serine/threonine-protein kinase At4g27290 isoform X1
Uncharacterized protein
G-type lectin S-receptor-like serine/threonine-protein kinase
G-type lectin S-receptor-like serine/threonine-protein kinase
G-type lectin S-receptor-like serine/threonine-protein kinase
Uncharacterized protein
G-type lectin S-receptor-like serine/threonine-protein kinase At4g27290
G-type lectin S-receptor-like serine/threonine-protein kinase At4g27290

Uncharacterized protein
epidermis-specific secreted glycoprotein EP1-like
G-type lectin S-receptor-like serine/threonine-protein kinase
Uncharacterized protein (Fragment)
G-type lectin S-receptor-like serine/threonine-protein kinase
Uncharacterized protein
Uncharacterized protein
G-type lectin S-receptor-like serine/threonine-protein kinase
G-type lectin S-receptor-like serine/threonine-protein kinase At4g27290
Uncharacterized protein
receptor-like serine/threonine-protein kinase SD1-8 isoform X2
G-type lectin S-receptor-like serine/threonine-protein kinase At4g27290 isoform X2
G-type lectin S-receptor-like serine/threonine-protein kinase At1g61370 isoform X1
G-type lectin S-receptor-like serine/threonine-protein kinase At1g11330
Uncharacterized protein
G-type lectin S-receptor-like serine/threonine-protein kinase At4g27290
G-type lectin S-receptor-like serine/threonine-protein kinase
Uncharacterized protein
G-type lectin S-receptor-like serine/threonine-protein kinase SD3-1 isoform X2
Uncharacterized protein
G-type lectin S-receptor-like serine/threonine-protein kinase At1g61390
Uncharacterized protein
receptor-like serine/threonine-protein kinase SD1-8 isoform X1
G-type lectin S-receptor-like serine/threonine-protein kinase
G-type lectin S-receptor-like serine/threonine-protein kinase CES101
G-type lectin S-receptor-like serine/threonine-protein kinase At1g11330
G-type lectin S-receptor-like serine/threonine-protein kinase At4g27290
G-type lectin S-receptor-like serine/threonine-protein kinase At4g27290 isoform X1
Uncharacterized protein
Uncharacterized protein
Uncharacterized protein
Uncharacterized protein
Uncharacterized protein
G-type lectin S-receptor-like serine/threonine-protein kinase
Uncharacterized protein
Uncharacterized protein
G-type lectin S-receptor-like serine/threonine-protein kinase
Uncharacterized protein
G-type lectin S-receptor-like serine/threonine-protein kinase SD1-1
G-type lectin S-receptor-like serine/threonine-protein kinase
Uncharacterized protein

G-type lectin S-receptor-like serine/threonine-protein kinase
G-type lectin S-receptor-like serine/threonine-protein kinase
G-type lectin S-receptor-like serine/threonine-protein kinase
G-type lectin S-receptor-like serine/threonine-protein kinase
Uncharacterized protein
Uncharacterized protein
PAN domain-containing protein
G-type lectin S-receptor-like serine/threonine-protein kinase
G-type lectin S-receptor-like serine/threonine-protein kinase
Uncharacterized protein
Uncharacterized protein
Uncharacterized protein
Uncharacterized protein
Uncharacterized protein
G-type lectin S-receptor-like serine/threonine-protein kinase
Uncharacterized protein
Uncharacterized protein
G-type lectin S-receptor-like serine/threonine-protein kinase
Uncharacterized protein
G-type lectin S-receptor-like serine/threonine-protein kinase
Uncharacterized protein
G-type lectin S-receptor-like serine/threonine-protein kinase
G-type lectin S-receptor-like serine/threonine-protein kinase
Uncharacterized protein
Uncharacterized protein
G-type lectin S-receptor-like serine/threonine-protein kinase
Uncharacterized protein
G-type lectin S-receptor-like serine/threonine-protein kinase
Uncharacterized protein
Uncharacterized protein
Uncharacterized protein
G-type lectin S-receptor-like serine/threonine-protein kinase SD1-1
G-type lectin S-receptor-like serine/threonine-protein kinase
Uncharacterized protein
Uncharacterized protein
Uncharacterized protein
G-type lectin S-receptor-like serine/threonine-protein kinase
G-type lectin S-receptor-like serine/threonine-protein kinase
G-type lectin S-receptor-like serine/threonine-protein kinase
Uncharacterized protein
Uncharacterized protein

G-type lectin S-receptor-like serine/threonine-protein kinase
Uncharacterized protein
Uncharacterized protein
Uncharacterized protein
G-type lectin S-receptor-like serine/threonine-protein kinase
G-type lectin S-receptor-like serine/threonine-protein kinase
Uncharacterized protein
G-type lectin S-receptor-like serine/threonine-protein kinase
Receptor-like serine/threonine-protein kinase SD1-8
Uncharacterized protein
Uncharacterized protein
G-type lectin S-receptor-like serine/threonine-protein kinase SD2-2
Uncharacterized protein
Uncharacterized protein
G-type lectin S-receptor-like serine/threonine-protein kinase
Uncharacterized protein
Uncharacterized protein
Uncharacterized protein
Uncharacterized protein
G-type lectin S-receptor-like serine/threonine-protein kinase RKS1
Uncharacterized protein
Uncharacterized protein (Fragment)
G-type lectin S-receptor-like serine/threonine-protein kinase
Receptor-like serine/threonine-protein kinase SD1-8
G-type lectin S-receptor-like serine/threonine-protein kinase
G-type lectin S-receptor-like serine/threonine-protein kinase
G-type lectin S-receptor-like serine/threonine-protein kinase SD1-1
Uncharacterized protein
Uncharacterized protein
G-type lectin S-receptor-like serine/threonine-protein kinase
G-type lectin S-receptor-like serine/threonine-protein kinase
Uncharacterized protein
G-type lectin S-receptor-like serine/threonine-protein kinase
G-type lectin S-receptor-like serine/threonine-protein kinase
G-type lectin S-receptor-like serine/threonine-protein kinase
G-type lectin S-receptor-like serine/threonine-protein kinase
G-type lectin S-receptor-like serine/threonine-protein kinase
Uncharacterized protein
Uncharacterized protein
Uncharacterized protein
Uncharacterized protein

G-type lectin S-receptor-like serine/threonine-protein kinase
Uncharacterized protein
Uncharacterized protein
Uncharacterized protein
G-type lectin S-receptor-like serine/threonine-protein kinase
Uncharacterized protein
Uncharacterized protein
Uncharacterized protein
Uncharacterized protein
Uncharacterized protein
G-type lectin S-receptor-like serine/threonine-protein kinase
Uncharacterized protein
Uncharacterized protein
G-type lectin S-receptor-like serine/threonine-protein kinase
Uncharacterized protein
Uncharacterized protein
Uncharacterized protein
Uncharacterized protein
G-type lectin S-receptor-like serine/threonine-protein kinase
Uncharacterized protein
Uncharacterized protein
Uncharacterized protein
Receptor-like serine/threonine-protein kinase SD1-8
G-type lectin S-receptor-like serine/threonine-protein kinase
G-type lectin S-receptor-like serine/threonine-protein kinase RKS1
Uncharacterized protein
G-type lectin S-receptor-like serine/threonine-protein kinase
Uncharacterized protein
G-type lectin S-receptor-like serine/threonine-protein kinase
Uncharacterized protein
G-type lectin S-receptor-like serine/threonine-protein kinase
G-type lectin S-receptor-like serine/threonine-protein kinase
G-type lectin S-receptor-like serine/threonine-protein kinase
G-type lectin S-receptor-like serine/threonine-protein kinase SD1-1
Uncharacterized protein
PAN domain-containing protein
G-type lectin S-receptor-like serine/threonine-protein kinase SD1-1
Uncharacterized protein
G-type lectin S-receptor-like serine/threonine-protein kinase
Uncharacterized protein
Uncharacterized protein

Uncharacterized protein
G-type lectin S-receptor-like serine/threonine-protein kinase
G-type lectin S-receptor-like serine/threonine-protein kinase
Uncharacterized protein
Uncharacterized protein
Uncharacterized protein
Uncharacterized protein
Uncharacterized protein
Uncharacterized protein
Uncharacterized protein
G-type lectin S-receptor-like serine/threonine-protein kinase
Uncharacterized protein
G-type lectin S-receptor-like serine/threonine-protein kinase
G-type lectin S-receptor-like serine/threonine-protein kinase
G-type lectin S-receptor-like serine/threonine-protein kinase
G-type lectin S-receptor-like serine/threonine-protein kinase
Uncharacterized protein
G-type lectin S-receptor-like serine/threonine-protein kinase
G-type lectin S-receptor-like serine/threonine-protein kinase
Uncharacterized protein
Uncharacterized protein
Uncharacterized protein
Uncharacterized protein
G-type lectin S-receptor-like serine/threonine-protein kinase
G-type lectin S-receptor-like serine/threonine-protein kinase
G-type lectin S-receptor-like serine/threonine-protein kinase
G-type lectin S-receptor-like serine/threonine-protein kinase
G-type lectin S-receptor-like serine/threonine-protein kinase SD2-2
G-type lectin S-receptor-like serine/threonine-protein kinase
Receptor-like serine/threonine-protein kinase SD1-6
Uncharacterized protein
Uncharacterized protein
Uncharacterized protein
Uncharacterized protein
Uncharacterized protein
Receptor-like serine/threonine-protein kinase SD1-8
Uncharacterized protein
Uncharacterized protein
G-type lectin S-receptor-like serine/threonine-protein kinase
Uncharacterized protein
Uncharacterized protein

Macrophage stimulating 1 (Fragment)
Plasminogen (Fragment)
Plasminogen (Fragment)
Antigen b membrane (Fragment)
Hepatocyte growth factor
Plasminogen
Uncharacterized protein
Uncharacterized protein
Hepatocyte growth factor (Fragment)
Plasminogen (Fragment)
Coagulation factor XI (Fragment)
Plasminogen (Fragment)
Uncharacterized protein
Uncharacterized protein
Hepatocyte growth factor
Uncharacterized protein
Uncharacterized protein
Plasminogen
Uncharacterized protein
Hepatocyte growth factor
Uncharacterized protein
Uncharacterized protein
Uncharacterized protein
Plasminogen
Uncharacterized protein
Uncharacterized protein
Uncharacterized protein
Hepatocyte growth factor
PAN domain-containing protein
Uncharacterized protein
S_locus_glycop domain-containing protein/B_lectin domain-containing protein/PAN_2 domain-contain
Pkinase domain-containing protein/S_locus_glycop domain-containing protein/B_lectin domain-contair
Pkinase domain-containing protein/B_lectin domain-containing protein/PAN_2 domain-containing prot
S_locus_glycop domain-containing protein/B_lectin domain-containing protein/Pkinase_Tyr domain-co
Pkinase domain-containing protein/S_locus_glycop domain-containing protein/PAN_2 domain-containi
S_locus_glycop domain-containing protein/B_lectin domain-containing protein/PAN_2 domain-contain
Pkinase domain-containing protein/S_locus_glycop domain-containing protein/B_lectin domain-contair
Pkinase domain-containing protein/S_locus_glycop domain-containing protein/B_lectin domain-contair
B_lectin domain-containing protein
B_lectin domain-containing protein/Pkinase_Tyr domain-containing protein/PAN_2 domain-containing
Pkinase domain-containing protein/S_locus_glycop domain-containing protein/B_lectin domain-contair

B_lectin domain-containing protein (Fragment)
B_lectin domain-containing protein/PAN_2 domain-containing protein
Pkinase domain-containing protein/S_locus_glycop domain-containing protein/PAN_2 domain-containi
EF-hand domain pair
Uncharacterized protein
Uncharacterized protein
Uncharacterized protein
Uncharacterized protein (Fragment)
Uncharacterized protein (Fragment)
Uncharacterized protein (Fragment)
Uncharacterized protein
Uncharacterized protein (Fragment)
Low-density lipoprotein receptor-related protein 11
Uncharacterized protein
Uncharacterized protein
Uncharacterized protein
Uncharacterized protein
Uncharacterized protein
Hepatocyte growth factor
Uncharacterized protein
Plasminogen
Uncharacterized protein
Uncharacterized protein
Uncharacterized protein
Uncharacterized protein
Uncharacterized protein
Uncharacterized protein
Uncharacterized protein
Uncharacterized protein
Uncharacterized protein
Uncharacterized protein
Uncharacterized protein
Uncharacterized protein
Uncharacterized protein
Uncharacterized protein
Uncharacterized protein
Putative fungistatic metabolite
Uncharacterized protein
Uncharacterized protein
Uncharacterized protein
Uncharacterized protein

Uncharacterized protein
Uncharacterized protein
Uncharacterized protein
Uncharacterized protein
Uncharacterized protein
Uncharacterized protein
Uncharacterized protein
Uncharacterized protein
Uncharacterized protein
Uncharacterized protein
Uncharacterized protein
Uncharacterized protein
Galactose oxidase
Uncharacterized protein
Uncharacterized protein
Micronemal protein 4
Uncharacterized protein
Galactose oxidase
Uncharacterized protein
Putative CFEM domain protein
Uncharacterized protein
Galactose oxidase
PAN domain-containing protein
Uncharacterized protein
MST1
PLG (Fragment)
HGF
KLKB1
F11
KLKB1 (Fragment)
Uncharacterized protein
Uncharacterized protein (Fragment)
Uncharacterized protein (Fragment)
Uncharacterized protein
Uncharacterized protein
Uncharacterized protein
Plasminogen (Fragment)
Hepatocyte growth factor (Fragment)
Plasminogen (Fragment)
Plasma kallikrein (Fragment)
Peptidoglycan-binding domain 1

Alpha-2-macroglobulin-like protein
Uncharacterized protein
Hepatocyte growth factor-like protein
Plasminogen (Fragment)
Uncharacterized protein
Uncharacterized protein
Uncharacterized protein
Uncharacterized protein
Uncharacterized protein
Uncharacterized protein
Hepatocyte growth factor
Uncharacterized protein
Uncharacterized protein (Fragment)
Uncharacterized protein
Uncharacterized protein
Uncharacterized protein
Uncharacterized protein
Uncharacterized protein
Predicted protein
Uncharacterized protein
Predicted protein (Fragment)
Speract/scavenger receptor, transmembrane glycoprotein
Uncharacterized protein
Uncharacterized protein
Uncharacterized protein
Uncharacterized protein
Uncharacterized protein
Uncharacterized protein
Predicted protein
Predicted protein
Predicted protein
Uncharacterized protein
Uncharacterized protein
Uncharacterized protein
Predicted protein (Fragment)
Uncharacterized protein
Uncharacterized protein
Uncharacterized protein
Uncharacterized protein
Predicted protein
Uncharacterized protein

Uncharacterized protein
Uncharacterized protein (Fragment)
Predicted protein
Uncharacterized protein
Uncharacterized protein
Peptidyl-prolyl cis-trans isomerase, cyclophilin-type
Uncharacterized protein
Predicted protein
Predicted protein (Fragment)
Hepatocyte growth factor (Fragment)
Coagulation factor XI (Fragment)
Plasminogen (Fragment)
Hepatocyte growth factor
Uncharacterized protein
Uncharacterized protein
Uncharacterized protein (Fragment)
Histone-lysine N-methyltransferase NSD2
Lipoxygenase homology domain-containing 1 isoform X2 isoform B
Glycoside hydrolase family 16
Cytochrome P450 chloroplastic isoform B
Serine threonine-kinase receptor R831
Cytochrome P450 chloroplastic isoform C
Uncharacterized protein
Erythromycin esterase
Uncharacterized protein
Uncharacterized protein
Nitrilase 2 isoform B
Glycoside hydrolase
DNA-directed RNA polymerase II subunit RPB7
40S ribosomal S7 isoform A
Cellulose binding elicitor
Serine threonine-kinase CTR1
40S ribosomal S7 isoform B
EGF domain-containing
Uncharacterized protein
Cytochrome P450 chloroplastic isoform A
Serine threonine-kinase receptor R831
O-acetyl-ADP-ribose deacetylase MACROD2 isoform X2
Serine threonine-kinase receptor R831
Mitochondrial fission process 1
Serine threonine-kinase CTR1

Serine threonine-kinase receptor R831
Peptidase C14
Uncharacterized protein
Cysteine synthase
Nudix hydrolase 8
Nitrate reductase
Lipoxygenase homology domain-containing 1 isoform X2 isoform A
Nitrilase 2 isoform A
Uncharacterized protein
Serine threonine-kinase isoform A
Uncharacterized protein
Uncharacterized protein
Expressed protein
Uncharacterized protein
Uncharacterized protein
Uncharacterized protein (Fragment)
Uncharacterized protein
Uncharacterized protein
Uncharacterized protein
Uncharacterized protein
Uncharacterized protein
Uncharacterized protein
Uncharacterized protein
Uncharacterized protein
Uncharacterized protein
Uncharacterized protein
Uncharacterized protein
Uncharacterized protein
Uncharacterized protein
Plasminogen
Kallikrein B1
Hepatocyte growth factor
Uncharacterized protein
Uncharacterized protein
Uncharacterized protein
Uncharacterized protein
Uncharacterized protein
Uncharacterized protein
Uncharacterized protein
DNA helicase
Uncharacterized protein

Uncharacterized protein
Uncharacterized protein
Uncharacterized protein
Uncharacterized protein (Fragment)
Uncharacterized protein
Uncharacterized protein
Uncharacterized protein
Uncharacterized protein
Uncharacterized protein
Uncharacterized protein
Uncharacterized protein
Uncharacterized protein
Uncharacterized protein (Fragment)
Uncharacterized protein
Uncharacterized protein
Uncharacterized protein
Uncharacterized protein
Uncharacterized protein
Uncharacterized protein
Uncharacterized protein
Uncharacterized protein
PAN/APPLE-like domain containing protein
Uncharacterized protein
Uncharacterized protein
Uncharacterized protein
Uncharacterized protein
Alpha-glucosidase
Uncharacterized protein
Glycoside hydrolase family 3
G-type lectin S-receptor-like serine/threonine-protein kinase At4g03230 isoform X1
putative receptor protein kinase ZmPK1
G-type lectin S-receptor-like serine/threonine-protein kinase SD2-2
G-type lectin S-receptor-like serine/threonine-protein kinase At4g27290 isoform X1
G-type lectin S-receptor-like serine/threonine-protein kinase At4g27290 isoform X3
G-type lectin S-receptor-like serine/threonine-protein kinase At2g19130
PAN domain-containing protein At5g03700
G-type lectin S-receptor-like serine/threonine-protein kinase At4g27290 isoform X2
G-type lectin S-receptor-like serine/threonine-protein kinase At4g27290
G-type lectin S-receptor-like serine/threonine-protein kinase At4g27290 isoform X1
G-type lectin S-receptor-like serine/threonine-protein kinase SD2-5
G-type lectin S-receptor-like serine/threonine-protein kinase B120 isoform X2

G-type lectin S-receptor-like serine/threonine-protein kinase At4g27290
G-type lectin S-receptor-like serine/threonine-protein kinase At4g27290
G-type lectin S-receptor-like serine/threonine-protein kinase RKS1
G-type lectin S-receptor-like serine/threonine-protein kinase At4g03230 isoform X3
G-type lectin S-receptor-like serine/threonine-protein kinase At4g27290 isoform X1
putative receptor protein kinase ZmPK1
G-type lectin S-receptor-like serine/threonine-protein kinase At4g27290 isoform X2
G-type lectin S-receptor-like serine/threonine-protein kinase At2g19130
receptor-like serine/threonine-protein kinase SD1-8
putative receptor protein kinase ZmPK1
LOW QUALITY PROTEIN: G-type lectin S-receptor-like serine/threonine-protein kinase At1g11330
G-type lectin S-receptor-like serine/threonine-protein kinase B120 isoform X1
G-type lectin S-receptor-like serine/threonine-protein kinase At4g27290
receptor-like serine/threonine-protein kinase SD1-8 isoform X1
G-type lectin S-receptor-like serine/threonine-protein kinase B120
receptor-like serine/threonine-protein kinase SD1-8 isoform X2
G-type lectin S-receptor-like serine/threonine-protein kinase At1g11300
G-type lectin S-receptor-like serine/threonine-protein kinase At1g11330
G-type lectin S-receptor-like serine/threonine-protein kinase At4g27290 isoform X2
receptor-like serine/threonine-protein kinase SD1-8 isoform X1
G-type lectin S-receptor-like serine/threonine-protein kinase At4g27290 isoform X4
G-type lectin S-receptor-like serine/threonine-protein kinase At4g27290 isoform X1
G-type lectin S-receptor-like serine/threonine-protein kinase At4g27290 isoform X2
G-type lectin S-receptor-like serine/threonine-protein kinase At1g34300
G-type lectin S-receptor-like serine/threonine-protein kinase At4g27290 isoform X4
G-type lectin S-receptor-like serine/threonine-protein kinase At4g03230 isoform X2
G-type lectin S-receptor-like serine/threonine-protein kinase At4g27290
LOW QUALITY PROTEIN: G-type lectin S-receptor-like serine/threonine-protein kinase At2g19130
G-type lectin S-receptor-like serine/threonine-protein kinase B120
uncharacterized protein LOC101500771
G-type lectin S-receptor-like serine/threonine-protein kinase SD3-1
putative receptor protein kinase ZmPK1
G-type lectin S-receptor-like serine/threonine-protein kinase At4g27290 isoform X2
G-type lectin S-receptor-like serine/threonine-protein kinase At4g27290 isoform X1
putative receptor protein kinase ZmPK1
G-type lectin S-receptor-like serine/threonine-protein kinase At4g27290 isoform X3
G-type lectin S-receptor-like serine/threonine-protein kinase At2g19130
G-type lectin S-receptor-like serine/threonine-protein kinase At4g27290 isoform X1
G-type lectin S-receptor-like serine/threonine-protein kinase At4g27290
uncharacterized protein LOC101510596
G-type lectin S-receptor-like serine/threonine-protein kinase At4g27290 isoform X2

S-locus-specific glycoprotein (Fragment)
Receptor-like serine/threonine-protein kinase SD1-8
G-type lectin S-receptor-like serine/threonine-protein kinase SD2-5
Receptor-like serine/threonine-protein kinase SD1-8
G-type lectin S-receptor-like serine/threonine-protein kinase LECRK3
G-type lectin S-receptor-like serine/threonine-protein kinase
G-type lectin S-receptor-like serine/threonine-protein kinase RKS1
G-type lectin S-receptor-like serine/threonine-protein kinase
G-type lectin S-receptor-like serine/threonine-protein kinase CES101 isoform X2
G-type lectin S-receptor-like serine/threonine-protein kinase RKS1
G-type lectin S-receptor-like serine/threonine-protein kinase
G-type lectin S-receptor-like serine/threonine-protein kinase LECRK1
G-type lectin S-receptor-like serine/threonine-protein kinase SD2-5 isoform X1
Putative receptor protein kinase ZmPK1
G-type lectin S-receptor-like serine/threonine-protein kinase SD2-2
Putative G-type lectin S-receptor-like serine/threonine-protein kinase
Protein kinase domain-containing protein
Putative receptor protein kinase ZmPK1
Putative G-type lectin S-receptor-like serine/threonine-protein kinase
G-type lectin S-receptor-like serine/threonine-protein kinase isoform X1
Protein kinase domain-containing protein
G-type lectin S-receptor-like serine/threonine-protein kinase isoform X1
Putative G-type lectin S-receptor-like serine/threonine-protein kinase
Receptor-like serine/threonine-protein kinase SD1-8
G-type lectin S-receptor-like serine/threonine-protein kinase
Receptor-like serine/threonine-protein kinase SD1-8
G-type lectin S-receptor-like serine/threonine-protein kinase RKS1
G-type lectin S-receptor-like serine/threonine-protein kinase RKS1
G-type lectin S-receptor-like serine/threonine-protein kinase
G-type lectin S-receptor-like serine/threonine-protein kinase RKS1
Glycosyl transferase, family 51
G-type lectin S-receptor-like serine/threonine-protein kinase RKS1 isoform X1
G-type lectin S-receptor-like serine/threonine-protein kinase RKS1
Putative receptor protein kinase ZmPK1
G-type lectin S-receptor-like serine/threonine-protein kinase
G-type lectin S-receptor-like serine/threonine-protein kinase SD2-5
G-type lectin S-receptor-like serine/threonine-protein kinase RKS1 isoform X1
G-type lectin S-receptor-like serine/threonine-protein kinase SD3-1
Receptor-like serine/threonine-protein kinase SD1-8
Uncharacterized protein
G-type lectin S-receptor-like serine/threonine-protein kinase RKS1

G-type lectin S-receptor-like serine/threonine-protein kinase SD2-5
Putative receptor protein kinase ZmPK1
Protein kinase domain-containing protein
G-type lectin S-receptor-like serine/threonine-protein kinase RKS1
Putative G-type lectin S-receptor-like serine/threonine-protein kinase
Putative receptor protein kinase ZmPK1
G-type lectin S-receptor-like serine/threonine-protein kinase
Putative receptor protein kinase ZmPK1
G-type lectin S-receptor-like serine/threonine-protein kinase SD2-5
G-type lectin S-receptor-like serine/threonine-protein kinase SD3-1
Putative receptor protein kinase ZmPK1
G-type lectin S-receptor-like serine/threonine-protein kinase B120
Putative G-type lectin S-receptor-like serine/threonine-protein kinase
PAN domain-containing protein
G-type lectin S-receptor-like serine/threonine-protein kinase
G-type lectin S-receptor-like serine/threonine-protein kinase isoform X1
G-type lectin S-receptor-like serine/threonine-protein kinase
Putative receptor protein kinase ZmPK1
G-type lectin S-receptor-like serine/threonine-protein kinase RKS1
Putative receptor protein kinase ZmPK1
G-type lectin S-receptor-like serine/threonine-protein kinase RKS1
Putative receptor protein kinase ZmPK1
Receptor-like serine/threonine-protein kinase SD1-6
G-type lectin S-receptor-like serine/threonine-protein kinase RKS1
Putative G-type lectin S-receptor-like serine/threonine-protein kinase
Pkinase/S_locus_glycop/B_lectin/Pkinase_Tyr/PAN_2/DUF3403 domain-containing protein
G-type lectin S-receptor-like serine/threonine-protein kinase RKS1 isoform X1
Receptor-like serine/threonine-protein kinase SD1-8
Putative receptor protein kinase ZmPK1
G-type lectin S-receptor-like serine/threonine-protein kinase
G-type lectin S-receptor-like serine/threonine-protein kinase
G-type lectin S-receptor-like serine/threonine-protein kinase RKS1
Putative receptor protein kinase ZmPK1
Receptor-like serine/threonine-protein kinase SD1-8
G-type lectin S-receptor-like serine/threonine-protein kinase
S-locus glycoprotein
Putative G-type lectin S-receptor-like serine/threonine-protein kinase
G-type lectin S-receptor-like serine/threonine-protein kinase isoform X1
Putative receptor protein kinase ZmPK1
G-type lectin S-receptor-like serine/threonine-protein kinase isoform X1
G-type lectin S-receptor-like serine/threonine-protein kinase

G-type lectin S-receptor-like serine/threonine-protein kinase
uncharacterized protein LOC494398 precursor
neurotrypsin-like isoform X1
uncharacterized protein LOC494398 isoform X2
uncharacterized protein LOC494398 isoform X1
plasminogen-like
macrophage mannose receptor 1-like
uncharacterized protein LOC108950161
polycystic kidney disease protein 1-like 2
Uncharacterized protein
Uncharacterized protein
uncharacterized protein LOC100185865 isoform X1
Uncharacterized protein
uncharacterized protein LOC108950126
uncharacterized protein LOC100185865 isoform X2
Uncharacterized protein
Uncharacterized protein
PAN domain-containing protein
PAN domain protein
PAN domain-containing protein
PAN domain-containing protein (Fragment)
Uncharacterized protein
Uncharacterized protein
Uncharacterized protein
Uncharacterized protein
Uncharacterized protein
Uncharacterized protein
Uncharacterized protein
Uncharacterized protein
Uncharacterized protein
Uncharacterized protein
Uncharacterized protein
Uncharacterized protein
Uncharacterized protein
Uncharacterized protein
Uncharacterized protein
Uncharacterized protein
Uncharacterized protein
Uncharacterized protein
Uncharacterized protein
Uncharacterized protein (Fragment)

Uncharacterized protein
Uncharacterized protein (Fragment)
Uncharacterized protein
Uncharacterized protein
Uncharacterized protein
Uncharacterized protein
Uncharacterized protein (Fragment)
Uncharacterized protein
Uncharacterized protein
Uncharacterized protein (Fragment)
Uncharacterized protein
Uncharacterized protein
Uncharacterized protein
Uncharacterized protein (Fragment)
Uncharacterized protein
Uncharacterized protein (Fragment)
Uncharacterized protein
Uncharacterized protein
Uncharacterized protein
Uncharacterized protein
Uncharacterized protein
Uncharacterized protein
Uncharacterized protein
Uncharacterized protein
Uncharacterized protein (Fragment)
Uncharacterized protein (Fragment)
Uncharacterized protein
Uncharacterized protein
Uncharacterized protein
Uncharacterized protein
Uncharacterized protein
Uncharacterized protein
Uncharacterized protein
Uncharacterized protein
Uncharacterized protein
Uncharacterized protein
Uncharacterized protein
Uncharacterized protein
Uncharacterized protein
Uncharacterized protein
Uncharacterized protein

Uncharacterized protein
Uncharacterized protein
Uncharacterized protein
Uncharacterized protein
Uncharacterized protein
Uncharacterized protein
Uncharacterized protein
Uncharacterized protein
Uncharacterized protein
Uncharacterized protein
Uncharacterized protein
Uncharacterized protein
Uncharacterized protein
Uncharacterized protein
Uncharacterized protein
Uncharacterized protein (Fragment)
Uncharacterized protein
Uncharacterized protein
Uncharacterized protein
Uncharacterized protein
Uncharacterized protein
Uncharacterized protein (Fragment)
Uncharacterized protein
Uncharacterized protein (Fragment)
Uncharacterized protein
Uncharacterized protein
Uncharacterized protein
Uncharacterized protein
Uncharacterized protein
Uncharacterized protein
Uncharacterized protein
Uncharacterized protein
Uncharacterized protein
Uncharacterized protein
Uncharacterized protein
Uncharacterized protein
Uncharacterized protein
Uncharacterized protein
Uncharacterized protein
Uncharacterized protein (Fragment)
Uncharacterized protein

Uncharacterized protein (Fragment)
Uncharacterized protein
Uncharacterized protein
Uncharacterized protein
Uncharacterized protein
Uncharacterized protein
Putative lectin receptor kinase-like protein (Fragment)
Uncharacterized protein (Fragment)
Uncharacterized protein
Uncharacterized protein
Uncharacterized protein (Fragment)
Uncharacterized protein
Uncharacterized protein
Uncharacterized protein
Uncharacterized protein
Uncharacterized protein
Uncharacterized protein
Uncharacterized protein
Uncharacterized protein
Uncharacterized protein (Fragment)
Uncharacterized protein (Fragment)
Uncharacterized protein
Uncharacterized protein
Uncharacterized protein
Uncharacterized protein (Fragment)
Uncharacterized protein (Fragment)
Uncharacterized protein
Uncharacterized protein (Fragment)
Uncharacterized protein (Fragment)
Uncharacterized protein
Uncharacterized protein (Fragment)
Uncharacterized protein
Uncharacterized protein (Fragment)
Uncharacterized protein (Fragment)
Uncharacterized protein
Uncharacterized protein
Uncharacterized protein
Uncharacterized protein
Uncharacterized protein
Uncharacterized protein
Uncharacterized protein

Uncharacterized protein
Uncharacterized protein
Uncharacterized protein (Fragment)
Uncharacterized protein
Uncharacterized protein (Fragment)
Uncharacterized protein (Fragment)
Uncharacterized protein
Uncharacterized protein
Uncharacterized protein
Uncharacterized protein (Fragment)
Uncharacterized protein (Fragment)
Uncharacterized protein
Uncharacterized protein
Uncharacterized protein
Uncharacterized protein
Uncharacterized protein
Uncharacterized protein
Uncharacterized protein
Uncharacterized protein
Uncharacterized protein
Uncharacterized protein (Fragment)
Uncharacterized protein (Fragment)
Uncharacterized protein (Fragment)
Uncharacterized protein (Fragment)
Uncharacterized protein (Fragment)
Uncharacterized protein
Uncharacterized protein
Uncharacterized protein
Uncharacterized protein
Uncharacterized protein (Fragment)
Uncharacterized protein
Uncharacterized protein
Uncharacterized protein (Fragment)
Uncharacterized protein
Uncharacterized protein (Fragment)
Uncharacterized protein (Fragment)
Uncharacterized protein
Uncharacterized protein
Uncharacterized protein
Uncharacterized protein
Uncharacterized protein

Uncharacterized protein
Uncharacterized protein (Fragment)
Uncharacterized protein
Uncharacterized protein
Uncharacterized protein
Uncharacterized protein
Uncharacterized protein
Uncharacterized protein (Fragment)
Uncharacterized protein (Fragment)
Uncharacterized protein (Fragment)
Uncharacterized protein (Fragment)
Uncharacterized protein
Uncharacterized protein (Fragment)
Uncharacterized protein
Uncharacterized protein
Uncharacterized protein
Uncharacterized protein
Uncharacterized protein
Uncharacterized protein
Uncharacterized protein
Uncharacterized protein
Uncharacterized protein
Uncharacterized protein
Uncharacterized protein
Uncharacterized protein (Fragment)
Uncharacterized protein
Uncharacterized protein
Uncharacterized protein
Uncharacterized protein (Fragment)
Uncharacterized protein
Uncharacterized protein
Uncharacterized protein (Fragment)
Uncharacterized protein
Uncharacterized protein (Fragment)
Uncharacterized protein (Fragment)
Uncharacterized protein
Uncharacterized protein
Uncharacterized protein (Fragment)
Uncharacterized protein
Uncharacterized protein
Uncharacterized protein

Uncharacterized protein
Uncharacterized protein
Uncharacterized protein
Uncharacterized protein
Uncharacterized protein
Uncharacterized protein
Uncharacterized protein
Uncharacterized protein (Fragment)
Uncharacterized protein
Uncharacterized protein (Fragment)
Uncharacterized protein (Fragment)
Uncharacterized protein (Fragment)
Uncharacterized protein
Uncharacterized protein
Uncharacterized protein
Uncharacterized protein
Uncharacterized protein
Uncharacterized protein
Uncharacterized protein
Uncharacterized protein
Uncharacterized protein
Uncharacterized protein
Uncharacterized protein
Uncharacterized protein
Uncharacterized protein
Uncharacterized protein
Uncharacterized protein
Uncharacterized protein
Uncharacterized protein
Uncharacterized protein
Uncharacterized protein (Fragment)
Uncharacterized protein
Uncharacterized protein
Uncharacterized protein
Uncharacterized protein
Uncharacterized protein
Uncharacterized protein
Uncharacterized protein
Uncharacterized protein (Fragment)
Uncharacterized protein (Fragment)
Uncharacterized protein

Uncharacterized protein (Fragment)
Uncharacterized protein
Uncharacterized protein
Uncharacterized protein (Fragment)
Uncharacterized protein
Uncharacterized protein (Fragment)
Uncharacterized protein
Uncharacterized protein
Uncharacterized protein
Uncharacterized protein
Uncharacterized protein
Uncharacterized protein
Uncharacterized protein
Uncharacterized protein
Uncharacterized protein
Uncharacterized protein
Uncharacterized protein
Uncharacterized protein
Uncharacterized protein
Uncharacterized protein
Uncharacterized protein
Uncharacterized protein
Uncharacterized protein
Uncharacterized protein
Uncharacterized protein
Uncharacterized protein
Uncharacterized protein
Uncharacterized protein
Uncharacterized protein
Uncharacterized protein
Uncharacterized protein
Uncharacterized protein
Uncharacterized protein
Uncharacterized protein
Uncharacterized protein
Uncharacterized protein (Fragment)
Uncharacterized protein
Uncharacterized protein
Uncharacterized protein
Uncharacterized protein
Uncharacterized protein

Uncharacterized protein
Uncharacterized protein (Fragment)
Uncharacterized protein
Uncharacterized protein
Uncharacterized protein
Uncharacterized protein
Uncharacterized protein
Uncharacterized protein
Uncharacterized protein (Fragment)
Uncharacterized protein (Fragment)
Uncharacterized protein
Uncharacterized protein
Uncharacterized protein
Uncharacterized protein
Uncharacterized protein
Uncharacterized protein
Uncharacterized protein
Uncharacterized protein
Uncharacterized protein
Uncharacterized protein
Uncharacterized protein
Uncharacterized protein
Uncharacterized protein
Uncharacterized protein
Uncharacterized protein
Uncharacterized protein
Uncharacterized protein
Uncharacterized protein
Uncharacterized protein
Uncharacterized protein
Uncharacterized protein
Uncharacterized protein
Uncharacterized protein
Uncharacterized protein
Uncharacterized protein
Uncharacterized protein
Uncharacterized protein
Uncharacterized protein
Uncharacterized protein
Uncharacterized protein
Uncharacterized protein

Uncharacterized protein (Fragment)
Uncharacterized protein
Uncharacterized protein
Uncharacterized protein
Uncharacterized protein
Uncharacterized protein
Uncharacterized protein
Uncharacterized protein
Uncharacterized protein
Uncharacterized protein (Fragment)
Uncharacterized protein
Uncharacterized protein
Uncharacterized protein
Uncharacterized protein
Uncharacterized protein
Uncharacterized protein
Uncharacterized protein
Uncharacterized protein
Uncharacterized protein
Uncharacterized protein
Uncharacterized protein
Uncharacterized protein
Uncharacterized protein
Uncharacterized protein
Uncharacterized protein
Uncharacterized protein
Uncharacterized protein
Uncharacterized protein
Uncharacterized protein
Uncharacterized protein
Uncharacterized protein
Uncharacterized protein
Uncharacterized protein
Uncharacterized protein (Fragment)
Unplaced genomic scaffold supercont1.10, whole genome shotgun sequence
Unplaced genomic scaffold supercont1.20, whole genome shotgun sequence
Uncharacterized protein
Uncharacterized protein
Uncharacterized protein
Uncharacterized protein
Uncharacterized protein

Uncharacterized protein (Fragment)
Uncharacterized protein
Uncharacterized protein
Uncharacterized protein (Fragment)
Uncharacterized protein (Fragment)
Uncharacterized protein (Fragment)
Uncharacterized protein
Uncharacterized protein
Uncharacterized protein
Galactose oxidase-like protein
Uncharacterized protein
Uncharacterized protein
Uncharacterized protein
Centrosomal protein (Fragment)
Uncharacterized protein
Uncharacterized protein
Uncharacterized protein
Uncharacterized protein
Uncharacterized protein (Fragment)
Uncharacterized protein
Uncharacterized protein
Uncharacterized protein
Uncharacterized protein
Uncharacterized protein
CLUMA_CG012660, isoform A
CLUMA_CG002608, isoform A
CLUMA_CG017132, isoform A
CLUMA_CG015859, isoform A
CLUMA_CG008967, isoform A
CLUMA_CG003095, isoform A
Uncharacterized protein
Uncharacterized protein
Uncharacterized protein
Uncharacterized protein
Uncharacterized protein
Uncharacterized protein
Uncharacterized protein
PAN domain containing protein
PAN domain containing protein
Uncharacterized protein
Uncharacterized protein

Uncharacterized protein
Uncharacterized protein
Uncharacterized protein
Glycoside hydrolase
Uncharacterized protein
Uncharacterized protein
Uncharacterized protein
Uncharacterized protein
Uncharacterized protein
Uncharacterized protein
Uncharacterized protein
Uncharacterized protein
Uncharacterized protein
Uncharacterized protein
Uncharacterized protein
Uncharacterized protein
Uncharacterized protein
Uncharacterized protein
Uncharacterized protein
Uncharacterized protein
Uncharacterized protein
Uncharacterized protein
Uncharacterized protein
Uncharacterized protein
Carbohydrate-binding module family 18 protein
Uncharacterized protein
Uncharacterized protein
Uncharacterized protein
Uncharacterized protein
Uncharacterized protein
Uncharacterized protein
Carbohydrate-binding module family 18 protein
Uncharacterized protein
Uncharacterized protein
Uncharacterized protein
Uncharacterized protein
Uncharacterized protein
Uncharacterized protein
Uncharacterized protein
Uncharacterized protein
Uncharacterized protein

Uncharacterized protein
Uncharacterized protein
Uncharacterized protein
Uncharacterized protein
Uncharacterized protein
Uncharacterized protein
Uncharacterized protein
Uncharacterized protein
Uncharacterized protein
Uncharacterized protein
Uncharacterized protein
Uncharacterized protein
Uncharacterized protein
Uncharacterized protein
Uncharacterized protein
Uncharacterized protein
Uncharacterized protein
Uncharacterized protein
Uncharacterized protein
Uncharacterized protein
Uncharacterized protein
Uncharacterized protein
Uncharacterized protein
Uncharacterized protein
Uncharacterized protein
Uncharacterized protein
Uncharacterized protein
Uncharacterized protein
Uncharacterized protein
Uncharacterized protein
Uncharacterized protein
Uncharacterized protein
Uncharacterized protein
Uncharacterized protein
Uncharacterized protein
Uncharacterized protein
Uncharacterized protein
Uncharacterized protein
Uncharacterized protein
Uncharacterized protein
Uncharacterized protein

Uncharacterized protein
Uncharacterized protein
Uncharacterized protein
Uncharacterized protein
Uncharacterized protein
Uncharacterized protein
Uncharacterized protein
Uncharacterized protein
Uncharacterized protein
Uncharacterized protein
Uncharacterized protein
Uncharacterized protein
Uncharacterized protein
Uncharacterized protein
Uncharacterized protein
Uncharacterized protein
Uncharacterized protein
Uncharacterized protein
Uncharacterized protein
Uncharacterized protein
Uncharacterized protein
Uncharacterized protein
Uncharacterized protein
Uncharacterized protein
Putative peptidoglycan binding domain-containing protein
Uncharacterized protein
Uncharacterized protein
Uncharacterized protein
Uncharacterized protein
Uncharacterized protein
Uncharacterized protein
Uncharacterized protein
Uncharacterized protein
Uncharacterized protein
Uncharacterized protein
Uncharacterized protein
Uncharacterized protein
Uncharacterized protein
Uncharacterized protein
Uncharacterized protein (Fragment)
Hepatocyte growth factor

Uncharacterized protein
Plasma kallikrein (Fragment)
Plasminogen (Fragment)
Plasminogen (Fragment)
WSC domain-containing protein-like protein 5
Lectin-B
Galactose oxidase 1
Uncharacterized protein
Uncharacterized protein
Kelch domain-containing protein
Uncharacterized protein
Galactose oxidase
Uncharacterized protein
Wsc domain containing protein
Galactose oxidase
Uncharacterized protein
AAA family ATPase
Uncharacterized protein
Uncharacterized protein
Kelch domain-containing protein
Uncharacterized protein
Uncharacterized protein
Uncharacterized protein
Uncharacterized protein
Kelch domain-containing protein
Glycosyltransferase family 31
Kelch domain-containing protein
Uncharacterized protein
Kelch domain-containing protein
WSC domain-containing protein
Kelch domain-containing protein
Uncharacterized protein
Kelch domain-containing protein
Keratin-associated protein 5-4
Uncharacterized protein (Fragment)
Galactose oxidase
Uncharacterized protein
Uncharacterized protein (Fragment)
Kelch domain-containing protein
Uncharacterized protein
Agglutinin Isolectin 1

PAN domain containing protein
Uncharacterized protein (Fragment)
Agglutinin isolectin 1
Kelch domain-containing protein
Uncharacterized protein
Uncharacterized protein
Uncharacterized protein
Keratin-associated protein 5-4
Galactose oxidase
Uncharacterized protein
Copper radical oxidase
Copper radical oxidase
Uncharacterized protein
Kelch domain-containing protein
Uncharacterized protein
Uncharacterized protein
Kelch domain-containing protein
Uncharacterized protein
Lectin
Uncharacterized protein
Uncharacterized protein
Kelch domain-containing protein
Uncharacterized protein
Lectin
Putative kelch domain-containing protein
Putative agglutinin isolectin 1
Carbohydrate-binding protein
Galactose oxidase (Kelch domain-containing protein)
Uncharacterized protein
Uncharacterized protein
Hepatocyte growth factor
Uncharacterized protein
Uncharacterized protein
Plasminogen
Uncharacterized protein
Uncharacterized protein
Coagulation factor XI, transcript variant X4
Coagulation factor XI, transcript variant X6
Plasminogen
Plasminogen
Hepatocyte growth factor

Coagulation factor XI, transcript variant X2
Coagulation factor XI, transcript variant X5
Uncharacterized protein
PAN domain protein
Uncharacterized protein
PAN domain protein
PAN domain protein
PAN domain protein
Uncharacterized protein
PAN domain protein
PAN domain protein
Uncharacterized protein
Uncharacterized protein (Fragment)
Uncharacterized protein
Uncharacterized protein
Uncharacterized protein
Uncharacterized protein
Uncharacterized protein
Uncharacterized protein
Uncharacterized protein
Uncharacterized protein
Uncharacterized protein
Uncharacterized protein
Uncharacterized protein
Uncharacterized protein
Uncharacterized protein
S-locus glycoprotein
S-locus glycoprotein
Uncharacterized protein
Uncharacterized protein
Uncharacterized protein
S-locus glycoprotein
S-locus glycoprotein
S-locus glycoprotein
Uncharacterized protein
S-locus glycoprotein
Uncharacterized protein
Uncharacterized protein
Uncharacterized protein
Uncharacterized protein
Uncharacterized protein

Uncharacterized protein
Uncharacterized protein
Uncharacterized protein
Uncharacterized protein
Uncharacterized protein
Uncharacterized protein
Uncharacterized protein
Uncharacterized protein
S-locus glycoprotein
S-locus glycoprotein
S-locus glycoprotein
Uncharacterized protein
S-locus glycoprotein
S-locus glycoprotein
S-locus glycoprotein
Uncharacterized protein
S-locus glycoprotein
Uncharacterized protein
Uncharacterized protein
S-locus glycoprotein
Uncharacterized protein
Uncharacterized protein
Uncharacterized protein
Uncharacterized protein
Uncharacterized protein
S-locus glycoprotein
S-locus glycoprotein
Uncharacterized protein
S-locus glycoprotein
S-locus glycoprotein
Uncharacterized protein
S-locus glycoprotein
S-locus glycoprotein
S-locus glycoprotein
S-locus glycoprotein
S-locus glycoprotein
Uncharacterized protein
S-locus glycoprotein
Uncharacterized protein
S-locus glycoprotein
S-locus glycoprotein

Uncharacterized protein
S-locus glycoprotein
Uncharacterized protein
Uncharacterized protein
Uncharacterized protein
S-locus glycoprotein
Uncharacterized protein
Uncharacterized protein
S-locus glycoprotein
S-locus glycoprotein
S-locus glycoprotein
S-locus glycoprotein
S-locus glycoprotein
S-locus glycoprotein
Uncharacterized protein
Uncharacterized protein
S-locus glycoprotein
Uncharacterized protein
S-locus glycoprotein
Uncharacterized protein
S-locus glycoprotein
Uncharacterized protein
S-locus glycoprotein
S-locus glycoprotein
S-locus glycoprotein
S-locus glycoprotein
Uncharacterized protein
Uncharacterized protein
S-locus glycoprotein
S-locus glycoprotein
Uncharacterized protein
S-locus glycoprotein
S-locus glycoprotein
S-locus glycoprotein
S-locus glycoprotein
Uncharacterized protein
S-locus glycoprotein
Uncharacterized protein
Uncharacterized protein
S-locus glycoprotein
S-locus glycoprotein

S-locus glycoprotein
S-locus glycoprotein
Uncharacterized protein
S-locus glycoprotein
S-locus glycoprotein
S-locus glycoprotein
Uncharacterized protein
Fringe-like protein
Apple-like protein
Uncharacterized protein
Uncharacterized protein
Apple-like protein
Uncharacterized protein
Uncharacterized protein
Uncharacterized protein
Uncharacterized protein
Uncharacterized protein
GLEYA adhesin domain
Uncharacterized protein
Uncharacterized protein
Hepatocyte growth factor (Fragment)
Hepatocyte growth factor-like (Fragment)
Plasminogen (Fragment)
Plasma kallikrein (Fragment)
Plasminogen (Fragment)
Galactose oxidase-like protein
Uncharacterized protein
Uncharacterized protein
Uncharacterized protein
Uncharacterized protein
Uncharacterized protein
S-locus-related I (Fragment)
Uncharacterized protein
Fucolectin
Uncharacterized protein
Uncharacterized protein
Hemicentin-1
Uncharacterized protein
Uncharacterized protein
Uncharacterized protein
Fibronectin type III domain-containing protein 1

Ficolin-2
Uncharacterized protein
Ficolin-2
Uncharacterized protein
Uncharacterized protein
Uncharacterized protein
Uncharacterized protein
Uncharacterized protein
Uncharacterized protein
Uncharacterized protein
Ficolin-1
Uncharacterized protein
Sushi, von Willebrand factor type A, EGF and pentraxin domain-containing protein 1
Uncharacterized protein
Uncharacterized protein
Ficolin-1
Uncharacterized protein
Uncharacterized protein
Uncharacterized protein
Uncharacterized protein
Uncharacterized protein
Ficolin-2
Ficolin-2
Uncharacterized protein
Alpha-2C adrenergic receptor
Agrin
Thrombospondin-2
von Willebrand factor D and EGF domain-containing protein
Uncharacterized protein
Uncharacterized protein
Laminin-like protein epi-1
Low-density lipoprotein receptor-related protein 11
Ficolin-2
Uncharacterized protein
Uncharacterized protein
Dynein heavy chain 2, axonemal
Fucolectin-5
Fucolectin
Uncharacterized protein
Uncharacterized protein
Uncharacterized protein

Fucolectin-4
Neurotrypsin
Uncharacterized protein
Uncharacterized protein
Lectin BRA-3
Uncharacterized protein
Fibropellin-1
Uncharacterized protein
Uncharacterized protein
COBW domain-containing protein 1
Uncharacterized protein
Uncharacterized protein
Uncharacterized protein
Lectin
Fucolectin-1
Fucolectin
Uncharacterized protein
Uncharacterized protein
Uncharacterized protein
Uncharacterized protein
Uncharacterized protein
Aggrecan core protein
Uncharacterized protein
Uncharacterized protein
Uncharacterized protein
Ficolin-2
Neurogenic locus notch-like protein 1
Fucolectin
Plasminogen
Uncharacterized protein
Uncharacterized protein
Ficolin-2
Ficolin-2
Protein VPRBP
Uncharacterized protein
Fucolectin-4
Uncharacterized protein
Ig-like and fibronectin type-III domain-containing protein C25G4.10
Uncharacterized protein
Uncharacterized protein
Uncharacterized protein

Uncharacterized protein
PLG (Fragment)
Plasminogen
F11 (Fragment)
HGF (Fragment)
Plasma kallikrein-like protein
MST1
KLKB1
Hepatocyte growth factor
Hepatocyte growth factor
PAN domain-containing protein
PAN domain-containing protein
Uncharacterized protein
Calcium binding EGF domain-containing protein
Uncharacterized protein
Uncharacterized protein
Uncharacterized protein
Uncharacterized protein
Thrombospondin type 1 domain protein
PAN domain protein
Calcium-binding EGF domain protein
Uncharacterized protein
Calcium binding EGF domain-containing protein
PAN domain-containing protein (Fragment)
PAN domain-containing protein
TSP1 domain-containing protein TSP3
Extracellular protein with signal peptide, 5xEGF and apple domains
Mucin-like low complexity glycoprotein with a signal peptide and an apple domain
Cysteine-rich extracellular protein with a signal peptide and two apple domains
TSP1 domain-containing protein TSP3
Uncharacterized protein
Uncharacterized protein
Uncharacterized protein
Uncharacterized protein
Uncharacterized protein
Uncharacterized protein
Uncharacterized protein (Fragment)
Uncharacterized protein (Fragment)
Uncharacterized protein
Uncharacterized protein
Uncharacterized protein

Uncharacterized protein
Uncharacterized protein
Uncharacterized protein
Uncharacterized protein
Uncharacterized protein (Fragment)
Uncharacterized protein
Uncharacterized protein
Uncharacterized protein
Uncharacterized protein
Uncharacterized protein
Plasminogen
Plasminogen (Fragment)
Coagulation factor XI (Fragment)
Hepatocyte growth factor (Fragment)
Plasminogen (Fragment)
receptor-like serine/threonine-protein kinase SD1-8 isoform X3
G-type lectin S-receptor-like serine/threonine-protein kinase SD1-13 isoform X5
G-type lectin S-receptor-like serine/threonine-protein kinase At4g27290 isoform X1
G-type lectin S-receptor-like serine/threonine-protein kinase At1g11330
receptor-like serine/threonine-protein kinase SD1-8 isoform X4
uncharacterized protein LOC103485800
G-type lectin S-receptor-like serine/threonine-protein kinase SD1-13 isoform X4
G-type lectin S-receptor-like serine/threonine-protein kinase At4g27290 isoform X3
G-type lectin S-receptor-like serine/threonine-protein kinase At1g34300
G-type lectin S-receptor-like serine/threonine-protein kinase B120 isoform X2
G-type lectin S-receptor-like serine/threonine-protein kinase B120 isoform X1
G-type lectin S-receptor-like serine/threonine-protein kinase At4g27290
G-type lectin S-receptor-like serine/threonine-protein kinase At4g27290 isoform X1
G-type lectin S-receptor-like serine/threonine-protein kinase At1g11330
G-type lectin S-receptor-like serine/threonine-protein kinase At1g67520 isoform X1
G-type lectin S-receptor-like serine/threonine-protein kinase SD1-13 isoform X3
receptor-like serine/threonine-protein kinase SD1-8 isoform X2
G-type lectin S-receptor-like serine/threonine-protein kinase At4g27290 isoform X2
putative receptor protein kinase ZmPK1
receptor-like serine/threonine-protein kinase SD1-8 isoform X3
G-type lectin S-receptor-like serine/threonine-protein kinase At4g27290 isoform X2
G-type lectin S-receptor-like serine/threonine-protein kinase At4g27290 isoform X2
G-type lectin S-receptor-like serine/threonine-protein kinase At4g27290 isoform X1
putative receptor protein kinase ZmPK1
putative receptor protein kinase ZmPK1
G-type lectin S-receptor-like serine/threonine-protein kinase At4g27290 isoform X1

G-type lectin S-receptor-like serine/threonine-protein kinase At4g27290
receptor-like serine/threonine-protein kinase SD1-8 isoform X1
G-type lectin S-receptor-like serine/threonine-protein kinase At4g27290 isoform X2
G-type lectin S-receptor-like serine/threonine-protein kinase At4g27290 isoform X1
G-type lectin S-receptor-like serine/threonine-protein kinase At4g27290 isoform X2
G-type lectin S-receptor-like serine/threonine-protein kinase At4g27290
G-type lectin S-receptor-like serine/threonine-protein kinase At2g19130
receptor-like serine/threonine-protein kinase SD1-8
G-type lectin S-receptor-like serine/threonine-protein kinase SD2-2
G-type lectin S-receptor-like serine/threonine-protein kinase SD2-5
receptor-like serine/threonine-protein kinase SD1-8 isoform X2
PAN domain-containing protein At5g03700
LOW QUALITY PROTEIN: G-type lectin S-receptor-like serine/threonine-protein kinase CES101
G-type lectin S-receptor-like serine/threonine-protein kinase At4g03230
G-type lectin S-receptor-like serine/threonine-protein kinase SD1-13 isoform X1
G-type lectin S-receptor-like serine/threonine-protein kinase At4g27290
uncharacterized protein LOC103489252
receptor-like serine/threonine-protein kinase SD1-8 isoform X1
G-type lectin S-receptor-like serine/threonine-protein kinase SD2-5
G-type lectin S-receptor-like serine/threonine-protein kinase SD1-13 isoform X2
Uncharacterized protein
Uncharacterized protein
Uncharacterized protein
Uncharacterized protein
Uncharacterized protein
Uncharacterized protein
Uncharacterized protein
Uncharacterized protein
Uncharacterized protein
Uncharacterized protein
Uncharacterized protein
Uncharacterized protein
Uncharacterized protein
Uncharacterized protein
Uncharacterized protein
Uncharacterized protein
Uncharacterized protein
Uncharacterized protein
Uncharacterized protein
Uncharacterized protein
Uncharacterized protein

Uncharacterized protein
Uncharacterized protein
Uncharacterized protein
Uncharacterized protein
Uncharacterized protein
Uncharacterized protein
Uncharacterized protein
Uncharacterized protein
Uncharacterized protein
Uncharacterized protein
Uncharacterized protein
Uncharacterized protein
Uncharacterized protein
Uncharacterized protein
Uncharacterized protein
Uncharacterized protein
Uncharacterized protein
Uncharacterized protein (Fragment)
Uncharacterized protein (Fragment)
Uncharacterized protein (Fragment)
Uncharacterized protein (Fragment)
Uncharacterized protein (Fragment)
Uncharacterized protein (Fragment)
Uncharacterized protein (Fragment)
Uncharacterized protein (Fragment)
Uncharacterized protein (Fragment)
Uncharacterized protein
Uncharacterized protein (Fragment)
Uncharacterized protein (Fragment)
No-mechanoreceptor potential a
Uncharacterized protein
Uncharacterized protein
Uncharacterized protein
Uncharacterized protein
Uncharacterized protein
Uncharacterized protein
Uncharacterized protein
Uncharacterized protein
Uncharacterized protein
Uncharacterized protein
Uncharacterized protein

Uncharacterized protein
CSON013239 protein
CSON004351 protein
CSON002270 protein
CSON006616 protein
CSON002270 protein
CSON000530 protein
CSON001081 protein
CSON012649 protein
CSON008419 protein
CSON002270 protein
CSON006874 protein
CSON005481 protein
CSON004351 protein
CSON013241 protein
Uncharacterized protein
Uncharacterized protein
Uncharacterized protein
Uncharacterized protein
Uncharacterized protein
Uncharacterized protein
Uncharacterized protein
Uncharacterized protein
Uncharacterized protein
Uncharacterized protein
Uncharacterized protein
PAN domain-containing protein
PAN domain-containing protein
PAN domain-containing protein
PAN domain-containing protein
Microneme protein 5
PAN domain-containing protein
Microneme protein 5
Uncharacterized protein
Dullard family phosphatase domain-containing protein
PAN domain-containing protein
PAN domain-containing protein
PAN domain-containing protein
PAN domain-containing protein
Uncharacterized protein (Fragment)
Uncharacterized protein

Uncharacterized protein
Uncharacterized protein (Fragment)
Uncharacterized protein
Apple-like protein (Fragment)
Apple-like protein
Uncharacterized protein
Apple-like protein
Apple-like protein
Apple-like protein
Apple-like protein
Apple-like protein
Apple-like protein
Uncharacterized protein
Apple-like protein
Apple-like protein
Apple-like protein
Apple-like protein
Apple-like protein
Apple-like protein
Apple-like protein
Apple-like protein
Apple-like protein
Apple-like protein
Apple-like protein
Apple-like protein
Apple-like protein
Apple-like protein
Apple-like protein
Apple-like protein (Fragment)
Apple-like protein
Apple-like protein
Apple-like protein
Apple-like protein
Apple-like protein
Apple-like protein
Apple-like protein
Apple-like protein
Apple-like protein
Apple-like protein
Apple-like protein
Apple-like protein

Apple-like protein
Apple-like protein
Apple-like protein
Apple-like protein
Apple-like protein
Apple-like protein
Apple-like protein (Fragment)
Apple-like protein
Apple-like protein
Apple-like protein
Apple-like protein
Apple-like protein
Apple-like protein
Apple-like protein
Apple-like protein
Bulb-type lectin domain-containing protein (Fragment)
Apple-like protein (Fragment)
Uncharacterized protein
Uncharacterized protein
Apple-like protein
Apple-like protein
Apple-like protein
Uncharacterized protein
Hepatocyte growth factor
Uncharacterized protein
Uncharacterized protein
Uncharacterized protein
Plasminogen
Uncharacterized protein
Uncharacterized protein
Uncharacterized protein
Uncharacterized protein
Cuticlin-1
Uncharacterized protein
Uncharacterized protein
Circumsporozoite protein
Uncharacterized protein
Plasminogen
Uncharacterized protein
Uncharacterized protein
Uncharacterized protein

Uncharacterized protein
Uncharacterized protein
Uncharacterized protein
Uncharacterized protein
Uncharacterized protein
Uncharacterized protein
Uncharacterized protein
Macrophage stimulating 1 (Fragment)
Uncharacterized protein
Uncharacterized protein (Fragment)
Microneme protein mic4
Pan domain-containing protein
Pan domain-containing protein
Pan domain-containing protein
Pan domain protein
Microneme protein mic4
Microneme protein mic4
Pan domain-containing protein (Fragment)
Microneme protein mic4
Microneme protein mic4
Pan domain-containing protein
Pan domain-containing protein
Pan domain-containing protein
Pan domain protein
Pan domain-containing protein
Uncharacterized protein
Microneme protein mic4
Pan domain-containing protein
Microneme protein mic4
Pan domain-containing protein (Fragment)
Beta-glucanase
Pan domain-containing protein (Fragment)
Pan apple domain-containing protein
Pan domain-containing protein
Pan domain-containing protein (Fragment)
Microneme protein (Fragment)
Microneme protein
Uncharacterized protein
Uncharacterized protein
Uncharacterized protein
Uncharacterized protein

Plasma kallikrein
Uncharacterized protein
Uncharacterized protein
Uncharacterized protein
Uncharacterized protein
Uncharacterized protein
Uncharacterized protein
Uncharacterized protein
Uncharacterized protein
Glycosyltransferase family 31 protein
Uncharacterized protein
Uncharacterized protein
Uncharacterized protein
Uncharacterized protein
Uncharacterized protein
Uncharacterized protein
Uncharacterized protein
Adhesion G-protein coupled receptor V1
Hepatocyte growth factor
Plasminogen
Hepatocyte growth factor
Hepatocyte growth factor
Macrophage stimulating 1 (Hepatocyte growth factor-like)
Hepatocyte growth factor-like 1
Plasminogen (Fragment)
Hepatocyte growth factor
Plasminogen
Adhesion G-protein-coupled receptor V1
Coagulation factor XI
Chorion peroxidase
Coagulation factor XI (Fragment)
Coagulation factor XI
Chorion peroxidase
Coagulation factor XI
Coagulation factor XI (Fragment)
Chorion peroxidase-like protein
Chorion peroxidase
Chorion peroxidase
Uncharacterized protein
Brain chitinase and chia
Transmembrane protease serine

Coagulation factor XI
Uncharacterized protein
Transmembrane protease serine
Putative PAN/APPLE domain-containing protein
Putative PAN/APPLE domain-containing protein
Transmembrane protease serine
Chorion peroxidase (Fragment)
PAN/APPLE domain-containing protein (Fragment)
Putative PAN/APPLE domain-containing protein (Fragment)
Transmembrane protease serine
Chorion peroxidase
Putative Coagulation factor XI (Fragment)
Chorion peroxidase
Coagulation factor XI
Transmembrane protease serine
Coagulation factor XI
Coagulation factor XI
Transmembrane protease serine (Fragment)
Chorion peroxidase
Transmembrane protease serine
Brain chitinase and chia
Uncharacterized protein
Transmembrane protease serine
Coagulation factor XI
Chorion peroxidase
Transmembrane protease serine
Chorion peroxidase
Uncharacterized protein (Fragment)
Uncharacterized protein
Uncharacterized protein (Fragment)
Uncharacterized protein (Fragment)
Coagulation factor XI
Coagulation factor XI (Fragment)
PAN/APPLE domain-containing protein
Chorion peroxidase
Putative PAN/APPLE domain-containing protein
Uncharacterized protein (Fragment)
Chorion peroxidase
Chorion peroxidase
Putative No mechanoreceptor potential A
PAN/APPLE domain-containing protein (Fragment)

PAN/APPLE domain-containing protein (Fragment)
Putative Brain chitinase and chia
Coagulation factor XI
PAN/APPLE domain-containing protein
Putative Coagulation factor XI (Fragment)
PAN/APPLE domain-containing protein
Chorion peroxidase
Coagulation factor XI
Chorion peroxidase (Fragment)
Uncharacterized protein
Chorion peroxidase (Fragment)
Coagulation factor XI (Fragment)
Class A Scavenger Receptor (SRCR domain) with Serine Protease domain
Uncharacterized protein
PAN/APPLE domain-containing protein
Chorion peroxidase
Chorion peroxidase
Coagulation factor XI (Fragment)
Uncharacterized protein (Fragment)
Chorion peroxidase
Brain chitinase and chia
Uncharacterized protein (Fragment)
Brain chitinase and chia (Fragment)
Coagulation factor XI
PAN/APPLE domain-containing protein
Coagulation factor XI (Fragment)
Chorion peroxidase
Chorion peroxidase
Chorion peroxidase
Coagulation factor XI
Putative Coagulation factor XI (Fragment)
Chorion peroxidase
Transmembrane protease serine
Brain chitinase and chia (Fragment)
Chorion peroxidase
Transmembrane protease serine (Fragment)
PAN/APPLE domain-containing protein
Chorion peroxidase
Coagulation factor XI
Uncharacterized protein
Uncharacterized protein (Fragment)

Uncharacterized protein (Fragment)
Coagulation factor XI
Chorion peroxidase
Chorion peroxidase
Chorion peroxidase
Chorion peroxidase
Chorion peroxidase
Coagulation factor XI (Fragment)
Coagulation factor XI
Chorion peroxidase
Uncharacterized protein
PAN/APPLE domain-containing protein (Fragment)
Chorion peroxidase
PAN/APPLE domain-containing protein
Putative PAN/APPLE domain-containing protein (Fragment)
Chorion peroxidase (Fragment)
Coagulation factor XI
Chorion peroxidase-like protein (Fragment)
Chorion peroxidase
Chorion peroxidase
Chorion peroxidase
Chorion peroxidase
Chorion peroxidase
Chorion peroxidase
Putative PAN/APPLE domain-containing protein
PAN/APPLE domain-containing protein (Fragment)
PAN/APPLE domain-containing protein
Putative No mechanoreceptor potential A
Zona pellucida domain-containing protein
PAN/APPLE domain-containing protein
PAN/APPLE domain-containing protein
Chorion peroxidase
Uncharacterized protein
PAN/APPLE domain-containing protein
Transmembrane protease serine
PAN/APPLE domain-containing protein (Fragment)
PAN/APPLE domain-containing protein
Putative Transmembrane protease serine 5
Chorion peroxidase
Putative PAN/APPLE domain-containing protein
PAN/APPLE domain-containing protein

Brain chitinase and chia
PAN/APPLE domain-containing protein (Fragment)
Chorion peroxidase
Uncharacterized protein
PAN/APPLE domain-containing protein
PAN/APPLE domain-containing protein
PAN/APPLE domain-containing protein
PAN/APPLE domain-containing protein
Transmembrane protease serine
Chorion peroxidase
Chorion peroxidase
Chorion peroxidase
Uncharacterized protein (Fragment)
Uncharacterized protein (Fragment)
Chorion peroxidase
Chorion peroxidase-like protein (Fragment)
Coagulation factor XI (Fragment)
Coagulation factor XI (Fragment)
PAN/APPLE domain-containing protein
Coagulation factor XI
PAN/APPLE domain-containing protein (Fragment)
Uncharacterized protein
Coagulation factor XI (Fragment)
Chorion peroxidase
Putative PAN/APPLE domain-containing protein
Uncharacterized protein
Transmembrane protease serine
PAN/APPLE domain-containing protein
Chorion peroxidase
Chorion peroxidase
PAN/APPLE domain-containing protein
PAN/APPLE domain-containing protein
PAN/APPLE domain-containing protein
PAN/APPLE domain-containing protein (Fragment)
Uncharacterized protein
Zona pellucida domain-containing protein
Class A Scavenger Receptor (SRCR domain) with Serine Protease domain
Putative Zona pellucida domain-containing protein
No-mechanoreceptor potential A
Uncharacterized protein
Uncharacterized protein (Fragment)

Chorion peroxidase (Fragment)
Uncharacterized protein (Fragment)
Coagulation factor XI
Coagulation factor XI
Uncharacterized protein
Chorion peroxidase
Chorion peroxidase
Coagulation factor XI (Fragment)
Coagulation factor XI
Class A Scavenger Receptor (SRCR domain) with Serine Protease domain
Class A Scavenger Receptor (SRCR domain) with Serine Protease domain
Putative No mechanoreceptor potential A
Class A Scavenger Receptor (SRCR domain) with Serine Protease domain
Zona pellucida domain-containing protein
Putative No mechanoreceptor potential A
Uncharacterized protein
Chorion peroxidase
Chorion peroxidase
Chorion peroxidase
Uncharacterized protein
Putative No mechanoreceptor potential A
Putative Zona pellucida domain-containing protein (Fragment)
Uncharacterized protein
Class A Scavenger Receptor (SRCR domain) with Serine Protease domain
Putative No mechanoreceptor potential A
Uncharacterized protein
Zona pellucida domain-containing protein
Uncharacterized protein
Uncharacterized protein
Zona pellucida domain-containing protein (Fragment)
Class A Scavenger Receptor (SRCR domain) with Serine Protease domain
Uncharacterized protein
Chorion peroxidase
Chorion peroxidase
Coagulation factor XI
PAN/APPLE domain-containing protein (Fragment)
Coagulation factor XI (Fragment)
Coagulation factor XI
Coagulation factor XI (Fragment)
Uncharacterized protein
No mechanoreceptor potential A-like protein

Putative No mechanoreceptor potential A
Class A Scavenger Receptor (SRCR domain) with Serine Protease domain
Uncharacterized protein
Uncharacterized protein
Uncharacterized protein
Putative No mechanoreceptor potential A
Uncharacterized protein
Class A Scavenger Receptor (SRCR domain) with Serine Protease domain
Uncharacterized protein
Uncharacterized protein
Uncharacterized protein
Zona pellucida domain-containing protein
Putative Zona pellucida domain-containing protein
Class A Scavenger Receptor (SRCR domain) with Serine Protease domain
Putative No mechanoreceptor potential A
Uncharacterized protein
Putative No mechanoreceptor potential A
Zona pellucida domain-containing protein
Putative Zona pellucida domain-containing protein
Chorion peroxidase
Chorion peroxidase
Zona pellucida domain-containing protein
Coagulation factor XI
PAN/APPLE domain-containing protein
Uncharacterized protein
Uncharacterized protein
Putative Zona pellucida domain-containing protein
Uncharacterized protein
Putative Coagulation factor XI
PAN/APPLE domain-containing protein
Chorion peroxidase
PAN/APPLE domain-containing protein
Uncharacterized protein (Fragment)
Chorion peroxidase
Transmembrane protease serine
Chorion peroxidase
Uncharacterized protein (Fragment)
Coagulation factor XI
Chorion peroxidase
Chorion peroxidase
Peroxinectin-like protein

Chorion peroxidase
PAN/APPLE domain-containing protein
PAN/APPLE domain-containing protein (Fragment)
Uncharacterized protein (Fragment)
PAN/APPLE domain-containing protein (Fragment)
Coagulation factor XI
Chorion peroxidase-like protein (Fragment)
PAN/APPLE domain-containing protein
Chorion peroxidase
PAN/APPLE domain-containing protein (Fragment)
PAN/APPLE domain-containing protein
Chorion peroxidase
Transmembrane protease serine
Uncharacterized protein
Putative Coagulation factor XI (Fragment)
Coagulation factor XI
PAN/APPLE domain-containing-like protein
Brain chitinase and chia (Fragment)
Uncharacterized protein
Uncharacterized protein
Coagulation factor XI (Fragment)
Coagulation factor XI
Transmembrane protease serine
Chorion peroxidase-like protein
Uncharacterized protein
Uncharacterized protein (Fragment)
Chorion peroxidase
Coagulation factor XI (Fragment)
Chorion peroxidase
Coagulation factor XI
PAN/APPLE domain-containing protein (Fragment)
PAN/APPLE domain-containing protein
Uncharacterized protein
Uncharacterized protein
Putative Chorion peroxidase
Putative Chorion peroxidase
Transmembrane protease serine
Putative PAN/APPLE domain-containing protein (Fragment)
Uncharacterized protein
Transmembrane protease serine
Coagulation factor XI (Fragment)

Coagulation factor XI
Coagulation factor XI
Chorion peroxidase
Chorion peroxidase
PAN/APPLE domain-containing protein (Fragment)
PAN/APPLE domain-containing-like protein (Fragment)
Putative PAN/APPLE domain-containing protein
Chorion peroxidase
PAN/APPLE domain-containing protein (Fragment)
Coagulation factor XI
PAN/APPLE domain-containing protein (Fragment)
Putative No mechanoreceptor potential A
Uncharacterized protein
Zona pellucida domain-containing protein
Uncharacterized protein
Putative No mechanoreceptor potential A
No mechanoreceptor potential A-like protein
Uncharacterized protein
Putative Zona pellucida domain-containing protein
Putative No mechanoreceptor potential A
Uncharacterized protein
Putative No mechanoreceptor potential A
Zona pellucida domain-containing protein
Putative No mechanoreceptor potential A
Uncharacterized protein
Putative No mechanoreceptor potential A
Putative No mechanoreceptor potential A
Class A Scavenger Receptor (SRCR domain) with Serine Protease domain
Putative Zona pellucida domain-containing protein
Uncharacterized protein
Class A Scavenger Receptor (SRCR domain) with Serine Protease domain
Class A Scavenger Receptor (SRCR domain) with Serine Protease domain
Zona pellucida domain-containing protein (Fragment)
Uncharacterized protein (Fragment)
Uncharacterized protein
Putative No mechanoreceptor potential A
Putative No mechanoreceptor potential A
Uncharacterized protein
Uncharacterized protein
Uncharacterized protein (Fragment)
Uncharacterized protein

Putative No mechanoreceptor potential A
Uncharacterized protein (Fragment)
Uncharacterized protein
Class A Scavenger Receptor (SRCR domain) with Serine Protease domain
Uncharacterized protein
Uncharacterized protein
Putative No mechanoreceptor potential A
Class A Scavenger Receptor (SRCR domain) with Serine Protease domain
Putative No mechanoreceptor potential A
Uncharacterized protein
Uncharacterized protein
Zona pellucida domain-containing protein
Zona pellucida domain-containing protein
Uncharacterized protein
Putative No mechanoreceptor potential A
Putative No mechanoreceptor potential A
Putative No mechanoreceptor potential A (Fragment)
Uncharacterized protein
Uncharacterized protein
Uncharacterized protein
Uncharacterized protein
Uncharacterized protein
Uncharacterized protein
Uncharacterized protein
Uncharacterized protein
Uncharacterized protein
Uncharacterized protein
Uncharacterized protein
Uncharacterized protein
Uncharacterized protein
Uncharacterized protein
Uncharacterized protein
Uncharacterized protein
Uncharacterized protein
Uncharacterized protein
Uncharacterized protein
Uncharacterized protein
Uncharacterized protein
Uncharacterized protein
Uncharacterized protein
Uncharacterized protein

Uncharacterized protein
Uncharacterized protein
Uncharacterized protein
Uncharacterized protein
Uncharacterized protein
Uncharacterized protein
Uncharacterized protein
Uncharacterized protein
Uncharacterized protein
Uncharacterized protein
Uncharacterized protein
Uncharacterized protein
Uncharacterized protein
Uncharacterized protein
Uncharacterized protein
Uncharacterized protein
Uncharacterized protein
Uncharacterized protein
Uncharacterized protein
Uncharacterized protein
Uncharacterized protein
Uncharacterized protein
Uncharacterized protein
Uncharacterized protein
Uncharacterized protein
Uncharacterized protein
Uncharacterized protein
Uncharacterized protein
Uncharacterized protein
Uncharacterized protein
Uncharacterized protein
Uncharacterized protein
Uncharacterized protein
Uncharacterized protein
Uncharacterized protein
Uncharacterized protein
Uncharacterized protein
Uncharacterized protein
Uncharacterized protein
Uncharacterized protein
Uncharacterized protein

Uncharacterized protein
Uncharacterized protein
Uncharacterized protein
Uncharacterized protein
Uncharacterized protein
Uncharacterized protein
Uncharacterized protein
Pectate lyase
Uncharacterized protein
Uncharacterized protein
Uncharacterized protein
Uncharacterized protein
Uncharacterized protein
Uncharacterized protein
Uncharacterized protein
Uncharacterized protein
Uncharacterized protein
Uncharacterized protein
Uncharacterized protein
Uncharacterized protein
Uncharacterized protein
Uncharacterized protein
Uncharacterized protein
Uncharacterized protein
Uncharacterized protein
Uncharacterized protein
Uncharacterized protein
Uncharacterized protein
Uncharacterized protein
Uncharacterized protein
Uncharacterized protein
Uncharacterized protein
Uncharacterized protein
Uncharacterized protein
Uncharacterized protein
Uncharacterized protein
Uncharacterized protein
Uncharacterized protein
Uncharacterized protein
Uncharacterized protein
Uncharacterized protein

Uncharacterized protein
Uncharacterized protein
Uncharacterized protein
Uncharacterized protein
Uncharacterized protein
Cell attachment protein in somatic embryogenesis
Uncharacterized protein
Uncharacterized protein
Putative lipoprotein YfhM
Uncharacterized protein
Uncharacterized protein
Plasminogen
hepatocyte growth factor-like protein isoform X2
hepatocyte growth factor-like protein isoform X1
Hepatocyte growth factor
coagulation factor XI
plasminogen isoform X3
Hepatocyte growth factor
Uncharacterized protein
Uncharacterized protein
Uncharacterized protein
Uncharacterized protein
Uncharacterized protein
Uncharacterized protein
Uncharacterized protein
Uncharacterized protein
Uncharacterized protein
Uncharacterized protein
G-type lectin S-receptor-like serine/threonine-protein kinase RLK1
G-type lectin S-receptor-like serine/threonine-protein kinase RLK1
Receptor-like serine/threonine-protein kinase SD1-8
G-type lectin S-receptor-like serine/threonine-protein kinase
Receptor-like serine/threonine-protein kinase SD1-8
Receptor-like serine/threonine-protein kinase SD1-8
G-type lectin S-receptor-like serine/threonine-protein kinase
G-type lectin S-receptor-like serine/threonine-protein kinase RLK1
G-type lectin S-receptor-like serine/threonine-protein kinase SD3-1
Receptor-like serine/threonine-protein kinase SD1-6
Receptor protein kinase ZmPK1
G-type lectin S-receptor-like serine/threonine-protein kinase RLK1
G-type lectin S-receptor-like serine/threonine-protein kinase B120

Receptor-like serine/threonine-protein kinase SD1-8
G-type lectin S-receptor-like serine/threonine-protein kinase B120
Receptor-like serine/threonine-protein kinase SD1-8
PAN domain-containing protein
G-type lectin S-receptor-like serine/threonine-protein kinase SD2-2
G-type lectin S-receptor-like serine/threonine-protein kinase
G-type lectin S-receptor-like serine/threonine-protein kinase B120
G-type lectin S-receptor-like serine/threonine-protein kinase
G-type lectin S-receptor-like serine/threonine-protein kinase
G-type lectin S-receptor-like serine/threonine-protein kinase B120
G-type lectin S-receptor-like serine/threonine-protein kinase
G-type lectin S-receptor-like serine/threonine-protein kinase B120
G-type lectin S-receptor-like serine/threonine-protein kinase SD2-5
G-type lectin S-receptor-like serine/threonine-protein kinase B120
G-type lectin S-receptor-like serine/threonine-protein kinase RLK1
G-type lectin S-receptor-like serine/threonine-protein kinase
Receptor-like serine/threonine-protein kinase SD1-8
Uncharacterized protein
Uncharacterized protein
Uncharacterized protein
Uncharacterized protein
Uncharacterized protein
Uncharacterized protein
Uncharacterized protein
Uncharacterized protein
Uncharacterized protein (Fragment)
Uncharacterized protein (Fragment)
Uncharacterized protein (Fragment)
Uncharacterized protein (Fragment)
Uncharacterized protein (Fragment)
Uncharacterized protein (Fragment)
Uncharacterized protein (Fragment)
Uncharacterized protein
Uncharacterized protein
Uncharacterized protein
SH3 domain-containing protein,putative peptidoglycan binding protein
Uncharacterized protein
Uncharacterized protein
Uncharacterized protein
Uncharacterized protein
Uncharacterized protein

Uncharacterized protein
Uncharacterized protein
Uncharacterized protein
Uncharacterized protein
Uncharacterized protein
Uncharacterized protein
Uncharacterized protein
Uncharacterized protein
PAN domain protein
PAN domain protein
Uncharacterized protein
Uncharacterized protein
Alpha-2-macroglobulin domain protein
Uncharacterized protein
PAN domain protein
Uncharacterized protein
Uncharacterized protein
Uncharacterized protein
Uncharacterized protein
Uncharacterized protein
Uncharacterized protein
PAN domain-containing protein
LOW QUALITY PROTEIN: uncharacterized protein LOC113471719
uncharacterized protein LOC103517195
uncharacterized protein LOC103516866
uncharacterized protein LOC103508935
uncharacterized protein LOC113470503
uncharacterized protein LOC103510819
Putative galactose oxidase
Uncharacterized protein
Uncharacterized protein
Putative hydrophobin-like protein
Uncharacterized protein
Putative galactose oxidase
Uncharacterized protein
Uncharacterized protein
Kelch domain-containing protein
Kelch domain-containing protein
Uncharacterized protein
Uncharacterized protein
Uncharacterized protein

Kelch domain-containing protein
Receptor-like serine/threonine-protein kinase SD1-8
Putative receptor protein kinase ZmPK1
G-type lectin S-receptor-like serine/threonine-protein kinase B120
G-type lectin S-receptor-like serine/threonine-protein kinase
G-type lectin S-receptor-like serine/threonine-protein kinase
G-type lectin S-receptor-like serine/threonine-protein kinase
G-type lectin S-receptor-like serine/threonine-protein kinase
G-type lectin S-receptor-like serine/threonine-protein kinase
Receptor-like serine/threonine-protein kinase SD1-8 (Fragment)
G-type lectin S-receptor-like serine/threonine-protein kinase
Putative receptor-like protein kinase
G-type lectin S-receptor-like serine/threonine-protein kinase
Receptor-like serine/threonine-protein kinase SD1-8
Receptor-like serine/threonine-protein kinase SD1-8 (Fragment)
Receptor-like serine/threonine-protein kinase SD1-8
G-type lectin S-receptor-like serine/threonine-protein kinase
PAN domain-containing protein
PAN domain-containing protein
Receptor-like serine/threonine-protein kinase SD1-8
Putative G-type lectin S-receptor-like serine/threonine-protein kinase
Receptor-like serine/threonine-protein kinase SD1-8
Receptor-like serine/threonine-protein kinase SD1-8
Putative G-type lectin S-receptor-like serine/threonine-protein kinase
G-type lectin S-receptor-like serine/threonine-protein kinase
Receptor-like serine/threonine-protein kinase SD1-8
G-type lectin S-receptor-like serine/threonine-protein kinase
S-locus-specific glycoprotein S13
Putative receptor protein kinase ZmPK1
Receptor-like serine/threonine-protein kinase SD1-8
G-type lectin S-receptor-like serine/threonine-protein kinase
G-type lectin S-receptor-like serine/threonine-protein kinase
Receptor-like serine/threonine-protein kinase SD1-8
G-type lectin S-receptor-like serine/threonine-protein kinase
Putative receptor protein kinase ZmPK1
Receptor-like serine/threonine-protein kinase SD1-8
G-type lectin S-receptor-like serine/threonine-protein kinase (Fragment)
G-type lectin S-receptor-like serine/threonine-protein kinase
Cysteine-rich receptor-like protein kinase 10
Receptor-like serine/threonine-protein kinase SD1-8
G-type lectin S-receptor-like serine/threonine-protein kinase SD2-5

G-type lectin S-receptor-like serine/threonine-protein kinase
Receptor-like serine/threonine-protein kinase SD1-8
Putative receptor protein kinase ZmPK1
G-type lectin S-receptor-like serine/threonine-protein kinase
Putative receptor protein kinase ZmPK1
Uncharacterized protein
Putative receptor protein kinase ZmPK1
G-type lectin S-receptor-like serine/threonine-protein kinase B120
Putative receptor protein kinase ZmPK1
PAN domain-containing protein
G-type lectin S-receptor-like serine/threonine-protein kinase
G-type lectin S-receptor-like serine/threonine-protein kinase
Receptor-like serine/threonine-protein kinase SD1-8 (Fragment)
Putative receptor protein kinase ZmPK1
G-type lectin S-receptor-like serine/threonine-protein kinase RKS1
Receptor-like serine/threonine-protein kinase SD1-8
G-type lectin S-receptor-like serine/threonine-protein kinase
Uncharacterized protein (Fragment)
G-type lectin S-receptor-like serine/threonine-protein kinase
G-type lectin S-receptor-like serine/threonine-protein kinase
Uncharacterized protein
Putative inactive G-type lectin S-receptor-like serine/threonine-protein kinase SRK
PAN domain protein
PAN domain protein
PAN domain protein
Zona pellucida-like domain protein
PAN domain protein
PAN domain protein
PAN domain protein
Uncharacterized protein
PAN domain protein
PAN domain protein
PAN domain protein
PAN domain protein
PAN domain protein
PAN domain protein
PAN domain protein
PAN domain protein
PAN domain protein
PAN domain protein
PAN domain protein

PAN domain protein
PAN domain protein
Uncharacterized protein
Uncharacterized protein
Uncharacterized protein
Uncharacterized protein
Uncharacterized protein
Catalytic
Uncharacterized protein
Uncharacterized protein
Sequence-specific DNA binding RNA polymerase II transcription factor
Uncharacterized protein
Uncharacterized protein
Zona pellucida-like domain containing protein 3 (Fragment)
Uncharacterized protein (Fragment)
Uncharacterized protein
Uncharacterized protein (Fragment)
Uncharacterized protein (Fragment)
Uncharacterized protein
Uncharacterized protein
Uncharacterized protein (Fragment)
Uncharacterized protein
Galactose oxidase
Galactose oxidase
Apple protein
Uncharacterized protein
Copper radical oxidase
Carbohydrate-binding wsc
Uncharacterized protein
Kelch domain-containing protein
Uncharacterized protein
Putative kelch domain-containing protein
Putative carbohydrate-binding wsc
Putative Apple
Uncharacterized protein
Putative fungistatic metabolite (Fragment)
Uncharacterized protein
Uncharacterized protein
Uncharacterized protein
Uncharacterized protein
Uncharacterized protein

Uncharacterized protein
Uncharacterized protein
Uncharacterized protein
Uncharacterized protein
Uncharacterized protein
Uncharacterized protein
Uncharacterized protein
Uncharacterized protein
Uncharacterized protein
Uncharacterized protein
Uncharacterized protein
Uncharacterized protein
Uncharacterized protein
Uncharacterized protein
Uncharacterized protein
Uncharacterized protein
Uncharacterized protein
Uncharacterized protein
Uncharacterized protein
Uncharacterized protein
Uncharacterized protein
Uncharacterized protein
Uncharacterized protein
Uncharacterized protein
Uncharacterized protein
Uncharacterized protein
Uncharacterized protein
Uncharacterized protein
Uncharacterized protein
Uncharacterized protein
Uncharacterized protein
Uncharacterized protein
Uncharacterized protein
Uncharacterized protein
Uncharacterized protein
Uncharacterized protein
Uncharacterized protein
Uncharacterized protein (Fragment)
Uncharacterized protein
Uncharacterized protein
Uncharacterized protein

Uncharacterized protein
Uncharacterized protein
Uncharacterized protein
Uncharacterized protein
Uncharacterized protein
Uncharacterized protein
Uncharacterized protein
Uncharacterized protein
Uncharacterized protein
Uncharacterized protein
S-locus-related I (Fragment)
S-locus-related I (Fragment)
Hepatocyte growth factor
hepatocyte growth factor isoform X2
hepatocyte growth factor-like protein isoform X2
Hepatocyte growth factor
Plasminogen
hepatocyte growth factor-like protein isoform X1
plasma kallikrein
coagulation factor XI
Uncharacterized protein
S-locus receptor kinase (Fragment)
S-locus receptor kinase (Fragment)
S-locus receptor kinase (Fragment)
S-locus receptor kinase (Fragment)
S-locus receptor kinase (Fragment)
S-locus receptor kinase (Fragment)
Uncharacterized protein
Uncharacterized protein
Uncharacterized protein
G-type lectin S-receptor-like serine/threonine-protein kinase
Uncharacterized protein
Uncharacterized protein
G-type lectin S-receptor-like serine/threonine-protein kinase
Uncharacterized protein
Uncharacterized protein
PAN domain-containing protein-like
Uncharacterized protein
Receptor kinase 3 isoform 1
Uncharacterized protein
Uncharacterized protein

G-type lectin S-receptor-like serine/threonine-protein kinase-like
Uncharacterized protein
Uncharacterized protein (Fragment)
Uncharacterized protein
Uncharacterized protein (Fragment)
Uncharacterized protein
Uncharacterized protein
Uncharacterized protein
Uncharacterized protein
Uncharacterized protein
Uncharacterized protein
Uncharacterized protein
Uncharacterized protein
Uncharacterized protein
Uncharacterized protein
Uncharacterized protein
Uncharacterized protein
Uncharacterized protein
Uncharacterized protein
Uncharacterized protein
Uncharacterized protein
Uncharacterized protein
Uncharacterized protein
Uncharacterized protein
Uncharacterized protein
Uncharacterized protein
Glycosyltransferase family 31
Uncharacterized protein
Uncharacterized protein
Uncharacterized protein
Uncharacterized protein, isoform A
Uncharacterized protein
Uncharacterized protein, isoform B
Uncharacterized protein
Uncharacterized protein
Uncharacterized protein
Nyo
Mey
CG17111
CG42326
Tyn

CG6495
Neo
NompA
GG11812
Uncharacterized protein
Uncharacterized protein, isoform C
Uncharacterized protein, isoform B
Uncharacterized protein, isoform A
GG11699
GG11810
Uncharacterized protein
GG12464
uncharacterized protein LOC108091652
uncharacterized protein LOC108090568 isoform X2
uncharacterized protein LOC108100778
uncharacterized protein LOC108086442 isoform X2
uncharacterized protein LOC108090568 isoform X1
uncharacterized protein LOC108086442 isoform X4
uncharacterized protein LOC108086442 isoform X3
uncharacterized protein LOC108086442 isoform X1
uncharacterized protein LOC108092855
uncharacterized protein LOC108100781
uncharacterized protein LOC108088395
GH23882
GH18127
GH22147
GH19913
GH19340
GH24267
GH22146
GH19351
GH19413
Uncharacterized protein
Blast:Cuticlin-1
Uncharacterized protein
Uncharacterized protein
Uncharacterized protein
Uncharacterized protein
Uncharacterized protein
Blast:Ubiquitin-like domain-containing CTD phosphatase 1
Uncharacterized protein

Uncharacterized protein
Uncharacterized protein
No-mechanoreceptor potential A short isoform
RE41896p (Fragment)
No mechanoreceptor potential A, isoform F
Uncharacterized protein, isoform E
No-mechanoreceptor potential A long isoform
RE37929p
Uncharacterized protein, isoform D
IP14862p
FI03277p (Fragment)
SP71
IP22030p
RE59252p
IP16971p
RE15579p
FI17836p1
GH22837p
FI18240p1
Morpheyus, isoform A
No mechanoreceptor potential A, isoform C
No mechanoreceptor potential A, isoform A
Uncharacterized protein
Uncharacterized protein
Uncharacterized protein, isoform B
Uncharacterized protein
Uncharacterized protein, isoform A
Uncharacterized protein
Uncharacterized protein
Uncharacterized protein
Uncharacterized protein
Uncharacterized protein
GL11022
GL13829
GL27254
GL27256
GL17345
GL11021
GL14072
GL13362
Uncharacterized protein

Uncharacterized protein
Uncharacterized protein
Uncharacterized protein
Uncharacterized protein, isoform B
Uncharacterized protein, isoform A
Uncharacterized protein, isoform A
GM20678
GM11116
GM20446
GM20679
GM12828
GM16429
GM12947
GM23595
GD10156
Uncharacterized protein, isoform B
Uncharacterized protein, isoform B
GD21582
GD25914
GD15272
GD21581
GD21471
Uncharacterized protein, isoform C
GD16493
Uncharacterized protein, isoform C
Uncharacterized protein
Uncharacterized protein
Uncharacterized protein, isoform B
Uncharacterized protein, isoform A
Uncharacterized protein
Uncharacterized protein
Uncharacterized protein
Uncharacterized protein
Uncharacterized protein
Uncharacterized protein
Uncharacterized protein
Uncharacterized protein
Uncharacterized protein
Uncharacterized protein
Uncharacterized protein
Uncharacterized protein

Uncharacterized protein
Uncharacterized protein
Uncharacterized protein
Uncharacterized protein, isoform E
Uncharacterized protein, isoform A
Uncharacterized protein, isoform B
Uncharacterized protein
Uncharacterized protein
Uncharacterized protein, isoform D
Uncharacterized protein
Uncharacterized protein, isoform C
Uncharacterized protein, isoform C
Uncharacterized protein, isoform B
Plasma kallikrein (Fragment)
Hepatocyte growth factor-like (Fragment)
Uncharacterized protein
Uncharacterized protein
Uncharacterized protein
Neurotrypsin
Uncharacterized protein (Fragment)
Uncharacterized protein
Cuticlin-1
Uncharacterized protein
Ras-related protein RabJ (Fragment)
Fibrinogen/ficolin like FReD protein (Fragment)
Uncharacterized protein
Seven cysteines N terminal
Collagen alpha-3(VI) chain
Uncharacterized protein
Collagen alpha 1VII chain
Apple
Low density lipoprotein receptor class A cysteine rich
Apple
Collagen alpha 1VII chain
Low density lipoprotein receptor class A cysteine rich
Seven cysteines N terminal
Uncharacterized protein
Uncharacterized protein
Alpha-L-fucosidase, family GH95
Hepatocyte growth factor (Fragment)
Plasma kallikrein (Fragment)

Plasminogen (Fragment)
Plasminogen (Fragment)
Uncharacterized protein
Uncharacterized protein (Fragment)
PAN domain-containing protein, putative (Fragment)
Uncharacterized protein
Uncharacterized protein
PAN domain-containing protein, putative (Fragment)
Uncharacterized protein
PAN domain-containing protein, putative
Uncharacterized protein
Micronemal protein MIC4, related
PAN domain-containing protein, putative
Uncharacterized protein
PAN domain-containing protein, putative
Microneme protein 5 (Fragment)
Uncharacterized protein
PAN domain-containing protein, putative
Uncharacterized protein
Uncharacterized protein
PAN domain-containing protein, putative (Fragment)
PAN domain-containing protein, putative
Uncharacterized protein
PAN domain-containing protein, putative
PAN domain-containing protein, related
Uncharacterized protein (Fragment)
Uncharacterized protein
Microneme protein 5 (Fragment)
Uncharacterized protein
Uncharacterized protein
PAN domain-containing protein, putative
Microneme protein 5 (Fragment)
PAN domain-containing protein, putative
PAN domain-containing protein, related
Uncharacterized protein (Fragment)
Microneme protein 5 (Fragment)
Uncharacterized protein (Fragment)
PAN domain-containing protein, putative
PAN domain-containing protein, putative (Fragment)
Uncharacterized protein
Micronemal protein MIC4, related

Uncharacterized protein
Uncharacterized protein
Uncharacterized protein
Uncharacterized protein
Uncharacterized protein
Uncharacterized protein
Uncharacterized protein
Uncharacterized protein
Uncharacterized protein
Uncharacterized protein
Uncharacterized protein
Uncharacterized protein
Uncharacterized protein
Uncharacterized protein
Uncharacterized protein
Uncharacterized protein
Chitin deacetylase
Uncharacterized protein
Uncharacterized protein
Uncharacterized protein
Uncharacterized protein
Uncharacterized protein
Uncharacterized protein
Guanine nucleotide-binding protein subunit beta
Uncharacterized protein
Uncharacterized protein
Uncharacterized protein
WSC domain-containing protein 2
Uncharacterized protein
Uncharacterized protein
Uncharacterized protein
Protein TOS1
Uncharacterized protein
Kremen protein 1
Uncharacterized protein
Uncharacterized protein
Nucleoporin nup146
Uncharacterized protein
Uncharacterized protein
Uncharacterized protein (Fragment)
Uncharacterized protein

Uncharacterized protein
Uncharacterized protein
Uncharacterized protein
Uncharacterized protein (Fragment)
Uncharacterized protein
Uncharacterized protein
Uncharacterized protein
Uncharacterized protein
Uncharacterized protein
Uncharacterized protein
Uncharacterized protein
Uncharacterized protein
Uncharacterized protein
Uncharacterized protein
Uncharacterized protein (Fragment)
Uncharacterized protein
Uncharacterized protein
Uncharacterized protein
Uncharacterized protein
Uncharacterized protein
Uncharacterized protein
Uncharacterized protein
Uncharacterized protein
Uncharacterized protein
Uncharacterized protein
Uncharacterized protein
Uncharacterized protein
Uncharacterized protein
Putative pectin acetylesterase
Uncharacterized protein
Uncharacterized protein (Fragment)
Uncharacterized protein
Uncharacterized protein
Uncharacterized protein
Uncharacterized protein
Uncharacterized protein
Uncharacterized protein
Uncharacterized protein
Uncharacterized protein
Uncharacterized protein
Uncharacterized protein (Fragment)

Uncharacterized protein
Uncharacterized protein
Uncharacterized protein
Uncharacterized protein
coagulation factor XI isoform X1
plasma kallikrein
coagulation factor XI isoform X3
Plasminogen
Hepatocyte growth factor
Hepatocyte growth factor
hepatocyte growth factor-like protein
coagulation factor XI isoform X2
Uncharacterized protein
Uncharacterized protein
Uncharacterized protein
Uncharacterized protein
Uncharacterized protein
Uncharacterized protein
Uncharacterized protein
Uncharacterized protein
Uncharacterized protein
Uncharacterized protein
Uncharacterized protein
Uncharacterized protein
Uncharacterized protein
Uncharacterized protein
Uncharacterized protein
Uncharacterized protein
Uncharacterized protein
Uncharacterized protein
Uncharacterized protein
Uncharacterized protein
Uncharacterized protein
Uncharacterized protein
Uncharacterized protein
Uncharacterized protein
Uncharacterized protein
Uncharacterized protein
Uncharacterized protein
Uncharacterized protein
Uncharacterized protein (Fragment)

Uncharacterized protein
Uncharacterized protein (Fragment)
Uncharacterized protein
Uncharacterized protein
Uncharacterized protein
Uncharacterized protein
Uncharacterized protein
Uncharacterized protein
Uncharacterized protein
Uncharacterized protein
Uncharacterized protein
Uncharacterized protein (Fragment)
Uncharacterized protein
Uncharacterized protein
Uncharacterized protein
Uncharacterized protein
Uncharacterized protein
Uncharacterized protein
Uncharacterized protein
Uncharacterized protein
Uncharacterized protein
Uncharacterized protein
Uncharacterized protein
Uncharacterized protein
Uncharacterized protein
Uncharacterized protein
Uncharacterized protein
Uncharacterized protein
Uncharacterized protein
Uncharacterized protein
Uncharacterized protein
Uncharacterized protein
Uncharacterized protein
Uncharacterized protein
Uncharacterized protein
Uncharacterized protein
Uncharacterized protein
Putative inner membrane lipoprotein
Membrane protein
Uncharacterized protein
Uncharacterized protein

Uncharacterized protein (Fragment)
Uncharacterized protein
Uncharacterized protein
PAN domain protein
Uncharacterized protein
PAN domain-containing protein
Uncharacterized protein
Uncharacterized protein
Uncharacterized protein
Putative lipoprotein YfhM
Uncharacterized protein
Uncharacterized protein
Uncharacterized protein
Uncharacterized protein
Uncharacterized protein
Uncharacterized protein
Uncharacterized protein
Uncharacterized protein
Uncharacterized protein
Uncharacterized protein
Uncharacterized protein
Uncharacterized protein
Uncharacterized protein
Uncharacterized protein
Uncharacterized protein
Uncharacterized protein
Uncharacterized protein
Uncharacterized protein
Uncharacterized protein
Uncharacterized protein
Uncharacterized protein
Uncharacterized protein
Uncharacterized protein
Uncharacterized protein
Uncharacterized protein
Uncharacterized protein
Uncharacterized protein
Uncharacterized protein
Uncharacterized protein
Uncharacterized protein
Uncharacterized protein

Uncharacterized protein
Uncharacterized protein
Uncharacterized protein
Uncharacterized protein
Uncharacterized protein
Uncharacterized protein
Uncharacterized protein
Uncharacterized protein
Uncharacterized protein
Uncharacterized protein
Plasminogen
Plasminogen
Hepatocyte growth factor
Kallikrein B1
Macrophage stimulating 1
Macrophage stimulating 1
Macrophage stimulating 1
Coagulation factor XI
Plasminogen
Coagulation factor XI
hepatocyte growth factor isoform X3
coagulation factor XI isoform X1
Hepatocyte growth factor
hepatocyte growth factor-like protein
plasma kallikrein isoform X2
coagulation factor XI isoform X2
uncharacterized protein LOC103112208
Hepatocyte growth factor
plasma kallikrein isoform X1
Plasminogen
S-locus-related I (Fragment)
S-locus-related I (Fragment)
S-locus-related I (Fragment)
S-locus-related I (Fragment)
SLR1 protein (Fragment)
Uncharacterized protein (Fragment)
Uncharacterized protein
Uncharacterized protein
Uncharacterized protein (Fragment)
Uncharacterized protein
Uncharacterized protein

Uncharacterized protein
Uncharacterized protein
Uncharacterized protein (Fragment)
Uncharacterized protein
Uncharacterized protein
Uncharacterized protein
Uncharacterized protein
Uncharacterized protein
Uncharacterized protein
Uncharacterized protein
Uncharacterized protein
Uncharacterized protein
Uncharacterized protein
Uncharacterized protein (Fragment)
Uncharacterized protein
Uncharacterized protein
Uncharacterized protein
Uncharacterized protein
Uncharacterized protein (Fragment)
Uncharacterized protein
Uncharacterized protein
Uncharacterized protein
Uncharacterized protein
Uncharacterized protein
Uncharacterized protein
Uncharacterized protein (Fragment)
Uncharacterized protein
Uncharacterized protein
Uncharacterized protein
Uncharacterized protein
Uncharacterized protein (Fragment)
Uncharacterized protein (Fragment)
Uncharacterized protein
Uncharacterized protein
Uncharacterized protein
Uncharacterized protein (Fragment)
Uncharacterized protein
Uncharacterized protein (Fragment)
Uncharacterized protein (Fragment)
Uncharacterized protein
Uncharacterized protein (Fragment)

Uncharacterized protein
Uncharacterized protein
Uncharacterized protein
Hepatocyte growth factor-like protein (Fragment)
Uncharacterized protein
Plasminogen
Hepatocyte growth factor
Hepatocyte growth factor
Uncharacterized protein
Uncharacterized protein
Plasminogen
Uncharacterized protein
Uncharacterized protein
Uncharacterized protein
Uncharacterized protein
Uncharacterized protein
Uncharacterized protein
Uncharacterized protein
Uncharacterized protein
Uncharacterized protein
Uncharacterized protein
Uncharacterized protein
Uncharacterized protein
Uncharacterized protein
Uncharacterized protein (Fragment)
Uncharacterized protein
Uncharacterized protein
Uncharacterized protein
Uncharacterized protein
Uncharacterized protein
Uncharacterized protein
Uncharacterized protein (Fragment)
Uncharacterized protein
Uncharacterized protein
Uncharacterized protein
Uncharacterized protein
Uncharacterized protein
Uncharacterized protein
Uncharacterized protein (Fragment)
Uncharacterized protein
Uncharacterized protein

Uncharacterized protein
Uncharacterized protein
Uncharacterized protein
Uncharacterized protein (Fragment)
Uncharacterized protein
Uncharacterized protein
Uncharacterized protein
Uncharacterized protein
Uncharacterized protein
Uncharacterized protein
Uncharacterized protein
Uncharacterized protein
Uncharacterized protein (Fragment)
Uncharacterized protein
Uncharacterized protein
Uncharacterized protein
Uncharacterized protein
Uncharacterized protein
Uncharacterized protein
Uncharacterized protein
Uncharacterized protein
Uncharacterized protein
Uncharacterized protein (Fragment)
Uncharacterized protein
Uncharacterized protein
Uncharacterized protein
Uncharacterized protein
Uncharacterized protein
Uncharacterized protein (Fragment)
Uncharacterized protein
Uncharacterized protein
Uncharacterized protein
Uncharacterized protein (Fragment)
Uncharacterized protein
Uncharacterized protein
Uncharacterized protein
Uncharacterized protein
Uncharacterized protein
Uncharacterized protein (Fragment)
Uncharacterized protein
Uncharacterized protein

Uncharacterized protein
Uncharacterized protein
Uncharacterized protein
Uncharacterized protein
Uncharacterized protein
Uncharacterized protein
Uncharacterized protein
Uncharacterized protein
Uncharacterized protein
Uncharacterized protein
Uncharacterized protein
Uncharacterized protein
Uncharacterized protein (Fragment)
Uncharacterized protein (Fragment)
Uncharacterized protein (Fragment)
Uncharacterized protein
Uncharacterized protein
Uncharacterized protein
Uncharacterized protein
Uncharacterized protein
Uncharacterized protein
Uncharacterized protein (Fragment)
Uncharacterized protein
Uncharacterized protein
Uncharacterized protein (Fragment)
Uncharacterized protein
Uncharacterized protein
Uncharacterized protein
Uncharacterized protein
Uncharacterized protein (Fragment)
Uncharacterized protein
Uncharacterized protein
Uncharacterized protein
Uncharacterized protein
Uncharacterized protein
Uncharacterized protein
Uncharacterized protein
Uncharacterized protein
Uncharacterized protein
Uncharacterized protein
Uncharacterized protein (Fragment)

Uncharacterized protein
Uncharacterized protein
Uncharacterized protein
Uncharacterized protein
Uncharacterized protein
Uncharacterized protein
Uncharacterized protein
Uncharacterized protein (Fragment)
Uncharacterized protein
Uncharacterized protein
Uncharacterized protein
Uncharacterized protein (Fragment)
Uncharacterized protein
Uncharacterized protein
Uncharacterized protein
Uncharacterized protein
Uncharacterized protein
Uncharacterized protein
Uncharacterized protein (Fragment)
Uncharacterized protein
Uncharacterized protein
Uncharacterized protein
Uncharacterized protein
Uncharacterized protein
Uncharacterized protein
Uncharacterized protein
Uncharacterized protein
Uncharacterized protein
Uncharacterized protein
Uncharacterized protein
Uncharacterized protein
Uncharacterized protein
Uncharacterized protein (Fragment)
Uncharacterized protein
Uncharacterized protein
Uncharacterized protein
Uncharacterized protein
Uncharacterized protein
Uncharacterized protein (Fragment)
Uncharacterized protein
Uncharacterized protein

Uncharacterized protein
Uncharacterized protein
Uncharacterized protein
Uncharacterized protein
Uncharacterized protein
Uncharacterized protein
Uncharacterized protein
Uncharacterized protein (Fragment)
Uncharacterized protein
Uncharacterized protein
Uncharacterized protein
Uncharacterized protein (Fragment)
Uncharacterized protein
Uncharacterized protein
Uncharacterized protein
Uncharacterized protein
Uncharacterized protein
Uncharacterized protein
Uncharacterized protein
Uncharacterized protein
Uncharacterized protein
Uncharacterized protein
Uncharacterized protein
Uncharacterized protein (Fragment)
Uncharacterized protein (Fragment)
Uncharacterized protein
Uncharacterized protein
Uncharacterized protein
Uncharacterized protein
Uncharacterized protein
Uncharacterized protein
Uncharacterized protein
Uncharacterized protein
Uncharacterized protein
Uncharacterized protein
Uncharacterized protein
Uncharacterized protein
Uncharacterized protein
Uncharacterized protein
Uncharacterized protein
Uncharacterized protein

Uncharacterized protein
Uncharacterized protein
Uncharacterized protein
Uncharacterized protein
Uncharacterized protein
Uncharacterized protein
Uncharacterized protein
Uncharacterized protein
Uncharacterized protein
Uncharacterized protein
Uncharacterized protein
Uncharacterized protein
Uncharacterized protein
Uncharacterized protein
Uncharacterized protein
Uncharacterized protein (Fragment)
Uncharacterized protein
Uncharacterized protein
Uncharacterized protein
Uncharacterized protein
Uncharacterized protein
Uncharacterized protein (Fragment)
Uncharacterized protein (Fragment)
Uncharacterized protein
Uncharacterized protein
Uncharacterized protein
Uncharacterized protein
Uncharacterized protein
Uncharacterized protein
Uncharacterized protein
Uncharacterized protein
Uncharacterized protein
Uncharacterized protein
Uncharacterized protein
Uncharacterized protein
Uncharacterized protein
Uncharacterized protein (Fragment)
Uncharacterized protein
Uncharacterized protein
Uncharacterized protein
Uncharacterized protein

Uncharacterized protein
Uncharacterized protein
Uncharacterized protein
Uncharacterized protein
Uncharacterized protein
Uncharacterized protein
Uncharacterized protein (Fragment)
Uncharacterized protein
Uncharacterized protein
Uncharacterized protein
Uncharacterized protein (Fragment)
Uncharacterized protein
Uncharacterized protein
Uncharacterized protein (Fragment)
Uncharacterized protein
Uncharacterized protein
Uncharacterized protein
Uncharacterized protein
Uncharacterized protein
Uncharacterized protein
Uncharacterized protein
Uncharacterized protein
Uncharacterized protein
Uncharacterized protein
Uncharacterized protein
Uncharacterized protein (Fragment)
Uncharacterized protein
Uncharacterized protein
Uncharacterized protein
Uncharacterized protein
Uncharacterized protein
Uncharacterized protein
Uncharacterized protein
Uncharacterized protein
Uncharacterized protein
Uncharacterized protein
Uncharacterized protein (Fragment)
Uncharacterized protein
Uncharacterized protein
Uncharacterized protein
Uncharacterized protein

Uncharacterized protein
Uncharacterized protein
Uncharacterized protein
Uncharacterized protein (Fragment)
Uncharacterized protein
Uncharacterized protein (Fragment)
Uncharacterized protein
Uncharacterized protein
Uncharacterized protein
Uncharacterized protein
Uncharacterized protein
Uncharacterized protein (Fragment)
Uncharacterized protein
Uncharacterized protein
Uncharacterized protein
Uncharacterized protein
Uncharacterized protein
Uncharacterized protein (Fragment)
Uncharacterized protein (Fragment)
Uncharacterized protein
Uncharacterized protein
Uncharacterized protein
Uncharacterized protein
Ras-related protein RabJ
Deleted in malignant brain tumors 1 protein
Cuticlin-1
Uncharacterized protein (Fragment)
Zona pellucida-like and PAN domain containing protein
PAN domain containing protein (Fragment)
Uncharacterized protein
Uncharacterized protein (Fragment)
Uncharacterized protein
Neyo-like protein (Fragment)
Zona pellucida-like domain containing protein
Zona pellucida and PAN domain containing protein (Fragment)
Uncharacterized protein
Uncharacterized protein
Hepatocyte growth factor (Fragment)
Uncharacterized protein
Uncharacterized protein
Uncharacterized protein

Uncharacterized protein
Uncharacterized protein
Uncharacterized protein
Uncharacterized protein (Fragment)
Uncharacterized protein
Uncharacterized protein
Uncharacterized protein
Uncharacterized protein
Uncharacterized protein (Fragment)
Uncharacterized protein
Uncharacterized protein
Uncharacterized protein
Uncharacterized protein (Fragment)
Uncharacterized protein
Uncharacterized protein
Uncharacterized protein
Uncharacterized protein (Fragment)
Uncharacterized protein
Uncharacterized protein
Uncharacterized protein
Uncharacterized protein
Uncharacterized protein
Uncharacterized protein
Uncharacterized protein
Uncharacterized protein
Uncharacterized protein (Fragment)
Uncharacterized protein
Uncharacterized protein
Uncharacterized protein
Uncharacterized protein
Uncharacterized protein
Uncharacterized protein
Uncharacterized protein
Uncharacterized protein
Uncharacterized protein
Uncharacterized protein
Uncharacterized protein
Uncharacterized protein
Uncharacterized protein
Uncharacterized protein
Uncharacterized protein

Uncharacterized protein
Uncharacterized protein
Uncharacterized protein
Uncharacterized protein
Uncharacterized protein
Uncharacterized protein
Uncharacterized protein
Uncharacterized protein
Uncharacterized protein
Uncharacterized protein
Uncharacterized protein
Uncharacterized protein
Uncharacterized protein
Uncharacterized protein
Uncharacterized protein
Uncharacterized protein
Uncharacterized protein
Uncharacterized protein
Uncharacterized protein
Uncharacterized protein
Uncharacterized protein
Uncharacterized protein
Uncharacterized protein
Uncharacterized protein
Uncharacterized protein
Uncharacterized protein
Uncharacterized protein
Uncharacterized protein
Uncharacterized protein
Uncharacterized protein
Uncharacterized protein
Uncharacterized protein
Uncharacterized protein
Uncharacterized protein
Uncharacterized protein
Uncharacterized protein
Uncharacterized protein
Uncharacterized protein
Uncharacterized protein
Uncharacterized protein
Uncharacterized protein

Uncharacterized protein
Uncharacterized protein
Uncharacterized protein
Uncharacterized protein
Uncharacterized protein
Uncharacterized protein
Uncharacterized protein
Uncharacterized protein
Uncharacterized protein
Uncharacterized protein
Uncharacterized protein
Uncharacterized protein
Uncharacterized protein
Uncharacterized protein
Uncharacterized protein
Uncharacterized protein
Uncharacterized protein
Uncharacterized protein
Uncharacterized protein
Uncharacterized protein
Uncharacterized protein
Uncharacterized protein
Uncharacterized protein
Uncharacterized protein
Uncharacterized protein
Uncharacterized protein
Uncharacterized protein
Uncharacterized protein
Uncharacterized protein
Uncharacterized protein
Uncharacterized protein
Uncharacterized protein
Uncharacterized protein
Uncharacterized protein
Uncharacterized protein
Uncharacterized protein
Uncharacterized protein
Uncharacterized protein
Uncharacterized protein
Uncharacterized protein
Uncharacterized protein

Uncharacterized protein
Uncharacterized protein
Uncharacterized protein
Uncharacterized protein
Uncharacterized protein
Uncharacterized protein
Uncharacterized protein
Uncharacterized protein
Uncharacterized protein
Uncharacterized protein
Uncharacterized protein
Uncharacterized protein
Uncharacterized protein
Uncharacterized protein
Uncharacterized protein
Uncharacterized protein
Uncharacterized protein
Uncharacterized protein
Uncharacterized protein
Uncharacterized protein
Uncharacterized protein
Uncharacterized protein
Uncharacterized protein
Uncharacterized protein
Uncharacterized protein
Uncharacterized protein
Uncharacterized protein
Uncharacterized protein
Uncharacterized protein
Uncharacterized protein
Uncharacterized protein
Uncharacterized protein
Uncharacterized protein
Uncharacterized protein
Uncharacterized protein
Uncharacterized protein
Uncharacterized protein
Uncharacterized protein
Uncharacterized protein
Uncharacterized protein
Uncharacterized protein

Uncharacterized protein
Uncharacterized protein
Uncharacterized protein
Uncharacterized protein
Uncharacterized protein
Uncharacterized protein
Uncharacterized protein
Uncharacterized protein
Uncharacterized protein
Uncharacterized protein
Uncharacterized protein
Uncharacterized protein
Uncharacterized protein
Uncharacterized protein
Uncharacterized protein
Uncharacterized protein
Uncharacterized protein
Uncharacterized protein
Uncharacterized protein
Uncharacterized protein
Uncharacterized protein
Uncharacterized protein
Uncharacterized protein
Uncharacterized protein
Uncharacterized protein
Uncharacterized protein
Uncharacterized protein
Uncharacterized protein
Uncharacterized protein
Uncharacterized protein
Uncharacterized protein
Uncharacterized protein
Uncharacterized protein
Uncharacterized protein
Uncharacterized protein
Uncharacterized protein
Uncharacterized protein
Uncharacterized protein
Uncharacterized protein
Uncharacterized protein
Uncharacterized protein

Uncharacterized protein
Uncharacterized protein
Uncharacterized protein
Uncharacterized protein
Uncharacterized protein
Uncharacterized protein
Uncharacterized protein
Uncharacterized protein
Uncharacterized protein
Uncharacterized protein
Uncharacterized protein
Uncharacterized protein
Uncharacterized protein
Uncharacterized protein
Uncharacterized protein
Uncharacterized protein
Uncharacterized protein
Uncharacterized protein
Uncharacterized protein
Uncharacterized protein
Uncharacterized protein
Uncharacterized protein
Uncharacterized protein
Uncharacterized protein
Uncharacterized protein
Uncharacterized protein
Uncharacterized protein
Uncharacterized protein
Uncharacterized protein
Uncharacterized protein
Uncharacterized protein
Uncharacterized protein
Uncharacterized protein
Uncharacterized protein
Uncharacterized protein
Uncharacterized protein
Uncharacterized protein
Uncharacterized protein
Uncharacterized protein
Uncharacterized protein
Uncharacterized protein

Uncharacterized protein
Uncharacterized protein
Uncharacterized protein
Uncharacterized protein
Uncharacterized protein
Uncharacterized protein
Uncharacterized protein
Uncharacterized protein
Uncharacterized protein
Uncharacterized protein
Uncharacterized protein
Uncharacterized protein
Uncharacterized protein
Uncharacterized protein
Uncharacterized protein
Uncharacterized protein
Uncharacterized protein
Uncharacterized protein
Uncharacterized protein
Uncharacterized protein
Uncharacterized protein
Uncharacterized protein
Uncharacterized protein
Uncharacterized protein
Uncharacterized protein
Uncharacterized protein
Uncharacterized protein
Uncharacterized protein
Uncharacterized protein
Uncharacterized protein
Uncharacterized protein
Uncharacterized protein
Uncharacterized protein
Uncharacterized protein
Uncharacterized protein
Uncharacterized protein
Uncharacterized protein
Uncharacterized protein
Uncharacterized protein
Uncharacterized protein
Uncharacterized protein

Uncharacterized protein
Uncharacterized protein
Uncharacterized protein
Uncharacterized protein
Uncharacterized protein
Uncharacterized protein
Uncharacterized protein
Uncharacterized protein
Uncharacterized protein
Uncharacterized protein
Uncharacterized protein
Uncharacterized protein
Uncharacterized protein
Uncharacterized protein
Uncharacterized protein
Uncharacterized protein
Uncharacterized protein
PAN domain protein
PAN domain protein
Hepatocyte growth factor
Uncharacterized protein
Uncharacterized protein
Uncharacterized protein
Hepatocyte growth factor
Plasminogen
Hepatocyte growth factor
Uncharacterized protein
Uncharacterized protein
Plasminogen
Uncharacterized protein
Putative serine protease HhoB
Uncharacterized protein
Uncharacterized protein
Uncharacterized protein
Protein containing PAN domain
Protein containing PAN domain/Ricin-type beta-trefoil lectin domain
Fibroblast growth factor receptor 2
Uncharacterized protein
Endoglucanase-5
Tyrosine-protein kinase receptor UFO
Plasminogen

Cuticlin-1
Cuticlin-1
Zonadhesin
Plasminogen
Receptor-like tyrosine-protein kinase kin-16
Uncharacterized protein
Putative chitinase 3
Uncharacterized protein
Aldehyde dehydrogenase family 3 member B1
Uncharacterized protein
Glucan 1,3-beta-glucosidase
Uncharacterized protein
Uncharacterized protein
Uncharacterized protein
Uncharacterized protein
Uncharacterized protein
Uncharacterized protein
Uncharacterized protein
Uncharacterized protein
Unplaced genomic scaffold supercont1.1, whole genome shotgun sequence
LGALS3_3 protein
Cut-1_10 protein
Cut-1_2 protein
Prss12_0 protein (Fragment)
PRSS12 protein (Fragment)
Prss12_1 protein (Fragment)
GRP3_1 protein (Fragment)
GRP3_0 protein (Fragment)
PRM1_2 protein
PRM1_0 protein
Mannose binding lectin
Hepatocyte growth factor
Coagulation factor XI
Plasma kallikrein
Hepatocyte growth factor-like protein
Hepatocyte growth factor (Fragment)
Plasminogen (Fragment)
Plasma kallikrein (Fragment)
Uncharacterized protein
PAN domain-containing protein
Uncharacterized protein

Uncharacterized protein
Hepatocyte growth factor-like protein
Coagulation factor XI
Hepatocyte growth factor
Plasma kallikrein (Fragment)
Coagulation factor XI (Fragment)
Hepatocyte growth factor
Uncharacterized protein
Uncharacterized protein
Plasminogen
Uncharacterized protein
Hepatocyte growth factor
Coagulation factor XI
Hepatocyte growth factor-like protein
Hepatocyte growth factor-like protein
Uncharacterized protein
Plasminogen
Plasminogen
Uncharacterized protein
Hepatocyte growth factor-like protein
Hepatocyte growth factor (Fragment)
Hepatocyte growth factor-like protein
Uncharacterized protein
Plasminogen
Plasminogen
Uncharacterized protein
Coagulation factor XI
Coagulation factor XI
Thyroglobulin-like protein
Plasminogen
Plasma kallikrein (Fragment)
Plasma kallikrein (Fragment)
Coagulation factor XI
Coagulation factor XI
Uncharacterized protein
Hepatocyte growth factor (Fragment)
Coagulation factor XI (Fragment)
Uncharacterized protein
Coagulation factor XI
Plasminogen
Uncharacterized protein

Uncharacterized protein
Uncharacterized protein
Uncharacterized protein
Uncharacterized protein
Uncharacterized protein
Uncharacterized protein
Uncharacterized protein
Uncharacterized protein
Uncharacterized protein
Uncharacterized protein
Uncharacterized protein
Uncharacterized protein
Uncharacterized protein
Uncharacterized protein
Uncharacterized protein
Uncharacterized protein
Uncharacterized protein
Uncharacterized protein
Uncharacterized protein
Uncharacterized protein
Uncharacterized protein
Uncharacterized protein
Uncharacterized protein
Uncharacterized protein
Uncharacterized protein
Uncharacterized protein
Uncharacterized protein
Uncharacterized protein
Uncharacterized protein
Uncharacterized protein
Uncharacterized protein
Uncharacterized protein
Uncharacterized protein
Uncharacterized protein
Uncharacterized protein
Uncharacterized protein
Uncharacterized protein
Uncharacterized protein
Uncharacterized protein
Uncharacterized protein
Uncharacterized protein

Uncharacterized protein
Uncharacterized protein
Uncharacterized protein
Uncharacterized protein
Galactose oxidase
Uncharacterized protein
Uncharacterized protein
Uncharacterized protein
Uncharacterized protein
Uncharacterized protein
Uncharacterized protein
Uncharacterized protein
Uncharacterized protein
Uncharacterized protein
Uncharacterized protein
Uncharacterized protein
Uncharacterized protein
Uncharacterized protein
Uncharacterized protein
Uncharacterized protein
Uncharacterized protein
Uncharacterized protein
Uncharacterized protein
Uncharacterized protein
Uncharacterized protein
Uncharacterized protein
Uncharacterized protein
Uncharacterized protein
Uncharacterized protein
Uncharacterized protein
Uncharacterized protein
Uncharacterized protein
Uncharacterized protein
Uncharacterized protein
Uncharacterized protein
Uncharacterized protein
Uncharacterized protein
Uncharacterized protein
Uncharacterized protein
Uncharacterized protein
Uncharacterized protein

Uncharacterized protein
Uncharacterized protein
Uncharacterized protein
Uncharacterized protein
Uncharacterized protein
Uncharacterized protein
Muc1-extracellular alpha--glucan glucosidase
Uncharacterized protein
Glycosyltransferase family 31
Uncharacterized protein
Kelch protein
Uncharacterized protein (Fragment)
Uncharacterized protein
Uncharacterized protein (Fragment)
Pantothenate transporter liz1
Uncharacterized protein
Uncharacterized protein
Muc1-extracellular alpha-1,4-glucan glucosidase
Heterokaryon incompatibility protein
Uncharacterized protein
Uncharacterized protein
Uncharacterized protein
Uncharacterized protein
Pan domain-containing
Uncharacterized protein
Galactose oxidase
Uncharacterized protein
Uncharacterized protein (Fragment)
Uncharacterized protein
Uncharacterized protein
Uncharacterized protein
Uncharacterized protein
Uncharacterized protein
Uncharacterized protein
Uncharacterized protein
Uncharacterized protein
Uncharacterized protein
Uncharacterized protein
Uncharacterized protein
Uncharacterized protein
Uncharacterized protein

Uncharacterized protein
Uncharacterized protein
Uncharacterized protein
Uncharacterized protein
Uncharacterized protein
Uncharacterized protein
Related to galactose oxidase
Uncharacterized protein
Uncharacterized protein
Uncharacterized protein
Uncharacterized protein
Uncharacterized protein
Related to MUC1-Extracellular alpha-1,4-glucan glucosidase
Uncharacterized protein
Uncharacterized protein
Uncharacterized protein
Uncharacterized protein
Uncharacterized protein
Uncharacterized protein
Uncharacterized protein
Uncharacterized protein
Uncharacterized protein
Uncharacterized protein
Uncharacterized protein
Uncharacterized protein
Uncharacterized protein
Uncharacterized protein
Uncharacterized protein
Uncharacterized protein
Uncharacterized protein
Uncharacterized protein
Uncharacterized protein
Uncharacterized protein
Uncharacterized protein
Uncharacterized protein
Uncharacterized protein (Fragment)
Uncharacterized protein
Uncharacterized protein
Uncharacterized protein
Uncharacterized protein
Uncharacterized protein

Uncharacterized protein
Uncharacterized protein
Uncharacterized protein
Uncharacterized protein
Uncharacterized protein
Uncharacterized protein
Galactose oxidase
Uncharacterized protein
Uncharacterized protein
Uncharacterized protein
Uncharacterized protein
Uncharacterized protein
Uncharacterized protein
Uncharacterized protein
Uncharacterized protein
Uncharacterized protein
Uncharacterized protein
Uncharacterized protein
Uncharacterized protein
Uncharacterized protein
Uncharacterized protein
Uncharacterized protein
Uncharacterized protein
Uncharacterized protein
Uncharacterized protein
Uncharacterized protein
Uncharacterized protein
Uncharacterized protein
Uncharacterized protein
Uncharacterized protein
Uncharacterized protein
Uncharacterized protein
Uncharacterized protein
Uncharacterized protein
Uncharacterized protein
Uncharacterized protein
Uncharacterized protein
Uncharacterized protein (Fragment)
Uncharacterized protein
Uncharacterized protein
Uncharacterized protein

Uncharacterized protein
Uncharacterized protein
Uncharacterized protein
Uncharacterized protein
Uncharacterized protein
Uncharacterized protein
Uncharacterized protein
Uncharacterized protein
Uncharacterized protein
Galactose oxidase
Uncharacterized protein
Uncharacterized protein
Uncharacterized protein
Uncharacterized protein (Fragment)
Uncharacterized protein
Uncharacterized protein
Uncharacterized protein
Uncharacterized protein
Uncharacterized protein
Uncharacterized protein
Uncharacterized protein
Mucin-1
Uncharacterized protein
Uncharacterized protein
Uncharacterized protein
Uncharacterized protein (Fragment)
Uncharacterized protein
Galactose oxidase
Uncharacterized protein
Uncharacterized protein
Uncharacterized protein
Uncharacterized protein
Uncharacterized protein
Uncharacterized protein
Uncharacterized protein
Uncharacterized protein
Uncharacterized protein
Uncharacterized protein
Uncharacterized protein
Uncharacterized protein
Uncharacterized protein

Uncharacterized protein
Uncharacterized protein
Uncharacterized protein
Uncharacterized protein
Uncharacterized protein
Uncharacterized protein
Uncharacterized protein
Uncharacterized protein
Uncharacterized protein
Uncharacterized protein
Uncharacterized protein
Uncharacterized protein
Uncharacterized protein
Uncharacterized protein
Uncharacterized protein
Uncharacterized protein
Uncharacterized protein
Uncharacterized protein
Uncharacterized protein
Uncharacterized protein
Uncharacterized protein
Uncharacterized protein
Uncharacterized protein
Uncharacterized protein
Uncharacterized protein
Uncharacterized protein
Uncharacterized protein
Uncharacterized protein
Uncharacterized protein
Uncharacterized protein
Uncharacterized protein
Uncharacterized protein
Uncharacterized protein
Uncharacterized protein
Uncharacterized protein
Uncharacterized protein
Uncharacterized protein
Uncharacterized protein
Uncharacterized protein
Uncharacterized protein
Uncharacterized protein

Uncharacterized protein
Uncharacterized protein
Uncharacterized protein
Uncharacterized protein
Uncharacterized protein
Uncharacterized protein
Uncharacterized protein
Uncharacterized protein
Uncharacterized protein
Uncharacterized protein
Uncharacterized protein
Uncharacterized protein
Uncharacterized protein
Uncharacterized protein
Uncharacterized protein
Uncharacterized protein
Uncharacterized protein
Uncharacterized protein
Uncharacterized protein
Uncharacterized protein
Uncharacterized protein
Uncharacterized protein
Uncharacterized protein
Uncharacterized protein
Uncharacterized protein
Uncharacterized protein
Uncharacterized protein
Uncharacterized protein
Uncharacterized protein
Uncharacterized protein
Uncharacterized protein
Uncharacterized protein
Uncharacterized protein
Uncharacterized protein
Uncharacterized protein
Uncharacterized protein
Uncharacterized protein
Uncharacterized protein
Uncharacterized protein
Uncharacterized protein
Uncharacterized protein

Uncharacterized protein
Uncharacterized protein
Uncharacterized protein
Uncharacterized protein
Uncharacterized protein
Uncharacterized protein
Uncharacterized protein
Uncharacterized protein
Uncharacterized protein
Uncharacterized protein
Uncharacterized protein
Uncharacterized protein
Uncharacterized protein
Uncharacterized protein
Uncharacterized protein
Uncharacterized protein
Uncharacterized protein
Uncharacterized protein
Uncharacterized protein
Uncharacterized protein
Uncharacterized protein
Uncharacterized protein
Uncharacterized protein
Uncharacterized protein
Uncharacterized protein
Uncharacterized protein
Uncharacterized protein
Uncharacterized protein
Uncharacterized protein
Uncharacterized protein
Uncharacterized protein
Uncharacterized protein
Uncharacterized protein
Uncharacterized protein
Uncharacterized protein
Uncharacterized protein
Uncharacterized protein
Uncharacterized protein
Uncharacterized protein
Uncharacterized protein
Uncharacterized protein

Uncharacterized protein
Uncharacterized protein
Uncharacterized protein
Uncharacterized protein
Uncharacterized protein
Uncharacterized protein
Uncharacterized protein
Uncharacterized protein
Uncharacterized protein
Uncharacterized protein
Uncharacterized protein
Uncharacterized protein
Uncharacterized protein
Uncharacterized protein
Uncharacterized protein
Uncharacterized protein
Uncharacterized protein
Uncharacterized protein
Uncharacterized protein
Uncharacterized protein
Uncharacterized protein
Uncharacterized protein
Uncharacterized protein
Uncharacterized protein
Uncharacterized protein
Uncharacterized protein
Uncharacterized protein
Uncharacterized protein
Uncharacterized protein
Uncharacterized protein
Uncharacterized protein
Uncharacterized protein
Uncharacterized protein
Uncharacterized protein
Uncharacterized protein
Uncharacterized protein
Uncharacterized protein
Related to galactose oxidase
Uncharacterized protein
Uncharacterized protein
Uncharacterized protein

Uncharacterized protein
Uncharacterized protein
Uncharacterized protein
Uncharacterized protein
Uncharacterized protein
Uncharacterized protein
Uncharacterized protein
Uncharacterized protein
Uncharacterized protein
Uncharacterized protein
Uncharacterized protein
Uncharacterized protein
Uncharacterized protein
Galactose oxidase
Uncharacterized protein
Uncharacterized protein
Uncharacterized protein
Uncharacterized protein
Uncharacterized protein
Uncharacterized protein
Uncharacterized protein
Uncharacterized protein
Uncharacterized protein
Uncharacterized protein
Uncharacterized protein
Uncharacterized protein
Uncharacterized protein
Uncharacterized protein
Uncharacterized protein
Uncharacterized protein
Uncharacterized protein
Uncharacterized protein
Uncharacterized protein
Uncharacterized protein
Uncharacterized protein
Uncharacterized protein
Uncharacterized protein
Uncharacterized protein
Uncharacterized protein
Uncharacterized protein
Uncharacterized protein

Uncharacterized protein
Uncharacterized protein
Uncharacterized protein
Uncharacterized protein
Uncharacterized protein
Uncharacterized protein
Uncharacterized protein
Uncharacterized protein
Uncharacterized protein
Uncharacterized protein
Uncharacterized protein
Uncharacterized protein
Uncharacterized protein
Uncharacterized protein
Uncharacterized protein
Uncharacterized protein
Uncharacterized protein
Galactose oxidase
Uncharacterized protein
Uncharacterized protein
Uncharacterized protein
Uncharacterized protein
Uncharacterized protein
Uncharacterized protein
Uncharacterized protein
Uncharacterized protein
Uncharacterized protein
Uncharacterized protein
Uncharacterized protein
Uncharacterized protein
Uncharacterized protein
Uncharacterized protein
Uncharacterized protein
Uncharacterized protein
Uncharacterized protein
Uncharacterized protein
Uncharacterized protein
Uncharacterized protein
Uncharacterized protein
Uncharacterized protein
Uncharacterized protein

Uncharacterized protein
Uncharacterized protein
Uncharacterized protein
Uncharacterized protein
Uncharacterized protein
Uncharacterized protein
Uncharacterized protein
Uncharacterized protein
Uncharacterized protein
Uncharacterized protein
Uncharacterized protein
Related to galactose oxidase
Uncharacterized protein
Uncharacterized protein
Uncharacterized protein
Uncharacterized protein
Uncharacterized protein
Uncharacterized protein
Uncharacterized protein
Uncharacterized protein
Uncharacterized protein
Uncharacterized protein
Uncharacterized protein
Uncharacterized protein
Uncharacterized protein
Uncharacterized protein
Uncharacterized protein
Uncharacterized protein
Uncharacterized protein
Uncharacterized protein (Fragment)
Uncharacterized protein
Uncharacterized protein
Uncharacterized protein
Uncharacterized protein
Uncharacterized protein
Uncharacterized protein
Uncharacterized protein
Uncharacterized protein
Uncharacterized protein
Uncharacterized protein
Uncharacterized protein

Uncharacterized protein
Uncharacterized protein (Fragment)
WGS project CBME000000000 data, contig CS3487_c001953
Uncharacterized protein
Uncharacterized protein
Uncharacterized protein
Uncharacterized protein
Uncharacterized protein
Uncharacterized protein
Uncharacterized protein
Uncharacterized protein
Uncharacterized protein
Uncharacterized protein
Uncharacterized protein
Uncharacterized protein
Uncharacterized protein
Uncharacterized protein
Uncharacterized protein
Uncharacterized protein
Uncharacterized protein
Uncharacterized protein
Uncharacterized protein
Uncharacterized protein
Uncharacterized protein
Uncharacterized protein
Uncharacterized protein
Uncharacterized protein
Uncharacterized protein
Uncharacterized protein
Uncharacterized protein
Uncharacterized protein
Uncharacterized protein
Uncharacterized protein
Uncharacterized protein
Uncharacterized protein
Uncharacterized protein
Uncharacterized protein
Uncharacterized protein
Uncharacterized protein
Uncharacterized protein
Uncharacterized protein

Uncharacterized protein
Uncharacterized protein
Uncharacterized protein
Uncharacterized protein
Uncharacterized protein
Uncharacterized protein
Uncharacterized protein
Uncharacterized protein
Uncharacterized protein
Uncharacterized protein
Uncharacterized protein
Uncharacterized protein
Uncharacterized protein
Uncharacterized protein
Uncharacterized protein
Uncharacterized protein
Uncharacterized protein
Uncharacterized protein
Uncharacterized protein
Uncharacterized protein
Uncharacterized protein
Uncharacterized protein
Uncharacterized protein
Uncharacterized protein
Uncharacterized protein
Uncharacterized protein
Uncharacterized protein
Uncharacterized protein
Uncharacterized protein
Uncharacterized protein
Uncharacterized protein
Uncharacterized protein
Uncharacterized protein
Galactose oxidase
Uncharacterized protein
Uncharacterized protein
Uncharacterized protein
Uncharacterized protein
Uncharacterized protein
Uncharacterized protein
Uncharacterized protein

Muc1-extracellular alpha-1,4-glucan glucosidase
Uncharacterized protein
Uncharacterized protein
Uncharacterized protein
Uncharacterized protein
Uncharacterized protein
Uncharacterized protein
Uncharacterized protein (Fragment)
Uncharacterized protein
Uncharacterized protein
Uncharacterized protein
Uncharacterized protein
Glycosyltransferase family 31
Uncharacterized protein
Uncharacterized protein
Uncharacterized protein
Uncharacterized protein
Uncharacterized protein
Uncharacterized protein
Uncharacterized protein
Uncharacterized protein
Uncharacterized protein
Uncharacterized protein
Uncharacterized protein
Uncharacterized protein
Uncharacterized protein
Uncharacterized protein
Uncharacterized protein
Uncharacterized protein
Uncharacterized protein
Uncharacterized protein
Uncharacterized protein
WGS project CBMI000000000 data, contig CS3069_c002857
WGS project CBMI000000000 data, contig CS3069_c001115
WGS project CBMI000000000 data, contig CS3069_c002119
Uncharacterized protein
Uncharacterized protein
Uncharacterized protein
Uncharacterized protein
Uncharacterized protein
Uncharacterized protein

Uncharacterized protein
Uncharacterized protein
Uncharacterized protein
Uncharacterized protein
Uncharacterized protein
Uncharacterized protein
Uncharacterized protein
Uncharacterized protein
Uncharacterized protein
Uncharacterized protein
Uncharacterized protein
Uncharacterized protein
Uncharacterized protein
Uncharacterized protein
Uncharacterized protein
Uncharacterized protein
Uncharacterized protein
Uncharacterized protein
Galactose oxidase
Uncharacterized protein
Muc1-extracellular alpha-1,4-glucan glucosidase
Uncharacterized protein
Uncharacterized protein
Uncharacterized protein
Uncharacterized protein
Uncharacterized protein
Glycosyltransferase family 31
Uncharacterized protein
Uncharacterized protein
Uncharacterized protein
Uncharacterized protein
Uncharacterized protein
Uncharacterized protein
Uncharacterized protein
Uncharacterized protein
Uncharacterized protein
Uncharacterized protein
Uncharacterized protein
Uncharacterized protein
Uncharacterized protein
Uncharacterized protein

Uncharacterized protein
Uncharacterized protein
Uncharacterized protein
Uncharacterized protein
Uncharacterized protein
Uncharacterized protein
Uncharacterized protein
Uncharacterized protein
Galactose oxidase
Uncharacterized protein
Hepatocyte growth factor
Hepatocyte growth factor-like/macrophage stimulating protein
Hepatocyte growth factor
Plasminogen
Plasminogen
Uncharacterized protein
Uncharacterized protein
Hepatocyte growth factor
HGF protein ( SFprotein ) (Fragment)
Uncharacterized protein
Uncharacterized protein (Fragment)
Uncharacterized protein
Uncharacterized protein (Fragment)
Uncharacterized protein (Fragment)
Uncharacterized protein
Uncharacterized protein
Uncharacterized protein
Uncharacterized protein
Uncharacterized protein (Fragment)
Plasminogen
Hepatocyte growth factor a
Plasminogen
Uncharacterized protein (Fragment)
Plasminogen
Uncharacterized protein
Macrophage stimulating 1
Hepatocyte growth factor
Plasma kallikrein (Fragment)
Uncharacterized protein
Uncharacterized protein
Alpha-2-macroglobulin family protein

Uncharacterized protein
Uncharacterized protein
Alpha-2-macroglobulin family protein
Uncharacterized protein
Alpha-2-macroglobulin family protein
Uncharacterized protein (Fragment)
Uncharacterized protein (Fragment)
Uncharacterized protein (Fragment)
Uncharacterized protein (Fragment)
Uncharacterized protein
Uncharacterized protein
Uncharacterized protein
Uncharacterized protein
Uncharacterized protein
Uncharacterized protein
Uncharacterized protein
Uncharacterized protein
Uncharacterized protein
Uncharacterized protein
Related to galactose oxidase
Uncharacterized protein
Uncharacterized protein
Uncharacterized protein
Uncharacterized protein
Uncharacterized protein
Uncharacterized protein
Uncharacterized protein
Uncharacterized protein
Uncharacterized protein
MUC1-Extracellular alpha-1,4-glucan glucosidase
Uncharacterized protein
Uncharacterized protein
Uncharacterized protein
Galactose oxidase
Uncharacterized protein
Uncharacterized protein
Uncharacterized protein
Uncharacterized protein
Uncharacterized protein
Uncharacterized protein
Uncharacterized protein

Uncharacterized protein
Uncharacterized protein
Uncharacterized protein
Uncharacterized protein
Uncharacterized protein
Related to galactose oxidase
Uncharacterized protein
Uncharacterized protein
Uncharacterized protein
MUC1-Extracellular alpha-1,4-glucan glucosidase
Probable MUC1-Extracellular alpha-1,4-glucan glucosidase
Uncharacterized protein
Uncharacterized protein
Uncharacterized protein
Uncharacterized protein
Uncharacterized protein
Uncharacterized protein
Uncharacterized protein
Uncharacterized protein
Uncharacterized protein
Uncharacterized protein
Uncharacterized protein
Uncharacterized protein
Uncharacterized protein
Uncharacterized protein
Uncharacterized protein
Uncharacterized protein
Uncharacterized protein
Uncharacterized protein
Uncharacterized protein
Uncharacterized protein
Uncharacterized protein
Uncharacterized protein
Uncharacterized protein
Uncharacterized protein
Uncharacterized protein
Uncharacterized protein
Uncharacterized protein
Uncharacterized protein
Uncharacterized protein
Uncharacterized protein

Uncharacterized protein
Uncharacterized protein
Uncharacterized protein
Uncharacterized protein
Uncharacterized protein
Uncharacterized protein
Uncharacterized protein
Uncharacterized protein
Uncharacterized protein
Uncharacterized protein
Uncharacterized protein
Uncharacterized protein
Uncharacterized protein
Uncharacterized protein
Uncharacterized protein
Uncharacterized protein
Uncharacterized protein
Uncharacterized protein
Uncharacterized protein
Uncharacterized protein
Uncharacterized protein
Uncharacterized protein
Uncharacterized protein
Uncharacterized protein
Galactose oxidase
Uncharacterized protein
Uncharacterized protein
Uncharacterized protein
Uncharacterized protein
Uncharacterized protein
Uncharacterized protein
Uncharacterized protein
Uncharacterized protein
Uncharacterized protein
Uncharacterized protein
Uncharacterized protein
Uncharacterized protein
Uncharacterized protein
Uncharacterized protein
Uncharacterized protein
Uncharacterized protein

Uncharacterized protein
Uncharacterized protein
Uncharacterized protein
Uncharacterized protein
Uncharacterized protein
Uncharacterized protein
Uncharacterized protein
Uncharacterized protein
Uncharacterized protein
Uncharacterized protein
Uncharacterized protein
Uncharacterized protein
Uncharacterized protein
Uncharacterized protein
Uncharacterized protein
Uncharacterized protein
Uncharacterized protein
Uncharacterized protein
Uncharacterized protein
Uncharacterized protein
Uncharacterized protein
Uncharacterized protein
Uncharacterized protein
Uncharacterized protein
Uncharacterized protein
Uncharacterized protein
Uncharacterized protein
Uncharacterized protein
Uncharacterized protein
Uncharacterized protein
Uncharacterized protein
Uncharacterized protein
Uncharacterized protein
Uncharacterized protein
Uncharacterized protein
Uncharacterized protein
Uncharacterized protein
Uncharacterized protein
Uncharacterized protein
Uncharacterized protein
Uncharacterized protein

Uncharacterized protein
Uncharacterized protein
Uncharacterized protein
Uncharacterized protein
Uncharacterized protein
Uncharacterized protein
Uncharacterized protein
Uncharacterized protein
Uncharacterized protein
Uncharacterized protein
Uncharacterized protein
Uncharacterized protein
Uncharacterized protein
Uncharacterized protein
Uncharacterized protein
Uncharacterized protein
Uncharacterized protein
Uncharacterized protein
Uncharacterized protein
Uncharacterized protein
Uncharacterized protein
Uncharacterized protein
Uncharacterized protein
Uncharacterized protein
Uncharacterized protein
Uncharacterized protein
Uncharacterized protein
Uncharacterized protein
Uncharacterized protein
Uncharacterized protein
Uncharacterized protein
Uncharacterized protein
Uncharacterized protein
Uncharacterized protein
Uncharacterized protein
Uncharacterized protein
Uncharacterized protein
Uncharacterized protein
Uncharacterized protein
Uncharacterized protein
Uncharacterized protein

Uncharacterized protein
Uncharacterized protein
Uncharacterized protein
Uncharacterized protein
Uncharacterized protein
Uncharacterized protein
Uncharacterized protein
Uncharacterized protein
Uncharacterized protein
Uncharacterized protein
Uncharacterized protein
Uncharacterized protein
Uncharacterized protein
Uncharacterized protein
Uncharacterized protein
Uncharacterized protein
Uncharacterized protein
Uncharacterized protein
Uncharacterized protein
Uncharacterized protein
Uncharacterized protein
Uncharacterized protein
Uncharacterized protein
Uncharacterized protein
Uncharacterized protein
Uncharacterized protein
Uncharacterized protein (Fragment)
Uncharacterized protein
Uncharacterized protein
Uncharacterized protein
Uncharacterized protein
Uncharacterized protein
Uncharacterized protein
Uncharacterized protein
Uncharacterized protein
Uncharacterized protein
Uncharacterized protein
Uncharacterized protein
Uncharacterized protein
Uncharacterized protein
Uncharacterized protein

Glucanase
Uncharacterized protein
Uncharacterized protein
Uncharacterized protein
Uncharacterized protein
Uncharacterized protein
Uncharacterized protein
Uncharacterized protein
Uncharacterized protein
Uncharacterized protein
Uncharacterized protein
Uncharacterized protein
Uncharacterized protein
Uncharacterized protein
Uncharacterized protein
Uncharacterized protein
Uncharacterized protein
Uncharacterized protein
Uncharacterized protein
Uncharacterized protein
Uncharacterized protein
Uncharacterized protein
Uncharacterized protein
Uncharacterized protein
Uncharacterized protein
Uncharacterized protein
Uncharacterized protein
Uncharacterized protein
Uncharacterized protein
Uncharacterized protein
Uncharacterized protein
Uncharacterized protein
Uncharacterized protein
Uncharacterized protein
Uncharacterized protein
Uncharacterized protein
Uncharacterized protein
Uncharacterized protein
Uncharacterized protein
Uncharacterized protein
Uncharacterized protein

Uncharacterized protein
Uncharacterized protein
Uncharacterized protein
Uncharacterized protein
Uncharacterized protein
Uncharacterized protein
Uncharacterized protein
Uncharacterized protein
Uncharacterized protein
Uncharacterized protein
Uncharacterized protein
Uncharacterized protein
Uncharacterized protein
Uncharacterized protein
Uncharacterized protein
Uncharacterized protein
Uncharacterized protein
Uncharacterized protein
Uncharacterized protein
Uncharacterized protein
Uncharacterized protein
Uncharacterized protein
Uncharacterized protein
Uncharacterized protein
Uncharacterized protein
Uncharacterized protein
Uncharacterized protein
Uncharacterized protein
Uncharacterized protein
Uncharacterized protein
Uncharacterized protein
Uncharacterized protein
Uncharacterized protein
Uncharacterized protein
Uncharacterized protein
Uncharacterized protein
Uncharacterized protein
Uncharacterized protein
Uncharacterized protein
Uncharacterized protein
Uncharacterized protein

Uncharacterized protein
Uncharacterized protein
Uncharacterized protein
Uncharacterized protein
Uncharacterized protein
Uncharacterized protein
Uncharacterized protein
Receptor-like protein kinase like protein
TIR-NBS-LRR type disease resistance protein
Uncharacterized protein
Uncharacterized protein
Uncharacterized protein
Uncharacterized protein
Uncharacterized protein
Uncharacterized protein
Uncharacterized protein
Uncharacterized protein
Uncharacterized protein
Uncharacterized protein
Uncharacterized protein
Uncharacterized protein
Uncharacterized protein
Uncharacterized protein
Uncharacterized protein
Uncharacterized protein
Uncharacterized protein
Uncharacterized protein
Uncharacterized protein
Uncharacterized protein
Uncharacterized protein
Uncharacterized protein
Uncharacterized protein
Uncharacterized protein
Uncharacterized protein
Uncharacterized protein
Uncharacterized protein
Uncharacterized protein
Uncharacterized protein
Uncharacterized protein
Uncharacterized protein
Uncharacterized protein

Uncharacterized protein
Uncharacterized protein
Uncharacterized protein
Uncharacterized protein
Uncharacterized protein
Uncharacterized protein
Uncharacterized protein
Uncharacterized protein
Uncharacterized protein
Uncharacterized protein
Uncharacterized protein
Uncharacterized protein
Uncharacterized protein
Uncharacterized protein
Uncharacterized protein
Uncharacterized protein
Uncharacterized protein
Uncharacterized protein
Uncharacterized protein
Uncharacterized protein
Receptor-like protein kinase precusor-like protein
Uncharacterized protein
Uncharacterized protein
Uncharacterized protein
Uncharacterized protein
Uncharacterized protein
Uncharacterized protein
Uncharacterized protein
Uncharacterized protein
Uncharacterized protein
Uncharacterized protein
Uncharacterized protein
Uncharacterized protein
Uncharacterized protein
Uncharacterized protein
Uncharacterized protein
Uncharacterized protein
Uncharacterized protein
Uncharacterized protein
Uncharacterized protein
Uncharacterized protein

Uncharacterized protein
Uncharacterized protein
Uncharacterized protein
Uncharacterized protein
Uncharacterized protein
Uncharacterized protein
Uncharacterized protein
Uncharacterized protein
Uncharacterized protein
Uncharacterized protein
Uncharacterized protein
Uncharacterized protein
Uncharacterized protein
Uncharacterized protein
Uncharacterized protein
Uncharacterized protein
Uncharacterized protein
Uncharacterized protein
Uncharacterized protein
Uncharacterized protein
Uncharacterized protein
Uncharacterized protein
Uncharacterized protein
Uncharacterized protein
Uncharacterized protein
Uncharacterized protein
Uncharacterized protein
Uncharacterized protein
Uncharacterized protein
Uncharacterized protein
Uncharacterized protein
Uncharacterized protein
Uncharacterized protein
Uncharacterized protein (Fragment)
Uncharacterized protein
Uncharacterized protein
Uncharacterized protein
Uncharacterized protein
Uncharacterized protein
Uncharacterized protein
Uncharacterized protein

Uncharacterized protein
Uncharacterized protein
Uncharacterized protein
Uncharacterized protein
Uncharacterized protein
Uncharacterized protein
Uncharacterized protein
Uncharacterized protein
Uncharacterized protein
Uncharacterized protein
Uncharacterized protein
Uncharacterized protein
Uncharacterized protein
Uncharacterized protein
Uncharacterized protein
Uncharacterized protein
Uncharacterized protein
Uncharacterized protein
Uncharacterized protein
Uncharacterized protein
Uncharacterized protein
Uncharacterized protein
Uncharacterized protein
Uncharacterized protein
Uncharacterized protein
Uncharacterized protein
Uncharacterized protein
Uncharacterized protein
Uncharacterized protein
Uncharacterized protein
Uncharacterized protein
Uncharacterized protein
Uncharacterized protein
Uncharacterized protein
Uncharacterized protein
Uncharacterized protein
Uncharacterized protein
Uncharacterized protein
Uncharacterized protein
Uncharacterized protein
Uncharacterized protein

Uncharacterized protein
Uncharacterized protein
Uncharacterized protein
Uncharacterized protein
Uncharacterized protein
Uncharacterized protein
Uncharacterized protein
Uncharacterized protein
Uncharacterized protein
Uncharacterized protein
Uncharacterized protein
Uncharacterized protein
Uncharacterized protein
Uncharacterized protein
Uncharacterized protein
Uncharacterized protein
Uncharacterized protein
Uncharacterized protein
Uncharacterized protein
Uncharacterized protein
Uncharacterized protein
Uncharacterized protein
Uncharacterized protein
Uncharacterized protein
Uncharacterized protein
Uncharacterized protein
Uncharacterized protein
Uncharacterized protein
Uncharacterized protein
Uncharacterized protein
Uncharacterized protein
S-locus lectin protein kinase family protein
Uncharacterized protein
Uncharacterized protein
Uncharacterized protein (Fragment)
Uncharacterized protein
Uncharacterized protein
Uncharacterized protein
Uncharacterized protein
Uncharacterized protein
Uncharacterized protein

Uncharacterized protein
Uncharacterized protein
Uncharacterized protein
Uncharacterized protein
Uncharacterized protein
Uncharacterized protein
Uncharacterized protein
Uncharacterized protein
Uncharacterized protein
Uncharacterized protein
Uncharacterized protein
Uncharacterized protein
Uncharacterized protein
Uncharacterized protein
Uncharacterized protein
Uncharacterized protein
Uncharacterized protein
Uncharacterized protein
Uncharacterized protein
Uncharacterized protein
Uncharacterized protein
Uncharacterized protein
Uncharacterized protein
Uncharacterized protein
Uncharacterized protein
G-type lectin S-receptor-like serine/threonine-protein kinase SD3-1
Putative receptor protein kinase ZmPK1
G-type lectin S-receptor-like serine/threonine-protein kinase isoform B
G-type lectin S-receptor-like serine/threonine-protein kinase isoform B
G-type lectin S-receptor-like serine/threonine-protein kinase isoform A
G-type lectin S-receptor-like serine/threonine-protein kinase
G-type lectin S-receptor-like serine/threonine-protein kinase isoform B
G-type lectin S-receptor-like serine/threonine-protein kinase
Receptor-like serine/threonine-protein kinase SD1-8
Receptor-like serine/threonine-protein kinase SD1-8
G-type lectin S-receptor-like serine/threonine-protein kinase
G-type lectin S-receptor-like serine/threonine-protein kinase SD1-1
G-type lectin S-receptor-like serine/threonine-protein kinase SD1-1
Receptor-like serine/threonine-protein kinase SD1-8
G-type lectin S-receptor-like serine/threonine-protein kinase
G-type lectin S-receptor-like serine/threonine-protein kinase SD1-1

G-type lectin S-receptor-like serine/threonine-protein kinase
G-type lectin S-receptor-like serine/threonine-protein kinase
G-type lectin S-receptor-like serine/threonine-protein kinase
PAN domain-containing protein
G-type lectin S-receptor-like serine/threonine-protein kinase B120
G-type lectin S-receptor-like serine/threonine-protein kinase
G-type lectin S-receptor-like serine/threonine-protein kinase
Receptor-like serine/threonine-protein kinase SD1-8
G-type lectin S-receptor-like serine/threonine-protein kinase B120
Receptor-like serine/threonine-protein kinase SD1-8
G-type lectin S-receptor-like serine/threonine-protein kinase SD1-1 isoform A
G-type lectin S-receptor-like serine/threonine-protein kinase
G-type lectin S-receptor-like serine/threonine-protein kinase SD3-1
Receptor-like serine/threonine-protein kinase SD1-6
G-type lectin S-receptor-like serine/threonine-protein kinase
Receptor-like serine/threonine-protein kinase SD1-8
G-type lectin S-receptor-like serine/threonine-protein kinase
G-type lectin S-receptor-like serine/threonine-protein kinase isoform A
G-type lectin S-receptor-like serine/threonine-protein kinase SD2-5
G-type lectin S-receptor-like serine/threonine-protein kinase isoform B
G-type lectin S-receptor-like serine/threonine-protein kinase isoform A
G-type lectin S-receptor-like serine/threonine-protein kinase
G-type lectin S-receptor-like serine/threonine-protein kinase
G-type lectin S-receptor-like serine/threonine-protein kinase SD2-5
G-type lectin S-receptor-like serine/threonine-protein kinase
G-type lectin S-receptor-like serine/threonine-protein kinase
Putative serine/threonine-protein kinase receptor
Receptor-like serine/threonine-protein kinase SD1-8
G-type lectin S-receptor-like serine/threonine-protein kinase
G-type lectin S-receptor-like serine/threonine-protein kinase
Putative receptor protein kinase ZmPK1
G-type lectin S-receptor-like serine/threonine-protein kinase
Putative receptor protein kinase ZmPK1
G-type lectin S-receptor-like serine/threonine-protein kinase SD3-1
G-type lectin S-receptor-like serine/threonine-protein kinase
G-type lectin S-receptor-like serine/threonine-protein kinase
G-type lectin S-receptor-like serine/threonine-protein kinase SD1-1
G-type lectin S-receptor-like serine/threonine-protein kinase
G-type lectin S-receptor-like serine/threonine-protein kinase
G-type lectin S-receptor-like serine/threonine-protein kinase SD3-1
Putative receptor protein kinase ZmPK1

G-type lectin S-receptor-like serine/threonine-protein kinase
G-type lectin S-receptor-like serine/threonine-protein kinase isoform C (Fragment)
G-type lectin S-receptor-like serine/threonine-protein kinase SD1-1 isoform B
G-type lectin S-receptor-like serine/threonine-protein kinase
G-type lectin S-receptor-like serine/threonine-protein kinase
G-type lectin S-receptor-like serine/threonine-protein kinase isoform B
G-type lectin S-receptor-like serine/threonine-protein kinase
Putative receptor protein kinase ZmPK1
G-type lectin S-receptor-like serine/threonine-protein kinase SD3-1
Receptor-like serine/threonine-protein kinase SD1-8 isoform F
G-type lectin S-receptor-like serine/threonine-protein kinase
G-type lectin S-receptor-like serine/threonine-protein kinase
Receptor-like serine/threonine-protein kinase SD1-8 isoform B
G-type lectin S-receptor-like serine/threonine-protein kinase
Putative receptor protein kinase ZmPK1
G-type lectin S-receptor-like serine/threonine-protein kinase
G-type lectin S-receptor-like serine/threonine-protein kinase
G-type lectin S-receptor-like serine/threonine-protein kinase isoform A
G-type lectin S-receptor-like serine/threonine-protein kinase
G-type lectin S-receptor-like serine/threonine-protein kinase SD1-1 isoform B
G-type lectin S-receptor-like serine/threonine-protein kinase isoform B
G-type lectin S-receptor-like serine/threonine-protein kinase
Putative receptor protein kinase ZmPK1
G-type lectin S-receptor-like serine/threonine-protein kinase isoform B
G-type lectin S-receptor-like serine/threonine-protein kinase
G-type lectin S-receptor-like serine/threonine-protein kinase isoform C
G-type lectin S-receptor-like serine/threonine-protein kinase isoform C
G-type lectin S-receptor-like serine/threonine-protein kinase isoform A
G-type lectin S-receptor-like serine/threonine-protein kinase
G-type lectin S-receptor-like serine/threonine-protein kinase
G-type lectin S-receptor-like serine/threonine-protein kinase B120 isoform B
Receptor-like serine/threonine-protein kinase SD1-8
G-type lectin S-receptor-like serine/threonine-protein kinase
G-type lectin S-receptor-like serine/threonine-protein kinase isoform C
G-type lectin S-receptor-like serine/threonine-protein kinase RKS1
Uncharacterized protein
G-type lectin S-receptor-like serine/threonine-protein kinase
G-type lectin S-receptor-like serine/threonine-protein kinase isoform B
G-type lectin S-receptor-like serine/threonine-protein kinase isoform A
G-type lectin S-receptor-like serine/threonine-protein kinase
G-type lectin S-receptor-like serine/threonine-protein kinase isoform A

G-type lectin S-receptor-like serine/threonine-protein kinase SD1-1 isoform C
Receptor-like serine/threonine-protein kinase SD1-7
G-type lectin S-receptor-like serine/threonine-protein kinase B120
G-type lectin S-receptor-like serine/threonine-protein kinase (Fragment)
G-type lectin S-receptor-like serine/threonine-protein kinase
G-type lectin S-receptor-like serine/threonine-protein kinase
G-type lectin S-receptor-like serine/threonine-protein kinase
G-type lectin S-receptor-like serine/threonine-protein kinase
G-type lectin S-receptor-like serine/threonine-protein kinase (Fragment)
G-type lectin S-receptor-like serine/threonine-protein kinase LECRK1
G-type lectin S-receptor-like serine/threonine-protein kinase isoform B
G-type lectin S-receptor-like serine/threonine-protein kinase
G-type lectin S-receptor-like serine/threonine-protein kinase
G-type lectin S-receptor-like serine/threonine-protein kinase
G-type lectin S-receptor-like serine/threonine-protein kinase isoform B
G-type lectin S-receptor-like serine/threonine-protein kinase isoform B
G-type lectin S-receptor-like serine/threonine-protein kinase isoform A
G-type lectin S-receptor-like serine/threonine-protein kinase
G-type lectin S-receptor-like serine/threonine-protein kinase isoform A
G-type lectin S-receptor-like serine/threonine-protein kinase
G-type lectin S-receptor-like serine/threonine-protein kinase B120 isoform B
G-type lectin S-receptor-like serine/threonine-protein kinase isoform B
G-type lectin S-receptor-like serine/threonine-protein kinase isoform B (Fragment)
G-type lectin S-receptor-like serine/threonine-protein kinase isoform A
G-type lectin S-receptor-like serine/threonine-protein kinase
G-type lectin S-receptor-like serine/threonine-protein kinase isoform B
G-type lectin S-receptor-like serine/threonine-protein kinase isoform A
G-type lectin S-receptor-like serine/threonine-protein kinase isoform B
G-type lectin S-receptor-like serine/threonine-protein kinase SD1-1 isoform A
G-type lectin S-receptor-like serine/threonine-protein kinase isoform A
G-type lectin S-receptor-like serine/threonine-protein kinase isoform A
G-type lectin S-receptor-like serine/threonine-protein kinase isoform B
Putative receptor protein kinase ZmPK1
Receptor-like serine/threonine-protein kinase SD1-8
G-type lectin S-receptor-like serine/threonine-protein kinase
G-type lectin S-receptor-like serine/threonine-protein kinase
PAN domain-containing protein
G-type lectin S-receptor-like serine/threonine-protein kinase isoform D
G-type lectin S-receptor-like serine/threonine-protein kinase
G-type lectin S-receptor-like serine/threonine-protein kinase
G-type lectin S-receptor-like serine/threonine-protein kinase isoform C

Receptor-like serine/threonine-protein kinase SD1-7 isoform A
G-type lectin S-receptor-like serine/threonine-protein kinase
Putative receptor protein kinase ZmPK1
Putative receptor protein kinase ZmPK1
G-type lectin S-receptor-like serine/threonine-protein kinase isoform F
G-type lectin S-receptor-like serine/threonine-protein kinase SD3-1
G-type lectin S-receptor-like serine/threonine-protein kinase
G-type lectin S-receptor-like serine/threonine-protein kinase
G-type lectin S-receptor-like serine/threonine-protein kinase SD1-1
Receptor-like serine/threonine-protein kinase SD1-8
Uncharacterized protein
G-type lectin S-receptor-like serine/threonine-protein kinase isoform A
G-type lectin S-receptor-like serine/threonine-protein kinase
G-type lectin S-receptor-like serine/threonine-protein kinase isoform A
G-type lectin S-receptor-like serine/threonine-protein kinase isoform A
G-type lectin S-receptor-like serine/threonine-protein kinase
G-type lectin S-receptor-like serine/threonine-protein kinase isoform B
G-type lectin S-receptor-like serine/threonine-protein kinase
Putative receptor protein kinase ZmPK1 (Fragment)
G-type lectin S-receptor-like serine/threonine-protein kinase
G-type lectin S-receptor-like serine/threonine-protein kinase
G-type lectin S-receptor-like serine/threonine-protein kinase isoform A
Receptor-like serine/threonine-protein kinase SD1-8
G-type lectin S-receptor-like serine/threonine-protein kinase isoform B
G-type lectin S-receptor-like serine/threonine-protein kinase
G-type lectin S-receptor-like serine/threonine-protein kinase isoform B
G-type lectin S-receptor-like serine/threonine-protein kinase SD3-1
Putative receptor protein kinase ZmPK1
G-type lectin S-receptor-like serine/threonine-protein kinase isoform B
Putative serine/threonine-protein kinase WNK2 isoform B
G-type lectin S-receptor-like serine/threonine-protein kinase
G-type lectin S-receptor-like serine/threonine-protein kinase isoform A
G-type lectin S-receptor-like serine/threonine-protein kinase isoform B
G-type lectin S-receptor-like serine/threonine-protein kinase isoform A
G-type lectin S-receptor-like serine/threonine-protein kinase SD1-1 isoform A
Receptor-like serine/threonine-protein kinase SD1-8
G-type lectin S-receptor-like serine/threonine-protein kinase SD1-1 isoform B
G-type lectin S-receptor-like serine/threonine-protein kinase
G-type lectin S-receptor-like serine/threonine-protein kinase isoform C
G-type lectin S-receptor-like serine/threonine-protein kinase
G-type lectin S-receptor-like serine/threonine-protein kinase

G-type lectin S-receptor-like serine/threonine-protein kinase
G-type lectin S-receptor-like serine/threonine-protein kinase
Receptor-like serine/threonine-protein kinase SD1-8
G-type lectin S-receptor-like serine/threonine-protein kinase
G-type lectin S-receptor-like serine/threonine-protein kinase
Putative receptor protein kinase ZmPK1
G-type lectin S-receptor-like serine/threonine-protein kinase
G-type lectin S-receptor-like serine/threonine-protein kinase
G-type lectin S-receptor-like serine/threonine-protein kinase
G-type lectin S-receptor-like serine/threonine-protein kinase
G-type lectin S-receptor-like serine/threonine-protein kinase isoform B
Receptor-like serine/threonine-protein kinase SD1-8 isoform C
G-type lectin S-receptor-like serine/threonine-protein kinase
G-type lectin S-receptor-like serine/threonine-protein kinase
Receptor-like serine/threonine-protein kinase SD1-7 isoform B
G-type lectin S-receptor-like serine/threonine-protein kinase RKS1
G-type lectin S-receptor-like serine/threonine-protein kinase
G-type lectin S-receptor-like serine/threonine-protein kinase B120 isoform C
G-type lectin S-receptor-like serine/threonine-protein kinase
G-type lectin S-receptor-like serine/threonine-protein kinase
Putative receptor protein kinase ZmPK1
G-type lectin S-receptor-like serine/threonine-protein kinase isoform B
G-type lectin S-receptor-like serine/threonine-protein kinase isoform A
Receptor-like serine/threonine-protein kinase SD1-8 isoform C
Retrovirus-related Pol polyprotein from transposon TNT 1-94
G-type lectin S-receptor-like serine/threonine-protein kinase
G-type lectin S-receptor-like serine/threonine-protein kinase
G-type lectin S-receptor-like serine/threonine-protein kinase
G-type lectin S-receptor-like serine/threonine-protein kinase isoform B
G-type lectin S-receptor-like serine/threonine-protein kinase isoform D
G-type lectin S-receptor-like serine/threonine-protein kinase
Uncharacterized protein
G-type lectin S-receptor-like serine/threonine-protein kinase
G-type lectin S-receptor-like serine/threonine-protein kinase isoform E
Receptor-like serine/threonine-protein kinase SD1-8
G-type lectin S-receptor-like serine/threonine-protein kinase isoform A
Actin
G-type lectin S-receptor-like serine/threonine-protein kinase
G-type lectin S-receptor-like serine/threonine-protein kinase isoform A
G-type lectin S-receptor-like serine/threonine-protein kinase isoform A
G-type lectin S-receptor-like serine/threonine-protein kinase isoform B

G-type lectin S-receptor-like serine/threonine-protein kinase isoform B
G-type lectin S-receptor-like serine/threonine-protein kinase isoform A
Receptor-like serine/threonine-protein kinase SD1-8
G-type lectin S-receptor-like serine/threonine-protein kinase isoform A
PAN domain-containing protein
Putative serine/threonine-protein kinase WNK2 isoform A
G-type lectin S-receptor-like serine/threonine-protein kinase isoform A
G-type lectin S-receptor-like serine/threonine-protein kinase B120 isoform A
G-type lectin S-receptor-like serine/threonine-protein kinase isoform A
G-type lectin S-receptor-like serine/threonine-protein kinase
G-type lectin S-receptor-like serine/threonine-protein kinase B120 isoform A
Receptor-like serine/threonine-protein kinase SD1-8 isoform A
G-type lectin S-receptor-like serine/threonine-protein kinase isoform C
G-type lectin S-receptor-like serine/threonine-protein kinase
G-type lectin S-receptor-like serine/threonine-protein kinase isoform A
G-type lectin S-receptor-like serine/threonine-protein kinase
G-type lectin S-receptor-like serine/threonine-protein kinase isoform D
Receptor-like serine/threonine-protein kinase SD1-8
G-type lectin S-receptor-like serine/threonine-protein kinase
G-type lectin S-receptor-like serine/threonine-protein kinase isoform A
G-type lectin S-receptor-like serine/threonine-protein kinase B120 isoform B
G-type lectin S-receptor-like serine/threonine-protein kinase isoform D
Receptor-like serine/threonine-protein kinase SD1-8
G-type lectin S-receptor-like serine/threonine-protein kinase isoform B
G-type lectin S-receptor-like serine/threonine-protein kinase B120 isoform A
G-type lectin S-receptor-like serine/threonine-protein kinase isoform A
G-type lectin S-receptor-like serine/threonine-protein kinase isoform C
G-type lectin S-receptor-like serine/threonine-protein kinase
G-type lectin S-receptor-like serine/threonine-protein kinase isoform B
G-type lectin S-receptor-like serine/threonine-protein kinase
G-type lectin S-receptor-like serine/threonine-protein kinase isoform B
G-type lectin S-receptor-like serine/threonine-protein kinase isoform B
G-type lectin S-receptor-like serine/threonine-protein kinase isoform B
G-type lectin S-receptor-like serine/threonine-protein kinase SD2-2
G-type lectin S-receptor-like serine/threonine-protein kinase isoform C
G-type lectin S-receptor-like serine/threonine-protein kinase
G-type lectin S-receptor-like serine/threonine-protein kinase
G-type lectin S-receptor-like serine/threonine-protein kinase
G-type lectin S-receptor-like serine/threonine-protein kinase
G-type lectin S-receptor-like serine/threonine-protein kinase B120 isoform B
G-type lectin S-receptor-like serine/threonine-protein kinase

G-type lectin S-receptor-like serine/threonine-protein kinase SD1-1
G-type lectin S-receptor-like serine/threonine-protein kinase isoform C
G-type lectin S-receptor-like serine/threonine-protein kinase isoform A
G-type lectin S-receptor-like serine/threonine-protein kinase
Putative serine/threonine-protein kinase WNK2 isoform C
G-type lectin S-receptor-like serine/threonine-protein kinase B120 isoform A
G-type lectin S-receptor-like serine/threonine-protein kinase
G-type lectin S-receptor-like serine/threonine-protein kinase
G-type lectin S-receptor-like serine/threonine-protein kinase isoform C
G-type lectin S-receptor-like serine/threonine-protein kinase isoform A
G-type lectin S-receptor-like serine/threonine-protein kinase
Receptor-like serine/threonine-protein kinase SD1-8 isoform G
Putative receptor protein kinase ZmPK1
G-type lectin S-receptor-like serine/threonine-protein kinase
G-type lectin S-receptor-like serine/threonine-protein kinase
G-type lectin S-receptor-like serine/threonine-protein kinase isoform B
G-type lectin S-receptor-like serine/threonine-protein kinase isoform C
G-type lectin S-receptor-like serine/threonine-protein kinase B120 isoform F
G-type lectin S-receptor-like serine/threonine-protein kinase
G-type lectin S-receptor-like serine/threonine-protein kinase B120 isoform A
G-type lectin S-receptor-like serine/threonine-protein kinase isoform B
G-type lectin S-receptor-like serine/threonine-protein kinase
G-type lectin S-receptor-like serine/threonine-protein kinase isoform D
G-type lectin S-receptor-like serine/threonine-protein kinase
Receptor-like serine/threonine-protein kinase SD1-8 isoform A
Receptor-like serine/threonine-protein kinase SD1-8
G-type lectin S-receptor-like serine/threonine-protein kinase SD2-5 isoform C
G-type lectin S-receptor-like serine/threonine-protein kinase
G-type lectin S-receptor-like serine/threonine-protein kinase CES101 isoform B
G-type lectin S-receptor-like serine/threonine-protein kinase CES101
G-type lectin S-receptor-like serine/threonine-protein kinase B120 isoform C
G-type lectin S-receptor-like serine/threonine-protein kinase isoform A
G-type lectin S-receptor-like serine/threonine-protein kinase SD2-5 isoform A
G-type lectin S-receptor-like serine/threonine-protein kinase isoform D
Receptor-like serine/threonine-protein kinase SD1-8 isoform D
G-type lectin S-receptor-like serine/threonine-protein kinase B120 isoform C
Receptor-like serine/threonine-protein kinase SD1-8
G-type lectin S-receptor-like serine/threonine-protein kinase isoform E
Glycosyltransferase family 31 protein
Uncharacterized protein
Uncharacterized protein

Uncharacterized protein
Uncharacterized protein
Uncharacterized protein
Uncharacterized protein
Uncharacterized protein
Uncharacterized protein
Uncharacterized protein
Uncharacterized protein
Uncharacterized protein
Uncharacterized protein
Uncharacterized protein
Uncharacterized protein
Uncharacterized protein
Uncharacterized protein
Uncharacterized protein
Uncharacterized protein
Uncharacterized protein
Uncharacterized protein
Uncharacterized protein
Uncharacterized protein
Uncharacterized protein
Uncharacterized protein
Uncharacterized protein
Uncharacterized protein
Uncharacterized protein
Uncharacterized protein
Uncharacterized protein
Uncharacterized protein
Uncharacterized protein
Uncharacterized protein
Uncharacterized protein
Uncharacterized protein
Uncharacterized protein
Uncharacterized protein
Uncharacterized protein
Uncharacterized protein
Uncharacterized protein
Uncharacterized protein
Uncharacterized protein
Uncharacterized protein
Uncharacterized protein

Uncharacterized protein
Uncharacterized protein
Uncharacterized protein
Uncharacterized protein
Hepatocyte growth factor
Macrophage stimulating 1
Macrophage stimulating 1
Macrophage stimulating 1
Macrophage stimulating 1
Plasminogen
Coagulation factor XI
Kallikrein B1
Hepatocyte growth factor
Kallikrein B1
Hepatocyte growth factor
Hepatocyte growth factor
Uncharacterized protein
Uncharacterized protein
Uncharacterized protein
Uncharacterized protein
Receptor-like serine/threonine-protein kinase SD1-8
Receptor-like serine/threonine-protein kinase SD1-8
G-type lectin S-receptor-like serine/threonine-protein kinase
G-type lectin S-receptor-like serine/threonine-protein kinase
Uncharacterized protein
G-type lectin S-receptor-like serine/threonine-protein kinase SD2-5
G-type lectin S-receptor-like serine/threonine-protein kinase SD2-5
G-type lectin S-receptor-like serine/threonine-protein kinase SD1-1
Uncharacterized protein
Uncharacterized protein
Uncharacterized protein
Uncharacterized protein
Uncharacterized protein
Uncharacterized protein
Uncharacterized protein
Uncharacterized protein
Uncharacterized protein
Uncharacterized protein
Uncharacterized protein
Uncharacterized protein
Uncharacterized protein

Uncharacterized protein
Uncharacterized protein
Uncharacterized protein
Uncharacterized protein
Uncharacterized protein
Uncharacterized protein
Uncharacterized protein
Uncharacterized protein
Uncharacterized protein
Uncharacterized protein
Uncharacterized protein
Uncharacterized protein
Uncharacterized protein
Uncharacterized protein
Uncharacterized protein
Uncharacterized protein
Uncharacterized protein
Uncharacterized protein
Uncharacterized protein
Uncharacterized protein
Uncharacterized protein
Uncharacterized protein
Uncharacterized protein
Uncharacterized protein
Uncharacterized protein
Uncharacterized protein
Uncharacterized protein
Uncharacterized protein
Uncharacterized protein
Uncharacterized protein
Uncharacterized protein
Uncharacterized protein
Uncharacterized protein
Uncharacterized protein
Uncharacterized protein
Uncharacterized protein
Uncharacterized protein
Uncharacterized protein
Uncharacterized protein
Uncharacterized protein
Uncharacterized protein

Uncharacterized protein
Uncharacterized protein
Uncharacterized protein
Uncharacterized protein
Uncharacterized protein
Uncharacterized protein
Uncharacterized protein
Uncharacterized protein
Uncharacterized protein
Uncharacterized protein
Uncharacterized protein
Uncharacterized protein
Uncharacterized protein
Uncharacterized protein
Uncharacterized protein
Uncharacterized protein
Uncharacterized protein
Uncharacterized protein
Uncharacterized protein
Uncharacterized protein
Uncharacterized protein
Uncharacterized protein
Uncharacterized protein
Uncharacterized protein
Uncharacterized protein
Uncharacterized protein
Uncharacterized protein
Uncharacterized protein
Uncharacterized protein
Uncharacterized protein
Uncharacterized protein
Uncharacterized protein
Uncharacterized protein
Uncharacterized protein
Uncharacterized protein
Uncharacterized protein
Uncharacterized protein
Uncharacterized protein
Uncharacterized protein
Uncharacterized protein
Uncharacterized protein

Uncharacterized protein
Uncharacterized protein
Uncharacterized protein
Uncharacterized protein
Uncharacterized protein
Uncharacterized protein
Uncharacterized protein
Uncharacterized protein
Uncharacterized protein
Uncharacterized protein
Uncharacterized protein
Uncharacterized protein
Uncharacterized protein
Uncharacterized protein
Uncharacterized protein
Uncharacterized protein
Uncharacterized protein
Uncharacterized protein
Uncharacterized protein
Uncharacterized protein
Uncharacterized protein
Uncharacterized protein
Uncharacterized protein
Uncharacterized protein
Uncharacterized protein
Uncharacterized protein
Uncharacterized protein
Uncharacterized protein
Uncharacterized protein
Uncharacterized protein
Uncharacterized protein
Uncharacterized protein
Uncharacterized protein
Uncharacterized protein
Uncharacterized protein
Uncharacterized protein
Uncharacterized protein
Uncharacterized protein
Uncharacterized protein
Uncharacterized protein
G-type lectin S-receptor-like serine/threonine-protein kinase At4g27290

G-type lectin S-receptor-like serine/threonine-protein kinase RKS1
G-type lectin S-receptor-like serine/threonine-protein kinase SD2-5
G-type lectin S-receptor-like serine/threonine-protein kinase At1g11300 isoform X2
G-type lectin S-receptor-like serine/threonine-protein kinase At4g27290
G-type lectin S-receptor-like serine/threonine-protein kinase SD2-5
G-type lectin S-receptor-like serine/threonine-protein kinase At1g11330
G-type lectin S-receptor-like serine/threonine-protein kinase RLK1
G-type lectin S-receptor-like serine/threonine-protein kinase B120
G-type lectin S-receptor-like serine/threonine-protein kinase RLK1
G-type lectin S-receptor-like serine/threonine-protein kinase RLK1
G-type lectin S-receptor-like serine/threonine-protein kinase SD3-1
G-type lectin S-receptor-like serine/threonine-protein kinase SD2-5
receptor-like serine/threonine-protein kinase SD1-8 isoform X3
G-type lectin S-receptor-like serine/threonine-protein kinase RLK1
G-type lectin S-receptor-like serine/threonine-protein kinase At1g11410
G-type lectin S-receptor-like serine/threonine-protein kinase CES101 isoform X2
receptor-like serine/threonine-protein kinase SD1-7
G-type lectin S-receptor-like serine/threonine-protein kinase At4g27290
receptor-like serine/threonine-protein kinase SD1-7
G-type lectin S-receptor-like serine/threonine-protein kinase SD2-2
G-type lectin S-receptor-like serine/threonine-protein kinase At4g03230
G-type lectin S-receptor-like serine/threonine-protein kinase At4g27290
G-type lectin S-receptor-like serine/threonine-protein kinase SD2-5 isoform X2
G-type lectin S-receptor-like serine/threonine-protein kinase At1g11300 isoform X1
G-type lectin S-receptor-like serine/threonine-protein kinase CES101
G-type lectin S-receptor-like serine/threonine-protein kinase At1g11410 isoform X2
G-type lectin S-receptor-like serine/threonine-protein kinase B120 isoform X1
G-type lectin S-receptor-like serine/threonine-protein kinase SD2-5 isoform X2
G-type lectin S-receptor-like serine/threonine-protein kinase SD2-5 isoform X2
G-type lectin S-receptor-like serine/threonine-protein kinase B120
G-type lectin S-receptor-like serine/threonine-protein kinase RLK1
G-type lectin S-receptor-like serine/threonine-protein kinase RLK1
G-type lectin S-receptor-like serine/threonine-protein kinase At4g03230
epidermis-specific secreted glycoprotein EP1-like
G-type lectin S-receptor-like serine/threonine-protein kinase SD2-5
G-type lectin S-receptor-like serine/threonine-protein kinase At4g27290
G-type lectin S-receptor-like serine/threonine-protein kinase RLK1
putative receptor protein kinase ZmPK1
epidermis-specific secreted glycoprotein EP1-like
receptor-like serine/threonine-protein kinase SD1-8 isoform X1
G-type lectin S-receptor-like serine/threonine-protein kinase SD2-5

G-type lectin S-receptor-like serine/threonine-protein kinase At1g11330
epidermis-specific secreted glycoprotein EP1-like
G-type lectin S-receptor-like serine/threonine-protein kinase At2g19130
G-type lectin S-receptor-like serine/threonine-protein kinase At1g11330 isoform X3
G-type lectin S-receptor-like serine/threonine-protein kinase At4g27290
G-type lectin S-receptor-like serine/threonine-protein kinase CES101 isoform X3
G-type lectin S-receptor-like serine/threonine-protein kinase At1g11410 isoform X1
G-type lectin S-receptor-like serine/threonine-protein kinase At4g27290
G-type lectin S-receptor-like serine/threonine-protein kinase At4g27290
LOW QUALITY PROTEIN: epidermis-specific secreted glycoprotein EP1-like
G-type lectin S-receptor-like serine/threonine-protein kinase At1g11410 isoform X1
Epidermis-specific secreted glycoprotein EP1-like protein
G-type lectin S-receptor-like serine/threonine-protein kinase SD3-1
G-type lectin S-receptor-like serine/threonine-protein kinase At1g11330
G-type lectin S-receptor-like serine/threonine-protein kinase At4g27290
G-type lectin S-receptor-like serine/threonine-protein kinase CES101 isoform X1
receptor-like serine/threonine-protein kinase SD1-8
G-type lectin S-receptor-like serine/threonine-protein kinase At1g11330
G-type lectin S-receptor-like serine/threonine-protein kinase SD2-5 isoform X1
G-type lectin S-receptor-like serine/threonine-protein kinase B120
receptor-like serine/threonine-protein kinase SD1-8 isoform X2
G-type lectin S-receptor-like serine/threonine-protein kinase At1g11410 isoform X2
G-type lectin S-receptor-like serine/threonine-protein kinase At4g27290
G-type lectin S-receptor-like serine/threonine-protein kinase At4g27290
G-type lectin S-receptor-like serine/threonine-protein kinase At4g27290
G-type lectin S-receptor-like serine/threonine-protein kinase At2g19130
G-type lectin S-receptor-like serine/threonine-protein kinase At1g11330
G-type lectin S-receptor-like serine/threonine-protein kinase CES101 isoform X1
PAN domain-containing protein At5g03700-like
PAN domain-containing protein At5g03700-like
G-type lectin S-receptor-like serine/threonine-protein kinase B120 isoform X2
G-type lectin S-receptor-like serine/threonine-protein kinase At4g27290
uncharacterized protein LOC107917261
G-type lectin S-receptor-like serine/threonine-protein kinase At1g11330
S-locus-specific glycoprotein S13-like
putative receptor protein kinase ZmPK1
G-type lectin S-receptor-like serine/threonine-protein kinase At1g11330
G-type lectin S-receptor-like serine/threonine-protein kinase B120
G-type lectin S-receptor-like serine/threonine-protein kinase At4g27290
G-type lectin S-receptor-like serine/threonine-protein kinase At1g34300
G-type lectin S-receptor-like serine/threonine-protein kinase RLK1

G-type lectin S-receptor-like serine/threonine-protein kinase At4g27290 isoform X1
receptor-like serine/threonine-protein kinase SD1-8
G-type lectin S-receptor-like serine/threonine-protein kinase At4g27290
G-type lectin S-receptor-like serine/threonine-protein kinase At4g27290
G-type lectin S-receptor-like serine/threonine-protein kinase At1g11330 isoform X2
G-type lectin S-receptor-like serine/threonine-protein kinase At1g11300
G-type lectin S-receptor-like serine/threonine-protein kinase At1g11330 isoform X1
G-type lectin S-receptor-like serine/threonine-protein kinase At4g27290
G-type lectin S-receptor-like serine/threonine-protein kinase At4g27290 isoform X1
epidermis-specific secreted glycoprotein EP1-like
G-type lectin S-receptor-like serine/threonine-protein kinase SD2-5
receptor-like serine/threonine-protein kinase SD1-8
G-type lectin S-receptor-like serine/threonine-protein kinase At4g03230
Cell division control protein 7-5
G-type lectin S-receptor-like serine/threonine-protein kinase SD2-2
putative receptor protein kinase ZmPK1
G-type lectin S-receptor-like serine/threonine-protein kinase At2g19130
G-type lectin S-receptor-like serine/threonine-protein kinase CES101 isoform X2
G-type lectin S-receptor-like serine/threonine-protein kinase B120
G-type lectin S-receptor-like serine/threonine-protein kinase At4g27290 isoform X2
G-type lectin S-receptor-like serine/threonine-protein kinase SD2-5
G-type lectin S-receptor-like serine/threonine-protein kinase At4g27290 isoform X2
G-type lectin S-receptor-like serine/threonine-protein kinase At2g19130
G-type lectin S-receptor-like serine/threonine-protein kinase SD2-5
G-type lectin S-receptor-like serine/threonine-protein kinase SD2-5 isoform X1
G-type lectin S-receptor-like serine/threonine-protein kinase At4g27290 isoform X3
G-type lectin S-receptor-like serine/threonine-protein kinase SD2-5 isoform X1
Uncharacterized protein (Fragment)
Uncharacterized protein
Uncharacterized protein
Uncharacterized protein
Uncharacterized protein
Uncharacterized protein (Fragment)
Uncharacterized protein
Uncharacterized protein
Uncharacterized protein
Uncharacterized protein
Uncharacterized protein
Uncharacterized protein (Fragment)
Uncharacterized protein
Uncharacterized protein

Uncharacterized protein
Uncharacterized protein
Uncharacterized protein (Fragment)
Uncharacterized protein
Uncharacterized protein (Fragment)
Uncharacterized protein
Uncharacterized protein
Uncharacterized protein
Uncharacterized protein (Fragment)
Uncharacterized protein
Uncharacterized protein
Uncharacterized protein
Uncharacterized protein (Fragment)
Uncharacterized protein
Uncharacterized protein
Uncharacterized protein
Uncharacterized protein
Uncharacterized protein
Uncharacterized protein
Uncharacterized protein
Uncharacterized protein
Uncharacterized protein
Uncharacterized protein (Fragment)
Uncharacterized protein
Uncharacterized protein
Uncharacterized protein
Uncharacterized protein
Uncharacterized protein
Uncharacterized protein
Uncharacterized protein
Uncharacterized protein
Uncharacterized protein
Uncharacterized protein
Uncharacterized protein
Uncharacterized protein
Uncharacterized protein
Uncharacterized protein
Uncharacterized protein
Uncharacterized protein
Uncharacterized protein (Fragment)
Uncharacterized protein

Uncharacterized protein
Uncharacterized protein
Uncharacterized protein
Uncharacterized protein
Uncharacterized protein
Uncharacterized protein
Uncharacterized protein
Uncharacterized protein
Uncharacterized protein
Uncharacterized protein
Uncharacterized protein
Uncharacterized protein
Uncharacterized protein
Uncharacterized protein
Uncharacterized protein
Uncharacterized protein
Uncharacterized protein
Uncharacterized protein
Uncharacterized protein
Uncharacterized protein
Uncharacterized protein
Uncharacterized protein
Uncharacterized protein
Uncharacterized protein
Uncharacterized protein
Uncharacterized protein
Uncharacterized protein
Uncharacterized protein
Uncharacterized protein
Uncharacterized protein
Uncharacterized protein
Uncharacterized protein
Uncharacterized protein (Fragment)
Uncharacterized protein
Uncharacterized protein
Uncharacterized protein
Uncharacterized protein
Uncharacterized protein
Uncharacterized protein
Uncharacterized protein
Uncharacterized protein

Uncharacterized protein
Uncharacterized protein
Uncharacterized protein
Uncharacterized protein
Uncharacterized protein
Uncharacterized protein
Uncharacterized protein
Uncharacterized protein
Uncharacterized protein
Uncharacterized protein
Uncharacterized protein
Uncharacterized protein
Uncharacterized protein
Uncharacterized protein (Fragment)
Uncharacterized protein (Fragment)
Uncharacterized protein (Fragment)
Uncharacterized protein (Fragment)
Uncharacterized protein (Fragment)
Uncharacterized protein (Fragment)
Uncharacterized protein (Fragment)
Uncharacterized protein
Uncharacterized protein (Fragment)
Uncharacterized protein (Fragment)
PAN domain protein
Uncharacterized protein (Fragment)
PAN domain protein
PAN domain protein
PAN domain protein
Uncharacterized protein
Uncharacterized protein
Uncharacterized protein
Uncharacterized protein
Peptidylprolyl isomerase
Uncharacterized protein
Uncharacterized protein
Uncharacterized protein
Uncharacterized protein
Uncharacterized protein
Uncharacterized protein (Fragment)
Uncharacterized protein
Uncharacterized protein (Fragment)

Uncharacterized protein (Fragment)
Uncharacterized protein
Plasminogen
Uncharacterized protein
Neurotrypsin
Uncharacterized protein
Uncharacterized protein
Uncharacterized protein (Fragment)
Cuticlin-1
Uncharacterized protein
PAN domain-containing protein
Uncharacterized protein
PAN domain-containing protein
PAN-1 domain containing protein (Fragment)
PAN-1 domain containing protein
Uncharacterized protein
Uncharacterized protein
Uncharacterized protein
Uncharacterized protein
Uncharacterized protein
Uncharacterized protein
Uncharacterized protein
Uncharacterized protein
Uncharacterized protein
Uncharacterized protein
Uncharacterized protein
Uncharacterized protein
Uncharacterized protein
Uncharacterized protein
Uncharacterized protein
Uncharacterized protein
Uncharacterized protein
Uncharacterized protein
Uncharacterized protein
Uncharacterized protein
Uncharacterized protein
Uncharacterized protein
Uncharacterized protein
Uncharacterized protein
Uncharacterized protein
Uncharacterized protein

Uncharacterized protein
Uncharacterized protein
Uncharacterized protein
Uncharacterized protein
Coagulation factor XI (Fragment)
Plasminogen (Fragment)
Plasminogen (Fragment)
Hepatocyte growth factor (Fragment)
PAN domain-containing protein
Uncharacterized protein
Uncharacterized protein
1,3(4)-beta-glucanase
Subtilisin SUB12
PAN domain-containing protein
PAN domain-containing protein
PAN domain-containing protein
Microneme protein, putative
Microneme protein MIC4
Microneme protein, putative
Uncharacterized protein
Uncharacterized protein
Microneme protein MIC17C
PAN domain protein
PAN domain-containing protein
PAN/Apple domain-containing protein
PAN domain-containing protein
Microneme protein MIC17C
Serine/threonine protein kinase
Non-specific serine/threonine protein kinase
Serine/threonine protein kinase
Serine/threonine protein kinase
Non-specific serine/threonine protein kinase
Serine/threonine protein kinase
Non-specific serine/threonine protein kinase
Uncharacterized protein
Non-specific serine/threonine protein kinase
Serine/threonine protein kinase
Serine/threonine protein kinase
Serine/threonine protein kinase
Serine/threonine protein kinase
Non-specific serine/threonine protein kinase

Serine/threonine protein kinase
Serine/threonine protein kinase
Non-specific serine/threonine protein kinase
Non-specific serine/threonine protein kinase
Serine/threonine protein kinase
Serine/threonine protein kinase
Serine/threonine protein kinase
Non-specific serine/threonine protein kinase
Non-specific serine/threonine protein kinase
Serine/threonine protein kinase
Non-specific serine/threonine protein kinase
Serine/threonine protein kinase
Serine/threonine protein kinase
Serine/threonine protein kinase
Serine/threonine protein kinase
Serine/threonine protein kinase
Uncharacterized protein
Serine/threonine protein kinase
Serine/threonine protein kinase
Serine/threonine protein kinase/TGF-beta stimulated factor
Serine/threonine protein kinase
Non-specific serine/threonine protein kinase
Serine/threonine protein kinase
Serine/threonine protein kinase
Non-specific serine/threonine protein kinase
Non-specific serine/threonine protein kinase
Serine/threonine protein kinase
Serine/threonine protein kinase
Serine/threonine protein kinase
Non-specific serine/threonine protein kinase
Serine/threonine protein kinase
MEKK
Serine/threonine protein kinase
Serine/threonine protein kinase
Serine/threonine protein kinase
Non-specific serine/threonine protein kinase
Serine/threonine protein kinase
Serine/threonine protein kinase
Serine/threonine protein kinase
Non-specific serine/threonine protein kinase
Serine/threonine protein kinase

Uncharacterized protein
Non-specific serine/threonine protein kinase
Serine/threonine protein kinase
Serine/threonine protein kinase
Serine/threonine protein kinase
Serine/threonine protein kinase
Serine/threonine protein kinase
Non-specific serine/threonine protein kinase
Non-specific serine/threonine protein kinase
Serine/threonine protein kinase
Serine/threonine protein kinase
Non-specific serine/threonine protein kinase
Serine/threonine protein kinase
Serine/threonine protein kinase
Serine/threonine protein kinase
Non-specific serine/threonine protein kinase
Serine/threonine protein kinase
Non-specific serine/threonine protein kinase
Serine/threonine protein kinase
Serine/threonine protein kinase
Serine/threonine protein kinase
Serine/threonine protein kinase
Serine/threonine protein kinase
Serine/threonine protein kinase
Serine/threonine protein kinase
Non-specific serine/threonine protein kinase
Serine/threonine protein kinase
Serine/threonine protein kinase
Serine/threonine protein kinase
Serine/threonine protein kinase
Serine/threonine protein kinase
Serine/threonine protein kinase
Serine/threonine protein kinase
Serine/threonine protein kinase
Non-specific serine/threonine protein kinase
Non-specific serine/threonine protein kinase
Serine/threonine protein kinase
Serine/threonine protein kinase
Uncharacterized protein
Serine/threonine protein kinase
Serine/threonine protein kinase

Serine/threonine protein kinase
Serine/threonine protein kinase
Serine/threonine protein kinase
Serine/threonine protein kinase
Serine/threonine protein kinase
Serine/threonine protein kinase
Serine/threonine protein kinase
MEKK
Serine/threonine protein kinase
Serine/threonine protein kinase
Serine/threonine protein kinase
Serine/threonine protein kinase
Serine/threonine protein kinase
Uncharacterized protein
Hepatocyte growth factor
Plasminogen
Uncharacterized protein
Uncharacterized protein
Hepatocyte growth factor
Uncharacterized protein
Uncharacterized protein
Uncharacterized protein
Cuticlin-1
Ras-related protein RabJ (Fragment)
Uncharacterized protein
Neurotrypsin
Uncharacterized protein
Uncharacterized protein
Putative EGF-like domain-containing protein
Putative S-locus glycoprotein domain-containing protein
Putative S-locus glycoprotein domain-containing protein
Putative EGF-like domain-containing protein
Putative S-locus glycoprotein domain-containing protein
Putative S-locus glycoprotein domain-containing protein
Putative bulb-type lectin domain-containing protein
Putative EGF-like domain-containing protein
Putative S-locus glycoprotein domain-containing protein
Putative S-locus glycoprotein domain, Bulb-type lectin domain protein
Putative S-locus glycoprotein domain-containing protein
Putative S-locus glycoprotein domain-containing protein
Putative S-locus glycoprotein domain-containing protein

Putative EGF-like domain-containing protein
Putative S-locus glycoprotein domain-containing protein
Putative bulb-type lectin domain-containing protein
Putative bulb-type lectin domain, PAN/Apple domain protein
Putative S-locus glycoprotein domain, Bulb-type lectin domain, PAN/Apple domain protein
Putative S-locus glycoprotein domain-containing protein
Putative S-locus glycoprotein domain-containing protein
Putative S-locus glycoprotein domain-containing protein
Putative S-domain-1 29
Putative S-locus glycoprotein domain-containing protein
Putative S-locus glycoprotein domain-containing protein
Putative S-locus glycoprotein domain-containing protein
Putative S-locus glycoprotein domain-containing protein
Putative S-locus glycoprotein domain-containing protein
Putative S-locus glycoprotein domain-containing protein
Putative S-locus glycoprotein domain-containing protein
Putative lectin protein kinase family protein
Putative S-locus-specific glycoprotein S6
Putative S-locus glycoprotein domain-containing protein
Putative S-locus lectin protein kinase family protein
Putative S-locus glycoprotein domain-containing protein
Putative S-locus glycoprotein domain-containing protein
Putative S-locus glycoprotein domain-containing protein
Putative EGF-like domain, S-locus glycoprotein domain, PAN/Apple domain protein
Putative EGF-like domain-containing protein
Uncharacterized protein
Uncharacterized protein
Uncharacterized protein
Putative GPCR kinase, PAN/Apple domain, Concanavalin A-like lectin/glucanase domain protein
Putative S-locus glycoprotein domain-containing protein
Putative EGF-like domain-containing protein
Putative S-locus glycoprotein domain-containing protein
Putative S-locus glycoprotein domain-containing protein
Putative S-locus glycoprotein domain-containing protein
Putative bulb-type lectin domain-containing protein
Putative bulb-type lectin domain-containing protein
Putative S-locus glycoprotein domain-containing protein
Putative EGF-like domain-containing protein
Putative S-locus glycoprotein domain-containing protein
Putative EGF-like domain-containing protein
Putative PAN/Apple domain-containing protein

Putative EGF-like domain-containing protein
Putative bulb-type lectin domain-containing protein
Putative S-locus glycoprotein domain-containing protein
Putative receptor kinase 3
Putative bulb-type lectin domain-containing protein
Putative PAN/Apple domain, Protein kinase-like domain protein
Putative EGF-like domain-containing protein
Putative bulb-type lectin domain-containing protein
Putative S-locus lectin protein kinase family protein
Putative S-locus glycoprotein domain-containing protein
Putative GPCR kinase
Putative epidermis-specific secreted glycoprotein EP1
Putative S-locus glycoprotein domain-containing protein
Putative S-locus-specific glycoprotein BS29-2
Putative S-locus glycoprotein domain-containing protein
Putative EGF-like domain-containing protein
Putative bulb-type lectin domain-containing protein
Putative EGF-like domain-containing protein
Putative S-locus lectin protein kinase family protein
Putative S-locus lectin protein kinase family protein
Putative bulb-type lectin domain-containing protein
Putative S-locus glycoprotein domain-containing protein
Putative S-locus glycoprotein domain-containing protein
Putative S-locus glycoprotein domain-containing protein
Putative EGF-like domain-containing protein
Putative S-locus glycoprotein domain-containing protein
Putative S-locus glycoprotein domain-containing protein
Putative curculin-like (Mannose-binding) lectin family protein / PAN domain-containing protein
Putative S-locus glycoprotein domain, PAN/Apple domain protein
Putative S-locus glycoprotein domain-containing protein
Putative S-locus glycoprotein domain-containing protein
Putative S-locus glycoprotein domain-containing protein
Putative S-locus glycoprotein domain, Bulb-type lectin domain, PAN/Apple domain protein
Putative EGF-like domain-containing protein
Putative S-locus glycoprotein domain-containing protein
Putative bulb-type lectin domain-containing protein
Putative S-locus glycoprotein domain-containing protein
Putative S-domain-2 5
Putative curculin-like (Mannose-binding) lectin family protein
Putative EGF-like domain-containing protein
Putative G-type lectin S-receptor-like serine/threonine-protein kinase

Putative S-locus glycoprotein domain-containing protein
Putative S-locus glycoprotein domain-containing protein
Putative S-locus glycoprotein domain-containing protein
Putative receptor-like protein kinase 4
Putative S-locus glycoprotein domain-containing protein
Putative S-locus glycoprotein domain-containing protein
Putative S-locus glycoprotein domain-containing protein
Putative bulb-type lectin domain-containing protein
Putative S-locus glycoprotein domain-containing protein
Putative PAN/Apple domain-containing protein
Putative S-locus glycoprotein domain-containing protein
Putative S-locus glycoprotein domain-containing protein
Putative S-locus glycoprotein domain-containing protein
Putative S-locus glycoprotein domain-containing protein
Putative EGF-like domain-containing protein
Putative EGF-like domain-containing protein
Putative PAN/Apple domain-containing protein
Putative S-locus glycoprotein domain, PAN/Apple domain protein
Putative EGF-like domain-containing protein
Putative bulb-type lectin domain-containing protein
Putative S-locus glycoprotein domain-containing protein
Putative bulb-type lectin domain-containing protein
Putative S-locus glycoprotein domain-containing protein
Putative EGF-like domain-containing protein
Putative S-locus lectin protein kinase family protein
Putative bulb-type lectin domain-containing protein
Putative S-locus glycoprotein domain, Bulb-type lectin domain, PAN/Apple domain protein
Putative S-locus glycoprotein domain-containing protein
Putative S-locus glycoprotein domain, Bulb-type lectin domain, PAN/Apple domain protein
Putative EGF-like domain-containing protein
Putative S-locus glycoprotein domain-containing protein
Putative S-locus glycoprotein domain-containing protein
Putative bulb-type lectin domain-containing protein
Putative serine/threonine/dual specificity protein kinase, catalytic domain-containing protein
Putative S-locus glycoprotein domain-containing protein
Putative PAN/Apple domain-containing protein
Putative S-locus glycoprotein domain-containing protein
Putative S-locus glycoprotein domain-containing protein
Putative S-locus glycoprotein domain-containing protein
Putative S-locus glycoprotein domain-containing protein
Putative S-locus glycoprotein domain-containing protein

Putative G-type lectin S-receptor-like serine/threonine-protein kinase
Putative S-locus glycoprotein domain-containing protein
Putative S-locus glycoprotein domain-containing protein
Putative S-locus glycoprotein domain-containing protein
Putative S-locus glycoprotein domain-containing protein
Putative S-locus glycoprotein domain, Bulb-type lectin domain, PAN/Apple domain protein
Putative EGF-like domain-containing protein
Putative bulb-type lectin domain-containing protein
Putative S-locus glycoprotein domain-containing protein
Putative S-locus glycoprotein domain, PAN/Apple domain protein
Uncharacterized protein
Putative G-type lectin S-receptor-like serine/threonine-protein kinase
Putative S-locus lectin protein kinase family protein
Putative EGF-like domain-containing protein
Putative EGF-like domain-containing protein
Putative GPCR kinase
Putative receptor kinase 1
Putative bulb-type lectin domain-containing protein
Putative EGF-like domain-containing protein
Putative serine/threonine/dual specificity protein kinase, catalytic domain-containing protein
Putative S-locus glycoprotein domain-containing protein
Putative PAN/Apple domain-containing protein
Putative S-locus glycoprotein domain-containing protein
Putative S-locus glycoprotein domain-containing protein
Putative S-locus lectin protein kinase family protein
Putative S-locus glycoprotein domain-containing protein
Putative EGF-like domain-containing protein
Putative bulb-type lectin domain-containing protein
Putative S-locus glycoprotein domain-containing protein
Putative PAN/Apple domain-containing protein
Putative bulb-type lectin domain, PAN/Apple domain protein
Putative S-locus glycoprotein domain, Bulb-type lectin domain, PAN/Apple domain protein
Uncharacterized protein
Putative S-locus glycoprotein domain-containing protein
Putative S-locus glycoprotein domain-containing protein
Putative bulb-type lectin domain-containing protein
Putative S-locus glycoprotein domain-containing protein
Putative PAN/Apple domain-containing protein
Putative EGF-like domain-containing protein
Putative S-locus glycoprotein domain-containing protein
Putative PAN/Apple domain-containing protein

Putative PAN/Apple domain-containing protein
Putative PAN/Apple domain-containing protein
Putative S-locus glycoprotein domain-containing protein
Putative S-locus glycoprotein domain, Bulb-type lectin domain, PAN/Apple domain protein
Putative S-locus glycoprotein domain-containing protein
Putative S-locus glycoprotein domain-containing protein
Putative S-locus glycoprotein domain-containing protein
Putative S-locus glycoprotein domain-containing protein
Putative EGF-like domain-containing protein
Putative S-locus glycoprotein domain-containing protein
Putative S-locus glycoprotein domain-containing protein
Putative bulb-type lectin domain-containing protein
Putative S-locus glycoprotein domain-containing protein
Putative receptor protein kinase ZmPK1
Putative bulb-type lectin domain-containing protein
Putative S-locus glycoprotein domain, Bulb-type lectin domain, PAN/Apple domain protein
Putative bulb-type lectin domain-containing protein
Putative S-locus glycoprotein domain, PAN/Apple domain protein
Putative S-locus glycoprotein domain-containing protein
Putative S-locus glycoprotein domain-containing protein
Putative GPCR kinase
Putative S-locus lectin protein kinase family protein
Uncharacterized protein
Uncharacterized protein
Cofilin/tropomyosin-type actin-binding protein (Fragment)
Uncharacterized protein (Fragment)
Uncharacterized protein
Uncharacterized protein
Uncharacterized protein
Uncharacterized protein
Uncharacterized protein
Uncharacterized protein
Uncharacterized protein
Uncharacterized protein
Uncharacterized protein
Uncharacterized protein
Uncharacterized protein
Uncharacterized protein
Uncharacterized protein
Uncharacterized protein
Uncharacterized protein

Uncharacterized protein
Uncharacterized protein
Uncharacterized protein
Uncharacterized protein
Uncharacterized protein
Uncharacterized protein
Uncharacterized protein
Uncharacterized protein
Uncharacterized protein
Uncharacterized protein
Uncharacterized protein
Uncharacterized protein
Uncharacterized protein
Uncharacterized protein
Uncharacterized protein
Uncharacterized protein
Uncharacterized protein (Fragment)
Uncharacterized protein
Uncharacterized protein
Uncharacterized protein
Uncharacterized protein
Uncharacterized protein
Uncharacterized protein
Uncharacterized protein
Uncharacterized protein
Uncharacterized protein
Uncharacterized protein
Uncharacterized protein
Uncharacterized protein (Fragment)
Uncharacterized protein
Uncharacterized protein
Uncharacterized protein (Fragment)
Uncharacterized protein
Uncharacterized protein
Uncharacterized protein
Uncharacterized protein
Uncharacterized protein (Fragment)
Uncharacterized protein
Uncharacterized protein
Uncharacterized protein
Uncharacterized protein

Uncharacterized protein
Uncharacterized protein
Uncharacterized protein
Uncharacterized protein
Uncharacterized protein
Uncharacterized protein
Uncharacterized protein
Uncharacterized protein
Uncharacterized protein
Uncharacterized protein
Uncharacterized protein
Uncharacterized protein
Uncharacterized protein
Uncharacterized protein
Uncharacterized protein
Uncharacterized protein
Uncharacterized protein
Uncharacterized protein
Uncharacterized protein
Uncharacterized protein
Uncharacterized protein
Uncharacterized protein
Uncharacterized protein
Uncharacterized protein
Uncharacterized protein
Uncharacterized protein
Uncharacterized protein
Uncharacterized protein
Uncharacterized protein
Uncharacterized protein
Uncharacterized protein
Uncharacterized protein
Uncharacterized protein
Uncharacterized protein
Uncharacterized protein
Uncharacterized protein
Uncharacterized protein
Uncharacterized protein
Uncharacterized protein
Uncharacterized protein
Uncharacterized protein

Uncharacterized protein
Uncharacterized protein
Uncharacterized protein
Uncharacterized protein
Uncharacterized protein
Uncharacterized protein
Uncharacterized protein
Uncharacterized protein
Uncharacterized protein
Uncharacterized protein
Uncharacterized protein
Uncharacterized protein
Uncharacterized protein
Uncharacterized protein
Uncharacterized protein
Uncharacterized protein
Uncharacterized protein
Uncharacterized protein
Uncharacterized protein
Uncharacterized protein
Uncharacterized protein
Uncharacterized protein
Uncharacterized protein
Uncharacterized protein
Uncharacterized protein
Uncharacterized protein
Uncharacterized protein
Uncharacterized protein
Uncharacterized protein
Uncharacterized protein
Uncharacterized protein
Uncharacterized protein
Uncharacterized protein
Uncharacterized protein
Uncharacterized protein
Uncharacterized protein
Uncharacterized protein
Uncharacterized protein
Uncharacterized protein
Uncharacterized protein
Uncharacterized protein

Uncharacterized protein
Uncharacterized protein
Uncharacterized protein
Uncharacterized protein
Uncharacterized protein
Uncharacterized protein
Uncharacterized protein
Uncharacterized protein
Uncharacterized protein
Uncharacterized protein
Uncharacterized protein
Uncharacterized protein
Uncharacterized protein
Uncharacterized protein
Uncharacterized protein
Uncharacterized protein
Uncharacterized protein
Uncharacterized protein
Uncharacterized protein
Uncharacterized protein
Uncharacterized protein
Uncharacterized protein
Uncharacterized protein
Uncharacterized protein
Uncharacterized protein
Uncharacterized protein
Uncharacterized protein
Plasma kallikrein preproprotein
Hepatocyte growth factor-like protein
Plasma kallikrein
Coagulation factor XI
Hepatocyte growth factor
Plasminogen isoform 1
Hepatocyte growth factor isoform 3
Hepatocyte growth factor-like protein
Plasminogen
Uncharacterized protein
Uncharacterized protein
Uncharacterized protein
Uncharacterized protein
Uncharacterized protein

Uncharacterized protein
Uncharacterized protein
Uncharacterized protein
Uncharacterized protein
Uncharacterized protein
Uncharacterized protein
Uncharacterized protein
Uncharacterized protein
Carbohydrate-binding WSC
Uncharacterized protein
Uncharacterized protein
Plasminogen
Hepatocyte growth factor
Uncharacterized protein
Serine-rich adhesin for platelets-like (Fragment)
S-locus related (Fragment)
S-locus related 1 (Fragment)
S-locus related (Fragment)
S-locus related (Fragment)
S-locus related (Fragment)
S-locus related 1 (Fragment)
Uncharacterized protein
Putative PAN domain-containing protein
Putative PAN domain-containing protein
Uncharacterized protein
Uncharacterized protein
Uncharacterized protein
Uncharacterized protein
Uncharacterized protein
Uncharacterized protein
Large extracellular alpha-helical protein
Membrane protein
Membrane protein (Fragment)
Large extracellular alpha-helical protein
Uncharacterized protein
Uncharacterized protein
Uncharacterized protein (Fragment)
Uncharacterized protein (Fragment)
Uncharacterized protein (Fragment)
Coagulation factor XI
Plasminogen

Plasma kallikrein
Hepatocyte growth factor-like protein
Hepatocyte growth factor
Plasminogen-like protein B
Plasminogen-like protein A
Putative macrophage stimulating 1-like protein
Hepatocyte growth factor isoform 6
Hepatocyte growth factor-like protein
cDNA FLJ53768, highly similar to Hepatocyte growth factor-like protein
Plasminogen
Coagulation factor XI (Fragment)
cDNA FLJ51250, highly similar to Plasma kallikrein
Plasma kallikrein (Fragment)
Hepatocyte growth factor
Hepatocyte growth factor (Fragment)
Plasminogen
Hepatocyte growth factor isoform 1 preproprotein variant (Fragment)
Coagulation factor XI (Fragment)
Plasma kallikrein (Fragment)
Hepatocyte growth factor (Fragment)
Plasminogen-like protein B
Plasminogen-related protein A (Fragment)
Uncharacterized protein PLGL (Fragment)
Plasma kallikrein
cDNA FLJ61328, highly similar to Hepatocyte growth factor-like protein
cDNA FLJ61329, moderately similar to Hepatocyte growth factor-like protein
Hepatocyte growth factor
Plasminogen (Fragment)
cDNA FLJ60766, highly similar to Hepatocyte growth factor-like protein
cDNA FLJ77744, highly similar to Homo sapiens kallikrein B, plasma (Fletcher factor) 1 (KLKB1), mRNA
Plasma kallikrein (Fragment)
Coagulation factor XI
HCG2029799, isoform CRA_b
Hepatocyte growth factor isoform 6
cDNA FLJ56324, highly similar to Hepatocyte growth factor-like protein
Hepatocyte growth factor-like protein homolog
Macrophage stimulating 1 (Hepatocyte growth factor-like) variant (Fragment)
Uncharacterized protein
Uncharacterized protein
Protein kinase, putative
Uncharacterized protein

Uncharacterized protein (Fragment)
Uncharacterized protein
Alpha-1,3-fucosyltransferase 10
Integrin alpha-D
Uncharacterized protein
Matrilin-4
Uncharacterized protein (Fragment)
Collagen alpha-1XIV chain
Tubulin--tyrosine ligase
E3 ubiquitin-protein ligase HERC2
Uncharacterized protein
Collagen alpha-1XIV chain
Matrilin-2
Uncharacterized protein
Uncharacterized protein
Uncharacterized protein (Fragment)
Opioid growth factor receptor
Importin subunit alpha
Arylsulfatase
Uncharacterized protein
Collagen alpha-1XX chain
Uncharacterized protein
Uncharacterized protein (Fragment)
Uncharacterized protein
Tubulin polyglutamylase ttll6
G8 domain-containing protein DDB_G0286897
Uncharacterized protein
Uncharacterized protein
Uncharacterized protein (Fragment)
Uncharacterized protein
Disintegrin and metalloproteinase domain-containing protein B
Predicted protein
Uncharacterized protein
Uncharacterized protein
Predicted protein
Serine/threonine-protein kinase
Serine/threonine-protein kinase
Uncharacterized protein
Predicted protein
Uncharacterized protein
Uncharacterized protein

Uncharacterized protein
Uncharacterized protein
Uncharacterized protein
Uncharacterized protein
Serine/threonine-protein kinase
Uncharacterized protein
Uncharacterized protein
Uncharacterized protein
Uncharacterized protein
Uncharacterized protein
Predicted protein
Serine/threonine-protein kinase
Uncharacterized protein
Predicted protein
Serine/threonine-protein kinase
Uncharacterized protein
Uncharacterized protein
Uncharacterized protein
Serine/threonine-protein kinase
Uncharacterized protein
Serine/threonine-protein kinase
Serine/threonine-protein kinase
Serine/threonine-protein kinase
Uncharacterized protein
Serine/threonine-protein kinase
Uncharacterized protein
Uncharacterized protein
Uncharacterized protein
Uncharacterized protein
Serine/threonine-protein kinase
Uncharacterized protein
Serine/threonine-protein kinase
Uncharacterized protein
Serine/threonine-protein kinase
Serine/threonine-protein kinase
Serine/threonine-protein kinase
Uncharacterized protein
Serine/threonine-protein kinase
Serine/threonine-protein kinase
Predicted protein
Uncharacterized protein

Serine/threonine-protein kinase
Uncharacterized protein
Uncharacterized protein
Serine/threonine-protein kinase
Uncharacterized protein
Serine/threonine-protein kinase
Uncharacterized protein
Uncharacterized protein
Uncharacterized protein
Uncharacterized protein
Uncharacterized protein
Serine/threonine-protein kinase
Serine/threonine-protein kinase
Uncharacterized protein
Uncharacterized protein
Serine/threonine-protein kinase
Serine/threonine-protein kinase
Serine/threonine-protein kinase
Serine/threonine-protein kinase
Uncharacterized protein
Serine/threonine-protein kinase
Uncharacterized protein
Uncharacterized protein
Uncharacterized protein
Uncharacterized protein
Uncharacterized protein
Uncharacterized protein
Serine/threonine-protein kinase
Uncharacterized protein
Serine/threonine-protein kinase
Uncharacterized protein
Uncharacterized protein
Uncharacterized protein
Uncharacterized protein
Serine/threonine-protein kinase
Serine/threonine-protein kinase
Uncharacterized protein
Uncharacterized protein
Uncharacterized protein
Uncharacterized protein
Serine/threonine-protein kinase

Uncharacterized protein
Uncharacterized protein
Uncharacterized protein
Uncharacterized protein
Uncharacterized protein
Uncharacterized protein
Predicted protein
Uncharacterized protein
Uncharacterized protein
Predicted protein
Uncharacterized protein
Uncharacterized protein
Serine/threonine-protein kinase
Uncharacterized protein
Uncharacterized protein
Uncharacterized protein
Uncharacterized protein
Uncharacterized protein
Predicted protein
Uncharacterized protein
Serine/threonine-protein kinase
Serine/threonine-protein kinase
Uncharacterized protein
Uncharacterized protein
Serine/threonine-protein kinase
Predicted protein
Uncharacterized protein
Uncharacterized protein
Predicted protein
Uncharacterized protein
Uncharacterized protein
Uncharacterized protein
Uncharacterized protein
Uncharacterized protein
Uncharacterized protein
Uncharacterized protein
Serine/threonine-protein kinase
Uncharacterized protein
Uncharacterized protein
Uncharacterized protein
Uncharacterized protein

Uncharacterized protein
Uncharacterized protein
Uncharacterized protein
Uncharacterized protein
Uncharacterized protein
Uncharacterized protein
Uncharacterized protein
Uncharacterized protein
Uncharacterized protein
Predicted protein (Fragment)
Uncharacterized protein
Uncharacterized protein
Uncharacterized protein
Uncharacterized protein
Uncharacterized protein
Predicted protein
Serine/threonine-protein kinase
Serine/threonine-protein kinase
Uncharacterized protein
Uncharacterized protein
Uncharacterized protein
Uncharacterized protein
Serine/threonine-protein kinase
Uncharacterized protein
Uncharacterized protein
Uncharacterized protein
Uncharacterized protein
Serine/threonine-protein kinase
Uncharacterized protein
Serine/threonine-protein kinase
Serine/threonine-protein kinase
Uncharacterized protein
Serine/threonine-protein kinase
Uncharacterized protein
Predicted protein
Uncharacterized protein
Uncharacterized protein
Uncharacterized protein
Uncharacterized protein
Serine/threonine-protein kinase
Uncharacterized protein

Serine/threonine-protein kinase
Serine/threonine-protein kinase
Uncharacterized protein
Serine/threonine-protein kinase
Uncharacterized protein
Uncharacterized protein
Uncharacterized protein
Uncharacterized protein
Serine/threonine-protein kinase
Uncharacterized protein
Uncharacterized protein
Serine/threonine-protein kinase
Uncharacterized protein
Uncharacterized protein
Serine/threonine-protein kinase
Uncharacterized protein
Uncharacterized protein
Uncharacterized protein
Predicted protein
Uncharacterized protein
Uncharacterized protein
Uncharacterized protein
Uncharacterized protein
Uncharacterized protein
Uncharacterized protein
Uncharacterized protein
Uncharacterized protein
Uncharacterized protein
Uncharacterized protein
Uncharacterized protein
Serine/threonine-protein kinase
Uncharacterized protein
Uncharacterized protein
Uncharacterized protein
Uncharacterized protein
Uncharacterized protein
Predicted protein
Serine/threonine-protein kinase
Uncharacterized protein
Uncharacterized protein
Serine/threonine-protein kinase

Uncharacterized protein
Uncharacterized protein
Uncharacterized protein
Uncharacterized protein
Serine/threonine-protein kinase
Serine/threonine-protein kinase
Uncharacterized protein
Uncharacterized protein
Uncharacterized protein
Uncharacterized protein
Serine/threonine-protein kinase
Uncharacterized protein
Uncharacterized protein
Serine/threonine-protein kinase
Uncharacterized protein
Serine/threonine-protein kinase
Uncharacterized protein
Uncharacterized protein
Uncharacterized protein
Uncharacterized protein
Uncharacterized protein
Uncharacterized protein
Uncharacterized protein
Uncharacterized protein
Uncharacterized protein
Serine/threonine-protein kinase
Serine/threonine-protein kinase
Uncharacterized protein
Uncharacterized protein
Serine/threonine-protein kinase
Uncharacterized protein
Serine/threonine-protein kinase
Serine/threonine-protein kinase
Uncharacterized protein
Uncharacterized protein
Serine/threonine-protein kinase
Uncharacterized protein
Uncharacterized protein
Predicted protein
Uncharacterized protein
Uncharacterized protein

Uncharacterized protein
Uncharacterized protein
Uncharacterized protein
Uncharacterized protein
Uncharacterized protein
Uncharacterized protein
Uncharacterized protein
Predicted protein (Fragment)
Predicted protein
Uncharacterized protein
Uncharacterized protein
Serine/threonine-protein kinase
Uncharacterized protein
Uncharacterized protein
Uncharacterized protein
Uncharacterized protein
Uncharacterized protein
Uncharacterized protein
Uncharacterized protein
Uncharacterized protein
Predicted protein
Uncharacterized protein
Serine/threonine-protein kinase
Predicted protein
Uncharacterized protein
Uncharacterized protein
Uncharacterized protein
Uncharacterized protein
Uncharacterized protein
Uncharacterized protein
Uncharacterized protein
Uncharacterized protein
Predicted protein
Uncharacterized protein
Uncharacterized protein
Uncharacterized protein
Uncharacterized protein
Uncharacterized protein
Uncharacterized protein
Serine/threonine-protein kinase
Uncharacterized protein

Uncharacterized protein
Uncharacterized protein
Predicted protein
Uncharacterized protein
Uncharacterized protein
Predicted protein
Uncharacterized protein
Uncharacterized protein
Uncharacterized protein
Predicted protein
Predicted protein
Predicted protein
Uncharacterized protein
Uncharacterized protein
Uncharacterized protein
Predicted protein
Uncharacterized protein
Uncharacterized protein
Predicted protein
Uncharacterized protein
Uncharacterized protein
Predicted protein
Uncharacterized protein
Uncharacterized protein
Uncharacterized protein
Predicted protein
Uncharacterized protein
Uncharacterized protein
Serine/threonine-protein kinase
Uncharacterized protein
Uncharacterized protein
Uncharacterized protein
Uncharacterized protein
Uncharacterized protein
Predicted protein (Fragment)
Uncharacterized protein
Uncharacterized protein
Serine/threonine-protein kinase
Uncharacterized protein
Uncharacterized protein
Predicted protein

Uncharacterized protein
Uncharacterized protein
Predicted protein
Uncharacterized protein
Uncharacterized protein
Uncharacterized protein
Uncharacterized protein
Uncharacterized protein
Uncharacterized protein
Uncharacterized protein
Uncharacterized protein
Serine/threonine-protein kinase
Predicted protein (Fragment)
Uncharacterized protein
Serine/threonine-protein kinase
Uncharacterized protein
Uncharacterized protein
Uncharacterized protein
Uncharacterized protein
Serine/threonine-protein kinase
Uncharacterized protein
Uncharacterized protein
Uncharacterized protein
Uncharacterized protein
Uncharacterized protein
Uncharacterized protein
Serine/threonine-protein kinase
Uncharacterized protein
Serine/threonine-protein kinase
Uncharacterized protein
Uncharacterized protein
Serine/threonine-protein kinase
Uncharacterized protein
Uncharacterized protein
Uncharacterized protein
Uncharacterized protein
Predicted protein
Uncharacterized protein
Uncharacterized protein
Serine/threonine-protein kinase
Uncharacterized protein

Uncharacterized protein
Uncharacterized protein
Uncharacterized protein
Predicted protein
Uncharacterized protein
Uncharacterized protein
Uncharacterized protein
Serine/threonine-protein kinase
Uncharacterized protein
Uncharacterized protein
Uncharacterized protein
Serine/threonine-protein kinase
Serine/threonine-protein kinase
Uncharacterized protein
Predicted protein
Predicted protein
Uncharacterized protein
Serine/threonine-protein kinase
Serine/threonine-protein kinase
Uncharacterized protein
Uncharacterized protein
Uncharacterized protein
Serine/threonine-protein kinase
Uncharacterized protein
Uncharacterized protein
Serine/threonine-protein kinase
Predicted protein
Uncharacterized protein
Predicted protein (Fragment)
Serine/threonine-protein kinase
Uncharacterized protein
Uncharacterized protein
Uncharacterized protein
Uncharacterized protein
Serine/threonine-protein kinase
Uncharacterized protein
Serine/threonine-protein kinase
Serine/threonine-protein kinase
Uncharacterized protein
Uncharacterized protein
Uncharacterized protein

Uncharacterized protein
Uncharacterized protein
Uncharacterized protein
Uncharacterized protein
Uncharacterized protein
Serine/threonine-protein kinase
Uncharacterized protein
Serine/threonine-protein kinase
Uncharacterized protein
Serine/threonine-protein kinase
Uncharacterized protein
Uncharacterized protein
Uncharacterized protein
Uncharacterized protein
Serine/threonine-protein kinase
Uncharacterized protein
Uncharacterized protein
Uncharacterized protein
Uncharacterized protein
Uncharacterized protein
Uncharacterized protein
Uncharacterized protein
Uncharacterized protein
Serine/threonine-protein kinase
Uncharacterized protein
Uncharacterized protein
Serine/threonine-protein kinase
Uncharacterized protein
Uncharacterized protein
Uncharacterized protein
Uncharacterized protein
Predicted protein
Predicted protein
Serine/threonine-protein kinase
Serine/threonine-protein kinase
Uncharacterized protein
Uncharacterized protein
Uncharacterized protein
Serine/threonine-protein kinase
Uncharacterized protein
Uncharacterized protein

Uncharacterized protein
Uncharacterized protein
Uncharacterized protein
Uncharacterized protein
Uncharacterized protein
Uncharacterized protein
Uncharacterized protein
Uncharacterized protein
Uncharacterized protein
Uncharacterized protein
Uncharacterized protein
Uncharacterized protein
Predicted protein
Uncharacterized protein
Uncharacterized protein
Serine/threonine-protein kinase
Serine/threonine-protein kinase
Uncharacterized protein
Serine/threonine-protein kinase
Uncharacterized protein
Uncharacterized protein
Uncharacterized protein
Predicted protein
Predicted protein
Uncharacterized protein
Uncharacterized protein
Uncharacterized protein
Uncharacterized protein
Uncharacterized protein
Uncharacterized protein
Uncharacterized protein
Predicted protein
Serine/threonine-protein kinase
Serine/threonine-protein kinase
Serine/threonine-protein kinase
Uncharacterized protein
Predicted protein (Fragment)
Uncharacterized protein
Serine/threonine-protein kinase
Uncharacterized protein
Uncharacterized protein

Uncharacterized protein
Serine/threonine-protein kinase
Uncharacterized protein
Uncharacterized protein
Serine/threonine-protein kinase
Uncharacterized protein
Uncharacterized protein
Uncharacterized protein
Uncharacterized protein
Uncharacterized protein
Uncharacterized protein
Uncharacterized protein
Uncharacterized protein
Uncharacterized protein
Uncharacterized protein
Uncharacterized protein
Uncharacterized protein
Uncharacterized protein
Uncharacterized protein
Serine/threonine-protein kinase
Serine/threonine-protein kinase
Uncharacterized protein
Serine/threonine-protein kinase
Uncharacterized protein
Uncharacterized protein
Uncharacterized protein
Uncharacterized protein
Serine/threonine-protein kinase
Uncharacterized protein
Uncharacterized protein
Uncharacterized protein
Uncharacterized protein
Uncharacterized protein
Uncharacterized protein
Uncharacterized protein
Uncharacterized protein
Uncharacterized protein
Uncharacterized protein
Uncharacterized protein
Uncharacterized protein
Uncharacterized protein

Uncharacterized protein
Uncharacterized protein
Uncharacterized protein
Uncharacterized protein
Uncharacterized protein
Uncharacterized protein
Uncharacterized protein
Uncharacterized protein
Uncharacterized protein
Uncharacterized protein
Uncharacterized protein
Uncharacterized protein
Uncharacterized protein
Uncharacterized protein (Fragment)
Uncharacterized protein
Uncharacterized protein
Uncharacterized protein
Uncharacterized protein
Uncharacterized protein
Uncharacterized protein
Uncharacterized protein
Uncharacterized protein
Uncharacterized protein
Uncharacterized protein
Uncharacterized protein
Uncharacterized protein
Uncharacterized protein (Fragment)
Uncharacterized protein
Uncharacterized protein
Uncharacterized protein
Uncharacterized protein
Uncharacterized protein
Uncharacterized protein
Uncharacterized protein
Uncharacterized protein
Uncharacterized protein
Uncharacterized protein
Uncharacterized protein
Uncharacterized protein
Uncharacterized protein
Uncharacterized protein

Uncharacterized protein
Uncharacterized protein
Uncharacterized protein
Uncharacterized protein
Uncharacterized protein
Uncharacterized protein
Uncharacterized protein
Uncharacterized protein
Uncharacterized protein
Uncharacterized protein
Uncharacterized protein
Uncharacterized protein
Uncharacterized protein
Uncharacterized protein
Uncharacterized protein
Uncharacterized protein
Uncharacterized protein
Uncharacterized protein
Uncharacterized protein (Fragment)
Uncharacterized protein
Uncharacterized protein
Uncharacterized protein
Uncharacterized protein
Uncharacterized protein (Fragment)
Uncharacterized protein
Uncharacterized protein
Uncharacterized protein
Uncharacterized protein
Uncharacterized protein
Uncharacterized protein
Uncharacterized protein
Uncharacterized protein
Uncharacterized protein
Uncharacterized protein
Uncharacterized protein
Uncharacterized protein
Uncharacterized protein
Uncharacterized protein
Uncharacterized protein
Uncharacterized protein
Uncharacterized protein

Uncharacterized protein
Uncharacterized protein
High-affinity leucine-specific transport system, periplasmic binding protein LivK
Uncharacterized protein
Uncharacterized protein
Uncharacterized protein (Fragment)
Uncharacterized protein
Uncharacterized protein
Uncharacterized protein
Uncharacterized protein
Uncharacterized protein
Uncharacterized protein
Uncharacterized protein
Uncharacterized protein
CnPolydom (Fragment)
Uncharacterized protein
Uncharacterized protein (Fragment)
Uncharacterized protein
PAN domain protein
Uncharacterized protein
Uncharacterized protein
PAN domain protein
Uncharacterized protein
Uncharacterized protein
Uncharacterized protein
PAN domain-containing protein
Uncharacterized protein
Uncharacterized protein
Uncharacterized protein
Uncharacterized protein
Uncharacterized protein
Uncharacterized protein
Uncharacterized protein
Uncharacterized protein
Uncharacterized protein
Uncharacterized protein
Uncharacterized protein
Uncharacterized protein
Uncharacterized protein
Uncharacterized protein
Uncharacterized protein

Uncharacterized protein
Uncharacterized protein
Uncharacterized protein
Uncharacterized protein
Uncharacterized protein
Uncharacterized protein
Uncharacterized protein
Conserved domain protein
Uncharacterized protein
Uncharacterized protein
Uncharacterized protein
Uncharacterized protein (Fragment)
Uncharacterized protein
Uncharacterized protein
Uncharacterized protein
Uncharacterized protein
Uncharacterized protein
Uncharacterized protein
Uncharacterized protein
Uncharacterized protein
Uncharacterized protein
Uncharacterized protein
Uncharacterized protein
Uncharacterized protein
Glycosyltransferase family 31 protein
Uncharacterized protein
Uncharacterized protein
Family 31 glycosyltransferase
Glycosyltransferase family 31
Predicted protein
Uncharacterized protein
Glycosyltransferase family 31 protein
Uncharacterized protein
Uncharacterized protein
Glycosyltransferase family 31 protein
Uncharacterized protein (Fragment)
Uncharacterized protein
Uncharacterized protein
Uncharacterized protein
Glycosyltransferase family 31 protein
Glycosyltransferase family 31 protein

Uncharacterized protein
Uncharacterized protein
Uncharacterized protein
Uncharacterized protein
Putative Vascular endothelial growth factor receptor 2
Uncharacterized protein
Uncharacterized protein
Uncharacterized protein
Uncharacterized protein (Fragment)
Putative Vascular endothelial growth factor receptor 2
Uncharacterized protein
Uncharacterized protein
Uncharacterized protein
Uncharacterized protein
Uncharacterized protein
Uncharacterized protein
Uncharacterized protein
Uncharacterized protein
Uncharacterized protein
Uncharacterized protein
Uncharacterized protein
Uncharacterized protein
Uncharacterized protein
Uncharacterized protein
Uncharacterized protein
Metalloendopeptidase
Putative glucan 1,3-beta-glucosidase A
Uncharacterized protein
Uncharacterized protein
Uncharacterized protein
Endoglucanase-5
Uncharacterized protein
Uncharacterized protein
Uncharacterized protein (Fragment)
Uncharacterized protein (Fragment)
hepatocyte growth factor-like protein isoform X1
Plasminogen
plasminogen
coagulation factor XI-like isoform X4
plasma kallikrein-like isoform X2
hepatocyte growth factor-like protein isoform X2

plasma kallikrein-like isoform X1
G-protein coupled receptor 98
coagulation factor XI-like isoform X3
uncharacterized protein LOC108257189 isoform X1
coagulation factor XI-like isoform X5
coagulation factor XI-like isoform X2
Hepatocyte growth factor-like protein
Hepatocyte growth factor
Uncharacterized protein
Plasminogen
Hepatocyte growth factor
Macrophage stimulating 1
Uncharacterized protein
PAN domain protein
Uncharacterized protein
Uncharacterized protein
Receptor protein kinase
Secreted glycoprotein 3
Secreted glycoprotein 2
Secreted glycoprotein 1
Putative sortilin-related receptor (Fragment)
Putative conserved secreted protein
Uncharacterized protein (Fragment)
Uncharacterized protein (Fragment)
Putative conserved secreted protein (Fragment)
Putative neurotrypsin
Ixominogen
Uncharacterized protein (Fragment)
Putative antigen b membrane protein (Fragment)
Putative conserved secreted protein
No-mechanoreceptor potential A, putative
PAN domaincontaining protein
PAN domaincontaining protein (Fragment)
Uncharacterized protein (Fragment) (Fragment)
PAN domaincontaining protein
PAN domaincontaining protein
PAN domaincontaining protein
Antigen b membrane protein, putative
Serine protease family protein
Uncharacterized protein (Fragment) (Fragment)
PAN domaincontaining protein

Uncharacterized protein
Antigen B membrane protein, putative
Uncharacterized protein
Uncharacterized protein
Uncharacterized protein
Alpha-2-macroglobulin-like protein
Uncharacterized protein
PAN domain-containing protein
Uncharacterized protein
Uncharacterized protein
Uncharacterized protein
Uncharacterized protein
Uncharacterized protein
Uncharacterized protein
Uncharacterized protein
Uncharacterized protein
Uncharacterized protein
Uncharacterized protein
Uncharacterized protein
Uncharacterized protein
Uncharacterized protein
Uncharacterized protein
Uncharacterized protein
Uncharacterized protein
Uncharacterized protein
Uncharacterized protein
Uncharacterized protein
Uncharacterized protein
Uncharacterized protein
Uncharacterized protein
Uncharacterized protein
Uncharacterized protein
Uncharacterized protein
Uncharacterized protein
Uncharacterized protein
Uncharacterized protein
Uncharacterized protein
Uncharacterized protein
Uncharacterized protein
Uncharacterized protein
Uncharacterized protein

Uncharacterized protein
Uncharacterized protein
Uncharacterized protein
Uncharacterized protein
Uncharacterized protein
Uncharacterized protein
Uncharacterized protein
Uncharacterized protein
Uncharacterized protein
Uncharacterized protein
Uncharacterized protein
Uncharacterized protein
Uncharacterized protein
Uncharacterized protein
Uncharacterized protein
Uncharacterized protein
Uncharacterized protein
Uncharacterized protein
Uncharacterized protein
Uncharacterized protein
Uncharacterized protein
Uncharacterized protein
Uncharacterized protein
Uncharacterized protein
Uncharacterized protein
Uncharacterized protein
Uncharacterized protein
Uncharacterized protein
Uncharacterized protein
putative receptor protein kinase ZmPK1 isoform X1
G-type lectin S-receptor-like serine/threonine-protein kinase At1g11410
G-type lectin S-receptor-like serine/threonine-protein kinase At1g34300 isoform X1
G-type lectin S-receptor-like serine/threonine-protein kinase At4g03230 isoform X1
G-type lectin S-receptor-like serine/threonine-protein kinase At4g27290 isoform X1
G-type lectin S-receptor-like serine/threonine-protein kinase SD3-1
G-type lectin S-receptor-like serine/threonine-protein kinase LECRK1
G-type lectin S-receptor-like serine/threonine-protein kinase At4g27290 isoform X4
G-type lectin S-receptor-like serine/threonine-protein kinase RKS1
G-type lectin S-receptor-like serine/threonine-protein kinase At4g27290 isoform X5
G-type lectin S-receptor-like serine/threonine-protein kinase CES101 isoform X1
G-type lectin S-receptor-like serine/threonine-protein kinase At4g27290 isoform X1

G-type lectin S-receptor-like serine/threonine-protein kinase CES101 isoform X1
G-type lectin S-receptor-like serine/threonine-protein kinase CES101 isoform X2
G-type lectin S-receptor-like serine/threonine-protein kinase SD2-5
putative receptor protein kinase ZmPK1
G-type lectin S-receptor-like serine/threonine-protein kinase At4g03230 isoform X2
G-type lectin S-receptor-like serine/threonine-protein kinase At4g27290 isoform X1
G-type lectin S-receptor-like serine/threonine-protein kinase At4g27290 isoform X2
G-type lectin S-receptor-like serine/threonine-protein kinase LECRK3 isoform X1
putative receptor protein kinase ZmPK1
G-type lectin S-receptor-like serine/threonine-protein kinase RKS1 isoform X2
G-type lectin S-receptor-like serine/threonine-protein kinase At1g11330 isoform X3
G-type lectin S-receptor-like serine/threonine-protein kinase At4g27290
G-type lectin S-receptor-like serine/threonine-protein kinase LECRK1
putative receptor protein kinase ZmPK1
receptor-like serine/threonine-protein kinase SD1-6
receptor-like serine/threonine-protein kinase SD1-8
G-type lectin S-receptor-like serine/threonine-protein kinase At4g27290 isoform X2
receptor-like serine/threonine-protein kinase SD1-8
G-type lectin S-receptor-like serine/threonine-protein kinase LECRK1
G-type lectin S-receptor-like serine/threonine-protein kinase B120 isoform X1
G-type lectin S-receptor-like serine/threonine-protein kinase At4g27290 isoform X1
G-type lectin S-receptor-like serine/threonine-protein kinase At4g27290
receptor-like serine/threonine-protein kinase SD1-6 isoform X3
G-type lectin S-receptor-like serine/threonine-protein kinase At1g11410 isoform X1
PAN domain-containing protein At5g03700-like
G-type lectin S-receptor-like serine/threonine-protein kinase At1g11410 isoform X2
G-type lectin S-receptor-like serine/threonine-protein kinase At4g27290
G-type lectin S-receptor-like serine/threonine-protein kinase RKS1 isoform X4
G-type lectin S-receptor-like serine/threonine-protein kinase CES101 isoform X1
G-type lectin S-receptor-like serine/threonine-protein kinase At4g27290 isoform X2
G-type lectin S-receptor-like serine/threonine-protein kinase At4g27290 isoform X2
G-type lectin S-receptor-like serine/threonine-protein kinase RKS1 isoform X2
putative receptor protein kinase ZmPK1
PAN domain-containing protein At5g03700-like
G-type lectin S-receptor-like serine/threonine-protein kinase At4g27290 isoform X1
G-type lectin S-receptor-like serine/threonine-protein kinase CES101 isoform X2
G-type lectin S-receptor-like serine/threonine-protein kinase At1g11410 isoform X1
G-type lectin S-receptor-like serine/threonine-protein kinase At1g11410 isoform X2
putative receptor protein kinase ZmPK1
G-type lectin S-receptor-like serine/threonine-protein kinase At1g11330 isoform X1
G-type lectin S-receptor-like serine/threonine-protein kinase CES101 isoform X1

G-type lectin S-receptor-like serine/threonine-protein kinase SD2-5
G-type lectin S-receptor-like serine/threonine-protein kinase At1g11330
G-type lectin S-receptor-like serine/threonine-protein kinase At1g11330
G-type lectin S-receptor-like serine/threonine-protein kinase CES101 isoform X2
G-type lectin S-receptor-like serine/threonine-protein kinase At2g19130
G-type lectin S-receptor-like serine/threonine-protein kinase At1g11410 isoform X2
G-type lectin S-receptor-like serine/threonine-protein kinase At1g11330 isoform X2
putative receptor protein kinase ZmPK1
G-type lectin S-receptor-like serine/threonine-protein kinase RKS1
G-type lectin S-receptor-like serine/threonine-protein kinase At1g11410 isoform X3
G-type lectin S-receptor-like serine/threonine-protein kinase CES101 isoform X2
putative receptor protein kinase ZmPK1
G-type lectin S-receptor-like serine/threonine-protein kinase CES101 isoform X1
putative receptor protein kinase ZmPK1
G-type lectin S-receptor-like serine/threonine-protein kinase At1g11330
G-type lectin S-receptor-like serine/threonine-protein kinase At1g11410 isoform X2
uncharacterized protein LOC109013535
receptor-like serine/threonine-protein kinase SD1-8 isoform X1
G-type lectin S-receptor-like serine/threonine-protein kinase At4g03230 isoform X1
G-type lectin S-receptor-like serine/threonine-protein kinase SD2-2
G-type lectin S-receptor-like serine/threonine-protein kinase At1g34300
G-type lectin S-receptor-like serine/threonine-protein kinase At2g19130
G-type lectin S-receptor-like serine/threonine-protein kinase At4g03230 isoform X1
G-type lectin S-receptor-like serine/threonine-protein kinase SD2-5
G-type lectin S-receptor-like serine/threonine-protein kinase At2g19130
G-type lectin S-receptor-like serine/threonine-protein kinase At1g67520
G-type lectin S-receptor-like serine/threonine-protein kinase At1g11410 isoform X1
G-type lectin S-receptor-like serine/threonine-protein kinase CES101 isoform X1
G-type lectin S-receptor-like serine/threonine-protein kinase At4g03230 isoform X3
G-type lectin S-receptor-like serine/threonine-protein kinase At4g27290
G-type lectin S-receptor-like serine/threonine-protein kinase At4g03230
G-type lectin S-receptor-like serine/threonine-protein kinase At1g11330
G-type lectin S-receptor-like serine/threonine-protein kinase At4g03230 isoform X2
G-type lectin S-receptor-like serine/threonine-protein kinase At4g03230
putative receptor protein kinase ZmPK1
G-type lectin S-receptor-like serine/threonine-protein kinase At1g11410 isoform X1
G-type lectin S-receptor-like serine/threonine-protein kinase At1g11330
G-type lectin S-receptor-like serine/threonine-protein kinase At1g11330
G-type lectin S-receptor-like serine/threonine-protein kinase At2g19130
epidermis-specific secreted glycoprotein EP1-like
receptor-like serine/threonine-protein kinase SD1-6 isoform X1

putative receptor protein kinase ZmPK1
G-type lectin S-receptor-like serine/threonine-protein kinase CES101 isoform X3
putative receptor protein kinase ZmPK1
G-type lectin S-receptor-like serine/threonine-protein kinase LECRK3
G-type lectin S-receptor-like serine/threonine-protein kinase LECRK1
putative receptor protein kinase ZmPK1
putative receptor protein kinase ZmPK1
putative receptor protein kinase ZmPK1
G-type lectin S-receptor-like serine/threonine-protein kinase CES101
receptor-like serine/threonine-protein kinase SD1-8
putative receptor protein kinase ZmPK1
G-type lectin S-receptor-like serine/threonine-protein kinase At1g11330 isoform X4
G-type lectin S-receptor-like serine/threonine-protein kinase At4g03230
receptor-like serine/threonine-protein kinase SD1-8
G-type lectin S-receptor-like serine/threonine-protein kinase LECRK3
G-type lectin S-receptor-like serine/threonine-protein kinase At4g27290 isoform X3
receptor-like serine/threonine-protein kinase SD1-6 isoform X2
putative receptor protein kinase ZmPK1
G-type lectin S-receptor-like serine/threonine-protein kinase LECRK3
G-type lectin S-receptor-like serine/threonine-protein kinase At1g11330 isoform X2
G-type lectin S-receptor-like serine/threonine-protein kinase At4g03230
G-type lectin S-receptor-like serine/threonine-protein kinase At1g11410 isoform X1
G-type lectin S-receptor-like serine/threonine-protein kinase CES101 isoform X3
G-type lectin S-receptor-like serine/threonine-protein kinase At1g11330 isoform X2
putative receptor protein kinase ZmPK1
receptor-like serine/threonine-protein kinase SD1-8
G-type lectin S-receptor-like serine/threonine-protein kinase At4g27290
G-type lectin S-receptor-like serine/threonine-protein kinase At1g11410 isoform X1
G-type lectin S-receptor-like serine/threonine-protein kinase B120 isoform X2
putative receptor protein kinase ZmPK1
G-type lectin S-receptor-like serine/threonine-protein kinase SD2-5
G-type lectin S-receptor-like serine/threonine-protein kinase At1g11410
G-type lectin S-receptor-like serine/threonine-protein kinase At4g03230
receptor-like serine/threonine-protein kinase SD1-8
G-type lectin S-receptor-like serine/threonine-protein kinase At1g11330 isoform X1
receptor-like serine/threonine-protein kinase SD1-6
putative receptor protein kinase ZmPK1
putative receptor protein kinase ZmPK1
G-type lectin S-receptor-like serine/threonine-protein kinase LECRK1
G-type lectin S-receptor-like serine/threonine-protein kinase CES101 isoform X2
G-type lectin S-receptor-like serine/threonine-protein kinase LECRK1

G-type lectin S-receptor-like serine/threonine-protein kinase LECRK1
G-type lectin S-receptor-like serine/threonine-protein kinase At4g27290
G-type lectin S-receptor-like serine/threonine-protein kinase B120 isoform X4
G-type lectin S-receptor-like serine/threonine-protein kinase At1g11410 isoform X3
G-type lectin S-receptor-like serine/threonine-protein kinase CES101 isoform X2
G-type lectin S-receptor-like serine/threonine-protein kinase At1g11410 isoform X2
G-type lectin S-receptor-like serine/threonine-protein kinase At1g11410
G-type lectin S-receptor-like serine/threonine-protein kinase LECRK3 isoform X2
G-type lectin S-receptor-like serine/threonine-protein kinase At1g11300
G-type lectin S-receptor-like serine/threonine-protein kinase CES101 isoform X1
putative receptor protein kinase ZmPK1
G-type lectin S-receptor-like serine/threonine-protein kinase At1g11410 isoform X1
G-type lectin S-receptor-like serine/threonine-protein kinase At1g11410 isoform X3
G-type lectin S-receptor-like serine/threonine-protein kinase CES101 isoform X2
G-type lectin S-receptor-like serine/threonine-protein kinase At1g11330
G-type lectin S-receptor-like serine/threonine-protein kinase At4g27290 isoform X2
putative receptor protein kinase ZmPK1
G-type lectin S-receptor-like serine/threonine-protein kinase At1g11330
G-type lectin S-receptor-like serine/threonine-protein kinase At1g11410 isoform X3
Alpha-2-macroglobulin
Uncharacterized protein
Uncharacterized protein
Uncharacterized protein (Fragment)
Uncharacterized protein
Uncharacterized protein
PAN domain protein
Uncharacterized protein (Fragment)
Uncharacterized protein
Hepatocyte growth factor
Uncharacterized protein
Uncharacterized protein
Uncharacterized protein
Uncharacterized protein
Uncharacterized protein
Lipoprotein, putative
Uncharacterized protein
PAN domain protein
Uncharacterized protein
Uncharacterized protein
Uncharacterized protein
Uncharacterized protein

Uncharacterized protein
Alpha-2-macroglobulin
Uncharacterized protein
Alpha-2-macroglobulin
Alpha-2-macroglobulin
Uncharacterized protein
Alpha-2-macroglobulin
Uncharacterized protein
Uncharacterized protein
Uncharacterized protein
Uncharacterized protein
Uncharacterized protein
Plasminogen
Plasminogen
Uncharacterized protein
Uncharacterized protein
Uncharacterized protein
Hepatocyte growth factor
Uncharacterized protein
Plasminogen
Uncharacterized protein
Uncharacterized protein
Uncharacterized protein
Uncharacterized protein
Uncharacterized protein
Uncharacterized protein
Uncharacterized protein
Uncharacterized protein
Uncharacterized protein
Uncharacterized protein
Uncharacterized protein
Uncharacterized protein
Uncharacterized protein
Uncharacterized protein
Uncharacterized protein
Uncharacterized protein
Uncharacterized protein
Uncharacterized protein
Uncharacterized protein
Uncharacterized protein
Uncharacterized protein

Uncharacterized protein
Uncharacterized protein
Uncharacterized protein
Uncharacterized protein
Uncharacterized protein
Uncharacterized protein
Uncharacterized protein
Uncharacterized protein
Uncharacterized protein
Uncharacterized protein
Uncharacterized protein
Uncharacterized protein
Uncharacterized protein
Uncharacterized protein
Uncharacterized protein
Uncharacterized protein
Uncharacterized protein
Uncharacterized protein
Uncharacterized protein
Uncharacterized protein
Uncharacterized protein
Uncharacterized protein
Uncharacterized protein
Uncharacterized protein
Uncharacterized protein
Uncharacterized protein
Uncharacterized protein
Uncharacterized protein
Uncharacterized protein
Uncharacterized protein
Uncharacterized protein
Uncharacterized protein
Uncharacterized protein
Uncharacterized protein
Uncharacterized protein
Uncharacterized protein
Uncharacterized protein
Uncharacterized protein
Uncharacterized protein
Uncharacterized protein
Uncharacterized protein

Uncharacterized protein
Uncharacterized protein
Uncharacterized protein
Uncharacterized protein
Uncharacterized protein
Uncharacterized protein
Uncharacterized protein
Uncharacterized protein
Uncharacterized protein
Uncharacterized protein
Uncharacterized protein
Uncharacterized protein
Uncharacterized protein
Uncharacterized protein
Uncharacterized protein
Uncharacterized protein
Uncharacterized protein
Uncharacterized protein
Uncharacterized protein
Uncharacterized protein
Uncharacterized protein
Uncharacterized protein
Uncharacterized protein
Uncharacterized protein
Uncharacterized protein
Uncharacterized protein
Uncharacterized protein
Uncharacterized protein
Uncharacterized protein
Uncharacterized protein
Uncharacterized protein
Uncharacterized protein
Uncharacterized protein
Uncharacterized protein
Uncharacterized protein
Uncharacterized protein
Uncharacterized protein
Uncharacterized protein
Uncharacterized protein
Uncharacterized protein
Uncharacterized protein

Uncharacterized protein
Uncharacterized protein
Uncharacterized protein
Uncharacterized protein
Uncharacterized protein
Uncharacterized protein
Uncharacterized protein
Uncharacterized protein
Uncharacterized protein
Uncharacterized protein
Uncharacterized protein
Uncharacterized protein
Uncharacterized protein
Uncharacterized protein
Uncharacterized protein
Uncharacterized protein
Uncharacterized protein
Uncharacterized protein
Uncharacterized protein
Uncharacterized protein
Uncharacterized protein
CBEL
CBEL
Plasminogen
Hepatocyte growth factor
Uncharacterized protein (Fragment)
Ras-related protein
Uncharacterized protein (Fragment)
Uncharacterized protein
Uncharacterized protein
Ras-related protein
Uncharacterized protein
Cuticlin-1
Uncharacterized protein
Uncharacterized protein
Uncharacterized protein
Uncharacterized protein
Hepatocyte growth factor
Putative antigen B membrane protein (Fragment)
Putative receptor kinase (Fragment)
Putative receptor kinase Aly8/ARK3 (Fragment)

Putative receptor kinase (Fragment)
Putative receptor kinase (Fragment)
Putative receptor kinase (Fragment)
Putative receptor kinase (Fragment)
Putative receptor kinase (Fragment)
Putative receptor kinase (Fragment)
Putative receptor kinase Aly8/ARK3 (Fragment)
Putative receptor kinase (Fragment)
Putative S-receptor kinase (Fragment)
Putative receptor kinase (Fragment)
Putative receptor kinase (Fragment)
Putative receptor kinase Aly14 (Fragment)
Putative receptor kinase Aly8/ARK3 (Fragment)
S-locus receptor kinase (Fragment)
Putative receptor kinase (Fragment)
Putative receptor kinase Aly8/ARK3 (Fragment)
Putative receptor kinase (Fragment)
Lal2 S-domain receptor kinase
Putative receptor kinase (Fragment)
Putative receptor kinase 3
Putative receptor kinase (Fragment)
Putative receptor kinase Aly8/ARK3 (Fragment)
Putative receptor kinase (Fragment)
Putative receptor kinase (Fragment)
Putative receptor kinase (Fragment)
Putative receptor kinase Aly14 (Fragment)
Putative receptor kinase Aly8/ARK3 (Fragment)
Putative receptor kinase (Fragment)
Putative receptor kinase Aly8/ARK3 (Fragment)
Putative receptor kinase Aly8/ARK3 (Fragment)
Putative receptor kinase (Fragment)
Putative receptor kinase (Fragment)
Putative receptor kinase (Fragment)
Putative receptor kinase (Fragment)
Putative receptor kinase (Fragment)
S-domain receptor kinase (Fragment)
Putative receptor kinase (Fragment)
Putative receptor kinase (Fragment)
Putative receptor kinase (Fragment)
Putative receptor kinase Aly8/ARK3 (Fragment)
Lal2 S-domain receptor kinase

Putative receptor kinase Aly8/ARK3 (Fragment)
Putative receptor kinase (Fragment)
Lal2 S-domain receptor kinase
Putative receptor kinase (Fragment)
Putative receptor kinase (Fragment)
Putative receptor kinase (Fragment)
Putative receptor kinase (Fragment)
Putative receptor kinase Aly8/ARK3 (Fragment)
Putative receptor kinase (Fragment)
Putative receptor kinase (Fragment)
Putative receptor kinase (Fragment)
Putative receptor kinase Aly8/ARK3 (Fragment)
Putative receptor kinase (Fragment)
S-locus receptor kinase (Fragment)
Putative receptor kinase (Fragment)
Putative receptor kinase Aly8/ARK3 (Fragment)
Putative receptor kinase (Fragment)
Lal2 S-domain receptor kinase
Putative receptor kinase (Fragment)
Putative receptor kinase Aly14 (Fragment)
S-domain receptor kinase
Putative S-locus lectin protein kinase-like protein
Putative receptor kinase (Fragment)
Putative receptor kinase Aly8/ARK3 (Fragment)
Uncharacterized protein
Uncharacterized protein
Uncharacterized protein
Uncharacterized protein
Uncharacterized protein
Uncharacterized protein
Uncharacterized protein
Uncharacterized protein
Uncharacterized protein
Uncharacterized protein
Uncharacterized protein
Uncharacterized protein
Uncharacterized protein
Uncharacterized protein
Uncharacterized protein
Uncharacterized protein
Uncharacterized protein

Uncharacterized protein
Uncharacterized protein
Uncharacterized protein
Uncharacterized protein
Uncharacterized protein
Uncharacterized protein
Uncharacterized protein
Uncharacterized protein
Uncharacterized protein
Uncharacterized protein
Uncharacterized protein
Uncharacterized protein
Uncharacterized protein
Uncharacterized protein
Uncharacterized protein
Uncharacterized protein
Uncharacterized protein
Uncharacterized protein
Uncharacterized protein
Uncharacterized protein
Uncharacterized protein
Uncharacterized protein
Uncharacterized protein
Uncharacterized protein
Uncharacterized protein
Uncharacterized protein
Uncharacterized protein
Uncharacterized protein
Uncharacterized protein
Uncharacterized protein
Uncharacterized protein
Uncharacterized protein
Uncharacterized protein
Uncharacterized protein
Uncharacterized protein
Uncharacterized protein
Uncharacterized protein
Uncharacterized protein
Uncharacterized protein
Uncharacterized protein
Uncharacterized protein

Uncharacterized protein
Uncharacterized protein
Uncharacterized protein
Uncharacterized protein
Uncharacterized protein
Uncharacterized protein
Uncharacterized protein
Uncharacterized protein
Uncharacterized protein
Uncharacterized protein
Uncharacterized protein
Uncharacterized protein
Uncharacterized protein
Uncharacterized protein
Uncharacterized protein
Uncharacterized protein
Uncharacterized protein
Uncharacterized protein
Uncharacterized protein
Uncharacterized protein
Uncharacterized protein
Uncharacterized protein
Uncharacterized protein
Uncharacterized protein
Uncharacterized protein
Uncharacterized protein
Uncharacterized protein
Uncharacterized protein
Uncharacterized protein
Uncharacterized protein
Uncharacterized protein
Uncharacterized protein
Uncharacterized protein
Uncharacterized protein
Uncharacterized protein
Uncharacterized protein
Uncharacterized protein
Uncharacterized protein
Uncharacterized protein
Uncharacterized protein
Uncharacterized protein

Uncharacterized protein
Uncharacterized protein
Uncharacterized protein
Beta-fructofuranosidase
Beta-fructofuranosidase
Putative beta-fructofuranosidase
Beta-fructofuranosidase, putative
Putative unspecified product
Beta-fructofuranosidase_-_putative
Uncharacterized protein
Beta-fructofuranosidase, putative
Uncharacterized protein
PAN domain protein
Uncharacterized protein
Uncharacterized protein (Fragment)
Collagen alpha1(I) chainlike [Ficedula albicollis] (Fragment)
Uncharacterized protein
Putative LOC101887535 [Musca domestica]
Putative LOC100881972 [Megachile rotundata] (Fragment)
Uncharacterized protein (Fragment)
Putative LOC100160261 [Acyrthosiphon pisum]
Putative LOC100572421 [Acyrthosiphon pisum] (Fragment)
Uncharacterized protein (Fragment)
Putative LOC100908673 [Metaseiulus occidentalis]
Uncharacterized protein (Fragment)
Uncharacterized protein (Fragment)
Uncharacterized protein
Proclotting enzymelike [Megachile rotundata] (Fragment)
Uncharacterized protein
Uncharacterized protein (Fragment)
Uncharacterized protein (Fragment)
Putative LOC100169340 [Acyrthosiphon pisum]
Uncharacterized protein (Fragment)
Serine protease 79 [Nasonia vitripennis]
Uncharacterized protein (Fragment)
Uncharacterized protein
Uncharacterized protein (Fragment)
Uncharacterized protein (Fragment)
Uncharacterized protein (Fragment)
Uncharacterized protein
Uncharacterized protein (Fragment)

Uncharacterized protein (Fragment)
Uncharacterized protein (Fragment)
Uncharacterized protein (Fragment)
Uncharacterized protein (Fragment)
Hepatocyte growth factor
Uncharacterized protein
Hepatocyte growth factor
Plasminogen
Uncharacterized protein
Uncharacterized protein
coagulation factor XI isoform X3
Hepatocyte growth factor
hepatocyte growth factor-like protein
plasma kallikrein
coagulation factor XI isoform X1
coagulation factor XI isoform X2
Plasminogen
Plasminogen
Hepatocyte growth factor
Plasminogen (Fragment)
Plasminogen (Fragment)
Hepatocyte growth factor (Fragment)
Uncharacterized protein
Predicted protein
Similar to galactose oxidase
PAN domain containing protein 1-like protein (Fragment)
Uncharacterized protein (Fragment)
Uncharacterized protein
Uncharacterized protein
Uncharacterized protein (Fragment)
Uncharacterized protein
Uncharacterized protein (Fragment)
Uncharacterized protein (Fragment)
Uncharacterized protein
Uncharacterized protein (Fragment)
Uncharacterized protein
Uncharacterized protein
Uncharacterized protein
Uncharacterized protein
Hepatocyte growth factor/Macrophage stimulating 1 (Hepatocyte growth factor-like) A
Hepatocyte growth factor/Macrophage stimulating 1 (Hepatocyte growth factor-like) B

Uncharacterized protein
Hepatocyte growth factor
Uncharacterized protein
Hepatocyte growth factor-like protein
Coagulation factor xi-like
uncharacterized protein LOC106180733
angiopoietin-2-like isoform X1
uncharacterized protein LOC106151397
uncharacterized protein LOC106171828
olfactomedin-like protein 2A
uncharacterized protein LOC106154099
ficolin-2
uncharacterized protein LOC106175132
uncharacterized protein LOC106162950
ficolin-2
uncharacterized protein LOC106153009
chymotrypsin A-like
uncharacterized protein LOC106179404
uncharacterized protein LOC106151371
uncharacterized protein LOC106166054
uncharacterized protein LOC106162850
uncharacterized protein LOC106176197
uncharacterized protein LOC106161639
mucin-2-like
uncharacterized protein LOC106179412
signal peptide, CUB and EGF-like domain-containing protein 1
Uncharacterized protein
uncharacterized protein LOC106170457
pulmonary surfactant-associated protein A-like
uncharacterized protein LOC106162620 isoform X1
uncharacterized protein LOC106179407
uncharacterized protein LOC106155386
uncharacterized protein LOC106159394
neurocan core protein
fibrinogen-like protein A
uncharacterized protein LOC106166537
uncharacterized protein LOC106180625
uncharacterized protein LOC106180721
uncharacterized protein LOC106166018
fucolectin
uncharacterized protein LOC106150931

fibropellin-1-like
uncharacterized protein LOC106177541
techylectin-5A-like isoform X2
fucolectin-1
fibrinogen C domain-containing protein 1-A
uncharacterized protein LOC106180489
sushi, von Willebrand factor type A, EGF and pentraxin domain-containing protein 1-like
contactin-associated protein-like 2
fibrinogen-like protein A
uncharacterized protein LOC106172624 isoform X1
uncharacterized protein LOC106150688 isoform X2
fucolectin-like
E3 ubiquitin-protein ligase RNF34-like
integumentary mucin C.1
uncharacterized protein LOC106166081
uncharacterized protein LOC106153018 isoform X2
uncharacterized protein LOC106153018 isoform X1
mucin-13
uncharacterized protein LOC106158949
uncharacterized protein LOC106180613 isoform X1
Metalloendopeptidase
uncharacterized protein LOC106165633
uncharacterized protein LOC106153926
uncharacterized protein LOC106180720
contactin-associated protein-like 2
fucolectin-like
fibropellin-1
uncharacterized protein LOC106177863
uncharacterized protein LOC106179405
uncharacterized protein LOC106180017
uncharacterized protein LOC106175144 isoform X2
angiopoietin-2-like
fibrinogen-like protein A
fucolectin-like
uncharacterized protein LOC106180015
uncharacterized protein LOC106169032
uncharacterized protein LOC106170105
Metalloendopeptidase
uncharacterized protein LOC106177799
uncharacterized protein LOC106162620 isoform X2
uncharacterized protein LOC106176997

uncharacterized protein LOC106178942
signal peptide, CUB and EGF-like domain-containing protein 2
uncharacterized protein LOC106161214
uncharacterized protein LOC106156679
fucolectin-5-like
uncharacterized protein LOC106180490
uncharacterized protein LOC106177245
micronemal protein 4
cartilage intermediate layer protein 1-like
fucolectin-like
uncharacterized protein LOC106164830
uncharacterized protein LOC106177800
uncharacterized protein LOC106173188
uncharacterized protein LOC106171474
homeobox protein 6-like
uncharacterized protein LOC106163073
uncharacterized protein LOC106175114
angiopoietin-4-like
fibropellin-1
uncharacterized protein LOC106163219
trypsin
C-type lectin domain family 3 member A-like
Metalloendopeptidase
uncharacterized protein LOC106170487
uncharacterized protein LOC112041273
contactin-associated protein 1-like
ryncolin-4-like
uncharacterized protein LOC106180613 isoform X2
contactin-associated protein-like 2
SCO-spondin
coiled-coil domain-containing protein 80-like
uncharacterized protein LOC106150688 isoform X1
uncharacterized protein LOC112041434
uncharacterized protein LOC106179398
uncharacterized protein LOC106163484
uncharacterized protein LOC106164610
uncharacterized protein LOC106176998
chymotrypsin-like protease CTRL-1
angiopoietin-1-like
uncharacterized protein LOC106177244
uncharacterized protein LOC106167159

uncharacterized protein LOC106180274
Plasminogen
coagulation factor XI
hepatocyte growth factor-like protein
Hepatocyte growth factor
Hepatocyte growth factor
plasminogen-like isoform X6
Plasminogen
Plasminogen
Plasminogen
Plasminogen
Plasminogen
Uncharacterized protein
Uncharacterized protein (Fragment)
Uncharacterized protein (Fragment)
Uncharacterized protein (Fragment)
Uncharacterized protein
Uncharacterized protein
Uncharacterized protein
Uncharacterized protein
Uncharacterized protein
Uncharacterized protein
Uncharacterized protein (Fragment)
Uncharacterized protein
Uncharacterized protein
Uncharacterized protein
Uncharacterized protein
Uncharacterized protein
Uncharacterized protein
Uncharacterized protein
PAN domain protein
Uncharacterized protein
Uncharacterized protein
Uncharacterized protein
PAN domain-containing protein
PAN domain-containing protein
PAN domain-containing protein
Fucolectin-related protein
C-type lectin
C-type lectin
Uncharacterized protein

Uncharacterized protein
PAN domain-containing protein
Uncharacterized protein
Uncharacterized protein
Uncharacterized protein
Uncharacterized protein
Uncharacterized protein
Uncharacterized protein
Uncharacterized protein
Uncharacterized protein
PAN domain-containing protein
Uncharacterized protein
PAN domain-containing protein
Uncharacterized protein
Uncharacterized protein
Uncharacterized protein
Uncharacterized protein
Uncharacterized protein
Uncharacterized protein
Uncharacterized protein
PAN domain-containing protein
Uncharacterized protein
Uncharacterized protein
Uncharacterized protein
Uncharacterized protein
PAN domain-containing protein
Uncharacterized protein
Uncharacterized protein
PAN domain-containing protein
Uncharacterized protein
Uncharacterized protein
Uncharacterized protein
Uncharacterized protein
Alpha-2-macroglobulin family protein
Uncharacterized protein
Hepatocyte growth factor-like protein
Plasminogen
Uncharacterized protein
Goosefish kalliklectin
Uncharacterized protein
Uncharacterized protein

Uncharacterized protein
Uncharacterized protein
Uncharacterized protein (Fragment)
Uncharacterized protein
Uncharacterized protein
Uncharacterized protein
Uncharacterized protein
Uncharacterized protein
Uncharacterized protein
Uncharacterized protein
Uncharacterized protein
Uncharacterized protein
Uncharacterized protein
Uncharacterized protein
Uncharacterized protein
Uncharacterized protein
Uncharacterized protein (Fragment)
Uncharacterized protein
Uncharacterized protein
Uncharacterized protein
Uncharacterized protein
Uncharacterized protein
Uncharacterized protein
Uncharacterized protein
Uncharacterized protein
Uncharacterized protein
Uncharacterized protein
Uncharacterized protein
Uncharacterized protein
Uncharacterized protein
Uncharacterized protein
Uncharacterized protein
Uncharacterized protein
Uncharacterized protein
Uncharacterized protein
Uncharacterized protein
Uncharacterized protein
Uncharacterized protein (Fragment)
Uncharacterized protein
Uncharacterized protein
Uncharacterized protein

Uncharacterized protein
Uncharacterized protein
Uncharacterized protein
Uncharacterized protein
Uncharacterized protein
Uncharacterized protein
Uncharacterized protein
Uncharacterized protein
Uncharacterized protein
Uncharacterized protein
Uncharacterized protein
Uncharacterized protein
Uncharacterized protein
Uncharacterized protein
Uncharacterized protein
Uncharacterized protein
Uncharacterized protein
Uncharacterized protein
Uncharacterized protein
Uncharacterized protein
Uncharacterized protein
Uncharacterized protein
Uncharacterized protein
Uncharacterized protein
Uncharacterized protein
Uncharacterized protein
Uncharacterized protein
Uncharacterized protein
Uncharacterized protein
Uncharacterized protein
Uncharacterized protein (Fragment)
Uncharacterized protein
Uncharacterized protein
Uncharacterized protein
Uncharacterized protein
Uncharacterized protein
Uncharacterized protein
Uncharacterized protein
Uncharacterized protein
Uncharacterized protein
Uncharacterized protein

Uncharacterized protein
Uncharacterized protein
Uncharacterized protein
Uncharacterized protein (Fragment)
Uncharacterized protein
Uncharacterized protein
Uncharacterized protein
Uncharacterized protein
Uncharacterized protein
Uncharacterized protein
Uncharacterized protein
Uncharacterized protein
Uncharacterized protein
Uncharacterized protein
Uncharacterized protein
Uncharacterized protein
Uncharacterized protein
Uncharacterized protein
Uncharacterized protein
Uncharacterized protein
Uncharacterized protein
Uncharacterized protein
Uncharacterized protein
Uncharacterized protein
Uncharacterized protein
Uncharacterized protein
Uncharacterized protein
Uncharacterized protein
Uncharacterized protein
Uncharacterized protein
Uncharacterized protein
Uncharacterized protein
Uncharacterized protein
Uncharacterized protein
Uncharacterized protein
Uncharacterized protein
Uncharacterized protein
Uncharacterized protein
Uncharacterized protein
Uncharacterized protein (Fragment)
Uncharacterized protein

Uncharacterized protein
S-receptor kinase-like protein 3
CM0216.570.nc protein
S-receptor kinase-like protein 1
Uncharacterized protein
CM0216.590.nc protein (Fragment)
CM0216.580.nc protein
Uncharacterized protein
Uncharacterized protein
Uncharacterized protein (Fragment)
Kallikrein B1
Macrophage stimulating 1
Plasminogen
Uncharacterized protein
Uncharacterized protein
Uncharacterized protein
Uncharacterized protein
Uncharacterized protein
Uncharacterized protein
Uncharacterized protein
Uncharacterized protein
Uncharacterized protein (Fragment)
Uncharacterized protein
Uncharacterized protein
S-locus-related I (Fragment)
Uncharacterized protein
Uncharacterized protein
Uncharacterized protein
Uncharacterized protein
Uncharacterized protein
Uncharacterized protein
Uncharacterized protein
Uncharacterized protein
Uncharacterized protein
Uncharacterized protein
Uncharacterized protein
Uncharacterized protein
Uncharacterized protein
Uncharacterized protein
Uncharacterized protein
PAN domain protein

Uncharacterized protein
Uncharacterized protein
Uncharacterized protein
Uncharacterized protein
Uncharacterized protein
Neuropeptide receptor (GPRnpr2)
Uncharacterized protein
Uncharacterized protein (Fragment)
Uncharacterized protein (Fragment)
Uncharacterized protein (Fragment)
Thrombospondin type-1 domain-containing protein 7B (Fragment)
Uncharacterized protein (Fragment)
Cleavage and polyadenylation factor complex subunit C74.02c (Fragment)
Cuticlin-1
Pulmonary surfactant-associated protein A
Neurotrypsin
Uncharacterized protein
Low-density lipoprotein receptor-related protein 11
Plasminogen-related protein A
DNA-directed RNA polymerase subunit alpha (Fragment)
Uncharacterized protein
Uncharacterized protein
Uncharacterized protein (Fragment)
Uncharacterized protein (Fragment)
Translation initiation factor IF-2 (Fragment)
Type I phosphodiesterase / nucleotide pyrophosphatase family protein
Uncharacterized protein
Uncharacterized protein
Uncharacterized protein
Uncharacterized protein (Fragment)
Uncharacterized protein
Plasminogen
Uncharacterized protein
Hepatocyte growth factor
Uncharacterized protein
Uncharacterized protein
Plasminogen
Uncharacterized protein
Uncharacterized protein
Hepatocyte growth factor
Uncharacterized protein

Plasminogen
Uncharacterized protein
Uncharacterized protein
Uncharacterized protein
Uncharacterized protein (Fragment)
Plasminogen
Uncharacterized protein
Plasma kallikrein preproprotein (Fragment)
Uncharacterized protein
Uncharacterized protein
Uncharacterized protein
Uncharacterized protein
Uncharacterized protein
Hepatocyte growth factor
Plasminogen
Plasminogen
Uncharacterized protein
Uncharacterized protein
Uncharacterized protein
Uncharacterized protein
Uncharacterized protein
Uncharacterized protein
Uncharacterized protein
Plasminogen
Bulb-type lectin domain
Protein kinase domain
Protein kinase domain
Bulb-type lectin domain
Protein kinase domain
Protein kinase domain
Protein kinase domain
Protein kinase domain
Protein kinase domain
Bulb-type lectin domain
Protein kinase domain
Apple-like
Bulb-type lectin domain
Protein kinase domain
Protein kinase domain
Protein kinase domain
Protein kinase domain

Apple-like
Protein kinase domain
Protein kinase domain
Bulb-type lectin domain
Protein kinase domain
Protein kinase domain
S-locus glycoprotein
Protein kinase domain
S-locus glycoprotein
Uncharacterized protein
Uncharacterized protein
Uncharacterized protein
Uncharacterized protein
Apple
Uncharacterized protein
Plasminogen
Uncharacterized protein
Uncharacterized protein
Uncharacterized protein
Uncharacterized protein
Uncharacterized protein
Uncharacterized protein
Uncharacterized protein
Uncharacterized protein
Uncharacterized protein
Uncharacterized protein
Uncharacterized protein
Uncharacterized protein
Uncharacterized protein
Uncharacterized protein
Uncharacterized protein
Uncharacterized protein
Uncharacterized protein
Uncharacterized protein (Fragment)
Uncharacterized protein
Uncharacterized protein
Uncharacterized protein
Uncharacterized protein
Uncharacterized protein
Uncharacterized protein
Uncharacterized protein

Uncharacterized protein
Uncharacterized protein
Uncharacterized protein
Uncharacterized protein
Uncharacterized protein
Uncharacterized protein
Uncharacterized protein
Uncharacterized protein
Uncharacterized protein
Uncharacterized protein
Uncharacterized protein
Uncharacterized protein (Fragment)
Uncharacterized protein
Uncharacterized protein
Uncharacterized protein
Uncharacterized protein
Uncharacterized protein
Uncharacterized protein (Fragment)
Uncharacterized protein
Uncharacterized protein
Uncharacterized protein
Uncharacterized protein
Uncharacterized protein
Uncharacterized protein
Uncharacterized protein
Uncharacterized protein
Uncharacterized protein
Uncharacterized protein
Uncharacterized protein (Fragment)
Uncharacterized protein (Fragment)
Uncharacterized protein
Uncharacterized protein
Uncharacterized protein
Uncharacterized protein
Uncharacterized protein (Fragment)
Uncharacterized protein (Fragment)
Uncharacterized protein
Uncharacterized protein
Uncharacterized protein
Uncharacterized protein
Uncharacterized protein

Uncharacterized protein
Uncharacterized protein
Uncharacterized protein
Uncharacterized protein
Uncharacterized protein
Uncharacterized protein
Uncharacterized protein
Uncharacterized protein
Uncharacterized protein
Uncharacterized protein
Uncharacterized protein
Uncharacterized protein
Uncharacterized protein
Uncharacterized protein
Uncharacterized protein
Uncharacterized protein
Uncharacterized protein
Uncharacterized protein
Uncharacterized protein
Uncharacterized protein
Uncharacterized protein
Uncharacterized protein
Uncharacterized protein
Uncharacterized protein (Fragment)
Uncharacterized protein
Uncharacterized protein
Uncharacterized protein
Uncharacterized protein (Fragment)
Uncharacterized protein
Uncharacterized protein
Uncharacterized protein
Uncharacterized protein
Uncharacterized protein
Uncharacterized protein
Uncharacterized protein
Uncharacterized protein
Uncharacterized protein
Uncharacterized protein (Fragment)
Uncharacterized protein
Uncharacterized protein
Uncharacterized protein

Uncharacterized protein
Uncharacterized protein
Uncharacterized protein
Uncharacterized protein (Fragment)
Uncharacterized protein
Uncharacterized protein
Uncharacterized protein
Uncharacterized protein
Uncharacterized protein (Fragment)
Uncharacterized protein
Uncharacterized protein
Uncharacterized protein
Uncharacterized protein
Uncharacterized protein
Uncharacterized protein
Uncharacterized protein
Uncharacterized protein
Uncharacterized protein
Uncharacterized protein (Fragment)
Uncharacterized protein (Fragment)
Uncharacterized protein
Uncharacterized protein
Uncharacterized protein (Fragment)
Uncharacterized protein
Uncharacterized protein
Uncharacterized protein
Uncharacterized protein
Uncharacterized protein
Uncharacterized protein
Uncharacterized protein (Fragment)
Uncharacterized protein
Uncharacterized protein
Uncharacterized protein
Uncharacterized protein
Uncharacterized protein
Uncharacterized protein
Glycoprotein-N-acetylgalactosamine 3-beta-galactosyltransferase 1-B
Plasma kallikrein
Uncharacterized protein
Uncharacterized protein
Uncharacterized protein

Agglutinin isolectin 1
Agglutinin isolectin 1
Uncharacterized protein
Galactose oxidase
Agglutinin isolectin 1 (Fragment)
Uncharacterized protein
Galactose oxidase
Uncharacterized protein
Uncharacterized protein
Uncharacterized protein
Uncharacterized protein
PAN domain protein
PAN domain-containing protein
Uncharacterized protein
Uncharacterized protein
Hepatocyte growth factor (Fragment)
Plasma kallikrein (Fragment)
Uncharacterized protein
Uncharacterized protein
Uncharacterized protein
Hepatocyte growth factor
Uncharacterized protein
Plasminogen
Uncharacterized protein
Uncharacterized protein
Uncharacterized protein
Hepatocyte growth factor
Uncharacterized protein
Uncharacterized protein
Uncharacterized protein
Uncharacterized protein
Uncharacterized protein
Uncharacterized protein
Uncharacterized protein
Uncharacterized protein
Uncharacterized protein
Uncharacterized protein
Uncharacterized protein
Uncharacterized protein
Uncharacterized protein
Uncharacterized protein

Uncharacterized protein
Uncharacterized protein
Uncharacterized protein
Uncharacterized protein
Uncharacterized protein
Uncharacterized protein
Uncharacterized protein
Uncharacterized protein
Uncharacterized protein (Fragment)
Uncharacterized protein
Uncharacterized protein
Uncharacterized protein
Uncharacterized protein
Uncharacterized protein (Fragment)
Uncharacterized protein
Uncharacterized protein
Uncharacterized protein
Uncharacterized protein
Uncharacterized protein
Uncharacterized protein
Uncharacterized protein
Uncharacterized protein
Uncharacterized protein
Uncharacterized protein
Uncharacterized protein
Uncharacterized protein (Fragment)
Uncharacterized protein
Uncharacterized protein
Uncharacterized protein
Uncharacterized protein
Uncharacterized protein
Uncharacterized protein
Uncharacterized protein
Uncharacterized protein
Uncharacterized protein
Uncharacterized protein
Uncharacterized protein
Uncharacterized protein
Uncharacterized protein
Uncharacterized protein
Uncharacterized protein

Uncharacterized protein
Uncharacterized protein
Uncharacterized protein
Uncharacterized protein
Uncharacterized protein
Uncharacterized protein
Uncharacterized protein
Uncharacterized protein
Uncharacterized protein
Uncharacterized protein
Uncharacterized protein
Uncharacterized protein
Uncharacterized protein
Uncharacterized protein
Uncharacterized protein
Uncharacterized protein
Uncharacterized protein
Uncharacterized protein
Uncharacterized protein
Uncharacterized protein
Uncharacterized protein
Uncharacterized protein
Uncharacterized protein
Uncharacterized protein
Uncharacterized protein
Uncharacterized protein
Uncharacterized protein
Uncharacterized protein
Uncharacterized protein
Uncharacterized protein
Uncharacterized protein
Uncharacterized protein (Fragment)
Uncharacterized protein
Uncharacterized protein
Uncharacterized protein
Uncharacterized protein
Uncharacterized protein
Uncharacterized protein
Uncharacterized protein (Fragment)
Uncharacterized protein
Uncharacterized protein

Uncharacterized protein
Uncharacterized protein
Uncharacterized protein
Uncharacterized protein
Uncharacterized protein
Uncharacterized protein
Uncharacterized protein
Uncharacterized protein
Uncharacterized protein
Uncharacterized protein
Uncharacterized protein
Uncharacterized protein (Fragment)
Uncharacterized protein
Uncharacterized protein
Uncharacterized protein
Uncharacterized protein
Uncharacterized protein
Uncharacterized protein
Uncharacterized protein
Uncharacterized protein
Uncharacterized protein
Uncharacterized protein
Uncharacterized protein (Fragment)
Uncharacterized protein (Fragment)
Uncharacterized protein
Uncharacterized protein
Uncharacterized protein
Uncharacterized protein
Uncharacterized protein
Uncharacterized protein
Uncharacterized protein
Uncharacterized protein
Uncharacterized protein
Uncharacterized protein
PAN domain-containing protein
Uncharacterized protein
Uncharacterized protein
Uncharacterized protein
Uncharacterized protein
Alpha-2-macroglobulin
Alpha-2-macroglobulin

Uncharacterized protein
Uncharacterized protein
Uncharacterized protein
Uncharacterized protein
Large extracellular alpha-helical protein
Uncharacterized protein
Uncharacterized protein
PAN domain-containing protein
PAN domain-containing protein
Chitin recognition protein
Uncharacterized protein
Galactose oxidase
Uncharacterized protein
Kelch domain-containing protein
PAN domain-containing protein
Plasminogen
Hepatocyte growth factor
Plasminogen
Uncharacterized protein
Uncharacterized protein
Uncharacterized protein
Plasminogen
Plasminogen
Uncharacterized protein
Hepatocyte growth factor
Plasminogen
Plasminogen
Hepatocyte growth factor
Uncharacterized protein
Uncharacterized protein
Uncharacterized protein
Uncharacterized protein
S-locus lectin kinase family protein
G-type lectin S-receptor-like Serine/Threonine-kinase
G-type lectin S-receptor-like Serine/Threonine-kinase
G-type lectin S-receptor-like Serine/Threonine-kinase
G-type lectin S-receptor-like Serine/Threonine-kinase
G-type lectin S-receptor-like Serine/Threonine-kinase
G-type lectin S-receptor-like Serine/Threonine-kinase
G-type lectin S-receptor-like Serine/Threonine-kinase
S-locus lectin kinase family protein

S-locus lectin kinase family protein
G-type lectin S-receptor-like Serine/Threonine-kinase
G-type lectin S-receptor-like Serine/Threonine-kinase
S-locus lectin kinase family protein
G-type lectin S-receptor-like Serine/Threonine-kinase
G-type lectin S-receptor-like Serine/Threonine-kinase
S-locus lectin kinase family protein
G-type lectin S-receptor-like Serine/Threonine-kinase
G-type lectin S-receptor-like Serine/Threonine-kinase
S-locus lectin kinase family protein
S-locus lectin kinase family protein
G-type lectin S-receptor-like Serine/Threonine-kinase plant
G-type lectin S-receptor-like Serine/Threonine-kinase
G-type lectin S-receptor-like Serine/Threonine-kinase
S-locus lectin kinase family protein
G-type lectin S-receptor-like Serine/Threonine-kinase
G-type lectin S-receptor-like Serine/Threonine-kinase
G-type lectin S-receptor-like Serine/Threonine-kinase
D-mannose-binding lectin protein, putative
Serine/Threonine kinase family protein
Serine/Threonine kinase, plant-type protein
S-locus lectin kinase family protein
S-locus lectin kinase family protein
G-type lectin S-receptor-like Serine/Threonine-kinase
S-locus lectin kinase family protein
S-locus lectin kinase family protein
S-locus lectin kinase family protein
S-locus lectin kinase family protein, putative
S-locus lectin kinase family protein
S-locus lectin kinase family protein
Putative non-specific serine/threonine protein kinase
Serine/Threonine kinase family protein
S-locus lectin kinase family protein
S-locus lectin kinase family protein
S-locus lectin kinase family protein
S-locus lectin kinase family protein
G-type lectin S-receptor-like Serine/Threonine-kinase
Serine/Threonine kinase, plant-type protein
S-locus lectin kinase family protein
S-locus lectin kinase family protein
Protein kinase; S-locus glycoprotein; Curculin-like (Mannose-binding) lectin; Apple-like

Cysteine-rich RLK (Receptor-like kinase) protein
Serine/Threonine kinase, plant-type protein
Serine/Threonine kinase family protein
Serine/Threonine kinase family protein
S-locus lectin kinase family protein
Cysteine-rich RLK (Receptor-like kinase) protein
G-type lectin S-receptor-like Serine/Threonine-kinase
S-locus lectin kinase family protein
G-type lectin S-receptor-like Serine/Threonine-kinase
G-type lectin S-receptor-like Serine/Threonine-kinase
S-locus lectin kinase family protein
S-locus lectin kinase family protein
Cysteine-rich RLK (Receptor-like kinase) protein
S-locus lectin kinase family protein
G-type lectin S-receptor-like Serine/Threonine-kinase
S-locus lectin kinase family protein
Serine/Threonine kinase, plant-type protein
Serine/Threonine kinase, plant-type protein
Serine/Threonine kinase family protein
S-locus lectin kinase family protein
S-locus lectin kinase family protein
S-locus lectin kinase family protein
Serine/Threonine-kinase
Epidermis-specific secreted EP1-like glycoprotein
G-type lectin S-receptor-like Serine/Threonine-kinase
Serine/Threonine kinase family protein
Serine/Threonine kinase, plant-type protein
Serine/Threonine kinase, plant-type protein
Serine/Threonine-kinase
Cysteine-rich RLK (Receptor-like kinase) protein
G-type lectin S-receptor-like Serine/Threonine-kinase
S-locus lectin kinase family protein
G-type lectin S-receptor-like Serine/Threonine-kinase
Methyltransferase PMT14-like protein, putative
S-locus lectin kinase family protein
G-type lectin S-receptor-like Serine/Threonine-kinase
S-locus lectin kinase family protein
S-locus lectin kinase family protein
Serine/Threonine kinase domain protein
S-locus lectin kinase family protein
Cysteine-rich RLK (Receptor-like kinase) protein

G-type lectin S-receptor-like Serine/Threonine-kinase
S-locus lectin kinase family protein
S-locus lectin kinase family protein
G-type lectin S-receptor-like Serine/Threonine-kinase
G-type lectin S-receptor-like Serine/Threonine-kinase
S-locus lectin kinase family protein
S-locus lectin kinase family protein
G-type lectin S-receptor-like Serine/Threonine-kinase
S-locus lectin kinase family protein
S-locus lectin kinase family protein
S-locus lectin kinase family protein
S-locus lectin kinase family protein
Serine/Threonine kinase family protein
S-locus lectin kinase family protein
S-locus lectin kinase family protein
Serine/Threonine kinase family protein
Cysteine-rich RLK (Receptor-like kinase) protein
G-type lectin S-receptor-like Serine/Threonine-kinase
G-type lectin S-receptor-like Serine/Threonine-kinase
G-type lectin S-receptor-like Serine/Threonine-kinase
S-locus lectin kinase family protein
S-locus lectin kinase family protein
Serine/Threonine kinase, plant-type protein
S-locus lectin kinase family protein
G-type lectin S-receptor-like Serine/Threonine-kinase
G-type lectin S-receptor-like Serine/Threonine-kinase
S-locus lectin kinase family protein
S-locus lectin kinase family protein
S-locus lectin kinase family protein
G-type lectin S-receptor-like Serine/Threonine-kinase
S-locus lectin kinase family protein
Malectin/receptor-like kinase family protein
S-locus lectin kinase family protein
S-locus lectin kinase family protein
S-locus lectin kinase family protein
S-locus lectin kinase family protein
D-mannose-binding lectin protein
Serine/Threonine-kinase
S-locus lectin kinase family protein
Malectin/receptor-like kinase family protein
S-locus lectin kinase family protein

Serine/Threonine kinase family protein
G-type lectin S-receptor-like Serine/Threonine-kinase
G-type lectin S-receptor-like Serine/Threonine-kinase
G-type lectin S-receptor-like Serine/Threonine-kinase
G-type lectin S-receptor-like Serine/Threonine-kinase
Lectin kinase family protein
Uncharacterized protein
Putative non-specific protein-tyrosine kinase RLK-Pelle-DLSV family
Putative non-specific protein-tyrosine kinase RLK-Pelle-DLSV family
Uncharacterized protein
Uncharacterized protein
Putative S-locus glycoprotein
Uncharacterized protein
Putative non-specific serine/threonine protein kinase
Putative non-specific serine/threonine protein kinase
Uncharacterized protein
Uncharacterized protein
Uncharacterized protein
Uncharacterized protein
Putative non-specific serine/threonine protein kinase
Uncharacterized protein
Uncharacterized protein
Uncharacterized protein
Putative non-specific serine/threonine protein kinase
Putative S-locus glycoprotein
Putative non-specific serine/threonine protein kinase
Uncharacterized protein
Putative non-specific serine/threonine protein kinase
Uncharacterized protein
Uncharacterized protein
Putative non-specific serine/threonine protein kinase
Putative non-specific serine/threonine protein kinase
Uncharacterized protein
Uncharacterized protein
Uncharacterized protein
Putative non-specific serine/threonine protein kinase
Uncharacterized protein
Putative non-specific serine/threonine protein kinase
Uncharacterized protein
Uncharacterized protein
Uncharacterized protein

Uncharacterized protein
Putative non-specific protein-tyrosine kinase RLK-Pelle-DLSV family
Uncharacterized protein
Uncharacterized protein
Putative non-specific serine/threonine protein kinase
Uncharacterized protein
Uncharacterized protein
Uncharacterized protein
Uncharacterized protein
Uncharacterized protein
Uncharacterized protein
Uncharacterized protein
Putative non-specific serine/threonine protein kinase
Uncharacterized protein
Uncharacterized protein
Uncharacterized protein
Putative non-specific serine/threonine protein kinase
Uncharacterized protein
Uncharacterized protein
Uncharacterized protein
Uncharacterized protein
Putative non-specific serine/threonine protein kinase
Uncharacterized protein
Uncharacterized protein
Uncharacterized protein
Uncharacterized protein
Putative non-specific serine/threonine protein kinase
G-type lectin S-receptor-like Serine/Threonine-kinase plant
Apple; Protein kinase; EGF-like, subtype 2; Curculin-like (Mannose-binding) lectin
G-type lectin S-receptor-like Serine/Threonine-kinase, putative
S-locus lectin kinase family protein
G-type lectin S-receptor-like Serine/Threonine-kinase
Serine/Threonine kinase, plant-type protein
S-locus lectin kinase family protein
G-type lectin S-receptor-like Serine/Threonine-kinase
Serine/Threonine kinase family protein
S-locus lectin kinase family protein
S-locus lectin kinase family protein
Serine/Threonine kinase, plant-type protein
Cysteine-rich RLK (Receptor-like kinase) protein
S-locus lectin kinase family protein

S-locus lectin kinase family protein
Serine/Threonine-kinase
Serine/Threonine kinase family protein
S-locus lectin kinase family protein
PAN-like domain protein
G-type lectin S-receptor-like Serine/Threonine-kinase, putative
Cysteine-rich RLK (Receptor-like kinase) protein
S-locus lectin kinase family protein
Serine/Threonine kinase, plant-type protein
Cysteine-rich RLK (Receptor-like kinase) protein
S-locus lectin kinase family protein
Cysteine-rich RLK (Receptor-like kinase) protein
Serine/Threonine kinase family protein
S-locus lectin kinase family protein
G-type lectin S-receptor-like Serine/Threonine-kinase
PAN-like domain protein
Serine/Threonine kinase, plant-type protein
S-locus lectin kinase family protein
S-locus lectin kinase family protein
G-type lectin S-receptor-like Serine/Threonine-kinase
S-locus lectin kinase family protein
G-type lectin S-receptor-like Serine/Threonine-kinase
S-locus lectin kinase family protein
G-type lectin S-receptor-like Serine/Threonine-kinase
G-type lectin S-receptor-like Serine/Threonine-kinase
Serine/Threonine kinase family protein
G-type lectin S-receptor-like Serine/Threonine-kinase
S-locus lectin kinase family protein
G-type lectin S-receptor-like Serine/Threonine-kinase SD2-5
Receptor-like kinase
Uncharacterized protein
Uncharacterized protein
Uncharacterized protein
Uncharacterized protein
Uncharacterized protein
Uncharacterized protein
Uncharacterized protein
Uncharacterized protein
Uncharacterized protein
Uncharacterized protein
Uncharacterized protein

Uncharacterized protein
Putative non-specific serine/threonine protein kinase
Uncharacterized protein
Uncharacterized protein
Uncharacterized protein
Uncharacterized protein
Uncharacterized protein
Uncharacterized protein
Uncharacterized protein
Uncharacterized protein
Uncharacterized protein
Uncharacterized protein
Uncharacterized protein mg271
Putative pan domain-containing protein
Putative pan domain-containing protein
Uncharacterized protein
Putative pan domain-containing protein
Putative pan domain-containing protein
Putative PAN domain-containing protein
Putative PAN domain-containing protein
Uncharacterized protein
Putative pan domain-containing protein
Uncharacterized protein
Putative pan domain-containing protein
Putative PAN domain-containing protein
Putative pan domain-containing protein
PAN domain protein
Cuticlin-1
Galectin-3-binding protein A
Hepatocyte growth factor
Hepatocyte growth factor
Uncharacterized protein
Coagulation factor XI
Plasminogen
Uncharacterized protein
Uncharacterized protein
Uncharacterized protein
Uncharacterized protein
Uncharacterized protein
Uncharacterized protein
Uncharacterized protein

Cuticlin-1
Uncharacterized protein (Fragment)
Low-density lipoprotein receptor-related protein 11
Uncharacterized protein
Uncharacterized protein
Uncharacterized protein
Uncharacterized protein
Uncharacterized protein
Uncharacterized protein
Uncharacterized protein
Uncharacterized protein
Uncharacterized protein
Uncharacterized protein
Uncharacterized protein
Uncharacterized protein
Uncharacterized protein
Uncharacterized protein
Uncharacterized protein
Uncharacterized protein
Uncharacterized protein
Uncharacterized protein
Uncharacterized protein
Plasminogen (Fragment)
Hepatocyte growth factor (Fragment)
Plasma kallikrein (Fragment)
Plasminogen (Fragment)
Hepatocyte growth factor (Fragment)
Plasma kallikrein (Fragment)
Uncharacterized protein
plasma kallikrein isoform X1
Hepatocyte growth factor
hepatocyte growth factor-like protein isoform X1
plasma kallikrein isoform X2
hepatocyte growth factor-like protein isoform X2
coagulation factor XI isoform X4
Hepatocyte growth factor
Plasminogen
coagulation factor XI isoform X2
coagulation factor XI isoform X1
coagulation factor XI isoform X3
Uncharacterized protein

PAN domain-containing protein
Alpha-2-macroglobulin domain-containing protein
Uncharacterized protein
Alpha-2-macroglobulin
Uncharacterized protein
Uncharacterized protein
Large extracellular alpha-helical protein
PAN domain-containing protein
Uncharacterized protein
Alpha-2-macroglobulin domain protein
Uncharacterized protein
Uncharacterized protein
Uncharacterized protein
Uncharacterized protein
Uncharacterized protein
Uncharacterized protein
PAN domain protein
Alpha-2-macroglobulin N-terminal region family protein
Mll9164 protein
Mll9167 protein
Mlr8222 protein
Mlr1663 protein
Mll9162 protein
Uncharacterized protein
Uncharacterized protein
Uncharacterized protein
Uncharacterized protein
Uncharacterized protein
Uncharacterized protein
Uncharacterized protein
Uncharacterized protein
Peptidase C14 caspase catalytic subunit p20
Alpha-2-macroglobulin domain protein
Uncharacterized protein
Uncharacterized protein
Uncharacterized protein
Uncharacterized protein
Uncharacterized protein
Uncharacterized protein
Uncharacterized protein
Uncharacterized protein

Uncharacterized protein
Uncharacterized protein
Uncharacterized protein
Uncharacterized protein
Uncharacterized protein
Uncharacterized protein
Uncharacterized protein
Uncharacterized protein (Fragment)
Orf32 (Fragment)
Orf29
Orf30
Orf28
Uncharacterized protein
Uncharacterized protein
Membrane protein
Uncharacterized protein
Uncharacterized protein
Uncharacterized protein
Membrane protein
Uncharacterized protein
Membrane protein
Membrane protein
Membrane protein
Uncharacterized protein
Membrane protein
Uncharacterized protein
Uncharacterized protein
Uncharacterized protein
Membrane protein
Uncharacterized protein
Membrane protein
Uncharacterized protein
Membrane protein
Uncharacterized protein
Membrane protein
Uncharacterized protein
Membrane protein
Membrane protein
Membrane protein
Uncharacterized protein
Membrane protein

Uncharacterized protein
Uncharacterized protein
Uncharacterized protein
Membrane protein
Uncharacterized protein
Uncharacterized protein
Membrane protein
Membrane protein
Uncharacterized protein
Uncharacterized protein (Fragment)
Uncharacterized protein
Uncharacterized protein
Uncharacterized protein
Uncharacterized protein
Uncharacterized protein
Uncharacterized protein
Uncharacterized protein
Uncharacterized protein
Uncharacterized protein
Uncharacterized protein
Uncharacterized protein
Uncharacterized protein (Fragment)
Uncharacterized protein (Fragment)
Uncharacterized protein
Uncharacterized protein
Uncharacterized protein
Uncharacterized protein
Uncharacterized protein
Uncharacterized protein
Uncharacterized protein
Uncharacterized protein
Uncharacterized protein
Uncharacterized protein
Uncharacterized protein
Uncharacterized protein
Uncharacterized protein
Uncharacterized protein
Uncharacterized protein
Uncharacterized protein
Uncharacterized protein
Uncharacterized protein

Uncharacterized protein
Uncharacterized protein
Uncharacterized protein (Fragment)
Uncharacterized protein
Uncharacterized protein
Uncharacterized protein (Fragment)
Uncharacterized protein (Fragment)
Uncharacterized protein
Uncharacterized protein
Uncharacterized protein (Fragment)
Uncharacterized protein
Uncharacterized protein
Uncharacterized protein (Fragment)
Uncharacterized protein
Uncharacterized protein
Uncharacterized protein
Uncharacterized protein
Uncharacterized protein
Uncharacterized protein (Fragment)
Uncharacterized protein (Fragment)
Uncharacterized protein
Uncharacterized protein
Uncharacterized protein
Uncharacterized protein
Uncharacterized protein (Fragment)
Uncharacterized protein
Uncharacterized protein
Uncharacterized protein
Uncharacterized protein
Uncharacterized protein (Fragment)
Uncharacterized protein
Uncharacterized protein
Uncharacterized protein
Uncharacterized protein
Uncharacterized protein
Uncharacterized protein (Fragment)
Uncharacterized protein
Uncharacterized protein (Fragment)
Uncharacterized protein (Fragment)
Uncharacterized protein
Uncharacterized protein

Uncharacterized protein (Fragment)
Uncharacterized protein
Uncharacterized protein
Uncharacterized protein (Fragment)
Uncharacterized protein (Fragment)
Uncharacterized protein
Uncharacterized protein
Uncharacterized protein
Uncharacterized protein
Uncharacterized protein
Uncharacterized protein (Fragment)
Uncharacterized protein
Uncharacterized protein
Uncharacterized protein
Uncharacterized protein
Uncharacterized protein
Uncharacterized protein (Fragment)
Uncharacterized protein
Uncharacterized protein
Uncharacterized protein
Uncharacterized protein (Fragment)
Uncharacterized protein
Uncharacterized protein (Fragment)
Uncharacterized protein
Uncharacterized protein (Fragment)
Uncharacterized protein (Fragment)
Uncharacterized protein (Fragment)
Uncharacterized protein
Uncharacterized protein
Uncharacterized protein
Uncharacterized protein
Uncharacterized protein (Fragment)
Uncharacterized protein
Uncharacterized protein (Fragment)
Uncharacterized protein (Fragment)
Uncharacterized protein
Uncharacterized protein (Fragment)
Uncharacterized protein
Uncharacterized protein
Uncharacterized protein (Fragment)
Uncharacterized protein

Uncharacterized protein (Fragment)
Uncharacterized protein
Uncharacterized protein
Uncharacterized protein
Uncharacterized protein (Fragment)
Uncharacterized protein
Uncharacterized protein
Uncharacterized protein
Uncharacterized protein (Fragment)
Uncharacterized protein (Fragment)
Uncharacterized protein (Fragment)
Uncharacterized protein
Uncharacterized protein (Fragment)
Uncharacterized protein
Uncharacterized protein
Uncharacterized protein
Uncharacterized protein
Uncharacterized protein
Uncharacterized protein (Fragment)
Uncharacterized protein (Fragment)
Uncharacterized protein
Uncharacterized protein
Uncharacterized protein (Fragment)
Uncharacterized protein (Fragment)
Uncharacterized protein (Fragment)
Uncharacterized protein (Fragment)
Uncharacterized protein
Uncharacterized protein (Fragment)
Uncharacterized protein (Fragment)
Uncharacterized protein (Fragment)
Uncharacterized protein
Uncharacterized protein (Fragment)
Uncharacterized protein (Fragment)
Uncharacterized protein (Fragment)
Uncharacterized protein
Uncharacterized protein (Fragment)
Uncharacterized protein (Fragment)
Uncharacterized protein (Fragment)
Uncharacterized protein
Uncharacterized protein
Uncharacterized protein (Fragment)

Uncharacterized protein
Uncharacterized protein (Fragment)
Uncharacterized protein (Fragment)
Uncharacterized protein
Uncharacterized protein
Uncharacterized protein
Uncharacterized protein
Uncharacterized protein
Uncharacterized protein
Uncharacterized protein
Uncharacterized protein
Uncharacterized protein (Fragment)
Uncharacterized protein (Fragment)
Uncharacterized protein (Fragment)
Uncharacterized protein
Uncharacterized protein (Fragment)
Uncharacterized protein
Uncharacterized protein
Uncharacterized protein
Uncharacterized protein
Uncharacterized protein (Fragment)
Uncharacterized protein
Uncharacterized protein (Fragment)
Uncharacterized protein (Fragment)
Uncharacterized protein
Uncharacterized protein
Uncharacterized protein
Uncharacterized protein
Uncharacterized protein
Uncharacterized protein
Uncharacterized protein
Uncharacterized protein
Uncharacterized protein
Uncharacterized protein
Uncharacterized protein
Serine protease
Uncharacterized protein
Uncharacterized protein
Uncharacterized protein
Uncharacterized protein
Uncharacterized protein

Uncharacterized protein
Uncharacterized protein
Uncharacterized protein
Uncharacterized protein (Fragment)
Uncharacterized protein
Uncharacterized protein
Uncharacterized protein
Uncharacterized protein
Uncharacterized protein
Uncharacterized protein
Uncharacterized protein
Uncharacterized protein
Uncharacterized protein
Uncharacterized protein
Uncharacterized protein
Uncharacterized protein
Uncharacterized protein
Uncharacterized protein
Uncharacterized protein
Uncharacterized protein
Uncharacterized protein
Uncharacterized protein
Uncharacterized protein
Uncharacterized protein
Uncharacterized protein
Uncharacterized protein
Uncharacterized protein
Uncharacterized protein
Uncharacterized protein
Uncharacterized protein
Uncharacterized protein
Uncharacterized protein
Uncharacterized protein
Uncharacterized protein
Uncharacterized protein
Uncharacterized protein
Uncharacterized protein
Uncharacterized protein
Uncharacterized protein
Uncharacterized protein
Uncharacterized protein

Uncharacterized protein
Uncharacterized protein
Uncharacterized protein
Uncharacterized protein
Uncharacterized protein
Uncharacterized protein
Uncharacterized protein
Uncharacterized protein
Uncharacterized protein
Uncharacterized protein
Uncharacterized protein
Uncharacterized protein
Uncharacterized protein
Uncharacterized protein
Uncharacterized protein
Uncharacterized protein
Uncharacterized protein
Uncharacterized protein
Uncharacterized protein
Uncharacterized protein
Uncharacterized protein (Fragment)
Uncharacterized protein (Fragment)
Uncharacterized protein
Uncharacterized protein
Uncharacterized protein
Uncharacterized protein
Uncharacterized protein
Uncharacterized protein
Uncharacterized protein
Uncharacterized protein
Uncharacterized protein
Uncharacterized protein
Uncharacterized protein
Uncharacterized protein
Uncharacterized protein
Uncharacterized protein
Uncharacterized protein
Uncharacterized protein
Uncharacterized protein (Fragment)
Uncharacterized protein
Uncharacterized protein

Uncharacterized protein
Uncharacterized protein
Uncharacterized protein
Uncharacterized protein
Uncharacterized protein
Uncharacterized protein
Uncharacterized protein
Uncharacterized protein
Uncharacterized protein
Uncharacterized protein
Uncharacterized protein
Uncharacterized protein
Uncharacterized protein
Uncharacterized protein
Uncharacterized protein
Uncharacterized protein (Fragment)
Uncharacterized protein
Uncharacterized protein
Uncharacterized protein
Uncharacterized protein
Uncharacterized protein
Uncharacterized protein
Uncharacterized protein
Uncharacterized protein (Fragment)
Uncharacterized protein
Uncharacterized protein
Uncharacterized protein
Uncharacterized protein (Fragment)
Uncharacterized protein (Fragment)
Uncharacterized protein
Uncharacterized protein
Uncharacterized protein (Fragment)
Uncharacterized protein (Fragment)
Uncharacterized protein
Uncharacterized protein
Uncharacterized protein
Trypsin-like serine protease
Uncharacterized protein
Uncharacterized protein
Uncharacterized protein
Uncharacterized protein (Fragment)

Uncharacterized protein
Uncharacterized protein
Uncharacterized protein
Uncharacterized protein
Uncharacterized protein
Uncharacterized protein (Fragment)
Uncharacterized protein
Uncharacterized protein
Uncharacterized protein
Uncharacterized protein (Fragment)
Uncharacterized protein (Fragment)
Serine protease
Uncharacterized protein
Uncharacterized protein
Uncharacterized protein
Uncharacterized protein
Uncharacterized protein
Uncharacterized protein
Uncharacterized protein
Trypsin-like serine protease
Uncharacterized protein
Uncharacterized protein
Uncharacterized protein
Uncharacterized protein
Uncharacterized protein
Uncharacterized protein
Uncharacterized protein
Uncharacterized protein
Uncharacterized protein (Fragment)
Uncharacterized protein
Uncharacterized protein
Uncharacterized protein
Uncharacterized protein
Uncharacterized protein
Uncharacterized protein
Uncharacterized protein
Uncharacterized protein
Uncharacterized protein
Uncharacterized protein
Uncharacterized protein
Serine protease

Uncharacterized protein
Uncharacterized protein
Uncharacterized protein
Apple-like protein
Apple-like protein
Glycosyltransferase family 31 protein
Glycosyltransferase family 31 protein (Fragment)
Uncharacterized protein (Fragment)
PAN-2 domain protein
Uncharacterized protein
Uncharacterized protein
Jacalin-like lectin domain protein
Jacalin-like lectin domain protein
Uncharacterized protein
Peptidase S1 and S6, chymotrypsin/Hap
Uncharacterized protein
Uncharacterized protein
Uncharacterized protein
PAN domain protein
MG2 domain protein
Uncharacterized protein
Uncharacterized protein
Uncharacterized protein
Uncharacterized protein
Uncharacterized protein
Beta-glucan synthesis-associated SKN1
Ectonucleotide pyrophosphatase phosphodiesterase family member 2-like
Serine threonine-kinase
Argininosuccinate lyase
Low-CO2 inducible isoform B
Serine threonine-kinase CTR1
Uncharacterized protein
Extracellular basic protease
Peptidyl-prolyl cis-trans isomerase D
Inhibitor of Bruton tyrosine kinase
Serine threonine-kinase isoform B
Peptidyl-prolyl cis-trans cyclophilin-type
Cysteine isoform A
Serine threonine-kinase isoform A
Ribosomal RNA small subunit methyltransferase isoform A
Chitosanase

Cysteine isoform B
Band 7
Ribosomal RNA small subunit methyltransferase isoform B
Uncharacterized protein
Uncharacterized protein
Low-CO2 inducible isoform A
Serine threonine-kinase
BQ2448_6804 protein
Uncharacterized protein
Uncharacterized protein
Uncharacterized protein
Uncharacterized protein
Uncharacterized protein
Uncharacterized protein
Uncharacterized protein
Uncharacterized protein
Uncharacterized protein
Uncharacterized protein
Uncharacterized protein
Predicted protein
Predicted protein
Predicted protein
Uncharacterized protein
Putative PAN domain-containing protein
Putative PAN domain-containing protein
Uncharacterized protein
Putative pan domain-containing protein
Putative pan domain-containing protein
Uncharacterized protein
Uncharacterized protein
Uncharacterized protein
Uncharacterized protein
PAN domain-containing protein
Fibrinogen C domain-containing protein 1
Uncharacterized protein
Ig-like and fibronectin type-III domain-containing protein C25G4.10
Ficolin-3
CUB and sushi domain-containing protein 2
Lactadherin
Uncharacterized protein
Hemocytin

Fibropellin-1
Snaclec 3
Uncharacterized protein
Lactadherin
Uncharacterized protein
C3 and PZP-like alpha-2-macroglobulin domain-containing protein 8
Uncharacterized protein
Hemocytin
Sushi, von Willebrand factor type A, EGF and pentraxin domain-containing protein 1
Uncharacterized protein
Delta-like protein 1
Mucin-19
Brevican core protein
Uncharacterized protein
Tyrosine-protein phosphatase non-receptor type 9
Uncharacterized protein
Uncharacterized protein
IgGFc-binding protein
Snaclec rhinocetin subunit beta
Neuropilin-2
Coagulation factor VIII
Uncharacterized protein
von Willebrand factor D and EGF domain-containing protein
Hepatocyte growth factor a
Macrophage stimulating 1
Hepatocyte growth factor
Macrophage stimulating 1
Macrophage stimulating 1
Plasminogen
Uncharacterized protein
Uncharacterized protein
Uncharacterized protein
Uncharacterized protein (Fragment)
Uncharacterized protein
Uncharacterized protein
Uncharacterized protein
Plasminogen
Coagulation factor XI
Hepatocyte growth factor
Uncharacterized protein
Uncharacterized protein

Uncharacterized protein
Uncharacterized protein
Hepatocyte growth factor
Uncharacterized protein
Uncharacterized protein
Uncharacterized protein
Uncharacterized protein
Uncharacterized protein
Peptidylprolyl isomerase D (Cyclophilin D)
Uncharacterized protein
Uncharacterized protein
Uncharacterized protein
Predicted protein
Predicted protein
Predicted protein
Predicted protein
Predicted protein
Predicted protein
Predicted protein
Predicted protein
Predicted protein
Predicted protein
Predicted protein
Predicted protein
Predicted protein
Predicted protein
Uncharacterized protein
Predicted protein
Predicted protein
S-locus-related I (Fragment)
G-type lectin S-receptor-like serine/threonine-protein kinase
G-type lectin S-receptor-like serine/threonine-protein kinase SD2-2
Receptor-like serine/threonine-protein kinase SD1-8
G-type lectin S-receptor-like serine/threonine-protein kinase
Uncharacterized protein
G-type lectin S-receptor-like serine/threonine-protein kinase RKS1
G-type lectin S-receptor-like serine/threonine-protein kinase
G-type lectin S-receptor-like serine/threonine-protein kinase
Putative receptor protein kinase ZmPK1
G-type lectin S-receptor-like serine/threonine-protein kinase RKS1
Epidermis-specific secreted glycoprotein EP1

G-type lectin S-receptor-like serine/threonine-protein kinase RLK1
G-type lectin S-receptor-like serine/threonine-protein kinase SD1-1
G-type lectin S-receptor-like serine/threonine-protein kinase
G-type lectin S-receptor-like serine/threonine-protein kinase
Putative receptor protein kinase ZmPK1
G-type lectin S-receptor-like serine/threonine-protein kinase
G-type lectin S-receptor-like serine/threonine-protein kinase RLK1
G-type lectin S-receptor-like serine/threonine-protein kinase
G-type lectin S-receptor-like serine/threonine-protein kinase
G-type lectin S-receptor-like serine/threonine-protein kinase
G-type lectin S-receptor-like serine/threonine-protein kinase
G-type lectin S-receptor-like serine/threonine-protein kinase RKS1
G-type lectin S-receptor-like serine/threonine-protein kinase RKS1
PAN domain-containing protein
G-type lectin S-receptor-like serine/threonine-protein kinase
Receptor-like serine/threonine-protein kinase SD1-8
Putative receptor protein kinase ZmPK1
G-type lectin S-receptor-like serine/threonine-protein kinase SD2-5
PAN domain-containing protein
G-type lectin S-receptor-like serine/threonine-protein kinase
G-type lectin S-receptor-like serine/threonine-protein kinase RKS1
Putative receptor protein kinase ZmPK1
G-type lectin S-receptor-like serine/threonine-protein kinase RLK1
G-type lectin S-receptor-like serine/threonine-protein kinase SD1-1
G-type lectin S-receptor-like serine/threonine-protein kinase
Epidermis-specific secreted glycoprotein EP1
G-type lectin S-receptor-like serine/threonine-protein kinase SD3-1
G-type lectin S-receptor-like serine/threonine-protein kinase
G-type lectin S-receptor-like serine/threonine-protein kinase
G-type lectin S-receptor-like serine/threonine-protein kinase
Receptor-like serine/threonine-protein kinase SD1-8
G-type lectin S-receptor-like serine/threonine-protein kinase
Receptor-like serine/threonine-protein kinase SD1-8
Putative receptor protein kinase ZmPK1
G-type lectin S-receptor-like serine/threonine-protein kinase RKS1
G-type lectin S-receptor-like serine/threonine-protein kinase
Uncharacterized protein
PAN domain-containing protein
PAN domain-containing protein
Uncharacterized protein
Uncharacterized protein

Uncharacterized protein
Putative PAN domain-containing protein
Uncharacterized protein
Putative PAN domain-containing protein
Receptor-like serine/threonine-protein kinase SD1-8 (Fragment)
G-type lectin S-receptor-like serine/threonine-protein kinase (Fragment)
G-type lectin S-receptor-like serine/threonine-protein kinase (Fragment)
EP1-like glycoprotein 3 (Fragment)
G-type lectin S-receptor-like serine/threonine-protein kinase (Fragment)
G-type lectin S-receptor-like serine/threonine-protein kinase (Fragment)
G-type lectin S-receptor-like serine/threonine-protein kinase (Fragment)
G-type lectin S-receptor-like serine/threonine-protein kinase (Fragment)
G-type lectin S-receptor-like serine/threonine-protein kinase (Fragment)
G-type lectin S-receptor-like serine/threonine-protein kinase (Fragment)
Putative receptor protein kinase ZmPK1 (Fragment)
G-type lectin S-receptor-like serine/threonine-protein kinase (Fragment)
G-type lectin S-receptor-like serine/threonine-protein kinase (Fragment)
G-type lectin S-receptor-like serine/threonine-protein kinase (Fragment)
G-type lectin S-receptor-like serine/threonine-protein kinase (Fragment)
G-type lectin S-receptor-like serine/threonine-protein kinase (Fragment)
G-type lectin S-receptor-like serine/threonine-protein kinase (Fragment)
EP1-like glycoprotein 3 (Fragment)
G-type lectin S-receptor-like serine/threonine-protein kinase (Fragment)
G-type lectin S-receptor-like serine/threonine-protein kinase (Fragment)
G-type lectin S-receptor-like serine/threonine-protein kinase (Fragment)
G-type lectin S-receptor-like serine/threonine-protein kinase (Fragment)
G-type lectin S-receptor-like serine/threonine-protein kinase SD1-1 (Fragment)
G-type lectin S-receptor-like serine/threonine-protein kinase (Fragment)
G-type lectin S-receptor-like serine/threonine-protein kinase (Fragment)
G-type lectin S-receptor-like serine/threonine-protein kinase (Fragment)
Mitochondrial arginine transporter BAC2 (Fragment)
Putative receptor protein kinase ZmPK1 (Fragment)
Receptor-like serine/threonine-protein kinase SD1-8 (Fragment)
G-type lectin S-receptor-like serine/threonine-protein kinase (Fragment)
Receptor-like serine/threonine-protein kinase SD1-8 (Fragment)
G-type lectin S-receptor-like serine/threonine-protein kinase (Fragment)
G-type lectin S-receptor-like serine/threonine-protein kinase (Fragment)
G-type lectin S-receptor-like serine/threonine-protein kinase (Fragment)
G-type lectin S-receptor-like serine/threonine-protein kinase (Fragment)
G-type lectin S-receptor-like serine/threonine-protein kinase CES101 (Fragment)
G-type lectin S-receptor-like serine/threonine-protein kinase (Fragment)

Putative receptor protein kinase ZmPK1 (Fragment)
G-type lectin S-receptor-like serine/threonine-protein kinase (Fragment)
G-type lectin S-receptor-like serine/threonine-protein kinase LECRK1 (Fragment)
G-type lectin S-receptor-like serine/threonine-protein kinase (Fragment)
G-type lectin S-receptor-like serine/threonine-protein kinase SD2-2 (Fragment)
G-type lectin S-receptor-like serine/threonine-protein kinase (Fragment)
Receptor-like serine/threonine-protein kinase SD1-8 (Fragment)
Receptor-like serine/threonine-protein kinase SD1-8 (Fragment)
Receptor-like serine/threonine-protein kinase SD1-8 (Fragment)
G-type lectin S-receptor-like serine/threonine-protein kinase (Fragment)
G-type lectin S-receptor-like serine/threonine-protein kinase (Fragment)
G-type lectin S-receptor-like serine/threonine-protein kinase (Fragment)
Receptor-like serine/threonine-protein kinase SD1-8 (Fragment)
G-type lectin S-receptor-like serine/threonine-protein kinase CES101 (Fragment)
G-type lectin S-receptor-like serine/threonine-protein kinase (Fragment)
G-type lectin S-receptor-like serine/threonine-protein kinase (Fragment)
G-type lectin S-receptor-like serine/threonine-protein kinase (Fragment)
G-type lectin S-receptor-like serine/threonine-protein kinase (Fragment)
G-type lectin S-receptor-like serine/threonine-protein kinase (Fragment)
Putative receptor protein kinase ZmPK1 (Fragment)
G-type lectin S-receptor-like serine/threonine-protein kinase (Fragment)
G-type lectin S-receptor-like serine/threonine-protein kinase (Fragment)
Uncharacterized protein (Fragment)
Putative receptor protein kinase ZmPK1 (Fragment)
G-type lectin S-receptor-like serine/threonine-protein kinase SD1-1 (Fragment)
G-type lectin S-receptor-like serine/threonine-protein kinase (Fragment)
Putative receptor protein kinase ZmPK1 (Fragment)
G-type lectin S-receptor-like serine/threonine-protein kinase (Fragment)
G-type lectin S-receptor-like serine/threonine-protein kinase CES101 (Fragment)
G-type lectin S-receptor-like serine/threonine-protein kinase SD2-5 (Fragment)
G-type lectin S-receptor-like serine/threonine-protein kinase (Fragment)
G-type lectin S-receptor-like serine/threonine-protein kinase B120 (Fragment)
G-type lectin S-receptor-like serine/threonine-protein kinase (Fragment)
G-type lectin S-receptor-like serine/threonine-protein kinase (Fragment)
G-type lectin S-receptor-like serine/threonine-protein kinase (Fragment)
G-type lectin S-receptor-like serine/threonine-protein kinase (Fragment)
Putative receptor protein kinase ZmPK1 (Fragment)
Acidic endochitinase (Fragment)
Receptor-like serine/threonine-protein kinase SD1-8 (Fragment)
Receptor-like serine/threonine-protein kinase SD1-8 (Fragment)
Pentatricopeptide repeat-containing protein (Fragment)

G-type lectin S-receptor-like serine/threonine-protein kinase SD3-1 (Fragment)
G-type lectin S-receptor-like serine/threonine-protein kinase (Fragment)
G-type lectin S-receptor-like serine/threonine-protein kinase (Fragment)
G-type lectin S-receptor-like serine/threonine-protein kinase (Fragment)
Receptor-like serine/threonine-protein kinase SD1-8 (Fragment)
G-type lectin S-receptor-like serine/threonine-protein kinase (Fragment)
G-type lectin S-receptor-like serine/threonine-protein kinase (Fragment)
Receptor-like serine/threonine-protein kinase SD1-8 (Fragment)
G-type lectin S-receptor-like serine/threonine-protein kinase RKS1 (Fragment)
G-type lectin S-receptor-like serine/threonine-protein kinase (Fragment)
Receptor-like serine/threonine-protein kinase SD1-8 (Fragment)
G-type lectin S-receptor-like serine/threonine-protein kinase (Fragment)
Plasminogen
Hepatocyte growth factor
Hepatocyte growth factor-like protein
Plasma kallikrein
Plasminogen
Angiostatin
Uncharacterized protein
Hepatocyte growth factor-like protein
Plasma kallikrein
Uncharacterized protein
Macrophage stimulating 1 (Hepatocyte growth factor-like)
Hepatocyte growth factor
Coagulation factor XI
Uncharacterized protein
Uncharacterized protein
Uncharacterized protein
Uncharacterized protein
Uncharacterized protein
Uncharacterized protein
Uncharacterized protein
Uncharacterized protein
Uncharacterized protein
Uncharacterized protein
Uncharacterized protein
Uncharacterized protein
Uncharacterized protein
Uncharacterized protein
Uncharacterized protein
Uncharacterized protein

Uncharacterized protein
Uncharacterized protein
Uncharacterized protein
Uncharacterized protein
Uncharacterized protein
Uncharacterized protein
Uncharacterized protein
Uncharacterized protein
Uncharacterized protein
Uncharacterized protein
Uncharacterized protein
Uncharacterized protein
Uncharacterized protein
Uncharacterized protein
Uncharacterized protein
Uncharacterized protein
Uncharacterized protein
Uncharacterized protein
Uncharacterized protein
Uncharacterized protein
Uncharacterized protein
Uncharacterized protein
Hepatocyte growth factor (Fragment)
Uncharacterized protein
Hepatocyte growth factor
Uncharacterized protein
Uncharacterized protein
Plasminogen
Uncharacterized protein
Uncharacterized protein
PAN domain-containing protein
Plasminogen
Plasma kallikrein
Hepatocyte growth factor-like protein
Hepatocyte growth factor
Plasma kallikrein
Hepatocyte growth factor-like protein
Plasminogen
Plasminogen
Uncharacterized protein
Hepatocyte growth factor

Uncharacterized protein (Fragment)
Uncharacterized protein (Fragment)
Protein jagged-2 (Fragment)
Uncharacterized protein (Fragment)
Peptidyl-prolyl cis-trans isomerase
Apple domain-containing protein (Fragment)
Uncharacterized protein
Trypsin
Uncharacterized protein
Peptidyl-prolyl cis-trans isomerase
Uncharacterized protein
Uncharacterized protein
Uncharacterized protein
Uncharacterized protein
Uncharacterized protein
PAN domain protein (Fragment)
PAN domain protein
PAN domain protein (Fragment)
PAN domain protein (Fragment)
PAN domain protein
PAN domain protein (Fragment)
PAN domain protein (Fragment)
PAN domain protein
PAN domain protein
PAN domain protein
PAN domain protein
PAN domain protein
PAN domain protein
PAN domain protein (Fragment)
PAN domain protein
PAN domain protein
Uncharacterized protein
PAN domain protein
PAN domain protein
Uncharacterized protein
PAN domain protein
PAN domain protein
PAN domain protein
Predicted protein
Predicted protein
Glycosyltransferase family 31

Predicted protein
Predicted protein
Uncharacterized protein
Expressed protein
Uncharacterized protein
Uncharacterized protein (Fragment)
Predicted protein
Uncharacterized protein
Predicted protein
Uncharacterized protein
Uncharacterized protein
Predicted protein
G-type lectin S-receptor-like serine/threonine-protein kinase At4g03230 isoform X2
G-type lectin S-receptor-like serine/threonine-protein kinase RKS1
receptor-like serine/threonine-protein kinase SD1-8
G-type lectin S-receptor-like serine/threonine-protein kinase SD1-13
G-type lectin S-receptor-like serine/threonine-protein kinase SD2-5 isoform X3
putative receptor protein kinase ZmPK1
receptor-like serine/threonine-protein kinase SD1-8
G-type lectin S-receptor-like serine/threonine-protein kinase LECRK1
putative receptor protein kinase ZmPK1
putative receptor protein kinase ZmPK1
G-type lectin S-receptor-like serine/threonine-protein kinase At4g03230 isoform X1
G-type lectin S-receptor-like serine/threonine-protein kinase SD2-5 isoform X2
G-type lectin S-receptor-like serine/threonine-protein kinase At4g03230 isoform X3
G-type lectin S-receptor-like serine/threonine-protein kinase At4g27290
G-type lectin S-receptor-like serine/threonine-protein kinase SD2-2
putative receptor protein kinase ZmPK1
G-type lectin S-receptor-like serine/threonine-protein kinase At4g27290
G-type lectin S-receptor-like serine/threonine-protein kinase At4g27290 isoform X2
LOW QUALITY PROTEIN: G-type lectin S-receptor-like serine/threonine-protein kinase At4g27290
G-type lectin S-receptor-like serine/threonine-protein kinase At1g34300
G-type lectin S-receptor-like serine/threonine-protein kinase At2g19130 isoform X2
G-type lectin S-receptor-like serine/threonine-protein kinase LECRK3
G-type lectin S-receptor-like serine/threonine-protein kinase RKS1
G-type lectin S-receptor-like serine/threonine-protein kinase SD2-5 isoform X1
G-type lectin S-receptor-like serine/threonine-protein kinase At4g03230
putative receptor protein kinase ZmPK1
G-type lectin S-receptor-like serine/threonine-protein kinase At2g19130
G-type lectin S-receptor-like serine/threonine-protein kinase At4g27290
putative G-type lectin S-receptor-like serine/threonine-protein kinase At1g61610

G-type lectin S-receptor-like serine/threonine-protein kinase CES101 isoform X1
G-type lectin S-receptor-like serine/threonine-protein kinase SD2-5
G-type lectin S-receptor-like serine/threonine-protein kinase RKS1
G-type lectin S-receptor-like serine/threonine-protein kinase At4g03230
G-type lectin S-receptor-like serine/threonine-protein kinase At1g11330
PAN domain-containing protein At5g03700
epidermis-specific secreted glycoprotein EP1-like
G-type lectin S-receptor-like serine/threonine-protein kinase B120
G-type lectin S-receptor-like serine/threonine-protein kinase SD3-1
G-type lectin S-receptor-like serine/threonine-protein kinase LECRK4
LOW QUALITY PROTEIN: uncharacterized protein LOC104592606
G-type lectin S-receptor-like serine/threonine-protein kinase CES101 isoform X2
G-type lectin S-receptor-like serine/threonine-protein kinase At1g11300
G-type lectin S-receptor-like serine/threonine-protein kinase SD3-1
putative receptor protein kinase ZmPK1
G-type lectin S-receptor-like serine/threonine-protein kinase At4g03230 isoform X2
G-type lectin S-receptor-like serine/threonine-protein kinase At4g27290 isoform X1
G-type lectin S-receptor-like serine/threonine-protein kinase SD3-1
G-type lectin S-receptor-like serine/threonine-protein kinase SD2-5
G-type lectin S-receptor-like serine/threonine-protein kinase SD2-5
G-type lectin S-receptor-like serine/threonine-protein kinase At4g27290
putative receptor protein kinase ZmPK1
G-type lectin S-receptor-like serine/threonine-protein kinase At4g27290
G-type lectin S-receptor-like serine/threonine-protein kinase At1g11330
LOW QUALITY PROTEIN: uncharacterized protein LOC104605769
putative receptor protein kinase ZmPK1
G-type lectin S-receptor-like serine/threonine-protein kinase At2g19130
LOW QUALITY PROTEIN: G-type lectin S-receptor-like serine/threonine-protein kinase RLK1
epidermis-specific secreted glycoprotein EP1-like
uncharacterized protein LOC104585713
G-type lectin S-receptor-like serine/threonine-protein kinase SD2-5
receptor-like serine/threonine-protein kinase SD1-8 isoform X3
putative receptor protein kinase ZmPK1
G-type lectin S-receptor-like serine/threonine-protein kinase At2g19130 isoform X1
G-type lectin S-receptor-like serine/threonine-protein kinase At4g03230 isoform X1
LOW QUALITY PROTEIN: uncharacterized protein LOC104592220
G-type lectin S-receptor-like serine/threonine-protein kinase At4g03230 isoform X4
uncharacterized protein LOC104610592
G-type lectin S-receptor-like serine/threonine-protein kinase At1g11330
G-type lectin S-receptor-like serine/threonine-protein kinase SD3-1
Predicted protein

Predicted protein
Predicted protein
Predicted protein
Predicted protein
Predicted protein
Predicted protein
Predicted protein
Predicted protein
Predicted protein
Predicted protein
Predicted protein
Predicted protein
Predicted protein
Predicted protein
Predicted protein (Fragment)
Predicted protein
Predicted protein
Predicted protein
Predicted protein
Predicted protein
Predicted protein
Predicted protein
Predicted protein
Predicted protein
Predicted protein
Predicted protein
Predicted protein
Predicted protein
Predicted protein
Predicted protein
Predicted protein
Predicted protein
Predicted protein
Predicted protein
Predicted protein
Predicted protein
Predicted protein
Predicted protein
Predicted protein
Predicted protein
Predicted protein

Predicted protein
Predicted protein
Predicted protein
Predicted protein
Predicted protein
Predicted protein
Predicted protein
Predicted protein
Predicted protein
Predicted protein
Predicted protein
Predicted protein
Predicted protein
Predicted protein
Predicted protein
Predicted protein
Predicted protein
Predicted protein
Predicted protein
Predicted protein
Predicted protein
Predicted protein
Predicted protein
Predicted protein
Predicted protein
Predicted protein
Predicted protein
Predicted protein
Predicted protein
Predicted protein
Predicted protein
Predicted protein (Fragment)
Predicted protein
Predicted protein
Predicted protein
Predicted protein
Predicted protein
Predicted protein (Fragment)
Predicted protein
Predicted protein
Predicted protein

Predicted protein
Predicted protein
Predicted protein
Predicted protein
Predicted protein
Predicted protein (Fragment)
Predicted protein
Predicted protein
Predicted protein
Predicted protein
Predicted protein
Predicted protein
Predicted protein
Predicted protein
Predicted protein
Predicted protein
Uncharacterized protein
Uncharacterized protein
Plasminogen
Uncharacterized protein
Uncharacterized protein
Plasminogen
Uncharacterized protein
Plasminogen
Plasminogen
Hepatocyte growth factor
Uncharacterized protein
Hepatocyte growth factor
plasma kallikrein isoform X2
plasma kallikrein isoform X1
Plasminogen
coagulation factor XI isoform X1
Plasminogen
hepatocyte growth factor-like protein
coagulation factor XI isoform X2
Hepatocyte growth factor
Uncharacterized protein
Uncharacterized protein
Uncharacterized protein
Uncharacterized protein
Uncharacterized protein

Uncharacterized protein
Uncharacterized protein
Uncharacterized protein
Uncharacterized protein
Uncharacterized protein
Uncharacterized protein
Uncharacterized protein
Uncharacterized protein
Uncharacterized protein
Uncharacterized protein
Plasminogen
Plasminogen
hepatocyte growth factor isoform X1
Plasminogen
LOW QUALITY PROTEIN: hepatocyte growth factor-like protein
Hepatocyte growth factor
plasminogen isoform X5
coagulation factor XI
Plasminogen
Alpha-2-macroglobulin domain protein
Alpha-2-macroglobulin family protein
Alpha-2-macroglobulin family protein
Alpha-2-macroglobulin family protein
Alpha-2-macroglobulin domain protein
Alpha-2-macroglobulin family protein
Membrane protein
Uncharacterized protein
PAN domain
Uncharacterized protein
Uncharacterized protein
Uncharacterized protein
Uncharacterized protein
Uncharacterized protein
PAN domain-containing protein, putative
PAN domain-containing protein
Uncharacterized protein
Uncharacterized protein
Microneme protein, putative
PAN domain-containing protein, related
Microneme protein 5 (Precursor), related
PAN domain-containing protein, putative

Microneme protein MIC4, putative
Micronemal protein 4
PAN domain-containing protein, putative
Uncharacterized protein
PAN domain-containing protein
PAN domain-containing protein
Uncharacterized protein
PAN domain-containing protein
Putative PAN domain-containing protein
Putative PAN domain containing protein
PAN domain-containing protein, putative
Subtilisin-like serine protease, putative
PAN domain containing protein, putative
Putative subtilisin-like serine protease
Uncharacterized protein
PAN domain-containing protein, putative
PAN domain-containing protein
Microneme protein MIC17B
Microneme protein 17 B
Microneme protein 17 C
Microneme protein 17 A
Microneme protein 4
Uncharacterized protein
Uncharacterized protein
Uncharacterized protein
Plasminogen
Uncharacterized protein (Fragment)
Hepatocyte growth factor
Uncharacterized protein
Uncharacterized protein
Uncharacterized protein
Uncharacterized protein
Uncharacterized protein
Uncharacterized protein
Uncharacterized protein
Uncharacterized protein
Uncharacterized protein
Uncharacterized protein
AT4G21390-like protein (Fragment)
AT4G27290-like protein (Fragment)
Plasminogen (Fragment)

Coagulation factor XI (Fragment)
Plasminogen (Fragment)
Hepatocyte growth factor (Fragment)
Uncharacterized protein
Uncharacterized protein
Uncharacterized protein
Uncharacterized protein
Uncharacterized protein
Uncharacterized protein
Uncharacterized protein B7A16.095
Glycosyltransferase family 31 protein
Uncharacterized protein
WSC-domain-containing protein
Uncharacterized protein
Uncharacterized protein
Uncharacterized protein
Uncharacterized protein
Uncharacterized protein
Uncharacterized protein
WSC-domain-containing protein
Uncharacterized protein
Receptor-like serinethreonine-protein kinase sd1-8
G-type lectin s-receptor-like serinethreonine-protein kinase
G-type lectin s-receptor-like serinethreonine-protein kinase sd1-1
G-type lectin s-receptor-like serinethreonine-protein kinase
Receptor-like serinethreonine-protein kinase sd1-8
G-type lectin s-receptor-like serinethreonine-protein kinase
G-type lectin s-receptor-like serinethreonine-protein kinase
G-type lectin s-receptor-like serinethreonine-protein kinase
G-type lectin s-receptor-like serinethreonine-protein kinase
G-type lectin s-receptor-like serinethreonine-protein kinase
G-type lectin s-receptor-like serinethreonine-protein kinase b120
G-type lectin s-receptor-like serinethreonine-protein kinase
G-type lectin s-receptor-like serinethreonine-protein kinase
G-type lectin s-receptor-like serinethreonine-protein kinase sd1-29
G-type lectin s-receptor-like serinethreonine-protein kinase sd2-5
Putative receptor protein kinase zmpk1
G-type lectin s-receptor-like serinethreonine-protein kinase sd2-5 (Fragment)
G-type lectin s-receptor-like serinethreonine-protein kinase
Uncharacterized protein
G-type lectin s-receptor-like serinethreonine-protein kinase

G-type lectin s-receptor-like serinethreonine-protein kinase
G-type lectin s-receptor-like serinethreonine-protein kinase
G-type lectin s-receptor-like serinethreonine-protein kinase
G-type lectin s-receptor-like serinethreonine-protein kinase sd1-1
Receptor-like serinethreonine-protein kinase sd1-8
Receptor-like serinethreonine-protein kinase sd1-8
G-type lectin s-receptor-like serinethreonine-protein kinase sd2-5
G-type lectin s-receptor-like serinethreonine-protein kinase
G-type lectin s-receptor-like serinethreonine-protein kinase sd1-1
G-type lectin s-receptor-like serinethreonine-protein kinase
G-type lectin s-receptor-like serinethreonine-protein kinase
G-type lectin s-receptor-like serinethreonine-protein kinase
G-type lectin s-receptor-like serinethreonine-protein kinase sd2-5
G-type lectin s-receptor-like serinethreonine-protein kinase rks1
G-type lectin s-receptor-like serinethreonine-protein kinase sd1-1 (Fragment)
Putative receptor protein kinase zmpk1
Receptor-like serinethreonine-protein kinase sd1-8
G-type lectin s-receptor-like serinethreonine-protein kinase sd1-1
G-type lectin s-receptor-like serinethreonine-protein kinase
G-type lectin s-receptor-like serinethreonine-protein kinase sd2-5
G-type lectin s-receptor-like serinethreonine-protein kinase
G-type lectin s-receptor-like serinethreonine-protein kinase ces101
Receptor-like serinethreonine-protein kinase sd1-8
G-type lectin s-receptor-like serinethreonine-protein kinase sd2-5
G-type lectin s-receptor-like serinethreonine-protein kinase
G-type lectin s-receptor-like serinethreonine-protein kinase sd2-5
Lectin-domain receptor-like kinase
G-type lectin s-receptor-like serinethreonine-protein kinase
G-type lectin s-receptor-like serinethreonine-protein kinase
G-type lectin s-receptor-like serinethreonine-protein kinase sd2-5
G-type lectin s-receptor-like serinethreonine-protein kinase
Pan domain-containing protein
G-type lectin s-receptor-like serinethreonine-protein kinase lecrk4
G-type lectin s-receptor-like serinethreonine-protein kinase sd2-2
G-type lectin s-receptor-like serinethreonine-protein kinase sd3-1
S locus glycoprotein like protein
Receptor-like kinase
G-type lectin S-receptor-like serine/threonine-protein kinase At1g11330 isoform X2
receptor-like serine/threonine-protein kinase SD1-8
G-type lectin S-receptor-like serine/threonine-protein kinase At4g03230 isoform X2
G-type lectin S-receptor-like serine/threonine-protein kinase At4g27290 isoform X1

G-type lectin S-receptor-like serine/threonine-protein kinase At4g03230 isoform X3
G-type lectin S-receptor-like serine/threonine-protein kinase At4g27290 isoform X2
G-type lectin S-receptor-like serine/threonine-protein kinase At4g03230 isoform X1
G-type lectin S-receptor-like serine/threonine-protein kinase At4g27290 isoform X2
G-type lectin S-receptor-like serine/threonine-protein kinase B120
G-type lectin S-receptor-like serine/threonine-protein kinase RLK1
G-type lectin S-receptor-like serine/threonine-protein kinase At4g27290 isoform X1
G-type lectin S-receptor-like serine/threonine-protein kinase At4g27290 isoform X1
G-type lectin S-receptor-like serine/threonine-protein kinase At1g11330 isoform X2
G-type lectin S-receptor-like serine/threonine-protein kinase At4g03230 isoform X5
putative receptor protein kinase ZmPK1
G-type lectin S-receptor-like serine/threonine-protein kinase At4g27290 isoform X2
G-type lectin S-receptor-like serine/threonine-protein kinase At4g27290 isoform X2
G-type lectin S-receptor-like serine/threonine-protein kinase At1g11330 isoform X1
G-type lectin S-receptor-like serine/threonine-protein kinase At4g27290
G-type lectin S-receptor-like serine/threonine-protein kinase CES101 isoform X1
G-type lectin S-receptor-like serine/threonine-protein kinase SD2-5
G-type lectin S-receptor-like serine/threonine-protein kinase At4g27290 isoform X1
G-type lectin S-receptor-like serine/threonine-protein kinase SD2-5
G-type lectin S-receptor-like serine/threonine-protein kinase At4g03230 isoform X2
receptor-like serine/threonine-protein kinase SD1-8 isoform X6
receptor-like serine/threonine-protein kinase SD1-8 isoform X1
G-type lectin S-receptor-like serine/threonine-protein kinase CES101 isoform X2
receptor-like serine/threonine-protein kinase SD1-8 isoform X2
G-type lectin S-receptor-like serine/threonine-protein kinase At4g27290 isoform X1
G-type lectin S-receptor-like serine/threonine-protein kinase CES101 isoform X1
G-type lectin S-receptor-like serine/threonine-protein kinase SD2-2
LOW QUALITY PROTEIN: G-type lectin S-receptor-like serine/threonine-protein kinase At1g11330
G-type lectin S-receptor-like serine/threonine-protein kinase At4g27290 isoform X1
G-type lectin S-receptor-like serine/threonine-protein kinase At4g03230 isoform X1
G-type lectin S-receptor-like serine/threonine-protein kinase RKS1
G-type lectin S-receptor-like serine/threonine-protein kinase At4g27290 isoform X1
G-type lectin S-receptor-like serine/threonine-protein kinase At2g19130
G-type lectin S-receptor-like serine/threonine-protein kinase At4g27290
G-type lectin S-receptor-like serine/threonine-protein kinase At1g11330 isoform X3
G-type lectin S-receptor-like serine/threonine-protein kinase At4g27290 isoform X1
G-type lectin S-receptor-like serine/threonine-protein kinase At4g03230 isoform X2
G-type lectin S-receptor-like serine/threonine-protein kinase At4g27290
G-type lectin S-receptor-like serine/threonine-protein kinase At4g03230 isoform X1
receptor-like serine/threonine-protein kinase SD1-8 isoform X3
G-type lectin S-receptor-like serine/threonine-protein kinase At1g11410

G-type lectin S-receptor-like serine/threonine-protein kinase At4g27290 isoform X3
G-type lectin S-receptor-like serine/threonine-protein kinase SD2-5
receptor-like serine/threonine-protein kinase SD1-8
G-type lectin S-receptor-like serine/threonine-protein kinase At4g27290 isoform X2
G-type lectin S-receptor-like serine/threonine-protein kinase At4g27290 isoform X1
G-type lectin S-receptor-like serine/threonine-protein kinase At4g27290 isoform X2
G-type lectin S-receptor-like serine/threonine-protein kinase At4g27290 isoform X3
G-type lectin S-receptor-like serine/threonine-protein kinase SD1-29 isoform X2
G-type lectin S-receptor-like serine/threonine-protein kinase At4g27290
G-type lectin S-receptor-like serine/threonine-protein kinase At4g27290 isoform X3
LOW QUALITY PROTEIN: G-type lectin S-receptor-like serine/threonine-protein kinase At1g61500
G-type lectin S-receptor-like serine/threonine-protein kinase At4g03230 isoform X2
G-type lectin S-receptor-like serine/threonine-protein kinase At4g27290
G-type lectin S-receptor-like serine/threonine-protein kinase At4g03230
G-type lectin S-receptor-like serine/threonine-protein kinase At4g27290 isoform X2
receptor-like serine/threonine-protein kinase SD1-8 isoform X3
G-type lectin S-receptor-like serine/threonine-protein kinase At1g61550
G-type lectin S-receptor-like serine/threonine-protein kinase At4g03230 isoform X1
G-type lectin S-receptor-like serine/threonine-protein kinase SD1-1
putative G-type lectin S-receptor-like serine/threonine-protein kinase At1g61610 isoform X2
G-type lectin S-receptor-like serine/threonine-protein kinase At4g03230 isoform X4
uncharacterized protein LOC104227947
G-type lectin S-receptor-like serine/threonine-protein kinase At4g27290 isoform X2
putative receptor protein kinase ZmPK1
G-type lectin S-receptor-like serine/threonine-protein kinase SD2-5
G-type lectin S-receptor-like serine/threonine-protein kinase At1g11330 isoform X3
G-type lectin S-receptor-like serine/threonine-protein kinase SD1-29 isoform X1
G-type lectin S-receptor-like serine/threonine-protein kinase At1g34300
G-type lectin S-receptor-like serine/threonine-protein kinase At4g27290 isoform X3
receptor-like serine/threonine-protein kinase SD1-8
G-type lectin S-receptor-like serine/threonine-protein kinase SD2-5
PAN domain-containing protein At5g03700
G-type lectin S-receptor-like serine/threonine-protein kinase At1g11330 isoform X2
G-type lectin S-receptor-like serine/threonine-protein kinase At1g11330 isoform X1
G-type lectin S-receptor-like serine/threonine-protein kinase At4g27290 isoform X2
G-type lectin S-receptor-like serine/threonine-protein kinase At1g11330 isoform X1
G-type lectin S-receptor-like serine/threonine-protein kinase At4g03230
G-type lectin S-receptor-like serine/threonine-protein kinase B120
G-type lectin S-receptor-like serine/threonine-protein kinase CES101 isoform X2
G-type lectin S-receptor-like serine/threonine-protein kinase SD3-1
receptor-like serine/threonine-protein kinase SD1-7 isoform X1

G-type lectin S-receptor-like serine/threonine-protein kinase At2g19130
G-type lectin S-receptor-like serine/threonine-protein kinase RLK1
putative receptor protein kinase ZmPK1
G-type lectin S-receptor-like serine/threonine-protein kinase At4g27290 isoform X2
G-type lectin S-receptor-like serine/threonine-protein kinase SD2-5
G-type lectin S-receptor-like serine/threonine-protein kinase At4g27290 isoform X1
G-type lectin S-receptor-like serine/threonine-protein kinase At4g27290
G-type lectin S-receptor-like serine/threonine-protein kinase At4g27290 isoform X2
G-type lectin S-receptor-like serine/threonine-protein kinase At4g27290 isoform X1
G-type lectin S-receptor-like serine/threonine-protein kinase SD2-5
G-type lectin S-receptor-like serine/threonine-protein kinase At4g27290 isoform X1
G-type lectin S-receptor-like serine/threonine-protein kinase At4g27290
G-type lectin S-receptor-like serine/threonine-protein kinase At4g27290
G-type lectin S-receptor-like serine/threonine-protein kinase At4g03230
G-type lectin S-receptor-like serine/threonine-protein kinase At5g35370
G-type lectin S-receptor-like serine/threonine-protein kinase SD2-5
PAN domain-containing protein At5g03700-like
G-type lectin S-receptor-like serine/threonine-protein kinase At4g03230 isoform X2
G-type lectin S-receptor-like serine/threonine-protein kinase CES101 isoform X3
G-type lectin S-receptor-like serine/threonine-protein kinase At1g11330 isoform X2
G-type lectin S-receptor-like serine/threonine-protein kinase SD2-5 isoform X1
G-type lectin S-receptor-like serine/threonine-protein kinase SD2-5
G-type lectin S-receptor-like serine/threonine-protein kinase At4g27290 isoform X3
G-type lectin S-receptor-like serine/threonine-protein kinase At1g11330 isoform X1
G-type lectin S-receptor-like serine/threonine-protein kinase At1g11410
G-type lectin S-receptor-like serine/threonine-protein kinase RLK1
G-type lectin S-receptor-like serine/threonine-protein kinase CES101 isoform X1
S-locus-specific glycoprotein S13-like
G-type lectin S-receptor-like serine/threonine-protein kinase At4g27290
putative receptor protein kinase ZmPK1
G-type lectin S-receptor-like serine/threonine-protein kinase SD2-5
G-type lectin S-receptor-like serine/threonine-protein kinase At2g19130 isoform X4
G-type lectin S-receptor-like serine/threonine-protein kinase B120
G-type lectin S-receptor-like serine/threonine-protein kinase At4g03230
G-type lectin S-receptor-like serine/threonine-protein kinase At4g27290 isoform X1
PAN domain-containing protein At5g03700-like
receptor-like serine/threonine-protein kinase SD1-8
G-type lectin S-receptor-like serine/threonine-protein kinase At4g03230 isoform X4
G-type lectin S-receptor-like serine/threonine-protein kinase At1g11330
G-type lectin S-receptor-like serine/threonine-protein kinase At4g27290
G-type lectin S-receptor-like serine/threonine-protein kinase At4g03230 isoform X1

putative receptor protein kinase ZmPK1
G-type lectin S-receptor-like serine/threonine-protein kinase SD2-5
G-type lectin S-receptor-like serine/threonine-protein kinase SD2-5
G-type lectin S-receptor-like serine/threonine-protein kinase CES101 isoform X4
G-type lectin S-receptor-like serine/threonine-protein kinase At4g27290 isoform X1
uncharacterized protein LOC107830114
G-type lectin S-receptor-like serine/threonine-protein kinase SD2-5 isoform X2
G-type lectin S-receptor-like serine/threonine-protein kinase At4g27290
G-type lectin S-receptor-like serine/threonine-protein kinase SD2-5
G-type lectin S-receptor-like serine/threonine-protein kinase At4g27290 isoform X2
G-type lectin S-receptor-like serine/threonine-protein kinase At1g11330
putative G-type lectin S-receptor-like serine/threonine-protein kinase At1g61610 isoform X2
G-type lectin S-receptor-like serine/threonine-protein kinase At4g03230 isoform X1
G-type lectin S-receptor-like serine/threonine-protein kinase At4g27290 isoform X2
G-type lectin S-receptor-like serine/threonine-protein kinase At4g27290 isoform X1
G-type lectin S-receptor-like serine/threonine-protein kinase At4g27290 isoform X1
G-type lectin S-receptor-like serine/threonine-protein kinase At4g27290 isoform X3
G-type lectin S-receptor-like serine/threonine-protein kinase At4g27290 isoform X1
G-type lectin S-receptor-like serine/threonine-protein kinase At4g27290 isoform X1
G-type lectin S-receptor-like serine/threonine-protein kinase CES101 isoform X1
G-type lectin S-receptor-like serine/threonine-protein kinase At4g27290
G-type lectin S-receptor-like serine/threonine-protein kinase At4g03230 isoform X2
G-type lectin S-receptor-like serine/threonine-protein kinase SD1-1
G-type lectin S-receptor-like serine/threonine-protein kinase RLK1
G-type lectin S-receptor-like serine/threonine-protein kinase At4g27290 isoform X2
G-type lectin S-receptor-like serine/threonine-protein kinase At4g27290 isoform X1
receptor-like serine/threonine-protein kinase SD1-7
G-type lectin S-receptor-like serine/threonine-protein kinase At4g27290
receptor-like serine/threonine-protein kinase SD1-8 isoform X2
G-type lectin S-receptor-like serine/threonine-protein kinase CES101 isoform X2
G-type lectin S-receptor-like serine/threonine-protein kinase At4g27290 isoform X2
G-type lectin S-receptor-like serine/threonine-protein kinase At1g34300
G-type lectin S-receptor-like serine/threonine-protein kinase CES101 isoform X2
G-type lectin S-receptor-like serine/threonine-protein kinase At1g11330
G-type lectin S-receptor-like serine/threonine-protein kinase RKS1
G-type lectin S-receptor-like serine/threonine-protein kinase At4g27290
G-type lectin S-receptor-like serine/threonine-protein kinase At4g27290 isoform X1
G-type lectin S-receptor-like serine/threonine-protein kinase RKS1
G-type lectin S-receptor-like serine/threonine-protein kinase At4g27290 isoform X1
G-type lectin S-receptor-like serine/threonine-protein kinase CES101 isoform X1
G-type lectin S-receptor-like serine/threonine-protein kinase At4g03230 isoform X1

G-type lectin S-receptor-like serine/threonine-protein kinase At2g19130
G-type lectin S-receptor-like serine/threonine-protein kinase At1g11300
G-type lectin S-receptor-like serine/threonine-protein kinase At4g03230 isoform X3
uncharacterized protein LOC107830112
G-type lectin S-receptor-like serine/threonine-protein kinase At4g03230 isoform X2
G-type lectin S-receptor-like serine/threonine-protein kinase At4g27290 isoform X1
G-type lectin S-receptor-like serine/threonine-protein kinase At1g11330 isoform X3
G-type lectin S-receptor-like serine/threonine-protein kinase At2g19130
G-type lectin S-receptor-like serine/threonine-protein kinase At4g03230 isoform X1
receptor-like serine/threonine-protein kinase SD1-8
G-type lectin S-receptor-like serine/threonine-protein kinase At1g61500
G-type lectin S-receptor-like serine/threonine-protein kinase At1g34300
G-type lectin S-receptor-like serine/threonine-protein kinase RLK1
G-type lectin S-receptor-like serine/threonine-protein kinase At4g27290 isoform X1
G-type lectin S-receptor-like serine/threonine-protein kinase At4g03230
G-type lectin S-receptor-like serine/threonine-protein kinase SD1-29
G-type lectin S-receptor-like serine/threonine-protein kinase B120
G-type lectin S-receptor-like serine/threonine-protein kinase At4g27290 isoform X3
S locus glycoprotein like protein
G-type lectin S-receptor-like serine/threonine-protein kinase At1g11330 isoform X1
G-type lectin S-receptor-like serine/threonine-protein kinase At4g03230 isoform X2
G-type lectin S-receptor-like serine/threonine-protein kinase At4g27290
G-type lectin S-receptor-like serine/threonine-protein kinase At1g11410
G-type lectin S-receptor-like serine/threonine-protein kinase At1g11330
G-type lectin S-receptor-like serine/threonine-protein kinase SD2-5
G-type lectin S-receptor-like serine/threonine-protein kinase At4g27290 isoform X2
G-type lectin S-receptor-like serine/threonine-protein kinase At4g27290 isoform X2
G-type lectin S-receptor-like serine/threonine-protein kinase At1g61550
receptor-like serine/threonine-protein kinase SD1-8 isoform X3
G-type lectin S-receptor-like serine/threonine-protein kinase At4g27290 isoform X2
receptor-like serine/threonine-protein kinase SD1-8
G-type lectin S-receptor-like serine/threonine-protein kinase B120
G-type lectin S-receptor-like serine/threonine-protein kinase At4g27290
G-type lectin S-receptor-like serine/threonine-protein kinase At4g27290 isoform X2
G-type lectin S-receptor-like serine/threonine-protein kinase At4g27290
G-type lectin S-receptor-like serine/threonine-protein kinase At4g27290 isoform X2
G-type lectin S-receptor-like serine/threonine-protein kinase At4g27290
G-type lectin S-receptor-like serine/threonine-protein kinase At4g27290 isoform X1
uncharacterized protein LOC107822425
uncharacterized protein LOC107772540
G-type lectin S-receptor-like serine/threonine-protein kinase At4g27290 isoform X2

receptor-like serine/threonine-protein kinase SD1-8
G-type lectin S-receptor-like serine/threonine-protein kinase At1g11330 isoform X2
G-type lectin S-receptor-like serine/threonine-protein kinase At4g27290
receptor-like serine/threonine-protein kinase SD1-8
G-type lectin S-receptor-like serine/threonine-protein kinase B120
receptor-like serine/threonine-protein kinase SD1-8 isoform X3
G-type lectin S-receptor-like serine/threonine-protein kinase At4g03230 isoform X1
G-type lectin S-receptor-like serine/threonine-protein kinase At4g27290
G-type lectin S-receptor-like serine/threonine-protein kinase At4g27290
G-type lectin S-receptor-like serine/threonine-protein kinase SD1-29
G-type lectin S-receptor-like serine/threonine-protein kinase At4g27290 isoform X2
G-type lectin S-receptor-like serine/threonine-protein kinase SD2-2
G-type lectin S-receptor-like serine/threonine-protein kinase RKS1
receptor-like serine/threonine-protein kinase SD1-8 isoform X1
G-type lectin S-receptor-like serine/threonine-protein kinase At1g11330
G-type lectin S-receptor-like serine/threonine-protein kinase At4g27290
G-type lectin S-receptor-like serine/threonine-protein kinase CES101 isoform X2
S-domain receptor-like kinase
G-type lectin S-receptor-like serine/threonine-protein kinase At4g03230 isoform X2
G-type lectin S-receptor-like serine/threonine-protein kinase At4g03230 isoform X2
G-type lectin S-receptor-like serine/threonine-protein kinase At4g03230 isoform X2
G-type lectin S-receptor-like serine/threonine-protein kinase At4g27290
G-type lectin S-receptor-like serine/threonine-protein kinase SD2-2
G-type lectin S-receptor-like serine/threonine-protein kinase At4g27290
G-type lectin S-receptor-like serine/threonine-protein kinase At4g03230 isoform X1
receptor-like serine/threonine-protein kinase SD1-8 isoform X3
G-type lectin S-receptor-like serine/threonine-protein kinase At4g03230 isoform X3
receptor-like serine/threonine-protein kinase SD1-8 isoform X2
G-type lectin S-receptor-like serine/threonine-protein kinase SD3-1
putative G-type lectin S-receptor-like serine/threonine-protein kinase At1g61610 isoform X1
G-type lectin S-receptor-like serine/threonine-protein kinase At4g27290 isoform X1
G-type lectin S-receptor-like serine/threonine-protein kinase SD3-1
G-type lectin S-receptor-like serine/threonine-protein kinase At4g03230 isoform X1
G-type lectin S-receptor-like serine/threonine-protein kinase SD2-5
Uncharacterized protein
PAN/APPLE-like domain protein
PAN domain-containing protein
Coagulation factor XI (Fragment)
Hepatocyte growth factor (Fragment)
Plasminogen (Fragment)
Plasminogen (Fragment)

Uncharacterized protein
Uncharacterized protein
Uncharacterized protein
Uncharacterized protein
Uncharacterized protein
Uncharacterized protein
Uncharacterized protein
Uncharacterized protein
Uncharacterized protein
Uncharacterized protein
Uncharacterized protein
Uncharacterized protein
Uncharacterized protein
Uncharacterized protein
Uncharacterized protein
Uncharacterized protein
Uncharacterized protein
Uncharacterized protein
Uncharacterized protein
Uncharacterized protein
Uncharacterized protein
Uncharacterized protein
Uncharacterized protein
Uncharacterized protein
Uncharacterized protein
Uncharacterized protein
Uncharacterized protein
Uncharacterized protein
Uncharacterized protein
Uncharacterized protein
Uncharacterized protein
Uncharacterized protein
Uncharacterized protein
Uncharacterized protein
Uncharacterized protein
Uncharacterized protein
Uncharacterized protein
Uncharacterized protein
Uncharacterized protein
Alpha-2-macroglobulin-like protein
Alpha-2-macroglobulin-like protein

Uncharacterized protein
Uncharacterized protein
Uncharacterized protein
Uncharacterized protein
Apple domain-containing protein
Apple domain-containing protein
Uncharacterized protein
Apple domain-containing protein
Uncharacterized protein
PAN domain-containing protein
PAN domain-containing protein
PAN domain-containing protein
Uncharacterized protein
Uncharacterized protein
Receptor-like serine/threonine-protein kinase SD1-8
PAN domain-containing protein
G-type lectin S-receptor-like serine/threonine-protein kinase SD3-1
G-type lectin S-receptor-like serine/threonine-protein kinase
G-type lectin S-receptor-like serine/threonine-protein kinase (Fragment)
G-type lectin S-receptor-like serine/threonine-protein kinase SD3-1
G-type lectin S-receptor-like serine/threonine-protein kinase (Fragment)
G-type lectin S-receptor-like serine/threonine-protein kinase RKS1
Uncharacterized protein (Fragment)
G-type lectin S-receptor-like serine/threonine-protein kinase
G-type lectin S-receptor-like serine/threonine-protein kinase (Fragment)
G-type lectin S-receptor-like serine/threonine-protein kinase (Fragment)
G-type lectin S-receptor-like serine/threonine-protein kinase (Fragment)
G-type lectin S-receptor-like serine/threonine-protein kinase (Fragment)
G-type lectin S-receptor-like serine/threonine-protein kinase
Uncharacterized protein (Fragment)
Epidermis-specific secreted glycoprotein EP1 (Fragment)
Uncharacterized protein (Fragment)
G-type lectin S-receptor-like serine/threonine-protein kinase (Fragment)
G-type lectin S-receptor-like serine/threonine-protein kinase (Fragment)
G-type lectin S-receptor-like serine/threonine-protein kinase
G-type lectin S-receptor-like serine/threonine-protein kinase RLK1
G-type lectin S-receptor-like serine/threonine-protein kinase SD1-29 (Fragment)
G-type lectin S-receptor-like serine/threonine-protein kinase (Fragment)
G-type lectin S-receptor-like serine/threonine-protein kinase
G-type lectin S-receptor-like serine/threonine-protein kinase (Fragment)
Uncharacterized protein (Fragment)

G-type lectin S-receptor-like serine/threonine-protein kinase (Fragment)
G-type lectin S-receptor-like serine/threonine-protein kinase (Fragment)
G-type lectin S-receptor-like serine/threonine-protein kinase SD1-29 (Fragment)
G-type lectin S-receptor-like serine/threonine-protein kinase SD1-29
Uncharacterized protein (Fragment)
Putative G-type lectin S-receptor-like serine/threonine-protein kinase (Fragment)
G-type lectin S-receptor-like serine/threonine-protein kinase SD2-2 (Fragment)
Uncharacterized protein (Fragment)
G-type lectin S-receptor-like serine/threonine-protein kinase SD1-29 (Fragment)
G-type lectin S-receptor-like serine/threonine-protein kinase
G-type lectin S-receptor-like serine/threonine-protein kinase (Fragment)
G-type lectin S-receptor-like serine/threonine-protein kinase (Fragment)
G-type lectin S-receptor-like serine/threonine-protein kinase (Fragment)
Putative receptor protein kinase ZmPK1 (Fragment)
G-type lectin S-receptor-like serine/threonine-protein kinase (Fragment)
G-type lectin S-receptor-like serine/threonine-protein kinase SD1-29 (Fragment)
Uncharacterized protein (Fragment)
G-type lectin S-receptor-like serine/threonine-protein kinase SD2-2
G-type lectin S-receptor-like serine/threonine-protein kinase (Fragment)
Uncharacterized protein (Fragment)
G-type lectin S-receptor-like serine/threonine-protein kinase (Fragment)
G-type lectin S-receptor-like serine/threonine-protein kinase
G-type lectin S-receptor-like serine/threonine-protein kinase RLK1
G-type lectin S-receptor-like serine/threonine-protein kinase
G-type lectin S-receptor-like serine/threonine-protein kinase
G-type lectin S-receptor-like serine/threonine-protein kinase
G-type lectin S-receptor-like serine/threonine-protein kinase
G-type lectin S-receptor-like serine/threonine-protein kinase
PAN domain-containing protein
G-type lectin S-receptor-like serine/threonine-protein kinase SD3-1
G-type lectin S-receptor-like serine/threonine-protein kinase (Fragment)
G-type lectin S-receptor-like serine/threonine-protein kinase (Fragment)
G-type lectin S-receptor-like serine/threonine-protein kinase (Fragment)
Putative G-type lectin S-receptor-like serine/threonine-protein kinase (Fragment)
G-type lectin S-receptor-like serine/threonine-protein kinase (Fragment)
G-type lectin S-receptor-like serine/threonine-protein kinase (Fragment)
G-type lectin S-receptor-like serine/threonine-protein kinase SD1-29 (Fragment)
Epidermis-specific secreted glycoprotein EP1
G-type lectin S-receptor-like serine/threonine-protein kinase (Fragment)
Receptor-like serine/threonine-protein kinase SD1-8
G-type lectin S-receptor-like serine/threonine-protein kinase

G-type lectin S-receptor-like serine/threonine-protein kinase
PAN domain-containing protein (Fragment)
G-type lectin S-receptor-like serine/threonine-protein kinase
G-type lectin S-receptor-like serine/threonine-protein kinase (Fragment)
G-type lectin S-receptor-like serine/threonine-protein kinase (Fragment)
G-type lectin S-receptor-like serine/threonine-protein kinase (Fragment)
G-type lectin S-receptor-like serine/threonine-protein kinase (Fragment)
G-type lectin S-receptor-like serine/threonine-protein kinase SD1-29 (Fragment)
G-type lectin S-receptor-like serine/threonine-protein kinase
G-type lectin S-receptor-like serine/threonine-protein kinase RLK1
G-type lectin S-receptor-like serine/threonine-protein kinase
G-type lectin S-receptor-like serine/threonine-protein kinase RKS1
G-type lectin S-receptor-like serine/threonine-protein kinase
Receptor-like serine/threonine-protein kinase SD1-8
G-type lectin S-receptor-like serine/threonine-protein kinase
G-type lectin S-receptor-like serine/threonine-protein kinase (Fragment)
Putative G-type lectin S-receptor-like serine/threonine-protein kinase (Fragment)
G-type lectin S-receptor-like serine/threonine-protein kinase SD1-29 (Fragment)
G-type lectin S-receptor-like serine/threonine-protein kinase SD1-29 (Fragment)
Uncharacterized protein (Fragment)
G-type lectin S-receptor-like serine/threonine-protein kinase (Fragment)
G-type lectin S-receptor-like serine/threonine-protein kinase (Fragment)
G-type lectin S-receptor-like serine/threonine-protein kinase (Fragment)
G-type lectin S-receptor-like serine/threonine-protein kinase SD2-2
Uncharacterized protein (Fragment)
Uncharacterized protein (Fragment)
G-type lectin S-receptor-like serine/threonine-protein kinase (Fragment)
G-type lectin S-receptor-like serine/threonine-protein kinase SD1-29
G-type lectin S-receptor-like serine/threonine-protein kinase (Fragment)
G-type lectin S-receptor-like serine/threonine-protein kinase (Fragment)
G-type lectin S-receptor-like serine/threonine-protein kinase
Uncharacterized protein
G-type lectin S-receptor-like serine/threonine-protein kinase (Fragment)
G-type lectin S-receptor-like serine/threonine-protein kinase
G-type lectin S-receptor-like serine/threonine-protein kinase SD1-29 (Fragment)
G-type lectin S-receptor-like serine/threonine-protein kinase (Fragment)
G-type lectin S-receptor-like serine/threonine-protein kinase RKS1 (Fragment)
G-type lectin S-receptor-like serine/threonine-protein kinase RLK1
G-type lectin S-receptor-like serine/threonine-protein kinase (Fragment)
G-type lectin S-receptor-like serine/threonine-protein kinase RKS1 (Fragment)
G-type lectin S-receptor-like serine/threonine-protein kinase (Fragment)

PAN domain-containing protein (Fragment)
G-type lectin S-receptor-like serine/threonine-protein kinase (Fragment)
G-type lectin S-receptor-like serine/threonine-protein kinase SD2-2
G-type lectin S-receptor-like serine/threonine-protein kinase (Fragment)
Uncharacterized protein (Fragment)
G-type lectin S-receptor-like serine/threonine-protein kinase (Fragment)
G-type lectin S-receptor-like serine/threonine-protein kinase (Fragment)
PAN domain-containing protein (Fragment)
G-type lectin S-receptor-like serine/threonine-protein kinase (Fragment)
Putative G-type lectin S-receptor-like serine/threonine-protein kinase (Fragment)
G-type lectin S-receptor-like serine/threonine-protein kinase
G-type lectin S-receptor-like serine/threonine-protein kinase (Fragment)
G-type lectin S-receptor-like serine/threonine-protein kinase (Fragment)
G-type lectin S-receptor-like serine/threonine-protein kinase RKS1
Uncharacterized protein (Fragment)
Uncharacterized protein
Uncharacterized protein
Hepatocyte growth factor
Uncharacterized protein
Uncharacterized protein
Hepatocyte growth factor
Uncharacterized protein
Uncharacterized protein
Uncharacterized protein
Apple domain:N/apple PAN
Uncharacterized protein
Alpha-2-macroglobulin
Uncharacterized protein (Fragment)
Plasminogen (Fragment)
Uncharacterized protein (Fragment)
Uncharacterized protein
Hepatocyte growth factor
Uncharacterized protein
Uncharacterized protein
Hepatocyte growth factor (Hepapoietin A, scatter factor) (Fragment)
Uncharacterized protein (Fragment)
Hepatocyte growth factor (Hepapoietin A, scatter factor)
Uncharacterized protein
Uncharacterized protein (Fragment)
Uncharacterized protein
Uncharacterized protein (Fragment)

Hepatocyte growth factor
Uncharacterized protein
Uncharacterized protein
PAN domain-containing protein
Uncharacterized protein
Uncharacterized protein
Uncharacterized protein
Uncharacterized protein
PAN domain-containing protein
Alpha-2-macroglobulin N-terminal domain family protein
Alpha-2-macroglobulin domain protein
Membrane protein
Uncharacterized protein
Uncharacterized protein
Uncharacterized protein
Alpha-2-macroglobulin family protein
Uncharacterized protein
PAN domain protein
Uncharacterized protein
Alpha-2-macroglobulin domain-containing protein
Alpha-2-macroglobulin domain-containing protein
Alpha-2-macroglobulin family protein
Uncharacterized protein
Alpha-2-macroglobulin family protein
Uncharacterized protein
Alpha-2-macroglobulin family protein
Uncharacterized protein
Uncharacterized protein
Uncharacterized protein
PAN domain protein
Alpha-2-macroglobulin family protein
Uncharacterized protein
Membrane protein
Uncharacterized protein
Uncharacterized protein
Uncharacterized protein
Uncharacterized protein
Uncharacterized protein
Uncharacterized protein
Alpha-2-macroglobulin domain-containing protein
Alpha-2-macroglobulin family protein

Uncharacterized protein
Uncharacterized protein (Fragment)
Uncharacterized protein (Fragment)
Uncharacterized protein
Uncharacterized protein
Uncharacterized protein
Uncharacterized protein
Uncharacterized protein
Uncharacterized protein (Fragment)
Uncharacterized protein
Uncharacterized protein
Uncharacterized protein
Uncharacterized protein
Uncharacterized protein (Fragment)
Uncharacterized protein (Fragment)
Uncharacterized protein
Uncharacterized protein
Uncharacterized protein
Uncharacterized protein
Uncharacterized protein
Uncharacterized protein
Uncharacterized protein (Fragment)
Uncharacterized protein (Fragment)
Uncharacterized protein
Uncharacterized protein
Uncharacterized protein
Uncharacterized protein (Fragment)
Uncharacterized protein
Uncharacterized protein
Uncharacterized protein
Uncharacterized protein
Uncharacterized protein
Uncharacterized protein
Uncharacterized protein
Uncharacterized protein (Fragment)
Uncharacterized protein (Fragment)
Uncharacterized protein
Uncharacterized protein
Uncharacterized protein (Fragment)
Uncharacterized protein (Fragment)
Uncharacterized protein (Fragment)

Uncharacterized protein
Uncharacterized protein
Uncharacterized protein (Fragment)
Uncharacterized protein
Uncharacterized protein
Uncharacterized protein
Uncharacterized protein (Fragment)
Uncharacterized protein (Fragment)
Hepatocyte growth factor
Hepatocyte growth factor
hepatocyte growth factor-like protein
coagulation factor XI isoform X1
plasminogen isoform X3
Plasminogen
Plasminogen
plasma kallikrein
coagulation factor XI isoform X2
coagulation factor XI isoform X3
PAN domain protein
Uncharacterized protein
PAN domain protein
PAN domain protein
PAN domain protein
PAN domain protein
PAN domain protein (Fragment)
PAN domain protein
PAN domain protein
PAN domain protein
PAN domain protein
PAN domain protein
PAN domain protein
PAN domain protein
PAN domain protein
PAN domain protein
PAN domain protein
PAN domain protein
PAN domain protein
PAN domain protein
PAN domain protein
PAN domain protein
PAN domain protein (Fragment)

PAN domain protein (Fragment)
Glycosyltransferase family 31 protein
Uncharacterized protein
Uncharacterized protein
Uncharacterized protein
Uncharacterized protein
Uncharacterized protein
Uncharacterized protein
Uncharacterized protein
Uncharacterized protein
Uncharacterized protein
Uncharacterized protein
Uncharacterized protein
Uncharacterized protein
Uncharacterized protein
Uncharacterized protein
Uncharacterized protein
Uncharacterized protein
Uncharacterized protein
Uncharacterized protein
S-locus receptor kinase (Fragment)
S-locus glycoprotein (Fragment)
Uncharacterized protein
PAN domain protein
Uncharacterized protein
Uncharacterized protein
Uncharacterized protein
Uncharacterized protein
Uncharacterized protein
Uncharacterized protein
Uncharacterized protein
Uncharacterized protein
Uncharacterized protein
Uncharacterized protein
Uncharacterized protein
Uncharacterized protein
Uncharacterized protein
Uncharacterized protein
PAN domain protein
Uncharacterized protein
PAN domain protein

PAN domain protein
Uncharacterized protein (Fragment)
Uncharacterized protein
Uncharacterized protein
PAN domain protein
Uncharacterized protein
Uncharacterized protein (Fragment)
Uncharacterized protein (Fragment)
Uncharacterized protein
Uncharacterized protein (Fragment)
Uncharacterized protein
Uncharacterized protein
Uncharacterized protein
Uncharacterized protein (Fragment)
Uncharacterized protein (Fragment)
Uncharacterized protein
Uncharacterized protein
Uncharacterized protein
Uncharacterized protein
Uncharacterized protein
Uncharacterized protein
Uncharacterized protein
Cation-transporting ATPase
Uncharacterized protein
Uncharacterized protein
Uncharacterized protein
Uncharacterized protein
Uncharacterized protein
Uncharacterized protein
Uncharacterized protein
Uncharacterized protein
Uncharacterized protein
Uncharacterized protein
Uncharacterized protein
Uncharacterized protein
Uncharacterized protein
Uncharacterized protein
Uncharacterized protein
Uncharacterized protein
Uncharacterized protein
Uncharacterized protein (Fragment)

Uncharacterized protein (Fragment)
Uncharacterized protein
Uncharacterized protein
Uncharacterized protein
Uncharacterized protein
Trynity
Uncharacterized protein
Uncharacterized protein
Uncharacterized protein
Uncharacterized protein
Uncharacterized protein
Uncharacterized protein
Plasminogen
Uncharacterized protein
Uncharacterized protein
Uncharacterized protein
Cuticlin-1
Neurotrypsin
Uncharacterized protein
Uncharacterized protein
Uncharacterized protein
Uncharacterized protein
Uncharacterized protein
Uncharacterized protein
Uncharacterized protein
Uncharacterized protein
Uncharacterized protein
Uncharacterized protein
PAN/APPLE-like domain-containing protein
Uncharacterized protein
Uncharacterized protein (Fragment)
Uncharacterized protein
Ras-related protein RabJ
Cuticlin-1 (Fragment)
PAN domain containing protein
PAN domain containing protein
Uncharacterized protein
Uncharacterized protein
Plasma kallikrein (Fragment)
Hepatocyte growth factor-like protein (Fragment)
Tryptophan synthase alpha chain

Uncharacterized protein
Plasminogen (Fragment)
Hepatocyte growth factor (Fragment)
Coagulation factor XI (Fragment)
Plasminogen (Fragment)
PAN domain protein
Uncharacterized protein
Uncharacterized protein
PAN domain protein
PAN domain protein (Fragment)
Uncharacterized protein
Uncharacterized protein
PAN domain protein
Uncharacterized protein (Fragment)
Uncharacterized protein
Protein lifeguard 1
Cuticlin-1
EGF-like domain-containing protein 2 (Fragment)
Uncharacterized protein
Cuticlin-1
Uncharacterized protein
Plasminogen
Uncharacterized protein
Plasminogen
Hepatocyte growth factor
Uncharacterized protein
Uncharacterized protein
Uncharacterized protein
Plasminogen
Macrophage stimulating 1
Uncharacterized protein
Glycosyl transferase family 1
Uncharacterized protein (Fragment)
Uncharacterized protein (Fragment)
Uncharacterized protein
Uncharacterized protein
Hepatocyte growth factor
Uncharacterized protein
Hepatocyte growth factor
SLR1 (Fragment)
Uncharacterized protein (Fragment)

Uncharacterized protein
Carbohydrate-binding protein (Fragment)
Factor XI
Hepatocyte growth factor 2 protein
Uncharacterized protein
Uncharacterized protein
Uncharacterized protein
Uncharacterized protein
Hepatocyte growth factor
Hepatocyte growth factor
Hepatocyte growth factor
S-domain receptor-like protein kinase
S-domain receptor-like protein kinase
S-domain receptor-like protein kinase
Serine/threonine-protein kinase
Uncharacterized protein
Uncharacterized protein
Serine/threonine-protein kinase
Uncharacterized protein
Serine/threonine-protein kinase
Serine/threonine-protein kinase
Serine/threonine-protein kinase
Serine/threonine-protein kinase
Uncharacterized protein
Serine/threonine-protein kinase
Uncharacterized protein
Uncharacterized protein
Serine/threonine-protein kinase
Uncharacterized protein
Uncharacterized protein
Serine/threonine-protein kinase
Serine/threonine-protein kinase
Uncharacterized protein
Uncharacterized protein
Uncharacterized protein
Serine/threonine-protein kinase
Uncharacterized protein
Uncharacterized protein
Serine/threonine-protein kinase
Serine/threonine-protein kinase
Uncharacterized protein

Uncharacterized protein
Uncharacterized protein
Serine/threonine-protein kinase
Uncharacterized protein
Uncharacterized protein
Uncharacterized protein
Serine/threonine-protein kinase
Uncharacterized protein
Serine/threonine-protein kinase
Uncharacterized protein
Uncharacterized protein
Serine/threonine-protein kinase
Serine/threonine-protein kinase
Serine/threonine-protein kinase
Serine/threonine-protein kinase
Uncharacterized protein
Serine/threonine-protein kinase
Serine/threonine-protein kinase
Uncharacterized protein
Uncharacterized protein
Serine/threonine-protein kinase
Uncharacterized protein
Serine/threonine-protein kinase
Serine/threonine-protein kinase
Uncharacterized protein
Uncharacterized protein
Serine/threonine-protein kinase
Serine/threonine-protein kinase
Serine/threonine-protein kinase
Uncharacterized protein
Uncharacterized protein
Serine/threonine-protein kinase
Serine/threonine-protein kinase
Serine/threonine-protein kinase
Serine/threonine-protein kinase
Serine/threonine-protein kinase
Uncharacterized protein
Serine/threonine-protein kinase
Serine/threonine-protein kinase
Uncharacterized protein
Serine/threonine-protein kinase

Uncharacterized protein
Serine/threonine-protein kinase
Serine/threonine-protein kinase
Uncharacterized protein
Uncharacterized protein
Uncharacterized protein
Serine/threonine-protein kinase
Uncharacterized protein
Uncharacterized protein
Serine/threonine-protein kinase
Uncharacterized protein
Uncharacterized protein
Uncharacterized protein
Uncharacterized protein
Uncharacterized protein
Serine/threonine-protein kinase
Uncharacterized protein
Serine/threonine-protein kinase
Serine/threonine-protein kinase
Serine/threonine-protein kinase
Serine/threonine-protein kinase
Uncharacterized protein
Serine/threonine-protein kinase
Uncharacterized protein
Uncharacterized protein
Uncharacterized protein
Uncharacterized protein
Serine/threonine-protein kinase
Serine/threonine-protein kinase
Serine/threonine-protein kinase
Uncharacterized protein
Serine/threonine-protein kinase
Uncharacterized protein
Serine/threonine-protein kinase
Uncharacterized protein
Uncharacterized protein
Serine/threonine-protein kinase
Uncharacterized protein
Uncharacterized protein
Uncharacterized protein
Serine/threonine-protein kinase

Uncharacterized protein
Uncharacterized protein
Serine/threonine-protein kinase
Serine/threonine-protein kinase
Uncharacterized protein
Serine/threonine-protein kinase
Uncharacterized protein
Serine/threonine-protein kinase
S-domain receptor-like protein kinase
Uncharacterized protein
Serine/threonine-protein kinase
Serine/threonine-protein kinase
Serine/threonine-protein kinase
S-domain receptor-like protein kinase
Serine/threonine-protein kinase
Hypothetical_protein
Serine/threonine-protein kinase
Serine/threonine-protein kinase
Serine/threonine-protein kinase
Uncharacterized protein
Uncharacterized protein
Serine/threonine-protein kinase
Serine/threonine-protein kinase
Uncharacterized protein
Uncharacterized protein
Serine/threonine-protein kinase
Serine/threonine-protein kinase
Serine/threonine-protein kinase
Serine/threonine-protein kinase
Uncharacterized protein
Serine/threonine-protein kinase
Uncharacterized protein
Serine/threonine-protein kinase
Serine/threonine-protein kinase
Serine/threonine-protein kinase
Uncharacterized protein
Uncharacterized protein
Uncharacterized protein
Serine/threonine-protein kinase
Serine/threonine-protein kinase
Serine/threonine-protein kinase

Uncharacterized protein
Serine/threonine-protein kinase
Serine/threonine-protein kinase
Hypothetical_protein
Serine/threonine-protein kinase
Serine/threonine-protein kinase
Serine/threonine-protein kinase
Uncharacterized protein
Serine/threonine-protein kinase
Serine/threonine-protein kinase
Uncharacterized protein
Uncharacterized protein
Serine/threonine-protein kinase
Serine/threonine-protein kinase
Uncharacterized protein
Serine/threonine-protein kinase
Serine/threonine-protein kinase
Uncharacterized protein
Serine/threonine-protein kinase
Uncharacterized protein
Serine/threonine-protein kinase
Serine/threonine-protein kinase
Serine/threonine-protein kinase
Serine/threonine-protein kinase
Serine/threonine-protein kinase
Serine/threonine-protein kinase
Serine/threonine-protein kinase
Serine/threonine-protein kinase
Serine/threonine-protein kinase
Serine/threonine-protein kinase
Uncharacterized protein
Uncharacterized protein
Serine/threonine-protein kinase
Serine/threonine-protein kinase
Serine/threonine-protein kinase
Serine/threonine-protein kinase
Serine/threonine-protein kinase
Serine/threonine-protein kinase
Serine/threonine-protein kinase
Serine/threonine-protein kinase
Serine/threonine-protein kinase

Serine/threonine-protein kinase
Serine/threonine-protein kinase
Serine/threonine-protein kinase
Uncharacterized protein
Uncharacterized protein
Serine/threonine-protein kinase
Uncharacterized protein
Serine/threonine-protein kinase
Serine/threonine-protein kinase
Serine/threonine-protein kinase
Hypothetical_protein
Uncharacterized protein
Serine/threonine-protein kinase
Uncharacterized protein
Hypothetical_protein
Hypothetical_protein
Uncharacterized protein
Uncharacterized protein
Uncharacterized protein
Serine/threonine-protein kinase
Hypothetical_protein
Serine/threonine-protein kinase
Serine/threonine-protein kinase
Serine/threonine-protein kinase
Serine/threonine-protein kinase
Uncharacterized protein
Serine/threonine-protein kinase
Uncharacterized protein
Serine/threonine-protein kinase
Serine/threonine-protein kinase
Uncharacterized protein
Serine/threonine-protein kinase
Serine/threonine-protein kinase
Uncharacterized protein
Uncharacterized protein
Serine/threonine-protein kinase
Hypothetical_protein
Uncharacterized protein
Uncharacterized protein
Uncharacterized protein
Serine/threonine-protein kinase

Serine/threonine-protein kinase
Serine/threonine-protein kinase
Serine/threonine-protein kinase
Uncharacterized protein
Serine/threonine-protein kinase
Serine/threonine-protein kinase
Uncharacterized protein
Uncharacterized protein
Serine/threonine-protein kinase
Uncharacterized protein
Uncharacterized protein
Uncharacterized protein
Serine/threonine-protein kinase
Uncharacterized protein
Serine/threonine-protein kinase
Serine/threonine-protein kinase
Serine/threonine-protein kinase
Serine/threonine-protein kinase
Uncharacterized protein
Uncharacterized protein
Serine/threonine-protein kinase
Serine/threonine-protein kinase
Serine/threonine-protein kinase
Serine/threonine-protein kinase
Serine/threonine-protein kinase
Serine/threonine-protein kinase
Serine/threonine-protein kinase
Serine/threonine-protein kinase
Serine/threonine-protein kinase
Uncharacterized protein
Uncharacterized protein
Serine/threonine-protein kinase
Hypothetical_protein
Serine/threonine-protein kinase
Serine/threonine-protein kinase
Serine/threonine-protein kinase
Serine/threonine-protein kinase
Uncharacterized protein
Serine/threonine-protein kinase
Serine/threonine-protein kinase
Uncharacterized protein

Serine/threonine-protein kinase
Serine/threonine-protein kinase
Serine/threonine-protein kinase
Serine/threonine-protein kinase
Serine/threonine-protein kinase
Serine/threonine-protein kinase
Serine/threonine-protein kinase
Serine/threonine-protein kinase
Uncharacterized protein
Serine/threonine-protein kinase
Uncharacterized protein
Uncharacterized protein
Uncharacterized protein
Serine/threonine-protein kinase
Uncharacterized protein
Uncharacterized protein
Serine/threonine-protein kinase
Uncharacterized protein
Uncharacterized protein
Serine/threonine-protein kinase
Serine/threonine-protein kinase
Uncharacterized protein
Uncharacterized protein
Serine/threonine-protein kinase
Uncharacterized protein
Serine/threonine-protein kinase
Serine/threonine-protein kinase
Uncharacterized protein
Serine/threonine-protein kinase
Uncharacterized protein
Serine/threonine-protein kinase
Uncharacterized protein
Uncharacterized protein
Uncharacterized protein
Serine/threonine-protein kinase
Serine/threonine-protein kinase
Uncharacterized protein
Uncharacterized protein
Serine/threonine-protein kinase
Serine/threonine-protein kinase
Uncharacterized protein

Uncharacterized protein
Uncharacterized protein
Uncharacterized protein
Uncharacterized protein
Uncharacterized protein
Uncharacterized protein
Uncharacterized protein
Serine/threonine-protein kinase
Uncharacterized protein
Serine/threonine-protein kinase
Uncharacterized protein
Uncharacterized protein
Uncharacterized protein
Serine/threonine-protein kinase
Serine/threonine-protein kinase
Serine/threonine-protein kinase
Serine/threonine-protein kinase
Uncharacterized protein
Serine/threonine-protein kinase
Serine/threonine-protein kinase
Serine/threonine-protein kinase
Uncharacterized protein
Serine/threonine-protein kinase
Uncharacterized protein
Uncharacterized protein
Uncharacterized protein
Uncharacterized protein
Uncharacterized protein
Serine/threonine-protein kinase
Serine/threonine-protein kinase
Uncharacterized protein
Serine/threonine-protein kinase
Uncharacterized protein
Serine/threonine-protein kinase
Serine/threonine-protein kinase
Uncharacterized protein
Serine/threonine-protein kinase
Uncharacterized protein
Uncharacterized protein
Uncharacterized protein
Uncharacterized protein

Uncharacterized protein
Uncharacterized protein
Uncharacterized protein
Uncharacterized protein
Serine/threonine-protein kinase
Serine/threonine-protein kinase
Serine/threonine-protein kinase
Uncharacterized protein
Uncharacterized protein
Serine/threonine-protein kinase
Uncharacterized protein
Serine/threonine-protein kinase
Uncharacterized protein
Serine/threonine-protein kinase
Serine/threonine-protein kinase
Serine/threonine-protein kinase
Serine/threonine-protein kinase
Serine/threonine-protein kinase
Uncharacterized protein
Uncharacterized protein
Uncharacterized protein
Serine/threonine-protein kinase
Serine/threonine-protein kinase
Serine/threonine-protein kinase
Uncharacterized protein
Uncharacterized protein
Uncharacterized protein
Uncharacterized protein
Serine/threonine-protein kinase
Uncharacterized protein
Uncharacterized protein
Serine/threonine-protein kinase
Serine/threonine-protein kinase
Uncharacterized protein
Serine/threonine-protein kinase
Uncharacterized protein
Serine/threonine-protein kinase
Uncharacterized protein
Uncharacterized protein
Uncharacterized protein
Uncharacterized protein

Uncharacterized protein
Uncharacterized protein
Uncharacterized protein
Uncharacterized protein
Uncharacterized protein
Serine/threonine-protein kinase
Serine/threonine-protein kinase
Serine/threonine-protein kinase
Uncharacterized protein
Uncharacterized protein
Uncharacterized protein
Uncharacterized protein
Uncharacterized protein
Uncharacterized protein
Uncharacterized protein
Uncharacterized protein
Uncharacterized protein
Uncharacterized protein
Uncharacterized protein
Uncharacterized protein
Uncharacterized protein
Uncharacterized protein
Uncharacterized protein
Uncharacterized protein
Uncharacterized protein
Uncharacterized protein
Uncharacterized protein
Uncharacterized protein
Uncharacterized protein
Uncharacterized protein
Uncharacterized protein
Uncharacterized protein
Uncharacterized protein
Uncharacterized protein
Uncharacterized protein
Uncharacterized protein
Uncharacterized protein
Uncharacterized protein
Uncharacterized protein
Uncharacterized protein
Uncharacterized protein

Uncharacterized protein
Uncharacterized protein
Uncharacterized protein
Uncharacterized protein
Uncharacterized protein
Uncharacterized protein
Uncharacterized protein
Uncharacterized protein
Uncharacterized protein
Uncharacterized protein
Uncharacterized protein
Uncharacterized protein
Uncharacterized protein
Uncharacterized protein
Uncharacterized protein
Uncharacterized protein
Uncharacterized protein
Uncharacterized protein
Uncharacterized protein
Uncharacterized protein
Uncharacterized protein
Uncharacterized protein
Uncharacterized protein
Uncharacterized protein
Uncharacterized protein
Uncharacterized protein
Uncharacterized protein
Uncharacterized protein
Uncharacterized protein
Uncharacterized protein
Uncharacterized protein
Uncharacterized protein
Uncharacterized protein
Uncharacterized protein
Uncharacterized protein
Uncharacterized protein
Uncharacterized protein
Uncharacterized protein
Uncharacterized protein
Uncharacterized protein
Uncharacterized protein

Uncharacterized protein
Uncharacterized protein
Uncharacterized protein
Uncharacterized protein
Uncharacterized protein
Uncharacterized protein
Uncharacterized protein
Uncharacterized protein
Uncharacterized protein
Uncharacterized protein
Uncharacterized protein
Uncharacterized protein
Uncharacterized protein
Uncharacterized protein
Uncharacterized protein
Uncharacterized protein
Uncharacterized protein
Uncharacterized protein
S-domain receptor-like protein kinase
S-domain receptor-like protein kinase
Uncharacterized protein
Uncharacterized protein
Serine/threonine-protein kinase
S-domain receptor-like protein kinase
Serine/threonine-protein kinase
Uncharacterized protein
Serine/threonine-protein kinase
Uncharacterized protein
Serine/threonine-protein kinase
Uncharacterized protein
Serine/threonine-protein kinase
Serine/threonine-protein kinase
Serine/threonine-protein kinase
Serine/threonine-protein kinase
Serine/threonine-protein kinase
Serine/threonine-protein kinase
Serine/threonine-protein kinase
Uncharacterized protein
Serine/threonine-protein kinase
Serine/threonine-protein kinase
Uncharacterized protein

Uncharacterized protein
Uncharacterized protein
Uncharacterized protein
Serine/threonine-protein kinase
Uncharacterized protein
Uncharacterized protein
Serine/threonine-protein kinase
Uncharacterized protein
Serine/threonine-protein kinase
Uncharacterized protein
S-domain receptor-like protein kinase family-3
Serine/threonine-protein kinase
Serine/threonine-protein kinase
Uncharacterized protein
Uncharacterized protein
Serine/threonine-protein kinase
Serine/threonine-protein kinase
Uncharacterized protein
Serine/threonine-protein kinase
Serine/threonine-protein kinase
Serine/threonine-protein kinase
Uncharacterized protein
Serine/threonine-protein kinase
Uncharacterized protein
Serine/threonine-protein kinase
Uncharacterized protein
Uncharacterized protein
Uncharacterized protein
Uncharacterized protein
Serine/threonine-protein kinase
Serine/threonine-protein kinase
Serine/threonine-protein kinase
Serine/threonine-protein kinase
Serine/threonine-protein kinase
Uncharacterized protein
Serine/threonine-protein kinase
Serine/threonine-protein kinase
Uncharacterized protein
Serine/threonine-protein kinase
Uncharacterized protein
Uncharacterized protein

Uncharacterized protein
Uncharacterized protein
Serine/threonine-protein kinase
Uncharacterized protein
Uncharacterized protein
Serine/threonine-protein kinase
Serine/threonine-protein kinase
Serine/threonine-protein kinase
Serine/threonine-protein kinase
Uncharacterized protein
Serine/threonine-protein kinase
Serine/threonine-protein kinase
Serine/threonine-protein kinase
Serine/threonine-protein kinase
Uncharacterized protein
Uncharacterized protein
Serine/threonine-protein kinase
Uncharacterized protein
Serine/threonine-protein kinase
Serine/threonine-protein kinase
Uncharacterized protein
Serine/threonine-protein kinase
Uncharacterized protein
Uncharacterized protein
Uncharacterized protein
Uncharacterized protein
Uncharacterized protein
Serine/threonine-protein kinase
Serine/threonine-protein kinase
Serine/threonine-protein kinase
Serine/threonine-protein kinase
Uncharacterized protein
Uncharacterized protein
Serine/threonine-protein kinase
Uncharacterized protein
Serine/threonine-protein kinase
Serine/threonine-protein kinase
Uncharacterized protein
Serine/threonine-protein kinase
Serine/threonine-protein kinase
Uncharacterized protein

Serine/threonine-protein kinase
Serine/threonine-protein kinase
Serine/threonine-protein kinase
Uncharacterized protein
Serine/threonine-protein kinase
Uncharacterized protein
Uncharacterized protein
Serine/threonine-protein kinase
Serine/threonine-protein kinase
Serine/threonine-protein kinase
Serine/threonine-protein kinase
Serine/threonine-protein kinase
Serine/threonine-protein kinase
Uncharacterized protein
Uncharacterized protein
Uncharacterized protein
Uncharacterized protein
Uncharacterized protein
S-domain receptor-like protein kinase family-1
Uncharacterized protein
Serine/threonine-protein kinase
Serine/threonine-protein kinase
Uncharacterized protein
Uncharacterized protein
Uncharacterized protein
Uncharacterized protein
Uncharacterized protein
Serine/threonine-protein kinase
Serine/threonine-protein kinase
Serine/threonine-protein kinase
Uncharacterized protein
Uncharacterized protein
S-domain receptor-like protein kinase
Serine/threonine-protein kinase
Serine/threonine-protein kinase
Uncharacterized protein
Uncharacterized protein
Uncharacterized protein
Uncharacterized protein
Serine/threonine-protein kinase
Serine/threonine-protein kinase

Serine/threonine-protein kinase
Serine/threonine-protein kinase
Uncharacterized protein
Uncharacterized protein
Serine/threonine-protein kinase
Uncharacterized protein
Uncharacterized protein
Serine/threonine-protein kinase
Uncharacterized protein
Serine/threonine-protein kinase
Serine/threonine-protein kinase
Uncharacterized protein
Uncharacterized protein
Serine/threonine-protein kinase
Uncharacterized protein
Serine/threonine-protein kinase
Uncharacterized protein
Uncharacterized protein
Serine/threonine-protein kinase
Uncharacterized protein
Uncharacterized protein
Serine/threonine-protein kinase
Serine/threonine-protein kinase
Uncharacterized protein
Uncharacterized protein
Serine/threonine-protein kinase
Serine/threonine-protein kinase
S-domain receptor-like protein kinase
Uncharacterized protein
Serine/threonine-protein kinase
Uncharacterized protein
Serine/threonine-protein kinase
Uncharacterized protein
Serine/threonine-protein kinase
Serine/threonine-protein kinase
Uncharacterized protein
Serine/threonine-protein kinase
Serine/threonine-protein kinase
Serine/threonine-protein kinase
Serine/threonine-protein kinase
Serine/threonine-protein kinase

Uncharacterized protein
Serine/threonine-protein kinase
Serine/threonine-protein kinase
Serine/threonine-protein kinase
Serine/threonine-protein kinase
Serine/threonine-protein kinase
Serine/threonine-protein kinase
Serine/threonine-protein kinase
Uncharacterized protein
Serine/threonine-protein kinase
Uncharacterized protein
Uncharacterized protein
Serine/threonine-protein kinase
Uncharacterized protein
Uncharacterized protein
Serine/threonine-protein kinase
Serine/threonine-protein kinase
Uncharacterized protein
Serine/threonine-protein kinase
Serine/threonine-protein kinase
Serine/threonine-protein kinase
Serine/threonine-protein kinase
Serine/threonine-protein kinase
Serine/threonine-protein kinase
Serine/threonine-protein kinase
Uncharacterized protein
Uncharacterized protein
Serine/threonine-protein kinase
Serine/threonine-protein kinase
Uncharacterized protein
Serine/threonine-protein kinase
Uncharacterized protein
Serine/threonine-protein kinase
Uncharacterized protein
Serine/threonine-protein kinase
Uncharacterized protein
Uncharacterized protein
Serine/threonine-protein kinase
Uncharacterized protein
Serine/threonine-protein kinase
Uncharacterized protein

Serine/threonine-protein kinase
Uncharacterized protein
Uncharacterized protein
S-domain receptor-like protein kinase family-1
Uncharacterized protein
Uncharacterized protein
Uncharacterized protein
Uncharacterized protein
Uncharacterized protein
Uncharacterized protein
Uncharacterized protein
Uncharacterized protein
Uncharacterized protein
Uncharacterized protein
Uncharacterized protein
Uncharacterized protein
Uncharacterized protein
Uncharacterized protein
Uncharacterized protein
Uncharacterized protein
Uncharacterized protein
Uncharacterized protein
Uncharacterized protein
Uncharacterized protein
Uncharacterized protein
Uncharacterized protein
Uncharacterized protein
Uncharacterized protein
Uncharacterized protein
Uncharacterized protein
Uncharacterized protein
Uncharacterized protein
Uncharacterized protein
Uncharacterized protein
Uncharacterized protein
Uncharacterized protein
Uncharacterized protein
Uncharacterized protein
Uncharacterized protein
Uncharacterized protein
Uncharacterized protein

Uncharacterized protein
Uncharacterized protein
Uncharacterized protein
Uncharacterized protein
Uncharacterized protein
Uncharacterized protein
Uncharacterized protein
Uncharacterized protein
Uncharacterized protein
Uncharacterized protein
Uncharacterized protein
Uncharacterized protein
Uncharacterized protein
Uncharacterized protein
Uncharacterized protein
Uncharacterized protein
Uncharacterized protein
Uncharacterized protein
Uncharacterized protein
Uncharacterized protein
Uncharacterized protein
Uncharacterized protein
Uncharacterized protein
Uncharacterized protein
Uncharacterized protein
Uncharacterized protein
Uncharacterized protein
Uncharacterized protein
Uncharacterized protein
Uncharacterized protein
Uncharacterized protein
Uncharacterized protein
Uncharacterized protein
Uncharacterized protein
Uncharacterized protein
Uncharacterized protein
Uncharacterized protein
Uncharacterized protein
Uncharacterized protein
Uncharacterized protein
Uncharacterized protein

Uncharacterized protein
Uncharacterized protein
Uncharacterized protein
Uncharacterized protein
Uncharacterized protein
Uncharacterized protein
Uncharacterized protein
Uncharacterized protein
Uncharacterized protein
Uncharacterized protein
Uncharacterized protein
Uncharacterized protein
Uncharacterized protein
Uncharacterized protein
Uncharacterized protein
Uncharacterized protein
Uncharacterized protein
Uncharacterized protein
Uncharacterized protein
Uncharacterized protein
Uncharacterized protein
Uncharacterized protein
Uncharacterized protein
Uncharacterized protein
Uncharacterized protein
Uncharacterized protein
Uncharacterized protein
Uncharacterized protein
Uncharacterized protein
Uncharacterized protein
Uncharacterized protein
Uncharacterized protein
Uncharacterized protein
Uncharacterized protein
Uncharacterized protein
Uncharacterized protein
Uncharacterized protein
Uncharacterized protein
Uncharacterized protein
Uncharacterized protein
Uncharacterized protein

Uncharacterized protein
Uncharacterized protein
Uncharacterized protein
Uncharacterized protein
Uncharacterized protein
Uncharacterized protein
Uncharacterized protein
Uncharacterized protein
Uncharacterized protein
Uncharacterized protein
Uncharacterized protein
Uncharacterized protein
Uncharacterized protein
Uncharacterized protein
Uncharacterized protein
Uncharacterized protein
Uncharacterized protein
Uncharacterized protein
Uncharacterized protein
Uncharacterized protein
Uncharacterized protein
Uncharacterized protein
Uncharacterized protein
Uncharacterized protein
Uncharacterized protein
Uncharacterized protein
Uncharacterized protein
Uncharacterized protein
Uncharacterized protein
Uncharacterized protein
Uncharacterized protein
Uncharacterized protein
Uncharacterized protein
Uncharacterized protein
Uncharacterized protein
Uncharacterized protein
Uncharacterized protein
Uncharacterized protein
Uncharacterized protein
Uncharacterized protein
Uncharacterized protein

Uncharacterized protein
Uncharacterized protein
Uncharacterized protein
Uncharacterized protein
Uncharacterized protein
Uncharacterized protein
Uncharacterized protein
Uncharacterized protein
Uncharacterized protein
Uncharacterized protein
Uncharacterized protein
Uncharacterized protein
Uncharacterized protein
Uncharacterized protein
Uncharacterized protein
Uncharacterized protein
Uncharacterized protein
Uncharacterized protein
S-domain receptor-like protein kinase family-3
Uncharacterized protein
Uncharacterized protein
Uncharacterized protein
Stress-induced receptor-like kinase 1
Uncharacterized protein
Uncharacterized protein
Uncharacterized protein
Uncharacterized protein
Uncharacterized protein
Uncharacterized protein
Uncharacterized protein
Uncharacterized protein
Uncharacterized protein
Uncharacterized protein
Uncharacterized protein
Uncharacterized protein
Uncharacterized protein
Uncharacterized protein
Uncharacterized protein
Uncharacterized protein
Uncharacterized protein
Uncharacterized protein

Uncharacterized protein
Uncharacterized protein
Uncharacterized protein
Uncharacterized protein
Uncharacterized protein
Uncharacterized protein
Uncharacterized protein
Uncharacterized protein
Uncharacterized protein
Uncharacterized protein
Uncharacterized protein
Uncharacterized protein
Uncharacterized protein
Uncharacterized protein
Uncharacterized protein
S-domain receptor-like protein kinase family-1
Os04g0356600 protein
Putative receptor kinase 5
Uncharacterized protein
Uncharacterized protein
Uncharacterized protein
Os04g0419700 protein
Uncharacterized protein
Uncharacterized protein
Uncharacterized protein
Putative S-receptor kinase
Os12g0130800 protein
Uncharacterized protein
Uncharacterized protein
Os04g0654800 protein
Uncharacterized protein
Uncharacterized protein OSJNBa0073C10.13
Uncharacterized protein
Os01g0223800 protein
Os09g0551000 protein
Os04g0634400 protein
Os05g0166900 protein (Fragment)
Putative Ser/Thr protein kinase
Os04g0633200 protein
Os06g0165500 protein
Retrotransposon protein, putative, Ty3-gypsy subclass

Os09g0551201 protein
Os09g0550700 protein
Os01g0668400 protein
Putative S-domain receptor-like protein kinase
Os04g0633300 protein
Putative S-receptor kinase
Putative receptor kinase 5
Os04g0420900 protein (Fragment)
Uncharacterized protein
OSJNBa0080E14.3 protein
Os04g0303100 protein
Os11g0208700 protein
Uncharacterized protein
D-mannose binding lectin family protein
Uncharacterized protein
Uncharacterized protein
Uncharacterized protein
Os09g0550500 protein (Fragment)
Os09g0550600 protein
Os06g0574550 protein (Fragment)
Uncharacterized protein
Os04g0632700 protein
Os09g0551500 protein
OSJNBa0043L09.14 protein
Os01g0890600 protein
Uncharacterized protein
Os04g0303300 protein
D-mannose binding lectin family protein, expressed
Receptor protein kinase-like
Uncharacterized protein
Uncharacterized protein
Os12g0130200 protein (Fragment)
Uncharacterized protein
OSJNBb0015D13.18 protein
Os04g0632901 protein
Os04g0633300 protein
Uncharacterized protein
Uncharacterized protein
Os11g0133500 protein
Os06g0574650 protein
Uncharacterized protein

Os09g0551500 protein
Os12g0177800 protein
Os07g0302100 protein (Fragment)
Os01g0871000 protein (Fragment)
Os04g0655000 protein
Os09g0551251 protein
Os05g0163500 protein
Os04g0420033 protein
Uncharacterized protein
Os04g0631800 protein
Os04g0303500 protein
D-mannose binding lectin family protein, expressed
OSJNBa0079M09.4 protein
Uncharacterized protein
Uncharacterized protein
Uncharacterized protein
Os01g0890600 protein (Fragment)
Uncharacterized protein
Os01g0670600 protein (Fragment)
Os04g0155500 protein
Os04g0632300 protein
Uncharacterized protein
Os11g0133100 protein
Uncharacterized protein
Os04g0633900 protein
Receptor-like protein kinase-like
Os01g0668901 protein
Uncharacterized protein
OSJNBa0043L09.5 protein
Os01g0890200 protein
Uncharacterized protein
Os03g0422800 protein (Fragment)
OSJNBa0028I23.5 protein
Uncharacterized protein
Os04g0632500 protein
Os11g0208700 protein
Uncharacterized protein
Os02g0527900 protein
OSJNBa0079M09.1 protein
Uncharacterized protein
Uncharacterized protein

Uncharacterized protein
Uncharacterized protein
Uncharacterized protein
Os12g0130500 protein (Fragment)
Os09g0550300 protein
Os04g0356600 protein (Fragment)
Os04g0633200 protein
Os04g0634400 protein
Os01g0642700 protein
Os04g0633600 protein
Os04g0226600 protein
OSJNBa0043L09.1 protein
Uncharacterized protein
Os07g0550500 protein
Os04g0420300 protein
Os09g0551400 protein
Uncharacterized protein
Uncharacterized protein
Uncharacterized protein
Uncharacterized protein
Uncharacterized protein
Putative D-mannose binding lectin (Fragment)
OSJNBa0043L09.7 protein
Uncharacterized protein
Uncharacterized protein
Uncharacterized protein
Uncharacterized protein
Uncharacterized protein
Uncharacterized protein
Uncharacterized protein
Uncharacterized protein
Uncharacterized protein
Os04g0633800 protein
Receptor-like protein kinase ARK1
Os04g0161850 protein (Fragment)
Os04g0420600 protein
Uncharacterized protein
Uncharacterized protein
Os04g0631800 protein (Fragment)
OSJNBb0015G09.5 protein
Os04g0632500 protein

Uncharacterized protein
OSJNBa0043L09.10 protein
Os01g0669100 protein
Putative receptor kinase 5
Os07g0186200 protein
Os01g0224000 protein
D-mannose binding lectin family protein
D-mannose binding lectin family protein, expressed
Putative receptor protein kinase
D-mannose binding lectin family protein, expressed
S-receptor kinase-like
Os01g0670600 protein
Putative S-domain receptor-like protein kinase
Putative serine/threonine kinase
Os01g0885700 protein
Os11g0149900 protein
Os11g0132900 protein (Fragment)
Os10g0101000 protein
Os01g0870400 protein
OSJNBa0094O15.4 protein
Os01g0670100 protein
OSJNBa0094O15.5 protein
Putative receptor-like protein kinase ARK1
OSJNBa0043L09.4 protein
D-mannose binding lectin family protein, expressed
Lectin protein kinase family protein, putative, expressed
Os05g0166300 protein
Os07g0551300 protein
Os05g0501400 protein
Os01g0587400 protein
D-mannose binding lectin family protein, expressed
Os07g0553633 protein
Putative receptor-like protein kinase
OSJNBa0043L09.9 protein
Putative receptor-like protein kinase
OSJNBa0071I13.4 protein
OSJNBa0028I23.2 protein
D-mannose binding lectin family protein
Os06g0165500 protein
Putative receptor-like protein kinase
Os02g0761600 protein

Os06g0575400 protein
OSJNBa0071I13.3 protein
Os01g0223900 protein
D-mannose binding lectin family protein
OSJNBa0028I23.8 protein
Os06g0552000 protein
Os06g0602500 protein
D-mannose binding lectin family protein, expressed
Os04g0632100 protein
Putative receptor kinase 5
Os04g0419900 protein (Fragment)
D-mannose binding lectin family protein
Os09g0550500 protein
D-mannose binding lectin family protein
OSJNBa0028I23.14 protein
OSJNBa0028I23.16 protein
D-mannose binding lectin family protein, expressed
D-mannose binding lectin family protein
Os07g0302100 protein
D-mannose binding lectin family protein, expressed
Os01g0222800 protein
Os02g0527900 protein
D-mannose binding lectin family protein, expressed
OSJNBa0043L09.8 protein
cDNA clone:002-172-B01, full insert sequence
OSJNBa0028I23.4 protein
OSJNBa0028I23.17 protein
OSJNBb0018A10.15 protein
S-receptor kinase-like
OSJNBb0015D13.19 protein
Os07g0550900 protein
Os01g0670300 protein
Os01g0784200 protein
Os01g0668600 protein
Os01g0889900 protein
Os04g0303100 protein
Os05g0165900 protein
Os05g0166600 protein
Os01g0784700 protein
Os01g0783800 protein
Os05g0252400 protein

D-mannose binding lectin family protein, expressed
Os12g0130200 protein
Os01g0155200 protein
OSJNBa0028I23.12 protein
OSJNBb0002J11.8 protein
OSJNBb0108J11.18 protein
Putative receptor kinase 5
Os01g0223700 protein
OSJNBb0066J23.17 protein
OSJNBa0028I23.7 protein
OSJNBa0028I23.11 protein
H0613A10.7 protein
OSIGBa0105P02.3 protein
OSIGBa0145C12.7 protein
H0613A10.6 protein
S-locus receptor-like kinase RLK11
H0105C05.10 protein
H0215E01.10 protein
S-locus receptor-like kinase RLK13
H0525E10.8 protein
B0808H03.4 protein
OSIGBa0147J19.7 protein
H0215E01.7 protein
H0613A10.1 protein
S-locus receptor-like kinase RLK14
H0525E10.7 protein
H0525E10.1 protein
Receptor-like protein kinase
H0215E01.6 protein
OSIGBa0157N01.3 protein
OSIGBa0148I18.4 protein
H0525E10.2 protein
OSIGBa0148I18.5 protein
H0613A10.3 protein
H0613A10.4 protein
H0215E01.5 protein
H0525E10.6 protein
Putative receptor-like protein kinase
B0808H03.2 protein
OSIGBa0135L04.3 protein
B0808H03.3 protein

H0315F07.2 protein
H0215E01.11 protein
H0315F07.5 protein
H0315F07.1 protein
Uncharacterized protein
Uncharacterized protein
Uncharacterized protein
Hepatocyte growth factor
Uncharacterized protein
Uncharacterized protein
Plasminogen
Plasminogen
Uncharacterized protein
Uncharacterized protein
Hepatocyte growth factor
Hepatocyte growth factor
Uncharacterized protein
Uncharacterized protein
Uncharacterized protein
Uncharacterized protein
Uncharacterized protein
Uncharacterized protein
Macrophage stimulating 1
Uncharacterized protein
Hepatocyte growth factor
Uncharacterized protein
Hepatocyte growth factor b
Macrophage stimulating 1
Uncharacterized protein
Hepatocyte growth factor
Coagulation factor XI (Fragment)
Hepatocyte growth factor a
Uncharacterized protein
Plasminogen
Skin mucus lectin (Fragment)
Uncharacterized protein
Coagulation factor XI
Uncharacterized protein (Fragment)
Uncharacterized protein (Fragment)
Ovarian tumour, otubain
WD40/YVTN repeat-like-containing domain

Uncharacterized protein
Uncharacterized protein
Uncharacterized protein (Fragment)
Cyclophilin-like peptidyl-prolyl cis-trans isomerase domain
PAN-1 domain
SdiA-regulated-domain-containing protein
Unnamed product
Cyclophilin-like peptidyl-prolyl cis-trans isomerase domain
Plasminogen
Uncharacterized protein
Coagulation factor XI
Macrophage stimulating 1
Kallikrein B1
Uncharacterized protein
Plasminogen
Uncharacterized protein
Uncharacterized protein
Uncharacterized protein
PAN/Apple domain-containing protein
PAN/Apple domain-containing protein
Uncharacterized protein
Plasminogen
Uncharacterized protein
Uncharacterized protein
Hepatocyte growth factor
Uncharacterized protein
Plasminogen
Uncharacterized protein
Uncharacterized protein
Uncharacterized protein
F11 isoform 2
Plasminogen
KLKB1 isoform 4 (Fragment)
KLKB1 isoform 7 (Fragment)
KLKB1 isoform 2
Hepatocyte growth factor
Uncharacterized protein
F11 isoform 6
KLKB1 isoform 1
HGF isoform 4
Hepatocyte growth factor

Uncharacterized protein
PLG isoform 1
Uncharacterized protein
PLGLB1 isoform 2
HGF isoform 10 (Fragment)
HGF isoform 9 (Fragment)
Hepatocyte growth factor
HGF isoform 3
F11 isoform 5 (Fragment)
Hepatocyte growth factor
PLG isoform 3
Plasminogen
Uncharacterized protein
Uncharacterized protein
KLKB1 isoform 3
Coagulation factor XI
HGF isoform 7
Uncharacterized protein
Hepatocyte growth factor
HGF isoform 5
PLGLB1 isoform 1
MST1 isoform 3
Uncharacterized protein
Uncharacterized protein
Uncharacterized protein
Uncharacterized protein
Uncharacterized protein
Uncharacterized protein
Uncharacterized protein
Uncharacterized protein
Uncharacterized protein
Uncharacterized protein
Uncharacterized protein
Uncharacterized protein
Uncharacterized protein
Uncharacterized protein
Uncharacterized protein
Uncharacterized protein
Uncharacterized protein
Uncharacterized protein
Uncharacterized protein

Uncharacterized protein
Uncharacterized protein
Uncharacterized protein
Uncharacterized protein
Uncharacterized protein
Uncharacterized protein
Uncharacterized protein
Uncharacterized protein
Uncharacterized protein
Uncharacterized protein
Uncharacterized protein
Uncharacterized protein
Uncharacterized protein
Uncharacterized protein
Uncharacterized protein
Uncharacterized protein
Uncharacterized protein
Uncharacterized protein
Uncharacterized protein
Uncharacterized protein
Uncharacterized protein
Uncharacterized protein
Uncharacterized protein
Uncharacterized protein
Uncharacterized protein
Uncharacterized protein
Uncharacterized protein
Uncharacterized protein
Uncharacterized protein
Uncharacterized protein
Uncharacterized protein
Uncharacterized protein
Uncharacterized protein
Uncharacterized protein
Uncharacterized protein
Uncharacterized protein
Uncharacterized protein
Uncharacterized protein
Uncharacterized protein
Uncharacterized protein
Uncharacterized protein

Uncharacterized protein
Uncharacterized protein
Uncharacterized protein
Uncharacterized protein
Uncharacterized protein
Uncharacterized protein
Uncharacterized protein
Uncharacterized protein
Uncharacterized protein
Uncharacterized protein
Uncharacterized protein
Uncharacterized protein
Uncharacterized protein
Uncharacterized protein
Uncharacterized protein
Uncharacterized protein
Uncharacterized protein
Uncharacterized protein
Uncharacterized protein
Uncharacterized protein
Uncharacterized protein
Uncharacterized protein
Uncharacterized protein
Uncharacterized protein
Uncharacterized protein
Uncharacterized protein
Uncharacterized protein
Uncharacterized protein
Uncharacterized protein
Uncharacterized protein
Uncharacterized protein
Uncharacterized protein
Uncharacterized protein
Uncharacterized protein
Uncharacterized protein
Uncharacterized protein
Uncharacterized protein
Uncharacterized protein
Uncharacterized protein
Uncharacterized protein
Uncharacterized protein

Uncharacterized protein
Uncharacterized protein
Uncharacterized protein
Uncharacterized protein
Uncharacterized protein
Uncharacterized protein
Uncharacterized protein
Uncharacterized protein
Uncharacterized protein
Uncharacterized protein
Uncharacterized protein
Uncharacterized protein
Uncharacterized protein
Uncharacterized protein
Uncharacterized protein
Uncharacterized protein (Fragment)
Uncharacterized protein
Uncharacterized protein
Uncharacterized protein
Uncharacterized protein
Uncharacterized protein
Uncharacterized protein
Uncharacterized protein
Uncharacterized protein
Uncharacterized protein
Uncharacterized protein
Uncharacterized protein
Uncharacterized protein
Uncharacterized protein
Uncharacterized protein
Uncharacterized protein
Uncharacterized protein
Uncharacterized protein
Uncharacterized protein
Uncharacterized protein
Uncharacterized protein
Uncharacterized protein
Uncharacterized protein
Uncharacterized protein
Uncharacterized protein
Uncharacterized protein

Uncharacterized protein
Uncharacterized protein
Uncharacterized protein
Uncharacterized protein
Uncharacterized protein
Uncharacterized protein
Uncharacterized protein (Fragment)
Uncharacterized protein
Uncharacterized protein
Uncharacterized protein
Uncharacterized protein
Uncharacterized protein
Uncharacterized protein
Uncharacterized protein
Uncharacterized protein
Uncharacterized protein
Uncharacterized protein
Uncharacterized protein
Uncharacterized protein
Uncharacterized protein
Uncharacterized protein
Uncharacterized protein
Uncharacterized protein
Uncharacterized protein
Uncharacterized protein
Uncharacterized protein
Uncharacterized protein
Uncharacterized protein
Uncharacterized protein
Uncharacterized protein
Uncharacterized protein
Uncharacterized protein
Uncharacterized protein
Uncharacterized protein
Uncharacterized protein
Uncharacterized protein
Uncharacterized protein
Uncharacterized protein
Uncharacterized protein
Uncharacterized protein
Uncharacterized protein

Uncharacterized protein
Uncharacterized protein
Uncharacterized protein
Uncharacterized protein
Uncharacterized protein
Uncharacterized protein
Uncharacterized protein
Uncharacterized protein
Uncharacterized protein
Uncharacterized protein
Uncharacterized protein
Uncharacterized protein
Uncharacterized protein
Uncharacterized protein
Uncharacterized protein
Uncharacterized protein
Uncharacterized protein
Uncharacterized protein
Uncharacterized protein
Uncharacterized protein
Uncharacterized protein
Uncharacterized protein
Uncharacterized protein
Uncharacterized protein
Uncharacterized protein
Uncharacterized protein
Uncharacterized protein
Uncharacterized protein
Uncharacterized protein
Uncharacterized protein
Uncharacterized protein
Uncharacterized protein
Uncharacterized protein
Uncharacterized protein
Uncharacterized protein
Uncharacterized protein
Uncharacterized protein
Uncharacterized protein
Uncharacterized protein
Uncharacterized protein
Uncharacterized protein

Uncharacterized protein
Uncharacterized protein
Uncharacterized protein
Uncharacterized protein
Uncharacterized protein
Uncharacterized protein
Uncharacterized protein
Uncharacterized protein
Uncharacterized protein
Uncharacterized protein
Uncharacterized protein
Uncharacterized protein
Uncharacterized protein
Uncharacterized protein
Uncharacterized protein
Uncharacterized protein
Uncharacterized protein
Uncharacterized protein
Uncharacterized protein
Uncharacterized protein
Uncharacterized protein
Uncharacterized protein
Uncharacterized protein
Uncharacterized protein
Putative S-locus receptor-like protein kinase family protein
G-type lectin S-receptor-like serine/threonine-protein kinase SD2-5
G-type lectin S-receptor-like serine/threonine-protein kinase
Uncharacterized protein
Uncharacterized protein
Receptor-like serine/threonine-protein kinase SD1-8
Uncharacterized protein
Uncharacterized protein
G-type lectin S-receptor-like serine/threonine-protein kinase RLK1
G-type lectin S-receptor-like serine/threonine-protein kinase
Putative receptor protein kinase ZmPK1
Uncharacterized protein
Uncharacterized protein
Uncharacterized protein
Uncharacterized protein
Uncharacterized protein
Receptor-like serine/threonine-protein kinase SD1-8 isoform X1

Uncharacterized protein
G-type lectin S-receptor-like serine/threonine-protein kinase
Receptor-like serine/threonine-protein kinase SD1-8
Uncharacterized protein
Uncharacterized protein
Uncharacterized protein
G-type lectin S-receptor-like serine/threonine-protein kinase
Receptor-like serine/threonine-protein kinase SD1-8
G-type lectin S-receptor-like serine/threonine-protein kinase
Receptor-like serine/threonine-protein kinase SD1-8
Uncharacterized protein
G-type lectin S-receptor-like serine/threonine-protein kinase
Uncharacterized protein
G-type lectin S-receptor-like serine/threonine-protein kinase
Uncharacterized protein
G-type lectin S-receptor-like serine/threonine-protein kinase
Uncharacterized protein
G-type lectin S-receptor-like serine/threonine-protein kinase
G-type lectin S-receptor-like serine/threonine-protein kinase
Putative receptor protein kinase ZmPK1
Putative receptor protein kinase ZmPK1
G-type lectin S-receptor-like serine/threonine-protein kinase SD2-5
G-type lectin S-receptor-like serine/threonine-protein kinase
Uncharacterized protein
Uncharacterized protein
G-type lectin S-receptor-like serine/threonine-protein kinase
Uncharacterized protein
G-type lectin S-receptor-like serine/threonine-protein kinase
G-type lectin S-receptor-like serine/threonine-protein kinase
G-type lectin S-receptor-like serine/threonine-protein kinase
G-type lectin S-receptor-like serine/threonine-protein kinase
Putative receptor protein kinase ZmPK1 isoform X1
Uncharacterized protein
Uncharacterized protein
Uncharacterized protein
Putative receptor protein kinase ZmPK1 isoform X2
G-type lectin S-receptor-like serine/threonine-protein kinase
Uncharacterized protein
G-type lectin S-receptor-like serine/threonine-protein kinase
G-type lectin S-receptor-like serine/threonine-protein kinase
G-type lectin S-receptor-like serine/threonine-protein kinase

G-type lectin S-receptor-like serine/threonine-protein kinase
Uncharacterized protein
Receptor-like serine/threonine-protein kinase SD1-8
Putative receptor protein kinase ZmPK1
Putative D-mannose binding lectin receptor-like protein kinase family protein
G-type lectin S-receptor-like serine/threonine-protein kinase
Putative receptor protein kinase ZmPK1
Receptor-like serine/threonine-protein kinase SD1-8
G-type lectin S-receptor-like serine/threonine-protein kinase
Receptor-like serine/threonine-protein kinase SD1-8
Uncharacterized protein
Uncharacterized protein
G-type lectin S-receptor-like serine/threonine-protein kinase
Putative G-type lectin S-receptor-like serine/threonine-protein kinase
G-type lectin S-receptor-like serine/threonine-protein kinase
Uncharacterized protein
Uncharacterized protein
Uncharacterized protein
Uncharacterized protein
Receptor-like serine/threonine-protein kinase SD1-8
G-type lectin S-receptor-like serine/threonine-protein kinase
Uncharacterized protein
G-type lectin S-receptor-like serine/threonine-protein kinase
Uncharacterized protein
Receptor-like serine/threonine-protein kinase SD1-8
Putative G-type lectin S-receptor-like serine/threonine-protein kinase
Uncharacterized protein
Uncharacterized protein
Uncharacterized protein
Uncharacterized protein
Uncharacterized protein
Uncharacterized protein
PAN domain-containing protein
Putative receptor protein kinase ZmPK1
Uncharacterized protein
G-type lectin S-receptor-like serine/threonine-protein kinase SD2-5
Uncharacterized protein
G-type lectin S-receptor-like serine/threonine-protein kinase
PAN domain-containing protein
Putative receptor protein kinase ZmPK1
G-type lectin S-receptor-like serine/threonine-protein kinase

Receptor-like serine/threonine-protein kinase SD1-8 isoform X2
G-type lectin S-receptor-like serine/threonine-protein kinase
Putative receptor protein kinase ZmPK1
Uncharacterized conserved protein YfaS, alpha-2-macroglobulin family
Alpha-2-macroglobulin family N-terminal region
Alpha-2-macroglobulin
Putative trypsin-like serine protease
Putative rhoa gtpase effector dia/diaphanous (Fragment)
Uncharacterized protein (Fragment)
Putative conserved plasma membrane protein (Fragment)
Uncharacterized protein
Peptidyl-prolyl isomerase cwc27
Uncharacterized protein
Cuticlin-1
Uncharacterized protein
Uncharacterized protein
Uncharacterized protein
Uncharacterized protein
Cuticlin-1
Uncharacterized protein
Uncharacterized protein
Peptidyl-prolyl cis-trans isomerase CWC27-like
Uncharacterized protein
Uncharacterized protein
Uncharacterized protein
Uncharacterized protein
Uncharacterized protein
Plasminogen
Uncharacterized protein
Uncharacterized protein
Uncharacterized protein
Uncharacterized protein
Uncharacterized protein
Alpha-2-macroglobulin domain-containing protein
Uncharacterized protein
Uncharacterized protein
Alpha-2-macroglobulin
Alpha-2-macroglobulin domain protein
Alpha-2-macroglobulin
Alpha-2-macroglobulin
Uncharacterized protein

Uncharacterized protein
Alpha-2-macroglobulin
Uncharacterized protein
Alpha-2-macroglobulin
Uncharacterized protein
Uncharacterized protein
Uncharacterized protein
Uncharacterized protein
PAN domain-containing protein
Alpha-2-macroglobulin
Uncharacterized protein
Uncharacterized protein
Alpha-2-macroglobulin
Uncharacterized protein
Uncharacterized protein
Alpha-2-macroglobulin
Alpha-2-macroglobulin
Alpha-2-macroglobulin
Alpha-2-macroglobulin
PAN domain-containing protein
Macrophage stimulating-1 protein
Uncharacterized protein
Plasminogen
Hepatocyte growth factor
Uncharacterized protein
Plasminogen
Uncharacterized protein
Plasminogen
Hepatocyte growth factor
Plasminogen
Plasminogen
Plasminogen
Uncharacterized protein
Uncharacterized protein
Uncharacterized protein
Uncharacterized protein
Uncharacterized protein
Galactose oxidase
Membrane protein
S-receptor-like serine/threonine-protein kinase
S-receptor-like serine/threonine-protein kinase (Fragment)

S-locus glycoprotein
PAN/Apple domain containing protein (Fragment)
S-locus glycoprotein domain containing protein
S-locus glycoprotein domain containing protein
S-locus glycoprotein
Serine/threonine protein kinase
S-receptor-like serine/threonine-protein kinase
S-receptor-like serine/threonine-protein kinase
Lectin domain containing protein
S-receptor-like serine/threonine-protein kinase
S-receptor-like serine/threonine-protein kinase
S-receptor-like serine/threonine-protein kinase
S-receptor-like serine/threonine-protein kinase
S-receptor-like serine/threonine-protein kinase
S-locus glycoprotein
S-receptor-like serine/threonine-protein kinase
S-locus glycoprotein
Mitogen-activated protein kinase kinase kinase
PAN/Apple domain containing protein
Tyrosine-protein kinase
Mitogen-activated protein kinase kinase kinase
Mitogen-activated protein kinase kinase kinase (Fragment)
S-receptor-like serine/threonine-protein kinase
S-locus glycoprotein (Fragment)
S-locus glycoprotein
S-receptor-like serine/threonine-protein kinase
S-receptor-like serine/threonine-protein kinase
S-receptor-like serine/threonine-protein kinase
S-locus glycoprotein
S-receptor-like serine/threonine-protein kinase
Mitogen-activated protein kinase kinase kinase
S-locus glycoprotein
S-locus glycoprotein domain containing protein
S-receptor-like serine/threonine-protein kinase
S-locus glycoprotein
S-locus glycoprotein domain containing protein
S-locus glycoprotein
S-receptor-like serine/threonine-protein kinase
S-receptor-like serine/threonine-protein kinase
S-receptor-like serine/threonine-protein kinase
Serine/threonine protein kinase

S-locus glycoprotein
Tyrosine-protein kinase
S-receptor-like serine/threonine-protein kinase
S-locus glycoprotein
Phosphorylase kinase, gamma catalytic subunit
S-receptor-like serine/threonine-protein kinase (Fragment)
S-receptor-like serine/threonine-protein kinase
S-locus glycoprotein
Kinase-like protein
S-receptor-like serine/threonine-protein kinase (Fragment)
Tyrosine-protein kinase
S-receptor-like serine/threonine-protein kinase
S-locus glycoprotein
Serine/threonine protein kinase
S-receptor-like serine/threonine-protein kinase
Serine/threonine protein kinase
S-receptor-like serine/threonine-protein kinase
S-receptor-like serine/threonine-protein kinase
S-receptor-like serine/threonine-protein kinase
S-receptor-like serine/threonine-protein kinase
S-receptor-like serine/threonine-protein kinase
S-receptor-like serine/threonine-protein kinase
S-receptor-like serine/threonine-protein kinase
S-locus glycoprotein domain containing protein
Serine/threonine protein kinase
Serine/threonine protein kinase
Lectin domain containing protein
S-receptor-like serine/threonine-protein kinase
S-locus glycoprotein
S-receptor-like serine/threonine-protein kinase
PAN/Apple domain containing protein
S-receptor-like serine/threonine-protein kinase
Phosphorylase kinase, gamma catalytic subunit (Fragment)
S-receptor-like serine/threonine-protein kinase
Mitogen-activated protein kinase kinase kinase
S-receptor-like serine/threonine-protein kinase
Mitogen-activated protein kinase kinase kinase
Serine/threonine protein kinase
S-receptor-like serine/threonine-protein kinase
S-receptor-like serine/threonine-protein kinase
S-receptor-like serine/threonine-protein kinase

Mitogen-activated protein kinase kinase kinase (Fragment)
Serine/threonine protein kinase
S-receptor-like serine/threonine-protein kinase
Serine/threonine protein kinase
Serine/threonine protein kinase
S-receptor-like serine/threonine-protein kinase
S-locus glycoprotein
S-locus glycoprotein
PAN/Apple domain containing protein
Tyrosine-protein kinase
Lectin domain containing protein
S-locus glycoprotein
S-receptor-like serine/threonine-protein kinase
Putative antigen B membrane protein (Fragment)
Collectin-10 (Fragment)
Putative antigen B membrane protein (Fragment)
Putative antigen B membrane protein (Fragment)
Uncharacterized protein (Fragment)
Uncharacterized protein (Fragment)
Uncharacterized protein
Uncharacterized protein
Uncharacterized protein
Uncharacterized protein
Uncharacterized protein
Uncharacterized protein
Uncharacterized protein
Uncharacterized protein
Uncharacterized protein
Uncharacterized protein
Uncharacterized protein
Uncharacterized protein
Uncharacterized protein
Uncharacterized protein
Uncharacterized protein
Uncharacterized protein
Uncharacterized protein
Uncharacterized protein
Uncharacterized protein
Uncharacterized protein
Uncharacterized protein
Uncharacterized protein

Uncharacterized protein
Uncharacterized protein
Uncharacterized protein
Hepatocyte growth factor isoform A
Plasminogen
Coagulation factor XI
Plasminogen
Hepatocyte growth factor-like protein isoform B
Hepatocyte growth factor
Hepatocyte growth factor-like protein isoform A
Uncharacterized protein (Fragment)
Uncharacterized protein (Fragment)
Uncharacterized protein
No-mechanoreceptor potential A, putative
Uncharacterized protein
Uncharacterized protein
Uncharacterized protein
Uncharacterized protein
Uncharacterized protein
PAN domain protein
PAN domain-containing protein
Uncharacterized protein
Uncharacterized protein
Uncharacterized protein
Uncharacterized protein
Uncharacterized protein
Uncharacterized protein
Uncharacterized protein
PAN domain-containing protein
Plasminogen (Fragment)
Coagulation factor XI (Fragment)
Plasminogen (Fragment)
Hepatocyte growth factor (Fragment)
Hepatocyte growth factor (Fragment)
Macrophage stimulating 1
Uncharacterized protein
Uncharacterized protein
PAN domain-containing protein
Uncharacterized protein (Fragment)
Uncharacterized protein
Uncharacterized protein

Uncharacterized protein
Uncharacterized protein
Uncharacterized protein
Uncharacterized protein
Uncharacterized protein
Putative neurotrypsin-like
Uncharacterized protein
Uncharacterized protein
Uncharacterized protein
Uncharacterized protein
Uncharacterized protein
Uncharacterized protein
Uncharacterized protein
Uncharacterized protein
Uncharacterized protein
Uncharacterized protein
Uncharacterized protein
Uncharacterized protein
Uncharacterized protein
Uncharacterized protein
Uncharacterized protein
Apple-like
Uncharacterized protein
Uncharacterized protein
Uncharacterized protein
Uncharacterized protein
Uncharacterized protein
Uncharacterized protein
Uncharacterized protein
Uncharacterized protein
Uncharacterized protein
Uncharacterized protein
Uncharacterized protein
Uncharacterized protein
Putative beta-1,3-galactosyltransferase
Uncharacterized protein
Uncharacterized protein
Uncharacterized protein
Uncharacterized protein
Uncharacterized protein
Uncharacterized protein

Uncharacterized protein
Uncharacterized protein
Uncharacterized protein
Uncharacterized protein
Uncharacterized protein
Uncharacterized protein
Uncharacterized protein
Uncharacterized protein
Uncharacterized protein
Uncharacterized protein
Uncharacterized protein
Hepatocyte growth factor-like protein (Fragment)
Plasminogen (Fragment)
Uncharacterized protein
Uncharacterized protein (Fragment)
Uncharacterized protein (Fragment)
Uncharacterized protein
Uncharacterized protein
Galactose oxidase
Uncharacterized protein
Macrophage stimulating 1
Macrophage stimulating 1
Uncharacterized protein
Macrophage stimulating 1
Hepatocyte growth factor
Uncharacterized protein
Macrophage stimulating 1
Uncharacterized protein
Uncharacterized protein
Uncharacterized protein
Uncharacterized protein
Uncharacterized protein
Uncharacterized protein
Uncharacterized protein
Uncharacterized protein
Uncharacterized protein
Uncharacterized protein
Uncharacterized protein
Uncharacterized protein
Uncharacterized protein
Uncharacterized protein

Alpha-2-macroglobulin family protein
Uncharacterized protein
Uncharacterized protein
Uncharacterized protein
Predicted protein (Fragment)
Uncharacterized protein
Uncharacterized protein
Uncharacterized protein
Uncharacterized protein
Uncharacterized protein
Uncharacterized protein
Uncharacterized protein
Uncharacterized protein
Uncharacterized protein
Uncharacterized protein
Hepatocyte growth factor (Fragment)
Coagulation factor XI (Fragment)
Plasminogen (Fragment)
Plasminogen (Fragment)
Hepatocyte growth factor (Fragment)
Plasma kallikrein (Fragment)
Plasminogen (Fragment)
Plasminogen (Fragment)
Uncharacterized protein
Uncharacterized protein
Uncharacterized protein
Uncharacterized protein
Uncharacterized protein
Uncharacterized protein
Uncharacterized protein
Uncharacterized protein
Uncharacterized protein
Uncharacterized protein
Uncharacterized protein
Uncharacterized protein
Uncharacterized protein
Uncharacterized protein
Uncharacterized protein
Uncharacterized protein
Uncharacterized protein
Uncharacterized protein

Uncharacterized protein
Uncharacterized protein
Uncharacterized protein
Uncharacterized protein
Uncharacterized protein
Uncharacterized protein
Uncharacterized protein
Uncharacterized protein
Uncharacterized protein
Uncharacterized protein
Uncharacterized protein
Uncharacterized protein
Uncharacterized protein
Uncharacterized protein
Uncharacterized protein
Uncharacterized protein
Uncharacterized protein
Uncharacterized protein
Uncharacterized protein
Uncharacterized protein
Uncharacterized protein
Uncharacterized protein
Uncharacterized protein
Uncharacterized protein
Uncharacterized protein
Uncharacterized protein
Uncharacterized protein
Uncharacterized protein
Uncharacterized protein
Uncharacterized protein
Uncharacterized protein
Uncharacterized protein
Uncharacterized protein
Uncharacterized protein
Uncharacterized protein
Uncharacterized protein
Uncharacterized protein
Uncharacterized protein
Uncharacterized protein
Uncharacterized protein
Uncharacterized protein

Uncharacterized protein
Uncharacterized protein
Uncharacterized protein
Uncharacterized protein
Uncharacterized protein
Uncharacterized protein
Uncharacterized protein
Uncharacterized protein
Uncharacterized protein
Uncharacterized protein
Uncharacterized protein
Uncharacterized protein
Uncharacterized protein
Uncharacterized protein
Uncharacterized protein
Uncharacterized protein
Uncharacterized protein
Uncharacterized protein
Uncharacterized protein
Uncharacterized protein
Uncharacterized protein
Uncharacterized protein
Uncharacterized protein
Uncharacterized protein (Fragment)
Uncharacterized protein
Uncharacterized protein
Uncharacterized protein
Uncharacterized protein
Uncharacterized protein
Uncharacterized protein
Uncharacterized protein
Uncharacterized protein
Uncharacterized protein
Uncharacterized protein
Uncharacterized protein
Uncharacterized protein
Uncharacterized protein
Uncharacterized protein
Uncharacterized protein
Uncharacterized protein
Uncharacterized protein

Uncharacterized protein
Uncharacterized protein
Uncharacterized protein (Fragment)
Uncharacterized protein
Uncharacterized protein
Uncharacterized protein
Uncharacterized protein
Uncharacterized protein
Uncharacterized protein
Uncharacterized protein
Uncharacterized protein
Uncharacterized protein
Uncharacterized protein
Uncharacterized protein
Uncharacterized protein
Uncharacterized protein
Uncharacterized protein (Fragment)
Uncharacterized protein (Fragment)
Uncharacterized protein
Uncharacterized protein
Uncharacterized protein
Uncharacterized protein
Uncharacterized protein (Fragment)
Uncharacterized protein
Uncharacterized protein
Uncharacterized protein
Uncharacterized protein
Uncharacterized protein
Uncharacterized protein
Uncharacterized protein
Uncharacterized protein
Uncharacterized protein
Uncharacterized protein
Uncharacterized protein (Fragment)
Uncharacterized protein
Uncharacterized protein
Uncharacterized protein
Uncharacterized protein
Uncharacterized protein
Uncharacterized protein
Uncharacterized protein

Uncharacterized protein
Uncharacterized protein
Uncharacterized protein
Uncharacterized protein
Uncharacterized protein
Uncharacterized protein
Uncharacterized protein
Uncharacterized protein
Uncharacterized protein
Uncharacterized protein
Uncharacterized protein
Uncharacterized protein
Uncharacterized protein
Uncharacterized protein
Uncharacterized protein
Uncharacterized protein
Uncharacterized protein
Uncharacterized protein
Uncharacterized protein
Uncharacterized protein
Uncharacterized protein
Uncharacterized protein
Uncharacterized protein
Uncharacterized protein
Uncharacterized protein
Uncharacterized protein
Uncharacterized protein
Uncharacterized protein
Uncharacterized protein
Uncharacterized protein
Uncharacterized protein
Uncharacterized protein
Galactose oxidase
Uncharacterized protein
Uncharacterized protein
Uncharacterized protein
Uncharacterized protein
Uncharacterized protein
Uncharacterized protein
Uncharacterized protein
Uncharacterized protein

Uncharacterized protein
Uncharacterized protein
Uncharacterized protein
Uncharacterized protein
Plasminogen (Fragment)
Hepatocyte growth factor (Fragment)
receptor-like serine/threonine-protein kinase SD1-7
receptor-like serine/threonine-protein kinase SD1-8
G-type lectin S-receptor-like serine/threonine-protein kinase SD3-1 isoform X1
receptor-like serine/threonine-protein kinase SD1-8
receptor-like serine/threonine-protein kinase SD1-8 isoform X1
G-type lectin S-receptor-like serine/threonine-protein kinase At2g19130
G-type lectin S-receptor-like serine/threonine-protein kinase At2g19130
receptor-like serine/threonine-protein kinase SD1-8
receptor-like serine/threonine-protein kinase SD1-7
G-type lectin S-receptor-like serine/threonine-protein kinase B120
G-type lectin S-receptor-like serine/threonine-protein kinase At2g19130
S-locus-specific glycoprotein S13-like
G-type lectin S-receptor-like serine/threonine-protein kinase At4g27290
receptor-like serine/threonine-protein kinase SD1-7 isoform X2
G-type lectin S-receptor-like serine/threonine-protein kinase At1g34300
receptor-like serine/threonine-protein kinase SD1-8
G-type lectin S-receptor-like serine/threonine-protein kinase At4g27290
G-type lectin S-receptor-like serine/threonine-protein kinase At2g19130
G-type lectin S-receptor-like serine/threonine-protein kinase At2g19130
G-type lectin S-receptor-like serine/threonine-protein kinase At2g19130
S-locus-specific glycoprotein BS29-1-like
S-locus-specific glycoprotein S6-like
G-type lectin S-receptor-like serine/threonine-protein kinase At2g19130
S-locus-specific glycoprotein S13-like
G-type lectin S-receptor-like serine/threonine-protein kinase At1g34300
receptor-like serine/threonine-protein kinase SD1-7 isoform X1
receptor-like serine/threonine-protein kinase SD1-8
G-type lectin S-receptor-like serine/threonine-protein kinase LECRK3
receptor-like serine/threonine-protein kinase SD1-8
G-type lectin S-receptor-like serine/threonine-protein kinase SD2-5
G-type lectin S-receptor-like serine/threonine-protein kinase B120
uncharacterized protein LOC103695870
putative receptor protein kinase ZmPK1
putative receptor protein kinase ZmPK1
PAN domain-containing protein At5g03700

receptor-like serine/threonine-protein kinase SD1-8
G-type lectin S-receptor-like serine/threonine-protein kinase At2g19130
G-type lectin S-receptor-like serine/threonine-protein kinase SD3-1
G-type lectin S-receptor-like serine/threonine-protein kinase At4g27290
receptor-like serine/threonine-protein kinase SD1-8
putative receptor protein kinase ZmPK1
receptor-like serine/threonine-protein kinase SD1-8
G-type lectin S-receptor-like serine/threonine-protein kinase At2g19130
uncharacterized protein LOC103707212
G-type lectin S-receptor-like serine/threonine-protein kinase At2g19130
G-type lectin S-receptor-like serine/threonine-protein kinase At5g35370
receptor-like serine/threonine-protein kinase SD1-8 isoform X2
G-type lectin S-receptor-like serine/threonine-protein kinase At4g27290
receptor-like serine/threonine-protein kinase SD1-8
putative receptor protein kinase ZmPK1
uncharacterized protein LOC103717327
G-type lectin S-receptor-like serine/threonine-protein kinase At2g19130
G-type lectin S-receptor-like serine/threonine-protein kinase At5g35370
LOW QUALITY PROTEIN: G-type lectin S-receptor-like serine/threonine-protein kinase SD2-5
G-type lectin S-receptor-like serine/threonine-protein kinase At2g19130
G-type lectin S-receptor-like serine/threonine-protein kinase At4g27290 isoform X1
G-type lectin S-receptor-like serine/threonine-protein kinase At2g19130
G-type lectin S-receptor-like serine/threonine-protein kinase SD2-2
G-type lectin S-receptor-like serine/threonine-protein kinase B120
Uncharacterized protein
Uncharacterized protein (Fragment)
Uncharacterized protein
Uncharacterized protein (Fragment)
Uncharacterized protein (Fragment)
Uncharacterized protein (Fragment)
Uncharacterized protein (Fragment)
Uncharacterized protein
Uncharacterized protein (Fragment)
Uncharacterized protein
Uncharacterized protein (Fragment)
Uncharacterized protein
Uncharacterized protein
Uncharacterized protein
Uncharacterized protein
Uncharacterized protein
Serine protease

Uncharacterized protein
Uncharacterized protein
Uncharacterized protein
Uncharacterized protein
Uncharacterized protein
Uncharacterized protein
Uncharacterized protein
Large extracellular alpha-helical protein
Uncharacterized protein
Uncharacterized protein
Predicted protein (Fragment)
Uncharacterized protein
Uncharacterized protein
Uncharacterized protein
Uncharacterized protein
Predicted protein
Uncharacterized protein
Uncharacterized protein
Predicted protein
Predicted protein
Plasminogen
Plasminogen
micronemal protein 4-like
Plasminogen
coagulation factor XI
hepatocyte growth factor-like protein
Hepatocyte growth factor
Uncharacterized protein (Fragment)
Uncharacterized protein (Fragment)
Uncharacterized protein
Uncharacterized protein
Uncharacterized protein
Uncharacterized protein
Uncharacterized protein
Uncharacterized protein
Uncharacterized protein
Uncharacterized protein
Uncharacterized protein
Uncharacterized protein
Uncharacterized protein
Uncharacterized protein

Uncharacterized protein
Uncharacterized protein
Uncharacterized protein
Uncharacterized protein
Uncharacterized protein
Uncharacterized protein
Uncharacterized protein
Uncharacterized protein
Cellulose binding elicitor lectin (CBEL), putative
Uncharacterized protein
Uncharacterized protein
Uncharacterized protein
Cellulose binding elicitor lectin (CBEL), putative
Uncharacterized protein
Cellulose binding elicitor lectin (CBEL), putative
Uncharacterized protein
Uncharacterized protein
Uncharacterized protein
Uncharacterized protein
Carbohydrate-binding protein, putative
Uncharacterized protein
Uncharacterized protein
Carbohydrate-binding protein, putative
Cellulose binding elicitor lectin (CBEL), putative
Uncharacterized protein
Cellulose binding elicitor lectin, putative
Uncharacterized protein
Uncharacterized protein
Uncharacterized protein
Carbohydrate-binding protein, putative
CBEL-like protein
Small cellulose-binding protein
Uncharacterized protein (Fragment)
Uncharacterized protein
Uncharacterized protein
Uncharacterized protein
Uncharacterized protein
Uncharacterized protein
Uncharacterized protein
Uncharacterized protein
Uncharacterized protein

Uncharacterized protein
Uncharacterized protein
Uncharacterized protein
Uncharacterized protein
Uncharacterized protein
Uncharacterized protein
Uncharacterized protein
Uncharacterized protein
Uncharacterized protein
Uncharacterized protein
Uncharacterized protein
Uncharacterized protein
Uncharacterized protein
Uncharacterized protein
Uncharacterized protein
Glycoside hydrolase
Uncharacterized protein
Cellulose binding elicitor lectin
Uncharacterized protein
SCP-like extracellular protein
A cellulose-binding elicitor and lectin-like protein
A cellulose-binding elicitor and lectin-like protein
Cellulose binding elicitor lectin
Uncharacterized protein
Cellulose binding elicitor lectin (Fragment)
SCP-like extracellular protein
Cyst germination specific acidic repeat protein (Fragment)
Uncharacterized protein
Carbohydrate-binding protein
Carbohydrate-binding protein
Uncharacterized protein
Uncharacterized protein
Uncharacterized protein
Uncharacterized protein
Uncharacterized protein
Tartrate-resistant acid phosphatase type 5
SCP extracellular protein
Cellobiose dehydrogenase
Cellobiose dehydrogenase
SCP extracellular protein
Glucan 1

Uncharacterized protein
Uncharacterized protein
Tartrate-resistant acid phosphatase type 5
Uncharacterized protein
Uncharacterized protein
Uncharacterized protein
WASH complex subunit CCDC53
Glucan 1
Uncharacterized protein
Uncharacterized protein
Cellobiose dehydrogenase
CBEL protein
Cellulose binding elicitor lectin (CBEL)
Cellulose binding elicitor lectin (CBEL)
Cellulose binding elicitor lectin (CBEL)
Cellulose binding elicitor lectin (CBEL)
Cellulose binding elicitor lectin (CBEL)
Uncharacterized protein
Carbohydrate-binding protein
Carbohydrate-binding protein
SCP-like extracellular protein
CBEL-like protein
SCP-like extracellular protein
Cellulose binding elicitor lectin
SCP-like extracellular protein
Uncharacterized protein
Cellulose binding elicitor lectin (CBEL)
Uncharacterized protein
Uncharacterized protein
Uncharacterized protein
Uncharacterized protein
Uncharacterized protein
Uncharacterized protein
Uncharacterized protein
Uncharacterized protein
Uncharacterized protein
Uncharacterized protein
Uncharacterized protein
Uncharacterized protein
Uncharacterized protein
Uncharacterized protein

Uncharacterized protein
Uncharacterized protein
Uncharacterized protein
Uncharacterized protein
Uncharacterized protein
Uncharacterized protein
Uncharacterized protein
Uncharacterized protein
Uncharacterized protein
Uncharacterized protein (Fragment)
Uncharacterized protein
Uncharacterized protein
Uncharacterized protein
Uncharacterized protein
Uncharacterized protein
Uncharacterized protein
Uncharacterized protein
Uncharacterized protein
Uncharacterized protein
Uncharacterized protein (Fragment)
Uncharacterized protein
Uncharacterized protein
Uncharacterized protein
Uncharacterized protein
Uncharacterized protein
Uncharacterized protein
Uncharacterized protein (Fragment)
Uncharacterized protein
Uncharacterized protein
Uncharacterized protein
Uncharacterized protein
Uncharacterized protein
Uncharacterized protein
Uncharacterized protein
Uncharacterized protein
Uncharacterized protein
Uncharacterized protein
Uncharacterized protein
Uncharacterized protein (Fragment)
Uncharacterized protein (Fragment)
Uncharacterized protein

Uncharacterized protein (Fragment)
Uncharacterized protein
Uncharacterized protein
Uncharacterized protein (Fragment)
Uncharacterized protein
Uncharacterized protein
Uncharacterized protein
Uncharacterized protein
Uncharacterized protein
Uncharacterized protein
Uncharacterized protein
Uncharacterized protein (Fragment)
Uncharacterized protein
Uncharacterized protein
Uncharacterized protein
Uncharacterized protein
Uncharacterized protein (Fragment)
Uncharacterized protein
Uncharacterized protein (Fragment)
Uncharacterized protein
Uncharacterized protein
Uncharacterized protein (Fragment)
Uncharacterized protein
Uncharacterized protein
Uncharacterized protein
Uncharacterized protein
Uncharacterized protein (Fragment)
Uncharacterized protein (Fragment)
Uncharacterized protein
Uncharacterized protein (Fragment)
Uncharacterized protein
Uncharacterized protein
Uncharacterized protein
Uncharacterized protein
Uncharacterized protein
Uncharacterized protein
Uncharacterized protein (Fragment)
Uncharacterized protein
Uncharacterized protein (Fragment)
Uncharacterized protein
Uncharacterized protein

Uncharacterized protein
Uncharacterized protein
Uncharacterized protein
Uncharacterized protein
Uncharacterized protein
Uncharacterized protein (Fragment)
Uncharacterized protein
Uncharacterized protein (Fragment)
Uncharacterized protein
Uncharacterized protein
Uncharacterized protein (Fragment)
Uncharacterized protein (Fragment)
Uncharacterized protein (Fragment)
Uncharacterized protein
Uncharacterized protein
Uncharacterized protein
Uncharacterized protein
Uncharacterized protein
Uncharacterized protein (Fragment)
Uncharacterized protein
Uncharacterized protein
Uncharacterized protein (Fragment)
Uncharacterized protein (Fragment)
Uncharacterized protein
Uncharacterized protein
Uncharacterized protein
CBEL protein, formerly GP34
Uncharacterized protein
Uncharacterized protein
Uncharacterized protein
Uncharacterized protein
Uncharacterized protein (Fragment)
Uncharacterized protein
Uncharacterized protein
Uncharacterized protein
Uncharacterized protein (Fragment)
Uncharacterized protein (Fragment)
Uncharacterized protein
Uncharacterized protein
Uncharacterized protein
Uncharacterized protein

Uncharacterized protein
Uncharacterized protein (Fragment)
Uncharacterized protein
Uncharacterized protein
Uncharacterized protein
Uncharacterized protein
Uncharacterized protein
Uncharacterized protein (Fragment)
Uncharacterized protein (Fragment)
Uncharacterized protein
Uncharacterized protein
Uncharacterized protein
Uncharacterized protein
Uncharacterized protein
Uncharacterized protein
Uncharacterized protein
Uncharacterized protein
Uncharacterized protein
Uncharacterized protein
Uncharacterized protein
Uncharacterized protein
Uncharacterized protein
Uncharacterized protein
Uncharacterized protein
Uncharacterized protein
Uncharacterized protein
Uncharacterized protein
Uncharacterized protein
Cellulose binding elicitor lectin (Fragment)
Cellulose binding elicitor lectin (Fragment)
Cellulose binding elicitor lectin (Fragment)
Uncharacterized protein
Cellulose binding elicitor lectin (Fragment)
Uncharacterized protein
Uncharacterized protein
Uncharacterized protein
Cellulose binding elicitor lectin
Uncharacterized protein
Cellulose binding elicitor lectin (Fragment)
Uncharacterized protein
Cellulose binding elicitor lectin (Fragment)

Uncharacterized protein
Uncharacterized protein
Uncharacterized protein
Uncharacterized protein
Uncharacterized protein
Cellulose binding elicitor lectin (Fragment)
Uncharacterized protein
Cellulose binding elicitor lectin (Fragment)
Cellulose binding elicitor lectin (Fragment)
Cellulose binding elicitor lectin (Fragment)
Cellulose binding elicitor lectin (Fragment)
Uncharacterized protein
Cellulose binding elicitor lectin (Fragment)
Cellulose binding elicitor lectin (Fragment)
Cellulose binding elicitor lectin (Fragment)
Uncharacterized protein
Uncharacterized protein
Putative glycoside hydrolase
Putative CBEL-like protein
Uncharacterized protein
Uncharacterized protein
Uncharacterized protein
Uncharacterized protein
Uncharacterized protein
CBEL-like protein
Uncharacterized protein
Uncharacterized protein
Uncharacterized protein
Uncharacterized protein
Putative glycosyl hydrolase family 5 protein
Uncharacterized protein
Uncharacterized protein
Uncharacterized protein
Uncharacterized protein
Uncharacterized protein
Uncharacterized protein
Uncharacterized protein
Uncharacterized protein (Fragment)
Uncharacterized protein
Uncharacterized protein
Uncharacterized protein (Fragment)

Uncharacterized protein
Uncharacterized protein (Fragment)
Uncharacterized protein
Uncharacterized protein
Uncharacterized protein
Hepatocyte growth factor (Fragment)
Uncharacterized protein (Fragment)
Uncharacterized protein
Uncharacterized protein
Uncharacterized protein (Fragment)
Uncharacterized protein
Endochitinase-like
Efflux ABC transporter permease/ATP-binding protein
Endo-1,3(4)-beta-glucanase
Uncharacterized protein
Uncharacterized protein
Putative carbohydrate binding domain
Oryzain alpha chain-like
Endo-1,3(4)-beta-glucanase
Uncharacterized protein
Uncharacterized protein
Uncharacterized protein
Perforin-1-like
Putative glucan 1,3-beta-glucosidase
Glycoside hydrolase family 43 protein
Cathepsin L
Uncharacterized protein
Cathepsin L
Endo-1,3(4)-beta-glucanase
Glycoside hydrolase family 76 protein
Uncharacterized protein
Bactericidal permeability-increasing protein-like
Uncharacterized protein
Uncharacterized protein
Uncharacterized protein
Uncharacterized protein
Uncharacterized protein
Uncharacterized protein (Fragment)
Cell wall surface anchored protein
Uncharacterized protein
Glucan endo-1,3-alpha-glucosidase Agn1

Uncharacterized protein (Fragment)
Uncharacterized protein
Cathepsin L
Uncharacterized protein
Uncharacterized protein
Uncharacterized protein
Uncharacterized protein
Putative chitosanase
PAN domain-containing protein
MAP/microtubule affinity-regulating kinase 3-like
Putative glucan 1,3-beta-glucosidase
Amino acid transporter
Putative glucan 1,3-beta-glucosidase
Uncharacterized protein
Putative glucan 1,3-beta-glucosidase
Uncharacterized protein
Pathogenesis-related protein PRMS-like
Putative glucan 1,3-beta-glucosidase
3-carboxymuconate cyclase
Uncharacterized protein
Putative chitosanase
Cathepsin L
Uncharacterized protein
Chaperonin containing TCP1 epsilon subunit
Putative glucan 1,3-beta-glucosidase
Esterase, SGNH hydrolase-type
Uncharacterized protein
Uncharacterized protein
Uncharacterized protein
PAN-1 domain
Apple-like
Histone acetyltransferase
Skin mucus lectin
Skin mucus lectin
Matrix metalloproteinase 1_14 (Fragment)
Alpha-2-macroglobulin
Alpha-2-macroglobulin
Uncharacterized protein
PAN domain protein
WNT and FGF inhibitory regulator domain-containing protein
PAN domain-containing protein

Uncharacterized protein (Fragment)
Uncharacterized protein
Uncharacterized protein
Uncharacterized protein (Fragment)
Uncharacterized protein
Uncharacterized protein
Uncharacterized protein
Uncharacterized protein
Uncharacterized protein
Uncharacterized protein (Fragment)
Uncharacterized protein
Uncharacterized protein
Uncharacterized protein
Uncharacterized protein (Fragment)
Uncharacterized protein
Uncharacterized protein
Uncharacterized protein (Fragment)
Uncharacterized protein
Uncharacterized protein (Fragment)
Uncharacterized protein
Uncharacterized protein
Uncharacterized protein
Uncharacterized protein
Uncharacterized protein
Uncharacterized protein (Fragment)
Uncharacterized protein (Fragment)
Uncharacterized protein
Uncharacterized protein
Uncharacterized protein (Fragment)
Uncharacterized protein
Uncharacterized protein
Uncharacterized protein
Uncharacterized protein
Uncharacterized protein
Uncharacterized protein
Uncharacterized protein
Uncharacterized protein
Uncharacterized protein
Uncharacterized protein
Uncharacterized protein (Fragment)
Peptidylprolyl isomerase

Uncharacterized protein
Uncharacterized protein (Fragment)
Uncharacterized protein
Uncharacterized protein (Fragment)
Uncharacterized protein
Uncharacterized protein
Uncharacterized protein
Uncharacterized protein
Uncharacterized protein
Uncharacterized protein
Uncharacterized protein (Fragment)
Uncharacterized protein (Fragment)
Uncharacterized protein
Uncharacterized protein (Fragment)
Uncharacterized protein
Uncharacterized protein (Fragment)
Uncharacterized protein
Uncharacterized protein
Uncharacterized protein
Uncharacterized protein
Uncharacterized protein
Uncharacterized protein
Uncharacterized protein
Uncharacterized protein
Uncharacterized protein
Uncharacterized protein
Uncharacterized protein
Uncharacterized protein
Uncharacterized protein
Uncharacterized protein
Uncharacterized protein
Uncharacterized protein
Uncharacterized protein
Uncharacterized protein (Fragment)
Uncharacterized protein (Fragment)
Uncharacterized protein
Uncharacterized protein
Uncharacterized protein (Fragment)
Uncharacterized protein (Fragment)
Uncharacterized protein
Uncharacterized protein (Fragment)

Uncharacterized protein
Uncharacterized protein
Uncharacterized protein
Uncharacterized protein
Uncharacterized protein
Uncharacterized protein (Fragment)
Uncharacterized protein
Uncharacterized protein
Uncharacterized protein
Uncharacterized protein
Uncharacterized protein
Uncharacterized protein
Uncharacterized protein (Fragment)
Uncharacterized protein
Uncharacterized protein (Fragment)
Uncharacterized protein
Uncharacterized protein
Uncharacterized protein
Uncharacterized protein
Uncharacterized protein (Fragment)
Uncharacterized protein
Uncharacterized protein
Uncharacterized protein
Uncharacterized protein
Uncharacterized protein
Uncharacterized protein (Fragment)
Uncharacterized protein (Fragment)
Uncharacterized protein (Fragment)
Uncharacterized protein
Uncharacterized protein
Uncharacterized protein
Uncharacterized protein
Uncharacterized protein
Uncharacterized protein
Uncharacterized protein
Uncharacterized protein
Uncharacterized protein
Uncharacterized protein
Uncharacterized protein (Fragment)
Uncharacterized protein (Fragment)
Uncharacterized protein

Uncharacterized protein
Uncharacterized protein
Uncharacterized protein
Uncharacterized protein
Uncharacterized protein
Uncharacterized protein
Uncharacterized protein (Fragment)
Uncharacterized protein (Fragment)
Uncharacterized protein (Fragment)
Uncharacterized protein
Uncharacterized protein (Fragment)
Uncharacterized protein
Uncharacterized protein
Uncharacterized protein (Fragment)
Uncharacterized protein
Uncharacterized protein
Uncharacterized protein
Uncharacterized protein (Fragment)
Uncharacterized protein
Uncharacterized protein (Fragment)
Uncharacterized protein (Fragment)
Uncharacterized protein
Uncharacterized protein
Uncharacterized protein (Fragment)
Uncharacterized protein
Uncharacterized protein
Uncharacterized protein
Uncharacterized protein
Uncharacterized protein
Uncharacterized protein (Fragment)
Uncharacterized protein
Uncharacterized protein
Uncharacterized protein
Uncharacterized protein
Uncharacterized protein
Uncharacterized protein
Uncharacterized protein
Uncharacterized protein
Uncharacterized protein
Uncharacterized protein (Fragment)
Uncharacterized protein

Uncharacterized protein
Uncharacterized protein
Uncharacterized protein
Uncharacterized protein
Uncharacterized protein
Uncharacterized protein
Uncharacterized protein
Uncharacterized protein (Fragment)
Uncharacterized protein (Fragment)
Uncharacterized protein (Fragment)
Uncharacterized protein
Uncharacterized protein
Uncharacterized protein (Fragment)
Uncharacterized protein
Uncharacterized protein
Uncharacterized protein
Uncharacterized protein
Uncharacterized protein
Uncharacterized protein
Uncharacterized protein (Fragment)
Uncharacterized protein (Fragment)
Uncharacterized protein
Uncharacterized protein
Uncharacterized protein
Uncharacterized protein
Uncharacterized protein (Fragment)
Uncharacterized protein (Fragment)
Uncharacterized protein
Uncharacterized protein
Uncharacterized protein
Uncharacterized protein
Uncharacterized protein
Uncharacterized protein
Uncharacterized protein
Uncharacterized protein
Uncharacterized protein
Uncharacterized protein
Uncharacterized protein
Uncharacterized protein
Uncharacterized protein
Uncharacterized protein

Uncharacterized protein
Uncharacterized protein (Fragment)
Uncharacterized protein
Uncharacterized protein
Uncharacterized protein
Uncharacterized protein
Uncharacterized protein (Fragment)
Uncharacterized protein (Fragment)
Uncharacterized protein
Uncharacterized protein
Uncharacterized protein (Fragment)
Uncharacterized protein
Uncharacterized protein
Uncharacterized protein
Uncharacterized protein (Fragment)
Uncharacterized protein (Fragment)
Uncharacterized protein (Fragment)
Uncharacterized protein (Fragment)
Uncharacterized protein
Uncharacterized protein
Uncharacterized protein
Uncharacterized protein
Uncharacterized protein
Uncharacterized protein (Fragment)
Uncharacterized protein
Uncharacterized protein (Fragment)
Uncharacterized protein
Uncharacterized protein
Plasminogen (Fragment)
Coagulation factor XI (Fragment)
Hepatocyte growth factor (Fragment)
Plasminogen (Fragment)
Podospora anserina S mat+ genomic DNA chromosome 6, supercontig 2
Podospora anserina S mat+ genomic DNA chromosome 5, supercontig 1
Podospora anserina S mat+ genomic DNA chromosome 6, supercontig 1
Podospora anserina S mat+ genomic DNA chromosome 2, supercontig 2 (Fragment)
Putative Glycosyltransferase Family 31
Uncharacterized protein
Glycosyltransferase Family 31
Glycosyltransferase Family 31
Uncharacterized protein

Uncharacterized protein
Uncharacterized protein
Uncharacterized protein
Uncharacterized protein
Plasminogen
Plasminogen
Hepatocyte growth factor
Uncharacterized protein
Uncharacterized protein
Uncharacterized protein
Uncharacterized protein
Uncharacterized protein
Uncharacterized protein
Uncharacterized protein
Hepatocyte growth factor
Uncharacterized protein
Uncharacterized protein
Uncharacterized protein
Plasminogen
Plasminogen
Uncharacterized protein
Plasminogen
Uncharacterized protein
Plasminogen
Uncharacterized protein
Uncharacterized protein
Uncharacterized protein
Uncharacterized protein
Uncharacterized protein
Plasminogen
Uncharacterized protein
Uncharacterized protein
Hepatocyte growth factor
Uncharacterized protein
Uncharacterized protein
Uncharacterized protein
Uncharacterized protein
Uncharacterized protein
Uncharacterized protein
Uncharacterized protein
Uncharacterized protein

Uncharacterized protein
Uncharacterized protein
Uncharacterized protein
Uncharacterized protein
Uncharacterized protein
Uncharacterized protein
Hepatocyte growth factor
Uncharacterized protein
KLKB1 (Fragment)
KLKB1 (Fragment)
KLKB1 (Fragment)
Plasminogen
KLKB1 (Fragment)
KLKB1 (Fragment)
HGFL
KLKB1 (Fragment)
KLKB1 (Fragment)
KLKB1 (Fragment)
HGF (Fragment)
KLKB1 (Fragment)
KLKB1
G-type lectin-domain containing receptor kinase 4
G-type lectin-domain containing receptor kinase 6
Tunicate retinoic acid-inducible modular protease
Alpha-2-macroglobulin N-terminal domain family protein
Uncharacterized protein
Uncharacterized protein
Uncharacterized protein
Uncharacterized protein
Uncharacterized protein
Uncharacterized protein
Uncharacterized protein
Uncharacterized protein
Uncharacterized protein
Uncharacterized protein
Uncharacterized protein
Uncharacterized protein
Uncharacterized protein
Uncharacterized protein
Uncharacterized protein
Uncharacterized protein

Uncharacterized protein
Uncharacterized protein
Uncharacterized protein
Uncharacterized protein
Uncharacterized protein
Uncharacterized protein
Uncharacterized protein
Uncharacterized protein
Uncharacterized protein
Uncharacterized protein
Uncharacterized protein
Uncharacterized protein
Uncharacterized protein
Uncharacterized protein
Uncharacterized protein
Uncharacterized protein
Uncharacterized protein
Uncharacterized protein
Uncharacterized protein
Uncharacterized protein
Uncharacterized protein (Fragment)
Uncharacterized protein
Metalloendopeptidase
Uncharacterized protein
PLGLB1 isoform 1
Plasminogen
Hepatocyte growth factor
HGF isoform 3
KLKB1 isoform 4 (Fragment)
HGF isoform 4
F11 isoform 3
Plasminogen
KLKB1 isoform 1 (Fragment)
HGF isoform 10 (Fragment)
MST1 isoform 2
KLKB1 isoform 5
Macrophage stimulating 1
KLKB1 isoform 3
HGF isoform 5
HGF isoform 7
MST1 isoform 1

Uncharacterized protein
HGF isoform 9 (Fragment)
KLKB1 isoform 2 (Fragment)
F11 isoform 2
Uncharacterized protein DKFZp459G0539
F11 isoform 5 (Fragment)
Uncharacterized protein DKFZp459J2015 (Fragment)
F11 isoform 1
Hepatocyte growth factor
PLGLB1 isoform 2 (Fragment)
Plasminogen
Uncharacterized protein
Uncharacterized protein
Uncharacterized protein
Uncharacterized protein
Uncharacterized protein
PAN domain-containing protein
Uncharacterized protein
Uncharacterized protein
Uncharacterized protein
S-locus lectin protein kinase family protein
G-type lectin S-receptor-like serine/threonine-protein kinase (Fragment)
S-locus lectin protein kinase family protein
S-locus lectin protein kinase family protein
S-locus lectin protein kinase family protein
G-type lectin S-receptor-like serine/threonine-protein kinase (Fragment)
G-type lectin S-receptor-like serine/threonine-protein kinase (Fragment)
S-locus lectin protein kinase family protein B120
G-type lectin S-receptor-like serine/threonine-protein kinase (Fragment)
Uncharacterized protein
G-type lectin S-receptor-like serine/threonine-protein kinase (Fragment)
G-type lectin S-receptor-like serine/threonine-protein kinase (Fragment)
G-type lectin S-receptor-like serine/threonine-protein kinase (Fragment)
G-type lectin S-receptor-like serine/threonine-protein kinase (Fragment)
G-type lectin S-receptor-like serine/threonine-protein kinase (Fragment)
G-type lectin S-receptor-like serine/threonine-protein kinase (Fragment)
G-type lectin S-receptor-like serine/threonine-protein kinase (Fragment)
Uncharacterized protein
Uncharacterized protein
Uncharacterized protein
Uncharacterized protein

Uncharacterized protein
Uncharacterized protein
Uncharacterized protein
Uncharacterized protein
Uncharacterized protein
Uncharacterized protein
Uncharacterized protein
Uncharacterized protein
Uncharacterized protein
Uncharacterized protein
Uncharacterized protein
Uncharacterized protein
Uncharacterized protein (Fragment)
Uncharacterized protein
Uncharacterized protein (Fragment)
Uncharacterized protein
Uncharacterized protein
Uncharacterized protein
Uncharacterized protein
Uncharacterized protein
Uncharacterized protein
Uncharacterized protein
Uncharacterized protein
Uncharacterized protein
Uncharacterized protein
Uncharacterized protein
Uncharacterized protein (Fragment)
Uncharacterized protein
Uncharacterized protein
Uncharacterized protein
Uncharacterized protein
Uncharacterized protein
Uncharacterized protein
Uncharacterized protein
Uncharacterized protein
Uncharacterized protein
Uncharacterized protein
Uncharacterized protein
Uncharacterized protein (Fragment)
Uncharacterized protein
Uncharacterized protein

Uncharacterized protein
Uncharacterized protein
Uncharacterized protein
Uncharacterized protein
Uncharacterized protein
Uncharacterized protein (Fragment)
Uncharacterized protein
Uncharacterized protein
Uncharacterized protein
Uncharacterized protein
Uncharacterized protein
Uncharacterized protein
Uncharacterized protein (Fragment)
Uncharacterized protein
Uncharacterized protein
Uncharacterized protein
Uncharacterized protein
Uncharacterized protein (Fragment)
Uncharacterized protein
Uncharacterized protein
Uncharacterized protein
Uncharacterized protein
Uncharacterized protein
Uncharacterized protein (Fragment)
Uncharacterized protein
Uncharacterized protein
Uncharacterized protein
Uncharacterized protein
Uncharacterized protein
Uncharacterized protein
Uncharacterized protein
Uncharacterized protein
Uncharacterized protein
Uncharacterized protein
Uncharacterized protein
Uncharacterized protein
Uncharacterized protein
Uncharacterized protein
Uncharacterized protein (Fragment)
Uncharacterized protein (Fragment)
Uncharacterized protein
Uncharacterized protein

Uncharacterized protein
Uncharacterized protein
Uncharacterized protein
Uncharacterized protein (Fragment)
Uncharacterized protein
Uncharacterized protein
Uncharacterized protein
Uncharacterized protein
Uncharacterized protein
Uncharacterized protein (Fragment)
Uncharacterized protein
Uncharacterized protein
Uncharacterized protein (Fragment)
Uncharacterized protein
Uncharacterized protein
Uncharacterized protein
Uncharacterized protein
Uncharacterized protein
Uncharacterized protein
Uncharacterized protein
Uncharacterized protein
Uncharacterized protein
Uncharacterized protein
Uncharacterized protein
Uncharacterized protein (Fragment)
Uncharacterized protein
Uncharacterized protein
Uncharacterized protein
Uncharacterized protein
Uncharacterized protein
Uncharacterized protein
Uncharacterized protein
Uncharacterized protein
Uncharacterized protein
Uncharacterized protein
Uncharacterized protein
Uncharacterized protein
Uncharacterized protein
Uncharacterized protein
Uncharacterized protein
Uncharacterized protein

Uncharacterized protein (Fragment)
Uncharacterized protein
Uncharacterized protein
Uncharacterized protein
Uncharacterized protein
Uncharacterized protein
Uncharacterized protein
Uncharacterized protein
Uncharacterized protein
Uncharacterized protein
Uncharacterized protein
Uncharacterized protein
Uncharacterized protein
Uncharacterized protein
Uncharacterized protein
Uncharacterized protein
Uncharacterized protein
Uncharacterized protein
Uncharacterized protein
Uncharacterized protein
Uncharacterized protein
Uncharacterized protein
Uncharacterized protein
Uncharacterized protein
Uncharacterized protein
Uncharacterized protein
Uncharacterized protein
Uncharacterized protein
Uncharacterized protein
Uncharacterized protein
Uncharacterized protein
Uncharacterized protein
Uncharacterized protein
Uncharacterized protein
Uncharacterized protein
Uncharacterized protein
Uncharacterized protein
Uncharacterized protein
Uncharacterized protein
Uncharacterized protein
Uncharacterized protein

Uncharacterized protein
Uncharacterized protein
Uncharacterized protein
Uncharacterized protein
Uncharacterized protein
Uncharacterized protein
Uncharacterized protein
Uncharacterized protein
Uncharacterized protein
Uncharacterized protein
Uncharacterized protein
Uncharacterized protein
Uncharacterized protein
Uncharacterized protein
Uncharacterized protein
Uncharacterized protein
Uncharacterized protein
Uncharacterized protein (Fragment)
Uncharacterized protein
Uncharacterized protein
Uncharacterized protein
Uncharacterized protein
Uncharacterized protein
Uncharacterized protein
Uncharacterized protein
Uncharacterized protein
Uncharacterized protein
Uncharacterized protein
Uncharacterized protein
Uncharacterized protein
Uncharacterized protein
CCHC-type integrase
Uncharacterized protein
Uncharacterized protein
Uncharacterized protein
Uncharacterized protein
Uncharacterized protein
Uncharacterized protein
Uncharacterized protein
Uncharacterized protein
Uncharacterized protein

Uncharacterized protein
Uncharacterized protein
Uncharacterized protein
Uncharacterized protein
Uncharacterized protein
Uncharacterized protein
Uncharacterized protein
Uncharacterized protein
Uncharacterized protein
Uncharacterized protein
Uncharacterized protein
Uncharacterized protein (Fragment)
Uncharacterized protein
Uncharacterized protein
Uncharacterized protein
Uncharacterized protein
Uncharacterized protein
Uncharacterized protein
Uncharacterized protein
Uncharacterized protein
Uncharacterized protein
Uncharacterized protein
Uncharacterized protein (Fragment)
Uncharacterized protein
Uncharacterized protein
Uncharacterized protein
Uncharacterized protein
Uncharacterized protein
Uncharacterized protein
Uncharacterized protein
Uncharacterized protein
Uncharacterized protein
Uncharacterized protein
Uncharacterized protein
Uncharacterized protein
Uncharacterized protein
Uncharacterized protein
Uncharacterized protein
Uncharacterized protein
Uncharacterized protein
Uncharacterized protein

Uncharacterized protein
Uncharacterized protein
Uncharacterized protein
Uncharacterized protein
Uncharacterized protein
Uncharacterized protein
Uncharacterized protein
Putative pan domain-containing protein
Uncharacterized protein
Putative pan domain-containing protein
Putative lipoprotein YfhM
Uncharacterized protein
Uncharacterized protein
Uncharacterized protein
Uncharacterized protein
Uncharacterized protein
Uncharacterized protein
Uncharacterized protein
Uncharacterized protein
Cutl-17
Uncharacterized protein
Uncharacterized protein
Uncharacterized protein
Uncharacterized protein (Fragment)
Uncharacterized protein
Uncharacterized protein
Uncharacterized protein
Uncharacterized protein
Uncharacterized protein
Uncharacterized protein
Uncharacterized protein
Uncharacterized protein
Uncharacterized protein
Uncharacterized protein
Uncharacterized protein
Uncharacterized protein
Uncharacterized protein
Uncharacterized protein
Uncharacterized protein
Uncharacterized protein
Uncharacterized protein

Emb-4
Uncharacterized protein
Uncharacterized protein
Uncharacterized protein
Uncharacterized protein
Uncharacterized protein
Uncharacterized protein
Uncharacterized protein
Uncharacterized protein
Uncharacterized protein (Fragment)
Uncharacterized protein
Uncharacterized protein
Uncharacterized protein
Uncharacterized protein
Uncharacterized protein
Uncharacterized protein
Uncharacterized protein
Uncharacterized protein (Fragment)
Uncharacterized protein
Uncharacterized protein
Uncharacterized protein
Uncharacterized protein
Uncharacterized protein
Cutl-18
Uncharacterized protein (Fragment)
Uncharacterized protein
Uncharacterized protein
Uncharacterized protein
Uncharacterized protein
Uncharacterized protein
Uncharacterized protein
Uncharacterized protein
Uncharacterized protein (Fragment)
Uncharacterized protein
Uncharacterized protein (Fragment)
Mspn-1 (Fragment)
Uncharacterized protein
Uncharacterized protein
Uncharacterized protein
Uncharacterized protein (Fragment)
Uncharacterized protein

Uncharacterized protein
Uncharacterized protein
Uncharacterized protein
Uncharacterized protein
Uncharacterized protein
Uncharacterized protein
Uncharacterized protein
Uncharacterized protein
Zinc finger protein
Uncharacterized protein
Uncharacterized protein
Uncharacterized protein
Uncharacterized protein
Uncharacterized protein
Proliferating cell nuclear antigen
Uncharacterized protein
Uncharacterized protein
Uncharacterized protein (Fragment)
Uncharacterized protein
Serine/threonine-protein phosphatase
Uncharacterized protein
Uncharacterized protein
Srap-1
Uncharacterized protein
Uncharacterized protein
Uncharacterized protein
Uncharacterized protein
Uncharacterized protein
Uncharacterized protein
Major sperm protein
Serpentine receptor class gamma (Fragment)
Uncharacterized protein
Uncharacterized protein
Uncharacterized protein
Protein kinase (Fragment)
Noah-2
Uncharacterized protein
Uncharacterized protein
Uncharacterized protein
Hepatocyte growth factor
Plasminogen

Hepatocyte growth factor
Uncharacterized protein
Uncharacterized protein
Uncharacterized protein
Uncharacterized protein
Apple domain-containing protein
Uncharacterized protein
Uncharacterized protein (Fragment)
Uncharacterized protein
Apple domain-containing protein
Apple domain-containing protein
Uncharacterized protein
Uncharacterized protein
Protein kinase
Protein kinase
Uncharacterized protein (Fragment)
Uncharacterized protein
Uncharacterized protein
Uncharacterized protein
Uncharacterized protein
Uncharacterized protein (Fragment)
Uncharacterized protein
Uncharacterized protein
Uncharacterized protein (Fragment)
Uncharacterized protein
Uncharacterized protein
Uncharacterized protein
Uncharacterized protein (Fragment)
Uncharacterized protein
Uncharacterized protein
Uncharacterized protein (Fragment)
S-locus-like receptor protein kinase (Fragment)
Uncharacterized protein
Uncharacterized protein (Fragment)
Uncharacterized protein
Uncharacterized protein
Uncharacterized protein
Uncharacterized protein
Uncharacterized protein
Uncharacterized protein (Fragment)
Uncharacterized protein

Uncharacterized protein (Fragment)
Uncharacterized protein
Uncharacterized protein
Uncharacterized protein
Uncharacterized protein
Uncharacterized protein
Uncharacterized protein
Uncharacterized protein
Uncharacterized protein
Uncharacterized protein (Fragment)
Uncharacterized protein
Uncharacterized protein
Uncharacterized protein
Uncharacterized protein
Uncharacterized protein
Uncharacterized protein
Uncharacterized protein
Uncharacterized protein
Uncharacterized protein
Uncharacterized protein
Uncharacterized protein
Uncharacterized protein (Fragment)
Uncharacterized protein (Fragment)
Uncharacterized protein
Uncharacterized protein
Uncharacterized protein
Uncharacterized protein
Uncharacterized protein (Fragment)
Uncharacterized protein
Uncharacterized protein
Uncharacterized protein
Uncharacterized protein (Fragment)
Uncharacterized protein
Uncharacterized protein
Uncharacterized protein
Uncharacterized protein
Uncharacterized protein (Fragment)
Uncharacterized protein
Uncharacterized protein
Uncharacterized protein
Uncharacterized protein

Uncharacterized protein
Uncharacterized protein
Uncharacterized protein
Uncharacterized protein
Uncharacterized protein
Uncharacterized protein
Uncharacterized protein
Uncharacterized protein
Uncharacterized protein
Uncharacterized protein
Uncharacterized protein
Uncharacterized protein
Uncharacterized protein (Fragment)
Uncharacterized protein (Fragment)
Uncharacterized protein
Uncharacterized protein
Uncharacterized protein
Uncharacterized protein
Uncharacterized protein
Uncharacterized protein
Uncharacterized protein
Uncharacterized protein
Uncharacterized protein
Uncharacterized protein
Uncharacterized protein (Fragment)
Uncharacterized protein
Uncharacterized protein
Uncharacterized protein
Uncharacterized protein
Uncharacterized protein
Uncharacterized protein
Uncharacterized protein (Fragment)
Uncharacterized protein
Uncharacterized protein
Uncharacterized protein
Uncharacterized protein
Uncharacterized protein
Uncharacterized protein
Uncharacterized protein
Uncharacterized protein (Fragment)
Uncharacterized protein

Uncharacterized protein
Uncharacterized protein
Uncharacterized protein
Uncharacterized protein
Uncharacterized protein
Uncharacterized protein
Uncharacterized protein
Uncharacterized protein
Uncharacterized protein
Uncharacterized protein
Uncharacterized protein
Uncharacterized protein
Uncharacterized protein
Uncharacterized protein
Uncharacterized protein
Uncharacterized protein
Uncharacterized protein
Uncharacterized protein
Uncharacterized protein
Uncharacterized protein
Uncharacterized protein
Uncharacterized protein
Uncharacterized protein
Uncharacterized protein
Uncharacterized protein
Uncharacterized protein
Uncharacterized protein
Uncharacterized protein
Uncharacterized protein
Uncharacterized protein
Uncharacterized protein
Uncharacterized protein
Uncharacterized protein
Uncharacterized protein
Uncharacterized protein
Uncharacterized protein (Fragment)
Uncharacterized protein
Uncharacterized protein
Uncharacterized protein (Fragment)
Uncharacterized protein
Uncharacterized protein

Uncharacterized protein
Uncharacterized protein
Uncharacterized protein (Fragment)
Uncharacterized protein
Uncharacterized protein (Fragment)
Uncharacterized protein (Fragment)
Uncharacterized protein
Uncharacterized protein
Uncharacterized protein
Uncharacterized protein
Uncharacterized protein
Uncharacterized protein
Uncharacterized protein
Uncharacterized protein
Uncharacterized protein (Fragment)
Uncharacterized protein
Uncharacterized protein
Uncharacterized protein
Uncharacterized protein (Fragment)
Uncharacterized protein
Uncharacterized protein
Uncharacterized protein
Uncharacterized protein (Fragment)
Uncharacterized protein
Uncharacterized protein
Uncharacterized protein
Uncharacterized protein
Uncharacterized protein
Uncharacterized protein
Uncharacterized protein
Uncharacterized protein
Uncharacterized protein
Uncharacterized protein
Uncharacterized protein
Uncharacterized protein
Uncharacterized protein (Fragment)
G-type lectin S-receptor-like serine/threonine-protein kinase
Uncharacterized protein
Uncharacterized protein
G-type lectin S-receptor-like serine/threonine-protein kinase
Uncharacterized protein

G-type lectin S-receptor-like serine/threonine-protein kinase
Putative receptor protein kinase ZmPK1
Uncharacterized protein
Uncharacterized protein
Receptor-like serine/threonine-protein kinase SD1-8
G-type lectin S-receptor-like serine/threonine-protein kinase
Uncharacterized protein
G-type lectin S-receptor-like serine/threonine-protein kinase
G-type lectin S-receptor-like serine/threonine-protein kinase
Uncharacterized protein
G-type lectin S-receptor-like serine/threonine-protein kinase
Uncharacterized protein
G-type lectin S-receptor-like serine/threonine-protein kinase
Uncharacterized protein
G-type lectin S-receptor-like serine/threonine-protein kinase
G-type lectin S-receptor-like serine/threonine-protein kinase
G-type lectin S-receptor-like serine/threonine-protein kinase SD2-5
Receptor-like serine/threonine-protein kinase SD1-8
G-type lectin S-receptor-like serine/threonine-protein kinase
G-type lectin S-receptor-like serine/threonine-protein kinase
G-type lectin S-receptor-like serine/threonine-protein kinase
G-type lectin S-receptor-like serine/threonine-protein kinase
Uncharacterized protein
Uncharacterized protein
G-type lectin S-receptor-like serine/threonine-protein kinase
Uncharacterized protein
G-type lectin S-receptor-like serine/threonine-protein kinase RLK1
G-type lectin S-receptor-like serine/threonine-protein kinase CES101
G-type lectin S-receptor-like serine/threonine-protein kinase SD2-2
Uncharacterized protein
G-type lectin S-receptor-like serine/threonine-protein kinase
G-type lectin S-receptor-like serine/threonine-protein kinase SD2-5
G-type lectin S-receptor-like serine/threonine-protein kinase
Uncharacterized protein
G-type lectin S-receptor-like serine/threonine-protein kinase SD2-5
G-type lectin S-receptor-like serine/threonine-protein kinase
G-type lectin S-receptor-like serine/threonine-protein kinase
Uncharacterized protein
G-type lectin S-receptor-like serine/threonine-protein kinase
G-type lectin S-receptor-like serine/threonine-protein kinase
G-type lectin S-receptor-like serine/threonine-protein kinase

G-type lectin S-receptor-like serine/threonine-protein kinase
G-type lectin S-receptor-like serine/threonine-protein kinase
Receptor-like serine/threonine-protein kinase SD1-8
PAN domain-containing protein
Uncharacterized protein
G-type lectin S-receptor-like serine/threonine-protein kinase
G-type lectin S-receptor-like serine/threonine-protein kinase
G-type lectin S-receptor-like serine/threonine-protein kinase LECRK1
G-type lectin S-receptor-like serine/threonine-protein kinase
G-type lectin S-receptor-like serine/threonine-protein kinase
G-type lectin S-receptor-like serine/threonine-protein kinase CES101
G-type lectin S-receptor-like serine/threonine-protein kinase
Uncharacterized protein
G-type lectin S-receptor-like serine/threonine-protein kinase
G-type lectin S-receptor-like serine/threonine-protein kinase CES101 isoform X2
Uncharacterized protein
Uncharacterized protein
G-type lectin S-receptor-like serine/threonine-protein kinase
G-type lectin S-receptor-like serine/threonine-protein kinase
G-type lectin S-receptor-like serine/threonine-protein kinase
Receptor-like serine/threonine-protein kinase SD1-8
G-type lectin S-receptor-like serine/threonine-protein kinase
Putative receptor protein kinase ZmPK1
G-type lectin S-receptor-like serine/threonine-protein kinase
G-type lectin S-receptor-like serine/threonine-protein kinase
Putative receptor protein kinase ZmPK1
G-type lectin S-receptor-like serine/threonine-protein kinase
G-type lectin S-receptor-like serine/threonine-protein kinase SD1-29 isoform X4
G-type lectin S-receptor-like serine/threonine-protein kinase
Uncharacterized protein
G-type lectin S-receptor-like serine/threonine-protein kinase
Uncharacterized protein
Putative receptor protein kinase ZmPK1
G-type lectin S-receptor-like serine/threonine-protein kinase LECRK3
G-type lectin S-receptor-like serine/threonine-protein kinase
G-type lectin S-receptor-like serine/threonine-protein kinase
Uncharacterized protein
G-type lectin S-receptor-like serine/threonine-protein kinase
G-type lectin S-receptor-like serine/threonine-protein kinase
G-type lectin S-receptor-like serine/threonine-protein kinase B120
G-type lectin S-receptor-like serine/threonine-protein kinase

G-type lectin S-receptor-like serine/threonine-protein kinase
Uncharacterized protein
G-type lectin S-receptor-like serine/threonine-protein kinase
G-type lectin S-receptor-like serine/threonine-protein kinase RLK1
Uncharacterized protein
G-type lectin S-receptor-like serine/threonine-protein kinase SD2-5
G-type lectin S-receptor-like serine/threonine-protein kinase RKS1 isoform X1
Uncharacterized protein
PAN domain-containing protein
Uncharacterized protein
PAN domain-containing protein
Uncharacterized protein
Uncharacterized protein
Uncharacterized protein
Uncharacterized protein
Uncharacterized protein
Uncharacterized protein
Uncharacterized protein
Uncharacterized protein
Uncharacterized protein
Uncharacterized protein
Uncharacterized protein
PAN domain-containing protein
PAN domain-containing protein
PAN domain-containing protein
Uncharacterized protein
Uncharacterized protein
Uncharacterized protein
Uncharacterized protein
Uncharacterized protein
Uncharacterized protein
Uncharacterized protein
Uncharacterized protein (Fragment)
Uncharacterized protein
Uncharacterized protein (Fragment)
Uncharacterized protein
Uncharacterized protein
Uncharacterized protein
Uncharacterized protein
Uncharacterized protein
Uncharacterized protein

Uncharacterized protein
Uncharacterized protein
Uncharacterized protein
Uncharacterized protein
Uncharacterized protein
Uncharacterized protein
Uncharacterized protein
Insulin-like growth factor binding protein, N-terminal
Uncharacterized protein
Uncharacterized protein
Uncharacterized protein
Uncharacterized protein
Uncharacterized protein
Uncharacterized protein
Uncharacterized protein
Uncharacterized protein
Uncharacterized protein
Uncharacterized protein
Uncharacterized protein
Uncharacterized protein
Uncharacterized protein
Uncharacterized protein
Uncharacterized protein
Uncharacterized protein
Uncharacterized protein
Uncharacterized protein
Uncharacterized protein
Uncharacterized protein
Uncharacterized protein
Uncharacterized protein
Uncharacterized protein
Uncharacterized protein
Uncharacterized protein
Uncharacterized protein
Uncharacterized protein
Uncharacterized protein
Uncharacterized protein
Uncharacterized protein
Uncharacterized protein
Uncharacterized protein
Uncharacterized protein

Uncharacterized protein
Uncharacterized protein
Uncharacterized protein
Uncharacterized protein
Uncharacterized protein
Uncharacterized protein
Uncharacterized protein
Uncharacterized protein
Uncharacterized protein
Uncharacterized protein
Uncharacterized protein
Uncharacterized protein
Uncharacterized protein
Uncharacterized protein
Uncharacterized protein
Uncharacterized protein
Uncharacterized protein
Uncharacterized protein
Uncharacterized protein
Uncharacterized protein
Uncharacterized protein
Uncharacterized protein
Uncharacterized protein
Uncharacterized protein
Uncharacterized protein
Uncharacterized protein
Uncharacterized protein
Uncharacterized protein
Uncharacterized protein
Uncharacterized protein
Uncharacterized protein
Uncharacterized protein
Uncharacterized protein
Uncharacterized protein
Uncharacterized protein
Uncharacterized protein
Uncharacterized protein
Uncharacterized protein
Uncharacterized protein (Fragment)
Uncharacterized protein
Uncharacterized protein

Uncharacterized protein
Uncharacterized protein
Uncharacterized protein
Uncharacterized protein (Fragment)
Uncharacterized protein
Uncharacterized protein
Uncharacterized protein
Uncharacterized protein (Fragment)
Uncharacterized protein
Uncharacterized protein
Uncharacterized protein (Fragment)
Uncharacterized protein
Uncharacterized protein
Uncharacterized protein
Uncharacterized protein
Uncharacterized protein
Uncharacterized protein
Uncharacterized protein
Uncharacterized protein
Uncharacterized protein
Uncharacterized protein
Uncharacterized protein
Uncharacterized protein
Uncharacterized protein
Uncharacterized protein
Apple domain-containing protein
Apple domain-containing protein
Apple domain-containing protein
Uncharacterized protein
Uncharacterized protein
Uncharacterized protein
Uncharacterized protein
Uncharacterized protein
Uncharacterized protein
Uncharacterized protein
Galactose oxidase
PAN domain-containing protein
PAN domain-containing protein
Uncharacterized protein
PAN domain protein
Uncharacterized protein

Uncharacterized protein
Alpha-2-macroglobulin family protein
PAN domain-containing protein
PAN domain-containing protein
Uncharacterized protein
Membrane protein
Uncharacterized protein
Alpha-2-macroglobulin family protein
PAN domain-containing protein
PAN domain-containing protein
Apple domain-containing protein
Uncharacterized protein
Uncharacterized protein
Uncharacterized protein
Galactose oxidase
Plasminogen (Fragment)
Hepatocyte growth factor (Fragment)
Plasma kallikrein (Fragment)
Hepatocyte growth factor-like protein
Uncharacterized protein
Uncharacterized protein
Uncharacterized protein
Uncharacterized protein
Hepatocyte growth factor
Uncharacterized protein
Plasminogen
Uncharacterized protein
Uncharacterized protein
Uncharacterized protein (Fragment)
Uncharacterized protein
Uncharacterized protein
Uncharacterized protein
Uncharacterized protein
Uncharacterized protein
Uncharacterized protein
Uncharacterized protein
Uncharacterized protein (Fragment)
Uncharacterized protein
Uncharacterized protein
Uncharacterized protein
Uncharacterized protein

Uncharacterized protein
Uncharacterized protein
Uncharacterized protein
Uncharacterized protein
Uncharacterized protein
Uncharacterized protein
Uncharacterized protein
Uncharacterized protein
Uncharacterized protein (Fragment)
Uncharacterized protein
Uncharacterized protein
Uncharacterized protein (Fragment)
Uncharacterized protein
Uncharacterized protein
Uncharacterized protein (Fragment)
Uncharacterized protein
Uncharacterized protein
Uncharacterized protein
Uncharacterized protein
Uncharacterized protein
Uncharacterized protein
Uncharacterized protein
Uncharacterized protein (Fragment)
Uncharacterized protein
Uncharacterized protein
Uncharacterized protein (Fragment)
Uncharacterized protein
Uncharacterized protein
Uncharacterized protein
Uncharacterized protein
Uncharacterized protein
Uncharacterized protein
Uncharacterized protein
Uncharacterized protein
Uncharacterized protein
Uncharacterized protein (Fragment)
Uncharacterized protein (Fragment)
Uncharacterized protein (Fragment)
Uncharacterized protein
Uncharacterized protein
Uncharacterized protein

Uncharacterized protein
Uncharacterized protein
Uncharacterized protein
Uncharacterized protein
Uncharacterized protein
Uncharacterized protein
Uncharacterized protein
Uncharacterized protein
Uncharacterized protein
Uncharacterized protein
Uncharacterized protein
Uncharacterized protein
Uncharacterized protein
Uncharacterized protein
Uncharacterized protein
Uncharacterized protein
Uncharacterized protein
Uncharacterized protein
Uncharacterized protein
Uncharacterized protein
Uncharacterized protein
Uncharacterized protein
Uncharacterized protein
Uncharacterized protein
Uncharacterized protein (Fragment)
Uncharacterized protein
Uncharacterized protein
Uncharacterized protein (Fragment)
Uncharacterized protein
Uncharacterized protein
Uncharacterized protein
Uncharacterized protein (Fragment)
Uncharacterized protein
Uncharacterized protein (Fragment)
Uncharacterized protein (Fragment)
Uncharacterized protein
Uncharacterized protein
Uncharacterized protein
Uncharacterized protein
Uncharacterized protein
Uncharacterized protein

Uncharacterized protein
Uncharacterized protein
Uncharacterized protein
Uncharacterized protein
Uncharacterized protein (Fragment)
Uncharacterized protein
Uncharacterized protein
Uncharacterized protein
Uncharacterized protein
Uncharacterized protein
Uncharacterized protein
Uncharacterized protein
Uncharacterized protein
Uncharacterized protein
Uncharacterized protein
Uncharacterized protein
Uncharacterized protein
Uncharacterized protein
Uncharacterized protein
Uncharacterized protein
Uncharacterized protein
Uncharacterized protein
Uncharacterized protein
Uncharacterized protein (Fragment)
Uncharacterized protein
Uncharacterized protein
Uncharacterized protein
Uncharacterized protein
Uncharacterized protein
Uncharacterized protein
Uncharacterized protein (Fragment)
Uncharacterized protein
Uncharacterized protein (Fragment)
Uncharacterized protein
Uncharacterized protein
Uncharacterized protein
Uncharacterized protein
Uncharacterized protein
Uncharacterized protein
Uncharacterized protein
Uncharacterized protein

Uncharacterized protein
Uncharacterized protein
Uncharacterized protein
Uncharacterized protein
Uncharacterized protein
Uncharacterized protein
PAN domain-containing protein
Uncharacterized protein
Uncharacterized protein
PAN domain-containing protein
Uncharacterized protein
PAN domain-containing protein
PAN domain-containing protein
Uncharacterized protein
Uncharacterized protein
Uncharacterized protein
Hepatocyte growth factor
Uncharacterized protein
Uncharacterized protein
Plasminogen
Plasminogen
Hepatocyte growth factor
Hepatocyte growth factor (Fragment)
Coagulation factor XI (Fragment)
Plasminogen (Fragment)
Plasminogen (Fragment)
Uncharacterized protein
Uncharacterized protein
Uncharacterized protein
Uncharacterized protein
Galactose oxidase-like protein
Carbohydrate-binding wsc
Nucleolar gtp-binding
Lccl domain-containing
Kelch domain-containing
Uncharacterized protein
Uncharacterized protein
Uncharacterized protein
Uncharacterized protein
Uncharacterized protein
Galactose oxidase

Uncharacterized protein
Uncharacterized protein
Uncharacterized protein
DUF1929 multi-domain protein
Archaeal flagellin n-terminal-like domain-containing protein
PAN-4 domain containing protein
PAN-AP domain containing protein
PAN-4 domain containing protein
Uncharacterized protein
Uncharacterized protein
Carbohydrate-binding protein, putative (Fragment)
Papain-like cysteine protease C1 (Fragment)
Uncharacterized protein
Uncharacterized protein
Papain-like cysteine protease C1
Uncharacterized protein
Elicitor-like transglutaminase
Uncharacterized protein
Uncharacterized protein
Uncharacterized protein
Uncharacterized protein
Uncharacterized protein
Uncharacterized protein
Uncharacterized protein
Putative CBEL-like protein
CBEL protein, formerly GP34
Uncharacterized protein
Uncharacterized protein
Uncharacterized protein
Uncharacterized protein
Uncharacterized protein
Uncharacterized protein
Uncharacterized protein
Uncharacterized protein
Uncharacterized protein
Uncharacterized protein
Uncharacterized protein
Uncharacterized protein
Uncharacterized protein
Uncharacterized protein
Uncharacterized protein

Uncharacterized protein
Uncharacterized protein
Uncharacterized protein
Uncharacterized protein
Metalloendopeptidase
Uncharacterized protein
Uncharacterized protein
Metalloendopeptidase
Uncharacterized protein
Uncharacterized protein
Uncharacterized protein
Uncharacterized protein
Uncharacterized protein
Uncharacterized protein
Metalloendopeptidase
Uncharacterized protein
Uncharacterized protein
Uncharacterized protein
Uncharacterized protein
Uncharacterized protein
Uncharacterized protein
Uncharacterized protein
Uncharacterized protein
Uncharacterized protein
S-locus related 1 (Fragment)
S-locus related (Fragment)
S-locus related 1 (Fragment)
S-locus related (Fragment)
S-locus related (Fragment)
S-locus related 1 (Fragment)
S-locus receptor kinase (Fragment)
S-locus receptor kinase (Fragment)
S-locus-related I (Fragment)
S-locus receptor kinase (Fragment)
S-locus receptor kinase (Fragment)
S glycoprotein (Fragment)
S glycoprotein (Fragment)
S-locus receptor kinase-19 (Fragment)
S receptor kinase 1
S-locus receptor kinase (Fragment)
S glycoprotein (Fragment)

S-locus glycoprotein (Fragment)
S glycoprotein (Fragment)
S-locus glycoprotein
S-locus receptor kinase (Fragment)
S receptor protein kinase
S-locus glycoprotein (Fragment)
S-locus receptor kinase (Fragment)
S glycoprotein (Fragment)
S-locus glycoprotein (Fragment)
S-locus receptor kinase-2 (Fragment)
S-locus glycoprotein
S-locus glycoprotein
S-locus glycoprotein (Fragment)
S locus glycoprotein
S-locus receptor kinase-18 (Fragment)
S locus protein 1 (Fragment)
S locus protein 5 (Fragment)
S-locus glycoprotein (Fragment)
S-locus receptor kinase-6 (Fragment)
S-locus glycoprotein (Fragment)
S locus protein 2 (Fragment)
S-locus receptor kinase-21 (Fragment)
S-locus receptor kinase-9 (Fragment)
S-locus receptor kinase (Fragment)
S-locus receptor kinase-8 (Fragment)
SLR1 (Fragment)
S-locus glycoprotein (Fragment)
S-locus glycoprotein
S-locus glycoprotein (Fragment)
S glycoprotein (Fragment)
S-locus glycoprotein
S-locus glycoprotein
S locus protein 3 (Fragment)
S-locus glycoprotein
S-locus receptor kinase-1 (Fragment)
S-locus glycoprotein (Fragment)
S-locus receptor kinase (Fragment)
S glycoprotein (Fragment)
S receptor kinase 19
S glycoprotein (Fragment)
S locus protein 4 (Fragment)

S-locus related protein (Fragment)
S glycoprotein T63
S-locus receptor kinase (Fragment)
S-locus receptor kinase (Fragment)
S-locus receptor kinase-4 (Fragment)
S-locus glycoprotein
SLG protein (Fragment)
S-locus glycoprotein
Uncharacterized protein
Uncharacterized protein
Uncharacterized protein
Peptidyl-prolyl cis-trans isomerase D
Uncharacterized protein
Uncharacterized protein
Uncharacterized protein
Uncharacterized protein
Plasminogen
Hepatocyte growth factor
Plasma kallikrein
Coagulation factor XI
Macrophage-stimulating 1
Macrophage stimulating 1 (Hepatocyte growth factor-like)
Kallikrein B, plasma 1
Coagulation factor XI
Putative serine protease 68
Putative CTL16
Major sperm protein
Uncharacterized protein
Uncharacterized protein
Uncharacterized protein
Uncharacterized protein
Uncharacterized protein
Uncharacterized protein
Uncharacterized protein
Uncharacterized protein
Uncharacterized protein
Uncharacterized protein
Uncharacterized protein
Uncharacterized protein
Uncharacterized protein
Uncharacterized protein

Uncharacterized protein
Uncharacterized protein
Uncharacterized protein
Uncharacterized protein
Plasminogen
Hepatocyte growth factor
Uncharacterized protein
Uncharacterized protein
Uncharacterized protein
Uncharacterized protein
Uncharacterized protein
Uncharacterized protein
Uncharacterized protein
Uncharacterized protein
Uncharacterized protein
Plasminogen
Uncharacterized protein
Uncharacterized protein
Uncharacterized protein
Hepatocyte growth factor
Uncharacterized protein
Uncharacterized protein
Uncharacterized protein
Antigen B membrane protein
Peptidase C14 caspase catalytic subunit p20
Apple domain-containing protein
Apple domain-containing protein
Uncharacterized protein
Uncharacterized protein
Uncharacterized protein
Uncharacterized protein
Uncharacterized protein
Uncharacterized protein
Alpha-2-macroglobulin
Uncharacterized protein
Uncharacterized protein
Uncharacterized protein
Uncharacterized protein
Uncharacterized protein
Uncharacterized protein (Fragment)
Uncharacterized protein

Serine protease
Uncharacterized protein
Uncharacterized protein
Uncharacterized protein
Uncharacterized protein
Uncharacterized protein
Uncharacterized protein
Hypothetical conserved protein
Hypothetical conserved protein
Uncharacterized protein
Alpha-2-macroglobulin domain-containing protein
Alpha-2-macroglobulin domain-containing protein
Alpha-2-macroglobulin domain-containing protein
Uncharacterized protein
Uncharacterized protein
Large extracellular alpha-helical protein
Putative alpha-2-macroglobulin domain-containing protein
PAN/APPLE domain-containing protein
Serine protease
Alpha-2-macroglobulin domain-containing protein
Uncharacterized protein
Alpha-2-macroglobulin domain protein
Alpha-2-macroglobulin domain-containing protein
Alpha-2-macroglobulin domain-containing protein
Alpha-2-macroglobulin domain protein
Uncharacterized protein
Uncharacterized protein
Uncharacterized protein
Uncharacterized protein
Uncharacterized protein
Peptidase C14 caspase catalytic subunit p20
Inner membrane lipoprotein
Alpha-2-macroglobulin domain protein
Peptidase C14 caspase catalytic subunit p20
Alpha-2-macroglobulin domain protein
Membrane protein
Large extracellular alpha-helical protein
Large extracellular alpha-helical protein
Large extracellular alpha-helical protein
Trypsin-like serine protease with C-terminal PDZ domain
Membrane protein

Uncharacterized protein
Uncharacterized protein
Uncharacterized protein
Serine protease
Hypothetical exported protein
Uncharacterized protein
Uncharacterized protein
Large extracellular alpha-helical protein
Uncharacterized protein
Alpha-2-macroglobulin MG1 domain protein
Uncharacterized protein
Serine protease
Uncharacterized protein
Peptidase C14 caspase catalytic subunit p20
Uncharacterized protein
Uncharacterized protein
Uncharacterized protein
Uncharacterized protein
Uncharacterized protein
Peptidase C14 caspase catalytic subunit p20
Uncharacterized protein
Uncharacterized protein (Fragment)
Peptidase C14 caspase catalytic subunit p20
Uncharacterized protein
Uncharacterized protein
Uncharacterized protein
PAN domain family protein
Uncharacterized protein
Alpha-2-macroglobulin N-terminal region family protein
Uncharacterized protein
Uncharacterized protein
Uncharacterized protein
Uncharacterized protein
Uncharacterized protein
Alpha-2-macroglobulin domain-containing protein
Uncharacterized protein
Uncharacterized protein
Uncharacterized protein
Uncharacterized protein
Uncharacterized protein
Uncharacterized protein

Uncharacterized protein
Uncharacterized protein
Uncharacterized protein
Uncharacterized protein
Uncharacterized protein
Uncharacterized protein
Uncharacterized protein
Uncharacterized protein
Uncharacterized protein
Uncharacterized protein
Uncharacterized protein
Uncharacterized protein
Uncharacterized protein
Uncharacterized protein
Uncharacterized protein
Uncharacterized protein
Membrane protein
Uncharacterized protein
Uncharacterized protein
Uncharacterized protein
Peptidase PDZ domain-containing protein
Alpha-2-macroglobulin domain-containing protein
Uncharacterized protein
Alpha-2-macroglobulin domain-containing protein
Peptidase PDZ domain-containing protein
Alpha-2-macroglobulin domain-containing protein
Uncharacterized protein
Uncharacterized protein
Uncharacterized protein
Uncharacterized protein
Uncharacterized protein
Uncharacterized protein
Uncharacterized protein
Uncharacterized protein
Uncharacterized protein
Membrane protein
Uncharacterized protein
Uncharacterized protein
Uncharacterized protein
Membrane protein
UPF0192 protein

Uncharacterized protein
Uncharacterized protein
Membrane protein
Uncharacterized protein
Uncharacterized protein
Membrane protein
Serine protease
Uncharacterized protein
Serine protease
Uncharacterized protein
Uncharacterized protein
Uncharacterized protein
Uncharacterized protein
Uncharacterized protein
Uncharacterized protein
Uncharacterized protein
Uncharacterized protein
Uncharacterized protein
Uncharacterized protein
Uncharacterized protein
Uncharacterized protein
Uncharacterized protein
Uncharacterized protein
Uncharacterized protein
Uncharacterized protein
Uncharacterized protein
Uncharacterized protein
Trypsin-like serine protease with C-terminal PDZ domain-containing protein
Large extracellular alpha-helical protein (Fragment)
Serine protease
Uncharacterized protein
Serine protease
Uncharacterized protein
Alpha-2-macroglobulin domain protein
Uncharacterized protein
Alpha-2-macroglobulin MG1 domain-containing protein (Fragment)
Large extracellular alpha-helical protein
Large extracellular alpha-helical protein
Alpha-2-macroglobulin domain-containing protein
Uncharacterized protein
Membrane protein

Alpha-2-macroglobulin domain-containing protein
Putative bifunctional protein: Alpha-2-macroglobulin Terpenoid cylases/protein prenyltransferase alpha
Uncharacterized protein
Uncharacterized protein
Uncharacterized protein
Uncharacterized protein
Serine protease
Uncharacterized protein
Serine protease
Uncharacterized protein
Uncharacterized protein
Membrane protein
Uncharacterized protein
Uncharacterized protein
Serine protease
Uncharacterized protein
Uncharacterized protein
Uncharacterized protein
Uncharacterized protein
Uncharacterized protein
Uncharacterized protein
Uncharacterized protein
Alpha-2-macroglobulin domain-containing protein
Peptidase PDZ domain-containing protein
Alpha-2-macroglobulin domain-containing protein
Peptidase PDZ domain-containing protein
Uncharacterized protein
Uncharacterized protein
Uncharacterized protein
PAN domain-containing protein
Uncharacterized protein
Putative inner membrane lipoprotein
Peptidase PDZ domain-containing protein
Alpha-2-macroglobulin domain-containing protein
Alpha-2-macroglobulin domain-containing protein
Peptidase PDZ domain-containing protein
Uncharacterized protein
Alpha-2-macroglobulin domain protein
Alpha-2-macroglobulin domain protein
PAN domain-containing protein
Uncharacterized protein

Uncharacterized protein
Uncharacterized protein
Uncharacterized protein
Uncharacterized protein
Uncharacterized protein
Uncharacterized protein
Uncharacterized protein
Uncharacterized protein
Uncharacterized protein
Serine protease
Uncharacterized protein
Uncharacterized protein
Uncharacterized protein
Uncharacterized protein
Uncharacterized protein
Uncharacterized protein
Serine protease
Uncharacterized protein
Uncharacterized protein
Uncharacterized protein
Uncharacterized protein
Uncharacterized protein
Uncharacterized protein
Uncharacterized protein
Uncharacterized protein
Uncharacterized protein
Uncharacterized protein
Peptidase C14 caspase catalytic subunit p20
Uncharacterized protein
Uncharacterized protein
Uncharacterized protein
Uncharacterized protein
Membrane protein
Uncharacterized protein (Fragment)
Uncharacterized protein
Uncharacterized protein
Uncharacterized protein
Uncharacterized protein
Uncharacterized protein
Uncharacterized protein
Uncharacterized protein

Uncharacterized protein
Alpha-2-macroglobulin domain-containing protein
Uncharacterized protein
PAN domain-containing protein
Uncharacterized protein
Uncharacterized protein
Uncharacterized protein
Uncharacterized protein
Uncharacterized protein
Acid protease
L domain-like protein
Uncharacterized protein
Uncharacterized protein
Uncharacterized protein
Uncharacterized protein
Uncharacterized protein
Uncharacterized protein
Uncharacterized protein
Uncharacterized protein
Uncharacterized protein
Uncharacterized protein
Uncharacterized protein
Uncharacterized protein
Uncharacterized protein
Uncharacterized protein
Uncharacterized protein
Uncharacterized protein
Uncharacterized protein MANES_12G079200
Serine/threonine-protein kinase
Serine/threonine-protein kinase
Serine/threonine-protein kinase
Serine/threonine-protein kinase
Serine/threonine-protein kinase
Putative receptor protein kinase ZmPK1
Uncharacterized protein MANES_03G118000
Uncharacterized protein MANES_13G076800
Uncharacterized protein
Uncharacterized protein
Uncharacterized protein (Fragment)
Uncharacterized protein
Serine-type endopeptidase activity (Fragment)

Uncharacterized protein (Fragment)
Uncharacterized protein
Uncharacterized protein
Uncharacterized protein
PAN domain-containing protein
Alpha-2-macroglobulin domain protein
PAN domain protein
Uncharacterized protein
Alpha-2-macroglobulin family protein
Uncharacterized protein
Uncharacterized protein
Uncharacterized protein
Uncharacterized protein
Uncharacterized protein
Alpha-2-macroglobulin family protein
Alpha-2-macroglobulin domain protein
PAN domain-containing protein
Large extracellular alpha-helical protein
Alpha-2-macroglobulin domain protein
Alpha-2-macroglobulin domain protein
Alpha-2-macroglobulin domain protein
Alpha-2-macroglobulin family protein
PAN domain-containing protein
Alpha-2-macroglobulin family protein
Uncharacterized protein
PAN domain-containing protein
Uncharacterized protein
Uncharacterized protein
PAN domain-containing protein
Alpha-2-macroglobulin family protein
Alpha-2-macroglobulin family protein
Alpha-2-macroglobulin family protein
Alpha-2-macroglobulin family protein
Alpha-2-macroglobulin family protein
Uncharacterized protein
PAN domain-containing protein
Large extracellular alpha-helical protein
Alpha-2-macroglobulin N-terminal domain family protein
Uncharacterized protein
Uncharacterized protein
Alpha-2-macroglobulin family protein

PAN domain-containing protein
Contig2, whole genome shotgun sequence
Alpha-2-macroglobulin family protein
PAN domain-containing protein (Fragment)
PAN domain-containing protein
Uncharacterized protein
PAN domain-containing protein
PAN domain-containing protein (Fragment)
Uncharacterized protein
PAN domain-containing protein
PAN domain-containing protein (Fragment)
PAN domain-containing protein
PAN domain-containing protein
Alpha-2-macroglobulin (Fragment)
PAN domain-containing protein
PAN domain-containing protein
Large extracellular alpha-helical protein
Uncharacterized protein
Uncharacterized protein (Fragment)
Uncharacterized protein (Fragment)
PAN domain-containing protein
PAN domain-containing protein
Alpha-2-macroglobulin N-terminal domain family protein
PAN domain-containing protein
PAN domain-containing protein
PAN domain-containing protein
Serine protease
PAN domain-containing protein
Uncharacterized protein
Uncharacterized protein
PAN domain protein
Uncharacterized protein
Uncharacterized protein
Alpha-2-macroglobulin
Uncharacterized protein
Uncharacterized protein
Uncharacterized protein
Uncharacterized protein
Apple domain-containing protein
PAN domain-containing protein
Apple domain:N/apple PAN

Apple domain, N/apple PAN
Uncharacterized protein
Apple
N/apple PAN protein
Uncharacterized protein
Apple PAN
Apple
Uncharacterized protein
Apple domain-containing protein
Apple domain-containing protein
Apple domain, N/apple PAN
Apple domain-containing protein
PAN domain-containing protein
PAN domain-containing protein
Apple domain-containing protein
Alpha-2-macroglobulin family protein
Uncharacterized protein (Fragment)
Uncharacterized protein
Alpha-2-macroglobulin family protein
Alpha-2-macroglobulin family protein
PAN domain-containing protein
Uncharacterized protein
PAN domain-containing protein
PAN domain-containing protein
Uncharacterized protein
Related to galactose oxidase
Uncharacterized protein
Uncharacterized protein
Uncharacterized protein
Uncharacterized protein
Uncharacterized protein
Related to galactose oxidase
Uncharacterized protein
Uncharacterized protein
Related to galactose oxidase
Uncharacterized protein
Uncharacterized protein
Uncharacterized protein
Uncharacterized protein
Uncharacterized protein
Uncharacterized protein

Uncharacterized protein
S-receptor kinase, putative
Leucine-rich repeat receptor protein kinase EXS, putative
S-locus-specific glycoprotein S6, putative
B-Raf proto-oncogene serine/threonine-protein kinase, putative
Receptor protein kinase, putative
Uncharacterized protein
S-locus-specific glycoprotein S13, putative
S-locus-specific glycoprotein S6, putative
S-locus-specific glycoprotein S13, putative
Serine-threonine protein kinase, plant-type, putative
S-receptor kinase, putative
Serine-threonine protein kinase, plant-type, putative
Uncharacterized protein
Receptor protein kinase zmpk1, putative
Uncharacterized protein
Serine-threonine protein kinase, plant-type, putative
Uncharacterized protein
S-locus-specific glycoprotein S6, putative
S-locus-specific glycoprotein S6, putative
S-locus-specific glycoprotein, putative
S-locus-specific glycoprotein S13, putative
Receptor protein kinase zmpk1, putative
Uncharacterized protein (Fragment)
Negative regulator of the PHO system, putative
S-locus-specific glycoprotein S13, putative
ATP binding protein, putative
Serine/threonine-protein kinase PBS1, putative
ATP binding protein, putative
Serine-threonine protein kinase, plant-type, putative
Uncharacterized protein
ATP binding protein, putative
Uncharacterized protein
ATP binding protein, putative
Serine-threonine protein kinase, plant-type, putative
S-locus-specific glycoprotein S6, putative
Leucine-rich repeat receptor protein kinase EXS, putative
S-locus-specific glycoprotein S6, putative
S-locus-specific glycoprotein S13, putative
S-receptor kinase, putative
S-locus-specific glycoprotein S13, putative

S-locus-specific glycoprotein S13, putative
S-locus-specific glycoprotein S6, putative
Serine-threonine protein kinase, plant-type, putative
Sugar binding protein, putative
S-locus-specific glycoprotein S13, putative
Uncharacterized protein
S-locus-specific glycoprotein S13, putative
S-locus-specific glycoprotein S6, putative
S-locus-specific glycoprotein S6, putative
S-receptor kinase, putative
S-locus-specific glycoprotein S6, putative
S-locus-specific glycoprotein S6, putative
Uncharacterized protein
Uncharacterized protein
Uncharacterized protein
Uncharacterized protein
Uncharacterized protein
Uncharacterized protein
Protein DETOXIFICATION
Uncharacterized protein
Uncharacterized protein
Uncharacterized protein
Uncharacterized protein
Uncharacterized protein
Uncharacterized protein
Uncharacterized protein
Putative non-specific serine/threonine protein kinase
Uncharacterized protein
Putative non-specific serine/threonine protein kinase
Uncharacterized protein
Uncharacterized protein
Uncharacterized protein
Uncharacterized protein
Uncharacterized protein
Uncharacterized protein
Uncharacterized protein
Uncharacterized protein
Uncharacterized protein
Uncharacterized protein
Uncharacterized protein
Uncharacterized protein

Uncharacterized protein
Uncharacterized protein
Uncharacterized protein
Uncharacterized protein
Uncharacterized protein
Uncharacterized protein
Putative non-specific serine/threonine protein kinase
Uncharacterized protein
Putative non-specific serine/threonine protein kinase
Uncharacterized protein
Uncharacterized protein
Putative non-specific serine/threonine protein kinase
Putative S-locus glycoprotein
Uncharacterized protein
Uncharacterized protein
Uncharacterized protein
Uncharacterized protein
Uncharacterized protein
Uncharacterized protein
Uncharacterized protein
Uncharacterized protein
Uncharacterized protein
Uncharacterized protein
Uncharacterized protein
Uncharacterized protein
Putative non-specific serine/threonine protein kinase
Uncharacterized protein
Uncharacterized protein
Uncharacterized protein
Uncharacterized protein
Uncharacterized protein
Uncharacterized protein
Uncharacterized protein
Uncharacterized protein
Uncharacterized protein
Uncharacterized protein
Uncharacterized protein
Uncharacterized protein
Uncharacterized protein
Uncharacterized protein
Uncharacterized protein

Uncharacterized protein
Uncharacterized protein
Uncharacterized protein
Uncharacterized protein
Uncharacterized protein
Uncharacterized protein
Putative non-specific serine/threonine protein kinase
Uncharacterized protein
Putative non-specific serine/threonine protein kinase
Uncharacterized protein
Uncharacterized protein
Uncharacterized protein
Uncharacterized protein
Uncharacterized protein
Uncharacterized protein
Uncharacterized protein
Uncharacterized protein
Uncharacterized protein
Uncharacterized protein
Uncharacterized protein
Uncharacterized protein
Uncharacterized protein
Putative S-locus glycoprotein
Uncharacterized protein
Uncharacterized protein
Uncharacterized protein
Uncharacterized protein
Uncharacterized protein
Uncharacterized protein
Uncharacterized protein
Uncharacterized protein
Uncharacterized protein
Putative non-specific serine/threonine protein kinase
Uncharacterized protein
Uncharacterized protein
Uncharacterized protein
Uncharacterized protein
Uncharacterized protein
Uncharacterized protein
Putative non-specific serine/threonine protein kinase
Uncharacterized protein

Uncharacterized protein
Uncharacterized protein
Putative non-specific serine/threonine protein kinase
Uncharacterized protein
Uncharacterized protein
Uncharacterized protein
Uncharacterized protein
Uncharacterized protein
Uncharacterized protein
Uncharacterized protein
Uncharacterized protein
Uncharacterized protein
Uncharacterized protein
Uncharacterized protein
Uncharacterized protein
Uncharacterized protein
Uncharacterized protein
Uncharacterized protein
Uncharacterized protein
Uncharacterized protein
Uncharacterized protein
Putative non-specific serine/threonine protein kinase
Uncharacterized protein
Uncharacterized protein
Uncharacterized protein
Uncharacterized protein
Uncharacterized protein
Uncharacterized protein
Uncharacterized protein
Uncharacterized protein
Uncharacterized protein
Putative PAN/Apple domain-containing protein
Uncharacterized protein
Putative non-specific serine/threonine protein kinase
Uncharacterized protein
Putative non-specific serine/threonine protein kinase
Uncharacterized protein
Uncharacterized protein
Uncharacterized protein
Uncharacterized protein
Uncharacterized protein

Uncharacterized protein
Putative non-specific serine/threonine protein kinase
Uncharacterized protein
Uncharacterized protein
Uncharacterized protein
Uncharacterized protein
Uncharacterized protein
Uncharacterized protein
Uncharacterized protein
Uncharacterized protein
Uncharacterized protein
Uncharacterized protein
Uncharacterized protein
Putative non-specific serine/threonine protein kinase
Uncharacterized protein
Uncharacterized protein
Putative bulb-type lectin domain, PAN/Apple domain-containing protein
Putative non-specific serine/threonine protein kinase
Uncharacterized protein
Uncharacterized protein
Uncharacterized protein
Uncharacterized protein
Uncharacterized protein
Uncharacterized protein
Uncharacterized protein
Uncharacterized protein
Putative non-specific serine/threonine protein kinase
Uncharacterized protein
Uncharacterized protein
Putative non-specific serine/threonine protein kinase
Uncharacterized protein
Putative S-locus glycoprotein
Uncharacterized protein
Uncharacterized protein
Uncharacterized protein
Uncharacterized protein
Uncharacterized protein
Uncharacterized protein
Uncharacterized protein
Putative non-specific serine/threonine protein kinase
Uncharacterized protein

Uncharacterized protein
Uncharacterized protein
Uncharacterized protein
Uncharacterized protein
Uncharacterized protein
Uncharacterized protein
Uncharacterized protein
Uncharacterized protein
Uncharacterized protein
Putative non-specific serine/threonine protein kinase
Uncharacterized protein
Uncharacterized protein
Uncharacterized protein
Uncharacterized protein
Uncharacterized protein
Uncharacterized protein
Uncharacterized protein
Putative non-specific serine/threonine protein kinase
Putative non-specific serine/threonine protein kinase
Uncharacterized protein
Uncharacterized protein
Uncharacterized protein
Uncharacterized protein
Uncharacterized protein
Uncharacterized protein
Putative non-specific serine/threonine protein kinase
Uncharacterized protein
Uncharacterized protein
Uncharacterized protein
Uncharacterized protein
Uncharacterized protein
Putative bulb-type lectin domain, PAN/Apple domain-containing protein
Large extracellular alpha-helical protein
Putative lipoprotein YfhM
PAN domain protein
Alpha-2-macroglobulin domain-containing protein
Uncharacterized protein
Uncharacterized protein
PAN domain protein
Uncharacterized protein
Uncharacterized protein

Uncharacterized protein
Uncharacterized protein
Putative WSC domain protein
Alpha-2-macroglobulin N-terminal domain family protein
PAN domain protein
Uncharacterized protein
Uncharacterized protein
Uncharacterized protein
PAN domain-containing protein
PAN domain-containing protein
Uncharacterized protein
Uncharacterized protein
PAN domain protein
PAN domain-containing protein (Fragment)
Uncharacterized protein
PAN domain-containing protein
Alpha-2-macroglobulin family protein
Alpha-2-macroglobulin family protein
PAN domain-containing protein
Uncharacterized protein
PAN domain-containing protein
PAN domain protein
Uncharacterized protein
Uncharacterized protein
PAN domain-containing protein
Putative lipoprotein YfhM
PAN domain protein
Alpha-2-macroglobulin family protein
Uncharacterized protein
PAN domain-containing protein
Uncharacterized protein
Uncharacterized protein
HEG protein
Uncharacterized protein
Serine/threonine-protein kinase
PAN domain-containing protein
Uncharacterized protein
Macrophage stimulating 1
Hepatocyte growth factor
Kallikrein B1
Kallikrein B1

Macrophage stimulating 1
Hepatocyte growth factor
Hepatocyte growth factor
Plasminogen
Coagulation factor XI
Coagulation factor XI
Uncharacterized protein
Uncharacterized protein
Uncharacterized protein
Uncharacterized protein
PAN domain protein
Uncharacterized protein
Uncharacterized protein
hepatocyte growth factor-like protein isoform X2
Plasminogen
secretory phospholipase A2 receptor-like isoform X1
Plasminogen
secretory phospholipase A2 receptor-like isoform X2
uncharacterized protein LOC106609371 isoform X4
Hepatocyte growth factor
uncharacterized protein LOC106609370
hepatocyte growth factor-like
secretory phospholipase A2 receptor-like isoform X3
hepatocyte growth factor-like isoform X1
hepatocyte growth factor-like isoform X2
Hepatocyte growth factor
uncharacterized protein LOC106609376
hepatocyte growth factor-like protein isoform X1
hepatocyte growth factor-like protein
Chitin-binding protein
Uncharacterized protein
Uncharacterized protein
Uncharacterized protein
TK/RTKC protein kinase
Uncharacterized protein
Uncharacterized protein
Uncharacterized protein
Uncharacterized protein
Uncharacterized protein
Uncharacterized protein
TK/RTKC protein kinase

Uncharacterized protein
Uncharacterized protein
Uncharacterized protein
Uncharacterized protein
Uncharacterized protein
Uncharacterized protein
Uncharacterized protein
Uncharacterized protein
Uncharacterized protein
PAN domain-containing protein
Uncharacterized protein
Uncharacterized protein
Uncharacterized protein
Uncharacterized protein
Uncharacterized protein
Uncharacterized protein
Uncharacterized protein
Uncharacterized protein
Uncharacterized protein
Uncharacterized protein
Uncharacterized protein
Uncharacterized protein
Uncharacterized protein
Uncharacterized protein
Uncharacterized protein
Uncharacterized protein
Uncharacterized protein
Uncharacterized protein
Glucanase
Uncharacterized protein
Uncharacterized protein
Uncharacterized protein (Fragment)
Uncharacterized protein
Uncharacterized protein (Fragment)
Uncharacterized protein
Uncharacterized protein
Uncharacterized protein
Uncharacterized protein
Uncharacterized protein (Fragment)
Uncharacterized protein
Uncharacterized protein

Uncharacterized protein
Glucanase
Uncharacterized protein
Uncharacterized protein
Uncharacterized protein
Uncharacterized protein
Uncharacterized protein
Glucanase
Uncharacterized protein
Uncharacterized protein
Uncharacterized protein
Uncharacterized protein (Fragment)
Uncharacterized protein
Uncharacterized protein
Uncharacterized protein (Fragment)
Uncharacterized protein
Uncharacterized protein
Uncharacterized protein
Uncharacterized protein
Uncharacterized protein
Uncharacterized protein
Uncharacterized protein
Uncharacterized protein
DNA/RNA non-specific endonuclease
Uncharacterized protein
Uncharacterized protein
Uncharacterized protein
Uncharacterized protein
Uncharacterized protein
Uncharacterized protein
Uncharacterized protein
Uncharacterized protein
Uncharacterized protein
Uncharacterized protein
Uncharacterized protein
Uncharacterized protein
Glucanase
Uncharacterized protein
Uncharacterized protein
Uncharacterized protein
Uncharacterized protein (Fragment)

Uncharacterized protein
Uncharacterized protein
Uncharacterized protein
Uncharacterized protein
Uncharacterized protein
Uncharacterized protein
Uncharacterized protein
Uncharacterized protein
Uncharacterized protein
Uncharacterized protein
Uncharacterized protein (Fragment)
Uncharacterized protein
Uncharacterized protein
Uncharacterized protein (Fragment)
Uncharacterized protein
Uncharacterized protein
Uncharacterized protein
Uncharacterized protein (Fragment)
Uncharacterized protein
Uncharacterized protein
Uncharacterized protein
Uncharacterized protein
Uncharacterized protein
Uncharacterized protein
Uncharacterized protein
Uncharacterized protein
Uncharacterized protein
Glucanase
Uncharacterized protein
Uncharacterized protein
Uncharacterized protein
Uncharacterized protein
Uncharacterized protein
PAN domain protein
Uncharacterized protein
Major microneme antigen
Microneme antigen L2
Microneme antigen
Macrophage stimulating 1
Kallikrein B1
Macrophage stimulating 1

Uncharacterized protein
Uncharacterized protein
Uncharacterized protein
Hepatocyte growth factor
Uncharacterized protein
Neyo-like protein (Fragment)
Uncharacterized protein
Zona pellucida and PAN domain containing protein
PAN domain containing protein 2 (Fragment)
Zona pellucida-like and PAN domain containing protein
Zona pellucida-like domain containing protein 3
PAN domain containing protein 1
PAn domain protein
PAn domain protein
Uncharacterized protein
Uncharacterized protein
Uncharacterized protein
Uncharacterized protein
MANEC domain
Uncharacterized protein
DNA topoisomerase II
Uncharacterized protein
Uncharacterized protein
Uncharacterized protein
Uncharacterized protein
SJCHGC03220 protein
Putative von Willebrand factor type A domain-containing protein
Uncharacterized protein
Uncharacterized protein
Uncharacterized protein
Uncharacterized protein
Uncharacterized protein (Fragment)
Phosphoinositide-3-kinase-interacting protein 1 (Fragment)
Uncharacterized protein
Galectin-3-binding protein A (Fragment)
Uncharacterized protein
Uncharacterized protein
Uncharacterized protein (Fragment)
Uncharacterized protein
Uncharacterized protein
Uncharacterized protein (Fragment)

coagulation factor XI-like
Uncharacterized protein
plasma kallikrein-like
Plasminogen-like (Fragment)
Coagulation factor XI-like (Fragment)
Hepatocyte growth factor-like (Fragment)
Hepatocyte growth factor
Hepatocyte growth factor-like (Fragment)
Hepatocyte growth factor
hepatocyte growth factor-like protein
Hepatocyte growth factor
Hepatocyte growth factor-like protein-like (Fragment)
Plasminogen
Uncharacterized protein
Uncharacterized protein
Uncharacterized protein
Uncharacterized protein
Uncharacterized protein
Uncharacterized protein
Uncharacterized protein
Uncharacterized protein
Putative hepatocyte growth factor-like
Hepatocyte growth factor
Plasminogen
Putative coagulation factor XI-like isoform 4
Putative hepatocyte growth factor-like protein-like
Putative coagulation factor XI-like isoform 2
Putative coagulation factor XI-like
Putative coagulation factor XI-like isoform 3
Plasminogen
Putative coagulation factor XI-like isoform 5
Uncharacterized protein
Uncharacterized protein
Uncharacterized protein
Uncharacterized protein (Fragment)
Uncharacterized protein
Hepatocyte growth factor
Uncharacterized protein
Uncharacterized protein
Uncharacterized protein
Uncharacterized protein

Uncharacterized protein
Uncharacterized protein
Uncharacterized protein (Fragment)
Uncharacterized protein
Uncharacterized protein
Uncharacterized protein
S-receptor kinase 1 (Fragment)
Uncharacterized protein
Uncharacterized protein
Uncharacterized protein
Uncharacterized protein
Uncharacterized protein
Uncharacterized protein
Uncharacterized protein
Uncharacterized protein (Fragment)
Uncharacterized protein (Fragment)
Uncharacterized protein (Fragment)
Uncharacterized protein
Uncharacterized protein (Fragment)
Uncharacterized protein (Fragment)
Uncharacterized protein
Uncharacterized protein (Fragment)
Uncharacterized protein
Uncharacterized protein (Fragment)
Uncharacterized protein
Uncharacterized protein (Fragment)
Uncharacterized protein (Fragment)
Uncharacterized protein (Fragment)
Uncharacterized protein
Uncharacterized protein
Uncharacterized protein
Uncharacterized protein
Uncharacterized protein
S-receptor kinase-like protein 1
GON-SepT-1
GON-SepT-2
Uncharacterized protein
Uncharacterized protein
Uncharacterized protein
Uncharacterized protein
Hepatocyte growth factor

Uncharacterized protein
Plasminogen
Uncharacterized protein
Uncharacterized protein
Uncharacterized protein
Hepatocyte growth factor
Uncharacterized protein
Uncharacterized protein
Uncharacterized protein
Uncharacterized protein
Uncharacterized protein
Uncharacterized protein
Uncharacterized protein
Uncharacterized protein
Uncharacterized protein (Fragment)
Uncharacterized protein
Uncharacterized protein
Uncharacterized protein
Uncharacterized protein
Uncharacterized protein
Uncharacterized protein
Uncharacterized protein
Uncharacterized protein
Uncharacterized protein
Uncharacterized protein
Uncharacterized protein
Uncharacterized protein
Uncharacterized protein
Uncharacterized protein
Uncharacterized protein
Uncharacterized protein
Uncharacterized protein
Uncharacterized protein
Uncharacterized protein
Uncharacterized protein
Uncharacterized protein
Uncharacterized protein
Uncharacterized protein
Uncharacterized protein
Uncharacterized protein
Uncharacterized protein

Uncharacterized protein
Uncharacterized protein
Uncharacterized protein
Uncharacterized protein
Uncharacterized protein
Uncharacterized protein
Uncharacterized protein
Uncharacterized protein
Uncharacterized protein
Uncharacterized protein
Uncharacterized protein
Uncharacterized protein
Uncharacterized protein
Uncharacterized protein
Uncharacterized protein
Uncharacterized protein
Uncharacterized protein
Uncharacterized protein
Uncharacterized protein
Uncharacterized protein
Uncharacterized protein
Uncharacterized protein
Uncharacterized protein
Uncharacterized protein
Uncharacterized protein
Uncharacterized protein
Uncharacterized protein
Uncharacterized protein
Uncharacterized protein
Uncharacterized protein
Uncharacterized protein
Uncharacterized protein
Uncharacterized protein
Uncharacterized protein
Uncharacterized protein
Uncharacterized protein
Uncharacterized protein
Uncharacterized protein
Uncharacterized protein
Uncharacterized protein
Uncharacterized protein

Uncharacterized protein
Uncharacterized protein
Uncharacterized protein
Uncharacterized protein
Uncharacterized protein
Uncharacterized protein
Uncharacterized protein
Uncharacterized protein
Uncharacterized protein
Uncharacterized protein
Uncharacterized protein
Uncharacterized protein
Uncharacterized protein
Uncharacterized protein
Uncharacterized protein
Uncharacterized protein
Uncharacterized protein
Uncharacterized protein
Uncharacterized protein
Uncharacterized protein
Uncharacterized protein
Uncharacterized protein
Uncharacterized protein
Uncharacterized protein
Uncharacterized protein
Uncharacterized protein
Uncharacterized protein
Uncharacterized protein
Uncharacterized protein
Uncharacterized protein
Uncharacterized protein
Uncharacterized protein
Uncharacterized protein
Uncharacterized protein
Uncharacterized protein
Uncharacterized protein
Uncharacterized protein
Uncharacterized protein
Uncharacterized protein
Uncharacterized protein
Uncharacterized protein

Uncharacterized protein
Uncharacterized protein
Uncharacterized protein
Uncharacterized protein
Uncharacterized protein
Uncharacterized protein
Uncharacterized protein
Uncharacterized protein
Uncharacterized protein
Uncharacterized protein
Uncharacterized protein
Uncharacterized protein
Uncharacterized protein
Uncharacterized protein
Uncharacterized protein
Uncharacterized protein
Uncharacterized protein
Uncharacterized protein
Uncharacterized protein
Uncharacterized protein
Uncharacterized protein
Uncharacterized protein
Uncharacterized protein
Uncharacterized protein
Uncharacterized protein
Uncharacterized protein
Uncharacterized protein
Uncharacterized protein
Uncharacterized protein
Uncharacterized protein
Uncharacterized protein
Uncharacterized protein
Uncharacterized protein
Uncharacterized protein
Uncharacterized protein
Uncharacterized protein
Uncharacterized protein
Uncharacterized protein
Uncharacterized protein
Uncharacterized protein
Uncharacterized protein

Uncharacterized protein
Uncharacterized protein
Uncharacterized protein
Uncharacterized protein
Uncharacterized protein
Uncharacterized protein
Uncharacterized protein
Uncharacterized protein
Uncharacterized protein
Uncharacterized protein
Uncharacterized protein
Uncharacterized protein
Uncharacterized protein
Uncharacterized protein
Uncharacterized protein
Uncharacterized protein
Uncharacterized protein
Uncharacterized protein
Uncharacterized protein
Uncharacterized protein
Uncharacterized protein
Uncharacterized protein
Uncharacterized protein
Uncharacterized protein
Uncharacterized protein
Uncharacterized protein
Uncharacterized protein
Uncharacterized protein
Uncharacterized protein
Uncharacterized protein
Uncharacterized protein
Uncharacterized protein
Uncharacterized protein
Uncharacterized protein
Uncharacterized protein
Uncharacterized protein
Uncharacterized protein
Uncharacterized protein
Putative alpha-2-macroglobulin domain-containing protein
Uncharacterized protein
Uncharacterized protein

Uncharacterized protein
Uncharacterized protein
Serine protease
Uncharacterized protein
PAN domain protein
S-locus glycoprotein
S-locus-related I (Fragment)
S-locus-related I (Fragment)
S-locus related 1 (Fragment)
S-locus-related I (Fragment)
S-locus related (Fragment)
S-locus related 1 (Fragment)
Alpha-2-macroglobulin family protein
Alpha-2-macroglobulin family protein
Alpha-2-macroglobulin family protein
Alpha-2-macroglobulin family protein
Alpha-2-macroglobulin family protein
Alpha-2-macroglobulin family protein
PAN domain-containing protein
Inner membrane lipoprotein
Periplasmic protein-like protein
Uncharacterized protein
Uncharacterized protein
Periplasmic protein-like protein
Inner membrane lipoprotein
Uncharacterized protein (Fragment)
Putative inner membrane lipoprotein
PAN domain protein
Uncharacterized protein
PAN domain protein
Membrane protein
UPF0192 protein
Uncharacterized protein y4bJ
Alpha-2-macroglobulin domain protein 2
Uncharacterized protein
Alpha-2-macroglobulin domain-containing protein 2
Uncharacterized protein
Uncharacterized protein
Uncharacterized protein
Uncharacterized protein
PAN domain protein

Uncharacterized protein
Membrane protein
Membrane protein
Uncharacterized protein
Uncharacterized protein
Uncharacterized protein
Uncharacterized protein
Uncharacterized protein
Alpha-2-macroglobulin family protein
Caspase domain protein
Uncharacterized protein
Membrane protein
Uncharacterized protein
Uncharacterized protein
Galectin-3-binding protein A
Uncharacterized protein (Fragment)
Uncharacterized protein
Uncharacterized protein
Cuticlin-1
Uncharacterized protein
Putative receptor kinase 3
Putative S-locus receptor kinase
Putative S-locus lectin protein kinase-like protein
Uncharacterized protein
Uncharacterized protein
Uncharacterized protein
Uncharacterized protein
Uncharacterized protein
Uncharacterized protein
Uncharacterized protein
Uncharacterized protein
Uncharacterized protein
Uncharacterized protein
Uncharacterized protein
Uncharacterized protein
Uncharacterized protein
Uncharacterized protein
Uncharacterized protein
Uncharacterized protein
Uncharacterized protein
Uncharacterized protein

Uncharacterized protein
Uncharacterized protein
Uncharacterized protein
Uncharacterized protein
Receptor protein kinase, putative
Putative ovule protein (Fragment)
Putative ovule protein (Fragment)
Putative ovule protein
Putative G-type lectin S-receptor-like serine/threonine-protein kinase-like
Putative ovule protein (Fragment)
Putative ovule protein (Fragment)
Putative G-type lectin S-receptor-like serine/threonine-protein kinase-like
Putative G-type lectin S-receptor-like serine/threonine-protein kinase-like
Putative G-type lectin S-receptor-like serine/threonine-protein kinase-like
Putative ovule protein
Putative ovule protein (Fragment)
Putative G-type lectin S-receptor-like serine/threonine-protein kinase-like
Putative ovule protein
Putative ovule protein (Fragment)
Putative PAN domain-containing protein-like (Fragment)
Putative G-type lectin S-receptor-like serine/threonine-protein kinase-like
Putative G-type lectin S-receptor-like serine/threonine-protein kinase SD3-1-like
Putative ovule protein
Putative ovule protein
Putative G-type lectin S-receptor-like serine/threonine-protein kinase SD2-2-like (Fragment)
Putative G-type lectin S-receptor-like serine/threonine-protein kinase-like
Uncharacterized protein
Uncharacterized protein
Uncharacterized protein
Serine/threonine-protein kinase
Uncharacterized protein
Serine/threonine-protein kinase
Uncharacterized protein
Uncharacterized protein
Uncharacterized protein
Uncharacterized protein
Uncharacterized protein
Uncharacterized protein
Uncharacterized protein
Uncharacterized protein
Serine/threonine-protein kinase

Serine/threonine-protein kinase
Serine/threonine-protein kinase
Uncharacterized protein
Uncharacterized protein
Serine/threonine-protein kinase
Uncharacterized protein
Serine/threonine-protein kinase
Uncharacterized protein
Uncharacterized protein
Serine/threonine-protein kinase
Uncharacterized protein
Serine/threonine-protein kinase
Uncharacterized protein
Uncharacterized protein
Uncharacterized protein
Uncharacterized protein
Uncharacterized protein
Uncharacterized protein
Uncharacterized protein
Uncharacterized protein
Serine/threonine-protein kinase
Serine/threonine-protein kinase
Uncharacterized protein
Uncharacterized protein
Uncharacterized protein
Serine/threonine-protein kinase
Uncharacterized protein
Serine/threonine-protein kinase
Uncharacterized protein
Uncharacterized protein
Uncharacterized protein
Serine/threonine-protein kinase
Uncharacterized protein
Uncharacterized protein
Uncharacterized protein
Uncharacterized protein
Uncharacterized protein
S-receptor-like kinase 6
S-receptor-like kinase 5
S-receptor-like kinase 4
Uncharacterized protein

Uncharacterized protein
Uncharacterized protein
Uncharacterized protein
Uncharacterized protein
Uncharacterized protein
Uncharacterized protein
Uncharacterized protein
Uncharacterized protein
Uncharacterized protein
Uncharacterized protein
Uncharacterized protein
Uncharacterized protein
Uncharacterized protein
Uncharacterized protein
Uncharacterized protein
Uncharacterized protein
Uncharacterized protein
Uncharacterized protein
Uncharacterized protein
Uncharacterized protein
Uncharacterized protein
Uncharacterized protein
Uncharacterized protein
Uncharacterized protein
Uncharacterized protein
Uncharacterized protein
Uncharacterized protein
Uncharacterized protein
Uncharacterized protein
Putative serine/threonine-protein kinase RLK1 (Fragment)
Uncharacterized protein
Uncharacterized protein
Uncharacterized protein
Uncharacterized protein
Uncharacterized protein
Uncharacterized protein
Uncharacterized protein
Uncharacterized protein
Uncharacterized protein
Uncharacterized protein
Uncharacterized protein

Uncharacterized protein
Uncharacterized protein
Uncharacterized protein
Uncharacterized protein
Uncharacterized protein
Uncharacterized protein
Uncharacterized protein
Uncharacterized protein
Uncharacterized protein
Uncharacterized protein
Uncharacterized protein
Uncharacterized protein
Uncharacterized protein
Uncharacterized protein
Uncharacterized protein
Uncharacterized protein
Uncharacterized protein
Uncharacterized protein
Uncharacterized protein
Uncharacterized protein
Uncharacterized protein
Uncharacterized protein
Uncharacterized protein
Uncharacterized protein
Uncharacterized protein
Uncharacterized protein
Uncharacterized protein
Uncharacterized protein
Uncharacterized protein
Uncharacterized protein
Uncharacterized protein
Uncharacterized protein
Uncharacterized protein
Uncharacterized protein
Uncharacterized protein
Uncharacterized protein
Uncharacterized protein
Uncharacterized protein
Uncharacterized protein
Uncharacterized protein
Uncharacterized protein

Uncharacterized protein
Uncharacterized protein
Uncharacterized protein
Uncharacterized protein
Uncharacterized protein
Uncharacterized protein
Uncharacterized protein
Uncharacterized protein
Uncharacterized protein
Uncharacterized protein (Fragment)
Uncharacterized protein (Fragment)
Uncharacterized protein (Fragment)
Uncharacterized protein (Fragment)
Uncharacterized protein (Fragment)
Uncharacterized protein (Fragment)
Uncharacterized protein (Fragment)
Uncharacterized protein (Fragment)
Uncharacterized protein
WGS project CABT00000000 data, contig 2.14
WGS project CABT00000000 data, contig 2.4
WGS project CABT00000000 data, contig 2.22
Uncharacterized protein
Uncharacterized protein
Uncharacterized protein
Uncharacterized protein
Uncharacterized protein
Uncharacterized protein
Uncharacterized protein
Uncharacterized protein
Uncharacterized protein
Uncharacterized protein
Uncharacterized protein
Uncharacterized protein
Uncharacterized protein
Uncharacterized protein
Uncharacterized protein
Uncharacterized protein
Uncharacterized protein
Uncharacterized protein
Uncharacterized protein
Uncharacterized protein

Uncharacterized protein
Uncharacterized protein
Uncharacterized protein
Uncharacterized protein (Fragment)
Uncharacterized protein
Uncharacterized protein
Uncharacterized protein
Uncharacterized protein
Uncharacterized protein
Uncharacterized protein
Uncharacterized protein
Uncharacterized protein
Uncharacterized protein
Uncharacterized protein
Uncharacterized protein
Uncharacterized protein
Uncharacterized protein
Uncharacterized protein
Uncharacterized protein
Uncharacterized protein
Uncharacterized protein
Uncharacterized protein
Uncharacterized protein (Fragment)
Uncharacterized protein
Uncharacterized protein
Uncharacterized protein
Uncharacterized protein
Uncharacterized protein
Uncharacterized protein
Uncharacterized protein
Uncharacterized protein
Uncharacterized protein
Uncharacterized protein
Uncharacterized protein
Uncharacterized protein
Uncharacterized protein
Uncharacterized protein
Uncharacterized protein
Uncharacterized protein
Uncharacterized protein
Uncharacterized protein

Uncharacterized protein
Uncharacterized protein
Uncharacterized protein
Uncharacterized protein
Uncharacterized protein
Uncharacterized protein
Uncharacterized protein
Uncharacterized protein
Uncharacterized protein
Uncharacterized protein
Uncharacterized protein
Uncharacterized protein
Uncharacterized protein
Uncharacterized protein
Uncharacterized protein
Uncharacterized protein
Uncharacterized protein
Uncharacterized protein
Uncharacterized protein
Uncharacterized protein
Uncharacterized protein
Uncharacterized protein
Uncharacterized protein
Uncharacterized protein
Uncharacterized protein
Uncharacterized protein
Uncharacterized protein
Uncharacterized protein
Uncharacterized protein
Uncharacterized protein
Uncharacterized protein
Uncharacterized protein
Uncharacterized protein
Uncharacterized protein
Uncharacterized protein
Uncharacterized protein
Receptor-like kinase
Uncharacterized protein
Uncharacterized protein
Uncharacterized protein
Uncharacterized protein

Uncharacterized protein
Uncharacterized protein
Uncharacterized protein
Uncharacterized protein
Uncharacterized protein
Uncharacterized protein
Uncharacterized protein
26S proteasome non-ATPase regulatory subunit 9
Protein TOS1
WSC domain-containing protein 2
Uncharacterized protein
Uncharacterized protein
Uncharacterized protein
Uncharacterized protein
Uncharacterized protein (Fragment)
Uncharacterized protein
Uncharacterized protein
Serine protease
Uncharacterized protein
Uncharacterized protein
Uncharacterized protein
Uncharacterized protein
Uncharacterized protein
Uncharacterized protein
Uncharacterized protein
Uncharacterized protein
Uncharacterized protein (Fragment)
Uncharacterized protein
Uncharacterized protein
Uncharacterized protein
Uncharacterized protein (Fragment)
Uncharacterized protein
Uncharacterized protein
Uncharacterized protein
Uncharacterized protein
Uncharacterized protein
Uncharacterized protein
Uncharacterized protein
Uncharacterized protein
Uncharacterized protein
Uncharacterized protein

Uncharacterized protein
Uncharacterized protein
Uncharacterized protein
Uncharacterized protein
Uncharacterized protein
Uncharacterized protein
Uncharacterized protein
Uncharacterized protein
SFRICE_003747
SFRICE_003772 (Fragment)
SFRICE_004066
SFRICE_012857 (Fragment)
SFRICE_006749
Uncharacterized protein
Uncharacterized protein
Uncharacterized protein
Apple-like protein
PAN-1 domain protein
Uncharacterized protein
Uncharacterized protein
Uncharacterized protein
Uncharacterized protein
Uncharacterized protein
Uncharacterized protein
Uncharacterized protein
Uncharacterized protein
Uncharacterized protein
Uncharacterized protein
Uncharacterized protein
Uncharacterized protein
Uncharacterized protein
Uncharacterized protein
Galactose oxidase-like protein
Uncharacterized protein
Uncharacterized protein
Uncharacterized protein
Galactose oxidase-like protein
Uncharacterized protein
Uncharacterized protein
Uncharacterized protein
Large extracellular alpha-helical protein

Uncharacterized protein
Uncharacterized protein
Alpha-2-macroglobulin
Uncharacterized protein
Uncharacterized protein
Plasminogen
Uncharacterized protein
Hepatocyte growth factor
Uncharacterized protein
Uncharacterized protein
Uncharacterized protein (Fragment)
Uncharacterized protein (Fragment)
Uncharacterized protein (Fragment)
Uncharacterized protein (Fragment)
Uncharacterized protein (Fragment)
Uncharacterized protein (Fragment)
Uncharacterized protein (Fragment)
Uncharacterized protein (Fragment)
Uncharacterized protein (Fragment)
Uncharacterized protein (Fragment)
Uncharacterized protein (Fragment)
Uncharacterized protein (Fragment)
Uncharacterized protein (Fragment)
Uncharacterized protein (Fragment)
Uncharacterized protein (Fragment)
Uncharacterized protein (Fragment)
Uncharacterized protein (Fragment)
Uncharacterized protein (Fragment)
Uncharacterized protein (Fragment)
Uncharacterized protein (Fragment)
Uncharacterized protein (Fragment)
Uncharacterized protein
Uncharacterized protein
Uncharacterized protein
Uncharacterized protein
Uncharacterized protein
Uncharacterized protein
Uncharacterized protein
Uncharacterized protein
Uncharacterized protein
Uncharacterized protein

Uncharacterized protein
Uncharacterized protein
Uncharacterized protein
Uncharacterized protein
Uncharacterized protein
Uncharacterized protein
Uncharacterized protein
Uncharacterized protein
Uncharacterized protein
Uncharacterized protein
Uncharacterized protein
Uncharacterized protein
Uncharacterized protein
Uncharacterized protein
Uncharacterized protein
Uncharacterized protein
Uncharacterized protein
Uncharacterized protein
Uncharacterized protein
Uncharacterized protein
Copper radical oxidase
Cytochrome p450 4a10
Sequence-specific DNA binding RNA polymerase II transcription factor
Putative microfibril-associated glycoprotein 4
Uncharacterized protein
Uncharacterized protein
Uncharacterized protein (Fragment)
Uncharacterized protein
Putative deleted in malignant brain tumors 1 protein isoform X2
Putative macrophage mannose receptor 1-like
Uncharacterized protein
Uncharacterized protein
Putative plasminogen-like
Uncharacterized protein
Uncharacterized protein
Uncharacterized protein
Uncharacterized protein
Uncharacterized protein
Uncharacterized protein
Uncharacterized protein
Uncharacterized protein

Putative microfibril-associated glycoprotein 4
Putative macrophage mannose receptor 1
Uncharacterized protein
Uncharacterized protein
Uncharacterized protein
Uncharacterized protein
Uncharacterized protein
Uncharacterized protein
Uncharacterized protein
Uncharacterized protein
Uncharacterized protein
Uncharacterized protein
Uncharacterized protein
Uncharacterized protein
Uncharacterized protein
Uncharacterized protein
Uncharacterized protein
Uncharacterized protein
Uncharacterized protein
Uncharacterized protein
Uncharacterized protein
Uncharacterized protein
Uncharacterized protein
Uncharacterized protein
Uncharacterized protein
Uncharacterized protein
Uncharacterized protein
Uncharacterized protein
Uncharacterized protein
Uncharacterized protein
Uncharacterized protein
Uncharacterized protein
Uncharacterized protein
Uncharacterized protein
Uncharacterized protein
Uncharacterized protein
Uncharacterized protein
Uncharacterized protein
Uncharacterized protein
Uncharacterized protein
Uncharacterized protein

Uncharacterized protein
Uncharacterized protein
Uncharacterized protein
Uncharacterized protein
Uncharacterized protein
Uncharacterized protein
Uncharacterized protein
Uncharacterized protein
Uncharacterized protein
Uncharacterized protein
Uncharacterized protein
Uncharacterized protein
Uncharacterized protein
Uncharacterized protein
Uncharacterized protein
Uncharacterized protein
Uncharacterized protein
Uncharacterized protein
Uncharacterized protein
Uncharacterized protein
Uncharacterized protein
Uncharacterized protein
Uncharacterized protein
Uncharacterized protein
Uncharacterized protein
Uncharacterized protein
Uncharacterized protein
Uncharacterized protein
Uncharacterized protein
Uncharacterized protein
Uncharacterized protein
Uncharacterized protein
Uncharacterized protein
Uncharacterized protein
Uncharacterized protein
Uncharacterized protein
Uncharacterized protein
Uncharacterized protein
Uncharacterized protein
Uncharacterized protein
Uncharacterized protein

Uncharacterized protein
Uncharacterized protein
Uncharacterized protein
Uncharacterized protein
Uncharacterized protein
Uncharacterized protein
Uncharacterized protein
Uncharacterized protein
Uncharacterized protein
Uncharacterized protein
Uncharacterized protein
Uncharacterized protein
Uncharacterized protein
Uncharacterized protein
Uncharacterized protein
Uncharacterized protein
Uncharacterized protein
Uncharacterized protein
Uncharacterized protein
Uncharacterized protein
Uncharacterized protein
PAN-1 domain and Apple-like domain-containing protein
Major sperm protein
PAN-1 domain and Apple-like domain-containing protein
PAN-1 domain and Apple-like domain-containing protein
PAN-1 domain and Apple-like domain-containing protein
Thrombospondin, type 1 repeat and Apple-like domain-containing protein
Zona pellucida domain and PAN-1 domain and Apple-like domain-containing protein
PAN-1 domain and Apple-like domain-containing protein
No mechanoreceptor potential A
PAN-1 domain and Apple-like domain-containing protein
PAN-1 domain and Apple-like domain-containing protein
Tequila
Apple-like domain-containing protein
C-type lectin domain and von Willebrand factor, type A domain and PAN-1 domain and Apple-like doma
PAN-1 domain and Apple-like domain-containing protein
PAN-1 domain and Apple-like domain-containing protein
No mechanoreceptor potential A
PAN-1 domain and Apple-like domain-containing protein
PAN-1 domain and Apple-like domain-containing protein
No mechanoreceptor potential A

Jitterbug
Uncharacterized protein
Uncharacterized protein
Uncharacterized protein
Uncharacterized protein
Uncharacterized protein
Uncharacterized protein
Uncharacterized protein
Uncharacterized protein
Uncharacterized protein
Uncharacterized protein
Uncharacterized protein
Uncharacterized protein
Uncharacterized protein
Uncharacterized protein
Uncharacterized protein
Uncharacterized protein
Uncharacterized protein
Uncharacterized protein
Uncharacterized protein
Uncharacterized protein
Uncharacterized protein
Uncharacterized protein
Uncharacterized protein
Uncharacterized protein
Uncharacterized protein
Uncharacterized protein
Uncharacterized protein
Uncharacterized protein
Uncharacterized protein
Uncharacterized protein
Uncharacterized protein
HGF/MSP/plasminogen-like protein (inferred by orthology to a C. elegans protein)
Uncharacterized protein
Uncharacterized protein
Uncharacterized protein
Properdin (inferred by orthology to a human protein)
Uncharacterized protein
Let-653 protein (projected from Caenorhabditis elegans ortholog let-653)
No mechanoreceptor potential A (inferred by orthology to a D. melanogaster protein)
Uncharacterized protein

Uncharacterized protein
Uncharacterized protein (Fragment)
Uncharacterized protein (Fragment)
Uncharacterized protein
Uncharacterized protein
Uncharacterized protein
Uncharacterized protein (Fragment)
Uncharacterized protein
Uncharacterized protein
Uncharacterized protein
Uncharacterized protein (Fragment)
Uncharacterized protein (Fragment)
Uncharacterized protein
Uncharacterized protein (Fragment)
Uncharacterized protein (Fragment)
Uncharacterized protein
Uncharacterized protein
Uncharacterized protein
Uncharacterized protein
Uncharacterized protein
Uncharacterized protein
Uncharacterized protein
Plasminogen (Fragment)
Plasma kallikrein (Fragment)
Hepatocyte growth factor (Fragment)
Lactadherin
Lactadherin
Uncharacterized protein
Uncharacterized protein (Fragment)
Ubiquitin carboxyl-terminal hydrolase 10
Uncharacterized protein
Uncharacterized protein
Ficolin-1
Matrilin-4
Nidogen-1
Fibrinogen beta chain
Uncharacterized protein
Uncharacterized protein
Uncharacterized protein (Fragment)
Contactin-associated protein-like 2 (Fragment)
Techylectin-5B

Uncharacterized protein
Neurogenic locus notch-like protein 3
Uncharacterized protein
Uncharacterized protein
Ficolin-1
Neuropilin-2
Ryncolin-2
Pentraxin-4
EGF-like domain-containing protein
Uncharacterized protein
Olfactomedin-like protein 2A
Uncharacterized protein
Uncharacterized protein
Ficolin-1
Uncharacterized protein
Uncharacterized protein
Ficolin-1
Sushi, von Willebrand factor type A, EGF and pentraxin domain-containing protein 1
Fibrinogen C domain-containing protein 1 (Fragment)
Uncharacterized protein
Uncharacterized protein
Uncharacterized protein
Ficolin-1
Neurogenic locus notch-like protein 1
Ryncolin-1
Uncharacterized protein
Uncharacterized protein
Angiopoietin-4
Uncharacterized protein
Neuronal pentraxin-2
Tetratricopeptide repeat protein 28
Fibrillin-2
Uncharacterized protein
Fibrinogen beta chain
Uncharacterized protein
Pikachurin
EGF-like repeat and discoidin I-like domain-containing protein 3
Uromodulin
Uncharacterized protein
Protocadherin Fat 3
Uncharacterized protein

Neuropilin-2
EGF-like domain-containing protein
Ficolin-1
Sushi, von Willebrand factor type A, EGF and pentraxin domain-containing protein 1
Pikachurin
Ficolin-2
Lactadherin
Uncharacterized transposon-derived protein F52C9.6 (Fragment)
Uncharacterized protein
Uncharacterized protein
Hemicentin-1
Fibrillin-1
Fibroleukin
Uncharacterized protein
Neurogenic locus Notch protein
Fibrillin-2
Uncharacterized protein (Fragment)
Fibrillin-2 (Fragment)
Uncharacterized protein
Uncharacterized protein
Uncharacterized protein
Neurocan core protein
Uncharacterized protein
Neurogenic locus Notch protein
EGF-like repeat and discoidin I-like domain-containing protein 3
Fibrillin-2
Uromodulin
Ryncolin-1
Uncharacterized protein
Fibrillin-1
Uncharacterized protein
Fibroleukin
Uncharacterized protein
C-type lectin domain family 12 member B
Uncharacterized protein
Neurogenic locus Notch protein
Uncharacterized protein
Ficolin-2
Signal peptide, CUB and EGF-like domain-containing protein 2
Oncoprotein-induced transcript 3 protein
Collagen alpha-2(V) chain

Ficolin-1
Ficolin-1
Fibrillin-1
Neuronal pentraxin-2
Uncharacterized protein
Uncharacterized protein
Protein HEG-like 1
Uncharacterized protein
Uncharacterized protein
Versican core protein
Uncharacterized protein
Uncharacterized protein
Contactin-associated protein-like 2
Uncharacterized protein
Uncharacterized protein
EGF-like repeat and discoidin I-like domain-containing protein 3
Fibrillin-1
Aggrecan core protein
Ficolin-1
Uncharacterized protein
Fibrinogen beta chain
Uncharacterized protein
Sushi, von Willebrand factor type A, EGF and pentraxin domain-containing protein 1
Fibrillin-2
Scavenger receptor cysteine-rich type 1 protein M130
EGF-like repeat and discoidin I-like domain-containing protein 3
A disintegrin and metalloproteinase with thrombospondin motifs 9
WD repeat-containing protein 5
Pancreatic secretory granule membrane major glycoprotein GP2
Lactadherin
Perivitellin-2 67 kDa subunit
Uncharacterized protein
Uncharacterized protein
Uncharacterized protein
Venom prothrombin activator oscutarin-C non-catalytic subunit
Ficolin-1
C-reactive protein 1.4
Ficolin-2
Uncharacterized protein
Perlucin-like protein
Sushi, von Willebrand factor type A, EGF and pentraxin domain-containing protein 1

Uncharacterized protein
Sphingomyelinase phosphodiesterase D
Epidermal growth factor-like protein 6
Beta-1,3-N-acetylglucosaminyltransferase lunatic fringe
Uncharacterized protein
Coagulation factor V
Fibrillin-1
Fibrinogen beta chain
Uncharacterized protein
Phosphoinositide phospholipase C
RNA-directed DNA polymerase from mobile element jockey
Sushi, von Willebrand factor type A, EGF and pentraxin domain-containing protein 1
Galectin-3-binding protein A
Contactin-associated protein-like 2
Uncharacterized protein
Uncharacterized protein
PAN domain protein
PAN domain-containing protein
Uncharacterized protein
Uncharacterized protein
Uncharacterized protein
Putative lipoprotein YfhM
PAN domain-containing protein
PAN domain-containing protein
PAN domain-containing protein
Plasminogen
Hepatocyte growth factor-like protein
Hepatocyte growth factor
Uncharacterized protein
Uncharacterized protein
Kallikrein
Uncharacterized protein
Plasminogen
Uncharacterized protein
Uncharacterized protein
Uncharacterized protein
Uncharacterized protein
Protein aardvark
Uncharacterized protein
Endoglucanase
Uncharacterized protein

Uncharacterized protein
Tubulin beta chain
Uncharacterized protein
Uncharacterized protein
Multidrug resistance protein MdtL
Metalloendopeptidase
Putative membrane protein YlbC
Uncharacterized protein
Serine/threonine-protein kinasePKR2
Uncharacterized protein
Abnormal spindle-like microcephaly-associated protein-like
Carbohydrate-binding domain-containing protein C2E1P3.05c
Uncharacterized protein
Metalloendopeptidase
Uncharacterized protein
Uncharacterized protein
Uncharacterized protein
Uncharacterized protein
Uncharacterized protein
Uncharacterized protein
Uncharacterized protein
Putative glycosidase
Glutaredoxin-C3
Uncharacterized protein
Uncharacterized protein
Uncharacterized protein
Uncharacterized protein
Uncharacterized protein
Alkali-sensitive linkage protein 1
Uncharacterized protein
Formamidopyrimidine-DNA glycosylase
Uncharacterized protein
Uncharacterized protein
Uncharacterized protein
Calcium-dependent protein kinase 1
Uncharacterized protein
Uncharacterized protein
Uncharacterized protein
Uncharacterized protein
Uncharacterized protein
Leucine-rich repeat-containing protein LOC400891-like

Translation initiation factor IF-2
tRNA 2'-phosphotransferase 1
Uncharacterized protein
Inter-alpha-trypsin inhibitor heavy chain H4
Uncharacterized protein
Uncharacterized protein
Uncharacterized protein
Vegetative incompatibility protein HET-E-1
Uncharacterized protein
Uncharacterized protein
26S protease regulatory subunit 6A
Uncharacterized protein
Cation channel sperm-associated protein 1
Pentatricopeptide repeat-containing protein, mitochondrial
Metalloendopeptidase
Uncharacterized protein
Uncharacterized protein
Uncharacterized protein (Fragment)
Uncharacterized protein
Uncharacterized protein
Uncharacterized protein
Uncharacterized protein
Uncharacterized protein
Uncharacterized protein
Uncharacterized protein
Uncharacterized protein
Uncharacterized protein
Uncharacterized protein
Uncharacterized protein
Uncharacterized protein
Uncharacterized protein
Uncharacterized protein
Uncharacterized protein
Uncharacterized protein
Uncharacterized protein
Uncharacterized protein
Uncharacterized protein
Uncharacterized protein
Uncharacterized protein
Uncharacterized protein
Uncharacterized protein
Alpha-2-macroglobulin family protein

Alpha-2-macroglobulin family protein
PAN domain-containing protein
Uncharacterized protein
Uncharacterized protein
Uncharacterized protein
Uncharacterized protein
Hepatocyte growth factor
Uncharacterized protein
Plasminogen
Uncharacterized protein
Macrophage stimulating 1
Uncharacterized protein
Kalliklectin
Uncharacterized protein
Uncharacterized protein
Uncharacterized protein
Uncharacterized protein
Uncharacterized protein
Uncharacterized protein
Uncharacterized protein
Hepatocyte growth factor
Uncharacterized protein
Hepatocyte growth factor
Uncharacterized protein
Uncharacterized protein
Uncharacterized protein
Uncharacterized protein
Uncharacterized protein
Uncharacterized protein
Uncharacterized protein
Uncharacterized protein
Uncharacterized protein
Apple domain-containing protein
PAN domain-containing protein
PAN domain-containing protein
PAN domain-containing protein
PAN domain-containing protein
PAN domain-containing protein
hepatocyte growth factor isoform X5
Hepatocyte growth factor
plasma kallikrein

Plasminogen
hepatocyte growth factor-like protein
hepatocyte growth factor isoform X6
Hepatocyte growth factor
coagulation factor XI
Hepatocyte growth factor
Hepatocyte growth factor
PAN domain-containing protein
PAN domain-containing protein
Plasminogen (Fragment)
Plasminogen (Fragment)
Hepatocyte growth factor (Fragment)
Coagulation factor XI (Fragment)
PAN domain protein
PAN domain protein
PAN domain protein
PAN domain protein
PAN domain protein
PAN domain protein (Fragment)
PAN domain protein
Uncharacterized protein (Fragment)
PAN domain protein
PAN domain protein
PAN domain protein
PAN domain protein (Fragment)
PAN domain protein
PAN domain protein
Uncharacterized protein
PAN domain protein
PAN domain protein
PAN domain protein
PAN domain protein
PAN domain protein
PAN domain protein
Uncharacterized protein
Uncharacterized protein
Apple domain-containing protein
PAN/apple domain protein
Uncharacterized protein
Uncharacterized protein
Uncharacterized protein

Uncharacterized protein (Fragment)
Uncharacterized protein
Leucine-rich repeat receptor-like protein kinase PXL2
Uncharacterized protein
Peptidyl-prolyl cis-trans isomerase D
Peptidyl-prolyl cis-trans isomerase cyp5
Uncharacterized protein (Fragment)
Uncharacterized protein
Uncharacterized protein (Fragment)
Uncharacterized protein (Fragment)
Uncharacterized protein
Uncharacterized protein (Fragment)
Uncharacterized protein
Uncharacterized protein
Uncharacterized protein
Uncharacterized protein
Uncharacterized protein
Uncharacterized protein
Uncharacterized protein
Uncharacterized protein
Uncharacterized protein
Uncharacterized protein
Uncharacterized protein
Uncharacterized protein
Uncharacterized protein
Uncharacterized protein
Carboxy-terminal crystallin fold protein 12p
C-terminal crystallin fold containing protein 12p
Antigen B membrane protein
Uncharacterized protein
Uncharacterized protein
Uncharacterized protein
Uncharacterized protein
Uncharacterized protein
Uncharacterized protein
Uncharacterized protein
Uncharacterized protein
Uncharacterized protein
Uncharacterized protein
Uncharacterized protein
Uncharacterized protein

Uncharacterized protein
Uncharacterized protein
Uncharacterized protein
Uncharacterized protein
Uncharacterized protein
Uncharacterized protein
Uncharacterized protein
Uncharacterized protein
Uncharacterized protein
Uncharacterized protein
Uncharacterized protein
Uncharacterized protein
Uncharacterized protein
Uncharacterized protein
Uncharacterized protein
Uncharacterized protein
Uncharacterized protein
Uncharacterized protein
Uncharacterized protein
Uncharacterized protein
Uncharacterized protein
Uncharacterized protein
Uncharacterized protein
Uncharacterized protein
Uncharacterized protein
Uncharacterized protein
Uncharacterized protein
Uncharacterized protein
Uncharacterized protein
Uncharacterized protein
Uncharacterized protein
Uncharacterized protein
Uncharacterized protein
Uncharacterized protein
Uncharacterized protein
Uncharacterized protein
Uncharacterized protein
Uncharacterized protein
Uncharacterized protein
Uncharacterized protein
Uncharacterized protein

Uncharacterized protein
Uncharacterized protein
Uncharacterized protein
Uncharacterized protein
Uncharacterized protein
Uncharacterized protein
Uncharacterized protein
Uncharacterized protein
Uncharacterized protein
Uncharacterized protein
Chromosome 13 SCAF15122, whole genome shotgun sequence (Fragment)
Macrophage stimulating 1
Chromosome 11 SCAF14979, whole genome shotgun sequence
Plasminogen
Macrophage stimulating 1
Hepatocyte growth factor a
Chromosome 13 SCAF15122, whole genome shotgun sequence (Fragment)
Chromosome 21 SCAF14577, whole genome shotgun sequence
Chromosome 19 SCAF14664, whole genome shotgun sequence
Plasminogen
Hepatocyte growth factor
Uncharacterized protein
Plasminogen
Dihydropyridine-sensitive l-type calcium channel (Fragment)
Uncharacterized protein (Fragment)
Uncharacterized protein (Fragment)
Uncharacterized protein
Uncharacterized protein
Uncharacterized protein
Uncharacterized protein
Uncharacterized protein (Fragment)
Uncharacterized protein
Uncharacterized protein (Fragment)
M6 family peptidase
Uncharacterized protein
Uncharacterized protein
Uncharacterized protein
Uncharacterized protein (Fragment)
Uncharacterized protein
PAN domain-containing protein
PAN domain-containing protein

Uncharacterized protein (Fragment)
Uncharacterized protein (Fragment)
Uncharacterized protein
Uncharacterized protein (Fragment)
Uncharacterized protein (Fragment)
Uncharacterized protein (Fragment)
Uncharacterized protein
Uncharacterized protein (Fragment)
Uncharacterized protein (Fragment)
Uncharacterized protein
Uncharacterized protein
Alpha-2-macroglobulin family protein
Uncharacterized protein
Uncharacterized protein
Uncharacterized protein
Uncharacterized protein
Uncharacterized protein
Uncharacterized protein
Uncharacterized protein
Uncharacterized protein
Uncharacterized protein
Uncharacterized protein
Uncharacterized protein
Uncharacterized protein
Uncharacterized protein
Uncharacterized protein
Uncharacterized protein
Uncharacterized protein
Uncharacterized protein
Uncharacterized protein
Uncharacterized protein
Uncharacterized protein
Uncharacterized protein
Uncharacterized protein
Uncharacterized protein
Uncharacterized protein
Uncharacterized protein
Uncharacterized protein
S-locus lectin protein kinase family protein, putative isoform 1
S-locus lectin protein kinase family protein isoform 2
S-locus lectin protein kinase family protein, putative isoform 1

S-locus lectin protein kinase family protein isoform 1
Serine/threonine kinases,protein kinases,ATP binding,sugar binding,kinases,carbohydrate binding, puta
S-locus lectin protein kinase family protein, putative isoform 2
Serine/threonine kinases,protein kinases,ATP binding,sugar binding,kinases,carbohydrate binding, puta
Receptor-like protein kinase 4 isoform 1
Serine/threonine kinases,protein kinases,ATP binding,sugar binding,kinases,carbohydrate binding
S-locus lectin protein kinase family protein, putative
S-locus lectin protein kinase family protein, putative
Receptor-like protein kinase 1
S-locus-specific glycoprotein S6
Serine/threonine kinases,protein kinases,ATP binding,sugar binding,kinases,carbohydrate binding
S-locus lectin protein kinase family protein, putative
S-locus lectin protein kinase family protein, putative
Receptor-like protein kinase 4 isoform 2
S-locus lectin protein kinase family protein, putative
S-locus lectin protein kinase family protein, putative
S-locus lectin protein kinase family protein, putative
S-locus lectin protein kinase family protein
S-locus lectin protein kinase family protein, putative
S-locus lectin protein kinase family protein, putative isoform 2 (Fragment)
Receptor-like protein kinase 1
S-locus lectin protein kinase family protein
Serine/threonine kinases,protein kinases,ATP binding,sugar binding,kinases,carbohydrate binding, puta
S-domain-2 5 isoform 1
S-domain-2 5 isoform 2
S-locus lectin protein kinase family protein, putative isoform 6
S-domain-2 5, putative
Receptor protein kinase 4
S-locus lectin protein kinase family protein, putative
CCHC-type integrase, putative
Serine/threonine kinases,protein kinases,ATP binding,sugar binding,kinases,carbohydrate binding, puta
Serine/threonine kinases,protein kinases,ATP binding,sugar binding,kinases,carbohydrate binding, puta
Serine/threonine-protein kinase receptor, putative
S-locus lectin protein kinase family protein, putative
S-locus-specific glycoprotein S6, putative
Serine/threonine kinases,protein kinases,ATP binding,sugar binding,kinases,carbohydrate binding, puta
S-locus lectin protein kinase family protein, putative isoform 5
Receptor kinase 3 isoform 1
S-locus lectin protein kinase family protein, putative
S-locus lectin protein kinase family protein, putative
Uncharacterized protein

Serine/threonine-protein kinase PBS1, putative
S-locus-specific glycoprotein S6, putative
S-locus lectin protein kinase family protein, putative
Receptor protein kinase 4
S-locus lectin protein kinase family protein isoform 2 (Fragment)
S-locus lectin protein kinase family protein, putative isoform 3
S-locus lectin protein kinase family protein isoform 1
S-locus lectin protein kinase family protein, putative
S-locus-specific glycoprotein S6
Serine/threonine kinases,protein kinases,ATP binding,sugar binding,kinases,carbohydrate binding, puta
S-locus lectin protein kinase family protein, putative
G-type lectin S-receptor serine/threonine-protein kinase SD3-1
S-locus lectin protein kinase family protein, putative
S-domain-2 5, putative
S-locus-specific glycoprotein S6, putative
S-locus lectin protein kinase family protein
S-locus lectin protein kinase family protein isoform 4
S-locus lectin protein kinase family protein, putative
S-locus lectin protein kinase family protein
Receptor kinase 3
S-locus lectin protein kinase family protein
Serine-threonine protein kinase, plant-type
Curculin-like lectin family protein / PAN domain-containing protein, putative
S-locus lectin protein kinase family protein
Serine/threonine kinases,protein kinases,ATP binding,sugar binding,kinases,carbohydrate binding, puta
S-locus lectin protein kinase family protein, putative isoform 1
Serine/threonine kinases,protein kinases,ATP binding,sugar binding,kinases,carbohydrate binding, puta
S-locus lectin protein kinase family protein
S-locus lectin protein kinase family protein, putative
S-locus lectin protein kinase family protein, putative isoform 1
S-locus lectin protein kinase family protein
S-locus lectin protein kinase family protein isoform 1
S-locus lectin protein kinase family protein, putative
S-locus lectin protein kinase family protein isoform 1
S-locus lectin protein kinase family protein, putative
S-locus lectin protein kinase family protein, putative isoform 1
S-locus lectin protein kinase family protein, putative isoform 2
S-locus lectin protein kinase family protein, putative
S-locus lectin protein kinase family protein, putative isoform 2 (Fragment)
S-locus lectin protein kinase family protein
S-locus lectin protein kinase family protein, putative

S-locus lectin protein kinase family protein
S-locus lectin protein kinase family protein
Receptor kinase 3 isoform 2 (Fragment)
S-locus lectin protein kinase family protein, putative
S-locus lectin protein kinase family protein, putative
S-locus lectin protein kinase family protein, putative
S-locus lectin protein kinase family protein, putative isoform 2
Uncharacterized protein (Fragment)
S-locus lectin protein kinase family protein, putative
S-locus lectin protein kinase family protein isoform 2
S-domain-2 5
Uncharacterized protein
Serine/threonine kinases,protein kinases,ATP binding,sugar binding,kinases,carbohydrate binding
S-locus lectin protein kinase family protein, putative
S-locus lectin protein kinase family protein, putative isoform 4
S-locus lectin protein kinase family protein, putative isoform 3
Serine/threonine-protein kinase PBS1, putative
D-mannose binding lectin protein with Apple-like carbohydrate-binding domain, putative
S-locus lectin protein kinase family protein, putative isoform 1
Receptor kinase 3, putative
Serine-threonine protein kinase, plant-type, putative
Uncharacterized protein
Uncharacterized protein
Uncharacterized protein
Uncharacterized protein
B70aff9a-902b-487a-8f75-5a5b7bcaae2e
Alpha-2-macroglobulin domain protein
Uncharacterized protein
Uncharacterized protein
Large extracellular alpha-helical protein
Large extracellular alpha-helical protein
Alpha-2-macroglobulin domain protein
PAN domain protein
Secreted protein
Secreted protein
Secreted protein
Carbohydrate-binding protein
Secreted protein (Fragment)
Uncharacterized protein
Carbohydrate-binding protein
Uncharacterized protein

Secreted protein
Uncharacterized protein (Fragment)
Secreted protein
Uncharacterized protein
Endonuclease
Uncharacterized protein
Secreted protein (Fragment)
Uncharacterized protein
Carbohydrate-binding protein (Fragment)
Secreted protein (Fragment)
Secreted protein
Secreted protein
Uncharacterized protein
Glucanase
Secreted protein
Uncharacterized protein
Secreted protein
Acyl-CoA dehydrogenase/oxidase
Uncharacterized protein
Carbohydrate-binding protein (Fragment)
Secreted protein
Uncharacterized protein
Uncharacterized protein
Uncharacterized protein
Glucanase
Secreted protein
Uncharacterized protein (Fragment)
Uncharacterized protein
Uncharacterized protein (Fragment)
Uncharacterized protein
Carbohydrate-binding protein
Uncharacterized protein
Cysteine protease family C01A
GPI anchor protein
Secreted protein
Secreted protein
Uncharacterized protein
Uncharacterized protein
Uncharacterized protein
Hepatocyte growth factor (Fragment)
Plasma kallikrein (Fragment)

Plasminogen (Fragment)
Plasminogen (Fragment)
Putative atp-dependent rrna helicase rrp3 protein
Putative glycosyltransferase family 31 protein
Uncharacterized protein
Uncharacterized protein
Uncharacterized protein
Uncharacterized protein
Uncharacterized protein
Uncharacterized protein
Uncharacterized protein
Uncharacterized protein
Uncharacterized protein
Uncharacterized protein
Uncharacterized protein
Uncharacterized protein
Uncharacterized protein
Uncharacterized protein
Uncharacterized protein
Uncharacterized protein
Uncharacterized protein
Uncharacterized protein
Uncharacterized protein
Uncharacterized protein
Uncharacterized protein
Uncharacterized protein
Uncharacterized protein
Uncharacterized protein
Uncharacterized protein
Uncharacterized protein
Uncharacterized protein
Uncharacterized protein
Uncharacterized protein
Uncharacterized protein F52C9.5
Uncharacterized protein F52C9.5
Paternally-expressed protein
Transient receptor potential cation channel trpm
Uncharacterized protein F52C9.5
Uncharacterized protein C34C12.7
Uncharacterized protein (Fragment)
Uncharacterized protein F52C9.5

Uncharacterized protein
Uncharacterized protein
Uncharacterized protein
Uncharacterized protein
CRAL-TRIO domain-containing protein C34C12.6
Uncharacterized protein
Uncharacterized protein
Transmembrane protease serine 9
Uncharacterized protein
Uncharacterized protein
Uncharacterized protein
Uncharacterized protein
Uncharacterized protein F52C9.5
Uncharacterized protein F52C9.5
Uncharacterized protein
Uncharacterized protein
Uncharacterized protein
Uncharacterized protein
Uncharacterized protein (Fragment)
Uncharacterized protein
Uncharacterized protein
Uncharacterized protein
Microneme protein, putative
PAN domain-containing protein
Microneme protein MIC4
Microneme protein MIC17C
PAN domain-containing protein
Uncharacterized protein
Subtilisin SUB12
Uncharacterized protein
PAN domain-containing protein
PAN domain-containing protein
PAN domain-containing protein
Microneme protein MIC17B
1,3(4)-beta-glucanase
PAN domain-containing protein
PAN domain-containing protein
Microneme protein MIC17A
PAN/Apple domain-containing protein
PAN/Apple domain-containing protein
PAN domain-containing protein

Microneme protein MIC17A
PAN domain-containing protein
PAN domain-containing protein
PAN domain-containing protein (Fragment)
Subtilisin SUB12
1,3(4)-beta-glucanase
PAN domain-containing protein
Uncharacterized protein
PAN/Apple domain-containing protein
PAN domain-containing protein
PAN domain protein
PAN/Apple domain-containing protein
PAN domain-containing protein
PAN domain-containing protein
Uncharacterized protein
PAN domain-containing protein (Fragment)
Putative microneme protein
Microneme protein MIC17B
Microneme protein MIC4
Microneme protein MIC17C
PAN domain-containing protein
PAN domain-containing protein, putative
Microneme protein, putative
Microneme protein MIC17B
Microneme protein MIC4
PAN domain-containing protein
PAN domain-containing protein
PAN domain-containing protein
Putative microneme protein
PAN domain protein
Subtilisin SUB12
PAN domain-containing protein
PAN/Apple domain-containing protein
PAN domain-containing protein (Fragment)
Microneme protein MIC17A
PAN domain protein
PAN domain-containing protein
PAN domain-containing protein
PAN domain-containing protein
PAN domain-containing protein
PAN domain protein

PAN domain containing protein
Subtilisin-like serine protease, putative
PAN domain-containing protein
PAN domain-containing protein
1,3(4)-beta-glucanase
PAN domain-containing protein (Fragment)
PAN domain-containing protein
PAN domain-containing protein
PAN domain-containing protein
PAN domain-containing protein (Fragment)
PAN domain-containing protein
PAN domain protein
1,3(4)-beta-glucanase
PAN domain-containing protein
Putative microneme protein
PAN domain-containing protein (Fragment)
PAN domain protein
Subtilisin SUB12
PAN domain-containing protein
PAN domain protein (Fragment)
PAN domain protein
Microneme protein MIC4
Putative microneme protein (Fragment)
PAN domain-containing protein (Fragment)
PAN domain-containing protein (Fragment)
PAN domain-containing protein
PAN/Apple domain-containing protein
PAN domain-containing protein
PAN domain-containing protein (Fragment)
PAN/Apple domain-containing protein (Fragment)
PAN domain-containing protein
Microneme protein MIC17A (Fragment)
PAN domain-containing protein (Fragment)
PAN/Apple domain-containing protein
Putative microneme protein (Fragment)
PAN domain-containing protein
Microneme protein MIC17B
PAN domain-containing protein
PAN domain-containing protein
PAN/Apple domain-containing protein
Subtilisin SUB12

PAN domain-containing protein
PAN domain-containing protein
PAN domain protein
Putative microneme protein
PAN domain-containing protein (Fragment)
PAN domain protein
1,3(4)-beta-glucanase
Microneme protein MIC4
PAN domain protein
PAN domain-containing protein (Fragment)
Putative microneme protein
PAN domain-containing protein
PAN domain-containing protein
PAN domain-containing protein
Subtilisin SUB12
1,3(4)-beta-glucanase
PAN domain protein
Microneme protein MIC17A (Fragment)
PAN domain-containing protein (Fragment)
PAN domain-containing protein
PAN/Apple domain-containing protein
Microneme protein MIC4
PAN domain-containing protein (Fragment)
PAN domain protein
Putative microneme protein (Fragment)
PAN domain protein
PAN domain-containing protein
PAN domain protein (Fragment)
Putative microneme protein
Microneme protein MIC4
PAN/Apple domain-containing protein (Fragment)
PAN/Apple domain-containing protein
PAN domain-containing protein (Fragment)
PAN domain-containing protein
PAN domain protein
PAN domain-containing protein
PAN domain-containing protein
PAN domain protein
1,3(4)-beta-glucanase
PAN domain protein
PAN domain-containing protein (Fragment)

Subtilisin SUB12
PAN domain-containing protein
PAN domain protein
1,3(4)-beta-glucanase
PAN domain-containing protein
PAN domain-containing protein
PAN domain protein (Fragment)
Putative microneme protein
PAN domain-containing protein (Fragment)
PAN domain-containing protein
Putative PAN domain protein
Subtilisin SUB12
PAN domain protein (Fragment)
PAN/Apple domain-containing protein
PAN domain protein (Fragment)
PAN domain-containing protein
Microneme protein MIC4
PAN domain-containing protein (Fragment)
PAN domain-containing protein
PAN domain protein
PAN domain protein
PAN domain-containing protein (Fragment)
Putative microneme protein
PAN domain-containing protein
1,3(4)-beta-glucanase
PAN domain-containing protein
PAN domain protein (Fragment)
PAN domain-containing protein (Fragment)
Subtilisin SUB12 (Fragment)
PAN/Apple domain-containing protein
Microneme protein MIC4
PAN domain-containing protein (Fragment)
PAN domain-containing protein
PAN domain protein
Uncharacterized protein
PAN domain-containing protein
PAN domain-containing protein
PAN domain-containing protein (Fragment)
Putative microneme protein
PAN domain protein
Microneme protein MIC4

1,3(4)-beta-glucanase
Putative microneme protein (Fragment)
PAN domain-containing protein
PAN domain protein
Putative microneme protein (Fragment)
PAN domain-containing protein
Subtilisin SUB12
PAN/Apple domain-containing protein
Microneme protein MIC17C (Fragment)
PAN domain-containing protein (Fragment)
PAN domain protein
PAN domain-containing protein
PAN domain-containing protein (Fragment)
Microneme protein MIC4
PAN domain protein (Fragment)
PAN domain protein
PAN domain protein (Fragment)
Subtilisin SUB12 (Fragment)
1,3(4)-beta-glucanase
PAN domain protein
PAN domain protein
Putative microneme protein
PAN domain-containing protein
PAN domain-containing protein
Putative microneme protein (Fragment)
PAN/Apple domain-containing protein
PAN domain-containing protein (Fragment)
PAN domain-containing protein
PAN domain-containing protein (Fragment)
PAN domain protein
Microneme protein MIC17A
PAN/Apple domain-containing protein
PAN domain-containing protein (Fragment)
PAN domain-containing protein
PAN domain protein
PAN domain protein
Putative microneme protein
PAN domain protein
PAN domain-containing protein
PAN domain protein (Fragment)
PAN domain-containing protein (Fragment)

PAN/Apple domain-containing protein
PAN domain-containing protein
Putative microneme protein (Fragment)
Microneme protein MIC4
1,3(4)-beta-glucanase
PAN domain-containing protein
Subtilisin SUB12
PAN domain protein
PAN/Apple domain-containing protein
PAN domain-containing protein
PAN domain-containing protein (Fragment)
Subtilisin SUB12 (Fragment)
PAN domain-containing protein (Fragment)
PAN domain-containing protein (Fragment)
PAN domain-containing protein
PAN domain-containing protein
Putative microneme protein
PAN domain protein (Fragment)
PAN domain protein
PAN domain-containing protein (Fragment)
PAN domain protein
PAN domain-containing protein
1,3(4)-beta-glucanase
Microneme protein MIC4
Subtilisin SUB12
1,3(4)-beta-glucanase
PAN domain-containing protein
Microneme protein MIC17A
PAN domain-containing protein (Fragment)
PAN domain-containing protein (Fragment)
PAN domain protein
PAN/Apple domain-containing protein
PAN domain-containing protein (Fragment)
Putative microneme protein (Fragment)
PAN domain protein
PAN domain protein (Fragment)
PAN/Apple domain-containing protein
PAN domain protein (Fragment)
Microneme protein MIC4
PAN domain-containing protein (Fragment)
Putative microneme protein

PAN domain protein
PAN domain-containing protein
PAN domain-containing protein
Genomic DNA chromosome Ib
Micronemal protein MIC4
Micronemal protein MIC4
Micronemal protein MIC4
Micronemal protein MIC4
Micronemal protein MIC4
Micronemal protein MIC4
Micronemal protein MIC4
Micronemal protein MIC4
Micronemal protein 4, putative
Micronemal protein MIC4
Micronemal protein MIC4
Micronemal protein 4
Micronemal protein MIC4
Hepatocyte growth factor homologue (Fragment)
Uncharacterized protein
Uncharacterized protein
Uncharacterized protein
Uncharacterized protein
Cuticlin-1 (Fragment)
Neurotrypsin
Uncharacterized protein (Fragment)
Uncharacterized protein
Uncharacterized protein
Cuticlin-1
Neurotrypsin (Fragment)
Uncharacterized protein
Uncharacterized protein
Uncharacterized protein
Uncharacterized protein
Neurotrypsin (Fragment)
Uncharacterized protein
Uncharacterized protein
Circumsporozoite protein (Fragment)
Cuticlin-1
Alpha-2-macroglobulin family protein
S-receptor-like serine/threonine-protein kinase
Mitogen-activated protein kinase kinase kinase

S-receptor-like serine/threonine-protein kinase
S-receptor-like serine/threonine-protein kinase
S-receptor-like serine/threonine-protein kinase
S-receptor-like serine/threonine-protein kinase
S-receptor-like serine/threonine-protein kinase
S-receptor-like serine/threonine-protein kinase
S-locus glycoprotein
S-receptor-like serine/threonine-protein kinase
S-locus glycoprotein
GPCR kinase
S-receptor-like serine/threonine-protein kinase
Phosphorylase kinase, gamma catalytic subunit
S-locus glycoprotein
S-locus glycoprotein
S-receptor-like serine/threonine-protein kinase
S-receptor-like serine/threonine-protein kinase
S-receptor-like serine/threonine-protein kinase
Phosphorylase kinase, gamma catalytic subunit
Serine/threonine protein kinase
S-locus glycoprotein
S-receptor-like serine/threonine-protein kinase
Serine/threonine protein kinase
S-locus glycoprotein
S-receptor-like serine/threonine-protein kinase
GPCR kinase
S-receptor-like serine/threonine-protein kinase
PAN/Apple domain containing protein
Mitogen-activated protein kinase kinase kinase
Tyrosine-protein kinase
S-locus glycoprotein (Fragment)
Serine/threonine protein kinase
S-receptor-like serine/threonine-protein kinase (Fragment)
Serine/threonine protein kinase (Fragment)
Tyrosine-protein kinase
Serine/threonine protein kinase
S-locus glycoprotein (Fragment)
S-receptor-like serine/threonine-protein kinase
S-locus glycoprotein
S-receptor-like serine/threonine-protein kinase
Mitogen-activated protein kinase kinase kinase
S-locus glycoprotein

S-locus glycoprotein domain containing protein
S-receptor-like serine/threonine-protein kinase
S-receptor-like serine/threonine-protein kinase
PAN/Apple domain containing protein
Lectin domain containing protein (Fragment)
PAN/Apple domain containing protein
S-receptor-like serine/threonine-protein kinase
S-receptor-like serine/threonine-protein kinase
S-locus glycoprotein
S-receptor-like serine/threonine-protein kinase
Uncharacterized protein (Fragment)
S-receptor-like serine/threonine-protein kinase
S-receptor-like serine/threonine-protein kinase
Phosphorylase kinase, gamma catalytic subunit
S-receptor-like serine/threonine-protein kinase
S-receptor-like serine/threonine-protein kinase
S-receptor-like serine/threonine-protein kinase
Serine/threonine protein kinase
S-receptor-like serine/threonine-protein kinase
S-locus receptor kinase
S-locus glycoprotein
S-receptor-like serine/threonine-protein kinase
Serine/threonine protein kinase
S-receptor-like serine/threonine-protein kinase
Lectin domain containing protein
Mitogen-activated protein kinase kinase kinase
S-locus glycoprotein
S-receptor-like serine/threonine-protein kinase
S-locus glycoprotein
S-receptor-like serine/threonine-protein kinase
Uncharacterized protein (Fragment)
S-locus glycoprotein
Serine/threonine protein kinase
PAN/Apple domain containing protein (Fragment)
Serine/threonine protein kinase
S-receptor-like serine/threonine-protein kinase
S-locus glycoprotein
S-locus glycoprotein
S-locus glycoprotein
Serine/threonine protein kinase
S-receptor-like serine/threonine-protein kinase

S-receptor-like serine/threonine-protein kinase
S-locus glycoprotein
S-receptor-like serine/threonine-protein kinase
S-receptor-like serine/threonine-protein kinase
S-receptor-like serine/threonine-protein kinase
Tyrosine-protein kinase
Mitogen-activated protein kinase kinase kinase
S-receptor-like serine/threonine-protein kinase
S-receptor-like serine/threonine-protein kinase
S-receptor-like serine/threonine-protein kinase
S-locus glycoprotein
S-receptor-like serine/threonine-protein kinase
S-locus glycoprotein (Fragment)
S-receptor-like serine/threonine-protein kinase
S-receptor-like serine/threonine-protein kinase
S-receptor-like serine/threonine-protein kinase
Lectin domain containing protein
S-locus glycoprotein domain containing protein
S-receptor-like serine/threonine-protein kinase
S-receptor-like serine/threonine-protein kinase
Uncharacterized protein (Fragment)
Uncharacterized protein (Fragment)
Putative metacaspase-1 (Fragment)
Uncharacterized protein
Uncharacterized protein
Serine protease P153
Uncharacterized protein
Uncharacterized protein
Uncharacterized protein
Uncharacterized protein
Uncharacterized protein
plasminogen isoform X4
coagulation factor XI
Plasminogen
hepatocyte growth factor isoform X1
plasma kallikrein
hepatocyte growth factor isoform X3
hepatocyte growth factor-like protein
Plasminogen
Plasminogen
hepatocyte growth factor isoform X2

Uncharacterized protein
Uncharacterized protein (Fragment)
Uncharacterized protein
Uncharacterized protein
Putative tyrosinase-like protein tyr-3 (Fragment)
Protein dyf-8
Uncharacterized protein
tRNA (guanine-N(7)-)-methyltransferase non-catalytic subunit
Uncharacterized protein
Uncharacterized protein
Uncharacterized protein
Uncharacterized protein
Endochitinase B (Fragment)
Uncharacterized protein
Uncharacterized protein (Fragment)
Uncharacterized protein (Fragment)
Uncharacterized protein
Uncharacterized protein
Uncharacterized protein
Uncharacterized protein
Uncharacterized protein
Uncharacterized protein
Uncharacterized protein (Fragment)
Uncharacterized protein
Uncharacterized protein (Fragment)
Neurotrypsin
Uncharacterized protein
Protein dyf-8
Uncharacterized protein
Uncharacterized protein
Uncharacterized protein
Neurotrypsin
Uncharacterized protein
Uncharacterized protein (Fragment)
Uncharacterized protein (Fragment)
tRNA (guanine-N(7)-)-methyltransferase non-catalytic subunit
Neurotrypsin
Uncharacterized protein
Uncharacterized protein (Fragment)
Uncharacterized protein (Fragment)
Uncharacterized protein (Fragment)

Uncharacterized protein
Uncharacterized protein
Uncharacterized protein
Uncharacterized protein
Lysine-specific demethylase 6A (Fragment)
Uncharacterized protein
Uncharacterized protein
Uncharacterized protein
Uncharacterized protein (Fragment)
Uncharacterized protein
tRNA (Guanine-N(7)-)-methyltransferase non-catalytic subunit wuho
Neurotrypsin
Uncharacterized protein (Fragment)
Uncharacterized protein
Neurotrypsin
Uncharacterized protein (Fragment)
Uncharacterized protein
Uncharacterized protein
Uncharacterized protein (Fragment)
Uncharacterized protein
Protein dyf-8
Uncharacterized protein
Uncharacterized protein
tRNA (guanine-N(7)-)-methyltransferase non-catalytic subunit
Uncharacterized protein (Fragment)
Uncharacterized protein
Uncharacterized protein (Fragment)
Putative tyrosinase-like protein tyr-3
Neurotrypsin
Hemicentin-1
Uncharacterized protein
Uncharacterized protein
Uncharacterized protein
Uncharacterized protein
Uncharacterized protein (Fragment)
Putative tyrosinase-like protein tyr-3
Lysine-specific demethylase 6A (Fragment)
Uncharacterized protein
Uncharacterized protein
Protein dyf-8
Uncharacterized protein

Uncharacterized protein
Uncharacterized protein
Uncharacterized protein (Fragment)
Neurotrypsin
Neurotrypsin
Protein dyf-8
Uncharacterized protein
Uncharacterized protein
Uncharacterized protein (Fragment)
Uncharacterized protein
Uncharacterized protein (Fragment)
Uncharacterized protein
Uncharacterized protein
Uncharacterized protein (Fragment)
Uncharacterized protein (Fragment)
Uncharacterized protein
Uncharacterized protein
Uncharacterized protein
Uncharacterized protein (Fragment)
Uncharacterized protein
Uncharacterized protein
Uncharacterized protein
Uncharacterized protein
Uncharacterized protein
Uncharacterized protein
Uncharacterized protein (Fragment)
Uncharacterized protein
Uncharacterized protein
Uncharacterized protein
Uncharacterized protein (Fragment)
Uncharacterized protein (Fragment)
Uncharacterized protein
Uncharacterized protein (Fragment)
Uncharacterized protein
Uncharacterized protein
Uncharacterized protein
Uncharacterized protein
Uncharacterized protein
Uncharacterized protein
Uncharacterized protein
Uncharacterized protein

Uncharacterized protein
Uncharacterized protein
Short-chain dehydrogenase TIC 32, chloroplastic
Uncharacterized protein
Uncharacterized protein (Fragment)
Uncharacterized protein
Endochitinase B
Uncharacterized protein
Uncharacterized protein
Uncharacterized protein
Uncharacterized protein
Uncharacterized protein
Uncharacterized protein
Uncharacterized protein
Uncharacterized protein
Uncharacterized protein
Uncharacterized protein
Uncharacterized protein
Uncharacterized protein
PAN domain protein
Uncharacterized protein (Fragment)
Uncharacterized protein
Uncharacterized protein (Fragment)
PAN domain protein
PAN domain protein
PAN domain protein
Uncharacterized protein (Fragment)
PAN domain protein
PAN domain protein (Fragment)
PAN domain protein (Fragment)
Uncharacterized protein
Uncharacterized protein
Uncharacterized protein
Protein dyf-8
PAN domain protein
Uncharacterized protein
PAN domain protein (Fragment)
Neurotrypsin
PAN domain protein
Uncharacterized protein
Neurotrypsin

Chitinase class I
Uncharacterized protein
Uncharacterized protein
Uncharacterized protein
PAN domain protein
Uncharacterized protein (Fragment)
Uncharacterized protein
PAN domain protein
Uncharacterized protein (Fragment)
Uncharacterized protein
Uncharacterized protein (Fragment)
tRNA (Guanine-N(7)-)-methyltransferase non-catalytic subunit wuho
Uncharacterized protein
Uncharacterized protein
Neurotrypsin
Uncharacterized protein (Fragment)
PAN domain protein
Putative PAN domain protein (Fragment)
Uncharacterized protein
Protein dyf-8
Uncharacterized protein (Fragment)
PAN domain protein
Uncharacterized protein
Putative tyrosinase-like protein tyr-3
Uncharacterized protein
Uncharacterized protein
Uncharacterized protein
Trypsin
PAN domain protein
Uncharacterized protein (Fragment)
Uncharacterized protein
PAN domain protein
Uncharacterized protein (Fragment)
PAN domain protein
Uncharacterized protein
PAN domain protein (Fragment)
Uncharacterized protein
Hemicentin-1
PAN domain protein
Uncharacterized protein (Fragment)
PAN domain protein (Fragment)

PAN domain protein
PAN domain protein
Uncharacterized protein
PAN domain protein
Uncharacterized protein (Fragment)
Uncharacterized protein
Uncharacterized protein
PAN domain protein
Endochitinase B (Fragment)
Endochitinase B (Fragment)
Uncharacterized protein
Uncharacterized protein
Lysine-specific demethylase 6A
PAN domain protein
Uncharacterized protein
Uncharacterized protein
PAN domain protein (Fragment)
Tetratricopeptide repeat protein
Uncharacterized protein (Fragment)
PAN domain protein
Uncharacterized protein
Uncharacterized protein
PAN domain protein
PAN domain protein
Uncharacterized protein
Uncharacterized protein
Uncharacterized protein (Fragment)
Uncharacterized protein (Fragment)
Uncharacterized protein
Uncharacterized protein
Uncharacterized protein
Uncharacterized protein
Endochitinase B (Fragment)
Uncharacterized protein (Fragment)
Uncharacterized protein
Uncharacterized protein (Fragment)
tRNA (guanine-N(7)-)-methyltransferase non-catalytic subunit
Uncharacterized protein
Uncharacterized protein
Neurotrypsin
Neurotrypsin

Uncharacterized protein
Putative tyrosinase-like protein tyr-3
Uncharacterized protein
Uncharacterized protein
Uncharacterized protein (Fragment)
Uncharacterized protein
Uncharacterized protein
Uncharacterized protein
Uncharacterized protein
Uncharacterized protein (Fragment)
Uncharacterized protein
Uncharacterized protein
Uncharacterized protein (Fragment)
Lysine-specific demethylase 6A
Uncharacterized protein
Uncharacterized protein
Uncharacterized protein
Uncharacterized protein
Lysine-specific demethylase 6A
Uncharacterized protein (Fragment)
Uncharacterized protein
Uncharacterized protein
Uncharacterized protein
Uncharacterized protein
Uncharacterized protein
Lysine-specific demethylase 6A
Uncharacterized protein
Uncharacterized protein (Fragment)
Uncharacterized protein
Uncharacterized protein
Uncharacterized protein
Uncharacterized protein
Uncharacterized protein
Uncharacterized protein
Uncharacterized protein
Uncharacterized protein
Uncharacterized protein
Uncharacterized protein
Protein dyf-8
Uncharacterized protein
Neurotrypsin

Uncharacterized protein (Fragment)
Uncharacterized protein
Microsomal triglyceride transfer protein large subunit
Protein dyf-8
Uncharacterized protein
Uncharacterized protein (Fragment)
Protein dyf-8
Uncharacterized protein
Uncharacterized protein
Uncharacterized protein
Uncharacterized protein
Uncharacterized protein (Fragment)
Uncharacterized protein
Uncharacterized protein (Fragment)
Uncharacterized protein
Hemicentin-1
Uncharacterized protein (Fragment)
Uncharacterized protein
Uncharacterized protein
Uncharacterized protein (Fragment)
Uncharacterized protein
Uncharacterized protein
Uncharacterized protein (Fragment)
Uncharacterized protein
Lysine-specific demethylase 6A
Uncharacterized protein (Fragment)
Uncharacterized protein (Fragment)
Uncharacterized protein
Uncharacterized protein
High-affinity choline transporter 1
Uncharacterized protein (Fragment)
Uncharacterized protein
Uncharacterized protein
Uncharacterized protein
Uncharacterized protein
Uncharacterized protein
Uncharacterized protein (Fragment)
Uncharacterized protein
Uncharacterized protein
Uncharacterized protein
Uncharacterized protein

Uncharacterized protein
Uncharacterized protein (Fragment)
Uncharacterized protein
Uncharacterized protein (Fragment)
Uncharacterized protein
Neurotrypsin
Uncharacterized protein (Fragment)
Uncharacterized protein
Uncharacterized protein
Putative tyrosinase-like protein tyr-3 (Fragment)
Uncharacterized protein
Uncharacterized protein (Fragment)
Protein dyf-8
Uncharacterized protein
Uncharacterized protein
Uncharacterized protein
Uncharacterized protein
Uncharacterized protein
Uncharacterized protein
Uncharacterized protein
Uncharacterized protein
Uncharacterized protein
Protein dyf-8
Uncharacterized protein
Uncharacterized protein
Uncharacterized protein (Fragment)
Uncharacterized protein (Fragment)
Uncharacterized protein (Fragment)
Uncharacterized protein
Neurotrypsin
Uncharacterized protein (Fragment)
Uncharacterized protein
Uncharacterized protein (Fragment)
Uncharacterized protein
Uncharacterized protein
Uncharacterized protein (Fragment)
Uncharacterized protein
Uncharacterized protein
Neurotrypsin
Uncharacterized protein
Uncharacterized protein

Neurotrypsin
Uncharacterized protein
tRNA (guanine-N(7)-)-methyltransferase non-catalytic subunit
Putative tyrosinase-like protein tyr-3 (Fragment)
Neurotrypsin
Uncharacterized protein
Protein dyf-8
Uncharacterized protein
Uncharacterized protein
Uncharacterized protein
Uncharacterized protein
Uncharacterized protein (Fragment)
Neurotrypsin (Fragment)
Uncharacterized protein
Uncharacterized protein
Uncharacterized protein
Uncharacterized protein
Uncharacterized protein
Uncharacterized protein
Uncharacterized protein
Uncharacterized protein
Protein dyf-8
Uncharacterized protein
Uncharacterized protein
Uncharacterized protein (Fragment)
Uncharacterized protein
Uncharacterized protein
Uncharacterized protein
Uncharacterized protein (Fragment)
Uncharacterized protein
Lysine-specific demethylase 6A
Uncharacterized protein
Uncharacterized protein
Uncharacterized protein
Uncharacterized protein
Uncharacterized protein
Uncharacterized protein
Uncharacterized protein (Fragment)
Hemicentin-1
Endochitinase B (Fragment)
Lysine-specific demethylase 6A

Uncharacterized protein (Fragment)
Uncharacterized protein
Uncharacterized protein (Fragment)
Uncharacterized protein (Fragment)
Uncharacterized protein
Uncharacterized protein (Fragment)
Lysine-specific demethylase 6A
Uncharacterized protein
Uncharacterized protein (Fragment)
Lysine-specific demethylase 6A
Uncharacterized protein
Microsomal triglyceride transfer protein large subunit
Uncharacterized protein (Fragment)
Uncharacterized protein
Uncharacterized protein
Neurotrypsin
Lysine-specific demethylase 6A
Microsomal triglyceride transfer protein large subunit
Uncharacterized protein
Uncharacterized protein (Fragment)
Uncharacterized protein
Putative tyrosinase-like protein tyr-3 (Fragment)
Uncharacterized protein
Uncharacterized protein
Uncharacterized protein
Uncharacterized protein (Fragment)
Uncharacterized protein
Uncharacterized protein
Uncharacterized protein
Uncharacterized protein
Neurotrypsin
Uncharacterized protein
Uncharacterized protein
Protein dyf-8
Uncharacterized protein
Uncharacterized protein (Fragment)
Protein dyf-8
Putative tyrosinase-like protein tyr-3
Uncharacterized protein (Fragment)
Uncharacterized protein
Uncharacterized protein (Fragment)

Uncharacterized protein
Neurotrypsin
Uncharacterized protein (Fragment)
Uncharacterized protein
Uncharacterized protein (Fragment)
Uncharacterized protein
Uncharacterized protein
Uncharacterized protein (Fragment)
Uncharacterized protein (Fragment)
Uncharacterized protein
Uncharacterized protein (Fragment)
Uncharacterized protein
Uncharacterized protein
Uncharacterized protein
Uncharacterized protein
Uncharacterized protein
Uncharacterized protein
Uncharacterized protein
Uncharacterized protein
Uncharacterized protein
Uncharacterized protein
Uncharacterized protein (Fragment)
Uncharacterized protein (Fragment)
Uncharacterized protein (Fragment)
Neurotrypsin (Fragment)
Neurotrypsin
Uncharacterized protein
Uncharacterized protein
Uncharacterized protein (Fragment)
Uncharacterized protein
Lysine-specific demethylase 6A
Uncharacterized protein
Uncharacterized protein (Fragment)
Uncharacterized protein
Uncharacterized protein
Lysine-specific demethylase 6A (Fragment)
Uncharacterized protein
Neurotrypsin
Uncharacterized protein
Protein dyf-8
Uncharacterized protein

Uncharacterized protein (Fragment)
Uncharacterized protein (Fragment)
Uncharacterized protein (Fragment)
Uncharacterized protein
Uncharacterized protein
Uncharacterized protein
Uncharacterized protein
Uncharacterized protein
Uncharacterized protein
Uncharacterized protein
Endochitinase B
Uncharacterized protein
Uncharacterized protein
Uncharacterized protein
Uncharacterized protein
Protein dyf-8
Uncharacterized protein
Uncharacterized protein
Neurotrypsin
Uncharacterized protein
Uncharacterized protein
Uncharacterized protein (Fragment)
Uncharacterized protein
Endochitinase B
Uncharacterized protein
Uncharacterized protein (Fragment)
GILT-like protein F37H8.5
Uncharacterized protein
Short-chain dehydrogenase TIC 32, chloroplastic
Uncharacterized protein (Fragment)
Uncharacterized protein
Uncharacterized protein (Fragment)
Uncharacterized protein (Fragment)
Uncharacterized protein
Uncharacterized protein
Neurotrypsin
Uncharacterized protein
Uncharacterized protein
Lysine-specific demethylase 6A (Fragment)
Lysine-specific demethylase 6A
Uncharacterized protein

Uncharacterized protein
Lysine-specific demethylase 6A
Uncharacterized protein
Uncharacterized protein (Fragment)
Uncharacterized protein (Fragment)
Uncharacterized protein
Protein dyf-8
Uncharacterized protein
Uncharacterized protein
Uncharacterized protein
tRNA (guanine-N(7)-)-methyltransferase non-catalytic subunit
Lysine-specific demethylase 6A (Fragment)
Endochitinase B
Uncharacterized protein
Microsomal triglyceride transfer protein large subunit
Uncharacterized protein
Uncharacterized protein
Uncharacterized protein (Fragment)
Uncharacterized protein (Fragment)
Uncharacterized protein
Uncharacterized protein (Fragment)
Uncharacterized protein
Protein dyf-8
Neurotrypsin
Uncharacterized protein
Uncharacterized protein
Uncharacterized protein (Fragment)
Uncharacterized protein
Uncharacterized protein
Uncharacterized protein
Uncharacterized protein
Uncharacterized protein
Uncharacterized protein (Fragment)
Neurotrypsin
Uncharacterized protein
Uncharacterized protein
Uncharacterized protein (Fragment)
Uncharacterized protein (Fragment)
Microsomal triglyceride transfer protein large subunit
Uncharacterized protein
Uncharacterized protein

Uncharacterized protein (Fragment)
Uncharacterized protein (Fragment)
Uncharacterized protein (Fragment)
Uncharacterized protein (Fragment)
Uncharacterized protein (Fragment)
Uncharacterized protein
Uncharacterized protein
Uncharacterized protein (Fragment)
Uncharacterized protein
Lysine-specific demethylase 6A (Fragment)
Uncharacterized protein
Uncharacterized protein
Uncharacterized protein
Uncharacterized protein
Uncharacterized protein
Uncharacterized protein
Uncharacterized protein
Uncharacterized protein (Fragment)
Uncharacterized protein (Fragment)
Uncharacterized protein
Uncharacterized protein (Fragment)
Uncharacterized protein
Uncharacterized protein (Fragment)
Uncharacterized protein
Uncharacterized protein
Uncharacterized protein
Uncharacterized protein
Uncharacterized protein (Fragment)
Uncharacterized protein (Fragment)
Uncharacterized protein
Uncharacterized protein
Uncharacterized protein (Fragment)
Uncharacterized protein
Uncharacterized protein
Putative tyrosinase-like protein tyr-3
Lysine-specific demethylase 6A
Uncharacterized protein
GILT-like protein F37H8.5
Uncharacterized protein
Uncharacterized protein
Putative tyrosinase-like protein tyr-3

Uncharacterized protein
Uncharacterized protein (Fragment)
Uncharacterized protein (Fragment)
Putative tyrosinase-like protein tyr-3
Uncharacterized protein (Fragment)
Uncharacterized protein
Uncharacterized protein
Uncharacterized protein (Fragment)
Uncharacterized protein
Uncharacterized protein
Uncharacterized protein (Fragment)
Uncharacterized protein (Fragment)
Neurotrypsin
Brain-specific angiogenesis inhibitor 1
Uncharacterized protein (Fragment)
Uncharacterized protein
Uncharacterized protein (Fragment)
Uncharacterized protein
Uncharacterized protein
Uncharacterized protein
Uncharacterized protein
Uncharacterized protein (Fragment)
Endochitinase B
Uncharacterized protein
Uncharacterized protein
Neurotrypsin
Putative tyrosinase-like protein tyr-3
Uncharacterized protein
Uncharacterized protein
Uncharacterized protein
Uncharacterized protein (Fragment)
Uncharacterized protein
Neurotrypsin
Uncharacterized protein
Myb-binding protein 1A-like protein
Myb-binding protein 1A-like protein
Uncharacterized protein
Uncharacterized protein
tRNA (guanine-N(7)-)-methyltransferase non-catalytic subunit
Uncharacterized protein (Fragment)
Uncharacterized protein

Uncharacterized protein
Uncharacterized protein
Uncharacterized protein
Uncharacterized protein
Uncharacterized protein
Uncharacterized protein
Uncharacterized protein (Fragment)
Myb-binding protein 1A-like protein
Microsomal triglyceride transfer protein large subunit
Uncharacterized protein (Fragment)
Neurotrypsin
Uncharacterized protein (Fragment)
Protein dyf-8
Uncharacterized protein
Uncharacterized protein
Uncharacterized protein
Uncharacterized protein
Uncharacterized protein
Endochitinase B (Fragment)
Lysine-specific demethylase 6A (Fragment)
Uncharacterized protein
Uncharacterized protein
Uncharacterized protein
Uncharacterized protein
Uncharacterized protein
Uncharacterized protein
Uncharacterized protein
Uncharacterized protein
Uncharacterized protein
Uncharacterized protein
Uncharacterized protein
Uncharacterized protein
Uncharacterized protein
Lysine-specific demethylase 6A (Fragment)
Uncharacterized protein
Uncharacterized protein (Fragment)
Putative tyrosinase-like protein tyr-3 (Fragment)
Uncharacterized protein
Uncharacterized protein
Microsomal triglyceride transfer protein large subunit
Putative tyrosinase-like protein tyr-3 (Fragment)

Uncharacterized protein (Fragment)
Microsomal triglyceride transfer protein large subunit
Uncharacterized protein
Uncharacterized protein (Fragment)
Uncharacterized protein
Uncharacterized protein
Protein dyf-8
Uncharacterized protein
Uncharacterized protein
Uncharacterized protein (Fragment)
Uncharacterized protein
Uncharacterized protein
Uncharacterized protein (Fragment)
Uncharacterized protein (Fragment)
Uncharacterized protein (Fragment)
Uncharacterized protein
Uncharacterized protein
Uncharacterized protein
Uncharacterized protein (Fragment)
Uncharacterized protein
Lysine-specific demethylase 6A (Fragment)
Uncharacterized protein
Lysine-specific demethylase 6A (Fragment)
Lysine-specific demethylase 6A (Fragment)
Uncharacterized protein
Uncharacterized protein
Uncharacterized protein
Uncharacterized protein
Uncharacterized protein
Uncharacterized protein (Fragment)
Uncharacterized protein (Fragment)
Uncharacterized protein
Uncharacterized protein (Fragment)
Uncharacterized protein
Uncharacterized protein
Uncharacterized protein
Short-chain dehydrogenase TIC 32, chloroplastic (Fragment)
Uncharacterized protein
Uncharacterized protein
Uncharacterized protein (Fragment)
Uncharacterized protein (Fragment)

Uncharacterized protein
Uncharacterized protein (Fragment)
Uncharacterized protein (Fragment)
Uncharacterized protein (Fragment)
Protein dyf-8
Neurotrypsin (Fragment)
Uncharacterized protein
Uncharacterized protein
Microsomal triglyceride transfer protein large subunit
Uncharacterized protein (Fragment)
Uncharacterized protein (Fragment)
Uncharacterized protein
Uncharacterized protein
Uncharacterized protein
Uncharacterized protein
Protein dyf-8
Uncharacterized protein
Uncharacterized protein
Uncharacterized protein
Uncharacterized protein
Uncharacterized protein
Uncharacterized protein
Uncharacterized protein
Uncharacterized protein
Neurotrypsin (Fragment)
Uncharacterized protein (Fragment)
Uncharacterized protein (Fragment)
Uncharacterized protein
Uncharacterized protein
Uncharacterized protein (Fragment)
Protein dyf-8
Endochitinase B (Fragment)
Uncharacterized protein (Fragment)
Uncharacterized protein
Hemicentin-1
Uncharacterized protein
Neurotrypsin
Uncharacterized protein (Fragment)
Uncharacterized protein (Fragment)
Uncharacterized protein
Lysine-specific demethylase 6A

Uncharacterized protein
Uncharacterized protein (Fragment)
Uncharacterized protein
Uncharacterized protein
Uncharacterized protein
Uncharacterized protein
Uncharacterized protein
Uncharacterized protein (Fragment)
Uncharacterized protein
Uncharacterized protein (Fragment)
Uncharacterized protein (Fragment)
Uncharacterized protein
Uncharacterized protein
Neurotrypsin
Uncharacterized protein
Uncharacterized protein (Fragment)
Uncharacterized protein
Protein dyf-8
Uncharacterized protein
Uncharacterized protein
Uncharacterized protein
Uncharacterized protein
Uncharacterized protein
Uncharacterized protein (Fragment)
Uncharacterized protein
Uncharacterized protein (Fragment)
Uncharacterized protein
Uncharacterized protein
Uncharacterized protein
Protein dyf-8
Putative tyrosinase-like protein tyr-3 (Fragment)
Uncharacterized protein
Uncharacterized protein
Uncharacterized protein
Uncharacterized protein (Fragment)
Uncharacterized protein
Uncharacterized protein
Uncharacterized protein (Fragment)
Uncharacterized protein
Uncharacterized protein
Uncharacterized protein (Fragment)

Uncharacterized protein
Uncharacterized protein (Fragment)
Uncharacterized protein
tRNA (guanine-N(7)-)-methyltransferase non-catalytic subunit
Uncharacterized protein (Fragment)
Uncharacterized protein (Fragment)
Protein dyf-8
Uncharacterized protein
Uncharacterized protein
Uncharacterized protein
Hemicentin-1
Protein dyf-8
Uncharacterized protein (Fragment)
Uncharacterized protein
Lysine-specific demethylase 6A (Fragment)
Uncharacterized protein (Fragment)
Uncharacterized protein
Uncharacterized protein (Fragment)
Uncharacterized protein
Uncharacterized protein
Uncharacterized protein
Neurotrypsin
Uncharacterized protein
Uncharacterized protein (Fragment)
Neurotrypsin
Uncharacterized protein
Uncharacterized protein
Uncharacterized protein (Fragment)
Uncharacterized protein
Uncharacterized protein
Uncharacterized protein
Uncharacterized protein
Putative tyrosinase-like protein tyr-3
Uncharacterized protein
Protein dyf-8
Uncharacterized protein
Uncharacterized protein
Uncharacterized protein (Fragment)
Uncharacterized protein
Uncharacterized protein
Uncharacterized protein

Uncharacterized protein
Uncharacterized protein (Fragment)
Putative tyrosinase-like protein tyr-3 (Fragment)
Neurotrypsin
Uncharacterized protein (Fragment)
Uncharacterized protein
Uncharacterized protein
Uncharacterized protein (Fragment)
Uncharacterized protein
Uncharacterized protein
Uncharacterized protein
Uncharacterized protein
Neurotrypsin
Uncharacterized protein
Uncharacterized protein (Fragment)
Uncharacterized protein
Lysine-specific demethylase 6A (Fragment)
Uncharacterized protein
Uncharacterized protein (Fragment)
Uncharacterized protein
Protein dyf-8
Uncharacterized protein
Uncharacterized protein (Fragment)
Neurotrypsin
Uncharacterized protein
Uncharacterized protein
Uncharacterized protein (Fragment)
Uncharacterized protein (Fragment)
Uncharacterized protein (Fragment)
Uncharacterized protein
Uncharacterized protein
Uncharacterized protein (Fragment)
Lysine-specific demethylase 6A (Fragment)
Uncharacterized protein
Uncharacterized protein (Fragment)
Uncharacterized protein
Uncharacterized protein
Uncharacterized protein (Fragment)
Uncharacterized protein
tRNA (Guanine-N(7)-)-methyltransferase non-catalytic subunit wuho
Uncharacterized protein

Uncharacterized protein
Lysine-specific demethylase 6A (Fragment)
Uncharacterized protein
Uncharacterized protein
Semaphorin-5A
Lysine-specific demethylase 6A (Fragment)
Uncharacterized protein
Lysine-specific demethylase 6A (Fragment)
Uncharacterized protein
Endochitinase B
Uncharacterized protein
Uncharacterized protein
Uncharacterized protein
Neurotrypsin
Uncharacterized protein
Uncharacterized protein
Uncharacterized protein
Uncharacterized protein (Fragment)
Lysine-specific demethylase 6A (Fragment)
Uncharacterized protein
Uncharacterized protein
Uncharacterized protein (Fragment)
Uncharacterized protein
Uncharacterized protein
Tigger transposable element-derived protein 4
Uncharacterized protein
Neurotrypsin
Uncharacterized protein (Fragment)
Uncharacterized protein
Neurotrypsin
Uncharacterized protein (Fragment)
Protein dyf-8
Uncharacterized protein (Fragment)
Uncharacterized protein
Uncharacterized protein
Uncharacterized protein (Fragment)
Uncharacterized protein (Fragment)
N/apple PAN-like protein (Fragment)
Uncharacterized protein
Uncharacterized protein
Uncharacterized protein

Uncharacterized protein (Fragment)
Uncharacterized protein
Uncharacterized protein
Uncharacterized protein
Uncharacterized protein (Fragment)
Uncharacterized protein
tRNA (guanine-N(7)-)-methyltransferase non-catalytic subunit
Uncharacterized protein
Lysine-specific demethylase 6A (Fragment)
Uncharacterized protein
Uncharacterized protein
Uncharacterized protein
Uncharacterized protein
Uncharacterized protein
Uncharacterized protein
Putative tyrosinase-like protein tyr-3
Uncharacterized protein
Uncharacterized protein
Uncharacterized protein
Uncharacterized protein
Uncharacterized protein
Lysine-specific demethylase 6A (Fragment)
Uncharacterized protein (Fragment)
Protein dyf-8
Neurotrypsin
Tigger transposable element-derived protein 4
Uncharacterized protein
Uncharacterized protein
Uncharacterized protein (Fragment)
Uncharacterized protein
Uncharacterized protein (Fragment)
Uncharacterized protein
Uncharacterized protein
Semaphorin-5A
Uncharacterized protein
Uncharacterized protein
Uncharacterized protein
Uncharacterized protein
Uncharacterized protein
Uncharacterized protein
Uncharacterized protein (Fragment)

Uncharacterized protein
Uncharacterized protein
Uncharacterized protein
Uncharacterized protein
tRNA (Guanine-N(7)-)-methyltransferase non-catalytic subunit wuho (Fragment)
Uncharacterized protein (Fragment)
Uncharacterized protein
Uncharacterized protein
Uncharacterized protein
Uncharacterized protein
Uncharacterized protein
Uncharacterized protein
Uncharacterized protein
Uncharacterized protein (Fragment)
Putative tyrosinase-like protein tyr-3 (Fragment)
Uncharacterized protein
Uncharacterized protein
Acetyl-CoA carboxylase 1 (Fragment)
Uncharacterized protein
Uncharacterized protein
Uncharacterized protein
Uncharacterized protein
Uncharacterized protein (Fragment)
Uncharacterized protein
tRNA (Guanine-N(7)-)-methyltransferase non-catalytic subunit wuho (Fragment)
Uncharacterized protein
Uncharacterized protein
Uncharacterized protein (Fragment)
Uncharacterized protein
Microsomal triglyceride transfer protein large subunit
Uncharacterized protein
Uncharacterized protein
Uncharacterized protein (Fragment)
Uncharacterized protein
Uncharacterized protein
Uncharacterized protein (Fragment)
Uncharacterized protein
Endochitinase B (Fragment)
Uncharacterized protein (Fragment)
Uncharacterized protein
Uncharacterized protein

Uncharacterized protein
Uncharacterized protein
Uncharacterized protein
Uncharacterized protein
Protein dyf-8
Uncharacterized protein
Uncharacterized protein
Uncharacterized protein
Uncharacterized protein
Uncharacterized protein
Glycosyltransferase family 31
Glycosyltransferase family 31 protein
Uncharacterized protein
Uncharacterized protein
Uncharacterized protein
Uncharacterized protein
Uncharacterized protein
Uncharacterized protein
Uncharacterized protein
Glycosyltransferase family 31 protein
Uncharacterized protein
GT31 glycosyltransferase
Uncharacterized protein
Uncharacterized protein (Fragment)
Uncharacterized protein
Glycosyltransferase family 31 protein
Uncharacterized protein
Uncharacterized protein
Uncharacterized protein
Uncharacterized protein
Uncharacterized protein (Fragment)
Uncharacterized protein
Uncharacterized protein
Uncharacterized protein
Uncharacterized protein
Uncharacterized protein
Glycosyltransferase family 31 protein
Uncharacterized protein
Uncharacterized protein
Uncharacterized protein (Fragment)
Uncharacterized protein

Uncharacterized protein
Uncharacterized protein
Uncharacterized protein
Uncharacterized protein
Uncharacterized protein
Uncharacterized protein
Uncharacterized protein
Uncharacterized protein
Macrophage mannose receptor 1
Uncharacterized protein
Uncharacterized protein
Uncharacterized protein
Uncharacterized protein
Uncharacterized protein
Uncharacterized protein
Uncharacterized protein
Uncharacterized protein
Uncharacterized protein
Uncharacterized protein
Uncharacterized protein
Uncharacterized protein
Uncharacterized protein
Uncharacterized protein
Uncharacterized protein
Uncharacterized protein
Uncharacterized protein
Uncharacterized protein
Uncharacterized protein
Uncharacterized protein
Uncharacterized protein
Uncharacterized protein (Fragment)
Uncharacterized protein
Lectin C-type domain protein
Uncharacterized protein (Fragment)
Uncharacterized protein
Uncharacterized protein
PAN domain protein
Uncharacterized protein
Uncharacterized protein (Fragment)
Uncharacterized protein
Uncharacterized protein

PAN domain protein
PAN domain protein
Uncharacterized protein (Fragment)
Uncharacterized protein (Fragment)
Uncharacterized protein
Uncharacterized protein (Fragment)
Uncharacterized protein
PAN domain protein
Uncharacterized protein
Uncharacterized protein (Fragment)
Uncharacterized protein (Fragment)
Uncharacterized protein
Uncharacterized protein
PAN domain protein
Uncharacterized protein
Uncharacterized protein
Uncharacterized protein
Uncharacterized protein
Uncharacterized protein
Uncharacterized protein
PAN domain protein
Uncharacterized protein
Trypsin
PAN 1 domain containing protein
Coagulation factor IX
PAN 1 domain containing protein
PAN 1 and Zona pellucida domain containing protei n
PAN 1 domain containing protein
Plasminogen protein B
PAN 1 domain containing protein
PAN 1 domain containing protein
Uncharacterized protein
PAN 1 domain containing protein
PAN 1 domain containing protein
PAN 1 domain containing protein
PAN 1 domain containing protein
Trypsin and PAN 1 domain containing protein
PAN 1 domain containing protein
Tyrosinase domain containing protein
PAN 1 domain containing protein
PAN 1 domain containing protein

G-type lectin S-receptor-like serine/threonine-protein kinase (Fragment)
G-type lectin S-receptor-like serine/threonine-protein kinase (Fragment)
G-type lectin S-receptor-like serine/threonine-protein kinase (Fragment)
G-type lectin S-receptor-like serine/threonine-protein kinase (Fragment)
G-type lectin S-receptor-like serine/threonine-protein kinase (Fragment)
G-type lectin S-receptor-like serine/threonine-protein kinase (Fragment)
G-type lectin S-receptor-like serine/threonine-protein kinase (Fragment)
G-type lectin S-receptor-like serine/threonine-protein kinase (Fragment)
G-type lectin S-receptor-like serine/threonine-protein kinase (Fragment)
G-type lectin S-receptor-like serine/threonine-protein kinase (Fragment)
G-type lectin S-receptor-like serine/threonine-protein kinase (Fragment)
G-type lectin S-receptor-like serine/threonine-protein kinase (Fragment)
Serine/threonine protein kinase (Fragment)
G-type lectin S-receptor-like serine/threonine-protein kinase (Fragment)
G-type lectin S-receptor-like serine/threonine-protein kinase (Fragment)
S-locus receptor kinase
G-type lectin S-receptor-like serine/threonine-protein kinase (Fragment)
G-type lectin S-receptor-like serine/threonine-protein kinase
Serine/threonine protein kinase
G-type lectin S-receptor-like serine/threonine-protein kinase (Fragment)
G-type lectin S-receptor-like serine/threonine-protein kinase (Fragment)
Serine/threonine protein kinase (Fragment)
G-type lectin S-receptor-like serine/threonine-protein kinase (Fragment)
G-type lectin S-receptor-like serine/threonine-protein kinase (Fragment)
G-type lectin S-receptor-like serine/threonine protein kinase SD3-1-like (Fragment)
G-type lectin S-receptor-like serine/threonine-protein kinase (Fragment)
G-type lectin S-receptor-like serine/threonine-protein kinase (Fragment)
G-type lectin S-receptor-like serine/threonine-protein kinase (Fragment)
G-type lectin S-receptor-like serine/threonine-protein kinase (Fragment)
G-type lectin S-receptor-like serine/threonine-protein kinase (Fragment)
Receptor-like serine/threonine-protein kinase SD1-8-like (Fragment)
Epidermis-specific secreted glycoprotein ep1-like
G-type lectin S-receptor-like serine/threonine-protein kinase (Fragment)
G-type lectin S-receptor-like serine/threonine-kinase
G-type lectin S-receptor-like serine/threonine-protein kinase (Fragment)
G-type lectin S-receptor-like serine/threonine-kinase (Fragment)
G-type lectin S-receptor-like serine/threonine-protein kinase
G-type lectin S-receptor-like serine/threonine-protein kinase (Fragment)
G-type lectin S-receptor-like serine/threonine-kinase (Fragment)
G-type lectin S-receptor-like serine/threonine-kinase
Receptor-like serine/threonine-protein kinase SD1-8-like protein

G-type lectin S-receptor-like serine/threonine-kinase
Receptor protein kinase 1-like protein
G-type lectin S-receptor-like serine/threonine-protein kinase
PAN domain-containing protein
Receptor-like serine/threonine-protein kinase SD1-8-like protein
G-type lectin S-receptor-like serine/threonine-protein kinase
G-type lectin S-receptor-like serine/threonine-protein kinase
Receptor-like serine/threonine-protein kinase SD1-8-like protein
G-type lectin S-receptor-like serine/threonine-protein kinase
G-type lectin S-receptor-like serine/threonine-kinase
G-type lectin S-receptor-like serine/threonine-protein kinase
G-type lectin S-receptor-like serine/threonine-protein kinase (Fragment)
G-type lectin S-receptor-like serine/threonine-protein kinase (Fragment)
Serine/threonine protein kinase (Fragment)
G-type lectin S-receptor-like serine/threonine-protein kinase (Fragment)
G-type lectin S-receptor-like serine/threonine-kinase (Fragment)
G-type lectin S-receptor-like serine/threonine-protein kinase (Fragment)
G-type lectin S-receptor-like serine/threonine-protein kinase
Receptor-like serine/threonine-protein kinase SD1-8-like protein
G-type lectin S-receptor-like serine/threonine-kinase (Fragment)
G-type lectin S-receptor-like serine/threonine-protein kinase
G-type lectin S-receptor-like serine/threonine-kinase
Serine/threonine protein kinase (Fragment)
G-type lectin S-receptor-like serine/threonine-kinase (Fragment)
G-type lectin S-receptor-like serine/threonine-protein kinase (Fragment)
G-type lectin S-receptor-like serine/threonine-kinase
Serine/threonine protein kinase
Receptor-like serine/threonine-protein kinase SD1-8-like protein (Fragment)
Uncharacterized protein
G-type lectin S-receptor-like serine/threonine-kinase
Serine/threonine protein kinase (Fragment)
G-type lectin S-receptor-like serine/threonine-protein kinase (Fragment)
G-type lectin S-receptor-like serine/threonine-protein kinase
Serine/threonine protein kinase (Fragment)
G-type lectin S-receptor-like serine/threonine-protein kinase (Fragment)
Receptor protein kinase 1-like protein
Serine/threonine protein kinase (Fragment)
G-type lectin S-receptor-like serine/threonine-protein kinase (Fragment)
G-type lectin S-receptor-like serine/threonine-kinase
G-type lectin S-receptor-like serine/threonine-protein kinase
Serine/threonine kinase family protein (Fragment)

Cysteine-rich receptor-like protein kinase
G-type lectin S-receptor-like serine/threonine-protein kinase
Serine/threonine protein kinase (Fragment)
G-type lectin S-receptor-like serine/threonine-protein kinase (Fragment)
Serine/threonine protein kinase
G-type lectin S-receptor-like serine/threonine-protein kinase (Fragment)
G-type lectin S-receptor-like serine/threonine-protein kinase
G-type lectin S-receptor-like serine/threonine-kinase (Fragment)
G-type lectin S-receptor-like serine/threonine-protein kinase (Fragment)
Receptor-like serine/threonine-protein kinase SD1-8-like protein
G-type lectin S-receptor-like serine/threonine-protein kinase (Fragment)
G-type lectin S-receptor-like serine/threonine-kinase
G-type lectin S-receptor-like serine/threonine-protein kinase (Fragment)
G-type lectin S-receptor-like serine/threonine-kinase
G-type lectin S-receptor-like serine/threonine-kinase (Fragment)
G-type lectin S-receptor-like serine/threonine-protein kinase (Fragment)
G-type lectin S-receptor-like serine/threonine-kinase
G-type lectin S-receptor-like serine/threonine-protein kinase (Fragment)
G-type lectin S-receptor-like serine/threonine-kinase (Fragment)
G-type lectin S-receptor-like serine/threonine-protein kinase (Fragment)
G-type lectin S-receptor-like serine/threonine-kinase (Fragment)
Cysteine-rich receptor-like protein kinase (Fragment)
Serine/threonine protein kinase (Fragment)
G-type lectin S-receptor-like serine/threonine-protein kinase (Fragment)
G-type lectin S-receptor-like serine/threonine-protein kinase (Fragment)
Receptor-like serine/threonine-protein kinase SD1-8-like protein (Fragment)
G-type lectin S-receptor-like serine/threonine-protein kinase (Fragment)
G-type lectin S-receptor-like serine/threonine-protein kinase
G-type lectin S-receptor-like serine/threonine-kinase
G-type lectin S-receptor-like serine/threonine-protein kinase
G-type lectin S-receptor-like serine/threonine-protein kinase (Fragment)
G-type lectin S-receptor-like serine/threonine-kinase
Serine/threonine protein kinase
Serine/threonine protein kinase
G-type lectin S-receptor-like serine/threonine-protein kinase (Fragment)
G-type lectin S-receptor-like serine/threonine-kinase (Fragment)
G-type lectin S-receptor-like serine/threonine-protein kinase
G-type lectin S-receptor-like serine/threonine-protein kinase
G-type lectin S-receptor-like serine/threonine-protein kinase (Fragment)
Uncharacterized protein
Uncharacterized protein

Uncharacterized protein
Uncharacterized protein
Uncharacterized protein
Uncharacterized protein (Fragment)
Uncharacterized protein
Uncharacterized protein
Uncharacterized protein
Uncharacterized protein
Uncharacterized protein
Uncharacterized protein
Uncharacterized protein
Uncharacterized protein
Uncharacterized protein
Uncharacterized protein
Uncharacterized protein
Uncharacterized protein
Uncharacterized protein
Uncharacterized protein
Uncharacterized protein
Uncharacterized protein
Uncharacterized protein
Uncharacterized protein
Uncharacterized protein
Uncharacterized protein
Uncharacterized protein
Uncharacterized protein
Uncharacterized protein
Uncharacterized protein
Uncharacterized protein
Uncharacterized protein
Uncharacterized protein
Uncharacterized protein
Uncharacterized protein
Uncharacterized protein
Uncharacterized protein
Uncharacterized protein
Uncharacterized protein
Uncharacterized protein
Uncharacterized protein
Uncharacterized protein
Uncharacterized protein

Uncharacterized protein
Uncharacterized protein
Uncharacterized protein
Uncharacterized protein
Uncharacterized protein
Uncharacterized protein
Uncharacterized protein
Uncharacterized protein
Uncharacterized protein
Uncharacterized protein
Uncharacterized protein
Uncharacterized protein
Uncharacterized protein
Uncharacterized protein
Uncharacterized protein
Uncharacterized protein
Uncharacterized protein
Uncharacterized protein
Uncharacterized protein
Uncharacterized protein
Uncharacterized protein
Uncharacterized protein
Uncharacterized protein
Uncharacterized protein
Uncharacterized protein
Uncharacterized protein
Uncharacterized protein
Uncharacterized protein
Uncharacterized protein
Uncharacterized protein
Serine/threonine-protein kinase
Serine/threonine-protein kinase
Serine/threonine-protein kinase
Serine/threonine-protein kinase
Serine/threonine-protein kinase
Uncharacterized protein
Uncharacterized protein
Serine/threonine-protein kinase
Serine/threonine-protein kinase
Serine/threonine-protein kinase
Serine/threonine-protein kinase

Uncharacterized protein
Serine/threonine-protein kinase
Serine/threonine-protein kinase
Uncharacterized protein
Serine/threonine-protein kinase
Uncharacterized protein
Uncharacterized protein
Serine/threonine-protein kinase
Serine/threonine-protein kinase
Serine/threonine-protein kinase
Serine/threonine-protein kinase
Serine/threonine-protein kinase
Serine/threonine-protein kinase
Uncharacterized protein
Uncharacterized protein
Serine/threonine-protein kinase
Uncharacterized protein
Uncharacterized protein
Serine/threonine-protein kinase
Uncharacterized protein
Uncharacterized protein
Serine/threonine-protein kinase
Uncharacterized protein
Uncharacterized protein
Uncharacterized protein
Uncharacterized protein
Uncharacterized protein
Uncharacterized protein
Uncharacterized protein
Uncharacterized protein
Uncharacterized protein
Uncharacterized protein
Serine/threonine-protein kinase
Uncharacterized protein
Uncharacterized protein
Serine/threonine-protein kinase
Uncharacterized protein
Uncharacterized protein
Uncharacterized protein
Serine/threonine-protein kinase
Serine/threonine-protein kinase

Serine/threonine-protein kinase
Uncharacterized protein
Uncharacterized protein
Serine/threonine-protein kinase
Uncharacterized protein
Uncharacterized protein
S-locus receptor kinase
Uncharacterized protein
Serine/threonine-protein kinase
Serine/threonine-protein kinase
Uncharacterized protein
Uncharacterized protein
Serine/threonine-protein kinase
Uncharacterized protein
Serine/threonine-protein kinase
Uncharacterized protein
Uncharacterized protein
Uncharacterized protein
Serine/threonine-protein kinase
Serine/threonine-protein kinase
Uncharacterized protein
Uncharacterized protein
Serine/threonine-protein kinase
Uncharacterized protein
Uncharacterized protein
Uncharacterized protein
Uncharacterized protein
Serine/threonine-protein kinase
Uncharacterized protein
Serine/threonine-protein kinase
Serine/threonine-protein kinase
Serine/threonine-protein kinase
Uncharacterized protein
Serine/threonine-protein kinase
Uncharacterized protein
Uncharacterized protein
Serine/threonine-protein kinase
Serine/threonine-protein kinase
Uncharacterized protein
Serine/threonine-protein kinase
Uncharacterized protein

Uncharacterized protein
Serine/threonine-protein kinase
Serine/threonine-protein kinase
Uncharacterized protein
Uncharacterized protein
Uncharacterized protein
Uncharacterized protein
Uncharacterized protein
Serine/threonine-protein kinase
Serine/threonine-protein kinase
Uncharacterized protein
Uncharacterized protein
Uncharacterized protein
Serine/threonine-protein kinase
Uncharacterized protein
Uncharacterized protein
Uncharacterized protein
Uncharacterized protein
Uncharacterized protein
Serine/threonine-protein kinase
Serine/threonine-protein kinase
Serine/threonine-protein kinase
Serine/threonine-protein kinase
Serine/threonine-protein kinase
Uncharacterized protein
Serine/threonine-protein kinase
Uncharacterized protein
Serine/threonine-protein kinase
Uncharacterized protein
Uncharacterized protein
Uncharacterized protein
Serine/threonine-protein kinase
Serine/threonine-protein kinase
Uncharacterized protein
Uncharacterized protein
Serine/threonine-protein kinase
Serine/threonine-protein kinase
Uncharacterized protein
Serine/threonine-protein kinase
Uncharacterized protein
Serine/threonine-protein kinase

Serine/threonine-protein kinase
Serine/threonine-protein kinase
Serine/threonine-protein kinase
Serine/threonine-protein kinase
Uncharacterized protein
Serine/threonine-protein kinase
Serine/threonine-protein kinase
Serine/threonine-protein kinase
Serine/threonine-protein kinase
Serine/threonine-protein kinase
Serine/threonine-protein kinase
Uncharacterized protein
Uncharacterized protein
Serine/threonine-protein kinase
Uncharacterized protein
Uncharacterized protein
Serine/threonine-protein kinase
Uncharacterized protein
Serine/threonine-protein kinase
Uncharacterized protein
Serine/threonine-protein kinase
Serine/threonine-protein kinase
Uncharacterized protein
Uncharacterized protein
Uncharacterized protein
Uncharacterized protein
Serine/threonine-protein kinase
Serine/threonine-protein kinase
Uncharacterized protein
Serine/threonine-protein kinase
Uncharacterized protein
Serine/threonine-protein kinase
Uncharacterized protein
Uncharacterized protein
Uncharacterized protein
Uncharacterized protein
Serine/threonine-protein kinase
Serine/threonine-protein kinase
Serine/threonine-protein kinase
Serine/threonine-protein kinase
Serine/threonine-protein kinase

Uncharacterized protein
Serine/threonine-protein kinase
Uncharacterized protein
Serine/threonine-protein kinase
Uncharacterized protein
Serine/threonine-protein kinase
Uncharacterized protein
Uncharacterized protein
Serine/threonine-protein kinase
Uncharacterized protein
Uncharacterized protein
Uncharacterized protein
Uncharacterized protein
Uncharacterized protein
Serine/threonine-protein kinase
Uncharacterized protein
Uncharacterized protein
Serine/threonine-protein kinase
Serine/threonine-protein kinase
Uncharacterized protein
Uncharacterized protein
Serine/threonine-protein kinase
Uncharacterized protein
Uncharacterized protein
Uncharacterized protein
Uncharacterized protein
Serine/threonine-protein kinase
Uncharacterized protein
Uncharacterized protein
Uncharacterized protein
Serine/threonine-protein kinase
Uncharacterized protein
Uncharacterized protein
Uncharacterized protein
Serine/threonine-protein kinase
Uncharacterized protein
Uncharacterized protein
Serine/threonine-protein kinase
Serine/threonine-protein kinase
Serine/threonine-protein kinase
Uncharacterized protein

Serine/threonine-protein kinase
Serine/threonine-protein kinase
Uncharacterized protein
Serine/threonine-protein kinase
Serine/threonine-protein kinase
Serine/threonine-protein kinase
Serine/threonine-protein kinase
Serine/threonine-protein kinase
Uncharacterized protein
Serine/threonine-protein kinase
Uncharacterized protein
Serine/threonine-protein kinase
Serine/threonine-protein kinase
Serine/threonine-protein kinase
Uncharacterized protein
Uncharacterized protein
Serine/threonine-protein kinase
Uncharacterized protein
Serine/threonine-protein kinase
Uncharacterized protein
Uncharacterized protein
Uncharacterized protein
Serine/threonine-protein kinase
Uncharacterized protein
Serine/threonine-protein kinase
Uncharacterized protein
Uncharacterized protein
Serine/threonine-protein kinase
Uncharacterized protein
Uncharacterized protein
Uncharacterized protein
Serine/threonine-protein kinase
Serine/threonine-protein kinase
Uncharacterized protein
Uncharacterized protein
Uncharacterized protein
Uncharacterized protein
Uncharacterized protein
Serine/threonine-protein kinase
Serine/threonine-protein kinase
Uncharacterized protein

Uncharacterized protein
Serine/threonine-protein kinase
Uncharacterized protein
Serine/threonine-protein kinase
Serine/threonine-protein kinase
Uncharacterized protein
Uncharacterized protein
Uncharacterized protein
Uncharacterized protein
Uncharacterized protein
Uncharacterized protein
Serine/threonine-protein kinase
Uncharacterized protein
Uncharacterized protein
Serine/threonine-protein kinase
Uncharacterized protein
Uncharacterized protein
Uncharacterized protein
Uncharacterized protein
Uncharacterized protein
Uncharacterized protein
Uncharacterized protein
Uncharacterized protein
Serine/threonine-protein kinase
Uncharacterized protein
Uncharacterized protein
Uncharacterized protein
Serine/threonine-protein kinase
Uncharacterized protein
Uncharacterized protein
Serine/threonine-protein kinase
Serine/threonine-protein kinase
Uncharacterized protein
Uncharacterized protein
Uncharacterized protein
Uncharacterized protein
Uncharacterized protein
Serine/threonine-protein kinase
Uncharacterized protein
Serine/threonine-protein kinase
Uncharacterized protein

Uncharacterized protein
Uncharacterized protein
Uncharacterized protein
Uncharacterized protein
Serine/threonine-protein kinase
Uncharacterized protein
Uncharacterized protein
Serine/threonine-protein kinase
Serine/threonine-protein kinase
Serine/threonine-protein kinase
Serine/threonine-protein kinase
Uncharacterized protein
Serine/threonine-protein kinase
Serine/threonine-protein kinase
Serine/threonine-protein kinase
Uncharacterized protein
Serine/threonine-protein kinase
Uncharacterized protein
Serine/threonine-protein kinase
Uncharacterized protein
Serine/threonine-protein kinase
Serine/threonine-protein kinase
Uncharacterized protein
Serine/threonine-protein kinase
Uncharacterized protein
Uncharacterized protein
Uncharacterized protein
Serine/threonine-protein kinase
Uncharacterized protein
Serine/threonine-protein kinase
Serine/threonine-protein kinase
Serine/threonine-protein kinase
Uncharacterized protein
Serine/threonine-protein kinase
Serine/threonine-protein kinase
Uncharacterized protein
Uncharacterized protein
Serine/threonine-protein kinase
Uncharacterized protein
Stress-induced receptor-like kinase A1b (Fragment)
Uncharacterized protein

Uncharacterized protein
Serine/threonine-protein kinase
Uncharacterized protein
Serine/threonine-protein kinase
Serine/threonine-protein kinase
Putative S-locus receptor kinase (Fragment)
Uncharacterized protein
Serine/threonine-protein kinase
Uncharacterized protein
Uncharacterized protein
Serine/threonine-protein kinase
Uncharacterized protein
Serine/threonine-protein kinase
Serine/threonine-protein kinase
Protein kinase 5, putative
Serine/threonine-protein kinase
Serine/threonine-protein kinase
Serine/threonine-protein kinase
Chromosome 3B-specific BAC library, contig ctg0954b
Uncharacterized protein
Uncharacterized protein
Uncharacterized protein
Uncharacterized protein
Serine/threonine-protein kinase
Uncharacterized protein
Uncharacterized protein
Serine/threonine-protein kinase
Uncharacterized protein
Uncharacterized protein
Serine/threonine-protein kinase
Uncharacterized protein
Uncharacterized protein
Uncharacterized protein
Uncharacterized protein
Uncharacterized protein
Uncharacterized protein
Serine/threonine-protein kinase
Uncharacterized protein
Uncharacterized protein
Uncharacterized protein
Uncharacterized protein

Uncharacterized protein
Uncharacterized protein
Serine/threonine-protein kinase
Uncharacterized protein
Uncharacterized protein
Uncharacterized protein
Serine/threonine-protein kinase
Uncharacterized protein
Serine/threonine-protein kinase
Serine/threonine-protein kinase
Uncharacterized protein
Serine/threonine-protein kinase
Uncharacterized protein
Uncharacterized protein
Uncharacterized protein
Serine/threonine-protein kinase
Serine/threonine-protein kinase
Uncharacterized protein
Serine/threonine-protein kinase
Uncharacterized protein
Uncharacterized protein
Uncharacterized protein
Uncharacterized protein
Uncharacterized protein
Serine/threonine-protein kinase
Serine/threonine-protein kinase
Serine/threonine-protein kinase
Uncharacterized protein
Serine/threonine-protein kinase
Serine/threonine-protein kinase
Uncharacterized protein
Uncharacterized protein
Serine/threonine-protein kinase
Serine/threonine-protein kinase
Uncharacterized protein
Serine/threonine-protein kinase
Uncharacterized protein
Serine/threonine-protein kinase
Stress-induced receptor-like kinase A1a (Fragment)
Uncharacterized protein
Serine/threonine-protein kinase

Serine/threonine-protein kinase
Serine/threonine-protein kinase
Serine/threonine-protein kinase
Serine/threonine-protein kinase
Serine/threonine-protein kinase
Uncharacterized protein
Uncharacterized protein
Uncharacterized protein
Serine/threonine-protein kinase
Uncharacterized protein
Serine/threonine-protein kinase
Uncharacterized protein
Serine/threonine-protein kinase
Serine/threonine-protein kinase
Uncharacterized protein
Serine/threonine-protein kinase
Uncharacterized protein
Serine/threonine-protein kinase
Serine/threonine-protein kinase
Serine/threonine-protein kinase
Serine/threonine-protein kinase
Serine/threonine-protein kinase
Uncharacterized protein
Serine/threonine-protein kinase
Serine/threonine-protein kinase
Serine/threonine-protein kinase
Serine/threonine-protein kinase
Serine/threonine-protein kinase
Uncharacterized protein
Serine/threonine-protein kinase
Serine/threonine-protein kinase
Uncharacterized protein
Serine/threonine-protein kinase
Serine/threonine-protein kinase
Serine/threonine-protein kinase
Serine/threonine-protein kinase
Uncharacterized protein
Serine/threonine-protein kinase
Serine/threonine-protein kinase
Serine/threonine-protein kinase
Serine/threonine-protein kinase

Uncharacterized protein
Serine/threonine-protein kinase
Serine/threonine-protein kinase
Uncharacterized protein
Serine/threonine-protein kinase
Serine/threonine-protein kinase
Uncharacterized protein
Serine/threonine-protein kinase
Serine/threonine-protein kinase
Serine/threonine-protein kinase
Serine/threonine-protein kinase
Serine/threonine-protein kinase
Serine/threonine-protein kinase
Serine/threonine-protein kinase
Serine/threonine-protein kinase
Uncharacterized protein
Serine/threonine-protein kinase
Uncharacterized protein
Uncharacterized protein
Serine/threonine-protein kinase
Serine/threonine-protein kinase
Uncharacterized protein
Uncharacterized protein
Serine/threonine-protein kinase
Uncharacterized protein
Serine/threonine-protein kinase
Uncharacterized protein
Serine/threonine-protein kinase
Serine/threonine-protein kinase
Serine/threonine-protein kinase
Uncharacterized protein
Uncharacterized protein
Uncharacterized protein
Serine/threonine-protein kinase
Uncharacterized protein
Uncharacterized protein
Serine/threonine-protein kinase
Uncharacterized protein
Serine/threonine-protein kinase
Serine/threonine-protein kinase
Serine/threonine-protein kinase

Uncharacterized protein
Serine/threonine-protein kinase
Serine/threonine-protein kinase
Serine/threonine-protein kinase
Uncharacterized protein
Uncharacterized protein
Serine/threonine-protein kinase
Uncharacterized protein
Serine/threonine-protein kinase
Uncharacterized protein
Uncharacterized protein
Uncharacterized protein
Uncharacterized protein
Serine/threonine-protein kinase
Uncharacterized protein
Uncharacterized protein
Serine/threonine-protein kinase
Uncharacterized protein
Serine/threonine-protein kinase
Uncharacterized protein
Serine/threonine-protein kinase
Serine/threonine-protein kinase
Uncharacterized protein
Uncharacterized protein
Serine/threonine-protein kinase
Serine/threonine-protein kinase
Uncharacterized protein
Serine/threonine-protein kinase
Uncharacterized protein
Uncharacterized protein
Serine/threonine-protein kinase
Serine/threonine-protein kinase
Uncharacterized protein
Serine/threonine-protein kinase
Serine/threonine-protein kinase
Serine/threonine-protein kinase
Uncharacterized protein
Serine/threonine-protein kinase
Uncharacterized protein
Uncharacterized protein
Uncharacterized protein

Serine/threonine-protein kinase
Serine/threonine-protein kinase
Uncharacterized protein
Uncharacterized protein
Uncharacterized protein
Uncharacterized protein
Uncharacterized protein
Serine/threonine-protein kinase
Uncharacterized protein
Serine/threonine-protein kinase
Uncharacterized protein
Serine/threonine-protein kinase
Uncharacterized protein
Uncharacterized protein
Serine/threonine-protein kinase
Uncharacterized protein
Uncharacterized protein
Serine/threonine-protein kinase
Uncharacterized protein
Serine/threonine-protein kinase
Uncharacterized protein
Uncharacterized protein
Uncharacterized protein
Uncharacterized protein
Uncharacterized protein
Uncharacterized protein
Uncharacterized protein
Serine/threonine-protein kinase
Serine/threonine-protein kinase
Serine/threonine-protein kinase
Uncharacterized protein
Serine/threonine-protein kinase
Serine/threonine-protein kinase
Uncharacterized protein
Uncharacterized protein
Serine/threonine-protein kinase
Serine/threonine-protein kinase
Serine/threonine-protein kinase
Serine/threonine-protein kinase
Serine/threonine-protein kinase
Serine/threonine-protein kinase

Uncharacterized protein
Uncharacterized protein
Serine/threonine-protein kinase
Uncharacterized protein
Uncharacterized protein
Uncharacterized protein
Uncharacterized protein
Serine/threonine-protein kinase
Uncharacterized protein
Serine/threonine-protein kinase
Serine/threonine-protein kinase
Uncharacterized protein
Serine/threonine-protein kinase
Serine/threonine-protein kinase
Uncharacterized protein
Uncharacterized protein
Uncharacterized protein
Uncharacterized protein
Uncharacterized protein
Uncharacterized protein
Uncharacterized protein
Uncharacterized protein
Uncharacterized protein
Uncharacterized protein
Serine/threonine-protein kinase
Uncharacterized protein
Uncharacterized protein
Uncharacterized protein
Serine/threonine-protein kinase
Uncharacterized protein
Serine/threonine-protein kinase
Uncharacterized protein
Serine/threonine-protein kinase
Uncharacterized protein
Serine/threonine-protein kinase
Uncharacterized protein
Serine/threonine-protein kinase
Uncharacterized protein
Serine/threonine-protein kinase
Serine/threonine-protein kinase
Serine/threonine-protein kinase

Serine/threonine-protein kinase
Serine/threonine-protein kinase
Serine/threonine-protein kinase
Serine/threonine-protein kinase
Serine/threonine-protein kinase
Uncharacterized protein
Serine/threonine-protein kinase
Serine/threonine-protein kinase
Uncharacterized protein
Serine/threonine-protein kinase
Serine/threonine-protein kinase
Uncharacterized protein
Serine/threonine-protein kinase
Uncharacterized protein
Uncharacterized protein
Uncharacterized protein
Uncharacterized protein
Uncharacterized protein
Uncharacterized protein
Uncharacterized protein
Uncharacterized protein
Uncharacterized protein
Uncharacterized protein
Uncharacterized protein
Uncharacterized protein
Uncharacterized protein
Uncharacterized protein
Uncharacterized protein
Uncharacterized protein
Uncharacterized protein
Uncharacterized protein
Uncharacterized protein
Uncharacterized protein
Uncharacterized protein
Uncharacterized protein
Uncharacterized protein
Uncharacterized protein
Uncharacterized protein
Uncharacterized protein
Uncharacterized protein
Uncharacterized protein

Uncharacterized protein
Uncharacterized protein
Uncharacterized protein
Uncharacterized protein
Uncharacterized protein
Uncharacterized protein
Uncharacterized protein
Uncharacterized protein
Uncharacterized protein
Uncharacterized protein
Uncharacterized protein
Uncharacterized protein
Uncharacterized protein
Uncharacterized protein
Uncharacterized protein
Uncharacterized protein
Uncharacterized protein
Uncharacterized protein
Uncharacterized protein
Uncharacterized protein
Uncharacterized protein
Uncharacterized protein
Uncharacterized protein
Uncharacterized protein
Uncharacterized protein
Uncharacterized protein
Uncharacterized protein
Uncharacterized protein
Uncharacterized protein
Uncharacterized protein
Uncharacterized protein
Uncharacterized protein
Uncharacterized protein
Uncharacterized protein
Uncharacterized protein
Uncharacterized protein
Uncharacterized protein
Uncharacterized protein
Uncharacterized protein
Uncharacterized protein
Uncharacterized protein

Uncharacterized protein
Uncharacterized protein
Uncharacterized protein
Uncharacterized protein
Uncharacterized protein
Uncharacterized protein
Uncharacterized protein
Uncharacterized protein
Uncharacterized protein
Uncharacterized protein
Uncharacterized protein
Uncharacterized protein
Uncharacterized protein
Uncharacterized protein
Uncharacterized protein
Uncharacterized protein
Uncharacterized protein
Uncharacterized protein
Uncharacterized protein
Uncharacterized protein
Uncharacterized protein
Uncharacterized protein
Uncharacterized protein
Uncharacterized protein
Uncharacterized protein
Uncharacterized protein
Uncharacterized protein
Uncharacterized protein
Uncharacterized protein
Uncharacterized protein
Uncharacterized protein
Uncharacterized protein
Uncharacterized protein
Uncharacterized protein
Uncharacterized protein
Uncharacterized protein
Uncharacterized protein
Uncharacterized protein
Uncharacterized protein
Uncharacterized protein
Uncharacterized protein

Uncharacterized protein
Uncharacterized protein
Uncharacterized protein
Uncharacterized protein
Uncharacterized protein
Uncharacterized protein
Uncharacterized protein
Uncharacterized protein
Uncharacterized protein
Uncharacterized protein
Uncharacterized protein
Uncharacterized protein
Uncharacterized protein
Uncharacterized protein
Uncharacterized protein
Uncharacterized protein
Uncharacterized protein
Uncharacterized protein
Uncharacterized protein
Uncharacterized protein
Uncharacterized protein
Uncharacterized protein
Uncharacterized protein
Uncharacterized protein
Uncharacterized protein
Uncharacterized protein
Uncharacterized protein
Uncharacterized protein
Uncharacterized protein
Uncharacterized protein
Uncharacterized protein
Uncharacterized protein
Uncharacterized protein
Uncharacterized protein
Uncharacterized protein
Uncharacterized protein
Uncharacterized protein
Uncharacterized protein
Uncharacterized protein
Uncharacterized protein
Uncharacterized protein

Uncharacterized protein
Uncharacterized protein
Uncharacterized protein
Uncharacterized protein
Uncharacterized protein
Uncharacterized protein
Uncharacterized protein
Uncharacterized protein
Uncharacterized protein
Uncharacterized protein
Uncharacterized protein
Uncharacterized protein
Uncharacterized protein
Uncharacterized protein
Uncharacterized protein
Uncharacterized protein
Uncharacterized protein
Uncharacterized protein
Uncharacterized protein
Uncharacterized protein
Uncharacterized protein
Uncharacterized protein
Uncharacterized protein
Uncharacterized protein
Uncharacterized protein
Uncharacterized protein
Uncharacterized protein
Uncharacterized protein
Uncharacterized protein
Uncharacterized protein
Uncharacterized protein
Uncharacterized protein
Uncharacterized protein
Uncharacterized protein
Uncharacterized protein
Uncharacterized protein
Uncharacterized protein
Uncharacterized protein
Uncharacterized protein
Uncharacterized protein
Uncharacterized protein

Uncharacterized protein
Uncharacterized protein
Uncharacterized protein
Uncharacterized protein
Uncharacterized protein
Uncharacterized protein
Uncharacterized protein
Uncharacterized protein
Uncharacterized protein
Uncharacterized protein
Uncharacterized protein
Uncharacterized protein
Uncharacterized protein
Uncharacterized protein
Uncharacterized protein
Uncharacterized protein
Uncharacterized protein
Uncharacterized protein
Uncharacterized protein
Uncharacterized protein
Uncharacterized protein
Uncharacterized protein
Uncharacterized protein
Uncharacterized protein
Uncharacterized protein
Uncharacterized protein
Uncharacterized protein
Uncharacterized protein
Uncharacterized protein
Uncharacterized protein
Uncharacterized protein
Uncharacterized protein
Uncharacterized protein
Uncharacterized protein
Uncharacterized protein
Uncharacterized protein
Uncharacterized protein
Uncharacterized protein
Uncharacterized protein
Uncharacterized protein
Uncharacterized protein

Uncharacterized protein
Uncharacterized protein
Uncharacterized protein
Uncharacterized protein
Uncharacterized protein
Uncharacterized protein
Uncharacterized protein
Uncharacterized protein
Uncharacterized protein
Uncharacterized protein
Uncharacterized protein
Uncharacterized protein
Uncharacterized protein
Uncharacterized protein
Uncharacterized protein
Uncharacterized protein
Uncharacterized protein
Uncharacterized protein
Uncharacterized protein
Uncharacterized protein
Uncharacterized protein
Uncharacterized protein
Uncharacterized protein
Uncharacterized protein
Uncharacterized protein
Uncharacterized protein
Uncharacterized protein
Uncharacterized protein
Uncharacterized protein
Uncharacterized protein
Uncharacterized protein
Uncharacterized protein
Uncharacterized protein
Uncharacterized protein
Uncharacterized protein
Uncharacterized protein
Uncharacterized protein
Uncharacterized protein
Uncharacterized protein
Uncharacterized protein
Uncharacterized protein

Uncharacterized protein
Uncharacterized protein
Uncharacterized protein
Uncharacterized protein
Uncharacterized protein
Uncharacterized protein
Uncharacterized protein
Uncharacterized protein
Uncharacterized protein
Uncharacterized protein
Uncharacterized protein
Uncharacterized protein
Uncharacterized protein
Uncharacterized protein
Uncharacterized protein
Uncharacterized protein
Uncharacterized protein
Uncharacterized protein
Uncharacterized protein
Uncharacterized protein
Uncharacterized protein
Uncharacterized protein
Uncharacterized protein
Uncharacterized protein
Uncharacterized protein
Uncharacterized protein
Uncharacterized protein
Uncharacterized protein
Uncharacterized protein
Uncharacterized protein
Uncharacterized protein
Uncharacterized protein
Uncharacterized protein
Uncharacterized protein
Uncharacterized protein
Uncharacterized protein
Uncharacterized protein
Uncharacterized protein
Uncharacterized protein
Uncharacterized protein
Uncharacterized protein

Uncharacterized protein
Uncharacterized protein
Uncharacterized protein
Uncharacterized protein
Uncharacterized protein
Uncharacterized protein
Uncharacterized protein
Uncharacterized protein
Uncharacterized protein
Uncharacterized protein
Uncharacterized protein
Uncharacterized protein
Uncharacterized protein
Uncharacterized protein
Uncharacterized protein
Uncharacterized protein
Uncharacterized protein
Uncharacterized protein
Uncharacterized protein
Uncharacterized protein
Uncharacterized protein
Uncharacterized protein
Uncharacterized protein
Uncharacterized protein
Uncharacterized protein
Uncharacterized protein
Uncharacterized protein
Uncharacterized protein
Uncharacterized protein
Uncharacterized protein
Uncharacterized protein
Uncharacterized protein
Uncharacterized protein
Uncharacterized protein
Uncharacterized protein
Uncharacterized protein
Uncharacterized protein
Uncharacterized protein
Uncharacterized protein
Uncharacterized protein
Uncharacterized protein

Uncharacterized protein
Uncharacterized protein
Uncharacterized protein
Uncharacterized protein
Uncharacterized protein
Uncharacterized protein
Uncharacterized protein
Uncharacterized protein
Uncharacterized protein
Uncharacterized protein
Uncharacterized protein
Uncharacterized protein
Uncharacterized protein
Uncharacterized protein
Uncharacterized protein
Uncharacterized protein
Uncharacterized protein
Uncharacterized protein
Uncharacterized protein
Uncharacterized protein
Uncharacterized protein
Uncharacterized protein
Uncharacterized protein
Uncharacterized protein
Uncharacterized protein
Uncharacterized protein
Uncharacterized protein
Uncharacterized protein
Uncharacterized protein
Uncharacterized protein
Uncharacterized protein
Uncharacterized protein
Uncharacterized protein
Uncharacterized protein
Uncharacterized protein
Uncharacterized protein
Uncharacterized protein
Uncharacterized protein
Uncharacterized protein
Uncharacterized protein
Uncharacterized protein

Uncharacterized protein
Uncharacterized protein
Uncharacterized protein
Uncharacterized protein
Uncharacterized protein
Uncharacterized protein
Uncharacterized protein
Uncharacterized protein
Uncharacterized protein
Uncharacterized protein
Uncharacterized protein
Uncharacterized protein
Uncharacterized protein
Uncharacterized protein
Uncharacterized protein
Uncharacterized protein
Uncharacterized protein
Uncharacterized protein
Uncharacterized protein
Uncharacterized protein
Uncharacterized protein
Uncharacterized protein
Uncharacterized protein
Uncharacterized protein
Uncharacterized protein
Uncharacterized protein
Uncharacterized protein
Uncharacterized protein
Uncharacterized protein
Uncharacterized protein
Uncharacterized protein
Uncharacterized protein
Uncharacterized protein
Uncharacterized protein
Uncharacterized protein
Uncharacterized protein
Uncharacterized protein
Uncharacterized protein
Uncharacterized protein
Uncharacterized protein
Uncharacterized protein

Uncharacterized protein
Uncharacterized protein
Uncharacterized protein
Uncharacterized protein
Uncharacterized protein
Uncharacterized protein
Uncharacterized protein
Uncharacterized protein
Uncharacterized protein
Uncharacterized protein
Uncharacterized protein
Uncharacterized protein
Uncharacterized protein
Uncharacterized protein
Uncharacterized protein
Uncharacterized protein
Uncharacterized protein
Uncharacterized protein
Uncharacterized protein
Uncharacterized protein
Uncharacterized protein
Uncharacterized protein
Uncharacterized protein
Uncharacterized protein
Uncharacterized protein
Uncharacterized protein
Uncharacterized protein
Uncharacterized protein
Uncharacterized protein
Uncharacterized protein
Uncharacterized protein
Uncharacterized protein
Uncharacterized protein
Uncharacterized protein
Uncharacterized protein
Stress-induced receptor-like kinase A1du
Uncharacterized protein
Uncharacterized protein
Uncharacterized protein
Uncharacterized protein
Uncharacterized protein

Uncharacterized protein
Uncharacterized protein
Uncharacterized protein
Uncharacterized protein
Uncharacterized protein
Uncharacterized protein
Uncharacterized protein
Uncharacterized protein
Uncharacterized protein
Uncharacterized protein
Uncharacterized protein
Uncharacterized protein
Uncharacterized protein
Uncharacterized protein
Uncharacterized protein
Uncharacterized protein
Uncharacterized protein
Uncharacterized protein
Uncharacterized protein
Uncharacterized protein
Uncharacterized protein
Uncharacterized protein
Uncharacterized protein
Uncharacterized protein
Uncharacterized protein
Uncharacterized protein
Uncharacterized protein
Uncharacterized protein
Uncharacterized protein
Uncharacterized protein
Uncharacterized protein
Uncharacterized protein
Uncharacterized protein
Uncharacterized protein
Uncharacterized protein
Uncharacterized protein
Uncharacterized protein
Uncharacterized protein
Uncharacterized protein
Uncharacterized protein
Uncharacterized protein

Uncharacterized protein
Uncharacterized protein
Uncharacterized protein
Uncharacterized protein
Uncharacterized protein
Uncharacterized protein
Uncharacterized protein
Uncharacterized protein
Uncharacterized protein
Uncharacterized protein
Uncharacterized protein
Uncharacterized protein
Uncharacterized protein
Uncharacterized protein
Uncharacterized protein
Uncharacterized protein
Uncharacterized protein
Uncharacterized protein
Uncharacterized protein
Uncharacterized protein
Uncharacterized protein
Uncharacterized protein
Uncharacterized protein
Uncharacterized protein
Uncharacterized protein
Uncharacterized protein
Uncharacterized protein
Uncharacterized protein
Uncharacterized protein
Uncharacterized protein
Uncharacterized protein
Uncharacterized protein
Uncharacterized protein
Uncharacterized protein
Uncharacterized protein
Uncharacterized protein
Uncharacterized protein
Uncharacterized protein
Uncharacterized protein
Uncharacterized protein
Uncharacterized protein

Uncharacterized protein
Uncharacterized protein
Uncharacterized protein
Uncharacterized protein
Uncharacterized protein
Uncharacterized protein
Uncharacterized protein
Uncharacterized protein
Uncharacterized protein
Uncharacterized protein
Uncharacterized protein
Uncharacterized protein
Uncharacterized protein
Uncharacterized protein
Uncharacterized protein
Uncharacterized protein
Uncharacterized protein
Uncharacterized protein
Uncharacterized protein
Uncharacterized protein
Uncharacterized protein
Uncharacterized protein
Uncharacterized protein
Uncharacterized protein
Uncharacterized protein
Uncharacterized protein
Uncharacterized protein
Uncharacterized protein
Uncharacterized protein
Uncharacterized protein
Uncharacterized protein
Uncharacterized protein
Uncharacterized protein
Uncharacterized protein
Uncharacterized protein
Uncharacterized protein
Uncharacterized protein
Uncharacterized protein
Uncharacterized protein
Uncharacterized protein
Uncharacterized protein

Uncharacterized protein
Uncharacterized protein
Uncharacterized protein
Uncharacterized protein
Uncharacterized protein
Uncharacterized protein
Uncharacterized protein
Uncharacterized protein
Uncharacterized protein
Uncharacterized protein
Uncharacterized protein
Uncharacterized protein
Uncharacterized protein
Uncharacterized protein
Uncharacterized protein
Uncharacterized protein
Uncharacterized protein
Uncharacterized protein
Uncharacterized protein
Uncharacterized protein
Uncharacterized protein
Uncharacterized protein
Uncharacterized protein
Uncharacterized protein
Uncharacterized protein
Uncharacterized protein
Uncharacterized protein
Uncharacterized protein
Uncharacterized protein
Uncharacterized protein
Uncharacterized protein
Uncharacterized protein
Uncharacterized protein
Uncharacterized protein
Uncharacterized protein
Uncharacterized protein
Uncharacterized protein
Uncharacterized protein
Uncharacterized protein
Uncharacterized protein
Uncharacterized protein

Uncharacterized protein
Uncharacterized protein
Uncharacterized protein
Uncharacterized protein
Uncharacterized protein
Uncharacterized protein
Uncharacterized protein
Uncharacterized protein
Uncharacterized protein
Uncharacterized protein
Uncharacterized protein
Uncharacterized protein
Uncharacterized protein
Uncharacterized protein
Uncharacterized protein
Uncharacterized protein
Uncharacterized protein
Uncharacterized protein
Uncharacterized protein
Uncharacterized protein
Uncharacterized protein
Uncharacterized protein
Uncharacterized protein
Uncharacterized protein
Uncharacterized protein
Uncharacterized protein
Uncharacterized protein
Uncharacterized protein
Uncharacterized protein
Uncharacterized protein
Uncharacterized protein
Uncharacterized protein
Uncharacterized protein
Uncharacterized protein
Uncharacterized protein
Uncharacterized protein
Uncharacterized protein
Uncharacterized protein
Uncharacterized protein
Uncharacterized protein
Uncharacterized protein

Uncharacterized protein
Uncharacterized protein
Uncharacterized protein
Uncharacterized protein
Uncharacterized protein
Uncharacterized protein
Uncharacterized protein
Uncharacterized protein
Uncharacterized protein
Uncharacterized protein
Uncharacterized protein
Uncharacterized protein
Uncharacterized protein
Uncharacterized protein
Uncharacterized protein
Uncharacterized protein
Uncharacterized protein
Uncharacterized protein
Uncharacterized protein
Uncharacterized protein
Uncharacterized protein
Uncharacterized protein
Uncharacterized protein
Uncharacterized protein
Uncharacterized protein
Uncharacterized protein
Uncharacterized protein
Uncharacterized protein
Uncharacterized protein
Uncharacterized protein
Uncharacterized protein
Uncharacterized protein
Uncharacterized protein
Uncharacterized protein
Uncharacterized protein
Uncharacterized protein
Uncharacterized protein
Uncharacterized protein
Uncharacterized protein
Uncharacterized protein
Uncharacterized protein

Uncharacterized protein
Uncharacterized protein
Receptor-like serine/threonine-protein kinase SD1-8
Receptor-like serine/threonine-protein kinase SD1-8
G-type lectin S-receptor-like serine/threonine-protein kinase SD2-5
Receptor-like serine/threonine-protein kinase SD1-8
Receptor-like serine/threonine-protein kinase SD1-8
Receptor-like serine/threonine-protein kinase SD1-8
Putative receptor protein kinase ZmPK1
G-type lectin S-receptor-like serine/threonine-protein kinase SD2-5
Putative receptor protein kinase ZmPK1
Putative receptor protein kinase ZmPK1
Putative receptor protein kinase ZmPK1
Putative receptor protein kinase ZmPK1
Putative receptor protein kinase ZmPK1
Putative receptor protein kinase ZmPK1
Receptor-like serine/threonine-protein kinase SD1-8
Putative receptor protein kinase ZmPK1
G-type lectin S-receptor-like serine/threonine-protein kinase SD2-5
G-type lectin S-receptor-like serine/threonine-protein kinase SD2-5
Putative receptor protein kinase ZmPK1
G-type lectin S-receptor-like serine/threonine-protein kinase RKS1
G-type lectin S-receptor-like serine/threonine-protein kinase SD2-5
Putative receptor protein kinase ZmPK1
Putative receptor protein kinase ZmPK1
Receptor-like serine/threonine-protein kinase SD1-8
Putative receptor protein kinase ZmPK1
Putative receptor protein kinase ZmPK1
Putative receptor protein kinase ZmPK1
Putative receptor protein kinase ZmPK1
Uncharacterized protein
Serine/threonine-protein kinase
Serine/threonine-protein kinase
Uncharacterized protein
Uncharacterized protein
Serine/threonine-protein kinase
Serine/threonine-protein kinase
Serine/threonine-protein kinase
Uncharacterized protein
Uncharacterized protein
Serine/threonine-protein kinase

Uncharacterized protein
Uncharacterized protein
Serine/threonine-protein kinase
Uncharacterized protein
Uncharacterized protein
Uncharacterized protein
Uncharacterized protein
Uncharacterized protein
Serine/threonine-protein kinase
Uncharacterized protein
Serine/threonine-protein kinase
Uncharacterized protein
Serine/threonine-protein kinase
Serine/threonine-protein kinase
Uncharacterized protein
Uncharacterized protein
Serine/threonine-protein kinase
Serine/threonine-protein kinase
Serine/threonine-protein kinase
Uncharacterized protein
Uncharacterized protein
Serine/threonine-protein kinase
Serine/threonine-protein kinase
Serine/threonine-protein kinase
Serine/threonine-protein kinase
Serine/threonine-protein kinase
Serine/threonine-protein kinase
Uncharacterized protein
Serine/threonine-protein kinase
Uncharacterized protein
Serine/threonine-protein kinase
Serine/threonine-protein kinase
Serine/threonine-protein kinase
Uncharacterized protein
Serine/threonine-protein kinase
Uncharacterized protein
Serine/threonine-protein kinase
Uncharacterized protein
Serine/threonine-protein kinase
Serine/threonine-protein kinase
Serine/threonine-protein kinase

Serine/threonine-protein kinase
Uncharacterized protein
Serine/threonine-protein kinase
Serine/threonine-protein kinase
Serine/threonine-protein kinase
Serine/threonine-protein kinase
Serine/threonine-protein kinase
Uncharacterized protein
Uncharacterized protein
Uncharacterized protein
Uncharacterized protein
Serine/threonine-protein kinase
Serine/threonine-protein kinase
Uncharacterized protein
Serine/threonine-protein kinase
Serine/threonine-protein kinase
Serine/threonine-protein kinase
Serine/threonine-protein kinase
Uncharacterized protein
Serine/threonine-protein kinase
Uncharacterized protein
Serine/threonine-protein kinase
Serine/threonine-protein kinase
Uncharacterized protein
Uncharacterized protein
Serine/threonine-protein kinase
Serine/threonine-protein kinase
Serine/threonine-protein kinase
Serine/threonine-protein kinase
Uncharacterized protein
Serine/threonine-protein kinase
Serine/threonine-protein kinase
Uncharacterized protein
Uncharacterized protein
Serine/threonine-protein kinase
Serine/threonine-protein kinase
Serine/threonine-protein kinase
Putative receptor protein kinase ZmPK1
Putative serine/threonine-protein kinase receptor
Putative receptor protein kinase ZmPK1
Uncharacterized protein

PAN domain protein
Uncharacterized protein
Uncharacterized protein
Uncharacterized protein
Uncharacterized protein (Fragment)
Uncharacterized protein (Fragment)
Uncharacterized protein (Fragment)
Uncharacterized protein (Fragment)
Uncharacterized protein
Uncharacterized protein
Uncharacterized protein (Fragment)
Hepatocyte growth factor
Coagulation factor XI
Cytochrome P450 4V2
Hepatocyte growth factor-like protein
Uncharacterized protein
Uncharacterized protein
Uncharacterized protein
Uncharacterized protein
Uncharacterized protein
Putative PAN domain-containing protein
Uncharacterized protein
Uncharacterized protein
Uncharacterized protein
Uncharacterized protein
Plasminogen
hepatocyte growth factor-like protein isoform X1
Plasminogen
coagulation factor XI isoform X2
Plasminogen
Plasminogen
Hepatocyte growth factor
coagulation factor XI isoform X1
Plasminogen (Fragment)
Plasminogen (Fragment)
PAN-4 domain
Apple-like
Alpha-2-macroglobulin protein (Fragment)
Large extracellular alpha-helical protein
Coagulation factor XI
Uncharacterized protein

Plasminogen
Kallikrein B1
plasma kallikrein isoform X4
Plasminogen
plasma kallikrein isoform X2
Hepatocyte growth factor
plasma kallikrein isoform X3
hepatocyte growth factor-like protein isoform X1
hepatocyte growth factor-like protein isoform X2
coagulation factor XI
Hepatocyte growth factor
plasma kallikrein isoform X1
Plasminogen
plasma kallikrein
Hepatocyte growth factor
hepatocyte growth factor-like protein
Hepatocyte growth factor
LOW QUALITY PROTEIN: coagulation factor XI
Uncharacterized protein
Uncharacterized protein
Uncharacterized protein
Uncharacterized protein
Uncharacterized protein
Uncharacterized protein
Uncharacterized protein
Uncharacterized protein
Uncharacterized protein
Uncharacterized protein
Uncharacterized protein
Uncharacterized protein
Uncharacterized protein
Uncharacterized protein
Uncharacterized protein
Uncharacterized protein
Uncharacterized protein
Uncharacterized protein
Uncharacterized protein
Uncharacterized protein
Uncharacterized protein
Uncharacterized protein
PAN domain protein

Uncharacterized protein
Uncharacterized protein
N/apple PAN domain protein
Uncharacterized protein
PAN domain-containing protein
Type I phosphodiesterase / nucleotide pyrophosphatase
Uncharacterized protein
Uncharacterized protein
Uncharacterized protein
Uncharacterized protein
Uncharacterized protein
Uncharacterized protein (Fragment)
Uncharacterized protein
Uncharacterized protein
Galactose oxidase
Galactose oxidase
Uncharacterized protein
Uncharacterized protein
Uncharacterized protein
Uncharacterized protein (Fragment)
Uncharacterized protein
Uncharacterized protein (Fragment)
Uncharacterized protein
Uncharacterized protein
Uncharacterized protein
Uncharacterized protein
Uncharacterized protein
Uncharacterized protein
Uncharacterized protein
Uncharacterized protein
Uncharacterized protein
Uncharacterized protein
Uncharacterized protein
Uncharacterized protein
Uncharacterized protein
Uncharacterized protein
Uncharacterized protein
Uncharacterized protein
Uncharacterized protein
Uncharacterized protein
Uncharacterized protein

Uncharacterized protein
Uncharacterized protein
Uncharacterized protein
Uncharacterized protein
Uncharacterized protein
Uncharacterized protein
Uncharacterized protein
Uncharacterized protein
Uncharacterized protein
Uncharacterized protein
Uncharacterized protein
Uncharacterized protein
Uncharacterized protein
Uncharacterized protein
Uncharacterized protein
Uncharacterized protein
Uncharacterized protein
Uncharacterized protein
Uncharacterized protein
Uncharacterized protein
Uncharacterized protein
Uncharacterized protein
Uncharacterized protein
Uncharacterized protein
Uncharacterized protein
Uncharacterized protein
Uncharacterized protein
Uncharacterized protein
Uncharacterized protein
Uncharacterized protein
Uncharacterized protein
Uncharacterized protein
Uncharacterized protein
Uncharacterized protein
Uncharacterized protein
Uncharacterized protein
Uncharacterized protein
Uncharacterized protein
Uncharacterized protein
Uncharacterized protein
Uncharacterized protein

Uncharacterized protein
Uncharacterized protein
Uncharacterized protein
Uncharacterized protein
Uncharacterized protein
G-type lectin S-receptor-like serine/threonine-protein kinase At1g11300
G-type lectin S-receptor-like serine/threonine-protein kinase SD3-1
G-type lectin S-receptor-like serine/threonine-protein kinase B120
receptor-like serine/threonine-protein kinase SD1-8
G-type lectin S-receptor-like serine/threonine-protein kinase At2g19130 isoform X2
G-type lectin S-receptor-like serine/threonine-protein kinase At4g27290 isoform X1
G-type lectin S-receptor-like serine/threonine-protein kinase At2g19130 isoform X1
G-type lectin S-receptor-like serine/threonine-protein kinase At4g27290
G-type lectin S-receptor-like serine/threonine-protein kinase At5g35370
G-type lectin S-receptor-like serine/threonine-protein kinase At4g27290 isoform X1
G-type lectin S-receptor-like serine/threonine-protein kinase At4g27290 isoform X1
G-type lectin S-receptor-like serine/threonine-protein kinase At4g27290
G-type lectin S-receptor-like serine/threonine-protein kinase At2g19130
PAN domain-containing protein At5g03700
G-type lectin S-receptor-like serine/threonine-protein kinase At4g27290
receptor-like serine/threonine-protein kinase SD1-8
G-type lectin S-receptor-like serine/threonine-protein kinase CES101 isoform X1
G-type lectin S-receptor-like serine/threonine-protein kinase CES101 isoform X4
G-type lectin S-receptor-like serine/threonine-protein kinase At4g03230 isoform X2
G-type lectin S-receptor-like serine/threonine-protein kinase At1g67520 isoform X1
EP1-like glycoprotein 4
putative receptor protein kinase ZmPK1
G-type lectin S-receptor-like serine/threonine-protein kinase At2g19130
G-type lectin S-receptor-like serine/threonine-protein kinase At1g11300
G-type lectin S-receptor-like serine/threonine-protein kinase At4g27290 isoform X2
G-type lectin S-receptor-like serine/threonine-protein kinase At4g03230 isoform X2
G-type lectin S-receptor-like serine/threonine-protein kinase At1g11300
G-type lectin S-receptor-like serine/threonine-protein kinase At4g27290
G-type lectin S-receptor-like serine/threonine-protein kinase At1g11300 isoform X1
G-type lectin S-receptor-like serine/threonine-protein kinase At4g27290 isoform X1
G-type lectin S-receptor-like serine/threonine-protein kinase At4g27290 isoform X2
G-type lectin S-receptor-like serine/threonine-protein kinase At4g27290
G-type lectin S-receptor-like serine/threonine-protein kinase At1g11300
G-type lectin S-receptor-like serine/threonine-protein kinase SD2-2
epidermis-specific secreted glycoprotein EP1
putative receptor protein kinase ZmPK1

G-type lectin S-receptor-like serine/threonine-protein kinase At4g27290 isoform X2
G-type lectin S-receptor-like serine/threonine-protein kinase At4g27290 isoform X2
G-type lectin S-receptor-like serine/threonine-protein kinase At1g67520 isoform X2
putative serine/threonine-protein kinase receptor
G-type lectin S-receptor-like serine/threonine-protein kinase At4g27290 isoform X3
putative receptor protein kinase ZmPK1
G-type lectin S-receptor-like serine/threonine-protein kinase SD2-5
putative receptor protein kinase ZmPK1
G-type lectin S-receptor-like serine/threonine-protein kinase RKS1
G-type lectin S-receptor-like serine/threonine-protein kinase At4g03230 isoform X1
G-type lectin S-receptor-like serine/threonine-protein kinase At1g11300
G-type lectin S-receptor-like serine/threonine-protein kinase SD1-1 isoform X2
G-type lectin S-receptor-like serine/threonine-protein kinase At4g27290 isoform X1
G-type lectin S-receptor-like serine/threonine-protein kinase At4g27290
G-type lectin S-receptor-like serine/threonine-protein kinase At4g27290 isoform X2
G-type lectin S-receptor-like serine/threonine-protein kinase At4g27290 isoform X2
G-type lectin S-receptor-like serine/threonine-protein kinase At1g11300
G-type lectin S-receptor-like serine/threonine-protein kinase At4g27290
putative receptor protein kinase ZmPK1
G-type lectin S-receptor-like serine/threonine-protein kinase B120 isoform X1
G-type lectin S-receptor-like serine/threonine-protein kinase At4g27290
G-type lectin S-receptor-like serine/threonine-protein kinase At4g27290 isoform X2
G-type lectin S-receptor-like serine/threonine-protein kinase At4g27290 isoform X1
G-type lectin S-receptor-like serine/threonine-protein kinase RKS1
epidermis-specific secreted glycoprotein EP1-like
G-type lectin S-receptor-like serine/threonine-protein kinase At1g11330
G-type lectin S-receptor-like serine/threonine-protein kinase At1g11330 isoform X2
uncharacterized protein LOC106759721
G-type lectin S-receptor-like serine/threonine-protein kinase At4g03230 isoform X2
G-type lectin S-receptor-like serine/threonine-protein kinase At4g03230 isoform X1
G-type lectin S-receptor-like serine/threonine-protein kinase At4g27290 isoform X1
G-type lectin S-receptor-like serine/threonine-protein kinase At4g03230 isoform X1
G-type lectin S-receptor-like serine/threonine-protein kinase At1g11330 isoform X1
putative receptor protein kinase ZmPK1
G-type lectin S-receptor-like serine/threonine-protein kinase At4g27290 isoform X1
G-type lectin S-receptor-like serine/threonine-protein kinase At4g27290 isoform X2
G-type lectin S-receptor-like serine/threonine-protein kinase CES101 isoform X2
G-type lectin S-receptor-like serine/threonine-protein kinase CES101 isoform X3
G-type lectin S-receptor-like serine/threonine-protein kinase At1g11410
G-type lectin S-receptor-like serine/threonine-protein kinase B120 isoform X2
G-type lectin S-receptor-like serine/threonine-protein kinase At1g34300

G-type lectin S-receptor-like serine/threonine-protein kinase At4g27290 isoform X1
G-type lectin S-receptor-like serine/threonine-protein kinase At1g67520
G-type lectin S-receptor-like serine/threonine-protein kinase At1g11300
putative receptor protein kinase ZmPK1
receptor-like serine/threonine-protein kinase SD1-6 isoform X3
G-type lectin S-receptor-like serine/threonine-protein kinase At4g27290
Uncharacterized protein
G-type lectin S-receptor-like serine/threonine-protein kinase
Uncharacterized protein
G-type lectin S-receptor-like serine/threonine-protein kinase CES101
G-type lectin S-receptor-like serine/threonine-protein kinase
Uncharacterized protein
Uncharacterized protein
Receptor-like serine/threonine-protein kinase SD1-8
Uncharacterized protein
Uncharacterized protein
Uncharacterized protein
Uncharacterized protein
Uncharacterized protein
G-type lectin S-receptor-like serine/threonine-protein kinase
G-type lectin S-receptor-like serine/threonine-protein kinase SD3-1
G-type lectin S-receptor-like serine/threonine-protein kinase
EP1-like glycoprotein 2
Uncharacterized protein
Uncharacterized protein
Uncharacterized protein
Uncharacterized protein
Uncharacterized protein
Uncharacterized protein
Uncharacterized protein
Uncharacterized protein
Uncharacterized protein
Uncharacterized protein
Uncharacterized protein
Uncharacterized protein
Uncharacterized protein
Uncharacterized protein
Uncharacterized protein
Uncharacterized protein (Fragment)
Uncharacterized protein
Uncharacterized protein

Uncharacterized protein
Uncharacterized protein
Uncharacterized protein
Uncharacterized protein
Uncharacterized protein
Uncharacterized protein
Uncharacterized protein
G-type lectin S-receptor-like serine/threonine-protein kinase
G-type lectin S-receptor-like serine/threonine-protein kinase
G-type lectin S-receptor-like serine/threonine-protein kinase LECRK3
Uncharacterized protein
Uncharacterized protein
Uncharacterized protein (Fragment)
Uncharacterized protein
Uncharacterized protein
Uncharacterized protein
Uncharacterized protein
Uncharacterized protein
Uncharacterized protein
Uncharacterized protein
Uncharacterized protein
Uncharacterized protein
Uncharacterized protein
Uncharacterized protein
Uncharacterized protein
Uncharacterized protein
Uncharacterized protein
Uncharacterized protein
Uncharacterized protein
Uncharacterized protein
G-type lectin S-receptor-like serine/threonine-protein kinase SD2-5
Uncharacterized protein
Uncharacterized protein
Uncharacterized protein
Uncharacterized protein
Uncharacterized protein
Uncharacterized protein
Uncharacterized protein
Uncharacterized protein
Uncharacterized protein
Uncharacterized protein

Uncharacterized protein
Uncharacterized protein
Uncharacterized protein
Uncharacterized protein
Uncharacterized protein
Uncharacterized protein
Uncharacterized protein
Uncharacterized protein
Uncharacterized protein
Uncharacterized protein
Uncharacterized protein
Uncharacterized protein
Uncharacterized protein
Uncharacterized protein
G-type lectin S-receptor-like serine/threonine-protein kinase RKS1
Receptor-like serine/threonine-protein kinase SD1-8
G-type lectin S-receptor-like serine/threonine-protein kinase
Uncharacterized protein
G-type lectin S-receptor-like serine/threonine-protein kinase
G-type lectin S-receptor-like serine/threonine-protein kinase
G-type lectin S-receptor-like serine/threonine-protein kinase
G-type lectin S-receptor-like serine/threonine-protein kinase SD2-5
Receptor-like serine/threonine-protein kinase SD1-6
Uncharacterized protein
G-type lectin S-receptor-like serine/threonine-protein kinase
G-type lectin S-receptor-like serine/threonine-protein kinase LECRK3
G-type lectin S-receptor-like serine/threonine-protein kinase B120
G-type lectin S-receptor-like serine/threonine-protein kinase
G-type lectin S-receptor-like serine/threonine-protein kinase LECRK4
G-type lectin S-receptor-like serine/threonine-protein kinase
G-type lectin S-receptor-like serine/threonine-protein kinase B120
G-type lectin S-receptor-like serine/threonine-protein kinase
G-type lectin S-receptor-like serine/threonine-protein kinase
G-type lectin S-receptor-like serine/threonine-protein kinase B120
Receptor-like serine/threonine-protein kinase SD1-8
Uncharacterized protein
Uncharacterized protein
G-type lectin S-receptor-like serine/threonine-protein kinase RKS1
Uncharacterized protein
G-type lectin S-receptor-like serine/threonine-protein kinase RKS1
G-type lectin S-receptor-like serine/threonine-protein kinase LECRK1 (Fragment)

G-type lectin S-receptor-like serine/threonine-protein kinase
G-type lectin S-receptor-like serine/threonine-protein kinase
G-type lectin S-receptor-like serine/threonine-protein kinase
Putative receptor protein kinase ZmPK1
G-type lectin S-receptor-like serine/threonine-protein kinase
G-type lectin S-receptor-like serine/threonine-protein kinase
PAN domain-containing protein
Putative receptor protein kinase ZmPK1
G-type lectin S-receptor-like serine/threonine-protein kinase
G-type lectin S-receptor-like serine/threonine-protein kinase
G-type lectin S-receptor-like serine/threonine-protein kinase
G-type lectin S-receptor-like serine/threonine-protein kinase
G-type lectin S-receptor-like serine/threonine-protein kinase RKS1
Uncharacterized protein
G-type lectin S-receptor-like serine/threonine-protein kinase
G-type lectin S-receptor-like serine/threonine-protein kinase
Putative receptor protein kinase ZmPK1
Receptor-like serine/threonine-protein kinase SD1-8
G-type lectin S-receptor-like serine/threonine-protein kinase SD1-1
Uncharacterized protein
G-type lectin S-receptor-like serine/threonine-protein kinase SD2-5
Receptor-like serine/threonine-protein kinase SD1-8
G-type lectin S-receptor-like serine/threonine-protein kinase RKS1
Uncharacterized protein
Putative receptor protein kinase ZmPK1
G-type lectin S-receptor-like serine/threonine-protein kinase
Uncharacterized protein
G-type lectin S-receptor-like serine/threonine-protein kinase
Uncharacterized protein
G-type lectin S-receptor-like serine/threonine-protein kinase SD2-5
G-type lectin S-receptor-like serine/threonine-protein kinase
Receptor-like serine/threonine-protein kinase SD1-8
Putative receptor protein kinase ZmPK1
G-type lectin S-receptor-like serine/threonine-protein kinase SD2-5
G-type lectin S-receptor-like serine/threonine-protein kinase SD2-5
Receptor-like serine/threonine-protein kinase SD1-8
G-type lectin S-receptor-like serine/threonine-protein kinase
G-type lectin S-receptor-like serine/threonine-protein kinase
G-type lectin S-receptor-like serine/threonine-protein kinase
G-type lectin S-receptor-like serine/threonine-protein kinase LECRK3
Putative receptor protein kinase ZmPK1

G-type lectin S-receptor-like serine/threonine-protein kinase
G-type lectin S-receptor-like serine/threonine-protein kinase
Uncharacterized protein
Receptor-like serine/threonine-protein kinase SD1-7
G-type lectin S-receptor-like serine/threonine-protein kinase
G-type lectin S-receptor-like serine/threonine-protein kinase
G-type lectin S-receptor-like serine/threonine-protein kinase
Uncharacterized protein
G-type lectin S-receptor-like serine/threonine-protein kinase B120
G-type lectin S-receptor-like serine/threonine-protein kinase CES101
G-type lectin S-receptor-like serine/threonine-protein kinase B120
Receptor-like serine/threonine-protein kinase SD1-8
Receptor-like serine/threonine-protein kinase SD1-8
G-type lectin S-receptor-like serine/threonine-protein kinase RKS1
EP1-like glycoprotein 2
G-type lectin S-receptor-like serine/threonine-protein kinase
G-type lectin S-receptor-like serine/threonine-protein kinase (Fragment)
G-type lectin S-receptor-like serine/threonine-protein kinase
Uncharacterized protein
G-type lectin S-receptor-like serine/threonine-protein kinase RKS1
G-type lectin S-receptor-like serine/threonine-protein kinase
G-type lectin S-receptor-like serine/threonine-protein kinase
G-type lectin S-receptor-like serine/threonine-protein kinase
G-type lectin S-receptor-like serine/threonine-protein kinase
G-type lectin S-receptor-like serine/threonine-protein kinase
G-type lectin S-receptor-like serine/threonine-protein kinase
G-type lectin S-receptor-like serine/threonine-protein kinase SD3-1
G-type lectin S-receptor-like serine/threonine-protein kinase
G-type lectin S-receptor-like serine/threonine-protein kinase
G-type lectin S-receptor-like serine/threonine-protein kinase B120
Receptor-like serine/threonine-protein kinase SD1-8
G-type lectin S-receptor-like serine/threonine-protein kinase
G-type lectin S-receptor-like serine/threonine-protein kinase CES101
Receptor-like serine/threonine-protein kinase SD1-8
G-type lectin S-receptor-like serine/threonine-protein kinase
G-type lectin S-receptor-like serine/threonine-protein kinase
G-type lectin S-receptor-like serine/threonine-protein kinase
Receptor-like serine/threonine-protein kinase SD1-8
G-type lectin S-receptor-like serine/threonine-protein kinase RKS1
Uncharacterized protein
Receptor-like serine/threonine-protein kinase SD1-8

G-type lectin S-receptor-like serine/threonine-protein kinase
G-type lectin S-receptor-like serine/threonine-protein kinase
Uncharacterized protein
G-type lectin S-receptor-like serine/threonine-protein kinase SD3-1
G-type lectin S-receptor-like serine/threonine-protein kinase
G-type lectin S-receptor-like serine/threonine-protein kinase
G-type lectin S-receptor-like serine/threonine-protein kinase
G-type lectin S-receptor-like serine/threonine-protein kinase
G-type lectin S-receptor-like serine/threonine-protein kinase
Receptor-like serine/threonine-protein kinase SD1-8
G-type lectin S-receptor-like serine/threonine-protein kinase
G-type lectin S-receptor-like serine/threonine-protein kinase
G-type lectin S-receptor-like serine/threonine-protein kinase SD2-5
Receptor-like serine/threonine-protein kinase SD1-8
G-type lectin S-receptor-like serine/threonine-protein kinase
Receptor-like serine/threonine-protein kinase SD1-8
G-type lectin S-receptor-like serine/threonine-protein kinase
G-type lectin S-receptor-like serine/threonine-protein kinase
G-type lectin S-receptor-like serine/threonine-protein kinase SD2-5
G-type lectin S-receptor-like serine/threonine-protein kinase
G-type lectin S-receptor-like serine/threonine-protein kinase RKS1
G-type lectin S-receptor-like serine/threonine-protein kinase LECRK1
Cysteine-rich receptor-like protein kinase 7
G-type lectin S-receptor-like serine/threonine-protein kinase
G-type lectin S-receptor-like serine/threonine-protein kinase
G-type lectin S-receptor-like serine/threonine-protein kinase
G-type lectin S-receptor-like serine/threonine-protein kinase
Uncharacterized protein
G-type lectin S-receptor-like serine/threonine-protein kinase
G-type lectin S-receptor-like serine/threonine-protein kinase LECRK1
EP1-like glycoprotein 2
Receptor-like serine/threonine-protein kinase SD1-8
G-type lectin S-receptor-like serine/threonine-protein kinase
G-type lectin S-receptor-like serine/threonine-protein kinase
Uncharacterized protein
G-type lectin S-receptor-like serine/threonine-protein kinase RKS1
G-type lectin S-receptor-like serine/threonine-protein kinase RKS1
G-type lectin S-receptor-like serine/threonine-protein kinase
G-type lectin S-receptor-like serine/threonine-protein kinase
G-type lectin S-receptor-like serine/threonine-protein kinase SD2-5
Uncharacterized protein

Uncharacterized protein
Uncharacterized protein
G-type lectin S-receptor-like serine/threonine-protein kinase
Uncharacterized protein
Uncharacterized protein
Uncharacterized protein
Uncharacterized protein
Uncharacterized protein
Uncharacterized protein
Uncharacterized protein
Uncharacterized protein
Uncharacterized protein
Uncharacterized protein
Uncharacterized protein
Uncharacterized protein
Uncharacterized protein
Uncharacterized protein
Uncharacterized protein
Uncharacterized protein
Uncharacterized protein
Uncharacterized protein
Uncharacterized protein
Uncharacterized protein
Uncharacterized protein
Uncharacterized protein
Uncharacterized protein
Uncharacterized protein
Uncharacterized protein
Uncharacterized protein
Uncharacterized protein
Uncharacterized protein
Uncharacterized protein
Uncharacterized protein
Uncharacterized protein
Uncharacterized protein (Fragment)
G-type lectin S-receptor-like serine/threonine-protein kinase SD1-29
G-type lectin S-receptor-like serine/threonine-protein kinase SD1-1
G-type lectin S-receptor-like serine/threonine-protein kinase
G-type lectin S-receptor-like serine/threonine-protein kinase
Receptor-like serine/threonine-protein kinase SD1-8
G-type lectin S-receptor-like serine/threonine-protein kinase LECRK1

Uncharacterized protein
Uncharacterized protein
G-type lectin S-receptor-like serine/threonine-protein kinase SD2-5
Uncharacterized protein
Uncharacterized protein
Uncharacterized protein
Uncharacterized protein
Uncharacterized protein
Receptor-like serine/threonine-protein kinase SD1-8
Uncharacterized protein
Uncharacterized protein
Uncharacterized protein
Uncharacterized protein
Putative receptor protein kinase ZmPK1
G-type lectin S-receptor-like serine/threonine-protein kinase
G-type lectin S-receptor-like serine/threonine-protein kinase
Putative receptor protein kinase ZmPK1
G-type lectin S-receptor-like serine/threonine-protein kinase
G-type lectin S-receptor-like serine/threonine-protein kinase
G-type lectin S-receptor-like serine/threonine-protein kinase
G-type lectin S-receptor-like serine/threonine-protein kinase LECRK3
G-type lectin S-receptor-like serine/threonine-protein kinase
G-type lectin S-receptor-like serine/threonine-protein kinase RKS1
G-type lectin S-receptor-like serine/threonine-protein kinase
G-type lectin S-receptor-like serine/threonine-protein kinase
G-type lectin S-receptor-like serine/threonine-protein kinase
Uncharacterized protein
G-type lectin S-receptor-like serine/threonine-protein kinase
G-type lectin S-receptor-like serine/threonine-protein kinase
G-type lectin S-receptor-like serine/threonine-protein kinase
G-type lectin S-receptor-like serine/threonine-protein kinase
G-type lectin S-receptor-like serine/threonine-protein kinase CES101
G-type lectin S-receptor-like serine/threonine-protein kinase
Putative receptor protein kinase ZmPK1
Uncharacterized protein
G-type lectin S-receptor-like serine/threonine-protein kinase
G-type lectin S-receptor-like serine/threonine-protein kinase
G-type lectin S-receptor-like serine/threonine-protein kinase
G-type lectin S-receptor-like serine/threonine-protein kinase B120
G-type lectin S-receptor-like serine/threonine-protein kinase RKS1
G-type lectin S-receptor-like serine/threonine-protein kinase

G-type lectin S-receptor-like serine/threonine-protein kinase
G-type lectin S-receptor-like serine/threonine-protein kinase
Putative receptor protein kinase ZmPK1
G-type lectin S-receptor-like serine/threonine-protein kinase
Putative receptor protein kinase ZmPK1
G-type lectin S-receptor-like serine/threonine-protein kinase LECRK3
G-type lectin S-receptor-like serine/threonine-protein kinase
G-type lectin S-receptor-like serine/threonine-protein kinase
Receptor-like serine/threonine-protein kinase SD1-7
Receptor-like serine/threonine-protein kinase SD1-8
Putative receptor protein kinase ZmPK1
S-locus-specific glycoprotein S13
Uncharacterized protein
Receptor-like serine/threonine-protein kinase SD1-8
Uncharacterized protein
Uncharacterized protein
Uncharacterized protein
Uncharacterized protein
Uncharacterized protein
Uncharacterized protein
Uncharacterized protein
Uncharacterized protein
G-type lectin S-receptor-like serine/threonine-protein kinase
Putative receptor protein kinase ZmPK1
Uncharacterized protein (Fragment)
Uncharacterized protein
Uncharacterized protein
Uncharacterized protein
Uncharacterized protein
Uncharacterized protein
Uncharacterized protein
Uncharacterized protein
Uncharacterized protein
Uncharacterized protein
Uncharacterized protein
Uncharacterized protein
Uncharacterized protein
Uncharacterized protein
Uncharacterized protein
Uncharacterized protein
Uncharacterized protein

Uncharacterized protein
Uncharacterized protein
Uncharacterized protein
Uncharacterized protein
Uncharacterized protein
Uncharacterized protein
Uncharacterized protein
Uncharacterized protein
Uncharacterized protein
Uncharacterized protein
Uncharacterized protein
Uncharacterized protein
Uncharacterized protein
Uncharacterized protein
Uncharacterized protein
Uncharacterized protein
Uncharacterized protein
Uncharacterized protein
Uncharacterized protein
Uncharacterized protein
Uncharacterized protein
Uncharacterized protein
Uncharacterized protein
Uncharacterized protein
Uncharacterized protein
Uncharacterized protein
Uncharacterized protein
Uncharacterized protein
Uncharacterized protein
Uncharacterized protein
Uncharacterized protein
Uncharacterized protein
Uncharacterized protein
Uncharacterized protein
Uncharacterized protein
Uncharacterized protein
Uncharacterized protein
Uncharacterized protein
Uncharacterized protein
Uncharacterized protein
Uncharacterized protein

Uncharacterized protein
Uncharacterized protein
Uncharacterized protein
Uncharacterized protein
Uncharacterized protein
Uncharacterized protein
Uncharacterized protein
Uncharacterized protein
Uncharacterized protein
Uncharacterized protein
Uncharacterized protein
Uncharacterized protein
Uncharacterized protein
Uncharacterized protein
Uncharacterized protein
Uncharacterized protein
Uncharacterized protein
Uncharacterized protein
Uncharacterized protein
Uncharacterized protein
Uncharacterized protein
Uncharacterized protein
Uncharacterized protein
Uncharacterized protein
Uncharacterized protein
Uncharacterized protein
Uncharacterized protein
Uncharacterized protein
Uncharacterized protein
Uncharacterized protein
Uncharacterized protein
Uncharacterized protein
Uncharacterized protein
Uncharacterized protein
Uncharacterized protein
Uncharacterized protein
Uncharacterized protein
Uncharacterized protein
Uncharacterized protein
Uncharacterized protein
Uncharacterized protein

Uncharacterized protein
Uncharacterized protein (Fragment)
Uncharacterized protein
Uncharacterized protein
Uncharacterized protein
Uncharacterized protein
Uncharacterized protein
Uncharacterized protein
Uncharacterized protein
Uncharacterized protein
Uncharacterized protein
Uncharacterized protein
Uncharacterized protein
Uncharacterized protein
VMP4 protein
Uncharacterized protein
Uncharacterized protein
Uncharacterized protein
Uncharacterized protein
Uncharacterized protein
Uncharacterized protein
Uncharacterized protein
Uncharacterized protein
Uncharacterized protein
Uncharacterized protein
Uncharacterized protein
Uncharacterized protein
Uncharacterized protein
Uncharacterized protein
Uncharacterized protein
Uncharacterized protein
Uncharacterized protein
Uncharacterized protein
Uncharacterized protein
Uncharacterized protein
Uncharacterized protein
Uncharacterized protein
coagulation factor XI
Hepatocyte growth factor
hepatocyte growth factor-like protein isoform X1
plasma kallikrein isoform X3

Hepatocyte growth factor
plasma kallikrein isoform X2
plasma kallikrein isoform X1
Plasminogen
Large extracellular alpha-helical protein
Wenxma_12, whole genome shotgun sequence
Uncharacterized protein
TSA: Wollemia nobilis Ref_Wollemi_Transcript_14214_3070 transcribed RNA sequence
TSA: Wollemia nobilis Ref_Wollemi_Transcript_14839_2719 transcribed RNA sequence
TSA: Wollemia nobilis Ref_Wollemi_Transcript_11276_3376 transcribed RNA sequence
TSA: Wollemia nobilis Ref_Wollemi_Transcript_14539_2766 transcribed RNA sequence
TSA: Wollemia nobilis Ref_Wollemi_Transcript_14348_2869 transcribed RNA sequence
TSA: Wollemia nobilis Ref_Wollemi_Transcript_14840_2775 transcribed RNA sequence
TSA: Wollemia nobilis Ref_Wollemi_Transcript_14540_2769 transcribed RNA sequence
Uncharacterized protein
Uncharacterized protein (Fragment)
Uncharacterized protein
Uncharacterized protein
Uncharacterized protein (Fragment)
Uncharacterized protein
PAN domain-containing protein (Fragment)
Uncharacterized protein
PAN domain-containing protein
Uncharacterized protein
Uncharacterized protein (Fragment)
PAN domain-containing protein (Fragment)
Uncharacterized protein (Fragment)
Uncharacterized protein
Uncharacterized protein (Fragment)
Uncharacterized protein
Uncharacterized protein
Uncharacterized protein (Fragment)
Uncharacterized protein
Uncharacterized protein
Uncharacterized protein
Uncharacterized protein
Uncharacterized protein
Uncharacterized protein
Uncharacterized protein
Uncharacterized protein (Fragment)
PAN domain-containing protein (Fragment)

Uncharacterized protein
Uncharacterized protein
Uncharacterized protein
Uncharacterized protein
PAN domain-containing protein (Fragment)
PAN domain-containing protein (Fragment)
PAN domain-containing protein
Uncharacterized protein (Fragment)
Uncharacterized protein
Uncharacterized protein
Uncharacterized protein (Fragment)
Uncharacterized protein (Fragment)
Uncharacterized protein
PAN domain-containing protein (Fragment)
Uncharacterized protein
Uncharacterized protein
PAN domain-containing protein (Fragment)
PAN domain-containing protein
Uncharacterized protein (Fragment)
Uncharacterized protein (Fragment)
Uncharacterized protein
Uncharacterized protein
Uncharacterized protein
Uncharacterized protein
PAN domain-containing protein
PAN domain-containing protein (Fragment)
Uncharacterized protein
Uncharacterized protein
Growth factor Livertine
Uncharacterized protein
Uncharacterized protein (Fragment)
Uncharacterized protein
MGC54016 protein
Hepatocyte growth factor
Hepatocyte growth factor
LOC397993 protein
Klkb1-prov protein
Hepatocyte growth factor-like protein
Macrophage-stimulating 1
Kallikrein B1
Hepatocyte growth factor

Hepatocyte growth factor
Uncharacterized protein
Uncharacterized protein
Uncharacterized protein
Uncharacterized protein
Plasminogen
Macrophage stimulating 1
Uncharacterized protein
Uncharacterized protein
Uncharacterized protein
Hepatocyte growth factor
Macrophage stimulating 1
Macrophage stimulating 1
Uncharacterized protein
Plasminogen
Uncharacterized protein
Uncharacterized protein
Uncharacterized protein
Uncharacterized protein
Uncharacterized protein
Uncharacterized protein
Plasminogen
Uncharacterized protein
Uncharacterized protein
Hepatocyte growth factor
Uncharacterized protein
Uncharacterized protein
Uncharacterized protein
Uncharacterized protein
PAN domain-containing protein
Uncharacterized protein
Uncharacterized protein
Putative receptor protein kinase ZmPK1
G-type lectin S-receptor-like serine/threonine-protein kinase B120
G-type lectin S-receptor-like serine/threonine-protein kinase SD2-2
Putative receptor protein kinase ZmPK1
G-type lectin S-receptor-like serine/threonine-protein kinase
Putative receptor protein kinase ZmPK1
G-type lectin S-receptor-like serine/threonine-protein kinase RKS1
G-type lectin S-receptor-like serine/threonine-protein kinase
Uncharacterized protein

G-type lectin S-receptor-like serine/threonine-protein kinase
Putative receptor protein kinase ZmPK1
G-type lectin S-receptor-like serine/threonine-protein kinase B120
Serine/threonine-protein kinase
G-type lectin S-receptor-like serine/threonine-protein kinase
Receptor-like serine/threonine-protein kinase SD1-8
Receptor-like protein kinase 4
G-type lectin S-receptor-like serine/threonine-protein kinase
G-type lectin S-receptor-like serine/threonine-protein kinase B120
Receptor-like protein kinase 4
G-type lectin S-receptor-like serine/threonine-protein kinase CES101
Receptor-like serine/threonine-protein kinase SD1-8
PAN domain-containing protein
S-locus-specific glycoprotein S13
Serine/threonine-protein kinase
Kinase interacting kinase1
Serine/threonine-protein kinase
Receptor-like serine/threonine-protein kinase SD1-8
Putative S-locus receptor-like protein kinase family protein
Serine/threonine-protein kinase
G-type lectin S-receptor-like serine/threonine-protein kinase B120
PAN domain-containing protein
Serine/threonine-protein kinase
Serine/threonine-protein kinase
G-type lectin S-receptor-like serine/threonine-protein kinase SD2-5
Putative S-locus-like receptor protein kinase family protein
Receptor-like serine/threonine-protein kinase SD1-8
Receptor-like protein kinase 4
Serine/threonine-protein kinase
G-type lectin S-receptor-like serine/threonine-protein kinase B120
G-type lectin S-receptor-like serine/threonine-protein kinase
S-domain class receptor-like kinase3
Serine/threonine-protein kinase
Serine/threonine-protein kinase
S-locus lectin kinase family protein
Kinase interacting kinase1
Serine/threonine-protein kinase
Serine/threonine-protein kinase
G-type lectin S-receptor-like serine/threonine-protein kinase B120
G-type lectin S-receptor-like serine/threonine-protein kinase B120
G-type lectin S-receptor-like serine/threonine-protein kinase B120

Receptor-like serine/threonine-protein kinase SD1-8
Receptor-like protein kinase 4
Serine/threonine-protein kinase
Putative S-locus receptor-like protein kinase family protein
G-type lectin S-receptor-like serine/threonine-protein kinase B120
G-type lectin S-receptor-like serine/threonine-protein kinase B120
Receptor-like protein kinase 4
Receptor-like protein kinase 4
Receptor-like protein kinase 4
Receptor-like protein kinase 4
Methionine aminopeptidase 1D chloroplastic/mitochondrial
Putative S-locus receptor-like protein kinase family protein
G-type lectin S-receptor-like serine/threonine-protein kinase B120
Receptor-like protein kinase 4
Serine/threonine-protein kinase
Serine/threonine-protein kinase
Receptor-like protein kinase 4
Receptor-like protein kinase 4
Uncharacterized protein
Serine/threonine-protein kinase
Receptor-like protein kinase 4
Receptor-like serine/threonine-protein kinase SD1-8
G-type lectin S-receptor-like serine/threonine-protein kinase
Receptor-like serine/threonine-protein kinase SD1-8
Uncharacterized protein
G-type lectin S-receptor-like serine/threonine-protein kinase B120
Putative receptor protein kinase ZmPK1
Serine/threonine-protein kinase receptor
G-type lectin S-receptor-like serine/threonine-protein kinase
G-type lectin S-receptor-like serine/threonine-protein kinase
Putative receptor protein kinase ZmPK1
G-type lectin S-receptor-like serine/threonine-protein kinase
Receptor-like kinase
G-type lectin S-receptor-like serine/threonine-protein kinase
G-type lectin S-receptor-like serine/threonine-protein kinase
Receptor-like serine/threonine-protein kinase SD1-8
Putative receptor protein kinase ZmPK1
G-type lectin S-receptor-like serine/threonine-protein kinase SD2-5
G-type lectin S-receptor-like serine/threonine-protein kinase B120
Uncharacterized protein
G-type lectin S-receptor-like serine/threonine-protein kinase

Ser/Thr protein kinase (Fragment)
G-type lectin S-receptor-like serine/threonine-protein kinase RKS1
Receptor-like protein kinase
Receptor-like serine/threonine-protein kinase SD1-8
PAN domain-containing protein
Uncharacterized protein
S-locus lectin protein kinase
Uncharacterized protein
Receptor-like protein kinase 4
Receptor-like protein kinase 4
G-type lectin S-receptor-like serine/threonine-protein kinase CES101
Serine/threonine-protein kinase
PAN domain-containing protein
Serine/threonine-protein kinase
Serine/threonine-protein kinase
G-type lectin S-receptor-like serine/threonine-protein kinase SD2-5
G-type lectin S-receptor-like serine/threonine-protein kinase RKS1
Receptor-like serine/threonine-protein kinase SD1-8
PAN domain-containing protein
Receptor-like serine/threonine-protein kinase SD1-8
Kinase interacting kinase1
Receptor-like protein kinase 4
PAN domain-containing protein
Receptor-like protein kinase 4
G-type lectin S-receptor-like serine/threonine-protein kinase B120
Serine/threonine-protein kinase
Serine/threonine-protein kinase
Putative S-locus-like receptor protein kinase family protein
Serine/threonine-protein kinase
PAN domain-containing protein
Cysteine-rich receptor-like protein kinase 6
G-type lectin S-receptor-like serine/threonine-protein kinase B120
G-type lectin S-receptor-like serine/threonine-protein kinase B120
PAN domain-containing protein
G-type lectin S-receptor-like serine/threonine-protein kinase B120
Receptor-like protein kinase 4
G-type lectin S-receptor-like serine/threonine-protein kinase B120
Serine/threonine-protein kinase
PAN domain-containing protein
G-type lectin S-receptor-like serine/threonine-protein kinase
G-type lectin S-receptor-like serine/threonine-protein kinase

Serine/threonine-protein kinase
Serine/threonine-protein kinase
Receptor-like serine/threonine-protein kinase SD1-8
G-type lectin S-receptor-like serine/threonine-protein kinase B120
G-type lectin S-receptor-like serine/threonine-protein kinase
G-type lectin S-receptor-like serine/threonine-protein kinase
G-type lectin S-receptor-like serine/threonine-protein kinase
KI domain interacting kinase 1
G-type lectin S-receptor-like serine/threonine-protein kinase
PAN domain-containing protein
G-type lectin S-receptor-like serine/threonine-protein kinase
G-type lectin S-receptor-like serine/threonine-protein kinase
G-type lectin S-receptor-like serine/threonine-protein kinase
G-type lectin S-receptor-like serine/threonine-protein kinase
Uncharacterized protein
G-type lectin S-receptor-like serine/threonine-protein kinase
Receptor-like serine/threonine-protein kinase SD1-8
Receptor-like serine/threonine-protein kinase SD1-8
Uncharacterized protein
Receptor-like serine/threonine-protein kinase SD1-8
G-type lectin S-receptor-like serine/threonine-protein kinase
PAN domain-containing protein
G-type lectin S-receptor-like serine/threonine-protein kinase
S-domain receptor-like protein kinase
G-type lectin S-receptor-like serine/threonine-protein kinase SD2-5
Uncharacterized protein
PAN domain-containing protein
Receptor-like serine/threonine-protein kinase SD1-8
Putative G-type lectin S-receptor-like serine/threonine-protein kinase
Uncharacterized protein
PAN domain-containing protein
G-type lectin S-receptor-like serine/threonine-protein kinase
G-type lectin S-receptor-like serine/threonine-protein kinase
Epidermis-specific secreted glycoprotein EP1
Receptor-like serine/threonine-protein kinase SD1-8
G-type lectin S-receptor-like serine/threonine-protein kinase
Receptor-like serine/threonine-protein kinase SD1-8
Receptor-like serine/threonine-protein kinase SD1-8
G-type lectin S-receptor-like serine/threonine-protein kinase B120
Putative receptor protein kinase ZmPK1
G-type lectin S-receptor-like serine/threonine-protein kinase

Putative receptor protein kinase ZmPK1
Putative receptor protein kinase ZmPK1
Uncharacterized protein
Cuticlin-1
Uncharacterized protein (Fragment)
Uncharacterized protein (Fragment)
N-(5'-phosphoribosyl)anthranilate isomerase
Cuticlin-1
Cuticlin-1 (Fragment)
Cuticlin-1
Uncharacterized protein F52C9.5
Uncharacterized protein
N-(5'-phosphoribosyl)anthranilate isomerase
Uncharacterized protein
Cuticlin-1
Uncharacterized protein
Uncharacterized protein
Uncharacterized protein
Neurotrypsin
S-locus receptor kinase (SRK)
S-locus receptor kinase (SRK)
S-locus receptor kinase (SRK)
S-locus receptor kinase (SRK)
S-locus receptor kinase (SRK)
S-locus receptor kinase (SRK)
S-locus receptor kinase (SRK)
Receptor-like protein kinase like protein
S-locus receptor kinase (SRK)
S-locus receptor kinase (SRK)
S-locus receptor kinase (SRK)
S-locus receptor kinase (SRK)
S-locus receptor kinase (SRK)
Uncharacterized protein
S-locus receptor kinase (SRK)
S-locus receptor kinase (SRK)
S-locus receptor kinase (SRK)
S-locus receptor kinase (SRK)
S-locus receptor kinase (SRK)
S-locus receptor kinase (SRK)
S-locus receptor kinase (SRK)
S-locus receptor kinase (SRK)

S-locus receptor kinase (SRK)
S-locus receptor kinase (SRK)
S-locus receptor kinase (SRK)
S-locus receptor kinase (SRK)
S-locus receptor kinase (SRK)
S-locus receptor kinase (SRK)
S-locus receptor kinase (SRK)
S-locus receptor kinase (SRK)
S-locus receptor kinase (SRK)
S-locus receptor kinase (SRK)
S-locus receptor kinase (SRK)
S-locus receptor kinase (SRK)
S-locus receptor kinase (SRK)
S-locus receptor kinase (SRK)
S-locus receptor kinase (SRK)
Receptor-like kinase
S-locus receptor kinase (SRK)
S-locus receptor kinase (SRK)
S-locus receptor kinase (SRK)
Uncharacterized protein
Uncharacterized protein
Uncharacterized protein
Uncharacterized protein
Uncharacterized protein
Uncharacterized protein
Uncharacterized protein
Uncharacterized protein
Uncharacterized protein
Uncharacterized protein
Uncharacterized protein
Uncharacterized protein
Uncharacterized protein
Uncharacterized protein
Hydrophobin-like protein
Uncharacterized protein
Uncharacterized protein
Ca2+-modulated nonselective cation channel polycystin
Uncharacterized protein
Uncharacterized protein
Uncharacterized protein
Ca2+-modulated nonselective cation channel polycystin

Uncharacterized protein
Uncharacterized protein
Uncharacterized protein
Uncharacterized protein
Uncharacterized protein
Uncharacterized protein
Uncharacterized protein
Uncharacterized protein
Uncharacterized protein
Uncharacterized protein
Uncharacterized protein
Uncharacterized protein
Uncharacterized protein
Uncharacterized protein
Uncharacterized protein
Uncharacterized protein
Uncharacterized protein
Uncharacterized protein
Uncharacterized protein
Uncharacterized protein
Uncharacterized protein
Uncharacterized protein
Uncharacterized protein
Uncharacterized protein
Uncharacterized protein
Uncharacterized protein
Uncharacterized protein
Uncharacterized protein
Uncharacterized protein
Uncharacterized protein
Uncharacterized protein
Uncharacterized protein
Uncharacterized protein
Uncharacterized protein
Uncharacterized protein
Uncharacterized protein
Uncharacterized protein
Uncharacterized protein
Uncharacterized protein
Uncharacterized protein
Uncharacterized protein

Uncharacterized protein
Uncharacterized protein
Uncharacterized protein
Uncharacterized protein
Uncharacterized protein
Uncharacterized protein
Uncharacterized protein
Uncharacterized protein
Uncharacterized protein
Uncharacterized protein
Uncharacterized protein
Uncharacterized protein

ıe 18C07, contigA1

ing protein
ning protein
tein
ntaining protein/PAN_2 domain-containing protein/DUF3403 domain-containing protein
ng protein
ing protein
ning protein
ning protein/Pkinase_Tyr domain-containing protein/PAN_2 domain-containing protein/DUF340

protein
ning protein/PAN_2 domain-containing protein

ng protein

a-alpha toroid

ain and C-type lectin-like domain and C-type lectin fold domain-containing protein

tive isoform 1

tive

tive

tive
tive

tive

tive

tive isoform 2

tive

)3 domain-conta...

**Table S2: GO terms exhibiting significant enrichment among 28,300 PAN domain-containing pro**

| GO.ID | Term | Annotated | Significant | Expected |
|---|---|---|---|---|
| GO:0005975 | carbohydrate metabolic process | 308 | 41 | 9.13 |
| GO:0006030 | chitin metabolic process | 97 | 17 | 2.88 |
| GO:0043170 | macromolecule metabolic process | 16450 | 547 | 487.81 |
| GO:0008152 | metabolic process | 16627 | 559 | 493.06 |
| GO:0006807 | nitrogen compound metabolic process | 16453 | 547 | 487.9 |
| GO:0071704 | organic substance metabolic process | 16544 | 558 | 490.6 |
| GO:1901564 | organonitrogen compound metabolic proces... | 16406 | 544 | 486.51 |
| GO:0044238 | primary metabolic process | 16510 | 558 | 489.59 |
| GO:0019538 | protein metabolic process | 16367 | 544 | 485.35 |
| GO:0006508 | proteolysis | 4803 | 543 | 142.43 |
| GO:0006865 | amino acid transport | 1 | 1 | 0 |
| GO:0006198 | cAMP catabolic process | 1 | 1 | 0 |
| GO:0005975 | carbohydrate metabolic process | 308 | 15 | 0.62 |
| GO:0009166 | nucleotide catabolic process | 1 | 1 | 0 |
| GO:0006508 | proteolysis | 4803 | 29 | 9.65 |
| GO:0006195 | purine nucleotide catabolic process | 1 | 1 | 0 |
| GO:0006163 | purine nucleotide metabolic process | 1 | 1 | 0 |
| GO:0009154 | purine ribonucleotide catabolic process | 1 | 1 | 0 |
| GO:0009150 | purine ribonucleotide metabolic process | 1 | 1 | 0 |
| GO:0055085 | transmembrane transport | 20 | 2 | 0.04 |
| GO:0035556 | intracellular signal transduction | 10 | 6 | 0.81 |
| GO:0043170 | macromolecule metabolic process | 16450 | 1529 | 1331.27 |
| GO:0008152 | metabolic process | 16627 | 1531 | 1345.59 |
| GO:0006807 | nitrogen compound metabolic process | 16453 | 1527 | 1331.51 |
| GO:0071704 | organic substance metabolic process | 16544 | 1530 | 1338.88 |
| GO:1901564 | organonitrogen compound metabolic proces... | 16406 | 1522 | 1327.71 |
| GO:0000160 | phosphorelay signal transduction system | 5 | 5 | 0.4 |
| GO:0044238 | primary metabolic process | 16510 | 1529 | 1336.12 |
| GO:0019538 | protein metabolic process | 16367 | 1521 | 1324.55 |
| GO:0006508 | proteolysis | 4803 | 1519 | 388.7 |
| GO:0005975 | carbohydrate metabolic process | 308 | 10 | 0.18 |
| GO:0005975 | carbohydrate metabolic process | 308 | 2 | 0.1 |
| GO:0006040 | amino sugar metabolic process | 97 | 80 | 10.06 |
| GO:0006022 | aminoglycan metabolic process | 98 | 81 | 10.16 |
| GO:1901135 | carbohydrate derivative metabolic proces... | 103 | 84 | 10.68 |
| GO:0006030 | chitin metabolic process | 97 | 80 | 10.06 |
| GO:0017144 | drug metabolic process | 97 | 80 | 10.06 |
| GO:1901071 | glucosamine-containing compound metaboli... | 97 | 80 | 10.06 |
| GO:0006508 | proteolysis | 4803 | 1687 | 498.12 |
| GO:0006979 | response to oxidative stress | 79 | 76 | 8.19 |
| GO:0050896 | response to stimulus | 113 | 96 | 11.72 |

| | | | | |
|---|---|---|---|---|
| GO:0006950 | response to stress | 82 | 78 | 8.5 |
| GO:0005975 | carbohydrate metabolic process | 308 | 141 | 15.5 |
| GO:0043170 | macromolecule metabolic process | 16450 | 939 | 827.8 |
| GO:0008152 | metabolic process | 16627 | 945 | 836.71 |
| GO:0006807 | nitrogen compound metabolic process | 16453 | 939 | 827.96 |
| GO:0071704 | organic substance metabolic process | 16544 | 943 | 832.53 |
| GO:1901564 | organonitrogen compound metabolic proces... | 16406 | 934 | 825.59 |
| GO:0055114 | oxidation-reduction process | 181 | 78 | 9.11 |
| GO:0044238 | primary metabolic process | 16510 | 943 | 830.82 |
| GO:0019538 | protein metabolic process | 16367 | 934 | 823.63 |
| GO:0006508 | proteolysis | 4803 | 932 | 241.7 |
| GO:0007154 | cell communication | 12011 | 11972 | 8768.58 |
| GO:0008037 | cell recognition | 11970 | 11970 | 8738.64 |
| GO:0009987 | cellular process | 14064 | 13773 | 10267.36 |
| GO:0006464 | cellular protein modification process | 11577 | 11539 | 8451.74 |
| GO:0006796 | phosphate-containing compound metabolic ... | 11538 | 11505 | 8423.26 |
| GO:0006793 | phosphorus metabolic process | 11538 | 11505 | 8423.26 |
| GO:0009875 | pollen-pistil interaction | 11970 | 11970 | 8738.64 |
| GO:0009856 | pollination | 11970 | 11970 | 8738.64 |
| GO:0006468 | protein phosphorylation | 11526 | 11501 | 8414.5 |
| GO:0000003 | reproduction | 12005 | 11970 | 8764.2 |
| GO:0006508 | proteolysis | 4803 | 28 | 7.11 |

oteins.

| classicFisher | Catergory |
|---|---|
| 7.00E-16 | Alveolata |
| 3.50E-09 | Alveolata |
| 4.00E-19 | Alveolata |
| 3.60E-29 | Alveolata |
| 4.30E-19 | Alveolata |
| 6.30E-29 | Alveolata |
| 1.90E-17 | Alveolata |
| 2.00E-29 | Alveolata |
| 6.50E-18 | Alveolata |
| 1.00E-30 | Alveolata |
| 0.00201 | Amoebozoa |
| 0.00201 | Amoebozoa |
| 1.20E-17 | Amoebozoa |
| 0.00201 | Amoebozoa |
| 6.70E-11 | Amoebozoa |
| 0.00201 | Amoebozoa |
| 0.00201 | Amoebozoa |
| 0.00201 | Amoebozoa |
| 0.00201 | Amoebozoa |
| 0.00073 | Amoebozoa |
| 4.40E-05 | Bacteria |
| 1.00E-30 | Bacteria |
| 1.00E-30 | Bacteria |
| 1.00E-30 | Bacteria |
| 1.00E-30 | Bacteria |
| 1.00E-30 | Bacteria |
| 3.50E-06 | Bacteria |
| 1.00E-30 | Bacteria |
| 1.00E-30 | Bacteria |
| 1.00E-30 | Bacteria |
| 1.20E-17 | Euglenozoa |
| 0.0038 | Haptophyceae |
| 1.00E-30 | Opisthokonta |
| 1.00E-30 | Opisthokonta |
| 1.00E-30 | Opisthokonta |
| 1.00E-30 | Opisthokonta |
| 1.00E-30 | Opisthokonta |
| 1.00E-30 | Opisthokonta |
| 1.00E-30 | Opisthokonta |
| 1.00E-30 | Opisthokonta |
| 1.00E-30 | Opisthokonta |

| | |
|---|---|
| 1.00E-30 | Opisthokonta |
| 1.00E-30 | Stramenopilles |
| 1.00E-30 | Stramenopilles |
| 1.00E-30 | Stramenopilles |
| 1.00E-30 | Stramenopilles |
| 1.00E-30 | Stramenopilles |
| 1.00E-30 | Stramenopilles |
| 1.00E-30 | Stramenopilles |
| 1.00E-30 | Stramenopilles |
| 1.00E-30 | Stramenopilles |
| 1.00E-30 | Stramenopilles |
| 1.00E-30 | Viridplantae |
| 1.00E-30 | Viridplantae |
| 1.00E-30 | Viridplantae |
| 1.00E-30 | Viridplantae |
| 1.00E-30 | Viridplantae |
| 1.00E-30 | Viridplantae |
| 1.00E-30 | Viridplantae |
| 1.00E-30 | Viridplantae |
| 1.00E-30 | Viridplantae |
| 1.00E-30 | Viridplantae |
| 2.00E-17 | Viruses |

**Table S3 - Proteins with PAN domain in alignment.**

| Organism | Gene name | Uniprot identifier |
|---|---|---|
| *H. sapiens* | HGF | P14210 |
| *A. thaliana* | SD16 | Q9S972 |
| *C. elegans* | LET-653 | Q27394 |
| *M. musculus* | HGF | Q08048 |
| *R. norvegicus* | PLG | Q01177 |
| *Z. mays* | PK1 | P17801 |
| *B. taurus* | PLG | P06868 |
| *C. lupus* | HGF | Q867B7 |
| *D. melanogaster* | NOMPA | A1Z8J4 |
| *D. rerio* | ADGRV1 | Q6JAN0 |
| *F. catus* | HGF | Q9BH09 |
| *O. sativa* | H0315F07.2 | Q25A00 |
| *P. patens* | PHYPA 026294 | A0A2K1ISB0 |
| *P. trichocarpa* | POPTR 001G409300 | A0A2K2CBN5 |
